# Single-molecule behavior and cell-growth regulation in human RTKs

**DOI:** 10.64898/2025.12.29.696957

**Authors:** Mitsuhiro Abe, Masataka Yanagawa, Yotaro Koizumi, Ryoji Kise, Asuka Inoue, Yasushi Sako

## Abstract

Receptor tyrosine kinases (RTKs) are a major family of cell surface receptor proteins responsible for various cellular functions in animal cells, including fate decisions, metabolism, polarization and migration. Lateral mobility, dimerization, clustering, and oligomerization are crucial behaviors in the activation process of RTKs on the cell surface. However, relationships between these molecular behaviors and molecular function remain to be elucidated, except for a few RTK members. Here, using an automated live-cell single-molecule imaging and analysis system, we studied the behavior of 52 of the 58 human RTK species on living cells over time during stimulation with ligands. We extracted 72 single-molecule parameters for each RTK species to examine their relationship to function, structure, and evolution. We noticed that RTKs’ ability to inhibit or support cell growth, as observed in a large-scale loss-of-function experiment in the public domain, significantly relates to their behavior. Growth-inhibitory signaling was coupled with the immediate formation of immobile clusters, followed by the enlargement of immobile and slow-mobile domains. In contrast, growth-supportive signaling coupled with higher lateral diffusivity and delayed clustering of immobile molecules. The relationship between structure and function suggests that functional differences are related to partitioning into membrane rafts and changes in mobility associated with phosphatidylinositol turnover. In multiple linear regression models, molecular behavior explained half or more of the molecular function related to cell growth. This level of explainability is comparable to that of evolutionary grouping.

## Introduction

Most of the activations and activities of membrane proteins require molecular processes across the lipid bilayer. Because rearranging inter- and intramolecular assemblies is a simple method to transmit information across membranes, membrane proteins often alter their clustering state depending on their activity. Molecular rearrangement may affect the lateral mobility of proteins. On the other hand, the structure of the bio-membrane is heterogeneous and changes dynamically on the nm–μm scale (Maxfiled, 2002; Lingwood and Simons, 2010). The function of various membrane proteins has been observed to depend on the membrane structure (Levental and Lyman, 2023). Interactions with specific membrane lipid molecule regulate the function and behavior of membrane proteins. The bilateral regulation between the protein function and membrane environment causes complex changes in the protein behavior (Lajoie et al, 2009).

However, the general relationships between the behaviors and functions of membrane proteins are not substantially understood as long as we noticed. The structure of a protein must be a major factor in determining protein behavior, and the structure is determined by evolutionary processes. It would be worthwhile to consider the correlations and causalities among the evolution, structure, behavior, and function of membrane proteins. To obtain a general understanding, a comparison of a wide range of related membrane proteins is necessary. Here, we performed a large-scale behavioral measurement of the receptor tyrosine kinase (RTK) superfamily. In human, RTKs, which consist of 58 species, are the second largest superfamily of membrane proteins in the plasma membrane. We measured the single-molecule behavior of 52 of them on the surface of living cells.

RTK is a single-membrane spanning protein and processes information from extracellular ligands, which are usually peptides or proteins. Functions of RTKs are wide, including the regulations of cell proliferation and differentiation, carbohydrate metabolism, cell polarization and migration, and adhesion to substrates (Wintheiser and Silberstein, 2022). Dysregulations of RTKs can result in various genetic and de novo diseases (Robertson et al, 2000; Tomuleasa et al, 2014). Despite their diverse functions, RTKs have a general activation process: Since they are single-membrane spanning, homodimerization and/or heterodimerization are essential for activating their cytoplasmic tyrosine kinase. After that, RTKs usually form oligomers and clusters on the cell surface. Several RTKs are known to accumulate in the cholesterol/sphingomyelin-rich membrane domains known as membrane rafts. RTKs interact with membrane lipids, including glycolipids, cholesterol, sphingomyelin, and acidic lipids (Kim et al, 2021). These properties must influence both molecular assembly and lateral mobility during RTK activation. Actually, for the EGF receptor (EGFR/ERBB1), the most extensively studied RTK, complex behavioral changes with activation progression have been reported in detail (Chung et al, 2010; Needham et al, 2016; Hiroshima et al, 2018, Kozer and Clayton, 2018; Mundumbi et al, 2023, Abe et al, 2024).

Remaining questions include whether the observed behavioral changes in several RTKs after cell stimulation can be generalized to other RTKs, how RTK behaviors vary, what causes these variations, and, most importantly, whether these behavioral variations relate to functional variations between RTKs. Since RTKs are a membrane protein family with a strong expected behavior/function relationship, studies on RTKs will be a good starting point for studying other membrane protein families.

One reason for the delay in behavioral studies of membrane proteins on the cell surface seems to be the lack of methods to detect molecular behavior with the same level of resolution achieved in structural and evolutionary analyses. Although single-molecule imaging and tracking (Sako et al, 2000; Schutz et al, 2000) enabled detailed study of membrane protein behavior, this technique was time-consuming due to the lengthy sample preparation, measurement, and analysis processes until recently. Recent progress in molecular genetics, protein probes, robotics, and computation has greatly improved single-molecule imaging measurements, overcoming the past limitations (Yasui et al, 2018, Yanagawa et al, 2021; Walther et al, 2024) and enabling the analysis of membrane protein behavior on the meso- to large-scale (Yanagawa et al, 2018; Takebayashi et al, 2023; Watanabe et al, 2024). This study also employed an automatic in-cell single-molecule imaging system with a deep learning filter (Yasui et al, 2018) and a single-molecule analysis pipeline (Yanagawa et al, 2021).

Based on single-molecule measurement data covering most (∼90%) of the human RTK species, in this study, we aimed to clarify the general relationships between the evolution, structure, behavior, and function of RTKs. We observed both commonalities and variations in RTK behavior in resting state and response state over time following cell stimulation. The variation related to the RTK’s function in cell growth and is thought to be regulated by the dynamic membrane structure.

## Results

### Single-molecule tracking of human RTKs on the living cell surface

We observed the single-molecule behavior of 52 of the 58 human RTKs (Fig. 1A). The structure of the human RTK family is diverse. They are classified into 20 subgroups based on the sequence similarity in the tyrosine kinase (TK) domain (Robinson et al, 2000). The 52 RTKs in this study distributed across 19 subgroups. cDNAs for EPHA5, EPHA7, EPHA10, RYK, and LMR3 were not obtainable. MER protein was not expressed in the cells for unknown reasons. To image single molecules, the cDNA of each RTK was fused to Halo-tag (Fig. 1B) and expressed in HEK293A cells. The Halo protein was conjugated with the fluorophore SF650 in living cells to track the movement and fluorescence intensity changes of individual fluorescent particles on the basal plasma membrane (Fig. 1C and Supplement Movie S1) using an automated live-cell single-molecule imaging microscope (Yasui et al, 2018). Movies were acquired for the same fields before (resting condition) and 2, 12, and 22 min after cell stimulation with the appropriate RTK ligands (Table 1) or vehicle solution (response condition) (Fig. 1D).

**Figure 1.**
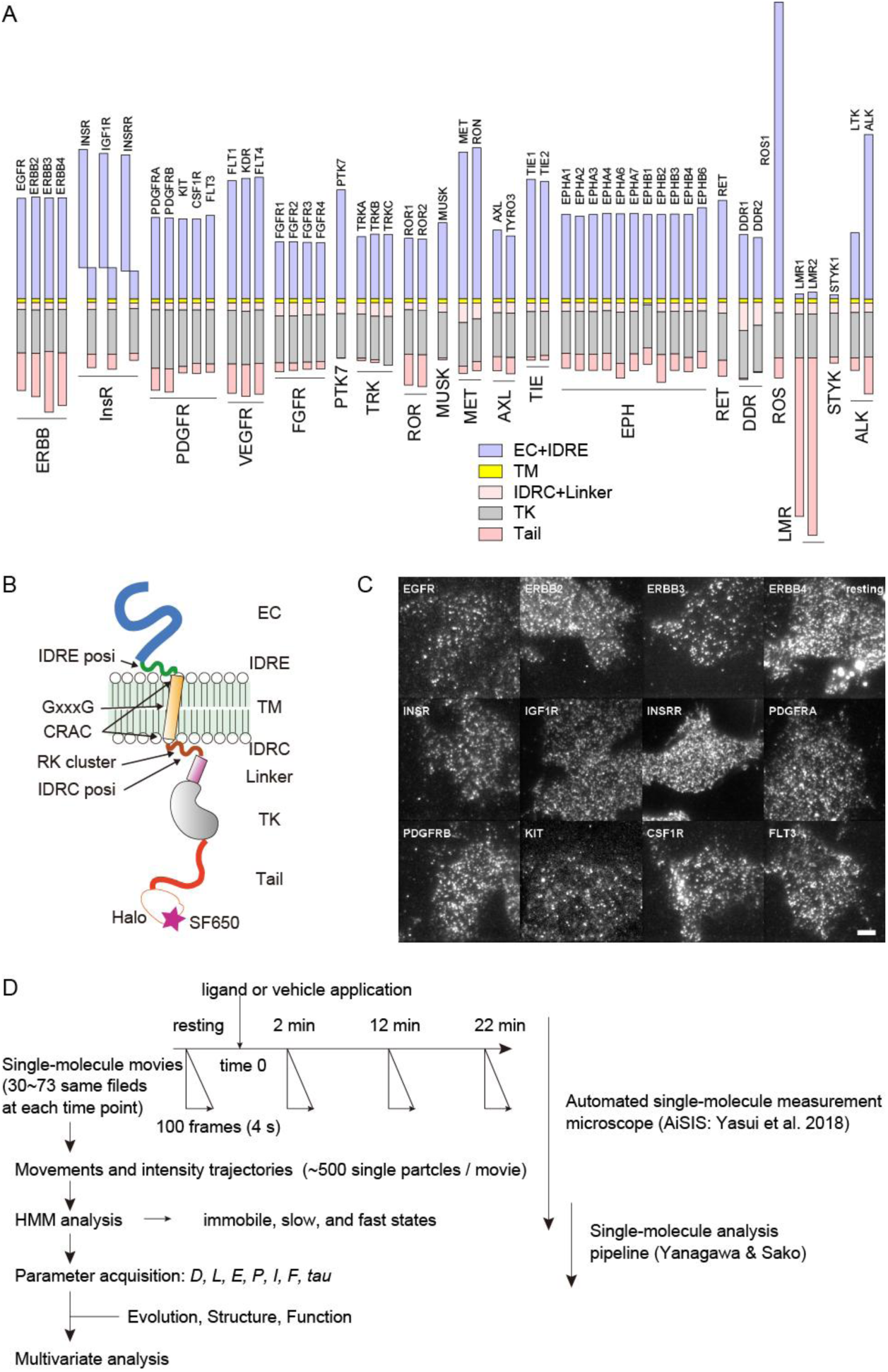
Single-molecule measurements of human RTK behavior. **A.** Structures of 52 human RTKs in this study. The names (in abbreviation) and subfamilies are indicated. EC: extracellular domain, IDRE: extracellular intrinsically disordered region, TM: trans-membrane domain, IDRC: cytoplasmic intrinsically disordered region, Linker: linker region between IDRC and TK, TK: tyrosine kinase domain, Tail: cytoplasmic tail after TK. EC domain of LMR1 and LMR2 may contain signal sequence being removed after maturation. **B.** RTK tagged with Halo-SF650 at the end of cytoplasmic tail. Positions of the structural regions and motifs are indicated. **C.** Examples of single-molecule images in living cells. Bar: 20 μm. (See Supplement Movie S1). **D.** Measurement and analysis procedure. Meanings of the parameters are listed in Figure 2A.

### Classification of the mobility state based on the hidden Markov model

To determine the parameters that should be extracted, we first applied a hidden Markov model (HMM) to classify the single-particle movements (Chung et al, 2010; Hiroshima et al, 2018), which assumes that there are temporal transitions among discrete random walk modes, each with a different lateral diffusion coefficient (*D*). The model was solved using variational Bayesian inference (Bishop, 2006). By varying the number of states (NoS) from one to five, the most likely NoS was determined that would provide the highest lower bound value on the average in movies under the same conditions (Supplement Fig. S1A). Interestingly, the 3-state model was the most probable for the 45 RTK species (87% of those examined), irrespective of ligand stimulation (Supplement Fig. S1B, and Supplement Table S1). For DDR1, DDR2, and ROS, 2-state was the most probable under all conditions. For ERBB2, INSRR, MET, and RON, 2- or 3-state was the most probable. Since the difference in lower-bound values between the 2- and 3-state models was not large for these RTKs, we adopted the 3-state model for all RTKs in this study to simplify comparison between species. According to the size of *D*, hereafter, we call the three mobility states as the immobile (state 1), slow (state 2), and fast (state 3).

### Single-molecule parameters for RTK behaviors

After the 3-state separation according to the result of HMM analysis, we extracted single-molecule behavioral parameters: lateral diffusion coefficient (*D*), fraction of movies exhibited no confinement (*E*), molecular density on the cell surface (*F*), single-particle fluorescence intensity (*I*), confinement length (*L*), particle fraction of the mobility mode (*P*), and state lifetime (*tau*). These parameter values were first determined for each single state in a single movie. Then they were averaged across all movies under the same conditions (Fig.2 and Supplement Table S2). Detailed explanations of the parameters can be found in the Supplement text, “Single-molecule parameters”. Please note that some of the parameter values are apparent, but we adopted them because they provide information about RTK behavior.

**Figure 2.**
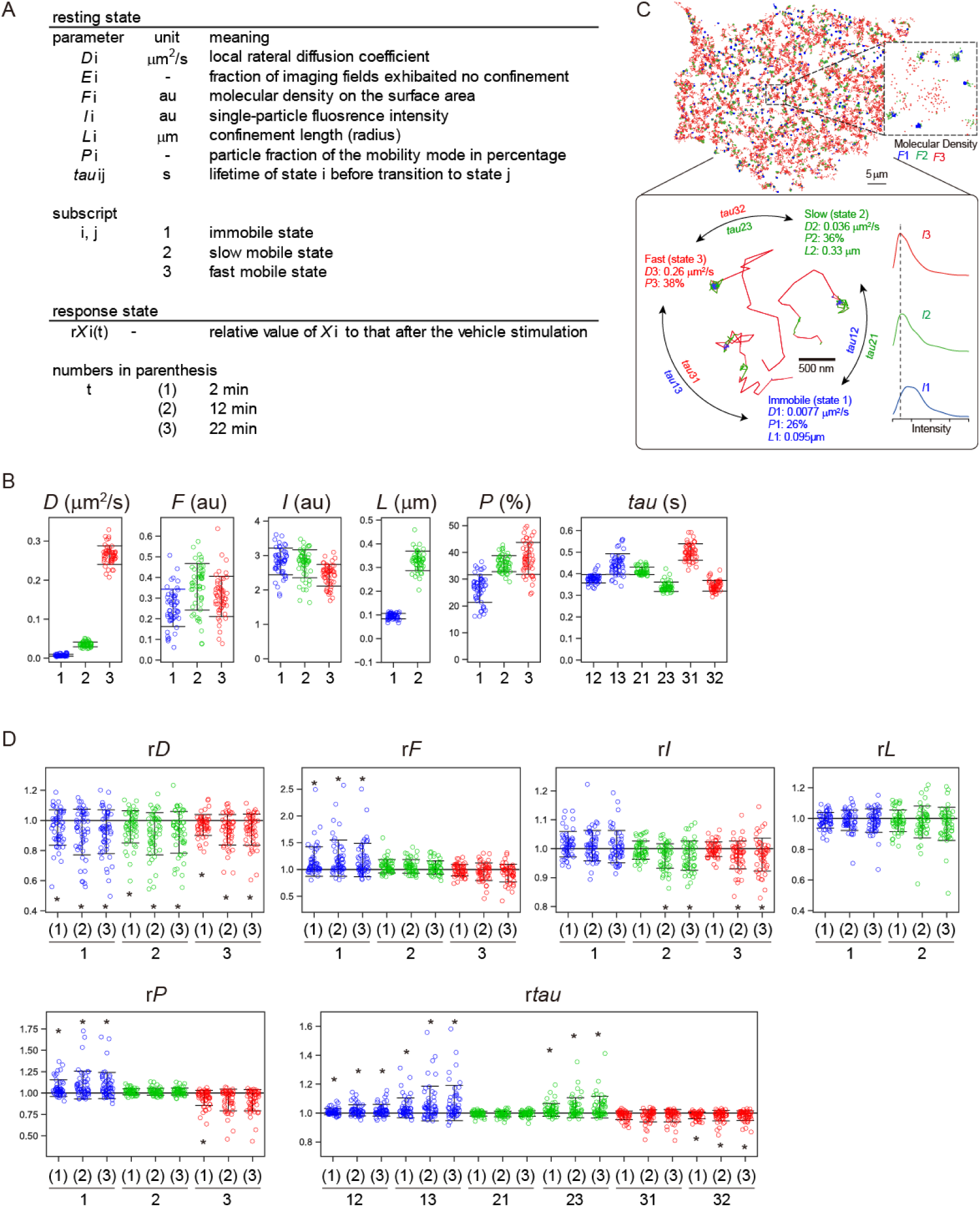
Single-molecule behavioral parameters. **A.** Meanings of single-molecule parameters. Low reproducible parameters *E* and *L*3 were not used for further analysis (see text and Supplement Fig. S2). Detailed explanation of the parameters is provided in Supplement text “Single-molecule parameters”. au: arbitrary unit. **B, D.** Distributions of the parameter values among RTK species in the resting (**B**) and response (**D**) states. See Supplement Table S2 for the values. Error bars represent the range of ±1SD. Asterisks in (**D**) indicate statistical difference (p < 0.01) from the vehicle stimulation in t-test after one-way ANOVA for the 3 timepoints. **C**. Cartoon showing the average resting state behavior.

Hereafter, parameter *X*i (i = 1, 2, or 3) refers to the value of parameter *X* in the immobile (i = 1), slow (i = 2), or fast (i = 3) state. *tau*ij refers the lifetime of state i before transitioning to state j. In the resting state, these parameters were indicated in the units listed in Figure 2A (upper panel). After ligand stimulation (the response state), they were indicated as ratios to the values in cells stimulated with vehicle. This makes it easier to compare RTKs by removing variations in basal levels. The numbers in the parentheses for the response state parameters, such as r*X*i(j), indicate the time point: 2 (j = 1), 12 (j = 2), and 22 (j = 3) min. Here, the prefix, r, indicates that it is a relative value.

### Measurement reproducibility

Before conducting detailed analyses, we verified the reproducibility of the parameter values by comparing the results of two resting state measurements using basically the same population of transfectants (Supplement Fig. S3 and Supplement Table S3). Most of the parameters were sufficiently reproducible. The exceptions were *L*3 and *E*1-*E*3. For the other parameters, the ratio of the parameter values between the two measurements was within the range of 0.996-1.109 on average across RTKs, with SD < 0.204. The correlation coefficients (R) between the two measurements were > 0.833. Since *F* (molecular density) is a secondary parameter derived from the particle number, particle intensity, and the cell area, the calculation noise for *F* should be greater. Excluding *F*1-*F*3 resulted in parameter ratios 0.996-1.021 (SD < 0.082), and R > 0.837. The reproduction of *L*3 and *E*1 was poor (R < 0.16) due to the large (*E*1) and small (*L*3) confinement effects for the immobile and fast mobile movements, respectively. The reproduction of *E*2 and *E*3 was modest (R = 0.680 and 0.666, respectively) due to stronger confinement effects for the slow and fast movements. We did not use *E*1-*E*3 nor *L*3, as well as their response-state counterparts (r*E*1*–*r*E*3 and r*L*3) in the subsequent analysis.

### General behavior of RTKs

We obtained the distributions of single-molecule parameters of RTKs in their resting and response states (Fig. 2B-D and Supplement Table S2). In the resting state, the three mobility states were characterized with *D* = 0.0077 (immobile), 0.036 (slow), and 0.26 (fast) μm^2^/s with *P* = 26% (immobile), 36% (slow), and 38% (fast), and *L =* 95 (immobile) and 330 (slow) nm. These observations are basically similar to those we have reported for EGFR (Hiroshima et al, 2018). *I* was largest and smallest for the immobile and fast states, respectively. *tau* was long for the transitions between the immobile and fast states (*tau*13 and *tau*31), which probably reflects the local membrane structure, i.e., immobile regions are surrounded by the slow regions and segregated from the fast region of the bulk membrane, as was observed for EGFR (Hiroshima et al, 2018; Abe et al, 2024).

On average, *D* decreased for all three mobility states after stimulation. *F*1 and *P*1 increased with the decrease in *P*3. *I*2 and *I*3 decreased in the later stages (times 2 and 3). *tau* increased for transitions from the immobile state to the slow and fast states, and from the slow to the fast state, but decreased for the transition from the fast to the slow state. These changes in *tau* values indicate a shift in the population to slower states, particularly the immobile state after stimulation. This tendency is also similar to that observed for EGFR (Hiroshima et al, 2018; Maeda et al, 2022).

### Function/behavior relationship

To analyze the relationship between the behavioral variation and functional variation, we used a quantitative dataset from the DepMap project (BROAD institute; Arafeh et al. 2025) to indicate RTK function. This dataset contains genome-wide CRISPR loss-of-function experiments for individual genes performed in 1,095 human cancer-derived cell lines to determine the effects on cell growth. We accumulated the results for the 52 RTK genes (Supplement Table S4). Positive values of the CRISPR factor indicate that the loss of function was beneficial to cell growth, i.e., the RTK is inhibitory to cell growth. Negative values indicate the opposite.

The simple correlation coefficients between the behavioral parameters and the CRISPR factors in single cell lines were distributed -0.61 < R < 0.50 (Supplement Table S5A). Although unignorable correlations (R < -0.6) were observed for two cases, given the large number of combinations, these R values could be the result of chance. To explore the possibility that multiple behavioral parameters collectively determine the RTK function, we performed a multiple correlation analysis to construct linear regression models for single cell lines using the resting or response state parameters (Fig. 3 and Table 2). In these models, we needed to reduce the number of behavioral parameters (explanatory variables) for several reasons in statistics (see Supplement text “Parameter selection for multivariate analyses”). We used the same sets of behavioral parameters for all cell lines expecting that the function of individual RTKs would be fundamentally similar across most cell lines, with variations appearing in cell-line-specific values of the partial regression coefficients (pRs) for each behavioral parameter. The optimal parameter sets were determined by striking a balance between the penalty for using a large number of parameters and the loss of explainability that occurs when meaningful parameters are discarded. As a result, 5 or 12 parameters were selected for the final models using resting or response state parameters, respectively, regardless of the statistical significance threshold (p = 0.01 ∼ 0.1; Fig. 3A, C).

**Figure 3.**
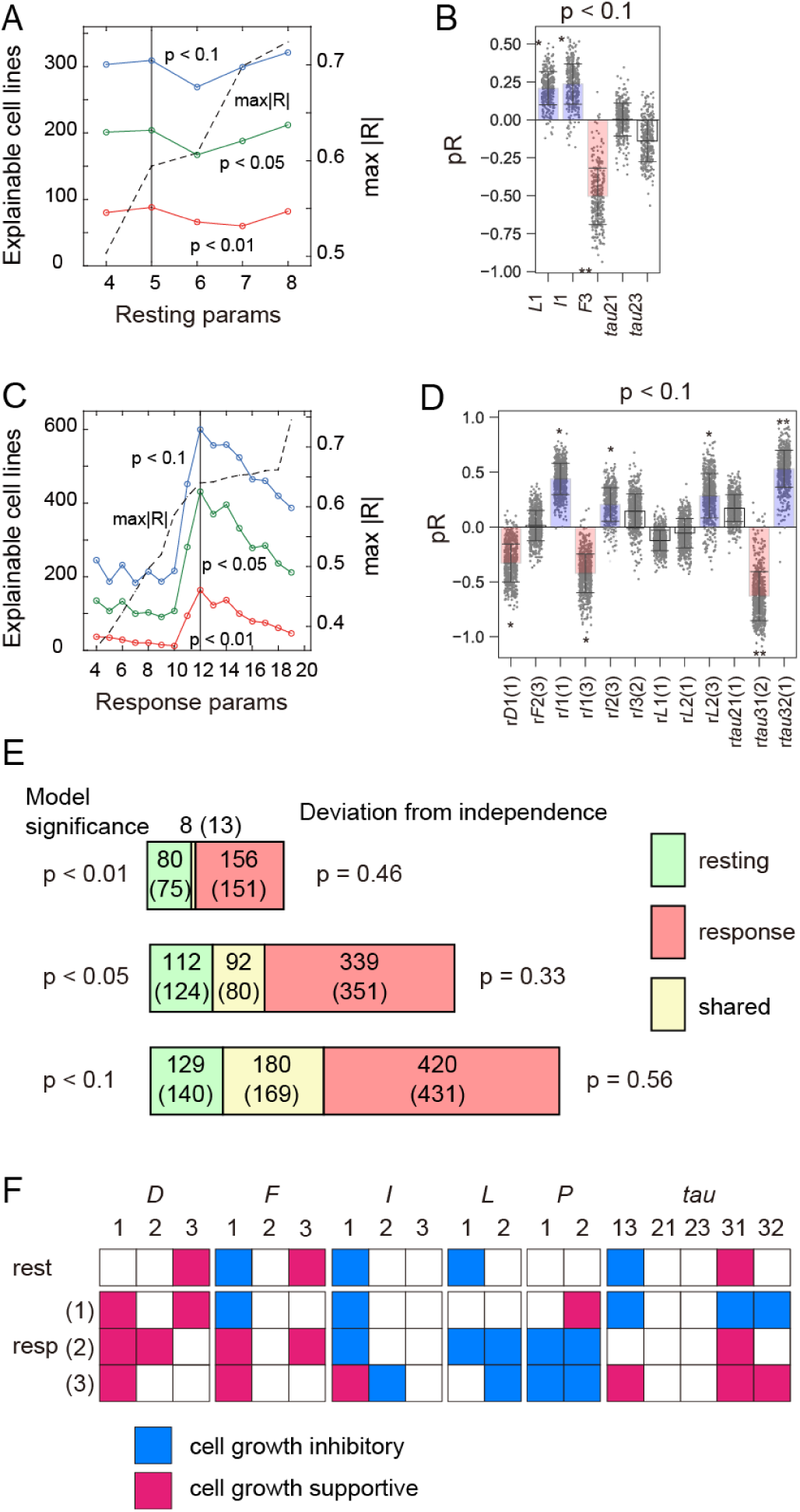
Single-molecule behavior models to explain RTK function. **A, C.** Parameter selection for constructing linear regression models from the resting (**A**) or response (**C**) behavior to the CRISPR factors in single cell lines. The vertical line indicates the condition used in the final models. See Supplement text for the parameter selection. **B, D.** Distributions of pRs in the final resting (**B**) or response (**D**) models with the statistical significance, p < 0.1. Bar: average. Asterisks: average |pR| * > 0.2 and **| > 0.5. **E.** Coincidence of the explainable cell lines between the resting and response models. Numbers in parentheses indicate expectations assuming independence between the two models. p-values (right) shows deviation of the observation from the assumption of independence (χ^2^-test). **F.** Significant (|pR| > 0.2) parameters in the final models, and parameters highly correlated with them but dropped in the process of modellings are indicated.

These parameter sets constructed statistically significant models (p < 0.1) for a large population of cell lines: 309 and 600 out of 1,095 in the resting and response state models, respectively. These population numbers far exceed the expectations of the null hypothesis (1,095 x 0.1 cell lines), indicating a strong relationship between behavior and function. The average explainability, or the average of the determination coefficient (R^2^) in these models, was 0.25 and 0.42 for the resting and response models, respectively. In other words, RTK behaviors in the resting and response states explained their function at least partially (25 or 42% on average) in these cell lines. Although the models did not show statistical significance for other cell lines, the profile of pRs was qualitatively similar, regardless of the model significance (Table 2). In terms of both the number of significant models and the size of explainability, the response state behavior explained RTK functions better than the resting state behavior. The coincidence between the cell lines explained by the resting state and response state models suggested that these two models almost independently explained RTK function (Fig. 3E).

### Growth supportive and growth inhibitory behaviors

As shown by the positive and negative average values of the CRISPR factors across cell lines, there are growth-inhibitory and growth-supportive RTKs (Supplement Table S4). The behavioral parameters associated with these signalings were identified as having significantly large absolute pR values in the significant models to the CRISPR factors (Fig. 3B, D). In addition to these parameters that remained in the final models, the other parameters that were correlated with them but dropped during the parameter selection (Supplement Table S6A) are the candidates responsible for signaling (Fig. 3F).

The behavioral parameters that contribute to growth-supportive and growth-inhibitory signaling appear to follow certain rules rather than occurring randomly (Fig. 3F): In the response state, pRs for r*D* (relative diffusion coefficient) generally positive, meaning larger *D* values were generally supportive of cell growth. In contrast, larger r*L* values (relative confinement length) were inhibitory. Longer lifetimes of immobile and fast states (r*tau*13, 31, and 32) resulted in separation between immobile and fast states. This separation was inhibitory at the early stage (2 min) but supportive of cell growth at the late stage (22 min). Roles (i.e., signs of pR) of *F*1 (immobile molecular density), *I*1 (immobile signal intensity), and *P*2 (slow particle fraction) also exhibited stage-specific changes. All responsible parameters in the resting state were also responsible in the response state. The signs of pR in these resting state parameters were similar to those for their relative values observed in the early to middle stages of the response state. Such systematic difference between the RTK behavior responsible for growth-inhibitory and growth-supporting signaling suggests that specific behavioral modes are linked to RTK function. Based on the responsibilities of r*D* (relative diffusion coefficient) and r*L*, (relative confinement length) we temporarily refer to growth-supportive and -inhibitory signaling as *D*-type and *L*-type, respectively.

### Structure and function

To examine the relationship between structure and function, we extracted 11 structural parameters from the amino acid sequence of RTK. These parameters included sequence lengths in the 6 regions (EC, extracellular domain; IDRE, extracellular intrinsically disordered region; TM, transmembrane domain; IDRC, cytoplasmic intrinsically disordered region; Linker, linker between IDRC and tyrosine kinase domain; Tail, cytoplasmic tail), as well as 5 parameters for the numbers of functional amino acids and functional motifs (IDRE posi, CRAC, GxxxG, RK cluster; IDRC posi; see Supplement text “Structural parameters”, Fig. 1B, Table 1, and Supplement Table S7). We did not consider deviation in the structures of the tyrosine kinase (TK) domain, since it is small among RTKs.

Despite the modest simple correlation (−0.50 < R < 0.49) between structural and functional parameters (Supplement Table S5B), statistically significant (p < 0.1) linear regression models using multiple structural parameters were constructed for RTK functions in 304 cell lines with the average determination factor of 0.23 (Fig. 4A, B and Supplement Table S8A). This result was comparable to that of the resting state models but worse than the response state models for function (Fig. 3). The profile of pR values in most models was similar, regardless of model significance (Supplement Table S8A).

**Figure 4.**
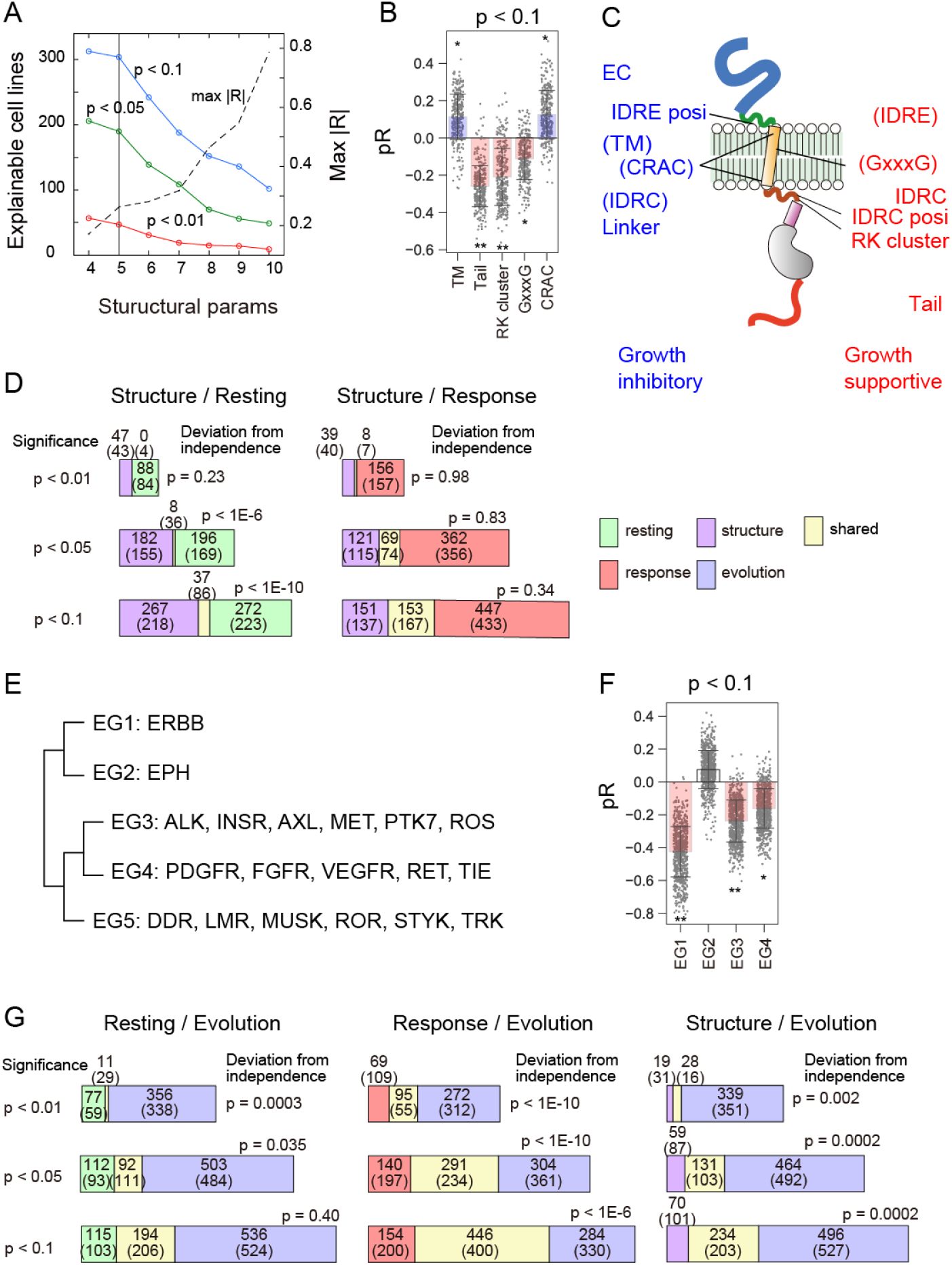
Structure and evolution models to explain RTK functions. **A.** Parameter selection for the linear regression model from structure to function. The vertical line indicates the condition used for the final modeling (See Supplement text for the parameter selection). **B, F.** Distributions of pRs in the structure (**B**) or evolution (**F**) models to function with the statistical significance, p < 0.1. Bar: average. Asterisk indicate average |pR| * > 0.1 and ** > 0.2. **C.** Structural parameters with |pR| > 0.2 (without parentheses) and > 0.1 (in the parentheses), respectively. **D, G.** Coincidence of the explainable cell lines between the structure models and behavior models (**D**) or between the evolution models and other models (**G**) to function. Numbers in parentheses indicate expectations assuming independence between the two models. p-values (right) shows deviation of the observation from the assumption of independence (χ^2^-test). **E.** Evolutionary grouping of RTKs based on the phylogenetic tree.

The structural parameters responsible for explaining RTK function were distributed systematically (Fig. 4C): For the growth-supportive (*D*-type) signaling, parameters in the regions of TM to Tail, especially those in the IDRC were mainly responsible. In EGFR, one of the representative growth-supportive RTK, involvement of these parameters in structure and dynamics changes after ligand stimulation has been reported (see Discussion). On the other hand, growth negative (*L*-type) parameters were mainly distributed in the EC to TM region. These parameters are important for distributing the protein into cholesterol (CRAC) and GM3 (IDRE posi)-rich membrane domains (membrane raft). Such tendencies were not limited to the models with high explainability but common to all models (Supplement Table S8A). The explainable cell lines were exclusive between the structure and resting state models, but independent between the structure and response state models (Fig. 4D).

### Evolution and function

The explainability of the structural models for RTK function was limited. This may be because the chosen structural parameters are too simple to describe complex behavior. However, it is difficult to parametrize the complicated structure of proteins. Instead, we used evolutionary relationships between RTK species, which contain higher-order structural relationships. Here, based on a phylogenetic tree of human tyrosine kinases (modified from Robinson et al. 2000; see Supplement text “Evolutionary parameters”), we categorized RTKs into 5 evolutionary groups (EG1–EG5; Fig. 4E).

After encoding the evolutionary categories as dummy variables (Supplement text “Evolutionary parameters”), we found that the range of simple correlations between the evolutionary parameters and functions was -0.77 < R < 0.49 (Supplement Table S5C). Highly negative correlations (R < -0.6) were observed between EG1 (EGFR family) and 82 (7.5%) cell lines. It may be worth noting that even evolutionary lineage explains less than R^2^ = 59% of RTK function. Evolutionary parameters could construct significant linear regression models (p < 0.1) in 730 cell lines, with an average R^2^ = 0.27 (Fig. 4F and Supplement Table S8B). As observed in other multiple linear regression models, the model structure (i.e., the profile of pRs) was similar in most cell lines, regardless of statistical significance. This indicates a commonality in the evolution/function relationship across most cell lines (Supplement Table S8B). The group of explainable cell lines in the evolution models was independent of that in the resting state models, but coincided significantly with those in the response state and structure models (Fig. 4G).

### Regression models for generalized RTK function

So far, we have constructed regression models to explain RTK function for individual cell lines. However, it would be desirable to model for the generalized cell response to the loss of RTK function. To this end, we applied principal component analysis (PCA) on the CRISPR data (Supplement Fig. S3A). The PCA score for PC1 is particularly high (47%), and its loading vector is nearly proportional to the cell-to-cell average value of each RTK’s CRISPR factors (Supplement Fig. S3B). The following three components (PC2-4) are rather large but smaller than 6% for each PC, which reflect cell line-specific RTK function. We used the PC1 loading vector as the generalized response.

Using the parameter sets employed in the final models for single cell lines (Figs. 3 and 4), we constructed linear regression models from the resting, response, and structural parameters for the general RTK function as observed in PC1 (Supplement Fig. S3C). The parameters that significantly contributed to the resting state model (|pR| > 0.2) were the same as those in the models for single-cell lines. Although significant parameters in the response and structure models changed slightly, they again suggested *D*- and *L*-type signaling (Supplement Fig. S3D, E). The explainability scores for the resting, response, and structure models were 0.19, 0.49, and 0.17, respectively.

### Structure, evolution, and behavior

The simple correlation coefficients between the structural and behavioral parameters were within the range of -0.39 < R < 0.60 indicating that no single structural parameter strongly correlates with any RTK behavior (Supplement Table S5D). To clarify the collective correlation between structural and behavioral parameter groups, we carried out a canonical correlation analysis (CCA). For simplicity, we only focused on the largest component, CC1 (Table 3A, B). Significant parameters (absolute values of the canonical correlation coefficient, |CR1| > 0.2) and their highly correlated parameters (|R| > 0.7) were detected (Fig. 5A and Supplement Table S6). Structural parameters in the IDRs and TM primarily influence molecular behavior, either positively or negatively. The behavioral parameters significantly involved in CC1 included 56% (10/18) of the resting state parameters and 31% (17/54) of the response state parameters. These fraction sizes of parameters indicate that the resting state behavior were correlated more closely with the structure. Conversely, 36% (4/11) and 73% (8/11) of the structural parameters were significantly correlated with the resting and response behaviors, respectively. It may be interesting that the parameters in IDRE and IDRC were correlated in the opposite direction to the response state parameters. Correlation between the resting state and response state parameters was also analyzed in CCA (Fig. 5B and Table 3C).

**Figure 5.**
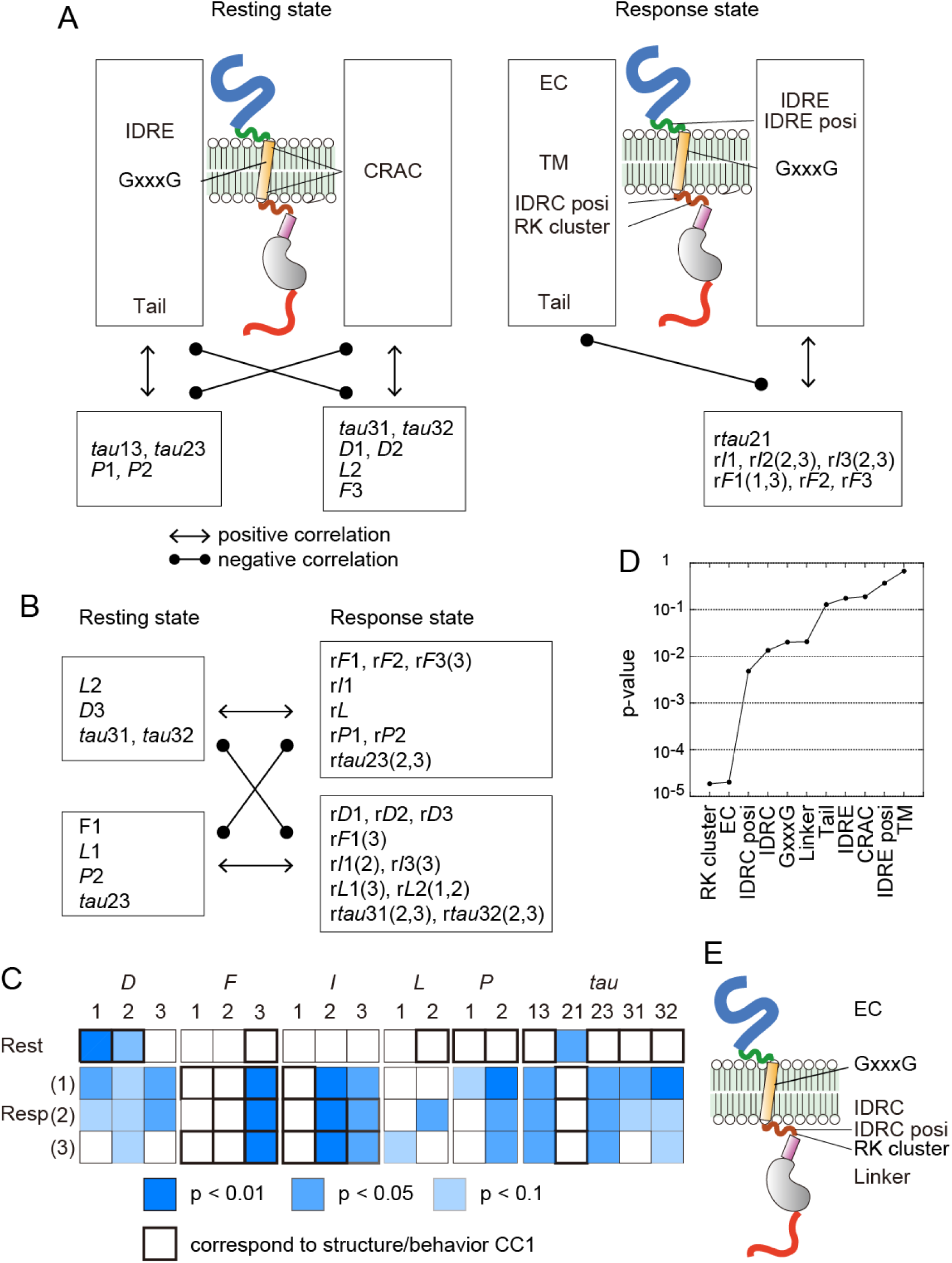
Relationships between structural, behavioral, and evolutionary parameters. **A, B.** Significant parameters found in CCA (|CR1| > 0.2, Table 3) between structure and behavior (**A**), and between resting and response state behaviors (**B**). Arrows indicate positive or negative correlation between the parameter sets. **C.** Behavioral parameters significantly explainable (p < 0.1) using a linear regression model from the evolutionary parameters. **D.** Significance of the evolution models to the structural parameters. **E.** Map of the structural parameters with p < 0.05 in (**D**).

Next, the relationships between evolution and behavior were considered. Since simple correlations were not high (−0.52 < R < 0.50; Supplement Table S5E), we constructed linear regression models from evolution to RTK behaviors after dropping EG5 from the evolutionary parameters to avoid multicollinearity (Fig. 5C and Table 4). In the resting state, statistically significant (p < 0.1) models could be composed for 3 of the 18 parameters (*D*1, *D*2, *tau*21). The average determination coefficient (explainability) was 0.19 (0.10 for all parameters). In contrast, significant models were obtainable for 33 of the 54 response state parameters, with an average determination coefficient of 0.22 (0.17 for all parameters). Evolution better explained the response behavior than the resting behavior of RTKs, which contrasts with the explanation using structural parameters (Fig. 5A). Notably, the behavioral parameters responsible for the CCA between structure and behavior were largely distinct from those explained by evolution (Fig. 5C).

Finaly, the relationship between structure and evolution was examined. The single correlation coefficients between the structural and evolutionary parameters were in the range of -0.50 < R < 0.62. High correlation (R > 0.6) was observed only between EG1 and RK cluster (Supplement Table S5F). However, using multiple parameters collectively, statistically significant (p < 0.05) linear regression models could be constructed from evolution for 6 of the 11 structural parameters, with the average determination coefficient of 0.30 (Fig. 5D, E, and Supplement Table S8C). This result indicates that these 6 parameters are highly conserved throughout evolution. Of the 6 parameters, 4, excluding IDRC and Linker, supported CC1 between structural and behavioral parameters (Fig. 5A). All 4 of these parameters were found in the CC1 between the response state parameters. However, only one of them, GxxxG, was also found in the CC1 between the resting state parameters.

### Comparison of the explainability to RTK function

The explainability to the RTK function (PC1 of the CRISPR factors) was compared across evolutionary, structural, and behavioral parameter sets (Fig. 6A and Supplement Fig. S4). The resting and response state models explained R^2^ = 19% and 49%, respectively, of the RTK function in cell growth. Simultaneously using the resting and response state parameters increased explainability to 57%. This value is smaller than the sum of the explainability of resting and response state behaviors because there was redundant information between the two parameter sets. The evolution model and the structure model explained 37% and 16%, respectively, of the RTK function. In the above analyses, we categorized RTKs into 5 evolutionary groups. Therefore, we did not use all the information about RTK evolution. Explainability of evolution increased to 60% when RTKs were categorized into the full 19 groups.

**Figure 6.**
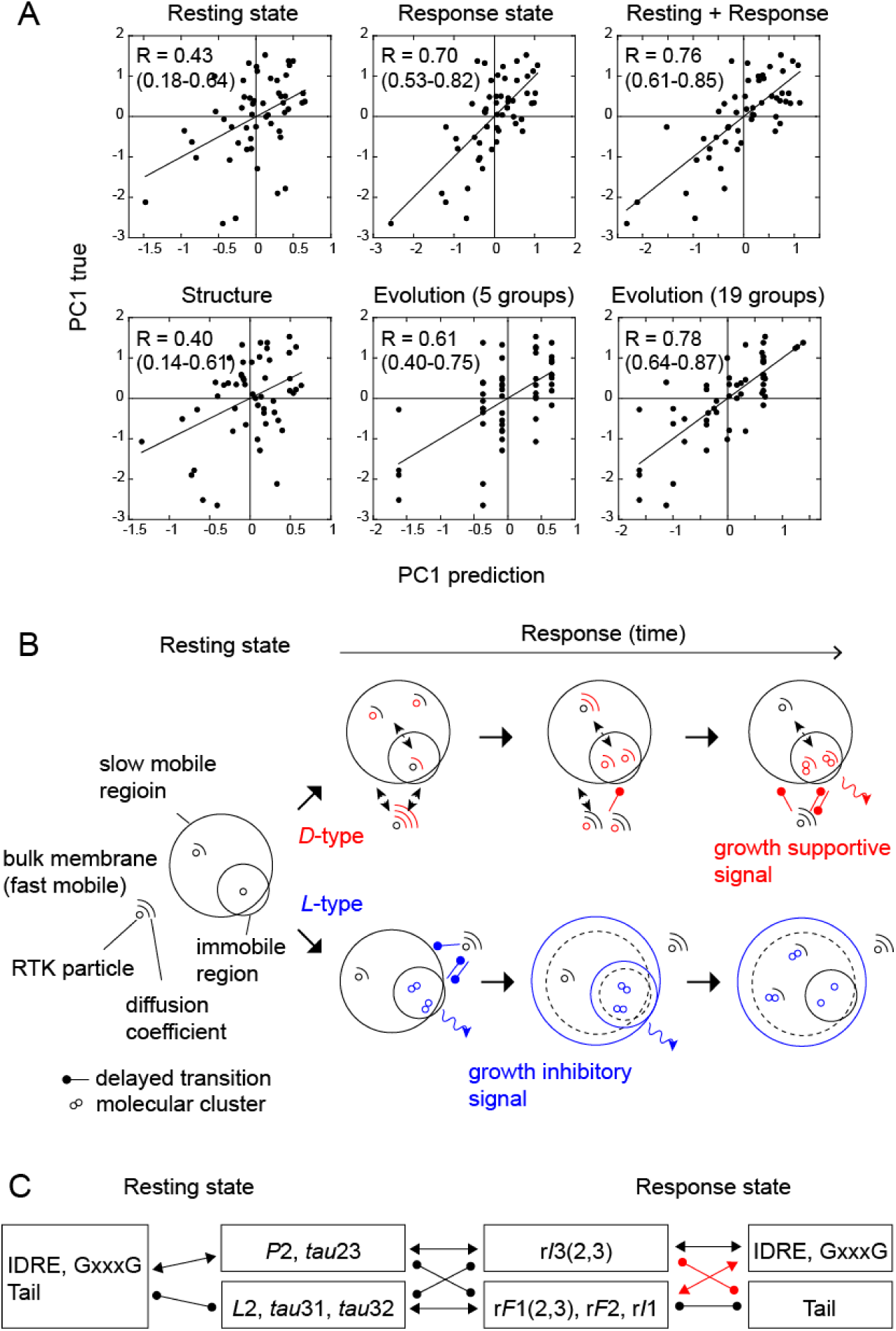
Behavior and function of RTKs. **A.** Explanation of PC1 of the CRISPR factors using various parameter sets. Solid lines show correlation between the true and predicted values. R: correlation coefficient. Numbers in parentheses indicate the 95% confidence intervals. **B.** Models of RTK behavior in growth-supportive (*D*-type) and inhibitory (*L*-type) signaling. *D*-type signaling involves stimulated diffusion coupled with PI dynamics, while *L*-type signaling occurs after immediate molecular clustering in the immobile region. Red or blue color indicate parameters responsible to explain RTK function. Increases in *F* and *P* are not distinguished in this cartoon for simplification. (See text for details.) **C.** Parameter linkage suggested by CCA between structural and behavioral parameters. Red arrows indicate changes with RTK signaling.

## Discussion

### General behavior of RTKs

Applying HMM, we have previously reported that the 3-state transition is the basic nature of the lateral diffusion of EGFR on the cell surface (Hiroshima et al, 2018). Here, we found that the 3-state transition is common to most RTK species. For some GPCRs, the 4-state model was the most probable (Yanagawa et al, 2018; Kawakami et al, 2022; Kuramoto et al, 2025). This suggests that the protein structure, e.g., the number of membrane-spanning helices, determines the basic nature of mobility. However, we also observe a 3-state transition in the movements of the TRPV1 and TRPV4 ion channels, which are 6 membrane-spanning proteins forming tetramers (Kuwashima et al, 2021, Kuwashima et al, 2024). The number of membrane-spanning helices is not the sole determinant. In the *Dictyostelium* cell membrane, it has been reported that membrane proteins with one to ten membrane-spanning helices commonly exhibited a 3-state mobility mode (Takebayashi et al, 2023). The membrane structure, which is distinct between mammals and *Dictyostelium*, could be another determinant of NoS. Additionally, we found confinement of the moving area in the two slower states (immobile and slow) as the common feature of RTK movements. This has previously been found in the movements of EGFR (Hiroshima et al, 2018). Since HMM only considers local movement (the distribution of single-step size), confinement is not a natural consequence of slow diffusion. The 3-state transition and the confinement of slower states indicate that RTK movements belong to a specific class of membrane protein dynamics.

Note that these observations are based on the view of HMM, i.e., transitioning among discretized states. The real nature of the RTK mobility may be continuous and gradual change in the mobility parameters. Methods for addressing continuous variations in single-particle trajectories have been proposed (Kowlek et al, 2019; Heckert et al, 2022). We used HMM in this study because it provides a simple way to organize the complex properties of the membrane protein movements. Additionally, the algorithm to solve HMM is available for large-scale analysis (Persson et al, 2013).

In the response state after stimulation with ligands, the majority of RTKs decreased their diffusion coefficient (*D*) in all mobility states, while increasing the particle fraction (*P*) and molecular density (*F*) of the immobile state. Though it was not detected on average, several RTKs evidently increased the size of molecular clustering (*I*) in the immobile state. On the contrary, *I* in the slow and fast states decreased in the later stage of signaling. These shifts in the behavioral parameter values were coupled with the increases and decreases in the state lifetimes for transitions to the slower states (*tau*12, 13, and 23) and from the fast state to the slow state (*tau*32), respectively. In other words, the slower mobility states were stabilized while the fast state was destabilized. These phenomena suggest that RTKs become entrapped in stable membrane domains in correlation with signaling. Clustering in the immobile state was observed for the singling of EGFR to a cytoplasmic protein, GRB2 (Hiroshima et al. 2018).

### Signaling modes specific to cell growth regulation

Multiple linear regression models identified behavioral parameters that positively and negatively impact cell growth (Fig. 3). Based on the parameters that contributed in the models, we refer to the growth-supportive and inhibitory signaling as *D*-type and *L*-type, respectively. Large r*I*1 (immobile clustering), which is crucial for signaling, was observed in *D*- and *L*-type signaling in the late and early stages, respectively, together with large r*F*1 (immobile molecular density). This temporal difference was coupled with that in the separation of the fast state from the slower states. The contribution of resting state parameters to cell growth regulation was similar to that of the response state parameters in the early to middle stages, suggesting priming for signaling in the resting state.

The structural parameters positively responsible to the growth-supportive (*D*-type) signaling included positive charges in the IDRC (IDRC posi and RK cluster), GxxxG, and the IDRC length (Fig. 4C). The positively charged R and K residues in IDRC (juxtamembrane, JM, domain) are expected to interact with the negative charge in the headgroups of PS and PIP_2_. In particular, R and K in this region of EGFR interact with PIP_2_ (Matsushita et al, 2013; Abd Halim et al, 2015; Maeda et al, 2018). This interaction is believed to induce the JM dimerization, which stabilizes the TK dimer in the kinase active form (Jura et al, 2009; Macdonald-Obermann and Pike, 2009: Endres et al, 2013). GxxxG in the TM helix supports this process by inducing a crossed TM dimer in the membrane (Yano et al, 2015). The crossed form is shorter in the direction of membrane thickness and disrupts the alignment of lipid tails, thus, will cause it to distribute in the flexible and thin liquid-disordered phase. We observed transient increase in *D* for EGFR dimers after EGF stimulation (Hiroshima et al, 2018). This increase could be due to invasion of kinase-active dimers into disordered membrane regions. An important phenomenon is that PI dynamics caused by PLCγ activity under EGFR reduces the PIP_2_ interaction with time (Abe et al, 2024). Then, the JM dimers dissolved and EGFR formed clusters for signal transduction to cytoplasmic proteins (Hiroshima et al, 2018; Maeda et al, 2022). EGFR is a typical growth-supportive RTK, and this scenario may be applicable to other growth-supportive RTKs. Molecular dynamics simulations predict interaction between PIP_2_ and R and K in the JM region of many RTK species (Hedger et al, 2015). Signaling to PLCγ is also observed for multiple RTKs (Wintheiser and Silberstein, 2025). This process will take time, as shown by the delayed increase in *I*1 in *D*-type signaling (Fig. 6B).

On the other hand, the structural parameters positively responsible to the growth-inhibitory (*L*-type) signaling included GM3 (IDRE posi) and cholesterol-binding (CRAC) motifs, as well as the longer TM helix (Fig. 4C). These features should distribute the molecule into the GM3/cholesterol/sphingomyelin-rich domains (membrane rafts). In the membrane rafts, cholesterol and the acyl chains of sphingomyelins form a complex (Fantini and Barrantes, 2013), which gives lipid membranes a rigid, thick, ordered state. The long TM region standing perpendicular to the membrane plane will tend to distribute itself within the thick membrane raft. An increase in raft-philic tendency after signaling, for example due to structural changes in the EC domain, causes the accumulation of molecules, which may result in an increase in or maintenance of the domain size *L* (Fig. 6B). Whether the relatively small *D* after signaling is purposely involved in growth-inhibitory signaling is a future question. The functions of growth-inhibitory RTKs could include inducing cell differentiation and/or directional sensing, as has been observed for TRKB, C, and EPH families. The preservation of spatial information may be important for directional sensing.

### RTK behavior influenced by evolution and structure

Although molecular behavior should be determined by, or at least correlated with, the molecular structure, none of the structural parameters we chose were strongly correlated with any behavioral parameters. The weak simple correlation was commonly observed between almost all of the parameters examined in different categories in this study, suggesting that the causalities and correlations between evolution, structure, behavior, and function are supported by multiple factors. For this reason, we used CCA and multiple linear regression modeling to analyze the relationships among evolution, structure, and behavior (Fig. 5).

CCA detected significant parameters that contributed to the correlation between structure and behavior in the resting and response states. However, in the multiple regression models using evolutionary parameters, response behavior was explained more effectively than resting behavior. The evolution models for structure mainly explained the parameters in the TM to IDRC region. These structural parameters were significant in CC1 between structure and response state behavior. These results suggest that the 3-way relationship among evolution, structure, and behavior was primarily supported by the correlation between the structure in the TM to IDRC region and the response state behavior. Structural parameters that were correlated with the resting state behavior and with the response behavior independent of the evolution were also primarily distributed in the membrane region (IDRE to IDRC). Overall, the importance of interaction with membrane lipids for RTK behavior was suggested, aligning with the roles of structural parameters in the *D*- and *L*-type signaling. The size of tail domain (Tail) was another factor significantly correlated with the molecular behavior, likely due to its interaction with the membrane skeleton and/or cytoplasmic proteins.

As discussed above, evolution has preserved the structural parameters closely related to response behavior. However, an examination of all response parameters has revealed a substantial difference between the explainability by evolution and the correlation with the structure (Fig. 5C). For instance, r*I*1, which is crucial for activation, could not be well explained by evolution. However, r*I*1 exhibited a significant correlation with structure. r*I*1 and other behavioral parameters were collectively associated with many structural parameters (Fig. 5A), suggesting that multiple mechanisms produce response behavior that transcends evolutionary specificities. Figure 6B suggest that two distinct pathways lead to increased r*I*1.

### Parameter linkage

In the linkage between structural and behavioral parameters in CCA (Fig. 5A, B), we discovered that some structural parameters underwent role reversal after cell stimulation (Fig. 6C): Resting state parameters *P*2 and *tau*23 were positively correlated with the response state parameters r*I*3(2, 3). Tail was a structural parameter that was positively correlated with *P*2 and *tau*23, however, it was negatively correlated with r*I*3(2, 3). This discrepancy means that the role of Tail was reversed after cell stimulation. A similar role reversal was observed for IDRE and GxxxG after cell stimulation. The tail domain is the major phosphorylation site in most RTKs. And as discussed above, GxxxG could be involved in RTK dimerization. Ligand stimulation can cause the role reversal in these parameters to the behavior.

There was a strong mutual exclusivity between the cell lines that could be explained by structural and resting state models for function (Fig. 4D). Role reversal was observed in Tail when examining the linkage of parameters that significantly contributed to models of cell lines that could be explained by structure or response alone (Supplement Fig. S5A). This could explain why the two models were exclusive.

### Explainability of RTK function

We used regression models to analyze the explainability of the RTK function. Unlike correlation analysis, regression models contain directional relations that express explainability. In a multiple linear regression model, pR indicates the dependency of the response variable (e.g., the CRISPR factor) on a given explanatory variable (e.g., one of the behavioral, structural, or evolutionary parameters), holding constant all other explanatory variables. In other words, pR is determined through multiple relations between the explanatory variables.

The average behavior of RTKs was a decrease in *D* after ligand stimulation (Fig. 2C). Considering that growth-supportive signaling is a function of the majority of RTKs, the significantly negative contribution of r*D* to the CRISPR factor (Fig. 3F) appears inconsistent. However, r*D* is the relative value to that with vehicle stimulation, and the simple correlations between *D* and the PC1 of CRISPR factors were not strong (for example R = -0.31 for r*D*1(1); Supplement Fig. S5B). Moreover, as mentioned above, pR is determined by the relationship between multiple explanatory variables; therefore, the profiles of R and pR are distinct (Supplement Fig. S5B, C). The parameter contribution to the model of CRISPR PC (Supplement Fig. S5D) clearly shows that the model is supported by interactions between multiple behavioral properties.

As a result of collective effects of multiple parameters, the single-molecule behavior of RTKs analyzed in this study explained about half or more of the molecular functions, as measured by the determinant coefficient (Fig. 6A and Supplement Fig. S4). This value was comparable to that of the evolutionary parameters. This explainability may be limited by the accuracy of the measurements, the parametrization, and/or the model structure. Alternatively, the limit could stem from the intrinsic nature of the relationship between molecular behavior and higher-order cellular functions, such as cell growth.

## Materials and Methods

### Plasmid construction

cDNAs of human INSRR, CSF1R, FLT3, TRKA, TRKB, ROR1, AXL, MER, EPHB3, RET, LMR2, and ALK were obtained from plasmids #70392, #23928, #23895, #23891, #23883, #116789, #105932, #23900, #65443, #23906, #23914, and #23917 from Addgene, respectively. Other cDNAs were obtained from the MegaMan Human Transcriptome Library (Agilent Technologies) or Human Universal QUICK-Clone™ cDNA II (TaKaRa Bio) by PCR amplification. RTK–Halo was generated by replacing the EGFR fragment in EGFR–Halo (Abe et al., 2024) with each RTK fragment, in which a GGGGSGGGGS linker was inserted between RTK and Halo. We totally constructed expression vectors for 53 RTKs fused with Halo.

### Cell preparation

HEK293A cells (purchased from Thermo Fisher) were maintained in DMEM/F12 medium containing 10% FBS (cell culture medium) at 37°C, 5% CO_2_. Cells were transferred to a φ-10 cm dish and cultured to 100% confluence, then, treated with trypsin/EDTA and suspended in the cell culture medium containing 15 mM HEPES (pH 7.3). After adjusting the cell density to 2 x 10^5^ /μl, 300 μl of the cell suspension was mixed with 24-72 μl of a mixture containing cDNA of each RTK fused with Halo-tag (1-3 ng) and Lipofectamine 3000 reagent (Thermo Fisher). An aliquot of 108 μl mixture was transferred to each well of a 96-well poly-L-lysin-coated glass bottom plate (PLL View plate; Perkin Elmer) and incubated overnight at 37°C, 5% CO_2_. Before observations, cells were stained with 80 μl/well of 0.5 nM SaraFluor 650 (SF650) Halo-tag ligand (Goryo Kayaku) in DMEM/F12 without phenol red and NaHCO_3_ but supplemented with 15 mM HEPES (pH 7.3) and 10% FBS (Medium B) for 15 min at 37°C. Cells were washed thoroughly with Medium B and incubated in DMEM/F12 medium containing 15 mM HEPES (pH 7.3) and 0.01% BSA at 25°C. Since MER protein was not expressed in the cells, other 52 RTKs were examined.

### Acquisition of single-molecule movies

An improved version of the automated in-cell single-molecule imaging system (AiSIS; Yasui et al, 2018; Watanabe et al, 2024) was used for single-molecule observation on the living cell surface. The system is based on an inverted fluorescence microscope (Ti2E, Nikon) in the total internal reflection mode equipped with a 60x, NA 1.49 oil-immersion objective (PlanApo, Nikon) and an autofocus system (PAF, Zido). SF650 conjugated to Halo-tag was excited by a 637-nm diode laser (OBIS, Coherent). The fluorescence signal from SF650 was selected by a dichroic mirror (ZT-488/640rpc, Chroma) and an emission filter (FF02-676/29, Opto-line) and imaged onto an EM-CCD camera (C9100-23B; Hamamatsu) with a frame rate of 25 s^-1^. AiSIS first search 15 fields suitable for single-molecule imaging in each well (containing cells with ∼1 fluorescent spot μm^-2^ on the surface) using a deep-learning filter. Then, it records 100 frame movies at the selected fields successively before cell stimulation (resting state). AiSIS applies ligand or vehicle solution 15 min after the resting state imaging, using Omni Robot (Tecan), and acquires 100 frame movies at 2, 12, and 22 min after the application at the same fields (usually containing 1 or 2 cells). Images were acquired at 25 °C.

### Data availability

All movies and the csv data sets showing the results of tracking and HMM analysis (obtained by AAS) are available at ssbd-repos-000536, https://doi.org/10.24631/ssbd.repos.2026.08.536 in the SSBD repository (Kyoda et al. 2025).

A Python program for HMM analysis (Yanagawa et al. 2026) is available on GitHub (https://github.com/yanagawamasataka5z-oss/smDA-HMM). As far as we know, the algorithm in this program is analogous to that utilize in AAS. Raw movies for two individual imaging fields of cells expressing EGFR-Halo are presented as examples. These movies were recorded at four distinct time points before and after EGF stimulation. For a detailed overview of the experimental conditions and the configuration of the HMM analysis parameters, please refer to the README file available on GitHub.

### Parameter extraction and analysis

Tracking of the single-particle movements and fluorescence intensities was performed using AAS software (Zido). See Supplement text “Single-particle tracking” for details. Typically, more than 500 particles/movie were detected on the basal cell surface in 37-73 movies under each condition in the resting and response states. All movies were used, except for those in which particle detection failed due to defocusing. Trajectories of single particles were analyzed using a hidden Markov model (HMM) based on the difference in the lateral diffusion coefficients (Hiroshima et al, 2018), employing the variational Bayes inference algorithm built in AAS (Supplement text “VB-HMM analysis”). HMM analysis subdivided single trajectories into the preset numbers of motional modes. After determination of the most probable NoS, using the single-molecule analysis pipeline (Yanagawa and Sako, 2021), parameters of molecular movements, density, and clustering were estimated for each mobility mode as the average of particles in single movies. To calculate the particle density, cell surface area was estimated from the single-particle image (Yanagawa and Sako, 2021). The parameter values were averaged over the movies in the same condition for further analyses (Table 2). See supplement text for the details of single-molecule parameters.

Further statistical and multivariate analyses have been performed for the standardized parameters using scikit-learn library for Python. Multiple regression analysis was used when directional relations to the response variables were anticipated. However, canonical correlation analysis (CCA) was used when no such directionality could be assumed a priori. CCA evaluates correlations between parameter groups neutrally. CCA was also used to evaluate the relationship between structure and behavior. Although directional relationships could be anticipated in this case, there was no logical way to select parameters. In multiple regression modeling for function, the number of cell lines for which a statistically significant model was composed was counted to determine the optimal parameter set (see Figs. 3A, 3C, and 4A). This assumes commonality in cellular responses indicating RTK function. However, such commonality cannot be expected between structure models for different behavioral parameters.

## Legends for Tables

**Table 1.** Structure and ligand of 52 human RTKs. The amino-acid length, numbers of positively charged amino acids (R and K), and numbers of functional motifs are listed with the ligands and their concentrations used in this study. See Supplement text “Structural parameters” for the estimation of each region. Orphan RTKs and the collagen receptor, DDR1 and DDR2, were stimulated by EGF as the reference ligand.

**Table 2.** Multiple regression from behavior to function. Multiple regression models from the resting (**A**) or response (**B**) state parameters to the CRISPR factors. pR values in the final models for 1,095 cell lines are listed according to the p-values of the models in F-test. determ: determination coefficient (R^2^). Averages of the pRs and determination coefficients for the models with p < 0.01, 0.05, 0.1 and all models are shown at the end of the lists. pR > 0.2 and < -0.2 are indicated in blue and red, respectively. p-values in t-test for each pR are also shown. p < 0.05 are colored yellow. na: t-test not applicable since pR < -1.

**Table 3.** Canonical correlation between parameter groups. Results of canonical correlation analysis between structural and behavioral parameters in the resting (**A**) or response (**B**) state, and between resting and response state parameters (**C**). Canonical correlation coefficients in the canonical component 1 (CR1) are listed. CC and determ are the canonical correlation and the determination coefficient (CC^2^).

**Table 4.** Multiple regression from evolution to behavior. Patial regression coefficients (pRs) in the linear regression models are listed. determ: determination coefficient (R^2^), p-value: p-value of the model in F-test. p-values in t-test for each pR are also shown. p < 0.05 are colored yellow.

## Acknowledgement

The authors thank Kesu Dong and Hiromi Sato for technical assistance. YS is funded by Grants-in-Aid for Scientific Research, MEXT (19H05647 and 24K01997). MA is funded by Grants-in-Aid for Scientific Research, MEXT (22K06609). MY is funded by Grants-in-Aid for Scientific Research, MEXT (24K01982, 24H01266, 25H01328) and Kobayashi Foundation. We thank the Support Unit for Bio-Material Analysis, RRD, RIKEN for DNA sequence.

The authors declare no competing interest.

