## Supplement text for "Single-molecule behavior and cell-growth regulation in human RTKs"

**Single-particle tracking**

The tracking of single-particle movements and fluorescence intensities was performed using AAS version 2 software (Zido, https://eng.zido.co.jp). This software utilized a scan-based detection and nearest-neighbor linking algorithm. The algorithm is comprised of two stages: (i) raster-scan detection with two-dimensional (2D) Gaussian fitting, (ii) frame-to-frame spot linking with gap closing, and trajectory filtering.

**(i)** Each frame of a movie was sampled on a regular grid with spacing $s$ (scan step) and a square region-of-interest (ROI) of side $r$ pixels. At each grid position, the ROI was fitted with a 2D Gaussian model on a background plane:

$$I(\mathrm{x,y}) = I_{\mathrm{max}} \exp[-(\mathrm{x} - \mathrm{x_g})^2 / (2\sigma_{\mathrm{x}}{}^2) - (\mathrm{y} - \mathrm{y_g})^2 / (2\sigma_{\mathrm{y}}{}^2)] + a(\mathrm{x} - \mathrm{x_g}) + b(\mathrm{y} - \mathrm{y_g}) + I_{\mathrm{back}} \ ,$$

where $I_{\mathrm{max}}$ is the peak intensity, ($\mathrm{x_g}$, $\mathrm{y_g}$) is the sub-pixel center, $\sigma_{\mathrm{x}}$ and $\sigma_{\mathrm{y}}$ are the standard deviations, $a$ and $b$ are the linear background slope coefficients, and $I_{\mathrm{back}}$ is the local background intensity at ($\mathrm{x_g}$, $\mathrm{y_g}$). The parameters were optimized through the implementation of a Levenberg–Marquardt least-squares minimization of the sum of squared residuals. In instances where the estimated peak position was found to be more than the distance $R$ away from the ROI center, the ROI was moved to center the estimated peak, and the estimation was repeated. Spots exhibiting peak intensity below a predetermined threshold $I_{\mathrm{thr}}$ were rejected. Duplicate detections of a single spot resulting from overlapping ROIs were eliminated through a process known as non-maximum suppression, i.e., spots were sorted based on their $I_{\mathrm{max}}$ in descending order, and any spot within the distance $s$ of a higher-ranked spot was discarded.

**(ii)** The detected spots were then linked into trajectories using a frame-by-frame nearest-neighbor algorithm with gap closing. The maximum linking distance $d_{\mathrm{link}}$, was set. A trajectory was permitted to skip up to $G_{\mathrm{max}}$ consecutive frames without an assigned spot (gap closing); if the gap exceeded $G_{\mathrm{max}}$, the tracking was terminated. The resulting gap was filled by linear interpolation of the flanking coordinates. The unmatched spots initiated new trajectories. Trajectories shorter than $N_{\mathrm{min}}$ frames were discarded.

The parameter values used in the present study were $s$ = 3 pix, $r$ = 12 pix, $R$ = 3 pix, $I_{\mathrm{thr}}$ = 200 (This is the raw output from the camera. In text, signal intensity was normalized to the single-molecule intensity.), $d_{\mathrm{link}}$ = 8 pix, $G_{\mathrm{max}}$ = 1 frame, and $N_{\mathrm{min}}$ = 15 frame (14 steps). The pixel size was 67 nm and the frame interval was 40 ms.

**VB-HMM analysis**

The variational Bayesian (VB)-HMM analysis was performed using AAS version 2 (ZIDO). VB-HMM is a standard technique in various fields of science and technology. Basic information about the method can be found in textbooks such as Bishop (2006). The model used to generate single-particle step sizes and the model parameters for the application to single-particle tracking have been previously described (Persson et al. 2013, 2014, Hiroshima et al. 2018, Yanagawa et al. 2018). The objective of the analysis is to ascertain the most probable sequence of the state transitions $\boldsymbol{Z}$ and the parameter values $\boldsymbol{\theta}$, under the observation of the step size trajectories $\boldsymbol{X}$. The procedure of the analysis is as follows:

**(i) Setting initial distributions of the parameters:**

$\boldsymbol{\theta}$ is composed by the initial state $\boldsymbol{\pi}_i$, the state transition matrix $\boldsymbol{A}_{ij}$, and the diffusion rate parameter $\boldsymbol{\beta}_i \equiv 1/(4\boldsymbol{D}_i \Delta t)$. Here, i and j denote the hidden states. $\boldsymbol{D}_i$ is the diffusion coefficient of the state i (i = 1 to $K$). Δt is the step time. In the VB paradigm, these parameters are represented as distributions. The step size between the n-th and n+1-th frames, $x_n$, is generated from the state in the n-th frame $z_n$, according to the distribution $\boldsymbol{\beta}_i$.

The parameters $\boldsymbol{\pi}$ and each row of $\boldsymbol{A}$ were assigned to follow Dirichlet distributions, and each $\boldsymbol{\beta}$ to follow a gamma distribution with the shape $\tilde{a}_i$ and the rate $\tilde{c}_i$. Let $N_i$ and $N_{ij}$ denote the number of the steps classified into state i, and the number of transitions from state i to j, respectively. The Dirichlet hyperparameters for the initial $\boldsymbol{\pi}_i$, $w_{\pi,i}^{(0)}$ and $\boldsymbol{A}_{ij}$, $w_{B,ij}^{(0)}$, and the gamma hyperparameters for the initial $\boldsymbol{\beta}_i$, ($\tilde{a}_i$, $\tilde{c}_i$) were parameterized by the user-settable values $\tilde{w}_\pi$, $\tilde{w}_B$, $k$, $\tilde{a}$, and $\tilde{c}$ in AAS as follows:

$$w_{\pi,i}{}^{(0)} = \tilde{w}_\pi \cdot N_{i,}$$

$$w_{B,ij}{}^{(0)} = \tilde{w}_B \cdot N_{ij} \qquad (i \neq j),$$

$$w_{B,ii}{}^{(0)} = k \cdot \tilde{w}_B \cdot N_{ii,}$$

$$\tilde{a}_i{}^{(0)} = \tilde{a} \cdot N_i, \;\; \tilde{c}_i{}^{(0)} = \tilde{c} \cdot N_{i.}.$$

The diagonal inflation factor $k$, exerts a significant influence on the prior, favoring persistent states and playing a crucial role in suppressing spurious fast-switching solutions. In the present analysis we used $\tilde{w}_\pi = 1$, $\tilde{w}_B = 1$, $k = 10$, $\tilde{a}_i = 1$, and $\tilde{c} = 0.001$ μm$^2$. $\Delta t = 40$ ms.

The numbers of $N_i$ and $N_{ij}$ were estimated by means of the K-means++ clustering method (Arthur and Vassilvitskii, 2007). AAS employs a deterministic variant of K-means++, wherein the conventional algorithm's random seeding is substituted with median and farthest-point selection, ensuring reproducible clustering.

**(ii) VB-E step:**

Under the current (prior) distribution of $\boldsymbol{\theta}$, $\boldsymbol{Z}$ was optimized through the utilization of the forward-backward algorithm incorporating scaling factors (Bishop, 2006).

**(iii) VB-M step:**

According to the optimized $\boldsymbol{Z}$, parameters of the $\boldsymbol{\theta}$ distribution were updated as follows:

$$w_{\pi,i} = w_{\pi,i}{}^{(0)} + \sum_m \gamma_{1,i}^{(m)},$$

$$w_{B,ij} = w_{B,ij}{}^{(0)} + \sum_{m,n} \xi_{n,ij}^{(m)},$$

$$a_i = \tilde{a}_i^{(0)} + \sum_{m,n} \gamma_{n,i}^{(m)},$$
$$c_i = \tilde{c}_i^{(0)} + \frac{1}{4\Delta t} \sum_{m,n} \gamma_{n,i}^{(m)} x_i^2.$$

$\gamma_{n,i} \equiv E[z_{n,i}]$ and $\xi_{n,ij} \equiv E[z_{n-1,i}\, z_{n,j}]$ are the posterior responsibilities obtained in the VB-E step. E[y] denotes the expectation of observing event y. $x_i$ is the average of the step sizes emerged from i state. The expectations $\gamma_{n,i}$ and $\xi_{n,i}$ were aggregated with all trajectories m for parameter update. The lower bound $L_q$ of the log-evidence for observation $\boldsymbol{X}$, $ln\, p(\boldsymbol{X})$, can be derived by utilizing the scaling factors present within the forward-backward algorithm and the expectations of the updated posterior parameter distributions. $L_q$ is the value for the total set of trajectories.

**(iv) Convergence judgment of $L_q$ and parameter estimation:**

The steps (ii) and (iii) were iterated until the difference from the previous $L_q$ became < 0.001% or until the maximum of $I_{max}$ iterations (= 100) was reached. The updated parameters in the VB-M step were employed for the prior distributions in the subsequent VB-E step. Following the final VB-M step, the VB-E step was reiterated, and the ensuing state sequence was reported as the per-frame argmax of $\gamma_{n,i}$. The best parameter values were reported as the expectations of distributions estimated from the optimal $\boldsymbol{Z}$. The best value of $\boldsymbol{D}_i$ is $c_i$ / [4($a_i$ − 1)Δ$t$] for $a_i$ > 1.

**(v) Model selection:**

We performed VB-HMM inferences for the models with the number of states, $K$ = 1 to 5. The models were compared based on the value of $L_q$*, which is the sum of the lower bounds for individual trajectories calculated based on the optimal $\boldsymbol{Z}$ and $\boldsymbol{\theta}$. Typically, $L_q$* is more susceptible to the impact of parameter distributions (due to the variations in $\gamma_{n,i}$ and $\xi_{n,ij}$) than $L_q$ employed in the convergence judgment. Consequently, the possibility of overfitting due to the assumption of a substantial number of states can be mitigated through the usage of $L_q$*. The optimal number of states $K$* was defined as $\text{argmax}_K$ $L_q$*($K$).

**Single-molecule parameters**

We extracted 7 kinds of single-molecule behavioral parameters from the single-particle trajectories either in the resting or response state of RTKs as follows:

**Lateral diffusion coefficient: *D***

The HMM analysis subdivides single-particle trajectories into fragments belonging to each of the three mobility states: immobile, slow, and fast. The mean squared displacement (MSD) was calculated over time intervals for each fragment and averaged for the fragments in each mobility state within a single movie (usually containing one or two cells, mostly one cell) to construct MSD-Δt plots for Δt = 0–0.56 s (14 steps). The plot was fitted by the equation:

$$\text{MSD}(\Delta t) = \frac{L^2}{3}\left(1 - \exp\left(\frac{-12 D_0 \Delta t}{L^2}\right)\right) + \varepsilon^2, \quad (1)$$

to estimate $D$ values (Xiao et al, 2008). Here, ε indicates the offset caused by the random movements of cells and membrane domains, and the measurement noise.

**Fraction of cells without confinement: *E***

When the MSD value at $\Delta t = 0.56$ s (the end of our MSD calculation) in eq. (1) was smaller than 70% of the value expected for simple diffusion with the estimated $D$ value, the cells in the movie were determined to show confinement effects for the particle movements. The fraction of movies without the confinement effect is $E$. This parameter was unstable (Supplement Figure S2) and was not used for further analyses. However, detection of the confinement effect was required to determine $L$ values (see below).

**Molecular density on the cell surface: *F***

The sum of fluorescence intensities from single particles was calculated for each mobility state in single movies. The values were divided by the focused area of the cells (Yanagawa and Sako, 2021) to obtain the values proportional to the density of fluorescent molecules. Note that $F$ is an indication of molecular density, not particle density.

**Single-particle fluorescence intensity: *I***

$I$ is the mean of the fluorescence intensity from single particles in each mobility state. Due to the limited spatial resolution of fluorescence microscopy (~300 nm), single particles contain both protein oligomers firmly attached at specific molecular surfaces and loosely assembled protein molecules, such as those in a single membrane domain. In this manuscript, “molecular clusters” includes both cases. Additionally, due to the imperfect labeling, $I$ relates quantitatively to the overall clustering degree but does not simply relate to the number of molecules in the clusters including unlabeled molecules. The labeling ratio of Halo-tag with SF650 was small (~5%) in our condition (Yanagawa et al, 2018).

**Confinement length: *L***

For cells and mobility states with confinement effects, $L$ in eq. (1) was estimated as the confinement length. The value of $L$ for the fast state ($L3$) was unstable and was not used for further analysis (Supplement Fig. S2).

**Particle fraction of the mobility state: *P***

$P$ is the percentage of particle numbers (not molecular numbers) in each mobility state. Therefore, the $P$ value for one mobility state is determined by the other two. In the multivariate analyses, $P3$ (percentage of fast state particles) was omitted from the explanatory variables.

**State lifetime: *tau***

*tau* represents the lifetime of individual states along the trajectory and can be calculated from the state transition matrix estimated in the HMM. *tau* values are apparent because they are affected by limited trajectory length (Hiroshima et al, 2018). In addition, our HMM did not account for the appearance (exocytosis) and disappearance (endocytosis) of molecules, as it is in the usual single-molecule HMM analysis. However, because we analyzed movements during a short period (0.56 s), this simplification may not be strongly problematic. Although the *tau* values are apparent, we can semi-quantitatively compare the stabilities of the mobility states based on the *tau* values. HMM assumes quasi-steady states, assuming local equilibrium exists between the mobility states. This implies that one of the lifetimes in a cyclic transition is determined by the others. *tau*12 was omitted in the multivariate analyses.

The above parameters were initially determined using a single movie and subsequently averaged across a total of 37-76 movies. For the response states, the parameters are expressed as a relative value compared to those under vehicle stimulation and are indicated as r*D*, r*E*, etc. It is not surprising that some of these parameters are correlated with each other. In fact, we found a high correlation coefficient (R > 0.8) between many of them. Nonetheless, it is not a prerequisite for high correlations to occur. One of the highly correlated parameter pairs was dropped in the multiple correlation analysis and canonical correlation analysis (CCA) to avoid multicollinearity (see below).

**Dependence on the particle density**

If two trajectories intersect, our algorithm assigns the intersection point to the trajectory that was closer in the previous frame, and interrupts tracking of the more distant trajectory. Accurately assessing the impact of this issue is difficult. As an approach, we evaluated the correlation between the parameter values for particle mobility, *D*, *L*, *P* and *tau,* and the particle density (Supplement Table S9). Taking the average and SD of Rs under the 416 measurement conditions (52 RTKs x 2 ligand conditions x 4 time points) and considering parameters with $|R_{ave}|/SD > 1$ to be biased, we found that *P*2 exhibited a positive bias with increasing density, while *tau*12, *tau*13, *tau*31, and *tau*32 exhibited negative biases. The biases in *tau*12 and *tau*32 contributed to the *P*2 bias because their small values mean faster transitions to the slow state. However, $|R_{ave}|$ for the biased parameters ranged from 0.33 to 0.24 (0.26 for *P*2), which were not particularly high.

Significant density dependence was observed for a limited number of RTK species. Of the 132 cases showing |R| > 0.6 among 5,824 (416 x 14 parameters), 72 cases (55%) were concentrated in FLT1 (29 cases), LMR2 (15 cases), MUSK (11 cases), PTK7 (9 cases), and EPHB2 (8 cases). These RTKs indeed had high particle density or large diffusion coefficients (Supplement Table S2). Excluding these five RTKs from Figure 6A resulted in either unchanged or slightly increased $R^2$ values (Supplement Fig. S4).

Based on the above, it cannot be concluded that the density dependence of the measured values, and therefore, the trajectory intersections, significantly impacts the overall results of this study.

**Structural parameters**

We divided the RTK structure into 7 regions and counted the number of amino acids in each region (Table 1). The regions were determined as follows: First, the transmembrane (TM) region was predicted using two programs, SOSUI (https://harrier.nagahama-i-bio.ac.jp/sosui/mobile/) and TMHMM2.0 (https://services.healthtech.dtu.dk/services/TMHMM-2.0). These two programs typically predict partially overlapping, though not identical, regions. One reason for this difference seems to be that they search for a fixed length (24 amino acids) of TM regions based on different criteria. In a few cases, one of the programs predicted no TM region. We began with the union of two predictions and removed either hydrophilic (R, K, N, D, Q, E) or extended hydrophobic (I, L, F, V, A, M, W) amino acids (Eisenberg et al, 1984) from the N- and C-ends to create the final prediction. Next, intrinsically disordered regions at the extracellular (IDRE) and the cytoplasmic (IDRC) sides of the TM region were predicted by a program, PONDR (https://www.pondr.com). We adopted the tyrosine kinase domain (TK) from UniProt (https://www.uniplot.org). Then, the region from the N-terminus to before the IDRE was designated the extracellular region (EC). The region between IDRC and TK was designated the linker region (Linker). The region after TK to the C-terminus was designated the cytoplasmic tail (Tail). For simplification, gaps between IDRs and TM, as well as within IDRs, were incorporated into the IDR regions. The predicted amino acid sequences from IDRE to Linker are listed in Supplement Table S4.

Additionally, we counted the number of several putative functional amino acids and motifs. The positively charged amino acid (R or K) in the IDRE (referred to as “IDRE posi” in this study) is thought to interact with the sialic acid in the ganglioside, GM3, which accumulates in membrane rafts (Kim et al, 2021). The single TM region of RTK is believed to form an $\alpha$-helix in the lipid bilayer. The G/A-x(x)x-G/A motif in the TM region brings two amino acids with small side chains (G and A) together on the same side of the TM $\alpha$−helix facilitating formation of crossed dimers. Such a structure has been observed in the EGFR TM-JM peptide dimer (Arkhipov et al, 2013) as well as in artificial $\alpha$-helices in lipid membranes (Yano et al, 2015). The CRAC and CARC (reverse CRAC) motifs, L/V-x(1-5)-Y/F-x(1-5)-R/K, between the TM and IDR regions are the putative cholesterol-binding sites (Fantini et al, 2016). The inner leaflet of the cell membrane contains acidic lipids, such as phosphatidylinositol bisphosphates ($PIP_2$) and phosphatidylserine (PS). Positively charged amino acids (R, K) in the flexible IDRC may interact with these acidic lipids (Hedger et al, 2014) as has been reported in EGFR (Jura et al, 2009; Matsushita et al, 2013). $PIP_2$ regulates EGFR activity (Abe et al, 2024).

**Evolutionary parameters**

The amino acid sequences of RTKs vary greatly between subfamilies, except in the TK region. For this reason, the evolutionary tree of RTKs has been constructed based on the TK sequences (Robinson et al., 2000). Their tree does not include STYK1, but it has recently been updated as shown on the Cell Signaling Technology website (www.cellsignal.com). We adopted this new version to categorize RTKs into 5 groups as follows.

EG1: ERBB (EGFR) family
EG2: EPH family
EG3: ALK, ROS (SEV), InsR, AXL, and MET families (RYK belongs to this group but was not included in this study.)
EG4: FGFR, RET, VEGFR, PDGFR, and TIE families
EG5: PTK7 (CCK4), DDR, TRK, MUSK, ROR, LMR, and STYK1 (SuRTK106) families

This grouping is rough, but more precise grouping may be too excessive, considering the variations and measurement noise in single-molecule parameters. For the multivariate analyses, the evolutionary grouping was converted into 5-component dummy parameters: (1, 0, 0, 0, 0) for EG1, (0, 1, 0, 0, 0) for EG2, and so on. Note that, because the degree of freedom in this grouping is four, 4-dimensional dummy parameters of EG1-EG4, dropping EG5, were used in multiple regression modeling.

**Parameter selection for multivariate analyses**

In the CCA and multiple linear regression modeling, the number of explanatory variables must be smaller than the data size. Otherwise, some variables must be dropped. We dropped one of the variables in the most highly correlated pair one after another because using a set of highly correlated parameters causes instability in the models (multicollinearity problem), looking at the correlation matrix of the explanatory variables. In the linear regression, explainability ($R^2$) can be increased at will by increasing the number of explanatory variables. Therefore, statistical significance, rather than the explainability, should be the criterion for obtaining the final model. In general, as the number of variables decreases, the statistical significance of the models increases, but the models become less explainable. This is because statistical significance pays a penalty to increase the number of variables. However, models with a small number of explanatory variables are not informative.

In multiple linear regression modeling, we sought the most powerful set of variables within a range of more than four variables and less than 0.7 of the maximum R between the variables (Figs. 3A, C and 4A). We evaluated the power of the variable sets as the number of statistically significant models that the sets constructed. This criterion is based on the expectation that the molecular functions are similar across the cell lines. Model significance was evaluated using the F-test. As the result, we chose 4 to 12 parameters for the multiple regression models, which is not particularly large for a dataset of 52 RTKs. In the response behavior models for function, which contains 12 parameters, the maximum R was 0.617 between r*I*1(1) and r*I*1(3) (Supplement Table S6A). Further reduction of the number of parameters in this case caused the number of significant models to decrease sharply (Fig. 3C), indicating that these two parameters carried distinct information and that multicollinearity was not an issue overall. Examining individual models revealed very few cases where pR < -1 (6/1095 cases from response behavior to function; the minimum pR = -1.088; Table 2B), suggesting multicollinearity. However, since we are discussing the overall trend rather than the details of individual models, no special treatment was given.

Unfortunately, to the best of our knowledge, there is no standard method for evaluating the significance of CCA models or the size of CR. CCA assumes multicollinearity from the outset and therefore does not typically address this issue. However, if two parameters have the same meaning, the same issue is weighted twice. To be on the safe side, we reduced the number of parameters so that no parameter pair had an R value greater than 0.7. We then discussed only the largest component, CC1.

Remember that the parameters omitted in the above procedure likely have a comparable role to the explanatory parameters in the final sets in both the multiple regression models and CCA.

**Supplement Figures**

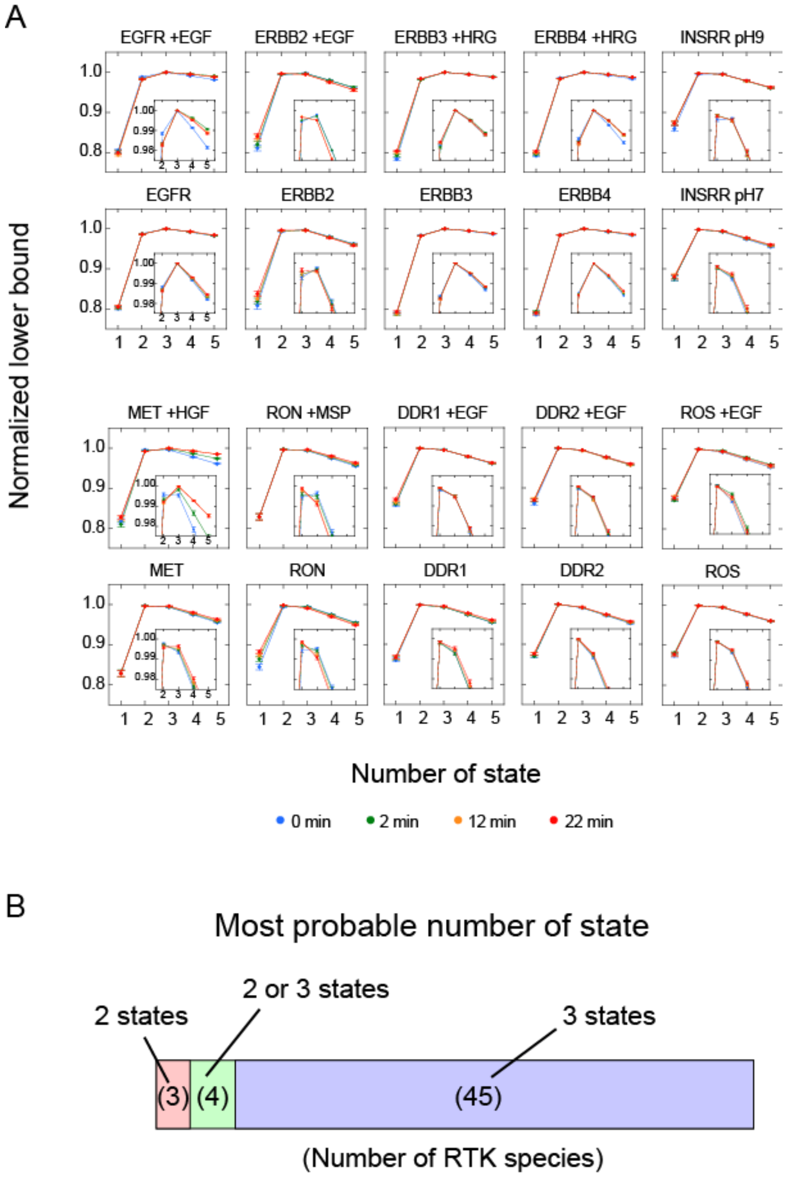


**Supplement Figure S1. HMM analysis**

**A.** Representative results of number of state (NoS) determination. See Supplement Table S1 for all results. The lower bound values for each trajectory were normalized to the maximum within each movie, then averaged across movies within the same condition. Error bars show SE among movies.

**B.** Distribution of the most probable NoS.

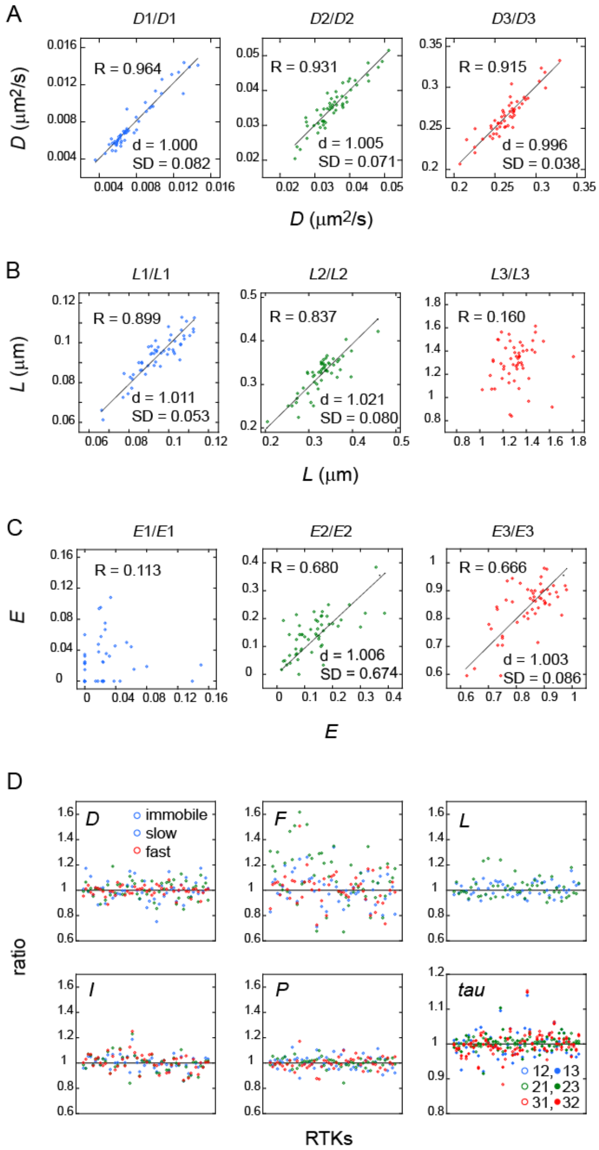


**Supplement Figure S2. Reproducibility of single-molecule measurements**

**A-C**. Comparisons of *D* (**A**), *L* (**B**), and *E* (**C**) values in two independent measurements. For each RTK species, the parameters in the first (x axis) and the second (y axis) trials were plotted for 52 RTKs. Lines indicate fitting to the liner function, $y = d * x$. R: correlation coefficient between the two trials. d: slope of the fitting line, SD: standard deviation of the ratios between two measurements. *L*3 and *E*1-3 were less reproducible and not used for further analysis.

**D.** Ratios of parameter values from two trials. Only parameters used in further analysis are plotted. Plots for *D* and *L* are replots from (**A**) and (**B**), respectively. RTKs are aligned as shown in Table 1 from left to right. For *D, F, I, L, P*, blue: immobile state, green: slow mobile state, red: fast state. For *tau*, open blue: immobile to slow, filled blue: immobile to fast, open green: slow to immobile, filled green: slow to fast, open red: fast to immobile, filled red: fast to slow. See Supplement Table S3 for details.

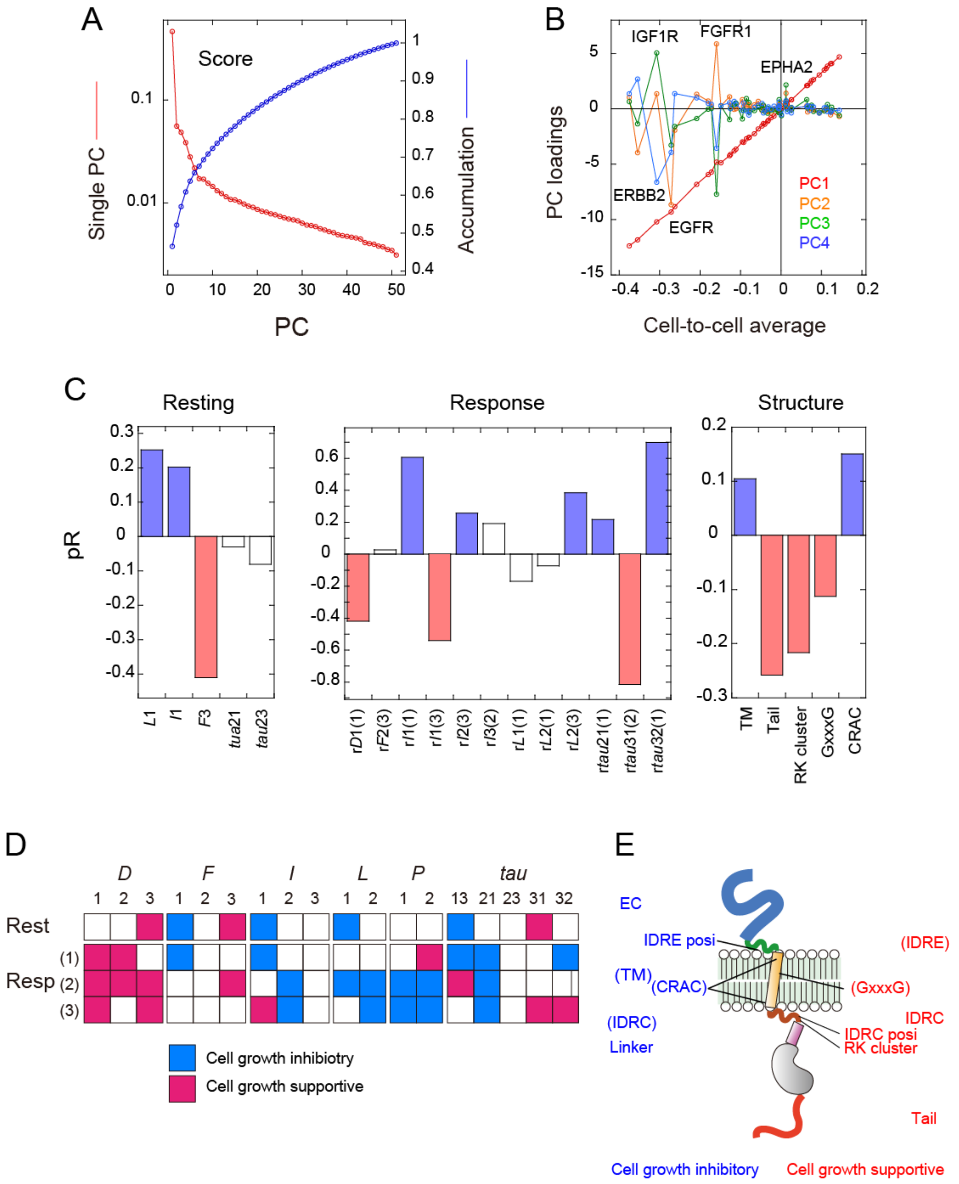

**Supplement Figure S3. Models for generalized RTK function**

**A.** Single PC scores (red) of the CRISPR factors and their accumulation (blue).

**B.** PC loadings of 52 RTKs plotted as the function of their average CRISPER factors over cell lines. RTKs with significant deviation in PC2-4 are indicated.

**C.** pR values in the resting state, response state, and structure models for CRISPR factor PC1. The explanatory variables in the final models for single cell lines (Figs. 3 and 4) were used.

**D.** Diagram of significant behavioral parameters (|pR| > 0.2) and parameters correlated with them but dropped in the parameter selection.

**E.** Structural parameters significantly contributed to explain RTK functions are mapped with the parameters correlated with them but dropped during the parameter selections. |pR|s without and with the parentheses were > 0.2 and > 0.1, respectively.

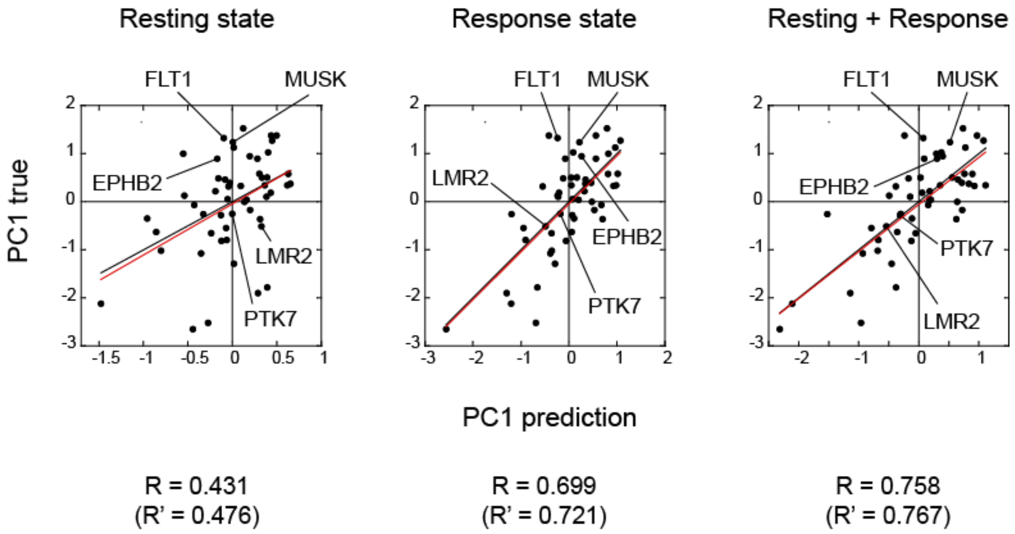


**Supplement Figure S4. Effects of the dependence of parameter values on particle density**

The positions of the RTKs with intense dependence of the mobility parameters on the particle density are indicated on the plots shown in Figure 6A. The red lines and R' show the correlation between the true and predicted values after the removal of the five indicated RTKs.

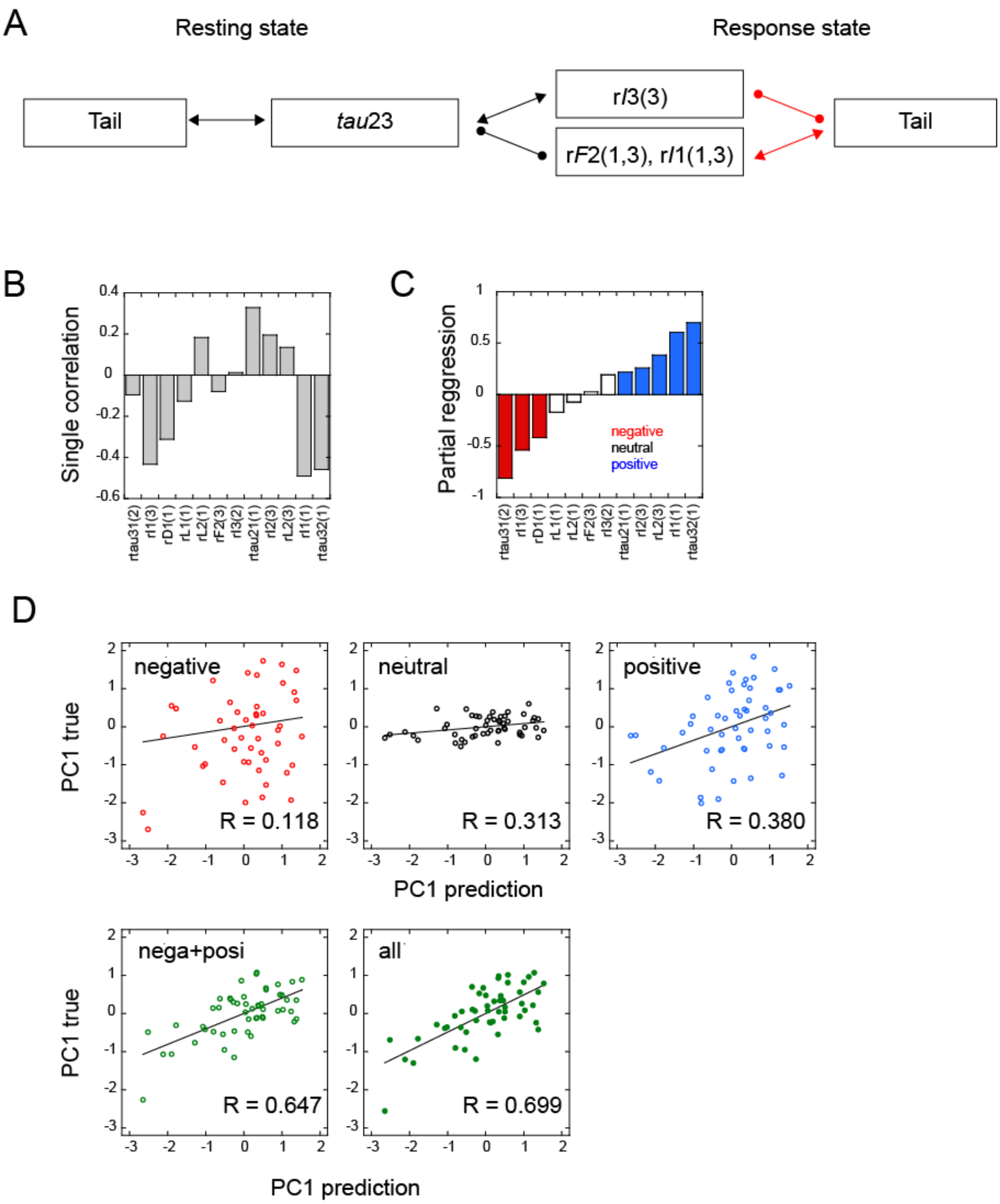


**Supplement Figure S5. Roles of parameters in the multiple regression models**

**A.** Parameter linkage involved in the exclusive resting state and structure models for RTK function (Fig. 4D). Roles of Tail changed after cell stimulation (red arrows).

**B, C.** Values of R (**B**) and pR (**C**) between the response state parameters and the CRISPR factor PC1. In (**C**), negative (pR < -0.2), neutral (-0.1 < pR < 0.2), and positive (pR > 0.2) parameters are colored differently.

**D.** Explanation of CRISPR PC1 values using indicated response state parameters. Multiple parameters explain PC1 values in combination.

**Legends for Supplement Tables and Movie**

**Supplement Table S1. HMM analysis**

Values of lower bound were normalized to the maximum for each movie and averaged across all movies in the same conditions. n: number of examined movies.

**Supplement Table S2. Single-molecule behavioral parameters**

**Supplement Table S3. Reproducibility in single-molecule measurements**

**Supplement Table S4. CRISPER factors from DepMap**

Average for individual RTKs in all cell lines are shown at the end.

**Supplement Table S5. Correlation matrices between parameters in different categories**

Correlation matrix between behavioral and functional parameters (**A**), structural and functional parameters (**B**), evolutionary and functional parameters (**C**), structural and behavioral parameters (**D**), evolutionary and behavioral parameters (**E**), and evolutionary and structural parameters (**F**).

**Supplement Table S6. Correlation matrices within the behavioral parameters**

Correlation metrices within the behavioral (**A**) and structural (**B**) parameters.

**Supplement Table S7. RTK sequences around the TM region**

The amino acid sequence from the estimated IDRE to Lineker are shown. Cerulean: IDR regions, Yellow: TM region. R and K are colored in red. GxxxG motifs in the TM region are underlined. Multiple motifs sometime overlap. Blue colored Y/Fs are at the middle of CRAC or CARC motives. Amino acids in boxes are included in the TK domain but considered to be included in IDRC.

**Supplement Table S8. Multiple regression models between structure and evolution to function**

**A. B.** Multiple correlations from structural parameters (**A**) and evolutionary parameters (**B**) to the CRISPR factors in individual cell lines. pR values in the final models for 1,095 cell lines are listed according to the p-values in F-test. determ: determination coefficient. Averages of the pRs and determination coefficients for the models with $p < 0.01$, 0.05, 0.1 and for all models are shown at the end of the lists. $pR > 0.1$ and $< -0.1$ are indicated in blue and red, respectively. **C.** Multiple correlations from evolutionary parameters to individual structural parameters. pR values in the models are listed according to the p-values in F-test. **A-C.** p-values in t-test for each pR are also shown. $p < 0.05$ is colored yellow.

**Supplement Table S9. Dependence on the particle density of the mobility parameters**

The correlation coefficient (R) was calculated between the indicated parameter values and particle densities in 37-73 single movies under the same conditions. The average ($R_{ave}$) and standard deviation (SD) of Rs across 416 conditions are listed.

**Supplement Movie 1.** Representative movies for EGFR-FLT3 (**A**), FLT1-ROR1 (**B**), ROR2-EPHA4 (**C**), and EPHA6-SYTK1 (**D**) in the resting and response states. Original movies in a 512 x 512 pixel format for each imaging field with a 16-bit depth used for parameter extraction are compressed to a 256 x 256 pixel format with an 8-bit depth for convenience. Times after cell stimulation are indicated in the upper left corner. Bar: 20 μm.
