## Supplement Table S1 for "Single-molecule behavior and cell-growth regulation in human RTKs"

ERBB

| EGFR | EGF | | | | | | | | |
|---|---|---|---|---|---|---|---|---|---|
| | 0min | | 2 min | | 12 min | | 22 min | | |
| NoS | lower bou | SE | lower bou | SE | lower bou | SE | lower bou | SE | n |
| 1 | 0.80359 | 0.00456 | 0.80089 | 0.00346 | 0.79422 | 0.00306 | 0.79962 | 0.00374 | 63 |
| 2 | 0.98833 | 0.00081 | 0.98305 | 0.00065 | 0.98292 | 0.00053 | 0.98287 | 0.00069 | |
| 3 | 0.99977 | 0.00017 | 1 | 5.2E-07 | 1 | 0 | 0.99996 | 4.1E-05 | |
| 4 | 0.99135 | 0.00041 | 0.9962 | 0.00021 | 0.99552 | 0.00023 | 0.99516 | 0.00028 | |
| 5 | 0.98147 | 0.00064 | 0.99062 | 0.00036 | 0.98897 | 0.00041 | 0.98846 | 0.00045 | |
| | vehicle | | | | | | | | |
| NoS | | | | | | | | | |
| 1 | 0.80141 | 0.00403 | 0.80211 | 0.00396 | 0.80489 | 0.00428 | 0.80487 | 0.00316 | 67 |
| 2 | 0.98798 | 0.00066 | 0.98676 | 0.00064 | 0.98665 | 0.00071 | 0.98635 | 0.0006 | |
| 3 | 0.99997 | 2.1E-05 | 0.99989 | 7.6E-05 | 1 | 0 | 1 | 0 | |
| 4 | 0.9916 | 0.00025 | 0.99239 | 0.00026 | 0.99257 | 0.00026 | 0.99306 | 0.00024 | |
| 5 | 0.98187 | 0.00046 | 0.98324 | 0.00044 | 0.9835 | 0.00048 | 0.98442 | 0.00044 | |

| ERBB2 | EGF | | | | | | | | |
|---|---|---|---|---|---|---|---|---|---|
| NoS | lower bou | SE | lower bou | SE | lower bou | SE | lower bou | SE | n |
| 1 | 0.81128 | 0.00756 | 0.82013 | 0.00653 | 0.83627 | 0.00686 | 0.8403 | 0.00611 | 45 |
| 2 | 0.99422 | 0.00095 | 0.99493 | 0.00095 | 0.99695 | 0.00083 | 0.99659 | 0.00076 | |
| 3 | 0.99781 | 0.00068 | 0.99712 | 0.00078 | 0.99501 | 0.0012 | 0.99505 | 0.0013 | |
| 4 | 0.97998 | 0.00141 | 0.98012 | 0.00156 | 0.97601 | 0.00211 | 0.97568 | 0.00224 | |
| 5 | 0.961 | 0.00218 | 0.96222 | 0.00233 | 0.95664 | 0.00296 | 0.95584 | 0.00311 | |
| | vehicle | | | | | | | | |
| NoS | | | | | | | | | |
| 1 | 0.80813 | 0.00854 | 0.81922 | 0.00726 | 0.8253 | 0.00786 | 0.83786 | 0.00708 | 42 |
| 2 | 0.99309 | 0.00133 | 0.99455 | 0.00112 | 0.99482 | 0.00133 | 0.99664 | 0.00091 | |
| 3 | 0.99737 | 0.00066 | 0.99689 | 0.00085 | 0.99615 | 0.00081 | 0.99632 | 0.00084 | |
| 4 | 0.9805 | 0.00139 | 0.97967 | 0.00156 | 0.97832 | 0.00156 | 0.97782 | 0.00145 | |
| 5 | 0.9624 | 0.00205 | 0.96135 | 0.00223 | 0.95968 | 0.00224 | 0.95857 | 0.00205 | |

| ERBB3 | HRG | | | | | | | | |
|---|---|---|---|---|---|---|---|---|---|
| NoS | lower bou | SE | lower bou | SE | lower bou | SE | lower bou | SE | n |
| 1 | 0.78348 | 0.00301 | 0.79325 | 0.00299 | 0.79952 | 0.00245 | 0.80347 | 0.00241 | 56 |
| 2 | 0.98289 | 0.00059 | 0.98169 | 0.00067 | 0.98366 | 0.00047 | 0.98406 | 0.00052 | |
| 3 | 1 | 0 | 1 | 0 | 1 | 0 | 1 | 0 | |
| 4 | 0.99461 | 0.00022 | 0.99536 | 0.00018 | 0.99463 | 0.00019 | 0.99498 | 0.00016 | |
| 5 | 0.98734 | 0.00039 | 0.98873 | 0.0003 | 0.98743 | 0.00035 | 0.98814 | 0.00028 | |
| | vehicle | | | | | | | | |
| NoS | | | | | | | | | |
| 1 | 0.78903 | 0.00338 | 0.78741 | 0.00331 | 0.78802 | 0.00302 | 0.79228 | 0.00335 | 54 |
| 2 | 0.98344 | 0.00065 | 0.98284 | 0.00061 | 0.98211 | 0.00051 | 0.98259 | 0.0006 | |
| 3 | 1 | 0 | 1 | 0 | 1 | 0 | 1 | 0 | |
| 4 | 0.99433 | 0.0002 | 0.99501 | 0.00018 | 0.99523 | 0.00014 | 0.99511 | 0.00017 | |
| 5 | 0.98685 | 0.00034 | 0.98806 | 0.00031 | 0.98846 | 0.00027 | 0.98837 | 0.0003 | |

| ERBB4 | HRG | | | | | | | | |
|---|---|---|---|---|---|---|---|---|---|
| NoS | lower bou | SE | lower bou | SE | lower bou | SE | lower bou | SE | n |
| 1 | 0.792 | 0.00384 | 0.79546 | 0.00345 | 0.7989 | 0.00308 | 0.80182 | 0.00277 | 43 |
| 2 | 0.98564 | 0.00066 | 0.98414 | 0.00058 | 0.98339 | 0.0006 | 0.98388 | 0.00052 | |
| 3 | 1 | 0 | 1 | 0 | 1 | 0 | 1 | 0 | |
| 4 | 0.99277 | 0.00032 | 0.9948 | 0.00023 | 0.99468 | 0.00024 | 0.99486 | 0.00021 | |
| 5 | 0.98382 | 0.00058 | 0.98769 | 0.00041 | 0.98751 | 0.00045 | 0.98795 | 0.00039 | |
| | vehicle | | | | | | | | |
| NoS | | | | | | | | | |
| 1 | 0.78556 | 0.00339 | 0.79253 | 0.00362 | 0.78873 | 0.00377 | 0.7902 | 0.00345 | 41 |
| 2 | 0.98504 | 0.00069 | 0.98467 | 0.00082 | 0.9844 | 0.00063 | 0.98409 | 0.0006 | |
| 3 | 1 | 0 | 1 | 0 | 1 | 0 | 1 | 0 | |
| 4 | 0.99286 | 0.00034 | 0.99335 | 0.00028 | 0.99368 | 0.00024 | 0.99408 | 0.00025 | |
| 5 | 0.98397 | 0.00063 | 0.98488 | 0.00052 | 0.98549 | 0.00044 | 0.98622 | 0.00047 | |

INSR

| INSR | INS | | | | | | | | |
|---|---|---|---|---|---|---|---|---|---|
| | 0min | | 2 min | | 12 min | | 22 min | | |
| NoS | lower bou | SE | lower bou | SE | lower bou | SE | lower bou | SE | n |
| 1 | 0.83769 | 0.00491 | 0.83501 | 0.00476 | 0.82633 | 0.00472 | 0.83041 | 0.00443 | 45 |
| 2 | 0.99046 | 0.00084 | 0.9882 | 0.00073 | 0.98772 | 0.0008 | 0.98734 | 0.00068 | |
| 3 | 0.99999 | 6.7E-06 | 1 | 0 | 0.99992 | 7.6E-05 | 1 | 0 | |
| 4 | 0.9908 | 0.00034 | 0.99229 | 0.0003 | 0.99272 | 0.00038 | 0.99304 | 0.00034 | |
| 5 | 0.98027 | 0.00061 | 0.98307 | 0.00054 | 0.98385 | 0.00065 | 0.9845 | 0.00064 | |
| | vehicle | | | | | | | | |
| NoS | | | | | | | | | |
| 1 | 0.8411 | 0.00422 | 0.84329 | 0.0035 | 0.8401 | 0.00319 | 0.83957 | 0.00304 | 42 |
| 2 | 0.99078 | 0.00078 | 0.99025 | 0.00065 | 0.98918 | 0.00071 | 0.98929 | 0.00058 | |
| 3 | 1 | 0 | 1 | 0 | 1 | 0 | 1 | 0 | |
| 4 | 0.99055 | 0.00037 | 0.99143 | 0.00031 | 0.99221 | 0.00031 | 0.99256 | 0.00027 | |
| 5 | 0.97981 | 0.00068 | 0.98159 | 0.00058 | 0.98306 | 0.00055 | 0.98378 | 0.0005 | |

| IGF1R | IGF | | | | | | | | |
|---|---|---|---|---|---|---|---|---|---|
| NoS | lower bou | SE | lower bou | SE | lower bou | SE | lower bou | SE | n |
| 1 | 0.85577 | 0.00528 | 0.851 | 0.00451 | 0.85372 | 0.00442 | 0.85213 | 0.00506 | 44 |
| 2 | 0.99637 | 0.0009 | 0.99559 | 0.0008 | 0.99552 | 0.00079 | 0.99473 | 0.00085 | |
| 3 | 0.99681 | 0.00065 | 0.99817 | 0.00044 | 0.99808 | 0.00053 | 0.99841 | 0.00053 | |
| 4 | 0.98056 | 0.00131 | 0.98345 | 0.00099 | 0.98344 | 0.00112 | 0.98447 | 0.00109 | |
| 5 | 0.96379 | 0.00196 | 0.968 | 0.00156 | 0.96797 | 0.00172 | 0.96981 | 0.00165 | |
| | vehicle | | | | | | | | |
| NoS | | | | | | | | | |
| 1 | 0.84937 | 0.00712 | 0.85454 | 0.00661 | 0.85648 | 0.00592 | 0.85869 | 0.0064 | 44 |
| 2 | 0.9961 | 0.00076 | 0.99595 | 0.00086 | 0.99551 | 0.00094 | 0.99574 | 0.00086 | |
| 3 | 0.997 | 0.00077 | 0.99706 | 0.00069 | 0.99705 | 0.00073 | 0.99751 | 0.00068 | |
| 4 | 0.98091 | 0.00137 | 0.98196 | 0.00128 | 0.9821 | 0.00134 | 0.98285 | 0.00122 | |
| 5 | 0.96395 | 0.00192 | 0.96598 | 0.00181 | 0.96652 | 0.00187 | 0.96753 | 0.00171 | |

| INSRR | pH9 (NaOH) | | | | | | | | |
|---|---|---|---|---|---|---|---|---|---|
| NoS | lower bou | SE | lower bou | SE | lower bou | SE | lower bou | SE | n |
| 1 | 0.85927 | 0.00692 | 0.8727 | 0.00492 | 0.87305 | 0.00505 | 0.87292 | 0.00493 | 57 |
| 2 | 0.99555 | 0.00103 | 0.99753 | 0.00054 | 0.99745 | 0.00055 | 0.99715 | 0.0007 | |
| 3 | 0.99563 | 0.00079 | 0.99513 | 0.00079 | 0.99477 | 0.0008 | 0.99528 | 0.00075 | |
| 4 | 0.97873 | 0.00146 | 0.97822 | 0.00142 | 0.97842 | 0.00143 | 0.97901 | 0.00133 | |
| 5 | 0.96115 | 0.00207 | 0.9609 | 0.002 | 0.96172 | 0.00201 | 0.96243 | 0.0019 | |
| | vehicle | | | | | | | | |
| NoS | | | | | | | | | |
| 1 | 0.87679 | 0.00671 | 0.87746 | 0.00666 | 0.88193 | 0.00549 | 0.88007 | 0.00525 | 40 |
| 2 | 0.998 | 0.00073 | 0.99775 | 0.00066 | 0.99806 | 0.00067 | 0.99826 | 0.00057 | |
| 3 | 0.99249 | 0.00095 | 0.99341 | 0.00097 | 0.99387 | 0.00092 | 0.99447 | 0.0009 | |
| 4 | 0.97384 | 0.00164 | 0.97576 | 0.00172 | 0.97611 | 0.0016 | 0.97739 | 0.00146 | |
| 5 | 0.95503 | 0.00229 | 0.95781 | 0.0024 | 0.95813 | 0.00224 | 0.9602 | 0.00201 | |

PDGFR

| PDGFRA | PDGF AA | | | | | | | | |
|---|---|---|---|---|---|---|---|---|---|
| | 0min | | 2 min | | 12 min | | 22 min | | |
| NoS | lower bou | SE | lower bou | SE | lower bou | SE | lower bou | SE | n |
| 1 | 0.78227 | 0.00451 | 0.79122 | 0.00418 | 0.78765 | 0.00394 | 0.79185 | 0.00421 | 50 |
| 2 | 0.98648 | 0.0007 | 0.98612 | 0.00065 | 0.98318 | 0.00065 | 0.98314 | 0.00078 | |
| 3 | 1 | 0 | 1 | 0 | 1 | 0 | 1 | 0 | |
| 4 | 0.99112 | 0.0003 | 0.99156 | 0.00027 | 0.99429 | 0.00023 | 0.99419 | 0.00027 | |
| 5 | 0.98061 | 0.00055 | 0.98132 | 0.00048 | 0.98657 | 0.00044 | 0.9864 | 0.00049 | |
| | vehicle | | | | | | | | |
| NoS | | | | | | | | | |
| 1 | 0.7841 | 0.00318 | 0.79451 | 0.0033 | 0.79254 | 0.00376 | 0.79791 | 0.00399 | 53 |
| 2 | 0.98625 | 0.00055 | 0.98611 | 0.00062 | 0.98515 | 0.00058 | 0.98572 | 0.00075 | |
| 3 | 1 | 0 | 1 | 0 | 1 | 0 | 1 | 0 | |
| 4 | 0.99091 | 0.00035 | 0.9914 | 0.00031 | 0.99166 | 0.00029 | 0.99198 | 0.0003 | |
| 5 | 0.98019 | 0.00065 | 0.98108 | 0.00057 | 0.98154 | 0.00054 | 0.98217 | 0.00056 | |

| PDGFRB | PDGF BB | | | | | | | | |
|---|---|---|---|---|---|---|---|---|---|
| NoS | lower bou | SE | lower bou | SE | lower bou | SE | lower bou | SE | n |
| 1 | 0.81351 | 0.00474 | 0.82342 | 0.00358 | 0.8092 | 0.00389 | 0.80818 | 0.00429 | 45 |
| 2 | 0.9892 | 0.00092 | 0.9866 | 0.00076 | 0.98315 | 0.00051 | 0.98287 | 0.00061 | |
| 3 | 0.99999 | 1E-05 | 0.99999 | 8.2E-06 | 1 | 0 | 1 | 0 | |
| 4 | 0.98929 | 0.0004 | 0.99293 | 0.00042 | 0.99563 | 0.00025 | 0.99609 | 0.00015 | |
| 5 | 0.97722 | 0.00074 | 0.98413 | 0.00078 | 0.98935 | 0.00041 | 0.99002 | 0.00026 | |
| | vehicle | | | | | | | | |
| NoS | | | | | | | | | |
| 1 | 0.81214 | 0.00467 | 0.81785 | 0.0041 | 0.82242 | 0.00385 | 0.8273 | 0.00456 | 39 |
| 2 | 0.9911 | 0.00078 | 0.98972 | 0.00081 | 0.98929 | 0.00089 | 0.99048 | 0.00076 | |
| 3 | 0.99988 | 9.6E-05 | 0.99994 | 5.4E-05 | 1 | 2.8E-06 | 1 | 0 | |
| 4 | 0.98956 | 0.00048 | 0.99011 | 0.00039 | 0.99059 | 0.00043 | 0.99119 | 0.0004 | |
| 5 | 0.97787 | 0.00082 | 0.9789 | 0.00071 | 0.97975 | 0.00078 | 0.98106 | 0.00074 | |

| KIT | SCF | | | | | | | | |
|---|---|---|---|---|---|---|---|---|---|
| NoS | lower bou | SE | lower bou | SE | lower bou | SE | lower bou | SE | n |
| 1 | 0.76689 | 0.00853 | 0.7821 | 0.00837 | 0.78393 | 0.00773 | 0.79304 | 0.00811 | 41 |
| 2 | 0.99515 | 0.00077 | 0.99512 | 0.00082 | 0.99498 | 0.00095 | 0.99492 | 0.00106 | |
| 3 | 0.9969 | 0.00088 | 0.99741 | 0.00078 | 0.99621 | 0.00098 | 0.99698 | 0.0009 | |
| 4 | 0.98055 | 0.00171 | 0.98092 | 0.00153 | 0.97917 | 0.00166 | 0.9803 | 0.00166 | |
| 5 | 0.96386 | 0.00251 | 0.96392 | 0.00226 | 0.96203 | 0.00234 | 0.96319 | 0.00237 | |
| | vehicle | | | | | | | | |
| NoS | | | | | | | | | |
| 1 | 0.77366 | 0.0085 | 0.78176 | 0.00776 | 0.78534 | 0.00726 | 0.78486 | 0.00645 | 42 |
| 2 | 0.99468 | 0.00106 | 0.99471 | 0.00101 | 0.99532 | 0.00102 | 0.99453 | 0.00091 | |
| 3 | 0.99709 | 0.0008 | 0.99749 | 0.00066 | 0.99681 | 0.00084 | 0.9981 | 0.00064 | |
| 4 | 0.97992 | 0.00144 | 0.98074 | 0.00127 | 0.98001 | 0.00149 | 0.98231 | 0.00127 | |
| 5 | 0.96205 | 0.00208 | 0.96359 | 0.00187 | 0.96269 | 0.00211 | 0.96583 | 0.00188 | |

| CSF1R | CSF1 | | | | | | | | |
|---|---|---|---|---|---|---|---|---|---|
| NoS | lower bou | SE | lower bou | SE | lower bou | SE | lower bou | SE | n |
| 1 | 0.7839 | 0.00363 | 0.79898 | 0.00324 | 0.80312 | 0.00397 | 0.79863 | 0.00467 | 43 |
| 2 | 0.98591 | 0.00057 | 0.98273 | 0.00051 | 0.98331 | 0.00057 | 0.98256 | 0.00088 | |
| 3 | 1 | 0 | 1 | 0 | 1 | 0 | 1 | 0 | |
| 4 | 0.99222 | 0.00028 | 0.996 | 0.0002 | 0.99595 | 0.00023 | 0.99559 | 0.00023 | |
| 5 | 0.98269 | 0.00054 | 0.98989 | 0.00038 | 0.98981 | 0.0004 | 0.98923 | 0.0004 | |
| | vehicle | | | | | | | | |
| NoS | | | | | | | | | |
| 1 | 0.77352 | 0.00492 | 0.77593 | 0.00409 | 0.78165 | 0.00452 | 0.78258 | 0.00474 | 40 |
| 2 | 0.98556 | 0.00089 | 0.9826 | 0.00077 | 0.98355 | 0.00072 | 0.98422 | 0.00074 | |
| 3 | 1 | 0 | 1 | 0 | 1 | 0 | 1 | 0 | |
| 4 | 0.99158 | 0.00039 | 0.99265 | 0.00033 | 0.99291 | 0.0003 | 0.99288 | 0.00033 | |
| 5 | 0.98125 | 0.0007 | 0.9832 | 0.00059 | 0.98383 | 0.00054 | 0.98365 | 0.00058 | |

| FLT3 | FLT3 | | | | | | | | |
|---|---|---|---|---|---|---|---|---|---|
| NoS | lower bou | SE | lower bou | SE | lower bou | SE | lower bou | SE | n |
| 1 | 0.80538 | 0.00315 | 0.81441 | 0.00346 | 0.81899 | 0.00334 | 0.83122 | 0.0035 | 41 |
| 2 | 0.98493 | 0.00064 | 0.98464 | 0.00072 | 0.98336 | 0.00062 | 0.9848 | 0.00059 | |
| 3 | 1 | 0 | 1 | 0 | 1 | 0 | 1 | 0 | |
| 4 | 0.99342 | 0.0003 | 0.99443 | 0.00028 | 0.99589 | 0.0002 | 0.99558 | 0.00021 | |
| 5 | 0.98493 | 0.00052 | 0.98693 | 0.00046 | 0.98977 | 0.00032 | 0.9894 | 0.00038 | |
| | vehicle | | | | | | | | |
| NoS | | | | | | | | | |
| 1 | 0.81317 | 0.00312 | 0.81353 | 0.00357 | 0.81568 | 0.00409 | 0.81824 | 0.00351 | 41 |
| 2 | 0.98553 | 0.00059 | 0.98522 | 0.00068 | 0.98556 | 0.00071 | 0.98515 | 0.00064 | |
| 3 | 1 | 0 | 0.99996 | 4E-05 | 1 | 0 | 1 | 0 | |
| 4 | 0.9931 | 0.00028 | 0.99378 | 0.00024 | 0.99399 | 0.00023 | 0.99397 | 0.00023 | |
| 5 | 0.98454 | 0.00051 | 0.98582 | 0.00042 | 0.98629 | 0.00044 | 0.98624 | 0.00044 | |

VEGFR

| FLT1 | VEGF A165 | | | | | | | | |
|---|---|---|---|---|---|---|---|---|---|
| | 0min | | 2 min | | 12 min | | 22 min | | |
| NoS | lower bou | SE | lower bou | SE | lower bou | SE | lower bou | SE | n |
| 1 | 0.7456 | 0.00568 | 0.7591 | 0.00646 | 0.76161 | 0.00631 | 0.77177 | 0.00667 | 41 |
| 2 | 0.98179 | 0.00082 | 0.98219 | 0.00074 | 0.98108 | 0.00073 | 0.982 | 0.00079 | |
| 3 | 1 | 0 | 1 | 0 | 1 | 0 | 0.99996 | 3.8E-05 | |
| 4 | 0.99535 | 0.00026 | 0.99555 | 0.00029 | 0.99559 | 0.0003 | 0.99515 | 0.00029 | |
| 5 | 0.98787 | 0.00043 | 0.98778 | 0.00046 | 0.98783 | 0.00044 | 0.9874 | 0.00043 | |
| | vehicle | | | | | | | | |
| NoS | | | | | | | | | |
| 1 | 0.74106 | 0.0069 | 0.73706 | 0.00619 | 0.74196 | 0.00735 | 0.74126 | 0.00613 | 48 |
| 2 | 0.98265 | 0.00095 | 0.98355 | 0.00081 | 0.98129 | 0.00083 | 0.9808 | 0.00077 | |
| 3 | 1 | 0 | 0.99995 | 5.3E-05 | 0.99999 | 6E-06 | 1 | 0 | |
| 4 | 0.9958 | 0.00022 | 0.99539 | 0.00032 | 0.99618 | 0.00026 | 0.99587 | 0.00024 | |
| 5 | 0.98914 | 0.00039 | 0.98801 | 0.00048 | 0.98888 | 0.0004 | 0.98842 | 0.00038 | |

| KDR | VEGF A165 | | | | | | | | |
|---|---|---|---|---|---|---|---|---|---|
| NoS | lower bou | SE | lower bou | SE | lower bou | SE | lower bou | SE | n |
| 1 | 0.81153 | 0.0046 | 0.82753 | 0.00448 | 0.84751 | 0.00378 | 0.84923 | 0.00391 | 73 |
| 2 | 0.98912 | 0.00059 | 0.98843 | 0.00058 | 0.98887 | 0.00064 | 0.98857 | 0.00057 | |
| 3 | 0.99999 | 1.2E-05 | 0.99993 | 6.2E-05 | 1 | 0 | 0.99998 | 1.5E-05 | |
| 4 | 0.99117 | 0.00036 | 0.99278 | 0.0003 | 0.99434 | 0.00021 | 0.99438 | 0.00017 | |
| 5 | 0.98091 | 0.00069 | 0.98429 | 0.00053 | 0.98744 | 0.00035 | 0.98765 | 0.00029 | |
| | vehicle | | | | | | | | |
| NoS | | | | | | | | | |
| 1 | 0.81158 | 0.0046 | 0.81096 | 0.0052 | 0.81795 | 0.00485 | 0.81885 | 0.00434 | 56 |
| 2 | 0.98942 | 0.00067 | 0.98875 | 0.00091 | 0.98964 | 0.00074 | 0.98845 | 0.0007 | |
| 3 | 0.99991 | 9.4E-05 | 0.99996 | 4E-05 | 0.99986 | 0.00011 | 1 | 0 | |
| 4 | 0.99084 | 0.00042 | 0.99101 | 0.00037 | 0.99108 | 0.00046 | 0.99114 | 0.00034 | |
| 5 | 0.98036 | 0.00075 | 0.98074 | 0.00068 | 0.98073 | 0.00077 | 0.98091 | 0.00064 | |

| FLT4 | VEGF D | | | | | | | | |
|---|---|---|---|---|---|---|---|---|---|
| NoS | lower bou | SE | lower bou | SE | lower bou | SE | lower bou | SE | n |
| 1 | 0.80371 | 0.00314 | 0.81145 | 0.00371 | 0.80817 | 0.00314 | 0.81151 | 0.00291 | 45 |
| 2 | 0.98517 | 0.00071 | 0.98589 | 0.00056 | 0.98495 | 0.00049 | 0.98474 | 0.00052 | |
| 3 | 1 | 0 | 1 | 0 | 1 | 0 | 1 | 0 | |
| 4 | 0.99346 | 0.00026 | 0.99386 | 0.00027 | 0.99443 | 0.00023 | 0.99452 | 0.00019 | |
| 5 | 0.98514 | 0.00047 | 0.98593 | 0.00046 | 0.98704 | 0.00041 | 0.98743 | 0.00035 | |
| | vehicle | | | | | | | | |
| NoS | | | | | | | | | |
| 1 | 0.80498 | 0.00436 | 0.81198 | 0.00419 | 0.80733 | 0.00385 | 0.81144 | 0.00332 | 40 |
| 2 | 0.98625 | 0.00081 | 0.98599 | 0.00078 | 0.98413 | 0.00081 | 0.98565 | 0.00072 | |
| 3 | 0.99995 | 5.2E-05 | 1 | 0 | 1 | 0 | 1 | 0 | |
| 4 | 0.99318 | 0.00029 | 0.99372 | 0.00026 | 0.99444 | 0.00025 | 0.99418 | 0.00025 | |
| 5 | 0.98476 | 0.00047 | 0.98579 | 0.00045 | 0.98697 | 0.00042 | 0.98675 | 0.00044 | |

FGFR

| FGFR1 | FGF1 | | | | | | | | | |
|---|---|---|---|---|---|---|---|---|---|---|
| | | 0min | | 2 min | | 12 min | | 22 min | | |
| NoS | | lower bou | SE | lower bou | SE | lower bou | SE | lower bou | SE | n |
| | 1 | 0.79283 | 0.00383 | 0.8098 | 0.00224 | 0.81133 | 0.00331 | 0.81442 | 0.00304 | 44 |
| | 2 | 0.98614 | 0.00066 | 0.98549 | 0.0005 | 0.98512 | 0.00056 | 0.98544 | 0.00052 | |
| | 3 | 1 | 0 | 1 | 0 | 1 | 0 | 1 | 0 | |
| | 4 | 0.99204 | 0.00032 | 0.99515 | 0.00027 | 0.99567 | 0.00021 | 0.99545 | 0.00021 | |
| | 5 | 0.98256 | 0.0006 | 0.9887 | 0.00051 | 0.98964 | 0.00038 | 0.98943 | 0.00041 | |
| | | | | | | | | | | 44 |
| | | vehicle | | | | | | | | |
| NoS | | | | | | | | | | |
| | 1 | 0.79777 | 0.00415 | 0.80572 | 0.0036 | 0.80558 | 0.00382 | 0.81374 | 0.00345 | 42 |
| | 2 | 0.98641 | 0.00079 | 0.98579 | 0.00073 | 0.98518 | 0.0007 | 0.9864 | 0.0008 | |
| | 3 | 1 | 0 | 1 | 0 | 1 | 0 | 1 | 0 | |
| | 4 | 0.99216 | 0.00031 | 0.99248 | 0.0003 | 0.993 | 0.00032 | 0.99284 | 0.00028 | |
| | 5 | 0.98267 | 0.00057 | 0.98336 | 0.00057 | 0.98412 | 0.00057 | 0.98411 | 0.0005 | |

| FGFR2 | FGF1 | | | | | | | | | |
|---|---|---|---|---|---|---|---|---|---|---|
| NoS | | lower bou | SE | lower bou | SE | lower bou | SE | lower bou | SE | n |
| | 1 | 0.80421 | 0.00364 | 0.81405 | 0.00322 | 0.82542 | 0.00278 | 0.82657 | 0.00364 | 48 |
| | 2 | 0.98623 | 0.00063 | 0.98513 | 0.00053 | 0.98617 | 0.00044 | 0.98632 | 0.00069 | |
| | 3 | 0.99995 | 4.8E-05 | 1 | 0 | 1 | 0 | 1 | 0 | |
| | 4 | 0.99231 | 0.00032 | 0.995 | 0.00017 | 0.99554 | 0.00014 | 0.99511 | 0.00017 | |
| | 5 | 0.98309 | 0.00058 | 0.98831 | 0.00032 | 0.98947 | 0.00025 | 0.9888 | 0.00029 | |
| | | vehicle | | | | | | | | |
| NoS | | | | | | | | | | |
| | 1 | 0.80675 | 0.0038 | 0.81609 | 0.00436 | 0.81708 | 0.00294 | 0.82096 | 0.00372 | 50 |
| | 2 | 0.98653 | 0.00076 | 0.98686 | 0.00079 | 0.98656 | 0.00069 | 0.9878 | 0.00065 | |
| | 3 | 0.99999 | 1.1E-05 | 0.99991 | 6.1E-05 | 1 | 0 | 0.99999 | 1.4E-05 | |
| | 4 | 0.99238 | 0.0003 | 0.9924 | 0.00033 | 0.99299 | 0.00024 | 0.99324 | 0.00022 | |
| | 5 | 0.98311 | 0.00054 | 0.98336 | 0.00055 | 0.98432 | 0.00043 | 0.98502 | 0.00038 | |

| FGFR3 | FGF1 | | | | | | | | | |
|---|---|---|---|---|---|---|---|---|---|---|
| NoS | | lower bou | SE | lower bou | SE | lower bou | SE | lower bou | SE | n |
| | 1 | 0.80613 | 0.00658 | 0.81865 | 0.00433 | 0.82386 | 0.00385 | 0.82652 | 0.00399 | 40 |
| | 2 | 0.98655 | 0.00097 | 0.98527 | 0.00067 | 0.98472 | 0.00072 | 0.98552 | 0.00062 | |
| | 3 | 0.99999 | 7.1E-06 | 1 | 0 | 1 | 0 | 1 | 0 | |
| | 4 | 0.99202 | 0.00037 | 0.99403 | 0.00029 | 0.99516 | 0.00022 | 0.99517 | 0.0002 | |
| | 5 | 0.98221 | 0.00065 | 0.98617 | 0.00053 | 0.98862 | 0.0004 | 0.98873 | 0.00035 | |
| | | vehicle | | | | | | | | |
| NoS | | | | | | | | | | |
| | 1 | 0.79826 | 0.00499 | 0.81102 | 0.00458 | 0.81746 | 0.00396 | 0.81533 | 0.00389 | 44 |
| | 2 | 0.98638 | 0.00094 | 0.9874 | 0.00071 | 0.98743 | 0.00075 | 0.98735 | 0.00065 | |
| | 3 | 0.99996 | 3.5E-05 | 1 | 0 | 0.99998 | 2E-05 | 1 | 0 | |
| | 4 | 0.99159 | 0.00043 | 0.99178 | 0.00038 | 0.99197 | 0.00036 | 0.99222 | 0.00029 | |
| | 5 | 0.98145 | 0.00075 | 0.98178 | 0.00068 | 0.9822 | 0.00064 | 0.98273 | 0.00053 | |

| FGFR4 | FGF1 | | | | | | | | | |
|---|---|---|---|---|---|---|---|---|---|---|
| NoS | | lower bou | SE | lower bou | SE | lower bou | SE | lower bou | SE | n |
| | 1 | 0.82189 | 0.00344 | 0.81953 | 0.00247 | 0.82698 | 0.00248 | 0.82834 | 0.00232 | 53 |
| | 2 | 0.986 | 0.00066 | 0.98483 | 0.00054 | 0.98567 | 0.00054 | 0.98585 | 0.00046 | |
| | 3 | 0.99999 | 1.1E-05 | 1 | 0 | 1 | 0 | 1 | 0 | |
| | 4 | 0.99428 | 0.00021 | 0.99537 | 0.00017 | 0.99542 | 0.00016 | 0.99542 | 0.00016 | |
| | 5 | 0.98695 | 0.00036 | 0.98908 | 0.00029 | 0.98936 | 0.00028 | 0.9894 | 0.0003 | |
| | | vehicle | | | | | | | | |
| NoS | | | | | | | | | | |
| | 1 | 0.81769 | 0.00393 | 0.82309 | 0.00315 | 0.82599 | 0.00351 | 0.82906 | 0.00388 | 44 |
| | 2 | 0.98553 | 0.00062 | 0.98611 | 0.00067 | 0.98695 | 0.00055 | 0.98631 | 0.00063 | |
| | 3 | 1 | 0 | 1 | 0 | 1 | 0 | 1 | 0 | |
| | 4 | 0.9944 | 0.00025 | 0.99461 | 0.00022 | 0.99468 | 0.00025 | 0.99468 | 0.00022 | |
| | 5 | 0.98717 | 0.00046 | 0.98774 | 0.0004 | 0.98787 | 0.00044 | 0.98789 | 0.00041 | |

PTK7

| PTK7 | Wnt5a | | | | | | | | | |
|---|---|---|---|---|---|---|---|---|---|---|
| | | 0min | | 2 min | | 12 min | | 22 min | | |
| NoS | | lower bou | SE | lower bou | SE | lower bou | SE | lower bou | SE | n |
| | 1 | 0.7515 | 0.00666 | 0.76397 | 0.00779 | 0.77115 | 0.00856 | 0.77571 | 0.00709 | 32 |
| | 2 | 0.98488 | 0.00098 | 0.98347 | 0.00112 | 0.98498 | 0.00137 | 0.98626 | 0.00107 | |
| | 3 | 1 | 0 | 0.9998 | 0.00016 | 0.99965 | 0.00026 | 0.99997 | 3.1E-05 | |
| | 4 | 0.99354 | 0.00033 | 0.99278 | 0.00046 | 0.99284 | 0.00066 | 0.99328 | 0.00052 | |
| | 5 | 0.98541 | 0.00063 | 0.98398 | 0.00076 | 0.98441 | 0.00105 | 0.98478 | 0.0009 | |
| | | vehicle | | | | | | | | |
| NoS | | | | | | | | | | |
| | 1 | 0.75175 | 0.00691 | 0.77138 | 0.0064 | 0.76773 | 0.0065 | 0.77682 | 0.00642 | 39 |
| | 2 | 0.98517 | 0.00087 | 0.98489 | 0.00104 | 0.98429 | 0.00101 | 0.98425 | 0.0011 | |
| | 3 | 0.99995 | 4.8E-05 | 1 | 0 | 1 | 0 | 1 | 0 | |
| | 4 | 0.99243 | 0.00041 | 0.99202 | 0.00048 | 0.99248 | 0.00047 | 0.99253 | 0.00043 | |
| | 5 | 0.98302 | 0.0008 | 0.98219 | 0.00087 | 0.98291 | 0.00084 | 0.98318 | 0.00077 | |

TRK

| TRKA | NGF | | | | | | | | | |
|---|---|---|---|---|---|---|---|---|---|---|
| | | 0min | | 2 min | | 12 min | | 22 min | | |
| NoS | | lower bou | SE | lower bou | SE | lower bou | SE | lower bou | SE | n |
| | 1 | 0.78756 | 0.00413 | 0.80494 | 0.00352 | 0.80687 | 0.0028 | 0.81516 | 0.00321 | 43 |
| | 2 | 0.98152 | 0.00087 | 0.98343 | 0.00069 | 0.98397 | 0.00058 | 0.98486 | 0.00069 | |
| | 3 | 1 | 0 | 1 | 1.5E-06 | 1 | 0 | 1 | 0 | |
| | 4 | 0.99442 | 0.00033 | 0.99486 | 0.00024 | 0.99486 | 0.00026 | 0.99483 | 0.00021 | |
| | 5 | 0.98659 | 0.00058 | 0.98773 | 0.00041 | 0.98782 | 0.0005 | 0.98802 | 0.00037 | |
| | | vehicle | | | | | | | | |
| NoS | | | | | | | | | | |
| | 1 | 0.78396 | 0.00404 | 0.79337 | 0.00327 | 0.80128 | 0.00302 | 0.80874 | 0.00307 | 31 |
| | 2 | 0.98124 | 0.00094 | 0.98228 | 0.0008 | 0.98257 | 0.00077 | 0.98369 | 0.00086 | |
| | 3 | 1 | 0 | 1 | 0 | 1 | 0 | 1 | 0 | |
| | 4 | 0.99389 | 0.00039 | 0.99448 | 0.00027 | 0.99381 | 0.00027 | 0.99394 | 0.00028 | |
| | 5 | 0.98521 | 0.00065 | 0.98643 | 0.00052 | 0.98554 | 0.0005 | 0.98593 | 0.00051 | |

| TRKB | BDNF | | | | | | | | | |
|---|---|---|---|---|---|---|---|---|---|---|
| NoS | | lower bou | SE | lower bou | SE | lower bou | SE | lower bou | SE | n |
| | 1 | 0.80149 | 0.00438 | 0.81627 | 0.00322 | 0.82194 | 0.00268 | 0.82194 | 0.00268 | 53 |
| | 2 | 0.98718 | 0.00076 | 0.98646 | 0.00061 | 0.98685 | 0.00055 | 0.98685 | 0.00055 | |
| | 3 | 0.99995 | 5.1E-05 | 1 | 0 | 1 | 0 | 1 | 0 | |
| | 4 | 0.99125 | 0.00036 | 0.99304 | 0.00024 | 0.99374 | 0.00019 | 0.99374 | 0.00019 | |
| | 5 | 0.98098 | 0.00062 | 0.98454 | 0.00045 | 0.98587 | 0.00036 | 0.98587 | 0.00036 | |
| | | vehicle | | | | | | | | |
| NoS | | | | | | | | | | |
| | 1 | 0.79899 | 0.00473 | 0.80315 | 0.00439 | 0.8082 | 0.00392 | 0.81648 | 0.00434 | 40 |
| | 2 | 0.98698 | 0.00091 | 0.98665 | 0.00077 | 0.9867 | 0.00089 | 0.98733 | 0.00076 | |
| | 3 | 1 | 0 | 1 | 0 | 1 | 0 | 1 | 0 | |
| | 4 | 0.99201 | 0.0003 | 0.99217 | 0.00031 | 0.99253 | 0.00028 | 0.99235 | 0.00032 | |
| | 5 | 0.98224 | 0.00054 | 0.98263 | 0.00056 | 0.98327 | 0.00051 | 0.98315 | 0.00056 | |

| TRKC | NT-3 | | | | | | | | | |
|---|---|---|---|---|---|---|---|---|---|---|
| NoS | | lower bou | SE | lower bou | SE | lower bou | SE | lower bou | SE | n |
| | 1 | 0.7932 | 0.00429 | 0.81543 | 0.00404 | 0.82 | 0.00229 | 0.82235 | 0.00257 | 50 |
| | 2 | 0.98543 | 0.00078 | 0.98648 | 0.00075 | 0.98612 | 0.00055 | 0.98602 | 0.00059 | |
| | 3 | 0.9999 | 0.0001 | 0.99991 | 8.6E-05 | 1 | 0 | 1 | 0 | |
| | 4 | 0.99164 | 0.00038 | 0.99256 | 0.00034 | 0.99396 | 0.00027 | 0.99407 | 0.00022 | |
| | 5 | 0.98152 | 0.00066 | 0.98363 | 0.0006 | 0.98627 | 0.00049 | 0.98643 | 0.00044 | |
| | | vehicle | | | | | | | | |
| NoS | | | | | | | | | | |
| | 1 | 0.80038 | 0.0032 | 0.8106 | 0.00324 | 0.81449 | 0.00305 | 0.81754 | 0.00307 | 67 |
| | 2 | 0.98651 | 0.00063 | 0.98701 | 0.00061 | 0.98685 | 0.00057 | 0.9872 | 0.00055 | |
| | 3 | 1 | 0 | 1 | 0 | 1 | 0 | 1 | 0 | |
| | 4 | 0.99195 | 0.00024 | 0.99234 | 0.00023 | 0.9923 | 0.00025 | 0.99231 | 0.00021 | |
| | 5 | 0.9822 | 0.00042 | 0.98295 | 0.00041 | 0.98292 | 0.00044 | 0.98311 | 0.00039 | |

ROR

| ROR1 | Wnt5a | | | | | | | | |
|---|---|---|---|---|---|---|---|---|---|
| | 0min | | 2 min | | 12 min | | 22 min | | |
| NoS | lower bou | SE | lower bou | SE | lower bou | SE | lower bou | SE | n |
| 1 | 0.78212 | 0.0062 | 0.79083 | 0.00593 | 0.79658 | 0.00611 | 0.79167 | 0.00529 | 45 |
| 2 | 0.9854 | 0.00074 | 0.98529 | 0.00077 | 0.98542 | 0.00087 | 0.98414 | 0.00082 | |
| 3 | 0.99982 | 0.00013 | 0.99996 | 4.3E-05 | 0.99995 | 4.9E-05 | 1 | 0 | |
| 4 | 0.9935 | 0.00036 | 0.99385 | 0.00028 | 0.99395 | 0.00027 | 0.99424 | 0.00025 | |
| 5 | 0.98532 | 0.00053 | 0.98575 | 0.00041 | 0.98609 | 0.00039 | 0.9865 | 0.00041 | |
| | vehicle | | | | | | | | |
| NoS | | | | | | | | | |
| 1 | 0.77788 | 0.00689 | 0.77709 | 0.00608 | 0.78322 | 0.00566 | 0.78953 | 0.00635 | 50 |
| 2 | 0.98644 | 0.00084 | 0.98537 | 0.00073 | 0.9837 | 0.00088 | 0.98506 | 0.00092 | |
| 3 | 0.99974 | 0.00015 | 1 | 0 | 0.99984 | 0.00011 | 0.99995 | 2.6E-05 | |
| 4 | 0.99321 | 0.00041 | 0.994 | 0.00026 | 0.99368 | 0.00028 | 0.99383 | 0.00033 | |
| 5 | 0.98497 | 0.0006 | 0.98573 | 0.0004 | 0.98534 | 0.00039 | 0.98565 | 0.00052 | |

MET

| MET | HGF | | | | | | | | |
|---|---|---|---|---|---|---|---|---|---|
| | 0min | | 2 min | | 12 min | | 22 min | | |
| NoS | lower bou | SE | lower bou | SE | lower bou | SE | lower bou | SE | n |
| 1 | 0.81743 | 0.00665 | 0.81001 | 0.00632 | 0.82526 | 0.00506 | 0.82759 | 0.00544 | 58 |
| 2 | 0.99583 | 0.00071 | 0.99321 | 0.00085 | 0.9917 | 0.0007 | 0.99218 | 0.00063 | |
| 3 | 0.99541 | 0.0007 | 0.99822 | 0.00044 | 0.99987 | 9.2E-05 | 0.99985 | 7.8E-05 | |
| 4 | 0.97836 | 0.00121 | 0.9865 | 0.00104 | 0.99288 | 0.00037 | 0.99274 | 0.00037 | |
| 5 | 0.96089 | 0.00165 | 0.97402 | 0.00161 | 0.98534 | 0.00064 | 0.98511 | 0.00066 | |
| | vehicle | | | | | | | | |
| NoS | | | | | | | | | |
| 1 | 0.8279 | 0.00835 | 0.8286 | 0.0082 | 0.82767 | 0.00856 | 0.8287 | 0.00772 | 47 |
| 2 | 0.99756 | 0.00066 | 0.99717 | 0.00077 | 0.9966 | 0.00083 | 0.9961 | 0.00088 | |
| 3 | 0.99362 | 0.00091 | 0.99453 | 0.00085 | 0.99533 | 0.00076 | 0.99642 | 0.00069 | |
| 4 | 0.97467 | 0.00151 | 0.97668 | 0.00147 | 0.97836 | 0.00139 | 0.98009 | 0.00126 | |
| 5 | 0.95547 | 0.00206 | 0.95838 | 0.00199 | 0.96074 | 0.00193 | 0.96326 | 0.0018 | |

| ROR2 | Wnt5a | | | | | | | | |
|---|---|---|---|---|---|---|---|---|---|
| NoS | lower bou | SE | lower bou | SE | lower bou | SE | lower bou | SE | n |
| 1 | 0.81436 | 0.00828 | 0.81551 | 0.00856 | 0.81773 | 0.00844 | 0.82003 | 0.00799 | 50 |
| 2 | 0.99116 | 0.00165 | 0.99111 | 0.00151 | 0.99195 | 0.00143 | 0.99149 | 0.00135 | |
| 3 | 0.99699 | 0.00078 | 0.99741 | 0.00066 | 0.99692 | 0.0007 | 0.99695 | 0.00073 | |
| 4 | 0.9821 | 0.00167 | 0.98287 | 0.00142 | 0.98221 | 0.00142 | 0.98262 | 0.00148 | |
| 5 | 0.96558 | 0.00245 | 0.96677 | 0.0021 | 0.96616 | 0.00207 | 0.96686 | 0.00212 | |
| | vehicle | | | | | | | | |
| NoS | | | | | | | | | |
| 1 | 0.81559 | 0.00831 | 0.81633 | 0.00847 | 0.81605 | 0.00857 | 0.81728 | 0.0082 | 42 |
| 2 | 0.99033 | 0.00198 | 0.99042 | 0.00186 | 0.99098 | 0.00158 | 0.99047 | 0.0017 | |
| 3 | 0.99804 | 0.00051 | 0.99868 | 0.00045 | 0.998 | 0.00054 | 0.99792 | 0.00068 | |
| 4 | 0.98358 | 0.0013 | 0.9849 | 0.00114 | 0.98392 | 0.00125 | 0.98417 | 0.00137 | |
| 5 | 0.96728 | 0.00201 | 0.96921 | 0.00175 | 0.96815 | 0.00187 | 0.96882 | 0.00201 | |

| RON | MSP-MST1 | | | | | | | | |
|---|---|---|---|---|---|---|---|---|---|
| NoS | lower bou | SE | lower bou | SE | lower bou | SE | lower bou | SE | n |
| 1 | 0.84775 | 0.0057 | 0.86213 | 0.0055 | 0.87809 | 0.00479 | 0.88642 | 0.00488 | 57 |
| 2 | 0.99441 | 0.00097 | 0.99594 | 0.00093 | 0.99796 | 0.00061 | 0.9987 | 0.00057 | |
| 3 | 0.99643 | 0.00079 | 0.99476 | 0.00089 | 0.99179 | 0.00102 | 0.99114 | 0.00104 | |
| 4 | 0.97688 | 0.0015 | 0.97525 | 0.0016 | 0.97096 | 0.0017 | 0.9702 | 0.00171 | |
| 5 | 0.95647 | 0.00216 | 0.955 | 0.00223 | 0.95 | 0.00234 | 0.94926 | 0.00236 | |
| | vehicle | | | | | | | | |
| NoS | | | | | | | | | |
| 1 | 0.84461 | 0.00724 | 0.86425 | 0.00566 | 0.8737 | 0.00499 | 0.88209 | 0.00492 | 59 |
| 2 | 0.99436 | 0.00121 | 0.99698 | 0.00078 | 0.99813 | 0.00056 | 0.99853 | 0.00058 | |
| 3 | 0.99521 | 0.00083 | 0.99416 | 0.00081 | 0.99215 | 0.00085 | 0.99084 | 0.00098 | |
| 4 | 0.97561 | 0.00145 | 0.97447 | 0.00136 | 0.97171 | 0.00136 | 0.96986 | 0.0016 | |
| 5 | 0.95525 | 0.00201 | 0.95431 | 0.00187 | 0.95103 | 0.00182 | 0.9488 | 0.00216 | |

MUSK

| MUSK | agrin | | | | | | | | |
|---|---|---|---|---|---|---|---|---|---|
| | 0min | | 2 min | | 12 min | | 22 min | | |
| NoS | lower bou | SE | lower bou | SE | lower bou | SE | lower bou | SE | n |
| 1 | 0.7719 | 0.00597 | 0.78032 | 0.0085 | 0.79223 | 0.00774 | 0.78455 | 0.00592 | 30 |
| 2 | 0.98561 | 0.00112 | 0.98494 | 0.00143 | 0.98688 | 0.00125 | 0.98398 | 0.00106 | |
| 3 | 1 | 0 | 0.99988 | 0.00011 | 0.99962 | 0.00029 | 1 | 0 | |
| 4 | 0.99201 | 0.00039 | 0.99253 | 0.00046 | 0.99185 | 0.00061 | 0.9932 | 0.00029 | |
| 5 | 0.98227 | 0.00069 | 0.98322 | 0.00071 | 0.98248 | 0.00085 | 0.98427 | 0.00048 | |
| | vehicle | | | | | | | | |
| NoS | | | | | | | | | |
| 1 | 0.76195 | 0.009 | 0.77494 | 0.00842 | 0.77637 | 0.00824 | 0.77637 | 0.01015 | 30 |
| 2 | 0.98414 | 0.0009 | 0.98341 | 0.00161 | 0.98455 | 0.00105 | 0.98392 | 0.00147 | |
| 3 | 1 | 0 | 1 | 0 | 1 | 0 | 1 | 0 | |
| 4 | 0.99247 | 0.00039 | 0.99276 | 0.00047 | 0.99313 | 0.00035 | 0.9934 | 0.00046 | |
| 5 | 0.98292 | 0.00065 | 0.98349 | 0.00075 | 0.98415 | 0.00059 | 0.98435 | 0.0007 | |

AXL

| AXL | GAS6 | | | | | | | | |
|---|---|---|---|---|---|---|---|---|---|
| | 0min | | 2 min | | 12 min | | 22 min | | |
| NoS | lower bou | SE | lower bou | SE | lower bou | SE | lower bou | SE | n |
| 1 | 0.76742 | 0.00568 | 0.78037 | 0.00664 | 0.7718 | 0.00524 | 0.76987 | 0.00499 | 33 |
| 2 | 0.98121 | 0.00102 | 0.98166 | 0.00126 | 0.9804 | 0.00101 | 0.97873 | 0.001 | |
| 3 | 1 | 0 | 1 | 0 | 1 | 0 | 1 | 0 | |
| 4 | 0.99314 | 0.00036 | 0.99382 | 0.00041 | 0.99518 | 0.00033 | 0.99536 | 0.00031 | |
| 5 | 0.98383 | 0.00065 | 0.98559 | 0.00069 | 0.98776 | 0.00053 | 0.98825 | 0.00059 | |
| | vehicle | | | | | | | | |
| NoS | | | | | | | | | |
| 1 | 0.77624 | 0.00638 | 0.78382 | 0.00515 | 0.77928 | 0.00595 | 0.77549 | 0.00534 | 31 |
| 2 | 0.98396 | 0.00091 | 0.98375 | 0.00113 | 0.97951 | 0.00106 | 0.98072 | 0.00081 | |
| 3 | 1 | 0 | 0.99996 | 4.4E-05 | 0.99986 | 0.00014 | 1 | 0 | |
| 4 | 0.99305 | 0.00042 | 0.99347 | 0.00047 | 0.99406 | 0.00049 | 0.9945 | 0.00038 | |
| 5 | 0.98412 | 0.0008 | 0.98496 | 0.00084 | 0.98591 | 0.00078 | 0.98657 | 0.0007 | |

| TYRO3 | GAS6 | | | | | | | | |
|---|---|---|---|---|---|---|---|---|---|
| NoS | lower bou | SE | lower bou | SE | lower bou | SE | lower bou | SE | n |
| 1 | 0.801 | 0.00574 | 0.80694 | 0.00542 | 0.79513 | 0.00491 | 0.79082 | 0.00518 | 43 |
| 2 | 0.98886 | 0.00093 | 0.98844 | 0.00098 | 0.98473 | 0.00083 | 0.984 | 0.00088 | |
| 3 | 1 | 0 | 0.99995 | 4.2E-05 | 1 | 0 | 1 | 0 | |
| 4 | 0.98951 | 0.00045 | 0.99032 | 0.00045 | 0.99185 | 0.0004 | 0.99256 | 0.00038 | |
| 5 | 0.97762 | 0.00085 | 0.97917 | 0.0008 | 0.98176 | 0.00072 | 0.98316 | 0.00068 | |
| | vehicle | | | | | | | | |
| NoS | | | | | | | | | |
| 1 | 0.81367 | 0.00465 | 0.81732 | 0.00465 | 0.81554 | 0.00473 | 0.82236 | 0.00449 | 55 |
| 2 | 0.99107 | 0.00075 | 0.9907 | 0.00065 | 0.98955 | 0.00087 | 0.99011 | 0.00077 | |
| 3 | 0.9998 | 0.00013 | 0.99989 | 7.4E-05 | 0.99993 | 3.3E-05 | 0.99967 | 0.0002 | |
| 4 | 0.98847 | 0.00042 | 0.98936 | 0.00036 | 0.98969 | 0.00034 | 0.98953 | 0.0005 | |
| 5 | 0.97601 | 0.00074 | 0.9774 | 0.00063 | 0.97812 | 0.00061 | 0.97802 | 0.00078 | |

TIE

| TIE1 | angiopoietin | | | | | | | | |
|---|---|---|---|---|---|---|---|---|---|
| | 0min | | 2 min | | 12 min | | 22 min | | |
| NoS | lower bou | SE | lower bou | SE | lower bou | SE | lower bou | SE | n |
| 1 | 0.77869 | 0.00885 | 0.79468 | 0.0088 | 0.8148 | 0.00811 | 0.8148 | 0.00811 | 38 |
| 2 | 0.98644 | 0.00152 | 0.99069 | 0.00115 | 0.99234 | 0.00106 | 0.99234 | 0.00106 | |
| 3 | 0.99973 | 0.00027 | 0.99965 | 0.0002 | 0.99918 | 0.0003 | 0.99918 | 0.0003 | |
| 4 | 0.98842 | 0.00079 | 0.98747 | 0.00079 | 0.98699 | 0.00082 | 0.98699 | 0.00082 | |
| 5 | 0.97522 | 0.0013 | 0.97382 | 0.00139 | 0.97394 | 0.00134 | 0.97394 | 0.00134 | |
| | vehicle | | | | | | | | |
| NoS | | | | | | | | | |
| 1 | 0.80525 | 0.00526 | 0.81481 | 0.00502 | 0.82127 | 0.00462 | 0.82331 | 0.00449 | 51 |
| 2 | 0.98924 | 0.00096 | 0.99231 | 0.00077 | 0.99356 | 0.00077 | 0.99376 | 0.0007 | |
| 3 | 0.99974 | 0.00022 | 0.9994 | 0.00025 | 0.99935 | 0.00023 | 0.99934 | 0.00025 | |
| 4 | 0.98818 | 0.00058 | 0.98744 | 0.00057 | 0.98781 | 0.00056 | 0.98771 | 0.00059 | |
| 5 | 0.9751 | 0.00097 | 0.97446 | 0.00091 | 0.97534 | 0.0009 | 0.97536 | 0.00095 | |

| TIE2 | angiopoietin | | | | | | | | |
|---|---|---|---|---|---|---|---|---|---|
| NoS | lower bou | SE | lower bou | SE | lower bou | SE | lower bou | SE | n |
| 1 | 0.80587 | 0.00422 | 0.78952 | 0.00471 | 0.81718 | 0.00446 | 0.83207 | 0.00404 | 71 |
| 2 | 0.98694 | 0.00133 | 0.98063 | 0.00109 | 0.97984 | 0.00077 | 0.98226 | 0.00074 | |
| 3 | 0.99982 | 0.0001 | 0.99996 | 3.6E-05 | 0.99997 | 3.3E-05 | 1 | 0 | |
| 4 | 0.98871 | 0.00056 | 0.99299 | 0.00041 | 0.99666 | 0.00018 | 0.99641 | 0.00016 | |
| 5 | 0.97526 | 0.00097 | 0.98322 | 0.00074 | 0.99061 | 0.0003 | 0.99058 | 0.00026 | |
| | vehicle | | | | | | | | |
| NoS | | | | | | | | | |
| 1 | 0.80825 | 0.0058 | 0.81534 | 0.0046 | 0.8234 | 0.00417 | 0.82516 | 0.00414 | 61 |
| 2 | 0.98808 | 0.00135 | 0.98867 | 0.00125 | 0.98992 | 0.00118 | 0.99116 | 0.00095 | |
| 3 | 0.99936 | 0.0003 | 0.99971 | 0.00016 | 0.99941 | 0.0003 | 0.99959 | 0.00022 | |
| 4 | 0.98817 | 0.00075 | 0.98886 | 0.00054 | 0.98843 | 0.00068 | 0.98879 | 0.00056 | |
| 5 | 0.97508 | 0.00114 | 0.97625 | 0.0009 | 0.97602 | 0.00103 | 0.97673 | 0.00087 | |

EPHA

| EPH A1 | ephrin A1 | | | | | | | | |
|---|---|---|---|---|---|---|---|---|---|
| | 0min | | 2 min | | 12 min | | 22 min | | |
| NoS | lower bou | SE | lower bou | SE | lower bou | SE | lower bou | SE | n |
| 1 | 0.80547 | 0.00438 | 0.82138 | 0.00355 | 0.82616 | 0.00278 | 0.83048 | 0.00426 | 31 |
| 2 | 0.98661 | 0.00078 | 0.98698 | 0.00064 | 0.98539 | 0.00053 | 0.98561 | 0.00064 | |
| 3 | 1 | 0 | 1 | 0 | 1 | 0 | 1 | 0 | |
| 4 | 0.99283 | 0.00025 | 0.99361 | 0.00021 | 0.99507 | 0.00019 | 0.99541 | 0.00019 | |
| 5 | 0.98418 | 0.00044 | 0.98586 | 0.00036 | 0.98855 | 0.00032 | 0.98933 | 0.00034 | |
| | vehicle | | | | | | | | |
| NoS | | | | | | | | | |
| 1 | 0.80789 | 0.00437 | 0.81498 | 0.00386 | 0.81235 | 0.00313 | 0.8147 | 0.00326 | 38 |
| 2 | 0.98712 | 0.0007 | 0.98815 | 0.00071 | 0.98649 | 0.00063 | 0.98683 | 0.00071 | |
| 3 | 1 | 0 | 1 | 0 | 1 | 0 | 0.99999 | 1.5E-05 | |
| 4 | 0.99288 | 0.00035 | 0.99289 | 0.00027 | 0.99329 | 0.00022 | 0.99322 | 0.0003 | |
| 5 | 0.98426 | 0.00065 | 0.98437 | 0.0005 | 0.98517 | 0.00045 | 0.98496 | 0.00058 | |

| EPH A2 | ephrin A1 | | | | | | | | |
|---|---|---|---|---|---|---|---|---|---|
| NoS | lower bou | SE | lower bou | SE | lower bou | SE | lower bou | SE | n |
| 1 | 0.80921 | 0.00682 | 0.82339 | 0.00477 | 0.83231 | 0.00339 | 0.84073 | 0.00291 | 38 |
| 2 | 0.98871 | 0.00099 | 0.98955 | 0.00083 | 0.98657 | 0.00066 | 0.98798 | 0.00076 | |
| 3 | 1 | 0 | 0.99996 | 4.1E-05 | 1 | 0 | 1 | 0 | |
| 4 | 0.99138 | 0.00038 | 0.99219 | 0.00035 | 0.99413 | 0.00024 | 0.99431 | 0.0002 | |
| 5 | 0.98148 | 0.00067 | 0.98321 | 0.0006 | 0.9868 | 0.00044 | 0.98744 | 0.00037 | |
| | vehicle | | | | | | | | |
| NoS | | | | | | | | | |
| 1 | 0.80372 | 0.00564 | 0.81346 | 0.00418 | 0.81638 | 0.00438 | 0.83071 | 0.00597 | 30 |
| 2 | 0.98834 | 0.00092 | 0.98873 | 0.00071 | 0.98802 | 0.00081 | 0.98871 | 0.00064 | |
| 3 | 0.99991 | 8.6E-05 | 1 | 0 | 1 | 0 | 1 | 0 | |
| 4 | 0.9919 | 0.00041 | 0.99222 | 0.00045 | 0.992 | 0.00045 | 0.99227 | 0.00046 | |
| 5 | 0.98251 | 0.00078 | 0.98311 | 0.00088 | 0.98258 | 0.00088 | 0.98323 | 0.00091 | |

| EPHA3 | ephrin A1 | | | | | | | | |
|---|---|---|---|---|---|---|---|---|---|
| | 0min | | 2 min | | 12 min | | 22 min | | |
| NoS | lower bou | SE | lower bou | SE | lower bou | SE | lower bou | SE | n |
| 1 | 0.8137 | 0.00344 | 0.8136 | 0.00347 | 0.8199 | 0.00261 | 0.82005 | 0.00264 | 56 |
| 2 | 0.98849 | 0.00075 | 0.98846 | 0.00079 | 0.98798 | 0.00076 | 0.98755 | 0.00077 | |
| 3 | 1 | 0 | 1 | 0 | 1 | 0 | 1 | 0 | |
| 4 | 0.99137 | 0.0003 | 0.99173 | 0.00031 | 0.99225 | 0.00032 | 0.99219 | 0.00032 | |
| 5 | 0.98132 | 0.00056 | 0.9822 | 0.00056 | 0.98314 | 0.00057 | 0.98297 | 0.00058 | |
| | vehicle | | | | | | | | |
| NoS | | | | | | | | | |
| 1 | 0.80566 | 0.00379 | 0.81249 | 0.00411 | 0.81787 | 0.00325 | 0.82242 | 0.00314 | 50 |
| 2 | 0.98755 | 0.00068 | 0.98849 | 0.00081 | 0.98808 | 0.00072 | 0.98881 | 0.00081 | |
| 3 | 1 | 0 | 0.99993 | 6.7E-05 | 1 | 0 | 0.9998 | 0.0002 | |
| 4 | 0.99183 | 0.00034 | 0.99182 | 0.00035 | 0.9919 | 0.00037 | 0.99153 | 0.00054 | |
| 5 | 0.98223 | 0.00063 | 0.98235 | 0.00062 | 0.98242 | 0.00067 | 0.98209 | 0.00088 | |

| EPHA4 | ephrin A1 | | | | | | | | |
|---|---|---|---|---|---|---|---|---|---|
| NoS | lower bou | SE | lower bou | SE | lower bou | SE | lower bou | SE | n |
| 1 | 0.81433 | 0.00438 | 0.82458 | 0.00354 | 0.81947 | 0.0032 | 0.82109 | 0.00392 | 30 |
| 2 | 0.98889 | 0.00085 | 0.98963 | 0.00084 | 0.98742 | 0.00061 | 0.9874 | 0.00083 | |
| 3 | 1 | 0 | 0.99993 | 6.6E-05 | 1 | 0 | 1 | 0 | |
| 4 | 0.99115 | 0.00041 | 0.99155 | 0.0005 | 0.99286 | 0.00033 | 0.99298 | 0.00034 | |
| 5 | 0.98094 | 0.00076 | 0.98192 | 0.00087 | 0.98432 | 0.00065 | 0.9847 | 0.00064 | |
| | vehicle | | | | | | | | |
| NoS | | | | | | | | | |
| 1 | 0.80286 | 0.00576 | 0.80689 | 0.00593 | 0.80902 | 0.00545 | 0.81127 | 0.00563 | 37 |
| 2 | 0.98651 | 0.00123 | 0.98689 | 0.00126 | 0.98642 | 0.00102 | 0.98737 | 0.00102 | |
| 3 | 0.99997 | 2E-05 | 0.99995 | 4.8E-05 | 1 | 0 | 0.99997 | 3.2E-05 | |
| 4 | 0.99207 | 0.00041 | 0.99243 | 0.00036 | 0.99229 | 0.00032 | 0.99251 | 0.00037 | |
| 5 | 0.98262 | 0.00068 | 0.98334 | 0.00059 | 0.98309 | 0.00057 | 0.98356 | 0.00064 | |

| EPHA6 | ephrin A2 | | | | | | | | |
|---|---|---|---|---|---|---|---|---|---|
| NoS | lower bou | SE | lower bou | SE | lower bou | SE | lower bou | SE | n |
| 1 | 0.7976 | 0.00325 | 0.80816 | 0.00316 | 0.81739 | 0.00288 | 0.82184 | 0.00309 | 49 |
| 2 | 0.98482 | 0.00063 | 0.98538 | 0.00064 | 0.98606 | 0.00052 | 0.9862 | 0.00058 | |
| 3 | 1 | 0 | 1 | 0 | 1 | 0 | 1 | 0 | |
| 4 | 0.99489 | 0.00023 | 0.9948 | 0.00026 | 0.99477 | 0.00021 | 0.99462 | 0.00022 | |
| 5 | 0.98818 | 0.00041 | 0.98807 | 0.00047 | 0.98807 | 0.00038 | 0.98782 | 0.00041 | |
| | vehicle | | | | | | | | |
| NoS | | | | | | | | | |
| 1 | 0.79652 | 0.00304 | 0.79808 | 0.00314 | 0.80713 | 0.00265 | 0.80598 | 0.00276 | 42 |
| 2 | 0.98434 | 0.00064 | 0.98349 | 0.00068 | 0.9838 | 0.00065 | 0.98337 | 0.00059 | |
| 3 | 1 | 0 | 1 | 0 | 1 | 0 | 1 | 0 | |
| 4 | 0.99534 | 0.0002 | 0.99554 | 0.00021 | 0.99529 | 0.00025 | 0.99557 | 0.00018 | |
| 5 | 0.98904 | 0.00035 | 0.98932 | 0.00035 | 0.9889 | 0.00043 | 0.98947 | 0.00031 | |

| EPHA7 | ephrin A2 | | | | | | | | |
|---|---|---|---|---|---|---|---|---|---|
| NoS | lower bou | SE | lower bou | SE | lower bou | SE | lower bou | SE | n |
| 1 | 0.80838 | 0.00278 | 0.81243 | 0.0037 | 0.81522 | 0.00353 | 0.81142 | 0.00428 | 60 |
| 2 | 0.98777 | 0.00069 | 0.98815 | 0.00065 | 0.98801 | 0.00067 | 0.9875 | 0.00048 | |
| 3 | 0.99998 | 1.5E-05 | 0.99993 | 7.1E-05 | 0.99994 | 4.5E-05 | 1 | 0 | |
| 4 | 0.99152 | 0.00037 | 0.99188 | 0.00033 | 0.99243 | 0.00031 | 0.99284 | 0.00028 | |
| 5 | 0.9816 | 0.00068 | 0.98244 | 0.0006 | 0.98346 | 0.00057 | 0.9842 | 0.00054 | |
| | vehicle | | | | | | | | |
| NoS | | | | | | | | | |
| 1 | 0.80034 | 0.00409 | 0.80851 | 0.00363 | 0.80848 | 0.00399 | 0.80796 | 0.00317 | 55 |
| 2 | 0.98786 | 0.00065 | 0.98706 | 0.00079 | 0.98703 | 0.00078 | 0.9861 | 0.00072 | |
| 3 | 0.99996 | 4.2E-05 | 0.99987 | 8.5E-05 | 0.99996 | 3.6E-05 | 0.99991 | 7.9E-05 | |
| 4 | 0.99159 | 0.00032 | 0.99186 | 0.00038 | 0.99222 | 0.00033 | 0.99275 | 0.00033 | |
| 5 | 0.98183 | 0.00058 | 0.98242 | 0.00065 | 0.98297 | 0.00059 | 0.98391 | 0.00058 | |

EPHB

| EPH B1 | | ephrin B1 | | | | | | | | |
|---|---|---|---|---|---|---|---|---|---|---|
| | | 0min | | 2 min | | 12 min | | 22 min | | |
| NoS | | lower bou | SE | lower bou | SE | lower bou | SE | lower bou | SE | n |
| | 1 | 0.80255 | 0.00495 | 0.80167 | 0.00414 | 0.80732 | 0.00549 | 0.81654 | 0.00403 | 36 |
| | 2 | 0.98592 | 0.00085 | 0.9855 | 0.00071 | 0.98536 | 0.00077 | 0.9867 | 0.00074 | |
| | 3 | 0.99999 | 1.3E-05 | 1 | 0 | 0.99986 | 0.00015 | 0.99992 | 7.7E-05 | |
| | 4 | 0.99404 | 0.00034 | 0.99439 | 0.00027 | 0.9944 | 0.00028 | 0.99416 | 0.00031 | |
| | 5 | 0.98659 | 0.00061 | 0.98711 | 0.00047 | 0.98729 | 0.00046 | 0.98705 | 0.00053 | |
| | | vehicle | | | | | | | | |
| NoS | | | | | | | | | | |
| | 1 | 0.7995 | 0.00401 | 0.8057 | 0.00356 | 0.80256 | 0.00333 | 0.80476 | 0.00334 | 37 |
| | 2 | 0.98477 | 0.00085 | 0.9855 | 0.00065 | 0.98433 | 0.00089 | 0.98535 | 0.00074 | |
| | 3 | 1 | 0 | 1 | 0 | 1 | 0 | 1 | 0 | |
| | 4 | 0.99346 | 0.00035 | 0.99387 | 0.00028 | 0.99412 | 0.00026 | 0.99415 | 0.00023 | |
| | 5 | 0.98512 | 0.00062 | 0.98625 | 0.00052 | 0.98651 | 0.00044 | 0.98672 | 0.00042 | |

| EPHB2 | | ephrin B2 | | | | | | | | |
|---|---|---|---|---|---|---|---|---|---|---|
| NoS | | lower bou | SE | lower bou | SE | lower bou | SE | lower bou | SE | n |
| | 1 | 0.78029 | 0.00439 | 0.7912 | 0.00414 | 0.80293 | 0.00341 | 0.80723 | 0.00322 | 50 |
| | 2 | 0.98522 | 0.00076 | 0.9856 | 0.00068 | 0.98479 | 0.00061 | 0.98541 | 0.00054 | |
| | 3 | 1 | 0 | 1 | 0 | 1 | 0 | 1 | 0 | |
| | 4 | 0.99215 | 0.00033 | 0.993 | 0.00025 | 0.99341 | 0.00022 | 0.99373 | 0.00022 | |
| | 5 | 0.98241 | 0.00063 | 0.98407 | 0.00047 | 0.98485 | 0.0004 | 0.98563 | 0.0004 | |
| | | vehicle | | | | | | | | |
| NoS | | | | | | | | | | |
| | 1 | 0.79282 | 0.00382 | 0.79828 | 0.00354 | 0.79034 | 0.00356 | 0.79437 | 0.00386 | 52 |
| | 2 | 0.98616 | 0.00062 | 0.9862 | 0.00062 | 0.98443 | 0.00064 | 0.98478 | 0.00067 | |
| | 3 | 1 | 0 | 1 | 0 | 1 | 0 | 1 | 0 | |
| | 4 | 0.99191 | 0.00029 | 0.99226 | 0.00027 | 0.99278 | 0.00025 | 0.99311 | 0.00026 | |
| | 5 | 0.98193 | 0.00053 | 0.98281 | 0.00048 | 0.98365 | 0.00044 | 0.98433 | 0.00046 | |

| EPH B3 | | ephrin B1 | | | | | | | | |
|---|---|---|---|---|---|---|---|---|---|---|
| NoS | | lower bou | SE | lower bou | SE | lower bou | SE | lower bou | SE | n |
| | 1 | 0.8017 | 0.00388 | 0.81259 | 0.00483 | 0.81289 | 0.00348 | 0.80861 | 0.00396 | 54 |
| | 2 | 0.98875 | 0.00077 | 0.98863 | 0.00069 | 0.98709 | 0.00059 | 0.98654 | 0.00057 | |
| | 3 | 1 | 1E-06 | 0.99996 | 3.7E-05 | 1 | 5.1E-07 | 1 | 0 | |
| | 4 | 0.99162 | 0.00031 | 0.99184 | 0.00035 | 0.99289 | 0.00027 | 0.9931 | 0.00026 | |
| | 5 | 0.9819 | 0.00058 | 0.98242 | 0.00063 | 0.98432 | 0.00052 | 0.98471 | 0.00049 | |
| | | vehicle | | | | | | | | |
| NoS | | | | | | | | | | |
| | 1 | 0.80686 | 0.00514 | 0.8102 | 0.0058 | 0.81757 | 0.00519 | 0.81391 | 0.00707 | 32 |
| | 2 | 0.98978 | 0.00081 | 0.98885 | 0.00095 | 0.98803 | 0.00103 | 0.98807 | 0.00097 | |
| | 3 | 0.99996 | 4.3E-05 | 0.99988 | 0.00012 | 1 | 0 | 0.99985 | 0.00015 | |
| | 4 | 0.99197 | 0.00052 | 0.99198 | 0.00056 | 0.99243 | 0.0005 | 0.99247 | 0.00053 | |
| | 5 | 0.98272 | 0.00097 | 0.98273 | 0.00097 | 0.98336 | 0.00089 | 0.98368 | 0.00091 | |

| EPH B4 | | ephrin B2 | | | | | | | | |
|---|---|---|---|---|---|---|---|---|---|---|
| NoS | | lower bou | SE | lower bou | SE | lower bou | SE | lower bou | SE | n |
| | 1 | 0.79781 | 0.00293 | 0.80724 | 0.0026 | 0.80695 | 0.00297 | 0.80894 | 0.00279 | 48 |
| | 2 | 0.98501 | 0.00061 | 0.98501 | 0.00063 | 0.98484 | 0.00057 | 0.98455 | 0.00059 | |
| | 3 | 1 | 0 | 1 | 0 | 1 | 0 | 1 | 0 | |
| | 4 | 0.99349 | 0.00027 | 0.99371 | 0.00031 | 0.9939 | 0.00028 | 0.99393 | 0.00027 | |
| | 5 | 0.98528 | 0.00053 | 0.98565 | 0.00057 | 0.98603 | 0.00051 | 0.98619 | 0.00051 | |
| | | vehicle | | | | | | | | |
| NoS | | | | | | | | | | |
| | 1 | 0.80277 | 0.00292 | 0.80878 | 0.00293 | 0.80757 | 0.00215 | 0.80586 | 0.00233 | 51 |
| | 2 | 0.98501 | 0.00064 | 0.98493 | 0.00074 | 0.98529 | 0.00052 | 0.98393 | 0.00072 | |
| | 3 | 1 | 0 | 1 | 0 | 1 | 0 | 1 | 0 | |
| | 4 | 0.99337 | 0.00022 | 0.99364 | 0.00025 | 0.99386 | 0.00021 | 0.99431 | 0.00022 | |
| | 5 | 0.98504 | 0.00042 | 0.9856 | 0.00045 | 0.98603 | 0.00041 | 0.98676 | 0.0004 | |

| EPH B6 | | ephrin B1 | | | | | | | | |
|---|---|---|---|---|---|---|---|---|---|---|
| NoS | | lower bou | SE | lower bou | SE | lower bou | SE | lower bou | SE | n |
| | 1 | 0.80862 | 0.00366 | 0.81104 | 0.0034 | 0.81409 | 0.00336 | 0.81587 | 0.0038 | 37 |
| | 2 | 0.98595 | 0.00078 | 0.98554 | 0.0008 | 0.98604 | 0.00074 | 0.98616 | 0.00075 | |
| | 3 | 1 | 0 | 1 | 0 | 1 | 0 | 1 | 0 | |
| | 4 | 0.99377 | 0.00027 | 0.99421 | 0.00026 | 0.99434 | 0.00025 | 0.99442 | 0.00023 | |
| | 5 | 0.98601 | 0.00048 | 0.98683 | 0.00044 | 0.98733 | 0.00042 | 0.98743 | 0.00039 | |
| | | vehicle | | | | | | | | |
| NoS | | | | | | | | | | |
| | 1 | 0.80767 | 0.003 | 0.81003 | 0.00278 | 0.81395 | 0.00277 | 0.81509 | 0.00307 | 40 |
| | 2 | 0.98512 | 0.0008 | 0.98453 | 0.00073 | 0.9858 | 0.00076 | 0.98577 | 0.0007 | |
| | 3 | 1 | 4.2E-06 | 1 | 0 | 1 | 0 | 1 | 0 | |
| | 4 | 0.99394 | 0.0003 | 0.99444 | 0.00024 | 0.99455 | 0.00023 | 0.99461 | 0.00024 | |
| | 5 | 0.9863 | 0.00052 | 0.98727 | 0.00042 | 0.98764 | 0.0004 | 0.98783 | 0.00041 | |

RET

| RET | | GDNF | | | | | | | | |
|---|---|---|---|---|---|---|---|---|---|---|
| | | 0min | | 2 min | | 12 min | | 22 min | | |
| NoS | | lower bou | SE | lower bou | SE | lower bou | SE | lower bou | SE | n |
| | 1 | 0.80261 | 0.00398 | 0.8077 | 0.00312 | 0.81453 | 0.00299 | 0.81734 | 0.00308 | 53 |
| | 2 | 0.98649 | 0.00078 | 0.98609 | 0.00067 | 0.98606 | 0.00059 | 0.98664 | 0.00069 | |
| | 3 | 1 | 0 | 1 | 0 | 1 | 0 | 1 | 0 | |
| | 4 | 0.99247 | 0.00028 | 0.9931 | 0.00026 | 0.99337 | 0.00022 | 0.99336 | 0.00023 | |
| | 5 | 0.98333 | 0.0005 | 0.98462 | 0.00046 | 0.98533 | 0.0004 | 0.98525 | 0.00042 | |
| | | vehicle | | | | | | | | |
| NoS | | | | | | | | | | |
| | 1 | 0.79597 | 0.00294 | 0.80921 | 0.00252 | 0.81278 | 0.00271 | 0.81985 | 0.00322 | 57 |
| | 2 | 0.98501 | 0.0006 | 0.98632 | 0.00061 | 0.98676 | 0.00058 | 0.98766 | 0.00068 | |
| | 3 | 1 | 0 | 1 | 0 | 1 | 0 | 0.99999 | 6.3E-06 | |
| | 4 | 0.99288 | 0.00028 | 0.99271 | 0.00025 | 0.99287 | 0.00026 | 0.99272 | 0.00027 | |
| | 5 | 0.98382 | 0.0005 | 0.98378 | 0.00046 | 0.98426 | 0.00047 | 0.9841 | 0.0005 | |

DDR

| DDR1 | | EGF | | | | | | | | |
|---|---|---|---|---|---|---|---|---|---|---|
| | | 0min | | 2 min | | 12 min | | 22 min | | |
| NoS | | lower bou | SE | lower bou | SE | lower bou | SE | lower bou | SE | n |
| | 1 | 0.85721 | 0.00438 | 0.85973 | 0.0039 | 0.86338 | 0.00346 | 0.87084 | 0.00312 | 45 |
| | 2 | 0.99804 | 0.00081 | 0.99861 | 0.00072 | 0.9989 | 0.00046 | 0.99886 | 0.00049 | |
| | 3 | 0.99477 | 0.00075 | 0.99476 | 0.00062 | 0.99468 | 0.00066 | 0.99442 | 0.00061 | |
| | 4 | 0.97761 | 0.00131 | 0.97814 | 0.00112 | 0.97809 | 0.0011 | 0.97846 | 0.00106 | |
| | 5 | 0.96016 | 0.0018 | 0.96133 | 0.00156 | 0.96143 | 0.00152 | 0.96244 | 0.00145 | |
| | | vehicle | | | | | | | | |
| NoS | | | | | | | | | | |
| | 1 | 0.86084 | 0.00455 | 0.86509 | 0.00502 | 0.86636 | 0.0051 | 0.86739 | 0.00423 | 44 |
| | 2 | 0.99831 | 0.0006 | 0.99804 | 0.00067 | 0.99839 | 0.00056 | 0.99844 | 0.00048 | |
| | 3 | 0.99317 | 0.00107 | 0.99273 | 0.00107 | 0.99399 | 0.001 | 0.9951 | 0.00087 | |
| | 4 | 0.97383 | 0.00173 | 0.97398 | 0.00179 | 0.97606 | 0.00165 | 0.978 | 0.00144 | |
| | 5 | 0.95425 | 0.00232 | 0.95501 | 0.00244 | 0.95788 | 0.00225 | 0.96067 | 0.00198 | |

| DDR2 | | EGF | | | | | | | | |
|---|---|---|---|---|---|---|---|---|---|---|
| | | 0min | | 2 min | | 12 min | | 22 min | | |
| NoS | | lower bou | SE | lower bou | SE | lower bou | SE | lower bou | SE | n |
| | 1 | 0.86169 | 0.00474 | 0.86901 | 0.0045 | 0.87154 | 0.00409 | 0.86965 | 0.00357 | 44 |
| | 2 | 0.99876 | 0.00055 | 0.99912 | 0.00044 | 0.9995 | 0.00026 | 0.99954 | 0.00029 | |
| | 3 | 0.99387 | 0.00089 | 0.99389 | 0.00085 | 0.99303 | 0.00078 | 0.99427 | 0.00071 | |
| | 4 | 0.97529 | 0.00147 | 0.97585 | 0.00136 | 0.9749 | 0.00123 | 0.97696 | 0.00116 | |
| | 5 | 0.95663 | 0.00204 | 0.95776 | 0.00187 | 0.95678 | 0.00166 | 0.95968 | 0.00162 | |
| | | vehicle | | | | | | | | |
| NoS | | | | | | | | | | |
| | 1 | 0.86962 | 0.00331 | 0.87061 | 0.00283 | 0.87666 | 0.00326 | 0.87639 | 0.00269 | 44 |
| | 2 | 0.99948 | 0.00026 | 0.99977 | 0.00017 | 0.99954 | 0.00024 | 0.99996 | 2.5E-05 | |
| | 3 | 0.99089 | 0.00084 | 0.99158 | 0.00074 | 0.99197 | 0.00081 | 0.99252 | 0.00063 | |
| | 4 | 0.97144 | 0.00136 | 0.97272 | 0.00119 | 0.97332 | 0.00131 | 0.9746 | 0.00099 | |
| | 5 | 0.95211 | 0.00185 | 0.95407 | 0.0016 | 0.95492 | 0.00179 | 0.9568 | 0.00134 | |

ROS

| ROS1 | | EGF | | | | | | | | |
|---|---|---|---|---|---|---|---|---|---|---|
| | | 0min | | 2 min | | 12 min | | 22 min | | |
| NoS | | lower bou | SE | lower bou | SE | lower bou | SE | lower bou | SE | n |
| | 1 | 0.87487 | 0.00448 | 0.87121 | 0.00344 | 0.87921 | 0.00387 | 0.87638 | 0.00393 | 44 |
| | 2 | 0.99941 | 0.00032 | 0.99923 | 0.00028 | 0.99888 | 0.00054 | 0.99882 | 0.00052 | |
| | 3 | 0.99153 | 0.00087 | 0.99491 | 0.00066 | 0.99228 | 0.0009 | 0.99354 | 0.00074 | |
| | 4 | 0.97247 | 0.00137 | 0.9779 | 0.00103 | 0.97364 | 0.00143 | 0.97611 | 0.00124 | |
| | 5 | 0.95346 | 0.00181 | 0.96074 | 0.00138 | 0.95497 | 0.00192 | 0.95854 | 0.0017 | |
| | | vehicle | | | | | | | | |
| NoS | | | | | | | | | | |
| | 1 | 0.87443 | 0.00363 | 0.88008 | 0.00312 | 0.87617 | 0.00372 | 0.87742 | 0.00345 | 57 |
| | 2 | 0.99932 | 0.00026 | 0.99929 | 0.00027 | 0.99892 | 0.00037 | 0.99885 | 0.00039 | |
| | 3 | 0.99375 | 0.00067 | 0.99427 | 0.00066 | 0.9944 | 0.00068 | 0.99478 | 0.00066 | |
| | 4 | 0.97645 | 0.0011 | 0.97739 | 0.0011 | 0.97727 | 0.00117 | 0.97805 | 0.0011 | |
| | 5 | 0.95912 | 0.00153 | 0.96046 | 0.00152 | 0.96005 | 0.00165 | 0.96122 | 0.00153 | |

LMR

| LMR1 | EGF | | | | | | | | |
|---|---|---|---|---|---|---|---|---|---|
| | 0min | | 2 min | | 12 min | | 22 min | | |
| NoS | lower bou | SE | lower bou | SE | lower bou | SE | lower bou | SE | n |
| 1 | 0.74693 | 0.00827 | 0.7605 | 0.00751 | 0.77196 | 0.00739 | 0.78506 | 0.00642 | 43 |
| 2 | 0.9818 | 0.00165 | 0.98446 | 0.00172 | 0.98649 | 0.00164 | 0.98838 | 0.00145 | |
| 3 | 0.99931 | 0.00043 | 0.99951 | 0.00025 | 0.99845 | 0.00052 | 0.99897 | 0.0005 | |
| 4 | 0.98867 | 0.00133 | 0.98834 | 0.0011 | 0.98675 | 0.00141 | 0.98713 | 0.00135 | |
| 5 | 0.97532 | 0.00216 | 0.97528 | 0.00193 | 0.97312 | 0.00225 | 0.97358 | 0.00217 | |
| | vehicle | | | | | | | | |
| NoS | | | | | | | | | |
| 1 | 0.75688 | 0.00757 | 0.77256 | 0.00757 | 0.79279 | 0.00624 | 0.80068 | 0.00727 | 42 |
| 2 | 0.98279 | 0.0021 | 0.98453 | 0.00181 | 0.98856 | 0.00141 | 0.99062 | 0.00132 | |
| 3 | 0.99826 | 0.00075 | 0.99738 | 0.00096 | 0.9977 | 0.00075 | 0.99676 | 0.00097 | |
| 4 | 0.98553 | 0.00203 | 0.98414 | 0.00225 | 0.98487 | 0.00183 | 0.98316 | 0.00221 | |
| 5 | 0.97046 | 0.00326 | 0.96918 | 0.00348 | 0.97053 | 0.00285 | 0.96859 | 0.00339 | |

| LMR2 | EGF | | | | | | | | |
|---|---|---|---|---|---|---|---|---|---|
| NoS | lower bou | SE | lower bou | SE | lower bou | SE | lower bou | SE | n |
| 1 | 0.82372 | 0.00386 | 0.82085 | 0.00398 | 0.81952 | 0.00382 | 0.82343 | 0.00368 | 57 |
| 2 | 0.98566 | 0.00078 | 0.98652 | 0.00066 | 0.98551 | 0.00075 | 0.98599 | 0.0008 | |
| 3 | 1 | 0 | 1 | 4.6E-06 | 1 | 0 | 1 | 0 | |
| 4 | 0.99477 | 0.00024 | 0.99508 | 0.00017 | 0.99507 | 0.0002 | 0.99506 | 0.00024 | |
| 5 | 0.98833 | 0.00044 | 0.98901 | 0.00032 | 0.98882 | 0.00037 | 0.98899 | 0.00044 | |
| | vehicle | | | | | | | | |
| NoS | | | | | | | | | |
| 1 | 0.79918 | 0.0057 | 0.80847 | 0.00623 | 0.80645 | 0.00628 | 0.80363 | 0.00608 | 42 |
| 2 | 0.98418 | 0.00081 | 0.98494 | 0.00087 | 0.98328 | 0.00093 | 0.98209 | 0.00091 | |
| 3 | 1 | 0 | 1 | 0 | 1 | 0 | 1 | 0 | |
| 4 | 0.99509 | 0.0002 | 0.99518 | 0.00023 | 0.99566 | 0.00022 | 0.99562 | 0.00021 | |
| 5 | 0.98889 | 0.00035 | 0.98901 | 0.00042 | 0.98966 | 0.00033 | 0.98969 | 0.00036 | |

STYK

| STYK1 | EGF | | | | | | | | |
|---|---|---|---|---|---|---|---|---|---|
| | 0min | | 2 min | | 12 min | | 22 min | | |
| NoS | lower bou | SE | lower bou | SE | lower bou | SE | lower bou | SE | n |
| 1 | 0.85302 | 0.00545 | 0.84212 | 0.00575 | 0.8516 | 0.00498 | 0.84558 | 0.00535 | 44 |
| 2 | 0.99595 | 0.00083 | 0.99389 | 0.00075 | 0.99564 | 0.00068 | 0.99437 | 0.00077 | |
| 3 | 0.998 | 0.00042 | 0.9994 | 0.00023 | 0.99882 | 0.00039 | 0.99889 | 0.00043 | |
| 4 | 0.98548 | 0.00079 | 0.98916 | 0.00051 | 0.98752 | 0.00072 | 0.9881 | 0.0008 | |
| 5 | 0.97243 | 0.00112 | 0.97804 | 0.00075 | 0.97571 | 0.00103 | 0.97657 | 0.00114 | |
| | vehicle | | | | | | | | |
| NoS | | | | | | | | | |
| 1 | 0.8389 | 0.00684 | 0.84394 | 0.00572 | 0.84799 | 0.00651 | 0.84568 | 0.00511 | 41 |
| 2 | 0.99457 | 0.00091 | 0.99443 | 0.00089 | 0.99397 | 0.00107 | 0.99392 | 0.00093 | |
| 3 | 0.99866 | 0.00043 | 0.99855 | 0.0004 | 0.99851 | 0.0005 | 0.99907 | 0.00032 | |
| 4 | 0.98704 | 0.00089 | 0.98746 | 0.0008 | 0.98759 | 0.00097 | 0.98885 | 0.00074 | |
| 5 | 0.97467 | 0.0013 | 0.97566 | 0.00115 | 0.97604 | 0.00141 | 0.97784 | 0.00112 | |

ALK

| LTK | EGF | | | | | | | | |
|---|---|---|---|---|---|---|---|---|---|
| | 0min | | 2 min | | 12 min | | 22 min | | |
| NoS | lower bou | SE | lower bou | SE | lower bou | SE | lower bou | SE | n |
| 1 | 0.82459 | 0.00315 | 0.83352 | 0.00293 | 0.83568 | 0.00274 | 0.84359 | 0.00278 | 60 |
| 2 | 0.99288 | 0.00081 | 0.99376 | 0.00076 | 0.99471 | 0.00065 | 0.99569 | 0.0006 | |
| 3 | 0.99948 | 0.00021 | 0.99962 | 0.00015 | 0.99958 | 0.00015 | 0.9991 | 0.00025 | |
| 4 | 0.98763 | 0.00053 | 0.98846 | 0.00044 | 0.98823 | 0.00041 | 0.98694 | 0.00054 | |
| 5 | 0.97471 | 0.00084 | 0.97645 | 0.00071 | 0.976 | 0.00065 | 0.97409 | 0.00082 | |
| | vehicle | | | | | | | | |
| NoS | | | | | | | | | |
| 1 | 0.82166 | 0.00307 | 0.82935 | 0.00261 | 0.83196 | 0.0026 | 0.84039 | 0.00261 | 58 |
| 2 | 0.99226 | 0.00074 | 0.99379 | 0.00063 | 0.99353 | 0.00061 | 0.99517 | 0.00056 | |
| 3 | 0.99993 | 3.5E-05 | 0.9996 | 0.00016 | 0.99973 | 0.0001 | 0.99956 | 0.00013 | |
| 4 | 0.98858 | 0.00034 | 0.98813 | 0.00041 | 0.98858 | 0.00033 | 0.98816 | 0.00033 | |
| 5 | 0.97611 | 0.00063 | 0.97577 | 0.00064 | 0.97658 | 0.00056 | 0.9761 | 0.00052 | |

| ALK | Heparin | | | | | | | | |
|---|---|---|---|---|---|---|---|---|---|
| | 0min | | 2 min | | 12 min | | 22 min | | |
| NoS | lower bou | SE | lower bou | SE | lower bou | SE | lower bou | SE | n |
| 1 | 0.81328 | 0.00297 | 0.80901 | 0.00253 | 0.81232 | 0.00255 | 0.81499 | 0.00246 | 64 |
| 2 | 0.98452 | 0.0006 | 0.98407 | 0.00059 | 0.98466 | 0.00055 | 0.98464 | 0.00046 | |
| 3 | 1 | 4.5E-06 | 1 | 0 | 1 | 0 | 1 | 0 | |
| 4 | 0.99563 | 0.00021 | 0.99553 | 0.00019 | 0.99554 | 0.00015 | 0.99573 | 0.00013 | |
| 5 | 0.98962 | 0.00035 | 0.98942 | 0.00032 | 0.98959 | 0.00026 | 0.98998 | 0.00023 | |
| | vehicle | | | | | | | | |
| NoS | | | | | | | | | |
| 1 | 0.80839 | 0.00307 | 0.81538 | 0.00322 | 0.81808 | 0.00284 | 0.8188 | 0.00318 | 63 |
| 2 | 0.98417 | 0.00056 | 0.98456 | 0.00063 | 0.98539 | 0.0006 | 0.98536 | 0.00067 | |
| 3 | 1 | 0 | 1 | 0 | 1 | 0 | 0.99999 | 9.8E-06 | |
| 4 | 0.99565 | 0.00021 | 0.99558 | 0.00021 | 0.99542 | 0.00021 | 0.99546 | 0.00023 | |
| 5 | 0.9897 | 0.00037 | 0.98959 | 0.00038 | 0.98941 | 0.00038 | 0.98946 | 0.0004 | |
