## Supplement Table S2 for "Single-molecule behavior and cell-growth regulation in human RTKs"

| RTK | resting state parameters | | | | | | | | | | | | | | | | | | | | | | | |
|---|---|---|---|---|---|---|---|---|---|---|---|---|---|---|---|---|---|---|---|---|---|---|---|---|
| | D1 | D2 | D3 | P1 | P2 | P3 | L1 | L2 | L3 | E1 | E2 | E3 | I1 | I2 | I3 | F1 | F2 | F3 | tau12 | tau13 | tau21 | tau23 | tau31 | tau32 |
| EGFR | 0.0069 | 0.0343 | 0.2594 | 27.0614 | 34.0069 | 38.9317 | 0.0931 | 0.3268 | 1.2064 | 0.0635 | 0.1587 | 0.8889 | 2.8317 | 2.9006 | 2.5273 | 0.1958 | 0.2702 | 0.2532 | 0.3788 | 0.4320 | 0.4029 | 0.3210 | 0.4940 | 0.3416 |
| ERBB2 | 0.0132 | 0.0454 | 0.2902 | 19.5409 | 34.9593 | 45.4998 | 0.0814 | 0.2980 | 1.4737 | 0.0000 | 0.0444 | 0.6222 | 2.1996 | 2.1996 | 1.9294 | 0.1010 | 0.1925 | 0.2229 | 0.3646 | 0.4040 | 0.4415 | 0.3289 | 0.5486 | 0.3675 |
| ERBB3 | 0.0059 | 0.0298 | 0.2471 | 32.6443 | 37.2950 | 30.0608 | 0.0926 | 0.3097 | 1.1812 | 0.0179 | 0.0714 | 0.7857 | 3.5663 | 3.2816 | 2.7094 | 0.4424 | 0.4849 | 0.2978 | 0.3835 | 0.4836 | 0.3917 | 0.3469 | 0.4541 | 0.3141 |
| ERBB4 | 0.0065 | 0.0310 | 0.2616 | 28.6658 | 35.9034 | 35.4308 | 0.0934 | 0.3354 | 1.2656 | 0.0238 | 0.1395 | 0.8140 | 2.8496 | 2.9675 | 2.5624 | 0.3265 | 0.4503 | 0.3479 | 0.3785 | 0.4455 | 0.3983 | 0.3322 | 0.4774 | 0.3278 |
| INSR | 0.0069 | 0.0363 | 0.2450 | 24.2139 | 34.3218 | 41.4643 | 0.1004 | 0.3561 | 1.1535 | 0.0222 | 0.0889 | 0.9111 | 2.6348 | 2.7285 | 2.4147 | 0.2776 | 0.4330 | 0.4135 | 0.3778 | 0.4199 | 0.4225 | 0.3121 | 0.5250 | 0.3417 |
| IGF1R | 0.0099 | 0.0420 | 0.2483 | 18.3213 | 31.9576 | 49.7212 | 0.0858 | 0.3383 | 1.2511 | 0.0000 | 0.0682 | 0.9091 | 3.0114 | 2.9505 | 2.7310 | 0.2404 | 0.4402 | 0.6363 | 0.3633 | 0.4048 | 0.4412 | 0.3259 | 0.5914 | 0.3935 |
| INSRR | 0.0118 | 0.0451 | 0.2559 | 17.9618 | 35.6618 | 46.3764 | 0.0934 | 0.3418 | 1.3479 | 0.0000 | 0.0702 | 0.8421 | 2.8705 | 2.8692 | 2.5623 | 0.2129 | 0.4559 | 0.5091 | 0.3552 | 0.3865 | 0.4440 | 0.3274 | 0.5447 | 0.3640 |
| PDGFRA | 0.0057 | 0.0306 | 0.2872 | 27.2019 | 31.7936 | 41.0044 | 0.0947 | 0.3397 | 1.6215 | 0.0800 | 0.3000 | 0.9000 | 2.5429 | 2.5934 | 2.2394 | 0.2181 | 0.2760 | 0.2852 | 0.3886 | 0.4556 | 0.4044 | 0.3262 | 0.5349 | 0.3636 |
| PDGFRB | 0.0082 | 0.0403 | 0.2855 | 22.8091 | 33.8114 | 43.3795 | 0.0938 | 0.4589 | 1.3327 | 0.0000 | 0.3556 | 0.8889 | 2.7858 | 2.7858 | 2.6779 | 0.2212 | 0.3416 | 0.3832 | 0.3665 | 0.3938 | 0.4137 | 0.3082 | 0.5020 | 0.3445 |
| KIT | 0.0103 | 0.0347 | 0.3064 | 23.6246 | 29.0118 | 47.3636 | 0.0770 | 0.2713 | 1.5465 | 0.0244 | 0.1463 | 0.7317 | 1.8294 | 1.6926 | 1.6875 | 0.0619 | 0.0807 | 0.1355 | 0.3806 | 0.4341 | 0.4072 | 0.3429 | 0.5717 | 0.4170 |
| CSF1R | 0.0058 | 0.0325 | 0.2702 | 28.5765 | 33.1237 | 38.1670 | 0.0896 | 0.3322 | 1.3718 | 0.0233 | 0.1628 | 0.8837 | 2.7927 | 2.8194 | 2.3588 | 0.2787 | 0.3439 | 0.3036 | 0.3863 | 0.4578 | 0.3969 | 0.3287 | 0.5113 | 0.3487 |
| FLT3 | 0.0063 | 0.0317 | 0.2540 | 29.6185 | 35.8475 | 34.5340 | 0.0899 | 0.3855 | 1.3717 | 0.0000 | 0.3171 | 0.9024 | 2.6160 | 2.5474 | 2.1929 | 0.2871 | 0.3509 | 0.2775 | 0.3920 | 0.4744 | 0.4087 | 0.3372 | 0.4939 | 0.3296 |
| FLT1 | 0.0037 | 0.0243 | 0.2710 | 34.2690 | 30.2344 | 35.4966 | 0.0670 | 0.2761 | 1.4191 | 0.1500 | 0.1463 | 0.9756 | 2.6855 | 2.0794 | 2.0070 | 0.1899 | 0.1858 | 0.1668 | 0.4368 | 0.5586 | 0.3943 | 0.3480 | 0.5469 | 0.3718 |
| KDR | 0.0067 | 0.0351 | 0.2700 | 27.5408 | 32.9832 | 39.4760 | 0.0866 | 0.3341 | 1.0885 | 0.0278 | 0.2603 | 0.9041 | 2.0811 | 2.0017 | 1.7624 | 0.1574 | 0.1999 | 0.2019 | 0.3911 | 0.4651 | 0.4128 | 0.3402 | 0.5347 | 0.3646 |
| FLT4 | 0.0063 | 0.0326 | 0.2552 | 29.2421 | 36.9785 | 33.7794 | 0.0962 | 0.3970 | 1.4048 | 0.0227 | 0.1333 | 0.8889 | 2.8575 | 2.7941 | 2.4117 | 0.3496 | 0.4539 | 0.3428 | 0.3806 | 0.4609 | 0.4054 | 0.3398 | 0.4799 | 0.3247 |
| FGFR1 | 0.0063 | 0.0360 | 0.2711 | 28.7211 | 35.3688 | 35.9101 | 0.0993 | 0.3282 | 1.3357 | 0.0455 | 0.0909 | 0.8409 | 2.7606 | 2.9804 | 2.6026 | 0.3097 | 0.4304 | 0.3498 | 0.3814 | 0.4537 | 0.4021 | 0.3339 | 0.4848 | 0.3343 |
| FGFR2 | 0.0068 | 0.0336 | 0.2627 | 27.5176 | 36.3640 | 36.1184 | 0.0939 | 0.3695 | 1.3119 | 0.0000 | 0.1875 | 0.8542 | 2.7886 | 2.8728 | 2.4734 | 0.2853 | 0.4090 | 0.3266 | 0.3753 | 0.4318 | 0.4065 | 0.3267 | 0.4740 | 0.3249 |
| FGFR3 | 0.0064 | 0.0344 | 0.2496 | 26.0521 | 34.9800 | 38.9680 | 0.1025 | 0.3298 | 1.4774 | 0.0000 | 0.2000 | 0.9000 | 2.9988 | 2.8476 | 2.4541 | 0.2692 | 0.3566 | 0.3063 | 0.3805 | 0.4378 | 0.4171 | 0.3225 | 0.5142 | 0.3406 |
| FGFR4 | 0.0062 | 0.0321 | 0.2252 | 29.4207 | 39.6945 | 30.8848 | 0.1124 | 0.3782 | 1.2958 | 0.0392 | 0.1509 | 0.8679 | 3.2403 | 3.0607 | 2.6394 | 0.3829 | 0.4983 | 0.3117 | 0.3727 | 0.4405 | 0.4081 | 0.3372 | 0.4418 | 0.3038 |
| PTK7 | 0.0064 | 0.0309 | 0.3107 | 36.5217 | 32.5962 | 30.8820 | 0.0795 | 0.3108 | 1.8050 | 0.0263 | 0.2308 | 0.9487 | 1.8820 | 1.6310 | 1.9163 | 0.0934 | 0.0781 | 0.0799 | 0.4235 | 0.5574 | 0.3915 | 0.3841 | 0.4997 | 0.3655 |
| TRKA | 0.0058 | 0.0324 | 0.2580 | 29.7865 | 38.8605 | 31.3530 | 0.0927 | 0.3285 | 1.2214 | 0.0233 | 0.0233 | 0.7674 | 3.4607 | 3.4736 | 3.0921 | 0.3548 | 0.4944 | 0.3349 | 0.3725 | 0.4588 | 0.4026 | 0.3479 | 0.4551 | 0.3119 |
| TRKB | 0.0078 | 0.0380 | 0.2735 | 26.2435 | 36.4185 | 37.3380 | 0.0976 | 0.3476 | 1.3006 | 0.0000 | 0.1132 | 0.8868 | 2.9020 | 3.0259 | 2.7017 | 0.2360 | 0.3707 | 0.3139 | 0.3741 | 0.4298 | 0.4088 | 0.3303 | 0.4870 | 0.3352 |
| TRKC | 0.0066 | 0.0357 | 0.2728 | 26.5529 | 34.8496 | 38.5975 | 0.0909 | 0.3569 | 1.3274 | 0.0000 | 0.1800 | 0.8600 | 2.6690 | 2.8421 | 2.5238 | 0.2391 | 0.3551 | 0.3234 | 0.3774 | 0.4339 | 0.4073 | 0.3236 | 0.5000 | 0.3410 |
| ROR1 | 0.0054 | 0.0278 | 0.2721 | 33.7011 | 32.0678 | 34.2311 | 0.0865 | 0.3287 | 1.1599 | 0.0600 | 0.1600 | 0.9200 | 2.2754 | 2.1453 | 1.9432 | 0.2116 | 0.2146 | 0.1911 | 0.4153 | 0.5305 | 0.3893 | 0.3524 | 0.5168 | 0.3571 |
| ROR2 | 0.0098 | 0.0426 | 0.2751 | 18.9715 | 37.5508 | 43.4778 | 0.0852 | 0.2729 | 1.3659 | 0.0238 | 0.0476 | 0.7857 | 3.0044 | 2.8125 | 2.5250 | 0.2041 | 0.3941 | 0.3958 | 0.3521 | 0.3986 | 0.4419 | 0.3402 | 0.5385 | 0.3621 |
| MUSK | 0.0067 | 0.0350 | 0.3116 | 29.7743 | 33.5526 | 36.6730 | 0.0883 | 0.3650 | 1.1855 | 0.0000 | 0.1667 | 0.8667 | 2.4064 | 2.4453 | 2.2568 | 0.1942 | 0.2441 | 0.2285 | 0.3873 | 0.4679 | 0.3953 | 0.3395 | 0.4980 | 0.3471 |
| MET | 0.0108 | 0.0416 | 0.3041 | 19.6432 | 32.7946 | 47.6477 | 0.0916 | 0.3291 | 1.4782 | 0.0000 | 0.0862 | 0.8276 | 2.4821 | 2.7393 | 2.4996 | 0.2114 | 0.3994 | 0.4847 | 0.3584 | 0.3953 | 0.4179 | 0.3289 | 0.5557 | 0.3899 |
| RON | 0.0132 | 0.0512 | 0.2574 | 16.1445 | 34.7126 | 49.1429 | 0.0807 | 0.2874 | 1.3121 | 0.0000 | 0.0351 | 0.7018 | 2.7427 | 3.0776 | 2.5341 | 0.1134 | 0.2927 | 0.3358 | 0.3585 | 0.3708 | 0.4546 | 0.3090 | 0.5484 | 0.3591 |
| AXL | 0.0063 | 0.0330 | 0.2673 | 28.4852 | 34.2548 | 37.2599 | 0.0817 | 0.3287 | 1.3196 | 0.0000 | 0.1212 | 0.8788 | 3.2694 | 3.2314 | 2.7898 | 0.2804 | 0.3581 | 0.3151 | 0.3854 | 0.4607 | 0.4064 | 0.3269 | 0.5085 | 0.3387 |
| TYRO3 | 0.0080 | 0.0380 | 0.2769 | 24.8646 | 33.8830 | 41.2523 | 0.0913 | 0.3337 | 1.4555 | 0.0000 | 0.1395 | 0.8605 | 2.8612 | 3.2628 | 2.9239 | 0.2326 | 0.4060 | 0.4227 | 0.3725 | 0.4142 | 0.4089 | 0.3169 | 0.4973 | 0.3436 |
| TIE1 | 0.0070 | 0.0336 | 0.2657 | 24.5988 | 32.8206 | 42.5806 | 0.0776 | 0.2615 | 1.1132 | 0.0000 | 0.0789 | 0.8684 | 3.1633 | 2.8408 | 2.5676 | 0.2834 | 0.3962 | 0.4297 | 0.3882 | 0.4762 | 0.4239 | 0.3569 | 0.5917 | 0.4006 |
| TIE2 | 0.0090 | 0.0378 | 0.2517 | 21.9690 | 40.5732 | 37.4578 | 0.0822 | 0.2524 | 1.1144 | 0.0141 | 0.0282 | 0.8028 | 3.3272 | 2.8529 | 2.4315 | 0.3082 | 0.5112 | 0.3942 | 0.3593 | 0.4385 | 0.4517 | 0.3675 | 0.5212 | 0.3438 |
| EPHA1 | 0.0072 | 0.0354 | 0.2685 | 28.7605 | 38.1040 | 33.1356 | 0.1078 | 0.3658 | 1.2498 | 0.0000 | 0.1613 | 0.7097 | 3.0136 | 3.1635 | 2.7851 | 0.3042 | 0.4453 | 0.3250 | 0.3700 | 0.4317 | 0.4005 | 0.3339 | 0.4500 | 0.3153 |
| EPHA2 | 0.0067 | 0.0362 | 0.2605 | 27.4003 | 35.1506 | 37.4491 | 0.1029 | 0.3520 | 1.2723 | 0.0263 | 0.1579 | 0.7632 | 2.8213 | 2.8642 | 2.5216 | 0.2107 | 0.2997 | 0.2721 | 0.3744 | 0.4304 | 0.4019 | 0.3301 | 0.4785 | 0.3365 |
| EPHA3 | 0.0069 | 0.0379 | 0.2615 | 26.1804 | 36.1705 | 37.6491 | 0.1060 | 0.3424 | 1.3635 | 0.0000 | 0.1429 | 0.8393 | 2.9735 | 2.9733 | 2.5356 | 0.2395 | 0.3450 | 0.2960 | 0.3694 | 0.4247 | 0.4091 | 0.3286 | 0.4839 | 0.3344 |
| EPHA4 | 0.0080 | 0.0392 | 0.2718 | 25.7492 | 36.5445 | 37.7063 | 0.1012 | 0.3723 | 1.3399 | 0.0333 | 0.1000 | 0.9333 | 3.0953 | 3.2818 | 2.8467 | 0.2643 | 0.4183 | 0.3469 | 0.3678 | 0.4132 | 0.4105 | 0.3220 | 0.4751 | 0.3309 |
| EPHA6 | 0.0046 | 0.0249 | 0.2256 | 34.9460 | 37.2400 | 27.8140 | 0.1021 | 0.3183 | 1.1278 | 0.0204 | 0.0816 | 0.8571 | 2.9902 | 2.7086 | 2.3629 | 0.3977 | 0.4011 | 0.2437 | 0.3968 | 0.5265 | 0.3941 | 0.3717 | 0.4601 | 0.3210 |
| EPHA7 | 0.0073 | 0.0381 | 0.2682 | 26.5814 | 37.0083 | 36.4103 | 0.1060 | 0.3328 | 1.3393 | 0.0169 | 0.1500 | 0.8500 | 3.3041 | 3.3815 | 2.9774 | 0.3135 | 0.4702 | 0.3707 | 0.3640 | 0.4212 | 0.4052 | 0.3324 | 0.4720 | 0.3305 |
| EPHB1 | 0.0052 | 0.0276 | 0.2256 | 31.2844 | 37.4404 | 31.2752 | 0.1063 | 0.2966 | 1.1909 | 0.1389 | 0.1622 | 0.7297 | 3.2422 | 3.1033 | 2.7000 | 0.3158 | 0.3785 | 0.2671 | 0.3784 | 0.4676 | 0.3971 | 0.3528 | 0.4584 | 0.3218 |
| EPHB2 | 0.0059 | 0.0326 | 0.2736 | 27.7494 | 32.0586 | 40.1919 | 0.0864 | 0.3392 | 1.2937 | 0.0208 | 0.2200 | 0.9800 | 2.6150 | 2.6242 | 2.2443 | 0.2386 | 0.3132 | 0.2986 | 0.3915 | 0.4602 | 0.4009 | 0.3231 | 0.5292 | 0.3557 |
| EPHB3 | 0.0059 | 0.0330 | 0.2618 | 27.7095 | 32.5229 | 39.7677 | 0.0958 | 0.3214 | 1.2600 | 0.0185 | 0.1852 | 0.9630 | 2.5320 | 2.4648 | 2.0887 | 0.2024 | 0.2571 | 0.2407 | 0.3878 | 0.4583 | 0.4028 | 0.3331 | 0.5282 | 0.3660 |
| EPHB4 | 0.0054 | 0.0274 | 0.2440 | 30.0527 | 35.9137 | 34.0336 | 0.0998 | 0.3451 | 1.2520 | 0.0000 | 0.1667 | 0.8958 | 2.9149 | 2.7520 | 2.3485 | 0.3024 | 0.3595 | 0.2720 | 0.3841 | 0.4762 | 0.4027 | 0.3470 | 0.4929 | 0.3370 |
| EPHB6 | 0.0063 | 0.0316 | 0.2348 | 30.1993 | 38.7517 | 31.0490 | 0.1031 | 0.3382 | 1.4212 | 0.0541 | 0.0811 | 0.9189 | 3.0351 | 2.8443 | 2.5597 | 0.3177 | 0.4043 | 0.2766 | 0.3759 | 0.4574 | 0.4029 | 0.3519 | 0.4523 | 0.3161 |
| RET | 0.0070 | 0.0343 | 0.2538 | 27.7946 | 37.3711 | 34.8343 | 0.1001 | 0.3228 | 1.3363 | 0.0377 | 0.0755 | 0.9434 | 2.9522 | 2.9417 | 2.5484 | 0.2685 | 0.3739 | 0.2836 | 0.3715 | 0.4380 | 0.4072 | 0.3371 | 0.4721 | 0.3251 |
| DDR1 | 0.0125 | 0.0453 | 0.2686 | 17.9142 | 39.3032 | 42.7826 | 0.1066 | 0.2975 | 1.2605 | 0.0000 | 0.0889 | 0.7333 | 2.8407 | 2.6194 | 2.2277 | 0.1929 | 0.4030 | 0.3402 | 0.3370 | 0.3667 | 0.4370 | 0.3409 | 0.4976 | 0.3505 |
| DDR2 | 0.0136 | 0.0493 | 0.2728 | 16.9671 | 38.5487 | 44.4843 | 0.1128 | 0.3132 | 1.4500 | 0.0000 | 0.0455 | 0.7500 | 3.1168 | 2.8580 | 2.4035 | 0.2213 | 0.2052 | 0.4014 | 0.3355 | 0.3587 | 0.4398 | 0.3357 | 0.5081 | 0.3581 |
| ROS1 | 0.0149 | 0.0482 | 0.2523 | 16.1174 | 38.4752 | 45.4074 | 0.1089 | 0.3087 | 1.2042 | 0.0000 | 0.0682 | 0.7500 | 2.8046 | 2.8442 | 2.1680 | 0.1262 | 0.3030 | 0.2638 | 0.3319 | 0.3421 | 0.4381 | 0.3277 | 0.4937 | 0.3536 |
| LMR1 | 0.0109 | 0.0361 | 0.3289 | 29.0781 | 41.0740 | 29.8479 | 0.0664 | 0.2049 | 1.3551 | 0.0000 | 0.0233 | 0.7442 | 2.4096 | 2.0057 | 1.8725 | 0.1045 | 0.1414 | 0.1010 | 0.3920 | 0.5510 | 0.4446 | 0.4233 | 0.5000 | 0.3392 |
| LMR2 | 0.0058 | 0.0258 | 0.2153 | 34.3168 | 41.2620 | 24.4213 | 0.1056 | 0.3048 | 1.0204 | 0.0175 | 0.0702 | 0.8772 | 2.5339 | 2.2688 | 1.9739 | 0.3907 | 0.4164 | 0.2024 | 0.3861 | 0.5161 | 0.4080 | 0.3974 | 0.4304 | 0.3078 |
| LTK | 0.0094 | 0.0389 | 0.2454 | 23.0037 | 42.7457 | 34.2506 | 0.1106 | 0.2741 | 1.2864 | 0.0000 | 0.0167 | 0.6500 | 3.6059 | 3.1594 | 2.5938 | 0.3402 | 0.5743 | 0.3716 | 0.3478 | 0.4104 | 0.4344 | 0.3595 | 0.4676 | 0.3251 |
| ALK | 0.0055 | 0.0277 | 0.2073 | 33.6439 | 41.5001 | 24.8559 | 0.1076 | 0.3128 | 1.1018 | 0.0317 | 0.1250 | 0.7500 | 3.3774 | 2.9924 | 2.5738 | 0.5072 | 0.5471 | 0.2711 | 0.3762 | 0.4810 | 0.3978 | 0.3675 | 0.4163 | 0.2932 |
| STYK1 | 0.0092 | 0.0400 | 0.2830 | 21.3508 | 33.6021 | 45.0471 | 0.1123 | 0.3977 | 1.3221 | 0.0233 | 0.3864 | 0.8636 | 2.3279 | 2.2905 | 2.0270 | 0.1349 | 0.2226 | 0.2534 | 0.3643 | 0.3912 | 0.4181 | 0.3163 | 0.5201 | 0.3627 |

| RTK | response state parameters | | | | | | | | | | | | | | | | | | | | | | | |
|---|---|---|---|---|---|---|---|---|---|---|---|---|---|---|---|---|---|---|---|---|---|---|---|---|
| | rD1(1) | rD1(2) | rD1(3) | rD2(1) | rD2(2) | rD2(3) | rD3(1) | rD3(2) | rD3(3) | rP1(1) | rP1(2) | rP1(3) | rP2(1) | rP2(2) | rP2(3) | rP3(1) | rP3(2) | rP3(3) | rL1(1) | rL1(2) | rL1(3) | rL2(1) | rL2(2) | rL2(3) |
| EGFR | 0.5599 | 0.6626 | 0.6916 | 0.5973 | 0.7166 | 0.7117 | 0.8159 | 0.8863 | 0.8914 | 1.3694 | 1.2542 | 1.1967 | 1.0846 | 1.0101 | 1.0258 | 0.6385 | 0.7931 | 0.8165 | 0.9854 | 0.9746 | 0.9918 | 0.9057 | 0.8781 | 0.8917 |
| ERBB2 | 0.8602 | 0.8925 | 0.8529 | 0.9129 | 0.8891 | 0.9086 | 0.9957 | 1.0025 | 1.0392 | 1.0245 | 0.9508 | 0.9966 | 1.0158 | 0.9919 | 0.9746 | 0.9750 | 1.0259 | 1.0186 | 1.0406 | 1.0169 | 1.0244 | 0.9235 | 1.0245 | 0.9055 |
| ERBB3 | 0.9680 | 1.0987 | 1.0524 | 1.0090 | 1.0399 | 1.0689 | 0.9195 | 0.9284 | 0.9496 | 1.0036 | 0.9317 | 0.9606 | 1.0402 | 1.0365 | 1.0439 | 0.9433 | 1.0325 | 0.9868 | 1.0274 | 1.0874 | 1.0893 | 1.1010 | 1.1666 | 1.1656 |
| ERBB4 | 0.7785 | 0.9118 | 0.9118 | 0.8710 | 0.9386 | 0.9011 | 0.9419 | 0.9259 | 0.9332 | 1.1308 | 1.0952 | 1.1131 | 1.0165 | 1.0418 | 1.0306 | 0.8686 | 0.8729 | 0.8635 | 1.0228 | 1.0077 | 1.0071 | 1.1189 | 1.0576 | 1.0015 |
| INSR | 0.9881 | 0.9426 | 1.0209 | 1.0237 | 1.0383 | 0.9680 | 1.0080 | 1.0517 | 1.0091 | 1.0460 | 1.0269 | 1.0117 | 1.0190 | 0.9716 | 1.0081 | 0.9518 | 1.0065 | 0.9801 | 0.9595 | 0.9626 | 0.9385 | 1.0189 | 0.9835 | 1.0613 |
| IGF1R | 1.0124 | 0.9506 | 0.9578 | 1.0584 | 1.0606 | 0.9855 | 0.9949 | 0.9862 | 0.9976 | 1.0659 | 1.0367 | 1.0432 | 1.0273 | 1.0124 | 1.0209 | 0.9520 | 0.9716 | 0.9590 | 0.9665 | 1.0615 | 0.9891 | 1.0065 | 1.0077 | 0.9394 |
| INSRR | 1.0548 | 1.0796 | 1.0998 | 0.9605 | 0.9386 | 0.8989 | 1.0013 | 0.9597 | 0.8988 | 0.9994 | 1.0551 | 0.9801 | 0.9613 | 0.9493 | 0.9402 | 1.0283 | 1.0163 | 1.0545 | 1.0730 | 1.0195 | 0.9981 | 1.0599 | 1.0328 | 1.0357 |
| PDGFRA | 1.0512 | 0.7518 | 0.7564 | 1.0671 | 0.8375 | 0.9491 | 1.0203 | 0.9282 | 0.9173 | 0.9916 | 1.1977 | 1.1892 | 1.0058 | 1.0421 | 1.0301 | 1.0008 | 0.8285 | 0.8379 | 0.9986 | 1.1058 | 0.9983 | 0.9876 | 0.9050 | 0.9026 |
| PDGFRB | 0.8224 | 0.5814 | 0.4961 | 0.8359 | 0.5965 | 0.6352 | 0.8857 | 0.7530 | 0.8045 | 1.2078 | 1.5339 | 1.4694 | 1.0521 | 1.0224 | 0.9922 | 0.8402 | 0.6813 | 0.7168 | 0.9752 | 0.9022 | 0.8316 | 0.9814 | 0.8127 | 0.8672 |
| KIT | 1.0888 | 1.1248 | 1.0287 | 1.1359 | 0.9616 | 1.2331 | 1.1406 | 1.1123 | 1.0714 | 0.9062 | 0.9346 | 0.8777 | 1.0421 | 1.0371 | 1.0483 | 1.0207 | 1.0094 | 1.0321 | 0.9723 | 1.0059 | 1.0169 | 0.8376 | 0.9950 | 0.9911 |
| CSF1R | 0.7281 | 0.5938 | 0.6548 | 0.7084 | 0.6336 | 0.6770 | 0.8326 | 0.8357 | 0.8712 | 1.2726 | 1.2390 | 1.2165 | 1.0526 | 1.0164 | 1.0118 | 0.7119 | 0.7596 | 0.7863 | 0.9906 | 0.9738 | 0.9131 | 1.0102 | 0.8986 | 0.8793 |
| FLT3 | 0.8693 | 0.6607 | 0.7900 | 0.8949 | 0.7393 | 0.7729 | 0.9397 | 0.8007 | 0.7920 | 1.0137 | 1.1330 | 1.1419 | 1.0309 | 1.0329 | 1.0504 | 0.9533 | 0.8323 | 0.8093 | 1.0828 | 0.9852 | 1.0096 | 0.9674 | 0.9021 | 0.8699 |
| FLT1 | 1.0947 | 1.0529 | 1.1184 | 0.9821 | 0.9732 | 0.9884 | 0.9601 | 0.8987 | 0.9947 | 1.0374 | 1.0102 | 1.0558 | 1.1092 | 1.0730 | 1.0809 | 0.8697 | 0.9177 | 0.8695 | 1.0158 | 1.0273 | 1.0841 | 0.9035 | 0.9052 | 1.0070 |
| KDR | 0.9538 | 0.8212 | 0.8602 | 0.9410 | 0.8251 | 0.7670 | 0.9294 | 0.8029 | 0.7791 | 1.0221 | 1.1611 | 1.1569 | 1.0872 | 1.1297 | 1.1293 | 0.9083 | 0.7792 | 0.7772 | 1.0461 | 1.0523 | 1.0428 | 0.9630 | 1.0271 | 0.9396 |
| FLT4 | 1.0643 | 1.0403 | 1.0544 | 0.9719 | 0.9834 | 0.9219 | 0.9928 | 0.9569 | 0.9569 | 0.9942 | 1.0237 | 1.0213 | 0.9922 | 0.9825 | 1.0269 | 1.0137 | 0.9970 | 0.9455 | 1.0165 | 1.0435 | 1.0614 | 0.8626 | 0.9088 | 0.9295 |
| FGFR1 | 0.8186 | 0.8232 | 0.7830 | 0.7831 | 0.7583 | 0.6733 | 0.8442 | 0.7906 | 0.7979 | 1.2228 | 1.2497 | 1.2482 | 1.0412 | 1.0252 | 1.0054 | 0.7711 | 0.7521 | 0.7671 | 1.0103 | 0.9565 | 0.8869 | 0.9889 | 0.9887 | 0.9729 |
| FGFR2 | 0.8272 | 0.7968 | 0.9486 | 0.7900 | 0.8454 | 0.8770 | 0.7914 | 0.8209 | 0.8591 | 1.2256 | 1.2203 | 1.1896 | 1.0089 | 1.0308 | 1.0268 | 0.8138 | 0.7839 | 0.8022 | 0.9943 | 0.9774 | 0.9577 | 0.9402 | 0.9107 | 0.9163 |
| FGFR3 | 0.8798 | 0.7346 | 0.8180 | 0.8888 | 0.7789 | 0.7488 | 0.9574 | 0.8308 | 0.8378 | 1.1549 | 1.2716 | 1.2440 | 1.0637 | 1.0539 | 1.0566 | 0.8400 | 0.7612 | 0.7674 | 0.9257 | 0.8966 | 0.8901 | 1.1020 | 1.0169 | 1.0323 |
| FGFR4 | 0.8767 | 0.8290 | 0.8591 | 0.9228 | 0.8502 | 0.8481 | 0.9102 | 0.9348 | 0.9555 | 1.1014 | 1.1022 | 1.1321 | 0.9933 | 1.0081 | 1.0011 | 0.9083 | 0.8855 | 0.8652 | 0.9264 | 0.9349 | 0.9342 | 0.9637 | 0.9327 | 0.8951 |
| PTK7 | 1.0105 | 1.0410 | 0.9008 | 1.1301 | 1.1759 | 0.9837 | 1.0946 | 1.0403 | 1.0198 | 1.0427 | 1.0486 | 1.0601 | 0.9670 | 0.9564 | 0.9340 | 1.0030 | 1.0076 | 1.0213 | 0.9499 | 0.9896 | 1.0182 | 0.9468 | 0.9212 | 0.9442 |
| TRKA | 1.0193 | 0.9890 | 0.9738 | 0.9036 | 0.8082 | 0.8554 | 0.9468 | 0.8653 | 0.9085 | 0.9911 | 1.0495 | 1.0537 | 0.9910 | 0.9664 | 0.9893 | 1.0095 | 0.9808 | 0.9457 | 1.0136 | 1.0729 | 1.1509 | 1.0222 | 1.0753 | 1.0313 |
| TRKB | 0.8473 | 0.8060 | 0.8308 | 0.8683 | 0.8754 | 0.8864 | 0.9562 | 0.9001 | 0.8390 | 1.0670 | 1.0987 | 1.1165 | 1.0258 | 1.0225 | 0.9948 | 0.9215 | 0.8971 | 0.9089 | 1.0665 | 1.0143 | 1.0451 | 1.0971 | 1.0366 | 1.0130 |
| TRKC | 0.9395 | 0.8575 | 0.8686 | 0.9910 | 0.8606 | 0.9311 | 0.9415 | 0.8644 | 0.9083 | 1.0418 | 1.1291 | 1.0953 | 1.0279 | 1.0265 | 1.0408 | 0.9440 | 0.8827 | 0.8904 | 1.0531 | 0.9918 | 1.0140 | 0.9717 | 0.9625 | 0.9187 |
| ROR1 | 1.0554 | 1.0537 | 1.1870 | 1.0457 | 1.0938 | 1.0485 | 0.9843 | 1.0113 | 0.9435 | 0.9735 | 0.9652 | 0.9704 | 0.9650 | 0.9798 | 1.0089 | 1.0642 | 1.0605 | 1.0242 | 0.9349 | 0.8974 | 0.9598 | 0.8835 | 0.9399 | 1.0448 |
| ROR2 | 1.0270 | 1.0775 | 1.1004 | 0.9685 | 1.0099 | 0.9912 | 0.9753 | 0.9874 | 0.9875 | 1.0111 | 1.0145 | 1.0513 | 0.9982 | 0.9993 | 0.9740 | 0.9964 | 0.9944 | 1.0001 | 1.0281 | 0.9532 | 1.0420 | 0.9807 | 0.9961 | 0.9775 |
| MUSK | 1.0173 | 0.9909 | 0.8627 | 1.0001 | 0.9045 | 1.0259 | 0.9452 | 0.9219 | 0.8840 | 0.9799 | 0.9532 | 0.9780 | 1.0029 | 1.0290 | 1.0043 | 1.0129 | 1.0100 | 1.0126 | 0.9969 | 1.1286 | 0.9688 | 0.9467 | 0.8231 | 0.8577 |
| MET | 0.8769 | 0.5927 | 0.6756 | 0.7665 | 0.5445 | 0.5546 | 0.8918 | 0.6560 | 0.6376 | 1.3299 | 1.7265 | 1.6554 | 1.0832 | 1.1338 | 1.1161 | 0.7957 | 0.5559 | 0.5717 | 0.9875 | 0.9525 | 0.9183 | 0.9171 | 0.7713 | 0.7243 |
| RON | 1.0008 | 1.0752 | 1.0446 | 0.9636 | 0.9087 | 0.8829 | 0.9900 | 0.9286 | 0.9978 | 1.0358 | 1.0370 | 1.0214 | 1.0054 | 1.0013 | 1.0145 | 0.9840 | 0.9870 | 0.9812 | 1.0218 | 1.0534 | 1.0691 | 1.0479 | 1.0055 | 1.0614 |
| AXL | 0.9836 | 0.9264 | 0.9435 | 1.0001 | 0.8030 | 0.7799 | 0.9554 | 0.9556 | 0.9239 | 1.0508 | 1.1123 | 1.0620 | 1.0422 | 1.0087 | 1.0312 | 0.9208 | 0.9035 | 0.9197 | 0.9651 | 1.0340 | 0.9637 | 0.8666 | 0.9327 | 0.8762 |
| TYRO3 | 1.0674 | 0.9044 | 0.8943 | 1.0493 | 1.0049 | 0.9425 | 0.9660 | 0.9443 | 0.9983 | 1.0284 | 1.0463 | 1.1406 | 1.0244 | 1.0306 | 1.0188 | 0.9634 | 0.9443 | 0.8977 | 1.0548 | 0.9785 | 0.9241 | 1.0822 | 0.9460 | 0.9175 |
| TIE1 | 1.1872 | 1.1512 | 1.0841 | 1.1039 | 1.0837 | 1.0987 | 0.9886 | 0.9803 | 0.9763 | 0.9466 | 0.9326 | 0.9222 | 1.0343 | 1.0616 | 1.0536 | 1.0032 | 0.9944 | 1.0063 | 0.9579 | 1.0448 | 0.9783 | 0.9860 | 1.0104 | 1.0001 |
| TIE2 | 0.7518 | 0.5596 | 0.5989 | 0.8259 | 0.5577 | 0.6453 | 0.8784 | 0.6516 | 0.6047 | 1.2646 | 1.6528 | 1.6343 | 1.0342 | 1.0995 | 1.1109 | 0.7997 | 0.4616 | 0.4372 | 0.8718 | 0.7085 | 0.6679 | 0.8203 | 0.5746 | 0.5137 |
| EPHA1 | 0.8749 | 0.8690 | 0.7489 | 0.9336 | 0.7958 | 0.8373 | 0.9562 | 0.8566 | 0.8398 | 1.0212 | 1.0964 | 1.1656 | 1.0326 | 1.0479 | 1.0465 | 0.9424 | 0.8493 | 0.7938 | 0.9753 | 0.9437 | 0.9510 | 0.9957 | 0.9707 | 0.9860 |
| EPHA2 | 1.1233 | 0.8737 | 0.9620 | 1.0384 | 0.8492 | 0.8546 | 0.9813 | 0.9077 | 0.8189 | 1.0480 | 1.1718 | 1.2075 | 1.0602 | 1.0865 | 1.0639 | 0.9088 | 0.7924 | 0.7792 | 0.9625 | 1.0176 | 0.9938 | 0.9553 | 1.0799 | 0.9655 |
| EPHA3 | 1.1001 | 1.0354 | 0.9443 | 0.9371 | 0.9405 | 0.9530 | 1.0252 | 0.9741 | 0.9953 | 1.0486 | 1.0762 | 1.0522 | 0.9932 | 1.0047 | 0.9994 | 0.9731 | 0.9437 | 0.9645 | 0.8589 | 0.8979 | 0.9235 | 1.0355 | 0.9307 | 0.9674 |
| EPHA4 | 0.8231 | 0.8447 | 0.8070 | 0.9639 | 0.8850 | 0.9912 | 0.9186 | 0.8896 | 0.9251 | 1.0167 | 1.1141 | 1.1058 | 1.0024 | 1.0205 | 1.0391 | 0.9879 | 0.9031 | 0.8884 | 1.0922 | 0.9874 | 1.0773 | 0.9426 | 0.9501 | 0.9598 |
| EPHA6 | 1.1256 | 1.0430 | 1.2009 | 0.9926 | 1.1017 | 1.0983 | 0.9453 | 0.9646 | 0.9866 | 0.9664 | 0.9961 | 0.9431 | 1.0055 | 1.0066 | 1.0222 | 1.0368 | 0.9952 | 1.0442 | 0.9277 | 1.0098 | 1.0500 | 1.0172 | 1.0060 | 0.9705 |
| EPHA7 | 0.9689 | 0.9596 | 0.9404 | 0.9399 | 0.9395 | 0.9382 | 1.0146 | 0.9933 | 0.9433 | 1.0326 | 1.0413 | 1.0341 | 0.9687 | 0.9771 | 0.9640 | 1.0078 | 0.9916 | 1.0113 | 1.0136 | 1.0388 | 1.0168 | 1.1139 | 1.1900 | 1.0568 |
| EPHB1 | 1.0899 | 1.0020 | 1.0527 | 1.0737 | 0.9554 | 1.0967 | 1.1349 | 1.0505 | 1.1138 | 0.9786 | 0.9763 | 0.9994 | 0.9928 | 0.9785 | 0.9943 | 1.0254 | 1.0464 | 1.0007 | 0.9158 | 0.9624 | 1.0267 | 1.1086 | 1.2190 | 1.2070 |
| EPHB2 | 0.9476 | 0.9529 | 1.0070 | 1.0530 | 1.0153 | 1.0711 | 1.0258 | 0.9338 | 0.9348 | 1.0109 | 0.9818 | 1.0096 | 1.0365 | 1.0571 | 1.0656 | 0.9622 | 0.9656 | 0.9360 | 1.0206 | 1.0229 | 1.0683 | 0.9679 | 1.1348 | 1.1869 |
| EPHB3 | 0.9848 | 0.9237 | 0.9299 | 0.8443 | 0.8832 | 0.8604 | 1.0081 | 0.9517 | 1.0185 | 1.0206 | 1.1353 | 1.0914 | 1.0169 | 1.0265 | 1.0485 | 0.9717 | 0.8860 | 0.8952 | 0.9668 | 0.9588 | 0.9786 | 0.9033 | 0.9992 | 0.8841 |
| EPHB4 | 0.8932 | 0.9956 | 1.0194 | 1.0640 | 0.9925 | 1.0666 | 0.9778 | 0.9771 | 0.9662 | 0.9853 | 0.9884 | 0.9924 | 0.9945 | 1.0019 | 0.9995 | 1.0241 | 1.0110 | 1.0103 | 0.9685 | 1.0730 | 1.0549 | 0.9733 | 1.1020 | 1.0906 |
| EPHB6 | 0.9087 | 0.9424 | 0.9208 | 0.9820 | 0.9119 | 0.9877 | 0.9946 | 0.9830 | 0.9500 | 0.9650 | 0.9817 | 0.9925 | 0.9934 | 1.0095 | 0.9984 | 1.0464 | 1.0068 | 1.0098 | 0.9792 | 0.9686 | 1.0356 | 0.9962 | 0.9477 | 0.8986 |
| RET | 0.9234 | 1.0011 | 0.9497 | 0.9546 | 1.0127 | 0.9683 | 0.9692 | 0.9598 | 0.9543 | 1.0465 | 1.0560 | 1.0584 | 0.9939 | 1.0054 | 0.9946 | 0.9692 | 0.9452 | 0.9568 | 0.9625 | 0.9404 | 0.9306 | 0.9865 | 0.9459 | 1.0101 |
| DDR1 | 1.0518 | 1.1257 | 1.0924 | 0.9801 | 1.0152 | 1.0623 | 1.0203 | 1.0610 | 1.0599 | 1.0166 | 1.0013 | 0.9485 | 0.9670 | 0.9712 | 0.9741 | 1.0212 | 1.0198 | 1.0400 | 1.0105 | 1.0023 | 1.0834 | 1.0891 | 1.0509 | 1.0206 |
| DDR2 | 0.9826 | 1.0885 | 1.0580 | 0.9270 | 0.9254 | 0.9833 | 0.9847 | 0.9871 | 1.0426 | 0.9578 | 0.9466 | 0.9755 | 0.9921 | 0.9889 | 0.9868 | 1.0218 | 1.0288 | 1.0181 | 0.9875 | 0.9692 | 0.9549 | 0.9828 | 0.9845 | 0.9205 |
| ROS1 | 0.8926 | 1.0409 | 0.9247 | 0.9757 | 1.0814 | 0.9649 | 1.0224 | 0.9921 | 1.0335 | 1.1157 | 1.0227 | 1.0603 | 0.9935 | 1.0059 | 0.9988 | 0.9653 | 0.9882 | 0.9815 | 1.0477 | 1.0055 | 0.9959 | 1.0075 | 1.0185 | 1.0352 |
| LMR1 | 1.0138 | 1.0614 | 0.9192 | 0.9492 | 1.0741 | 0.9502 | 1.0728 | 1.0908 | 1.0089 | 1.0332 | 1.0294 | 0.9527 | 0.9954 | 0.9647 | 1.0311 | 0.9722 | 1.0232 | 0.9965 | 0.9783 | 0.9222 | 0.9730 | 0.9852 | 0.9847 | 1.0179 |
| LMR2 | 0.8979 | 0.9534 | 0.9535 | 1.0050 | 1.0080 | 1.0676 | 1.0057 | 1.0517 | 1.0520 | 1.0660 | 1.0047 | 0.9970 | 0.9472 | 0.9580 | 0.9932 | 0.9970 | 1.0714 | 1.0227 | 0.9028 | 0.9559 | 0.9143 | 0.9672 | 0.9627 | 0.8796 |
| LTK | 1.0005 | 1.0217 | 1.0411 | 1.0355 | 1.0296 | 0.9607 | 0.9938 | 1.0618 | 1.0606 | 1.0260 | 0.9994 | 0.9566 | 1.0358 | 0.9997 | 0.9953 | 0.9371 | 1.0010 | 1.0383 | 0.9240 | 0.9715 | 0.9581 | 0.9778 | 0.9848 | 1.0396 |
| ALK | 0.9684 | 0.9274 | 0.9496 | 1.0427 | 0.9162 | 0.9215 | 1.0633 | 1.0512 | 1.0352 | 1.0360 | 1.0352 | 1.0489 | 0.9662 | 0.9706 | 0.9809 | 1.0073 | 1.0011 | 0.9648 | 0.9813 | 0.9659 | 0.9395 | 1.0264 | 1.0252 | 1.0038 |
| STYK1 | 0.8754 | 0.9912 | 0.9617 | 1.0942 | 1.0044 | 1.0892 | 0.9318 | 1.1063 | 1.0516 | 1.1250 | 1.0474 | 1.0305 | 1.0069 | 0.9918 | 0.9799 | 0.9350 | 0.9868 | 1.0080 | 0.9674 | 0.9546 | 0.9859 | 0.9694 | 0.9590 | 0.9988 |

| rL3(1) | rL3(2) | rL3(3) | rE1(1) | rE1(2) | rE1(3) | rE2(1) | rE2(2) | rE2(3) | rE3(1) | rE3(2) | rE3(3) | rI1(1) | rI1(2) | rI1(3) | rI2(1) | rI2(2) | rI2(3) | rI3(1) | rI3(2) | rI3(3) | rF1(1) | rF1(2) | rF1(3) | rF2(1) |
|---|---|---|---|---|---|---|---|---|---|---|---|---|---|---|---|---|---|---|---|---|---|---|---|---|
| 0.8184 | 1.0216 | 1.0190 | 0.8889 | 0.9919 | 0.9072 | 0.9298 | 1.0000 | 0.9455 | 0.9935 | 1.1555 | 1.1927 | 1.1299 | 1.0777 | 1.0332 | 0.9713 | 0.9071 | 0.8685 | 1.0148 | 0.8924 | 0.8570 | 2.4934 | 1.9167 | 1.5641 | 1.5903 |
| 0.9595 | 1.0424 | 1.0986 | 0.9545 | 0.9762 | 0.9328 | 0.9231 | 0.9956 | 0.8372 | 0.8036 | 0.7776 | 0.8036 | 0.9717 | 1.0093 | 1.0193 | 0.9717 | 1.0093 | 1.0193 | 0.9768 | 0.9597 | 0.9787 | 1.0121 | 1.0279 | 0.9839 | 0.9806 |
| 1.0605 | 1.1964 | 1.1138 | 0.9830 | 1.0004 | 0.9812 | 0.9812 | 0.9427 | 1.0008 | 1.1339 | 1.3853 | 1.1854 | 0.9376 | 0.8935 | 0.9132 | 0.9446 | 0.9112 | 0.9362 | 0.9983 | 0.9540 | 0.9689 | 0.9505 | 0.8674 | 0.8944 | 0.9937 |
| 0.9042 | 0.9660 | 0.9018 | 0.9280 | 1.0526 | 1.0263 | 1.0357 | 0.9174 | 0.8919 | 1.1478 | 1.1159 | 1.1460 | 1.0118 | 1.0385 | 0.9877 | 0.9797 | 0.9728 | 0.9454 | 0.9845 | 0.9774 | 0.9424 | 1.2311 | 1.3347 | 1.3572 | 1.0616 |
| 1.3946 | 1.0975 | 1.3413 | 1.0477 | 1.0227 | 1.0244 | 0.9813 | 0.9784 | 0.8780 | 1.0041 | 0.9769 | 0.9268 | 0.9611 | 0.9891 | 0.9745 | 0.9758 | 0.9431 | 0.9665 | 0.9913 | 0.9554 | 0.9787 | 0.9879 | 1.0398 | 1.0337 | 0.9338 |
| 1.0298 | 1.1947 | 0.8025 | 1.0000 | 1.0000 | 1.0233 | 1.0013 | 0.9756 | 0.9499 | 1.0750 | 1.1622 | 1.0226 | 0.9798 | 0.9753 | 0.9401 | 0.9830 | 0.9486 | 0.9276 | 0.9777 | 0.9359 | 0.9454 | 1.0184 | 1.0226 | 0.9164 | 1.0088 |
| 1.1004 | 1.1073 | 1.1605 | 1.0256 | 0.9825 | 1.0256 | 0.9722 | 0.9144 | 1.0089 | 1.0208 | 1.0000 | 1.1538 | 1.0020 | 0.9796 | 0.9783 | 1.0125 | 0.9798 | 1.0004 | 1.0049 | 0.9896 | 1.0161 | 0.9970 | 0.9856 | 0.9992 | 0.9747 |
| 0.5073 | 0.3221 | 0.4863 | 1.1074 | 1.0163 | 0.9507 | 1.1673 | 0.9897 | 1.5619 | 1.1556 | 1.0862 | 1.1310 | 1.0127 | 1.0615 | 1.0315 | 1.0169 | 0.9294 | 0.9009 | 1.0193 | 1.1158 | 1.1452 | 1.0266 | 1.2572 | 1.2019 | 1.0190 |
| 0.9809 | 0.8027 | 1.0490 | 0.9556 | 0.9318 | 0.8780 | 1.0690 | 0.7990 | 0.7172 | 0.9957 | 0.9788 | 0.9208 | 1.0452 | 1.0061 | 1.0044 | 1.0452 | 1.0061 | 1.0044 | 1.0033 | 0.8349 | 0.8294 | 1.4577 | 1.8491 | 1.6051 | 1.2685 |
| 0.9425 | 0.8851 | 0.9767 | 1.0500 | 1.0250 | 1.0000 | 0.9189 | 0.8180 | 0.9452 | 0.8578 | 1.0960 | 1.0960 | 0.9960 | 0.9500 | 0.9706 | 1.0009 | 0.9819 | 1.0026 | 1.0105 | 0.9737 | 0.9986 | 1.0279 | 0.9892 | 0.9556 | 1.1086 |
| 0.7767 | 0.7681 | 0.7901 | 0.9509 | 0.8702 | 0.9252 | 1.0495 | 0.9371 | 0.9688 | 1.0000 | 1.0263 | 0.9453 | 1.1071 | 1.0534 | 1.0079 | 0.9707 | 0.8628 | 0.8821 | 1.0149 | 0.8906 | 0.8942 | 1.6874 | 1.3659 | 1.2487 | 1.1970 |
| 1.0273 | 0.8805 | 0.8288 | 0.8775 | 0.7944 | 0.9994 | 0.8658 | 1.1786 | 1.1429 | 0.9722 | 1.0270 | 1.1129 | 1.0273 | 1.0597 | 1.0195 | 0.9905 | 0.9322 | 0.9149 | 0.9955 | 0.9621 | 0.8988 | 1.1250 | 1.2370 | 1.0435 | 1.0783 |
| 0.7233 | 1.0166 | 0.6109 | 1.1513 | 1.0951 | 1.0310 | 0.9011 | 0.9037 | 1.1475 | 1.0446 | 0.9250 | 0.9708 | 1.0127 | 0.9641 | 0.9358 | 1.0014 | 0.9462 | 0.9068 | 1.0025 | 0.9876 | 0.9463 | 0.8012 | 0.8329 | 0.8125 | 0.8644 |
| 0.9458 | 0.7045 | 0.7233 | 0.8995 | 0.9269 | 0.9618 | 0.9225 | 0.9972 | 1.0129 | 1.0505 | 0.9510 | 0.9697 | 0.9877 | 0.9751 | 0.9973 | 0.9501 | 0.9242 | 0.9409 | 0.9904 | 0.9694 | 0.9824 | 1.0387 | 1.2546 | 1.2459 | 1.0136 |
| 0.8463 | 0.8844 | 0.9893 | 1.0811 | 1.0556 | 1.0000 | 0.8416 | 0.8416 | 0.7708 | 0.8972 | 1.1176 | 1.0897 | 1.0179 | 0.9880 | 0.9722 | 1.0098 | 0.9915 | 0.9879 | 1.0135 | 0.9894 | 1.0221 | 1.0007 | 1.0574 | 1.1062 | 1.0170 |
| 0.5859 | 0.8228 | 0.7306 | 0.9728 | 0.9502 | 0.9977 | 1.0324 | 1.0588 | 0.9000 | 0.9715 | 0.7900 | 0.9459 | 1.0314 | 1.0038 | 1.0325 | 0.9423 | 0.8960 | 0.9051 | 0.9698 | 0.8919 | 0.9105 | 1.4148 | 1.4707 | 1.4186 | 1.1022 |
| 0.8005 | 0.8546 | 1.0057 | 0.9991 | 0.9991 | 1.0643 | 1.0012 | 1.0035 | 0.9744 | 1.0440 | 0.8590 | 0.9408 | 1.0370 | 0.9651 | 0.9665 | 0.9654 | 0.8996 | 0.9138 | 0.9694 | 0.9247 | 0.9215 | 1.5065 | 1.4270 | 1.4254 | 1.1641 |
| 0.8796 | 1.0094 | 0.8053 | 0.9474 | 0.9425 | 0.9699 | 1.1250 | 1.0625 | 0.8818 | 0.9202 | 0.7917 | 0.8661 | 1.0273 | 1.0251 | 0.9927 | 0.9745 | 0.9351 | 0.9226 | 0.9926 | 0.9365 | 0.9281 | 1.3290 | 1.4359 | 1.3751 | 1.1615 |
| 0.9516 | 0.8112 | 0.7905 | 0.9976 | 0.9788 | 0.9830 | 1.1259 | 1.0667 | 0.9527 | 1.0062 | 1.1604 | 0.8478 | 0.9752 | 0.9858 | 0.9872 | 0.9640 | 0.9479 | 0.9426 | 0.9702 | 0.9592 | 0.9598 | 1.1930 | 1.2350 | 1.2568 | 1.0603 |
| 0.9542 | 0.9349 | 0.6223 | 0.9750 | 1.0385 | 0.8750 | 1.1000 | 1.0302 | 1.0927 | 0.8193 | 0.9221 | 0.9346 | 0.9599 | 1.0142 | 0.9744 | 0.9989 | 1.0135 | 1.0151 | 0.9928 | 1.0048 | 1.0314 | 1.1009 | 1.0994 | 1.0554 | 1.1154 |
| 1.0236 | 1.2558 | 1.1401 | 1.0099 | 1.0000 | 1.0000 | 1.0011 | 0.8571 | 0.8078 | 1.0985 | 1.3358 | 1.0741 | 1.0134 | 1.0319 | 1.0535 | 0.9760 | 0.9945 | 1.0345 | 0.9884 | 1.0352 | 1.0609 | 1.1500 | 1.4184 | 1.4643 | 1.0832 |
| 0.8976 | 1.1250 | 1.0734 | 0.9623 | 1.0270 | 1.0256 | 0.9199 | 1.0570 | 1.0239 | 0.9115 | 1.0213 | 1.1316 | 1.1027 | 1.0732 | 1.0912 | 1.0551 | 1.0352 | 1.0693 | 1.0533 | 1.0328 | 1.0689 | 1.4686 | 1.5430 | 1.5291 | 1.3370 |
| 1.1014 | 1.0406 | 1.1335 | 1.0106 | 0.9800 | 0.9947 | 1.0679 | 0.8915 | 1.0553 | 1.0241 | 0.9729 | 1.1080 | 1.0685 | 1.0826 | 1.0863 | 1.0088 | 0.9804 | 0.9859 | 1.0126 | 1.0066 | 1.0004 | 1.2144 | 1.5842 | 1.5231 | 1.1239 |
| 1.2536 | 1.4262 | 1.4518 | 1.0245 | 1.0930 | 1.1469 | 1.0105 | 1.1287 | 0.8346 | 0.9317 | 0.9332 | 1.0702 | 1.0343 | 1.0294 | 0.9825 | 1.0245 | 1.0459 | 1.0231 | 1.0374 | 1.0592 | 1.0198 | 1.0884 | 1.1339 | 1.0185 | 1.0579 |
| 1.2235 | 0.9726 | 0.9279 | 1.0453 | 1.0671 | 1.0870 | 0.9551 | 1.0400 | 1.0435 | 1.1550 | 1.0488 | 1.0806 | 0.9984 | 1.0136 | 0.9907 | 1.0271 | 1.0037 | 1.0228 | 1.0288 | 0.9904 | 0.9887 | 1.0139 | 1.0317 | 1.0228 | 0.9953 |
| 1.3888 | 1.5742 | 1.0358 | 0.9655 | 1.0000 | 0.9655 | 1.1600 | 0.9744 | 1.3432 | 0.9985 | 1.1183 | 1.1739 | 1.0412 | 0.9714 | 1.0725 | 1.0585 | 1.0308 | 1.0893 | 1.0254 | 1.0058 | 1.0857 | 1.1122 | 1.0815 | 1.1376 | 1.1646 |
| 1.0795 | 0.8147 | 1.0509 | 1.0444 | 0.9649 | 0.9828 | 1.0092 | 1.1529 | 1.0277 | 1.1536 | 1.0056 | 0.8671 | 1.0876 | 1.0921 | 1.1343 | 0.9817 | 0.9112 | 0.9297 | 0.9832 | 0.8822 | 0.8926 | 1.7914 | 2.5730 | 2.5025 | 1.1617 |
| 1.0398 | 1.0484 | 0.9447 | 1.0000 | 0.9994 | 0.9825 | 0.9470 | 0.9091 | 0.9596 | 1.0257 | 1.0063 | 1.2984 | 1.0276 | 1.0034 | 1.0077 | 0.9876 | 1.0021 | 1.0058 | 0.9671 | 0.9745 | 0.9789 | 1.0022 | 1.0084 | 1.1010 | 0.9494 |
| 0.7662 | 1.0790 | 0.9456 | 0.9073 | 0.9375 | 1.0045 | 1.0345 | 0.9606 | 0.9275 | 0.8966 | 0.9655 | 0.8966 | 1.0326 | 1.0516 | 1.0757 | 1.0131 | 1.0240 | 1.0682 | 0.9951 | 0.9860 | 0.9961 | 1.0661 | 1.1808 | 1.2293 | 1.0696 |
| 1.1266 | 0.9050 | 0.8346 | 1.0000 | 1.0000 | 0.9762 | 0.9908 | 1.0270 | 0.9851 | 0.9932 | 1.0233 | 1.0835 | 1.0064 | 1.0881 | 1.1622 | 0.9905 | 0.9952 | 1.0097 | 0.9941 | 0.9885 | 0.9943 | 0.9319 | 1.1165 | 1.2729 | 0.8989 |
| 0.7888 | 0.9355 | 1.0082 | 1.0000 | 1.0000 | 1.0200 | 1.1074 | 0.9909 | 1.0000 | 1.1057 | 1.0370 | 1.1282 | 0.9845 | 0.9626 | 0.9508 | 0.9859 | 0.9618 | 0.9901 | 0.9937 | 0.9747 | 0.9855 | 0.8752 | 0.8064 | 0.8286 | 0.9447 |
| 1.1291 | 0.6702 | 0.6881 | 1.0165 | 0.9857 | 1.0000 | 0.9973 | 0.9807 | 0.9973 | 1.0827 | 0.8421 | 0.6550 | 1.1304 | 1.2224 | 1.1932 | 0.9683 | 0.9651 | 0.9430 | 0.9350 | 0.9199 | 0.9124 | 1.4148 | 2.2177 | 2.1127 | 0.9534 |
| 0.9428 | 0.7822 | 0.6584 | 1.0000 | 1.0000 | 1.0000 | 1.2870 | 0.6593 | 1.0385 | 0.8504 | 0.7827 | 0.8348 | 1.0365 | 1.0130 | 1.0162 | 1.0018 | 0.9412 | 0.9391 | 1.0026 | 0.9502 | 0.9763 | 0.9962 | 1.2008 | 1.1557 | 0.9461 |
| 0.7024 | 0.6036 | 0.5717 | 0.8808 | 0.9655 | 0.9333 | 1.1372 | 0.8789 | 1.0385 | 1.2633 | 1.1818 | 1.1818 | 1.0272 | 1.0390 | 1.0258 | 0.9863 | 0.9874 | 0.9445 | 0.9860 | 1.0085 | 0.9521 | 1.1869 | 1.6813 | 1.5427 | 1.1408 |
| 0.8405 | 0.8657 | 0.9853 | 1.0468 | 0.9617 | 0.9249 | 1.1422 | 0.9932 | 1.1143 | 0.8936 | 1.0213 | 0.9347 | 0.9781 | 0.9518 | 0.9784 | 0.9854 | 0.9860 | 0.9654 | 0.9974 | 0.9848 | 0.9784 | 0.9614 | 0.9590 | 0.9921 | 0.9264 |
| 1.1035 | 0.9512 | 0.9818 | 0.8778 | 0.8919 | 0.9483 | 0.9259 | 0.9338 | 0.9558 | 1.2610 | 0.9567 | 1.1724 | 0.9534 | 1.0328 | 1.0124 | 0.9549 | 0.9910 | 0.9793 | 0.9643 | 1.0000 | 0.9955 | 0.9676 | 1.1550 | 1.1214 | 0.9210 |
| 0.9966 | 0.9429 | 0.9657 | 0.9242 | 0.8673 | 0.9739 | 0.9697 | 1.0065 | 0.8000 | 1.0049 | 1.0156 | 0.9524 | 0.9800 | 0.9481 | 0.9371 | 0.9783 | 0.9548 | 0.9886 | 0.9750 | 0.9881 | 1.0125 | 0.9493 | 0.9484 | 0.9052 | 1.0061 |
| 1.0262 | 0.9770 | 1.0906 | 1.0188 | 1.0172 | 1.0188 | 1.1214 | 0.9804 | 1.0458 | 1.0695 | 1.1653 | 1.0756 | 0.9697 | 0.9669 | 0.9775 | 0.9817 | 0.9890 | 0.9606 | 0.9815 | 0.9969 | 0.9582 | 1.0996 | 1.1644 | 1.0723 | 1.0773 |
| 0.9191 | 0.9514 | 0.7219 | 1.0671 | 1.0645 | 1.0671 | 1.0260 | 1.0625 | 1.0282 | 1.1481 | 1.0774 | 1.1852 | 0.9960 | 0.9831 | 0.9727 | 1.0160 | 0.9919 | 0.9676 | 0.9798 | 0.9857 | 0.9348 | 1.0417 | 1.0823 | 1.0115 | 1.0659 |
| 0.9821 | 1.2607 | 1.0795 | 0.9412 | 0.9027 | 1.0021 | 0.9487 | 1.1026 | 0.8974 | 0.8249 | 0.8593 | 0.7482 | 1.0073 | 0.9699 | 0.9955 | 0.9897 | 0.9430 | 0.9637 | 1.0036 | 0.9454 | 0.9667 | 1.1131 | 1.0303 | 1.0953 | 1.1111 |
| 0.9328 | 0.6710 | 0.6419 | 0.9181 | 0.9727 | 0.9178 | 0.9164 | 0.9960 | 1.0649 | 0.9038 | 0.9800 | 0.7167 | 1.0508 | 1.0556 | 1.0437 | 0.9789 | 0.9866 | 0.9408 | 1.0104 | 1.0130 | 0.9966 | 1.0304 | 1.0237 | 1.0035 | 0.9464 |
| 0.9713 | 0.7882 | 0.8312 | 0.8898 | 0.9091 | 0.9563 | 0.8880 | 0.8413 | 0.9528 | 1.1717 | 0.9509 | 1.0443 | 1.0064 | 1.0556 | 1.0546 | 1.0531 | 1.0490 | 1.0807 | 1.0407 | 1.0549 | 1.0773 | 1.1258 | 1.0586 | 1.0571 | 1.1713 |
| 0.8638 | 0.9540 | 0.9111 | 1.0029 | 0.9744 | 1.0014 | 0.9403 | 0.8627 | 0.8358 | 1.0000 | 1.1667 | 1.2500 | 0.9910 | 1.0038 | 0.9869 | 0.9937 | 1.0000 | 1.0070 | 0.9854 | 0.9764 | 0.9829 | 0.9989 | 1.0283 | 0.9016 | 0.9943 |
| 1.0409 | 1.0735 | 0.9621 | 1.0378 | 1.0392 | 1.0378 | 0.9618 | 0.9257 | 0.9357 | 0.9757 | 0.8713 | 1.0912 | 1.0129 | 1.0044 | 1.0071 | 1.0198 | 1.0142 | 0.9993 | 1.0173 | 1.0306 | 1.0142 | 1.1308 | 1.1344 | 1.1420 | 1.0730 |
| 1.0896 | 1.0687 | 1.0505 | 1.0000 | 1.0238 | 0.9773 | 0.9756 | 1.0470 | 1.0732 | 0.9655 | 0.8089 | 0.8485 | 1.0138 | 0.9549 | 0.9444 | 1.0276 | 0.9859 | 0.9700 | 1.0230 | 1.0033 | 0.9762 | 1.0675 | 0.9267 | 0.8448 | 1.0361 |
| 1.0130 | 0.8911 | 0.9360 | 0.9773 | 1.0005 | 0.9773 | 0.9762 | 0.9994 | 0.9286 | 1.0020 | 1.1338 | 0.8485 | 1.0473 | 0.9895 | 1.0074 | 1.0183 | 1.0214 | 1.0267 | 0.9975 | 0.9892 | 1.0041 | 1.0648 | 0.9560 | 0.9905 | 1.0398 |
| 0.9819 | 1.1034 | 0.9980 | 1.0364 | 1.0000 | 1.0000 | 1.0139 | 1.0689 | 0.9636 | 0.8300 | 1.0253 | 0.9697 | 0.9427 | 0.9807 | 0.9666 | 0.9284 | 0.9695 | 0.9386 | 0.9455 | 0.9730 | 0.9267 | 0.9941 | 0.9983 | 0.9676 | 0.8759 |
| 1.1021 | 1.0944 | 1.1370 | 1.0000 | 1.0244 | 1.0244 | 0.9988 | 1.0000 | 0.9524 | 0.7234 | 0.7004 | 0.6911 | 0.9998 | 0.9160 | 0.9388 | 0.9556 | 0.9265 | 0.9561 | 0.9684 | 0.9643 | 0.9921 | 0.9546 | 0.9539 | 1.0111 | 0.9278 |
| 1.1312 | 1.2310 | 1.0991 | 1.0191 | 1.0386 | 1.0035 | 0.9087 | 0.8094 | 0.9340 | 0.8690 | 0.8323 | 0.8471 | 0.9539 | 1.0022 | 0.9486 | 1.0515 | 1.0268 | 0.9863 | 1.0284 | 1.0202 | 0.9924 | 1.0185 | 1.0338 | 0.9274 | 1.0376 |
| 0.9595 | 1.0268 | 1.0724 | 0.9825 | 0.9825 | 1.0182 | 1.0572 | 0.9828 | 1.0539 | 0.9717 | 0.9717 | 0.8205 | 1.0030 | 1.0124 | 0.9916 | 0.9831 | 0.9973 | 0.9791 | 0.9891 | 0.9926 | 0.9868 | 1.0296 | 1.0169 | 0.9365 | 1.0236 |
| 1.0524 | 0.9349 | 0.9317 | 1.0871 | 0.9836 | 0.9682 | 1.1700 | 1.0357 | 1.0770 | 0.9392 | 0.8137 | 0.9632 | 1.0158 | 1.0136 | 1.0251 | 1.0489 | 1.0194 | 1.0414 | 1.0379 | 1.0164 | 1.0511 | 0.9930 | 0.9973 | 1.0124 | 0.9898 |
| 0.9775 | 0.8678 | 1.3314 | 1.0000 | 1.0256 | 1.0519 | 1.6593 | 1.0415 | 1.3078 | 1.0754 | 0.9986 | 1.0498 | 1.0738 | 1.0168 | 1.0011 | 1.0465 | 1.0111 | 1.0215 | 1.0647 | 1.0107 | 1.0101 | 1.3760 | 1.1972 | 1.2011 | 1.2459 |

| rF2(2) | rF2(3) | rF3(1) | rF3(2) | rF3(3) | rtau12(1) | rtau12(2) | rtau12(3) | rtau13(1) | rtau13(2) | rtau13(3) | rtau21(1) | ra21(2) | rtau21(3) | rtau23(1) | rtau23(2) | rtau23(3) | rtau31(1) | rtau31(2) | rtau31(3) | rtau32(1) | rtau32(2) | rt32(3) |
|---|---|---|---|---|---|---|---|---|---|---|---|---|---|---|---|---|---|---|---|---|---|---|
| 1.2964 | 1.1230 | 0.9618 | 1.0248 | 0.8787 | 1.0824 | 1.0894 | 1.0914 | 1.3111 | 1.2691 | 1.2336 | 0.9665 | 0.9875 | 1.0118 | 1.1948 | 1.1297 | 1.1270 | 0.9170 | 1.0219 | 1.0198 | 0.9232 | 0.9920 | 0.9973 |
| 1.0214 | 0.9719 | 0.9612 | 1.0192 | 0.9895 | 0.9836 | 0.9759 | 1.0023 | 1.0001 | 0.9522 | 0.9970 | 0.9980 | 1.0005 | 0.9964 | 1.0091 | 0.9933 | 0.9867 | 0.9973 | 1.0055 | 1.0037 | 0.9956 | 1.0189 | 1.0092 |
| 0.9849 | 1.0233 | 0.9565 | 1.0681 | 1.0446 | 1.0076 | 0.9848 | 0.9894 | 1.0064 | 0.9729 | 0.9851 | 1.0147 | 1.0244 | 1.0172 | 1.0212 | 1.0121 | 1.0172 | 0.9807 | 1.0123 | 0.9855 | 0.9758 | 1.0030 | 0.9893 |
| 1.1473 | 1.1339 | 0.9129 | 0.9524 | 0.9410 | 1.0278 | 1.0151 | 1.0264 | 1.0889 | 1.0828 | 1.0974 | 0.9910 | 1.0054 | 1.0053 | 1.0341 | 1.0579 | 1.0557 | 0.9644 | 0.9750 | 0.9779 | 0.9676 | 0.9686 | 0.9772 |
| 0.9238 | 0.9667 | 0.8925 | 0.9401 | 0.9106 | 1.0087 | 1.0127 | 1.0007 | 1.0278 | 1.0432 | 1.0333 | 1.0020 | 0.9968 | 1.0086 | 1.0273 | 1.0139 | 1.0304 | 0.9885 | 1.0394 | 1.0148 | 0.9898 | 1.0267 | 1.0104 |
| 1.0321 | 0.9576 | 0.9058 | 0.9343 | 0.8843 | 1.0191 | 1.0196 | 1.0129 | 1.0265 | 1.0301 | 1.0187 | 0.9927 | 1.0045 | 0.9934 | 1.0166 | 1.0108 | 1.0233 | 0.9670 | 0.9974 | 0.9837 | 0.9733 | 0.9953 | 0.9900 |
| 0.8864 | 0.9486 | 1.0603 | 0.9380 | 1.0634 | 0.9980 | 1.0039 | 0.9996 | 0.9743 | 1.0072 | 0.9868 | 0.9853 | 0.9709 | 0.9824 | 0.9744 | 0.9956 | 0.9782 | 0.9868 | 0.9961 | 1.0147 | 1.0085 | 1.0247 | 1.0283 |
| 0.9199 | 0.8761 | 1.0437 | 0.8957 | 0.9042 | 1.0068 | 1.0647 | 1.0845 | 1.0089 | 1.1720 | 1.2108 | 1.0052 | 0.9971 | 1.0182 | 0.9978 | 1.1118 | 1.1084 | 1.0069 | 0.9823 | 1.0047 | 0.9964 | 0.9836 | 0.9899 |
| 1.2360 | 1.0622 | 0.9772 | 0.6967 | 0.6799 | 1.0159 | 1.1443 | 1.1562 | 1.1094 | 1.3913 | 1.4194 | 0.9785 | 0.9723 | 0.9868 | 1.0735 | 1.1696 | 1.1539 | 0.9570 | 0.9661 | 1.0260 | 0.9635 | 0.9635 | 0.9699 |
| 1.0482 | 1.0642 | 1.0555 | 0.9430 | 1.0012 | 0.9741 | 0.9793 | 0.9585 | 0.9734 | 0.9692 | 0.9358 | 1.0274 | 1.0093 | 1.0195 | 0.9892 | 0.9970 | 0.9938 | 1.0045 | 0.9825 | 0.9850 | 0.9809 | 0.9792 | 0.9921 |
| 0.8992 | 0.8600 | 0.8332 | 0.6870 | 0.6455 | 1.0709 | 1.1019 | 1.1118 | 1.2461 | 1.2762 | 1.2814 | 0.9854 | 1.0106 | 1.0247 | 1.1428 | 1.1470 | 1.1329 | 0.9358 | 0.9988 | 1.0251 | 0.9531 | 0.9985 | 1.0140 |
| 0.9944 | 0.8612 | 1.0081 | 0.8194 | 0.6885 | 1.0028 | 1.0760 | 1.0645 | 1.0319 | 1.1824 | 1.1719 | 1.0196 | 1.0353 | 1.0297 | 1.0277 | 1.1083 | 1.1206 | 1.0036 | 1.0146 | 0.9852 | 0.9875 | 0.9834 | 0.9756 |
| 0.8527 | 0.7923 | 0.7651 | 0.8324 | 0.7474 | 0.9906 | 0.9828 | 0.9635 | 1.0122 | 0.9807 | 0.9940 | 1.0147 | 1.0076 | 0.9886 | 1.0625 | 1.0268 | 1.0777 | 0.9298 | 0.9359 | 0.9048 | 0.9403 | 0.9584 | 0.9719 |
| 1.0743 | 1.0645 | 0.8660 | 0.7720 | 0.7162 | 0.9710 | 1.0106 | 1.0059 | 0.9896 | 1.0452 | 1.0440 | 1.0095 | 1.0082 | 1.0031 | 1.0012 | 1.0524 | 1.0521 | 0.9466 | 0.8832 | 0.8842 | 0.9369 | 0.8965 | 0.8941 |
| 1.0342 | 1.1117 | 1.0088 | 1.0151 | 1.0283 | 0.9957 | 1.0158 | 0.9986 | 0.9853 | 1.0183 | 1.0125 | 0.9908 | 0.9892 | 0.9969 | 0.9937 | 0.9992 | 1.0231 | 0.9983 | 1.0006 | 0.9839 | 1.0109 | 1.0111 | 0.9919 |
| 1.0765 | 1.0132 | 0.8269 | 0.7713 | 0.7730 | 1.0406 | 1.0567 | 1.0767 | 1.1187 | 1.1473 | 1.1775 | 0.9661 | 0.9613 | 0.9664 | 1.0700 | 1.0746 | 1.0764 | 0.9116 | 0.9146 | 0.9413 | 0.9332 | 0.9353 | 0.9526 |
| 1.1066 | 1.1233 | 0.9191 | 0.8510 | 0.8720 | 1.0533 | 1.0525 | 1.0454 | 1.1423 | 1.1388 | 1.1254 | 0.9690 | 0.9751 | 0.9893 | 1.0592 | 1.0708 | 1.0601 | 0.9567 | 0.9386 | 0.9482 | 0.9544 | 0.9438 | 0.9452 |
| 1.1032 | 1.1014 | 0.9352 | 0.7876 | 0.7872 | 1.0164 | 1.0473 | 1.0329 | 1.0701 | 1.1482 | 1.1219 | 0.9839 | 0.9713 | 0.9676 | 1.0442 | 1.0763 | 1.0885 | 0.9395 | 0.9268 | 0.9183 | 0.9393 | 0.9295 | 0.9361 |
| 1.0769 | 1.0587 | 0.9655 | 0.9570 | 0.9455 | 1.0341 | 1.0255 | 1.0337 | 1.0824 | 1.0832 | 1.1169 | 0.9855 | 0.9861 | 0.9847 | 1.0432 | 1.0620 | 1.0652 | 0.9963 | 0.9802 | 0.9758 | 1.0016 | 0.9952 | 0.9907 |
| 1.0840 | 1.0676 | 1.2274 | 1.3029 | 1.2800 | 1.0279 | 1.0215 | 1.0213 | 1.0322 | 1.0309 | 1.0338 | 0.9789 | 0.9750 | 0.9684 | 0.9725 | 0.9759 | 0.9677 | 0.9908 | 0.9796 | 0.9891 | 0.9930 | 0.9921 | 1.0044 |
| 1.2446 | 1.3195 | 1.1060 | 1.2937 | 1.3100 | 1.0153 | 1.0288 | 1.0201 | 1.0050 | 1.0349 | 1.0485 | 1.0150 | 0.9962 | 0.9946 | 0.9984 | 0.9853 | 1.0192 | 1.0190 | 1.0169 | 1.0012 | 1.0064 | 1.0018 | 1.0088 |
| 1.3399 | 1.3249 | 1.1981 | 1.1946 | 1.2023 | 1.0023 | 1.0091 | 1.0368 | 1.0325 | 1.0479 | 1.0540 | 0.9992 | 0.9942 | 0.9941 | 1.0098 | 1.0350 | 1.0004 | 0.9809 | 0.9680 | 0.9855 | 0.9562 | 0.9675 | 0.9642 |
| 1.3046 | 1.2931 | 1.0908 | 1.1729 | 1.1249 | 1.0040 | 1.0275 | 1.0130 | 1.0437 | 1.1094 | 1.0872 | 0.9999 | 0.9895 | 0.9964 | 1.0392 | 1.0606 | 1.0663 | 0.9976 | 1.0007 | 0.9989 | 0.9958 | 0.9791 | 0.9798 |
| 1.1201 | 1.0253 | 1.1091 | 1.2306 | 1.0381 | 0.9939 | 1.0048 | 0.9972 | 0.9801 | 0.9903 | 0.9888 | 0.9944 | 1.0144 | 1.0078 | 0.9587 | 0.9602 | 0.9884 | 1.0244 | 1.0339 | 1.0097 | 1.0096 | 1.0018 | 1.0031 |
| 1.0409 | 0.9863 | 1.0045 | 1.0247 | 0.9939 | 0.9995 | 1.0086 | 1.0167 | 1.0160 | 1.0286 | 1.0314 | 0.9969 | 1.0031 | 0.9883 | 1.0040 | 1.0083 | 1.0054 | 1.0073 | 1.0045 | 1.0028 | 0.9961 | 0.9962 | 1.0114 |
| 1.2183 | 1.2340 | 1.1001 | 1.0602 | 1.1215 | 0.9846 | 0.9518 | 0.9832 | 0.9927 | 0.9464 | 0.9826 | 0.9867 | 0.9878 | 1.0036 | 1.0061 | 1.0179 | 0.9988 | 1.0176 | 0.9838 | 1.0175 | 1.0024 | 1.0143 | 1.0070 |
| 1.2540 | 1.2600 | 0.8272 | 0.5672 | 0.5775 | 1.0152 | 1.0914 | 1.0758 | 1.1364 | 1.3625 | 1.3327 | 0.9565 | 0.9351 | 0.9277 | 1.1007 | 1.2140 | 1.2197 | 0.8986 | 0.8157 | 0.8106 | 0.9407 | 0.8477 | 0.8693 |
| 0.9730 | 1.0738 | 0.9070 | 0.9102 | 1.0024 | 1.0079 | 1.0022 | 0.9986 | 1.0306 | 1.0055 | 1.0075 | 1.0052 | 0.9931 | 0.9998 | 1.0123 | 1.0158 | 1.0009 | 0.9997 | 0.9868 | 0.9846 | 0.9963 | 1.0028 | 0.9872 |
| 1.0760 | 1.2119 | 0.9767 | 0.9448 | 0.9943 | 0.9983 | 0.9958 | 0.9987 | 1.0114 | 1.0506 | 1.0065 | 0.9972 | 0.9541 | 0.9744 | 1.0332 | 1.0465 | 1.0190 | 0.9590 | 0.9529 | 0.9583 | 0.9844 | 0.9841 | 0.9605 |
| 0.9732 | 0.9891 | 0.8833 | 0.8950 | 0.8775 | 1.0077 | 1.0041 | 1.0285 | 1.0063 | 1.0346 | 1.0953 | 0.9999 | 0.9994 | 0.9942 | 1.0118 | 1.0289 | 1.0401 | 0.9732 | 0.9874 | 0.9826 | 0.9785 | 0.9816 | 0.9763 |
| 0.8849 | 0.9171 | 0.9534 | 0.8755 | 0.9299 | 0.9785 | 0.9807 | 0.9842 | 0.9850 | 0.9800 | 0.9718 | 1.0212 | 1.0354 | 1.0334 | 1.0026 | 1.0115 | 0.9989 | 1.0009 | 1.0017 | 1.0042 | 0.9898 | 0.9832 | 0.9733 |
| 1.1224 | 1.0713 | 0.7385 | 0.4559 | 0.4129 | 1.0513 | 1.1258 | 1.1281 | 1.1908 | 1.5579 | 1.5815 | 0.9939 | 0.9766 | 0.9876 | 1.1160 | 1.3541 | 1.4123 | 0.9796 | 0.8750 | 0.8691 | 0.9744 | 0.8829 | 0.8859 |
| 1.0299 | 0.9556 | 0.8782 | 0.8312 | 0.7266 | 1.0010 | 1.0123 | 1.0400 | 1.0108 | 1.0706 | 1.1375 | 1.0117 | 1.0034 | 0.9999 | 1.0364 | 1.0589 | 1.0793 | 0.9842 | 0.9587 | 0.9514 | 0.9864 | 0.9608 | 0.9405 |
| 1.4171 | 1.2465 | 0.9786 | 1.0392 | 0.8920 | 0.9923 | 1.0204 | 1.0220 | 1.0260 | 1.1019 | 1.1091 | 0.9987 | 0.9980 | 0.9818 | 1.0461 | 1.0722 | 1.0787 | 0.9704 | 0.9419 | 0.9223 | 0.9796 | 0.9348 | 0.9370 |
| 0.9437 | 0.9310 | 0.9312 | 0.9054 | 0.9124 | 1.0171 | 1.0196 | 1.0140 | 1.0152 | 1.0348 | 1.0330 | 0.9915 | 0.9921 | 0.9927 | 1.0070 | 1.0228 | 1.0157 | 0.9943 | 0.9835 | 0.9977 | 0.9948 | 0.9837 | 0.9962 |
| 0.9671 | 0.9779 | 0.8921 | 0.8547 | 0.8433 | 1.0012 | 1.0013 | 1.0061 | 1.0008 | 1.0524 | 1.0622 | 0.9993 | 0.9801 | 0.9931 | 1.0249 | 1.0496 | 1.0552 | 1.0042 | 0.9709 | 0.9755 | 1.0076 | 0.9784 | 0.9706 |
| 0.9779 | 1.0262 | 1.0370 | 1.0000 | 1.0468 | 0.9820 | 0.9882 | 0.9757 | 0.9513 | 0.9605 | 0.9166 | 1.0046 | 0.9946 | 1.0189 | 0.9740 | 0.9805 | 0.9643 | 0.9957 | 0.9688 | 0.9753 | 0.9955 | 0.9834 | 0.9803 |
| 1.1150 | 1.0202 | 1.1003 | 1.1082 | 1.0216 | 1.0173 | 1.0076 | 1.0195 | 1.0026 | 1.0103 | 1.0238 | 0.9828 | 0.9848 | 0.9820 | 0.9934 | 0.9983 | 0.9967 | 0.9997 | 1.0008 | 1.0174 | 1.0105 | 1.0073 | 1.0304 |
| 1.0698 | 1.0202 | 1.0407 | 1.0966 | 0.9481 | 1.0028 | 0.9873 | 0.9979 | 1.0110 | 0.9884 | 0.9992 | 1.0126 | 0.9968 | 1.0010 | 1.0059 | 0.9857 | 0.9952 | 1.0336 | 1.0332 | 1.0038 | 1.0203 | 1.0194 | 1.0099 |
| 1.0584 | 1.1316 | 1.0552 | 0.9690 | 0.9988 | 0.9974 | 0.9817 | 0.9960 | 0.9991 | 0.9779 | 0.9811 | 1.0014 | 1.0158 | 1.0154 | 1.0161 | 1.0085 | 1.0108 | 0.9881 | 0.9742 | 0.9513 | 0.9776 | 0.9644 | 0.9519 |
| 0.8834 | 0.9045 | 0.9386 | 0.8102 | 0.7777 | 1.0061 | 1.0274 | 1.0122 | 1.0094 | 1.0657 | 1.0611 | 0.9953 | 0.9831 | 0.9952 | 1.0128 | 1.0344 | 1.0532 | 0.9852 | 0.9549 | 0.9717 | 0.9911 | 0.9551 | 0.9712 |
| 1.0560 | 1.0590 | 1.2043 | 1.0726 | 1.0467 | 1.0034 | 1.0166 | 1.0146 | 0.9931 | 1.0054 | 1.0063 | 1.0083 | 1.0197 | 1.0064 | 0.9819 | 0.9937 | 0.9874 | 1.0022 | 1.0024 | 1.0068 | 0.9932 | 0.9956 | 0.9889 |
| 1.0191 | 0.9194 | 1.0486 | 1.0330 | 0.9214 | 1.0018 | 1.0007 | 1.0147 | 0.9733 | 0.9951 | 1.0092 | 1.0169 | 1.0108 | 1.0159 | 0.9743 | 0.9976 | 0.9917 | 1.0137 | 1.0125 | 1.0169 | 1.0009 | 0.9948 | 1.0011 |
| 1.0812 | 1.0617 | 1.0521 | 1.0451 | 1.0672 | 1.0233 | 1.0235 | 1.0185 | 1.0331 | 1.0424 | 1.0395 | 0.9984 | 1.0034 | 0.9953 | 0.9930 | 0.9953 | 1.0020 | 0.9951 | 0.9809 | 0.9923 | 0.9846 | 0.9711 | 0.9808 |
| 0.9276 | 0.8854 | 1.0746 | 0.9913 | 0.9400 | 1.0005 | 0.9935 | 0.9890 | 1.0008 | 0.9825 | 0.9781 | 0.9870 | 0.9812 | 0.9943 | 0.9871 | 0.9824 | 0.9909 | 1.0153 | 0.9974 | 1.0088 | 1.0208 | 1.0101 | 1.0234 |
| 0.9700 | 0.9954 | 1.0804 | 1.0745 | 1.0255 | 0.9974 | 0.9886 | 0.9972 | 0.9850 | 0.9760 | 0.9923 | 1.0016 | 0.9914 | 0.9960 | 0.9826 | 0.9808 | 0.9847 | 1.0080 | 1.0021 | 1.0137 | 0.9957 | 0.9984 | 1.0024 |
| 0.9740 | 0.8772 | 0.8617 | 0.9842 | 0.8784 | 1.0117 | 0.9998 | 1.0017 | 1.0334 | 1.0007 | 1.0240 | 0.9904 | 0.9964 | 0.9920 | 1.0168 | 1.0139 | 1.0139 | 1.0002 | 0.9922 | 0.9994 | 1.0003 | 1.0011 | 1.0016 |
| 0.8849 | 0.9566 | 0.9424 | 0.9281 | 0.9218 | 0.9983 | 1.0173 | 0.9754 | 1.0375 | 1.0424 | 1.0105 | 0.9803 | 0.9890 | 1.0082 | 1.0264 | 0.9925 | 1.0500 | 0.9970 | 1.0253 | 1.0304 | 0.9979 | 1.0132 | 1.0083 |
| 1.0994 | 1.0108 | 1.0949 | 1.1709 | 1.0469 | 0.9917 | 0.9732 | 0.9500 | 0.9998 | 0.9682 | 0.9687 | 0.9434 | 0.9577 | 0.9614 | 0.9708 | 0.9590 | 0.9833 | 0.9748 | 1.0119 | 0.9947 | 1.0071 | 1.0316 | 1.0124 |
| 0.9980 | 0.9571 | 0.9348 | 1.0001 | 0.9935 | 0.9951 | 0.9923 | 0.9932 | 1.0133 | 0.9963 | 0.9846 | 1.0040 | 0.9969 | 1.0077 | 1.0179 | 0.9941 | 0.9798 | 0.9772 | 1.0031 | 1.0150 | 0.9753 | 0.9978 | 1.0015 |
| 0.9710 | 1.0005 | 1.0286 | 0.9741 | 0.9969 | 1.0227 | 1.0204 | 1.0220 | 1.0345 | 1.0354 | 1.0371 | 0.9882 | 0.9900 | 0.9883 | 0.9915 | 0.9986 | 1.0024 | 1.0139 | 1.0217 | 1.0020 | 1.0134 | 1.0162 | 1.0022 |
| 1.1732 | 1.1366 | 1.1213 | 1.1191 | 1.0836 | 1.0337 | 1.0082 | 1.0103 | 1.0713 | 1.0199 | 1.0156 | 0.9909 | 0.9906 | 0.9972 | 1.0075 | 0.9909 | 0.9900 | 0.9896 | 1.0051 | 1.0155 | 0.9800 | 0.9925 | 1.0024 |
