## Supplement Table S3 for "Single-molecule behavior and cell-growth regulation in human RTKs"

| parameter | 1st trial | | 2nd trial | | | reproducibility | | |
|---|---|---|---|---|---|---|---|---|
| | ave1 | sd1 | ave2 | sd2 | unit | ratio | SD | R |
| D1 | 0.00771 | 0.00253 | 0.00779 | 0.00271 | um2/s | 1.00012 | 0.08246 | 0.96481 |
| D2 | 0.035563 | 0.006101 | 0.035548 | 0.006438 | | 1.005388 | 0.071468 | 0.930797 |
| D3 | 0.264232 | 0.023733 | 0.265508 | 0.025126 | | 0.996277 | 0.037995 | 0.914787 |
| P1 | 26.4819 | 5.11357 | 26.4638 | 5.3521 | % | 1.00375 | 0.05345 | 0.96391 |
| P2 | 35.7696 | 3.01192 | 35.6448 | 2.89869 | | 1.00403 | 0.03879 | 0.88678 |
| P3 | 37.3476 | 5.97782 | 37.8914 | 6.2383 | | 0.9983 | 0.04937 | 0.9578 |
| L1 | 0.09456 | 0.01122 | 0.09364 | 0.01092 | um | 1.01123 | 0.05346 | 0.89873 |
| L2 | 0.328279 | 0.041235 | 0.322736 | 0.042169 | | 1.02112 | 0.07996 | 0.83711 |
| L3 | 1.309 | 0.14043 | 1.298 | 0.17928 | | 1.02832 | 0.19258 | 0.16006 |
| E1 | 0.02168 | 0.0314 | 0.02173 | 0.02851 | % | 1.399869 | 1.873224 | 0.113154 |
| E2 | 0.13483 | 0.08298 | 0.14371 | 0.07307 | | 1.006264 | 0.674049 | 0.680238 |
| E3 | 0.84208 | 0.08296 | 0.84338 | 0.11303 | | 1.003064 | 0.086453 | 0.66628 |
| I1 | 2.82491 | 0.388599 | 2.82245 | 0.386826 | au | 1.002076 | 0.053916 | 0.91463 |
| I2 | 2.759517 | 0.407539 | 2.763786 | 0.38589 | | 0.999345 | 0.070106 | 0.88638 |
| I3 | 2.423231 | 0.315163 | 2.410859 | 0.30978 | | 1.007187 | 0.068823 | 0.85419 |
| F1 | 0.25313 | 0.09026 | 0.25228 | 0.08899 | au | 1.01018 | 0.126187 | 0.95069 |
| F2 | 0.35476 | 0.11252 | 0.35338 | 0.11297 | | 1.1091 | 0.20418 | 0.83382 |
| F3 | 0.30807 | 0.09756 | 0.30563 | 0.09454 | | 1.01258 | 0.12766 | 0.95357 |
| tau21 | 0.412924 | 0.017335 | 0.412147 | 0.016463 | s | 1.001865 | 0.010827 | 0.96645 |
| tau31 | 0.500567 | 0.03842 | 0.500827 | 0.037186 | | 0.999924 | 0.036802 | 0.87681 |
| tau12 | 0.376178 | 0.019148 | 0.376354 | 0.019845 | | 0.999861 | 0.022419 | 0.90776 |
| tau32 | 0.34412 | 0.024622 | 0.344406 | 0.024106 | | 0.999522 | 0.031856 | 0.89618 |
| tau13 | 0.444141 | 0.047915 | 0.443889 | 0.049595 | | 1.001676 | 0.039386 | 0.93097 |
| tau23 | 0.33927 | 0.021438 | 0.338355 | 0.021018 | | 1.002973 | 0.026227 | 0.90562 |

| | |
|---|---|
| ave | average across 52 RTKs |
| sd | standard deviation across 52 RTKs |
| ratio | average of the ratio of two trials |
| SD | sd of ratio |
| R | R between the 1t and 2nd trials |
