## Supplement Table S4 for "Single-molecule behavior and cell-growth regulation in human RTKs"

| Cell line | ALK | AXL | CSF1R | DDR1 | DDR2 | EGFR | EPHA1 | EPHA2 | EPHA3 | EPHA4 | EPHA6 | EPHA7 | EPHB1 | EPHB2 | EPHB3 | EPHB4 | EPHB6 | ERBB2 | ERBB3 | ERBB4 | FGFR1 | FGFR2 | FGFR3 | FGFR4 | FLT1 |
|---|---|---|---|---|---|---|---|---|---|---|---|---|---|---|---|---|---|---|---|---|---|---|---|---|---|
| NIHOVCAR3 | -0.02129 | 0.181169 | -0.19039 | 0.09667 | -0.30646 | 0.02066 | 0.087502 | -0.0021 | -0.04822 | -0.04315 | 0.09182 | 0.19567 | 0.02918 | 0.15565 | -0.11104 | 0.014557 | -0.07001 | -0.50935 | -0.06365 | -0.14829 | -0.00838 | -0.20339 | 0.19874 | 0.03249 | 0.18521 |
| HEL | -0.16529 | -0.02687 | -0.01161 | 0.04493 | -0.01682 | -0.01211 | -0.02057 | 0.05011 | 0.05238 | 0.10583 | 0.10213 | 0.17142 | -0.01442 | -0.06521 | 0.036752 | 0.113381 | 0.047537 | -0.39955 | -0.07324 | -0.25216 | 0.039357 | -0.05572 | 0.0088 | -0.05538 | 0.36701 |
| HEL9217 | -0.06352 | -0.06452 | 0.11887 | 0.2506 | 0.18066 | -0.02174 | -0.13516 | -0.21491 | -0.09621 | 0.22213 | -0.07148 | -0.02895 | -0.19455 | 0.047864 | 0.2717 | -0.09843 | 0.065367 | -0.4919 | -0.08907 | -0.29042 | 0.055654 | 0.10331 | 0.10566 | 0.09006 | -0.12714 |
| LS513 | 0.097097 | -0.04203 | 0.13788 | 0.11452 | -0.04199 | -0.48925 | 0.035592 | 0.09087 | -0.06163 | -0.27409 | 0.11037 | 0.07721 | -0.16219 | 0.191818 | -0.05632 | 0.237166 | -0.08466 | -0.71154 | -0.10871 | -0.0811 | 0.061829 | 0.04018 | 0.08908 | -0.00914 | 0.3352 |
| C2BBE1 | 0.014821 | 0.023458 | -0.02027 | -0.15255 | -0.02392 | -1.18348 | 0.03139 | 0.1395 | 0.02232 | 0.05143 | 0.13419 | 0.04087 | 0.075591 | 0.051525 | -0.07557 | 0.051399 | 0.071116 | -1.15072 | -0.14228 | -0.12703 | -0.02851 | -0.20241 | 0.09251 | -0.09375 | 0.04502 |
| 253J | 0.102561 | 0.205062 | -0.228 | 0.0068 | 0.04855 | -0.23238 | -0.14979 | 0.09933 | 0.00536 | -0.02742 | 0.12774 | 0.08264 | 0.000562 | 0.058269 | -0.1038 | 0.004682 | 0.141173 | -0.34561 | -0.32657 | -0.11609 | -0.0691 | -0.14996 | -0.02183 | 0.02083 | 0.15694 |
| HCC827 | 0.059558 | 0.051001 | 0.02433 | -0.0576 | 0.04397 | -0.12848 | -0.00089 | -0.23374 | 0.0311 | 0.11711 | 0.18324 | 0.16203 | 0.098247 | 0.027009 | 0.032664 | -0.01686 | 0.281642 | -0.26206 | -0.06379 | -0.19057 | -0.12418 | 0.04553 | 0.04117 | 0.05229 | 0.04537 |
| ONCODG1 | -0.08617 | 0.102732 | -0.00462 | -0.02605 | -0.04214 | -0.17562 | 0.089099 | 0.05119 | 0.17495 | 0.17337 | 0.11008 | 0.05327 | -0.0183 | 0.049777 | -0.1739 | -0.13688 | 0.039382 | -0.324 | -0.26373 | -0.17451 | 0.04207 | -0.00236 | 0.00416 | 0.10304 | 0.16886 |
| HS294T | 0.012794 | -0.21757 | 0.05182 | 0.16189 | -0.04366 | -0.18584 | 0.15999 | 0.04101 | -0.2096 | 0.05791 | 0.011 | -0.00491 | -0.05029 | -0.13657 | 0.040539 | 0.189185 | -0.20643 | -0.32592 | -0.25896 | 0.076724 | -0.07906 | -0.11361 | -0.01723 | 0.0148 | 0.119 |
| NCIH1581 | -0.0587 | -0.0288 | -0.02181 | 0.02793 | -0.28813 | -0.0915 | -0.12819 | 0.02902 | -0.02694 | 0.04464 | 0.18986 | -0.02305 | 0.04028 | 0.108597 | -0.06341 | 0.072369 | 0.219228 | -0.32285 | -0.22836 | -0.05694 | 0.058515 | -0.78504 | 0.1354 | 0.01471 | 0.16112 |
| SKBR3 | -0.11761 | 0.001457 | 0.0199 | -0.05805 | -0.09087 | -0.05123 | -0.06422 | -0.0612 | -0.05395 | 0.03159 | 0.11954 | 0.10526 | 0.11004 | -0.01051 | 0.068845 | 0.081957 | 0.067957 | -1.75958 | -1.25547 | -0.0856 | -0.05371 | 0.03277 | -0.03675 | -0.0791 | 0.11701 |
| T24 | -0.01631 | -0.03364 | 0.01122 | 0.21087 | 0.04489 | 0.01913 | -0.08897 | 0.08891 | 0.06625 | 0.09212 | 0.24405 | 0.04982 | 0.090442 | -0.10606 | -0.01754 | -0.01777 | 0.314487 | -0.23196 | -0.18014 | -0.39212 | -0.06081 | -0.11109 | 0.11206 | 0.10627 | 0.0901 |
| MCF7 | -0.23048 | 0.014416 | 0.00397 | -0.04316 | -0.1234 | -0.22695 | 0.100628 | -0.07272 | -0.05086 | 0.17995 | 0.29261 | 0.07201 | -0.20811 | 0.07726 | 0.116936 | -0.02279 | 0.194032 | -0.28621 | -0.21348 | -0.199 | -0.12939 | -0.1015 | -0.02604 | -0.00403 | -0.13616 |
| NCIH1693 | 0.024642 | 0.042362 | -0.07955 | -0.04757 | -0.15461 | -0.10048 | -0.07565 | 0.22586 | -0.04131 | 0.00793 | 0.04926 | 0.05693 | 0.005363 | 0.030843 | -0.09744 | 0.110813 | 0.056773 | -0.44151 | -0.20385 | -0.07841 | -0.00418 | 0.01333 | 0.09003 | -0.06392 | 0.18159 |
| PATU8988S | -0.09585 | 0.061826 | 0.01007 | -0.06578 | -0.03973 | -0.45387 | -0.02932 | -0.05429 | -0.08295 | 0.08285 | -0.00529 | 0.13468 | -0.03429 | 0.012269 | -0.00995 | -0.11875 | 0.105369 | -0.40202 | -0.21236 | -0.27925 | -0.30063 | -0.01017 | 0.09976 | 0.0196 | 0.22932 |
| PATU8988T | 0.007847 | -0.13657 | 0.0984 | -0.01205 | -0.04726 | -0.27998 | 0.0562 | 0.16206 | 0.0854 | -0.0193 | -0.18038 | 0.01991 | -0.04611 | -0.01837 | -0.02105 | 0.038787 | 0.118446 | -0.05749 | -0.09579 | 0.095017 | -0.18256 | 0.07235 | 0.09405 | -0.07911 | 0.13053 |
| OPM2 | 0.06012 | -0.03191 | 0.00104 | -0.10011 | -0.09704 | 0.03162 | 0.041933 | -0.07462 | 0.12853 | 0.04978 | 0.17905 | 0.09346 | 0.029386 | 0.043386 | -0.10103 | 0.04177 | -0.11307 | -0.33732 | -0.14734 | -0.11668 | 0.065838 | -0.10858 | -0.979 | -0.08989 | 0.16054 |
| CH157MN | 0.088067 | -0.26176 | 0.09632 | 0.09684 | -0.01116 | 0.04297 | 0.076744 | 0.20416 | -0.0783 | 0.07078 | 0.20645 | 0.13412 | -0.04137 | 0.22182 | 0.032886 | -0.08726 | -0.05057 | -0.26984 | 0.02024 | -0.1905 | -0.08208 | -0.01501 | 0.12558 | -0.06932 | 0.22527 |
| KPL1 | 0.05897 | 0.059172 | 0.23556 | -0.13076 | -0.18041 | -0.01851 | -0.10217 | 0.04788 | -0.2869 | 0.06228 | 0.20843 | -0.01959 | 0.089103 | 0.162425 | 0.019861 | 0.102532 | 0.059816 | -0.07475 | -0.2559 | -0.04969 | 0.107674 | 0.00816 | 0.09534 | -0.17486 | 0.14727 |
| HCC827GR5 | 0.059674 | -0.01036 | 0.01637 | -0.04103 | 0.07754 | -0.17509 | -0.03391 | 0.04789 | 0.16853 | -0.01424 | 0.04755 | 0.04928 | -0.07711 | 0.102754 | -0.09706 | 0.097529 | 0.034321 | -0.14858 | -0.28549 | -0.10917 | -0.1323 | 0.08484 | 0.13421 | 0.02558 | 0.06311 |
| PC14 | -0.00086 | -0.02033 | -0.08621 | -0.04353 | -0.0105 | -1.37953 | -0.05786 | -0.13743 | -0.06837 | -0.04172 | 0.11565 | 0.09919 | 0.116589 | 0.096162 | -0.15836 | -0.04107 | 0.01172 | -0.28003 | -0.35093 | -0.052 | -0.14369 | 0.04467 | -0.00051 | 0.0518 | 0.09105 |
| NCIH1650 | 0.060461 | 0.033785 | -0.03942 | 0.01838 | -0.21559 | -0.14213 | -0.04289 | 0.16992 | -0.06938 | 0.11661 | 0.13071 | 0.11613 | 0.020919 | 0.044931 | 0.003214 | 0.031 | 0.117072 | -0.5021 | -0.2652 | 0.143837 | -0.45337 | 0.11184 | -0.02775 | 0.0029 | 0.13964 |
| U343 | -0.08383 | 0.114771 | -0.0239 | -0.04007 | 0.00046 | -0.25923 | 0.123838 | 0.21004 | -0.02296 | 0.08119 | -0.0012 | 0.065 | 0.125368 | 0.012555 | 0.078675 | 0.057159 | 0.297668 | -0.17313 | -0.23218 | -0.08734 | -0.46795 | -0.13761 | 0.0243 | 0.00343 | 0.0323 |
| S117 | -0.11136 | 0.130829 | -0.14035 | -0.00099 | 0.18503 | -0.04588 | -0.17048 | -0.13494 | 0.09497 | 0.06369 | 0.11091 | 0.16108 | 0.053921 | 0.033739 | -0.02452 | -0.15513 | 0.06662 | -0.23347 | -0.41638 | -0.11761 | -0.08383 | 0.10399 | -0.02163 | 0.05662 | 0.07765 |
| SKNMC | -0.11221 | -0.14912 | 0.11324 | 0.04522 | 0.24338 | -0.13727 | 0.090339 | -0.03404 | -0.11362 | 0.11246 | -0.00369 | -0.0781 | -0.31695 | -0.10772 | 0.109086 | 0.028673 | 0.115765 | -0.22512 | -0.07095 | -0.57708 | -0.16819 | -0.00259 | -0.32855 | 0.14649 | -0.1652 |
| U118MG | 0.052567 | -0.10132 | -0.00295 | 0.06581 | 0.07072 | -0.01232 | -0.10273 | -0.08867 | 0.09433 | 0.08706 | -0.02421 | 0.19586 | -0.02486 | 0.053245 | -0.04666 | -0.08233 | 0.038494 | -0.18778 | -0.16378 | -0.16196 | -0.64738 | 0.09424 | -0.02355 | -0.02415 | -0.02757 |
| RDES | -0.07409 | -0.00541 | -0.1574 | -0.07943 | -0.12966 | 0.05521 | -0.10847 | -0.09977 | -0.08238 | 0.13893 | 0.23261 | 0.03581 | -0.03801 | 0.069987 | -0.04745 | 0.163739 | 0.162462 | -0.34711 | -0.31778 | 0.019982 | -0.04802 | -0.25556 | -0.15607 | 0.08482 | -0.03368 |
| PANC0203 | -0.0098 | -0.152 | -0.00359 | -0.14613 | 0.02899 | -0.55521 | -0.01444 | -0.04191 | -0.09377 | -0.03681 | 0.23449 | 0.1076 | -0.11582 | 0.141727 | -0.20835 | 0.070155 | 0.117276 | -0.44447 | -0.164 | 0.063806 | -0.2542 | -0.07583 | 0.02684 | -0.02371 | 0.21128 |
| MV411 | 0.148516 | 0.242031 | -0.28503 | 0.16695 | -0.08642 | -0.0261 | -0.25934 | -0.1404 | -0.10485 | 0.06385 | 0.15237 | 0.02123 | 0.088009 | -0.06331 | 0.004566 | 0.033295 | -0.04251 | -0.1944 | -0.17466 | 0.069148 | -0.02554 | -0.11299 | 0.03688 | -0.09819 | 0.42178 |
| GCIY | 0.042823 | 0.095879 | -0.04866 | -0.03943 | -0.04708 | -0.47713 | 0.133873 | 0.07388 | 0.05449 | 0.01896 | 0.22153 | 0.07181 | 0.061883 | 0.196311 | -0.22008 | -0.16078 | 0.091619 | -0.26305 | -0.2128 | -0.13673 | 0.088873 | -0.08241 | -0.02406 | -0.02358 | 0.17241 |
| TOV112D | -0.10482 | 0.034505 | -0.00777 | -0.08019 | -0.15927 | -0.10304 | 0.018522 | 0.04315 | -0.02096 | 0.02 | 0.11476 | 0.13993 | 0.04334 | -0.00891 | 0.047182 | -0.05775 | 0.191573 | -0.56211 | -0.35647 | -0.16265 | -0.45069 | 0.05865 | -0.06965 | 0.10095 | 0.09729 |
| A673 | 0.008384 | -0.24817 | -0.08146 | -0.01728 | -0.01393 | -0.1894 | -0.40111 | -0.17382 | -0.2856 | -0.07895 | -0.05222 | 0.20781 | 0.287908 | 0.01399 | 0.046092 | 0.077179 | 0.20602 | -0.23876 | -0.21654 | -0.23751 | -0.04041 | -0.02938 | 0.22702 | 0.09031 | 0.11519 |
| KARPAS299 | 0.00737 | -0.00563 | 0.07562 | -0.07988 | 0.0733 | -0.23672 | -0.03426 | -0.16853 | -0.0232 | 0.07417 | 0.11655 | 0.12294 | -0.00056 | 0.021856 | -0.05135 | -0.02759 | 0.124784 | -0.30997 | -0.13586 | -0.20077 | 0.026545 | -0.06338 | -0.10407 | -0.1149 | 0.11129 |
| HT1080 | -0.03049 | 0.027461 | 0.00606 | -0.27794 | -0.10464 | -0.1184 | 0.039088 | -0.02028 | 0.06675 | -0.07908 | 0.19206 | -0.02857 | 0.066257 | 0.055226 | 0.034437 | 0.100912 | 0.101714 | -0.04892 | -0.2097 | -0.08695 | 0.091791 | 0.04993 | 0.09811 | 0.07717 | 0.15823 |
| D283MED | 0.186179 | -0.06701 | -0.01417 | -0.0436 | -0.17748 | -0.05964 | 0.117362 | -0.21705 | -0.00525 | 0.19404 | 0.13959 | 0.00187 | 0.077584 | 0.062788 | -0.03569 | 0.009056 | 0.059802 | -0.08316 | -0.15057 | 0.06203 | 0.003112 | 0.00645 | 0.0598 | 0.21847 | 0.02218 |
| PANC1005 | 0.05832 | -0.37148 | -0.02974 | -0.00273 | -0.04713 | -0.50585 | -0.01248 | 0.06229 | 0.00167 | 0.17786 | 0.14453 | 0.10529 | -0.12403 | 0.107126 | -0.14709 | 0.114976 | 0.030094 | -0.39357 | -0.32107 | 0.064018 | -0.22191 | -0.20022 | -0.03283 | 0.08207 | 0.17954 |
| HS683 | -0.14763 | 0.001234 | 0.04779 | 0.01974 | -0.00547 | -0.12633 | 0.050099 | 0.13348 | -0.37679 | 0.00069 | 0.12678 | 0.1254 | 0.140916 | 0.0575 | -0.02673 | -0.07206 | 0.101087 | -0.251 | -0.28397 | -0.06841 | -0.68972 | -0.00524 | 0.07967 | -0.12553 | 0.00207 |
| 697 | -0.02276 | 0.06798 | 0.10203 | 0.04432 | 0.10632 | -0.10665 | -0.13379 | -0.18816 | 0.10171 | 0.05549 | 0.17249 | 0.16924 | 0.113523 | 0.053529 | -0.0664 | -0.00379 | 0.110279 | -0.29787 | -0.12494 | -0.33172 | 0.043975 | -0.32519 | -0.00258 | 0.06206 | 0.2184 |
| KU812 | 0.040238 | 0.078215 | -0.04998 | -0.08956 | 0.0408 | 0.01134 | 0.006977 | 0.06728 | 0.37172 | 0.30069 | 0.10703 | 0.20481 | 0.010629 | 0.028739 | -0.08177 | 0.126084 | 0.090836 | -0.35378 | -0.2098 | 0.017855 | -0.15761 | 0.01624 | 0.24867 | 0.17979 | -0.05913 |
| U87MG | -0.22183 | -0.07817 | -0.05077 | 0.05305 | -0.02192 | -0.10384 | 0.02828 | -0.12562 | 0.00239 | -0.13492 | 0.25084 | 0.0801 | 0.092433 | 0.119581 | -0.08483 | -0.00252 | 0.049955 | -0.21665 | -0.15641 | -0.19565 | -0.04235 | 0.12368 | -0.0364 | 0.10612 | -0.00913 |
| NCO2 | -0.1232 | 0.149051 | 0.17702 | -0.04997 | -0.04445 | -0.18648 | 0.0197 | 0.00456 | -0.1763 | 0.08821 | 0.16212 | 0.10429 | -0.11076 | -0.02064 | -0.21514 | 0.048846 | -0.03039 | -0.04835 | -0.10392 | -0.29255 | -0.08732 | 0.14918 | 0.07634 | 0.14035 | -0.11609 |
| MJ | -0.03019 | 0.062716 | -0.00405 | 0.16242 | -0.03787 | -0.27272 | -0.33522 | -0.34046 | -0.26142 | -0.10329 | 0.0288 | -0.15004 | -0.23438 | -0.08819 | -0.07508 | 0.06997 | -0.19951 | -0.54254 | -0.36219 | -0.15543 | -0.09437 | -0.02139 | -0.05712 | -0.03582 | 0.63686 |
| MHHNB11 | -0.27665 | -0.0541 | -0.04229 | 0.0507 | -0.03249 | 0.03973 | -0.10226 | -0.01035 | -0.09297 | 0.1235 | 0.14474 | 0.06583 | -0.06971 | 0.150567 | -0.20579 | 0.083518 | 0.201162 | -0.07774 | -0.18352 | -0.13907 | -0.13196 | -0.08685 | -0.03884 | -0.07622 | 0.24081 |
| G292CLONEA141B1 | 0.131697 | -0.04287 | 0.07906 | 0.0104 | 0.03791 | 0.00897 | -0.03317 | -0.07915 | -0.0993 | -0.00294 | 0.17353 | 0.16557 | 0.0071 | 0.027609 | -0.06409 | -0.0504 | 0.004174 | -0.15013 | -0.21763 | -0.12471 | -1.30402 | -0.22892 | -0.17985 | 0.07344 | -0.00178 |
| T3M4 | -0.07992 | -0.04667 | -0.01583 | 0.00193 | 0.20314 | -0.09927 | -0.11223 | -0.04478 | 0.07925 | 0.14759 | 0.0804 | 0.13266 | 0.130079 | -0.08061 | -0.02834 | 0.019004 | -0.05739 | -0.64869 | -0.42604 | -0.26535 | -0.03289 | -0.20587 | 0.01811 | 0.12172 | 0.18581 |
| ACCMESO1 | -0.00364 | 0.01575 | 0.05741 | 0.04157 | -0.09532 | -0.18392 | -0.02125 | 0.17767 | 0.02854 | 0.05736 | 0.06835 | 0.09573 | 0.02464 | 0.139261 | -0.00407 | 0.064444 | 0.100941 | -0.09235 | -0.21566 | -0.1068 | -0.6382 | 0.06322 | -0.0727 | 0.04212 | 0.23879 |
| PC3 | -0.08889 | -0.15964 | 0.10486 | 0.03928 | -0.15645 | -0.21585 | -0.06463 | 0.31719 | -0.15079 | 0.04526 | -0.09735 | 0.10463 | -0.15312 | -0.03747 | -0.04207 | -0.05844 | -0.18416 | -0.24096 | -0.07475 | -0.09143 | -0.03413 | -0.04267 | 0.03827 | -0.09952 | -0.34762 |
| NCIH2452 | 0.098968 | 0.052787 | 0.08345 | -0.09651 | 0.12366 | -0.06849 | -0.14105 | 0.11575 | -0.01604 | 0.18733 | 0.15891 | 0.25452 | -0.03205 | 0.02395 | -0.16631 | -0.04272 | 0.00292 | -0.07549 | -0.23143 | 0.034304 | -0.52161 | -0.24507 | -0.01147 | -0.12638 | -0.02164 |
| PANC0504 | -0.15584 | 0.23362 | -0.08587 | 0.22095 | -0.24592 | -0.78079 | -0.12301 | 0.3107 | -0.23748 | 0.05005 | -0.01459 | -0.03335 | -0.15074 | 0.026795 | -0.07172 | 0.301658 | -0.02181 | -0.54527 | -0.18022 | 0.138546 | 0.046553 | 0.05042 | -0.10887 | -0.01359 | 0.45028 |
| HPAFII | 0.070612 | 0.134942 | 0.16722 | -0.31256 | 0.03417 | -0.31563 | 0.011136 | 0.04646 | 0.13613 | 0.15619 | 0.20635 | 0.1641 | -0.10167 | 0.038929 | -0.12005 | -0.10829 | -0.09523 | -0.4912 | -0.03759 | 0.10017 | -0.08326 | 0.05181 | 0.07946 | 0.05228 | 0.05092 |
| D341Med | -0.00912 | 0.086879 | 0.01707 | -0.017 | 0.02936 | -0.10216 | -0.01549 | -0.11661 | -0.0291 | 0.10616 | 0.14891 | 0.06688 | 0.08384 | 0.035496 | -0.10443 | -0.05204 | -0.02984 | -0.17467 | -0.10748 | 0.031946 | -0.102 | -0.19345 | 0.1742 | 0.12971 | -0.04736 |
| ZR751 | 0.0136 | 0.053569 | 0.02066 | -0.13059 | -0.17226 | -0.34156 | 0.016995 | 0.06753 | -0.03128 | 0.17101 | 0.13481 | -0.0188 | 0.019661 | -0.01904 | -0.10568 | -0.00513 | 0.013553 | -0.6395 | -0.39253 | -0.09482 | -0.09482 | 0.05956 | 0.01195 | -0.0121 | 0.07554 |
| GAMG | 0.03782 | -0.19849 | 0.00451 | -0.07187 | -0.04052 | -0.11257 | 0.162128 | 0.22033 | 0.02555 | 0.10479 | 0.16202 | 0.00726 | 0.000841 | 0.061456 | -0.04516 | -0.01723 | 0.053443 | -0.22266 | -0.23958 | -0.10835 | -1.41363 | 0.06988 | 0.02842 | -0.03739 | -0.07702 |
| SIMA | -0.05292 | 0.20946 | -0.07565 | 0.06744 | -0.019 | -0.05325 | 0.09713 | -0.09496 | -0.00842 | 0.08262 | 0.18446 | 0.11513 | -0.0091 | 0.10304 | -0.14411 | 0.142613 | 0.132561 | -0.14297 | -0.0997 | -0.08991 | -0.42478 | -0.39452 | 0.03102 | -0.17793 | 0.20047 |
| KE37 | 0.030928 | -0.10587 | 0.00846 | -0.01769 | -0.17876 | -0.02948 | -0.1329 | -0.18151 | -0.00307 | 0.14834 | 0.2609 | 0.0372 | -0.02758 | -0.0172 | -0.24754 | -0.1601 | -0.0622 | -0.17681 | -0.10727 | 0.129329 | -0.13014 | 0.00402 | -0.06368 | -0.04968 | 0.29742 |
| CAOV4 | -0.02103 | -0.15734 | 0.1101 | 0.0764 | -0.02281 | -0.05668 | 0.020865 | 0.0165 | 0.15046 | 0.08909 | 0.33045 | 0.07802 | -0.00962 | 0.085102 | -0.38193 | -0.12143 | 0.001513 | -0.32563 | -0.2041 | -0.08417 | -0.07655 | -0.35365 | -0.01587 | -0.32824 | 0.20438 |
| KP3 | 0.028601 | -0.128 | 0.01088 | -0.06585 | -0.04808 | -0.43981 | 0.00437 | 0.1312 | -0.10829 | 0.04722 | 0.01153 | 0.0694 | -0.07741 | -0.00756 | 0.00318 | 0.121224 | 0.084728 | -0.37207 | -0.26086 | -0.14259 | -0.15187 | -0.01796 | -0.09009 | -0.01823 | 0.14713 |
| HCC1187 | -0.09143 | 0.071599 | 0.06142 | -0.22814 | 0.22711 | 0.06601 | 0.138237 | 0.23675 | 0.17917 | 0.22829 | 0.14381 | 0.10534 | -0.01538 | 0.214227 | 0.065885 | 0.034187 | 0.254334 | -0.13184 | -0.05096 | -0.03575 | 0.065283 | -0.057 | -0.07107 | -0.14323 | 0.1352 |
| OCIAML2 | 0.059963 | 0.075115 | -0.06616 | 0.02151 | -0.0965 | 0.00679 | -0.00273 | -0.12132 | 0.01802 | -0.0764 | 0.12475 | 0.08683 | -0.01945 | -0.02922 | -0.17447 | -0.00436 | 0.057178 | -0.12831 | -0.19082 | -0.23636 | 0.057013 | -0.06708 | 0.03677 | -0.12496 | 0.12891 |
| SU8686 | 0.064071 | -0.06097 | -0.05603 | -0.33515 | -0.10746 | -0.33256 | 0.026352 | 0.08748 | -0.10326 | 0.14554 | 0.16429 | 0.07922 | 0.025144 | -0.018 | -0.19176 | 0.100219 | 0.066648 | -0.2743 | -0.13449 | -0.03539 | -0.20029 | -0.06149 | -0.08649 | 0.0995 | 0.15783 |
| VCAP | -0.01647 | 0.040828 | 0.04922 | 0.06913 | 0.16158 | -0.08935 | -0.07968 | -0.07891 | 0.06485 | -0.18173 | 0.08026 | -0.07275 | 0.00111 | -0.00452 | -0.06308 | -0.03636 | 0.231798 | -0.25059 | -0.33048 | -0.22228 | -0.02584 | -0.03012 | -0.14087 | 0.03785 | 0.13871 |
| HUPT3 | -0.0121 | -0.01956 | -0.00601 | 0.08462 | 0.00846 | -0.65261 | 0.050717 | 0.08012 | -0.08193 | 0.03081 | 0.10661 | 0.15329 | 0.015544 | 0.059093 | -0.01014 | -0.02915 | 0.075542 | -0.22837 | -0.13232 | -0.18918 | -0.00984 | -0.19843 | -0.10069 | 0.03358 | 0.13851 |
| CHP212 | -0.02086 | -0.11779 | 0.06282 | -0.06002 | -0.07528 | 0.14452 | -0.21232 | -0.27815 | -0.19786 | 0.05952 | 0.05181 | -0.05285 | -0.00032 | -0.14786 | -0.08115 | 0.026303 | -0.06024 | 0.01923 | -0.08496 | -0.09931 | 0.070964 | -0.07794 | 0.10685 | 0.1379 | 0.5036 |
| COV434 | -0.08128 | 0.117336 | -0.07043 | 0.08659 | -0.02016 | -0.08638 | 0.062823 | -0.07274 | -0.01878 | 0.06961 | 0.0268 | 0.02522 | -0.06957 | 0.176532 | -0.03355 | -0.05986 | 0.105137 | -0.27652 | -0.13537 | -0.37052 | -0.08033 | 0.0467 | -0.08006 | -0.01911 | -0.01529 |
| OCILY19 | -0.04375 | 0.028322 | 0.00402 | -0.18889 | 0.06363 | 0.34189 | -0.05031 | -0.10335 | 0.13641 | -0.01751 | 0.28002 | 0.15255 | -0.02429 | -0.02119 | -0.14997 | -0.04307 | 0.13929 | -0.19321 | -0.3008 | -0.21568 | -0.52743 | -0.04811 | -0.02996 | -0.0736 | 0.04638 |
| SLR20 | 0.002872 | 0.030561 | 0.04358 | 0.02025 | -0.00664 | -0.01808 | -0.04089 | 0.03387 | -0.03069 | -0.02851 | 0.11757 | 0.04124 | 0.044242 | 0.050005 | -0.05291 | -0.03005 | 0.113531 | -0.15001 | -0.18722 | -0.14228 | -0.08671 | 0.06635 | -0.02897 | -0.08301 | 0.17991 |
| LN319 | -0.19668 | 0.020497 | 0.06131 | 0.23453 | -0.02471 | -0.17176 | 0.053325 | -0.11303 | 0.05846 | -0.00257 | -0.16549 | 0.108 | 0.09146 | 0.01295 | 0.048124 | -0.00571 | 0.000103 | -0.84481 | -0.66189 | -0.18895 | 0.02805 | -0.18449 | -0.02614 | 0.15409 | 0.13709 |
| JHOS2 | -0.00071 | -0.25986 | -0.08917 | -0.15563 | -0.01235 | -0.66713 | -0.03551 | 0.10856 | -0.04574 | 0.02628 | 0.0645 | 0.05278 | 0.126291 | 0.159679 | -0.14007 | 0.122163 | 0.055564 | -0.26359 | -0.34321 | -0.02457 | 0.021495 | -0.10956 | 0.0249 | 0.02524 | 0.20987 |
| HS729 | 0.086845 | -0.04258 | 0.09693 | 0.11001 | 0.013 | -0.11226 | -0.17683 | 0.00594 | 0.02682 | -0.00679 | 0.13664 | 0.2097 | -0.00363 | 0.161845 | -0.0788 | 0.105087 | 0.028621 | -0.2074 | -0.16361 | -0.14389 | -1.06663 | -0.04344 | -0.01514 | 0.12274 | -0.004 |
| 8MGBA | -0.06764 | -0.05788 | 0.14832 | 0.07846 | 0.0812 | -0.00089 | 0.108617 | 0.05678 | 0.02566 | -0.00708 | 0.03651 | 0.01826 | -0.04956 | -0.04845 | 0.126185 | -0.15478 | 0.148294 | -0.28194 | -0.13521 | -0.26006 | -0.12924 | -0.09665 | 0.08309 | 0.09291 | -0.07326 |
| CFPAC1 | -0.03747 | 0.037655 | -0.08877 | 0.12839 | 0.05211 | -0.21804 | -0.03246 | 0.13559 | -0.18774 | 0.06908 | 0.12963 | 0.17315 | -0.03511 | 0.053856 | 0.041361 | 0.070934 | 0.074128 | -0.87384 | -0.50136 | -0.32728 | -0.14052 | -0.15825 | 0.02008 | 0.03683 | 0.11542 |
| PANC0327 | 0.253642 | -0.03173 | -0.13758 | -0.08169 | 0.06932 | -0.90227 | 0.111131 | 0.56952 | 0.18154 | 0.14424 | 0.34708 | 0.16845 | 0.114838 | 0.219717 | -0.12598 | -7.7E-05 | 0.182024 | -0.18865 | -0.27795 | 0.26344 | -0.15265 | 0.3111 | -0.20412 | -0.1145 | 0.19876 |
| SNU308 | -0.10242 | 0.019679 | -0.19739 | -0.01299 | -0.1708 | -0.25128 | 0.04604 | 0.10365 | 0.03277 | 0.04578 | 0.04976 | 0.06 | -0.05299 | 0.128185 | 0.245188 | 0.106828 | 0.013382 | -2.43247 | -0.84863 | -0.06669 | -0.02101 | 0.07159 | 0.02918 | -0.20317 | 0.0115 |
| CAL29 | 0.014787 | 0.007073 | -0.06439 | 0.00993 | -0.12635 | -0.53245 | 0.149518 | 0.36004 | -0.01065 | 0.22369 | 0.18793 | 0.1213 | 0.030909 | 0.151211 | -0.18478 | 0.034487 | 0.000354 | -0.78364 | -0.58841 | 0.017941 | 0.079189 | -0.26828 | -0.01833 | -0.19841 | 0.29286 |

| Cell line | ALK | AXL | CSF1R | DDR1 | DDR2 | EGFR | EPHA1 | EPHA2 | EPHA3 | EPHA4 | EPHA6 | EPHA7 | EPHB1 | EPHB2 | EPHB3 | EPHB4 | EPHB6 | ERBB2 | ERBB3 | ERBB4 | FGFR1 | FGFR2 | FGFR3 | FGFR4 | FLT1 |
|---|---|---|---|---|---|---|---|---|---|---|---|---|---|---|---|---|---|---|---|---|---|---|---|---|---|
| RERFGC1B | -0.01203 | 0.210363 | 0.0794 | 0.05354 | -0.12046 | -0.25083 | 0.041222 | -0.13757 | -0.1837 | 0.1182 | 0.41501 | 0.09455 | 0.065957 | 0.063161 | -0.14063 | 0.06216 | -0.15435 | -0.29763 | -0.25596 | -0.26444 | -0.0201 | -0.54652 | -0.21104 | 0.11045 | 0.1581 |
| SKLMS1 | -0.03434 | -0.17141 | 0.09252 | 0.02675 | -0.14905 | -0.14574 | -0.02613 | -0.01771 | -0.0524 | -0.03681 | 0.1167 | 0.11546 | 0.01313 | 0.129416 | -0.26141 | 0.015308 | -0.07347 | -0.38992 | -0.41271 | -0.03472 | -1.01933 | -0.01484 | 0.10421 | 0.00671 | -0.02738 |
| THP1 | -0.07771 | 0.003523 | 0.03679 | -0.1516 | -0.08769 | -0.10089 | 0.136169 | -0.13648 | 0.04037 | -0.10348 | 0.10556 | 0.04598 | 0.096946 | 0.045793 | -0.13348 | -0.09561 | 0.053436 | -0.4171 | -0.11557 | -0.151 | -0.04321 | -0.05538 | 0.05529 | 0.11145 | 0.03894 |
| T47D | 0.025149 | 0.042054 | 0.00837 | -0.02952 | -0.02488 | -0.36436 | -0.03322 | -0.06164 | -0.09203 | -0.01191 | 0.18734 | -0.02471 | -0.03202 | -0.0006 | -0.31619 | 0.071854 | 0.039605 | -0.60841 | -0.22385 | -0.13261 | -0.07902 | -0.01114 | -0.04392 | -0.06532 | 0.24047 |
| HS578T | 0.067609 | 0.039575 | 0.02219 | -0.06625 | -0.26542 | 0.0362 | 0.006865 | -0.00353 | 0.09457 | 0.24987 | 0.16022 | 0.01991 | 0.0044 | 0.175462 | -0.10471 | 0.071823 | 0.049494 | -0.22595 | -0.25722 | 0.094837 | -0.42911 | 0.04053 | 0.11946 | -0.20361 | 0.07399 |
| SKNSH | -0.15649 | 0.147009 | -0.07523 | 0.15822 | -0.1499 | -0.33689 | 0.224095 | -0.23449 | -0.01975 | 0.08412 | 0.19026 | 0.24096 | -0.08199 | 0.018383 | 0.075158 | 0.080525 | 0.329325 | -0.0738 | -0.08638 | -0.00974 | 0.036668 | -0.12053 | -0.06957 | 0.22401 | 0.24875 |
| HCC2935 | 0.027881 | -0.02207 | -0.10541 | -0.22542 | 0.0722 | 0.05803 | -0.17855 | -0.04571 | 0.16434 | 0.04956 | 0.07106 | 0.07103 | 0.015328 | 0.147903 | -0.17802 | 0.143105 | 0.160944 | -0.00336 | -0.26927 | -0.0411 | -0.84973 | -0.1395 | -0.03857 | -0.26062 | 0.24973 |
| JM1 | -0.08057 | 0.075836 | 0.15189 | -0.04715 | -0.03697 | -0.03846 | -0.03576 | -0.19487 | 0.08481 | 0.04312 | 0.06847 | 0.09399 | 0.104329 | -0.01232 | 0.138117 | 0.049365 | 0.170929 | -0.19243 | -0.44546 | -0.11184 | -0.14079 | 0.00963 | 0.17524 | -0.04413 | 0.14126 |
| M059K | 0.151887 | 0.090254 | -0.01668 | 0.1181 | 0.04822 | -0.19485 | 0.094973 | 0.15603 | 0.06695 | 0.11909 | 0.13055 | 0.00534 | 0.000646 | 0.060827 | 0.016838 | -0.04117 | 0.043663 | -0.35429 | -0.25664 | -0.19939 | -0.6478 | -0.17207 | -0.10551 | -0.01209 | 0.07461 |
| NCIH2052 | -0.22676 | 0.033817 | 0.05128 | 0.01535 | -0.0123 | -0.36026 | -0.04353 | 0.11125 | -0.0234 | 0.17451 | 0.16016 | -0.17071 | 0.196984 | -0.00461 | 0.142437 | 0.011827 | 0.275787 | -0.21882 | -0.38636 | -0.13679 | -0.38948 | -0.07277 | -0.15118 | 0.14744 | 0.21863 |
| SW1990 | 0.051731 | 0.120053 | -0.16729 | 0.05403 | 0.02163 | -0.29748 | 0.00715 | 0.2366 | 0.13065 | 0.02329 | 0.22578 | -0.00491 | 0.063562 | -0.12943 | -0.04496 | 0.07442 | 0.139708 | -0.35133 | -0.46286 | 0.023885 | 0.095048 | 0.25508 | -0.00029 | 0.13953 | 0.26254 |
| OSRC2 | 0.010375 | 0.175581 | 0.05948 | -0.05985 | -0.0886 | -0.23491 | 0.008474 | 0.35056 | -0.02896 | 0.0683 | 0.15919 | 0.04251 | 0.001645 | 0.13069 | -0.14276 | 0.043551 | 0.179986 | -0.13504 | -0.25602 | -0.02577 | 0.039129 | -0.00346 | 0.14668 | -0.0629 | 0.08265 |
| BT12 | -0.0426 | 0.130548 | 0.03418 | 0.13082 | -0.16381 | -0.15152 | -0.07746 | 0.0154 | 0.05026 | 0.09546 | 0.16133 | 0.1571 | 0.033558 | 0.077765 | -0.19664 | 0.002647 | -0.01194 | -0.08442 | -0.42257 | -0.03987 | -1.13604 | 0.05852 | -0.11242 | 0.09277 | 0.00725 |
| CORL105 | -0.12854 | -0.0319 | 0.02195 | 0.06767 | -0.18098 | -0.8934 | -0.33223 | -0.12539 | -0.08566 | -0.03292 | 0.24702 | 0.09855 | 0.061536 | -0.00949 | -0.22735 | 0.024938 | 0.053075 | -0.2692 | -0.21744 | -0.1746 | -0.03386 | -0.03082 | 0.17423 | 0.01059 | 0.29567 |
| SW579 | -0.04371 | -0.17435 | 0.24478 | -0.11658 | -0.07694 | -0.14498 | 0.181255 | 0.14342 | 0.00991 | -0.01221 | 0.12786 | 0.18428 | -0.07748 | 0.125987 | -0.07897 | -0.00468 | 0.128283 | -0.14296 | -0.25421 | -0.17457 | -0.82926 | -0.03309 | 0.29334 | 0.07032 | -0.00195 |
| PANC1 | 0.162819 | -0.01101 | 0.00261 | -0.03445 | -0.05049 | -0.10781 | 0.020215 | -0.16018 | -0.10443 | -0.0063 | 0.10489 | 0.09552 | 0.02681 | 0.106632 | -0.20756 | -0.05825 | 0.072654 | -0.3129 | -0.20391 | -0.08211 | -0.009 | -0.08499 | 0.02202 | -0.04822 | 0.10598 |
| NOMO1 | -0.00655 | -0.03307 | -0.05512 | -0.05321 | -0.23807 | 0.22265 | -0.0731 | -0.28457 | -0.02943 | 0.21035 | 0.35933 | 0.06001 | 0.29816 | 0.073825 | -0.34575 | 0.089136 | 0.171333 | -0.48099 | -0.30104 | 0.391038 | -0.01422 | -0.23087 | -0.0406 | -0.24737 | 0.07533 |
| RD | -0.00745 | 0.047538 | 0.12137 | -0.09607 | 0.03526 | -0.0044 | 0.00977 | -0.02329 | -0.06247 | -0.03824 | 0.13745 | 0.04533 | 0.022914 | -0.02485 | 0.0105 | -0.024 | 0.05774 | -0.24715 | -0.17254 | -0.15407 | -0.13819 | -0.13669 | 0.10296 | 0.01517 | 0.17453 |
| CAL62 | -0.04954 | 0.024995 | -0.08868 | -0.21194 | -0.04161 | -0.07776 | 0.063357 | 0.06751 | -0.08742 | 0.06271 | 0.10771 | 0.07272 | 0.026988 | 0.067609 | -0.04319 | 0.0489 | 0.028138 | -0.22043 | -0.12125 | 0.004055 | -0.05668 | -0.03248 | -0.04901 | -0.03148 | 0.13203 |
| LOUNH91 | -0.07599 | -0.08754 | -0.01023 | 0.1207 | -0.11723 | -1.40456 | -0.02589 | -0.05152 | -0.00971 | 0.06947 | 0.22147 | -0.04666 | 0.008087 | 0.018144 | -0.22315 | 0.041173 | -0.21145 | -0.47876 | -0.13812 | -0.18609 | 0.086712 | 0.02328 | -0.07365 | 0.06162 | 0.16899 |
| HS766T | -0.09752 | -0.25392 | -0.08255 | 0.09671 | 0.01132 | -0.26511 | -0.11417 | 0.42363 | -0.11633 | 0.20786 | 0.1299 | -0.08279 | 0.156945 | 0.073858 | -0.11267 | 0.190375 | 0.117122 | -0.22233 | -0.22457 | 0.063381 | 0.125243 | -0.07513 | 0.04628 | -0.08788 | 0.12257 |
| SCC9 | 0.186282 | 0.122178 | 0.06833 | 0.09949 | -0.01844 | -1.63673 | 0.011316 | -0.05108 | -0.2544 | 0.09889 | -0.26686 | 0.1231 | 0.079702 | 0.202802 | -0.01935 | 0.158705 | 0.044033 | -0.36067 | -0.24729 | -0.21217 | -0.12659 | 0.13293 | 0.22228 | 0.17723 | 0.17047 |
| SNU869 | 0.105322 | 0.122781 | -0.04148 | -0.06298 | 0.01324 | -0.22596 | 0.168058 | -0.13569 | 0.00161 | 0.0272 | 0.16902 | 0.04704 | -0.00505 | 0.144784 | -0.13458 | -0.04105 | 0.135415 | -0.1433 | -0.14518 | -0.128 | 0.030001 | -0.22885 | 0.0846 | 0.02429 | 0.08465 |
| L363 | 0.124361 | 0.105523 | 0.09641 | 0.11066 | -0.02704 | -0.25279 | 0.199886 | -0.1626 | 0.2288 | 0.12585 | 0.14357 | 0.1542 | -0.15065 | 0.036037 | -0.00668 | -0.11499 | 0.178955 | -0.35019 | -0.23769 | -0.01896 | -0.1233 | -0.10379 | 0.08962 | -0.02381 | 0.17579 |
| CORL311 | 0.02165 | 0.095059 | 0.01926 | 0.03103 | -0.10909 | -0.10236 | -0.01879 | -0.00189 | 0.00241 | -0.00498 | 0.24141 | -0.04934 | 0.076178 | 0.144527 | 0.080023 | 0.1718 | 0.183694 | -0.29225 | -0.07117 | -0.08424 | -0.05017 | -0.12876 | 0.04011 | -0.00387 | 0.12302 |
| SCC25 | 0.141435 | 0.081915 | -0.16913 | -0.05985 | 0.03203 | -0.58354 | 0.032502 | 0.01308 | 0.0053 | 0.04066 | 0.17369 | 0.04434 | 0.006518 | 0.208261 | -0.02638 | 0.009379 | 0.111624 | -0.36344 | -0.46765 | -0.0411 | -0.25022 | -0.08755 | 0.03495 | -0.16438 | 0.15618 |
| RCC10RGB | -0.112 | -0.20428 | 0.24616 | -0.09533 | 0.04735 | -0.48405 | 0.101976 | 0.08973 | -0.22399 | 0.43481 | 0.0311 | 0.20125 | -0.04751 | -0.02588 | 0.359985 | 0.075367 | 0.391022 | -0.20983 | 0.07044 | -0.32967 | -0.08056 | 0.10327 | -0.21995 | 0.11797 | 0.08787 |
| HDMYZ | -0.05566 | -0.09315 | -0.04622 | -0.00017 | -0.09219 | -0.12739 | -0.00417 | -0.08148 | -0.1016 | -0.11356 | 0.20017 | 0.04013 | -0.18293 | 0.083751 | -0.36478 | 0.100633 | 0.205943 | -0.1094 | -0.14481 | 0.06747 | 0.071988 | 0.17148 | 0.01049 | -0.00904 | 0.35996 |
| BHT101 | 0.078177 | 0.053015 | 0.01671 | 0.0813 | -0.00513 | -0.10231 | -0.00078 | 0.06843 | 0.00937 | 0.09005 | 0.19175 | -0.15784 | -0.07703 | 0.046586 | -0.03566 | 0.239894 | 0.01706 | -0.30353 | -0.25852 | 0.009744 | -0.09499 | 0.08305 | 0.02734 | -0.05849 | 0.29652 |
| MFE280 | 0.060111 | -0.05652 | -0.09003 | 0.24695 | -0.19098 | 0.00877 | 0.004249 | -0.38011 | 0.10767 | 0.19776 | 0.15128 | 0.01766 | 0.085547 | 0.172155 | -0.18428 | -0.19317 | 0.150394 | -0.68416 | -0.88393 | -0.21283 | 0.239895 | -0.01609 | 0.09396 | 0.24084 | 0.09071 |
| SET2 | 0.083313 | -0.21029 | 0.14858 | 0.16774 | 0.14071 | -0.07326 | -0.10336 | -0.16712 | -0.09885 | 0.21512 | 0.02423 | -0.02689 | 0.040052 | -0.04939 | -0.01896 | -0.09689 | -0.0176 | -0.53057 | -0.2938 | 0.102844 | -0.05857 | 0.27438 | 0.24808 | -0.03472 | 0.28167 |
| EOL1 | 0.065977 | -0.08811 | -0.0731 | -0.12271 | -0.0917 | -0.10938 | -0.02771 | -0.2488 | 0.02813 | -0.08296 | 0.1393 | 0.12605 | 0.081784 | -0.02416 | -0.08867 | -0.02037 | -0.02163 | -0.10202 | -0.26834 | -0.12529 | -0.12681 | -0.09821 | -0.09818 | 0.02441 | 0.12805 |
| NMCG1 | 0.093563 | -0.07777 | 0.01639 | 0.07944 | 0.06159 | 0.20794 | -0.14132 | 0.01421 | -0.12199 | 0.08669 | 0.2282 | 0.06766 | 0.111159 | 0.126751 | -0.16687 | 0.186227 | 0.243001 | 0.00679 | -0.15926 | -0.35486 | 0.00634 | -0.06204 | 0.02188 | 0.04034 | 0.02627 |
| COLO320 | -0.26435 | -0.01669 | 0.21743 | 0.07221 | 0.28017 | -0.0178 | 0.271582 | -0.13131 | -0.15188 | 0.20283 | -0.14044 | 0.07901 | 0.034979 | -0.20823 | 0.110615 | -0.36922 | 0.094959 | -0.55648 | -0.07859 | -0.35638 | -0.23089 | -0.10393 | 0.27559 | 0.11452 | -0.17292 |
| LP1 | -0.12939 | 0.130478 | -0.12216 | -0.278 | -0.19624 | 0.0453 | -0.02771 | -0.22814 | -0.06612 | -0.00226 | 0.28971 | 0.12752 | 0.087735 | 0.033515 | -0.00753 | 0.348828 | 0.129642 | -0.04842 | 0.002 | -0.03064 | -0.00291 | -0.13708 | 0.05211 | -0.03833 | 0.22861 |
| C8166 | -0.05843 | 0.092488 | -0.17873 | 0.16209 | 0.0266 | -0.08632 | 0.130804 | -0.14719 | -0.20194 | 6.9E-06 | 0.09311 | -0.09641 | 0.050897 | -0.08936 | 0.127431 | 0.028477 | 0.262972 | -0.33609 | -0.18378 | -0.29825 | 0.051498 | -0.11944 | 0.03291 | 0.01221 | 0.13621 |
| DETROIT562 | 0.072195 | -0.02782 | -0.06468 | -0.09522 | 0.00054 | -0.75869 | -0.03248 | 0.3485 | -0.11439 | 0.02931 | 0.0923 | 0.10451 | 0.07985 | 0.139052 | 0.059809 | 0.099917 | 0.137125 | -0.55698 | -0.54846 | 0.016051 | -0.14715 | -0.10462 | 0.00163 | 0.03468 | -0.0478 |
| SNU1079 | -0.09261 | 0.016513 | 0.08838 | -0.20022 | -0.12831 | -0.13122 | 0.065481 | 0.09372 | -0.1642 | -0.03928 | 0.15086 | 0.11859 | -0.15468 | 0.013496 | 0.115426 | 0.046957 | 0.029117 | -0.50982 | -0.64202 | -0.07131 | -0.14305 | 0.10747 | 0.11617 | -0.30988 | 0.10731 |
| DAOY | 0.099679 | -0.02553 | 0.04741 | -0.09185 | -0.0583 | -0.0875 | 0.056111 | 0.20556 | -0.01318 | 0.19532 | 0.12542 | 0.12877 | 0.027044 | 0.138018 | -0.04951 | 0.15364 | 0.104664 | -0.19578 | -0.23667 | -0.08305 | -0.57231 | -0.02526 | -0.00824 | -0.01649 | 0.19885 |
| CAL120 | -0.02011 | 0.035873 | 0.00746 | -0.02762 | 0.1386 | -0.11464 | 0.120042 | 0.06063 | 0.04915 | 0.02793 | 0.24358 | 0.02793 | -0.02511 | 0.143724 | -0.19974 | 0.09212 | 0.188422 | -0.25946 | -0.31023 | -0.1523 | -1.53258 | -0.00723 | -0.16465 | -0.09357 | 0.37957 |
| HUPT4 | 0.184484 | -0.34369 | 0.16275 | -0.20331 | 0.1078 | -1.12307 | 0.033807 | 0.18988 | -0.47753 | -0.05639 | -0.01196 | 0.00656 | 0.084891 | 0.118899 | -0.10911 | 0.102498 | 0.209843 | -0.73783 | -0.1449 | -0.00102 | -0.46362 | -0.05628 | -0.0788 | 0.09384 | 0.11468 |
| LN382 | 0.049229 | 0.000723 | -0.0041 | -0.10285 | 0.04058 | -0.14315 | 0.001315 | -0.03124 | 0.29676 | 0.1258 | 0.17248 | 0.17096 | 0.140801 | 0.115677 | -0.11817 | -0.01148 | 0.154073 | -0.21082 | -0.31157 | 0.007591 | -0.09467 | -0.04206 | -0.08202 | 0.06932 | 0.11638 |
| JHESOAD1 | 0.056179 | -0.00466 | -0.02676 | -0.02094 | -0.06369 | -0.75741 | -0.02522 | 0.1897 | 0.07792 | 0.04517 | 0.11302 | -0.00416 | 0.012539 | 0.067787 | -0.11313 | -0.06284 | -0.00833 | -0.23705 | -0.20501 | -0.03553 | -0.08564 | -0.04678 | -0.01553 | 0.03326 | 0.10198 |
| JHH6 | 0.019461 | -0.18088 | -0.04337 | -0.08525 | -0.0298 | -0.16385 | 0.104747 | 0.08674 | -0.16479 | -0.04329 | 0.14392 | 0.00851 | 0.015418 | -0.03866 | 0.015955 | -0.0479 | 0.147475 | -0.28651 | -0.19652 | -0.13939 | -0.49945 | 0.04724 | 0.03404 | 0.04818 | -0.00101 |
| A375 | 0.080218 | 0.075734 | -0.01559 | -0.01282 | -0.03232 | -0.10635 | -0.06149 | 0.09519 | 0.0789 | 0.05588 | 0.18037 | 0.04431 | -0.01408 | 0.064062 | -0.02093 | -0.03347 | 0.103109 | -0.1613 | -0.13923 | 0.019302 | 0.065909 | -0.02208 | 0.0785 | -0.02833 | 0.22153 |
| SNU398 | -0.00551 | -0.05484 | 0.02302 | 0.01042 | -0.10537 | -0.08865 | -0.00048 | -0.00162 | 0.03204 | -0.0913 | 0.06197 | 0.01469 | -0.0465 | 0.015831 | -0.14275 | -0.13441 | 0.25192 | -0.27768 | -0.19455 | -0.23994 | -0.16238 | 0.02549 | 0.01526 | -0.20211 | 0.30643 |
| ASPC1 | 0.020344 | -0.03226 | -0.15525 | -0.07316 | 0.01075 | -0.55475 | 0.059431 | -0.07196 | -0.01817 | 0.11415 | 0.08528 | 0.05708 | -0.13364 | -0.06904 | -0.01678 | 0.076901 | 0.018255 | -0.27703 | -0.06286 | -0.07358 | 0.000672 | -0.04159 | 0.0017 | -0.04684 | 0.15203 |
| HCC1937 | -0.05337 | -0.02138 | 0.06105 | -0.03956 | -0.02655 | -0.31146 | -0.01798 | 0.06995 | -0.00081 | 0.1474 | 0.12051 | 0.04769 | 0.011575 | 0.140554 | 0.130224 | -0.0108 | 0.138427 | -0.23371 | -0.2852 | -0.00539 | -0.05128 | -0.1305 | 0.1252 | -0.05518 | -0.01181 |
| SUPM2 | -0.05449 | 0.058755 | 0.00488 | 0.08199 | 0.05823 | -0.13815 | -0.02315 | -0.1139 | -0.10217 | 0.01383 | 0.06322 | -0.0032 | -0.20629 | 0.057921 | -0.02422 | -0.03188 | 0.229802 | -0.11132 | -0.10202 | -0.05737 | -0.02451 | 0.03427 | 0.1583 | 0.03597 | 0.26989 |
| KPNYN | -0.059 | 0.090934 | -0.00865 | 0.16303 | -0.31405 | 0.13051 | -0.02316 | -0.04479 | -0.12529 | 0.28375 | 0.08266 | -0.12617 | 0.04531 | 0.174739 | -0.0793 | 0.00899 | 0.143298 | -0.32882 | -0.06597 | 0.007478 | -0.16784 | -0.07669 | 0.17877 | -0.27261 | 0.13203 |
| BICR31 | 0.135447 | 0.081709 | 0.03284 | 0.03429 | -0.08254 | -0.64887 | 0.051365 | 0.05188 | -0.11689 | 0.03013 | 0.09065 | 0.00735 | 0.002919 | -0.02393 | 0.121755 | 0.05721 | 0.221254 | -0.49072 | -0.27067 | -0.08723 | -0.09338 | -0.04883 | 0.0056 | -0.10233 | 0.03882 |
| KALS1 | 0.074834 | -0.02079 | -0.00103 | 0.05583 | -0.02309 | -0.0912 | 0.009639 | -0.0348 | 0.08668 | 0.01201 | 0.26097 | 0.13111 | 0.062893 | 0.104017 | -0.09917 | -0.14614 | 0.101027 | -0.10603 | -0.17357 | -0.05972 | -0.27794 | -0.1639 | -0.13802 | -0.08547 | 0.15175 |
| U251MG | 0.135552 | -0.17238 | -0.04513 | 0.05698 | -0.17784 | -0.27542 | -0.04244 | 0.62751 | -0.06825 | 0.13438 | 0.16167 | -0.08928 | 0.061753 | 0.084325 | -0.11962 | 0.118022 | 0.01154 | -0.15077 | -0.26159 | 0.001773 | -0.2732 | -0.19056 | 0.13748 | -0.03817 | 0.28233 |
| DEL | 0.077866 | 0.000453 | -0.06037 | -0.05247 | -0.00916 | -0.05096 | -0.04759 | -0.22132 | 0.04423 | -0.03779 | 0.10712 | 0.22062 | 0.061092 | -0.07962 | -0.10007 | 0.101933 | 0.131593 | -0.22106 | -0.1382 | -0.13367 | -0.12811 | 0.01112 | -0.18832 | -0.24697 | 0.25043 |
| CAKI2 | -0.08653 | 0.092457 | 0.09516 | -0.28818 | 0.01098 | -0.50953 | 0.198253 | 0.05113 | -0.03911 | -0.07425 | 0.05766 | 0.12746 | 0.106644 | -0.06945 | 0.202067 | -0.08462 | 0.186578 | -0.50531 | -0.36175 | -0.35414 | -0.1008 | -0.10759 | 0.06735 | 0.11583 | 0.01543 |
| PANC0403 | -0.01505 | -0.09989 | -0.0137 | -0.02272 | 0.02903 | -0.92244 | -0.00333 | -0.06937 | -0.00497 | 0.08216 | 0.02364 | 0.09919 | 0.019299 | 0.092823 | -0.10525 | 0.021975 | 0.033209 | -0.31145 | -0.20753 | -0.0504 | -0.01371 | -0.13382 | 0.01006 | 0.04089 | 0.16492 |
| JHOM1 | 0.087093 | -0.06006 | 0.04688 | 0.03775 | -0.0931 | -0.14664 | -0.16428 | 0.14623 | -0.11857 | 0.08409 | 0.14833 | 0.00505 | -0.0204 | 0.216235 | -0.18681 | 0.04892 | 0.098279 | -0.31126 | -0.14845 | 0.01159 | -0.25633 | 0.09374 | 0.05468 | -0.06685 | 0.15737 |
| SCC4 | -0.1047 | 0.021685 | -0.13815 | -0.08198 | -0.05621 | -0.67606 | -0.02468 | 0.15482 | 0.04082 | 0.16601 | 0.26765 | 0.04338 | 0.024212 | 0.127471 | -0.1737 | 0.225783 | 0.09679 | -0.52709 | -0.3621 | -0.06019 | -0.09856 | 0.03579 | -0.16622 | -0.00914 | 0.25476 |
| DANG | 0.029299 | 0.037557 | 0.02507 | -0.0622 | 0.02181 | -0.08964 | -0.0245 | -0.20723 | -0.09303 | 0.1184 | 0.03799 | 0.08146 | 0.059329 | 0.026094 | -0.07545 | -0.15996 | 0.059207 | -0.64388 | -0.35086 | -0.02023 | -0.05119 | -0.12216 | 0.02971 | 0.04745 | 0.07083 |
| DKMG | -0.16197 | 0.210797 | -0.00121 | -0.24764 | 0.10187 | 0.24818 | 0.187933 | -0.06737 | 0.04666 | 0.09748 | 0.19484 | 0.01731 | 0.16137 | -0.10327 | -0.1953 | -0.00055 | 0.05306 | -0.08222 | -0.26817 | -0.39153 | -0.30456 | -0.14518 | 0.11249 | -0.04452 | 0.00777 |
| SLR23 | 0.042531 | 0.046423 | -0.01417 | -0.03007 | -0.02743 | -0.33562 | 0.001807 | 0.22596 | -0.0381 | 0.15833 | 0.13317 | 0.21285 | 0.056154 | 0.072786 | -0.02157 | 0.056937 | 0.204211 | -0.17768 | -0.33751 | -0.04233 | -0.02812 | 0.03467 | 0.01431 | 0.08574 | 0.04532 |
| OCUM1 | 0.048335 | 0.044991 | -0.1442 | -0.17195 | 0.04256 | -0.20725 | 0.181771 | -0.07284 | -0.08493 | -0.0908 | 0.05363 | -0.08415 | 0.095971 | 0.026598 | 0.107557 | 0.069605 | 0.113295 | -0.16651 | -0.39255 | -0.11904 | -0.04713 | 0.09454 | -0.00244 | 0.20853 | 0.14274 |
| AU565 | 0.018788 | -0.14972 | -0.08524 | -0.01509 | 0.03683 | -0.09133 | -0.03009 | 0.1685 | -0.21903 | 0.20665 | 0.29251 | 0.00652 | 0.108163 | 0.035343 | 0.07482 | 0.186105 | 0.078094 | -2.24301 | -1.88214 | -0.08577 | 0.001547 | -0.04365 | -0.07354 | -0.10251 | -0.00929 |
| CL11 | 0.049362 | 0.167327 | 0.33025 | -0.00612 | -0.14254 | -0.3985 | -0.02613 | 0.28635 | 0.04996 | 0.09258 | 0.0433 | 0.11572 | 0.079669 | -0.11136 | -0.06204 | 0.123583 | 0.13961 | -0.47344 | -0.19937 | 0.160998 | -0.04279 | -0.16792 | 0.20213 | -0.08671 | 0.15379 |
| KMRC20 | 0.08716 | 0.025854 | -0.06404 | -0.22909 | -0.01387 | -0.46991 | 0.048033 | 0.0447 | -0.01354 | 0.13496 | 0.1451 | 0.11122 | -0.00295 | 0.099262 | -0.12105 | 0.054214 | -0.0364 | -0.33456 | -0.26329 | -0.05821 | -0.33837 | -0.05185 | -0.09582 | -0.0448 | 0.18827 |
| NCIH2887 | -0.05928 | -0.09445 | 0.11459 | -0.11841 | -0.43055 | -0.04569 | 0.164297 | 0.01473 | -0.04844 | 0.12361 | 0.25937 | 0.05861 | -0.16496 | 0.199042 | -0.1543 | 0.131643 | -0.26959 | -0.32768 | -0.19613 | 0.068807 | 0.039969 | 0.04076 | 0.00782 | -0.11378 | -0.17933 |
| LS1034 | -0.14694 | -0.03019 | 0.07094 | 0.18129 | -0.18052 | 0.08843 | -0.00409 | -0.12189 | -0.0478 | 0.32741 | 0.22485 | 0.05985 | 0.035821 | 0.142686 | -0.27732 | -0.29062 | 0.110456 | -0.2503 | -0.26512 | 0.001119 | -0.02617 | -0.15713 | -0.04959 | -0.01629 | 0.07015 |
| COLO201 | 0.164703 | 0.049522 | -0.00635 | -0.05223 | 0.06354 | -0.12049 | 0.015765 | -0.10621 | -0.02059 | 0.10115 | 0.04449 | 0.09404 | 0.097667 | 0.127393 | -0.12009 | 0.049073 | 0.02433 | -0.15348 | -0.12503 | -0.13093 | 0.061402 | -0.07969 | -0.12967 | 0.02386 | 0.1167 |
| LMSU | -0.04207 | -0.10571 | -0.10565 | -0.17225 | 0.02142 | -0.14644 | -0.07187 | 0.02338 | 0.01224 | 0.04884 | 0.14404 | 0.03624 | -0.0355 | 0.108426 | -0.19664 | -0.00508 | 0.091106 | -0.08635 | -0.18143 | -0.03404 | -0.86647 | -0.13612 | -0.08197 | 0.04972 | 0.09618 |
| COV318 | -0.02513 | 0.036319 | -0.24662 | 0.02743 | -0.16291 | 0.00351 | -0.20825 | -0.01317 | -0.03113 | 0.07556 | 0.21639 | -0.01393 | 0.153537 | 0.142072 | -0.25295 | 0.163942 | -0.06136 | 0.0813 | -0.24073 | 0.202285 | 0.016444 | 0.00102 | 0.30786 | -0.31823 | 0.16793 |
| CORL279 | 0.043491 | 0.191905 | 0.03418 | -0.14428 | -0.06214 | 0.10419 | -0.10524 | -0.18224 | 0.00984 | -0.00425 | 0.20799 | 0.17697 | 0.111927 | 0.286595 | -0.14803 | 0.165249 | -0.16485 | -0.28683 | -0.10231 | 0.243916 | -0.34532 | -0.11651 | 0.14016 | -0.01058 | 0.26901 |
| DU4475 | -0.18127 | 0.140591 | 0.16881 | -0.00027 | -0.20918 | -0.10457 | 0.006712 | -0.09978 | -0.00333 | -0.01126 | 0.12386 | -0.00368 | -0.07643 | 0.137006 | 0.241336 | 0.246086 | 0.100045 | -0.34509 | -0.03307 | -0.26395 | -0.00867 | 0.03755 | 0.07631 | 0.06118 | -0.00174 |
| KELLY | -0.53237 | 0.111594 | -0.02045 | -0.03857 | -0.02023 | -0.09875 | 0.055368 | -0.10606 | -0.01568 | -0.01579 | 0.16181 | 0.13023 | -0.04855 | -0.00748 | 0.020477 | 0.019842 | 0.022973 | -0.2204 | -0.11741 | -0.09607 | 0.007287 | -0.06468 | -0.02992 | -0.0684 | 0.10638 |
| SKNAS | 0.036517 | -0.05823 | -0.06932 | -0.02257 | -0.00021 | -0.03962 | 0.004495 | -0.14219 | -0.02081 | 0.12828 | 0.12602 | 0.03755 | 0.093872 | 0.013306 | -0.0627 | 0.008985 | -0.0042 | -0.19444 | -0.11322 | -0.06894 | -0.0288 | -0.03236 | 0.006 | 0.01756 | 0.07215 |

| Cell line | ALK | AXL | CSF1R | DDR1 | DDR2 | EGFR | EPHA1 | EPHA2 | EPHA3 | EPHA4 | EPHA6 | EPHA7 | EPHB1 | EPHB2 | EPHB3 | EPHB4 | EPHB6 | ERBB2 | ERBB3 | ERBB4 | FGFR1 | FGFR2 | FGFR3 | FGFR4 | FLT1 |
|---|---|---|---|---|---|---|---|---|---|---|---|---|---|---|---|---|---|---|---|---|---|---|---|---|---|
| RERFLCAI | 0.045276 | -0.12346 | -0.05142 | -0.00184 | 0.0149 | -0.10832 | 0.089574 | 0.06506 | -0.03201 | 0.0382 | 0.11169 | 0.09846 | 0.025911 | 0.065941 | -0.04177 | -0.08321 | 0.056912 | -0.2542 | -0.1839 | -0.18163 | -0.45579 | -0.01028 | 0.15958 | 0.04551 | 0.17012 |
| UOK101 | 0.219233 | 0.062976 | 0.0638 | 0.0305 | -0.06761 | -0.43829 | -0.24995 | 0.0725 | 0.02689 | 0.22879 | 0.11079 | 0.31331 | 0.010932 | 0.156869 | -0.1206 | -0.00374 | 0.093306 | -0.2637 | -0.09532 | -0.15742 | -0.56713 | -0.19811 | 0.41582 | 0.02794 | 0.1853 |
| KASUMI1 | -0.07411 | -0.04646 | -0.18633 | 0.11879 | 0.14521 | -0.063 | -0.00943 | -0.20427 | 0.03656 | 0.26106 | 0.1367 | 0.2048 | -0.01388 | -0.23614 | -0.39065 | -0.00366 | 0.033579 | -0.27536 | -0.20699 | -0.37668 | 0.112846 | 0.18173 | -0.00445 | -0.15739 | -0.0242 |
| CALU6 | 0.020593 | -0.03233 | 0.06375 | -0.02378 | -0.03877 | -0.21619 | 0.132877 | -0.08796 | 0.01796 | 0.04289 | 0.12247 | 0.12787 | 0.032599 | 0.062352 | -0.07039 | 0.014042 | 0.117003 | -0.17687 | -0.14731 | -0.16764 | -0.13824 | -0.00103 | 0.01127 | 0.11501 | 0.11912 |
| KP4 | 0.043188 | -0.00961 | -0.01023 | -0.02321 | -0.08638 | -0.05657 | -0.01355 | -0.00526 | 0.03428 | 0.03925 | 0.05099 | 0.07639 | -0.02074 | 0.02906 | -0.09943 | 0.060419 | 0.065983 | -0.17817 | -0.06104 | -0.04176 | -0.23364 | -0.08892 | 0.02904 | 0.05308 | 0.12851 |
| SNU213 | -0.07709 | -0.10259 | 0.03314 | -0.15004 | 0.04923 | -1.10509 | 0.074125 | 0.03619 | -0.01721 | 0.01734 | 0.11697 | 0.03731 | 0.056713 | 0.066795 | -0.17427 | 0.041223 | 0.056601 | -0.49412 | -0.166 | -0.15881 | 0.008714 | -0.12076 | 0.03977 | -0.0103 | 0.09496 |
| AM38 | 0.033091 | -0.26049 | -0.05978 | 0.00386 | -0.00499 | -0.00921 | 0.208594 | -0.289 | -0.13931 | -0.03206 | 0.38157 | 0.25658 | -0.01285 | 0.023629 | -0.06909 | 0.082183 | 0.0702 | -0.4171 | -0.21284 | -0.1162 | 0.105223 | -0.09113 | -0.02942 | 0.06912 | -0.00548 |
| SUDHL10 | 0.043824 | 0.103787 | -0.09706 | -0.21855 | -0.02725 | 0.12551 | -0.09047 | -0.15138 | 0.00523 | 0.01339 | 0.27068 | 0.16265 | -0.01816 | -0.11178 | -0.08487 | -0.00702 | 0.254443 | -0.2721 | -0.00772 | 0.025456 | -0.01543 | -0.12875 | -0.0988 | -0.06284 | 0.15744 |
| SLR24 | 0.044063 | 0.053258 | -0.07189 | 0.03398 | -0.11762 | -0.6552 | 0.021163 | 0.20975 | -0.1506 | 0.05287 | 0.1118 | 0.23955 | 0.017725 | 0.174771 | -0.11104 | 0.083079 | 0.159734 | -0.49268 | -0.66452 | 0.007577 | 0.021352 | -0.00623 | 0.16518 | -0.02958 | 0.11324 |
| SF539 | 0.02385 | 0.115685 | 0.02456 | -0.19 | -0.14139 | -0.01515 | -0.07148 | 0.15843 | 0.05705 | 0.10073 | 0.18122 | -0.00489 | 0.078744 | 0.023391 | -0.03916 | 0.009045 | 0.051991 | -0.08956 | -0.24459 | -0.04327 | -0.22238 | 0.01473 | -0.08105 | -0.03666 | 0.10866 |
| HS852T | 0.095376 | 0.021228 | -0.08515 | -0.0244 | 0.06812 | -0.40684 | 0.067277 | 0.19386 | 0.11058 | 0.18105 | 0.24386 | 0.10619 | 0.097286 | 0.051885 | -0.08637 | 0.023288 | 0.093983 | -0.25618 | -0.24839 | 0.019902 | -0.0807 | -0.14493 | -0.32747 | -0.09474 | 0.16084 |
| HCC38 | 0.101673 | -0.05416 | 0.00014 | -0.13855 | -0.00469 | -0.24308 | 0.135059 | 0.27107 | -0.30175 | 0.1867 | 0.07001 | 0.19556 | -0.19579 | 0.212884 | -0.02178 | -0.01142 | 0.395923 | -0.54168 | -0.27706 | -0.0811 | -0.57328 | -0.00707 | 0.08246 | -0.07928 | 0.24829 |
| HCC1419 | -0.26881 | 0.053011 | 0.20967 | 0.03389 | 0.05463 | -0.03546 | 0.061054 | -0.03374 | -0.02038 | 0.08794 | 0.12502 | 0.0937 | 0.035611 | 0.139164 | 0.00672 | -0.08245 | 0.030383 | -1.44979 | -1.11586 | -0.3832 | -0.0585 | -0.17361 | 0.08703 | 0.05801 | 0.04352 |
| COV362 | -0.11014 | -0.09028 | 0.0704 | -0.02824 | -0.25723 | -0.15224 | 0.081676 | 0.2671 | -0.09855 | 0.1949 | 0.16473 | -0.07275 | -0.00037 | 0.205382 | -0.07785 | 0.157274 | 0.178364 | -0.21439 | -0.21132 | 0.014759 | -0.00499 | 0.09144 | -0.01937 | -0.24774 | 0.02517 |
| EWS502 | 0.064774 | 0.051418 | 0.00751 | 0.08308 | 0.00137 | -0.14305 | 0.013982 | -0.09636 | -0.01093 | 0.12488 | 0.14108 | 0.00528 | 0.150818 | -0.02543 | -0.22209 | 0.058778 | 0.056427 | -0.31859 | -0.21404 | -0.12566 | -0.09978 | 0.02228 | -0.11842 | 0.06791 | 0.29681 |
| SNU840 | -0.13884 | 0.073054 | 0.07414 | 0.11031 | -0.28255 | -0.27981 | -0.22366 | 0.50293 | 0.13695 | 0.05607 | 0.3885 | -0.1155 | 0.081444 | 0.068468 | -0.27731 | 0.091536 | 0.019709 | -0.9919 | -0.57399 | 0.03637 | -0.00977 | 0.02333 | -0.03207 | 0.00467 | 0.20386 |
| KP2 | -0.17246 | 0.01488 | -0.28411 | -0.50289 | -0.12375 | 0.12598 | -0.2055 | 0.1492 | 0.03628 | -0.10708 | 0.10198 | 0.07903 | -0.11714 | 0.160865 | -0.32745 | 0.157436 | 0.1415 | -0.72839 | -0.33922 | 0.057267 | -0.0148 | -0.02742 | 0.11737 | -0.14866 | 0.21667 |
| NCIH1755 | 0.052334 | -0.05467 | -0.01806 | -0.08326 | -0.24283 | -0.1347 | -0.01314 | 0.261 | -0.11164 | 0.03993 | 0.05094 | 0.1579 | 0.019178 | 0.158139 | -0.18334 | 0.012915 | -0.11284 | -0.12228 | -0.23365 | -0.20631 | -0.42883 | -0.10827 | -0.00879 | -0.01767 | 0.03702 |
| SNU1033 | -0.05479 | 0.307763 | -0.0265 | 0.00649 | -0.07179 | -0.13199 | 0.100293 | 0.09148 | -0.22694 | 0.01899 | 0.19463 | 0.0369 | -0.02196 | -0.03304 | 0.177166 | -0.09573 | 0.339395 | -0.21861 | 0.1972 | -0.21958 | -0.01784 | -0.15808 | 0.14037 | -0.09761 | 0.02676 |
| BT549 | 0.113049 | -0.14935 | 0.00498 | -0.03693 | 0.15808 | -0.03667 | 0.004705 | 0.54304 | 0.01207 | 0.19658 | 0.10453 | 0.02424 | 0.166189 | 0.000351 | -0.10766 | 0.130969 | 0.020025 | -0.22098 | -0.21937 | 0.053262 | -0.66747 | -0.15457 | -0.00203 | -0.28566 | 0.29961 |
| NCIH209 | 0.071412 | 0.117382 | 0.03919 | 0.04313 | -0.024 | -0.18711 | -0.01022 | -0.0906 | -0.00753 | 0.02283 | 0.14416 | 0.05312 | 0.029213 | 0.006564 | -0.04784 | -0.05931 | 0.077054 | -0.25355 | -0.16292 | -0.07825 | -0.09138 | -0.05973 | 0.0347 | 0.01155 | 0.05658 |
| OV90 | -0.0433 | 0.106561 | 0.04875 | -0.0649 | -0.05059 | -0.12194 | -0.14931 | -0.08967 | -0.11523 | 0.01148 | 0.09319 | 0.08042 | -0.02673 | 0.013811 | -0.22462 | 0.004915 | -0.03211 | -0.20834 | -0.22505 | -0.2027 | -0.03832 | -0.09187 | -0.0424 | -0.0107 | 0.01299 |
| NCIH841 | -0.03389 | 0.091901 | -0.04764 | -0.08982 | -0.14023 | -0.07692 | 0.031003 | -0.0184 | -0.05342 | 0.13424 | 0.08736 | 0.07745 | 0.07784 | 0.075906 | -0.07888 | 0.207762 | 0.145443 | -0.211 | -0.11147 | -0.20707 | -0.24869 | -0.03777 | -0.02908 | 0.11339 | 0.41832 |
| KLE | -0.04527 | 0.176603 | 0.16149 | 0.21607 | 0.01277 | 0.13302 | 0.140337 | 0.12089 | 0.02419 | 0.18999 | -0.00741 | 0.22766 | 0.186013 | 0.188652 | 0.19358 | 0.034061 | 0.276477 | -0.20014 | -0.0761 | -0.37803 | -0.04147 | -0.07153 | 0.10178 | 0.19572 | 0.13091 |
| NB4 | -0.05671 | 0.156543 | 0.0217 | -0.13354 | 0.01655 | -0.05786 | -0.12388 | -0.05296 | 0.00397 | 0.08365 | 0.01074 | 0.12169 | 0.082091 | -0.06891 | 0.080029 | 0.027433 | 0.042853 | -0.72694 | -0.11285 | -0.21988 | -0.0356 | 0.02994 | 0.03396 | -0.09549 | -0.06979 |
| EM2 | -0.10613 | 0.066681 | 0.06353 | 0.05463 | 0.00466 | -0.06272 | 0.057848 | -0.22639 | 0.04039 | -0.01147 | 0.10464 | 0.28163 | -0.1235 | -0.09285 | 0.128938 | -0.25454 | 0.006804 | -0.35147 | -0.00784 | -0.54691 | -0.02804 | 0.04729 | 0.12772 | 0.22385 | -0.11107 |
| OUMS23 | -0.04268 | -0.10355 | 0.09198 | 0.05211 | -0.05357 | -0.0248 | 0.109941 | 0.17391 | -0.01887 | 0.00574 | 0.05035 | 0.01135 | 0.038487 | 0.158543 | -0.09717 | -0.06941 | 0.237102 | -0.30829 | -0.29418 | -0.06983 | -0.04706 | -0.18977 | -0.14859 | -0.05382 | 0.30762 |
| SNU1077 | 0.002666 | 0.120102 | 0.01562 | -0.02127 | 0.02762 | -0.60819 | 0.022043 | 0.17195 | -0.00631 | -0.00257 | -0.00177 | 0.00251 | -0.01489 | 0.023905 | 0.044293 | -0.13633 | 0.187799 | -0.25252 | -0.19032 | -0.27248 | -0.01786 | 0.00924 | -0.17745 | -0.15018 | 0.04001 |
| SNU5 | 0.043164 | 0.261967 | 0.08339 | -0.00615 | -0.01975 | -0.19461 | -0.1075 | -0.07921 | -0.24757 | 0.02262 | 0.08559 | -0.04183 | -0.07811 | 0.099698 | -0.29135 | 0.141318 | 0.049763 | -0.35065 | -0.31079 | -0.0106 | -0.03453 | 0.03038 | -0.05741 | 0.20119 | 0.01734 |
| WM115 | -0.04502 | -0.08478 | 0.10058 | 0.03989 | -0.20339 | -0.07921 | 0.149532 | -0.05416 | 0.13033 | -0.18417 | 0.12353 | -0.01682 | -0.19642 | -0.00523 | 0.091899 | 0.296873 | 0.183468 | -0.31549 | -0.36594 | -0.5591 | 0.077242 | 0.00744 | -0.41589 | -0.01488 | 0.34342 |
| ECGI10 | 0.081562 | 0.156564 | 0.02785 | -0.05414 | -0.0817 | -0.6529 | -0.04827 | 0.06057 | 0.01774 | 0.08374 | 0.11593 | 0.04583 | 0.01861 | 0.09212 | -0.17422 | -0.03759 | 0.098096 | -1.46159 | -1.01547 | -0.15448 | 0.020626 | -0.00025 | 0.03144 | -0.03457 | 0.15026 |
| PK1 | 0.050451 | 0.05005 | -0.10605 | -0.06753 | -0.08155 | -0.60794 | -0.09665 | 0.14688 | 0.17064 | 0.15146 | 0.05712 | 0.17158 | 0.029507 | 0.129227 | -0.17029 | 0.047755 | 0.176841 | -1.42655 | -0.83943 | -0.08936 | -0.11875 | -0.04427 | 0.05823 | 0.08066 | 0.17352 |
| EFO21 | 0.114819 | -0.04931 | 0.0373 | -0.04401 | -0.11921 | 0.06068 | -0.00151 | 0.17298 | 0.05427 | 0.09822 | 0.11511 | -0.0026 | 0.039104 | 0.123449 | -0.11326 | 0.059356 | 0.073658 | -0.14992 | -0.25766 | -0.05045 | 0.035775 | -0.02552 | 0.05839 | -0.08846 | 0.12783 |
| IMR32 | -0.0189 | 0.077146 | -0.02417 | 0.13393 | 0.08948 | 0.09512 | 0.015592 | -0.09046 | -0.17332 | 0.03191 | 0.04815 | 0.2444 | -0.04245 | 0.079706 | -0.07163 | 0.03732 | 0.133297 | -0.09011 | 0.08231 | -0.16247 | 0.088926 | -0.18372 | 0.11257 | 0.15122 | 0.1012 |
| NCIH2122 | 0.006195 | -0.00211 | -0.03869 | 0.02336 | 0.15207 | -0.05785 | -0.15723 | -0.13873 | -0.08167 | 0.07131 | 0.03731 | 0.21518 | -0.09014 | 0.02298 | -0.18405 | 0.058636 | 0.336544 | -0.25174 | 0.06725 | -0.20482 | 0.039383 | -0.13322 | 0.06035 | 0.03864 | 0.00897 |
| SKNBE2 | -0.25415 | -0.0902 | -0.21377 | 0.01293 | 0.09684 | -0.04364 | 0.051706 | -0.18901 | -0.164 | 0.17658 | 0.11173 | 0.16356 | -0.09214 | -0.10021 | -0.14448 | -0.02837 | 0.050608 | -0.23126 | 0.05284 | -0.13352 | -0.10068 | -0.0636 | 0.07747 | 0.0376 | 0.18849 |
| KMRC3 | 0.121539 | -0.36094 | -0.02152 | 0.10885 | 0.00048 | -0.3179 | -0.02988 | 0.22689 | -0.0636 | 0.01612 | 0.10942 | 0.06471 | 0.076868 | 0.141485 | -0.16125 | -0.02948 | 0.143563 | -0.24624 | -0.21422 | -0.05992 | -0.55507 | -0.05018 | 0.17228 | 0.02248 | 0.14142 |
| KARPAS422 | -0.10609 | 0.320508 | 0.13997 | -0.05887 | -0.06156 | -0.06141 | -0.2055 | -0.18925 | 0.01403 | -0.09503 | 0.11112 | 0.20427 | 0.060415 | -0.11314 | -0.0192 | -0.10682 | -0.02836 | -0.42846 | -0.20017 | -0.14713 | -0.06971 | -0.04941 | -0.02421 | 0.14394 | 0.14231 |
| SNU886 | 0.053522 | -0.03398 | -0.05798 | -0.18189 | 0.03752 | -0.51122 | 0.082359 | 0.05093 | -0.08618 | -0.02909 | 0.3125 | 0.0932 | -0.12046 | 0.114622 | 0.070879 | -0.00171 | 0.096674 | -0.40689 | -0.26152 | -0.02284 | -0.86597 | 0.13748 | -0.10327 | 0.02205 | -0.04397 |
| TUHR14TKB | -0.06174 | 0.052265 | 0.10605 | -0.00895 | -0.25985 | -0.21365 | 0.125563 | 0.1581 | -0.18145 | -0.14731 | 0.19729 | -0.00796 | -0.08357 | 0.157895 | -0.08668 | 0.167247 | -0.09877 | -0.67836 | -0.41483 | -0.10927 | -0.08418 | 0.09123 | -0.00384 | -0.03729 | -0.04642 |
| TE10 | -0.15749 | -0.03643 | 0.0001 | -0.09974 | -0.04583 | -0.58727 | 0.121959 | 0.69249 | -0.04656 | 0.06946 | 0.09199 | -0.00045 | 0.131914 | 0.014791 | -0.01606 | -0.02263 | 0.181271 | -0.2775 | -0.45027 | -0.11643 | -0.03455 | 0.15739 | -0.05362 | 0.01976 | 0.06107 |
| MPP89 | -0.00786 | -0.16849 | -0.05249 | 0.16857 | -0.05101 | -0.14837 | -0.06716 | -0.02818 | 0.05862 | 0.10981 | -0.01067 | 0.08522 | -0.04847 | -0.00533 | 0.010356 | -0.16642 | 0.167941 | -0.36383 | -0.22735 | -0.29131 | -0.68809 | -0.12786 | 0.01343 | 0.06372 | 0.03508 |
| PSN1 | -0.06026 | -0.05514 | 0.05456 | 0.01316 | -0.01669 | 0.06628 | 0.132706 | 0.18001 | -0.17654 | 0.19347 | 0.02803 | -0.03079 | 0.028236 | 0.055821 | 0.008606 | 0.060129 | 0.156759 | -0.29676 | -0.04343 | -0.00119 | -0.12531 | 0.06187 | 0.06923 | 0.02731 | 0.07654 |
| HT144 | -0.33683 | -0.14701 | 0.24058 | 0.04375 | -0.158 | 0.04164 | 0.085647 | -0.03136 | -0.00777 | 0.05098 | 0.20524 | 0.18235 | -0.05564 | 0.164064 | -0.00949 | 0.26922 | 0.226392 | -0.22665 | -0.29713 | 0.099856 | -0.04173 | 0.18359 | -0.11565 | -0.10776 | 0.17783 |
| 42MGBA | -0.0082 | -4.5E-06 | 0.10554 | -0.18971 | 0.00356 | -0.05692 | 0.014402 | 0.15893 | 0.01127 | 0.10189 | 0.02509 | -0.01323 | 0.071258 | 0.172763 | 0.014454 | 0.197049 | 0.063434 | -0.26187 | -0.05002 | -0.11903 | -0.58589 | 0.16868 | -0.05886 | -0.06485 | 0.04906 |
| JHOC5 | -0.06551 | 0.159049 | 0.0227 | 0.00173 | -0.12217 | -0.50082 | 0.118243 | 0.22706 | -0.1023 | 0.0296 | 0.27517 | 0.09794 | 0.124919 | 0.223146 | -0.07247 | -0.00417 | 0.153234 | -1.16406 | -0.71892 | -0.07802 | -0.08269 | -0.00675 | 0.13461 | 0.00018 | 0.01326 |
| SNU620 | 0.009065 | 0.107064 | -0.12292 | 0.054 | -0.00099 | -0.14559 | 0.022056 | -0.01439 | -0.01388 | -0.07822 | 0.08952 | 0.05894 | -0.13022 | 0.116375 | -0.03068 | -0.03286 | 0.020707 | -0.35601 | -0.49437 | -0.00353 | -0.09914 | -0.22378 | 0.20873 | 0.29803 | 0.1615 |
| JURLMK1 | 0.136018 | -0.04799 | 0.12619 | 0.09484 | 0.01918 | -0.04694 | 0.068318 | -0.00032 | -0.02113 | 0.08774 | 0.05082 | 0.10351 | -0.20339 | -0.00184 | -0.17031 | -0.08227 | 0.038354 | -0.2746 | -0.22795 | -0.09701 | 0.004268 | -0.13991 | -0.16895 | 0.27474 | -0.23179 |
| CCFSTTG1 | 0.05112 | -0.1074 | 0.15413 | 0.03614 | -0.06901 | -0.12174 | 0.023289 | 0.11359 | -0.06085 | 0.0805 | 0.14314 | 0.10751 | 0.196163 | -0.02369 | -0.15639 | 0.034413 | 0.095323 | -0.18118 | -0.21188 | -0.0411 | -0.31915 | -0.00788 | 0.04877 | -0.12751 | -0.15787 |
| EFM19 | 0.121082 | 0.201428 | 0.02911 | 0.09752 | -0.22051 | -0.13441 | -0.02192 | -0.15091 | -0.10182 | 0.26211 | 0.16732 | -0.08903 | 0.085311 | 0.037494 | -0.05366 | 0.153439 | 0.002653 | -0.55417 | -0.19382 | 0.151739 | 0.206022 | -0.20361 | -0.04206 | 0.04985 | 0.16111 |
| ISTMES2 | 0.047154 | 0.047732 | -0.03856 | -0.00208 | 0.03897 | -0.18865 | -0.01614 | 0.06763 | -0.09866 | 0.00083 | 0.1104 | 0.13934 | 0.038647 | 0.042913 | -0.03228 | -0.03845 | 0.068495 | -0.06855 | -0.27766 | -0.11375 | -0.47731 | -0.2025 | -0.00817 | 0.01263 | 0.10567 |
| YAPC | 0.037621 | 0.095741 | -0.12529 | 0.04145 | -0.01048 | -0.43062 | 0.043766 | -0.07522 | -0.16159 | 0.02403 | 0.1258 | 0.08177 | 0.042482 | 0.05552 | -0.21783 | 0.029331 | 0.036821 | -0.3076 | -0.31226 | -0.05632 | -0.05478 | -0.12316 | -0.00924 | 0.00171 | 0.1299 |
| DB | -0.09432 | 0.075034 | 0.15842 | 0.12985 | 0.15121 | 0.03337 | -0.06954 | -0.28822 | 0.04327 | 0.06911 | 0.09103 | 0.01682 | 0.007491 | 0.013783 | 0.072904 | 0.093027 | 0.193346 | -0.02154 | 0.06787 | -0.37568 | -0.0357 | -0.01254 | 0.20308 | 0.28167 | 0.09955 |
| MSTO211H | -0.19355 | 0.047014 | -0.06869 | -0.22897 | 0.00823 | -0.06326 | -0.17869 | 0.02742 | 0.13019 | 0.1471 | 0.21734 | 0.16247 | -0.15103 | 0.093629 | -0.09165 | 0.255794 | 0.078187 | -0.1591 | -0.10832 | 0.080307 | -0.5839 | 0.13819 | 0.1742 | 0.36476 | 0.54097 |
| OCIAML3 | -0.06359 | -0.0247 | 0.04818 | 0.12255 | 0.00779 | 0.07317 | 0.051111 | -0.28579 | -0.04757 | -0.02665 | -0.07083 | 0.05196 | -0.00679 | -0.01959 | -0.10718 | -0.15782 | -0.16916 | -0.13992 | 0.04834 | -0.2145 | 0.04213 | 0.09113 | 0.22322 | 0.04283 | -0.01542 |
| NCIH3122 | 0.015004 | -0.02805 | -0.04801 | 0.18263 | -0.01089 | -0.0735 | 0.115448 | -0.11332 | -0.11475 | 0.03823 | 0.05662 | 0.29182 | -0.12952 | 0.203401 | 0.083028 | -0.01962 | 0.095113 | 0.03259 | -0.17996 | -0.16555 | 0.065366 | 0.09246 | 0.13658 | -0.07309 | 0.03404 |
| HCC461 | 0.03635 | 0.04348 | -0.03423 | 0.00241 | 0.0447 | -0.43511 | 0.020312 | 0.18393 | -0.11322 | 0.04948 | 0.13471 | 0.10225 | 0.01427 | 0.087408 | -0.08128 | 0.142749 | 0.10821 | -0.18474 | -0.28621 | -0.04573 | -0.0695 | -0.11635 | 0.05629 | -0.03197 | 0.23293 |
| SKNFI | 0.024889 | -0.03017 | 0.05357 | -0.09966 | -0.08265 | -0.06235 | 0.010728 | -0.02656 | 0.02637 | 0.08038 | 0.08363 | 0.17139 | 0.044206 | -0.04054 | -0.16122 | -0.00945 | -0.04292 | -0.05809 | -0.10065 | -0.11182 | -0.15872 | -0.03896 | 0.13251 | 0.00502 | 0.15237 |
| NCIH522 | 0.04263 | 0.117253 | -0.03681 | 0.12684 | -0.05141 | -0.08918 | -0.17266 | -0.08895 | 0.04824 | 0.10073 | 0.2154 | 0.03598 | 0.021601 | 0.043624 | -0.04063 | -0.0311 | 0.021856 | -0.79165 | -0.21609 | -0.34148 | -0.43002 | 0.08756 | -0.10742 | -0.01225 | 0.28241 |
| SNU668 | -0.02543 | -0.01202 | 0.01523 | 0.05388 | -0.05806 | -0.15512 | -0.06257 | -0.08247 | 0.07572 | 0.1042 | 0.16172 | 0.19599 | 0.128221 | -0.00677 | -0.13158 | 0.086271 | 0.203113 | -0.47696 | -0.23521 | -0.12758 | -0.03975 | 0.05312 | 0.08701 | -0.00099 | -0.03769 |
| JVM3 | -0.11972 | -0.21931 | -0.20373 | 0.06874 | -0.09735 | 0.02756 | 0.09664 | -0.14497 | -0.03685 | 0.0759 | 0.01304 | 0.08273 | -0.11069 | -0.02887 | -0.33177 | -0.04978 | -0.04106 | -0.47253 | -0.06637 | -6.7E-05 | -0.16491 | 0.18163 | -0.20245 | -0.01149 | 0.25228 |
| RPMI7951 | -0.04515 | -0.20537 | 0.07426 | -0.13193 | -0.05209 | -0.2105 | 0.053995 | -0.00442 | 0.02784 | 0.00667 | 0.06599 | 0.0372 | 0.08535 | 0.053437 | -0.05686 | -0.07631 | 0.115049 | -0.10299 | -0.19399 | -0.22793 | -0.05359 | -0.13858 | 0.06963 | 0.17673 | -0.02002 |
| COLO678 | -0.07222 | 0.016572 | -0.28549 | -0.05591 | -0.17191 | -0.69638 | -0.21138 | 0.02803 | 0.16034 | 0.12453 | 0.04771 | 0.04079 | -0.14267 | 0.056748 | -0.36813 | 0.112128 | -0.03291 | -0.77952 | -0.17787 | 0.067576 | -0.00146 | -0.08074 | 0.02766 | -0.19158 | 0.35193 |
| HCC1428 | -0.06214 | -0.01558 | -0.10637 | -0.00873 | 0.05956 | -0.03398 | 0.156421 | -0.04938 | 0.05408 | 0.10689 | -0.01056 | 0.0313 | 0.087957 | 0.097567 | -0.1085 | 0.069472 | -0.11941 | -0.04234 | -0.05781 | -0.20788 | -0.18733 | -0.2766 | 0.03482 | -0.21428 | 0.01836 |
| CAPAN1 | 0.148013 | -0.07072 | 0.09819 | -0.1829 | -0.08072 | -0.83106 | -0.12543 | -0.0815 | 0.14024 | 0.19601 | 0.17847 | 0.07712 | -0.13884 | 0.051803 | -0.05999 | -0.03659 | 0.223574 | -0.66541 | -0.20576 | 0.229542 | -0.1467 | 0.04272 | -0.10343 | 0.17229 | 0.02532 |
| NCIH82 | -0.20263 | 0.044843 | -0.03844 | -0.01533 | -0.00098 | -0.32828 | 0.042058 | -0.002 | -0.15601 | 0.24505 | -0.05375 | 0.29651 | 0.151244 | -0.05637 | -0.11387 | -0.02835 | -0.01657 | -0.37988 | -0.15847 | -0.14656 | -0.01172 | -0.0897 | 0.07222 | 0.31186 | -0.1431 |
| MKN45 | -0.10003 | 0.095714 | -0.10721 | 0.16962 | -0.1503 | -0.02571 | -0.09991 | -0.06989 | -0.03628 | 0.14849 | 0.07516 | 0.26537 | -0.04113 | 0.080515 | 0.090459 | 0.004874 | 0.187799 | -0.33919 | -0.35073 | -0.10686 | 0.054452 | -0.0804 | -0.00934 | 0.05405 | -0.08488 |
| MG63 | 0.068163 | -0.07866 | 0.10079 | 0.00619 | -0.06937 | -0.16691 | 0.020461 | 0.1008 | -0.20814 | 0.06698 | 0.04779 | -0.03497 | -0.05755 | 0.075096 | 0.003121 | 0.007644 | 0.085773 | -0.27803 | -0.18523 | -0.15929 | -0.64856 | -0.0498 | 0.00568 | 0.04245 | 0.21237 |
| SKHEP1 | 0.044302 | -0.03839 | -0.32146 | 0.06479 | 0.1218 | -0.10805 | -0.04246 | 0.07263 | 0.03048 | 0.08089 | 0.02924 | 0.03701 | -0.12472 | 0.052051 | -0.1028 | 0.319174 | -0.02685 | -0.28815 | -0.3918 | 0.067035 | -0.60639 | -0.07115 | 0.16807 | -0.06432 | 0.13619 |
| MOLM13 | 0.073758 | -0.10673 | -0.11549 | 0.13597 | -0.03569 | -0.07665 | -0.1231 | -0.11093 | 0.00172 | 0.03384 | 0.08028 | 0.14375 | 0.044688 | 0.015658 | 0.007248 | -0.03034 | 0.195262 | -0.2438 | -0.16734 | -0.07181 | -0.02359 | -0.01824 | -0.01124 | -0.03859 | 0.20363 |
| SKMM2 | 0.065891 | 0.124268 | 0.02365 | -0.0049 | 0.05063 | -0.14058 | 0.03713 | -0.33478 | 0.02859 | -0.06425 | 0.1385 | -0.01995 | -0.11598 | -0.00344 | -0.16279 | -0.08178 | 0.220509 | -0.0912 | -0.12559 | -0.11392 | -0.11455 | -0.20976 | 0.01947 | -0.04607 | 0.04378 |
| U2OS | -0.10628 | -0.10832 | 0.02308 | -0.07693 | 0.08818 | -0.19162 | 0.032838 | 0.21793 | 0.04355 | 0.14956 | 0.24101 | 0.03654 | 0.097314 | -0.02808 | -0.04598 | 0.04158 | 0.256794 | -0.36901 | -0.18421 | -0.19907 | -0.53452 | -0.25412 | -0.18499 | 0.05961 | 0.02876 |
| SUDHL4 | 0.094468 | 0.049853 | -0.12135 | -0.07298 | -0.03339 | 0.09179 | -0.06634 | -0.12191 | 0.1582 | 0.11876 | 0.24896 | -0.03219 | -0.00542 | -0.09769 | -0.2229 | 0.094244 | 0.16557 | -0.20839 | -0.27936 | -0.05418 | -0.16596 | -0.04226 | -0.0892 | -0.01423 | 0.25506 |
| Cell line | ALK | AXL | CSF1R | DDR1 | DDR2 | EGFR | EPHA1 | EPHA2 | EPHA3 | EPHA4 | EPHA6 | EPHA7 | EPHB1 | EPHB2 | EPHB3 | EPHB4 | EPHB6 | ERBB2 | ERBB3 | ERBB4 | FGFR1 | FGFR2 | FGFR3 | FGFR4 | FLT1 |

| | | | | | | | | | | | | | | | | | | | | | | | | | |
|---|---|---|---|---|---|---|---|---|---|---|---|---|---|---|---|---|---|---|---|---|---|---|---|---|---|
| SKNDZ | 0.013058 | 0.000696 | -0.05962 | -0.06613 | -0.14158 | -0.07448 | -0.04596 | -0.04476 | 0.05359 | 0.26393 | 0.16205 | -0.01996 | 0.014122 | 0.095221 | -0.1455 | -0.03083 | -0.0171 | -0.20293 | -0.15308 | -0.04189 | -0.18056 | -0.20814 | 0.10228 | -0.08368 | 0.14093 |
| NCIH226 | 0.053815 | -0.02031 | -0.05874 | 0.29793 | -0.01658 | 0.12411 | 0.00958 | 0.25544 | 0.24856 | -0.06881 | 0.10322 | 0.23661 | 0.322325 | 0.46698 | -0.03992 | 0.068694 | 0.115706 | -0.10584 | 0.032 | -0.2261 | -1.20287 | 0.12076 | -0.11315 | 0.00205 | 0.39911 |
| SNU1105 | -0.22626 | -0.08906 | -0.10029 | -0.03501 | -0.02729 | -0.03868 | -0.20293 | 0.5605 | -0.03639 | 0.11748 | 0.05824 | 0.05259 | 0.103095 | 0.107398 | 0.129342 | 0.154191 | 0.206514 | -0.05947 | -0.15681 | -0.03213 | -0.85017 | -0.10223 | -0.39605 | 0.08975 | -0.08555 |
| SNU626 | 0.173922 | 0.064361 | -0.10396 | -0.19089 | -0.13047 | -0.11135 | 0.258765 | 0.17622 | 0.02047 | 0.2496 | 0.08768 | 0.13994 | 0.273289 | -0.0327 | -0.24951 | -0.25415 | 0.098137 | 0.05104 | -0.0542 | -0.32388 | 0.205423 | 0.27092 | 0.01022 | 0.13936 | -0.26352 |
| RL | -0.05567 | -0.03478 | 0.01713 | 0.00205 | 0.02002 | -0.00077 | -0.03932 | -0.21598 | -0.06617 | 0.07511 | 0.0905 | 0.12731 | -0.0067 | 0.056515 | -0.11335 | 0.154562 | 0.121465 | -0.16303 | -0.111 | -0.25166 | -0.06811 | -0.17922 | 0.0168 | 0.11322 | 0.16197 |
| HCC1143 | 0.108347 | 0.228249 | -0.01031 | -0.22879 | -0.14546 | -0.26639 | -0.08003 | 0.11414 | -0.17671 | 0.16093 | 0.14407 | 0.04842 | 0.049488 | 0.054508 | -0.15966 | 0.034352 | 0.107912 | -0.93045 | -0.59307 | 0.248464 | -0.35236 | 0.04574 | 0.16805 | 0.03439 | 0.03008 |
| G402 | 0.183773 | 0.072348 | -0.09684 | 0.00521 | -0.06508 | -0.12944 | 0.080735 | 0.09314 | -0.22026 | 0.04779 | 0.25984 | 0.07558 | 0.114189 | 0.060063 | 0.125573 | -0.02354 | 0.171945 | -0.24164 | -0.22351 | -0.22587 | -0.60494 | 0.10437 | -0.15487 | -0.14015 | 0.23307 |
| SF295 | 0.03018 | -0.01161 | 0.08731 | -0.01053 | 0.01304 | -0.08654 | 0.047031 | 0.05018 | 0.06862 | -0.05053 | 0.20215 | 0.04597 | 0.06283 | -0.0524 | -0.11007 | 0.056269 | 0.056859 | -0.15384 | -0.19144 | -0.05245 | -0.0565 | -0.08432 | 0.07401 | 0.04155 | 0.10558 |
| SNU478 | -0.01301 | 0.000374 | -0.0495 | -0.09573 | -0.03066 | -1.1077 | 0.302898 | 0.11285 | -0.14063 | 0.06752 | 0.04361 | 0.05904 | 0.144946 | -0.09828 | -0.21501 | -0.01832 | 0.151477 | -0.14392 | -0.10827 | 0.099681 | -0.23087 | 0.12016 | -0.03168 | -0.19247 | 0.29077 |
| NCIH647 | -0.11786 | -0.11334 | 0.02065 | 0.09507 | 0.11974 | 0.03007 | -0.00535 | 0.09121 | -0.05414 | 0.19279 | 0.16024 | 0.17317 | 0.057089 | 0.017502 | 0.119699 | 0.022355 | 0.19568 | -0.32179 | -0.23171 | -0.19589 | 0.0442 | 0.15938 | -0.09205 | 0.11169 | -0.03309 |
| T84 | 0.086175 | 0.175282 | 0.09361 | 0.00975 | 0.11756 | -0.26879 | -0.05385 | -0.13206 | -0.07482 | -0.01734 | 0.00646 | 0.07965 | 0.082862 | 0.057813 | -0.15325 | -0.14235 | 0.298131 | -0.26719 | -0.00432 | -0.13426 | 0.104007 | -0.02653 | 0.01795 | 0.12203 | -0.01779 |
| OE33 | 0.029429 | 0.018378 | -0.01661 | -0.0796 | -0.11764 | -0.42231 | 0.017063 | 0.00641 | 0.00459 | 0.20912 | 0.14173 | -0.07686 | -0.00486 | 0.045356 | -0.02072 | 0.087051 | 0.161003 | -0.74513 | -0.2228 | -0.04361 | 0.002952 | 0.00381 | -0.01856 | -0.057 | 0.07732 |
| SKRC20 | -0.24709 | 0.189281 | 0.1276 | 0.05774 | -0.13819 | -0.55742 | -0.302 | 0.09489 | 0.11228 | 0.024 | 0.2005 | 0.10524 | -0.01036 | -0.01258 | 0.062342 | -0.1475 | 0.107801 | -0.7853 | -0.47764 | -0.1063 | -0.14222 | -0.1293 | -0.02019 | 0.15142 | 0.01494 |
| TF1 | -0.03514 | 0.077454 | -0.03733 | -0.10535 | -0.01438 | -0.13284 | 0.096743 | -0.08514 | -0.04152 | 0.09308 | 0.17864 | -0.05178 | 0.046544 | 0.032817 | -0.20695 | 0.212136 | 0.050683 | -0.32432 | -0.1513 | -0.06683 | -0.02678 | 0.0999 | 0.01075 | 0.0076 | 0.21315 |
| H4 | 0.030593 | 0.171212 | -0.09872 | 0.17422 | 0.12897 | -0.00902 | -0.17886 | 0.14916 | 0.04888 | 0.01746 | 0.19985 | 0.13372 | -0.05385 | 0.343682 | -0.32386 | 0.017594 | 0.180045 | -0.12349 | -0.12636 | -0.14586 | -0.28253 | -0.12795 | -0.11525 | 0.08068 | 0.1022 |
| LUDLU1 | 0.118097 | 0.177665 | -0.06448 | 0.0599 | -0.05699 | -0.78752 | -0.03019 | 0.1531 | 0.09546 | 0.23585 | 0.11055 | 0.07761 | 0.095753 | 0.102672 | -0.08739 | 0.054228 | 0.167542 | -0.29841 | -0.30739 | -0.09675 | -0.03775 | -0.02426 | 0.099 | 0.01964 | 0.09326 |
| MHHES1 | 0.035334 | 0.018819 | 0.00547 | 0.12949 | -0.13058 | 0.0436 | 0.121552 | -0.11077 | 0.01042 | 0.08445 | 0.09497 | 0.06083 | -0.06932 | -0.02 | -0.15827 | -0.01181 | 0.040574 | -0.21624 | -0.14504 | -0.10847 | -0.24231 | 0.08164 | 0.0812 | -0.12946 | 0.13279 |
| HLF | -0.00266 | -0.03092 | -0.0318 | -0.18094 | 0.11828 | 0.07142 | -0.04651 | 0.22269 | -0.06478 | 0.03569 | -0.01029 | 0.1353 | 0.045115 | 0.120018 | -0.13231 | 0.06715 | 0.16401 | -0.31398 | -0.30225 | -0.09786 | -1.20296 | -0.07715 | -0.0564 | -0.12327 | 0.13267 |
| NCIH520 | -0.09303 | -0.04206 | -0.04353 | 0.09448 | 0.03341 | -0.22135 | 0.038866 | -0.08439 | -0.07183 | 0.06822 | 0.13551 | 0.16524 | -0.03743 | -0.06017 | -0.05157 | 0.014538 | -0.12774 | -0.33992 | -0.10141 | -0.00058 | -0.52793 | 0.1454 | 0.01764 | -0.11205 | 0.18572 |
| J82 | -0.20871 | -0.11795 | 0.16213 | -0.14423 | 0.062 | -0.05534 | 0.279953 | 0.15191 | 0.11149 | -0.10538 | -0.10643 | 0.24393 | 0.287741 | 0.079523 | 0.083445 | -0.10329 | 0.183935 | -0.35185 | -0.20295 | -0.26168 | -0.16458 | 0.11005 | 0.09392 | 0.11732 | -0.12423 |
| TEN | 0.017717 | -0.03762 | -0.0036 | -0.06998 | -0.05106 | -1.1704 | -0.03463 | 0.05092 | -0.0927 | 0.15264 | -0.00668 | 0.0707 | 0.017451 | 0.126081 | -0.23358 | -0.06247 | -0.12233 | -0.56304 | -0.28352 | -0.11789 | -0.04354 | -0.12804 | 0.07541 | 0.07712 | 0.15089 |
| RI1 | 0.02336 | 0.095771 | -0.15133 | 0.06682 | -0.03857 | 0.03999 | -0.12832 | -0.16392 | -0.0168 | 0.05489 | 0.24551 | 0.01475 | 0.083662 | 0.007752 | -0.03638 | 0.101295 | -0.12218 | -0.12003 | -0.30914 | 0.147081 | -0.03897 | 0.11593 | -0.21258 | 0.03701 | 0.17622 |
| COLO800 | -0.26096 | 0.114677 | -0.01174 | 0.28779 | 0.04211 | -0.11813 | -0.3385 | -0.16775 | -0.00177 | 0.26331 | 0.02528 | 0.15532 | -0.10117 | 0.170938 | 6.06E-05 | 0.049997 | 0.049108 | -0.2951 | -0.22425 | -0.17757 | -0.04013 | -0.07233 | -0.06713 | 0.18956 | 0.10992 |
| BL70 | -0.06596 | 0.001367 | 0.15622 | -0.06707 | -0.10536 | -0.05039 | 0.008236 | -0.09691 | -0.14287 | 0.07127 | -0.01163 | 0.07752 | 0.06342 | -0.07333 | 0.063783 | -0.04494 | -0.17911 | -0.16066 | -0.29979 | 0.118427 | -0.00393 | 0.06699 | -0.15261 | -0.02288 | -0.01681 |
| NCIH747 | 0.113659 | 0.037848 | 0.00213 | 0.06423 | -0.15193 | -0.34371 | 0.024017 | 0.1976 | -0.03592 | 0.14843 | 0.19542 | 0.01944 | -0.05795 | 0.154212 | -0.10827 | -0.01864 | -0.03666 | -0.5254 | -0.29181 | 0.042898 | 0.081111 | -0.07145 | -0.06302 | -0.12335 | 0.21651 |
| K029AX | -0.07609 | -0.17703 | -0.25398 | -0.06982 | 0.18648 | -0.12984 | -0.05017 | -0.02739 | -0.19205 | 0.0771 | 0.10286 | 0.12784 | 0.049302 | 0.174921 | 0.287273 | -0.03016 | 0.119776 | -0.2393 | -0.19022 | 0.022063 | 0.149563 | -0.10859 | 0.0163 | 0.1588 | -0.18061 |
| MEC1 | 0.007291 | 0.065002 | 0.03211 | 0.12374 | -0.09118 | -0.22599 | -0.14298 | -0.16625 | -0.02353 | -0.06136 | 0.31085 | 0.11913 | 0.032169 | -0.11317 | -0.05392 | -0.11207 | 0.057908 | -0.35005 | -0.27016 | -0.0411 | 0.051554 | -0.06346 | 0.09902 | -0.18012 | 0.11245 |
| U937 | -0.01042 | 0.025212 | 0.02716 | -0.04269 | 0.03087 | -0.1428 | 0.058299 | -0.11688 | 0.04956 | 0.07638 | 0.07161 | 0.1071 | 0.008799 | -0.02698 | -0.09703 | 0.004779 | 0.123601 | -0.40567 | -0.16634 | -0.11158 | -0.05146 | -0.02786 | -0.07523 | 0.03966 | 0.03192 |
| SNU685 | -0.04429 | -0.03014 | -0.02846 | -0.10694 | 0.01831 | -0.11632 | -0.01093 | -0.03199 | 0.06574 | 0.03758 | 0.11721 | 0.07603 | -0.05992 | -0.02109 | -0.04793 | 0.077537 | 0.045905 | -0.02061 | -0.19491 | -0.15148 | -0.37193 | 0.18453 | -0.02741 | 0.12433 | 0.04639 |
| TE5 | -0.02344 | 0.091146 | 0.08445 | 0.16048 | -0.14628 | -0.14849 | 0.135501 | 0.52829 | -0.0958 | 0.3267 | -0.10325 | 0.0372 | -0.01265 | 0.223471 | 0.089412 | 0.087475 | 0.01679 | -0.64004 | -0.40591 | -0.12532 | -0.08178 | 0.14784 | 0.16737 | -0.06373 | 0.26554 |
| SAOS2 | -0.10678 | 0.02788 | 0.10051 | -0.19712 | -0.1164 | -0.1298 | -0.00149 | 0.038 | -0.01791 | -0.04031 | 0.16678 | 0.04376 | 0.052138 | 0.141897 | 0.055791 | 0.015876 | 0.033314 | -0.24909 | -0.20143 | -0.13667 | 0.040856 | -0.13648 | 0.15278 | 0.13965 | 0.07941 |
| 769P | -0.08935 | 0.154722 | -0.06446 | -0.07972 | -0.11784 | -0.86587 | 0.065353 | 0.10708 | -0.04096 | 0.10899 | 0.17207 | 0.16104 | 0.039572 | 0.2172 | -0.11935 | 0.086858 | 0.09368 | -0.38801 | -0.17552 | -0.06972 | 0.039881 | -0.20729 | 0.05979 | 0.00443 | 0.14069 |
| NCIH1944 | 0.009953 | 0.063073 | 0.00219 | 0.07674 | 3.3E-05 | -0.18346 | -0.03385 | 0.00055 | -0.02156 | 0.16554 | 0.09814 | 0.1141 | 0.069539 | 0.105994 | -0.06192 | 0.058257 | 0.257227 | -0.58189 | -0.20973 | -0.1543 | 0.003988 | -0.0504 | 0.01802 | 0.04243 | 0.10938 |
| BICR6 | 0.092671 | 0.103469 | -0.04333 | -0.03486 | 0.00288 | -0.43995 | 0.142188 | 0.32941 | -0.02012 | 0.04934 | 0.29872 | 0.18751 | 0.008 | 0.096865 | -0.12037 | 0.079526 | 0.14468 | -1.07212 | -0.60758 | -0.09322 | -0.06748 | -0.02688 | -0.01159 | 0.01481 | 0.05688 |
| NCIH838 | 0.024781 | -0.01189 | 0.03942 | 0.00771 | -0.02907 | -0.13779 | -0.01152 | 0.04304 | 0.05747 | 0.06297 | 0.09006 | 0.05273 | 0.092169 | 0.022811 | -0.1234 | -0.06139 | 0.055515 | -0.22452 | -0.20357 | -0.08163 | -0.50972 | -0.10435 | -0.19695 | -0.01714 | 0.0999 |
| PANC0813 | -0.04464 | -0.09493 | -0.03121 | -0.11605 | 0.00928 | -0.46222 | -0.07932 | -0.06602 | -0.01924 | 0.05828 | 0.08336 | 0.08317 | -0.05085 | 0.10026 | -0.091 | -0.0017 | 0.033854 | -0.361 | -0.3067 | 0.075344 | 0.076059 | -0.04962 | -0.02196 | -0.04482 | 0.13945 |
| SNU449 | 0.08882 | -0.17945 | 0.05907 | -0.08509 | 0.05764 | -0.37049 | -0.12961 | 0.16684 | -0.0588 | 0.00188 | 0.04978 | 0.18343 | 0.20104 | 0.120618 | -0.17148 | -0.03746 | 0.005515 | -0.1292 | 0.01007 | -0.12087 | -0.23604 | -0.21206 | -0.04454 | 0.12443 | 0.0556 |
| SW837 | -0.01527 | -0.03871 | 0.0183 | 0.00783 | -0.12662 | -0.49024 | -0.01482 | 0.0301 | -0.04382 | 0.14729 | 0.12857 | -0.02583 | 0.021607 | 0.134625 | -0.08082 | -0.0476 | 0.131144 | -0.6003 | -0.28865 | -0.04411 | -0.06906 | -0.04844 | 0.12784 | -0.10359 | 0.17573 |
| SNU475 | -0.03344 | 0.208482 | 0.11499 | -0.12703 | -0.03105 | -0.14602 | 0.155077 | -0.02324 | 0.07112 | -0.02897 | 0.12465 | 0.05663 | -0.11924 | 0.097851 | -0.05837 | 0.041093 | 0.180669 | -0.44141 | -0.34245 | -0.1416 | -0.58408 | 0.13053 | -0.12409 | 0.05426 | 0.01872 |
| TC71 | -0.03992 | -0.04546 | 0.08491 | -0.07101 | -0.08701 | 0.03679 | -0.01665 | 0.07455 | -0.21423 | 0.12728 | 0.12104 | 0.05467 | -0.09358 | 0.123738 | 0.13015 | 0.049875 | 0.108883 | -0.26089 | -0.28571 | -0.11463 | -0.22747 | -0.06488 | 0.14462 | 0.04299 | 0.16142 |
| UACC62 | -0.03164 | -0.02564 | 0.11303 | -0.13719 | 0.08656 | -0.10575 | -0.00427 | -0.00925 | -0.08272 | 0.09187 | 0.24737 | -0.02429 | 0.088961 | 0.070188 | -0.13266 | 0.091326 | 0.212122 | -0.13079 | -0.26003 | -0.10425 | -0.06751 | 0.04167 | 0.09806 | -0.12416 | -0.09015 |
| KMS20 | -0.11159 | -0.08111 | 0.10417 | 0.27555 | -0.06294 | -0.12008 | -0.21621 | -0.20006 | 0.0133 | 0.20856 | 0.04945 | 0.20503 | -0.00059 | -0.06755 | 0.033066 | -0.17011 | 0.160063 | -0.36841 | -0.2863 | -0.09873 | -0.16641 | 0.00195 | -0.01643 | 0.11802 | -0.03641 |
| NCIN87 | -0.06603 | 0.258556 | 0.0697 | 0.12462 | -0.03831 | -0.4549 | 0.073726 | 0.13785 | -0.03317 | 0.16807 | -0.34279 | 0.26456 | 0.119922 | 0.129503 | 0.058056 | 0.04918 | 0.304877 | -1.30572 | -0.48099 | -0.14355 | -0.09265 | -0.15063 | 0.14588 | 0.12178 | 0.07411 |
| TYKNU | -0.19809 | -0.03108 | 0.09892 | -0.08683 | -0.15585 | 0.14087 | -0.11281 | 0.18071 | -0.12806 | 0.04893 | 0.26975 | -0.02432 | -0.10238 | -0.12177 | -0.21336 | 0.029533 | -0.14236 | -0.14935 | -0.14483 | -0.12065 | -0.05767 | 0.13204 | 0.02085 | -0.04569 | 0.10384 |
| NCIH1694 | -0.10317 | 0.065469 | 0.03005 | 0.09402 | 0.01367 | -0.25217 | 0.057071 | -0.05049 | -0.08668 | 0.01479 | 0.02963 | 0.08695 | -0.10089 | 0.084254 | 0.156243 | 0.080915 | 0.152752 | -0.35401 | -0.01532 | -0.2286 | -0.02361 | -0.28009 | 0.0924 | 0.22005 | 0.03683 |
| CAKI1 | 0.07065 | 0.182624 | 0.08223 | -0.10145 | -0.28906 | -0.84721 | 0.070445 | 0.25409 | 0.01644 | 0.19858 | -0.02074 | 0.18231 | 0.08894 | 0.225629 | 0.032002 | 0.073056 | 0.336231 | -0.37945 | -0.35948 | -0.04081 | -0.09572 | 0.09792 | -0.1553 | -0.10729 | 0.82311 |
| NCIH1915 | -0.09722 | -0.10212 | -0.01678 | -0.10641 | 0.00733 | 0.02293 | 0.157748 | -0.05822 | -0.12287 | 0.03007 | 0.1806 | -0.06557 | -0.15484 | 0.015909 | -0.18576 | 0.074071 | 0.018391 | -0.29194 | -0.35046 | -0.12602 | -0.35692 | 0.02559 | 0.06061 | -0.02949 | 0.11785 |
| EFE184 | -0.20719 | 0.026596 | 0.00502 | 0.12342 | 0.04021 | -0.3757 | -0.06732 | 0.03463 | -0.13463 | 0.13139 | -0.04888 | 0.17862 | 0.091259 | 0.040974 | 0.213693 | -0.11768 | 0.396906 | -0.37896 | -0.18885 | -0.27363 | 0.094976 | -0.10214 | 0.26222 | 0.29011 | -0.1089 |
| OCIMY7 | 0.023593 | -0.00181 | 0.03197 | -0.08516 | -0.16966 | -0.00461 | -0.08465 | -0.28617 | -0.30384 | -0.13815 | 0.261 | 0.05923 | 0.102598 | 0.06207 | -0.05403 | 0.119625 | -0.04572 | -0.39676 | -0.23862 | -0.05869 | 0.014283 | 0.02947 | -0.04031 | 0.11933 | 0.29053 |
| SW1088 | 0.104057 | -0.04448 | 0.01389 | -0.01201 | -0.016 | -0.05013 | 0.098157 | -0.06076 | 0.0348 | 0.09063 | 0.14616 | 0.07123 | 0.181271 | 0.031266 | -0.10156 | 0.043634 | -0.05326 | -0.09408 | -0.18223 | -0.1279 | -0.29294 | 0.00199 | -0.00853 | -0.00156 | 0.02211 |
| LU65 | -0.04361 | 0.125417 | -0.04065 | -0.09586 | -0.11298 | -0.35118 | -0.23787 | -0.13019 | -0.12986 | 0.07138 | 0.16286 | 0.01907 | -0.00293 | 0.115546 | -0.07937 | 0.09991 | 0.100512 | -0.1245 | -0.24012 | -0.12569 | -0.03343 | -0.12528 | -0.00952 | 0.10394 | 0.13414 |
| SH4 | 0.257558 | 0.038811 | -0.03462 | 0.07547 | -0.04375 | -0.06947 | 0.012996 | -0.04402 | -0.07329 | -0.04412 | 0.18521 | -0.02204 | -0.02089 | 0.100492 | -0.06267 | 0.166246 | 0.102653 | -0.24441 | -0.28296 | 0.01621 | -0.08874 | -0.00101 | 0.0158 | 0.02526 | 0.13923 |
| RERFLCSQ1 | -0.18639 | 0.053951 | -0.10006 | 0.09126 | -0.0217 | -1.22434 | 0.099102 | -0.04791 | -0.13965 | 0.28191 | 0.19402 | 0.00372 | -0.01988 | 0.093795 | -0.6823 | -0.08676 | 0.024066 | -0.96051 | -1.09955 | -0.01005 | -0.00047 | -0.02067 | -0.14953 | -0.12073 | -0.02368 |
| LU99 | -0.06129 | -0.03357 | 0.03827 | -0.06373 | -0.0175 | -0.28857 | 0.104008 | -0.13649 | -0.0751 | -0.00493 | 0.08676 | 0.17023 | 0.015098 | 0.124027 | -0.06295 | 0.04251 | 0.005505 | -0.12414 | -0.30025 | -0.1756 | -0.59513 | -0.04506 | -0.04734 | -0.01195 | -0.04715 |
| KNS60 | -0.10611 | -0.56317 | 0.23001 | -0.32204 | -0.06522 | -0.00837 | -0.16101 | -0.14382 | -0.04433 | 0.12277 | 0.16128 | 0.08449 | -0.00889 | 0.197271 | 0.013148 | 0.104514 | 0.110139 | -0.11516 | -0.37415 | -0.13669 | -1.05171 | -0.1203 | -0.27179 | 0.09997 | -0.03179 |
| NCIH1666 | 0.010362 | 0.086104 | -0.13441 | -0.02217 | -0.09771 | -1.58902 | -0.00919 | 0.09153 | 0.01204 | 0.11653 | 0.11559 | 0.0137 | 0.082582 | 0.115913 | 0.035593 | 0.140191 | 0.188069 | -0.08405 | -0.18875 | -0.02366 | -0.05534 | -0.02464 | 0.08219 | 0.0396 | 0.2027 |
| MELHO | -0.04672 | 0.032299 | 0.18555 | 0.14423 | 0.20088 | -0.21355 | -0.10173 | -0.01005 | 0.01027 | -0.27075 | 0.17838 | 0.29781 | 0.190679 | -0.09918 | 0.224064 | -0.01305 | 0.049459 | -0.20178 | -0.19927 | -0.4604 | 0.03566 | -0.07561 | -0.06208 | 0.06026 | 0.32 |
| TE8 | 0.130514 | 0.005123 | -0.01826 | 0.09539 | -0.08719 | -0.90534 | 0.10204 | -0.00951 | -0.0607 | 0.16991 | 0.09578 | 0.02984 | -0.04367 | 0.07732 | -0.08345 | 0.041257 | 0.161005 | -0.29699 | -0.24249 | -0.16356 | -0.51931 | -0.0933 | -0.03141 | 0.04048 | 0.11098 |
| HCC95 | -0.0545 | 0.159076 | -0.06875 | 0.11863 | -0.11415 | -0.06829 | -0.05708 | 0.17416 | -0.11424 | 0.06486 | 0.20515 | 0.0125 | 0.036077 | 0.171048 | -0.06624 | 0.059627 | 0.151081 | -0.99943 | -0.89255 | 0.085519 | -0.28429 | 0.00608 | 0.04529 | -0.10219 | 0.12273 |
| LN428 | 0.109219 | 0.062416 | -0.01709 | 0.10501 | -0.15205 | -0.33142 | 0.017326 | 0.03867 | 0.01141 | -0.06574 | 0.13603 | 0.09397 | 0.016718 | 0.102403 | -0.18473 | 0.067737 | 0.002846 | -0.25289 | -0.16128 | 0.036841 | -0.0805 | -0.03664 | -0.01252 | 0.07275 | 0.26734 |
| BCPAP | 0.042056 | -0.11479 | 0.0854 | -0.05921 | -0.02897 | -0.14004 | -0.10968 | -0.13138 | -0.01618 | -0.00245 | 0.10161 | 0.04562 | 0.008262 | 0.013081 | -0.03636 | 0.16819 | 0.049929 | -0.23093 | -0.19678 | -0.064 | 0.02685 | 0.00817 | -0.08133 | -0.0205 | 0.19864 |
| CJM | 0.081577 | -0.00582 | -0.09818 | -0.04779 | -0.09269 | -0.0054 | 0.043999 | 0.03642 | -0.13754 | 0.06906 | 0.00355 | -0.0233 | 0.154119 | -0.0213 | 0.078123 | -0.02794 | 0.355241 | -0.34119 | -0.33242 | -0.18469 | 0.157269 | -0.12992 | -0.04642 | -0.10054 | 0.04621 |
| TUHR10TKB | -0.0135 | -0.12141 | -0.22257 | -0.17878 | -0.3267 | -0.11015 | -0.06679 | 0.16169 | 0.10589 | 0.14788 | 0.0473 | -0.004 | -0.01083 | 0.122126 | -0.49918 | 0.341799 | 0.002695 | -0.26543 | -0.13923 | -0.18687 | -0.11978 | -0.29504 | 0.05729 | -0.13843 | 0.00338 |
| SNU8 | -0.03151 | 0.003696 | -0.00142 | 0.10895 | 0.07571 | -1.08108 | 0.094467 | 0.38779 | -0.06571 | 0.04944 | 0.116 | 0.07171 | 0.072795 | 0.127677 | -0.0694 | 0.111163 | 0.044751 | -0.904 | -0.64854 | -0.11148 | -0.11123 | -0.14255 | 0.08394 | 0.05258 | 0.09323 |
| SNU1196 | 0.079017 | 0.001852 | -0.17947 | 0.14495 | -0.0429 | -0.46222 | 0.005494 | 0.01624 | 0.02417 | 0.12766 | 0.09203 | 0.23644 | 0.053432 | 0.058784 | -0.0495 | -0.04781 | -0.01298 | -0.3695 | -0.27645 | -0.21706 | -0.20665 | 0.02993 | 0.14616 | 0.1342 | -0.03692 |
| NCIH460 | -0.05301 | -0.02325 | 0.09554 | 0.03214 | -0.06453 | -0.08954 | -0.01014 | 0.06708 | -0.06194 | 0.03433 | 0.22754 | 0.00271 | 0.151445 | 0.142084 | -0.09262 | 0.06544 | -0.03023 | -0.20665 | -0.19898 | 0.136105 | 0.040544 | -0.03293 | -0.0355 | 0.06813 | 0.10345 |
| CAS1 | 0.16657 | -0.12275 | 0.04633 | 0.06487 | -0.02345 | 0.04566 | 0.080444 | 0.10798 | 0.13644 | -0.00553 | -0.04514 | 0.27294 | 0.19539 | 0.099782 | -0.08445 | 0.091683 | 0.09936 | -0.25611 | -0.2876 | -0.2998 | -0.39628 | -0.05896 | -0.05255 | 0.10026 | 0.24274 |
| SNU216 | -0.15268 | -0.07965 | 0.16395 | -0.04399 | -0.15187 | -0.45688 | -0.04479 | 0.13252 | -0.01083 | -0.00335 | 0.23016 | -0.03615 | -0.00314 | 0.126989 | -0.04071 | 0.032176 | -0.04709 | -1.75326 | -0.65718 | -0.30507 | -0.07161 | 0.08224 | 0.02411 | 0.08996 | 0.03368 |
| HCC56 | -0.04102 | 0.239144 | 0.03512 | -0.2003 | -0.14242 | -0.33723 | 0.235457 | 0.08361 | -0.08276 | 0.16252 | 0.13045 | -0.11539 | -0.0214 | -0.03739 | -0.24638 | 0.168919 | 0.057369 | -0.12357 | -0.1877 | -0.08546 | 0.038081 | -0.16039 | 0.08158 | 0.01746 | 0.20407 |
| PK45H | -0.1348 | -0.49705 | -0.03226 | -0.17702 | -0.03047 | -0.02248 | -0.10162 | 0.34929 | 0.10268 | 0.2328 | 0.19103 | 0.11016 | -0.16488 | -0.12001 | -0.174 | -0.07481 | -0.03602 | -0.39878 | -0.17879 | -0.17202 | -0.18926 | 0.03244 | 0.00625 | -0.07403 | 0.02983 |
| YH13 | -0.01989 | 0.031997 | -0.01587 | -0.08362 | 0.01481 | -0.02248 | 0.044708 | 0.09758 | 0.00312 | 0.15822 | 0.02451 | 0.20144 | 0.001274 | 0.116644 | -0.08089 | 0.117091 | 0.06765 | -0.11551 | -0.08125 | -0.06564 | -0.92824 | -0.02455 | 0.14373 | -0.00331 | 0.10921 |
| SW1463 | 0.168945 | 0.040146 | 0.03893 | -0.11641 | 0.05658 | -0.37733 | 0.080133 | 0.02 | -0.03247 | 0.21737 | 0.24753 | -0.18951 | 0.104829 | 0.028634 | -0.05159 | 0.014347 | 0.069653 | -0.04562 | -0.21128 | 0.034897 | -0.00213 | -0.09214 | 0.18371 | -0.05426 | 0.30025 |
| LI7 | -0.09861 | 0.141106 | 0.06356 | -0.09816 | -0.08415 | -0.18487 | -0.13283 | 0.17175 | -0.04338 | 0.07327 | 0.04434 | 0.06681 | 0.115953 | 0.107523 | 0.122851 | 0.022929 | 0.077582 | -0.26744 | -0.14701 | 0.022968 | -2.48857 | -0.0331 | 0.02475 | 0.00756 | 0.16136 |
| HSC2 | -0.04296 | 0.026446 | -0.07743 | 0.01635 | -0.01789 | -1.54666 | 0.088159 | 0.00122 | 0.01443 | 0.06385 | 0.08588 | 0.03969 | 0.033673 | -0.0231 | -0.03289 | 0.033661 | 0.026279 | -0.22643 | -0.30375 | -0.09385 | -0.00159 | -0.02251 | 0.00656 | 0.03533 | 0.20902 |
| RT112 | 0.116827 | 0.053302 | -0.00268 | 0.05631 | -0.13311 | -0.06399 | 0.063559 | 0.42126 | 0.03368 | 0.31094 | 0.16067 | -0.05576 | 0.044276 | 0.242994 | -0.17121 | -0.07615 | 0.257576 | -0.32583 | -0.13594 | 0.108801 | 0.099543 | 0.15999 | -0.82543 | -0.19378 | -0.20298 |

| Cell line | ALK | AXL | CSF1R | DDR1 | DDR2 | EGFR | EPHA1 | EPHA2 | EPHA3 | EPHA4 | EPHA6 | EPHA7 | EPHB1 | EPHB2 | EPHB3 | EPHB4 | EPHB6 | ERBB2 | ERBB3 | ERBB4 | FGFR1 | FGFR2 | FGFR3 | FGFR4 | FLT1 |
|---|---|---|---|---|---|---|---|---|---|---|---|---|---|---|---|---|---|---|---|---|---|---|---|---|---|
| HUH1 | 0.076016 | -0.03292 | -0.09285 | -0.01811 | -0.10299 | -0.42776 | -0.1188 | -0.05324 | -0.06621 | 0.06489 | 0.0333 | 0.0809 | 0.035886 | 0.054271 | -0.19358 | 0.009189 | 0.036781 | -0.37729 | -0.32901 | 0.013231 | -0.08552 | -0.0375 | 0.03754 | -0.26556 | 0.116 |
| JHH4 | 0.095201 | -0.08598 | 0.01271 | -0.05972 | 0.07217 | -0.19555 | 0.039781 | 0.00262 | -0.04018 | -0.02139 | 0.12942 | 0.23742 | -0.02175 | -0.00858 | -0.07548 | -0.13278 | 0.078711 | -0.53583 | -0.66213 | -0.08377 | -0.26225 | -0.25208 | 0.05308 | 0.08235 | 0.20791 |
| MALME3M | -0.32861 | 0.137075 | 0.1477 | 0.03229 | -0.00599 | 0.13226 | -0.17357 | -0.19759 | 0.00908 | -0.11064 | 0.37252 | 0.12331 | -0.0614 | -0.07884 | 0.361931 | 0.222298 | 0.125672 | -0.25148 | -0.32021 | -0.49402 | -0.09696 | 0.15221 | -0.2119 | 0.10352 | 0.01311 |
| SNU387 | 0.049196 | -0.05891 | 0.06822 | 0.02946 | -0.03725 | -0.16084 | -0.06705 | 0.07487 | 0.0373 | 0.14235 | 0.12436 | 0.06724 | 0.031588 | 0.032541 | -0.158 | -0.13228 | 0.131037 | -0.32991 | -0.10291 | -0.05733 | -0.83568 | -0.06399 | -0.09579 | 0.04995 | 0.09806 |
| KNS81 | -0.03443 | -0.16407 | -0.0047 | 0.03894 | -0.00052 | 0.01382 | 0.092902 | 0.30755 | -0.06507 | 0.10093 | 0.26961 | 0.11633 | 0.002592 | 0.183212 | 0.356055 | 0.280168 | 0.258184 | -0.23268 | -0.18812 | -0.29268 | 0.074875 | 0.09525 | -0.02459 | -0.1342 | 0.07197 |
| HUH7 | -0.03494 | -0.07748 | 0.03681 | -0.00791 | -0.13714 | -0.03691 | 0.174791 | 0.18679 | 0.04231 | 0.18907 | 0.12089 | -0.0476 | 0.022392 | 0.213275 | -0.14015 | -0.05297 | 0.12647 | -0.25858 | -0.26469 | 0.038134 | -0.03482 | -0.10815 | -0.03114 | -0.85697 | 0.05973 |
| NCIH2170 | 0.052981 | -0.04438 | 0.01652 | 0.04511 | -0.14336 | -0.0926 | -0.10298 | 0.13663 | -0.07155 | 0.24622 | 0.15185 | -0.06913 | 0.021018 | 0.001356 | -0.06895 | -0.05143 | -0.04584 | -1.91503 | -0.59986 | -0.00066 | -0.06029 | 0.03074 | 0.1211 | -0.03418 | 0.12435 |
| SNU182 | 0.014887 | 0.098233 | 0.07522 | 0.13902 | 0.00256 | -0.32088 | -0.0533 | 0.04136 | 0.08033 | 0.06028 | -0.02544 | -0.0356 | 0.032896 | 0.004607 | 0.123937 | 0.002901 | 0.478729 | -0.35051 | -0.2282 | -0.30653 | -0.10832 | 0.05393 | -0.15502 | 0.05135 | 0.00687 |
| VMRCRCW | -0.09563 | 0.281315 | -0.02584 | 0.0652 | -0.11017 | 0.05093 | -0.00579 | -0.3155 | 0.01071 | 0.065 | 0.11238 | 0.05443 | 0.035217 | -0.12331 | -0.04316 | 0.045657 | -0.153 | -0.01494 | -0.3294 | -0.27531 | 0.079166 | 0.13701 | 0.19742 | -0.05805 | 0.15631 |
| GSU | 0.095003 | -0.02438 | 0.07337 | -0.026 | 0.06908 | -0.56273 | 0.055681 | -0.07715 | 0.1503 | 0.14162 | 0.16557 | 0.1286 | 0.032294 | 0.067201 | 0.032362 | 0.00908 | 0.006729 | -0.51006 | -0.10191 | -0.10284 | -0.04213 | -0.13148 | 0.04173 | -0.04246 | 0.06898 |
| KU1919 | -0.07217 | 0.047621 | 0.08809 | -0.13528 | -0.10811 | -0.13431 | -0.23196 | -0.04349 | -0.04852 | 0.02302 | 0.05008 | 0.0977 | 0.042004 | 0.010811 | 0.014168 | -0.05999 | 0.078326 | -0.15028 | -0.28086 | -0.11114 | -0.094 | 0.04695 | 0.04737 | 0.07566 | 0.1939 |
| F36P | 0.079895 | -1.05821 | -0.12568 | -0.04853 | -0.17708 | -0.20651 | -0.00383 | -0.23733 | -0.00339 | 0.06646 | 0.09949 | 0.20064 | 0.057265 | -0.14063 | -0.05233 | 0.032556 | 0.047945 | -0.15639 | -0.1134 | 0.026138 | -0.0863 | -0.03997 | -0.05758 | 0.01244 | 0.11893 |
| TE11 | 0.012877 | 0.064204 | 0.00303 | -0.05977 | -0.00347 | -0.42388 | -0.01559 | 0.05075 | 0.11682 | 0.03183 | 0.1182 | 0.05578 | -0.03986 | 0.070491 | -0.07215 | 0.050301 | 0.152315 | -0.499 | -0.65689 | -0.11086 | -0.00121 | 0.00458 | -0.05964 | -0.07832 | 0.14545 |
| SW1116 | 0.096159 | -0.09013 | -0.00151 | -0.08714 | 0.01049 | -0.70553 | 0.091409 | 0.1279 | 0.00602 | -0.08514 | 0.17588 | 0.10007 | -0.00536 | 0.029159 | -0.14062 | -0.01092 | -0.00828 | -0.87772 | -0.20886 | -0.00682 | -0.06595 | -0.07314 | -0.09988 | 0.04502 | 0.1874 |
| SF767 | 0.07246 | -0.00134 | -0.01643 | 0.08656 | 0.04836 | -0.45652 | 0.236654 | 0.05177 | -0.06979 | 0.07269 | 0.26215 | 0.07472 | 0.09854 | -0.03852 | -0.02909 | -0.01977 | 0.039679 | -0.78201 | -0.87983 | -0.20963 | -0.07614 | -0.2483 | -0.17643 | 0.10122 | -0.12374 |
| NCIH716 | 0.080029 | 0.053104 | -0.06588 | -0.07679 | -0.34702 | 0.14849 | -0.27115 | -0.11129 | -0.11441 | -0.05293 | 0.20898 | 0.11529 | 0.115654 | 0.024062 | -0.20179 | 0.169988 | -0.12033 | -0.34914 | -0.31089 | 0.052869 | 0.002446 | -0.31542 | 0.13167 | -0.11397 | 0.24208 |
| SNU423 | -0.16878 | -0.3698 | -0.06427 | -0.05172 | 0.05135 | -0.03796 | -0.14659 | 0.09968 | -0.08273 | 0.1199 | 0.18558 | -0.02861 | -0.13102 | 0.149167 | 0.030334 | -0.08725 | 0.070479 | -0.28689 | -0.29049 | -0.0701 | -0.91493 | -0.06646 | 0.05018 | 0.11959 | 0.23617 |
| TUHR4TKB | 0.05414 | 0.031062 | -0.06954 | 0.00694 | -0.03503 | -0.19388 | 0.092874 | 0.13278 | -0.0414 | -0.09934 | 0.14447 | 0.06163 | 0.028218 | -0.12524 | 0.011868 | 0.022441 | 0.00978 | -0.31774 | -0.24898 | -0.23349 | -0.21872 | 0.01652 | 0.00377 | 0.1415 | 0.01911 |
| NCIH1792 | 0.014002 | 0.009704 | -0.21048 | 0.01152 | 0.05357 | -0.59956 | 0.021549 | 0.08549 | -0.23704 | 0.03404 | 0.15186 | 0.10912 | 0.051786 | 0.092466 | -0.10009 | -0.26787 | 0.093789 | -0.21942 | -0.10724 | -0.10619 | -0.44356 | -0.01547 | 0.11567 | 0.03132 | -0.10663 |
| EW8 | 0.048508 | 0.041818 | 0.18226 | -0.03587 | 0.01821 | -0.21834 | 0.230157 | 0.09385 | -0.12071 | -0.12378 | -0.02471 | -0.15944 | -0.03964 | 0.036582 | 0.089075 | 0.221694 | 0.094401 | -0.50112 | -0.13739 | -0.2384 | 0.159757 | 0.05932 | -0.04338 | 0.11565 | -0.08335 |
| SNU46 | -0.02299 | 0.107411 | 0.00042 | 0.06914 | 0.10757 | -0.12521 | 0.148707 | -0.28633 | -0.03959 | -0.15881 | 0.26047 | 0.22435 | 0.087192 | 0.031576 | -0.23025 | -0.22551 | 0.148688 | -0.58385 | 0.02241 | -0.19159 | -0.25479 | -0.27612 | -0.12029 | -0.1298 | -0.26117 |
| LS123 | 0.208165 | 0.006674 | 0.03159 | 0.17271 | -0.16568 | -0.36781 | -0.043 | 0.29596 | -0.07107 | 0.19831 | -0.12275 | 0.34332 | -0.05526 | 0.420948 | 0.159969 | 0.116821 | 0.263174 | -0.24217 | -0.20096 | 0.10274 | -0.21956 | -0.00903 | 0.22102 | 0.19532 | 0.27932 |
| TCCPAN2 | 0.097479 | 0.083928 | -0.08802 | -0.03299 | 0.02129 | -0.20752 | 0.061754 | -0.10626 | -0.08001 | 0.02533 | 0.12335 | 0.18832 | 0.006988 | -0.03244 | 0.041375 | -0.01382 | 0.046861 | -0.23927 | -0.17217 | -0.05134 | -0.03555 | -0.00828 | -0.15282 | -0.04158 | 0.10995 |
| BICR16 | -0.0619 | 0.118753 | -0.09494 | 0.00147 | -0.01744 | -0.30333 | -0.07922 | 0.23118 | -0.1194 | 0.15901 | -0.15747 | -0.07992 | 0.284043 | 0.165826 | 0.314065 | 0.12338 | 0.127363 | -0.86993 | -0.70236 | -0.05701 | -0.12552 | 0.02637 | -0.01641 | 0.19184 | 0.12313 |
| SNB75 | -0.17151 | -0.25332 | 0.07699 | -0.13061 | 0.03592 | -0.07822 | 0.145492 | -0.00017 | -0.04544 | 0.01397 | -0.0185 | -0.00442 | -0.17004 | 0.066474 | -0.27353 | 0.072556 | 0.019544 | -0.23817 | -0.35617 | -0.2702 | -0.21746 | -0.12829 | -0.08507 | 0.09598 | 0.06755 |
| RKN | -0.10443 | -0.10074 | -0.03723 | -0.0138 | 0.04554 | -0.10565 | 0.084246 | -0.01069 | -0.10354 | 0.03963 | 0.12657 | 0.1398 | -0.01443 | 0.065281 | -0.04153 | 0.023904 | 0.141804 | -0.19137 | -0.13025 | -0.19468 | -0.10174 | -0.06929 | -0.06002 | 0.01881 | 0.05067 |
| KE39 | -0.03857 | 0.153747 | -0.02736 | -0.05307 | 0.02798 | -0.39349 | 0.070219 | -0.19385 | 0.01424 | 0.10803 | 0.07278 | 0.09767 | 0.061825 | 0.170306 | 0.10538 | 0.102361 | 0.251156 | -0.42577 | -0.22177 | -0.08839 | -0.18992 | 0.04252 | 0.08072 | 0.17349 | -0.05056 |
| NCIH1299 | -0.00991 | -0.15797 | 0.01679 | -0.09192 | -0.11514 | -0.12248 | 0.008199 | 0.01611 | -0.02367 | 0.18168 | 0.05202 | 0.02594 | -0.02318 | 0.128817 | -0.16587 | -0.11275 | 0.099054 | -0.31095 | -0.09313 | -0.10338 | 0.011557 | -0.01502 | -0.06631 | 0.0193 | 0.12289 |
| CALU1 | 0.050788 | -0.21383 | 0.09796 | -0.14743 | 0.12094 | -0.23393 | 0.094332 | -0.13248 | 0.04944 | 0.15781 | 0.06496 | 0.06971 | 0.085546 | -0.05415 | 0.007509 | 0.081229 | 0.19153 | -0.14538 | -0.10109 | -0.25338 | -0.1606 | -0.03786 | 0.00471 | -0.0437 | 0.05135 |
| INA6 | -0.04316 | 0.353013 | -0.21433 | -0.05893 | -0.18928 | -0.08428 | 0.135267 | -0.19655 | -0.11598 | 0.13582 | 0.21744 | -0.13145 | 0.09218 | -0.12067 | -0.36508 | -0.14449 | 0.143073 | -0.33477 | -0.25249 | 0.148501 | -0.01762 | 0.05356 | -0.32577 | -0.02989 | 0.08261 |
| NCIH1092 | -0.16515 | -0.01972 | -0.04258 | -0.12102 | 0.06974 | -0.22801 | 0.029265 | -0.3254 | -0.02211 | 0.17113 | 0.02478 | 0.13165 | -0.01884 | -0.11885 | -0.11737 | 0.097808 | 0.004762 | -0.29937 | -0.1501 | -0.17482 | -0.18844 | -0.03794 | -0.0458 | 0.19767 | -0.05426 |
| CAL78 | -0.05774 | -0.16973 | 0.08289 | -0.04832 | 0.01903 | -0.19224 | -0.07689 | 0.06052 | 0.02035 | -0.00243 | 0.15283 | 0.08264 | 0.009214 | 0.101081 | 0.168177 | 0.021475 | 0.069683 | -0.14943 | -0.29355 | -0.12075 | -0.23665 | -0.00896 | -0.01064 | 0.0109 | -0.01064 |
| SNU410 | -0.0087 | -0.15461 | -0.00109 | 0.00124 | -0.02836 | -0.14658 | 0.003963 | 0.00026 | 0.00864 | 0.01155 | 0.0895 | 0.09762 | 0.068496 | 0.058425 | -0.06726 | -0.0482 | 0.028122 | -0.15576 | -0.21685 | -0.09694 | -0.35163 | -0.0772 | -0.04346 | 0.02306 | 0.06328 |
| CAL33 | -0.01915 | 0.066954 | 0.01141 | 0.13237 | -0.02512 | -0.74479 | 0.086608 | 0.20893 | 0.05954 | 0.16419 | -0.11249 | 0.23201 | 0.005915 | 0.048716 | 0.105781 | 0.056632 | 0.040533 | -0.3482 | -0.28455 | -0.20572 | 0.086031 | -0.04747 | 0.0253 | 0.09203 | 0.07601 |
| 59M | 0.136204 | 0.038456 | -0.06349 | 0.13122 | -0.00887 | -0.2152 | 0.114555 | -0.10583 | -0.08 | 0.18542 | 0.07398 | 0.11429 | -0.0801 | -0.05042 | -0.01985 | -0.01098 | 0.100853 | -0.26687 | -0.21704 | -0.17746 | -0.89975 | 0.00682 | -0.02767 | 0.16254 | 0.05476 |
| NCIH2030 | -0.09671 | 0.022907 | 0.09617 | 0.04307 | 0.00248 | -0.4511 | 0.130111 | 0.41702 | -0.11707 | -0.04996 | 0.15498 | -0.07119 | -0.05405 | -0.12868 | -0.05063 | 0.047908 | 0.106526 | -0.60933 | -0.37517 | -0.1207 | -0.33531 | -0.02373 | 0.02295 | -0.02772 | 0.18753 |
| UMUC3 | 0.013936 | -0.01024 | -0.0145 | -0.21464 | -0.01981 | -0.21777 | -0.02071 | 0.1694 | -0.17026 | 0.05053 | 0.06119 | -0.05271 | 0.01346 | 0.123734 | -0.16587 | 0.044715 | 0.121705 | -0.22118 | -0.26423 | -0.09997 | -0.13135 | -0.16514 | -0.06211 | 0.0379 | 0.04634 |
| KURAMOCHI | -0.04749 | -0.00566 | 0.01488 | -0.04698 | 0.02174 | -0.30153 | 0.027806 | -0.04374 | 0.04428 | 0.05311 | 0.16693 | 0.18083 | 0.06462 | 0.061469 | -0.08791 | -0.0777 | 0.009689 | -0.41141 | -0.14358 | -0.26128 | -0.00279 | -0.06697 | 0.06698 | -0.03443 | 0.03475 |
| NCIH2171 | 0.115344 | 0.010546 | 0.02856 | 0.01479 | -0.03932 | -0.17667 | 0.027682 | -0.23977 | 0.0846 | 0.15874 | 0.14931 | 0.08715 | -0.0133 | 0.027571 | -0.13049 | 0.058539 | 0.084125 | -0.18392 | -0.15578 | 0.00111 | 0.032527 | -0.03217 | -0.03415 | -0.07413 | 0.12041 |
| OVISE | 0.067792 | 0.121332 | 0.07922 | -0.06218 | -0.01233 | -0.33618 | 0.021718 | -0.00139 | -0.01752 | 0.0949 | 0.13286 | -0.06358 | 0.04474 | 0.153541 | -0.08359 | 0.042551 | 0.022655 | -0.37258 | -0.41284 | -0.05914 | -0.0075 | 0.03499 | 0.13804 | -0.02774 | 0.23833 |
| ABC1 | 0.058202 | 0.092168 | -0.06652 | -0.00648 | -0.10521 | -0.19707 | 0.080527 | 0.09576 | 0.14971 | 0.06918 | 0.14375 | 0.05844 | 0.047107 | 0.128787 | -0.07465 | 0.01544 | 0.140893 | -0.11573 | -0.12881 | -0.09748 | -0.04258 | -0.03673 | 0.02448 | 0.02895 | 0.0316 |
| SNU61 | -0.04013 | 0.061272 | 0.00973 | 0.05734 | -0.02282 | -1.04616 | 0.115602 | 0.10173 | -0.21465 | -0.03487 | 0.29761 | 0.15974 | -0.04852 | 0.110781 | -0.06448 | 0.115448 | 0.014247 | -0.58229 | -0.13231 | 0.016505 | -0.04597 | 0.0231 | 0.05008 | -0.04205 | 0.06534 |
| NCIH2004RT | -0.06032 | 0.036508 | 0.01507 | -0.01164 | -0.02117 | -0.17711 | -0.0324 | 0.02775 | 0.16957 | -0.06042 | 0.24828 | 0.10025 | 0.001408 | 0.1506 | 0.014579 | 0.131543 | -0.09208 | -0.22595 | -0.18811 | 0.292286 | -0.72693 | 0.13091 | 0.10634 | 0.01309 | 0.04618 |
| BXPC3 | -0.09124 | 0.092092 | -0.16492 | 0.04274 | -0.03731 | -0.05105 | 0.087008 | 0.06218 | -0.00811 | 0.11715 | 0.11318 | 0.15446 | 0.029461 | 0.065148 | -0.19539 | -0.04125 | 0.103146 | -1.05445 | -0.84716 | -0.07951 | 0.11481 | -0.08961 | -0.01553 | 0.08348 | 0.1394 |
| SNU761 | 0.100613 | 0.03259 | 0.08487 | -0.00613 | -0.06239 | -1.41905 | -0.04882 | 0.36812 | 0.10425 | 0.06008 | 0.05086 | 0.06198 | -0.01301 | 0.042506 | 0.047003 | -0.02077 | 0.22088 | -0.26824 | -0.15046 | -0.01732 | -0.02391 | -0.05156 | 0.03435 | 0.08832 | 0.14154 |
| KMS34 | -0.04508 | 0.007758 | 0.20705 | -0.02855 | 0.09736 | -0.17803 | -0.00584 | -0.16526 | -0.01486 | 0.03442 | 0.30028 | 0.02227 | -0.06962 | -0.08144 | -0.22736 | 0.140618 | 0.220734 | -0.12003 | -0.22298 | 0.029167 | 0.077595 | -0.20607 | -0.7996 | 0.11211 | 0.21246 |
| HEYA8 | -0.08805 | -0.14291 | -0.06137 | -0.01721 | -0.16421 | -0.22474 | 0.011068 | 0.02141 | 0.01064 | -0.00947 | 0.2 | -0.0808 | 0.100652 | 0.186445 | -0.23105 | 0.045812 | 0.090267 | -0.15914 | -0.15196 | 0.056173 | -0.06962 | -0.02734 | 0.11108 | -0.15263 | 0.22606 |
| OE21 | 0.096012 | -0.03609 | -0.19912 | 0.01137 | -0.12235 | -0.95565 | 0.029611 | 0.07566 | 0.02701 | 0.10147 | -0.0105 | 0.13318 | 0.08877 | 0.098723 | 0.047604 | 0.041664 | 0.157831 | -0.2906 | -0.88613 | 0.029746 | -0.06715 | -0.04085 | 0.10995 | -0.05078 | 0.25408 |
| VMCUB1 | -0.03443 | 0.084004 | 0.05016 | 0.07989 | 0.0035 | -0.47482 | 0.130191 | 0.14316 | -0.12593 | 0.26372 | 0.02657 | 0.03119 | 0.003599 | 0.2483 | 0.082132 | -0.10142 | 0.155724 | -0.57901 | -0.30796 | -0.1577 | -0.07943 | -0.12908 | 0.04011 | 0.19356 | -0.06895 |
| HSC4 | -0.00192 | 0.174957 | -0.14642 | 0.08275 | -0.22795 | -1.28505 | 0.056891 | 0.00945 | -0.19925 | 0.18368 | 0.04462 | 0.2259 | 0.099082 | 0.057781 | 0.210902 | 0.13536 | 0.156339 | 0.0038 | 0.00415 | -0.17677 | -0.10059 | -0.19193 | 0.27972 | 0.11977 | 0.20586 |
| HT1197 | -0.1867 | -0.00377 | 0.3043 | -0.21258 | 0.14714 | -0.06535 | 0.2718 | 0.08994 | -0.02704 | 0.03431 | 0.2068 | 0.062 | 0.169507 | 0.004099 | -0.16377 | 0.062704 | 0.408554 | -0.44703 | -0.27275 | -0.13666 | 0.051596 | 0.10813 | 0.10214 | 0.02448 | 0.02829 |
| BHY | 0.085276 | 0.02222 | -0.08339 | -0.03117 | -0.05085 | -1.91973 | 0.042014 | 0.29284 | -0.06363 | 0.10159 | 0.10595 | 0.06556 | 0.010758 | 0.119054 | 0.021326 | 0.116889 | 0.186677 | -0.32968 | -0.23853 | -0.16623 | -0.01711 | 0.02914 | -0.05352 | 0.05365 | 0.11059 |
| SNU1076 | 0.142884 | 0.065006 | -0.20176 | -0.10144 | -0.22184 | -0.45653 | 0.117346 | 0.03343 | -0.1451 | 0.04372 | 0.3547 | 0.00499 | 0.015497 | 0.101366 | -0.08308 | 0.167555 | 0.26541 | -0.82637 | -0.56608 | -0.04758 | -0.02944 | -0.07205 | 0.06685 | 0.12018 | 0.0945 |
| IGR39 | -0.02708 | -0.10568 | 0.06742 | 0.02919 | -0.00744 | -0.15177 | 0.116445 | 0.34546 | -0.0031 | 0.2386 | 0.19681 | -0.12478 | -0.08522 | 0.065103 | -0.06773 | -0.07436 | 0.067038 | -0.38414 | -0.40769 | -0.12349 | -0.06944 | -0.06715 | -0.05788 | -0.04843 | 0.05057 |
| K562 | -0.12178 | 0.157982 | -0.01494 | 0.16628 | -0.08109 | -0.1803 | -0.14786 | -0.15964 | -0.08818 | 0.14093 | 0.10998 | 0.11962 | -0.05468 | -0.0226 | -0.00896 | 0.007566 | 0.022251 | -0.20635 | -0.21518 | -0.00758 | -0.08612 | 0.01513 | -0.07829 | 0.08353 | 0.05762 |
| HT29 | 0.034211 | 0.048599 | 0.06394 | -0.02332 | 0.05601 | -0.44534 | 0.052573 | 0.05511 | 0.08831 | -0.03866 | 0.13384 | 0.09523 | 0.025691 | -0.0381 | -0.14107 | -0.01446 | -0.01781 | -0.74602 | -0.13271 | -0.03783 | -0.0358 | -0.1426 | 0.04982 | 0.03336 | 0.09529 |
| UACC893 | 0.189278 | 0.077422 | -0.06616 | 0.07745 | -0.06536 | -0.30061 | -3.5E-05 | 0.13553 | -0.09778 | 0.19621 | -0.03249 | -0.04588 | 0.01766 | 0.158069 | 0.015257 | 0.225508 | 0.254494 | -2.44515 | -0.37731 | 0.029074 | -0.05137 | -0.31741 | 0.10307 | -0.00477 | 0.15849 |
| SIHA | 0.152909 | -0.08978 | 0.03486 | -0.04274 | 0.01594 | -0.21319 | -0.0184 | 0.01596 | -0.06721 | 0.02742 | 0.06476 | 0.04349 | 0.128323 | 0.108305 | -0.17999 | -0.08075 | 0.140462 | -0.29963 | -0.31477 | -0.09462 | -0.93031 | -0.01192 | 0.01003 | -0.07229 | 0.12782 |
| AML193 | 0.072218 | -0.01983 | -0.24821 | -0.10611 | -0.10606 | -0.11369 | -0.04414 | -0.33522 | -0.08655 | 0.02214 | 0.14909 | 0.14395 | 0.021276 | -0.12578 | -0.23311 | -0.02267 | -0.00838 | -0.25415 | -0.26197 | 0.059115 | -0.38522 | -0.16303 | -0.01449 | -0.03747 | 0.19064 |
| A172 | 0.015323 | -0.34579 | -0.02146 | -0.23239 | -0.06966 | -0.09434 | 0.065979 | 0.09171 | 0.0347 | 0.12774 | 0.21898 | 0.08473 | 0.06439 | 0.010225 | 0.072224 | 0.081303 | 0.070166 | -0.20293 | -0.32384 | -0.13381 | -1.80957 | 0.12109 | -0.0009 | -0.01291 | 0.08189 |
| NCIH1836 | 0.216307 | 0.058094 | -0.03412 | 0.08102 | -0.13394 | -0.02931 | -0.11881 | -0.14963 | 0.04207 | -0.06422 | 0.23348 | -0.12466 | 0.121014 | 0.083708 | -0.05692 | 0.142131 | 0.060568 | -0.38733 | -0.21925 | -0.16719 | -0.12384 | 0.0409 | 0.25173 | 0.17909 | 0.25178 |
| TDOTT | 0.092336 | -0.11379 | 0.01038 | -0.00678 | -0.13488 | -0.62565 | 0.187987 | 0.02665 | 0.00627 | 0.1337 | 0.19304 | 0.00571 | 0.006533 | 0.181569 | 0.052682 | 0.049703 | 0.157957 | -0.42372 | -0.45678 | -0.02385 | -0.09688 | 0.01323 | 0.16432 | 0.03147 | 0.32915 |
| HCC78 | -0.07873 | 0.182361 | -0.02525 | -0.0336 | 0.00198 | -0.30536 | -0.11552 | -0.22121 | 0.17435 | 0.00192 | 0.36681 | 0.16487 | 0.060266 | -0.12177 | -0.26022 | -0.17491 | 0.017091 | -0.55298 | -0.35327 | -0.23558 | -0.00021 | -0.10946 | -0.19729 | -0.02479 | 0.145 |
| EBC1 | -0.4612 | -0.13797 | -0.12594 | -0.22388 | -0.14347 | 0.03638 | 0.182375 | -0.36262 | 0.0307 | 0.21048 | 0.29492 | 0.04747 | -0.05579 | -0.10361 | -0.22763 | -0.2862 | 0.221249 | -0.24122 | -0.10001 | -0.15452 | 0.004144 | 0.08922 | 0.03936 | -0.15432 | 0.07329 |
| RCM1 | -0.05415 | 0.145985 | 0.06777 | 0.2134 | 0.00867 | -0.9922 | -0.02889 | 0.07685 | 0.09725 | 0.17388 | -0.07532 | 0.13942 | 0.020701 | -0.00532 | -0.07284 | -0.06232 | 0.00913 | -0.5876 | -0.29104 | 0.344822 | 0.128634 | 0.22301 | 0.04619 | 5E-05 | 0.10442 |
| ST486 | 0.311678 | 0.028934 | -0.08075 | 0.14046 | -0.07153 | -0.11719 | -0.30423 | -0.25823 | -0.05817 | -0.06688 | 0.18463 | 0.15099 | 0.100142 | 0.068785 | -0.13477 | -0.09065 | 0.040365 | -0.15531 | -0.18601 | 0.021978 | -0.1916 | -0.00902 | -0.02083 | 0.03922 | 0.16898 |
| YKG1 | -0.05083 | -0.24059 | 0.00247 | 0.00864 | -0.0973 | -0.16199 | 0.073318 | 0.47622 | 0.03565 | 0.14756 | 0.1105 | 0.15548 | 0.043847 | 0.193967 | 0.025466 | -0.04386 | 0.062872 | -0.24392 | -0.16601 | 0.055242 | -0.55129 | -0.05927 | 0.0666 | 0.10697 | -0.02114 |
| T98G | -0.01326 | -0.08225 | -0.08362 | -0.03218 | -0.04117 | -0.07276 | -0.10539 | 0.02406 | -0.02985 | 0.11657 | 0.10509 | 0.14618 | -0.0598 | 0.063286 | -0.01267 | -0.0715 | 0.025238 | -0.14286 | -0.17216 | -0.05466 | -0.72822 | -0.00728 | -0.03311 | -0.03713 | 0.00956 |
| MDAMB436 | -0.03632 | 0.034514 | -0.01275 | -0.06039 | -0.01902 | -0.14537 | -0.07933 | -0.00162 | -0.09066 | 0.03317 | 0.10051 | 0.05684 | 0.086866 | 0.126922 | -0.27747 | 0.141175 | -0.01673 | -0.14021 | -0.24478 | -0.06971 | -0.20864 | 0.07435 | -0.02823 | 0.11148 | 0.10961 |
| KMS27 | 0.01804 | -0.01861 | -0.15714 | 0.1102 | 0.01101 | -0.14821 | 0.071672 | -0.23282 | -0.01149 | -0.0309 | 0.19015 | 0.053 | 0.139075 | -0.10901 | 0.031883 | 0.073872 | 0.158895 | -0.1228 | -0.22975 | -0.07457 | -0.06427 | -0.24167 | 0.04423 | 0.01236 | 0.18331 |
| JHH2 | 0.051591 | -0.0236 | 0.0476 | 0.0128 | -0.0612 | -0.29021 | 0.108832 | 0.17815 | 0.04912 | 0.115 | -0.10974 | 0.07745 | 0.081848 | 0.011758 | 0.058716 | -0.07079 | 0.102416 | -0.45861 | -0.09775 | -0.04379 | -0.89011 | -0.18283 | 0.08485 | 0.18071 | 0.15721 |
| UACC257 | 0.304852 | -0.01262 | -0.10481 | 0.01587 | -0.13918 | 0.02321 | -0.00819 | -0.12735 | -0.20572 | 0.32844 | 0.14432 | -0.02358 | 0.32171 | -0.00938 | -0.10207 | 0.149362 | 0.04526 | -0.27357 | -0.2042 | -0.18848 | -0.03896 | -0.03291 | -0.01975 | -0.0814 | 0.13618 |
| C32 | 0.208938 | 0.113778 | -0.03537 | 0.29908 | -0.02492 | 0.07808 | 0.169427 | -0.19427 | -0.09782 | -0.13009 | 0.24423 | 0.08939 | 0.127718 | -0.00509 | 0.116833 | 0.294977 | 0.031834 | -0.33031 | -0.36891 | -0.17004 | 0.040741 | -0.02538 | 0.04413 | -0.00928 | 0.1813 |
| SNU16 | 0.206189 | 0.050461 | -0.13728 | 0.05661 | -0.02842 | 0.07636 | 0.020333 | -0.04559 | -0.08113 | 0.02094 | 0.05608 | 0.0405 | 0.047744 | -0.1315 | -0.0796 | 0.045247 | 0.009613 | -0.10383 | 0.0769 | -0.16891 | -0.27236 | -2.2789 | -0.14742 | 0.05675 | 0.04144 |

| Cell line | ALK | AXL | CSF1R | DDR1 | DDR2 | EGFR | EPHA1 | EPHA2 | EPHA3 | EPHA4 | EPHA6 | EPHA7 | EPHB1 | EPHB2 | EPHB3 | EPHB4 | EPHB6 | ERBB2 | ERBB3 | ERBB4 | FGFR1 | FGFR2 | FGFR3 | FGFR4 | FLT1 |
|---|---|---|---|---|---|---|---|---|---|---|---|---|---|---|---|---|---|---|---|---|---|---|---|---|---|
| JHOS4 | 0.07765 | 0.085775 | -0.06555 | -0.01152 | -0.17729 | -0.07044 | -0.05295 | 0.04807 | 0.11302 | 0.12722 | 0.14992 | 0.00454 | 0.084724 | 0.081502 | -0.1725 | -0.01135 | 0.007214 | -0.55839 | -0.42215 | 0.02302 | 0.006545 | 0.10553 | -0.00404 | -0.13145 | 0.16043 |
| EPLC272H | -0.00248 | -0.02225 | -0.05494 | -0.01137 | -0.10937 | -0.33322 | 0.058368 | 0.08928 | 0.0372 | 0.15971 | 0.23761 | 0.03834 | -0.01201 | 0.166731 | -0.10334 | 0.061039 | 0.091223 | -0.46451 | -0.55064 | -0.12484 | -0.05695 | -0.57453 | 0.02837 | -0.31374 | 0.20805 |
| NCIH1975 | -0.03163 | 0.015502 | 0.01218 | -0.02154 | -0.10734 | -2.95552 | 0.080367 | 0.18336 | -0.12348 | 0.02198 | 0.11615 | 0.04738 | -0.06063 | 0.044442 | -0.09949 | -0.18279 | -0.07034 | -0.4758 | -0.33391 | -0.26456 | -0.02315 | 0.03372 | -0.00973 | 0.0238 | 0.06481 |
| KMS26 | 0.010795 | 0.141561 | -0.03406 | -0.02812 | 0.03468 | -0.16894 | -0.04673 | -0.05943 | -0.03012 | 0.23208 | -0.01084 | 0.00165 | 0.262067 | 0.034604 | -0.02056 | -0.27507 | -0.04348 | -0.34066 | -0.37674 | -0.11303 | 0.070627 | 0.11522 | -0.40968 | -0.05214 | -0.04103 |
| NCIH1437 | 0.014282 | 0.09976 | -0.0043 | -0.07261 | 0.16081 | -0.10553 | 0.084788 | -0.17439 | -0.04095 | -0.04204 | 0.0611 | 0.05979 | 0.018115 | 0.021131 | -0.07828 | 0.089959 | 0.029701 | -0.18138 | -0.12655 | -0.03493 | 0.064958 | -0.09243 | 0.0007 | -0.06786 | 0.16455 |
| LN235 | -0.09403 | -0.14529 | -0.04438 | 0.02037 | 0.00639 | -0.02337 | 0.202173 | 0.19041 | 0.04785 | 0.21382 | 0.10147 | 0.15681 | -0.15778 | 0.024949 | 0.041028 | -0.13232 | 0.225917 | -0.12848 | -0.25204 | -0.33712 | -0.56404 | 0.06615 | -0.09081 | -0.12718 | -0.052 |
| TM31 | 0.082112 | -0.13684 | 0.04704 | -0.19217 | -0.06111 | -0.12807 | 0.057774 | -0.01193 | -0.00076 | 0.13253 | 0.08784 | 0.0461 | -0.0254 | 0.016378 | -0.22125 | -0.06388 | 0.234656 | -0.36972 | -0.23475 | -0.16522 | -0.67952 | -0.06665 | -0.00888 | 0.0408 | 0.12494 |
| BC3C | 0.035455 | 0.039948 | -0.14762 | -0.05564 | -0.10202 | -0.74421 | -0.09062 | 0.3965 | 0.07762 | 0.21293 | 0.27408 | 0.04786 | -0.17933 | 0.122729 | -0.10963 | 0.008517 | 0.158416 | -0.23945 | -0.17773 | -0.03046 | -0.2038 | -0.10379 | -0.01574 | -0.02899 | 0.13452 |
| LN229 | 0.052398 | -0.12425 | 0.05343 | 0.03532 | 0.02185 | -0.32292 | 0.008 | 0.05382 | 0.07753 | 0.10622 | 0.14848 | 0.0403 | 0.080565 | 0.019592 | -0.06384 | -0.02039 | -0.03321 | -1.71635 | -0.63856 | -0.09812 | -0.05784 | -0.01012 | -0.02549 | 0.01173 | 0.23507 |
| LCLC97TM1 | 0.186004 | -0.01677 | 0.10864 | 0.1227 | -0.05847 | -0.10303 | -0.26006 | 0.02077 | 0.01192 | 0.09648 | 0.02985 | 0.10914 | 0.055825 | 0.314518 | 0.015607 | -0.00103 | 0.103296 | -0.22707 | -0.08705 | -0.25355 | 0.09788 | 0.08012 | -0.23796 | 0.01083 | -0.02637 |
| TTC709 | -0.082 | 0.146236 | 0.12453 | 0.05531 | -0.05898 | -0.14443 | -0.08531 | -0.09735 | 0.04126 | 0.04972 | 0.15901 | -0.54048 | 0.00709 | 0.07128 | 0.030828 | 0.022806 | 0.089373 | -0.12442 | -0.32072 | -0.05363 | 0.138207 | -0.71262 | 0.05788 | 0.14618 | -0.05329 |
| PATU8902 | -4.9E-05 | -0.07302 | 0.00694 | -0.08809 | 0.05552 | -0.21763 | 0.044558 | -0.07595 | -0.22568 | 0.02679 | 0.30503 | 0.04658 | 0.053367 | 0.0804 | -0.35129 | 0.136768 | 0.109903 | -0.41197 | -0.08157 | 0.015386 | 0.062516 | -0.06543 | 0.14248 | -0.11897 | 0.25913 |
| SLR26 | 0.15289 | 0.212554 | -0.02931 | -0.14977 | -0.10963 | -0.32745 | -0.10274 | 0.26691 | 0.10638 | 0.12221 | 0.1854 | 0.30184 | 0.01155 | 0.234929 | -0.11888 | -0.02822 | -0.05083 | -0.02486 | -0.46509 | 0.000258 | -0.29162 | -0.01232 | 0.07099 | 0.06473 | 0.09795 |
| MIAPACA2 | -0.02834 | 0.039277 | -0.0111 | 0.00247 | 0.04704 | -0.10712 | -0.00165 | -0.04655 | 0.02186 | 0.07945 | 0.19099 | 0.11938 | 0.042233 | 0.173456 | -0.07318 | 0.058081 | 0.040729 | -0.12196 | -0.14825 | -0.00651 | -0.08352 | -0.1345 | -0.07644 | -0.05831 | 0.11831 |
| M07E | -0.12485 | 0.001435 | -0.05635 | -0.08045 | -0.01247 | -0.2529 | -0.12207 | -0.18036 | -0.03415 | -0.03617 | 0.13009 | 0.06045 | -0.07229 | 0.046075 | -0.02266 | 0.086321 | -0.09639 | -0.24308 | -0.23717 | 0.082352 | -0.01697 | -0.00613 | 0.0664 | -0.03811 | 0.16532 |
| KYO1 | 0.160755 | 0.330972 | 0.20965 | -0.07511 | 0.02334 | -0.11847 | -0.00379 | 0.106 | 0.0621 | -0.21249 | 0.27082 | 0.00917 | 0.05981 | 0.240605 | -0.21439 | 0.309132 | -0.10199 | -0.32767 | -0.36717 | -0.26697 | -0.11738 | 0.13763 | 0.16489 | -0.13641 | 0.00632 |
| TE6 | -0.15108 | -0.09877 | -0.01358 | 0.07866 | -0.02079 | -0.42367 | -0.12892 | 0.01508 | -0.11264 | 0.05329 | 0.1303 | 0.22356 | 0.008572 | -0.08303 | -0.17053 | -0.00767 | 0.144033 | -0.89689 | -0.72402 | -0.30898 | 0.038183 | 0.05162 | -0.02087 | 0.00922 | 0.1074 |
| PECAPJ34CLONEC12 | 0.145534 | 0.058753 | -0.04559 | -0.03505 | -0.06831 | -0.64061 | -0.0785 | 0.16048 | -0.33494 | 0.21233 | 0.13314 | -0.14232 | 0.104532 | 0.071772 | -0.07057 | 0.035345 | -0.01611 | -0.57155 | -0.74817 | 0.06289 | 0.095865 | -0.0534 | 0.0012 | -0.09843 | 0.098 |
| KYM1 | -0.03038 | 0.141208 | -0.01533 | -0.1712 | -0.03118 | -0.04558 | -0.20379 | -0.15459 | 0.06875 | 0.06269 | 0.19689 | 0.18569 | -0.31159 | -0.02832 | 0.127623 | 0.143331 | 0.220669 | -0.07662 | -0.32948 | -0.20786 | -0.74332 | -0.12363 | -0.2296 | -0.02709 | 0.05424 |
| COV644 | 0.021412 | -0.14591 | -0.03264 | 0.07735 | -0.17892 | -1.32535 | -0.05835 | 0.17236 | -0.04264 | 0.17205 | 0.18942 | -0.01888 | 0.058776 | 0.071315 | 0.042191 | -0.07538 | 0.024403 | -0.20528 | -0.32184 | 0.025716 | -0.0309 | -0.01034 | 0.05431 | -0.06247 | 0.03092 |
| SF126 | 0.038347 | 0.082535 | -0.03966 | 0.04018 | -0.04415 | -0.0344 | 0.067686 | 0.13881 | -0.16618 | 0.14903 | 0.07316 | 0.09947 | 0.013889 | 0.063962 | 0.003256 | 0.023309 | 0.091194 | -0.20153 | -0.11587 | -0.15276 | -0.96022 | -0.09975 | 0.07848 | 0.11605 | -0.02284 |
| RVH421 | -0.08783 | -0.02937 | 0.09267 | 0.10764 | -0.09348 | -0.14818 | 0.036819 | -0.03297 | -0.09579 | 0.01682 | 0.08624 | 0.0854 | 0.04518 | 0.134564 | -0.06218 | 0.097314 | 0.282784 | -0.27869 | -0.2281 | -0.08377 | 0.085444 | -0.03326 | -0.08923 | 0.07207 | 0.28214 |
| HS746T | -0.03672 | -0.01177 | -0.10333 | 0.08321 | 0.00421 | -0.28465 | 0.083558 | -0.11304 | -0.08019 | 0.1287 | 0.25787 | 0.26865 | -0.2309 | 0.077413 | 0.054159 | -0.04313 | 0.146246 | -0.16467 | -0.3192 | -0.01949 | -0.16558 | 0.02974 | 0.09427 | -0.16294 | 0.11585 |
| SNU1041 | 0.03398 | -0.0798 | -0.02814 | 0.01209 | -0.09778 | -0.37869 | -0.00129 | -0.19456 | 0.06451 | -0.05638 | 0.0493 | 0.04377 | -0.06145 | 0.049439 | 0.11008 | -0.07102 | 0.01638 | -0.4412 | -0.35058 | -0.13289 | 0.013385 | 0.03122 | -0.06831 | -0.03789 | 0.18463 |
| PECAPJ15 | -0.04726 | 0.049577 | -0.04503 | -0.03778 | 0.02585 | -0.42569 | 0.203262 | 0.1518 | -0.11623 | 0.04931 | 0.12632 | 0.05299 | -0.0838 | 0.04653 | 0.015284 | 0.008645 | 0.146828 | -0.7259 | -0.53185 | -0.09312 | -0.06484 | -0.12345 | 0.03993 | -0.07178 | 0.04619 |
| JHH1 | -0.01272 | 0.099964 | -0.05573 | -0.14242 | -0.03759 | -1.25874 | 0.106626 | 0.05181 | -0.1515 | 0.09303 | 0.18073 | 0.07815 | -0.02986 | 0.044536 | -0.02104 | 0.05018 | 0.088124 | -0.31148 | -0.16189 | -0.00484 | -0.05804 | -0.16217 | -0.13397 | 0.04335 | 0.19243 |
| MDAMB157 | 0.036007 | -0.15418 | -0.0082 | -0.05216 | -0.08435 | -0.05257 | -0.04315 | 0.04178 | -0.04723 | 0.08718 | 0.13874 | -0.00346 | -0.00599 | 0.104386 | -0.21981 | -0.00178 | 0.086613 | -0.24776 | -0.27376 | -0.01622 | -0.60131 | 0.06337 | 0.11781 | -0.14736 | 0.13798 |
| KNS42 | -0.18488 | -0.04482 | 0.00754 | -0.07092 | 0.00662 | 0.01518 | 0.019976 | -0.02521 | 0.08637 | 0.13526 | 0.29743 | 0.06626 | 0.004093 | -0.06116 | -0.22877 | -0.05497 | 0.328199 | -0.32658 | -0.1491 | -0.50289 | -0.01104 | 0.07557 | -0.31045 | -0.01129 | 0.01743 |
| SNU201 | -0.06095 | 0.334135 | 0.03194 | 0.02188 | -0.00888 | -0.12012 | 0.108978 | 0.01682 | 0.03747 | 0.05116 | 0.17198 | 0.01655 | 0.026315 | 0.179226 | -0.15709 | -0.00037 | -0.1109 | -0.17185 | -0.34378 | -0.13052 | -0.13878 | -0.10307 | -0.08239 | -0.01234 | 0.25539 |
| HCC1806 | -0.09494 | 0.044869 | -0.07003 | -0.0338 | -0.03957 | -0.343 | 0.074668 | 0.0476 | 0.05974 | 0.01665 | 0.14085 | 0.13817 | -0.04348 | 0.119082 | 0.01042 | 0.001506 | 0.11189 | -0.24773 | -0.32909 | -0.12834 | -0.02847 | -0.10917 | 0.02358 | 0.0008 | 0.14686 |
| LCLC103H | 0.050022 | -0.25902 | -0.02792 | 0.03526 | 0.12786 | -0.23157 | 0.100672 | -0.06476 | 0.06518 | -0.00476 | 0.11395 | -0.07731 | -0.05108 | 0.172425 | -0.18742 | 0.079498 | 0.038194 | -0.3838 | -0.26135 | -0.09427 | -0.4934 | 0.00537 | 0.00519 | -0.23248 | 0.01613 |
| YD8 | 0.053331 | 0.12558 | 0.09809 | 0.00209 | -0.05211 | -0.3187 | -0.12184 | 0.24975 | -0.08485 | -0.02234 | 0.14367 | -0.03239 | 0.073606 | 0.17972 | -0.05897 | -0.03163 | -0.00973 | -1.52851 | -0.94454 | -0.05614 | -1.20303 | -0.16187 | -0.00075 | -0.11554 | 0.02491 |
| HS944T | 0.055076 | -0.08048 | 0.0521 | -0.01253 | 0.00503 | -0.02877 | -0.09514 | -0.04869 | 0.00139 | -0.00848 | 0.09089 | 0.16068 | 0.006248 | -0.07825 | -0.10212 | -0.08433 | 0.210162 | -0.16246 | -0.24798 | -0.07567 | -0.22679 | -0.01894 | 0.07534 | -0.02872 | 0.16858 |
| FU97 | -0.12397 | 0.14746 | -0.17278 | -0.06597 | 0.07314 | -0.00452 | -0.07694 | 0.06027 | 0.14722 | 0.03933 | 0.10475 | 0.18555 | 0.110506 | 0.057339 | -0.21954 | -0.04128 | 0.0039 | -0.06835 | -0.22365 | 0.016045 | -0.07498 | -0.0886 | 0.11336 | -0.59133 | 0.06957 |
| LN340 | -0.02536 | -0.12652 | 0.02791 | 0.07077 | 0.24228 | -0.12049 | 0.078057 | 0.02421 | 0.12485 | 0.26872 | 0.07155 | -0.02419 | 0.070201 | -0.07303 | -0.11093 | 0.156368 | -0.20882 | -0.25221 | -0.08022 | -0.03416 | -0.54645 | 0.09146 | 0.13313 | 0.07093 | 0.24119 |
| KYSE520 | -0.10295 | 0.04343 | -0.03562 | -0.12946 | -0.13469 | -0.44259 | 0.087551 | 0.02644 | 0.02792 | 0.08734 | 0.17575 | 0.0516 | 0.061717 | -0.02366 | -0.08309 | -0.03333 | 0.023673 | -0.0995 | -0.26671 | -0.00575 | -0.17846 | -0.04129 | 0.04975 | -0.06384 | 0.1693 |
| NCIH441 | 0.117928 | 0.05559 | -0.135 | 0.07727 | 0.01656 | -0.2816 | -0.06461 | -0.18227 | -0.03713 | 0.10033 | 0.13027 | 0.1189 | -0.0536 | 0.030426 | 0.028966 | 0.082076 | 0.178318 | -0.31659 | -0.24886 | 0.068307 | -0.1006 | -0.02952 | 0.26091 | -0.02425 | 0.01464 |
| NCIH211 | -0.00077 | 0.084911 | 0.0518 | -0.01088 | -0.01036 | -0.16723 | 0.039384 | -0.17396 | -0.07097 | 0.07877 | 0.11939 | 0.02554 | 0.093664 | -0.01149 | -0.08364 | 0.00289 | -0.03948 | -0.22694 | -0.13827 | -0.16053 | -0.20789 | -0.19044 | -0.02648 | -0.12002 | 0.0999 |
| JL1 | 0.142803 | 0.006983 | 0.09843 | 0.07346 | 0.03982 | -0.42519 | -0.11594 | -0.10255 | -0.09897 | 0.00157 | 0.15547 | 0.15559 | 0.177908 | 0.103124 | -0.12703 | -0.0486 | 0.272232 | -0.06833 | -0.43836 | -0.07126 | -0.78958 | -0.06467 | -0.02414 | -0.12815 | -0.06336 |
| OVMANA | -0.35663 | 0.215188 | 0.14166 | 0.05029 | -0.04237 | -0.44776 | -0.05722 | -0.06409 | 0.01615 | -0.01369 | 0.18109 | 0.3335 | -0.12922 | 0.166508 | -0.29243 | -0.10533 | 0.096104 | -0.8714 | -0.61594 | -0.14851 | -0.00597 | 0.09728 | 0.19959 | -0.13609 | -0.18706 |
| TE1 | 0.001789 | 0.004491 | -0.08201 | -0.08134 | -0.0642 | -0.26523 | -0.01867 | 0.07721 | -0.1759 | 0.06697 | 0.11505 | -0.01836 | -0.04014 | 0.06139 | -0.11709 | -0.04631 | 0.11634 | -0.32749 | -0.36737 | -0.04838 | -0.06944 | 0.03771 | 0.02006 | 0.00296 | 0.10068 |
| NCIH28 | -0.17305 | 0.02027 | 0.18352 | 0.0586 | 0.01347 | -0.52173 | 0.082673 | 0.28628 | 0.04502 | -0.14266 | 0.13667 | 0.08882 | 0.182911 | 0.284569 | 0.087401 | 0.083679 | 0.187677 | -0.43953 | -0.1499 | -0.20821 | 0.261837 | 0.08988 | 0.06926 | -0.01945 | 0.0725 |
| 786O | -0.03761 | -0.11389 | 0.02289 | -0.00887 | -0.08728 | -0.2574 | -0.16606 | 0.49604 | 0.01925 | 0.0471 | 0.09887 | 0.1218 | 0.156503 | 0.079987 | -0.20477 | 0.101854 | 0.0706 | -0.26224 | -0.26218 | -0.02005 | 0.006905 | 0.04353 | 0.05305 | -0.1844 | 0.11304 |
| SW620 | -0.07287 | 0.040223 | -0.07458 | -0.06706 | -0.1362 | -0.00486 | -0.09964 | -0.11733 | 0.03774 | 0.11819 | 0.15311 | 0.00235 | -0.01214 | 0.013293 | -0.08611 | 0.053464 | 0.083559 | -0.33694 | -0.18927 | 0.033922 | 0.037758 | -0.06669 | -0.01304 | 0.02605 | 0.30655 |
| SUIT2 | 0.007397 | -0.00619 | -0.00282 | -0.03679 | -0.11004 | -0.4781 | 0.010947 | 0.37018 | 0.20807 | 0.14798 | 0.16857 | 0.10752 | -0.08042 | 0.024714 | -0.07734 | -0.02404 | 0.071821 | -0.36919 | -0.18947 | -0.05941 | -0.02717 | -0.23382 | -0.00043 | -0.07027 | 0.07884 |
| JJN3 | 0.185797 | -0.11057 | 0.01064 | 0.02871 | -0.1051 | -0.20547 | 0.045507 | -0.16143 | 0.03194 | -0.05579 | 0.15758 | -0.05938 | 0.160448 | 0.062157 | 0.097242 | -0.00516 | 0.082189 | -0.06162 | -0.14334 | -0.13155 | -0.09314 | -0.01018 | 0.04906 | -0.19164 | 0.09553 |
| RAJI | 0.266009 | 0.238529 | -0.02062 | 0.2252 | -0.02337 | -0.16561 | 0.062008 | 0.0002 | -0.04674 | 0.02469 | 0.14193 | 0.12714 | 0.156133 | 0.123599 | -0.18852 | -0.23085 | 0.195611 | 0.06132 | -0.26748 | 0.003375 | -0.06367 | 0.04367 | 0.08246 | -0.08864 | 0.08459 |
| SF268 | 0.012064 | 0.031616 | 0.0115 | -0.08164 | -0.05486 | -0.18946 | -0.03348 | -0.09012 | -0.00437 | 0.07983 | 0.10898 | 0.02955 | 0.032684 | 0.063205 | 0.112599 | -0.0223 | -0.00338 | -0.30995 | -0.29499 | -0.0948 | -0.21781 | -0.06862 | -0.09232 | 0.03305 | 0.11887 |
| SUDHL8 | 0.027317 | 0.061933 | 0.02139 | 0.01225 | -0.00117 | -0.00753 | 0.11385 | -0.07882 | 0.01478 | 0.02614 | 0.3344 | 0.08287 | 0.215731 | -0.18558 | -0.11514 | 0.139157 | -0.01345 | -0.13291 | 0.03934 | -0.11016 | -0.07814 | 0.05093 | -0.20586 | -0.01643 | 0.07162 |
| A2780 | 0.003853 | -0.13013 | -0.15977 | -0.09652 | -0.14449 | -0.08787 | -0.03877 | -0.01018 | -0.02891 | 0.14906 | 0.19458 | -0.04886 | 0.061598 | 0.073153 | -0.0872 | 0.009694 | 0.036968 | -0.2071 | -0.24016 | 0.108435 | -0.70302 | -0.0273 | 0.04562 | -0.05189 | 0.22507 |
| KMS18 | -0.00358 | -0.00044 | -0.0526 | 0.1442 | -0.20397 | -0.13657 | -0.0916 | -0.15079 | -0.06842 | -0.00151 | 0.10052 | 0.13542 | 0.01272 | 0.0157 | -0.13353 | 0.004696 | 0.002206 | -0.31775 | -0.29999 | -0.11049 | -0.06152 | -0.19549 | -1.79234 | -0.07278 | 0.13244 |
| SUDHL5 | 0.098876 | 0.223858 | 0.08789 | 0.23808 | -0.01569 | 0.07354 | 0.051029 | -0.06804 | 0.29375 | 0.18516 | 0.16739 | 0.14594 | 0.024727 | 0.134559 | 0.025043 | 0.013952 | 0.246205 | -0.17307 | 0.05036 | 0.024479 | -0.22001 | 0.03484 | -0.05009 | 0.13573 | 0.20049 |
| WM1799 | 0.015521 | -0.0697 | 0.10304 | 0.05381 | -0.13237 | -0.35029 | 0.033129 | -0.00818 | -0.10866 | 0.07934 | 0.03213 | 0.23289 | -0.01822 | -0.00516 | -0.08019 | 0.149055 | 0.166918 | -0.32494 | -0.2895 | -0.30675 | 0.11957 | -0.06653 | 0.15394 | 0.06991 | -0.20475 |
| CORL23 | 0.062353 | 0.332422 | -0.33611 | -0.11503 | 0.09937 | -0.15945 | -0.23966 | -0.16904 | -0.18946 | 0.07628 | 0.0822 | -0.1302 | 0.070932 | -0.1499 | -0.02238 | 0.129684 | -0.02195 | -0.10711 | -0.19928 | -0.05442 | 0.184273 | 0.19287 | -0.0195 | -0.03014 | 0.16292 |
| OVTOKO | -0.21169 | 0.049975 | 0.16309 | -0.17236 | -0.14125 | -0.64818 | -0.39068 | 0.33724 | -0.09282 | 0.29818 | 0.24433 | -0.12759 | -0.01487 | 0.248823 | -0.23034 | 0.107818 | 0.059991 | -0.8448 | -0.35836 | 0.088534 | 0.164865 | 0.0692 | -0.0466 | -0.04133 | 0.17342 |
| SUDHL1 | 0.095944 | -0.05968 | 0.02428 | 0.08172 | 0.00093 | -0.17256 | -0.0742 | -0.11446 | -0.04944 | 0.05361 | 0.09469 | 0.07458 | -0.03704 | 0.023756 | -0.17491 | 0.03724 | 0.035474 | -0.24572 | -0.19393 | -0.08294 | -0.01903 | 0.0441 | -0.0757 | -0.00524 | 0.16119 |
| SKMES1 | 0.026651 | 0.039899 | -0.04269 | -0.02199 | -0.04056 | -0.94071 | 0.062044 | -0.02151 | 0.01915 | -0.01479 | 0.12815 | 0.02996 | -0.06108 | 0.029571 | -0.14479 | -0.13896 | 0.054668 | -0.39723 | -0.48141 | -0.10864 | -0.0444 | 0.04036 | 0.00131 | 0.08244 | 0.04012 |
| NCIH1355 | 0.222254 | -0.01776 | 0.09229 | 0.07011 | -0.00966 | 0.1663 | -0.1471 | -0.03632 | 0.175 | -0.08075 | 0.06683 | 0.28688 | 0.090753 | 0.016345 | -0.02647 | 0.049427 | 0.086149 | -0.16197 | -0.21948 | -0.1269 | -0.0814 | -0.07836 | 0.03924 | 0.0821 | 0.19172 |
| HCC44 | 0.175698 | -0.15831 | -0.04772 | -0.14808 | -0.05167 | -0.17337 | -0.07325 | 0.20116 | 0.09502 | 0.05355 | 0.08797 | 0.20628 | 0.039996 | 0.119491 | -0.45985 | 0.158432 | 0.080688 | -0.27107 | -0.12131 | -0.13769 | -0.16089 | -0.06148 | 0.12681 | 0.04487 | 0.16094 |
| HCC70 | 0.068486 | -0.00698 | 0.05384 | 0.0731 | 0.0103 | 0.06869 | 0.06097 | -0.04932 | 0.01378 | 0.00797 | 0.56893 | 0.11327 | 0.104682 | -0.09355 | -0.09309 | 0.119093 | 0.162458 | -0.26409 | -0.21497 | 0.042695 | 0.102041 | -0.06147 | -0.13342 | -0.09023 | 0.08605 |
| HUH6 | 0.079667 | -0.0121 | -0.04804 | 0.0554 | 0.04548 | -0.12894 | 0.070411 | 0.11238 | 0.04816 | 0.05272 | 0.02805 | -0.02679 | 0.007097 | 0.140678 | -0.07762 | 0.031815 | 0.052923 | -0.24383 | -0.20525 | -0.15898 | -0.36422 | -0.11659 | -0.11299 | -0.34481 | 0.2615 |
| LN443 | 0.060969 | 0.162148 | -0.00627 | -0.11277 | 0.04687 | -0.01721 | 0.009231 | -0.03174 | 0.01274 | 0.08005 | 0.14813 | 0.09384 | -0.02216 | 0.092133 | 0.00199 | -0.02774 | 0.133212 | -0.3069 | -0.25079 | -0.03813 | -0.93763 | -0.06543 | 0.02409 | 0.0177 | 0.13323 |
| LN464 | -0.00259 | 0.138362 | -0.03625 | -0.11486 | -0.09615 | -0.09808 | -0.03679 | -0.11211 | -0.10294 | 0.06131 | 0.05884 | -0.08211 | 0.101574 | 0.078068 | -0.15311 | 0.02414 | -0.03273 | -0.25703 | -0.21403 | -0.06006 | -0.09578 | 0.02321 | 0.20832 | 0.06713 | 0.16198 |
| SW1573 | 0.01798 | -0.196 | -0.22588 | 0.07315 | -0.00234 | -0.07009 | -0.08439 | 0.06757 | -0.06955 | 0.18112 | 0.11685 | 0.05762 | -0.00678 | 0.122411 | -0.09337 | 0.00547 | 0.135023 | -0.23206 | -0.47011 | -0.11226 | -0.04261 | 0.04489 | 0.05253 | 0.10267 | 0.09762 |
| SW948 | -0.11223 | 0.141765 | 0.26077 | -0.39313 | -0.09272 | -0.25198 | 0.103034 | -0.21274 | -0.03905 | -0.03905 | 0.20791 | 0.08769 | -0.05009 | 0.204544 | -0.02967 | -0.26099 | 0.250482 | -0.30576 | -0.25758 | 0.066646 | -0.27502 | -0.08072 | -0.09314 | 0.147 | 0.16908 |
| A549 | -0.09588 | -0.03577 | -0.05706 | -0.0488 | -0.04112 | -0.28354 | -0.03954 | 0.18654 | -0.09689 | 0.09434 | 0.0854 | 0.17297 | 0.02804 | -0.15499 | 0.157091 | 0.041775 | 0.170615 | -0.9046 | -0.54979 | -0.00372 | 0.034664 | -0.1424 | 0.1112 | 0.14067 | 0.19153 |
| SNU1066 | 0.215714 | 0.102127 | -0.10305 | -0.06519 | 0.03491 | -0.72447 | 0.033538 | 0.2487 | 0.00969 | 0.08366 | 0.15975 | 0.13512 | 0.003492 | 0.081025 | -0.08612 | 0.034883 | 0.083309 | -0.84575 | -0.73137 | 0.020854 | -0.0225 | 0.0599 | 0.04244 | -0.05624 | 0.04271 |
| SNU503 | 0.023882 | 0.014236 | -0.08899 | -0.05529 | -0.3978 | -0.60899 | 0.03126 | -0.01933 | 0.00527 | 0.1796 | 0.16046 | -0.07007 | 0.066759 | 0.129358 | -0.1841 | 0.047168 | 0.027056 | -0.78081 | -0.37876 | 0.053074 | -0.16182 | 0.06892 | 0.06028 | -0.07463 | 0.2475 |
| KMRC1 | -0.14385 | 0.012014 | -0.15014 | -0.08079 | -0.40811 | -0.3684 | 0.096777 | 1.36624 | 0.01573 | 0.02298 | 0.09103 | -0.07299 | 0.098189 | -0.01446 | -0.03969 | 0.230548 | 0.166987 | -0.29383 | -0.33831 | 0.437965 | -0.35528 | -0.05301 | 0.18255 | -0.06769 | 0.15622 |
| L33 | -0.26404 | 0.211777 | 0.01892 | 0.11418 | -0.15463 | -0.18174 | 0.158695 | -0.06473 | -0.09304 | 0.18422 | 0.00422 | 0.13956 | -0.17288 | 0.067044 | 0.023593 | -0.11195 | 0.205001 | -1.22142 | -0.70484 | -0.1292 | 0.057357 | -0.14089 | 0.13524 | 0.0619 | 0.14466 |
| OV7 | -0.17264 | -0.15246 | 0.12682 | -0.08146 | -0.11948 | -0.0604 | 0.028674 | 0.0953 | -0.10779 | 0.19948 | 0.09878 | -0.10228 | 0.032458 | 0.098999 | 0.001502 | 0.050164 | -0.01266 | -0.30052 | -0.28501 | -0.10135 | -0.57487 | 0.04807 | 0.23274 | -0.13184 | 0.01421 |
| KYSE180 | 0.066079 | 0.010785 | -0.01937 | -0.01661 | 0.00577 | -0.30799 | 0.006012 | -0.08442 | 0.01735 | 0.11317 | 0.16379 | 0.16903 | 0.045901 | 0.207399 | -0.02563 | -0.05444 | 0.067327 | -0.68809 | -0.88995 | 0.023388 | 0.095442 | -0.03354 | 0.00462 | 0.00591 | -0.03088 |
| TE9 | -0.03651 | 0.061185 | 0.00564 | -0.01374 | -0.06029 | -0.19548 | 0.0614 | 0.00535 | 0.22619 | 0.14402 | 0.13723 | 0.05053 | -0.02459 | 0.113758 | -0.06766 | -0.0108 | 0.085865 | -0.78486 | -0.49033 | -0.14255 | -0.14288 | 0.00059 | 0.00302 | -0.05597 | 0.23204 |
| CORL47 | -0.008 | 0.200896 | 0.09028 | 0.25426 | -0.09015 | -0.18155 | -0.08577 | -0.4459 | 0.05692 | 0.24618 | -0.03346 | -0.04876 | 0.250459 | 0.075371 | 0.082882 | 0.011463 | 0.17065 | -0.25558 | -0.07315 | -0.28675 | -0.13574 | -0.04214 | -0.16219 | 0.00943 | -0.05384 |

| Cell line | ALK | AXL | CSF1R | DDR1 | DDR2 | EGFR | EPHA1 | EPHA2 | EPHA3 | EPHA4 | EPHA6 | EPHA7 | EPHB1 | EPHB2 | EPHB3 | EPHB4 | EPHB6 | ERBB2 | ERBB3 | ERBB4 | FGFR1 | FGFR2 | FGFR3 | FGFR4 | FLT1 |
|---|---|---|---|---|---|---|---|---|---|---|---|---|---|---|---|---|---|---|---|---|---|---|---|---|---|
| OVCAR8 | 0.020773 | -0.08476 | 0.07355 | -0.02107 | -0.11863 | -0.20588 | -0.02127 | -0.02379 | -0.05519 | 0.0983 | 0.20569 | -0.00403 | 0.089788 | 0.130688 | -0.02974 | 0.017208 | 0.032731 | -0.39791 | -0.34366 | -0.01844 | -0.00407 | -0.05989 | 0.08312 | -0.06094 | 0.10276 |
| A3KAW | 0.02347 | 0.172144 | 0.0305 | 0.101 | -0.13119 | 0.05773 | 0.063403 | -0.09103 | -0.05656 | 0.07095 | 0.13017 | 0.09726 | 0.126379 | 0.061751 | -0.12406 | -0.0461 | 0.172948 | -0.21068 | -0.181 | -0.16922 | 0.126532 | -0.02966 | 0.03371 | 0.18411 | -0.01878 |
| DMS53 | 0.090298 | -0.08314 | -0.12742 | -0.03252 | -0.1732 | -0.1688 | -0.07377 | -0.07507 | -0.06769 | 0.02275 | 0.05996 | -0.04452 | 0.070625 | 0.185469 | -0.11015 | 0.056395 | 0.036071 | -0.1269 | -0.31146 | 0.095517 | -0.48342 | -0.15576 | 0.04779 | -0.0633 | 0.16754 |
| HCC1395 | -0.16123 | 0.059905 | 0.02232 | 0.04394 | -0.16759 | -0.05682 | 0.070326 | 0.02273 | 0.09457 | 0.03263 | 0.28526 | 0.09875 | -0.00196 | -0.0043 | 0.022067 | 0.125704 | 0.135637 | -0.2195 | -0.24093 | 0.084761 | -0.31933 | 0.12069 | -0.00785 | 0.04699 | 0.12192 |
| NCIH2882 | 0.114089 | 0.031294 | 0.08183 | 0.02162 | 0.10674 | -0.00987 | -0.35603 | 0.15622 | 0.12096 | 0.06893 | -0.06968 | 0.28978 | 0.252669 | -0.03302 | -0.10501 | -0.06791 | 0.235999 | -0.34479 | -0.12765 | -0.30723 | -0.28342 | 0.115 | -0.03449 | -0.01275 | -0.14767 |
| RMUGS | 0.012677 | -0.26523 | 0.09053 | -0.03933 | -0.28919 | -0.34555 | -0.12849 | 0.25636 | -0.14841 | 0.37633 | 0.19434 | -0.07242 | 0.097742 | 0.148856 | 0.05125 | 0.005556 | 0.206195 | -0.54993 | -0.16739 | -0.04163 | 0.02319 | 0.11182 | 0.16769 | -0.18055 | 0.29236 |
| L1236 | 0.019927 | 0.10089 | 0.14797 | 0.00264 | -0.05379 | -0.07844 | 0.016682 | -0.12286 | -0.04888 | 0.07833 | 0.31471 | 0.06814 | 0.23601 | -0.11049 | -0.31705 | -0.01134 | -0.11032 | -0.36972 | -0.12847 | -0.00887 | -0.08786 | -0.03777 | 0.03841 | 0.15556 | 0.30414 |
| OAW42 | -0.15821 | -0.10775 | -0.19196 | -0.09559 | 0.03986 | -0.01429 | 0.036153 | -0.27932 | 0.11026 | 0.07348 | 0.36434 | 0.1949 | 0.025483 | -0.15396 | -0.51186 | 0.001594 | -0.33552 | -0.54426 | -0.08511 | -0.50696 | 0.032324 | -0.13735 | -0.06361 | -0.2891 | 0.1238 |
| EKVX | 0.130309 | 0.055162 | 0.03856 | -0.02967 | -0.09514 | -0.80833 | -0.01897 | 0.05551 | -0.1208 | 0.24229 | -0.01601 | 0.05199 | 0.083462 | 0.007128 | 0.00811 | -0.00858 | 0.214669 | -0.53887 | -0.31025 | -0.10994 | 0.058672 | -0.01194 | -0.09027 | 0.10081 | 0.23197 |
| KMRC2 | -0.10238 | -0.263 | -0.04864 | 0.07437 | 0.08209 | -0.54761 | -0.04887 | 0.08816 | -0.25694 | 0.19142 | 0.27922 | 0.05863 | -0.09116 | 0.022064 | 0.033582 | 0.166209 | 0.185383 | -0.27831 | -0.33893 | 0.009722 | 0.121368 | 0.05558 | -0.09002 | 0.15539 | 0.24718 |
| JIMT1 | 0.075153 | 0.086137 | 0.04785 | -0.04086 | 0.0234 | -0.5045 | 0.045893 | -0.09682 | 0.09304 | 0.13805 | 0.09381 | 0.10228 | 0.029074 | 0.121315 | -0.07166 | 0.050934 | 0.057341 | -0.29905 | -0.28296 | -0.10825 | -0.00609 | -0.05811 | -0.02372 | 0.01621 | 0.13242 |
| CAOV3 | 0.037144 | -0.16662 | -0.11702 | 0.03296 | -0.11518 | -0.20522 | -0.08536 | 0.15856 | -0.05184 | 0.04358 | 0.18852 | 0.01076 | 0.184451 | 0.117728 | -0.0576 | 0.092037 | 0.065562 | -0.18116 | -0.3486 | 0.221701 | -0.05995 | -0.09297 | -0.21647 | -0.058 | 0.20031 |
| KMS11 | -0.0238 | -0.02728 | 0.03337 | 0.02836 | -0.05854 | 0.13288 | -0.01868 | -0.1685 | -0.03258 | 0.05483 | 0.19478 | 0.08179 | 0.083465 | -0.05712 | -0.10001 | 0.055829 | -0.01734 | -0.12985 | -0.45393 | -0.13024 | 0.079885 | -0.08115 | -1.90641 | 0.10828 | 0.0675 |
| TT2609C02 | 0.120573 | -0.13667 | -0.04002 | 0.09185 | -0.10479 | -0.17311 | -0.14289 | -0.02914 | 0.01112 | 0.01149 | 0.07159 | 0.03773 | -7.7E-06 | 0.05318 | -0.22759 | 0.08282 | -0.08238 | -0.30956 | -0.28163 | -0.07022 | -0.16121 | -0.06177 | 0.18321 | 0.11097 | 0.03503 |
| COLO680N | -0.09304 | 0.075335 | -0.02757 | -0.07822 | -0.23821 | -0.71963 | 0.038907 | 0.18513 | 0.06679 | 0.12912 | 0.08597 | 0.03034 | 0.079136 | 0.098186 | -0.02634 | -0.10478 | 0.087155 | -0.24979 | -0.13903 | 0.077333 | -0.05208 | -0.07003 | 0.01942 | 0.09448 | -0.1325 |
| NCIH2291 | 0.057741 | -0.38609 | -0.05164 | 0.05452 | -0.07796 | -0.22851 | 0.175309 | -0.13137 | 0.05324 | -0.03077 | 0.16452 | 0.16781 | 0.212967 | 0.203912 | 0.046064 | 0.168785 | 0.290507 | -0.17021 | -0.3055 | -0.01092 | 0.048328 | -0.01194 | 0.08351 | -0.05456 | 0.10684 |
| RMGI | -0.03703 | 0.057038 | 0.07355 | 0.07362 | -0.00105 | -0.81363 | -0.02005 | 0.23489 | -0.06481 | 0.04136 | 0.17678 | 0.0279 | 0.029793 | 0.041349 | -0.08428 | 0.089557 | -0.00594 | -0.59501 | -0.30931 | -0.12815 | 0.050662 | 0.08796 | -0.24692 | 0.00421 | 0.3069 |
| TCCSUP | 0.099245 | 0.023381 | 0.01174 | 0.039 | -0.00678 | -0.17039 | 0.021559 | -0.04972 | 0.04075 | 0.12379 | 0.03167 | 0.1595 | 0.042967 | 0.056412 | 0.074208 | -0.04942 | 0.109283 | -0.15256 | -0.20887 | -0.05232 | -1.2008 | -0.07754 | -0.06243 | 0.09736 | 0.1146 |
| HMC18 | 0.055707 | 0.102372 | 0.01511 | 0.0426 | -0.08061 | -0.16375 | -0.07731 | -0.08574 | 0.01332 | 0.07255 | 0.11294 | 0.04947 | -0.00998 | -0.05922 | 0.00113 | 0.047064 | 0.069807 | -0.2448 | -0.1344 | -0.00713 | 0.00386 | -0.01736 | 0.03116 | -0.04816 | 0.12253 |
| SNUC1 | 0.274046 | 0.041059 | 0.26383 | 0.33947 | 0.06602 | -0.09924 | -0.18731 | 0.1026 | 0.05337 | -0.03839 | 0.54929 | 0.51403 | 0.084076 | 0.064595 | 0.071092 | -0.12156 | 0.127663 | -0.43103 | -0.06454 | 0.127853 | -0.27061 | 0.01653 | 0.16134 | -0.01877 | 0.19859 |
| HT1376 | -0.16208 | -0.00268 | 0.0666 | 0.18794 | 0.07056 | -1.07251 | 0.108352 | 0.0103 | 0.18051 | 0.21633 | 0.06278 | 0.13864 | 0.035897 | 0.167143 | -0.07055 | 0.065538 | 0.189909 | -0.38497 | -0.27725 | -0.24758 | 0.01789 | -0.10721 | 0.00094 | 0.08968 | -0.24361 |
| HCC202 | 0.055338 | 0.071491 | 0.02898 | -0.08258 | -0.11887 | -0.19404 | 0.256055 | 0.1444 | 0.06435 | 0.12856 | 0.19208 | 0.07746 | -0.13492 | 0.036537 | -0.05575 | 0.075087 | 0.18017 | -1.77742 | -0.16428 | 0.086043 | -0.03878 | -0.04013 | -0.08574 | 0.01109 | 0.08909 |
| PECAPJ41CLONED2 | 0.192325 | -0.09601 | -0.07543 | 0.00317 | -0.17998 | -0.7266 | -0.09023 | -0.20811 | -0.08806 | -0.04324 | 0.05846 | 0.15468 | -0.06676 | 0.160752 | -0.17579 | -0.06734 | 0.183651 | -0.26177 | -0.15413 | -0.03347 | -0.04037 | 0.01444 | 0.03872 | -0.05078 | -0.05139 |
| JHH5 | -0.07992 | -0.01919 | -0.02909 | -0.0741 | -0.10548 | -0.64082 | -0.1843 | 0.14584 | -0.02455 | 0.06095 | 0.08031 | -0.01636 | 0.078595 | 0.033085 | -0.03241 | -0.09868 | 0.07101 | -0.45612 | -0.33931 | 0.074411 | 0.039992 | -0.05964 | 0.0714 | -0.03876 | 0.12098 |
| PECAPJ49 | 0.121892 | -0.15826 | -0.05582 | -0.11734 | 0.02551 | -1.76995 | 0.045208 | 0.10469 | 0.01719 | 0.10569 | 0.22987 | 0.08775 | -0.05607 | 0.157362 | 0.166887 | 0.104933 | 0.121685 | -0.58423 | -0.28775 | -0.07922 | -0.00013 | 0.02885 | -0.02325 | 0.01907 | 0.28282 |
| SNU601 | 0.071246 | 0.127467 | -0.13841 | 0.08386 | -0.01151 | -0.29926 | 0.00822 | -0.08807 | -0.04456 | 0.09723 | 0.07394 | 0.1189 | 0.084643 | 0.182571 | -0.1078 | 0.018651 | 0.136882 | -0.09566 | -0.1797 | -0.12139 | -0.00115 | 0.06527 | -0.12438 | 0.05192 | 0.06136 |
| GB1 | 0.033411 | 0.024873 | 0.00367 | -0.05914 | -0.06535 | -0.1088 | -0.09799 | 0.1001 | 0.10723 | 0.24749 | 0.07137 | 0.13881 | -0.09727 | 0.040193 | -0.11754 | -0.09228 | 0.144404 | -0.10853 | -0.20458 | 0.045573 | -0.33863 | 0.01645 | -0.0278 | 0.00625 | 0.19799 |
| HEPG2 | -0.06118 | -0.0328 | -0.21778 | -0.11434 | -0.03292 | 0.00913 | -0.00565 | 0.12225 | 0.08752 | 0.1162 | 0.10899 | 0.16316 | 0.076155 | -0.0399 | -0.14538 | 0.178283 | 0.167785 | -0.30613 | -0.16311 | 0.030628 | -0.17259 | -0.11869 | 0.15317 | 0.18298 | 0.0891 |
| A253 | -0.05793 | 0.077362 | -0.025 | -0.11459 | 0.00398 | -0.65763 | 0.107176 | 0.171 | -0.08038 | 0.16053 | 0.08934 | -0.09534 | 0.090816 | 0.187322 | -0.11591 | 0.024477 | 0.114814 | -0.51659 | -0.63063 | -0.06554 | -0.00906 | -0.07353 | 0.1921 | -0.17805 | 0.04451 |
| UBLC1 | -0.20876 | 0.045004 | -0.02646 | -0.18868 | -0.20748 | -1.05038 | 0.142114 | 0.24446 | -0.11068 | 0.01716 | 0.2115 | -0.01562 | -0.05743 | 0.199929 | -0.16864 | 0.134854 | 0.103197 | -0.30965 | -0.34415 | -0.06751 | -0.0444 | 0.04335 | -0.03681 | 0.0358 | 0.27161 |
| GSS | -0.02785 | -0.06933 | 0.10715 | 0.13846 | -0.11354 | -0.18366 | -0.33002 | 0.00204 | 0.09458 | 0.07218 | 0.27185 | 0.00167 | 0.20031 | 0.014926 | -0.04595 | -0.08081 | 0.128664 | -0.36342 | -0.20291 | -0.16665 | 0.170829 | -0.42481 | -0.0131 | -0.25498 | 0.05742 |
| NCIH1703 | -0.01062 | 0.041981 | -0.03837 | 0.07271 | -0.11331 | -0.02936 | 0.095135 | 0.06035 | -0.0363 | -0.01127 | 0.08811 | 0.02927 | 0.084487 | 0.017745 | -0.08656 | 0.099503 | 0.175698 | -0.23838 | -0.22198 | -0.25635 | -0.10235 | -0.08794 | 0.13734 | -0.05486 | 0.1406 |
| SJSA1 | 0.000582 | -0.11021 | -0.08373 | 0.08229 | -0.0984 | -0.09501 | -0.08841 | -0.02947 | -0.00966 | -0.03452 | 0.06777 | 0.03671 | 0.024449 | -0.02764 | -0.10285 | -0.025 | 0.035981 | -0.14847 | -0.04942 | -0.12143 | -0.0471 | -0.1533 | 0.01115 | -0.05295 | 0.06264 |
| JMSU1 | 0.058448 | -0.03498 | 0.00766 | -0.16841 | -0.06274 | -0.1558 | 0.070812 | 0.03962 | 0.07012 | -0.03142 | 0.10088 | 0.07307 | 0.044962 | 0.069632 | -0.23403 | -0.01589 | 0.218956 | -0.25334 | -0.17691 | -0.24891 | -1.68774 | -0.0722 | 0.05476 | -0.02459 | -0.04487 |
| L428 | 0.116048 | 0.188652 | -0.03453 | 0.14834 | -0.04103 | -0.2473 | -0.18266 | -0.15971 | -0.04911 | 0.05579 | 0.05339 | 0.06965 | -0.00385 | 0.031599 | -0.08066 | 0.049022 | 0.150549 | -0.37951 | -0.17622 | 0.002564 | 0.102087 | -0.08214 | 0.20305 | -0.24903 | 0.44812 |
| GI1 | 0.17172 | -0.06213 | -0.02016 | -0.12912 | 0.11163 | -0.06467 | -0.04639 | -0.04553 | 0.05506 | -0.02747 | 0.16985 | 0.17685 | 0.023842 | 0.046629 | -0.16073 | 0.004705 | 0.027858 | -0.19796 | -0.19446 | -0.04619 | -0.14116 | 0.1174 | -0.07146 | -0.14184 | 0.29806 |
| A427 | -0.04048 | 0.078207 | -0.09737 | 0.07244 | -0.14671 | -0.08035 | -0.17107 | -0.09796 | -0.1164 | 0.34794 | -0.0068 | 0.04339 | -0.00598 | -0.0678 | 0.007318 | 0.119339 | 0.070904 | -0.04449 | -0.19696 | -0.18625 | -0.60291 | -0.04829 | 0.06281 | -0.01423 | 0.0843 |
| MKN74 | 0.098349 | -0.05842 | -0.02949 | -0.0838 | 0.00244 | -0.48734 | -0.10667 | -0.05025 | -0.05639 | 0.03838 | -0.01226 | 0.14763 | 0.002009 | 0.208651 | 0.044919 | 0.126708 | 0.096879 | -0.13323 | -0.10497 | -0.02317 | -0.00878 | 0.07267 | 0.21851 | 0.02092 | -0.05632 |
| LNZ308 | -0.07587 | 0.048625 | 0.04943 | 0.0075 | -0.04807 | -0.05985 | 0.056472 | -0.02712 | 0.06924 | 0.01774 | 0.19545 | 0.06376 | 0.02154 | 0.007007 | -0.16173 | 0.00589 | -0.03471 | -0.18832 | -0.27832 | -0.12121 | -0.42507 | -0.02346 | 0.0648 | -0.01722 | 0.08413 |
| YD38 | 0.15943 | 0.184495 | -0.04276 | -0.12746 | -0.06584 | -0.68669 | 0.057749 | 0.17593 | 0.12811 | 0.0459 | 0.18276 | -0.02471 | -0.17874 | 0.087178 | -0.03542 | 0.082665 | 0.047563 | -0.44562 | -0.4703 | 0.038911 | -0.1721 | -0.03833 | -0.04317 | 0.11062 | 0.39532 |
| MM1S | -0.20057 | 0.150304 | 0.15759 | 0.03275 | -0.12941 | -0.08883 | -0.05109 | -0.17448 | -0.08958 | -0.05786 | 0.20818 | 0.10158 | -0.03874 | 0.019348 | -0.01276 | 0.076876 | 0.021988 | -0.37798 | -0.23565 | -0.12072 | -0.01005 | -0.15014 | 0.00681 | 0.14612 | 0.14829 |
| SH10TC | -0.12908 | 0.13707 | 0.22719 | -0.05566 | -0.10631 | -0.1423 | -0.04245 | 0.13357 | 0.09349 | 0.22038 | 0.00517 | 0.01672 | 0.014169 | -0.10484 | 0.060332 | -0.10824 | 0.041832 | -0.3278 | -0.25258 | -0.27333 | -0.02336 | 0.04414 | -0.03799 | 0.15989 | -0.02686 |
| WM983B | -0.07666 | 0.202223 | -0.00051 | -0.10733 | -0.01907 | -0.07606 | 0.06185 | -0.08142 | -0.01927 | 0.07178 | 0.03746 | 0.06353 | -0.01677 | 0.026791 | -0.15107 | 0.039654 | 0.065275 | -0.35621 | -0.26969 | -0.17971 | 0.04346 | 0.05055 | 0.02638 | -0.09179 | 0.3201 |
| NCIH1648 | 0.103229 | -0.03812 | 0.01122 | 0.09494 | -0.03652 | -0.79744 | 0.063124 | -0.1428 | -0.17509 | 0.10352 | 0.02848 | 0.07707 | 0.175055 | 0.131441 | -0.15512 | 0.002429 | -0.04522 | -0.16083 | -0.14407 | 0.192479 | -0.59742 | -0.07683 | 0.10015 | -0.10108 | 0.43113 |
| NCIH526 | 0.050302 | -0.01168 | -0.1544 | -0.08641 | -0.29174 | -0.05998 | -0.01821 | -0.09751 | 0.00125 | 0.02186 | 0.04276 | 0.22771 | 0.002618 | -0.06178 | -0.35166 | -0.02154 | 0.090839 | -0.15711 | -0.26291 | 0.006942 | -0.03188 | -0.09499 | 0.06359 | 0.02572 | 0.08126 |
| MDAMB231 | -0.02257 | -0.10125 | -0.04636 | -0.15401 | -0.12138 | -0.06508 | 0.063317 | 0.0619 | -0.20516 | 0.136 | 0.24969 | -0.02533 | -0.00544 | 0.137328 | -0.06212 | 0.108354 | 0.172109 | -0.25069 | -0.2774 | 0.044137 | -0.21074 | -0.18465 | 0.05302 | -0.04791 | 0.07126 |
| LK2 | -0.04059 | -0.05638 | 0.00028 | -0.02383 | -0.08872 | -0.04487 | 0.128417 | -0.08395 | -0.07146 | 0.08028 | 0.23595 | 0.04496 | 0.10569 | 0.021093 | -0.1219 | -0.06828 | 0.081327 | -0.291 | -0.1239 | -0.13166 | -1.20803 | 0.00989 | -0.02371 | -0.07205 | 0.18053 |
| P31FUJ | 0.058904 | -0.00288 | -0.11226 | 0.06896 | -0.02864 | -0.10495 | -0.09355 | -0.23658 | -0.03413 | 0.04829 | 0.0484 | 0.15835 | 0.013239 | -0.05346 | -0.12458 | 0.062692 | 0.086439 | -0.28729 | -0.0754 | -0.0657 | -0.00878 | -0.05148 | -0.04319 | 0.02537 | 0.22985 |
| BICR56 | 0.154063 | -0.01844 | -0.0778 | 0.07249 | 0.09939 | -1.01676 | 0.071485 | 0.5855 | 0.00192 | 0.18471 | -0.01415 | 0.2006 | 0.083546 | 0.173379 | 0.095089 | 0.057731 | 0.066039 | -0.71854 | -0.52192 | -0.1555 | -0.09077 | -0.13485 | 0.05893 | 0.03781 | 0.12212 |
| KIJK | -0.03461 | 0.152934 | 0.0818 | -0.00262 | 0.02565 | -0.16915 | 0.018935 | -0.13353 | -0.18627 | 0.00725 | 0.10612 | 0.22896 | 0.026213 | 0.003178 | -0.0764 | -0.03549 | 0.140719 | -0.22663 | -0.0803 | -0.06128 | -0.08487 | 0.17547 | 0.25393 | -0.15968 | 0.09925 |
| RERFLCAD2 | -0.04729 | -0.00933 | -0.21598 | -0.00625 | -0.28998 | 0.02867 | 0.171082 | -0.11742 | -0.05728 | 0.10544 | 0.01129 | 0.10526 | -0.0728 | 0.192581 | -0.13676 | 0.047001 | 0.037265 | -0.30189 | -0.15383 | -0.04928 | -0.1683 | 0.03624 | 0.00116 | 0.19281 | 0.25493 |
| NCIH727 | -0.04496 | 0.091775 | 0.09067 | -0.01924 | -0.02495 | -0.31898 | -0.04206 | -0.05045 | -0.0478 | 0.0857 | 0.03861 | 0.15285 | -0.0357 | -0.13623 | -0.02632 | 0.027938 | -0.01525 | -0.33473 | -0.18612 | -0.0708 | 0.033582 | -0.06934 | -0.02201 | -0.10225 | 0.20087 |
| ONS76 | 0.189006 | -0.09415 | -0.08122 | -0.01477 | 0.09195 | -0.01647 | -0.05232 | -0.08722 | -0.20918 | 0.16818 | 0.16407 | 0.09184 | 0.02208 | 0.035753 | -0.07736 | 0.144899 | 0.129704 | -0.20687 | -0.11329 | -0.1627 | -0.09393 | -0.13165 | -0.0193 | 0.12085 | 0.05471 |
| KYSE30 | -0.04047 | 0.037531 | -0.06543 | -0.29826 | 0.05658 | -0.75406 | -0.09156 | 0.19389 | 0.00864 | -0.09654 | 0.1525 | 0.07842 | -0.01148 | 0.121985 | -0.35544 | -0.00439 | 0.150947 | -0.6529 | -0.67262 | -0.00051 | -0.05146 | -0.0925 | 0.13857 | -0.0699 | 0.08434 |
| HSC3 | -0.15128 | 0.042256 | 0.0249 | -0.03716 | -0.1557 | -0.77102 | -0.08996 | 0.17435 | -0.16102 | 0.07038 | 0.0598 | -0.10023 | -0.05498 | 0.165382 | -0.1514 | 0.011016 | 0.144328 | -0.4227 | -0.22572 | 0.008905 | 0.132893 | -0.12927 | -0.00666 | -0.0961 | 0.07811 |
| NCIH2023 | 0.044314 | 0.085666 | -0.16043 | -0.00137 | 0.02932 | -0.26661 | 0.009902 | -0.0375 | -0.02015 | 0.00177 | 0.07283 | 0.0441 | -0.10495 | 0.059605 | -0.11708 | -0.01112 | -0.08602 | -0.24985 | -0.25054 | -0.0847 | -0.17749 | -0.05303 | 0.07302 | -0.01698 | 0.08532 |
| SEM | -0.13186 | -0.07402 | 0.08051 | -0.16088 | 0.02566 | 0.03648 | -0.10099 | -0.26206 | 0.01939 | 0.10209 | 0.20788 | 0.18464 | 0.110321 | -0.12624 | -0.02854 | -0.10209 | 0.034355 | -0.07927 | -0.18143 | -0.02155 | 0.02447 | 0.04036 | 0.03379 | -0.05043 | 0.13282 |
| CAMA1 | 0.008531 | 0.088118 | -0.07328 | -0.18169 | -0.08869 | 0.06452 | -0.14886 | 0.03468 | 0.02052 | 0.19628 | 0.20355 | -0.01214 | 0.059385 | 0.104625 | -0.16506 | 0.147136 | 0.010968 | -0.32366 | -0.30684 | -0.05121 | -0.08096 | 0.01059 | -0.00276 | -0.2109 | 0.10733 |
| KYSE70 | 0.086528 | 0.01226 | -0.07809 | 0.0015 | -0.02015 | -0.28739 | -0.05139 | 0.07892 | 0.0257 | 0.02779 | 0.24743 | -0.00556 | 0.06848 | 0.07407 | -0.48179 | 0.00173 | 0.103834 | -0.15346 | -0.21654 | -0.08045 | -0.13863 | -0.00488 | 0.04681 | -0.06027 | 0.1526 |
| NCIH2126 | -0.05054 | -0.00875 | 0.02293 | -0.01498 | -0.0272 | -0.55069 | 0.125185 | 0.03645 | -0.19639 | 0.03221 | 0.0531 | 0.03143 | 0.047723 | 0.044114 | 0.054378 | 0.077369 | 0.163194 | -0.43385 | -0.12086 | -0.14475 | -0.0701 | -0.0585 | 0.23424 | 0.05678 | 0.24107 |
| LXF289 | 0.060492 | -0.08481 | -0.05716 | -0.07304 | 0.01422 | -0.20683 | 0.0181 | 0.12153 | 0.07562 | 0.05186 | 0.11975 | 0.00308 | -0.04941 | 0.118674 | -0.08685 | -0.0623 | -0.03224 | -0.40385 | -0.34006 | -0.04665 | -0.08216 | -0.00955 | 0.02532 | 0.02791 | -0.00281 |
| A2058 | 0.041109 | -0.02116 | 0.0052 | -0.01969 | -0.05164 | -0.11852 | 0.019334 | -0.01143 | 0.05556 | 0.06489 | 0.06787 | 0.10378 | 0.015636 | 0.013562 | -0.06357 | 0.086671 | 0.051193 | -0.32714 | -0.33105 | -0.00371 | 0.061404 | -0.03168 | 0.06984 | 0.04907 | 0.08579 |
| SHP77 | 0.114321 | 0.016542 | 0.10211 | -0.03579 | -0.20541 | -0.05796 | -0.07785 | -0.13806 | 0.01155 | 0.15612 | 0.07621 | -0.00351 | 0.224309 | 0.009558 | -0.05018 | 0.040498 | 0.253147 | -0.16591 | -0.05608 | -0.09729 | -0.02784 | -0.08504 | -0.10365 | 0.09665 | -0.00679 |
| RERFLCAD1 | -0.03476 | -0.06058 | -0.13904 | -0.07551 | 0.1383 | -0.35123 | 0.01557 | 0.00373 | -0.09912 | 0.20609 | 0.19434 | 0.09979 | 0.015716 | 0.081984 | -0.30421 | 0.104258 | 0.133161 | -0.31406 | -0.14509 | -0.00164 | -0.18421 | -0.18371 | -0.08422 | 0.05015 | 0.20372 |
| BFTC909 | -0.03581 | -0.16462 | 0.0525 | 0.1497 | -0.01914 | -0.32277 | 0.15651 | 0.41387 | -0.03629 | 0.11527 | 0.11572 | 0.10544 | 0.043832 | 0.2256 | 0.021383 | -0.06833 | 0.105529 | -0.15958 | -0.19033 | -0.1309 | -0.56176 | -0.00255 | -0.07788 | 0.01541 | 0.12568 |
| KATOIII | 0.110487 | 0.057995 | -0.01663 | -0.09021 | 0.03 | -0.19833 | -0.01032 | -0.13607 | -0.01345 | 0.01868 | 0.07311 | 0.09341 | 0.082581 | 0.100556 | -0.16186 | -0.01357 | 0.072818 | -0.13157 | -0.46324 | -0.18068 | -0.01494 | -2.47872 | 0.16984 | -0.01784 | 0.10683 |
| BICR22 | 0.107373 | 0.027304 | -0.04 | -0.05214 | -0.05307 | -0.70865 | -0.0209 | 0.18284 | -0.10787 | 0.07825 | 0.2642 | 0.14264 | 0.119955 | 0.250239 | 0.041368 | -0.01559 | 0.009678 | -1.07166 | -0.79022 | 0.067361 | -0.11797 | -0.07151 | -0.06971 | -0.024 | 0.22616 |
| MCAS | -0.07362 | -0.26873 | -0.07622 | -0.1426 | -0.03235 | -0.75578 | -0.09334 | 0.02438 | 0.03111 | 0.05728 | 0.31453 | 0.13417 | 0.042776 | 0.133099 | 0.000974 | 0.0464 | 0.001001 | -0.35778 | -0.21992 | -0.17838 | -0.10118 | -0.03044 | -0.04453 | -0.07418 | 0.1744 |
| HS695T | -0.4045 | -0.05145 | 0.22027 | 0.14736 | -0.20061 | -0.15762 | -0.14729 | 0.01349 | 0.03324 | 0.17317 | 0.21976 | -0.00704 | 0.013036 | 0.237206 | 0.062216 | 0.272485 | 0.116314 | -0.31798 | -0.32439 | -0.14086 | 0.03011 | -0.01703 | 0.0158 | 0.02526 | 0.18394 |
| NCIH446 | -0.16 | 0.26753 | 0.00666 | 0.03222 | -0.1584 | -0.01528 | 0.204825 | -0.15277 | -0.32895 | 0.08914 | 0.04285 | -0.2022 | 0.018559 | 0.126405 | -0.04093 | 0.003075 | 0.074686 | -0.54999 | -0.01473 | 0.026704 | -0.00055 | 0.02979 | -0.26092 | -0.06413 | 0.16411 |
| HS936T | -0.09199 | 0.204088 | 0.15908 | 0.08354 | 0.01048 | -0.26199 | 0.083486 | 0.04789 | -0.1714 | 0.04349 | 0.06967 | 0.15761 | 0.191042 | 0.026317 | -0.19334 | 0.199924 | -0.02437 | -0.0231 | -0.28511 | -0.0627 | -0.04427 | -0.12071 | -0.08638 | 0.03491 | 0.00695 |
| BFTC905 | -0.04275 | 0.074506 | 0.05238 | -0.06297 | 0.03784 | -0.06174 | 0.03777 | -0.08106 | -0.00936 | 0.06232 | 0.06487 | 0.15946 | -0.06721 | -0.12754 | -0.0707 | -0.05758 | 0.141007 | -0.22536 | -0.31227 | -0.00537 | -0.01668 | -0.09046 | 0.05398 | -0.1292 | 0.32863 |
| NB1 | -1.16295 | 0.046511 | -0.08206 | -0.13998 | 0.02611 | -0.09848 | 0.330578 | -0.07131 | -0.23336 | -0.06893 | -0.02504 | 0.15603 | 0.041986 | -0.0392 | -0.02004 | -0.10844 | 0.032902 | -0.27477 | -0.06432 | -0.12119 | 0.156079 | 0.07983 | 0.0849 | 0.09695 | 0.12348 |
| COLO679 | -0.17379 | 0.062041 | 0.0378 | -0.09593 | -0.07503 | -0.1817 | 0.121663 | -0.04202 | -0.13625 | 0.23014 | 0.14594 | 0.18354 | 0.100706 | -0.05667 | 0.158669 | -0.0597 | 0.295993 | -0.33154 | -0.46694 | -0.23981 | -0.06366 | -0.20879 | -0.01337 | 0.14986 | 0.06112 |

| Cell line | ALK | AXL | CSF1R | DDR1 | DDR2 | EGFR | EPHA1 | EPHA2 | EPHA3 | EPHA4 | EPHA6 | EPHA7 | EPHB1 | EPHB2 | EPHB3 | EPHB4 | EPHB6 | ERBB2 | ERBB3 | ERBB4 | FGFR1 | FGFR2 | FGFR3 | FGFR4 | FLT1 |
|---|---|---|---|---|---|---|---|---|---|---|---|---|---|---|---|---|---|---|---|---|---|---|---|---|---|
| SNU738 | 0.104946 | -0.00062 | -0.10935 | -0.12138 | 0.05744 | -0.0607 | -0.02147 | -0.11253 | -0.09074 | -0.08976 | 0.22496 | 0.09602 | 0.067241 | 0.213575 | -0.13945 | 0.051981 | 0.138603 | -0.44742 | -0.25908 | -0.01488 | -0.22301 | -0.30273 | 0.07603 | 0.08231 | 0.18715 |
| KYSE410 | -0.06986 | 0.15786 | -0.02793 | -0.04096 | -0.06726 | -0.23287 | -0.02683 | 0.08319 | -0.01843 | 0.14917 | 0.10842 | 0.21087 | -0.01753 | 0.021611 | -0.0056 | 0.018874 | -0.17316 | -2.16649 | -0.94523 | -0.12362 | 0.081385 | -0.0545 | 0.04605 | 0.03735 | 0.03453 |
| SKMEL30 | -0.01702 | 0.207722 | 0.03991 | -0.00278 | -0.0014 | -0.02834 | -0.18258 | -0.14248 | -0.17725 | 0.20703 | -0.43588 | -0.07235 | 0.132177 | -0.08534 | -0.22076 | 0.098682 | 0.032162 | -0.34854 | -0.19018 | -0.2095 | -0.02199 | 0.31104 | -0.16732 | -0.15521 | 0.37435 |
| SKOV3 | -0.03973 | -0.059 | 0.10929 | 0.06687 | -0.21605 | -0.07622 | -0.02155 | 0.55809 | 0.00422 | 0.19203 | 0.15373 | -0.03696 | 0.087619 | 0.230951 | -0.18559 | 0.090868 | 0.024276 | -1.67053 | -0.28545 | -0.07593 | -0.11691 | -0.05723 | -0.04498 | -0.0903 | 0.07753 |
| RPMI8226 | 0.074816 | 0.114933 | 0.07568 | -0.00654 | 0.02136 | -0.08534 | 0.005016 | -0.05898 | -0.15999 | 0.14215 | 0.23797 | 0.09532 | 0.088831 | -0.17588 | 0.032117 | 0.089775 | 0.090427 | -0.1289 | -0.26138 | -0.01694 | -0.03817 | -0.38085 | 0.12099 | 0.05657 | 0.12476 |
| LN18 | 0.014593 | 0.010276 | -0.02503 | -0.06406 | -0.10896 | -0.09797 | 0.031099 | -0.10872 | 0.12437 | 0.03795 | 0.10368 | 0.19844 | 0.022009 | -0.12218 | 0.05287 | -0.09607 | 0.142746 | -0.2747 | -0.19507 | -0.15912 | -0.1539 | -0.03969 | 0.07471 | 0.03771 | 0.05398 |
| SW403 | -0.24234 | 0.004674 | 0.16271 | 0.09872 | -0.22966 | 0.03681 | -0.55929 | 0.1013 | 0.02788 | 0.10042 | -0.03047 | 0.0908 | 0.096669 | 0.026219 | -0.08368 | 0.077251 | -0.18946 | -0.43118 | -0.16949 | 0.149835 | -0.05447 | 0.03125 | 0.13892 | -0.09572 | -0.00929 |
| EJM | -0.06182 | -0.0284 | 0.10258 | -0.18681 | -0.05596 | -0.09406 | 0.148644 | -0.19632 | -0.04752 | 0.06738 | 0.18084 | -0.01196 | -0.01977 | 0.070754 | 0.014466 | 0.009724 | 0.283092 | -0.39596 | -0.14919 | 0.036341 | -0.01278 | 0.13377 | -0.08362 | 0.06925 | -0.12851 |
| SKMEL24 | 0.017383 | -0.04474 | 0.05871 | 0.01176 | 0.063 | -0.26129 | 0.071046 | -0.12767 | -0.17711 | 0.05097 | 0.1337 | 0.02824 | 0.145134 | -0.08004 | -0.0822 | 0.078445 | -0.00239 | -0.31179 | -0.24611 | -0.08432 | -0.02791 | 0.06917 | -0.06692 | 0.06133 | 0.08077 |
| KYSE140 | 0.080496 | 0.000523 | -0.12366 | -0.0185 | -0.044 | -0.7542 | 0.044777 | -0.01895 | 0.06443 | 0.16446 | 0.15862 | 0.1152 | -0.00381 | 0.080628 | -0.05704 | 0.000545 | 0.125146 | -0.36238 | -0.51859 | -0.09747 | -0.04738 | -0.06175 | 0.01149 | -0.05778 | -0.0975 |
| KYSE510 | 0.021613 | 0.050542 | -0.17627 | 0.03274 | -0.07389 | -0.20601 | -0.03209 | 0.09191 | 0.10567 | 0.08283 | 0.12144 | 0.07923 | 0.042046 | 0.061 | -0.07815 | -0.0215 | -0.03012 | -0.45009 | -0.45233 | -0.16983 | -0.08288 | -0.00763 | 0.09797 | -0.07043 | 0.03392 |
| WM793 | 0.003636 | -0.02334 | -0.01158 | -0.05204 | -0.19147 | 0.10228 | -0.0102 | -0.02622 | -0.04081 | 0.18411 | 0.16924 | -0.04771 | 0.058493 | 0.097848 | -0.08633 | 0.045914 | 0.13948 | -0.4308 | -0.42253 | 0.006836 | 0.105053 | -0.07856 | 0.04097 | -0.02496 | 0.14969 |
| HUNS1 | 0.041959 | -0.0406 | 0.00606 | 0.07486 | 0.10012 | -0.13466 | 0.149717 | -0.16807 | 0.04964 | 0.14718 | 0.05844 | 0.16735 | 0.139024 | 0.056927 | -0.14591 | 0.020245 | 0.024709 | -0.27388 | -0.14277 | -0.00249 | -0.00219 | -0.13341 | -0.06819 | -0.16009 | 0.03026 |
| HEC50B | 0.119844 | 0.100854 | -0.04673 | -0.04065 | -0.05753 | -0.46247 | -0.1654 | 0.06327 | -0.13107 | -0.00559 | 0.20652 | 0.12355 | 0.125366 | -0.0195 | -0.00794 | -0.03147 | 0.226638 | -0.02739 | -0.2328 | -0.05096 | 0.012267 | 0.00139 | 0.07551 | -0.03846 | 0.07205 |
| CAL27 | 0.091996 | -0.00159 | -0.0317 | 0.05052 | -0.08038 | -0.66494 | 0.016498 | 0.27655 | -0.23936 | 0.10911 | -0.02075 | 0.03152 | -0.01948 | 0.102566 | -0.05088 | 0.023495 | 0.065728 | -0.58752 | -0.85135 | -0.09917 | -0.10691 | -0.11481 | 0.05756 | 0.10052 | 0.18318 |
| RH30 | 0.098453 | 0.259478 | -0.02977 | 0.00138 | -0.10761 | -0.32005 | -0.00577 | -0.0848 | 0.18024 | -0.25975 | 0.183 | 0.19462 | -0.03177 | 0.015908 | 0.17792 | 0.100307 | 0.183127 | -0.16304 | -0.14547 | -0.21006 | 0.176401 | 0.08788 | 0.26907 | -0.14539 | 0.10787 |
| UMUC1 | 0.099779 | -0.13937 | -0.0045 | -0.03533 | 0.05252 | -0.14883 | -0.06722 | 0.42107 | 0.03326 | 0.18113 | 0.15926 | 0.15462 | 0.022738 | -0.10757 | -0.11447 | -0.17005 | 0.079917 | -0.19702 | -0.13415 | -0.16161 | -0.12368 | -0.56149 | -0.23038 | 0.07219 | 0.15452 |
| GCT | -0.0021 | -0.23534 | 0.07866 | 0.02982 | 0.01803 | -0.05073 | -0.10679 | 0.0003 | -0.00659 | -0.04528 | 0.23218 | 0.02382 | 0.064444 | 0.134385 | -0.02117 | 0.126265 | -0.04654 | -0.23212 | -0.19346 | -0.1502 | -0.03798 | 0.00697 | 0.09311 | -0.02793 | 0.06343 |
| YD15 | 0.149284 | 0.101491 | -0.01172 | -0.03328 | -0.09567 | -0.81012 | -0.03812 | 0.24445 | -0.06357 | 0.08257 | 0.28859 | 0.071 | -0.09909 | 0.104488 | -0.15191 | 0.078428 | 0.133543 | -0.38533 | -0.41747 | -0.10141 | -0.06461 | 0.1233 | 0.00963 | 0.01044 | 0.22774 |
| NCIH322 | -0.09262 | 0.022368 | -0.02702 | -0.00426 | 0.00315 | -0.59824 | 0.053005 | -0.0512 | -0.05779 | 0.1483 | 0.0446 | 0.03959 | 0.043808 | 0.012157 | 0.125146 | 0.06728 | -0.0438 | -0.49117 | -0.10208 | -0.17365 | 0.023031 | 0.12545 | 0.03335 | -0.15507 | -0.05097 |
| AMO1 | 0.005548 | -0.15159 | 0.21671 | 0.16721 | 0.13331 | -0.32847 | -0.00873 | -0.04313 | 0.20616 | 0.03796 | 0.0099 | 0.02414 | -0.10487 | 0.003422 | -0.05892 | -0.26472 | 0.291921 | -0.30561 | -0.16578 | -0.22645 | -0.04119 | -0.01325 | 0.02704 | 0.04293 | 0.01333 |
| SCABER | 0.024857 | 0.117342 | 0.00041 | -0.03988 | 0.05868 | -0.84394 | 0.103993 | 0.21972 | 0.03488 | 0.03293 | 0.20416 | 0.08 | -0.13201 | 0.021503 | -0.05142 | 0.012704 | -0.15403 | -0.75253 | -0.50653 | -0.12235 | -0.20479 | -0.04563 | -0.00112 | -0.05569 | 0.23134 |
| HCC366 | 0.009756 | -0.09784 | -0.24359 | -0.17143 | -0.1847 | -0.44205 | -0.15827 | 0.38022 | -0.01903 | 0.02596 | 0.14188 | 0.02221 | -0.12269 | 0.187428 | -0.08341 | -0.185 | 0.090555 | -0.32938 | -0.15653 | -0.03286 | -0.24865 | -0.25012 | 0.13422 | 0.18335 | 0.04674 |
| NCIH2087 | 0.02548 | 0.03662 | -0.12793 | 0.00084 | 0.01406 | -0.1221 | 0.071341 | -0.1133 | -0.14429 | 0.06817 | 0.13434 | 0.10335 | 0.037491 | 0.098767 | -0.18732 | 0.072719 | 0.082423 | -0.30597 | -0.15245 | -0.09637 | -0.09444 | -0.1859 | 0.01805 | -0.0208 | 0.10584 |
| HARA | -0.04289 | -0.02199 | -0.00097 | -0.09604 | 0.00415 | -0.30881 | -0.02261 | -0.09472 | -0.1753 | 0.0821 | 0.15014 | 0.05814 | 0.042648 | -0.08043 | 0.003484 | 0.033152 | -0.03009 | -1.08113 | -1.03147 | -0.09225 | -0.00234 | -0.06575 | -0.06346 | -0.08431 | 0.11804 |
| NCIH1373 | -0.11368 | 0.126795 | -0.04388 | -0.08082 | -0.1156 | -0.05713 | -0.06341 | -0.04403 | -0.24454 | -0.16398 | 0.15065 | 0.02534 | -0.13144 | 0.213588 | -0.29777 | 0.100396 | 0.098208 | -0.25264 | -0.37657 | 0.146893 | 0.078457 | 0.01899 | 0.14919 | -0.03428 | 0.22125 |
| FADU | 0.084623 | 0.098312 | 0.01756 | -0.0266 | -0.11358 | -0.17949 | 0.080806 | 0.01465 | -0.01786 | 0.11028 | 0.07247 | 0.0959 | 0.039153 | 0.115445 | -0.06597 | 0.112596 | 0.148954 | -0.28407 | -0.29589 | -0.10649 | -0.11769 | -0.02228 | 0.08146 | 0.02597 | 0.03709 |
| HGC27 | -0.05111 | -0.04715 | -0.1398 | -0.11004 | -0.03445 | -0.05641 | 0.022409 | 0.10767 | 0.13383 | 0.18732 | 0.00803 | 0.12674 | 0.001872 | -0.00166 | -0.04581 | 0.016883 | 0.031356 | -0.31966 | -0.17029 | -0.14717 | 0.008514 | -0.05116 | -0.08499 | 0.08596 | 0.1091 |
| JHH7 | 0.138811 | -0.03613 | -0.10027 | -0.16141 | 0.03664 | -0.12172 | -0.06123 | -0.09383 | -0.06958 | 0.03862 | 0.1339 | 0.13085 | 0.007196 | 0.172998 | 0.050205 | -0.06709 | 0.17029 | -0.15571 | -0.30801 | -0.16388 | -0.10334 | -0.21705 | -0.07077 | -0.45544 | 0.12204 |
| MDAMB468 | 0.009631 | 0.147556 | -0.09024 | -0.0642 | -0.03286 | -0.13617 | 0.151737 | 0.22177 | -0.15536 | 0.1624 | 0.15705 | -0.0064 | 0.027433 | 0.210759 | -0.00277 | 0.098657 | -0.01327 | -0.09352 | -0.36458 | 0.05781 | -0.07535 | -0.04581 | 0.05881 | -0.1735 | 0.19091 |
| MORCPR | 0.054342 | -0.00016 | 0.02193 | 0.03651 | -0.02604 | -0.00923 | 0.158757 | -0.07393 | -0.01845 | 0.14409 | -0.03053 | 0.10023 | -0.16274 | 0.059523 | -0.1496 | 0.001238 | 0.117123 | -0.20401 | -0.04957 | -0.14402 | -0.20877 | -0.04134 | 0.04317 | 0.24755 | 0.05791 |
| NCIH661 | 0.035115 | -0.11862 | -0.04047 | -0.12529 | 0.00545 | -0.22767 | -0.07152 | 0.00129 | -0.08791 | 0.08548 | 0.13692 | 0.10005 | 0.009651 | 0.137252 | -0.0312 | -0.00278 | 0.180512 | -0.71951 | -0.50413 | -0.09608 | -0.1082 | -0.00216 | 0.08937 | 0.04071 | 0.22225 |
| OCIMY5 | -0.23131 | 0.200475 | 0.10325 | -0.07825 | -0.18518 | 0.11963 | 0.144321 | -0.09953 | -0.12064 | 0.08886 | 0.15067 | 0.31678 | 0.093987 | 0.013118 | -0.00866 | 0.093512 | -0.01076 | -0.27488 | -0.19994 | -0.30332 | -0.21778 | -0.2146 | -0.08659 | 0.05392 | 0.17508 |
| KYSE150 | -0.13224 | 0.099957 | 0.08098 | 0.03239 | -0.0832 | -0.36428 | -0.00333 | 0.04135 | 0.01668 | 0.08472 | 0.28185 | 0.0226 | -0.03812 | -0.12757 | 0.10739 | 0.060123 | 0.036364 | -0.22568 | -0.31046 | -0.07202 | -0.05571 | 0.18689 | 0.05775 | -0.05521 | 0.21352 |
| CAL51 | 0.058243 | 0.101836 | 0.01616 | 0.00217 | -0.00343 | -0.10416 | 0.043335 | -0.02025 | 0.01074 | 0.12932 | 0.1472 | 0.19411 | 0.05152 | 0.02343 | -0.25023 | 0.070052 | 0.089815 | -0.31476 | -0.13544 | 0.007286 | 0.027993 | -0.19798 | 0.00445 | 0.09487 | 0.00583 |
| KNS62 | 0.041177 | 0.029856 | -0.12106 | -0.11925 | -0.09139 | -0.13466 | -0.0569 | -0.17275 | -0.06546 | -0.03245 | -0.01038 | 0.04802 | -0.01805 | 0.080398 | 0.04616 | -0.00764 | 0.143834 | -0.287 | -0.26072 | -0.18393 | 0.015428 | 0.02593 | -0.02458 | 0.06477 | 0.12523 |
| HCC1954 | 0.001111 | 0.002652 | -0.04993 | 0.08509 | -0.07669 | -0.62112 | -0.01898 | 0.11319 | 0.08954 | 0.31241 | 0.11907 | -0.10864 | 0.074577 | 0.034012 | -0.17112 | 0.053378 | 0.17975 | -0.63752 | -0.29011 | 0.164165 | 0.047333 | -0.17516 | -0.00056 | -1.36252 | -0.05743 |
| NCIH358 | -0.02357 | -0.02853 | -0.01135 | 0.06865 | -0.03374 | -0.43629 | -0.17722 | -0.06769 | 0.0501 | 0.08306 | 0.18391 | 0.12688 | 0.026593 | 0.025597 | -0.13095 | -0.0378 | -0.04567 | -0.32614 | -0.43544 | 0.070201 | -0.09129 | 0.0155 | -0.02938 | -0.10037 | 0.17457 |
| HOP62 | 0.030636 | -0.04121 | 0.06004 | 0.12704 | -0.0063 | -0.05728 | -0.05383 | -0.0052 | -0.09691 | 0.03978 | 0.06271 | 0.17316 | 0.044413 | 0.21967 | 0.125177 | 0.016227 | 0.153983 | -0.25949 | -0.10574 | -0.26108 | -0.27243 | 0.02752 | -0.04158 | -0.01862 | -0.03235 |
| KMBC2 | -0.17165 | 0.058342 | -0.14368 | -0.03252 | -0.14489 | -0.85331 | 0.139096 | 0.24023 | -0.06416 | -0.04692 | 0.2017 | 0.07384 | -0.15309 | 0.080518 | 0.001075 | 0.067301 | 0.047954 | -0.50287 | -0.32401 | -0.04026 | -0.03484 | 0.03644 | 0.01061 | -0.06352 | 0.08579 |
| DBTRG05MG | -0.01462 | -0.27087 | 0.14984 | -0.02903 | -0.11162 | 0.02788 | 0.092785 | 0.24947 | 0.09824 | 0.02968 | 0.00433 | 0.04724 | 0.049126 | 0.178052 | -0.02012 | 0.072827 | 0.428652 | -0.10879 | 0.00464 | 0.12359 | 0.038088 | 0.02168 | -0.22428 | -0.09371 | 0.17539 |
| COLO684 | 0.228011 | 0.244876 | -0.04512 | 0.29324 | -0.07803 | 0.15002 | -0.0971 | -0.12351 | 0.03398 | -0.07782 | 0.01707 | 0.0668 | 0.286509 | 0.091461 | 0.202191 | 0.167143 | 0.161474 | 0.07719 | 0.0657 | -0.26649 | -0.22065 | -0.03839 | 0.30322 | 0.22928 | -0.01055 |
| KYSE450 | 0.115348 | 0.022873 | -0.02694 | -0.09333 | -0.03289 | -0.78702 | 0.00428 | 0.01958 | 0.02137 | -0.02782 | 0.16327 | 0.07521 | 0.071242 | -0.01447 | -0.08822 | -0.14232 | 0.106776 | -0.24833 | -0.51508 | -0.1178 | -0.07173 | -0.07642 | -0.08287 | 0.04218 | 0.10607 |
| NCIH1048 | 0.082978 | 0.002736 | -0.02596 | 0.00635 | -0.041 | -0.04561 | 0.121972 | -0.07529 | -0.00059 | 0.03653 | 0.19932 | 0.08993 | 0.057564 | 0.019607 | 0.002336 | -0.07726 | 0.069514 | -0.16937 | -0.16758 | -0.1859 | 0.044106 | -0.84363 | -0.25606 | -0.01866 | 0.06443 |
| CHAGOK1 | 0.134378 | 0.095804 | -0.04337 | -0.02014 | 0.00349 | -0.1441 | 0.013564 | -0.03643 | -0.04049 | 0.08258 | 0.08115 | 0.07204 | 0.054354 | 0.00393 | -0.08106 | -0.01366 | 0.052662 | -0.21212 | -0.22458 | -0.10331 | -0.06911 | -0.0343 | 0.00264 | -0.01219 | 0.16156 |
| NCIH1568 | -0.00278 | 0.11924 | 0.08791 | -0.13558 | -0.08449 | -0.20623 | 0.106996 | -0.01165 | 0.08122 | 0.06627 | 0.20153 | -0.20219 | -0.01125 | 0.036172 | -0.22283 | 0.088394 | 0.022706 | -0.18691 | -0.23862 | -0.15765 | 0.017127 | -0.28264 | 0.03116 | -0.01015 | 0.05303 |
| NCIH510 | -0.01104 | 0.08113 | -0.08349 | 0.05413 | -0.08351 | -0.00804 | -0.05662 | -0.13536 | -0.01017 | 0.05798 | 0.15327 | -0.15342 | -0.0182 | 0.060617 | 0.035576 | 0.024648 | 0.181089 | -0.33549 | -0.25865 | -0.06668 | -0.08856 | 0.00835 | 0.0107 | 0.08068 | 0.11511 |
| HCC515 | 0.030883 | 0.029303 | 0.17219 | -0.05671 | -0.25899 | -0.46636 | -0.0342 | -0.00149 | -0.18601 | 0.21436 | 0.04058 | -0.11894 | -0.01306 | 0.197029 | -0.18652 | -0.06394 | 0.086566 | -0.26628 | -0.19306 | -0.03317 | -0.12459 | -0.01496 | 0.13616 | -0.07418 | 0.14428 |
| KYSE270 | -0.03071 | 0.199639 | -0.00214 | -0.13805 | -0.03268 | -0.86856 | -0.07801 | 0.09829 | 0.12104 | 0.08114 | 0.16882 | -0.00508 | 0.002545 | 0.158479 | 0.010003 | 0.202135 | 0.146321 | -0.47705 | -0.48477 | -0.01778 | -0.03309 | -0.05886 | -0.00169 | -0.07049 | 0.37595 |
| MDAMB415 | 0.04147 | -0.00647 | -0.11006 | -0.01096 | -0.2576 | -0.00392 | 0.074982 | 0.22664 | -0.01281 | 0.18608 | 0.0525 | 0.04658 | 0.050463 | 0.023654 | -0.01437 | 0.139023 | 0.024032 | -0.30092 | -0.16856 | 0.062928 | -0.04064 | -1.17146 | 0.09329 | -0.82324 | -0.02584 |
| HCC15 | -0.12918 | 0.084387 | -0.14661 | -0.04449 | -0.05024 | -0.0183 | -0.22961 | -0.31788 | -0.05292 | -0.01997 | 0.1358 | 0.18199 | -0.07453 | -0.10437 | -0.14092 | 0.092779 | 0.011 | -0.16851 | -0.42665 | 0.032487 | -0.07973 | -0.25167 | 0.14174 | -0.17947 | 0.18516 |
| MFE296 | 0.094035 | 0.023628 | 0.08667 | 0.004 | -0.08677 | 0.20442 | -0.24599 | -0.08036 | -0.01834 | 0.22421 | 0.00964 | 0.29926 | 0.104286 | 0.348353 | 0.109198 | 0.138178 | 0.140629 | -0.19778 | -0.12731 | -0.02747 | -0.14074 | -1.48753 | 0.17471 | 0.14078 | 0.21329 |
| AGS | 0.060592 | 0.165014 | -0.02357 | 0.01817 | 0.01088 | -0.20588 | 0.05006 | 0.06964 | -0.04941 | 0.1278 | -0.02358 | 0.25475 | 0.194516 | 0.148535 | 0.03254 | 0.046304 | -0.17583 | -0.25631 | -0.05999 | 0.054726 | 0.025485 | -0.06399 | -0.05994 | -0.05078 | 0.31366 |
| MELJUSO | -0.01269 | -0.00529 | 0.01416 | -0.07211 | -0.10658 | 0.11278 | 0.027803 | 0.0666 | -0.02494 | 0.09641 | 0.11975 | -0.00748 | 0.061712 | 0.084193 | -0.03273 | 0.118376 | 0.162696 | -0.16404 | -0.21063 | -0.00793 | -0.04342 | -0.0551 | 0.11841 | 0.01821 | 0.06669 |
| IGR1 | 0.047679 | 0.013088 | -0.03237 | -0.0925 | -0.06011 | 0.08146 | -0.01371 | -0.11516 | 0.04067 | -0.01318 | 0.09517 | 0.09132 | 0.022158 | -0.03426 | -0.0926 | 0.086319 | 0.081287 | -0.05222 | -0.08032 | -0.09204 | -0.26196 | -0.10376 | 0.16397 | -0.03066 | 0.13534 |
| MDAMB435S | -0.05889 | -0.06008 | 0.0942 | -0.18433 | 0.05146 | -0.09672 | 0.030905 | -0.23712 | -0.08282 | 0.07455 | 0.18442 | 0.09152 | 0.121295 | -0.03992 | -0.19071 | -0.06084 | -0.01801 | -0.13013 | -0.28536 | -0.0947 | -0.05766 | -0.02895 | -0.0748 | 0.0214 | 0.04179 |
| TOV21G | -0.06309 | 0.144403 | -0.00767 | -0.13014 | 0.07257 | -0.1477 | -0.05472 | 0.00556 | 0.08433 | 0.05643 | 0.14195 | 0.00432 | 0.068823 | 0.097281 | 0.074579 | 0.036857 | 0.063806 | -0.21486 | -0.09988 | -0.11026 | -0.06218 | -0.09777 | 0.05855 | 0.02716 | 0.35249 |
| NCIH2009 | 0.066661 | -0.02966 | -0.02715 | -0.02838 | -0.01687 | -0.20223 | -0.12265 | -0.0763 | -0.09215 | -0.0478 | 0.17469 | 0.08745 | -0.01587 | 0.093948 | -0.02386 | 0.119071 | 0.022849 | -0.14461 | -0.0914 | 0.016815 | -0.00515 | -0.07709 | 0.07299 | 0.04581 | 0.19854 |
| SF172 | 0.094208 | -0.14061 | -0.07601 | -0.0342 | 0.05147 | -0.69795 | 0.011534 | 0.08928 | 0.10195 | -0.00753 | 0.18225 | 0.08727 | 0.136351 | 0.091563 | -0.19083 | -0.10488 | 0.085675 | -0.14765 | -0.28878 | -0.12673 | -0.06728 | -0.08718 | -0.01723 | -0.01941 | 0.14536 |
| NCIH1793 | -0.03822 | 0.005834 | 0.02676 | -0.04413 | 0.00583 | -0.28178 | -0.08325 | 0.00377 | -0.03132 | 0.01619 | 0.19834 | 0.02302 | 0.018053 | 0.092511 | -0.07769 | -0.00643 | 0.166149 | -0.3997 | -0.56921 | 0.058621 | -0.47729 | -0.08407 | 0.03189 | -0.02046 | 0.13269 |
| KMM1 | 0.129899 | 0.040764 | 0.03662 | -0.07299 | -0.14334 | -0.12335 | -0.05916 | -0.0674 | 0.09316 | 0.12372 | 0.14306 | 0.08497 | 0.076682 | 0.044998 | -0.10422 | 0.036764 | 0.098869 | -0.3617 | -0.21412 | -0.05945 | -0.2675 | -0.03743 | 0.03858 | 0.15199 | 0.14533 |
| SW1271 | -0.08708 | 0.148466 | -0.00083 | 0.02204 | -0.05883 | -0.03684 | 0.139899 | 0.25483 | -0.03609 | -0.02887 | 0.08176 | 0.12025 | -0.0373 | 0.151561 | 0.060416 | 0.093704 | 0.117506 | -0.30478 | -0.17663 | -0.01907 | -0.11896 | -0.02764 | 0.08575 | -0.02366 | 0.16182 |
| NCIH1869 | -0.17578 | -0.04936 | 0.11544 | -0.01378 | -0.12086 | -0.51995 | 0.002275 | -0.23189 | 0.00083 | 0.16917 | 0.35059 | -0.10276 | 0.05397 | -0.16678 | -0.07259 | -0.07717 | 0.026497 | -0.62139 | -0.08792 | -0.03492 | -0.02951 | -0.03568 | 0.02435 | -0.05473 | 0.05843 |
| 647V | 0.030586 | -0.42263 | -0.13673 | 0.00902 | -0.07244 | -0.54672 | 0.142996 | 0.2233 | -0.10944 | 0.06313 | 0.05958 | 0.22103 | 0.027819 | 0.082325 | 0.127376 | 0.007785 | 0.229533 | -0.33127 | -0.28053 | -0.15383 | -0.24414 | -0.1356 | 0.1001 | 0.01651 | 0.1263 |
| SNU719 | -0.06328 | -0.03785 | 0.10078 | 0.12203 | 0.01475 | -0.6077 | 0.20828 | -0.11838 | -0.15845 | 0.00029 | 0.14472 | 0.01833 | 0.066789 | -0.03873 | -0.1607 | 0.077577 | 0.177087 | -0.27205 | -0.25914 | -0.05204 | -0.20174 | -0.00147 | 0.06201 | 0.01295 | 0.09876 |
| WM88 | -0.0085 | -0.04329 | -0.30614 | 0.05441 | -0.22509 | -0.17438 | 0.092852 | -0.16888 | -0.14978 | -0.02567 | 0.14296 | -0.07711 | 0.199438 | -0.03748 | 0.267756 | 0.045578 | 0.050479 | -0.4992 | -0.10525 | 0.080164 | 0.014307 | 0.24692 | -0.05194 | 0.30718 | 0.09409 |
| NCIH23 | 0.066938 | 0.105337 | 0.02081 | 0.02912 | 0.07277 | 0.025 | 0.033379 | -0.16781 | -0.15404 | 0.07095 | 0.16033 | -0.0041 | -0.00073 | 0.039954 | -0.03101 | 0.029572 | 0.09177 | -0.13566 | -0.30069 | 0.008175 | -0.00129 | 0.09345 | 0.03977 | -0.00076 | 0.07248 |
| HCC1359 | -0.03595 | -0.01168 | -0.08008 | 0.01347 | 0.0159 | -0.0611 | 0.013718 | -0.10011 | -0.06011 | 0.03853 | 0.09209 | 0.09067 | 0.057661 | 0.04616 | -0.04419 | -0.04005 | 0.167509 | -0.12073 | -0.16779 | -0.11113 | -1.51788 | -0.03577 | 0.10138 | -0.05611 | 0.02994 |
| FTC133 | 0.083624 | -0.04612 | -0.05913 | -0.08211 | 0.05851 | -0.24721 | 0.079056 | 0.08821 | -0.10593 | 0.0631 | 0.08845 | 0.15646 | 0.049756 | 0.010059 | -0.13937 | 0.022225 | 0.050081 | -0.2863 | -0.14113 | -0.06185 | -0.31469 | -0.05265 | 0.05859 | 0.06524 | 0.15024 |
| 5637 | 0.203483 | 0.085451 | 0.05069 | -0.05678 | -0.10131 | -0.62362 | 0.09415 | 0.15021 | -0.10598 | 0.10661 | 0.13151 | 0.06792 | 0.036462 | 0.169023 | 0.020537 | 0.009414 | 0.124256 | -0.5865 | -0.29967 | -0.04824 | -0.10841 | -0.1795 | 0.01127 | 0.02997 | 0.0936 |
| ES2 | -0.02554 | -0.12946 | -0.06913 | -0.10245 | -0.02237 | -0.13002 | -0.10287 | -0.12881 | 0.01401 | 0.07485 | 0.13895 | 0.14033 | 0.018182 | 0.096166 | -0.08612 | -0.03725 | 0.026897 | -0.26257 | -0.12769 | -0.13902 | -0.07184 | -0.07016 | -0.1533 | -0.01626 | 0.14903 |
| SNU349 | 0.03336 | -0.49085 | 0.1029 | 0.02064 | -0.18714 | -0.22364 | -0.09212 | 0.11834 | -0.12278 | 0.08206 | 0.29185 | -0.07931 | 0.068181 | 0.307224 | -0.05 | 0.188956 | 0.195445 | -0.12623 | -0.25116 | 0.100577 | -0.72234 | -0.19976 | -0.05835 | -0.19526 | 0.10099 |
| MDAMB453 | -0.10249 | 0.025445 | -0.07355 | 0.00522 | -0.1499 | 0.09012 | -0.13486 | 0.02054 | 0.07009 | 0.18191 | 0.27585 | 0.04394 | 0.052036 | 0.044957 | -0.14281 | 0.091942 | -0.01976 | -2.40526 | -1.24062 | -0.00907 | 0.057027 | 0.1217 | 0.04766 | -1.86684 | 0.07972 |
| NUGC3 | 0.039298 | 0.039766 | 0.04327 | -0.0171 | 0.07913 | -2.02214 | 0.015928 | -0.22911 | 0.00067 | 0.02155 | 0.09372 | 0.07592 | 0.0939 | 0.041049 | -0.10548 | -0.03769 | 0.185595 | -0.20598 | -0.16442 | -0.10192 | -0.12132 | 0.00902 | -0.018 | -0.01041 | -0.02409 |

| Cell line | ALK | AXL | CSF1R | DDR1 | DDR2 | EGFR | EPHA1 | EPHA2 | EPHA3 | EPHA4 | EPHA6 | EPHA7 | EPHB1 | EPHB2 | EPHB3 | EPHB4 | EPHB6 | ERBB2 | ERBB3 | ERBB4 | FGFR1 | FGFR2 | FGFR3 | FGFR4 | FLT1 |
|---|---|---|---|---|---|---|---|---|---|---|---|---|---|---|---|---|---|---|---|---|---|---|---|---|---|
| NCIH2286 | 0.025295 | 0.043529 | -0.04278 | 0.03654 | 0.03874 | -0.21492 | -0.01906 | -0.04055 | -0.02849 | 0.13159 | 0.0187 | 0.13905 | -0.02165 | 0.095672 | -0.12089 | -0.00215 | 0.050618 | -0.09703 | -0.27457 | -0.03244 | -0.21839 | -0.11534 | 0.02794 | -0.03745 | 0.041 |
| ESS1 | 0.145703 | -0.16085 | -0.03538 | -0.10227 | -0.05994 | -0.13821 | 0.096095 | -0.00747 | 0.06246 | 0.04848 | 0.23692 | 0.08829 | 0.016709 | 0.070295 | -0.01002 | 0.04267 | -0.01842 | -0.2802 | -0.25328 | -0.04281 | -0.47567 | -0.04376 | 0.09923 | 0.06311 | 0.11775 |
| HT | -0.04035 | 0.012375 | -0.14643 | 0.03808 | -0.13748 | -0.14926 | 0.032455 | -0.14374 | -0.04326 | 0.06862 | 0.07684 | -0.01937 | 0.009109 | -0.01309 | 0.004161 | -0.01314 | -0.00735 | -0.27892 | -0.08342 | -0.07946 | -0.00714 | -0.07945 | 0.03878 | -0.04586 | 0.17315 |
| IPC298 | 0.076711 | -0.04642 | 0.09186 | -0.09884 | -0.03111 | -0.25838 | 0.058618 | -0.0807 | 0.16192 | 0.00494 | 0.04763 | 0.04825 | 0.127746 | -0.03304 | 0.046132 | 0.054784 | 0.225504 | -0.08062 | -0.22657 | -0.09011 | 0.016162 | -0.19832 | 0.10978 | 0.08953 | 0.02938 |
| NCIH1573 | 0.625734 | -0.07159 | 0.03786 | -0.32371 | -0.15794 | 0.0257 | -0.18678 | -0.2093 | -0.00193 | 0.12354 | 0.14984 | -0.14575 | -0.24998 | 0.064024 | 0.334452 | 0.293243 | -0.28581 | -0.12663 | -0.06921 | 0.024049 | 0.141318 | 0.02879 | -0.27695 | 0.23735 | 0.0508 |
| TE4 | 0.006407 | 0.002139 | 0.10748 | -0.00898 | 0.04122 | -0.06525 | 0.072685 | -0.02077 | -0.02136 | 0.09191 | 0.08726 | 0.04075 | 0.071208 | -0.09317 | -0.12061 | -0.14343 | -0.00144 | -1.74824 | -0.25364 | -0.21098 | 0.00832 | 0.03513 | 0.09361 | 0.07424 | -0.00061 |
| IM95 | 0.18404 | 0.01923 | 0.06843 | 0.05073 | -0.33922 | -0.78557 | 0.108288 | -0.0111 | -0.28025 | 0.13153 | -0.24154 | 0.07062 | 0.153387 | 0.298991 | 0.079512 | 0.240838 | 0.167221 | -0.99543 | -0.01704 | -0.27369 | -0.23754 | -0.08282 | 0.13734 | 0.13785 | 0.08773 |
| CMLT1 | -0.01138 | -0.14992 | -0.20241 | 0.14143 | 0.01853 | 0.16576 | 0.351171 | -0.0701 | 0.01141 | 0.12298 | 0.04166 | 0.17931 | -0.04821 | -0.13418 | 0.132549 | -0.05614 | 0.273769 | -0.49209 | 0.01244 | -0.3062 | 0.073992 | 0.02053 | 0.15534 | 0.2793 | -0.28132 |
| NCIH157DM | -0.0861 | 0.039039 | -0.04995 | 0.00281 | -0.15969 | -0.22722 | 0.079425 | 0.03933 | 0.01595 | 0.09634 | 0.17642 | 0.109 | 0.003845 | 0.023821 | 0.124633 | 0.004554 | 0.153782 | -0.14696 | -0.12473 | -0.12352 | -0.02037 | -0.07732 | -0.16469 | 0.09021 | 0.08907 |
| RCHACV | 0.137293 | -0.04856 | 0.06302 | 0.00386 | 0.02955 | -0.21467 | -0.20788 | -0.06562 | 0.05582 | 0.07358 | 0.12563 | -0.13331 | -0.10759 | -0.1131 | -0.18094 | 0.199786 | 0.139045 | -0.15683 | -0.14621 | -0.04113 | -0.22787 | -0.07244 | -0.06752 | 0.02414 | 0.35283 |
| NCIH2172 | -0.07612 | 0.080646 | -0.00845 | -0.01587 | -0.04172 | -0.2739 | -0.11148 | -0.05198 | -0.16444 | 0.03789 | 0.11911 | 0.07103 | 0.125845 | 0.102287 | -0.06144 | 0.00261 | 0.018383 | -0.14553 | -0.09807 | 0.055577 | -0.09924 | -0.09543 | -0.02376 | -0.01557 | 0.1592 |
| HT55 | 0.026343 | 0.019879 | 0.09791 | 0.02739 | -0.22155 | -0.47774 | -0.05056 | 0.07723 | 0.02946 | 0.17475 | 0.19645 | -0.0435 | 0.25424 | 0.110843 | -0.08105 | 0.140286 | 0.290374 | -0.47113 | -0.11327 | -0.06968 | 0.055563 | 0.00592 | -0.01087 | -0.14078 | 0.19754 |
| JHUEM1 | -0.31992 | 0.316641 | 0.13659 | 0.05504 | -0.20761 | -0.2268 | -0.01053 | 0.01423 | -0.13429 | 0.06514 | 0.01792 | 0.04981 | 0.143056 | 0.198102 | -0.12273 | 0.249761 | 0.136472 | -0.38382 | -0.26002 | -0.23081 | 0.162935 | 0.01523 | -0.11075 | 0.01069 | 0.16528 |
| NCIH2110 | 0.12435 | 0.142017 | 0.05857 | 0.11524 | -0.03343 | -0.30282 | -0.02312 | -0.12393 | -0.13583 | 0.19076 | 0.07784 | 0.12803 | -0.01171 | 0.087934 | 0.011948 | -0.01005 | 0.224532 | -0.26116 | -0.14878 | -0.23739 | -0.06389 | -0.12332 | 0.19343 | 0.08492 | 0.10078 |
| SNU1 | 0.014384 | -0.01012 | -0.12265 | -0.02308 | -0.10907 | -0.11354 | 0.038865 | -0.05776 | 0.12509 | 0.17728 | 0.08142 | 0.06422 | 0.121312 | -0.06284 | -0.101 | 0.007891 | 0.101385 | -0.1878 | -0.28936 | -0.00165 | -0.07144 | -0.05953 | 0.08024 | -0.00753 | 0.20535 |
| MDAMB361 | 0.151735 | 0.003828 | -0.15794 | -0.01925 | 0.02686 | -0.16004 | 0.076326 | 0.09215 | 0.0545 | 0.01329 | -0.2586 | 0.10958 | 0.078318 | 0.087814 | -0.2758 | 0.073071 | 0.041123 | -3.37447 | -2.52229 | 0.03982 | -0.02836 | 0.09649 | 0.00987 | -0.09219 | 0.107 |
| MDST8 | -0.15252 | 0.007338 | 0.14746 | 0.08905 | -0.00541 | -0.12386 | 0.111402 | 0.07182 | -0.00978 | 0.07843 | 0.05259 | 0.17627 | 0.182094 | -0.04753 | -0.18234 | 0.060813 | -0.10657 | -0.04977 | -0.08027 | 0.001363 | -0.07265 | -0.09473 | -0.2081 | 0.03831 | 0.22293 |
| EFO27 | -0.07242 | 0.118324 | -0.00442 | -0.01205 | -0.23453 | -1.73189 | 0.010667 | -0.00385 | -0.14359 | 0.11403 | 0.16208 | 0.05378 | 0.107809 | 0.001708 | -0.06515 | -0.01173 | -0.06368 | -0.36736 | -0.13583 | -0.08641 | -0.11304 | 0.00958 | 0.20054 | 0.02661 | 0.12439 |
| PF382 | 0.101986 | -0.00178 | -0.08205 | 0.06114 | -0.0522 | -0.16418 | -0.17162 | -0.14212 | 0.0267 | 0.15467 | 0.17617 | 0.0246 | 0.037365 | -0.02571 | -0.21664 | -0.18937 | 0.086238 | -0.24605 | -0.25503 | 0.054351 | -0.05454 | -0.03136 | -0.01722 | -0.09308 | 0.15641 |
| NALM6 | 0.157527 | -0.03675 | 0.08413 | 0.05367 | -0.08627 | -0.20539 | -0.22945 | -0.06857 | -0.07989 | 0.10347 | 0.04332 | -0.00792 | -0.01139 | -0.1212 | -0.15818 | 1.97E-05 | 0.183062 | -0.21235 | -0.27497 | -0.08561 | -0.17046 | -0.04244 | -0.15906 | 0.10669 | 0.05225 |
| SKUT1 | -0.12882 | -0.11517 | 0.06755 | -0.04845 | -0.00958 | -0.24737 | -0.00223 | 0.03062 | -0.01463 | 0.19582 | 0.19114 | 0.01584 | -0.09408 | 0.002779 | -0.12051 | -0.00907 | -0.00605 | -0.38972 | -0.37647 | -0.11914 | -0.18349 | 0.08299 | -0.11504 | -0.10947 | 0.24776 |
| HEC1B | 0.068218 | 0.037776 | 0.06814 | -0.05942 | -0.16131 | -0.25098 | 0.095448 | 0.06368 | -0.00647 | 0.16694 | 0.05192 | 0.09394 | 0.030916 | -0.06713 | -0.09539 | -0.07655 | 0.183017 | -0.7718 | -0.58914 | -0.17601 | -0.01684 | 0.04337 | 0.02303 | 0.07357 | 0.04189 |
| RKO | 0.057317 | -0.01372 | 0.06444 | 0.07158 | -0.04422 | -0.06066 | -0.04016 | -0.04213 | -0.14587 | 0.07295 | 0.08542 | -0.01545 | 0.055764 | 0.123844 | -0.00534 | 0.048341 | 0.089303 | -0.20077 | -0.16116 | -0.1009 | 0.066038 | -0.03719 | 0.13506 | -0.02681 | 0.04763 |
| NAMALWA | 0.200923 | 0.116622 | -0.00148 | -0.0482 | -0.02517 | -0.30562 | -0.00935 | -0.15234 | 0.0473 | 0.02769 | 0.14349 | 0.07293 | 0.038206 | 0.039519 | -0.05346 | 0.065998 | -0.00722 | -0.1167 | -0.22996 | 0.095471 | -0.02739 | 0.1014 | 0.06383 | -0.06102 | 0.11552 |
| NCIH650 | -0.35496 | 0.067724 | 0.05167 | -0.28136 | -0.31198 | 0.02651 | -0.04736 | -0.28067 | 0.2461 | 0.12111 | 0.43416 | -0.05888 | -0.09679 | -0.1524 | -0.09287 | -0.26647 | -0.11156 | -0.32382 | -0.17314 | -0.25471 | 0.07579 | -0.04327 | -0.13672 | -0.11466 | 0.05558 |
| HEC265 | -0.08033 | -0.03519 | -0.04464 | -0.02388 | -0.17525 | -0.49654 | -0.05893 | -0.17005 | -0.21105 | 0.06321 | 0.20773 | 0.0411 | 0.0732 | 0.205085 | -0.10046 | 0.065596 | 0.117369 | -0.16444 | -0.33853 | -0.03664 | 0.084481 | -0.01342 | 0.04116 | 0.00348 | 0.13176 |
| OVK18 | 0.037152 | 0.071863 | -0.02758 | -0.08375 | -0.17578 | -0.05846 | -0.01637 | -0.19088 | -0.07211 | 0.11343 | 0.05834 | -0.10187 | 0.0125 | 0.106796 | -0.14635 | 0.058474 | 0.14766 | -0.16889 | -0.12447 | -0.04683 | 0.081961 | -0.11063 | 0.09979 | -0.09782 | 0.11497 |
| 2313287 | 0.041316 | 0.115278 | -0.0393 | 0.03159 | -0.17271 | -0.30021 | -0.00789 | 0.14589 | -0.17237 | 0.07967 | 0.05544 | -0.00683 | 0.045531 | 0.145085 | 0.142073 | 0.053498 | 0.159683 | -0.3686 | -0.15844 | -0.06675 | -0.06995 | -0.05903 | 0.13668 | -0.3033 | 0.1773 |
| LOVO | 0.004547 | -0.16709 | 0.29282 | -0.05517 | -0.20752 | -0.13359 | -0.06661 | 0.1944 | -0.20848 | 0.13687 | 0.10104 | -0.10932 | -0.02435 | 0.119674 | 0.044088 | 0.007744 | 0.300995 | -0.50377 | -0.15981 | 0.032284 | 0.006028 | -0.13735 | 0.00556 | 0.05275 | 0.14638 |
| SUPT1 | 0.141483 | -0.02503 | -0.15242 | 0.03791 | -0.11846 | -0.09275 | -0.3144 | -0.09867 | 0.07576 | -0.00164 | 0.21996 | 0.09576 | 0.058986 | 0.002578 | -0.20344 | -0.04934 | 0.004413 | -0.34562 | -0.30373 | 0.035965 | -0.12886 | -0.04854 | -0.09785 | -0.09673 | 0.30172 |
| 22RV1 | -0.17727 | 0.048619 | -0.16605 | 0.01049 | -0.07898 | 0.07952 | 0.07116 | -0.24982 | -0.1409 | -0.10252 | 0.00663 | -0.08683 | 0.252922 | 0.185957 | 0.132625 | 0.132878 | 0.163436 | -0.09692 | -0.11929 | -0.37582 | 0.226623 | -0.07806 | 0.41805 | 0.09628 | 0.12892 |
| LS180 | -0.0274 | 0.070234 | -0.00932 | -0.00899 | -0.12616 | -0.54242 | 0.08242 | 0.0451 | -0.19446 | 0.17278 | 0.12572 | 0.07823 | 0.109005 | 0.072868 | -0.05655 | 0.08694 | 0.133014 | -0.37921 | -0.091 | 0.038087 | -0.02085 | -0.0724 | 0.00103 | -0.12249 | 0.06172 |
| SW48 | -0.04737 | 0.117 | -0.00749 | 0.05013 | -0.0913 | -0.72334 | 0.078657 | 0.026 | -0.21661 | 0.06148 | 0.01893 | 0.08719 | -0.01323 | 0.029752 | -0.02066 | 0.121871 | 0.218586 | -0.09688 | -0.03563 | -0.01523 | -0.08908 | 0.02183 | 0.02698 | 0.03545 | 0.09939 |
| SNUC4 | -0.10045 | 0.138417 | -0.03076 | 0.06735 | -0.0826 | -0.92834 | 0.066971 | -0.05176 | -0.04456 | 0.03535 | 0.12504 | 0.11566 | 0.07473 | 0.307632 | 0.064989 | 0.091885 | 0.025549 | -0.67945 | -0.21947 | -0.17238 | -0.0751 | 0.01917 | 0.31451 | 0.1418 | 0.21278 |
| REH | 0.11437 | -0.0449 | -0.03989 | 0.09781 | 0.02896 | -0.09518 | -0.12527 | -0.11768 | 0.0879 | 0.0671 | 0.10928 | 0.15185 | -0.04469 | -0.04198 | -0.15661 | -0.01408 | -0.00017 | -0.14668 | -0.16822 | -0.0747 | 0.001395 | -0.08661 | -0.14541 | 0.05564 | 0.09739 |
| ISHIKAWAHERAKLIO0 | -0.16086 | 0.047566 | 0.21889 | 0.22405 | -0.01127 | -0.143 | 0.147388 | 0.03159 | -0.04639 | -0.0951 | 0.175 | -0.1317 | 0.051253 | 0.203571 | 0.190436 | 0.031016 | 0.174703 | -0.54549 | -0.04169 | -0.07088 | -0.11241 | -0.00239 | 0.01853 | 0.0126 | 0.28413 |
| OC314 | -0.05469 | 0.018542 | 0.00506 | -0.21595 | 0.03246 | -0.02285 | 0.042778 | -0.14129 | 0.16843 | -0.01415 | 0.42811 | -0.03136 | -0.15302 | 0.198309 | 0.142542 | -0.04631 | 0.064738 | -0.30436 | -0.84773 | 0.129589 | 0.123178 | 0.0014 | 0.14552 | 0.18585 | 0.06186 |
| CCK81 | -0.23759 | -0.18857 | 0.0688 | 0.33727 | -0.34769 | -0.2839 | -0.11301 | 0.26194 | -0.14486 | 0.30607 | 0.2533 | -0.2159 | 0.256425 | 0.001273 | 0.02037 | 0.068304 | 0.249439 | -0.6738 | -0.03772 | -0.02166 | 0.180285 | 0.04527 | 0.02243 | 0.00757 | 0.22175 |
| RL952 | -0.0245 | 0.013337 | -0.10086 | -0.19849 | 0.08819 | -0.51009 | -0.09828 | 0.04686 | 0.03724 | 0.1044 | 0.04095 | 0.13429 | -0.02953 | 0.082888 | -0.04484 | 0.024506 | 0.130762 | -0.51682 | -0.47733 | -0.1033 | 0.045769 | -0.10213 | 0.0442 | -0.00937 | 0.17945 |
| IGROV1 | 0.091164 | 0.123869 | -0.04712 | -0.12275 | -0.15949 | -0.21936 | -0.03732 | -0.43755 | -0.18534 | -0.0912 | 0.40669 | 0.1334 | -0.01992 | 0.018602 | -0.14934 | -0.05817 | -0.08858 | -0.33916 | -0.17989 | -0.09946 | 0.185484 | -0.18902 | 0.33579 | 0.06561 | -0.20358 |
| COLO792 | -0.04244 | 0.087077 | -0.0305 | -0.04286 | 0.06927 | -0.1628 | -0.07503 | -0.01985 | -0.16059 | -0.00812 | 0.11326 | 0.03882 | 0.136257 | 0.098545 | -0.14567 | 0.075799 | 0.066773 | -0.22881 | -0.31585 | -0.09099 | 0.062636 | -0.06664 | -0.04071 | 0.02789 | 0.18601 |
| KM12 | -0.06072 | 0.07085 | -0.04492 | 0.03911 | -0.02502 | -0.02767 | 0.017865 | -0.13464 | -0.08219 | 0.09507 | 0.10038 | 0.12906 | -0.00998 | 0.121218 | -0.08096 | 0.007441 | 0.079221 | -0.18354 | -0.07927 | -0.04744 | -0.04531 | 0.02902 | 0.05806 | 0.08198 | 0.2405 |
| SNUC5 | 0.011868 | 0.004755 | 0.10581 | -0.1097 | -0.0656 | -0.12916 | 0.031055 | -0.00903 | -0.03984 | 0.16386 | 0.04128 | 0.06774 | 0.07623 | 0.079471 | 0.076346 | 0.068823 | 0.109902 | -0.29142 | -0.07013 | 0.058548 | 0.069563 | -0.01104 | 0.07622 | 0.05601 | -0.01718 |
| HCT116 | 0.1439 | 0.005483 | 0.10888 | 0.03953 | -0.11686 | 0.05347 | 0.128623 | 0.16521 | -0.05977 | 0.11934 | -0.13631 | 0.04326 | 0.098896 | 0.143438 | -0.13589 | 0.049028 | 0.086319 | -0.33943 | 0.00181 | 0.094987 | -0.10515 | -0.26438 | 0.1864 | 0.17507 | 0.15612 |
| 639V | 0.061642 | 0.015813 | -0.01167 | -0.04413 | -0.03701 | -0.32838 | -0.03702 | 0.07489 | 0.0242 | 0.02719 | 0.06023 | 0.00587 | 0.014473 | 0.019691 | -0.06091 | -0.06223 | 0.084565 | -0.2499 | -0.32081 | -0.1188 | -0.66951 | -0.04804 | 0.01402 | 0.02426 | 0.05507 |
| SNGM | -0.02491 | 0.029931 | -0.07513 | -0.00974 | -0.10704 | -0.298 | -0.14754 | 0.09561 | -0.03934 | 0.04601 | 0.11276 | 0.07631 | 0.075171 | 0.112988 | -0.08958 | 0.021163 | 0.112444 | -0.20859 | -0.23599 | -0.0093 | -0.02912 | -0.11893 | -0.09527 | 0.01245 | 0.05959 |
| HCC2450 | 0.047273 | 0.041971 | 0.04926 | -0.10847 | 0.0388 | -0.09656 | -0.00072 | 0.21145 | -0.04723 | 0.17373 | 0.10777 | 0.04687 | 0.041452 | 0.123473 | -0.01569 | -0.02878 | 0.008252 | -0.49348 | -0.47424 | -0.00194 | -0.01282 | -0.04322 | 0.22317 | -0.00782 | 0.25203 |
| HUCCT1 | -0.10144 | -0.23968 | -0.16338 | -0.14022 | -0.1853 | -1.2599 | -0.06759 | 0.1988 | 0.0267 | 0.02589 | 0.20708 | -0.04068 | -0.02729 | 0.126702 | -0.13218 | 0.032448 | 0.096187 | -0.373 | -0.25287 | -0.06158 | -0.1729 | 0.01871 | 0.09557 | -0.04434 | 0.21938 |
| LNCAPCLONEFGC | 0.038972 | 0.124198 | -0.0425 | 0.00106 | -0.11555 | -0.06487 | 0.037855 | 0.01983 | -0.0711 | 0.05367 | 0.04892 | 0.06948 | 0.091375 | 0.212732 | -0.0045 | 0.10733 | 0.088462 | -0.11445 | -0.25491 | -0.00878 | -0.09144 | -0.09457 | 0.0877 | 0.0201 | 0.22116 |
| EN | -0.22092 | 0.254831 | -0.03388 | 0.13294 | -0.11274 | -0.14944 | -0.26599 | -0.17729 | -0.16846 | -0.00882 | -0.00428 | 0.19631 | 0.586872 | 0.010605 | -0.03238 | -0.13298 | 0.187087 | -0.5297 | -0.14142 | -0.40624 | 0.008136 | -0.16805 | -0.08544 | 0.05161 | -0.19647 |
| DU145 | -0.00388 | -0.18097 | 0.11139 | -0.10138 | -0.14052 | -0.88089 | 0.085492 | 0.0446 | -0.06706 | 0.10466 | 0.19844 | -0.04331 | -0.06556 | 0.0959 | -0.17543 | 0.050733 | 0.103214 | -0.18209 | -0.10485 | -0.06274 | -0.08682 | 0.15554 | 0.07917 | 0.03942 | 0.04775 |
| NCIH1155 | -0.08471 | 0.054994 | -0.00053 | -0.05502 | -0.00578 | -0.09257 | 0.00643 | -0.12852 | -0.01812 | 0.02351 | 0.1193 | 0.03536 | 0.080213 | 0.000502 | -0.09817 | 0.058985 | -0.01765 | -0.11107 | -0.09417 | -0.11841 | -0.03104 | -0.06452 | 0.05317 | -0.06565 | 0.23309 |
| KCL22 | 0.060989 | 0.024022 | -0.1013 | -0.00105 | 0.04247 | -0.09926 | 0.06776 | -0.17424 | 0.07128 | 0.02907 | 0.09587 | 0.03686 | 0.057362 | 0.080035 | 0.002278 | -0.11574 | 0.216487 | -0.18427 | -0.24917 | -0.04726 | 0.06624 | -0.13406 | 0.0302 | -0.01304 | -0.0293 |
| HEC6 | -0.16687 | 0.068333 | -0.24259 | -0.12071 | -0.25296 | -0.05831 | 0.065839 | -0.05048 | -0.0777 | 0.15876 | 0.12402 | 0.06593 | -0.10186 | 0.22704 | -0.05222 | 0.180351 | 0.030623 | -0.34564 | -0.23959 | -0.06837 | -0.12437 | 0.08304 | 0.12397 | 0.03615 | 0.1722 |
| LS411N | 0.301674 | 0.0242 | 0.02592 | 0.1114 | -0.12819 | 0.02399 | 0.151499 | -0.11343 | -0.30007 | 0.16928 | 0.0218 | 0.03326 | 0.117761 | 0.212912 | -0.12154 | -0.04529 | 0.140185 | -0.20193 | 0.04549 | -0.03668 | 0.259863 | 0.05949 | -0.13861 | 0.0271 | 0.28678 |
| HT115 | -0.05755 | -0.03226 | -0.0519 | 0.04429 | 0.10144 | -0.75541 | 0.016429 | -0.07482 | -0.09882 | 0.03816 | 0.17423 | 0.13703 | 0.004182 | 0.048466 | 0.122959 | -0.08602 | 0.077397 | -0.80053 | -0.1334 | -0.05807 | -0.0822 | -0.0146 | -0.20607 | 0.12014 | -0.00673 |
| MFE319 | -0.37605 | 0.044939 | 0.0425 | -0.00388 | -0.04662 | -0.50825 | 0.063717 | 0.16541 | 0.11866 | 0.14226 | 0.117 | -0.07159 | -0.07211 | 0.075087 | 0.073497 | 0.061712 | 0.149813 | -0.8318 | -0.04514 | -0.25461 | -0.01381 | 0.00185 | 0.03296 | 0.05866 | 0.00024 |
| SNU81 | -0.03114 | -0.16113 | -0.01937 | 0.08299 | -0.07214 | -0.67853 | 0.144073 | 0.2178 | 0.02464 | 0.0809 | -0.30444 | 0.07122 | 0.033273 | 0.163113 | 0.068391 | 0.153389 | 0.209278 | -0.59199 | 0.00781 | 0.178695 | 0.053072 | 0.27335 | -0.14148 | 0.15664 | 0.13979 |
| JHUEM7 | -0.25846 | -0.02378 | 0.1271 | -0.23879 | -0.05175 | -0.09289 | 0.036996 | 0.13905 | 0.11404 | 0.12665 | 0.11077 | 0.14155 | 0.149782 | 0.143359 | -0.13242 | 0.10394 | 0.006456 | -0.25859 | -0.19389 | -0.24286 | 0.029296 | -0.01605 | 0.09823 | -0.17863 | 0.13495 |
| HEC59 | -0.24914 | 0.12702 | 0.15611 | 0.06458 | -0.08205 | 0.0381 | 0.128714 | 0.01541 | 0.05036 | 0.14505 | 0.04534 | 0.22847 | 0.05006 | 0.104654 | 0.007179 | 0.095527 | 0.219767 | -0.20696 | -0.32076 | -0.16522 | 0.069982 | -0.04905 | 0.1328 | 0.04139 | 0.06379 |
| JURKAT | 0.029933 | 0.03821 | 0.02426 | -0.16427 | -0.05144 | 0.00749 | 0.056925 | -0.13827 | 0.10957 | 0.29289 | 0.05126 | -0.04564 | -0.17873 | -0.13521 | 0.001216 | 0.073429 | -0.06704 | -0.24645 | -0.17273 | 0.017125 | -0.04782 | 0.22165 | -0.13919 | 0.20179 | 0.12126 |
| HEC251 | -0.10862 | 0.049292 | 0.11224 | 0.21907 | -0.09454 | -0.00311 | 0.124228 | -0.32705 | -0.15887 | 0.13385 | 0.21638 | -0.12067 | -0.07472 | 0.326811 | 0.060198 | -0.09971 | -0.01577 | -0.29338 | -0.36946 | -0.06263 | 0.11717 | -0.0961 | -0.04778 | -0.04949 | 0.23749 |
| HCT15 | 0.126938 | -0.00059 | 0.00929 | -0.06794 | -0.13019 | 0.03701 | 0.088352 | -0.07323 | -0.0843 | 0.14164 | 0.14934 | 0.27153 | 0.01589 | 0.086584 | -0.03963 | -0.01702 | 0.113062 | -0.05715 | -0.14516 | 0.018815 | -0.04768 | 0.14284 | 0.03417 | 0.09057 | -0.072 |
| 143B | 0.037258 | 0.043437 | -0.04868 | 0.03039 | 0.00636 | -0.12844 | 0.070387 | -0.10405 | 0.12365 | 0.0304 | 0.18661 | 0.00467 | -0.0251 | 0.119013 | -0.04551 | 0.136668 | 0.121867 | -0.17068 | -0.17669 | -0.11888 | -0.10931 | 0.02384 | 0.07535 | -0.06466 | 0.22729 |
| BECKER | 0.034598 | -0.08556 | -0.0542 | 0.15667 | -0.26258 | -0.05647 | 0.034944 | -0.06993 | 0.08874 | 0.08749 | -0.02296 | 0.03728 | 0.224854 | 0.362422 | 0.150748 | -0.08745 | 0.151706 | -0.1678 | -0.18446 | 0.117355 | -0.34099 | 0.01944 | 0.30664 | -0.04624 | 0.45571 |
| BT16 | -0.00131 | 0.179911 | -0.0112 | 0.02304 | -0.06173 | -0.04309 | -0.54583 | -0.21821 | -0.01828 | 0.19515 | 0.08371 | -0.08175 | 0.106107 | 0.058949 | 0.158627 | -0.23192 | -0.36882 | -0.78223 | -0.25623 | -0.51227 | 0.053371 | 0.2367 | -0.03426 | 0.00025 | 0.13899 |
| CHLA06ATRT | 0.044463 | 0.058508 | -0.0489 | 0.01919 | 0.03219 | -0.21253 | -0.0292 | -0.09345 | 0.07151 | 0.05632 | 0.12617 | 0.10824 | 0.102871 | -0.02028 | 0.009227 | -0.04356 | 0.048982 | -0.21545 | -0.17299 | 0.009394 | -0.06422 | -0.01468 | -0.04471 | -0.09868 | 0.20743 |
| CHLA10 | 0.201883 | 0.060399 | -0.02411 | 0.14322 | -0.00058 | -0.17989 | -0.01981 | -0.14908 | -0.02285 | 0.04897 | 0.03783 | 0.09671 | -0.00779 | -0.00459 | -0.12813 | 0.023848 | -0.05408 | -0.32908 | -0.25777 | 0.008971 | 0.008896 | 0.00475 | 0.03088 | -0.05691 | 0.15764 |
| CHLA266 | 0.09025 | 0.075833 | -0.01737 | -0.01745 | -0.07653 | -0.10386 | 0.018974 | -0.0314 | 0.02548 | -0.07439 | 0.17696 | 0.00201 | 0.032858 | 0.124987 | -0.12029 | -0.17023 | 0.158487 | -0.19558 | -0.22472 | -0.1989 | 0.022703 | -0.66442 | 0.03186 | -0.00775 | 0.24397 |
| CHLA57 | -0.14347 | 0.305735 | 0.00325 | 0.23352 | -0.34535 | -0.10364 | 0.300478 | -0.01703 | -0.11257 | 0.25269 | 0.01351 | -0.02177 | 0.133273 | 0.012832 | 0.280481 | 0.27865 | 0.048484 | -0.28713 | 0.01697 | -0.368 | 0.055726 | -0.20081 | -0.08938 | 0.0218 | -0.2163 |
| CMK115 | 0.064215 | 0.103422 | 0.01066 | 0.10539 | -0.2831 | -0.03942 | -0.24596 | -0.17456 | 0.09652 | 0.02127 | 0.03181 | 0.1253 | -0.0613 | -0.03955 | -0.16736 | 0.170917 | 0.162805 | -0.33283 | -0.1532 | 0.180337 | 0.082602 | 0.21319 | 0.02993 | 0.15128 | 0.00287 |
| COGE352 | 0.138655 | 0.015209 | -0.05715 | -0.04438 | -0.00792 | -0.16936 | 0.09827 | 0.01639 | -0.02332 | 0.09976 | 0.01177 | 0.07216 | -0.02853 | 0.054599 | 0.009649 | 0.070099 | -0.08541 | -0.22902 | -0.18507 | -0.0802 | -0.03022 | -0.10871 | -0.05028 | 0.08377 | -0.04475 |
| COLO205 | 0.116142 | -0.09348 | -0.01221 | 0.03036 | 0.05913 | 0.17366 | -0.08617 | 0.14799 | -0.22489 | 0.17586 | 0.18051 | 0.02615 | 0.194286 | 0.214943 | -0.08384 | -0.16442 | 0.091437 | -0.18559 | 0.04426 | 0.002073 | 0.008799 | -0.26266 | 0.13727 | -0.08288 | 0.12751 |
| CHL1DM | -0.02406 | -0.01155 | 0.10191 | -0.04198 | 0.00748 | -0.2763 | -0.18311 | -0.11294 | -0.05642 | 0.15496 | 0.12286 | 0.00171 | 0.023448 | -0.01328 | -0.02424 | -0.07461 | -0.09916 | -0.5558 | -0.49264 | -0.06451 | -0.29887 | -0.39181 | -0.31967 | -0.00074 | 0.24236 |

| Cell line | ALK | AXL | CSF1R | DDR1 | DDR2 | EGFR | EPHA1 | EPHA2 | EPHA3 | EPHA4 | EPHA6 | EPHA7 | EPHB1 | EPHB2 | EPHB3 | EPHB4 | EPHB6 | ERBB2 | ERBB3 | ERBB4 | FGFR1 | FGFR2 | FGFR3 | FGFR4 | FLT1 |
|---|---|---|---|---|---|---|---|---|---|---|---|---|---|---|---|---|---|---|---|---|---|---|---|---|---|
| COV504 | 0.037301 | -0.08244 | 0.0173 | 0.13744 | -0.18157 | -0.33623 | -0.01296 | 0.38147 | -0.0929 | 0.15511 | 0.165 | -0.15844 | 0.042201 | 0.227842 | -0.30453 | 0.073072 | -0.04065 | -0.39574 | -0.2063 | 0.087416 | -0.50714 | -0.01527 | 0.1056 | -0.09126 | 0.15921 |
| CW9019 | -0.03138 | -0.32195 | 0.0189 | -0.12238 | 0.04272 | 0.00121 | 0.061341 | 0.05264 | 0.10611 | 0.11638 | 0.11556 | 0.09438 | 0.059032 | 0.212205 | -0.01913 | 0.032169 | 0.056345 | -0.43873 | -0.3049 | -0.21514 | -0.1204 | -0.1452 | -0.06058 | -0.25171 | 0.17592 |
| D425 | -0.14536 | 0.415835 | 0.09443 | -0.03296 | -0.09716 | -0.00998 | -0.21221 | 0.00774 | -0.1067 | 0.02813 | 0.16983 | -0.3311 | -0.20504 | 0.123878 | 0.124277 | -0.07476 | 0.204287 | -0.42296 | -0.2333 | 0.000606 | 0.024575 | 0.0041 | -0.04527 | -0.16509 | -0.02065 |
| D458 | 0.04972 | -0.07878 | -0.04783 | 0.00436 | -0.32268 | 0.07696 | -0.24736 | -0.06829 | -0.0041 | 0.27206 | 0.11786 | -0.11262 | 0.031079 | 0.115654 | -0.05833 | 0.117116 | 0.044512 | -0.19191 | -0.1775 | 0.086644 | 0.080636 | -0.04372 | 0.07918 | 0.04482 | 0.2689 |
| DLD1 | -0.04969 | 0.061972 | 0.0679 | 0.02231 | -0.15668 | -0.38184 | -0.06044 | 0.02329 | -0.09976 | 0.03614 | 0.1657 | -0.02133 | 0.033647 | 0.117686 | 0.027385 | -0.08193 | -0.03724 | -0.46008 | -0.26267 | 0.077972 | 0.002827 | 0.03368 | 0.11077 | -0.01729 | 0.20408 |
| DOV13 | -0.04175 | -0.02577 | -0.07928 | -0.02861 | 0.03888 | -0.27019 | -0.02518 | 0.44867 | -0.07112 | 0.04117 | 0.30906 | -0.02951 | 0.029611 | -0.01452 | -0.01312 | 0.067411 | -0.02081 | -0.38677 | -0.3139 | -0.29293 | 0.095508 | -0.02346 | 0.0191 | 0.02812 | 0.06211 |
| EVSAT | -0.08296 | -0.05653 | 0.03762 | 0.00088 | -0.00178 | -0.0246 | 0.151259 | 0.0638 | -0.24277 | -0.00494 | 0.10966 | 0.0326 | 0.109542 | 0.041334 | 0.007294 | 0.104133 | 0.101987 | -0.4244 | -0.26291 | -0.02031 | -0.32321 | -0.39457 | 0.0376 | -0.33657 | -0.06816 |
| F5 | 0.043619 | 0.171885 | 0.01913 | -0.06054 | 0.04155 | -0.08106 | -0.0161 | -0.1379 | 0.05291 | 0.36908 | 0.14086 | -0.0044 | 0.031746 | 0.200409 | 0.148505 | 0.143545 | 0.118098 | -0.31096 | -0.21959 | 0.051754 | 0.078062 | 0.04684 | -0.00354 | -0.04016 | 0.40374 |
| NCIH292 | 0.023875 | 0.110418 | -0.06648 | -0.02377 | -0.10947 | -0.33877 | 0.006994 | 0.12053 | -0.0242 | 0.08622 | 0.11496 | -0.08689 | 0.01985 | 0.028232 | 0.025356 | 0.111647 | 0.284275 | -0.438 | -0.2193 | -0.07799 | -0.15861 | 0.07034 | 0.13233 | -0.01733 | 0.13468 |
| HCC2998 | 0.095807 | -0.12382 | 0.05312 | 0.04271 | 0.01821 | 0.1225 | 0.101248 | -0.0505 | 0.11762 | -0.04572 | 0.02245 | 0.07853 | 0.074446 | 0.179863 | -0.14427 | -0.01701 | 0.103455 | -0.38886 | -0.11284 | 0.179628 | -0.09646 | 0.15152 | -0.05896 | 0.05134 | 0.21387 |
| JR | 0.007849 | -0.01439 | 0.03168 | -0.10696 | -0.18532 | -0.09657 | 0.095645 | 0.04952 | -0.06218 | 0.05868 | 0.18797 | 0.06697 | 0.00284 | 0.050539 | -0.12227 | 0.055174 | 0.175142 | -0.32709 | -0.16248 | -0.25506 | -0.01254 | -0.13114 | -0.02949 | -0.68059 | 0.08133 |
| KCIMOH1 | -0.00936 | -0.13674 | -0.11188 | 0.01987 | -0.12746 | -0.67232 | -0.0353 | -0.04106 | 0.26513 | -0.10253 | 0.04697 | 0.01692 | 0.015162 | 0.016927 | -0.02835 | 0.042823 | 0.108274 | -0.25936 | -0.30057 | 0.037081 | -0.01262 | -0.0033 | -0.0084 | -0.16417 | 0.09409 |
| KD | 0.234865 | 0.018656 | 0.13396 | 0.16345 | -0.13031 | -0.04979 | 0.062406 | 0.03228 | -0.0295 | 0.08962 | 0.13927 | 0.03013 | 0.104676 | 0.136455 | -0.01153 | 0.116147 | 0.224453 | -0.14368 | -0.18936 | -0.07172 | -0.6971 | 0.20664 | 0.16092 | -0.07724 | 0.074 |
| KP1N | 0.142359 | -0.02658 | -0.03087 | -0.07968 | 0.03857 | -0.01051 | 0.078811 | 0.07347 | -0.39221 | 0.071 | -0.09128 | 0.1141 | -0.08212 | 0.054401 | 0.000999 | 0.093726 | 0.144788 | -0.37498 | -0.2309 | -0.01869 | -0.15502 | 0.11314 | -0.02231 | -0.03445 | 0.15819 |
| MAC2A | -0.0087 | -0.23702 | -0.07215 | 0.1857 | -0.1171 | -0.36029 | -0.29707 | 0.04233 | 0.01576 | 0.18589 | 0.06452 | 0.0306 | -0.10283 | 0.189544 | -0.2222 | 0.049625 | 0.105882 | -0.13708 | -0.30293 | -0.09236 | -0.0202 | -0.00745 | 0.14307 | -0.26711 | 0.12043 |
| MOGGUVW | -0.00033 | 0.012019 | 0.08821 | 0.26169 | -0.00669 | -0.01612 | 0.01696 | 0.07162 | -0.02053 | 0.08314 | -0.01762 | 0.0341 | 0.030532 | 0.249783 | 0.100047 | 0.083167 | 0.199368 | -0.09267 | 0.0368 | -0.02916 | -0.32883 | -0.16577 | 0.36307 | 0.13106 | 0.35916 |
| MON | -0.11247 | 0.191346 | 0.1404 | 0.27073 | 0.21193 | -0.10149 | 0.039561 | 0.0791 | -0.15559 | 0.12585 | 0.23263 | 0.23122 | 0.044875 | 0.380938 | 0.062956 | 0.184259 | 0.170725 | -0.29637 | -0.32988 | 0.21563 | 0.098322 | 0.01137 | 0.05351 | 0.09851 | 0.01984 |
| MONOMAC1 | 0.0824 | -0.02729 | 0.08114 | 0.06774 | -0.01699 | 0.05075 | -0.04174 | -0.28663 | 0.07345 | 0.01538 | -0.00085 | 0.20463 | 0.003068 | -0.04131 | 0.22341 | 0.00349 | 0.12985 | -0.08474 | -0.07566 | -0.22559 | -0.0137 | -0.18856 | 0.01127 | 0.19621 | -0.15713 |
| MYLA | 0.04158 | 0.049585 | 0.11634 | 0.17103 | -0.12522 | -0.20552 | -0.00617 | -0.12142 | 0.06002 | 0.03713 | 0.15065 | -0.00303 | 0.193733 | 0.204607 | -0.30222 | -0.00448 | -0.02092 | -0.16806 | -0.2891 | 0.049448 | -0.09175 | -0.10792 | -0.01655 | -0.03706 | 0.28265 |
| NCIH1993 | -0.0438 | 0.093782 | 0.25953 | 0.1402 | -0.217 | -0.23614 | 0.080161 | -0.00728 | -0.04733 | -0.08358 | 0.41275 | -0.04334 | 0.207053 | 0.20812 | -0.37421 | -0.04684 | -0.08838 | -0.47514 | -0.12608 | -0.38762 | -0.01748 | -0.11679 | -0.09116 | -0.05995 | 0.18708 |
| OC316 | 0.022853 | 0.022561 | 0.00697 | 0.01727 | -0.18526 | -0.33365 | 0.145033 | -0.01333 | -0.08345 | 0.11583 | 0.16234 | 0.01771 | 0.160379 | 0.155432 | -0.06767 | 0.002531 | 0.0275 | -0.96239 | -0.76311 | -0.11657 | 0.037334 | 0.00376 | -0.07087 | 0.04603 | 0.09597 |
| OVCAR5 | -0.08048 | -0.03713 | 0.02897 | -0.03052 | -0.09046 | -0.54343 | 0.021355 | 0.2577 | -0.10533 | 0.06986 | 0.0135 | -0.02593 | -0.00438 | 0.032858 | -0.08732 | -0.06477 | 0.05341 | -0.32429 | -0.22108 | -0.03305 | -0.04901 | -0.04814 | 0.03039 | 0.05857 | 0.14908 |
| CCLFPEDS0001T | 0.07168 | -0.08131 | -0.03324 | -0.07596 | -0.25095 | -0.10737 | -0.05557 | 0.00172 | -0.03354 | 0.21145 | 0.07303 | -0.14419 | 0.074329 | 0.027487 | -0.20692 | 0.044179 | -0.04153 | -0.29567 | -0.40643 | -0.10625 | 0.113534 | -0.0291 | 0.07361 | -0.15185 | 0.1073 |
| CCLFPEDS0003T | 0.020765 | 0.214075 | 0.1039 | -0.12937 | 0.07869 | -0.05693 | -0.03837 | -0.04508 | 0.14361 | 0.29783 | 0.22031 | -0.22009 | 0.214866 | -0.10666 | 0.061119 | 0.014689 | 0.206244 | -0.09125 | -0.208 | -0.2702 | 0.051564 | 0.28141 | 0.19378 | -0.01845 | -0.24513 |
| U251MGDM | -0.12143 | -0.0558 | -0.02528 | -0.09964 | -0.10411 | -0.10578 | -0.05772 | 0.16755 | -0.03151 | 0.16469 | 0.23822 | -0.03194 | 0.104098 | 0.086988 | -0.07513 | -0.00084 | 0.037824 | -0.11948 | -0.19091 | -0.0632 | -0.07054 | -0.03448 | 0.00468 | -0.01356 | 0.05532 |
| RT11284 | -0.19572 | 0.016414 | 0.00549 | 0.04806 | 0.01457 | -0.30858 | 0.122227 | 0.30655 | 0.02017 | 0.05879 | 0.13998 | 0.05486 | 0.07383 | 0.028014 | -0.25782 | 0.025811 | 0.02518 | -0.48488 | -0.2409 | -0.17418 | -0.06973 | 0.1798 | -0.79887 | -0.00773 | 0.09078 |
| SCMCRM2 | -0.26681 | 0.118584 | 0.13813 | 0.02311 | -0.19712 | 0.0507 | 0.070767 | 0.04402 | -0.16339 | -0.01931 | 0.21814 | -0.10462 | -0.07156 | 0.155687 | 0.075249 | 0.297904 | 0.215065 | -0.05396 | -0.21043 | -0.17714 | 0.147208 | 0.06914 | 0.0661 | 0.24072 | 0.1616 |
| SHSY5Y | -1.59805 | 0.117946 | -0.07754 | 0.14837 | -0.2243 | -0.05884 | -0.04162 | -0.09153 | 0.05381 | 0.09625 | 0.15449 | 0.04137 | -0.10949 | -2.9E-05 | -0.10637 | 0.200711 | 0.108695 | -0.35548 | -0.15195 | -0.0411 | 0.185384 | 0.07927 | 0.14199 | 0.00109 | 0.33042 |
| SKMEL2 | 0.089147 | 0.076682 | -0.0177 | -0.00438 | -0.10181 | -0.04613 | 0.009233 | 0.09383 | 0.05385 | 0.1406 | -0.0357 | 0.06632 | -0.05632 | 0.096724 | 0.015484 | 0.121886 | 0.112849 | -0.22217 | 0.00775 | -0.12097 | -0.02084 | -0.01363 | 0.16217 | 0.07431 | 0.10026 |
| SKNEP1 | -0.09871 | -0.06565 | 0.13179 | -0.07339 | -0.16084 | -0.08325 | 0.141252 | -0.15725 | -0.02546 | 0.1835 | 0.11979 | 0.19998 | -0.04738 | -0.05329 | -0.10882 | 0.126121 | 0.455398 | -0.56017 | -0.14671 | -0.18625 | 0.018028 | -0.05444 | -0.12532 | 0.20937 | 0.01737 |
| SKPNDW | -0.18614 | 0.04693 | 0.19332 | -0.05022 | -0.14203 | 0.11953 | -0.02228 | 0.01765 | -0.02732 | 0.13653 | -0.13996 | 0.09237 | -0.07089 | 0.014435 | 0.018834 | 0.037963 | 0.018482 | -0.2561 | 0.04044 | -0.34134 | -0.05418 | 0.08876 | 0.10127 | 0.08748 | 0.1147 |
| SKRC31 | 0.087037 | 0.107311 | -0.01493 | -0.02099 | -0.08521 | -0.25857 | 0.048777 | 0.05158 | -0.21614 | 0.01488 | 0.2337 | 0.19479 | 0.007779 | 0.074786 | -0.22447 | 0.037005 | 0.059704 | -0.19059 | -0.23103 | -0.05125 | -0.06559 | 0.11359 | -0.12051 | 0.01024 | 0.14746 |
| SMSCTR | -0.19959 | 0.229926 | 0.18852 | 0.01221 | 0.13306 | -0.02195 | 0.133155 | -0.19406 | -0.2168 | 0.04121 | 0.24712 | 0.07349 | 0.015671 | 0.145899 | -0.04077 | 0.035876 | 0.239781 | -0.23035 | -0.28006 | -0.18615 | 0.080698 | 0.06424 | 0.01127 | 0.1844 | -0.16033 |
| SMZ1 | -0.0257 | 0.02248 | -0.17911 | -0.05524 | -0.09025 | -0.11783 | 0.007302 | -0.28422 | -0.01995 | 0.19491 | 0.03528 | 0.07237 | 0.201331 | -0.03689 | -0.13612 | -0.07011 | 0.103025 | -0.08188 | -0.21668 | -0.08255 | 0.011119 | -0.02449 | 0.01363 | -0.12734 | 0.31697 |
| TC32 | -0.07948 | -0.02916 | 0.0069 | 0.14853 | -0.19672 | 0.20429 | 0.206019 | -0.29401 | 0.05391 | -0.00889 | 0.21139 | 0.03385 | 0.211393 | 0.172221 | 0.049021 | 0.040331 | 0.33302 | -0.13926 | -0.04389 | -0.29498 | -0.00845 | -0.05513 | -0.14707 | 0.14702 | 0.10614 |
| TTC549 | 0.015068 | -0.01353 | 0.07184 | -0.00857 | -0.2408 | -0.03382 | -0.26049 | -0.07283 | -0.07031 | 0.02195 | 0.28712 | -0.04477 | 0.038231 | 0.016987 | -0.06277 | -0.08041 | 0.185348 | -0.10283 | -0.1146 | -0.14892 | -0.05368 | 0.00846 | -0.13561 | -0.0062 | -0.06832 |
| TTC642 | -0.31423 | -0.11867 | 0.14252 | 0.05799 | 0.10648 | -0.17378 | 0.270995 | 0.00543 | 0.14858 | -0.12224 | 0.22134 | -0.03323 | 0.066453 | 0.259154 | 0.135569 | 0.052472 | 0.015957 | 0.03887 | -0.01408 | -0.00505 | -0.23571 | 0.2935 | 0.34232 | 0.03407 | -0.04371 |
| UPCISCC152 | -0.13954 | 0.017668 | -0.09949 | 0.04585 | 0.18746 | -0.29251 | -0.04158 | -0.19251 | -0.06444 | 0.07986 | 0.11315 | 0.09985 | 0.029316 | -0.05792 | -0.13877 | 0.11077 | 0.081236 | -0.98562 | -0.28668 | -0.15497 | -0.06162 | -0.05625 | 0.0054 | -0.06525 | -0.07147 |
| UPCISCC154 | 0.052849 | -0.00576 | -0.17263 | 0.02071 | -0.04829 | -0.63063 | -0.07584 | 0.06554 | -0.09361 | 0.06647 | 0.22603 | 0.03043 | 0.075482 | 0.113101 | -0.09664 | 0.020768 | 0.083216 | -0.73732 | -0.71757 | -0.0441 | -0.10506 | -0.07079 | 0.04866 | -0.08948 | 0.18295 |
| UW228 | 0.080163 | -0.30755 | -0.05844 | -0.0074 | -0.0487 | -0.0644 | 0.054878 | 0.40351 | 0.05702 | 0.06209 | 0.19423 | 0.01693 | 0.043132 | 0.002026 | -0.01694 | -0.02295 | 0.118898 | -0.07418 | -0.2408 | -0.04688 | -0.25416 | -0.39741 | -0.09035 | -0.03292 | 0.15564 |
| VMRCLCD | 0.066632 | 0.029717 | 0.01551 | 0.10199 | -0.14485 | 0.11217 | -0.03185 | -0.23657 | 0.01193 | 0.08334 | 0.00974 | -0.02266 | 0.05139 | 0.054181 | -0.1073 | -0.02934 | -0.07807 | -0.27523 | -0.13394 | -0.14507 | 0.01778 | -0.01774 | -0.06842 | -0.10894 | 0.07573 |
| WM2664 | -0.03669 | -0.22676 | 0.02542 | -0.03293 | 0.00074 | -0.0298 | 0.000747 | -0.02158 | 0.08172 | 0.10993 | 0.14901 | 0.06548 | 0.034249 | 0.084281 | -0.10568 | 0.134661 | 0.138441 | -0.1994 | -0.14197 | -0.08824 | -0.06915 | 0.01496 | -0.05507 | 0.02121 | 0.00042 |
| 127399 | -0.13651 | -0.06133 | -0.0008 | -0.05361 | 0.21135 | -0.1614 | -0.12432 | 0.02033 | 0.17453 | 0.1808 | -0.07068 | 0.25839 | 0.26394 | 0.031531 | -0.28051 | -0.0754 | 0.177927 | -0.40376 | -0.10232 | -0.49765 | 0.142896 | -0.16813 | 0.23063 | 0.17043 | -0.15732 |
| SW982 | -0.09368 | 0.04196 | 0.06687 | 0.11124 | -0.13185 | -0.19338 | -0.23357 | 0.11727 | 0.11299 | 0.10239 | 0.24414 | 0.16535 | 0.064216 | -0.0265 | 0.047305 | 0.207032 | 0.017425 | -0.21141 | -0.12272 | -0.14096 | 0.01119 | -0.00431 | -0.19294 | 0.02831 | 0.16918 |
| SYO1 | -0.06305 | 0.136275 | -0.04345 | -0.1459 | -0.02313 | -0.06681 | -0.01262 | 0.28601 | 0.01615 | -0.03777 | 0.06492 | -0.10209 | 0.037165 | 0.013835 | -0.06727 | -0.08926 | 0.156369 | -0.24827 | -0.01207 | -0.07751 | -0.0563 | -0.16409 | -0.19689 | 0.17481 | -0.05023 |
| YAMATO | 0.086547 | -0.03772 | 0.08495 | 0.01834 | 0.03831 | 0.12224 | 0.015459 | 0.04674 | 0.11296 | 0.14452 | 0.01339 | 0.03023 | 0.166834 | -0.01056 | 0.164088 | 0.046326 | 0.078126 | -0.19413 | -0.17957 | -0.1069 | 0.11849 | -0.00626 | 0.00474 | 0.03496 | 0.1873 |
| BIN67 | -0.11921 | 0.057558 | -0.0012 | 0.04118 | 0.06407 | -0.11395 | 0.035316 | -0.24291 | -0.03304 | 0.10064 | 0.08249 | 0.31797 | 0.04295 | 0.051654 | 0.106213 | 0.06684 | 0.162707 | -0.75809 | -0.32272 | -0.70025 | -0.56631 | 0.05396 | 0.0912 | -0.06572 | -0.01252 |
| SCCOHT1 | 0.081385 | 0.010613 | 0.03993 | -0.01909 | -0.13593 | -0.08571 | -0.03456 | 0.00276 | -0.12011 | 0.23265 | 0.05633 | -0.18244 | 0.080413 | 0.15885 | 0.059338 | 0.174052 | -0.15067 | -0.13756 | -0.25234 | 0.117759 | -0.3308 | -0.60781 | 0.24251 | -0.11174 | 0.31331 |
| SCS214 | -0.10735 | 0.126626 | 0.33847 | 0.12698 | -0.0724 | -0.02001 | 0.246566 | -0.00556 | 0.00895 | 0.13067 | -0.0154 | 0.01046 | 0.063367 | 0.018556 | -0.03056 | 0.222468 | 0.187987 | -0.37197 | -0.04749 | -0.34672 | 0.095621 | -0.11564 | 0.10573 | 0.15619 | 0.06733 |
| TC106 | 0.019053 | -0.25468 | -0.13047 | 0.0822 | 0.07769 | -0.04179 | -0.04316 | -0.04567 | 0.04338 | 0.09273 | 0.21856 | 0.07396 | 0.00781 | -0.01696 | -0.02931 | -0.05073 | 0.114481 | -0.32391 | -0.13469 | -0.25944 | -0.17082 | -0.17881 | 0.01127 | 0.21319 | 0.22574 |
| COGAR359 | -0.20267 | 0.027919 | -0.29752 | -0.13585 | -0.01453 | -0.13752 | 0.056295 | -0.01125 | 0.05147 | 0.17455 | 0.09454 | 0.18796 | -0.07681 | 0.230277 | 0.097747 | -0.07708 | 0.133832 | -0.05469 | -0.19671 | -0.2461 | -0.04479 | -0.19542 | 0.19532 | 0.03782 | 0.16558 |
| Y79 | -0.06885 | 0.101537 | -0.07555 | -0.06336 | -0.0289 | -0.09031 | -0.06469 | -0.12027 | -0.0309 | 0.11525 | 0.18472 | -0.02391 | 0.03976 | 0.176079 | -0.0892 | 0.131062 | -0.01356 | -0.07013 | -0.17385 | -0.06679 | 0.061694 | -0.12222 | 0.07848 | -0.03318 | 0.10942 |
| CHLA15 | -1.29138 | 0.117665 | 0.02772 | 0.03601 | -0.06893 | -0.04586 | 0.013453 | -0.14551 | 0.13401 | 0.03969 | 0.24592 | 0.03348 | -0.03555 | 0.018415 | -0.15075 | 0.123874 | 0.116766 | -0.28899 | -0.12939 | -0.16797 | 0.110068 | -0.02342 | -0.13375 | -0.03621 | 0.0008 |
| COGN278 | 0.033912 | 0.252203 | -0.08575 | -0.10769 | -0.1146 | 0.00803 | -0.08676 | -0.0782 | 0.02544 | 0.10908 | 0.16215 | 0.18632 | -0.12979 | 0.187111 | -0.22119 | -0.04798 | 0.21346 | -0.13507 | -0.15407 | -0.15863 | 0.066329 | -0.12406 | 0.14659 | 0.16016 | 0.32075 |
| COGN305 | -0.22394 | 0.080318 | -0.05578 | -0.12948 | -0.09924 | -0.19065 | 0.009509 | -0.16879 | 0.14721 | 0.26742 | 0.23275 | -0.03585 | -0.0135 | -0.14204 | 0.065605 | -0.01428 | 0.071828 | -0.32954 | -0.09872 | -0.0836 | -0.14382 | -0.17156 | -0.33276 | 0.01975 | 0.43357 |
| NB1643 | -0.66435 | -0.08016 | -0.04246 | -0.02519 | -0.21164 | 0.14609 | -0.10696 | 0.05282 | 0.16954 | 0.09768 | 0.05285 | 0.0485 | 0.007008 | 0.166201 | -0.11541 | -0.05566 | 0.103652 | -0.26259 | 0.05253 | -0.05374 | -0.12161 | -0.11337 | -0.02425 | -0.0814 | 0.23533 |
| 8305C | -0.16331 | -0.09064 | 0.03363 | 0.10725 | 0.04258 | 0.03014 | -0.01148 | -0.0228 | -0.03328 | 0.17084 | 0.19681 | 0.06551 | 0.056713 | 0.066435 | -0.14502 | -0.07458 | 0.128409 | -0.23924 | -0.22055 | -0.00802 | -0.10414 | -0.04688 | 0.12194 | 0.04281 | 0.14478 |
| 8505C | 0.043564 | 0.11486 | 0.02218 | -0.0766 | -0.01494 | -0.14956 | -0.01208 | 0.16807 | -0.30556 | 0.08051 | 0.1191 | 0.06214 | 0.02095 | 0.109258 | -0.25026 | 0.025573 | 0.082211 | -0.14965 | -0.33452 | -0.03179 | 0.027335 | -0.021 | -0.14302 | 0.03915 | 0.3006 |
| HA1E | 0.089176 | -0.22577 | -0.08931 | -0.00554 | 0.06293 | -0.12507 | 0.014592 | -0.00522 | 0.0346 | -0.08982 | 0.18041 | 0.10543 | -0.208 | 0.230786 | -0.14003 | -0.04643 | 0.077308 | -0.62046 | -0.25696 | 0.037989 | -0.98051 | -0.06122 | 0.11683 | -0.06097 | 0.05858 |
| PLCPRF5 | 0.014696 | 0.009423 | -0.08947 | -0.0338 | -0.11228 | -0.85625 | -0.18742 | 0.10792 | 0.04283 | 0.06493 | 0.09923 | 0.06528 | -0.06406 | 0.071264 | -0.05939 | 0.068609 | -0.0222 | -0.57323 | -0.311 | -0.11941 | -0.11793 | -0.12484 | 0.09375 | 0.14769 | 0.21707 |
| CME1 | 0.112942 | -0.15644 | -0.13296 | -0.07219 | -0.06257 | -0.19205 | -0.04319 | 0.0416 | -0.03373 | 0.06798 | 0.28385 | 0.05143 | -0.00033 | 0.034989 | 0.121497 | -0.06789 | 0.214719 | -0.43148 | -0.16224 | -0.16897 | -0.04842 | -0.11204 | -0.04702 | -0.00088 | 0.17214 |
| A431 | 0.022419 | -0.01753 | -0.01847 | -0.19319 | 0.03166 | -0.88509 | 0.040338 | 0.3486 | 0.01453 | 0.18536 | 0.16343 | 0.12173 | -0.00071 | 0.181479 | -0.15766 | 0.01331 | 0.073472 | -0.96257 | -1.06746 | -0.11355 | -0.04689 | -0.1042 | -0.01967 | -0.01593 | 0.0968 |
| ANGMCSS | 0.025241 | -0.00996 | -0.06853 | 0.02422 | -0.0468 | -0.2533 | 0.100679 | 0.31563 | 0.1012 | 0.02728 | 0.07458 | 0.08122 | 0.003466 | 0.107065 | -0.02172 | 0.002771 | 0.174769 | -0.10338 | -0.35843 | -0.10736 | -0.23607 | -0.04768 | -0.00406 | -0.07576 | 0.12516 |
| BICR10 | -0.11192 | 0.162481 | 0.15159 | 0.08859 | -0.13516 | -0.0002 | 0.156012 | 0.12169 | -0.06023 | 0.06268 | 0.07206 | -0.06241 | 0.019 | 0.048745 | 0.062033 | -0.04914 | -0.01078 | -0.17506 | -0.09931 | -0.20563 | -0.71867 | 0.04342 | 0.06561 | 0.12239 | 0.11186 |
| BICR78 | 0.225922 | -0.01163 | 0.00466 | 0.01044 | -0.04428 | -0.84835 | 0.17824 | 0.09861 | -0.36687 | 0.15273 | 0.15805 | 0.19165 | 0.005038 | 0.167662 | 0.205668 | -0.17243 | 0.552664 | -0.43413 | -0.46132 | 0.070033 | -0.08405 | -0.04618 | 0.19596 | 0.55156 | 0.10856 |
| C33A | -0.37771 | -0.07171 | 0.03579 | 0.05832 | -0.06123 | -0.14355 | 0.08199 | 0.01174 | -0.10822 | 0.19384 | 0.10694 | 0.04741 | 0.053278 | 0.162191 | 0.061677 | 0.154307 | 0.152115 | -0.8828 | -0.23212 | -0.48979 | -0.06215 | -0.04171 | -0.05176 | 0.22293 | 0.11407 |
| C4I | 0.143106 | 0.122768 | -0.01956 | 0.06654 | 0.01178 | -0.80811 | -0.01442 | -0.14736 | -0.038 | 0.05375 | 0.0463 | 0.01931 | 0.044226 | 0.081644 | -0.10156 | -0.08237 | 0.10371 | -0.18241 | -0.20568 | -0.08842 | -0.01263 | -0.04592 | 0.09294 | 0.00607 | 0.09997 |
| C4II | 0.152803 | 0.179582 | -0.08788 | -0.16154 | 0.18727 | -0.78335 | -0.09029 | -0.08531 | 0.14205 | 0.25216 | 0.12315 | 0.03331 | -0.06638 | 0.076779 | -0.02464 | 0.026813 | 0.045282 | -0.49828 | -0.10247 | -0.15758 | 0.054635 | 0.14922 | 0.0911 | -0.03229 | 0.08001 |
| CASKI | 0.086119 | -0.06372 | -0.05818 | 0.02181 | -0.14322 | -0.41815 | 0.009671 | 0.01001 | -0.06918 | 0.2453 | 0.00204 | 0.02867 | 0.118513 | 0.097784 | -0.02906 | -0.08246 | 0.187452 | -0.37122 | -0.20194 | -0.06296 | 0.074645 | -0.08822 | 0.00899 | 0.0791 | 0.16891 |
| CHP134 | -0.14372 | 0.029963 | -0.18903 | -0.03708 | 0.06182 | -0.16802 | -0.58698 | -0.27156 | -0.09021 | 0.04625 | 0.035 | -0.12938 | 0.269312 | 0.005641 | 0.068818 | -0.21673 | -0.07009 | -0.27135 | -0.20993 | -0.22398 | -0.26193 | -0.06772 | 0.04569 | 0.13024 | 0.01069 |
| COV413A | 0.074609 | 0.019011 | -0.07368 | 0.00708 | 0.00859 | -1.02237 | 0.135782 | -0.12812 | 0.03306 | 0.04805 | 0.12645 | 0.12966 | 0.029422 | 0.070588 | -0.0696 | 0.118583 | 0.060093 | -0.53163 | -0.25731 | -0.15078 | -0.01543 | -0.04819 | 0.11017 | 0.00997 | 0.0871 |
| DOTC24510 | 0.040886 | 0.03619 | 0.02898 | 0.04171 | -0.03527 | -0.6771 | -0.07844 | -0.09626 | -0.00466 | 0.04587 | 0.17309 | 0.11971 | -0.02135 | -0.00685 | -0.04157 | -0.11204 | 0.032381 | -0.83623 | -0.68834 | -0.12578 | -0.11656 | 0.0425 | 0.06404 | 0.11717 | 0.01311 |
| GIMEN | -0.16004 | 0.023579 | 0.01349 | 0.25184 | -0.0781 | -0.53564 | -0.01088 | -0.10396 | -0.05516 | -0.00845 | 0.06674 | 0.05769 | -0.19581 | 0.221767 | 0.215677 | 0.104362 | -0.21926 | 0.05008 | 0.03888 | -0.22103 | -0.04721 | -0.15467 | 0.11689 | 0.20665 | -0.09265 |
| GP5D | -0.04414 | 0.029966 | 0.04982 | 0.00649 | 0.02842 | -0.25602 | 0.065702 | -0.22976 | -0.018 | 0.04498 | 0.2028 | 0.18608 | 0.102434 | 0.022183 | -0.16718 | 0.040953 | 0.123795 | -0.57655 | -0.13337 | -0.00381 | -0.04919 | 0.0564 | -0.03072 | -0.05408 | 0.20302 |

| Cell line | ALK | AXL | CSF1R | DDR1 | DDR2 | EGFR | EPHA1 | EPHA2 | EPHA3 | EPHA4 | EPHA6 | EPHA7 | EPHB1 | EPHB2 | EPHB3 | EPHB4 | EPHB6 | ERBB2 | ERBB3 | ERBB4 | FGFR1 | FGFR2 | FGFR3 | FGFR4 | FLT1 |
|---|---|---|---|---|---|---|---|---|---|---|---|---|---|---|---|---|---|---|---|---|---|---|---|---|---|
| H103 | -0.01829 | -0.00877 | -0.1264 | -0.1328 | -0.03502 | -0.72681 | 0.081463 | 0.11703 | 0.06024 | 0.00384 | 0.13806 | -0.00865 | 0.01974 | 0.127194 | -0.07635 | -0.07175 | 0.032844 | -0.56629 | -0.42954 | -0.10203 | -0.08582 | 0.04193 | 0.15262 | 0.0102 | 0.02658 |
| H157 | 0.084906 | 0.013401 | -0.19002 | -0.09509 | 0.00454 | -0.54086 | -0.07606 | 0.15411 | -0.01178 | 0.12772 | 0.15799 | 0.17257 | 0.010926 | 0.15469 | -0.23379 | 0.084128 | 0.033501 | -0.36995 | -0.48985 | -0.0431 | -0.30425 | -0.10481 | -0.0206 | -0.10282 | 0.05884 |
| JOPACA1 | 0.111503 | -0.26982 | -0.06248 | -0.02613 | 0.00924 | -0.59429 | 0.063976 | 0.27329 | -0.01012 | 0.02955 | 0.19268 | 0.03594 | -0.00786 | 0.107564 | -0.21722 | 0.065458 | 0.027199 | -0.36635 | -0.27252 | -0.07784 | -0.06042 | -0.08395 | -0.11832 | 0.02942 | -0.03356 |
| LAN2 | -0.0705 | 0.049122 | 0.03118 | -0.05576 | -0.15471 | -0.22084 | -0.13047 | 0.0145 | -0.1028 | 0.06973 | 0.15207 | 0.15004 | -0.12621 | 0.336749 | 0.201467 | 0.055542 | 0.035826 | -0.24507 | 0.02427 | 0.071221 | 0.19071 | 0.07394 | -0.06881 | -0.20969 | -0.01576 |
| MB1 | -0.29098 | 0.118201 | -0.02609 | 0.02591 | -0.02723 | -0.0423 | 0.012967 | -0.01854 | 0.02033 | 0.21387 | 0.12086 | 0.10108 | 0.100938 | 0.215002 | -0.15563 | 0.010198 | 0.018665 | -0.1863 | -0.11078 | -0.0741 | -0.3375 | -0.05642 | -0.04567 | -0.04356 | 0.11178 |
| MS751 | 0.1407 | -0.08758 | -0.10267 | 0.02708 | -0.03415 | -0.27097 | -0.14399 | -0.05139 | -0.11157 | 0.13563 | 0.14717 | 0.04464 | 0.05946 | 0.012053 | -0.03916 | -0.04234 | 0.055229 | -0.1622 | -0.30681 | 0.048906 | -0.08966 | -0.02836 | 0.00637 | -0.06208 | 0.20957 |
| NGP | -0.33571 | 0.12487 | 0.0041 | -0.08464 | -0.00726 | -0.0962 | -0.08409 | -0.09693 | -0.08751 | 0.08236 | 0.12582 | -0.10191 | 0.023934 | 0.047844 | 0.131921 | 0.044381 | 0.207774 | -0.4129 | -0.17735 | -0.17415 | -0.21762 | -0.06381 | -0.0869 | 0.02929 | 0.04601 |
| NMB | -0.30601 | 0.054905 | 0.10448 | 0.26608 | 0.15794 | 0.07837 | 0.052852 | -0.16381 | -0.03832 | 0.15446 | 0.45304 | 0.06902 | 0.11407 | 0.044492 | 0.003131 | -0.08362 | 0.044759 | -0.14459 | -0.25132 | -0.34231 | -0.05651 | -0.09407 | -0.15396 | 0.03942 | 0.11319 |
| OACM51 | 0.011237 | 0.039929 | -0.01108 | 0.03212 | 0.02596 | -0.13895 | -0.0006 | -0.09936 | 0.01602 | 0.05218 | 0.02048 | 0.07992 | 0.023452 | -0.02548 | -0.11881 | 0.075213 | 0.04739 | -0.083 | -0.22739 | -0.1096 | 0.018471 | -0.15419 | -0.03036 | 0.01488 | 0.17388 |
| OCIP5X | 0.107073 | -0.0372 | 0.0441 | 0.02545 | -0.08989 | -0.02401 | 0.054593 | -0.00205 | -0.14505 | 0.03339 | 0.20544 | -0.13864 | -0.12344 | 0.055739 | -0.12817 | -0.07909 | 0.094066 | -0.23088 | -0.04099 | -0.10776 | 0.087761 | -0.00969 | 0.05426 | 0.10187 | -0.04674 |
| PA1 | -0.18706 | 0.043077 | 0.15458 | 0.12928 | 0.14542 | 0.02315 | 0.103983 | -0.07652 | 0.10209 | -0.02588 | 0.11128 | -0.04399 | 0.256021 | 0.008864 | 0.18865 | -0.01779 | 0.195552 | -0.36852 | -0.07709 | -0.49012 | 0.017706 | -0.1714 | 0.02341 | 0.05188 | -0.07834 |
| PACADD119 | -0.06697 | -0.09912 | -0.02638 | 0.04007 | 0.1027 | -0.6042 | 0.132998 | 0.10232 | 0.05367 | -0.20504 | 0.10306 | 0.16379 | -0.12409 | -0.04245 | 0.097928 | -0.00221 | 0.251793 | -0.49207 | -0.32526 | -0.15694 | 0.045521 | -0.09155 | 0.07167 | 0.12547 | 0.14085 |
| PACADD137 | 0.058231 | 0.243602 | -0.03758 | -0.08938 | -0.22291 | -0.42756 | -0.01373 | -0.17517 | -0.03711 | 0.1238 | 0.16128 | 0.03762 | -0.05541 | -0.00534 | 0.019358 | 0.066764 | 0.031127 | -0.79913 | -0.23202 | -0.08106 | 0.035591 | -0.05114 | 0.06602 | 0.09669 | 0.01453 |
| PACADD161 | -0.08881 | -0.03811 | 0.10651 | 0.09323 | 0.02991 | -0.26733 | 0.141005 | 0.05959 | 0.07297 | -0.11535 | 0.14855 | 0.12584 | 0.225997 | 0.049879 | -0.16113 | -0.22367 | 0.046427 | -0.32211 | -0.37317 | -0.01673 | 0.108403 | -0.09863 | -0.13996 | 0.02839 | 0.02713 |
| PACADD165 | 0.102342 | 0.044419 | -0.04717 | -0.1159 | -0.04428 | -0.25603 | -0.06998 | -0.00269 | -0.04486 | 0.1162 | 0.13678 | 0.12554 | -0.07399 | 0.212974 | 0.025632 | 0.034685 | 0.05258 | -0.17215 | -0.20861 | -0.14421 | -1.25384 | -0.0962 | 0.00103 | 0.08435 | 0.10306 |
| PACADD188 | 0.057662 | -0.07829 | 0.07983 | 0.15286 | -0.10497 | -0.18447 | 0.153989 | -0.02756 | -0.16955 | 0.31872 | 0.05361 | 0.05163 | 0.005318 | 0.052342 | 0.301809 | -0.07764 | 0.057832 | -0.5539 | 0.06666 | -0.37913 | -0.12007 | 0.01609 | -0.05469 | 0.21681 | 0.03161 |
| RPMI2650 | 0.051522 | 0.099959 | 0.09488 | -0.07824 | -0.06079 | -0.14606 | -0.0599 | -0.05396 | 0.07726 | 0.01339 | 0.24055 | 0.00028 | 0.061998 | 0.120122 | -0.20393 | 0.183852 | 0.07326 | -0.39019 | -0.25404 | 0.060642 | -0.41603 | 0.05172 | 0.08697 | 0.03471 | 0.21494 |
| SCLC22H | 0.043442 | 0.018087 | -0.05725 | 0.09824 | 0.03867 | -0.08485 | -0.19355 | -0.09493 | 0.0186 | -0.01556 | 0.17535 | 0.06875 | -0.04766 | 0.068149 | -0.21905 | -0.02907 | 0.074841 | -0.19969 | -0.21744 | -0.15923 | -0.49715 | 0.08637 | -0.07673 | -0.00564 | 0.17248 |
| SUM102PT | 0.004635 | -0.05189 | -0.09003 | -0.02951 | -0.05619 | -0.33584 | -0.12697 | -0.03162 | 0.09993 | 0.07335 | 0.0827 | 0.0998 | 0.038802 | 0.038938 | -0.11725 | -0.01453 | 0.035348 | -0.17628 | -0.28138 | -0.01668 | -0.0644 | -0.05455 | -0.02713 | -0.0184 | 0.1388 |
| SUM1315MO2 | -0.06492 | -0.11108 | 0.09868 | -0.20165 | 0.04999 | -0.19058 | -0.0232 | 0.04618 | 0.02051 | 0.01608 | 0.01677 | 0.01852 | 0.02489 | 0.044459 | -0.20805 | -0.00329 | -0.03529 | -0.16559 | -0.3612 | -0.0681 | -1.48586 | 0.16991 | 0.15239 | -0.09549 | 0.04403 |
| SUM149PT | -0.00158 | -0.1884 | 0.10304 | -0.20879 | -0.04228 | -0.76428 | -0.20145 | 0.09851 | -0.01451 | 0.0484 | -0.1046 | 0.12695 | 0.132172 | 0.142489 | 0.336142 | -0.11924 | 0.384863 | -0.12421 | -0.31408 | -0.06167 | -0.05458 | -0.23406 | 0.27957 | -0.15136 | 0.5941 |
| SUM159PT | 0.121289 | 0.034252 | 0.09308 | -0.28579 | -0.08534 | -0.40248 | -0.31111 | -0.01919 | 0.07251 | 0.08752 | 0.06882 | -0.01791 | 0.032826 | 0.283554 | -0.02135 | 0.060649 | 0.193726 | -0.07405 | -0.14754 | -0.02324 | 0.015255 | -0.17612 | 0.23139 | 0.10014 | 0.20027 |
| SUM185PE | -0.25166 | -0.23907 | 0.27818 | 0.12084 | 0.14125 | 0.05823 | 0.149208 | 0.07264 | 0.02565 | 0.23227 | -0.12608 | 0.27252 | 0.027102 | 0.052504 | 0.25854 | -0.24753 | 0.254535 | -0.10908 | -0.16669 | -0.45235 | 0.006921 | 0.01717 | -0.43091 | -0.0789 | 0.11468 |
| SUM190PT | 0.079362 | -0.05998 | -0.0192 | 0.0725 | -0.0042 | -0.20409 | -0.08717 | -0.01992 | -0.01458 | 0.04931 | 0.11473 | 0.10216 | -0.09824 | 0.023991 | -0.11025 | 0.044019 | 0.019575 | -1.47677 | -0.42511 | -0.03953 | 0.012862 | 0.13607 | 0.09689 | -0.0097 | 0.05059 |
| SUM229PE | 0.062808 | -0.10752 | -0.03886 | -0.0961 | -0.01979 | -0.97511 | -0.01679 | -0.1433 | -0.03168 | 0.01369 | 0.20572 | -0.03753 | 0.252353 | 0.049128 | -0.12471 | 0.104316 | -0.19145 | -0.18361 | -0.16951 | 0.081751 | -0.04114 | -0.06775 | -0.01181 | 0.04386 | 0.24915 |
| SUM52PE | -0.15902 | -0.28456 | -0.14205 | 0.27264 | 0.06296 | -0.02294 | 0.070591 | -0.07475 | 0.07993 | -0.07693 | -0.23449 | 0.14706 | 0.173072 | 0.143083 | -0.14177 | -0.19393 | 0.243251 | 0.02213 | -0.40335 | -0.02504 | -0.20805 | -0.8505 | -0.37778 | -0.21044 | 0.08544 |
| SW156 | -0.10301 | -0.43804 | -0.08706 | -0.05003 | -0.11327 | -0.16492 | -0.49877 | -0.03323 | -0.07146 | 0.21073 | 0.34108 | 0.14539 | 0.061813 | -0.01587 | -0.10048 | -0.05284 | 0.115092 | -0.22136 | -0.30402 | -0.10959 | 0.022483 | 0.06495 | -0.09164 | -0.08025 | 0.22337 |
| SW626 | -0.01818 | -0.32087 | -0.04174 | -0.08578 | 0.02177 | -0.40128 | 0.015614 | 0.00056 | 0.00317 | 0.01128 | 0.39435 | -0.10966 | 0.008747 | -0.02227 | -0.1119 | -0.08757 | 0.04714 | -0.0133 | -0.09484 | -0.13182 | 0.187847 | 0.01901 | -0.09365 | -0.04426 | -0.01123 |
| SW954 | -0.24111 | -0.09046 | 0.0461 | 0.1396 | -0.06292 | -0.61289 | 0.151421 | 0.17349 | 0.07976 | 0.03738 | 0.20384 | -0.03173 | -0.22608 | 0.142879 | -0.13486 | 0.172533 | -0.25463 | -1.01665 | -0.72645 | 0.012545 | -0.09751 | 0.01011 | 0.47298 | 0.00838 | 0.33968 |
| SW756 | -0.05992 | 0.069935 | -0.04801 | 0.01413 | -0.15381 | -0.25426 | -0.01406 | -0.06072 | 0.07759 | 0.04543 | 0.28644 | 0.0686 | -0.08519 | 0.015404 | -0.14171 | 0.015021 | 0.133475 | -0.37869 | -0.19045 | -0.10233 | -0.00588 | -0.14434 | 0.11076 | -0.15911 | 0.29556 |
| TO14 | -0.1564 | 0.04691 | 0.00411 | -0.08766 | -0.00932 | -0.1784 | -0.08528 | -0.13921 | -0.03529 | 0.086 | 0.09886 | 0.09817 | 0.227423 | 0.108708 | -0.04249 | -0.05569 | -0.11408 | -0.17233 | -0.17874 | -0.07486 | 0.072472 | -0.06052 | 0.2451 | -0.05627 | 0.16457 |
| UMUC13 | -0.05508 | 0.03623 | -0.07215 | -0.08014 | -0.0097 | -0.07367 | 0.032707 | -0.04213 | -0.02685 | 0.01514 | 0.20113 | 0.04793 | -0.03302 | 0.005671 | -0.09393 | 0.070046 | 0.115074 | -0.13636 | -0.29881 | -0.23931 | -1.91046 | -0.01152 | 0.14254 | 0.01632 | 0.00662 |
| UMUC14 | 0.032758 | 0.063917 | -0.09873 | 0.10541 | -0.06163 | -0.10468 | 0.044163 | 0.05517 | 0.01343 | 0.22981 | 0.08238 | 0.08447 | 0.203732 | 0.076046 | -0.14082 | -0.15808 | 0.162907 | -0.36955 | -0.06713 | -0.19874 | -0.00794 | 0.03634 | -1.28591 | 0.03439 | 0.02104 |
| UMUC16 | -0.31604 | 0.028174 | 0.11941 | -0.14691 | -0.06232 | -0.22779 | 0.118095 | -0.05441 | -0.01973 | 0.03563 | 0.30336 | 0.04235 | 0.028493 | 0.000262 | 0.06887 | -0.09053 | 0.120291 | -0.24669 | 0.02154 | -0.16025 | 0.240128 | -0.08805 | 0.03472 | 0.18294 | 0.13759 |
| UMUC5 | -0.10196 | -0.145 | -0.14999 | 0.01764 | 0.0429 | -0.27408 | 0.044998 | 0.45156 | -0.14549 | 0.10889 | -0.12455 | 0.14383 | -0.17465 | -0.00451 | 0.175247 | -0.0757 | 0.043803 | -0.26017 | -0.1956 | -0.18168 | -0.11585 | 0.04047 | 0.18783 | 0.01517 | -0.09781 |
| UMUC10 | -0.09466 | 0.0055 | 0.00613 | -0.0278 | -0.10893 | -0.66446 | 0.102942 | 0.19951 | -0.03146 | 0.14529 | 0.1373 | -0.10015 | 0.136071 | 0.060709 | 0.001079 | 0.020965 | 0.166212 | -0.50715 | -0.38816 | -0.05196 | 0.054132 | -0.05727 | -0.0119 | 0.17617 | -0.00075 |
| UMUC11 | 0.057569 | 0.071159 | -0.11173 | 0.15121 | -0.01142 | -0.03543 | 0.063087 | -0.01662 | 0.05966 | 0.1035 | 0.22131 | 0.10403 | 0.112682 | 0.108766 | 0.06752 | 0.151971 | 0.095769 | -0.20556 | -0.35951 | -0.0526 | -1.11213 | 0.19955 | -0.04218 | -0.13757 | 0.11758 |
| UMUC6 | 0.055612 | 0.109287 | -0.08809 | -0.04775 | 0.01888 | -0.30992 | 0.012845 | -0.01201 | -0.01682 | -0.04701 | 0.19632 | 0.02536 | 0.020141 | 0.06801 | -0.01148 | -0.04565 | 0.179856 | -0.14992 | -0.30551 | 0.021804 | -0.02584 | -0.06102 | 0.08601 | -0.00913 | 0.11463 |
| UMUC7 | -0.05855 | 0.421064 | -0.00768 | -0.03846 | -0.24071 | -0.54422 | -0.02407 | 0.04883 | 0.11774 | -0.05211 | 0.18077 | 0.28937 | 0.062154 | -0.00293 | -0.0109 | -0.0721 | 0.255129 | -0.87454 | -0.29996 | -0.29863 | -0.08549 | 0.10186 | 0.06125 | 0.10735 | -0.01144 |
| UMUC9 | 0.158002 | 0.078658 | -0.20225 | 0.0033 | -0.14145 | -0.32708 | 0.298668 | 0.22952 | 0.11814 | 0.1438 | 0.07361 | -0.0056 | -0.00255 | 0.172861 | 0.032609 | 0.0053 | 0.15949 | -0.4747 | -0.237 | -0.25477 | -0.09351 | -0.11339 | 0.09921 | 0.12016 | 0.05149 |
| UWB1289 | 0.11917 | -0.04719 | 0.02648 | -0.13248 | -0.13229 | -0.53275 | 0.135414 | 0.18162 | -0.00844 | 0.12331 | 0.0635 | 0.07005 | 0.038543 | 0.08691 | -0.06368 | -0.04117 | -0.01567 | -0.61155 | -0.77049 | -0.02836 | -0.17485 | -0.07649 | 0.01233 | 0.05082 | 0.07901 |
| VP229 | 0.089194 | 0.070385 | 0.01776 | 0.04743 | -0.1485 | -0.38913 | 0.082797 | -0.18222 | -0.0439 | -0.0171 | 0.09159 | 0.0338 | -0.09002 | 0.07045 | -0.00027 | -0.03944 | 0.152703 | -0.76462 | -0.35421 | -0.27943 | -0.04464 | 0.06236 | -0.15692 | -0.01572 | 0.10731 |
| WERIRB1 | -0.16333 | 0.152184 | -0.19313 | 0.01597 | -0.14282 | -0.00964 | -0.13604 | -0.05519 | 0.01319 | 0.03031 | 0.17496 | -0.00318 | -0.18695 | 0.244976 | -0.23811 | 0.139168 | 0.17207 | -0.58724 | -0.17387 | -0.01283 | -0.02698 | -0.11586 | 0.09568 | 0.21494 | 0.05948 |
| WPE1NA22 | 0.032265 | -0.0296 | -0.06549 | 0.01222 | -0.07284 | -1.04852 | -0.22118 | -0.15354 | -0.12639 | 0.06295 | 0.13916 | -0.01673 | 0.029517 | 0.032276 | -0.04146 | -0.01021 | 0.042998 | -0.20036 | -0.24343 | -0.08407 | 0.058411 | 0.00829 | 0.01127 | 0.03096 | 0.1042 |
| TC138 | -0.30523 | 0.1091 | 0.02578 | 0.1223 | -0.0801 | 0.04963 | 0.233616 | -0.12431 | 0.0038 | 0.03937 | 0.16086 | 0.27821 | 0.148214 | 0.058978 | -0.04116 | 0.067772 | 0.092613 | -0.43647 | -0.03802 | -0.24801 | -0.01403 | 0.27694 | -0.06605 | 0.27657 | -0.23641 |
| TC205 | -0.07479 | 0.01025 | -0.13414 | -0.11795 | -0.14577 | -0.05639 | 0.012182 | -0.07727 | -0.06319 | 0.05006 | 0.18192 | 0.22353 | -0.01245 | 0.0116 | 0.094147 | 0.121407 | 0.135671 | -0.38576 | -0.02328 | -0.03273 | 0.158939 | -0.13332 | -0.03492 | 0.07788 | 0.26298 |
| CCLFPEDS0008T | 0.064167 | 0.036697 | 0.09168 | 0.18827 | 0.03786 | -0.24574 | 0.293082 | -0.0638 | 0.01219 | 0.13015 | 0.05401 | 0.13839 | -0.01174 | 0.007667 | -0.02369 | 0.561694 | 0.036371 | -0.09931 | -0.18444 | -0.05607 | -0.27476 | -0.09619 | -0.07583 | 0.05818 | 0.01486 |
| A388 | -0.05984 | 0.017411 | -0.0032 | -0.06175 | 0.00768 | -1.10563 | -0.00758 | -0.09269 | -0.0584 | -0.05525 | 0.09747 | 0.05129 | 0.101763 | 0.035156 | -0.19393 | 0.002894 | 0.21694 | -0.40073 | -0.28083 | -0.04725 | -0.04959 | -0.00686 | 0.04388 | -0.10252 | 0.02107 |
| ASH3 | 0.057073 | 0.131329 | 0.02547 | -0.16852 | -0.03958 | -0.38052 | 0.118686 | -0.13584 | -0.05155 | 0.02797 | 0.09345 | -0.08825 | 0.067571 | 0.031433 | 0.054903 | 0.090693 | -0.00659 | -0.19179 | -0.28595 | -0.15413 | 0.57957 | -0.03047 | 0.13615 | 0.02783 | 0.1032 |
| BLUE1 | -0.03306 | -0.0358 | -0.04981 | -0.2452 | -0.0624 | -0.1897 | -0.40444 | -0.28969 | 0.06422 | 0.21681 | 0.0197 | 0.13884 | -0.0228 | -0.13928 | -0.24248 | -0.01843 | -0.01216 | -0.19886 | -0.2908 | 0.045249 | 0.000758 | -0.19285 | -0.22936 | 0.01752 | -0.18233 |
| BOKU | -0.13869 | -0.21796 | -0.05348 | 0.06887 | 0.06288 | -0.1006 | 0.046683 | -0.04567 | 0.04302 | 0.02224 | 0.21986 | 0.22263 | -0.05275 | 0.026114 | -0.32059 | -0.08106 | 0.016728 | -0.24012 | -0.25638 | -0.1417 | -0.43059 | -0.19956 | 0.07016 | -0.05488 | 0.04242 |
| BPH1 | -0.11855 | 0.049217 | -0.01975 | -0.01682 | -0.00644 | -1.2563 | 0.209654 | 0.74611 | -0.03913 | 0.10281 | 0.076 | 0.01183 | -0.25627 | 0.169542 | -0.12264 | 0.094267 | 0.079104 | -1.37544 | -0.77661 | -0.08844 | -0.0975 | 0.10271 | 0.10637 | 0.03175 | 0.25844 |
| C10 | -0.09836 | -0.11659 | -0.25664 | -0.15999 | -0.05887 | -0.02425 | 0.074725 | 0.17391 | -0.03169 | 0.11403 | 0.2659 | 0.1886 | -0.01239 | 0.087676 | -0.13698 | 0.228492 | 0.035744 | -0.02317 | -0.34707 | 0.040863 | 0.014386 | 0.01912 | 0.05543 | -0.27199 | -0.03222 |
| C75 | -0.41244 | -0.00769 | -0.06515 | 0.0219 | -0.00973 | -1.04627 | -0.36285 | 0.1666 | 0.23201 | 0.21055 | 0.07315 | 0.0842 | 0.319783 | 0.140033 | -0.20101 | 0.170744 | 0.106152 | -1.22994 | -0.25901 | -0.53329 | 0.12343 | -0.08639 | -0.11207 | 0.25281 | 0.00731 |
| C80 | -0.07096 | -0.10387 | 0.10295 | 0.21237 | 0.01977 | -0.28826 | 0.051839 | 0.21137 | 0.04977 | 0.01637 | 0.27783 | 0.07972 | 0.318858 | 0.08766 | 0.308478 | 0.190581 | -0.17964 | -0.30407 | -0.13011 | -0.31387 | -0.23103 | -0.06491 | -0.03913 | 0.11394 | -0.22751 |
| C84 | -0.05635 | -0.06897 | -0.0472 | 0.15671 | -0.00899 | -0.94991 | 0.089923 | -0.13968 | -0.04705 | -0.01632 | 0.03558 | -0.15392 | -0.04597 | 0.158633 | -0.02307 | 0.180328 | 0.160379 | -0.48739 | -0.27121 | 0.077536 | -0.23215 | -0.25086 | 0.21228 | 0.12523 | -0.09468 |
| C99 | 0.041215 | 0.158894 | 0.20994 | -0.00239 | 0.00495 | -0.23621 | 0.035679 | -0.05792 | 0.18534 | 0.10296 | 0.17264 | 0.14182 | -0.00501 | -0.10707 | -0.09869 | 0.356893 | 0.10033 | -0.91157 | -0.07946 | -0.24839 | 0.0002 | -0.07671 | 0.12654 | 0.02527 | 0.06803 |
| CHLA90 | 0.078009 | 0.073081 | 0.11392 | -0.1806 | -0.05686 | -0.33289 | -0.09951 | -0.17877 | -0.02666 | 0.08229 | 0.28318 | 0.15735 | -0.00526 | -0.09057 | -0.19346 | -0.14684 | -0.10053 | -0.27953 | -0.34065 | -0.09906 | -0.90637 | 0.07043 | 0.19607 | 0.22212 | 0.12619 |
| EGI1 | 0.031853 | -0.24719 | -0.08463 | -0.15307 | -0.04499 | -0.43095 | 0.117576 | -0.09019 | -0.02457 | -0.00718 | 0.13886 | 0.04258 | 0.022395 | 0.12163 | -0.10241 | 0.016219 | 0.145307 | -0.19285 | -0.24279 | -0.01926 | -0.1085 | -0.18241 | -0.06777 | -0.12116 | 0.21009 |
| EMTOKA | 0.03909 | 0.053547 | -0.06402 | -0.20088 | 0.20065 | -0.72813 | 0.046677 | 0.00908 | -0.03189 | 0.27834 | 0.02165 | -0.01973 | 0.182697 | 0.158414 | -0.09071 | -0.03843 | -0.14084 | -0.37995 | -0.17126 | -0.0454 | -0.19243 | 0.09633 | 0.01374 | 0.05826 | 0.06454 |
| ESO26 | 0.076535 | 0.069509 | 0.0263 | 0.03931 | -0.05583 | -1.1275 | -0.09362 | -0.11041 | 0.05087 | 0.1033 | 0.05693 | 0.14668 | 0.038464 | 0.046603 | -0.15382 | 0.011218 | -0.1446 | -2.57494 | -0.72238 | -0.12208 | -0.08825 | -0.08458 | 0.00548 | -0.01362 | 0.04866 |
| ESO51 | 0.040609 | 0.002214 | -0.1372 | 0.1056 | -0.1188 | 0.13871 | 0.061386 | 0.08934 | 0.16159 | 0.12894 | -0.00303 | 0.16792 | 0.139462 | 0.022926 | -0.05554 | -0.25163 | 0.242087 | -0.12983 | -0.41052 | -0.08557 | -0.32461 | -0.11828 | 0.02241 | 0.01013 | 0.2379 |
| FARAGE | 0.178244 | 0.062911 | 0.12213 | 0.03515 | -0.08488 | -0.23251 | -0.10636 | -0.01024 | -0.02841 | -0.00955 | -0.11232 | 0.01155 | 0.181187 | 0.000816 | 0.089762 | 0.034282 | 0.071414 | -0.25196 | -0.222 | -0.01663 | -0.08612 | -0.05236 | 0.10377 | -0.20672 | 0.03435 |
| FLO1 | 0.039159 | -0.42457 | -0.04481 | -0.04577 | -0.15358 | -0.00918 | -0.01625 | 0.25037 | 0.0122 | 0.07138 | 0.21605 | 0.16669 | -0.12452 | 0.090997 | -0.04349 | 0.025658 | -0.06467 | -0.25974 | -0.22073 | -0.10916 | 0.049202 | 0.11296 | -0.09865 | -0.04825 | 0.15881 |
| H357 | -0.05325 | 0.058835 | -0.00724 | -0.05482 | 0.17623 | -0.88476 | -0.00651 | 0.51733 | 0.00083 | -0.0053 | 0.07136 | 0.01643 | 0.074873 | -0.0335 | -2.9E-05 | 0.145358 | -0.16153 | -0.99974 | -0.42098 | -0.23217 | 0.036422 | 0.03082 | 0.00724 | 0.10362 | 0.15176 |
| H376 | 0.104942 | 0.049493 | -0.10865 | -0.00927 | 0.03576 | -0.45333 | 0.136359 | 0.16736 | -0.10315 | -0.00081 | 0.13495 | 0.0943 | 0.090811 | 0.109615 | -0.11016 | 0.023256 | 0.064312 | -0.44161 | -0.35738 | 0.124194 | -0.20626 | -0.28838 | -0.01987 | -0.00708 | 0.22894 |
| H413 | -0.01021 | 0.036771 | -0.00778 | -0.12009 | -0.11451 | -0.42267 | 0.079188 | 0.09524 | -0.05038 | 0.18204 | -0.00263 | 0.19262 | -0.05228 | 0.214979 | -0.13809 | 0.19401 | 0.03797 | -1.03655 | -0.97151 | -0.14108 | -0.05325 | -0.0702 | -0.05174 | 0.07537 | 0.12706 |
| HCA1 | 0.081688 | -0.05219 | -0.0707 | 0.00437 | -0.07555 | -0.54351 | 0.074558 | -0.06503 | -0.0515 | 0.03341 | 0.13998 | -0.02456 | 0.033107 | 0.036582 | -0.07985 | 0.058699 | 0.192997 | -0.3977 | -0.25885 | -0.06638 | 0.020468 | -0.17745 | 0.03287 | -0.07213 | 0.06811 |
| HCS2 | 0.088595 | -0.0713 | -0.15836 | -0.08526 | -0.09602 | -0.6297 | 0.105796 | 0.13219 | -0.03009 | 0.09736 | 0.08098 | 0.06714 | -0.01994 | 0.1872 | -0.14032 | -0.01755 | -0.1091 | -0.31917 | -0.11565 | 0.070627 | -0.03284 | -0.13605 | 0.01345 | -0.07956 | 0.03272 |
| HEC1 | 0.061066 | 0.002011 | 0.04487 | -0.00444 | -0.13502 | -0.15764 | -0.04205 | -0.00508 | -0.03493 | 0.09033 | 0.05172 | 0.04176 | -0.05053 | 0.051684 | -0.08721 | 0.010999 | 0.048414 | -0.31054 | -0.30134 | -0.0822 | -0.02016 | -0.03615 | 0.0914 | 0.05273 | 0.13201 |
| HEC116 | -0.01125 | 0.161472 | -0.10287 | -0.0005 | -0.11003 | -0.31147 | -0.0414 | -0.0598 | -0.12388 | 0.17603 | 0.10492 | -0.08499 | 0.040527 | 0.056001 | -0.09873 | -0.08948 | 0.057378 | -0.35701 | -0.18906 | 0.036032 | -0.10769 | 0.14666 | 0.04039 | 0.14227 | 0.22911 |
| HG3 | 0.12087 | -0.16137 | -0.01268 | 0.09495 | 0.0865 | -0.09358 | -0.01226 | -0.07052 | -0.03323 | 0.06624 | 0.17133 | 0.21398 | 0.088139 | 0.008477 | -0.12482 | 0.037646 | 0.068929 | -0.49016 | -0.23825 | -0.10202 | -0.23165 | -0.09739 | 0.06036 | -0.0012 | 0.02866 |
| HKA1 | -0.0334 | -0.04243 | -0.27684 | -0.00431 | -0.04184 | -0.32961 | -0.01685 | -0.01822 | 0.08084 | -0.14686 | -0.01395 | 0.23003 | 0.051746 | 0.278975 | -0.28041 | -0.02272 | 0.286858 | -0.82818 | -0.42475 | 0.001326 | 0.00213 | -0.12494 | 0.21906 | -0.00341 | 0.22047 |
| HMY1 | -0.04569 | 0.083461 | -0.00486 | -0.04615 | -0.05511 | -0.03742 | -0.11789 | -0.03196 | -0.04236 | 0.0511 | 0.13807 | 0.04435 | 0.014422 | -0.00546 | -0.16739 | 0.09113 | 0.045715 | -0.25613 | -0.24563 | -0.06495 | 0.076012 | 0.00793 | -0.12173 | 0.03439 | 0.04067 |

| Cell line | ALK | AXL | CSF1R | DDR1 | DDR2 | EGFR | EPHA1 | EPHA2 | EPHA3 | EPHA4 | EPHA6 | EPHA7 | EPHB1 | EPHB2 | EPHB3 | EPHB4 | EPHB6 | ERBB2 | ERBB3 | ERBB4 | FGFR1 | FGFR2 | FGFR3 | FGFR4 | FLT1 |
|---|---|---|---|---|---|---|---|---|---|---|---|---|---|---|---|---|---|---|---|---|---|---|---|---|---|
| HSC1 | 0.080969 | -0.13688 | -0.09466 | 0.05818 | -0.03549 | -0.1945 | -0.02267 | 0.11185 | -0.1759 | 0.05136 | 0.39787 | -0.00505 | -0.01575 | 0.064668 | -0.00452 | 0.146388 | 0.090277 | -0.13538 | -0.55312 | -0.1097 | -0.06582 | -0.06139 | 0.05673 | 0.10613 | 0.01552 |
| HSC5 | -0.05936 | -0.02324 | 0.00418 | -0.07901 | -0.04775 | -0.61728 | 0.096318 | 0.2978 | 0.03352 | -0.00603 | 0.21065 | 0.11056 | 0.083965 | -0.02267 | -0.05449 | 0.041484 | 0.074938 | -0.93101 | -0.60942 | -0.21678 | -0.06067 | 0.03874 | -0.00832 | -0.14262 | 0.12763 |
| HT3 | 0.060472 | 0.120763 | -0.02817 | -0.07692 | -0.11028 | -0.0551 | 0.011452 | 0.00932 | -0.16508 | 0.17382 | 0.21852 | 0.05269 | -0.02277 | 0.235795 | -0.26193 | -0.03132 | 0.196856 | -0.07709 | -0.12038 | 0.055028 | -0.16485 | 0.00639 | 0.10889 | -0.0732 | 0.07711 |
| HUO9 | -0.32489 | -0.06674 | 0.02691 | 0.01028 | 0.18178 | -0.09805 | 0.169019 | 0.11424 | -0.02826 | -0.08938 | -0.04584 | 0.28664 | -0.12051 | 0.003317 | -0.23738 | -0.09709 | 0.083225 | -0.21294 | -0.01708 | -0.23086 | -0.05697 | 0.1855 | 0.17382 | -0.03616 | -0.13546 |
| IHH4 | 0.078226 | -0.15771 | 0.02355 | 0.01921 | -0.06501 | -0.16363 | -0.14191 | -0.20817 | 0.04753 | 0.02956 | 0.26485 | 0.0741 | 0.001365 | 0.026893 | -0.13916 | -0.01417 | 0.023268 | -0.27435 | -0.1908 | 0.056993 | -0.02503 | -0.04184 | 0.17394 | 0.05234 | 0.22217 |
| JAR | 0.220925 | 0.069057 | -0.23918 | 0.01589 | 0.03978 | -0.15875 | -0.0792 | -0.08881 | 0.13185 | 0.00773 | -0.06731 | 0.11075 | 0.129626 | 0.316645 | -0.11121 | -0.04434 | 0.019824 | -0.10586 | -0.23267 | 0.039368 | -0.08668 | 0.03269 | 0.05533 | -0.0533 | 0.04575 |
| JEG3 | 0.170647 | 0.146952 | 0.16452 | 0.14605 | -0.08593 | -0.11323 | -0.03534 | -0.06916 | -0.06819 | 0.04736 | 0.12163 | 0.20535 | 0.021752 | 0.034808 | -0.06122 | 0.202076 | 0.165551 | -0.39935 | -0.20501 | -0.23271 | -5.8E-05 | -0.22466 | -0.18097 | 0.12899 | 0.23966 |
| JMURTK2 | 0.045673 | 0.096597 | -0.07235 | 0.00743 | -0.09121 | -0.14853 | -0.05447 | -0.00167 | -0.08158 | 0.14001 | 0.15963 | 0.03515 | 0.096204 | 0.153013 | -0.12653 | 0.150155 | 0.073288 | -0.03753 | -0.1723 | 0.021418 | -1.38489 | -0.53782 | 0.03987 | 0.0012 | 0.1805 |
| KARPAS1718 | 0.133705 | 0.018479 | -0.08189 | 0.09303 | -0.0588 | -0.21206 | -0.14022 | -0.22363 | 0.03413 | -0.05847 | 0.15957 | 0.06245 | 0.014041 | -0.04314 | -0.07804 | 0.086418 | 0.003373 | -0.1286 | -0.29301 | 0.013292 | -0.10668 | 0.08233 | 0.10601 | 0.00627 | 0.3709 |
| KKU100 | -0.03943 | 0.031916 | -0.01358 | 0.09356 | -0.23624 | -0.24606 | 0.000797 | 0.03286 | 0.04645 | 0.10356 | 0.2726 | 0.23707 | 0.022411 | 0.176075 | 0.043041 | 0.217564 | 0.111212 | -0.21852 | -0.23424 | -0.20009 | -0.06298 | -0.10574 | 0.28974 | -0.03924 | 0.13575 |
| KKU213 | 0.063062 | -0.2505 | 0.13111 | 0.16005 | -0.13629 | -0.40123 | 0.08278 | 0.18142 | -0.12831 | 0.1166 | 0.19002 | 0.0527 | -0.0466 | 0.106546 | -0.13335 | 0.01059 | -0.03978 | -0.29219 | -0.17926 | -0.16912 | -0.07726 | 0.12182 | 0.1107 | 0.01731 | 0.00903 |
| KML1 | -0.03898 | 0.091663 | 0.08556 | -0.07893 | -0.23967 | -0.25055 | 0.10589 | 0.07338 | -0.02625 | 0.24814 | 0.05666 | 0.03837 | -0.19532 | -0.11382 | 0.227778 | -0.2741 | 0.327064 | -0.22903 | -0.12813 | -0.22957 | -0.00725 | -0.02402 | 0.12048 | 0.06167 | -0.16047 |
| KON | 0.003094 | 0.078803 | 0.01755 | -0.04159 | -0.19316 | -0.57276 | 0.161312 | 0.51043 | -0.12002 | 0.2061 | 0.09528 | 0.02954 | -0.03055 | 0.118491 | 0.085312 | 0.002855 | 0.177242 | -0.34252 | -0.37135 | 0.011371 | -0.40704 | -0.0729 | 0.16886 | -0.04034 | 0.01347 |
| KOSC2 | 0.207043 | 0.024779 | -0.07663 | -0.02794 | 0.10765 | -0.20034 | 0.162364 | 0.29758 | -0.08138 | 0.17442 | 0.10409 | -0.11793 | -0.07474 | 0.250911 | -0.14896 | 0.1499 | 0.16047 | -0.7987 | -0.54811 | 0.07411 | -0.08184 | -0.12313 | 0.10279 | -0.06065 | 0.16538 |
| KYAE1 | 0.120835 | 0.068321 | 0.08683 | 0.23181 | 0.07761 | -0.48756 | -0.0887 | 0.19809 | -0.11985 | 0.35813 | 0.14713 | 0.25809 | 0.124002 | 0.287139 | 0.217796 | 0.113208 | 0.16144 | -1.16797 | -0.64695 | -0.17078 | -0.03131 | -0.29521 | 0.25933 | 0.31592 | 0.35277 |
| LO68 | 0.090526 | -0.16748 | -0.13349 | 0.04712 | 0.12496 | -0.07527 | -0.0181 | 0.20258 | -0.18822 | 0.07475 | 0.09868 | 0.21573 | 0.040245 | 0.037136 | -0.41687 | -0.00963 | 0.134489 | -0.33367 | -0.12267 | 0.002574 | -1.19749 | 0.20767 | -0.19717 | -0.0214 | 0.1217 |
| LS | -0.46893 | -0.08393 | 0.05061 | 0.10362 | 0.05263 | -0.1459 | 0.064715 | -0.15532 | -0.1179 | 0.27544 | -0.13777 | 0.03778 | -0.02168 | -0.03866 | 0.160449 | 0.046226 | -0.21482 | -0.19852 | -0.12889 | -0.14401 | -0.29168 | -0.327 | 0.22599 | 0.06489 | -0.04284 |
| LU135 | 0.064502 | 0.030077 | -0.04581 | 0.06432 | -0.03616 | -0.01808 | 0.086466 | -0.09406 | -0.06741 | 0.0446 | 0.10674 | 0.0821 | 0.032636 | 0.105802 | -0.14901 | 0.034024 | 0.063231 | -0.17519 | -0.16866 | -0.03406 | -0.08391 | -0.24104 | -0.04436 | -0.12351 | 0.11078 |
| MCC13 | 0.093941 | -0.11159 | -0.12855 | 0.17573 | -0.11567 | -0.10688 | -0.16083 | -0.06907 | 0.02587 | 0.04072 | 0.05109 | 0.07105 | 0.00464 | 0.108287 | -0.14044 | 0.024272 | 0.088143 | -0.26206 | -0.28003 | -0.04731 | 0.141592 | 0.00144 | -0.08388 | 0.06842 | 0.0545 |
| MCC142 | -0.13922 | 0.061158 | -0.033 | -0.07938 | -0.03719 | -0.14261 | 0.009585 | 0.10379 | 0.0902 | 0.01138 | 0.0513 | 0.13959 | 0.14203 | 0.05667 | -0.22196 | -0.21427 | 0.088706 | -0.15424 | -0.33467 | 0.091661 | -0.64522 | -0.08965 | -0.08355 | -0.1438 | 0.19994 |
| MCC26 | -0.34791 | 0.130353 | 0.03437 | 0.04104 | -0.03002 | 0.02969 | 0.104674 | 0.25687 | -0.08123 | 0.04925 | 0.06975 | 0.19009 | -0.13658 | 0.194247 | 0.092921 | 0.134652 | 0.288206 | -0.13747 | -0.20355 | -0.09302 | -0.05302 | 0.02484 | 0.09933 | 0.1796 | 0.05706 |
| MEL202 | 0.019924 | 0.025911 | 0.04505 | -0.05985 | -0.05209 | -0.14361 | 0.065594 | -0.00928 | -0.05803 | -0.00221 | 0.06234 | 0.04185 | -0.03989 | -0.00528 | -0.05106 | 0.164884 | 0.107726 | -0.28646 | -0.29665 | -0.23586 | -0.19477 | -0.02637 | 0.12737 | 0.00862 | 0.09478 |
| MERO14 | 0.042853 | 0.294888 | -0.24156 | -0.11015 | -0.0341 | -0.3128 | 0.321113 | 0.01151 | -0.15972 | 0.08541 | 0.17298 | 0.01081 | 0.111549 | 0.194721 | -0.32163 | 0.073205 | -0.01206 | -0.58206 | -0.04312 | 0.083667 | -1.12758 | 0.0274 | 0.1276 | 0.28852 | 0.12313 |
| MERO25 | 0.092251 | 0.00139 | -0.11733 | 0.0292 | -0.1305 | -0.29361 | -0.04669 | -0.0628 | -0.09405 | 0.08245 | 0.22282 | 0.10257 | -0.0239 | 0.123157 | -0.13856 | 0.079642 | 0.065478 | -0.13025 | -0.31127 | 0.034892 | -0.06797 | -0.12261 | -0.14231 | -0.05882 | 0.24608 |
| MERO41 | 0.163147 | -0.05714 | 0.00143 | -0.1071 | 0.04992 | -0.01451 | -0.20224 | -0.05772 | 0.04267 | -0.07368 | 0.3047 | 0.03383 | 0.213739 | 0.054256 | -0.03961 | 0.114073 | -0.14379 | -0.24166 | -0.30373 | 0.093847 | -0.90961 | 0.01263 | -0.207 | -0.14328 | 0.25788 |
| MERO48A | -0.03612 | -0.33732 | -0.17082 | -0.04679 | -0.19532 | -0.15524 | -0.02948 | 0.08785 | -0.13371 | 0.10643 | 0.07853 | 0.01787 | -0.05034 | 0.158003 | -0.37088 | 0.110926 | -0.03285 | -0.31589 | -0.28137 | 0.164505 | -0.97675 | -0.05044 | 0.08665 | -0.06336 | 0.20871 |
| MERO82 | 0.041487 | 0.110651 | -0.10289 | 0.10369 | -0.03006 | -0.22488 | 0.145636 | 0.04927 | 0.04452 | 0.20562 | 0.03508 | -0.01177 | 0.019833 | 0.240063 | 0.142345 | 0.073975 | 0.11186 | -0.25338 | -0.30401 | -0.0398 | -1.24515 | -0.11527 | -0.01924 | -0.26549 | 0.12541 |
| MERO83 | 0.191361 | 0.110323 | -0.04827 | -0.17261 | -0.02995 | -0.51258 | -0.09639 | 0.01263 | 0.05283 | 0.24714 | 0.09887 | 0.12691 | 0.214668 | 0.080348 | -0.09309 | 0.344225 | 0.002976 | -0.34751 | -0.30909 | -0.12627 | -1.42302 | 0.0666 | 0.04024 | 0.04449 | -0.04706 |
| MERO95 | 0.026739 | -0.10541 | -0.07629 | -0.05698 | -0.00318 | -0.36823 | -0.26834 | 0.12149 | -0.18496 | 0.04167 | 0.10911 | 0.09733 | 0.025411 | 0.003421 | -0.04867 | -0.00398 | 0.203973 | -0.10191 | -0.212 | -0.08333 | -1.37048 | -0.00251 | 0.01902 | -0.05765 | 0.18038 |
| MM127 | -0.15551 | -0.00875 | -0.08598 | -0.05485 | 0.04331 | -0.15885 | 0.003414 | -0.02904 | -0.01068 | -0.08499 | 0.10332 | 0.01709 | 0.03887 | 0.122834 | -0.29713 | 0.104012 | -0.02261 | -0.10172 | -0.18946 | 0.098921 | -0.04434 | -0.01428 | 0.08115 | -0.017 | 0.08641 |
| MM370 | -0.01241 | -0.01768 | 0.0432 | -0.31429 | 0.06668 | -0.02026 | -0.06733 | -0.05702 | 0.03698 | 0.24556 | 0.05682 | 0.1509 | 0.001911 | 0.147519 | 0.08925 | 0.209957 | 0.112285 | -0.18674 | -0.19715 | -0.10882 | -0.09182 | -0.08439 | -0.00944 | -0.11091 | -0.05746 |
| MM383 | 0.136809 | 0.080846 | -0.0657 | 0.16663 | 0.2477 | -0.19964 | -0.0921 | -0.00179 | 0.26399 | 0.12994 | 0.15201 | 0.0587 | -0.10464 | 0.076377 | -0.02271 | -0.16329 | 0.311846 | -0.47127 | -0.56836 | -0.11972 | 0.191459 | 0.04624 | -0.19151 | -0.02738 | 0.1118 |
| MM386 | -0.12715 | 0.098428 | 0.01002 | -0.28737 | 0.00836 | 0.00698 | -0.05143 | -0.14667 | 0.01851 | -0.07516 | 0.15841 | 0.11669 | 0.021603 | -0.02307 | -0.03571 | 0.055156 | 0.051993 | -0.24615 | -0.25194 | -0.0738 | 0.183535 | -0.11126 | 0.10961 | -0.03018 | -0.03642 |
| MM426 | -0.12106 | 0.02765 | 0.06964 | -0.0881 | -0.02634 | 0.0292 | 0.192752 | -0.08905 | -0.22019 | 0.12698 | 0.20689 | 0.05328 | 0.058274 | 0.019613 | 0.060458 | 0.224287 | 0.375688 | -0.2367 | -0.21574 | -0.32366 | -0.17079 | -0.10752 | 0.03956 | 0.11343 | -0.08008 |
| MOLM14 | 0.21033 | 0.044012 | -0.23838 | -0.11904 | 0.17686 | -0.12965 | -0.1572 | -0.15592 | 0.01907 | -0.06565 | 0.1476 | 0.2087 | -0.07426 | -0.05867 | -0.01567 | 0.032996 | 0.211888 | -0.16082 | -0.07154 | 0.06637 | 0.049816 | -0.09261 | 0.03984 | 0.08389 | -0.03311 |
| MUTZ8 | -0.02019 | 0.013151 | -0.25635 | -0.01775 | 0.21794 | -0.04587 | -0.24709 | -0.17918 | 0.13883 | 0.27345 | 0.13272 | -0.01741 | 0.075219 | 0.105918 | -0.50991 | -0.09119 | 0.559128 | -0.27237 | -0.1714 | 0.083842 | 0.00796 | -0.01914 | 0.08525 | 0.2214 | 0.31302 |
| NH12 | -0.40046 | 0.066026 | 0.045 | -0.23614 | -0.04122 | -0.24276 | 0.029586 | 0.00357 | 0.06235 | -0.05675 | 0.42 | -0.06256 | -0.00414 | 0.023713 | -0.09826 | 0.017569 | 0.054998 | -0.37443 | -0.33218 | -0.06047 | 0.052226 | 0.00521 | 0.02165 | -0.03836 | 0.09671 |
| NO10 | 0.005316 | 0.060338 | 0.05358 | -0.00828 | -0.10829 | -0.12734 | -0.04159 | 0.23627 | 0.12695 | -0.00069 | 0.26056 | 0.07742 | -0.00917 | 0.110218 | -0.12784 | 0.097125 | 0.071884 | -0.28453 | -0.09953 | -0.09156 | 0.001048 | 0.11787 | -0.13769 | 0.00626 | 0.01188 |
| NO11 | 0.116755 | -0.28584 | -0.14304 | 0.0303 | -0.21019 | 0.12085 | -0.06204 | 0.11727 | 0.05868 | 0.20912 | -0.18943 | 0.1235 | -0.09889 | 0.146435 | -0.05165 | 0.038546 | 0.170007 | -0.14507 | -0.53732 | 0.172873 | -0.35418 | -0.03804 | -0.21797 | 0.10344 | 0.31304 |
| NOZ | 0.126488 | -0.30006 | -0.13083 | 0.01036 | -0.07721 | -0.03254 | -0.05207 | 0.07352 | 0.00628 | 0.0301 | 0.14385 | 0.04217 | 0.095157 | 0.030717 | -0.08241 | -0.00206 | -0.00335 | -0.235 | -0.11039 | -0.10545 | 0.002889 | 0.01971 | 0.17784 | 0.13348 | 0.34204 |
| NP2 | -0.02456 | -0.07581 | 0.02699 | -0.14477 | -0.13732 | -0.11806 | 0.070169 | 0.115 | -0.03525 | 0.08755 | 0.15551 | 0.03006 | 0.016981 | -0.02734 | -0.01903 | 0.062308 | 0.055316 | -0.19942 | -0.20285 | -0.0488 | -0.82496 | -0.02004 | -0.07446 | -0.00726 | 0.09967 |
| NP3 | 0.077246 | 0.080362 | 0.00242 | -0.0749 | -0.04308 | -0.13149 | 0.045857 | -0.14096 | -0.12939 | -0.03493 | 0.0971 | 0.05224 | 0.004513 | 0.063537 | -0.12662 | -0.05992 | 0.176246 | -0.19526 | -0.06892 | 0.040403 | -1.00472 | -0.10264 | -0.04137 | -0.11474 | 0.03391 |
| NP5 | -0.26095 | 0.019644 | 0.02227 | 0.01127 | -0.17283 | 0.10604 | -0.07939 | -0.12652 | -0.11028 | 0.09658 | 0.14093 | -0.05936 | -0.27458 | 0.039332 | -0.12536 | 0.213611 | -0.20075 | -0.32135 | -0.34039 | -0.04253 | -0.08911 | -0.08203 | 0.14878 | 0.036 | 0.37057 |
| NP8 | 0.105945 | 0.179272 | 0.06035 | 0.21621 | 0.10664 | -0.0729 | -0.18263 | -0.06995 | -0.18962 | 0.21049 | 0.01982 | -0.04035 | -0.09149 | 0.147392 | 0.088451 | -0.05811 | 0.099656 | -0.11731 | -0.29754 | -0.06117 | -0.67249 | 0.04513 | 0.15535 | -0.0001 | -0.04597 |
| OCILY18 | 0.079041 | -0.05779 | 0.03621 | -0.08688 | -0.03599 | -0.13795 | 0.049854 | -0.01175 | -0.05183 | 0.10924 | 0.12114 | 0.12849 | 0.022463 | -0.02759 | -0.21953 | -0.02676 | -0.06275 | -0.32299 | -0.15183 | 0.001196 | -0.05555 | 0.016 | 0.07778 | 0.03494 | 0.00722 |
| OCIM2 | 0.20408 | -0.22803 | 0.16946 | 0.12317 | -0.03987 | 0.0277 | -0.32029 | -0.03491 | 0.01833 | 0.17524 | 0.00453 | 0.22991 | -0.11701 | 0.019961 | -0.09641 | 0.02011 | 0.18502 | -0.22093 | -0.24675 | 0.059927 | -0.23771 | 0.03212 | 0.05337 | 0.32036 | 0.21036 |
| OCUG1 | -0.18727 | 0.044651 | 0.15782 | 0.19788 | -0.07402 | -0.35364 | 0.048883 | 0.21094 | 0.08199 | -0.03783 | 0.22319 | 0.10902 | -0.11632 | 0.186347 | -0.31867 | 0.293165 | -0.22196 | -0.28091 | -0.432 | -0.08436 | -0.01689 | 0.06176 | 0.02856 | -0.01293 | 0.27665 |
| ONDA7 | -0.04802 | -0.08906 | 0.06226 | 0.00966 | -0.01592 | -0.06467 | 0.087573 | -0.0131 | -0.08564 | 0.0137 | 0.11717 | 0.10345 | 0.003165 | -0.06339 | -0.01906 | 0.002171 | 0.161751 | -0.34162 | -0.28915 | -0.17521 | -0.4514 | -0.10283 | -0.00175 | -0.06241 | 0.08395 |
| ONDA8 | -0.13327 | -0.07653 | -0.21596 | 0.05294 | 0.05672 | -0.08717 | 0.033817 | 0.25284 | -0.16193 | 0.19126 | 0.07143 | -0.0058 | 0.159162 | 0.069084 | 0.145132 | -0.10063 | 0.055049 | -0.09089 | -0.16406 | -0.23758 | -1.23996 | -0.04349 | 0.19704 | 0.01677 | 0.10946 |
| ONDA9 | -0.07413 | 0.149428 | -0.0306 | -0.15302 | 0.06712 | -0.08519 | 0.223031 | -0.05502 | -0.42364 | 0.18453 | 0.14972 | 0.14204 | 0.141252 | 0.070018 | -0.26062 | 0.249959 | 0.037575 | -0.29621 | -0.0902 | -0.02134 | 0.038262 | -0.04959 | -0.02691 | -0.13024 | 0.24926 |
| OSC19 | 0.17628 | -0.0355 | 0.09962 | 0.13843 | -0.08245 | -0.78401 | 0.062002 | 0.32247 | -0.03335 | 0.05654 | -0.21843 | 0.0422 | 0.062701 | 0.348457 | 0.194602 | 0.196987 | 0.07535 | -0.75168 | -0.40966 | -0.0955 | -0.11656 | -0.00616 | 0.21497 | 0.14413 | 0.26951 |
| OSC20 | -0.04688 | -0.16765 | -0.16886 | 0.19272 | -0.23757 | -0.9483 | 0.043264 | 0.16577 | -0.4404 | 0.08047 | 0.15873 | 0.01146 | 0.103859 | 0.098503 | -0.2334 | -0.02223 | -0.06245 | -0.29734 | -0.24225 | -0.16881 | 0.043605 | -0.13063 | -0.28849 | -0.02699 | 0.08678 |
| P4E6 | 0.076497 | 0.049401 | -0.15048 | -0.05525 | 0.02024 | -1.02356 | -0.13079 | -0.01579 | -0.06279 | -0.03318 | 0.1003 | 0.06613 | 0.060586 | 0.118766 | -0.09794 | 0.021961 | 0.138531 | -0.16002 | -0.27182 | -0.11013 | -0.0713 | -0.01983 | -0.08796 | 0.00696 | 0.18386 |
| PEA1 | -0.04332 | -0.11046 | -0.17886 | 0.07217 | 0.06308 | -0.1234 | -0.11114 | 0.28033 | 0.05939 | 0.01224 | 0.15073 | 0.09104 | -0.0914 | -0.13885 | -0.16147 | 0.089085 | 0.076686 | -0.49737 | -0.23601 | 0.01582 | -0.04776 | 0.04645 | -0.05222 | -0.17092 | 0.11419 |
| PEO1 | 0.143616 | 0.02748 | -0.02385 | -0.05344 | 0.02531 | -0.33787 | 0.033801 | 0.08869 | 0.09197 | 0.01698 | 0.0703 | 0.09747 | -0.03516 | 0.088533 | -0.17112 | -0.01065 | 0.057871 | -0.72446 | -0.63648 | -0.04816 | -0.03951 | -0.07254 | -0.04579 | -0.06653 | 0.02795 |
| PEO4 | 0.017634 | 0.181844 | -0.11319 | 0.0211 | -0.11781 | -0.01565 | 0.000128 | 0.07807 | -0.04791 | 0.04413 | 0.03157 | -0.01972 | 0.149962 | 0.217754 | -0.11104 | 0.046674 | -0.12566 | -0.52272 | -0.36045 | -0.31172 | 0.186541 | 0.10904 | 0.04613 | 0.09648 | 0.17983 |
| PGA1 | -0.01763 | 0.062927 | -0.04729 | -0.04568 | 0.08392 | -0.14771 | 0.04413 | -0.22535 | 0.02775 | -0.01072 | 0.09498 | 0.06145 | -0.02335 | -0.00371 | -0.17434 | 0.053106 | 0.057469 | -0.08576 | -0.19446 | 0.1419 | -0.0218 | 0.01276 | 0.08747 | 0.0564 | 0.07596 |
| ROS50 | -0.01632 | -0.1545 | -0.11073 | -0.17254 | -0.19885 | 0.26344 | -0.13874 | -0.52952 | 0.01676 | -0.03255 | 0.25503 | -0.01962 | -0.08062 | -0.03971 | -0.18033 | -0.24979 | 0.235517 | -0.08892 | -0.31242 | 0.085418 | -0.01065 | 0.08375 | -0.41282 | -0.3863 | 0.13926 |
| SAT | 0.060381 | 0.037749 | -0.16482 | 0.05607 | -0.19678 | -1.03119 | -0.03093 | -0.10614 | -0.14959 | -0.05058 | 0.13877 | -0.06961 | -0.14663 | 0.099106 | -0.2206 | -0.04606 | 0.101317 | -0.76912 | -0.69465 | -0.04452 | -0.21077 | 0.09127 | 0.01563 | 0.04076 | 0.1205 |
| SEKI | 0.067339 | 0.135095 | -0.08265 | -0.11966 | -0.04139 | -0.08362 | -0.26098 | -0.20562 | 0.04885 | 0.00413 | 0.08859 | 0.1027 | -0.08694 | -0.09741 | -0.22279 | 0.095679 | 0.058571 | -0.1227 | -0.2214 | -0.03354 | 0.001321 | 0.01368 | 0.07996 | -0.12056 | 0.08396 |
| SHI1 | -0.10006 | 0.045236 | -0.08324 | -0.1277 | 0.17217 | 0.0143 | -0.07956 | -0.31925 | -0.00344 | 0.11152 | 0.09844 | 0.23641 | 0.083749 | -0.01183 | -0.11978 | -0.11848 | -0.00849 | -0.33468 | -0.16066 | -0.04813 | 0.039051 | -0.05876 | -0.02038 | -0.09267 | 0.12337 |
| SHMAC4 | 0.055045 | -0.01783 | -0.08447 | -0.0614 | -0.01301 | -0.98851 | -0.02089 | 0.03029 | 0.01056 | 0.0545 | 0.09923 | 0.09481 | 0.019611 | -0.01259 | -0.01198 | -0.02374 | 0.030366 | -0.22327 | -0.21408 | -0.05117 | -0.08772 | -0.05157 | 0.04134 | 0.01703 | 0.06579 |
| SHMAC5 | 0.081651 | -0.05972 | -0.07983 | 0.06208 | -0.04623 | -1.15693 | -0.19664 | -0.067 | -0.12572 | -0.09643 | 0.08446 | 0.00225 | -0.0346 | 0.04305 | -0.23277 | 0.137702 | 0.142036 | -0.12619 | -0.41398 | 0.123068 | -0.02282 | -0.03865 | 0.1083 | 0.1332 | 0.26193 |
| SISO | 0.001424 | 0.020326 | 0.01482 | 0.05159 | -0.01532 | -0.69511 | -0.023 | 0.0137 | -0.02604 | 0.05888 | 0.01869 | -0.01359 | 0.003455 | 0.153246 | -0.0881 | 0.054174 | 0.074527 | -0.28684 | -0.31667 | -0.12372 | -0.07223 | -0.09304 | 0.25135 | 0.03161 | 0.01852 |
| SKGI | -0.05079 | 0.059565 | -0.16581 | -0.02638 | -0.06416 | -0.37933 | 0.053073 | -0.03468 | -0.11011 | 0.07318 | 0.01289 | 0.08593 | -0.11961 | 0.16459 | -0.2678 | 0.124338 | 0.022467 | -0.19669 | -0.23279 | -0.04502 | -0.02605 | -0.20211 | 0.096 | -0.10494 | 0.09507 |
| SKGII | 0.093057 | -0.0876 | -0.03682 | -0.00854 | -0.18854 | -0.44329 | 0.047237 | 0.02435 | 0.07341 | 0.01094 | 0.03448 | -0.02524 | 0.095451 | 0.098278 | -0.11387 | -0.02267 | -0.003 | -0.56711 | -0.55624 | 0.01405 | -0.12154 | -0.10882 | 0.14954 | 0.04933 | 0.15928 |
| SKGT2 | -0.05276 | 0.041704 | -0.03303 | -0.06527 | -0.02286 | -0.17185 | 0.092074 | 0.07214 | -0.14696 | 0.00106 | 0.16856 | 0.13857 | -0.0307 | 0.035643 | 0.034782 | 0.118804 | 0.185485 | -0.64395 | -0.29162 | -0.15609 | -0.03272 | 0.04152 | 0.0919 | -0.02387 | 0.09586 |
| SKGT4 | -0.09567 | -0.0405 | -0.05129 | -0.127 | 0.02822 | -1.23862 | -0.05163 | -0.03042 | 0.04929 | 0.03535 | 0.17458 | 0.09317 | -0.01847 | -0.01986 | -0.09111 | -0.08899 | 0.006332 | -0.4394 | -0.73253 | -0.08114 | 0.020098 | -0.14027 | 0.00104 | -0.08327 | 0.19535 |
| SKN | -0.1737 | -0.20522 | 0.07163 | 0.14774 | -0.10775 | 0.00666 | 0.053867 | -0.09128 | 0.02453 | 0.02105 | 0.11238 | 0.15877 | 0.072181 | 0.045907 | -0.20333 | -0.00172 | 0.08947 | -0.54645 | -0.08382 | -0.29941 | -0.3672 | -0.2102 | 0.02498 | 0.10367 | 0.22624 |
| SKNO1 | 0.091725 | 0.085582 | -0.02579 | -0.02102 | -0.05284 | -0.2327 | -0.04766 | -0.28671 | -0.03636 | 0.10133 | 0.16342 | 0.00352 | -0.06644 | -0.08932 | -0.11188 | -0.19074 | 0.128226 | -0.29635 | -0.17899 | -0.02559 | -0.06968 | 0.05212 | -0.05123 | -0.06241 | 0.21932 |
| SNU638 | 0.064735 | -0.05105 | -0.04785 | -0.05327 | -0.10066 | -0.23658 | 0.117644 | -0.14506 | 0.03605 | 0.06025 | 0.04519 | -0.06159 | 0.017656 | 0.039302 | -0.19391 | -0.02716 | 0.096579 | -0.13393 | -0.18791 | -0.0723 | -0.05524 | -0.04719 | -0.07939 | 0.08802 | 0.11208 |
| SUSA | 0.012695 | -0.05013 | -0.01762 | -0.00625 | -0.14522 | -0.2494 | 0.188076 | 0.13778 | -0.14507 | -0.09678 | 0.02731 | -0.09161 | 0.156436 | 0.43345 | 0.090902 | -0.08738 | 0.252561 | -0.21103 | -0.19248 | -0.08215 | -0.20787 | 0.10343 | -0.02818 | 0.03653 | -0.00721 |
| TANOUE | 0.13256 | 0.060908 | -0.02241 | 0.11331 | -0.08072 | -0.09619 | -0.05372 | -0.13495 | -0.00333 | -0.0093 | 0.08052 | 0.02002 | 0.003301 | -0.02959 | -0.17213 | -0.03767 | -0.01538 | -0.21133 | -0.1915 | -0.03675 | 0.032673 | -0.01728 | 0.02124 | -0.00064 | 0.16349 |
| TASK1 | -0.14096 | 0.00242 | -0.0276 | 0.13238 | 0.03617 | -0.07659 | -0.14311 | -0.11981 | 0.01061 | 0.00909 | 0.11603 | 0.00501 | -0.09567 | 0.014866 | 0.035168 | -0.02373 | 0.024381 | -0.27347 | -0.16524 | -0.16433 | -0.08488 | 0.04109 | -0.11998 | 0.26858 | 0.04196 |

| Cell line | ALK | AXL | CSF1R | DDR1 | DDR2 | EGFR | EPHA1 | EPHA2 | EPHA3 | EPHA4 | EPHA6 | EPHA7 | EPHB1 | EPHB2 | EPHB3 | EPHB4 | EPHB6 | ERBB2 | ERBB3 | ERBB4 | FGFR1 | FGFR2 | FGFR3 | FGFR4 | FLT1 |
|---|---|---|---|---|---|---|---|---|---|---|---|---|---|---|---|---|---|---|---|---|---|---|---|---|---|
| TFK1 | 0.10666 | 0.034663 | -0.09191 | 0.07649 | -0.04664 | -0.54732 | -0.03239 | -0.05378 | -0.0126 | 0.06738 | 0.15031 | 0.17873 | 0.068626 | 0.071452 | -0.03025 | 0.000295 | 0.141429 | -0.2075 | -0.1561 | -0.04006 | 0.034082 | -0.08573 | 0.02806 | 0.02162 | 0.03783 |
| TGW | -0.33416 | 0.15727 | -0.03018 | 0.09518 | -0.11931 | -0.09636 | -0.15718 | -0.10008 | 0.05843 | 0.11147 | 0.1455 | 0.03364 | 0.009475 | 0.036856 | -0.08205 | -0.09517 | 0.129434 | -0.31433 | -0.28273 | -0.11024 | -0.02065 | 0.09772 | 0.00179 | 0.07983 | 0.18545 |
| TR146 | 0.048177 | 0.102835 | -0.05707 | 0.02498 | -0.10362 | -0.76262 | -0.07071 | 0.01151 | 0.03002 | 0.0629 | 0.11996 | -0.18942 | -0.06732 | 0.001959 | -0.21416 | -0.00794 | 0.16875 | -0.44113 | -0.31652 | -0.05571 | -0.02795 | 0.08421 | 0.0444 | -0.09138 | 0.04763 |
| U2904 | -0.01138 | -0.01311 | 0.02385 | 0.20119 | 0.14262 | 0.10835 | 0.1485 | -0.05091 | -0.07148 | 0.07423 | -0.09463 | 0.10768 | -0.07032 | 0.06562 | -0.05372 | 0.178237 | 0.130319 | -0.28575 | -0.2312 | -0.18912 | 0.02651 | 0.06309 | 0.04662 | 0.21672 | 0.09695 |
| UHO1 | -0.16771 | 0.135982 | 0.02732 | 0.0992 | -0.07429 | 0.04932 | 0.055192 | -0.15465 | -0.0356 | 0.1149 | 0.10915 | 0.13704 | 0.054942 | 0.012881 | -0.12102 | 0.058602 | 0.049189 | -0.24661 | -0.16982 | -0.00695 | -0.04284 | -0.10081 | 0.02509 | -0.01097 | 0.09494 |
| UMRC3 | -0.09144 | 0.029135 | -0.01707 | -0.15988 | -0.04838 | -0.38233 | -0.11722 | -0.02115 | 0.0622 | -0.03659 | 0.25663 | 0.06084 | 0.140746 | 0.046776 | -0.01797 | -0.12539 | 0.006949 | -0.16775 | -0.18901 | 0.187933 | -1.04041 | 0.15696 | -0.07579 | -0.16469 | 0.47025 |
| UMRC7 | 0.090147 | -0.54059 | -0.06513 | 0.22003 | -0.14564 | -0.38718 | -0.16563 | 0.77305 | -0.0533 | 0.08456 | 0.27416 | 0.01852 | 0.115444 | 0.085256 | -0.2675 | 0.001075 | -0.05754 | -0.1367 | -0.15701 | -0.09706 | -0.31787 | 0.22148 | 0.04503 | -0.11972 | 0.11808 |
| UPCISCC026 | 0.149922 | 0.135164 | 0.01111 | -0.10221 | 0.01494 | -0.73312 | -0.00404 | 0.22549 | -0.03391 | -0.00822 | 0.1601 | -0.04268 | -0.00301 | 0.233933 | -0.09719 | -0.00221 | 0.067611 | -0.39838 | -0.37978 | -0.04705 | 0.016243 | 0.07002 | 0.01665 | -0.02858 | 0.12439 |
| UPCISCC029A | 0.066894 | -0.17349 | 0.02398 | 0.08916 | 0.00351 | -0.06626 | -0.03962 | 0.21228 | 0.02908 | 0.01472 | 0.16908 | 0.07242 | -0.04552 | -0.01225 | -0.12001 | -0.10041 | 0.021623 | -0.32978 | -0.32773 | -0.09325 | -1.74038 | -0.07745 | 0.05087 | -0.02771 | 0.2129 |
| UPCISCC040 | 0.067956 | 0.071217 | -0.02241 | -0.02716 | -0.00158 | -0.39958 | -0.03085 | 0.03515 | -0.1098 | 0.03408 | 0.12054 | 0.03977 | -0.00066 | 0.117471 | -0.1183 | 0.012084 | 0.11983 | -0.64629 | -0.65109 | -0.04021 | -0.09079 | -0.09286 | 0.02142 | -0.05779 | 0.12825 |
| UPCISCC074 | 0.153698 | 0.012022 | -0.04929 | 0.04619 | 0.06469 | -1.26705 | -0.06548 | 0.00135 | -0.00502 | 0.09849 | 0.12697 | 0.07277 | -0.01004 | 0.108872 | -0.13824 | -0.01905 | 0.06401 | -0.52675 | -0.51858 | 0.039959 | -0.02812 | 0.04387 | -0.08732 | -0.09503 | 0.09993 |
| UPCISCC111 | 0.155587 | 0.00596 | 0.00666 | 0.03293 | -0.01666 | -0.57943 | 0.058223 | 0.38808 | -0.08919 | 0.04908 | 0.19165 | -0.05437 | -0.04273 | 0.012688 | -0.05015 | 0.047349 | 0.090354 | -0.84843 | -0.77619 | -0.05968 | -0.23664 | 0.02327 | 0.02982 | -0.02557 | 0.05948 |
| UPCISCC116 | 0.016388 | -0.17226 | -0.01128 | 0.01914 | 0.10069 | -0.86151 | -0.11125 | -0.06644 | -0.16909 | 0.23138 | 0.29683 | 0.09335 | 0.066055 | 0.252294 | 0.255514 | -0.09209 | 0.237774 | -0.61928 | -0.65592 | -0.08832 | -0.41471 | 0.00496 | 0.07751 | 0.09322 | -0.01933 |
| UPCISCC131 | 0.214835 | -0.02561 | -0.01829 | -0.07271 | -0.13972 | -0.51315 | 0.225099 | -0.01314 | -0.02125 | -0.10382 | 0.04499 | -0.04955 | 0.057788 | -0.00779 | -0.08263 | 0.020044 | 0.075984 | -0.64179 | -0.48922 | -0.01968 | -0.06858 | 0.02667 | -0.0257 | -0.06723 | 0.29676 |
| UPCISCC200 | 0.027087 | 0.108496 | -0.08794 | 0.04448 | -0.23906 | -0.44647 | -0.05653 | 0.33853 | -0.08043 | 0.09287 | 0.09373 | -0.05129 | 0.090143 | 0.136254 | -0.17985 | 0.203945 | -0.03451 | -1.10309 | -0.54419 | 0.041882 | -0.02743 | 0.09117 | -0.06612 | 0.0522 | 0.25402 |
| VAESBJ | 0.024291 | 0.133771 | -0.17704 | -0.12088 | -0.11616 | -0.13365 | -0.13033 | -0.04138 | -0.05113 | 0.15266 | 0.22174 | 0.02808 | -0.03548 | 0.118542 | -0.29189 | 0.111797 | -0.04082 | -0.28384 | -0.41537 | 0.074877 | -0.06335 | -0.13591 | 0.0209 | 0.03955 | 0.44527 |
| WAOSEL | 0.29478 | 0.143428 | -0.15258 | 0.2174 | -0.00237 | -0.21971 | -0.14212 | -0.29391 | -0.01238 | 0.09469 | 0.25908 | 0.19959 | 0.040933 | 0.096092 | -0.1086 | 0.083453 | 0.093387 | -0.29928 | -0.14087 | 0.20275 | 0.063974 | 0.0358 | -0.1009 | -0.2767 | 0.42504 |
| WSUNHL | 0.149024 | -0.04067 | 0.11103 | 0.15253 | -0.06252 | -0.18801 | 0.077648 | -0.17665 | -0.08249 | 0.06435 | 0.17439 | 0.1459 | 0.023203 | 0.022883 | -0.15729 | 0.099903 | -0.04223 | -0.29675 | -0.22694 | -0.08882 | 0.034984 | -0.01975 | 0.02986 | 0.09297 | 0.07529 |
| PFSK1 | -0.02985 | -0.20557 | -0.22632 | 0.25663 | 0.23163 | -0.24381 | -0.04976 | 0.13154 | -0.31813 | 0.30847 | 0.31682 | -0.03375 | 0.166565 | 0.188643 | -0.0381 | 0.016275 | -0.15304 | -0.66598 | -0.04919 | -0.09177 | -0.02988 | 0.17089 | 0.02019 | 0.16765 | 0.19138 |
| CAL72 | -0.35256 | 0.112751 | 0.16952 | 0.04563 | -0.19343 | 0.1346 | 0.048531 | -0.02256 | -0.24493 | 0.21579 | -0.02089 | 0.08619 | 0.083007 | 0.233225 | 0.070179 | 0.028189 | 0.060563 | -0.18974 | -0.1213 | -0.17303 | -1.49579 | 0.22262 | 0.07072 | 0.05328 | 0.07305 |
| OCIC4P | 0.056527 | 7.84E-05 | 0.01699 | 0.02588 | -0.14275 | -0.27898 | -0.09851 | -0.06693 | -0.03094 | 0.0349 | 0.03373 | 0.02132 | -0.03957 | 0.103161 | -0.13305 | -0.07583 | -0.04966 | -0.391 | -0.34392 | -0.02009 | -0.21155 | 0.00761 | 0.02752 | -0.02136 | 0.13868 |
| SEMK2 | 0.030412 | -0.10628 | -0.04652 | 0.00377 | -0.00883 | -0.14724 | -0.07965 | -0.15381 | 0.02179 | 0.01139 | 0.14918 | 0.1317 | -0.08183 | 0.039279 | -0.17226 | 0.116521 | 0.124715 | -0.14429 | -0.23678 | -0.06399 | 0.004926 | -0.0518 | 0.08907 | -0.18498 | 0.10481 |
| HB1119 | 0.0251 | -0.13464 | 0.07557 | 0.02492 | 0.04596 | -0.19248 | -0.0789 | -0.00912 | -0.00195 | 0.0184 | 0.13486 | 0.07865 | -0.09721 | -0.03861 | 0.028246 | 0.035203 | 0.115641 | -0.07915 | -0.07809 | -0.17243 | -0.06956 | 0.04681 | 0.00933 | -0.00741 | -0.10589 |
| CTV1DM | -0.07766 | 0.117247 | 0.20429 | 0.06415 | 0.03801 | 0.02867 | 0.396387 | -0.23731 | -0.09993 | 0.03961 | 0.01108 | 0.22618 | -0.1888 | -0.10579 | 0.416484 | 0.151477 | 0.187443 | -0.19699 | 0.07437 | -0.08746 | 0.246614 | 0.17391 | 0.12455 | 0.1007 | -0.04959 |
| RH28 | -0.11623 | 0.082709 | 0.20455 | -0.12873 | 0.00197 | -0.20396 | 0.098768 | -0.12934 | 0.05241 | 0.1738 | 0.01892 | 0.11183 | -0.01545 | 0.018362 | -0.11772 | 0.100639 | 0.198723 | -0.35007 | -0.13546 | -0.43974 | 0.170331 | 0.00229 | -0.11694 | -0.206 | 0.08978 |
| RHJT | -0.02592 | -0.03404 | 0.07689 | -0.13386 | 0.03725 | -0.08942 | 0.052986 | -0.06679 | 0.07613 | 0.11834 | 0.08466 | 0.0688 | 0.203407 | 0.133292 | -0.06792 | 0.186404 | -0.04428 | -0.34269 | -0.14946 | -0.2519 | 0.199684 | -0.064 | -0.09732 | -0.32779 | 0.20568 |
| TTC442 | -0.0007 | -0.23696 | -0.05171 | -0.13721 | 0.05862 | -0.24837 | -0.15808 | -0.06443 | 0.0177 | 0.10279 | 0.1424 | 0.2146 | 0.027555 | 0.142226 | -0.05043 | -0.05309 | 0.043494 | -0.50698 | -0.56002 | -0.27624 | -0.11366 | -0.08285 | 0.04549 | -0.06182 | 0.12126 |
| RH4 | -0.02438 | 0.084936 | 0.07201 | -0.13041 | -0.03117 | -0.10245 | -0.01552 | -0.07354 | 0.05169 | -0.04815 | 0.08529 | 0.05562 | 0.079826 | 0.042001 | 0.073473 | 0.048752 | 0.183334 | -0.14979 | -0.15694 | -0.10709 | -0.04029 | -0.17592 | -0.02404 | -0.24694 | 0.13783 |
| SNU1544 | -0.30433 | 0.219002 | 0.1277 | 0.03132 | -0.33413 | -0.60893 | 0.05373 | 0.05052 | 0.00994 | 0.14332 | 0.19482 | 0.04505 | -0.21027 | 0.109781 | 0.105538 | 0.11664 | 0.189406 | -0.64084 | -0.23617 | -0.13189 | -0.12394 | 0.0964 | 0.11502 | 0.14308 | -0.17036 |
| LPS6 | -0.21517 | -0.13396 | 0.04535 | 0.00068 | -0.13866 | -0.17933 | 0.256683 | -0.0631 | -0.10055 | -0.24369 | 0.17358 | 0.06425 | 0.216033 | 0.04633 | 0.052527 | 0.025021 | 0.143261 | -0.58494 | -0.36905 | 0.05024 | -0.74997 | 0.04775 | 0.00424 | -0.0599 | -0.17352 |
| LPS27 | 0.005257 | 0.018307 | -0.03054 | -0.10283 | -0.02247 | -0.06666 | -0.01721 | -0.00876 | -0.11201 | 0.00126 | 0.14781 | 0.13674 | 0.06914 | 0.151304 | -0.13439 | 0.098661 | 0.01969 | -0.17873 | -0.38878 | -0.10432 | -0.99596 | 0.02111 | 0.04705 | -0.2106 | 0.10913 |
| 93T449 | 0.10309 | -0.27887 | -0.03132 | -0.09656 | 0.03679 | -0.19091 | -0.09253 | 0.04145 | 0.04719 | 0.05512 | 0.114 | 0.0662 | 0.138108 | 0.140154 | -0.10958 | -0.10172 | -0.05566 | -0.2669 | -0.28966 | -0.05104 | -1.54447 | -0.07572 | -0.11215 | 0.0854 | 0.1476 |
| 94T778 | 0.033667 | -0.351 | -0.04749 | -0.12598 | -0.05409 | -0.32925 | 0.041721 | 0.41115 | 0.11915 | 0.03892 | 0.10885 | -0.10688 | 0.078181 | 0.083373 | -0.07718 | -0.0083 | 0.037731 | -0.08383 | -0.19975 | -0.09136 | -0.87606 | -0.33512 | 0.04698 | -0.05668 | 0.05195 |
| 95T1000 | 0.047916 | -0.44272 | 0.13138 | -0.06631 | -0.02999 | -0.15958 | -0.2987 | 0.08939 | -0.00333 | 0.06374 | 0.27178 | 0.04856 | -0.00031 | 0.063168 | -0.15493 | -0.00499 | -0.02278 | -0.28411 | -0.184 | -0.04856 | -1.20707 | -0.10243 | 0.02492 | 0.09977 | 0.04099 |
| LPS141 | -0.13046 | 0.111925 | 0.10783 | 0.02786 | 0.05802 | -0.07306 | 0.187091 | 0.04196 | -0.22484 | 0.06098 | 0.1404 | -0.05576 | -0.07575 | 0.112108 | -0.02058 | 0.172486 | 0.242682 | -0.42759 | -0.24466 | -0.19356 | -0.23311 | 0.11025 | 0.07277 | -0.13476 | 0.00606 |
| LPS853 | 0.123402 | 0.068659 | 0.13038 | -0.04394 | 0.03994 | -0.21732 | 0.002335 | -0.06301 | -0.02936 | 0.1357 | 0.0505 | 0.05602 | 0.064233 | 0.091075 | -0.20397 | -0.00714 | 0.064293 | -0.15361 | -0.19083 | -0.07067 | -0.17365 | 0.00696 | 0.00967 | -0.06207 | 0.09317 |
| LPS510 | 0.058453 | -0.28654 | -0.10732 | -0.22539 | -0.19407 | -0.22056 | 0.027154 | -0.19191 | 0.08195 | -0.08314 | 0.31527 | -0.07208 | -0.06369 | 0.104703 | 0.200775 | 0.081723 | -0.08485 | -0.30154 | -0.04563 | -0.34494 | -1.20279 | 0.14024 | 0.14179 | -0.20341 | 0.28019 |
| OS252 | -0.05067 | -0.12638 | 0.13154 | 0.1336 | -0.06587 | -0.05023 | 0.069774 | -0.16705 | 0.06003 | 0.19118 | 0.21811 | 0.22952 | 0.059771 | 0.025549 | 0.079162 | -0.05797 | 0.109029 | -0.42961 | -0.21366 | -0.22784 | -0.53573 | -0.07211 | -0.04528 | 0.05509 | 0.07134 |
| MFM223 | -0.00057 | -0.02085 | -0.0298 | -0.0483 | -0.06213 | -0.19947 | -0.04927 | -0.06576 | 0.01399 | 0.09401 | 0.17896 | 0.00574 | 0.056687 | 0.056611 | -0.13173 | 0.066943 | -0.02176 | -0.17431 | -0.14781 | -0.00257 | 0.087294 | -1.17899 | 0.0509 | -0.11142 | 0.112 |
| COLO824 | 0.00304 | -0.00694 | 0.08259 | 0.12188 | -0.08672 | -0.10114 | 0.036811 | 0.08052 | 0.15323 | 0.24012 | 0.01205 | 0.26033 | -0.06967 | 0.182932 | 0.228399 | 0.0934 | 0.095157 | -0.4864 | -0.34693 | -0.09767 | 0.038077 | 0.02235 | 0.14757 | -0.23168 | 0.00883 |
| ICC10 | 0.127712 | 0.144938 | -0.1881 | 0.01389 | 0.00753 | -0.76192 | 0.022407 | -0.01238 | 0.0463 | 0.0186 | 0.12017 | 0.10748 | -0.02239 | 0.085123 | -0.13679 | 0.075626 | 0.11926 | -0.095 | -0.28385 | 0.034041 | -0.05138 | -0.42229 | 0.04698 | -0.13427 | 0.12441 |
| ICC106 | 0.007015 | -0.00464 | -0.1114 | -0.08122 | -0.06232 | -0.64882 | 0.044619 | 0.01155 | 0.00951 | -0.02429 | 0.14974 | 0.1758 | 0.029189 | 0.087681 | -0.14455 | -0.02184 | 0.214922 | -0.29095 | -0.22541 | 0.012014 | -0.02677 | -0.45862 | 0.13146 | 0.11622 | 0.22639 |
| ICC108 | 0.014457 | 0.1413 | 0.11039 | 0.00909 | -0.04346 | -1.8253 | 0.150012 | 0.15442 | 0.04348 | 0.07746 | 0.11951 | 0.05729 | 0.00868 | 0.080169 | -0.29515 | 0.019678 | 0.172936 | -0.39607 | -0.20746 | -0.02957 | -0.07454 | -0.02853 | 0.07539 | 0.04428 | 0.06833 |
| ICC12 | 0.056048 | 0.036016 | -0.00549 | -0.11258 | -0.05274 | -0.2034 | 0.205535 | 0.257 | 0.03776 | 0.02795 | 0.26241 | 0.02136 | 0.199437 | 0.051925 | -0.31058 | 0.010341 | 0.20228 | -0.32864 | -0.12676 | -0.19379 | -0.09543 | -0.00589 | -0.04186 | 0.13375 | 0.19938 |
| ICC137 | 0.151066 | -0.13986 | -0.11487 | 0.06771 | -0.07454 | -0.18953 | 0.020791 | -0.00792 | -0.20478 | 0.06302 | 0.0302 | 0.06773 | -0.02875 | 0.004728 | -0.13341 | -0.12385 | 0.135834 | -0.48393 | -0.54127 | -0.1077 | -0.08977 | -0.69901 | -0.12056 | 0.19953 | 0.04203 |
| ICC15 | -0.01928 | 0.094796 | -0.0568 | -0.13591 | -0.02226 | -0.96851 | -0.04225 | -0.04904 | -0.06422 | 0.00157 | 0.10322 | 0.09169 | -0.00299 | 0.063526 | -0.07651 | -0.04504 | 0.00423 | -0.33718 | -0.29401 | -0.11477 | -0.04719 | -0.092 | 0.06715 | -0.0682 | 0.12932 |
| ICC2 | -0.30227 | -0.09048 | 0.19294 | 0.14678 | 0.09805 | -0.60336 | 0.167213 | 0.13463 | -0.14927 | -0.24188 | 0.08635 | 0.05052 | 0.011458 | -0.15785 | 0.251922 | 0.088423 | 0.049312 | -0.56352 | -0.29802 | -0.55943 | 0.058995 | 0.0419 | -0.09058 | 0.03269 | -0.1947 |
| ICC3 | -0.0883 | 0.003748 | -0.01543 | 0.1127 | 0.11492 | -0.22264 | -0.11649 | 0.38574 | 0.18738 | 0.00766 | 0.011 | -0.04118 | 0.140458 | 0.128744 | -0.14704 | 0.309578 | 0.055754 | -0.39012 | -0.22576 | 0.031076 | -0.2418 | -0.02206 | -0.00423 | -0.06877 | 0.12631 |
| ICC4 | 0.096163 | 0.074851 | 0.03842 | 0.12042 | -0.048 | -0.8514 | 0.000808 | 0.10021 | 0.00788 | -0.08549 | 0.33355 | 0.08653 | 0.013988 | 0.033216 | -0.0843 | -0.07443 | 0.076827 | -0.40913 | -0.32235 | -0.03079 | 0.127311 | 0.08554 | 0.14096 | 0.1147 | 0.19946 |
| ICC8 | 0.056493 | -0.01676 | -0.07755 | -0.00925 | -0.01018 | -0.11464 | 0.119488 | 0.14404 | -0.02402 | 0.03293 | 0.17862 | 0.12236 | 0.062197 | 0.122442 | -0.14252 | 0.081932 | 0.146097 | -0.13448 | -0.2325 | -0.01784 | 0.013293 | -0.04784 | -0.11885 | -0.06042 | 0.19087 |
| ICC9 | 0.08254 | -0.0541 | -0.23648 | 0.02972 | -0.12208 | -0.71557 | 0.083622 | 0.18846 | -0.08315 | 0.00939 | 0.17402 | 0.10747 | 0.063622 | 0.105032 | -0.03522 | 0.112039 | -0.05457 | -0.52402 | -0.26079 | 0.157617 | -0.00655 | -0.17863 | -0.0014 | 0.02609 | 0.29762 |
| G415 | 0.076877 | -0.14644 | -0.08688 | -0.08371 | -0.05073 | -0.21432 | -0.01543 | 0.06438 | -0.00854 | 0.14562 | 0.23881 | -0.1085 | 0.031742 | 0.045635 | -0.0719 | 0.013505 | -0.01425 | -0.17403 | -0.25557 | -0.03271 | -0.10793 | -0.03939 | -0.00651 | -0.08007 | 0.22238 |
| HKGZCC | 0.021023 | -0.00396 | 0.00635 | 0.00945 | -0.02662 | -0.89075 | 0.064658 | 0.02877 | -0.03501 | -0.00255 | 0.07563 | 0.05241 | 0.080639 | 0.102213 | -0.02889 | 0.073518 | 0.074602 | -0.57947 | -0.57945 | -0.19624 | 0.043429 | -0.04687 | -0.02815 | -0.00815 | 0.10711 |
| KMCH1 | -0.07662 | -0.07056 | -0.24602 | -0.08505 | -0.15446 | -1.34887 | 0.098224 | -0.04553 | 0.00771 | 0.23854 | 0.24694 | 0.09432 | 0.114512 | 0.0965 | -0.17558 | 0.092794 | 0.03443 | -0.2965 | -0.22577 | -0.07277 | -0.12664 | -0.13543 | -0.05287 | -0.07289 | 0.00699 |
| RBE | -0.07855 | -0.0132 | -0.03905 | -0.21336 | 0.06517 | -0.5914 | 0.11568 | 0.24973 | 0.01497 | 0.15842 | 0.15249 | 0.14475 | 0.06105 | 0.052873 | -0.03238 | -0.04861 | 0.088316 | -0.32002 | -0.48643 | 0.112248 | -0.02188 | -0.06639 | 0.11857 | 0.03668 | 0.17166 |
| SG231 | -0.17691 | -0.05261 | -0.14329 | 0.01069 | -0.04153 | -0.1279 | -0.0479 | -0.10719 | 0.047 | -0.20207 | 0.15035 | 0.13946 | 0.187697 | 0.105356 | 0.114977 | 0.187875 | 0.010178 | 0.08881 | -0.28806 | -0.16797 | -0.04354 | -0.08647 | 0.17699 | 0.08468 | 0.151 |
| SSP25 | -0.00327 | -0.09423 | -0.035 | -0.06527 | -0.0386 | -0.4805 | 0.040479 | 0.31297 | -0.04991 | 0.08889 | 0.12761 | 0.06286 | 0.001422 | 0.118358 | -0.17224 | 0.040446 | 0.142451 | -0.86999 | -0.7843 | -0.09671 | -0.15078 | -0.0477 | 0.04955 | 0.00982 | 0.14014 |
| TGBC1TKB | 0.265795 | -0.00121 | 0.01712 | 0.09004 | -0.01269 | -0.94251 | 0.011931 | 0.1373 | 0.02897 | 0.08539 | 0.22555 | -0.06547 | 0.062152 | 0.19058 | 0.001287 | 0.212196 | -0.05202 | -0.34054 | -0.24996 | -0.33552 | -0.09913 | -0.03777 | 0.07563 | -0.14009 | -0.0822 |
| TGBC52TKB | 0.09076 | -0.23017 | 0.0291 | -0.07604 | -0.0014 | -0.62941 | -0.08465 | -0.03423 | -0.15795 | 0.08918 | 0.07541 | 0.0273 | -0.01568 | -0.12007 | -0.03647 | 0.077975 | 0.079445 | -0.26937 | -0.33928 | -0.13646 | -0.11344 | -0.05441 | 0.24866 | 0.06043 | 0.37244 |
| TKKK | -0.07627 | 0.075332 | -0.04656 | 0.15624 | -0.2346 | -0.61723 | 0.004969 | 0.06789 | -0.10545 | -0.05185 | 0.02964 | -0.03041 | 0.10035 | -0.00844 | -0.07352 | -0.14569 | 0.107515 | -0.93798 | -0.2716 | -0.32895 | -0.14219 | 0.1507 | 0.04628 | -0.26715 | 0.20549 |
| YSCCC | -0.21359 | 0.051278 | 0.0192 | -0.32184 | -0.0554 | -0.34314 | 0.163921 | -0.0511 | -0.2501 | 0.08399 | -0.04012 | -0.05241 | -0.02231 | 0.120019 | 0.001592 | 0.090585 | 0.167699 | -0.27331 | -0.41116 | -0.14183 | -0.14072 | -0.00295 | 0.23045 | 0.0895 | 0.09812 |
| CCLP1 | 0.043542 | -0.06694 | -0.00108 | 0.15596 | -0.0795 | -0.08872 | 0.280772 | -0.02201 | -0.00887 | 0.09533 | -0.01825 | 0.0212 | 0.005681 | 0.18848 | -0.08895 | -0.05028 | -0.04024 | -0.14927 | -0.13369 | -0.12867 | -1.25587 | -0.12941 | 0.16622 | 0.08209 | 0.10669 |
| CCSW1 | 0.0233 | 0.026191 | -0.10912 | -0.00054 | -0.01628 | -0.36488 | -0.00513 | 0.24213 | 0.03604 | 0.15273 | 0.05072 | 0.09834 | 0.027028 | 0.077927 | -0.00139 | 0.085534 | -0.06109 | -0.92357 | -0.75612 | -0.05647 | -0.05038 | -0.11111 | 0.10538 | 0.09773 | 0.18917 |
| GB2 | 0.110586 | 0.016827 | -0.12179 | 0.04009 | -0.02998 | -1.12942 | -0.01084 | -0.22152 | 0.0571 | 0.11774 | 0.02862 | 0.15178 | 0.123194 | -0.0244 | -0.15189 | -0.0003 | -0.00614 | -0.23022 | -0.22921 | 0.003543 | -0.32038 | -0.07776 | 0.09082 | -0.1231 | 0.1942 |
| MM253 | -0.04094 | -0.36052 | 0.14795 | 0.21809 | -0.25539 | -0.19803 | -0.29201 | -0.08008 | -0.0834 | 0.07223 | 0.15215 | 0.05035 | -0.06455 | -0.05936 | -0.05557 | -0.20997 | 0.033082 | -0.48629 | -0.3068 | -0.28309 | 0.089614 | 0.04766 | 0.03164 | 0.08498 | 0.2152 |
| MM485 | 0.281188 | -0.29873 | -0.04009 | 0.1847 | 0.04734 | -0.25942 | -0.23842 | 0.03817 | -0.06658 | 0.07123 | 0.30512 | -0.00073 | 0.170472 | -0.0122 | -0.27763 | -0.20803 | -0.19584 | -0.34171 | -0.16928 | 0.114248 | -0.20412 | 0.00586 | 0.09115 | -0.09288 | -0.14121 |
| NO36 | 0.0166 | 0.024872 | -0.10325 | 0.0798 | -0.08542 | -0.38367 | -0.05794 | -0.01484 | -0.00723 | 0.12406 | 0.08037 | 0.2018 | -0.06881 | -0.079 | -0.0577 | -0.08454 | -0.00205 | -0.44932 | -0.3112 | -0.0374 | -0.58969 | 0.08265 | 0.14189 | -0.10522 | 0.09932 |
| NZM3 | 0.045634 | -0.03536 | -0.01303 | 0.04293 | -0.05831 | -0.17956 | -0.05202 | -0.0988 | -0.04688 | 0.00383 | 0.09611 | -0.01252 | -0.04637 | 0.030201 | -0.19564 | -0.04225 | 0.068769 | -0.32409 | -0.22121 | 0.032424 | -0.03124 | 0.06583 | 0.14129 | -0.02678 | 0.0016 |
| NZM42 | 0.005027 | -0.01239 | 0.131 | -0.08481 | -0.00977 | -0.20863 | 0.12881 | -0.11833 | -0.0017 | 0.14237 | 0.12571 | 0.03236 | 0.053726 | 0.082972 | -0.02548 | 0.06664 | 0.187014 | -0.24228 | -0.38955 | 0.031464 | -0.08715 | -0.02368 | 0.01127 | -0.0206 | 0.09195 |
| NZM7 | 0.0253 | -0.01043 | 0.04782 | -0.06218 | -0.00562 | -0.16756 | -0.02162 | -0.09688 | 0.06946 | 0.05357 | 0.12599 | 0.11668 | 0.022373 | -0.02397 | -0.03134 | 0.021931 | -0.04108 | -0.24255 | -0.2742 | -0.09446 | 0.034879 | 0.01038 | -0.10251 | 0.03509 | 0.11078 |
| NZOV9 | -0.0008 | -0.06957 | 0.02964 | 0.09575 | -0.11656 | -0.11709 | -0.06273 | -0.15912 | 0.13845 | -0.14251 | 0.22736 | 0.09057 | -0.0821 | 0.297216 | -0.03173 | -0.03057 | -0.2423 | -0.0989 | -0.16809 | -0.06028 | -0.76621 | -0.12784 | 0.3398 | -0.1043 | 0.23054 |
| ONE58 | 0.177517 | 0.022326 | 0.03012 | -0.04466 | -0.04805 | -0.26158 | -0.01457 | 0.02479 | -0.03784 | 0.11148 | 0.1596 | 0.04725 | -0.01867 | 0.055192 | -0.20269 | -0.06691 | 0.09465 | -0.2856 | -0.38214 | 0.058631 | -1.36462 | -0.01384 | -0.06103 | -0.1144 | 0.04646 |
| NALM16 | -0.01947 | -0.01396 | -0.00567 | 0.04621 | -0.12264 | -0.05828 | -0.23626 | -0.1927 | 0.03492 | 0.13522 | 0.06913 | -0.03335 | -0.08777 | 0.024344 | -0.0145 | -0.00305 | 0.064368 | -0.23324 | -0.07859 | 0.009125 | -0.17039 | 0.22322 | -0.01138 | 0.00718 | 0.08685 |
| ECC2 | 0.041719 | -0.21292 | 0.03482 | 0.00524 | -0.08352 | -0.65629 | 0.082208 | 0.4197 | -0.11358 | 0.01204 | 0.03061 | 0.1576 | 0.032805 | 0.031629 | -0.07855 | -0.01771 | 0.017256 | -0.45855 | -0.17965 | -0.14899 | -0.19528 | -0.08688 | 0.15547 | -0.03111 | 0.09635 |

| Cell line | ALK | AXL | CSF1R | DDR1 | DDR2 | EGFR | EPHA1 | EPHA2 | EPHA3 | EPHA4 | EPHA6 | EPHA7 | EPHB1 | EPHB2 | EPHB3 | EPHB4 | EPHB6 | ERBB2 | ERBB3 | ERBB4 | FGFR1 | FGFR2 | FGFR3 | FGFR4 | FLT1 |
|---|---|---|---|---|---|---|---|---|---|---|---|---|---|---|---|---|---|---|---|---|---|---|---|---|---|
| 9505BIK | -0.01731 | 0.242815 | -0.02336 | 0.15992 | 0.01771 | -0.37241 | 0.087486 | -0.10076 | 0.03095 | 0.15914 | 0.10908 | -0.02377 | 0.034874 | 0.045725 | 0.108962 | 0.179103 | -0.04569 | -0.26912 | -0.17341 | -0.09115 | 0.048161 | -0.19053 | 0.06675 | 0.11869 | -0.02513 |
| A375SKINCJ1 | 0.016702 | -0.002 | -0.06061 | 0.00593 | -0.1298 | -0.11713 | -0.26084 | 0.04143 | 0.0638 | 0.09731 | 0.23475 | -0.02408 | 0.181882 | 0.098307 | -0.15463 | -0.05993 | 0.112525 | -0.19485 | -0.27043 | -0.27003 | 0.132727 | -0.00915 | 0.06864 | 0.05591 | 0.2208 |
| A375SKINCJ2 | 0.127948 | -0.04421 | 0.01956 | 0.08239 | 0.01195 | -0.13666 | -0.03513 | -0.01316 | 0.00506 | -0.08968 | 0.092 | 0.07049 | -0.02174 | 0.086201 | 0.029727 | 0.053064 | -0.01507 | -0.09198 | -0.282 | -0.07532 | 0.037609 | -0.12713 | -0.04103 | 0.03163 | 0.08641 |
| A375SKINCJ3 | -0.20167 | -0.04751 | -0.00537 | 0.03794 | -0.12082 | -0.29233 | -0.17016 | 0.07217 | 0.01885 | 0.12395 | 0.01527 | -0.20712 | 0.149838 | -0.06126 | -0.13107 | 0.047328 | 0.025262 | -0.46302 | -0.04175 | 0.079934 | -0.22345 | -0.02022 | 0.11546 | -0.14989 | 0.08133 |
| SKMEL19 | -0.33009 | -0.06168 | 0.01578 | 0.17166 | -0.0943 | -0.13067 | -0.00336 | 0.06822 | 0.05645 | -0.12293 | 0.2884 | 0.02333 | 0.035028 | 0.071609 | 0.436661 | 0.04363 | 0.083981 | -0.46293 | -0.17417 | -0.16789 | -0.03847 | 0.10957 | 0.2115 | -0.02518 | -0.05404 |
| MEL270 | -0.30667 | 0.130835 | -0.18321 | 0.00586 | -0.37113 | -0.20484 | -0.01798 | 0.13224 | 0.31646 | -0.01742 | -0.16842 | 0.32412 | -0.0301 | 0.05941 | 0.048619 | 0.336627 | -0.00419 | -0.42091 | -0.34646 | -0.2633 | -0.16333 | 0.44912 | -0.20326 | -0.29281 | -0.18261 |
| MEL285 | 0.013906 | -0.03817 | -0.01255 | -0.04319 | -0.01595 | -0.08916 | -0.07316 | -0.06533 | 0.00233 | 0.11657 | 0.06311 | 0.02287 | -0.05978 | 0.24612 | -0.10353 | 0.074694 | -0.16393 | -0.27236 | -0.2041 | -0.033 | -1.35531 | -0.12857 | 0.10867 | -0.12759 | 0.15382 |
| MEL290 | -0.02049 | -0.11231 | -0.00479 | 0.09111 | -0.09096 | -0.35709 | -0.04552 | 0.6681 | -0.06767 | 0.14646 | 0.23845 | 0.14627 | 0.032536 | 0.176512 | -0.10059 | 0.14436 | 0.025457 | -0.33583 | -0.33024 | 0.061236 | -1.37763 | 0.07136 | 0.04308 | 0.03467 | 0.29634 |
| OMM25 | -0.07283 | 0.064118 | -0.00575 | -0.02032 | -0.05823 | -0.12999 | 0.059899 | -0.01283 | 0.01591 | 0.06728 | 0.10687 | -0.00718 | -0.07398 | -0.01529 | -0.06678 | 0.098935 | 0.070659 | -0.26692 | -0.2811 | -0.09011 | -0.01843 | 0.03654 | 0.16293 | 0.04128 | 0.1627 |
| HOKUG | 0.078762 | -0.02323 | -0.05306 | 0.00759 | -0.06178 | -0.86151 | -0.05462 | 0.07822 | 0.02164 | 0.07044 | 0.06951 | 0.04095 | -0.02587 | 0.188574 | -0.05436 | -0.04134 | -0.07465 | -0.24708 | -0.36547 | -0.00466 | -0.15332 | -0.12523 | 0.22446 | -0.05857 | 0.28719 |
| SKGIIIA | 0.038836 | 0.038653 | -0.05028 | 0.10083 | -0.05237 | -0.50324 | -0.13348 | -0.08169 | -0.10064 | -0.00662 | 0.12487 | -0.02372 | 0.006384 | 0.115444 | -0.04807 | -0.03802 | 0.11026 | -0.31676 | -0.36845 | -0.00307 | -0.02745 | 0.06684 | 0.06061 | -0.00978 | 0.26279 |
| T3M3 | -0.06299 | 0.049304 | 0.03673 | 0.00558 | -0.17533 | -0.16418 | -0.19981 | -0.14182 | 0.04272 | 0.14276 | 0.1009 | 0.04304 | 0.021119 | 0.101858 | 0.004982 | 0.063255 | 0.169039 | -0.28604 | -0.20428 | -0.0795 | -0.0106 | 0.1981 | 0.2534 | 0.19841 | 0.13574 |
| TGBC18TKB | 0.113807 | 0.056827 | -0.14789 | 0.03852 | -0.03595 | -1.43643 | 0.012086 | -0.07448 | -0.0624 | 0.02685 | 0.11907 | -0.00247 | 0.016781 | 0.088824 | -0.14608 | -0.01906 | 0.005659 | -1.76281 | -0.2331 | 0.056549 | -0.08029 | -0.07049 | 0.08219 | 0.03549 | 0.19796 |
| ECC4 | 0.040025 | -0.20781 | 0.28065 | 0.40135 | 0.07389 | 0.05777 | -0.20885 | -0.13235 | 0.15603 | -0.02895 | 0.22017 | -0.19584 | -0.2171 | -0.00688 | 0.168292 | -0.19789 | 0.230243 | -0.26345 | -0.07819 | -0.06818 | 0.036586 | 0.02694 | -0.19657 | 0.18817 | 0.35325 |
| TT1TKB | -0.01167 | 0.251161 | 0.12377 | -0.1004 | -0.16321 | -1.15209 | 0.011775 | 0.04219 | -0.03407 | 0.048 | 0.09013 | 0.17476 | -0.01181 | 0.145273 | -0.20934 | 0.031451 | -0.02018 | -0.42325 | -0.29374 | -0.01669 | 0.00273 | 0.12221 | -0.01698 | -0.01653 | 0.21547 |
| HHUA | 0.173386 | 0.064668 | -0.02904 | 0.01911 | -0.05494 | -0.24228 | 0.012018 | 0.17301 | 0.11052 | -0.00444 | 0.05034 | 0.01147 | 0.066333 | 0.109251 | -0.00767 | -0.08675 | -0.02098 | -0.42187 | -0.30521 | -0.01262 | -0.03265 | -0.0314 | 0.04852 | 0.05753 | 0.35219 |
| HOUAI | 0.110229 | 0.157371 | 0.0235 | 0.13555 | -0.01487 | -0.48401 | -0.10486 | -0.15594 | -0.00206 | 0.11659 | 0.09605 | -0.05149 | 0.042394 | 0.186037 | -0.04689 | 0.1973 | 0.093609 | -0.25513 | -0.23779 | -0.05673 | -0.21116 | 0.06383 | 0.08129 | 0.02272 | 0.21823 |
| SAS | 0.111792 | 0.097514 | 0.02744 | 0.05986 | -0.14598 | -1.45516 | -0.08396 | 0.01989 | 0.01155 | 0.00366 | 0.13263 | -0.02696 | -0.01953 | 0.122675 | -0.03353 | 0.016051 | 0.059476 | -0.25569 | -0.24329 | -0.05694 | -0.092 | 0.01533 | -0.0001 | 0.02386 | 0.27046 |
| LCAM1 | -0.06728 | -0.36885 | 0.1369 | 0.0478 | -0.02944 | -0.2041 | -0.19967 | -0.04748 | -0.0974 | -0.03043 | 0.07793 | -0.00329 | -0.11782 | -0.10938 | -0.1163 | 0.050772 | -0.0082 | -0.08682 | -0.18569 | 0.050789 | -0.88333 | 0.06649 | 0.01127 | 0.03874 | 0.0496 |
| PK8 | -0.02213 | 0.004211 | -0.11577 | -0.1294 | -0.15307 | -0.08162 | 0.080704 | 0.14829 | -0.04456 | 0.09148 | 0.27144 | 0.10637 | -0.10698 | 0.036044 | -0.04406 | 0.14275 | -0.08081 | -0.35171 | -0.32971 | 0.091663 | -0.23836 | -0.00152 | 0.18945 | -0.08371 | 0.05364 |
| HOTHC | 0.064982 | -0.08788 | -0.02495 | -0.03702 | -0.00458 | -0.1516 | 0.014367 | -0.02837 | -0.03009 | 0.03451 | 0.09358 | 0.07077 | -0.03998 | 0.081729 | -0.16755 | -0.0412 | 0.071717 | -0.21062 | -0.199 | -0.05205 | -0.17034 | 0.05148 | -0.0022 | -0.0276 | 0.19206 |
| T3M5 | 0.071151 | -0.10342 | -0.01611 | 0.17716 | -0.09325 | -0.29257 | -0.0589 | 0.02881 | -0.07143 | 0.12675 | 0.12344 | 0.01541 | -0.17212 | 0.171046 | 0.037829 | -0.16118 | 0.010073 | -0.41202 | -0.14818 | -0.13638 | 0.018792 | 0.13612 | -0.02265 | -0.06622 | -0.028 |
| CA922 | 0.05122 | -0.01785 | -0.01115 | -0.01888 | -0.05353 | -0.17393 | -0.01312 | -0.06165 | -0.12556 | 0.02996 | 0.08996 | 0.09741 | 0.043436 | 0.098971 | -0.05657 | -0.01152 | 0.092013 | -0.08554 | -0.24053 | -0.05048 | 0.016841 | -0.021 | 0.07517 | -0.00428 | 0.13543 |
| HSQ89 | 0.106887 | -0.02383 | 0.05109 | -0.00705 | 0.02213 | -0.18904 | -0.13672 | -0.10442 | -0.04816 | 0.18143 | 0.07843 | 0.06089 | -0.02432 | -0.02288 | -0.01959 | 0.052431 | 0.077996 | -0.11326 | -0.15656 | 0.052745 | -0.70469 | -0.05422 | 0.17976 | -0.04526 | 0.04332 |
| HO1U1 | 0.206116 | -0.01252 | -0.05888 | -0.04848 | -0.03386 | -0.1693 | 0.164528 | -0.01608 | 0.07965 | 0.17468 | 0.12435 | 0.00849 | 0.113133 | 0.03774 | -0.09855 | 0.011066 | 0.058941 | -0.41211 | -0.41453 | -0.1193 | -0.15967 | 0.10998 | -0.06105 | 0.07744 | 0.17309 |
| HTMMT | 0.028961 | 0.119557 | -0.022 | -0.10515 | -0.09126 | -0.21454 | -0.09991 | -0.03484 | 0.01518 | 0.05048 | 0.12699 | 0.00725 | -0.01748 | 0.054975 | -0.06194 | 0.036129 | -0.01737 | -0.31847 | -0.22962 | -0.06652 | -0.3353 | 0.02984 | 0.00121 | -0.02605 | 0.12978 |
| RMSYM | 0.092991 | 0.070519 | -0.07853 | 0.1328 | -0.00554 | -0.12312 | 0.008025 | -0.12007 | -0.02477 | 0.0993 | 0.18884 | 0.04545 | -0.09575 | -0.04653 | -0.13659 | 0.04919 | 0.122885 | -0.31582 | -0.28922 | -0.04545 | -0.8518 | 0.14476 | 0.16968 | -0.02234 | 0.06409 |
| LU134A | 0.116386 | 0.015595 | -0.05529 | 0.05189 | -0.08672 | -0.12401 | -0.17496 | -0.17274 | -0.12882 | -0.07929 | 0.18805 | 0.16713 | -0.01265 | -0.07465 | 0.08557 | 0.181294 | 0.237339 | -0.20122 | -0.22248 | 0.040695 | 0.053424 | -0.02902 | 0.00917 | 0.07775 | 0.3521 |
| P30OHK | 0.112561 | -0.02957 | -0.05664 | 0.06688 | -0.07254 | -0.12567 | 0.037026 | -0.09763 | 0.06372 | 0.06523 | 0.07883 | 0.12225 | -0.04677 | 0.002311 | -0.23603 | 0.001017 | 0.024533 | -0.1478 | -0.23291 | 0.040759 | -0.34423 | -0.2932 | 0.06391 | -0.00202 | 0.10138 |
| P2URK562 | 0.168078 | 0.067496 | -0.02801 | 0.13169 | -0.10813 | -0.44559 | -0.33446 | -0.03458 | 0.03446 | 0.19565 | -0.09941 | -0.03079 | -0.05633 | 0.039477 | -0.25568 | -0.15763 | 0.102832 | -0.14586 | -0.1922 | 0.066428 | -0.08632 | -0.10911 | -0.14875 | 0.05214 | 0.37387 |
| SLVL | -0.03562 | 0.090669 | -0.16772 | -0.31541 | -0.08927 | -0.14771 | 0.186863 | -0.03061 | 0.04374 | -0.0483 | 0.28185 | 0.18848 | 0.059356 | -0.16139 | -0.60685 | -0.35293 | 0.107971 | -0.18224 | -0.31822 | 0.073742 | -0.29144 | -0.0478 | 0.09865 | -0.23683 | 0.10208 |
| HSSCH2 | -0.01737 | 0.073248 | 0.09284 | -0.11085 | 0.00584 | -0.21318 | 0.012777 | 0.03202 | -0.03113 | 0.05222 | 0.2491 | -0.04091 | 0.039173 | -0.00458 | -0.06965 | 0.068374 | -0.00749 | -0.33193 | -0.35271 | 0.000261 | -1.46225 | -0.00562 | 0.04522 | 0.00917 | 0.03533 |
| NOS1 | 0.061637 | -0.10718 | 0.12386 | 0.1305 | -0.00253 | 0.00223 | -0.06491 | -0.10825 | -0.05622 | 0.19854 | 0.12187 | 0.09576 | 0.271656 | -0.05502 | -0.08123 | 0.026422 | 0.104055 | -0.40359 | -0.11351 | -0.0162 | -0.05434 | 0.10014 | -0.00919 | 0.04305 | 0.16472 |
| HSOS1 | 0.12101 | -0.02959 | 0.0981 | 0.10149 | 0.12413 | -0.20546 | -0.0544 | 0.09069 | -0.02419 | 0.11908 | 0.11034 | 0.1715 | -0.06828 | 0.044918 | -0.08095 | -0.11783 | 0.165387 | -0.25492 | -0.23434 | 0.089277 | -0.0743 | 0.06996 | 0.0166 | 0.07816 | 0.13865 |
| HKBMM | 0.004299 | -0.15836 | -0.05418 | 0.10198 | -0.04625 | -0.08959 | -0.02528 | 0.02579 | -0.09585 | 0.05096 | 0.01268 | 0.05451 | 0.00226 | 0.052841 | -0.18895 | 0.096499 | 0.118036 | -0.1511 | -0.28802 | 0.128558 | -0.20747 | -0.00352 | -0.10267 | 0.02963 | 0.23153 |
| LU165 | 0.068751 | -0.27818 | 0.12007 | 0.06358 | 0.04887 | -0.39166 | -0.1033 | -0.12008 | -0.18713 | -0.23167 | 0.10362 | 0.1722 | 0.120793 | -0.19439 | 0.088894 | -0.04674 | 0.167461 | -0.425 | -0.08377 | -0.17745 | -0.23601 | 0.2651 | 0.20607 | -0.08299 | 0.04665 |
| TN2 | 0.026499 | -0.01349 | 0.00902 | -0.01893 | -0.06844 | -0.09726 | -0.02815 | -0.11289 | -0.02499 | 0.00579 | 0.17724 | 0.13908 | 0.050023 | 0.029758 | -0.17218 | -0.01077 | 0.111497 | -0.13947 | -0.10355 | -0.1304 | -0.19051 | 0.01736 | -0.07111 | 0.01183 | 0.14489 |
| NB69 | -0.24535 | -0.37334 | 0.11257 | 0.00335 | -0.06213 | -0.70195 | -0.00384 | 0.12511 | 0.14151 | -0.16814 | 0.31894 | 0.03969 | 0.017774 | -0.05125 | -0.10697 | -0.08219 | 0.126914 | -0.14758 | -0.14132 | -0.17292 | -0.00579 | 0.04184 | -0.13471 | -0.06054 | 0.12917 |
| MMAC | -0.13318 | 0.1154 | 0.17297 | -0.01464 | -0.15367 | -0.08035 | 0.053564 | -0.20133 | 0.12078 | 0.01346 | 0.11542 | -0.12426 | -0.08261 | -0.02304 | -0.29177 | -0.05893 | 0.181091 | -0.41597 | -0.17119 | -0.03813 | 0.08755 | -0.01069 | -0.076 | 0.01461 | 0.10089 |
| ES4 | -0.03546 | 0.147791 | 0.09825 | 0.05185 | 0.02115 | -0.21793 | -0.05861 | -0.14076 | -0.22652 | 0.07569 | 0.14363 | 0.01846 | 0.052541 | 0.080074 | -0.09973 | -0.01826 | 0.33778 | -0.60911 | -0.61889 | -0.31693 | -0.14 | -0.25936 | -0.02744 | -0.17647 | 0.23413 |
| ES5 | 0.052922 | 0.017926 | -0.0116 | 0.11484 | 0.04554 | 0.08831 | -0.03068 | -0.27057 | 0.04937 | 0.08795 | 0.30349 | -0.05234 | 0.101115 | -0.06331 | -0.15486 | -0.09606 | 0.021922 | -0.26109 | -0.2164 | -0.08014 | 0.066307 | 0.16888 | -0.06238 | -0.01342 | -0.01823 |
| ES8 | 0.024571 | -0.12549 | 0.07711 | -0.22211 | 0.01839 | 0.10203 | 0.059829 | -0.15568 | 0.163 | 0.21812 | 0.05929 | 0.06047 | -0.0157 | -0.02358 | -0.12378 | 0.049408 | 0.1493 | -0.10402 | -0.17689 | 0.128306 | -0.07876 | -0.06206 | 0.01316 | 0.07376 | 0.1621 |
| EW1 | 0.083105 | 0.086303 | -0.07073 | 0.06744 | 0.0409 | -0.08057 | -0.05411 | -0.01737 | -0.21664 | 0.00683 | 0.23585 | 0.1695 | -0.06225 | 0.120993 | 0.114518 | 0.10511 | 0.044451 | -0.10682 | -0.04713 | -0.23209 | -0.123 | 0.21323 | 0.00715 | 0.07007 | -0.03658 |
| EW16 | -0.20547 | 0.044294 | 0.12932 | -0.01049 | -0.01413 | -0.15864 | 0.010806 | -0.06377 | 0.07795 | 0.1244 | 0.04565 | -0.01727 | -0.12025 | -0.05828 | -0.06427 | 0.037078 | 0.13276 | -0.42432 | -0.17625 | -0.3809 | 0.011931 | 0.10709 | -0.06395 | -0.0139 | 0.17717 |
| EW22 | 0.045304 | -0.094 | -0.11076 | 0.174 | -0.07585 | 0.04337 | 0.027617 | -0.15117 | -0.05958 | -0.15081 | 0.00858 | 0.03643 | 0.075472 | 0.067932 | -0.02465 | -0.00919 | -0.02123 | -0.15226 | -0.18715 | -0.14697 | -0.02848 | -0.11275 | 0.02979 | 0.08633 | 0.2421 |
| EW7 | -0.00803 | -0.09642 | 0.02233 | -0.06558 | 0.09563 | 0.11329 | 0.082485 | -0.15494 | 0.15234 | 0.04234 | 0.22569 | -0.05006 | -0.06882 | -0.15856 | 0.028645 | 0.035649 | 0.090609 | -0.14935 | -0.10612 | -0.2943 | -0.17616 | -0.2124 | 0.04325 | -0.03993 | 0.15594 |
| HEY | -0.12263 | -0.14042 | 0.10257 | -0.05994 | -0.09257 | -0.12539 | -0.01117 | 0.05348 | 0.09696 | -0.00936 | 0.24508 | 0.0158 | 0.085422 | 0.090365 | 0.00607 | 0.015916 | 0.123173 | -0.21579 | -0.14431 | -0.20543 | 0.037312 | 0.01052 | 0.10188 | -0.05631 | 0.10916 |
| HSC39 | 0.123333 | 0.204239 | -0.00748 | -0.03947 | -0.08362 | -0.0608 | -0.09835 | -0.07527 | -0.06212 | 0.16796 | 0.17331 | 0.07461 | 0.192311 | -0.01499 | -0.18996 | 0.066164 | 0.028242 | -0.34658 | -1.29509 | -0.06958 | -0.11918 | -1.49951 | -0.11151 | 0.00992 | -0.05759 |
| ISTMEL1 | 0.108647 | -0.02063 | -0.0115 | 0.04811 | -0.20593 | -0.00768 | -0.01952 | -0.03097 | -0.13714 | 0.0723 | -0.12643 | 0.4205 | -0.15832 | 0.109886 | -0.12586 | -0.07158 | -0.0509 | -0.39184 | -0.28596 | -0.00108 | 0.000483 | 0.15943 | 0.09286 | 0.05981 | 0.24151 |
| LB1047RCC | -0.08242 | 0.022345 | 0.05768 | -0.00273 | -0.08164 | -0.09915 | -0.0447 | 0.22998 | -0.11877 | 0.08249 | 0.05342 | 0.03667 | 0.122486 | 0.064936 | 0.054229 | 0.09869 | 0.223564 | -0.65629 | -0.63598 | -0.01427 | -0.11534 | 0.0245 | 0.07511 | 0.04107 | 0.16672 |
| LC1SQ | -0.00105 | 0.131037 | 0.11841 | 0.17466 | -0.11567 | -1.78236 | -0.03169 | -0.23355 | -0.2542 | 0.04498 | 0.17497 | 0.15313 | 0.016289 | 0.169892 | 0.074392 | -0.05632 | 0.074828 | -0.16872 | -0.75284 | -0.07127 | 0.169766 | 0.15245 | -0.20309 | -0.03177 | 0.16446 |
| MCIXC | 0.095574 | 0.19738 | 0.07855 | 0.01424 | -0.28949 | 0.09599 | -0.09413 | -0.34857 | -0.24325 | -0.0019 | 0.1391 | -0.04377 | 0.006848 | -0.13858 | 0.090465 | -0.13006 | -0.11267 | -0.39713 | -0.06486 | -0.32594 | -0.14914 | 0.23047 | 0.0759 | -0.26156 | 0.06972 |
| MKN28 | 0.070964 | 0.097053 | 0.11833 | 0.12676 | -0.16955 | -0.32263 | -0.05569 | -0.0409 | 0.08559 | 0.12548 | -0.03782 | 0.11416 | 0.122762 | 0.093854 | -0.0436 | -0.02607 | 0.008288 | -0.36639 | -0.14293 | -0.03823 | -0.20098 | 0.19703 | 0.02052 | -0.0726 | 0.04926 |
| OCUBM | 0.139744 | -0.08687 | -0.06996 | -0.12388 | 0.08266 | -0.08439 | 0.029956 | 0.02232 | -0.1868 | 0.00649 | 0.03066 | 0.24586 | -0.02884 | 0.077997 | 0.108606 | -0.09984 | 0.272902 | -2.58775 | -1.60369 | -0.13496 | -0.20278 | -0.18738 | 0.0681 | -0.19907 | 0.22525 |
| OVCA420 | -0.08831 | 0.036583 | 0.00991 | -0.0093 | -0.06326 | -0.17378 | -0.20074 | -0.0687 | 0.04652 | -0.06078 | 0.24414 | 0.01163 | 0.05343 | -0.18038 | -0.07973 | -0.01564 | -0.04161 | -0.97091 | -1.03666 | -0.07683 | 0.072642 | 0.03536 | 0.0051 | 0.09785 | 0.07327 |
| OVMIU | -0.10933 | -0.06505 | -0.15027 | -0.10652 | -0.09892 | -0.62515 | 0.086177 | 0.0014 | 0.03422 | 0.35419 | 0.01657 | -0.12203 | 0.093606 | 0.018507 | -0.17187 | 0.044494 | -0.28076 | -0.28638 | -0.33061 | -0.42245 | 0.072896 | 0.12389 | 0.15326 | -0.10476 | 0.20997 |
| PL4 | 0.225243 | -0.14275 | -0.04327 | -0.06622 | 0.16031 | 0.13002 | 0.062591 | 0.01661 | -0.06226 | -0.03634 | 0.43458 | 0.04771 | -0.00978 | -0.11096 | 0.008717 | -0.03269 | 0.188169 | -0.61871 | -0.29936 | 0.200881 | -0.40203 | -0.01602 | 0.14136 | 0.00388 | 0.1111 |
| RCCFG2 | 0.095692 | -0.18644 | -0.09243 | -0.00406 | -0.19058 | 0.05143 | 0.130697 | 0.09976 | -0.11209 | 0.07057 | 0.10628 | 0.03587 | 0.083157 | 0.19921 | -0.12775 | 0.135218 | -0.05822 | -0.04329 | -0.14387 | -0.06684 | -0.47456 | -0.00391 | 0.08328 | -0.00986 | 0.09471 |
| SCH | 0.205843 | 0.261251 | -0.03822 | 0.09247 | 0.00225 | 0.20155 | 0.02811 | 0.15751 | -0.07671 | 0.23763 | -0.05746 | 0.53711 | 0.051538 | 0.158434 | 0.022067 | 0.127387 | 0.42659 | -0.00778 | -0.17789 | -0.21407 | -0.18684 | 0.24268 | 0.31626 | 0.0882 | -0.09265 |
| TMK1 | 0.122902 | 0.036428 | 0.03279 | 0.11339 | -0.12724 | -0.51997 | -0.01404 | 0.30017 | -0.19316 | 0.06637 | -0.28178 | 0.11411 | 0.186064 | 0.302517 | 0.127988 | 0.151263 | 0.101297 | -0.36005 | 0.10612 | -0.15749 | -0.03005 | -0.03978 | 0.20922 | 0.06498 | 0.3144 |
| ARH77 | 0.133797 | 0.059392 | -0.05368 | 0.16205 | -0.19223 | 0.09091 | -0.03421 | 0.0383 | -0.02018 | 0.2232 | -0.05034 | 0.2128 | 0.052056 | 0.195642 | 0.090701 | 0.018354 | 0.171669 | -0.17763 | 0.06686 | -0.04268 | -0.03495 | 0.05358 | 0.19716 | -0.00981 | 0.15763 |
| BB30HNC | -0.00806 | 0.063571 | -0.19237 | -0.07363 | -0.29046 | -1.18992 | 0.148705 | -0.04037 | 0.23386 | 0.19618 | 0.057 | 0.08249 | 0.116161 | 0.136447 | 0.107114 | 0.136158 | 0.066351 | -0.30278 | -0.32017 | 0.059945 | -0.12971 | -0.13822 | 0.16774 | 0.00744 | 0.18507 |
| BE2M17 | -0.1283 | -0.01884 | -0.01998 | -0.00148 | -0.07607 | -0.24236 | -0.06661 | 0.05307 | -0.04999 | 0.13346 | 0.26423 | 0.05082 | -0.04195 | 0.024808 | -0.14732 | -0.02459 | -0.07917 | -0.23352 | -0.13564 | -0.41161 | -0.26273 | -0.08076 | -0.04765 | -0.10009 | 0.11163 |
| D423MG | 0.195888 | -0.05106 | 0.03033 | -0.30289 | -0.05842 | -0.26665 | 0.133123 | 0.09046 | 0.13447 | 0.03872 | 0.27209 | 0.00724 | 0.032542 | 0.097549 | -0.2347 | -0.06098 | 0.089157 | -0.38166 | -0.39591 | -0.02073 | -0.12457 | 0.07014 | 0.09017 | -0.02346 | 0.06858 |
| D502MG | -0.03827 | -0.10523 | 0.21522 | -0.04385 | -0.05395 | -0.47195 | -0.07602 | 0.39651 | -0.05035 | -0.12588 | 0.42529 | 0.13599 | 0.212405 | -0.00359 | 0.054852 | 0.054553 | 0.273327 | -0.22118 | -0.44966 | -0.16798 | -0.19378 | -0.08707 | -0.1513 | -0.02816 | 0.15584 |
| D542MG | 0.009309 | -0.11341 | -0.04075 | -0.12383 | -0.23014 | -0.00435 | 0.111192 | 0.05934 | 0.07424 | 0.01286 | 0.23524 | 0.02366 | 0.169767 | -0.05759 | -0.07298 | 0.08882 | 0.003283 | -0.16854 | -0.28078 | -0.03603 | -0.29953 | -0.01786 | -0.04227 | -0.17702 | -0.08831 |
| DIFI | 0.104297 | 0.157606 | -0.11056 | -0.21976 | 0.13259 | -0.97196 | -0.08028 | -0.18263 | 0.2347 | -0.05467 | 0.08267 | 0.18853 | 0.051919 | 0.095407 | -0.24736 | -0.2926 | 0.009489 | -0.25399 | -0.29254 | -0.02673 | 0.038074 | 0.05716 | 0.06562 | -0.02344 | 0.13404 |
| DOK | 0.132656 | 0.013253 | 0.00277 | -0.0573 | -0.23554 | -1.08068 | -0.04261 | 0.4338 | -0.16011 | 0.19592 | -0.1294 | 0.16901 | -0.05214 | 0.188607 | 0.146016 | 0.206225 | 0.063898 | -0.60224 | -1.10963 | 0.021125 | -0.00203 | -0.2432 | 0.24667 | 0.1539 | 0.25151 |
| H3118 | 0.053826 | -0.15793 | -0.09216 | -0.06808 | -0.11926 | -1.80345 | -0.00535 | 0.42595 | 0.04447 | 0.16982 | 0.02801 | 0.07512 | 0.115592 | 0.29268 | -0.06154 | -0.07549 | 0.092843 | -0.3146 | -0.21266 | 0.080589 | -0.13742 | 0.17073 | 0.02144 | -0.14565 | 0.15404 |
| HCE4 | 0.235034 | 0.083099 | -0.18117 | 0.13485 | 0.03482 | -0.12014 | -0.01402 | 0.21008 | 0.14439 | -0.02998 | -0.22661 | 0.14817 | 0.001673 | 0.309532 | 0.214671 | 0.006233 | 0.119946 | -0.13712 | -0.11908 | -0.27521 | -0.6762 | 0.1223 | 0.54698 | 0.15216 | 0.19643 |
| IM9 | 0.048045 | 0.137882 | 0.05656 | 0.17774 | -0.14487 | -0.02857 | 0.029814 | 0.0267 | 0.11497 | 0.03975 | 0.20203 | 0.07363 | -0.01891 | -0.06603 | -0.22853 | 0.147509 | 0.049096 | -0.03293 | 0.00798 | -0.37653 | -0.1082 | 0.09112 | 0.07624 | -0.00166 | 0.23345 |
| JHU011 | 0.103426 | 0.054428 | 0.11902 | 0.07093 | -0.08268 | -1.41832 | 0.00068 | 0.30977 | -0.19996 | 0.05827 | -0.17404 | 0.18332 | 0.110129 | 0.270163 | -0.05 | 0.073591 | -0.03253 | -0.35394 | -0.17666 | -0.03286 | -0.04826 | 0.05956 | 0.01693 | 0.04984 | 0.00341 |
| JHU022 | 0.100982 | 0.013763 | -0.11638 | 0.20752 | -0.28736 | -0.72287 | -0.08092 | 0.06542 | -0.02766 | 0.23783 | -0.0585 | -0.01815 | 0.24613 | -0.16958 | -0.1575 | -0.10897 | 0.034063 | -0.70531 | -0.61919 | -0.07538 | 0.168594 | 0.03326 | 0.07 | 0.03925 | 0.09966 |
| JHU029 | 0.079139 | -0.02383 | 0.05516 | -0.13394 | -0.00614 | -1.54414 | 0.213441 | -0.05292 | -0.13366 | 0.09074 | 0.14081 | 0.03175 | 0.123932 | -0.00661 | -0.03684 | 0.043541 | 0.116258 | -0.05665 | -0.03151 | -0.02685 | -0.06815 | -0.06295 | 0.0744 | 0.05747 | -0.00425 |

| Cell line | ALK | AXL | CSF1R | DDR1 | DDR2 | EGFR | EPHA1 | EPHA2 | EPHA3 | EPHA4 | EPHA6 | EPHA7 | EPHB1 | EPHB2 | EPHB3 | EPHB4 | EPHB6 | ERBB2 | ERBB3 | ERBB4 | FGFR1 | FGFR2 | FGFR3 | FGFR4 | FLT1 |
|---|---|---|---|---|---|---|---|---|---|---|---|---|---|---|---|---|---|---|---|---|---|---|---|---|---|
| KINGS1 | -0.0888 | -0.21075 | 0.07237 | 0.0712 | -0.03958 | 0.06748 | -0.1233 | 0.20602 | 0.06507 | -0.03374 | 0.28963 | 0.02349 | 0.286989 | 0.039594 | -0.20811 | 0.046121 | -0.15477 | -0.19364 | 0.0674 | -0.06471 | -0.01567 | -0.08246 | -0.10795 | -0.00741 | -0.02509 |
| KPNYS | 0.080802 | 0.17659 | 0.04434 | 0.1126 | -0.10312 | 0.05831 | 0.058417 | -0.27771 | -0.0891 | -0.01707 | 0.19427 | 0.0344 | 0.216136 | 0.051427 | -0.14078 | 0.016564 | -0.07982 | -0.05405 | -0.06525 | -0.14532 | 0.168877 | -0.16858 | -0.1833 | -0.01216 | 0.00371 |
| KYSE220 | -0.0668 | 0.199573 | -0.02618 | 0.07664 | -0.13001 | -0.63802 | 0.01004 | 0.24661 | 0.09523 | 0.11524 | 0.02968 | 0.03774 | 0.098402 | 0.030047 | -0.07612 | 0.09632 | -9E-05 | -0.15037 | 0.03981 | -0.11816 | -0.00565 | -0.08039 | 0.20884 | 0.12401 | 0.13163 |
| LB771HNC | 0.212482 | 0.127583 | -0.17138 | 0.16288 | -0.21748 | -0.61518 | -0.0566 | 0.05088 | 0.00473 | 0.1271 | 0.09809 | 0.07889 | 0.059808 | 0.110906 | -0.02922 | 0.077236 | 0.089613 | -0.30069 | -0.24295 | -0.09307 | -0.01893 | -0.11865 | -0.00216 | -0.03132 | 0.14 |
| LNZTA3WT4 | -0.04495 | 0.010531 | 0.06641 | 0.07025 | 0.05024 | 0.06337 | -0.02437 | -0.19402 | 0.00228 | -0.00011 | 0.13587 | 0.17199 | 0.04651 | 0.094171 | -0.12134 | 0.068987 | 0.040948 | -0.24702 | -0.14426 | 0.010608 | -1.56575 | -0.07613 | -0.05135 | -0.0252 | -0.02231 |
| NB10 | 0.033382 | -0.18884 | 0.01361 | -0.13334 | -0.0705 | 0.14964 | -0.02954 | -0.29558 | 0.00056 | -0.04149 | 0.23615 | 0.04985 | 0.019837 | -0.06141 | -0.21745 | 0.042015 | 0.096855 | -0.0766 | -0.1779 | -0.20967 | 0.055908 | 0.02138 | 0.00792 | -0.15772 | 0.16805 |
| NB13 | 0.095717 | 0.071909 | -0.08187 | 0.12985 | -0.08645 | 0.13566 | -0.01077 | -0.47245 | 0.22078 | 0.15886 | 0.03854 | -0.02439 | 0.103906 | -0.20657 | -0.32451 | 0.089727 | -0.15954 | -0.19558 | -0.4956 | -0.4388 | 0.051338 | 0.09584 | -0.15562 | 0.00904 | 0.02006 |
| NB17 | -0.20949 | -0.00654 | 0.08537 | 0.05872 | 0.07215 | 0.26426 | 0.13693 | -0.06791 | -0.01682 | -0.00902 | 0.37931 | 0.06776 | -0.00968 | -0.08746 | -0.15469 | 0.033615 | 0.03776 | 0.13301 | -0.14393 | -0.25863 | 0.006139 | -0.18153 | -0.31595 | -0.02483 | -0.12543 |
| NB5 | 0.127298 | -0.01395 | 0.06162 | -0.03619 | 0.19552 | -0.20853 | 0.04987 | -0.04026 | 0.0898 | 0.12704 | 0.35678 | 0.05958 | 0.093846 | 0.09779 | -0.3475 | 0.104879 | 0.113868 | -0.09899 | -0.10348 | -0.00657 | 0.217637 | 0.15805 | 0.12416 | -0.03219 | 0.29064 |
| NB6 | -0.06761 | 0.0713 | 0.02684 | 0.064 | -0.08755 | -0.0131 | -0.04762 | -0.19468 | 0.04509 | -0.16171 | 0.08574 | 0.17154 | 0.036326 | -0.02073 | -0.34283 | 0.005316 | 0.007783 | -0.25023 | -0.09981 | -0.10685 | -0.04598 | -0.02963 | -0.14489 | -0.104 | -0.0321 |
| NB7 | -0.23156 | 0.037486 | 0.04931 | -0.0395 | -0.00034 | 0.12516 | -0.04786 | 0.02953 | 0.08626 | -0.07242 | -0.08426 | 0.0457 | -0.07439 | -0.08667 | -0.16133 | 0.06868 | -0.09086 | -0.25478 | -0.3588 | -0.11317 | 0.030126 | -0.02823 | -0.06392 | 0.08958 | 0.11305 |
| NTERA2CLD1 | -0.06156 | 0.099227 | 0.00199 | 0.08145 | -0.06989 | -0.12922 | -0.17155 | 0.00484 | 0.00265 | 0.10066 | 0.1247 | 0.20713 | -0.0264 | 0.119635 | -0.21076 | -0.06366 | -0.05771 | -0.37625 | -0.28697 | -0.21149 | 0.166849 | 0.08236 | 0.11951 | 0.0613 | 0.14534 |
| PCI15A | 0.002393 | 0.086223 | 0.03148 | 0.04016 | -0.13998 | -0.9774 | 0.034981 | 0.05801 | -0.05645 | 0.00206 | 0.08895 | 0.13909 | -0.00803 | 0.269972 | -0.04558 | -0.0028 | 0.194641 | -0.26669 | -0.38626 | -0.08854 | -0.06899 | 0.0142 | 0.0369 | -0.10341 | -0.01043 |
| PCI30 | 0.061764 | -0.00589 | -0.02224 | -0.05134 | -0.15175 | 0.13766 | -0.02709 | 0.03702 | 0.04944 | 0.1376 | -0.05703 | 0.13841 | 0.071618 | 0.151931 | -0.04902 | 0.010882 | -0.17195 | -0.24625 | -0.18031 | -0.04896 | -0.59502 | 0.08531 | -0.11131 | -0.0456 | 0.10704 |
| PCI38 | -0.01991 | 0.019928 | -0.14034 | -0.1266 | -0.00611 | -0.6864 | 0.188728 | -0.01168 | -0.00028 | 0.05334 | 0.07803 | 0.02727 | 0.075314 | 0.085946 | -0.01303 | 0.087517 | 0.137165 | -0.41726 | -0.59974 | -0.05319 | -0.10637 | -0.09583 | 0.13092 | -0.07269 | 0.03088 |
| PCI4B | 0.105816 | 0.12631 | 0.14217 | -0.01186 | -0.05002 | -0.7714 | 0.198464 | 0.23774 | -0.13902 | -0.04165 | 0.15377 | -0.04111 | 0.118104 | 0.211893 | -0.16891 | 0.037918 | 0.081937 | -0.60183 | -0.56086 | -0.08921 | -0.08863 | 0.11491 | 0.17309 | -0.06277 | 0.15295 |
| PCI6A | 0.213283 | -0.01493 | -0.0569 | 0.07985 | 0.1651 | -0.70915 | 0.170779 | 0.17773 | -0.12299 | 0.14525 | 0.02486 | 0.26686 | 0.044313 | 0.10867 | 0.214037 | 0.097933 | 0.319632 | -0.39049 | -0.5471 | -0.18557 | 0.094407 | -0.12409 | -0.00107 | 0.00648 | 0.17026 |
| SKMG1 | -0.04281 | -0.20384 | 0.01962 | -0.05265 | -0.13975 | 0.13571 | -0.02099 | -0.02912 | 0.09392 | 0.04531 | 0.08626 | 0.16696 | 0.018028 | -0.11549 | 0.077653 | -0.07802 | 0.007847 | -0.12989 | -0.25879 | -0.01049 | 0.08986 | 0.17247 | 0.04991 | -0.0066 | 0.25324 |
| SKN3 | 0.094088 | 0.319926 | 0.13851 | 0.13001 | -0.25694 | -0.2703 | 0.152427 | 0.2139 | -0.19668 | 0.22262 | -0.22321 | 0.27024 | 0.009586 | 0.410592 | 0.150357 | 0.135531 | 0.219195 | -0.03992 | 0.04053 | -0.23022 | -0.06064 | 0.11478 | 0.18027 | 0.2567 | 0.12036 |
| 21NT | -0.03427 | 0.01614 | -0.11753 | 0.12761 | -0.11105 | -0.5791 | -0.12738 | -0.08454 | -0.16383 | 0.18273 | 0.11025 | 0.03226 | -0.01564 | 0.086661 | -0.14164 | 0.320981 | 0.117174 | -0.23947 | -0.50513 | 0.051768 | -0.10461 | -0.17221 | 0.06828 | -0.05517 | 0.14665 |
| CCLFUPGI0005T | 0.040142 | -0.04437 | -0.08219 | 0.0559 | 0.01624 | -0.89917 | -0.13094 | 0.06677 | 0.10891 | 0.07627 | 0.07894 | 0.07131 | 0.04108 | -0.05258 | -0.14874 | -0.15409 | 0.053315 | -0.54985 | -0.1902 | -0.01486 | -0.02558 | -0.16553 | -0.09245 | -0.06798 | 0.13275 |
| HT144SKINFV1 | -0.21543 | 0.217713 | 0.08934 | 0.05854 | 0.09958 | -0.08799 | -0.07355 | 0.1712 | -0.00673 | -0.0648 | 0.33832 | -0.1298 | -0.01621 | 0.215553 | -0.0187 | -0.03665 | -0.07758 | -0.43451 | -0.37679 | 0.105971 | -0.08086 | 0.08247 | 0.09075 | -0.06964 | 0.27658 |
| HT144SKINFV3 | 0.059688 | 0.060911 | -0.03234 | 0.09384 | -0.00043 | -0.24087 | -0.02633 | 0.01412 | -0.13915 | 0.09813 | 0.1217 | 0.08311 | -0.08521 | 0.147063 | -0.04718 | 0.062373 | 0.020893 | -0.02677 | -0.36064 | -0.10112 | -0.13966 | 0.07232 | 0.02635 | 0.06881 | 0.15575 |
| HT144SKINFV2 | -0.10512 | 0.074978 | 0.03017 | 0.01915 | 0.01525 | 0.06117 | -0.05435 | 0.00415 | -0.03281 | 0.08028 | 0.37132 | -0.03395 | -0.00177 | 0.154107 | 0.013557 | 0.182256 | 0.153054 | -0.37475 | -0.39355 | -0.166 | 0.04945 | 0.24454 | 0.10867 | -0.11811 | 0.15114 |
| RVH421SKINFV1 | 0.168473 | 0.099907 | -0.08726 | -0.12862 | -0.04985 | -0.12168 | 0.077452 | -0.07104 | -0.03369 | -0.06203 | 0.01663 | 0.01366 | 0.034718 | -0.15422 | -0.13271 | 0.015531 | -0.01489 | -0.35585 | -0.34629 | -0.07539 | -0.13476 | 0.05732 | 0.0712 | -0.01226 | 0.1529 |
| RPE1SS48 | -0.06134 | -0.09747 | -0.08643 | -0.08716 | -0.03462 | -0.11002 | 0.003642 | -0.01769 | 0.0107 | 0.02086 | 0.13962 | 0.0338 | 0.033158 | 0.131636 | -0.12981 | 0.166975 | -0.09875 | -0.19753 | -0.26424 | 0.056794 | -0.1277 | -0.0999 | 0.08484 | 0.0428 | 0.27537 |
| RPE1SS77 | 0.048893 | -0.37609 | -0.10848 | 0.0466 | -0.0494 | -0.15404 | -0.19236 | 0.15173 | -0.15594 | 0.03193 | 0.16192 | 0.09813 | 0.028134 | 0.118045 | -0.11409 | 0.154401 | 0.042374 | -0.16748 | -0.36996 | -0.05475 | -0.19982 | -0.08906 | 0.07032 | -0.04346 | 0.126 |
| RPE1SS6 | -0.00912 | -0.67013 | -0.02825 | -0.02173 | 0.00257 | -0.09608 | -0.05776 | -0.03918 | -0.13154 | -0.00378 | 0.10749 | -0.03481 | 0.025758 | 0.120286 | -0.10162 | 0.153429 | 0.079679 | -0.15256 | -0.01649 | -0.08064 | 0.032703 | -0.03127 | 0.16329 | -0.07494 | 0.1954 |
| RPE1SS119 | 0.025264 | -0.33438 | 0.01047 | 0.14972 | -0.1394 | -0.22822 | -0.16523 | -0.10923 | -0.14351 | 0.14296 | 0.17867 | 0.03721 | 0.097421 | 0.076694 | 0.107873 | -0.05491 | -0.05118 | -0.15709 | -0.16131 | 0.137625 | 0.086778 | 0.07383 | -0.00101 | -0.03659 | 0.00376 |
| RPE1SS51 | -0.10904 | -0.10068 | 0.11643 | -0.07838 | 0.08689 | -0.14621 | 0.036163 | -0.24298 | 0.00388 | 0.01942 | 0.13676 | -0.06412 | 0.20135 | 0.222661 | -0.03276 | 0.089275 | -0.11364 | -0.33516 | -0.1949 | 0.091639 | 0.21263 | -0.05093 | -0.26361 | -0.11563 | 0.01997 |
| PSS008 | -0.0094 | 0.014063 | 0.14507 | 0.0657 | 0.08075 | -0.25811 | 0.0532 | -0.05713 | -0.01074 | 0.0344 | 0.18231 | -0.05555 | 0.036441 | -0.10307 | -0.09391 | 0.143933 | 0.229065 | -0.56663 | -0.26941 | -0.32664 | 0.004277 | 0.09257 | 0.06062 | 0.0849 | 0.14413 |
| MAVER1 | 0.056607 | 0.159571 | 0.15695 | 0.07517 | -0.03232 | -0.02776 | 0.138914 | -0.02835 | -0.04688 | 0.04243 | 0.24816 | 0.05171 | 0.10613 | 0.213938 | -0.18212 | 0.160591 | 0.163309 | -0.12777 | -0.3113 | -0.06994 | -0.16924 | 0.12153 | 0.12722 | -0.0033 | 0.13201 |
| MESOV | 0.017652 | -0.00058 | -0.14488 | 0.01371 | -0.01877 | -0.0857 | -0.16324 | 0.16906 | 0.00847 | 0.04804 | 0.10529 | 0.04605 | -0.02688 | 0.184011 | -0.11276 | 0.013231 | -0.0461 | -0.42156 | -0.22197 | -0.01437 | 0.029017 | 0.07372 | 0.14899 | -0.01632 | 0.21597 |
| WM3211 | 0.023731 | -0.14248 | 0.08666 | -0.06004 | -0.07252 | -0.1074 | -0.04683 | 0.09955 | -0.08536 | -0.01341 | 0.16736 | 0.02152 | -0.005 | 0.025421 | -0.19679 | -0.01267 | -0.10289 | -0.41261 | -0.22732 | -0.02904 | -1.78532 | 0.06293 | 0.10996 | -0.01419 | 0.24747 |
| M040416 | -0.21829 | 0.103078 | 0.08968 | -0.06032 | -0.07317 | -0.27263 | -0.05635 | -0.24432 | -0.0584 | -0.18324 | 0.02557 | 0.14053 | 0.12087 | -0.0612 | 0.108776 | -0.16113 | 0.065979 | -0.095 | -0.2952 | -0.05285 | 0.003364 | -0.06699 | -0.18689 | -0.0696 | 0.17398 |
| M140325 | 0.144138 | 0.043939 | 0.0486 | -0.05571 | 0.00386 | -0.11035 | -0.03627 | -0.10045 | -0.10371 | -0.0327 | -0.02178 | 0.09409 | 0.112629 | 0.088021 | -0.07151 | -0.25362 | 0.072729 | -0.4629 | -0.37037 | -0.01497 | -0.08507 | 0.10545 | 0.06492 | -0.04297 | 0.10237 |
| MM160113 | -0.3273 | -0.1655 | 0.17634 | 0.15603 | 0.01331 | -0.27845 | -0.27246 | -0.00336 | 0.0054 | -0.08549 | 0.05202 | -0.17138 | -0.00122 | -0.0744 | 0.002455 | -0.21552 | 0.182176 | -0.42417 | -0.28482 | -0.20761 | -0.07268 | 0.32675 | -0.51109 | 0.26029 | -0.03714 |
| SNU739 | 0.018326 | 0.010763 | -0.02278 | -0.12291 | -0.09302 | -0.4133 | -0.00933 | 0.07961 | 0.05235 | 0.07598 | 0.08571 | 0.10875 | -0.06834 | 0.050463 | -0.06229 | 0.067013 | -0.02364 | -0.51448 | -0.13872 | -0.03011 | -0.28005 | -0.47877 | -0.06099 | 0.0141 | 0.30062 |
| SNU1327 | -0.04943 | -0.08905 | -0.08608 | -0.00035 | -0.09705 | -0.42032 | 0.061448 | -0.06968 | -0.0036 | 0.16053 | 0.24234 | 0.13898 | 0.049467 | -0.04501 | -0.18017 | -0.00208 | 0.100131 | -0.25634 | -0.15355 | 0.007419 | -0.00442 | -0.32463 | 0.12089 | -0.06176 | -0.03863 |
| SNU2535 | 0.182075 | 0.185101 | -0.14011 | 0.02594 | -0.07973 | -0.0737 | -0.28394 | 0.05192 | -0.3227 | 0.07431 | 0.12336 | 0.1145 | 0.145284 | 0.089984 | -0.20435 | 0.035939 | 0.095914 | -0.21954 | -0.30304 | -0.1277 | 0.02411 | 0.08201 | 0.12328 | -0.07923 | 0.33857 |
| CCC5 | 0.037962 | -0.02917 | 0.08694 | 0.04574 | -0.16205 | -0.50853 | -0.04865 | 0.19891 | 0.00465 | -0.03916 | 0.13088 | 0.02423 | -0.09086 | -0.17409 | -0.14016 | 0.006519 | 0.069285 | -0.33104 | -0.09275 | -0.11853 | -0.04863 | 0.24477 | -0.12819 | 0.08285 | 0.13261 |
| ETCC016 | 0.108226 | 0.082316 | -0.06947 | -0.05385 | -0.04825 | -0.21306 | -0.13095 | 0.21452 | -0.04364 | 0.07424 | 0.13693 | 0.03628 | 0.148429 | 0.033638 | -0.18306 | -0.02935 | -0.06941 | -0.41525 | -0.52543 | -0.03198 | -0.07571 | 0.08731 | -0.00549 | -0.02311 | 0.13694 |
| JVE015 | 0.009299 | -0.06661 | -0.34781 | -0.00666 | -0.02225 | -0.11215 | -0.17737 | -0.15741 | -0.2844 | -0.01458 | 0.2191 | 0.39922 | 0.215423 | -0.13206 | -0.41285 | -0.04542 | -0.10296 | -0.30416 | -0.20931 | 0.114722 | 0.024238 | -0.06681 | 0.04163 | -0.28643 | 0.03911 |
| JVE127 | 0.01371 | -0.01195 | -0.09172 | 0.15349 | 0.02085 | -0.11644 | -0.05246 | 0.01719 | 0.00196 | 0.16355 | 0.05458 | 0.06465 | 0.097597 | 0.020486 | -0.10306 | 0.015379 | 0.262227 | -0.25729 | -0.19893 | -0.08967 | -0.05289 | 0.07924 | 0.05278 | 0.12457 | 0.29652 |
| JVE253 | 0.12975 | 0.067883 | -0.03528 | 0.00807 | 0.04968 | -0.12503 | 0.045687 | -0.12938 | -0.04265 | 0.06014 | 0.09873 | 0.06477 | 0.000244 | 0.013539 | -0.13215 | -0.03194 | 0.013969 | -0.3556 | -0.22464 | -0.0929 | -0.13781 | 0.05567 | 0.01026 | -0.03158 | 0.10884 |
| KP363T | 0.04367 | -0.09023 | 0.04259 | 0.1214 | -0.07222 | -0.31316 | 0.102061 | 0.03888 | 0.01757 | -0.04935 | 0.11258 | 0.10701 | -0.02104 | 0.034704 | -0.07325 | -0.06126 | 0.035643 | -0.24868 | -0.12631 | 0.08966 | -0.23987 | -0.06152 | 0.03478 | -0.07874 | 0.37365 |
| MAPACHS77 | 0.115928 | -0.16381 | -0.01518 | -0.05509 | -0.04382 | -0.91415 | -0.06547 | 0.21827 | -0.00951 | -0.11871 | 0.1538 | 0.03516 | -0.07138 | -0.02871 | -0.08801 | -0.02316 | 0.141899 | -0.46207 | -0.11774 | -0.10107 | -0.15594 | 0.05414 | -0.03659 | 0.07126 | 0.26657 |
| 170MGBA | 0.136804 | -0.06263 | -0.02443 | -0.06698 | -0.05955 | -0.12062 | 0.029475 | -0.0237 | 0.03859 | 0.06174 | 0.15059 | 0.03139 | -0.06158 | 0.070279 | -0.08547 | 0.070542 | 0.030888 | -0.20314 | -0.3457 | -0.06379 | -0.16129 | 0.03432 | 0.02046 | -0.02497 | 0.08649 |
| WM3772F | 0.008414 | 0.038678 | 0.10095 | -0.2081 | -0.14638 | -0.12909 | -0.04226 | -0.07658 | -0.11933 | 0.02346 | 0.23654 | 0.02682 | 0.045351 | 0.048809 | -0.04848 | 0.18932 | 0.073318 | -0.45236 | -0.14632 | 0.1568 | -0.00264 | 0.04706 | -0.00445 | -0.01554 | 0.28057 |
| S462 | 0.174353 | 0.056127 | 0.12423 | -0.05384 | -0.1153 | -0.2654 | -0.16557 | -0.13265 | 0.09017 | -0.0023 | 0.23497 | -0.0727 | -0.03203 | -0.04265 | -0.11722 | -0.1403 | 0.006148 | -0.48471 | -0.36103 | 0.047828 | -0.40284 | 0.02378 | 0.06859 | 0.0214 | -0.19334 |
| MPNST724 | 0.178346 | 0.069862 | -0.00209 | -0.03203 | -0.01217 | -0.11248 | 0.019142 | 0.15865 | -0.02136 | 0.08498 | 0.11574 | 0.09452 | -0.0082 | 0.140811 | -0.15487 | -0.12422 | 0.067847 | -0.33046 | -0.23256 | 0.028333 | -1.6187 | 0.04224 | -0.00429 | 0.00876 | 0.13332 |
| NCCLMS1C1 | 0.172578 | 0.15351 | -0.07582 | -0.04528 | -0.06069 | -0.12739 | -0.00967 | -0.04646 | -0.23463 | 0.11129 | 0.18768 | -0.11629 | -0.01563 | -0.01728 | -0.12721 | 0.010147 | 0.082948 | -0.28662 | -0.30793 | 0.11504 | -0.61934 | -0.07971 | 0.03964 | -0.02685 | 0.13735 |
| NCCMPNST1C1 | 0.072637 | -0.05505 | -0.12164 | 0.13993 | -0.03513 | -0.02147 | 0.014276 | 0.07451 | -0.00488 | -0.03075 | 0.08392 | -0.03394 | -0.05002 | -0.03374 | -0.11876 | -0.11211 | 0.012979 | -0.1687 | -0.28691 | 0.005188 | -0.33159 | 0.03551 | 0.10775 | 0.05695 | 0.07055 |
| NCCMPNST2C1 | 0.032772 | -0.07468 | -0.03614 | -0.09911 | 0.08029 | -0.20115 | 0.178178 | -0.06155 | 0.12584 | -0.02279 | 0.08278 | 0.155 | -0.16836 | 0.048482 | -0.44366 | 0.124355 | 0.14352 | -0.37526 | -0.14315 | -0.20538 | -0.31898 | 0.10141 | 0.04973 | 0.10802 | 0.07166 |
| PSS131R | 0.119084 | -0.28226 | 0.02233 | 0.0361 | -0.10204 | -0.33906 | -0.1798 | 0.0846 | -0.02884 | 0.0583 | 0.12425 | 0.07868 | -0.10761 | -0.06559 | -0.18737 | 0.02704 | 0.101536 | -0.15835 | -0.27542 | 0.046274 | -0.19702 | 0.22454 | 0.02131 | -0.02046 | 0.0612 |
| YUHOIN0650 | 0.027433 | -0.22289 | 0.01823 | 0.08534 | 0.04537 | -0.37185 | -0.0898 | 0.39634 | -0.02272 | 0.08013 | 0.10998 | 0.05363 | 0.09463 | 0.0901 | -0.13059 | -0.08839 | -0.02332 | -0.31822 | -0.287 | -0.07189 | -0.06819 | -0.03276 | 0.10022 | -0.0041 | 0.18638 |
| SKNMM | 0.063931 | 0.044013 | -0.02673 | 0.07696 | 0.00771 | -0.20183 | -0.05491 | -0.1112 | -0.12101 | -0.00393 | 0.03386 | 0.02519 | 0.024762 | 0.092598 | -0.18715 | -0.0106 | 0.009611 | -0.21879 | -0.19747 | -0.02827 | -0.01893 | 0.03605 | 0.03725 | 0.03183 | 0.11134 |
| UPMD1 | 0.178259 | 0.112578 | 0.01983 | 0.03065 | -0.03471 | -0.14022 | 0.154547 | -0.11522 | -0.14392 | 0.05316 | 0.01057 | -0.04845 | -0.01239 | 0.013891 | -0.19136 | 0.022087 | 0.153312 | -0.14913 | -0.20738 | -0.05879 | -0.1996 | -0.01299 | 0.04791 | 0.04657 | 0.14817 |
| average | -0.00469 | 0.001021 | -0.00639 | -0.00486 | -0.04733 | -0.27148 | -0.00499 | 0.01215 | -0.02428 | 0.07161 | 0.12256 | 0.06566 | 0.025962 | 0.063496 | -0.06581 | 0.025396 | 0.080183 | -0.35461 | -0.26274 | -0.08563 | -0.15978 | -0.04389 | 0.01295 | -0.0043 | 0.11465 |

| Cell line | FLT3 | FLT4 | IGF1R | INSR | INSRR | KDR | KIT | LMR1 | LMR2 | LTK | MET | MUSK | PDGFRA | PDGFRB | PTK7 | RET | RON | ROR1 | ROR2 | ROS1 | STYK1 | TIE1 | TIE2 | TRKA | TRKB | TRKC | TYRO3 |
|---|---|---|---|---|---|---|---|---|---|---|---|---|---|---|---|---|---|---|---|---|---|---|---|---|---|---|---|
| NIHOVCAR3 | -0.0859 | 0.069397 | -0.29435 | -0.15649 | -0.0524 | -0.09882 | -0.08374 | -0.01948 | -0.10188 | -0.09651 | -0.04401 | 0.10725 | -0.1337 | -0.12162 | 0.171933 | 0.065869 | 0.03287 | 0.2317 | 0.048163 | 0.04651 | 0.12063 | -0.12324 | -0.29513 | 0.15918 | 0.26295 | -0.04778 | -0.27215 |
| HEL | -0.20633 | 0.088765 | -0.18856 | -0.07004 | -0.23612 | -0.14974 | 0.096286 | -0.00902 | 0.25645 | 0.0949 | -0.0929 | 0.10191 | -0.18267 | -0.04062 | 0.041041 | 0.057095 | -0.0815 | 0.26934 | 0.233863 | 0.11817 | 0.05287 | -0.52727 | 0.053522 | -0.10427 | 0.35643 | 0.412187 | -1.22504 |
| HEL9217 | 0.14323 | -0.21791 | -0.33696 | -0.17133 | -0.06929 | -0.29966 | 0.140651 | 0.03172 | -0.11505 | 0.13386 | -0.1446 | 0.47522 | -0.21882 | 0.06596 | 0.071155 | -0.15486 | -0.25669 | 0.40982 | -0.08196 | 0.2659 | -0.07129 | -0.1439 | -0.09807 | -0.02248 | 0.40256 | 0.164331 | -0.65848 |
| LS513 | -0.12379 | 0.116349 | -1.10943 | -0.36508 | -0.21198 | -0.11012 | -0.1689 | -0.11909 | -0.19118 | -0.08373 | -0.01141 | 0.05231 | -0.16202 | -0.20335 | -0.05385 | -0.07929 | -0.04427 | 0.09726 | 0.095967 | 0.09307 | 0.15816 | -0.12402 | 0.171672 | -0.00605 | 0.18206 | 0.027204 | -0.51781 |
| C2BBE1 | -0.10919 | -0.05868 | -0.4988 | -0.15364 | -0.04938 | -0.0916 | 0.015124 | -0.11003 | -0.15813 | -0.05256 | -0.05603 | 0.03344 | -0.30551 | -0.0817 | -0.04993 | -0.09143 | -0.03888 | 0.04618 | -0.0019 | 0.12234 | -0.05578 | -0.16287 | 0.042973 | -0.06365 | 0.19414 | 0.007965 | -0.557 |
| 253J | 0.01868 | 0.155799 | -0.12764 | 0.07565 | -0.10192 | -0.08176 | -0.11419 | -0.24775 | -0.0223 | -0.01233 | -0.06574 | -0.04234 | -0.08264 | -0.14808 | -0.23566 | -0.12408 | -0.11519 | 0.27724 | -0.02351 | 0.09459 | 0.03002 | -0.13047 | 0.195719 | -0.0831 | 0.03054 | -0.09235 | -0.22597 |
| HCC827 | -0.02762 | 0.01489 | -0.08191 | 0.05859 | -0.02018 | -0.1282 | -0.16145 | -0.17549 | -0.32667 | -0.13886 | -0.10499 | 0.09174 | -0.1364 | -0.16207 | -0.02676 | 0.052709 | -0.03777 | 0.18004 | -0.02485 | 0.11508 | 0.1333 | -0.23241 | 0.139586 | -0.05681 | 0.241 | -0.04421 | -0.33921 |
| ONCODG1 | -0.0122 | -0.03732 | -0.21503 | -0.04863 | -0.12579 | -0.11116 | 0.10689 | -0.18847 | -0.15585 | -0.08119 | -0.09901 | 0.17655 | 0.06912 | -0.11015 | -0.01727 | -0.09363 | -0.10332 | 0.20636 | -0.09559 | 0.05531 | 0.07817 | -0.14641 | 0.105576 | -0.02801 | 0.22868 | -0.02789 | -0.23645 |
| HS294T | -0.36449 | -0.02535 | -0.04549 | -0.14248 | -0.12543 | -0.02711 | -0.20571 | 0.10589 | -0.29701 | -0.03639 | 0.111825 | 0.09151 | -0.09296 | 0.07585 | -0.06292 | 0.084718 | -0.15032 | 0.29912 | -0.08059 | 0.12252 | 0.08078 | -0.07289 | 0.004705 | -0.04215 | 0.13524 | -0.00124 | -0.95331 |
| NCIH1581 | -0.03246 | 0.073085 | -0.35236 | -0.123 | -0.10674 | -0.21705 | -0.01233 | -0.22839 | -0.05327 | -0.06133 | -0.15334 | 0.09318 | -0.21454 | -0.16306 | -0.19666 | -0.18588 | -0.05809 | 0.19244 | -0.11402 | 0.02206 | 0.14044 | -0.20383 | 0.028533 | -0.05878 | 0.13948 | -0.20745 | -0.44493 |
| SKBR3 | -0.06912 | -0.00636 | -0.30797 | -0.04312 | -0.13785 | -0.27791 | -0.06834 | -0.31127 | -0.04801 | 0.05155 | 0.036583 | 0.15081 | -0.23511 | -0.08872 | -0.02867 | -0.078 | -0.07013 | 0.13362 | -0.12853 | 0.06055 | 0.05497 | -0.34879 | 0.064552 | 0.02459 | 0.17289 | 0.163038 | -0.51294 |
| T24 | -0.15494 | 0.045402 | -0.20423 | 0.18651 | -0.25694 | -0.30004 | 0.073495 | -0.16447 | -0.08711 | 0.11757 | -0.03851 | -0.07568 | -0.41907 | 0.03457 | -0.14074 | -0.09286 | -0.06536 | -0.02618 | -0.04192 | 0.29941 | 0.0121 | -0.1775 | -0.06449 | -0.22387 | 0.12705 | 0.038062 | -0.39279 |
| MCF7 | -0.01791 | -0.03989 | -0.42968 | -0.00627 | -0.08173 | -0.39994 | -0.01551 | -0.02184 | -0.03358 | 0.19112 | 0.099642 | 0.05991 | -0.35185 | 0.0279 | -0.00772 | -0.39947 | -0.022 | -0.01091 | -0.22456 | 0.069 | 0.06641 | -0.22455 | 0.095684 | -0.04383 | 0.17877 | 0.147695 | -0.17899 |
| NCIH1693 | -0.17495 | -0.00324 | -0.12086 | -0.20427 | -0.08662 | -0.12302 | -0.05082 | -0.11119 | -0.11758 | 0.00569 | -0.00674 | -0.00968 | -0.5478 | -0.07418 | -0.03619 | 0.000292 | -0.13115 | -0.09428 | 0.034215 | 0.09117 | 0.13439 | -0.29441 | 0.074453 | -0.02403 | 0.13309 | 0.08913 | -0.31682 |
| PATU8988S | -0.16933 | -0.08843 | -0.70908 | -0.22516 | -0.18153 | -0.34228 | -0.18127 | -0.09421 | -0.06902 | 0.05315 | -0.04265 | 0.0472 | -0.25544 | -0.07718 | -0.18958 | -0.11595 | -0.16995 | 0.06128 | -0.04735 | 0.03786 | 0.10324 | -0.14683 | 0.092702 | -0.11854 | 0.26851 | -0.13315 | -0.44766 |
| PATU8988T | -0.0713 | -0.03517 | -0.01548 | -0.0514 | -0.08271 | -0.09879 | 0.017458 | -0.08331 | -0.03704 | -0.163 | 0.018077 | -0.04154 | -0.22277 | -0.08146 | -0.13129 | -0.22178 | -0.08604 | -0.02137 | -0.07635 | 0.17188 | 0.06539 | -0.1318 | 0.044304 | -0.03214 | 0.09776 | 0.035379 | -0.3004 |
| OPM2 | -0.13806 | -0.15595 | -0.40324 | -0.06173 | -0.18163 | -0.03014 | 0.046846 | -0.31513 | 0.10378 | 0.06214 | -0.13217 | 0.08299 | -0.24336 | -0.17266 | -0.04751 | -0.00262 | -0.10223 | -0.06985 | -0.05913 | 0.17159 | 0.05213 | -0.11871 | 0.043768 | -0.12066 | 0.07739 | 0.041502 | -0.32701 |
| CH157MN | -0.08451 | 0.05195 | -0.09491 | -0.11827 | -0.24443 | -0.27641 | 0.0447 | -0.12546 | 0.086425 | -0.06302 | 0.097085 | 0.16899 | -0.16909 | -0.03698 | -0.11603 | -0.16753 | -0.17782 | 0.32889 | -0.04527 | 0.16822 | 0.19406 | -0.04593 | -0.0483 | 0.04763 | 0.27719 | 0.108461 | -0.49553 |
| KPL1 | -0.11957 | 0.00426 | -0.16313 | 0.0849 | -0.1544 | -0.11022 | -0.03799 | -0.15296 | -0.14374 | -0.01502 | 0.004336 | 0.02175 | -0.31994 | -0.19886 | -0.0491 | -0.12559 | -0.03118 | 0.24498 | -0.07991 | 0.14736 | 0.10271 | -0.11507 | -0.02395 | -0.14272 | 0.16657 | -0.05556 | -0.28363 |
| HCC827GR5 | -0.12664 | -0.04402 | -0.17981 | -0.08685 | -0.09585 | -0.27855 | 0.029095 | -0.02458 | -0.29134 | -0.00681 | -0.53822 | 0.1201 | -0.17138 | 0.07418 | -0.13068 | -0.15558 | 0.03946 | 0.33269 | 0.042535 | 0.19751 | 0.03416 | -0.19329 | 0.10059 | -0.05212 | 0.04647 | -0.09709 | -0.39642 |
| PC14 | -0.11793 | -0.00509 | -0.29093 | -0.03778 | -0.21206 | -0.23783 | -0.07767 | -0.17064 | -0.05957 | -0.11783 | -0.12412 | 0.10701 | -0.17358 | -0.07087 | -0.18109 | -0.26232 | -0.04136 | 0.14931 | -0.00533 | 0.05327 | 0.04402 | -0.19548 | 0.085827 | -0.1306 | 0.17557 | -0.01625 | -0.34497 |
| NCIH1650 | -0.32681 | -0.10779 | -0.05451 | 0.09062 | -0.10347 | -0.40084 | 0.010828 | -0.02108 | -0.20392 | -0.06704 | -0.08267 | -0.03046 | -0.34213 | -0.2152 | -0.12042 | -0.16498 | -0.03777 | 0.30128 | 0.017477 | 0.11832 | 0.06488 | -0.3026 | 0.017101 | -0.08393 | 0.09538 | 0.191506 | -0.56341 |
| U343 | -0.00725 | 0.000168 | -0.16228 | 0.16211 | -0.00293 | -0.17983 | -0.01656 | -0.35161 | 0.054344 | -0.07567 | -0.69172 | -0.01402 | -1.07488 | -0.50861 | 0.101735 | -0.09898 | -0.01723 | 0.02608 | -0.00361 | 0.07698 | 0.21249 | -0.32866 | 0.212138 | -0.05229 | 0.09593 | 0.010226 | -0.29343 |
| S117 | -0.24444 | -0.02404 | -0.35164 | -0.15766 | -0.25335 | -0.49513 | -0.13076 | -0.33161 | -0.18641 | -0.13659 | -0.07636 | 0.14067 | -0.48081 | -0.19727 | -0.03151 | -0.10093 | -0.04527 | 0.21738 | -0.22174 | -0.09359 | 0.19356 | -0.0279 | 0.157565 | -0.0375 | 0.11678 | -0.07587 | -0.38667 |
| SKNMC | -0.08025 | 0.072415 | -0.32108 | -0.56899 | -0.09501 | -0.32681 | -0.06237 | -0.26954 | -0.37539 | 0.10776 | -0.19299 | -0.00576 | -0.2717 | 0.08788 | -0.04795 | -0.10226 | -0.15411 | 0.13233 | -0.2151 | 0.03198 | 0.11425 | 0.088577 | -0.02617 | -0.01842 | 0.37977 | -0.0731 | -0.1492 |
| U118MG | -0.09928 | -0.02996 | -0.24405 | -0.06366 | -0.10429 | 0.02304 | -0.127 | -0.14254 | -0.03723 | 0.07079 | -0.34118 | 0.11093 | -0.82598 | -0.13339 | -0.20696 | -0.07086 | -0.01326 | 0.26636 | -0.1536 | 0.09774 | 0.07934 | -0.15536 | 0.085535 | 0.04109 | 0.33034 | 0.048683 | -0.27497 |
| RDES | -0.20755 | 0.082184 | -0.53793 | -0.02202 | -0.22242 | -0.13636 | -0.09732 | -0.11744 | -0.13088 | -0.15805 | -0.03089 | -0.01609 | -0.24247 | -0.21728 | 0.067458 | -0.07003 | -0.11458 | 0.34518 | 0.000725 | 0.22887 | 0.00652 | -0.22337 | -0.0046 | -0.0037 | 0.11978 | -0.07343 | -0.44519 |
| PANC0203 | -0.16117 | 0.056105 | -0.56645 | -0.07764 | -0.11812 | -0.25419 | -0.08913 | -0.32326 | -0.19258 | -0.07041 | -0.03088 | 0.06683 | -0.1999 | -0.09936 | -0.22694 | -0.14561 | 0.05745 | 0.03668 | -0.00806 | 0.19444 | 0.12947 | -0.03701 | 0.138488 | -0.02538 | 0.07702 | -0.05262 | -0.58041 |
| MV411 | -1.05165 | -0.05472 | -0.1066 | 0.13855 | -0.14542 | -0.12283 | 0.040003 | -0.26681 | 0.010858 | 0.07847 | -0.06045 | 0.02135 | -0.12317 | -0.29073 | -0.16014 | -0.09354 | -0.01662 | 0.18129 | 0.024349 | 0.07548 | 0.29009 | -0.12004 | 0.111486 | -0.02752 | 0.12185 | -0.04267 | -0.51772 |
| GCIY | -0.04449 | -0.01557 | -0.60106 | -0.19905 | -0.03944 | -0.0585 | -0.03891 | -0.27784 | -0.23008 | 0.08568 | -0.08983 | 0.11681 | -0.26203 | -0.06102 | -0.00205 | -0.11207 | 0.08485 | -0.0149 | 0.096597 | 0.21522 | 0.07543 | -0.00646 | 0.11394 | -0.02758 | 0.14102 | 0.009508 | -0.18104 |
| TOV112D | 0.00599 | -0.03326 | 0.0777 | 0.0973 | -0.22338 | -0.35354 | -0.13231 | -0.23871 | -0.24859 | 0.04222 | 0.020217 | 0.04757 | -0.33581 | -0.13366 | -0.05874 | -0.39981 | -0.14673 | 0.02597 | 0.015909 | 0.07951 | 0.05902 | -0.29086 | -0.00898 | -0.08945 | 0.14498 | -0.05172 | -0.45683 |
| A673 | -0.21672 | -0.01439 | -0.60501 | 0.09268 | -0.2169 | 0.25596 | 0.188314 | -0.087 | 0.020845 | 0.00325 | -0.25563 | 0.19563 | -0.14712 | 0.07288 | 0.036649 | -0.07191 | -0.07277 | 0.49235 | -0.18176 | 0.28468 | -0.01622 | -0.24698 | 0.024974 | 0.23813 | 0.23685 | -0.03917 | -0.29595 |
| KARPAS299 | -0.10189 | -0.01578 | -0.1838 | -0.04851 | -0.10684 | -0.25027 | 0.052158 | -0.14886 | -0.18279 | -0.01742 | -0.03166 | -0.00839 | -0.13968 | 0.08869 | -0.08955 | -0.05297 | -0.1627 | 0.07332 | -0.1247 | 0.09777 | 0.09295 | -0.17259 | 0.009187 | 0.02698 | 0.22693 | 0.023476 | -0.21327 |
| HT1080 | -0.28709 | 0.049895 | -0.17773 | -0.01628 | -0.11972 | 0.01648 | 0.021316 | -0.01806 | 0.071739 | 0.13548 | -0.04792 | 0.12186 | -0.21695 | -0.14476 | -0.07228 | -0.03949 | 0.00172 | 0.03039 | 0.079753 | 0.26201 | 0.17415 | -0.30626 | 0.11936 | 0.02277 | 0.04229 | 0.018326 | -0.35491 |
| D283MED | -0.20308 | 0.240921 | -0.18184 | -0.02802 | -0.14203 | -0.25639 | 0.047365 | -0.279 | -0.07997 | -0.14986 | -0.1241 | 0.09857 | -0.09891 | -0.06234 | -0.16314 | -0.24267 | -0.13562 | 0.17982 | -0.13976 | 0.04799 | 0.23777 | -0.07063 | 0.118455 | -0.02912 | 0.16475 | -0.08112 | -0.32851 |
| PANC1005 | -0.13796 | 0.05347 | -0.5876 | 0.08359 | -0.28906 | -0.40799 | -0.18174 | 0.08648 | -0.05668 | 0.05929 | -0.0524 | 0.20287 | -0.50662 | -0.15631 | -0.08092 | -0.11989 | -0.08983 | 0.27356 | -0.00834 | 0.12642 | 0.08844 | -0.00889 | 0.11413 | -0.08993 | -0.03604 | -0.15688 | -0.28377 |
| HS683 | -0.11851 | -0.03914 | -0.14793 | -0.04703 | -0.16213 | -0.14918 | -0.05956 | -0.20953 | -0.15402 | 0.06448 | -0.68443 | 0.20743 | -0.3606 | -0.24756 | -0.09282 | -0.10501 | -0.13795 | 0.07852 | -0.03917 | 0.17877 | 0.08026 | -0.16635 | 0.027748 | -0.03525 | 0.19221 | -0.11131 | -0.21771 |
| 697 | -0.1578 | -0.04118 | -0.58605 | 0.03961 | -0.06737 | -0.36656 | 0.191965 | -0.1659 | -0.11656 | 0.10084 | -0.0268 | 0.15682 | 0.07379 | -0.43866 | 0.171265 | -0.08157 | -0.06468 | 0.03034 | -0.19339 | -0.00984 | 0.34672 | 0.089236 | 0.017513 | 0.13691 | 0.24927 | 0.189967 | -0.15256 |
| KU812 | -0.08916 | 0.057935 | -0.19384 | -0.02504 | 0.11591 | 0.00606 | -0.0427 | -0.30699 | 0.0646 | -0.24771 | -0.25046 | 0.1294 | -0.13453 | 0.12535 | -0.22602 | -0.04967 | 0.14503 | -0.23114 | 0.057386 | 0.16949 | 0.14325 | -0.15352 | -0.01181 | -0.07739 | 0.20427 | -0.15173 | -0.5099 |
| U87MG | -0.17912 | -0.08537 | -0.18972 | -0.02143 | -0.19436 | -0.13881 | 0.013722 | -0.43637 | 0.002827 | 0.06867 | -0.76987 | 0.117 | -0.05917 | -0.04727 | 0.026026 | -0.05542 | -0.08877 | 0.02973 | -0.04853 | 0.25233 | 0.20983 | -0.09022 | 0.217432 | -0.04543 | 0.27466 | -0.07055 | -0.15634 |
| NCO2 | 0.067 | 0.069205 | -0.31024 | -0.09729 | 0.04292 | 0.1841 | 0.140246 | -0.21616 | -0.07041 | 0.27605 | -0.37415 | 0.04865 | -0.23863 | 0.01129 | -0.05998 | -0.05212 | -0.32627 | 0.29251 | -0.06075 | -0.09544 | -0.04804 | 0.024041 | -0.15547 | 0.0193 | 0.2915 | 0.293107 | -0.34854 |
| MJ | -0.33491 | -0.23093 | -0.26451 | 0.14722 | -0.23062 | 0.02046 | 0.254951 | -0.18038 | -0.33997 | -0.02273 | -0.24688 | 0.22865 | -0.34594 | -0.15614 | 0.026498 | -0.20085 | -0.16207 | -0.06287 | 0.211599 | -0.05712 | 0.02488 | -0.20231 | -0.05161 | -0.01413 | 0.1435 | -0.21263 | -0.27366 |
| MHHNB11 | -0.30196 | 0.057302 | -0.44578 | -0.18998 | -0.21211 | -0.18074 | -0.12948 | -0.4694 | -0.0683 | -0.04453 | 0.075265 | 0.07158 | -0.17196 | -0.01302 | -0.11798 | -0.10312 | -0.01051 | 0.31949 | -0.02121 | 0.06636 | 0.18534 | -0.34458 | -0.04921 | -0.12685 | 0.09283 | 0.058202 | -0.42463 |
| G292CLONEA141B1 | 0.04514 | -0.00362 | -0.1945 | -0.09851 | -0.21412 | -0.15825 | -0.01213 | 0.03218 | -0.02307 | -0.10309 | -0.14452 | -0.02629 | -0.40758 | -0.12905 | 0.062384 | -0.06077 | -0.09556 | 0.18824 | -0.03236 | 0.14038 | 0.17223 | -0.09023 | -0.00672 | -0.09158 | 0.02369 | -0.01182 | -0.29676 |
| T3M4 | -0.08471 | 0.109647 | -0.37306 | -0.02665 | -0.14329 | -0.15024 | -0.18113 | -0.17366 | -0.22256 | -0.06749 | -0.00729 | 0.02662 | -0.00957 | 0.06219 | -0.06347 | -0.03296 | 0.07316 | 0.12205 | -0.21532 | 0.03966 | 0.14262 | -0.00669 | -0.0408 | -0.31835 | 0.2139 | -0.09729 | -0.03885 |
| ACCMESO1 | -0.18272 | 0.005838 | -0.08293 | -0.00803 | -0.06432 | -0.2336 | 0.063341 | -0.09596 | -0.08879 | -0.13978 | -0.09152 | 0.02123 | -0.05574 | -0.11125 | -0.05985 | -0.225 | -0.08461 | 0.00284 | -0.06256 | 0.00959 | 0.08899 | -0.23326 | 0.050575 | -0.15161 | 0.00593 | 0.127977 | -0.2529 |
| PC3 | -0.17071 | -0.0288 | -0.10684 | 0.16244 | -0.00158 | -0.45717 | -0.1649 | -0.26151 | -0.12558 | 0.02518 | -0.04445 | -0.06283 | -0.57062 | -0.01244 | -0.19362 | -0.27268 | -0.02034 | -0.2326 | -0.15545 | 0.12792 | -0.01984 | 0.07618 | 0.24269 | -0.00711 | 0.3097 | 0.361635 | -0.20317 |
| NCIH2452 | -0.1518 | -0.00914 | -0.15769 | -0.03023 | -0.22514 | -0.1594 | 0.066877 | -0.17353 | -0.17183 | 0.02577 | -0.11063 | 0.07024 | -0.20377 | -0.18716 | -0.12792 | -0.2388 | -0.01493 | 0.16122 | -0.09117 | 0.11274 | 0.02927 | -0.24094 | 0.080265 | -0.13489 | 0.20936 | 0.09406 | -0.38072 |
| PANC0504 | 0.01746 | -0.08161 | -0.29616 | -0.19225 | -0.22784 | 0.09361 | 0.020359 | -0.34054 | 0.093678 | -0.17158 | -0.0545 | 0.08521 | -0.34027 | -0.21579 | -0.04634 | -0.07423 | -0.06505 | 0.01914 | -0.05593 | 0.30929 | 0.04678 | -0.55515 | -0.08058 | 0.11206 | 0.03426 | 0.166161 | -0.662 |
| HPAFII | 0.05798 | -0.04806 | -0.43517 | 0.10817 | -0.09405 | -0.23088 | 0.000328 | -0.15251 | -0.12237 | 0.08543 | 0.020108 | 0.40232 | -0.23586 | -0.17176 | -0.1456 | -0.14374 | 0.01567 | -0.00232 | -0.20567 | 0.32755 | 0.20798 | 0.015111 | 0.199141 | 0.12925 | 0.05829 | -0.03418 | 0.14456 |
| D341Med | -0.09992 | 0.030095 | -0.56687 | -0.10658 | -0.16764 | -0.01852 | 0.003942 | -0.09407 | -0.2232 | -0.01753 | -0.04823 | 0.0679 | -0.02422 | 0.03837 | -0.09012 | 0.008667 | -0.06602 | -0.01837 | -0.04224 | 0.1337 | 0.08335 | -0.14838 | 0.001061 | 0.03025 | 0.2348 | 0.017004 | -0.46943 |
| ZR751 | -0.03483 | 0.090414 | -0.25743 | -0.04167 | -0.0629 | -0.14499 | -0.05681 | -0.1916 | -0.02096 | -0.07625 | -0.08902 | 0.02792 | -0.05941 | -0.2016 | -0.15329 | -0.0539 | -0.07709 | 0.0977 | -0.06432 | 0.1557 | 0.26801 | 0.034878 | 0.14942 | -0.08138 | 0.08077 | -0.09046 | -0.43512 |
| GAMG | -0.08427 | -0.08819 | -0.25674 | -0.09664 | -0.11429 | -0.16334 | 0.025525 | -0.18027 | -0.08492 | 0.009 | -0.48312 | -0.02885 | -0.28108 | -0.17188 | -0.09053 | -0.05222 | -0.09965 | 0.09476 | -0.04358 | 0.06323 | 0.09131 | -0.25175 | 0.132749 | -0.00911 | 0.12012 | 0.079849 | -0.25412 |
| SIMA | -0.08196 | 0.030379 | -0.64229 | -0.06133 | -0.06097 | 0.00848 | -0.11073 | -0.40622 | -0.16342 | 0.02995 | 0.049495 | 0.01802 | -0.22973 | -0.04465 | -0.03092 | -0.14461 | -0.04837 | 0.147 | -0.01527 | 0.1165 | 0.14513 | -0.2352 | 0.120536 | -0.04179 | 0.14804 | -0.02988 | -0.56636 |
| KE37 | -0.09257 | -0.06056 | -0.09675 | -0.0612 | -0.14046 | -0.03744 | 0.081854 | -0.21133 | -0.23047 | 0.04815 | -0.02267 | 0.05943 | -0.18801 | -0.15289 | -0.03638 | -0.1821 | 0.02643 | 0.10336 | 0.071404 | 0.07883 | 0.06694 | -0.10813 | -0.07784 | -0.02825 | 0.24395 | -0.03788 | -0.49134 |
| CAOV4 | -0.05656 | -0.21325 | -0.04047 | -0.29757 | -0.2012 | -0.29403 | -0.10996 | -0.42247 | -0.30287 | -0.16437 | 0.136114 | 0.13431 | -0.44744 | -0.57565 | -0.01964 | -0.08845 | 0.01662 | 0.21028 | -0.26582 | 0.29554 | 0.06965 | -0.13968 | -0.0244 | -0.04575 | 0.16305 | 0.162565 | -0.60632 |
| KP3 | -0.03902 | 0.093844 | -0.28403 | 0.00245 | -0.09249 | -0.19105 | -0.02213 | -0.15555 | -0.13697 | -0.16027 | -0.0574 | 0.12902 | -0.26017 | 0.02669 | -0.15287 | -0.10105 | -0.08873 | 0.05711 | -0.01668 | 0.06638 | 0.10749 | -0.15188 | 0.070216 | -0.23166 | 0.14668 | -0.09746 | -0.54367 |
| HCC1187 | -0.00315 | -0.11154 | 0.24851 | -0.11248 | -0.0739 | -0.1682 | -0.07709 | -0.12763 | -0.0496 | -0.10057 | -0.00828 | 0.12051 | -0.48422 | -0.16912 | 0.126706 | -0.13932 | 0.13585 | 0.02453 | 0.068523 | 0.11972 | 0.1215 | -0.11215 | 0.121291 | -0.13863 | 0.20561 | 0.14754 | -0.41613 |
| OCIAML2 | -0.17143 | -0.10151 | -0.41042 | -0.20485 | -0.01473 | -0.0779 | 0.03751 | -0.2053 | -0.04779 | -0.10671 | 0.00186 | -0.05336 | -0.11573 | -0.02078 | 0.014661 | -0.12236 | -0.00687 | 0.00569 | -0.05359 | 0.06211 | 0.11421 | -0.1505 | -6.1E-05 | 0.00617 | 0.20625 | 0.099085 | -0.3802 |
| SU8686 | -0.15145 | 0.08959 | -0.24066 | 0.06978 | -0.07757 | 0.03441 | -0.0069 | -0.06687 | -0.04991 | -0.17975 | -0.15806 | 0.01959 | -0.27008 | -0.19948 | -0.38175 | -0.16568 | 0.00096 | 0.00403 | -0.00785 | 0.08396 | 0.09171 | -0.01666 | 0.103319 | -0.06547 | 0.12978 | 0.060561 | -0.31216 |
| VCAP | -0.25379 | -0.15385 | -0.90326 | -0.26276 | -0.04656 | -0.15477 | 0.053825 | -0.2126 | 0.101636 | -0.0784 | 0.029525 | 0.05779 | -0.25557 | -0.06507 | -0.14509 | -0.19345 | -0.09636 | 0.12879 | -0.03992 | -0.00782 | 0.15136 | -0.20851 | 0.151932 | 0.03369 | 0.18656 | 0.194359 | -0.40404 |
| HUPT3 | -0.10402 | 0.064003 | -0.20974 | -0.13768 | -0.1984 | -0.21335 | -0.10916 | -0.20785 | 0.027452 | -0.15157 | -0.06003 | 0.14765 | -0.06232 | 0.0224 | -0.1355 | -0.04707 | -0.05839 | -0.04029 | -0.07003 | 0.12819 | 0.1532 | -0.12685 | 0.046736 | -0.07885 | 0.0901 | -0.07789 | -0.27818 |
| CHP212 | -0.06798 | -0.01938 | -0.31213 | -0.22639 | 0.01177 | -0.08231 | -0.11149 | -0.0652 | -0.12883 | 0.06274 | -0.04924 | 0.00399 | -0.01509 | 0.03789 | -0.11907 | -0.00912 | -0.02949 | 0.1875 | -0.04357 | -0.04514 | 0.13096 | -0.24292 | 0.264759 | -0.07059 | 0.17245 | -0.06318 | -0.47919 |
| COV434 | -0.0498 | -0.04857 | -0.01609 | -0.15001 | -0.11368 | 0.00879 | 0.055173 | -0.13552 | -0.19868 | 0.03918 | -0.07562 | 0.10648 | -0.09106 | -0.01537 | -0.19897 | -0.06607 | -0.0487 | 0.12525 | -0.10297 | 0.04934 | 0.04258 | -0.13127 | -0.04516 | 0.00437 | 0.28559 | -0.01953 | -0.42815 |
| OCILY19 | -0.11758 | -0.09407 | -0.65094 | -0.15467 | -0.21693 | -0.28714 | 0.106659 | -0.27564 | -0.18686 | -0.19373 | 0.06759 | 0.20922 | -0.03102 | -0.205 | -0.39788 | -0.12038 | -0.13635 | 0.05559 | -0.01937 | 0.31298 | -0.00586 | -0.19304 | 0.124654 | -0.10685 | 0.27573 | -0.0637 | -0.54735 |
| SLR20 | -0.05066 | 0.020191 | -0.31248 | -0.23713 | -0.17922 | -0.14124 | 0.008434 | -0.1776 | 0.01172 | 0.15716 | -0.13095 | 0.08936 | -0.29839 | -0.08814 | -0.15597 | -0.12106 | -0.04077 | 0.11838 | 0.00304 | 0.07168 | 0.12397 | -0.09954 | 0.034753 | -0.07327 | 0.13937 | -0.23129 | -0.41868 |
| LN319 | -0.24717 | -0.06835 | -0.13979 | 0.04819 | -0.15596 | -0.13029 | -0.06205 | -0.16324 | -0.05363 | 0.20913 | 0.011267 | -0.05129 | -0.27163 | -0.16142 | 0.165189 | -0.18074 | -0.11109 | 0.11602 | 0.126885 | 0.17735 | 0.17545 | 0.048493 | -0.08692 | -0.0633 | 0.19533 | -0.08791 | -0.55836 |
| JHOS2 | -0.03352 | 0.015081 | -0.00561 | 0.10293 | -0.06463 | -0.26441 | -0.13117 | -0.14931 | -0.08666 | -0.0675 | -0.03828 | 0.08741 | -0.27909 | -0.06375 | -0.06831 | -0.10325 | -0.1401 | -0.0821 | -0.03668 | 0.20513 | 0.11239 | -0.22182 | 0.097332 | 0.02801 | 0.15237 | 0.022859 | -0.32645 |
| HS729 | -0.03913 | -0.02693 | -0.23816 | -0.28237 | -0.15078 | -0.05255 | -0.01828 | -0.11732 | 0.025193 | 0.06068 | -0.1271 | 0.09476 | -0.31456 | -0.03472 | -0.06825 | -0.12238 | -0.1299 | 0.0722 | -0.04477 | 0.13359 | -0.0728 | -0.009 | 0.121053 | 0.01353 | 0.10106 | 0.004501 | -0.27378 |
| 8MGBA | 0.02735 | 0.078543 | -0.3649 | -0.10983 | -0.14878 | -0.22003 | 0.150184 | -0.11182 | -0.22163 | -0.06543 | -0.03697 | 0.18053 | -0.15082 | -0.17913 | -0.06338 | -0.05806 | -0.13833 | 0.07277 | -0.03083 | 0.1805 | -0.00153 | -0.2944 | -0.03868 | -0.11184 | 0.2185 | -0.01146 | -0.21258 |
| CFPAC1 | -0.20302 | -0.03754 | -0.23545 | -0.01366 | -0.1059 | -0.11157 | -0.01697 | -0.03965 | 0.06166 | -0.01389 | -0.08933 | 0.0729 | -0.03954 | -0.21639 | -0.04353 | 0.066445 | 0.16098 | -0.00464 | 0.018163 | 0.08596 | 0.18123 | -0.22527 | -0.11428 | 0.00345 | 0.12999 | 0.003179 | -0.29614 |
| PANC0327 | 0.13243 | 0.057192 | 0.19065 | 0.06281 | -0.20893 | -0.10345 | 0.064652 | -0.02899 | -0.10069 | -0.15901 | -0.06733 | -0.12705 | -0.24584 | -0.12143 | -0.11129 | -0.08696 | 0.06373 | 0.38004 | -0.10558 | 0.20253 | 0.03606 | -0.24386 | 0.175332 | 0.01057 | 0.2437 | 0.094017 | -0.41889 |
| SNU308 | 0.08982 | 0.110746 | -0.16381 | -0.26079 | -0.25887 | -0.15776 | 0.232296 | -0.22334 | -0.27814 | -0.10521 | -0.01027 | 0.21103 | 0.14701 | -0.00932 | -0.1245 | -0.05618 | -0.20119 | 0.2249 | -0.17921 | 0.04906 | 0.06458 | -0.33507 | 0.153954 | -0.24998 | 0.32758 | -0.05044 | -0.37345 |
| CAL29 | -0.1115 | 0.135123 | -0.25552 | -0.01656 | -0.13044 | -0.16974 | -0.03289 | -0.44894 | -0.2649 | -0.08293 | -0.00124 | 0.04432 | -0.23301 | -0.15786 | -0.16397 | 0.014691 | -0.06951 | 0.14361 | -0.04585 | -0.01283 | 0.08678 | -0.22917 | 0.067804 | -0.21334 | 0.14899 | 0.099107 | -0.22406 |

| Cell line | FLT3 | FLT4 | IGF1R | INSR | INSRR | KDR | KIT | LMR1 | LMR2 | LTK | MET | MUSK | PDGFRA | PDGFRB | PTK7 | RET | RON | ROR1 | ROR2 | ROS1 | STYK1 | TIE1 | TIE2 | TRKA | TRKB | TRKC | TYRO3 |
|---|---|---|---|---|---|---|---|---|---|---|---|---|---|---|---|---|---|---|---|---|---|---|---|---|---|---|---|
| RERFGC1B | -0.19715 | -0.09578 | -1.57138 | -0.12328 | -0.33408 | -0.24805 | -0.30775 | 0.00499 | -0.43594 | -0.3694 | -0.3607 | 0.12103 | -0.20991 | -0.004 | -0.08352 | -0.29054 | -0.09833 | -0.06919 | 0.067124 | -0.06747 | -0.08564 | -0.4409 | 0.119136 | -0.26569 | 0.24053 | 0.186992 | -0.13012 |
| SKLMS1 | -0.1944 | -0.00363 | -0.2675 | -0.1324 | -0.12458 | -0.05115 | 0.004043 | -0.36777 | -0.16764 | 0.0231 | -0.14629 | 0.10375 | -0.40016 | -0.1268 | -0.09297 | -0.01219 | -0.17605 | -0.07054 | -0.05215 | 0.11068 | 0.17221 | -0.20814 | 0.182446 | -0.07453 | 0.16811 | 0.021978 | -0.39338 |
| THP1 | -0.05911 | 0.016431 | -0.11277 | -0.20209 | -0.02298 | -0.23347 | 0.03067 | -0.0627 | 0.051429 | 0.0402 | -0.15475 | 0.03739 | -0.21571 | 0.00337 | -0.03439 | -0.07113 | -0.13362 | -0.06308 | -0.00955 | 0.11473 | 0.13616 | -0.05041 | 0.082889 | -0.02463 | 0.06864 | -0.04406 | -0.40607 |
| T47D | -0.03881 | 0.003043 | -0.23445 | -0.12707 | -0.09101 | -0.05439 | 0.041786 | -0.20415 | -0.00847 | 0.05737 | -0.13254 | 0.19634 | -0.29821 | -0.16395 | -0.11998 | 0.104419 | 0.02057 | -0.08669 | 0.020295 | 0.14825 | 0.17184 | -0.13559 | 0.068198 | -0.05094 | 0.25539 | 0.052167 | -0.42562 |
| HS578T | 0.05718 | 0.009492 | -0.05515 | 0.03328 | -0.15735 | -0.30092 | -0.04109 | -0.24246 | -0.18267 | -0.15872 | -0.10156 | 0.05206 | -0.47628 | -0.48636 | -0.1238 | 0.013852 | 0.05986 | 0.17597 | -0.07095 | 0.08307 | 0.23818 | -0.07373 | 0.003958 | -0.05285 | 0.17656 | -0.05699 | -0.40392 |
| SKNSH | -0.07355 | -0.224 | -0.72342 | -0.05389 | -0.13315 | -0.26591 | -0.05544 | 0.05384 | 0.140964 | -0.02993 | -0.11051 | 0.31814 | 0.12318 | -0.3874 | 0.007516 | -0.07621 | 0.00905 | -0.04145 | 0.070738 | 0.37939 | 0.142 | -0.19336 | 0.244671 | 0.2387 | 0.1642 | -0.09339 | -0.55422 |
| HCC2935 | -0.15519 | 0.022257 | -0.21664 | -0.07113 | -0.13132 | -0.05132 | -0.00027 | -0.02746 | -0.0709 | -0.34191 | 0.039452 | 0.07945 | -0.36475 | -0.1717 | -0.2043 | 0.085412 | -0.27593 | 0.44553 | -0.05912 | 0.07306 | 0.0452 | -0.07304 | 0.111365 | -0.14744 | 0.11178 | -0.07616 | -0.482 |
| JM1 | 0.04568 | 0.048286 | -0.66806 | -0.05642 | -0.13276 | -0.23217 | -0.09415 | 0.10525 | -0.11965 | 0.01946 | -0.02003 | 0.1227 | -0.20924 | -0.05045 | -0.0528 | -0.10272 | -0.03094 | 0.06756 | -0.04463 | 0.27859 | 0.15072 | -0.31618 | -0.08593 | 0.02198 | 0.18028 | 0.045168 | -0.27688 |
| M059K | -0.02761 | -0.0983 | -0.33464 | 0.09196 | -0.23595 | -0.14223 | -0.0864 | -0.13374 | 0.009834 | -0.11775 | -0.11869 | 0.1091 | -0.23946 | -0.17949 | -0.30666 | -0.03647 | -0.07811 | 0.15778 | -0.07213 | 0.05375 | 0.25026 | -0.18635 | 0.15124 | 0.06648 | 0.13923 | -0.07517 | -0.18409 |
| NCIH2052 | -0.42736 | 0.218368 | -0.36529 | 0.13021 | -0.11915 | -0.38038 | 0.125905 | 0.03509 | 0.241279 | -0.23824 | -0.31305 | -0.04144 | -0.2578 | 0.324 | -0.136 | -0.15433 | -0.15784 | 0.28302 | 0.221644 | 0.07467 | 0.11148 | -0.08271 | 0.286925 | 0.14627 | 0.13242 | 0.064139 | -0.48559 |
| SW1990 | -0.06402 | -0.02086 | -0.14754 | 0.02758 | 0.01827 | -0.1705 | 0.049887 | -0.28219 | 0.024502 | -0.05822 | 0.02904 | 0.07684 | -0.20533 | 0.07762 | 0.010924 | -0.14715 | -0.15472 | 0.05834 | -0.16026 | 0.27392 | 0.07145 | -0.11852 | 0.156414 | -0.12134 | 0.08469 | 0.037294 | -0.23432 |
| OSRC2 | -0.27505 | -0.04523 | -0.21453 | -0.05196 | -0.08619 | -0.09165 | -0.04607 | -0.208 | -0.03844 | 0.11871 | -0.12797 | 0.03334 | -0.09771 | -0.10063 | -0.14849 | -0.03286 | -0.29543 | 0.21619 | -0.03587 | 0.17003 | 0.18395 | 0.020341 | 0.147638 | -0.22497 | 0.02542 | -0.14268 | -0.27437 |
| BT12 | -0.01423 | 0.070908 | -0.18719 | -0.15524 | -0.01657 | -0.12471 | 0.088743 | -0.21752 | -0.03163 | -0.1241 | -0.05916 | 0.21848 | 0.06031 | -0.005 | -0.02447 | -0.10808 | 0.02967 | 0.03592 | -0.17089 | 0.17399 | 0.15703 | -0.11747 | -0.12359 | -0.02965 | 0.08651 | 0.015417 | -0.47303 |
| CORL105 | -0.25197 | -0.07501 | -0.37084 | -0.07607 | -0.28322 | 0.02645 | 0.008939 | -0.23647 | -0.07303 | -0.1448 | -0.07886 | 0.16061 | -0.09038 | -0.05653 | -0.13302 | -0.12829 | 0.10808 | -0.09918 | 0.005344 | 0.13094 | 0.20599 | -0.05604 | 0.05197 | -0.03939 | 0.06718 | -0.04533 | -0.21527 |
| SW579 | -0.21143 | -0.05877 | -0.17683 | 0.04601 | -0.15434 | -0.1183 | 0.079621 | -0.13022 | 0.143904 | -0.05642 | 0.04666 | -0.01952 | -1.67159 | -0.07957 | -0.00455 | -0.07319 | -0.23411 | 0.09802 | 0.037838 | 0.32839 | 0.25378 | -0.1571 | 0.036296 | -0.10277 | 0.05157 | -0.10318 | -0.22288 |
| PANC1 | -0.13096 | 0.069003 | -0.276 | -0.04296 | -0.08516 | -0.16988 | -0.10181 | -0.14876 | 0.016073 | -0.04203 | -0.07655 | 0.13435 | -0.20431 | -0.02 | -0.08646 | -0.11547 | 0.0205 | 0.02402 | -0.05316 | 0.06097 | 0.04558 | -0.12455 | 0.163528 | -0.09479 | 0.17364 | 0.028597 | -0.32222 |
| NOMO1 | -0.46392 | -0.14399 | -0.28269 | -0.11185 | -0.13347 | -0.43081 | -0.01572 | 0.0131 | 0.023217 | 0.1422 | 0.023585 | 0.19094 | -0.24682 | -0.14221 | -0.08873 | -0.13672 | -0.06748 | 0.38123 | -0.08824 | 0.08982 | 0.33877 | -0.41173 | 0.299185 | -0.09589 | 0.40337 | 0.135117 | -0.68363 |
| RD | -0.20005 | 0.128482 | -0.69423 | -0.00086 | -0.10733 | -0.09351 | -0.1 | -0.24207 | -0.1598 | -0.06344 | -0.07901 | 0.08037 | -0.03455 | -0.00459 | -0.09375 | 0.024659 | -0.07775 | 0.09817 | 0.085808 | 0.01652 | 0.13044 | -0.13677 | 0.005588 | -0.11817 | 0.14098 | -0.00545 | -0.4628 |
| CAL62 | -0.0101 | 0.078156 | -0.36241 | -0.14075 | -0.12566 | -0.12791 | -0.00796 | -0.21944 | -0.16314 | -0.0356 | -0.16538 | 0.02946 | -0.14648 | -0.18407 | -0.21734 | -0.0637 | -0.01425 | 0.11045 | -0.03186 | 0.13174 | 0.04169 | -0.18831 | 0.059433 | -0.15897 | 0.07932 | -0.03023 | -0.33007 |
| LOUNH91 | -0.25786 | -0.29904 | -0.19036 | -0.23733 | -0.2626 | -0.38597 | -0.20063 | -0.30385 | -0.24262 | -0.07901 | -0.09574 | 0.12113 | -0.41998 | -0.21375 | -0.06379 | -0.23733 | -0.04974 | 0.15545 | -0.04608 | 0.02735 | 0.16432 | -0.21698 | -0.10122 | -0.02521 | 0.21503 | -0.08363 | -0.4282 |
| HS766T | 8.4E-05 | -0.00912 | -0.1696 | 0.09099 | -0.17436 | -0.10394 | -0.02415 | -0.26661 | 0.075297 | -0.06005 | -0.07193 | 0.08624 | -0.13068 | -0.14769 | -0.08275 | -0.06182 | -0.08184 | 0.15945 | -0.23422 | 0.08051 | 0.17446 | -0.10579 | 0.008628 | -0.09384 | 0.01757 | -0.032 | -0.53402 |
| SCC9 | 0.00673 | 0.203457 | -0.1314 | -0.06 | 0.12305 | -0.19452 | -0.03145 | 0.03434 | -0.07937 | 0.10079 | -0.09387 | 0.12614 | -0.19102 | -0.03693 | 0.102336 | -0.00428 | 0.04002 | 0.18651 | 0.114765 | 0.27816 | 0.15726 | -0.04829 | 0.114751 | 0.00114 | 0.06056 | 0.221706 | -0.34516 |
| SNU869 | -0.12725 | 0.076171 | -0.66369 | -0.0494 | -0.04886 | -0.13445 | -0.07168 | -0.16475 | 0.014827 | 0.04124 | -0.10126 | 0.18563 | -0.02591 | -0.06939 | 0.001962 | -0.03564 | -0.14982 | -0.06391 | -0.01633 | 0.12665 | 0.11633 | -0.26066 | 0.08193 | -0.10378 | 0.04127 | -0.11481 | -0.24506 |
| L363 | -0.10903 | -0.05812 | -0.74174 | -0.17163 | 0.07331 | -0.11949 | -0.14797 | -0.23323 | -0.06611 | 0.0835 | -0.20055 | 0.08804 | -0.17185 | -0.08795 | 0.027868 | -0.06589 | -0.16 | 0.27994 | 0.019657 | 0.10211 | 0.11228 | -0.06229 | 0.015793 | -0.13519 | -0.00473 | -0.00364 | -0.5368 |
| CORL311 | -0.07097 | 0.071118 | -0.78641 | -0.21261 | -0.1547 | -0.24365 | -0.1582 | -0.21855 | -0.38489 | -0.20872 | 0.038419 | 0.07407 | -0.74884 | 0.07274 | -0.02022 | 0.091799 | -0.03588 | 0.23027 | -0.13831 | 0.10387 | 0.07614 | -0.12026 | 0.013733 | 0.05462 | 0.29286 | -0.04887 | -0.64961 |
| SCC25 | -0.14236 | -0.03828 | -0.14269 | 0.03411 | -0.21418 | -0.16456 | 0.037792 | -0.14584 | -0.08556 | -0.08878 | -0.0733 | 0.09933 | -0.33808 | -0.13268 | -0.18891 | -0.15942 | 0.00032 | 0.01554 | -0.06987 | 0.1306 | 0.08437 | -0.36688 | 0.104193 | -0.15243 | 0.00268 | -0.01105 | -0.46288 |
| RCC10RGB | -0.24645 | 0.160334 | -0.30635 | -0.04775 | -0.09445 | -0.24587 | 0.082071 | -0.15966 | -0.11038 | -0.06205 | -0.24861 | -0.16719 | -0.1636 | 0.23246 | -0.00717 | 0.089397 | -0.11713 | -0.0448 | -0.0801 | 0.30859 | 0.38697 | -0.24271 | 0.115304 | -0.08073 | 0.30202 | -0.11981 | -0.41301 |
| HDMYZ | -0.01792 | -0.12048 | 0.0447 | -0.07222 | -0.1835 | -0.04181 | 0.004476 | -0.05848 | -0.20359 | -0.02399 | -0.13009 | 0.16558 | -0.22303 | -0.02352 | 0.013179 | -0.04319 | -0.36368 | -0.02937 | -0.21741 | -0.08266 | 0.05062 | -0.36912 | 0.196497 | 0.00122 | -0.02965 | 0.145087 | -0.30476 |
| BHT101 | -0.25486 | -0.01892 | -0.27 | -0.22273 | 0.04905 | -0.08064 | -0.06206 | -0.28185 | -0.20559 | -0.17943 | 0.043063 | 0.1762 | -0.22782 | -0.18594 | -0.06499 | -0.01532 | -0.02329 | 0.16854 | 0.087753 | 0.25569 | -0.0492 | -0.0511 | 0.017341 | 0.03597 | 0.04511 | 0.029529 | -0.40669 |
| MFE280 | -0.20645 | -0.01879 | -0.1244 | -0.11265 | 0.05972 | -0.12094 | 0.201597 | -0.08118 | -0.0828 | 0.07229 | -0.23559 | 0.24235 | -0.00702 | -0.1046 | -0.12093 | 0.197075 | 0.08262 | 0.27042 | 0.014942 | 0.33807 | 0.29101 | -0.24426 | -0.16485 | 0.04523 | -0.01606 | -0.01239 | -0.2204 |
| SET2 | 0.07239 | 0.125665 | -0.27662 | -0.17491 | -0.10195 | -0.06557 | 0.105831 | -0.01928 | 0.32673 | 0.06659 | -0.13009 | 0.1844 | 0.16799 | -0.50224 | 0.053752 | -0.1738 | 0.01603 | 0.0874 | -0.14704 | 0.14883 | 0.4443 | -0.16623 | 0.269218 | -0.04307 | 0.31496 | 0.204762 | -0.55591 |
| EOL1 | -0.10766 | 0.058215 | -0.30394 | 0.00508 | -0.07154 | -0.00913 | 0.061319 | -0.0965 | -0.25684 | -0.02815 | -0.13547 | 0.10119 | -0.02785 | -0.11239 | -0.00243 | -0.07683 | -0.15503 | 0.13047 | 0.117642 | 0.16595 | 0.04533 | -0.09895 | 0.048735 | -0.08359 | 0.11671 | -0.0108 | -0.34245 |
| NMCG1 | -0.12729 | -0.00029 | -0.30251 | -0.08272 | 0.01224 | -0.2887 | 0.145375 | -0.11917 | 0.035899 | 0.06328 | -0.01029 | 0.14417 | -0.19604 | 0.25939 | -0.06836 | -0.20062 | 0.00368 | 0.08503 | -0.12191 | -0.01584 | 0.26258 | -0.1163 | 0.081318 | 0.09273 | 0.04856 | 0.040821 | -0.39005 |
| COLO320 | -0.12368 | 0.135124 | -0.5931 | 0.08763 | 0.0186 | -0.52309 | 0.062798 | -0.14453 | -0.0049 | -0.04695 | -0.06913 | 0.20812 | -0.06596 | 0.40199 | -0.02001 | -0.03984 | 0.21073 | -0.03864 | -0.40121 | 0.04649 | 0.23603 | -0.30764 | 0.029828 | 0.04682 | 0.12853 | -0.03321 | -0.29066 |
| LP1 | -0.23154 | 0.105833 | -0.94488 | -0.42462 | -0.15387 | -0.17277 | 0.192331 | -0.11552 | -0.08784 | -0.04964 | -0.18294 | 0.27922 | -0.20471 | 0.03999 | -0.13909 | 0.159226 | -0.08474 | 0.38506 | -0.19883 | 0.14083 | 0.16355 | -0.13079 | 0.066289 | -0.34649 | 0.15851 | -0.11936 | -0.60034 |
| C8166 | -0.1911 | -0.2615 | -0.19057 | 0.03666 | -0.09709 | -0.31226 | 0.167436 | -0.08726 | -0.29649 | 0.1526 | 0.006399 | 0.15876 | -0.14307 | 0.07707 | 0.028538 | -0.31316 | -0.10239 | 0.35197 | 0.149856 | 0.23168 | -0.04222 | -0.55291 | -0.01363 | 0.09283 | 0.25138 | 0.108327 | -0.80541 |
| DETROIT562 | -0.06236 | -0.03019 | 0.07549 | -0.12223 | -0.10438 | -0.07602 | 0.036842 | -0.06116 | -0.14529 | -0.11275 | -0.01799 | 0.0953 | -0.19297 | -0.17696 | -0.18805 | -0.10768 | -0.07876 | 0.12545 | -0.12764 | -0.0406 | 0.01742 | -0.12733 | 0.119798 | -0.0822 | 0.22556 | 0.213762 | -0.53707 |
| SNU1079 | -0.07218 | 0.058251 | -0.0281 | 0.09017 | -0.09966 | -0.06902 | -0.05634 | 0.02298 | -0.17534 | 0.25546 | -0.02405 | 0.10429 | 0.0092 | -0.0837 | -0.04188 | -0.00848 | -0.22622 | 0.13799 | -0.09207 | 0.18857 | 0.14338 | -0.09667 | 0.136273 | -0.17143 | 0.17733 | 0.047575 | -0.25848 |
| DAOY | -0.09098 | -0.00682 | -0.14269 | -0.13064 | -0.02764 | -0.13365 | -0.11228 | -0.15824 | -0.26974 | 0.10953 | -0.0225 | 0.06548 | -0.09504 | -0.02668 | -0.02823 | -0.13872 | -0.16299 | 0.11961 | -0.02521 | 0.05507 | 0.20106 | -0.18449 | 0.138508 | -0.04891 | 0.14437 | -0.01462 | -0.29036 |
| CAL120 | 0.02143 | 0.034987 | -0.08389 | -0.04427 | -0.12224 | -0.16302 | 0.017616 | -0.36929 | 0.028647 | -0.15292 | -0.08375 | 0.12322 | -0.05108 | -0.02912 | -0.12031 | -0.01686 | -0.03071 | -0.16306 | -0.13974 | 0.32248 | 0.0817 | -0.34508 | 0.166808 | 0.01429 | -0.04398 | -0.01957 | -0.18207 |
| HUPT4 | -0.13805 | 0.138937 | -0.39402 | -0.04283 | -0.31255 | -0.09687 | 0.026103 | -0.10576 | 0.055129 | -0.15275 | -0.08594 | 0.15261 | -0.17852 | 0.03048 | -0.21366 | -0.19994 | -0.03495 | -0.01067 | 0.02788 | 0.0186 | 0.14553 | 0.097335 | -0.06763 | -0.08196 | 0.13441 | 0.07499 | -0.49918 |
| LN382 | -0.06699 | -0.10438 | -0.18434 | -0.07849 | -0.11499 | -0.15194 | -0.24554 | -0.06254 | -0.17825 | -0.05039 | -0.26186 | 0.05748 | -0.28308 | -0.61858 | -0.00097 | -0.07125 | -0.09818 | 0.15913 | 0.036995 | 0.13843 | 0.0991 | -0.09352 | 0.128663 | -0.1016 | 0.12044 | -0.11007 | -0.3057 |
| JHESOAD1 | -0.14451 | 0.001775 | -0.28182 | -0.07206 | -0.06191 | -0.15518 | -0.04446 | -0.2827 | -0.06388 | -0.10904 | -0.1401 | 0.12183 | -0.08985 | -0.0126 | -0.14445 | -0.03485 | -0.04179 | -0.15032 | -0.13544 | 0.17728 | 0.13377 | -0.25571 | 0.027348 | -0.00125 | 0.18583 | 0.078648 | -0.29529 |
| JHH6 | -0.03169 | 0.021731 | -0.20053 | -0.22163 | -0.16599 | -0.28965 | -0.00132 | -0.14115 | -0.18454 | 0.07313 | -0.03897 | 0.09721 | 0.03212 | -0.11637 | -0.00932 | -0.03076 | -0.06452 | 0.0992 | -0.05195 | 0.00034 | 0.14473 | -0.2845 | 0.013477 | -0.12038 | 0.18895 | 0.082016 | -0.48965 |
| A375 | -0.10514 | -0.01337 | -0.58694 | -0.07301 | -0.15134 | -0.07593 | -0.04259 | -0.11404 | -0.02694 | 0.06316 | -0.04141 | 0.05887 | -0.18636 | -0.16356 | -0.09038 | -0.06704 | -0.11061 | 0.15054 | -0.00145 | 0.1284 | 0.10486 | -0.15775 | 0.077502 | -0.07461 | 0.12093 | -0.00539 | -0.55488 |
| SNU398 | -0.33563 | 0.045752 | -0.10695 | 0.03778 | -0.18634 | -0.18385 | 0.099577 | -0.03937 | -0.44737 | -0.35643 | 0.030339 | 0.07015 | -0.19319 | -0.18563 | -0.08779 | -0.22198 | -0.08003 | 0.15283 | -0.28031 | 0.06142 | 0.05302 | -0.00709 | 0.208247 | 0.0826 | 0.07925 | -0.03172 | -0.41033 |
| ASPC1 | -0.0976 | 0.000511 | -0.85729 | -0.31544 | -0.12058 | -0.22635 | 0.010149 | -0.02203 | -0.48971 | -0.12715 | -0.21712 | 0.12078 | -0.09261 | 0.22774 | -0.0926 | -0.07289 | -0.13359 | -0.04822 | -0.14711 | 0.12605 | 0.10192 | -0.1533 | 0.179543 | 0.12392 | 0.02102 | 0.057613 | -0.28431 |
| HCC1937 | -0.04339 | -0.12379 | 0.00138 | -0.09598 | -0.15635 | -0.10759 | -0.01304 | -0.15978 | -0.10854 | 0.05852 | -0.07533 | 0.06009 | -0.22747 | -0.00968 | -0.04682 | -0.08128 | -0.08682 | -0.14742 | -0.11734 | 0.17389 | -0.05581 | -0.08025 | 0.058762 | -0.06967 | 0.06811 | 0.091376 | -0.26938 |
| SUPM2 | -0.20477 | 0.185622 | -0.01407 | 0.01251 | -0.04159 | -0.20004 | -0.19244 | -0.18761 | -0.20963 | -0.13856 | 0.025686 | -0.0049 | -0.21959 | -0.07848 | -0.06913 | -0.24319 | -0.07325 | -0.08632 | 0.040689 | -0.03416 | -0.02408 | 0.003865 | 0.126828 | -0.07918 | 0.04681 | 0.135998 | -0.34375 |
| KPNYN | 0.03557 | 0.092367 | -0.93419 | -0.18252 | -0.18136 | -0.25947 | 0.040388 | -0.09407 | -0.04824 | -0.15715 | -0.06367 | 0.07624 | -0.30287 | -0.30408 | -0.12757 | -0.22221 | -0.09125 | 0.26741 | -0.21234 | 0.036 | 0.41098 | -0.26598 | 0.004728 | 0.0036 | 0.36508 | -0.06555 | -0.69545 |
| BICR31 | -0.02986 | 0.029623 | -0.19562 | 0.01473 | -0.04037 | -0.16633 | 0.039092 | -0.07764 | -0.04608 | -0.05328 | -0.09337 | 0.19016 | -0.26941 | -0.08362 | -0.02393 | -0.21497 | 0.05622 | 0.08648 | 0.138467 | 0.00617 | 0.07135 | -0.12877 | 0.125065 | -0.0925 | 0.14711 | 0.148408 | -0.35757 |
| KALS1 | -0.06099 | -0.02586 | -0.13147 | -0.06896 | -0.13024 | -0.03842 | 0.008616 | -0.05403 | 0.02873 | 0.01756 | -0.24984 | 0.10714 | -0.23849 | -0.1343 | -0.13373 | -0.01925 | -0.18498 | 0.1667 | -0.00585 | 0.08472 | 0.10995 | -0.14868 | 0.000428 | -0.14691 | 0.10005 | 0.020127 | -0.48544 |
| U251MG | -0.08873 | -0.01479 | -0.12466 | -0.1208 | -0.14369 | -0.04215 | 0.07026 | -0.19108 | -0.09748 | -0.05608 | -0.00595 | 0.11243 | -0.19572 | -0.22464 | -0.16314 | -0.17997 | -0.04741 | 0.16222 | 0.028997 | 0.15336 | 0.09967 | -0.0508 | 0.051738 | -0.07982 | -0.02217 | -0.11491 | -0.40305 |
| DEL | -0.16402 | -0.16533 | -0.16436 | -0.07011 | -0.14672 | -0.20466 | 0.102293 | -0.21902 | -0.34787 | -0.08727 | 0.033716 | 0.06333 | -0.09319 | -0.08779 | -0.0934 | -0.08163 | -0.06648 | 0.10479 | -0.02868 | 0.03898 | 0.02781 | -0.16618 | 0.101759 | 0.03053 | 0.39529 | 0.031506 | -0.53337 |
| CAKI2 | -0.09967 | -0.21614 | -0.3189 | -0.00224 | -0.19666 | -0.27528 | 0.257874 | -0.17015 | -0.2483 | -0.03765 | -0.0371 | 0.08576 | 0.03706 | -0.02374 | -0.02224 | -0.04641 | -0.27862 | 0.2898 | -0.08911 | 0.13473 | 0.12011 | -0.14681 | -0.06374 | 0.13904 | 0.31297 | 0.06487 | -0.26716 |
| PANC0403 | -0.18445 | -0.00194 | -0.43894 | -0.1044 | -0.10227 | -0.04838 | 0.003616 | -0.03319 | -0.02343 | -0.06536 | -0.06974 | 0.13926 | -0.1511 | -0.08245 | -0.15384 | -0.06541 | 0.00567 | -0.00892 | -0.01478 | 0.12985 | 0.12214 | -0.14636 | 0.124061 | 0.01425 | 0.07962 | 0.021059 | -0.38158 |
| JHOM1 | -0.00546 | -0.03229 | -0.35024 | -0.11068 | -0.15046 | -0.34583 | 0.020262 | -0.22554 | 0.153848 | 0.01825 | -0.04009 | 0.19927 | -0.21177 | -0.29588 | -0.09347 | -0.12995 | -0.04217 | 0.15181 | -0.07815 | 0.20488 | 0.20706 | -0.24463 | 0.128602 | -0.0294 | 0.25842 | -0.2101 | -0.44748 |
| SCC4 | 0.06211 | -0.10487 | -0.28377 | 0.04522 | -0.07631 | -0.1541 | -0.02073 | -0.31099 | -0.02298 | -0.04793 | -0.10173 | 0.13686 | -0.22952 | -0.24126 | -0.10299 | -0.04569 | -0.02439 | 0.25605 | -0.07318 | 0.1275 | -0.01977 | -0.22966 | 0.128746 | 0.04243 | 0.44924 | 0.115786 | -0.19084 |
| DANG | -0.06126 | 0.075941 | -0.27484 | 0.07984 | -0.09499 | -0.10949 | -0.00717 | -0.12669 | -0.12799 | -0.0938 | -0.06757 | 0.03802 | -0.23172 | 0.04068 | -0.08808 | -0.05893 | -0.06162 | 0.00529 | 0.013771 | 0.07943 | 0.15678 | -0.11703 | 0.249213 | -0.08227 | 0.09374 | 0.017412 | -0.38665 |
| DKMG | -0.06369 | -0.10922 | 0.03483 | -0.2427 | -0.12301 | -0.21182 | 0.014768 | -0.39271 | -0.07354 | 0.06012 | -1.30759 | -0.01174 | -1.96199 | -0.71367 | 0.02688 | 0.101103 | -0.14371 | 0.2556 | -0.176 | 0.08604 | 0.03997 | -0.35301 | 0.000863 | -0.05516 | 0.2674 | 0.165581 | -0.28567 |
| SLR23 | -0.07223 | 0.035893 | -0.19937 | -0.10619 | -0.04887 | -0.09923 | 0.068784 | -0.21873 | 0.057375 | -0.1046 | -0.13768 | 0.12444 | -0.14977 | -0.17593 | -0.08883 | 0.077659 | -0.02007 | -0.09436 | -0.02881 | 0.11762 | 0.14373 | -0.11615 | 0.293988 | -0.09551 | -0.0067 | -0.04963 | -0.25241 |
| OCUM1 | -0.08508 | -0.15421 | -0.62088 | -0.42361 | -0.08792 | -0.1201 | -0.03465 | -0.06395 | -0.48114 | -0.24056 | -0.01459 | -0.12238 | -0.19171 | -0.14484 | -0.04042 | -0.03562 | -0.04507 | 0.00372 | -0.08188 | 0.07803 | 0.18764 | -0.0258 | 0.307608 | -0.05037 | -0.00069 | -0.0081 | -0.3932 |
| AU565 | -0.08821 | -0.02136 | -0.19329 | -0.03425 | -0.06873 | -0.18786 | -0.16354 | -0.50747 | -0.05401 | -0.13249 | -0.00408 | 0.14947 | -0.19886 | -0.27299 | -0.1357 | -0.07016 | -0.08005 | -0.10145 | -0.20408 | 0.19948 | 0.30023 | -0.32027 | 0.054909 | 0.01952 | 0.10884 | -0.1675 | -0.40324 |
| CL11 | -0.18332 | 0.189915 | 0.14827 | -0.19546 | 0.11236 | -0.10955 | -0.04776 | 0.06723 | -0.06798 | 0.09759 | 0.089716 | 0.12652 | -0.04759 | 0.02625 | 0.231409 | -0.17966 | 0.16704 | 0.19731 | -0.21425 | 0.25268 | 0.30376 | -0.05994 | 0.160006 | 0.10177 | 0.10848 | -0.00724 | -0.45027 |
| KMRC20 | -0.12852 | -0.02796 | -0.13472 | -0.09561 | -0.17295 | -0.08596 | 0.027572 | -0.28593 | -0.17472 | 0.02088 | -0.15146 | 0.08474 | -0.06833 | -0.12439 | -0.17763 | -0.18092 | -0.10657 | 0.06657 | 0.031072 | 0.13007 | 0.06429 | -0.14092 | 0.07865 | -0.07817 | 0.13156 | -0.06068 | -0.29115 |
| NCIH2887 | -0.10898 | -0.50139 | -0.0912 | 0.01108 | -0.0232 | -0.3534 | 0.070067 | -0.08422 | -0.10526 | -0.08787 | -0.00866 | 0.14469 | 0.09143 | -0.49328 | -0.04861 | -0.28704 | -0.14079 | 0.08691 | -0.26672 | 0.32065 | 0.15029 | -0.08366 | 0.115622 | 0.12515 | 0.09137 | -0.04978 | -0.65563 |
| LS1034 | -0.04269 | -0.05977 | -0.73599 | -0.0896 | -0.14559 | -0.2482 | -0.07011 | -0.38922 | -0.17195 | -0.04471 | -0.08225 | 0.09799 | 0.06408 | -0.21194 | 0.093205 | -0.14029 | -0.00921 | 0.01372 | 0.026424 | 0.21578 | 0.15234 | -0.27997 | 0.135226 | -0.22829 | 0.38873 | 0.030391 | -0.21654 |
| COLO201 | -0.05676 | -0.02319 | -0.49631 | -0.04565 | -0.18766 | -0.12033 | -0.01942 | -0.23726 | -0.14093 | -0.01862 | 0.053079 | 0.09633 | -0.10448 | 0.08398 | 0.073298 | -0.01911 | -0.02247 | 0.04135 | 0.01203 | 0.10584 | 0.22369 | -0.22126 | 0.073323 | -0.02982 | 0.19335 | 0.070192 | -0.32527 |
| LMSU | -0.15001 | 0.063561 | -0.188 | -0.1383 | -0.17987 | -0.15485 | -0.06187 | -0.38947 | -0.21594 | -0.03733 | -0.14463 | 0.03748 | -0.58104 | -0.25394 | -0.09417 | -0.45867 | -0.12049 | 0.05027 | -0.04804 | 0.1961 | 0.20143 | -0.28225 | 0.191983 | -0.07765 | 0.10356 | -0.1451 | -0.32541 |
| COV318 | -0.12909 | 0.045557 | -0.40475 | 0.16629 | -0.12412 | -0.30839 | -0.23043 | -0.34461 | -0.03727 | 0.04939 | 0.007667 | 0.14585 | -0.05465 | -0.20799 | -0.17192 | 0.009567 | 0.20113 | 0.1134 | 0.020813 | 0.28169 | 0.12254 | 0.016407 | 0.122843 | -0.10811 | 0.16806 | -0.05531 | -0.38025 |
| CORL279 | -0.00829 | 0.055562 | -0.13171 | 0.13762 | 0.07049 | -0.02472 | -0.23744 | -0.06914 | 0.390265 | 0.15421 | 0.058921 | 0.13753 | -0.17278 | -0.21993 | -0.03797 | -0.02142 | -0.27133 | 0.02501 | 0.098895 | 0.23761 | 0.14341 | -0.2246 | -0.00481 | 0.14362 | -0.01932 | 0.219933 | -0.54193 |
| DU4475 | -0.34839 | -0.17989 | -0.30186 | -0.10919 | 0.07729 | -0.08008 | 0.148169 | -0.28905 | -0.15914 | 0.0934 | 0.02013 | -0.16293 | -0.55661 | -0.049 | -0.04965 | -0.47579 | -0.07247 | 0.08412 | -0.08472 | 0.11144 | -0.14412 | -0.33191 | 0.03989 | 0.22927 | 0.3056 | -0.11698 | -0.44951 |
| KELLY | -0.13771 | 0.034172 | -0.78119 | -0.07603 | -0.10748 | -0.13737 | -0.03645 | -0.21825 | -0.08336 | -0.00946 | -0.05077 | 0.13267 | -0.25339 | -0.25671 | -0.0336 | 0.008696 | -0.09196 | 0.11308 | -0.02627 | 0.13239 | 0.17828 | 0.006619 | 0.041522 | -0.03425 | 0.19965 | 0.029046 | -0.23708 |
| SKNAS | -0.06679 | 0.026862 | -0.79937 | -0.08615 | -0.11546 | -0.10727 | -0.13447 | -0.2808 | -0.17814 | -0.14429 | -0.10795 | 0.08819 | -0.15196 | -0.04089 | -0.05479 | -0.05473 | -0.00469 | 0.02533 | -0.03286 | 0.12424 | -0.00707 | -0.18192 | 0.021339 | -0.0097 | 0.2148 | -0.04041 | -0.29915 |

| Cell line | FLT3 | FLT4 | IGF1R | INSR | INSRR | KDR | KIT | LMR1 | LMR2 | LTK | MET | MUSK | PDGFRA | PDGFRB | PTK7 | RET | RON | ROR1 | ROR2 | ROS1 | STYK1 | TIE1 | TIE2 | TRKA | TRKB | TRKC | TYRO3 |
|---|---|---|---|---|---|---|---|---|---|---|---|---|---|---|---|---|---|---|---|---|---|---|---|---|---|---|---|
| RERFLCAI | -0.11698 | 0.057142 | -0.47254 | -0.07919 | -0.09032 | -0.1467 | -0.03749 | -0.11219 | -0.0491 | -0.08253 | -0.1268 | 0.01935 | -0.09077 | -0.03513 | -0.12212 | -0.15386 | -0.11116 | 0.24817 | 0.049859 | 0.13492 | 0.10629 | -0.04529 | -0.06969 | -0.03874 | 0.04942 | 0.06162 | -0.67838 |
| UOK101 | -0.18587 | 0.120245 | -0.20304 | 0.17136 | -0.17965 | 0.03195 | -0.03091 | 0.00816 | -0.34051 | -0.03852 | -0.09836 | 0.12738 | -0.34805 | -0.44862 | 0.031445 | -0.03538 | -0.15981 | -0.24123 | -0.10392 | 0.06039 | 0.0674 | -0.32458 | 0.22451 | -0.01968 | 0.22969 | -0.07303 | -0.28158 |
| KASUMI1 | 0.15778 | -0.16325 | -0.2395 | -0.15492 | -0.06312 | -0.85344 | -0.67347 | 0.01089 | -0.19516 | -0.02634 | 0.069568 | 0.0916 | -0.58349 | 0.00427 | -0.04898 | -0.36203 | -0.15781 | 0.27598 | -0.4612 | 0.13222 | 0.1372 | -0.1443 | -0.07453 | -0.02457 | 0.20327 | 0.187763 | -0.20941 |
| CALU6 | -0.11741 | 0.003494 | -0.35458 | -0.09343 | -0.17035 | -0.19739 | -0.01714 | -0.31751 | -0.07265 | -0.03233 | -0.16223 | 0.07892 | -0.29293 | -0.19657 | -0.02999 | -0.00351 | -0.21738 | 0.01959 | -0.02355 | 0.0741 | 0.01563 | -0.10803 | -0.02666 | -0.16818 | 0.06276 | -0.05387 | -0.31225 |
| KP4 | -0.20267 | 0.052926 | -0.06047 | -0.10316 | -0.13616 | -0.25042 | -0.05448 | -0.03278 | 0.022611 | -0.04346 | -0.63559 | 0.01236 | -0.24075 | -0.01709 | -0.04716 | -0.16888 | -0.02831 | -0.01608 | 0.02513 | -0.01582 | 0.15707 | -0.14035 | 0.052087 | -0.05135 | 0.12898 | 0.017793 | -0.3489 |
| SNU213 | -0.05646 | 0.046232 | -0.40535 | -0.20233 | -0.07364 | -0.14573 | 0.031729 | -0.19681 | -0.27034 | -0.083 | -0.11956 | 0.08231 | -0.0407 | 0.12295 | -0.04755 | 0.004399 | -0.01549 | -0.06783 | -0.02411 | 0.11536 | 0.13721 | -0.15684 | 0.015761 | -0.03506 | 0.17352 | 0.043826 | -0.26849 |
| AM38 | -0.1991 | -0.07582 | -0.74957 | -0.27707 | -0.09556 | -0.31897 | 0.028163 | -0.17533 | -0.32457 | -0.0154 | -0.14749 | 0.22555 | -0.38448 | -0.217 | 0.136043 | -0.2085 | -0.05698 | 0.03775 | 0.181088 | 0.05078 | 0.0415 | -6.1E-05 | -0.01924 | -0.02229 | 0.17236 | 0.095098 | -0.33785 |
| SUDHL10 | -0.00209 | -0.07683 | 0.05527 | -0.11652 | -0.07617 | -0.15194 | 0.086862 | -0.20372 | -0.19372 | -0.01241 | 0.052014 | 0.1116 | -0.25436 | -0.11953 | -0.12974 | -0.11527 | -0.05452 | 0.16484 | 0.015315 | 0.23876 | 0.07575 | -0.14862 | 0.007919 | 0.04281 | 0.35351 | 0.02123 | -0.54207 |
| SLR24 | -0.01091 | -0.0086 | -0.09937 | -0.15067 | -0.06227 | -0.27411 | -0.0004 | -0.18949 | -0.15084 | -0.03752 | -0.15575 | -0.0689 | -0.22741 | -0.21282 | -0.1541 | -0.12403 | -0.09087 | 0.13955 | -0.01273 | 0.10433 | 0.15992 | -0.2913 | -0.0922 | -0.18861 | 0.14493 | -0.122 | -0.59302 |
| SF539 | -0.08142 | 0.055814 | -0.02003 | -0.00986 | -0.10369 | -0.14936 | 0.069866 | -0.17657 | 0.074227 | 0.04764 | -0.08566 | -0.0977 | -0.63645 | -0.17596 | -0.04019 | -0.0135 | -0.04703 | 0.02922 | -0.11349 | 0.086 | 0.03822 | -0.15653 | 0.041997 | -0.19058 | 0.09583 | 0.038051 | -0.20832 |
| HS852T | -0.14946 | -0.04118 | -0.48738 | -0.14491 | -0.0726 | -0.09637 | -0.07594 | -0.11416 | -0.06394 | -0.01056 | -0.07927 | 0.06782 | -0.24516 | -0.14518 | -0.12312 | -0.15913 | -0.17751 | 0.17849 | 0.045328 | 0.05668 | 0.05439 | -0.24178 | 0.056862 | -0.04648 | 0.11472 | 0.070135 | -0.40414 |
| HCC38 | -0.27139 | 0.151476 | 0.17226 | -0.04514 | -0.26943 | -0.2629 | -0.11049 | 0.00046 | -0.01706 | -0.07619 | -0.00348 | 0.1607 | -0.25661 | -0.10923 | -0.07861 | -0.22494 | -0.03199 | 0.31824 | -0.10432 | 0.26911 | 0.08958 | -0.02252 | 0.301396 | -0.13927 | 0.08821 | 0.199953 | -0.18803 |
| HCC1419 | -0.0851 | -0.10435 | -0.16899 | 0.01173 | -0.05156 | -0.12716 | -0.03932 | -0.22081 | -0.19945 | 0.00838 | -0.03874 | 0.21186 | -0.20638 | 0.04289 | 0.063309 | 0.032513 | -0.04078 | 0.00871 | -0.01346 | -0.03678 | 0.07209 | -0.19123 | 0.089479 | 0.0436 | 0.2009 | -0.08502 | -0.21582 |
| COV362 | -0.01701 | 0.029202 | -0.24089 | -0.13933 | -0.06708 | -0.228 | 0.019079 | -0.32074 | -0.09931 | 0.0096 | -0.08168 | 0.03734 | -0.27564 | -0.42029 | -0.11041 | -0.09211 | -0.06349 | 0.3318 | -0.04748 | 0.07533 | 0.22125 | -0.44098 | 0.171423 | -0.13058 | 0.37331 | 0.024335 | -0.37381 |
| EWS502 | -0.15253 | 0.11223 | -0.65677 | 0.03861 | 0.03073 | -0.1041 | -0.20585 | -0.17919 | -0.04445 | -0.02495 | -0.04948 | 0.12672 | -0.2248 | -0.20971 | 0.027686 | -0.15874 | -0.13859 | 0.22048 | 0.124783 | 0.09131 | 0.00321 | -0.15192 | 0.129932 | 0.01981 | 0.10698 | 0.008595 | -0.3975 |
| SNU840 | -0.3166 | 0.072793 | -0.16385 | 0.07218 | -0.13545 | -0.2671 | -0.02532 | -0.13616 | -0.09237 | -0.10932 | 0.037536 | 0.13516 | -0.17427 | -0.22422 | -0.08279 | -0.1303 | 0.07179 | 0.28723 | -0.18861 | 0.12336 | 0.03934 | -0.11881 | 0.044661 | -0.14873 | 0.26207 | 0.066993 | -0.72966 |
| KP2 | -0.00084 | -0.20514 | -0.54344 | -0.11122 | -0.2206 | -0.17588 | 0.087642 | -0.22287 | -0.31254 | 0.07248 | 0.108813 | 0.02868 | -0.30813 | -0.24524 | -0.03973 | 0.08094 | -0.05582 | 0.18667 | -0.04609 | 0.20411 | -0.26344 | -0.12417 | 0.022595 | -0.02838 | 0.0783 | -0.24295 | -0.39373 |
| NCIH1755 | -0.1423 | 0.149293 | -0.49125 | 0.01352 | -0.09542 | -0.12575 | 0.101848 | -0.18079 | 0.027047 | -0.121 | -0.17301 | 0.21364 | -0.10566 | -0.14762 | -0.22645 | 0.075392 | -0.15903 | 0.19623 | -0.01191 | 0.13019 | 0.20967 | -0.07338 | -0.05896 | 0.047 | 0.1228 | 0.068352 | -0.18776 |
| SNU1033 | -0.18901 | 0.200563 | -0.60653 | -0.02171 | -0.13046 | -0.33412 | 0.117828 | -0.23955 | -0.31943 | -0.25678 | -0.0281 | 0.07752 | -0.44322 | 0.01143 | -0.09024 | -0.07257 | -0.1801 | 0.42031 | -0.1636 | 0.10053 | 0.17851 | -0.36639 | 0.188206 | -0.01723 | 0.0414 | 0.111383 | -0.41504 |
| BT549 | -0.15647 | -0.07855 | -0.06849 | -0.11373 | -0.16408 | -0.15025 | -0.00563 | -0.33104 | -0.16391 | -0.05462 | -0.0634 | 0.06861 | -0.07744 | -0.28927 | -0.07861 | -0.08376 | -0.02873 | 0.23405 | -0.09943 | 0.1128 | 0.06248 | -0.13631 | 0.104541 | -0.09993 | 0.10506 | -0.13378 | -0.51473 |
| NCIH209 | -0.09691 | -0.00522 | -0.12421 | -0.08407 | -0.12336 | -0.09379 | 0.072294 | -0.15442 | -0.04482 | -0.07692 | -0.03786 | 0.11007 | -0.12412 | -0.14294 | 0.005916 | -0.06067 | -0.08696 | -0.0454 | -0.05413 | 0.07055 | 0.04331 | -0.14585 | 0.061295 | -0.08314 | 0.19868 | -0.01834 | -0.34071 |
| OV90 | -0.17681 | -0.02815 | -1.05643 | -0.20756 | -0.13595 | -0.09102 | 0.078105 | -0.20381 | -0.26051 | -0.05939 | 0.039396 | 0.15178 | -0.19772 | -0.08704 | -0.04479 | -0.02141 | -0.14006 | 0.15984 | 0.033764 | 0.06265 | 0.14777 | -0.06571 | 0.065098 | -0.13178 | 0.07256 | 0.024743 | -0.73698 |
| NCIH841 | -0.09325 | -0.1866 | 0.002 | 0.08544 | -0.22215 | -0.12457 | 0.067902 | -0.1305 | -0.14263 | -0.24192 | -0.15122 | 0.02135 | -0.08296 | -0.0164 | -0.17613 | -0.42938 | -0.27028 | 0.4399 | 0.076202 | 0.30867 | -0.1082 | -0.03848 | -0.05642 | -0.1693 | 0.19433 | 0.084776 | -0.66116 |
| KLE | -0.02985 | 0.271442 | -0.14485 | -0.10333 | -0.0281 | -0.16714 | 0.174226 | 0.06063 | -0.15423 | 0.03551 | -0.1995 | 0.02858 | 0.04839 | 0.18367 | -0.07221 | 0.123474 | -0.0781 | 0.17507 | 0.053066 | 0.25695 | 0.31897 | 0.082357 | -0.10946 | 0.04655 | 0.14549 | 0.140797 | -0.09224 |
| NB4 | -0.18577 | 0.055386 | -0.76939 | -0.10556 | -0.06909 | -0.31926 | 0.022328 | -0.01182 | -0.16141 | 0.04343 | -0.12113 | 0.12833 | 0.0068 | -0.02916 | -0.03882 | 0.18105 | 0.02782 | -0.02238 | -0.08219 | 0.06093 | 0.17897 | -0.2173 | -0.10011 | 0.13731 | 0.11633 | 0.188376 | -0.60385 |
| EM2 | -0.05618 | 0.145614 | -0.44528 | -0.27432 | -0.04746 | -0.44677 | 0.108487 | -0.29928 | 0.154646 | 0.08746 | -0.19663 | 0.11341 | -0.10491 | -0.07822 | -0.12058 | -0.19718 | -0.26036 | 0.43339 | -0.22859 | 0.0635 | 0.10684 | -0.53511 | 0.078359 | 0.03723 | 0.25682 | 0.038975 | -0.15165 |
| OUMS23 | -0.20946 | -0.24251 | -0.23952 | -0.05241 | -0.10655 | -0.15391 | 0.235276 | -0.10816 | -0.40822 | 0.06327 | -0.16908 | 0.30708 | -0.06928 | -0.10395 | 0.018236 | 0.227076 | -0.04456 | 0.33294 | 0.192304 | -0.10575 | -0.05582 | -0.07976 | 0.056501 | -0.04048 | 0.33699 | 0.103416 | -0.28554 |
| SNU1077 | 0.05007 | 0.049297 | -0.65628 | -0.09627 | -0.06091 | -0.24392 | -0.02056 | -0.08133 | -0.14987 | -0.02333 | -0.20173 | 0.08734 | -0.12401 | -0.08246 | 0.016251 | -0.15921 | -0.01401 | 0.07924 | -0.07291 | 0.13465 | 0.21366 | -0.09787 | 0.041531 | -0.13262 | 0.10088 | -0.07657 | -0.26121 |
| SNU5 | -0.17103 | 0.14706 | -0.12775 | 0.12633 | -0.177 | -0.07256 | -0.05439 | -0.2831 | 0.018647 | 0.00321 | 0.143936 | 0.11269 | -0.22963 | 0.0994 | -0.03777 | -0.03956 | -0.03919 | 0.15403 | -0.1494 | 0.33854 | 0.03738 | -0.13763 | 0.092016 | -0.01692 | -0.00093 | 0.156027 | -0.20669 |
| WM115 | -0.29614 | 0.051877 | -0.22148 | 0.11776 | 0.01437 | -0.04481 | -0.17912 | -0.17255 | -0.563 | -0.10199 | 0.043987 | 0.15247 | -0.29088 | 0.0303 | -0.10923 | 0.074307 | 0.0796 | 0.10364 | -0.1122 | 0.28162 | 0.09384 | -0.19012 | -0.06146 | 0.1817 | 0.22745 | 0.274506 | -0.61227 |
| ECGI10 | 0.01267 | -0.0879 | -0.18028 | -0.14466 | -0.0706 | -0.02274 | 0.035029 | -0.13965 | -0.33286 | -0.05132 | -0.12157 | 0.22547 | -0.14303 | -0.17478 | -0.12831 | 0.040932 | -0.0384 | 0.10122 | -0.03721 | 0.08156 | 0.04048 | -0.2271 | -0.04376 | 0.06587 | 0.22256 | 0.003328 | -0.4284 |
| PK1 | -0.19851 | -0.10076 | -0.49584 | -0.06885 | -0.15902 | -0.25031 | 0.058222 | -0.10311 | -0.08535 | -0.09884 | -0.15081 | 0.21278 | -0.32706 | -0.05323 | -0.10175 | -0.00791 | 0.02289 | 0.28335 | 0.144251 | 0.14955 | -0.07639 | -0.10729 | 0.181387 | -0.04949 | 0.26138 | -0.0859 | -0.29101 |
| EFO21 | -0.00637 | -0.04828 | -0.40439 | -0.18141 | -0.17669 | -0.15351 | -7E-05 | -0.17644 | -0.04419 | -0.04969 | 0.01371 | 0.1543 | -0.12199 | -0.13841 | -0.12828 | 0.002582 | -0.10548 | 0.06642 | -0.10181 | 0.12522 | 0.17785 | -0.11234 | 0.122857 | -0.16028 | 0.16363 | -0.06748 | -0.40948 |
| IMR32 | -0.05663 | -0.01509 | -0.25327 | -0.06144 | -0.13401 | -0.24571 | -0.1339 | -0.36539 | -0.24334 | 0.09404 | 0.062835 | -0.028 | -0.18666 | 0.11631 | 0.143177 | -0.03084 | -0.0674 | -0.0292 | -0.07291 | 0.26867 | 0.1371 | 0.012297 | 0.059159 | 0.16863 | 0.23351 | 0.053325 | -0.59244 |
| NCIH2122 | -0.21687 | 0.035465 | -0.93268 | -0.11797 | -0.15701 | -0.16051 | -0.10301 | 0.11055 | -0.53347 | -0.09921 | 0.046954 | 0.05829 | -0.20242 | 0.30471 | -0.01084 | 0.043298 | -0.10703 | -0.04955 | -0.01141 | 0.05123 | 0.13369 | -0.06659 | 0.050971 | -0.01348 | 0.15844 | 0.102909 | -0.38011 |
| SKNBE2 | -0.02012 | 0.102809 | -0.54532 | -0.11927 | -0.17435 | -0.27766 | -0.14632 | -0.10763 | -0.1077 | -0.11505 | -0.01281 | 0.0399 | -0.14572 | 0.00655 | -0.08054 | -0.01356 | -0.05932 | 0.15987 | -0.1721 | 0.19726 | 0.21878 | -0.1679 | 0.07017 | 0.09841 | 0.21346 | 0.003808 | -0.51333 |
| KMRC3 | -0.0306 | -0.01738 | -0.50711 | -0.17159 | -0.1795 | -0.0199 | -0.00826 | -0.13599 | 0.226991 | -0.14426 | -0.17629 | 0.05201 | -0.16756 | -0.11371 | -0.11693 | -0.06443 | -0.16342 | -0.09093 | 0.014668 | 0.10342 | 0.21487 | -0.10785 | 0.056358 | -0.06966 | 0.12047 | -0.10955 | -0.2999 |
| KARPAS422 | 0.00336 | -0.07089 | -0.3133 | 0.0222 | -0.10575 | 0.00525 | -0.0225 | -0.26449 | 0.098693 | 0.10745 | 0.134428 | 0.09002 | -0.26251 | -0.43252 | 0.070811 | -0.32286 | -0.03553 | 0.28447 | 0.172498 | 0.07934 | 0.01777 | -0.42542 | -0.00776 | 0.11753 | 0.0948 | 0.201085 | -0.92413 |
| SNU886 | -0.05825 | -0.01369 | -0.21134 | -0.08267 | -0.15778 | -0.20801 | -0.12467 | 0.05829 | -0.35859 | -0.13365 | -0.21108 | 0.18403 | -0.16741 | -0.08267 | -0.13798 | -0.0764 | -0.15727 | 0.09106 | 0.036792 | 0.1194 | 0.24976 | -0.21122 | -0.20139 | 0.02885 | 0.23545 | 0.066071 | -0.30343 |
| TUHR14TKB | -0.09055 | -0.09407 | -0.41188 | -0.06223 | -0.10666 | -0.02588 | 0.129113 | -0.16623 | -0.20059 | 0.18062 | -0.13403 | 0.14868 | -0.46691 | -0.17504 | 0.132303 | 0.07887 | 0.01247 | -0.05381 | 0.091662 | 0.17049 | 0.00996 | -0.53571 | -0.01137 | -0.0949 | 0.05742 | 0.045444 | -0.45142 |
| TE10 | 0.05582 | -0.10199 | -0.72275 | 0.02021 | -0.13147 | -0.287 | -0.01017 | -0.0543 | 0.012027 | 0.03638 | -0.03675 | 0.10052 | 0.01063 | -0.00579 | 0.034818 | -0.09393 | -0.00477 | 0.23023 | 0.051587 | 0.11339 | 0.20808 | -0.27586 | 0.061384 | 0.03037 | 0.19046 | 0.051482 | -0.31583 |
| MPP89 | 0.07047 | 0.160997 | -0.26493 | -0.00546 | -0.2081 | -0.30681 | 0.027786 | -0.19372 | 0.045521 | 0.09835 | -0.09269 | -0.03048 | -0.59606 | -0.32586 | -0.10043 | -0.03169 | -0.04071 | -0.09947 | 0.003118 | 0.05753 | 0.22324 | -0.07065 | 0.12225 | -0.13214 | 0.11432 | -0.02933 | -0.37818 |
| PSN1 | -0.1287 | 0.092956 | -0.33092 | 0.03916 | -0.03276 | -0.10871 | -0.05658 | -0.16131 | -0.04502 | -0.03927 | 0.029646 | -0.02493 | -0.2111 | 0.12072 | 0.052892 | -0.09153 | 0.04759 | 0.14653 | -0.0504 | 0.09472 | 0.1045 | -0.14816 | 0.005156 | 0.01354 | 0.03862 | -0.03693 | -0.16093 |
| HT144 | -0.21991 | -0.13161 | -0.20604 | -0.01834 | -0.02644 | -0.11313 | -0.02564 | -0.41711 | -0.05276 | 0.24522 | -0.02136 | 0.13913 | -0.21136 | -0.35018 | -0.05281 | -0.03118 | 0.02123 | 0.11153 | -0.09899 | 0.26173 | 0.19091 | 0.130964 | 0.233434 | -0.0352 | 0.07328 | 0.017301 | -0.60114 |
| 42MGBA | -0.08568 | -0.05163 | -0.09852 | -0.14239 | -0.08678 | -0.32431 | -0.16644 | -0.28061 | -0.06153 | 0.09413 | -0.1177 | 0.09738 | -1.62675 | -0.17481 | -0.05006 | -0.11865 | -0.24711 | 0.03345 | 0.144485 | 0.07516 | 0.02154 | -0.0111 | 0.111319 | -0.02972 | 0.25879 | -0.13713 | -0.37555 |
| JHOC5 | -0.04978 | 0.11448 | -0.12287 | -0.07572 | -0.03606 | -0.0663 | -0.03315 | -0.19392 | -0.14776 | 0.02033 | -0.01624 | 0.05101 | -0.07418 | -0.03815 | -0.05049 | -0.05379 | 0.0197 | 0.03324 | -0.0296 | 0.14144 | 0.15923 | -0.16434 | 0.062857 | 0.0498 | 0.22103 | -0.02165 | -0.41518 |
| SNU620 | -0.32932 | -0.03914 | -0.2654 | -0.13097 | -0.30716 | -0.35034 | 0.030617 | -0.14212 | 0.017261 | -0.20883 | -0.00732 | 0.07383 | -0.20247 | 0.00084 | -0.07995 | 0.026634 | -0.00459 | 0.33306 | -0.26138 | 0.35001 | 0.22816 | -0.18988 | 0.164391 | 0.14384 | 0.18511 | 0.086601 | -0.33949 |
| JURLMK1 | -0.2056 | -0.01292 | -0.19842 | -0.0433 | -0.15843 | -0.16826 | 0.067002 | -0.07255 | 0.025093 | -0.02935 | -0.11279 | 0.30645 | -0.31397 | 0.11936 | -0.19628 | -0.01715 | -0.15129 | 0.19736 | 0.106054 | 0.10271 | 0.01372 | -0.42393 | 0.103754 | -0.00702 | 0.25933 | 0.052326 | -0.28309 |
| CCFSTTG1 | -0.15095 | 0.03532 | -0.249 | 0.04819 | -0.14547 | -0.26757 | 0.006598 | -0.26468 | -0.151 | 0.10512 | -0.012 | 0.1278 | -0.43953 | -0.52854 | -0.00227 | -0.02099 | -0.07439 | 0.22537 | -0.0813 | 0.18884 | 0.02085 | -0.10033 | 0.222617 | -0.0835 | 0.16645 | -0.09941 | -0.23014 |
| EFM19 | 0.02238 | 0.019837 | -0.2444 | -0.23923 | -0.16768 | -0.50534 | -0.0803 | -0.10456 | 0.020702 | -0.15326 | -0.10694 | -0.00889 | -0.4509 | -0.37158 | -0.05073 | -0.20917 | -0.09632 | 0.07383 | 0.055515 | 0.10913 | 0.1069 | -0.04931 | 0.321403 | 0.09705 | -0.02813 | -0.05356 | -0.51501 |
| ISTMES2 | -0.09516 | 0.144902 | -0.06828 | -0.12721 | -0.12693 | -0.17934 | -0.01295 | -0.07909 | 0.007165 | -0.03001 | -0.0619 | 0.03101 | -0.06207 | -0.10615 | -0.05412 | -0.14601 | 0.0098 | 0.13746 | 0.004718 | 0.12038 | 0.08017 | -0.13028 | 0.221782 | -0.09826 | 0.0265 | -0.02348 | -0.27745 |
| YAPC | -0.1647 | -0.06044 | -0.70675 | -0.02698 | -0.13242 | -0.11893 | -0.09082 | -0.15616 | -0.03687 | -0.14149 | -0.07099 | 0.08355 | -0.11413 | -0.05818 | -0.32705 | -0.00014 | 0.00311 | 0.03517 | -0.05784 | 0.1051 | 0.11594 | -0.05358 | 0.247345 | -0.09777 | 0.14905 | 0.053697 | -0.40758 |
| DB | -0.04352 | 0.031103 | -0.12291 | 0.03423 | -0.03932 | -0.50664 | 0.003297 | -0.12821 | -0.16206 | -0.16768 | 0.00447 | 0.02262 | 0.04063 | 0.03427 | -0.12077 | -0.26833 | 0.0225 | 0.424 | 0.085581 | 0.12525 | -0.09984 | -0.22547 | 0.000979 | 0.1168 | 0.18297 | 0.175406 | -0.28396 |
| MSTO211H | 0.01057 | 0.089808 | -0.17614 | -0.20001 | -0.28669 | -0.13628 | 0.200758 | -0.16505 | 0.035352 | -0.01913 | -0.07418 | -0.16573 | -0.03582 | -0.27416 | -0.26749 | -0.38629 | -0.14228 | 0.25464 | -0.05968 | 0.20669 | -0.03735 | -0.03883 | 0.039669 | -0.11586 | 0.23119 | -0.13481 | -0.34374 |
| OCIAML3 | -0.11105 | -0.04155 | -0.19249 | -0.26611 | -0.18048 | -0.1593 | -0.0991 | -0.24074 | -0.0519 | -0.15942 | -0.09716 | 0.36601 | -0.18175 | -0.16406 | -0.06855 | -0.2609 | -0.15601 | 0.09616 | -0.07521 | 0.17172 | 0.02224 | -0.22304 | 0.045678 | -0.05868 | 0.24684 | 0.088971 | -0.24521 |
| NCIH3122 | 0.00401 | -0.07649 | -0.24013 | 0.03013 | 0.02707 | -0.19875 | 0.227897 | -0.10181 | -0.1656 | 0.00837 | -0.08588 | 0.11993 | -0.15879 | 0.13707 | -0.02981 | -0.12547 | -0.03563 | 0.24673 | -0.06808 | 0.05111 | 0.07894 | -0.21879 | -0.06622 | -0.05465 | 0.14202 | 0.080486 | -0.33456 |
| HCC461 | -0.09105 | 0.015419 | -0.07799 | 0.01489 | -0.15221 | -0.19214 | -0.19153 | -0.11789 | -0.01806 | 0.02864 | -0.09096 | 0.01034 | -0.39274 | -0.07971 | 0.012159 | -0.06918 | -0.09793 | -0.10662 | -0.01869 | 0.16961 | 0.05856 | -0.10459 | 0.073683 | -0.04923 | 0.14148 | -0.00299 | -0.35894 |
| SKNFI | -0.05145 | 0.063526 | -1.65471 | -0.16249 | -0.22263 | -0.12003 | 0.079095 | -0.27511 | -0.07755 | -0.05594 | -0.02679 | -0.04577 | 0.04085 | -0.24707 | -0.06507 | -0.31645 | -0.10401 | 0.07894 | -0.01358 | 0.11479 | 0.10487 | -0.14706 | -0.00607 | -0.09227 | 0.16301 | 0.08 | -0.18535 |
| NCIH522 | -0.3517 | 0.039909 | -0.11537 | 0.01569 | -0.15501 | -0.09499 | -0.12446 | -0.31732 | 0.04238 | 0.0686 | -0.13941 | 0.09532 | -0.29467 | -0.18851 | -0.02809 | -0.29247 | -0.16891 | 0.13084 | -0.00347 | 0.12981 | 0.10747 | -0.13573 | 0.115879 | -0.15205 | 0.18416 | 0.203349 | -0.54123 |
| SNU668 | -0.1212 | -0.02225 | -0.60402 | 0.11168 | -0.03642 | -0.37795 | -0.08491 | -0.03425 | -0.44574 | -0.13405 | -0.06152 | 0.17874 | -0.3183 | 0.08704 | -0.05449 | -0.13346 | -0.00922 | 0.30675 | -0.19674 | 0.05496 | 0.21574 | -0.17359 | 0.103745 | 0.12922 | 0.2001 | -0.00978 | -0.37766 |
| JVM3 | -0.12853 | -0.11627 | -0.08136 | -0.19925 | -0.20473 | -0.04148 | 0.028127 | -0.10439 | -0.15831 | 0.22775 | -0.28489 | 0.13022 | -0.24917 | -0.13505 | 0.052785 | 0.08984 | -0.05214 | -0.0402 | -0.04654 | 0.29744 | -0.02609 | -0.00804 | 0.120643 | -0.13092 | 0.50697 | 0.103027 | -0.33732 |
| RPMI7951 | -0.05415 | 0.064686 | -0.27564 | -0.03592 | -0.1264 | -0.20771 | 0.015943 | 0.04385 | -0.11095 | -0.11435 | -0.09878 | 0.08138 | -0.42633 | -0.20893 | -0.07161 | -0.00058 | -0.02618 | -0.07919 | -0.00254 | 0.00748 | 0.22485 | -0.28066 | 0.14501 | 0.06198 | 0.15475 | 0.050215 | -0.3495 |
| COLO678 | -0.13008 | -0.0281 | -0.41464 | -0.12631 | -0.23923 | -0.2573 | -0.22338 | -0.20619 | -0.05544 | 0.00566 | 0.087878 | 0.17165 | -0.11825 | -0.13772 | -0.12985 | -0.11121 | 0.24196 | 0.25628 | -0.02842 | 0.05718 | 0.01685 | -0.23286 | 0.051852 | -0.14584 | 0.12283 | -0.24425 | -0.68284 |
| HCC1428 | -0.08495 | 0.034219 | -0.01068 | 0.05425 | -0.107 | -0.20446 | 0.09942 | -0.25507 | -0.02455 | 0.0251 | -0.10171 | 0.13636 | -0.14809 | 0.05794 | -0.15318 | -0.01295 | -0.03322 | 0.29859 | -0.00946 | 0.05753 | 0.11367 | -0.19532 | 0.195109 | 0.10718 | 0.16223 | -0.0304 | -0.17624 |
| CAPAN1 | -0.10259 | 0.053108 | -0.14097 | -0.18084 | -0.15279 | 0.20661 | 0.021832 | -0.09533 | -0.05571 | -0.24971 | -0.02778 | 0.30102 | -0.19195 | -0.01627 | -0.06979 | -0.11476 | 0.06191 | 0.13588 | -0.08191 | 0.08733 | 0.04116 | -0.07935 | 0.086675 | -0.01284 | 0.11689 | 0.074787 | -0.29343 |
| NCIH82 | -0.30363 | -0.21083 | -0.20223 | 0.21115 | 0.06706 | -0.09367 | -0.00797 | -0.05711 | -0.34487 | 0.11367 | -0.01448 | -0.01047 | -0.02116 | 0.10036 | -0.12429 | 0.185109 | -0.25246 | -0.33859 | -0.12614 | 0.05579 | 0.16953 | -0.01523 | -0.03236 | -0.04967 | 0.21784 | 0.378681 | -0.65626 |
| MKN45 | -0.21437 | 0.215494 | -0.23668 | 0.21171 | -0.27863 | -0.33102 | -0.03066 | -0.21179 | -0.39018 | -0.08005 | -0.08363 | 0.20772 | -0.18149 | -0.09207 | -0.10185 | -0.15817 | -0.19911 | -0.00876 | -0.1042 | 0.08207 | 0.26642 | -0.28524 | -0.01881 | 0.01055 | 0.30738 | 0.169742 | -0.35845 |
| MG63 | -0.0434 | -0.05048 | -0.56466 | -0.19087 | -0.1655 | -0.03858 | -0.09036 | -0.41688 | 0.052201 | -0.08041 | -0.09277 | 0.06041 | -0.39965 | -0.40783 | -0.13875 | -0.04495 | -0.10933 | 0.14428 | -0.02674 | 0.20493 | 0.08735 | -0.45611 | 0.081534 | 0.04933 | 0.0209 | -0.12864 | -0.38293 |
| SKHEP1 | -0.0215 | 0.061903 | -0.08539 | 0.01913 | -0.27843 | -0.07939 | 0.18166 | -0.09015 | -0.00622 | -0.0488 | 0.139103 | -0.06221 | -0.28057 | -0.17295 | -0.12119 | -0.14171 | 0.08051 | 0.08554 | -0.11178 | 0.11724 | 0.17545 | 0.009372 | 0.030083 | 0.03631 | 0.06725 | -0.0473 | -0.37156 |
| MOLM13 | -1.76476 | 0.029995 | -0.08319 | -0.05287 | -0.14488 | -0.16126 | -0.10437 | -0.09945 | -0.10514 | -0.01198 | -0.02047 | 0.10519 | -0.19417 | -0.1915 | -0.09406 | -0.08219 | -0.04278 | 0.18794 | 0.021056 | 0.14584 | 0.08677 | -0.12773 | -0.08669 | -0.06597 | 0.28121 | 0.098237 | -0.48281 |
| SKMM2 | 0.11479 | -0.04778 | -0.76106 | -0.35356 | -0.1799 | -0.22226 | 0.020908 | -0.14174 | -0.11393 | 0.0479 | -0.06997 | 0.05876 | -0.13696 | 0.01651 | -0.09426 | 0.00022 | 0.06244 | -0.00065 | -0.0949 | 0.04619 | 0.06183 | -0.17687 | 0.032063 | -0.14822 | -0.0039 | 0.175626 | -0.29867 |
| U2OS | -0.01034 | 0.016449 | -0.74369 | -0.03157 | -0.11824 | 0.07402 | 0.13335 | -0.02217 | 0.114535 | -0.07761 | -0.11663 | -0.05461 | -0.21441 | -0.57744 | 0.043589 | -0.03761 | -0.23266 | 0.00232 | -0.0994 | 0.09909 | 0.08123 | -0.2824 | 0.169546 | 0.00499 | 0.20519 | 0.023889 | -0.31923 |
| SUDHL4 | -0.1458 | 0.017728 | -0.05723 | -0.00823 | -0.14891 | -0.19667 | 0.015156 | -0.17889 | -0.13806 | 0.19155 | 0.017644 | -0.08092 | -0.08381 | -0.07678 | -0.20444 | -0.1655 | 0.01303 | 0.09923 | -0.10682 | 0.12627 | -0.00214 | -0.06957 | -0.01109 | -0.15248 | 0.11706 | 0.024945 | -0.50662 |
| Cell line | FLT3 | FLT4 | IGF1R | INSR | INSRR | KDR | KIT | LMR1 | LMR2 | LTK | MET | MUSK | PDGFRA | PDGFRB | PTK7 | RET | RON | ROR1 | ROR2 | ROS1 | STYK1 | TIE1 | TIE2 | TRKA | TRKB | TRKC | TYRO3 |

| | | | | | | | | | | | | | | | | | | | | | | | | | | | |
|---|---|---|---|---|---|---|---|---|---|---|---|---|---|---|---|---|---|---|---|---|---|---|---|---|---|---|---|
| SKNDZ | -0.17701 | -0.03072 | -0.65843 | -0.20849 | -0.10473 | -0.31097 | -0.00679 | -0.34918 | -0.10495 | -0.03502 | -0.00758 | 0.04608 | -0.28149 | -0.41564 | -0.05591 | -0.2531 | -0.03985 | 0.15676 | 0.076182 | 0.16803 | 0.1342 | -0.04931 | 0.022461 | -0.119 | 0.29457 | -0.07865 | -0.56637 |
| NCIH226 | -0.12789 | 0.081294 | 0.03587 | -0.01264 | -0.16243 | -0.06237 | 0.341519 | -0.08493 | -0.13493 | 0.1386 | -0.41916 | 0.22531 | -0.1123 | -0.32848 | -0.11301 | -0.12046 | 0.17922 | 0.18039 | -0.05795 | -0.0492 | 0.22257 | -0.41588 | 0.075942 | -0.16111 | 0.06945 | 0.024995 | -0.08229 |
| SNU1105 | 0.15123 | -0.04421 | -0.10052 | -0.00566 | -0.13978 | -0.15549 | -0.32118 | -0.16101 | -0.06055 | 0.05791 | -0.37923 | 0.22546 | -0.31763 | -0.2931 | -0.01737 | -0.06262 | -0.05498 | 0.08236 | -0.1888 | 0.05662 | 0.18893 | -0.24036 | 0.136389 | -0.12247 | 0.22108 | -0.09117 | -0.40157 |
| SNU626 | -0.0753 | -0.11232 | -0.16096 | -0.1692 | -0.12087 | -0.25224 | 0.293398 | 0.00906 | -0.08891 | 0.03849 | -0.02267 | 0.17603 | 0.01088 | 0.0974 | -0.29429 | 0.040311 | 0.1931 | -0.02507 | -0.1424 | 0.1311 | 0.33263 | -0.07384 | 0.269846 | 0.02815 | 0.39684 | -0.09352 | -0.13445 |
| RL | -0.31437 | 0.001181 | -0.1789 | -0.08688 | -0.04037 | -0.05323 | -0.01955 | -0.1193 | -0.28701 | 0.32806 | -0.02716 | 0.0816 | -0.12665 | -0.2015 | -0.12881 | -0.22137 | -0.00721 | 0.23309 | -0.04916 | 0.10544 | 0.11731 | -0.04431 | 0.189178 | 0.01735 | 0.09926 | -0.03015 | -0.32412 |
| HCC1143 | -0.37182 | -0.02959 | 0.01338 | 0.08734 | -0.22602 | -0.29927 | -0.20165 | -0.32862 | -0.14056 | -0.04847 | -0.02278 | 0.07516 | -0.38081 | -0.11361 | 0.014831 | -0.16735 | -0.02222 | 0.07032 | -0.05482 | 0.06388 | 0.17212 | -0.10361 | 0.273648 | -0.03464 | 0.10083 | 0.014568 | -0.50711 |
| G402 | -0.04086 | 0.196798 | -0.18163 | -0.00076 | -0.10428 | 0.02179 | 0.024752 | 0.24721 | -0.3501 | 0.00506 | 0.04148 | 0.03246 | -0.1922 | -0.09871 | -0.02941 | -0.14382 | -0.04073 | -0.03609 | -0.08045 | 0.32798 | 0.02465 | -0.11824 | 0.037145 | -0.11472 | -0.03646 | 0.0567 | -0.48114 |
| SF295 | -0.01043 | -0.03741 | -0.12594 | -0.01357 | -0.21354 | -0.02228 | -0.13885 | -0.23215 | -0.09587 | 0.10347 | -0.67425 | 0.13372 | -0.05998 | -0.23282 | -0.06343 | -0.10302 | -0.09394 | 0.06203 | -0.04143 | 0.07358 | 0.13484 | -0.18178 | 0.028169 | -0.09981 | 0.16862 | 0.029793 | -0.28314 |
| SNU478 | -0.02502 | -0.0261 | -0.11303 | -0.17744 | -0.00398 | -0.01291 | -0.09915 | 0.06469 | -0.43789 | 0.02222 | -0.02269 | 0.20485 | -0.12456 | 0.07864 | -0.13742 | -0.07283 | 0.07866 | 0.18305 | 0.020617 | 0.19482 | -0.00104 | 0.045671 | 0.023727 | -0.41603 | 0.10468 | -0.04662 | -0.1519 |
| NCIH647 | -0.08121 | 0.044249 | -0.28717 | 0.17126 | -0.14674 | -0.18997 | 0.159709 | 0.04952 | -0.43374 | 0.22336 | -0.22245 | 0.09066 | -0.0041 | -0.05681 | -0.0766 | 0.171641 | 0.10598 | 0.22274 | -0.26407 | -0.07214 | 0.21309 | 0.002304 | 0.196156 | 0.06627 | -0.1432 | 0.128718 | -0.09151 |
| T84 | -0.12036 | 0.026205 | -0.61005 | -0.05724 | 0.09721 | -0.43439 | 0.164505 | -0.01786 | -0.22474 | 0.13339 | -0.07008 | 0.06202 | -0.31274 | 0.02483 | -0.04494 | -0.20366 | -0.16712 | -0.10677 | -0.00143 | 0.21435 | 0.26894 | -0.18531 | -0.20383 | -0.08922 | 0.31578 | 0.044136 | -0.28059 |
| OE33 | -0.06105 | 0.043485 | -0.19564 | -0.05208 | -0.15721 | -0.1555 | 0.025175 | -0.20102 | -0.26269 | -0.0198 | -1.44027 | 0.16824 | -0.04485 | -0.15138 | -0.1061 | 0.02949 | -0.09199 | 0.01246 | -0.14303 | 0.1195 | 0.26121 | -0.11933 | 0.060199 | -0.10447 | 0.1002 | -0.00837 | -0.35629 |
| SKRC20 | -0.0509 | -0.26446 | -0.0305 | 0.07415 | -0.15508 | -0.11278 | -0.08204 | -0.21906 | -0.11247 | 0.22259 | -0.47018 | 0.09308 | -0.26891 | -0.1899 | -0.07736 | -0.35704 | -0.00403 | 0.07344 | 0.201156 | -0.00093 | 0.115 | -0.44169 | 0.229443 | -0.09428 | 0.19624 | -0.21165 | -0.13605 |
| TF1 | -0.06413 | 0.07823 | -0.15311 | -0.08576 | 0.03265 | -0.14209 | -0.0592 | -0.33056 | -0.31285 | 0.06682 | -0.02732 | 0.04451 | -0.25011 | -0.12493 | -0.01172 | 0.033628 | -0.12719 | 0.17655 | -0.01166 | 0.03091 | 0.10223 | -0.48188 | -0.02118 | 0.04253 | 0.21069 | 0.002276 | -0.53391 |
| H4 | -0.20622 | -0.02482 | -0.16 | -0.16659 | -0.03519 | -0.02559 | 0.075761 | -0.58688 | -0.28132 | -0.0841 | -0.55663 | 0.0725 | -0.45787 | -0.32983 | -0.16274 | -0.08724 | -0.11554 | 0.11443 | 0.105162 | 0.04046 | -0.01955 | -0.12823 | 0.172507 | 0.0368 | 0.14121 | 0.045457 | -0.24657 |
| LUDLU1 | -0.16806 | 0.063444 | -0.79393 | 0.03324 | -0.15864 | -0.26819 | -0.04243 | -0.12422 | -0.19125 | -0.04514 | -0.00585 | -0.01529 | -0.12036 | -0.10558 | -0.25798 | -0.21328 | 0.04765 | 0.06142 | -0.26059 | 0.12524 | 0.20414 | -0.22168 | -0.03262 | -0.08063 | 0.18579 | -0.02635 | -0.55643 |
| MHHES1 | -0.1027 | -0.04757 | -0.95739 | -0.04598 | -0.09934 | -0.16911 | 0.011188 | -0.24438 | -0.01792 | -0.03139 | -0.13229 | 0.14362 | -0.31671 | -0.29092 | -0.03799 | -0.09632 | -0.06523 | -0.11001 | -0.03112 | 0.25224 | 0.14756 | -0.22348 | 0.051285 | 0.05374 | 0.18821 | 0.009907 | -0.44085 |
| HLF | -0.01107 | -0.01298 | -0.39164 | -0.04974 | -0.1317 | -0.15443 | -0.05675 | -0.15823 | -0.20913 | 0.01593 | -0.04438 | 0.07928 | -0.20293 | -0.15506 | -0.09264 | -0.07797 | -0.23202 | 0.26177 | 0.075677 | 0.18353 | 0.06817 | -0.13587 | 0.11201 | -0.12058 | 0.13435 | -0.04916 | -0.38612 |
| NCIH520 | -0.27031 | 0.016267 | -0.13327 | -0.26613 | -0.23085 | -0.37043 | -0.10488 | -0.26344 | -0.20256 | -0.06913 | 0.003932 | 0.05325 | -0.18462 | -0.07637 | -0.04601 | -0.14399 | -0.0341 | 0.08255 | -0.07293 | 0.04659 | 0.18319 | -0.20629 | 0.08121 | -0.06889 | 0.0629 | 0.047671 | -0.46871 |
| J82 | 0.08163 | 0.042533 | -0.09801 | 0.22157 | -0.04774 | -0.2676 | -0.23601 | 0.15986 | 0.039957 | -0.04179 | -0.14856 | 0.19055 | -0.42163 | -0.16512 | -0.1794 | -0.14497 | -0.08545 | 0.19649 | -0.1409 | 0.34474 | 0.01498 | -0.06112 | 0.102359 | -0.04923 | 0.10497 | 0.006174 | -0.33265 |
| TEN | -0.01581 | -0.05256 | -0.31561 | -0.15749 | -0.03373 | -0.39824 | -0.02018 | -0.08633 | -0.3157 | 0.02699 | 0.050523 | 0.12137 | -0.25231 | -0.1845 | 0.020441 | -0.02905 | -0.05297 | 0.09725 | -0.07905 | 0.02549 | 0.19375 | -0.00857 | 0.163817 | -0.08855 | 0.22593 | -0.0548 | -0.28284 |
| RI1 | 0.06258 | -0.08652 | -0.04149 | 0.01089 | -0.15306 | -0.01307 | -0.06664 | -0.29505 | -0.12785 | 0.02301 | 0.037479 | 0.04612 | -0.18267 | -0.22079 | -0.01627 | -0.09429 | -0.0912 | 0.08413 | -0.0583 | 0.21232 | 0.17704 | -0.10686 | 0.04173 | -0.04578 | 0.11445 | 0.073524 | -0.60245 |
| COLO800 | 0.02182 | -0.00373 | -0.0698 | -0.33099 | -0.03056 | -0.06246 | 0.006688 | -0.17171 | 0.11917 | -0.06184 | -0.15749 | 0.22045 | -0.53977 | -0.15343 | 0.154678 | -0.04743 | -0.27334 | -0.00231 | 0.054829 | 0.21018 | 0.13623 | -0.17888 | 0.022726 | 0.05746 | 0.14216 | -0.34238 | -0.31659 |
| BL70 | -0.02913 | -0.10657 | -0.02408 | 0.33406 | -0.31943 | -0.44219 | 0.007589 | 0.01824 | -0.09522 | -0.19732 | 0.040877 | 0.20579 | -0.0276 | -0.08155 | -0.32009 | -0.3676 | -0.20356 | 0.44493 | 0.020633 | 0.18288 | 0.11861 | -0.36271 | -0.08338 | 0.08891 | 0.18305 | 0.285974 | -0.36423 |
| NCIH747 | -0.04212 | -0.04289 | -0.31619 | -0.10721 | -0.07644 | -0.16502 | -0.03608 | -0.23096 | -0.2544 | -0.09782 | -0.01509 | 0.14061 | -0.17069 | -0.20428 | 0.016287 | -0.12079 | -0.01148 | 0.01399 | -0.12043 | 0.18227 | 0.21031 | -0.20057 | 0.180444 | -0.10949 | 0.23938 | 0.003345 | -0.50263 |
| K029AX | 0.19611 | -0.01827 | -0.39553 | 0.03983 | -0.01659 | 0.09542 | 0.005783 | 0.06596 | 0.022452 | -0.11785 | -0.00088 | 0.099 | 0.13365 | -0.26525 | -0.04109 | -0.05203 | -0.07938 | 0.08325 | -0.27794 | -0.13958 | 0.02499 | -0.48802 | 0.069996 | 0.18687 | 0.45829 | 0.031517 | -0.43186 |
| MEC1 | 0.04766 | -0.07339 | -0.07318 | -0.20212 | -0.05277 | -0.07652 | 0.051648 | -0.40694 | 0.130069 | 0.05828 | -0.32644 | 0.11391 | -0.17973 | -0.13106 | -0.04585 | -0.13569 | 0.04481 | 0.03952 | 0.077705 | 0.23661 | 0.17251 | -0.08489 | 0.007299 | 0.00795 | 0.293 | 0.000386 | -0.71694 |
| U937 | 0.01906 | -0.04612 | -0.30822 | -0.13613 | -0.0888 | -0.08311 | 0.032706 | -0.07702 | -0.09102 | -0.1198 | -0.118 | 0.13986 | -0.17644 | 0.01317 | -0.00909 | -0.05742 | -0.04368 | 0.09108 | -0.07834 | 0.13046 | 0.09757 | -0.26999 | 0.022441 | -0.00074 | 0.1806 | -0.10601 | -0.4439 |
| SNU685 | 0.04257 | 0.120539 | -0.13027 | -0.11684 | -0.14764 | -0.20335 | 0.128906 | 0.00837 | -0.08356 | 0.08868 | -0.03695 | 0.04281 | -0.17192 | -0.23821 | -0.05664 | -0.12283 | -0.06797 | 0.07282 | -0.18627 | 0.11242 | 0.15124 | -0.33969 | 0.044065 | 0.08204 | 0.19299 | -0.0143 | -0.39147 |
| TE5 | -0.25131 | 0.090106 | -0.12931 | 0.09792 | -0.13321 | -0.05913 | 0.211501 | -0.12552 | -0.06821 | 0.0281 | -0.04612 | 0.27328 | -0.15922 | -0.14942 | -0.0711 | -0.16723 | -0.0535 | 0.09494 | -0.03502 | 0.16316 | 0.24936 | -0.12956 | -0.00102 | 0.06967 | 0.14394 | 0.066536 | -0.23276 |
| SAOS2 | 0.04277 | 0.036801 | -0.56275 | 0.00282 | -0.11093 | -0.0769 | 0.003958 | -0.05325 | -0.12406 | -0.04767 | -0.03233 | 0.09687 | -0.4254 | -0.18702 | -0.05641 | -0.03402 | -0.00178 | 0.08263 | -0.04836 | 0.18956 | -0.02393 | 0.020998 | 0.120128 | -0.02424 | 0.13109 | -0.02083 | -0.31185 |
| 769P | -0.08089 | -0.02667 | -0.20041 | -0.07759 | 0.01419 | -0.09807 | 0.156763 | -0.19465 | -0.13753 | -0.00572 | -0.17154 | 0.14501 | -0.14922 | -0.17323 | -0.00791 | 0.080632 | -0.10043 | 0.1387 | -0.04818 | 0.11295 | 0.14903 | -0.10547 | 0.065853 | 0.08994 | 0.13282 | 0.019109 | -0.24097 |
| NCIH1944 | -0.21687 | -0.01319 | -0.37502 | -0.05564 | -0.05893 | -0.09425 | -0.0711 | -0.12209 | -0.2538 | 0.08344 | -0.13729 | 0.09914 | -0.14567 | -0.08057 | 0.004218 | -0.0326 | -0.07941 | -0.01183 | -0.03208 | 0.14207 | 0.09074 | -0.24004 | 0.132692 | -0.04403 | 0.17396 | 0.050806 | -0.47222 |
| BICR6 | -0.18388 | -0.00731 | -0.1879 | -0.00302 | -0.08757 | -0.18458 | -0.04093 | -0.06252 | -0.30194 | -0.13683 | 0.003301 | 0.17873 | -0.32013 | -0.12787 | -0.00061 | -0.30409 | -0.05003 | 0.17296 | 0.030059 | 0.19786 | -0.03013 | -0.08699 | 0.114144 | 0.04279 | 0.04522 | 0.12117 | -0.82376 |
| NCIH838 | -0.13521 | -0.02081 | -0.70008 | -0.03257 | -0.18139 | -0.18627 | -0.04618 | -0.20535 | -0.08398 | -0.06657 | -0.0711 | 0.04404 | -0.2051 | -0.02816 | -0.14392 | -0.05332 | -0.11468 | 0.16696 | 0.066839 | 0.06984 | 0.11291 | -0.09437 | 0.065697 | -0.10679 | 0.03741 | 0.00021 | -0.21691 |
| PANC0813 | -0.10889 | 0.005714 | -0.65709 | -0.04946 | -0.19853 | -0.09301 | -0.03275 | -0.04957 | -0.17602 | 0.04514 | -0.13882 | 0.1041 | -0.16101 | 0.00451 | -0.12648 | -0.19601 | -0.04026 | -0.03416 | 0.026279 | 0.11794 | 0.16228 | -0.23875 | 0.115469 | -0.02045 | 0.23791 | 0.018433 | -0.36516 |
| SNU449 | -0.13217 | 0.094331 | -0.33947 | -0.12999 | -0.23157 | -0.13365 | -0.16743 | -0.26447 | -0.0343 | -0.06155 | -0.00468 | 0.22725 | -0.10361 | 0.05847 | -0.00526 | -0.1251 | -0.01066 | 0.20329 | 0.019101 | 0.07871 | 0.23044 | -0.0101 | 0.003461 | -0.05052 | 0.19328 | -0.14218 | -0.35739 |
| SW837 | -0.07169 | 0.071027 | -0.79227 | -0.15644 | -0.09332 | -0.23026 | -0.04715 | -0.12018 | -0.03554 | -0.10467 | -0.04592 | 0.11744 | -0.20753 | -0.1319 | -0.17128 | -0.07646 | -0.06401 | 0.05657 | -0.10533 | 0.08955 | 0.24579 | -0.1175 | 0.052459 | -0.08812 | 0.19202 | -0.03208 | -0.49216 |
| SNU475 | -0.13196 | -0.09202 | -0.14546 | -0.01899 | -0.09048 | -0.15264 | 0.050141 | -0.35533 | -0.31935 | 0.09964 | -0.20148 | 0.07888 | -0.6304 | -0.57133 | -0.01908 | -0.05145 | -0.05407 | 0.11025 | 0.140759 | 0.12078 | -0.03386 | -0.0688 | 0.13528 | -0.07641 | -0.00385 | -0.00977 | -0.3061 |
| TC71 | -0.10371 | 0.108042 | -1.8046 | -0.06032 | -0.05436 | -0.30705 | 0.062391 | -0.11546 | -0.1003 | -0.03362 | -0.07772 | -0.01472 | -0.17431 | -0.20633 | -0.07374 | -0.18798 | -0.02306 | 0.10803 | -0.01055 | 0.09572 | -0.02217 | -0.16385 | -0.06325 | -0.06116 | 0.21709 | 0.028014 | -0.46749 |
| UACC62 | -0.21402 | -0.01156 | -0.14969 | -0.10414 | -0.01024 | -0.12599 | -0.06518 | -0.10247 | -0.27702 | 0.11676 | -0.00753 | 0.10899 | -0.19968 | -0.03665 | -0.02321 | -0.02994 | -0.15783 | 0.13942 | 0.007367 | 0.09858 | 0.08864 | -0.08747 | -0.01453 | 0.0019 | 0.16831 | 0.111571 | -0.47643 |
| KMS20 | -0.06848 | -0.20274 | -0.71599 | -0.05942 | -0.24756 | -0.35078 | 0.06127 | -0.3004 | 0.022334 | 0.00843 | -8.1E-05 | 0.30232 | -0.12791 | -0.29593 | 0.226228 | -0.03777 | -0.01337 | 0.01991 | 0.270664 | 0.16671 | 0.0472 | -0.38629 | -0.09791 | 0.04173 | 0.20374 | 0.165226 | -0.37801 |
| NCIN87 | -0.11839 | 0.252851 | -0.24377 | 0.02102 | 0.08524 | -0.0482 | 0.026477 | -0.00409 | -0.02283 | 0.01113 | -0.10716 | 0.16099 | -0.06316 | 0.0674 | -0.08503 | -0.04894 | -0.08461 | 0.19425 | 0.078631 | 0.0576 | 0.21628 | -0.0829 | -0.04125 | 0.03315 | 0.26606 | 0.248697 | -0.13035 |
| TYKNU | -0.23791 | -0.06105 | 0.04055 | -0.11465 | -0.35031 | -0.14964 | -0.0397 | -0.13947 | -0.15216 | -0.0526 | -0.0117 | 0.19723 | 0.00381 | -0.17689 | 0.010724 | -0.201 | -0.05355 | 0.06991 | -0.01361 | 0.07817 | 0.00379 | -0.12662 | 0.130279 | -0.2097 | 0.07107 | -0.0069 | -0.18749 |
| NCIH1694 | -0.13291 | 0.059212 | -0.37212 | -0.0787 | 0.02507 | -0.39194 | -0.11909 | -0.21948 | -0.19999 | -0.20048 | -0.041 | 0.10049 | -0.20215 | 0.02203 | -0.03327 | -0.1044 | -0.11702 | 0.00024 | -0.17636 | 0.21538 | 0.23394 | -0.04703 | 0.134038 | 0.08259 | 0.20816 | 0.117836 | -0.2918 |
| CAKI1 | -0.0682 | 0.023199 | -0.28286 | 0.05202 | -0.07366 | -0.14678 | 0.139019 | -0.29891 | -0.0705 | -0.06078 | 0.050704 | 0.03427 | 0.33772 | -0.2972 | -0.19275 | -0.00912 | -0.24063 | 0.09666 | -0.09505 | 0.14227 | 0.28084 | -0.25152 | 0.192356 | -0.06527 | 0.27849 | 0.028114 | -0.3594 |
| NCIH1915 | -0.11905 | 0.022749 | -0.03542 | -0.16313 | -0.15502 | -0.37016 | -0.22617 | -0.24242 | -0.21504 | -0.08862 | -0.04742 | 0.08823 | -0.2353 | -0.15755 | -0.28452 | -0.47269 | -0.01482 | -0.05501 | -0.1089 | 0.06476 | 0.08817 | -0.32926 | 0.173688 | -0.16801 | 0.0145 | -0.0598 | -0.78977 |
| EFE184 | -0.40488 | 0.109444 | -0.2533 | 0.04371 | 0.09317 | -0.09485 | 0.214249 | -0.35696 | -0.31482 | -0.05205 | -0.14543 | -0.02712 | -0.28178 | -0.05443 | -0.07472 | 0.155785 | 0.15389 | 0.23644 | 0.230793 | 0.06423 | -0.10848 | -0.2487 | 0.116661 | 0.05063 | 0.08681 | 0.061307 | -0.22229 |
| OCIMY7 | -0.41766 | 0.21935 | -0.39193 | -0.07996 | -0.04358 | -0.24656 | 0.005857 | -0.19196 | -0.08804 | 0.0889 | 0.09002 | 0.1863 | -0.08263 | -0.02685 | -0.12494 | -0.06476 | -0.12547 | 0.19978 | 0.170506 | 0.1214 | 0.1192 | 0.029921 | 0.021031 | -0.04277 | 0.26315 | -0.1291 | -0.28247 |
| SW1088 | -0.09023 | -0.19938 | -0.10431 | -0.12814 | -0.11579 | -0.0709 | 0.013469 | -0.1244 | -0.07799 | -0.02386 | -0.28922 | 0.1664 | -0.11598 | -0.51256 | -0.10207 | -0.12889 | -0.10889 | 0.17223 | -0.12994 | 0.044 | 0.13256 | -0.05569 | 0.230604 | -0.03846 | 0.02263 | 0.017775 | -0.18955 |
| LU65 | -0.13239 | 0.04791 | -0.48973 | 0.00792 | -0.13067 | -0.31456 | -0.22144 | -0.1522 | -0.03792 | 0.09503 | 0.139602 | 0.05686 | -0.21663 | -0.12299 | -0.22298 | -0.19245 | 0.07954 | 0.01082 | -0.1849 | -0.03061 | -0.08101 | -0.23867 | 0.07663 | -0.22844 | 0.02541 | 0.069643 | -0.56345 |
| SH4 | -0.19046 | 0.02644 | -0.23101 | -0.11985 | -0.03276 | -0.07159 | -0.15837 | -0.24717 | -0.0478 | 0.19503 | -0.09795 | 0.12095 | -0.17565 | -0.12395 | -0.09218 | -0.14237 | -0.0826 | 0.02581 | 0.001791 | 0.06812 | 0.1257 | -0.1763 | 0.04759 | -0.10889 | 0.05772 | -0.03905 | -0.46062 |
| RERFLCSQ1 | -0.19469 | -0.08153 | -0.06764 | 0.00138 | -0.17541 | -0.22652 | 0.00517 | -0.13478 | -0.29391 | -0.04576 | -0.13054 | 0.14566 | -0.08001 | -0.13605 | -0.01412 | -0.16601 | -0.18819 | 0.06589 | -0.08967 | 0.1708 | 0.03928 | -0.23472 | 0.064741 | -0.08691 | 0.19304 | -0.03816 | -0.37138 |
| LU99 | 0.10125 | 0.008139 | -0.4049 | -0.06524 | 0.00075 | -0.09868 | 0.145287 | -0.21374 | -0.06742 | 0.09072 | -0.01808 | 0.22574 | -0.21441 | -0.17813 | -0.08493 | -0.07032 | 0.0567 | 0.09716 | 0.122316 | 0.13152 | -0.14265 | -0.0794 | 0.094712 | -0.03435 | 0.23136 | 0.077213 | -0.29125 |
| KNS60 | 0.09882 | 0.134891 | -0.22663 | -0.01616 | -0.05878 | -0.3299 | 0.047721 | -0.17277 | -0.30247 | -0.01815 | -0.19972 | 0.13719 | -0.28374 | -0.39931 | -0.15646 | -0.06722 | 0.19311 | -0.00233 | 0.037898 | 0.04796 | 0.18461 | -0.15178 | -0.04267 | 0.2358 | 0.1086 | 0.056661 | -0.35874 |
| NCIH1666 | -0.15585 | 0.004037 | -0.55836 | -0.10988 | -0.05891 | -0.28652 | -0.14259 | -0.156 | -0.04712 | -0.03957 | -0.04125 | 0.13775 | -0.22348 | -0.30789 | -0.06873 | -0.03383 | 0.04874 | -0.0081 | 0.028925 | 0.10126 | 0.1944 | -0.16454 | 0.025786 | -0.14259 | 0.01448 | -0.06465 | -0.32224 |
| MELHO | 0.04109 | 0.125402 | 0.14184 | -0.0237 | -0.01156 | -0.30667 | -0.25611 | -0.1362 | -0.39677 | -0.19602 | -0.06565 | 0.08737 | -0.4809 | -0.23488 | -0.19325 | -0.05108 | 0.16388 | -0.28251 | 0.03809 | 0.2877 | 0.16786 | -0.19786 | 0.057222 | -0.05857 | 0.46555 | 0.1013 | -0.82631 |
| TE8 | -0.15864 | 0.047093 | -0.62666 | -0.0298 | -0.06618 | -0.22187 | 0.091365 | -0.12515 | -0.14369 | -0.0122 | -0.04241 | 0.08593 | -0.06269 | -0.16732 | 0.093712 | -0.01452 | -0.09144 | 0.11755 | -0.07006 | 0.09936 | 0.01694 | -0.14987 | 0.020488 | -0.04054 | 0.11276 | -0.07344 | -0.32717 |
| HCC95 | -0.04156 | -0.14111 | -0.32194 | -0.01486 | -0.24161 | -0.26976 | 0.005066 | -0.15429 | -0.03808 | -0.18745 | 0.057503 | 0.05202 | -0.25051 | -0.3107 | -0.07661 | -0.26326 | -0.32562 | 0.42513 | -0.28936 | 0.17503 | 0.24676 | -0.24359 | 0.244711 | -0.13996 | 0.11087 | 0.160947 | -0.53556 |
| LN428 | -0.10854 | -0.02847 | -0.23397 | -0.13317 | -0.14089 | 0.00679 | -0.04666 | -0.18173 | -0.16942 | -0.04392 | -0.05895 | 0.14426 | -0.19831 | -0.11728 | -0.1092 | -0.15787 | -0.13019 | -0.14098 | -0.05432 | 0.16531 | 0.05969 | -0.15505 | -0.03514 | -0.10844 | 0.17417 | 0.028284 | -0.38364 |
| BCPAP | -0.07926 | 0.008779 | -0.23558 | -0.17858 | -0.14715 | -0.23551 | -0.05184 | -0.2689 | 0.011685 | -0.09997 | -0.20548 | 0.08483 | -0.18753 | -0.1796 | -0.09405 | -0.11294 | -0.02587 | -0.08264 | -0.00131 | 0.14948 | 0.05802 | -0.08926 | 0.118412 | -0.06171 | 0.06776 | 0.057983 | -0.34477 |
| CJM | -0.18533 | -0.05701 | -0.54502 | -0.19079 | -0.18022 | -0.01854 | -0.14134 | -0.21049 | -0.25134 | 0.05475 | -0.0418 | 0.10746 | 0.02751 | -0.18847 | -0.05447 | -0.11069 | -0.1122 | 0.27274 | -0.08223 | 0.15495 | 0.01872 | -0.13828 | 0.095277 | -0.1821 | 0.11637 | -0.05374 | -0.12777 |
| TUHR10TKB | -0.07679 | -0.08393 | -0.23145 | 0.01755 | -0.42701 | -0.35623 | -0.25186 | -0.09569 | -0.28623 | 0.10321 | 0.169795 | -0.1523 | -0.34841 | -0.00835 | -0.22155 | -0.10978 | -0.32447 | 0.27943 | 0.140945 | 0.10287 | 0.10535 | -0.62774 | 0.163266 | -0.07408 | 0.1018 | 0.052436 | -0.16903 |
| SNU8 | -0.05463 | -0.04933 | -0.18026 | -0.09203 | -0.0725 | -0.1634 | -0.03185 | -0.13037 | -0.10692 | -0.05709 | -0.03526 | 0.02354 | -0.18613 | -0.01766 | 0.007761 | -0.05395 | 0.02128 | -0.0643 | -0.0738 | 0.02699 | 0.0787 | -0.17605 | 0.115732 | -0.05162 | 0.11493 | 0.073354 | -0.31882 |
| SNU1196 | -0.22977 | 0.123519 | -0.75937 | -0.09915 | -0.11557 | -0.17791 | -0.17725 | -0.19831 | -0.04492 | -0.0112 | -0.15576 | -0.00316 | -0.08926 | -0.12309 | 0.198321 | -0.24978 | -0.2184 | 0.05371 | 0.037355 | 0.07836 | 0.19519 | -0.03065 | 0.130045 | -0.25709 | 0.22019 | 0.172146 | -0.20792 |
| NCIH460 | -0.19387 | 0.123504 | -0.21235 | -0.09749 | -0.07136 | -0.19458 | 0.063157 | -0.27949 | -0.13941 | -0.04905 | 0.038086 | 0.18137 | -0.11766 | -0.25734 | -0.1904 | -0.24015 | -0.18878 | 0.12795 | -0.07756 | 0.13064 | 0.16117 | -0.03228 | -0.04658 | -0.05967 | 0.02906 | -0.08232 | -0.49294 |
| CAS1 | -0.03437 | 0.009581 | -0.05297 | -0.15762 | -0.13995 | -0.22413 | 0.014233 | -0.20949 | -0.06392 | 0.00295 | -0.18598 | 0.26371 | -0.48953 | -0.58136 | 0.010986 | 0.093543 | -0.02363 | 0.19213 | -0.03806 | 0.09811 | 0.16556 | -0.17957 | -0.0184 | 0.02169 | 0.21011 | 0.117766 | -0.39113 |
| SNU216 | -0.0454 | 0.045465 | -0.21886 | 0.12152 | -0.23149 | -0.21041 | 0.079001 | -0.34219 | -0.17337 | 0.10276 | -0.04528 | 0.10771 | -0.17975 | 0.02001 | 0.035964 | -0.17091 | -0.07087 | 0.08959 | -0.03312 | 0.06995 | 0.11692 | -0.24857 | -0.00957 | -0.0642 | 0.073 | 0.086934 | -0.36567 |
| HCC56 | -0.16213 | -0.02983 | -0.65647 | -0.33784 | -0.18496 | -0.23776 | 0.028908 | -0.11312 | 0.110834 | -0.09642 | -0.01351 | 0.26928 | -0.05886 | -0.29357 | -0.02937 | -0.14663 | -0.02042 | 0.22353 | -0.22354 | -0.06111 | 0.29319 | -0.09379 | 0.127486 | -0.21852 | 0.09761 | -0.03007 | -0.36145 |
| PK45H | -0.23403 | -0.1081 | -0.20007 | -0.07328 | -0.36934 | -0.1835 | -0.11502 | -0.183 | -0.29157 | -0.17256 | -0.03048 | 0.04817 | -0.43341 | -0.08263 | -0.13108 | -0.09881 | -0.08194 | 0.01884 | -0.05765 | -0.117 | -0.01444 | -0.13842 | -0.00052 | -0.05707 | 0.2516 | 0.124407 | -0.63563 |
| YH13 | -0.07608 | -0.03054 | -0.04373 | 0.05684 | -0.15428 | -0.07411 | -0.05954 | -0.21442 | -0.12892 | 0.02983 | -0.90786 | 0.17706 | -0.44043 | -0.31565 | -0.01125 | -0.05405 | 0.01713 | 0.00341 | 0.114093 | 0.11541 | 0.21975 | -0.09126 | 0.052898 | -0.1127 | 0.03541 | 0.091068 | -0.20235 |
| SW1463 | -0.0182 | -0.07595 | -0.28423 | 0.01335 | -0.04351 | -0.2582 | 0.083517 | -0.43274 | -0.30271 | -0.2929 | -0.01674 | 0.18591 | 0.01307 | -0.11067 | 0.052829 | -0.1838 | 0.09016 | 0.17318 | 0.006713 | 0.09049 | 0.19574 | 0.050269 | 0.256866 | -0.10728 | 0.10284 | -0.0032 | -0.52826 |
| LI7 | -0.14622 | -0.12598 | -0.5015 | -0.06182 | -0.07294 | -0.07823 | 0.045367 | -0.18682 | 0.113863 | -0.09261 | -0.07371 | 0.11628 | -0.00833 | -0.06903 | -0.06787 | -0.07688 | -0.10538 | 0.21126 | 0.051009 | -0.02036 | 0.10605 | -0.08756 | 0.133504 | -0.06443 | 0.0918 | 0.107178 | -0.27833 |
| HSC2 | 0.02817 | -0.00181 | -0.2755 | -0.118 | -0.07561 | -0.18982 | 0.084171 | -0.15648 | -0.16619 | 0.02308 | -0.12596 | 0.16756 | -0.24786 | -0.10807 | -0.14911 | -0.11157 | 0.0092 | -0.06323 | -0.09191 | 0.10923 | 0.00158 | -0.17841 | 0.014534 | 0.0251 | 0.14673 | 0.04414 | -0.42459 |
| RT112 | -0.13235 | 0.025874 | -0.75972 | 0.15753 | -0.20933 | -0.28226 | -0.0122 | -0.26613 | -0.01391 | -0.3957 | 0.014363 | 0.14664 | -0.06223 | -0.29114 | -0.02205 | -0.2297 | 0.12075 | 0.23819 | 0.038928 | 0.12927 | 0.33053 | -0.44179 | 0.133627 | -0.18722 | 0.28131 | -0.14199 | -0.60968 |

| Cell line | FLT3 | FLT4 | IGF1R | INSR | INSRR | KDR | KIT | LMR1 | LMR2 | LTK | MET | MUSK | PDGFRA | PDGFRB | PTK7 | RET | RON | ROR1 | ROR2 | ROS1 | STYK1 | TIE1 | TIE2 | TRKA | TRKB | TRKC | TYRO3 |
|---|---|---|---|---|---|---|---|---|---|---|---|---|---|---|---|---|---|---|---|---|---|---|---|---|---|---|---|
| HUH1 | -0.25044 | -0.04196 | -0.56744 | -0.20674 | -0.05257 | -0.16695 | -0.00405 | -0.21149 | -0.23888 | 0.02153 | -0.11493 | 0.15252 | -0.02175 | -0.1399 | -0.13378 | 0.05024 | -0.03205 | 0.08693 | 0.048306 | 0.03586 | 0.20652 | -0.05188 | 0.136077 | -0.21188 | 0.11986 | -0.08467 | -0.33907 |
| JHH4 | -0.18396 | 0.041932 | -0.13648 | -0.16859 | -0.11189 | -0.14122 | 0.000849 | -0.11005 | -0.09258 | -0.02662 | -0.06203 | 0.26254 | -0.12404 | 0.03714 | 0.05533 | -0.01716 | 0.21626 | 0.02453 | -0.01946 | 0.17 | 0.16105 | -0.25904 | 0.105676 | -0.00924 | 0.16066 | 0.092063 | -0.27057 |
| MALME3M | 0.00817 | -0.08626 | -0.46396 | -0.052 | 0.05422 | -0.1131 | 0.036624 | -0.12511 | -0.27223 | 0.03965 | -0.14295 | 0.20581 | -0.24985 | -0.05428 | 0.000577 | -0.2002 | -0.02699 | -0.06684 | -0.12094 | -0.10044 | 0.01529 | 0.134393 | -0.058 | -0.10034 | 0.34528 | -0.0308 | -0.76616 |
| SNU387 | 0.03051 | -0.01784 | -0.83364 | -0.17823 | -0.19765 | -0.33855 | 0.031837 | -0.01662 | -0.0042 | -0.07932 | -0.11747 | -0.04183 | -0.14649 | -0.04824 | 0.104208 | 0.091124 | -0.00103 | 0.25636 | -0.00295 | 0.09427 | 0.13828 | -0.14529 | 0.053316 | -0.00132 | 0.18351 | -0.12051 | -0.24658 |
| KNS81 | 0.03218 | 0.233385 | -0.25172 | -0.14282 | -0.18798 | -0.45837 | 0.070355 | 0.10656 | -0.17816 | 0.09422 | 0.066298 | -0.13727 | -0.27307 | -0.47151 | 0.060126 | -0.33007 | -0.14201 | 0.11934 | 0.038532 | 0.26764 | 0.09411 | -0.04427 | 0.048891 | -0.08088 | 0.20068 | -0.28185 | -0.51758 |
| HUH7 | 0.0047 | 0.008304 | -0.19056 | -0.39546 | -0.12577 | -0.31496 | 0.33686 | -0.27007 | -0.02246 | -0.25786 | -0.093 | 0.20301 | -0.04386 | -0.10595 | -0.15223 | -0.20666 | 0.05313 | 0.36973 | -0.09582 | 0.09609 | 0.14827 | -0.10517 | 0.199784 | -0.09854 | 0.01252 | -0.28758 | -0.32866 |
| NCIH2170 | -0.11155 | 0.036849 | -0.06332 | -0.08172 | -0.31005 | -0.4181 | -0.07454 | -0.24714 | -0.20506 | -0.17091 | 0.004437 | 0.18696 | -0.21908 | -0.30908 | -0.14401 | -0.27693 | -0.10827 | 0.08478 | -0.14973 | 0.08951 | 0.20156 | -0.05503 | -0.02826 | -0.14055 | 0.17039 | 0.006504 | -0.59338 |
| SNU182 | -0.04949 | -0.19245 | -0.2775 | -0.24441 | -0.17359 | -0.29049 | -0.11017 | -0.18087 | -0.41449 | -0.01425 | -0.12069 | 0.147 | -0.19644 | -0.1129 | -0.04053 | -0.02748 | -0.19381 | -0.11642 | -0.08947 | 0.18998 | 0.2395 | -0.22254 | -0.01468 | 0.00636 | 0.15626 | 0.298057 | -0.33026 |
| VMRCRCW | 0.01732 | 0.003499 | -0.09369 | 0.11289 | -0.33611 | 0.03883 | 0.239332 | -0.418 | 0.162313 | 0.1642 | 0.126486 | -0.02109 | -0.02804 | -0.35392 | -0.2765 | -0.02837 | -0.17085 | 0.2085 | 0.130884 | 0.05695 | 0.22119 | -0.26644 | 0.128198 | 0.04111 | 0.32591 | -0.13226 | -0.25993 |
| GSU | -0.348 | 0.084357 | -0.36863 | -0.03769 | -0.38193 | -0.31016 | -0.13796 | -0.10169 | -0.11951 | 0.08738 | -0.05708 | 0.19649 | -0.32249 | -0.44675 | -0.16166 | -0.05354 | 0.17513 | -0.10386 | 0.000657 | -0.0181 | 0.11268 | -0.02757 | 4.61E-05 | -0.11825 | -0.0105 | 0.043512 | -0.34889 |
| KU1919 | -0.21115 | 0.022356 | -0.13917 | 0.09458 | -0.14646 | -0.09896 | -0.10829 | -0.20107 | -0.10534 | 0.01445 | -0.01243 | 0.11864 | -0.26136 | -0.04088 | -0.05728 | -0.00298 | 0.03117 | 0.00351 | 0.020503 | 0.07869 | 0.00581 | -0.16015 | 0.044213 | -0.00043 | 0.05621 | 0.118724 | -0.30864 |
| F36P | -0.12063 | 0.098195 | -0.58737 | 0.08887 | -0.1328 | -0.10528 | -0.0321 | -0.23738 | 0.057038 | -0.06584 | -0.03161 | 0.06379 | -0.14056 | -0.10168 | -0.20793 | -0.09467 | -0.04191 | 0.27241 | 0.146904 | 0.17907 | -0.02351 | -0.23157 | 0.147625 | -0.12232 | 0.14369 | -0.07225 | -0.39412 |
| TE11 | -0.1025 | -0.0489 | -0.30659 | -0.00211 | -0.1463 | -0.10725 | -0.01162 | -0.22256 | -0.13816 | 0.01 | -0.09368 | 0.19814 | -0.13703 | -0.18236 | -0.10521 | -0.09372 | 0.06142 | 0.22987 | -0.12708 | 0.11324 | 0.09453 | -0.15777 | 0.049816 | -0.03754 | 0.1689 | -6E-05 | -0.24994 |
| SW1116 | -0.07834 | -0.03981 | -0.38517 | -0.16109 | -0.06141 | -0.10546 | -0.02875 | -0.27783 | -0.15131 | 0.08706 | -0.11671 | 0.16295 | -0.08291 | -0.18679 | -0.10923 | 0.001594 | -0.00095 | -0.01697 | -0.10444 | 0.12156 | 0.14321 | -0.18412 | 0.034254 | -0.04658 | -0.02445 | -0.03241 | -0.30142 |
| SF767 | -0.16716 | -0.03421 | -0.13432 | -0.01033 | -0.31709 | -0.04637 | 0.074647 | -0.23711 | -0.29346 | -0.15856 | -0.053 | 0.13784 | -0.1692 | -0.32237 | -0.08727 | -0.062 | -0.11014 | 0.21389 | -0.10103 | 0.08442 | -0.03355 | -0.11887 | 0.132307 | -0.04531 | -0.059 | 0.055295 | -0.40207 |
| NCIH716 | -0.23553 | 0.130137 | -0.19376 | -0.13794 | -0.2748 | -0.14811 | 0.002452 | 0.02459 | -0.05281 | -0.04363 | 0.09668 | -0.02378 | -0.30976 | -0.29558 | -0.10082 | -0.28854 | -0.3383 | 0.33312 | -0.1515 | 0.30767 | 0.11672 | -0.09201 | 0.114617 | 0.03286 | -0.04339 | -0.03306 | -0.52676 |
| SNU423 | -0.0845 | 0.018758 | -0.27698 | -0.11279 | -0.2 | -0.08594 | 0.008929 | -0.2098 | 0.123589 | -0.20887 | -0.13058 | 0.14818 | -0.35432 | -0.10351 | -0.07793 | -0.05826 | -0.09176 | 0.13074 | 0.009738 | 0.23025 | 0.14639 | -0.00361 | 0.261751 | 0.05782 | 0.18294 | 0.048246 | -0.22591 |
| TUHR4TKB | 0.00055 | -0.09143 | -0.19583 | -0.17471 | -0.22024 | -0.17549 | -0.00646 | -0.27485 | -0.12193 | -0.10131 | 0.087909 | 0.11471 | -0.00381 | -0.22245 | 0.037051 | -0.01507 | -0.21404 | -0.00886 | -0.11522 | 0.3178 | -0.01878 | -0.32161 | -0.00157 | -0.01094 | 0.08147 | 0.021563 | -0.26821 |
| NCIH1792 | -0.24964 | -0.15177 | -0.60108 | -0.0396 | -0.09016 | -0.2692 | -0.0687 | -0.04528 | -0.09127 | -0.08766 | -0.03702 | -0.00667 | -0.15188 | 0.12841 | -0.12477 | -0.045 | -0.13919 | 0.01894 | -0.06077 | 0.07143 | -0.00779 | -0.19532 | 0.057274 | -0.03052 | 0.01022 | 0.095875 | -0.44496 |
| EW8 | -0.12306 | -0.11478 | -0.62038 | -0.2631 | -0.14988 | -0.12108 | 0.087128 | -0.01351 | 0.037185 | 0.14113 | -0.14269 | 0.05228 | 0.04237 | 0.13777 | -0.10483 | -0.12308 | -0.15552 | 0.26518 | 0.041623 | 0.09161 | 0.07931 | -0.27684 | -0.03076 | 0.00846 | 0.18549 | -0.0075 | -0.48168 |
| SNU46 | 0.03127 | 0.105013 | -0.40267 | -0.08119 | -0.16903 | -0.40843 | 0.197698 | -0.15928 | -0.61244 | 0.04227 | 0.067067 | 0.12826 | -0.35855 | 0.02902 | 0.137478 | -0.38999 | -0.30941 | 0.2624 | 0.05455 | 0.18359 | -0.31134 | -0.31966 | -0.2024 | 0.0404 | 0.36935 | 0.237406 | -0.4079 |
| LS123 | 0.02919 | 0.478937 | 0.08115 | -0.01193 | 0.28142 | 0.04687 | 0.406679 | 0.2232 | 0.02235 | 0.12335 | -0.19072 | 0.04809 | 0.1028 | 0.37225 | 0.001892 | 0.358435 | 0.16328 | 0.1487 | -0.08329 | 0.24387 | 0.2812 | 0.259283 | -0.29194 | 0.20185 | -0.07659 | 0.356501 | 0.1313 |
| TCCPAN2 | -0.16628 | -0.01331 | -1.07127 | -0.15394 | -0.14306 | -0.20519 | -0.12242 | -0.36279 | -0.17516 | -0.08338 | -0.05133 | 0.11853 | -0.23641 | -0.09735 | -0.10849 | -0.11421 | 0.06576 | 0.03287 | -0.03762 | 0.12424 | 0.10279 | -0.14528 | 0.076623 | -0.06453 | 0.10257 | -0.00458 | -0.27562 |
| BICR16 | 0.11687 | 0.046421 | -0.14278 | -0.21336 | -0.07866 | -0.55083 | -0.03581 | -0.18362 | -0.16074 | 0.072 | -0.14609 | 0.10545 | 0.11054 | -0.5066 | -0.00465 | 0.002124 | 0.10785 | 0.12808 | 0.059961 | 0.02367 | 0.17193 | -0.34042 | 0.142588 | -0.00925 | 0.22408 | 0.006447 | -0.66911 |
| SNB75 | 0.02691 | 0.053476 | -0.06871 | -0.18366 | -0.23929 | -0.21518 | 0.166762 | -0.04797 | 0.05328 | -0.20514 | -0.02951 | 0.15358 | -0.45025 | -0.16139 | -0.05429 | 0.3271 | -0.02032 | 0.15986 | -0.11409 | 0.10735 | 0.23629 | -0.11373 | 0.077173 | -0.00075 | 0.18558 | 0.168403 | -0.37756 |
| RKN | -0.13133 | -0.05888 | -0.15368 | -0.04897 | -0.23972 | -0.27011 | 0.100454 | -0.10138 | -0.1713 | -0.09233 | -0.13174 | 0.09542 | -0.03314 | -0.06667 | -0.01799 | -0.07791 | -0.12006 | 0.05946 | -0.14488 | 0.13556 | 0.12178 | -0.18775 | 0.083751 | -0.14874 | 0.02672 | 0.049295 | -0.34752 |
| KE39 | -0.1523 | 0.032549 | -0.74676 | -0.35943 | -0.08798 | -0.17691 | -0.05209 | -0.05811 | -0.08667 | 0.03693 | -0.04294 | 0.12622 | -0.11834 | -0.06176 | -0.05804 | 0.102385 | 0.02298 | 0.136 | -0.08522 | 0.16388 | 0.17525 | -0.30408 | 0.139051 | -0.06262 | 0.01342 | 0.001947 | -0.30546 |
| NCIH1299 | -0.08555 | 0.07825 | -0.41647 | -0.0603 | -0.13217 | -0.09622 | 0.048401 | -0.1692 | -0.1575 | 0.03846 | -0.05049 | 0.04736 | -0.2279 | -0.2255 | -0.12356 | -0.07969 | -0.04421 | 0.09635 | -0.02277 | 0.01638 | 0.12763 | -0.1703 | 0.01223 | -0.08866 | 0.09789 | -0.05759 | -0.34973 |
| CALU1 | -0.08469 | 0.013837 | -0.34867 | -0.00102 | -0.14467 | -0.24494 | 0.071192 | -0.10778 | -0.13878 | -0.0887 | -0.18091 | 0.03462 | -0.128 | -0.0351 | -0.1082 | -0.19696 | -0.12914 | 0.17312 | 0.08248 | 0.18979 | 0.01661 | -0.04439 | 0.096346 | 0.01556 | 0.12211 | 0.05237 | -0.1471 |
| INA6 | -0.15926 | 0.0331 | -0.60158 | -0.1929 | -0.04696 | -0.11503 | -0.20861 | -0.03042 | -0.0825 | -0.03377 | -0.03741 | -0.12244 | -0.14848 | -0.18758 | 0.04082 | -0.2183 | -0.4809 | 0.26279 | -0.22688 | 0.07976 | 0.04873 | -0.34329 | -0.34122 | 0.02127 | 0.15504 | 0.11077 | -0.58416 |
| NCIH1092 | -0.22387 | 0.156167 | -0.1605 | -0.39459 | -0.13272 | -0.30857 | 0.017564 | -0.11734 | -0.22331 | -0.0616 | -0.01964 | 0.06942 | -0.11195 | 0.02702 | -0.04416 | -0.06753 | -0.1307 | 0.28657 | 0.129001 | 0.18624 | 0.06056 | -0.16464 | 0.169972 | -0.074 | 0.11628 | 0.037447 | -0.47715 |
| CAL78 | -0.07243 | 0.110526 | -0.28804 | 0.00719 | -0.10854 | -0.10578 | 0.080256 | -0.14213 | 0.016081 | 0.06687 | -0.13842 | 0.14203 | -0.01842 | -0.07898 | -0.09405 | -0.08959 | -0.21245 | 0.08732 | -0.12925 | 0.13342 | 0.06907 | -0.32847 | 0.026904 | -0.0243 | 0.14683 | 0.038282 | -0.33548 |
| SNU410 | -0.08513 | 0.010331 | -0.44839 | -0.11431 | -0.10398 | -0.17651 | 0.038929 | -0.16926 | -0.14146 | -0.09802 | -0.12983 | 0.13343 | -0.05005 | -0.14063 | -0.15512 | -0.06154 | -0.07675 | 0.03482 | -0.02155 | 0.08976 | 0.11181 | -0.18147 | 0.088923 | -0.06488 | 0.16263 | -0.03211 | -0.30314 |
| CAL33 | -0.18721 | 0.16536 | -0.20936 | 0.03419 | 0.00879 | -0.04084 | 0.141342 | -0.17691 | -0.11112 | -0.04664 | -0.009 | 0.1113 | -0.09792 | -0.06064 | -0.02861 | -0.1101 | 0.00213 | 0.2497 | -0.10249 | 0.17876 | 0.10395 | -0.1506 | -0.08227 | 0.09073 | 0.21044 | 0.02367 | -0.21998 |
| 59M | -0.18225 | 0.045524 | -0.34043 | -0.04675 | 0.00798 | -0.11148 | 0.20619 | -0.06366 | -0.00025 | -0.10563 | -0.11933 | 0.0703 | -0.04067 | -0.0496 | -0.1929 | 0.032546 | -0.22248 | 0.18135 | 0.144292 | -0.00531 | 0.20456 | -0.13629 | -0.12728 | 0.02441 | 0.1643 | -0.05085 | -0.34933 |
| NCIH2030 | -0.09554 | -0.0651 | -0.11472 | -0.16993 | -0.19218 | -0.1834 | -0.06233 | -0.51518 | 0.004777 | 0.00962 | -0.05555 | -0.03773 | -0.16832 | 0.04167 | -0.13587 | -0.10747 | 0.0225 | 0.01798 | -0.14725 | 0.15287 | -0.15536 | -0.16949 | 0.056694 | -0.03751 | 0.13169 | 0.03625 | -0.55836 |
| UMUC3 | -0.12591 | 0.127963 | -0.13085 | -0.07781 | -0.17004 | -0.25494 | -0.03463 | -0.11072 | -0.11842 | -0.0176 | -0.10475 | 0.05178 | -0.34394 | -0.14496 | -0.09772 | -0.1759 | -0.04665 | 0.20722 | 0.078689 | 0.18319 | 0.07412 | -0.05039 | 0.140475 | -0.09088 | 0.03926 | 0.089094 | -0.18299 |
| KURAMOCHI | -0.03684 | -0.02847 | -0.11368 | -0.05218 | -0.10104 | -0.13898 | -0.01824 | -0.20596 | -0.30598 | -0.00704 | -0.09816 | 0.14735 | -0.08589 | -0.09524 | -0.02932 | -0.01229 | -0.08532 | 0.06411 | 0.004622 | 0.06836 | 0.19159 | -0.14235 | 0.112832 | -0.04886 | 0.16745 | -0.01116 | -0.21229 |
| NCIH2171 | -0.1556 | -0.00216 | -0.00112 | -0.13202 | -0.09519 | -0.07603 | 0.029402 | -0.2106 | -0.11145 | -0.0912 | -0.07966 | 0.19585 | -0.10541 | -0.2817 | -0.17329 | -0.00921 | -0.12869 | 0.05474 | -0.04675 | 0.09812 | 0.20014 | -0.11094 | 0.024102 | -0.01687 | 0.16284 | 0.0369 | -0.45323 |
| OVISE | -0.10506 | -0.02659 | -0.23323 | -0.00494 | -0.13316 | -0.10263 | -0.04611 | -0.18653 | -0.16685 | -0.18187 | -0.1211 | 0.12897 | -0.13805 | -0.12069 | -0.06277 | -0.03424 | -0.00093 | 0.0144 | -0.02856 | 0.1376 | 0.0945 | -0.08721 | 0.067143 | -0.09999 | 0.20623 | -0.01165 | -0.2755 |
| ABC1 | -0.00544 | -0.01218 | -0.73841 | -0.10173 | -0.0526 | -0.05879 | -0.02854 | -0.12846 | -0.19547 | -0.22335 | -0.14642 | 0.10067 | -0.16653 | -0.12783 | -0.08343 | 0.017747 | -0.02504 | 0.12718 | -0.1253 | 0.13693 | -0.03374 | -0.22398 | 0.071023 | -0.13646 | 0.15077 | -0.00215 | -0.26353 |
| SNU61 | -0.07822 | -0.10237 | -0.74451 | -0.20501 | -0.11404 | -0.0125 | -0.14168 | -0.16974 | -0.06308 | -0.05236 | -0.02554 | 0.17689 | -0.14957 | -0.26081 | -0.02061 | 0.044942 | -0.0738 | -0.01031 | 0.000786 | 0.21698 | 0.00472 | -0.0395 | 0.118423 | 0.00038 | 0.14534 | 0.074839 | -0.48666 |
| NCIH2004RT | -0.26207 | -0.04812 | 0.1263 | -0.3878 | 0.04604 | -0.08901 | -0.2321 | -0.44389 | 0.073991 | -0.15548 | -0.15988 | 0.02687 | -1.73089 | -0.47688 | -0.08639 | -0.05801 | -0.11147 | 0.13307 | 0.30733 | 0.0234 | 0.05689 | -0.06245 | -0.01709 | 0.04127 | 0.23123 | 0.222946 | -0.4559 |
| BXPC3 | -0.125 | -0.0255 | -0.47262 | -0.04267 | -0.17529 | -0.16053 | -0.05221 | -0.03294 | -0.09583 | -0.05465 | -0.03967 | 0.05971 | -0.19278 | -0.04713 | -0.06461 | -0.1434 | 0.01854 | 0.25241 | -0.13692 | 0.073 | 0.19458 | -0.21681 | 0.103566 | -0.07814 | -0.04598 | -0.05476 | -0.62671 |
| SNU761 | -0.16808 | 0.078631 | -0.19849 | -0.12863 | -0.05786 | -0.20313 | 0.098089 | -0.02381 | -0.09189 | 0.01244 | -0.041 | 0.17641 | -0.1344 | -0.10239 | -0.0666 | -0.25631 | -0.14831 | 0.13967 | 0.099179 | 0.21461 | 0.22955 | -0.13855 | 0.186922 | -0.00905 | 0.05846 | 0.113529 | -0.26717 |
| KMS34 | -0.10032 | 0.139418 | -0.37949 | 0.09201 | -0.18243 | -0.07721 | 0.010388 | -0.16498 | -0.02267 | 0.13765 | -0.13216 | 0.02866 | 0.00029 | -0.0703 | -0.19615 | 0.084213 | -0.03497 | 0.16954 | 0.229834 | 0.03977 | 0.24415 | -0.19872 | 0.10565 | 0.01087 | 0.0876 | 0.109027 | -0.35348 |
| HEYA8 | -0.03538 | 0.123215 | -0.11837 | 0.01515 | -0.19685 | -0.1056 | -0.11552 | -0.24325 | -0.00523 | -0.01128 | -0.03735 | 0.0377 | -0.20946 | -0.31203 | -0.04449 | -0.21306 | -0.05787 | 0.16719 | 0.029543 | 0.1802 | 0.20013 | -0.32895 | 0.093265 | -0.14375 | 0.07729 | -0.0726 | -0.47273 |
| OE21 | -0.14239 | 0.133528 | -0.30237 | -0.02433 | 0.01855 | -0.12007 | -0.01003 | -0.19315 | -0.08556 | -0.01168 | -0.12142 | -0.14109 | -0.18271 | -0.15832 | -0.1332 | -0.19263 | 0.02996 | 0.29676 | -0.08741 | 0.17211 | 0.1838 | -0.03711 | 0.102546 | 0.03331 | 0.05087 | 0.004555 | -0.32815 |
| VMCUB1 | -0.22505 | 0.055695 | -0.16658 | 0.01498 | -0.00953 | -0.10532 | 0.016423 | -0.11732 | -0.1671 | -0.09463 | -0.15745 | 0.10144 | -0.10824 | -0.23251 | -0.13993 | -0.04813 | -0.1775 | -0.02671 | -0.0888 | 0.18308 | 0.0848 | -0.34979 | 0.123351 | -0.06393 | 0.16578 | -0.09326 | -0.30666 |
| HSC4 | -0.00872 | 0.231494 | 0.00417 | 0.11073 | -0.03384 | -0.13203 | 0.120979 | -0.1904 | -0.15298 | 0.16537 | -0.19465 | 0.16162 | -0.36665 | -0.15842 | -0.1716 | 0.116422 | 0.09415 | 0.19216 | 0.041511 | 0.16594 | 0.16693 | -0.40443 | 0.024982 | -0.05382 | 0.05075 | 0.022491 | -0.10002 |
| HT1197 | -0.22292 | -0.0401 | -0.25616 | 0.04981 | -0.1534 | -0.24316 | 0.059347 | -0.07507 | -0.52397 | -0.23058 | -0.20593 | 0.0537 | -0.02473 | -0.08075 | 0.016714 | 0.020072 | 0.2146 | 0.31475 | -0.1268 | 0.4127 | -0.14829 | -0.16473 | -0.21577 | -0.09984 | 0.18464 | 0.076311 | -0.39233 |
| BHY | -0.16932 | 0.042091 | -0.41777 | -0.16524 | -0.23114 | -0.21142 | -0.09268 | -0.21028 | -0.07904 | -0.06667 | -0.09021 | 0.07864 | -0.24073 | -0.18246 | -0.042 | 0.018324 | 0.0625 | 0.02162 | -0.04392 | 0.11999 | 0.1795 | -0.28096 | 0.045231 | -0.15016 | 0.05586 | -0.09293 | -0.25821 |
| SNU1076 | -0.24277 | 0.025839 | -0.24015 | -0.02898 | -0.24182 | -0.12772 | -0.00542 | -0.24629 | -0.1916 | -0.00846 | -0.00947 | 0.05458 | -0.44875 | -0.12795 | -0.043 | -0.2063 | -0.10162 | 0.35212 | -0.25636 | 0.07445 | 0.06972 | -0.14074 | 0.016774 | -0.16827 | 0.05262 | -0.20936 | -0.36698 |
| IGR39 | -0.03853 | 0.015304 | -0.30684 | -0.04686 | -0.11757 | -0.14104 | -0.10086 | -0.22305 | -0.00811 | 0.01887 | -0.03731 | 0.13391 | -0.14956 | -0.26218 | -0.23384 | -0.15269 | -0.12558 | -0.05082 | -0.06329 | 0.12123 | 0.12033 | -0.35127 | 0.200624 | -0.09883 | 0.18093 | -0.06249 | -0.4096 |
| K562 | -0.34178 | -0.07376 | 0.02924 | -0.01627 | -0.19406 | -0.0472 | -0.00082 | -0.157 | -0.04809 | 0.00751 | -0.18842 | -0.16751 | -0.34549 | -0.17732 | -0.18248 | -0.396 | 0.03108 | 0.33654 | -0.13395 | 0.14538 | 0.32354 | -0.07365 | 0.052697 | -0.00571 | 0.31718 | -0.02175 | -0.72024 |
| HT29 | -0.11412 | -0.04689 | -0.90612 | -0.10097 | -0.23718 | -0.12779 | -0.01854 | -0.0398 | -0.14598 | -0.00946 | -0.04087 | 0.10359 | -0.09184 | -0.00039 | -0.07134 | -0.19903 | -0.06176 | 0.03187 | -0.02076 | 0.0512 | 0.0663 | -0.10131 | 0.099763 | -0.08451 | 0.16155 | -0.05693 | -0.1689 |
| UACC893 | -0.3727 | 0.082725 | -0.14156 | -0.01028 | -0.0453 | -0.00423 | -0.13622 | 0.06324 | -0.14542 | 0.08314 | -0.09668 | -0.06502 | -0.20458 | 0.00505 | -0.0601 | -0.19469 | -0.09933 | 0.13862 | -0.16362 | 0.2205 | 0.17194 | -0.14338 | 0.154749 | 0.15805 | -0.00075 | 0.019509 | -0.29671 |
| SIHA | -0.17493 | 0.058123 | -0.17542 | 0.00901 | -0.1596 | -0.20685 | -0.06093 | -0.1734 | -0.31083 | -0.04134 | -0.05441 | 0.07557 | -0.20462 | -0.11176 | -0.20281 | -0.09842 | -0.03324 | 0.12099 | 0.079611 | 0.0648 | 0.06529 | -0.29426 | 0.116642 | -0.15525 | 0.13887 | -0.02322 | -0.2666 |
| AML193 | -0.09528 | -0.04674 | -0.1899 | -0.12929 | -0.2436 | -0.33148 | -0.0306 | -0.12749 | -0.02157 | 0.05058 | -0.11847 | -0.11429 | -0.09782 | -0.152 | -0.09431 | -0.09369 | -0.28049 | 0.25682 | -0.14244 | 0.09013 | 0.13707 | -0.16274 | -0.10218 | -0.08708 | 0.05083 | -0.08072 | -0.4192 |
| A172 | -0.16176 | -0.0116 | -0.11273 | -0.0532 | -0.08883 | -0.12334 | -0.12472 | -0.16956 | -0.23959 | -0.05304 | 0.188467 | 0.0782 | -0.22259 | -0.43895 | -0.07285 | -0.13984 | -0.1151 | -0.04759 | -0.02916 | -0.10884 | 0.08987 | -0.40261 | 0.11596 | 0.04237 | 0.25727 | -0.04301 | -0.68731 |
| NCIH1836 | -0.04506 | 0.096267 | -0.21417 | -0.32157 | -0.07194 | -0.2234 | -0.16724 | -0.30952 | 0.107618 | 0.23962 | -0.1079 | 0.06663 | -0.40567 | -0.17037 | -0.17256 | 0.093921 | -0.1569 | -0.07587 | -0.05014 | 0.07014 | 0.23583 | -0.11915 | -0.07898 | 0.11112 | 0.20382 | -0.06929 | -0.59173 |
| TDOTT | -0.22493 | 0.11726 | -0.53498 | -0.0704 | -0.14393 | -0.34314 | 0.122604 | -0.07822 | -0.15596 | -0.03372 | -0.15284 | 0.17326 | -0.09927 | -0.02634 | 0.144796 | -0.12598 | -0.04921 | -0.02691 | -0.35293 | 0.11281 | 0.08385 | -0.27684 | -0.00937 | -0.08026 | 0.12814 | -0.00579 | -0.17992 |
| HCC78 | -0.16361 | -0.31975 | -0.21523 | -0.12318 | -0.10704 | -0.20377 | 0.025077 | -0.33107 | -0.05492 | -0.02935 | -0.07143 | 0.19708 | -0.14578 | -0.51312 | -0.17563 | 0.137291 | -0.07875 | 0.04042 | -0.0855 | 0.22304 | 0.15784 | -0.14345 | 0.127357 | 0.06621 | 0.29768 | -0.24028 | -0.60953 |
| EBC1 | -0.16545 | -0.02337 | 0.03881 | -0.0092 | -0.30941 | -0.32041 | -0.074 | -0.17437 | -0.62892 | 0.02366 | -1.38507 | 0.13977 | -0.32796 | -0.2149 | -0.20885 | -0.17477 | 0.01864 | 0.25365 | -0.01988 | 0.12763 | 0.01762 | -0.30112 | 0.10022 | -0.17002 | 0.18248 | 0.108254 | -0.35257 |
| RCM1 | -0.24859 | 0.031782 | -0.6309 | -0.21922 | -0.18574 | -0.09138 | -0.18009 | -0.29499 | -0.13401 | 0.01292 | 0.057499 | 0.08454 | -0.14807 | -0.07921 | -0.16798 | -0.00377 | -0.14848 | 0.41182 | -0.16702 | 0.19246 | 0.03159 | -0.09679 | 0.13992 | 0.00946 | 0.0738 | -0.14138 | -0.39576 |
| ST486 | -0.08373 | 0.00316 | -0.16177 | 0.09785 | -0.15994 | -0.09429 | -0.05345 | -0.34406 | -0.09614 | -0.17641 | -0.09981 | 0.16901 | -0.2225 | -0.26067 | -0.10746 | -0.01149 | 0.02056 | 0.11681 | -0.02421 | 0.21157 | 0.20792 | -0.02915 | 0.069759 | -0.02397 | 0.08911 | 0.104129 | -0.42567 |
| YKG1 | -0.25036 | 0.016017 | -0.04674 | 0.00118 | -0.05419 | -0.23159 | -0.09641 | -0.21604 | 0.04357 | -0.21171 | -0.92326 | 0.06494 | -0.20117 | -0.23384 | -0.08731 | -0.08317 | -0.09902 | 0.10265 | 0.099813 | 0.22097 | 0.14905 | 0.021336 | 0.07215 | -0.06865 | 0.37892 | -0.00472 | -0.2825 |
| T98G | -0.0024 | -0.01899 | -0.23664 | -0.1044 | -0.23243 | -0.24861 | 0.061687 | -0.08544 | -0.03837 | -0.02227 | -0.06016 | -0.03839 | -0.16999 | -1.1911 | -0.07478 | -0.06398 | -0.04502 | -0.09701 | -0.02281 | 0.14211 | 0.10915 | -0.2506 | -0.07594 | -0.02554 | 0.09496 | -0.01414 | -0.24329 |
| MDAMB436 | -0.05957 | -0.05325 | -0.24884 | 0.01356 | -0.11244 | -0.10027 | -0.03002 | -0.32703 | 0.028945 | 0.02706 | -0.14565 | 0.06224 | -0.18856 | -0.30383 | -0.0073 | -0.18046 | -0.02902 | 0.00936 | -0.10516 | 0.10944 | 0.11728 | -0.37689 | 0.04351 | -0.10485 | 0.08765 | -0.15825 | -0.54727 |
| KMS27 | -0.11095 | 0.060404 | -0.57896 | -0.15926 | -0.30728 | -0.0747 | -0.08377 | -0.25295 | -0.03139 | 0.14898 | -0.1055 | 0.03232 | -0.07209 | -0.03917 | -0.04906 | -0.01033 | -0.09657 | 0.28136 | 0.060956 | 0.0734 | 0.12482 | -0.14719 | 0.144407 | -0.08532 | 0.18194 | 0.233571 | -0.25809 |
| JHH2 | -0.16833 | -0.00046 | -0.06424 | 0.10558 | -0.11483 | -0.16566 | -0.12265 | 0.03641 | -0.00579 | -0.06365 | -0.21017 | 0.1334 | -0.17859 | -0.03109 | -0.23423 | -0.30706 | -0.20399 | 0.23998 | -0.01611 | 0.25543 | -0.01226 | -0.42647 | 0.027046 | -0.20961 | 0.22102 | 0.072986 | -0.04022 |
| UACC257 | 0.18153 | 0.088189 | -0.48104 | -0.22951 | -0.18612 | 0.01026 | -0.08896 | -0.09144 | -0.32733 | 0.03499 | 0.072717 | -0.21096 | -0.21819 | -0.1118 | -0.15482 | -0.02305 | -0.01629 | -0.1616 | -0.25148 | 0.1261 | 0.07549 | -0.002 | 0.075681 | -0.02258 | 0.02431 | 0.064426 | -0.59191 |
| C32 | -0.25945 | 0.17231 | -0.35779 | -0.06961 | 0.00535 | -0.2746 | -0.15632 | -0.33652 | -0.32935 | 0.01749 | -0.01816 | 0.14603 | -0.2048 | 0.01841 | -0.08308 | 0.176744 | -0.10906 | -0.03165 | -0.04578 | 0.08184 | 0.00286 | 0.029445 | 0.177103 | 0.02605 | 0.02903 | -0.20764 | -0.3749 |
| SNU16 | -0.09736 | 0.126807 | -0.86121 | -0.12061 | 0.04824 | -0.26369 | 0.121595 | -0.09463 | -0.06347 | 0.00919 | -0.01475 | 0.28241 | -0.15628 | -0.11253 | -0.05014 | -0.19815 | 0.10263 | 0.21921 | 0.004594 | 0.031 | 0.27316 | -0.21551 | -0.04914 | -0.0224 | 0.16738 | 0.139436 | -0.3087 |

| Cell line | FLT3 | FLT4 | IGF1R | INSR | INSRR | KDR | KIT | LMR1 | LMR2 | LTK | MET | MUSK | PDGFRA | PDGFRB | PTK7 | RET | RON | ROR1 | ROR2 | ROS1 | STYK1 | TIE1 | TIE2 | TRKA | TRKB | TRKC | TYRO3 |
|---|---|---|---|---|---|---|---|---|---|---|---|---|---|---|---|---|---|---|---|---|---|---|---|---|---|---|---|
| JHOS4 | 0.00198 | -0.01831 | -0.23033 | 0.02672 | 0.00534 | -0.26649 | -0.15426 | -0.23599 | -0.14544 | -0.07171 | -0.06403 | 0.13914 | -0.23795 | -0.30241 | -0.02183 | -0.24575 | -0.04116 | 0.12149 | -0.19522 | 0.04243 | 0.12771 | -0.0058 | 0.160152 | -0.08608 | 0.16107 | -0.09328 | -0.2883 |
| EPLC272H | -0.14518 | -0.08927 | -0.54768 | -0.13995 | -0.08213 | -0.24901 | -0.01418 | -0.28739 | -0.48752 | -0.14496 | -0.12886 | 0.14695 | -0.27241 | -0.13609 | -0.12131 | -0.10291 | -0.04795 | 0.11218 | -0.09456 | 0.1626 | 0.1009 | -0.50385 | 0.07382 | -0.06039 | 0.1493 | -0.02717 | -0.49042 |
| NCIH1975 | -0.0365 | -0.05347 | -0.19056 | -0.11888 | -0.18473 | -0.2066 | -0.05502 | -0.12204 | -0.04311 | 0.02086 | -0.05216 | 0.06075 | -0.20292 | -0.133 | -0.13986 | -0.08673 | -0.061 | 0.05848 | -0.11668 | 0.06405 | 0.12317 | -0.05678 | 0.035921 | -0.07023 | 0.11173 | 0.010246 | -0.30812 |
| KMS26 | 0.05288 | 0.120877 | -0.53754 | -0.27441 | -0.1866 | -0.23673 | 0.186324 | -0.05876 | 0.023446 | -0.12949 | -0.06295 | 0.12037 | -0.11975 | -0.29592 | -0.12457 | -0.21668 | 0.04806 | 0.08882 | -0.11285 | 0.02298 | 0.22965 | -0.00289 | 0.109155 | -0.05747 | 0.27207 | -0.07321 | -0.2769 |
| NCIH1437 | -0.07309 | -0.0135 | -0.86514 | -0.08576 | -0.0728 | -0.26072 | -0.08465 | -0.2172 | -0.05055 | -0.02826 | -0.10732 | 0.07549 | -0.29612 | -0.00123 | -0.15436 | -0.05971 | -0.04595 | 0.04722 | 0.024803 | 0.06748 | 0.3336 | -0.20181 | 0.011547 | -0.02966 | 0.03061 | 0.01878 | -0.33197 |
| LN235 | -0.07485 | -0.01119 | -0.12576 | 0.17733 | -0.04074 | -0.50221 | -0.19091 | -0.34643 | -0.32668 | -0.12802 | -0.12643 | 0.06874 | -0.57687 | -0.35367 | -0.04763 | 0.033141 | -0.24728 | 0.11987 | -0.04696 | 0.20996 | 0.1894 | 0.152283 | -0.00796 | -0.05023 | 0.09277 | -0.06665 | -0.36201 |
| TM31 | -0.05701 | 0.093954 | -0.58572 | -0.07403 | -0.18218 | -0.29168 | -0.00935 | 0.00596 | -0.18958 | 0.04385 | -0.10197 | 0.05916 | -0.18918 | 0.08578 | -0.08533 | -0.14963 | -0.11983 | 0.03165 | -0.01671 | 0.07872 | 0.13695 | -0.0778 | 0.038707 | -0.24502 | 0.12248 | 0.059927 | -0.30516 |
| BC3C | -0.16683 | 0.03427 | -0.38879 | 0.0085 | -0.14124 | -0.10214 | -0.07861 | -0.35657 | -0.10196 | -0.12339 | -0.09453 | 0.09112 | -0.38632 | -0.16997 | -0.16165 | -0.19075 | 0.13485 | 0.08732 | 0.023662 | -0.02972 | 0.09015 | -0.14709 | -0.01345 | -0.09122 | 0.09139 | 0.079542 | -0.34425 |
| LN229 | -0.16293 | -0.07303 | -0.28789 | -0.08692 | -0.12236 | -0.01775 | -0.07557 | -0.36168 | 0.016715 | -0.0424 | -0.15151 | 0.09156 | -0.30435 | -0.18755 | -0.06701 | -0.07839 | -0.13647 | 0.02871 | -0.12935 | 0.17932 | 0.11326 | -0.15787 | 0.068029 | -0.06532 | 0.1721 | -0.015 | -0.29692 |
| LCLC97TM1 | -0.13946 | 0.077864 | -0.63727 | -0.02656 | 0.03514 | -0.13415 | 0.056452 | -0.15811 | -0.12059 | 0.02994 | -0.08945 | 0.04681 | -0.18927 | -0.05836 | -0.16124 | 0.094622 | -0.11366 | 0.15744 | -0.01101 | 0.07197 | 0.20504 | -0.08642 | -0.01468 | -0.03207 | 0.17247 | 0.201894 | -0.2317 |
| TTC709 | -0.15454 | 0.00911 | -0.20553 | -0.21621 | 0.00192 | -0.10725 | 0.126601 | -0.30732 | -0.07839 | -0.03458 | -8.4E-06 | 0.37979 | -0.37027 | -0.1516 | -0.07623 | -0.00486 | 0.05341 | 0.10718 | 0.069299 | 0.29501 | 0.18238 | -0.25087 | 0.030047 | 0.1299 | 0.20008 | 0.163064 | -0.24219 |
| PATU8902 | -0.08906 | -0.06127 | -1.06142 | -0.17668 | -0.30801 | -0.13743 | -0.14387 | -0.32249 | -0.19938 | -0.15736 | -0.06407 | 0.2027 | -0.18662 | -0.12689 | -0.29658 | 0.011743 | 0.09221 | 0.22487 | 0.087029 | 0.31252 | 0.137 | -0.29751 | 0.130842 | -0.15913 | 0.12587 | -0.13732 | -0.51652 |
| SLR26 | -0.05316 | 0.029927 | -0.0215 | -0.02802 | -0.27063 | -0.27242 | -0.08044 | -0.08961 | -0.18985 | -0.01474 | -0.71829 | 0.02122 | -0.31996 | -0.05127 | -0.13472 | 0.010531 | -0.07625 | 0.20983 | -0.16144 | 0.0461 | 0.10055 | -0.16106 | 0.083638 | -0.2081 | -0.00181 | -0.11086 | -0.26468 |
| MIAPACA2 | -0.14787 | -0.00987 | -0.51591 | -0.05195 | -0.08286 | -0.14877 | -0.02097 | -0.2516 | -0.00875 | -0.01361 | -0.08643 | 0.04927 | -0.14933 | 0.02359 | -0.12462 | 0.04595 | -0.01167 | 0.07514 | 0.007297 | 0.06411 | 0.08698 | -0.11866 | 0.084047 | -0.04747 | 0.11764 | -0.03461 | -0.33778 |
| M07E | -0.04384 | -0.03921 | -0.15333 | -0.08794 | -0.10466 | -0.07954 | 0.034164 | -0.21323 | 0.125961 | -0.0467 | -0.12116 | 0.21459 | -0.23067 | -0.06259 | -0.13528 | -0.06509 | -0.01024 | -0.0067 | 0.039866 | 0.15718 | 0.20223 | -0.28431 | 0.113399 | -0.01899 | 0.16299 | 0.112581 | -0.34946 |
| KYO1 | -0.20128 | 0.077505 | -0.03576 | 0.31771 | -0.0599 | -0.09919 | -0.08443 | -0.0731 | -0.20663 | -0.0176 | -0.16185 | 0.05471 | -0.13642 | -0.50245 | 0.030833 | -0.1351 | 0.12589 | -0.44597 | 0.256247 | 0.28767 | 0.16712 | -0.4686 | 0.135853 | 0.04982 | -0.1074 | -0.10904 | -0.54837 |
| TE6 | -0.18604 | -0.03407 | -0.23997 | -0.02222 | -0.01527 | -0.1552 | -0.00255 | -0.30427 | -0.10214 | 0.13907 | 0.067864 | 0.10791 | -0.10432 | 0.08296 | -0.14082 | -0.14626 | -0.2997 | 0.21229 | -0.0263 | 0.15494 | -0.0184 | -0.11543 | -0.14129 | 0.08015 | 0.21596 | 0.166588 | -0.3049 |
| PECAPJ34CLONEC12 | -0.16917 | 0.16098 | -0.15703 | 0.10854 | -0.12215 | -0.15761 | 0.007127 | -0.19044 | -0.1297 | -0.08321 | -0.0729 | 0.12851 | -0.14572 | -0.18736 | -0.17859 | -0.16288 | -0.03441 | 0.21313 | -0.11458 | 0.08737 | 0.17361 | -0.25366 | 0.070937 | -0.05825 | 0.11435 | -0.13543 | -0.42388 |
| KYM1 | 0.12766 | -0.21517 | -0.29153 | -0.09686 | -0.31735 | -0.38446 | -0.0307 | -0.26907 | -0.03447 | 0.05246 | -0.62625 | 0.42202 | -0.21542 | -0.20501 | 0.150986 | -0.05159 | -0.06297 | -0.07215 | -0.07487 | 0.06587 | 0.06545 | -0.19862 | 0.053992 | -0.00901 | 0.16909 | 0.216301 | -0.2291 |
| COV644 | 0.02043 | 0.053642 | -0.28434 | -0.079 | -0.20963 | -0.25455 | -0.01071 | -0.17927 | -0.01418 | -0.07593 | -0.05292 | 0.104 | -0.20136 | -0.29943 | -0.14427 | -0.10198 | -0.0304 | 0.12651 | -0.14108 | 0.04914 | 0.21951 | -0.21323 | 0.142483 | -0.13114 | 0.32737 | -0.17612 | -0.49979 |
| SF126 | -0.16567 | 0.076671 | -0.05377 | -0.08447 | -0.13082 | -0.17967 | 0.068582 | -0.151 | -0.06154 | -0.1266 | -0.11049 | 0.10693 | -0.24844 | -0.10193 | -0.01937 | -0.0819 | -0.12699 | 0.08901 | 0.01731 | 0.01644 | 0.17947 | -0.16053 | 0.047949 | -0.01501 | 0.16111 | 0.04821 | -0.15085 |
| RVH421 | -0.03109 | -0.09048 | -0.08351 | -0.05343 | -0.2113 | -0.063 | -0.03135 | -0.2249 | -0.2922 | 0.05003 | -0.01527 | 0.08139 | -0.01975 | -0.31045 | 0.045568 | -0.0374 | -0.18352 | 0.01189 | 0.05383 | 0.14641 | 0.11817 | -0.3655 | 0.009681 | -0.05297 | 0.20624 | -0.06691 | -0.33232 |
| HS746T | -0.08456 | 0.001413 | -0.2462 | -0.14164 | -0.08757 | -0.3175 | -0.03809 | -0.22412 | -0.20476 | -0.06705 | -0.31146 | 0.18761 | -0.28167 | -0.00903 | -0.23474 | -0.04424 | -0.14076 | 0.17191 | -0.07496 | 0.19352 | 0.08927 | -0.11012 | 0.097298 | -0.07708 | 0.05901 | -0.05875 | -0.38838 |
| SNU1041 | -0.1819 | 0.014313 | -0.11711 | 0.01583 | -0.13024 | -0.22427 | 0.089745 | -0.09442 | -0.21789 | -0.11507 | -0.07776 | 0.16816 | -0.18116 | -0.0958 | -0.204 | -0.17986 | -0.22056 | 0.12327 | -0.00148 | 0.09453 | 0.08461 | -0.33925 | 0.107974 | -0.01338 | 0.14159 | 0.243634 | -0.30976 |
| PECAPJ15 | -0.17244 | 0.04324 | -0.27908 | 0.00175 | -0.15017 | -0.28148 | -0.06252 | -0.29848 | -0.16964 | 0.06569 | -0.01747 | 0.14352 | -0.12532 | -0.17676 | -0.10918 | -0.17739 | -0.10294 | 0.21606 | -0.2103 | 0.14587 | 0.16828 | -0.17486 | 0.072522 | 0.00786 | 0.09113 | 0.061519 | -0.26038 |
| JHH1 | -0.03825 | 0.041544 | -0.24341 | -0.08313 | -0.18059 | -0.21708 | -0.01495 | -0.1755 | 0.006048 | -0.13866 | -0.00803 | 0.04338 | -0.16922 | -0.07111 | -0.07585 | -0.09898 | 0.06246 | 0.10409 | 0.067639 | 0.08917 | 0.0949 | -0.06131 | 0.051827 | 0.05285 | 0.0764 | -0.10487 | -0.26986 |
| MDAMB157 | -0.04227 | -0.03106 | -0.23713 | -0.12343 | -0.2227 | -0.23768 | -0.07845 | -0.35205 | -0.19888 | -0.09739 | 0.0393 | 0.08341 | -0.24347 | -0.64041 | -0.07881 | -0.12249 | -0.10436 | 0.28767 | -0.07432 | 0.07809 | 0.11768 | -0.26427 | 0.048692 | -0.13175 | 0.22365 | -0.08052 | -0.27515 |
| KNS42 | -0.2202 | -0.0314 | -0.56189 | 0.00619 | -0.20823 | -0.28672 | -0.07577 | -0.32851 | -0.08277 | 0.10723 | -0.0192 | 0.02378 | -0.59541 | -0.17033 | -0.15335 | -0.18014 | -0.1129 | 0.1701 | 0.021924 | 0.02879 | 0.12195 | -0.29864 | -0.09648 | 0.08056 | 0.23288 | -0.01108 | -0.6434 |
| SNU201 | -0.06091 | 0.066101 | -0.13274 | -0.10064 | -0.16585 | -0.10485 | -0.19213 | -0.43112 | 0.137949 | -0.15277 | -0.21659 | 0.15161 | -0.57198 | -0.36715 | -0.06524 | -0.09327 | -0.11152 | 0.18696 | -0.10074 | 0.10035 | 0.17631 | 0.094298 | 0.179962 | -0.08566 | 0.15125 | -0.10889 | -0.4592 |
| HCC1806 | -0.06587 | 0.040412 | -0.13495 | -0.03159 | -0.0948 | -0.04799 | -0.0611 | -0.24141 | -0.18754 | -0.06873 | -0.07283 | 0.13427 | -0.28711 | -0.06366 | -0.14864 | -0.0758 | -0.07821 | -0.04326 | -0.06763 | 0.11705 | 0.03176 | -0.0888 | 0.101647 | -0.05912 | 0.11352 | 0.018026 | -0.23978 |
| LCLC103H | 0.1061 | -0.03714 | -0.23839 | -0.04405 | -0.00868 | -0.10857 | -0.12169 | -0.19842 | -0.03125 | -0.10969 | -0.13375 | 0.20454 | -0.16745 | -0.03441 | -0.05502 | -0.02281 | -0.14822 | 0.03026 | -0.03212 | 0.09362 | 0.09269 | -0.29228 | 0.17853 | -0.05657 | 0.2529 | -0.05331 | -0.32514 |
| YD8 | -0.08908 | 0.026049 | -0.32053 | -0.05344 | -0.10182 | -0.01263 | -0.09229 | -0.16559 | -0.00099 | -0.0569 | -0.15756 | 0.05413 | -0.06761 | -0.06597 | -0.16073 | 0.053384 | -0.00035 | 0.03908 | 0.07246 | 0.06139 | 0.07836 | -0.0832 | 0.191619 | -0.04459 | 0.00364 | -0.00279 | -0.33773 |
| HS944T | -0.04364 | 0.009144 | -0.47026 | 0.01842 | -0.15326 | -0.24004 | 0.053621 | -0.00204 | -0.072 | -0.0298 | -0.02798 | 0.00257 | -0.16073 | -0.03359 | -0.0771 | -0.05229 | -0.0513 | 0.05165 | 0.084559 | 0.08089 | -0.01588 | -0.21657 | 0.0088 | 0.01601 | 0.14443 | 0.030289 | -0.29853 |
| FU97 | -0.05619 | -0.03514 | -0.13626 | -0.0396 | -0.09931 | -0.15123 | -0.09675 | -0.34633 | -0.06335 | -0.08455 | -0.06062 | 0.12774 | -0.81797 | -0.0954 | -0.02367 | -0.00852 | -0.0494 | 0.13103 | 0.04179 | 0.24588 | 0.10356 | 0.011657 | 0.062957 | -0.10965 | 0.08043 | 0.066489 | -0.24579 |
| LN340 | -0.04484 | 0.052419 | -0.13352 | 0.11141 | -0.10384 | -0.17587 | -0.21682 | -0.40847 | -0.10542 | -0.28941 | -0.55448 | 0.19884 | -0.08558 | 0.06031 | -0.0693 | -0.13986 | -0.06026 | -0.13778 | 0.040117 | 0.294 | 0.09135 | 0.075086 | 0.078361 | -0.06375 | 0.1978 | -0.10692 | -0.10926 |
| KYSE520 | -0.03343 | 0.031218 | -0.07802 | -0.12841 | -0.18588 | -0.2028 | -0.2147 | -0.1232 | -0.12117 | -0.04325 | 0.01632 | 0.08122 | -0.18861 | 0.03253 | -0.07069 | -0.16179 | -0.1248 | 0.14249 | -0.05548 | 0.16898 | 0.05321 | -0.21262 | 0.091188 | -0.14915 | 0.02574 | -0.01604 | -0.1862 |
| NCIH441 | -0.03839 | 0.074413 | -0.37344 | -0.13667 | -0.20482 | -0.09742 | -0.07185 | -0.02153 | -0.1985 | -0.03016 | -0.07079 | 0.01862 | -0.30102 | 0.14566 | -0.11242 | -0.02971 | -0.00874 | 0.04815 | 0.010501 | 0.09353 | 0.09828 | -0.30856 | 0.30017 | -0.01898 | 0.1171 | 0.048818 | -0.43765 |
| NCIH211 | -0.18229 | 0.053977 | -0.23213 | -0.01641 | -0.14059 | -0.18408 | -0.03745 | -0.17848 | -0.27247 | -0.06001 | -0.13 | 0.00863 | -0.2713 | -0.02793 | -0.06424 | -0.04149 | -0.11558 | 0.05845 | -0.0877 | 0.07916 | 0.10732 | -0.18663 | 0.016972 | 0.01781 | 0.15774 | -0.03136 | -0.23843 |
| JL1 | -0.10172 | -0.00435 | -0.30121 | -0.11539 | -0.24745 | -0.06715 | -0.13874 | -0.16346 | 0.028736 | 0.25176 | -0.13217 | 0.18276 | -0.16694 | -0.12784 | -0.17709 | -0.03951 | 0.02368 | -0.04343 | 0.169297 | 0.19873 | 0.23216 | 0.021545 | 0.111399 | 0.06651 | 0.07255 | 0.189351 | -0.24209 |
| OVMANA | -0.00247 | -0.1734 | -0.26621 | 0.12017 | -0.08773 | -0.14185 | 0.124843 | -0.39013 | -0.18642 | 0.08178 | 0.081339 | -0.0298 | -0.14661 | 0.03571 | 0.11418 | -0.2397 | -0.03494 | 0.20358 | -0.09296 | 0.15368 | 0.01106 | 0.038633 | 0.062362 | 0.07234 | 0.00363 | -0.1056 | -0.22612 |
| TE1 | -0.20088 | 0.005898 | -0.41676 | -0.13413 | -0.18161 | -0.13047 | -0.02517 | -0.13701 | -0.15305 | -0.19964 | -0.09513 | 0.1303 | -0.33193 | -0.07175 | -0.04017 | -0.03809 | -0.01232 | 0.03871 | -0.0123 | 0.10338 | 0.09209 | -0.21885 | 0.07048 | -0.07645 | 0.08188 | -0.02963 | -0.35786 |
| NCIH28 | -0.15369 | -0.16534 | -0.7556 | -0.01762 | -0.33114 | -0.19503 | 0.114931 | -0.16447 | -0.10793 | -0.04608 | -0.0915 | 0.01077 | -0.18484 | 0.06471 | -0.10383 | -0.10678 | -0.18386 | 0.1524 | -0.02144 | 0.14358 | 0.11605 | -0.22987 | 0.0349 | 0.06882 | -0.01217 | 0.092965 | -0.26732 |
| 786O | 0.01757 | -0.00274 | -0.1732 | -0.00112 | -0.06573 | -0.25912 | 0.041681 | -0.17703 | -0.07986 | -0.11834 | -0.07528 | 0.12446 | -0.27899 | -0.34333 | -0.12491 | -0.12097 | -0.10895 | 0.11149 | -0.07623 | 0.1018 | 0.24487 | -0.1764 | 0.018195 | -0.06419 | 0.19413 | -0.08336 | -0.55755 |
| SW620 | 0.00827 | 0.128559 | -0.32513 | -0.01413 | -0.24152 | -0.1816 | -0.02918 | -0.16778 | -0.07606 | -0.16713 | -0.05409 | 0.13654 | -0.25678 | -0.21181 | -0.19003 | -0.20583 | -0.00179 | 0.16833 | -0.01794 | 0.05875 | 0.11126 | -0.14162 | 0.143545 | -0.10294 | 0.11097 | 0.04307 | -0.48668 |
| SUIT2 | -0.18735 | -0.14716 | -0.98488 | -0.11873 | -0.13894 | -0.08913 | 0.089075 | -0.1045 | 0.018979 | 0.04448 | -0.10028 | -0.06398 | -0.1576 | -0.11355 | -0.24124 | -0.15694 | 0.01959 | 0.10715 | 0.038346 | -0.01582 | 0.04249 | -0.26637 | -0.03477 | -0.13649 | 0.29594 | 0.014835 | -0.31023 |
| JJN3 | 0.05173 | -0.10896 | -0.56456 | -0.22268 | -0.0349 | -0.1571 | 0.030354 | -0.2347 | -0.19672 | 0.06965 | 0.081045 | 0.15983 | 0.16641 | -0.06794 | -0.1076 | -0.05061 | 0.01108 | -0.07806 | 0.065969 | 0.11846 | -0.08526 | -0.19196 | -0.00252 | -0.09972 | 0.10626 | 0.100372 | -0.42487 |
| RAJI | -0.16998 | -0.03176 | -0.22922 | 0.02377 | -0.18843 | -0.18253 | 0.082653 | -0.15204 | -0.1572 | 0.17545 | -0.20931 | 0.16204 | -0.04858 | -0.23888 | -0.029 | -0.03746 | 0.0439 | -0.00127 | -0.08818 | 0.06524 | 0.16382 | 0.083717 | 0.143933 | -0.14171 | 0.06456 | -0.06049 | -0.19136 |
| SF268 | -0.0007 | 0.087872 | -0.04499 | -0.0821 | -0.24281 | -0.08306 | 0.067264 | -0.09182 | -0.15663 | 0.06252 | -0.09985 | 0.10115 | 0.02157 | -0.1976 | -0.17755 | -0.13902 | -0.26471 | -0.0106 | 0.110247 | 0.17314 | 0.03156 | -0.13289 | 0.01921 | -0.13092 | 0.06186 | -0.08029 | -0.15408 |
| SUDHL8 | -0.04785 | 0.053242 | 0.1223 | 0.13766 | -0.40163 | -0.19312 | -0.05222 | -0.36266 | -0.34385 | -0.21519 | 0.101034 | -0.01227 | 0.09406 | -0.21162 | -0.48655 | -0.23071 | -0.2381 | -0.00116 | -0.05646 | 0.18697 | 0.03266 | -0.09218 | 0.124961 | -0.06242 | 0.01104 | 0.041725 | -0.33381 |
| A2780 | -0.19675 | 0.013357 | -0.02793 | 0.10298 | -0.10406 | -0.15035 | 0.031487 | -0.33579 | -0.38673 | -0.07342 | -0.09649 | 0.05572 | -0.15657 | -0.26111 | -0.15194 | -0.16228 | -0.0617 | 0.07911 | -0.02028 | 0.20945 | 0.12358 | -0.10304 | 0.110536 | -0.18902 | 0.12733 | -0.00028 | -0.4919 |
| KMS18 | 0.06997 | 0.110117 | -0.76814 | -0.23195 | -0.15872 | -0.18511 | 0.056088 | -0.0934 | 0.073328 | 0.08258 | -0.0212 | 0.10105 | -0.06606 | -0.06683 | -0.01717 | -0.09575 | -0.08199 | 0.1327 | -0.02238 | 0.10268 | 0.13051 | -0.18221 | 0.069194 | -0.22395 | 0.15139 | -0.02106 | -0.54374 |
| SUDHL5 | 0.03249 | 0.13806 | 0.08224 | 0.17693 | -0.01592 | -0.39218 | 0.092291 | -0.00709 | 0.001942 | 0.01703 | -0.11163 | 0.18772 | -0.10021 | -0.08852 | -0.14273 | -0.26606 | -0.02536 | 0.1976 | 0.096101 | 0.11888 | 0.13442 | -0.09633 | 0.081507 | 0.00384 | 0.12713 | 0.163527 | -0.26917 |
| WM1799 | -0.26073 | -0.04405 | -0.35047 | -0.00568 | -0.19721 | -0.04754 | -0.05296 | -0.09939 | -0.32053 | -0.06251 | -0.08096 | 0.11356 | -0.05445 | 0.0232 | -0.00793 | 0.08154 | -0.26385 | 0.10104 | -0.02455 | 0.21312 | 0.16933 | -0.08995 | 0.062429 | 0.06519 | 0.14708 | 0.106991 | -0.47967 |
| CORL23 | -0.21344 | 0.059409 | -0.61292 | 0.00984 | -0.07579 | 0.07861 | -0.21986 | -0.05878 | -0.25949 | 0.08977 | 0.011311 | 0.18906 | -0.26066 | -0.1682 | -0.12437 | -0.16207 | 0.14469 | 0.17743 | 0.052868 | 0.26489 | 0.05629 | 0.051357 | 0.008985 | -0.10058 | -0.16708 | 0.120435 | -0.35921 |
| OVTOKO | 0.0069 | -0.02362 | -0.1361 | 0.04539 | -0.12678 | -0.36957 | 0.123181 | -0.15014 | -0.09989 | -0.49827 | 0.012396 | 0.23603 | -0.17018 | -0.17262 | 0.000224 | -0.25412 | -0.04978 | 0.21587 | -0.17217 | 0.15172 | 0.41321 | -0.15365 | 0.084044 | 0.03106 | 0.18385 | -0.10184 | -0.31814 |
| SUDHL1 | -0.14 | 0.002134 | -0.09664 | -0.02445 | -0.16166 | -0.07553 | -0.05763 | -0.15421 | -0.04729 | -0.00504 | -0.13762 | 0.10161 | -0.12557 | -0.31537 | -0.12891 | -0.10675 | -0.06229 | -0.00275 | 0.036193 | 0.11824 | 0.13379 | -0.14623 | 0.095942 | -0.04325 | 0.1951 | 0.016622 | -0.35886 |
| SKMES1 | -0.18379 | 0.029719 | -0.36402 | -0.0702 | -0.15971 | -0.28757 | -0.18907 | -0.19218 | -0.12104 | -0.126 | -0.08739 | 0.22467 | -0.30783 | -0.17992 | -0.11816 | -0.14177 | -0.05223 | 0.02606 | 0.018836 | 0.11211 | 0.11361 | -0.16642 | 0.068545 | -0.09295 | 0.10686 | -0.01628 | -0.35802 |
| NCIH1355 | -0.11999 | 0.185191 | -0.42124 | -0.02968 | 0.20338 | 0.07701 | 0.108049 | -0.04266 | 0.113578 | 0.02223 | -0.10568 | 0.27783 | -0.1637 | 0.10856 | 0.046669 | -0.16944 | 0.08653 | 0.23254 | -0.03516 | 0.30328 | 0.20639 | -0.30214 | -0.16386 | 0.16012 | 0.10194 | 0.00335 | -0.07587 |
| HCC44 | -0.07452 | -0.01178 | -0.57545 | 0.027 | -0.1662 | -0.19372 | -0.12638 | -0.04628 | -0.04253 | -0.2183 | -0.01771 | 0.10385 | -0.14559 | -0.15473 | -0.15353 | -0.0003 | 0.09114 | 0.18396 | -0.17322 | 0.15277 | 0.00263 | -0.17366 | 0.095406 | -0.0283 | 0.07979 | 0.029419 | -0.51443 |
| HCC70 | 0.03163 | -0.14609 | 0.07209 | 0.18294 | -0.13121 | -0.27356 | -0.07721 | -0.39791 | -0.17078 | -0.04822 | 0.150617 | 0.06062 | -0.32656 | -0.16924 | 0.084784 | -0.13285 | -0.1425 | 0.24793 | 0.034829 | 0.12995 | -0.03605 | -0.23992 | 0.18817 | 0.12956 | 0.16555 | -0.01817 | -0.40182 |
| HUH6 | -0.05077 | 0.047054 | -1.09143 | -0.44525 | -0.09858 | -0.31463 | -0.07794 | -0.08538 | 0.180886 | 0.09441 | -0.15613 | 0.15353 | -0.19639 | 0.03956 | 0.04422 | -0.04637 | -0.03309 | -0.07712 | -0.06956 | 0.14202 | 0.07268 | -0.20645 | 0.071498 | 0.05097 | 0.19934 | -0.06522 | -0.5609 |
| LN443 | -0.15247 | 0.011631 | -0.08521 | -0.07746 | -0.07055 | -0.39783 | -0.16013 | -0.37764 | -0.10568 | 0.04148 | -0.20031 | 0.07228 | -0.9113 | -0.10532 | -0.04843 | -0.01304 | -0.13406 | 0.1605 | 0.014731 | 0.18601 | 0.09659 | -0.19887 | 0.11762 | -0.0015 | 0.13957 | -0.01017 | -0.30598 |
| LN464 | -0.15738 | -0.04861 | -0.23088 | -0.16228 | -0.09393 | -0.19743 | 0.063249 | -0.14903 | 0.096457 | -0.03654 | -0.04892 | 0.14937 | -0.30291 | -0.11773 | -0.12919 | 0.103411 | -0.05632 | -0.0344 | -0.02399 | 0.15306 | 0.1187 | -0.11856 | -0.02335 | -0.08406 | 0.03974 | 0.042953 | -0.26238 |
| SW1573 | -0.13421 | -0.12251 | -0.27008 | -0.07739 | -0.23363 | -0.16636 | -0.0399 | -0.14444 | -0.10459 | -0.04298 | -0.02287 | 0.10501 | -0.26818 | -0.19079 | -0.22248 | -0.16348 | -0.02887 | 0.07528 | -0.11113 | 0.04046 | 0.03063 | -0.23696 | 0.034375 | -0.14088 | 0.15904 | -0.09928 | -0.19776 |
| SW948 | -0.00946 | 0.241287 | -0.35184 | -0.12874 | -0.1796 | -0.44035 | 0.317645 | -0.05981 | -0.48103 | -0.35472 | 0.113933 | 0.08013 | 0.17162 | 0.00717 | 0.324562 | 0.058114 | -0.16729 | 0.00721 | 0.011563 | 0.19457 | 0.15767 | -0.20594 | -0.12443 | -0.18994 | -0.00099 | 0.137595 | -0.56021 |
| A549 | -0.11204 | -0.03631 | -0.53599 | -0.10962 | -0.05704 | -0.22833 | -0.03492 | -0.20585 | -0.09118 | 0.16586 | -0.07529 | 0.10354 | -0.28415 | 0.09187 | 0.042071 | -0.11497 | -0.02211 | -0.01185 | -0.0463 | 0.31055 | 0.14469 | -0.26295 | -0.10008 | -0.00135 | 0.17528 | -0.1298 | -0.46381 |
| SNU1066 | -0.09764 | 0.044608 | -0.13148 | -0.01298 | -0.13218 | -0.16653 | 0.056493 | -0.28736 | -0.27852 | -0.08831 | 0.00262 | 0.06408 | -0.20524 | -0.14753 | -0.17369 | -0.16477 | -0.04569 | 0.19906 | -0.01293 | 0.13776 | -0.10879 | -0.20021 | 0.071346 | -0.11316 | 0.12078 | 0.086475 | -0.39687 |
| SNU503 | -0.04834 | 0.007558 | -0.1119 | -0.03244 | -0.22352 | -0.20403 | 0.010576 | -0.18949 | -0.13905 | -0.02117 | -0.0876 | 0.1355 | -0.20185 | -0.25402 | -0.10446 | -0.12731 | -0.14975 | 0.20524 | -0.16263 | 0.13244 | 0.11036 | -0.28656 | 0.079855 | -0.13833 | 0.09456 | -0.10133 | -0.45063 |
| KMRC1 | -0.06708 | 0.1238 | -0.0538 | -0.06315 | -0.15603 | -0.50361 | -0.12437 | -0.01032 | 0.316886 | -0.07079 | -0.01534 | -0.13006 | -0.38873 | -0.19859 | -0.28313 | -0.41299 | 0.15338 | 0.08185 | -0.23842 | 0.03391 | 0.24573 | -0.38218 | 0.051908 | 0.08976 | -0.05624 | 0.00356 | -0.67585 |
| L33 | -0.21411 | 0.071043 | -0.3573 | -0.07776 | 0.01078 | -0.07192 | 0.110963 | -0.214 | -0.24887 | -0.06414 | -0.16652 | 0.07806 | -0.25156 | -0.27162 | 0.02896 | -0.16681 | -0.17977 | 0.1111 | -0.1041 | 0.2218 | 0.09156 | -0.53054 | -0.08489 | 0.0687 | 0.12252 | 0.067485 | -0.41602 |
| OV7 | 0.26013 | 0.100139 | -0.37347 | -0.11815 | -0.18929 | -0.46864 | 0.038761 | -0.20243 | -0.25766 | -0.05122 | -0.07186 | 0.07608 | -0.48677 | -0.46565 | -0.04455 | -0.21078 | -0.11495 | 0.26255 | -0.12644 | 0.11898 | 0.2516 | -0.05431 | -0.16626 | -0.06834 | 0.38587 | -0.03513 | -0.47799 |
| KYSE180 | -0.12732 | -0.00603 | -0.04956 | -0.02647 | -0.20526 | -0.16161 | 0.059366 | -0.28882 | -0.23573 | -0.0628 | -0.01056 | 0.04201 | -0.28615 | -0.06576 | -0.14586 | -0.05343 | -0.01972 | 0.13367 | -0.04572 | 0.10851 | 0.0329 | -0.27309 | 0.04119 | -0.10952 | 0.09526 | -0.03573 | -0.42692 |
| TE9 | -0.13529 | -0.01708 | -0.12462 | -0.11467 | -0.13091 | -0.18978 | -0.06543 | -0.27096 | -0.11166 | 0.03549 | -0.09062 | 0.08761 | -0.23269 | -0.15264 | -0.04565 | -0.09536 | -0.01997 | -0.09671 | -0.1225 | 0.02088 | 0.06561 | -0.29685 | 0.102421 | -0.0314 | 0.06142 | -0.01295 | -0.54628 |
| CORL47 | -0.20647 | -0.0898 | -0.48208 | 0.03963 | -0.26701 | -0.22243 | 0.082869 | -0.10008 | -0.30665 | 0.13372 | -0.08183 | 0.06955 | -0.1299 | -0.0804 | -0.07504 | 0.05552 | -0.07377 | 0.30409 | -0.38471 | -0.00578 | 0.35207 | 0.002321 | -0.12813 | 0.04672 | 0.29729 | -0.14664 | -0.17144 |

| Cell line | FLT3 | FLT4 | IGF1R | INSR | INSRR | KDR | KIT | LMR1 | LMR2 | LTK | MET | MUSK | PDGFRA | PDGFRB | PTK7 | RET | RON | ROR1 | ROR2 | ROS1 | STYK1 | TIE1 | TIE2 | TRKA | TRKB | TRKC | TYRO3 |
|---|---|---|---|---|---|---|---|---|---|---|---|---|---|---|---|---|---|---|---|---|---|---|---|---|---|---|---|
| OVCAR8 | -0.07229 | 0.001663 | -0.22033 | -0.1082 | -0.15285 | -0.21683 | -0.03134 | -0.18164 | -0.09921 | -0.0391 | -0.14958 | 0.0748 | -0.19933 | -0.11559 | -0.13343 | -0.12588 | -0.07915 | 0.09618 | -0.09254 | 0.13479 | 0.16834 | -0.191 | 0.126745 | -0.04944 | 0.08184 | 0.017628 | -0.25279 |
| A3KAW | -0.07011 | 0.024347 | -0.51886 | 0.10413 | 0.06445 | -0.14827 | -0.20185 | -0.01787 | 0.012622 | 0.0157 | -0.04251 | 0.17097 | -0.38595 | 0.23583 | -0.02101 | -0.08711 | -0.05266 | 0.56983 | -0.02414 | 0.00383 | 0.08626 | -0.29726 | 0.08147 | -0.05433 | 0.04493 | 0.020244 | -0.33073 |
| DMS53 | -0.09894 | 0.006216 | -0.2198 | -0.15709 | -0.062 | -0.16897 | -0.07714 | -0.27762 | -0.09026 | 0.05589 | -0.01716 | 0.15343 | -0.10326 | -0.1689 | -0.1388 | 0.01988 | 0.0134 | 0.3765 | -0.02812 | 0.1888 | 0.10335 | -0.26786 | 0.122062 | -0.18077 | -0.08662 | -0.02564 | -0.37292 |
| HCC1395 | -0.05973 | -0.02842 | -0.18878 | -0.05923 | -0.11428 | -0.22208 | 0.022263 | -0.11125 | 0.078437 | 0.01152 | -0.03815 | 0.18016 | -0.24478 | -0.19881 | -0.12988 | -0.0128 | -0.11486 | 0.1389 | 0.004245 | 0.05346 | 0.07897 | -0.08026 | 0.066516 | -0.09526 | 0.01943 | -0.06708 | -0.46376 |
| NCIH2882 | 0.23189 | -0.10286 | -0.4182 | -0.05682 | -0.12416 | -0.1373 | -0.03195 | -0.11909 | -0.46072 | 0.10009 | 0.040819 | 0.03746 | 0.26062 | -0.00778 | -0.2057 | -0.07181 | -0.1234 | 0.24951 | 0.003038 | -0.12801 | 0.19776 | -0.32364 | 0.008333 | 0.05877 | 0.27806 | 0.307076 | -0.27167 |
| RMUGS | 0.01109 | 0.082871 | -0.5178 | 0.00256 | -0.15653 | -0.37148 | -0.00067 | -0.22705 | -0.17025 | -0.19044 | -0.0318 | 0.15501 | -0.03402 | -0.27127 | 0.064918 | -0.17154 | -0.13164 | 0.11241 | -0.14904 | 0.05022 | 0.20065 | -0.36311 | 0.196704 | 0.04181 | 0.36868 | -0.06545 | -0.44879 |
| L1236 | -0.12808 | -0.00128 | -0.2558 | -0.16018 | -0.08981 | -0.01247 | -0.2395 | -0.23512 | -0.19454 | -0.00779 | -0.018 | 0.07861 | -0.2838 | -0.08736 | 0.052007 | -0.05416 | -0.14018 | -0.04268 | -0.19872 | 0.08953 | -0.0564 | -0.1946 | -0.04016 | -0.09431 | 0.01918 | 0.031869 | -0.35111 |
| OAW42 | -0.28076 | -0.60819 | -0.25295 | -0.39958 | -0.23655 | -0.36256 | -0.1543 | -0.67202 | -0.39655 | -0.30329 | -0.1475 | 0.24049 | -0.36677 | -0.3382 | -0.12007 | 0.13492 | -0.03575 | -0.09668 | 0.151542 | 0.01175 | 0.13129 | -0.37102 | 0.129576 | -0.10432 | 0.13532 | -0.10889 | -0.20895 |
| EKVX | -0.08036 | -0.08693 | -0.56192 | -0.08415 | -0.14252 | -0.19149 | -0.03115 | -0.2327 | -0.01045 | -0.06061 | -0.17086 | 0.12741 | -0.08735 | -0.14216 | 0.007173 | -0.09454 | -0.26792 | 0.19529 | 0.039054 | 0.15591 | 0.10347 | -0.29015 | 0.094857 | -0.01818 | 0.21199 | -0.03386 | -0.52193 |
| KMRC2 | -0.33058 | 0.065094 | -0.26058 | 0.15561 | -0.03861 | 0.08126 | -0.3171 | -0.27584 | -0.10109 | 0.00992 | -0.11291 | 0.18515 | -0.12848 | -0.34468 | -0.26061 | -0.02379 | 0.08074 | 0.07174 | 0.111078 | -0.03454 | 0.30614 | -0.0656 | 0.148046 | -0.2252 | 0.13334 | 0.015727 | -0.17715 |
| JIMT1 | -0.10172 | 0.077348 | -0.04136 | -0.03233 | -0.15 | -0.12424 | -0.04166 | -0.2053 | -0.10578 | -0.02414 | -0.12388 | 0.12717 | -0.23167 | -0.08562 | -0.11297 | -0.05818 | -0.01831 | 0.05631 | -0.10522 | 0.10678 | 0.06471 | -0.23915 | 0.115961 | -0.04026 | 0.08205 | -0.02813 | -0.42045 |
| CAOV3 | -0.22196 | 0.150963 | -0.02942 | 0.06743 | -0.14896 | -0.42109 | -0.21905 | -0.29004 | -0.02899 | -0.19402 | 0.1016 | -0.00618 | -0.4038 | -0.15901 | -0.23513 | -0.11252 | 0.04664 | 0.242 | -0.09278 | 0.10952 | 0.18173 | -0.47495 | 0.189545 | -0.08696 | 0.04474 | -0.09768 | -0.57824 |
| KMS11 | -0.13284 | 0.006479 | -1.00048 | -0.28497 | -0.37453 | -0.02457 | -0.07489 | -0.18711 | -0.05369 | -0.03622 | -0.09339 | 0.16504 | -0.10068 | -0.25187 | -0.11761 | -0.15124 | -0.09479 | -0.0501 | -0.05515 | 0.14819 | 0.17848 | -0.06075 | 0.021297 | -0.3877 | 0.14077 | -0.01124 | -0.32954 |
| TT2609C02 | -0.13859 | -0.01661 | -0.24073 | -0.18662 | -0.13503 | -0.09268 | -0.12643 | -0.3187 | 0.050155 | 0.00108 | -0.09269 | 0.10725 | -0.06216 | -0.23643 | 0.010066 | -0.00535 | -0.11322 | -0.04805 | -0.09552 | 0.21632 | 0.12204 | -0.26077 | 0.140534 | -0.05691 | 0.06455 | 0.043558 | -0.32244 |
| COLO680N | -0.04674 | -0.11363 | -0.29983 | 0.03359 | -0.2754 | -0.38277 | 0.02611 | 0.08774 | -0.22161 | -0.20987 | -0.05876 | 0.35336 | -0.32075 | -0.15259 | -0.22022 | -0.39436 | -0.03865 | 0.1107 | -0.06702 | 0.03791 | 0.18597 | -0.1201 | 0.154119 | -0.11018 | 0.0672 | -0.01952 | -0.42391 |
| NCIH2291 | -0.04394 | -0.09483 | 0.02545 | -0.14296 | -0.12565 | 0.00887 | 0.080707 | -0.27351 | -0.20226 | 0.02467 | -0.18068 | -0.05945 | -0.32404 | 0.07865 | -0.1669 | -0.09844 | 0.00847 | -0.01665 | -0.02399 | 0.07741 | 0.04349 | -0.31903 | 0.157225 | -0.01452 | 0.26545 | 0.126856 | -0.47197 |
| RMGI | -0.11086 | -0.07412 | -0.15264 | -0.07665 | 0.0303 | -0.06571 | 0.033071 | -0.23231 | -0.08949 | -0.08903 | -0.16551 | 0.09734 | -0.1333 | -0.1439 | 0.008587 | -0.03132 | -0.00713 | 0.02671 | 0.033404 | 0.18962 | 0.23334 | -0.20028 | 0.00725 | -0.04096 | 0.1679 | -0.05126 | -0.41001 |
| TCCSUP | -0.09894 | -0.0047 | -0.24018 | 0.01032 | -0.13151 | -0.18507 | 0.004654 | -0.14559 | -0.2009 | 0.00945 | -0.07543 | 0.03171 | -0.11595 | -0.06996 | -0.11639 | -0.1738 | -0.12503 | 0.10601 | 0.091395 | 0.19457 | 0.03235 | -0.18706 | 0.026495 | -0.11998 | 0.04465 | 0.016837 | -0.17105 |
| HMC18 | -0.13162 | -0.04865 | -0.28873 | -0.0631 | -0.26703 | -0.21563 | 0.018576 | -0.08446 | -0.00497 | 0.09967 | -0.16718 | -0.0605 | -1.04944 | -0.02486 | -0.31493 | -0.14118 | -0.13816 | 0.03806 | -0.05716 | 0.09949 | 0.10596 | -0.08989 | 0.126593 | -0.0911 | 0.11076 | -0.01797 | -0.37481 |
| SNUC1 | -0.06006 | 0.25077 | -1.42545 | -0.0188 | -0.10468 | 0.03485 | 0.011423 | -0.03263 | -0.11936 | -0.29971 | -0.0148 | 0.22766 | -0.17843 | -0.03873 | -0.21655 | 0.046006 | 0.12907 | 0.20642 | 0.0742 | 0.17206 | 0.09993 | -0.45477 | 0.082262 | -0.02094 | 0.24915 | -0.07707 | -0.00284 |
| HT1376 | -0.00946 | 0.154538 | -0.70203 | -0.21327 | -0.20114 | -0.13978 | 0.098454 | -0.11117 | -0.12827 | -0.18445 | -0.09036 | 0.17042 | -0.37656 | -0.11561 | -0.03658 | -0.23563 | 0.03757 | 0.05531 | -0.10687 | 0.12701 | 0.12371 | -0.49473 | 0.158629 | -0.15959 | 0.26651 | 0.046852 | -0.17288 |
| HCC202 | -0.19254 | -0.00367 | -0.06951 | -0.08576 | -0.06428 | -0.33292 | -0.14057 | -0.29836 | -0.32152 | -0.07478 | -0.09167 | 0.01134 | -0.73582 | 0.0301 | -0.01766 | -0.14466 | 0.01454 | 0.01474 | -0.12924 | -0.08261 | 0.0488 | -0.20907 | 0.068247 | 0.00687 | -0.05823 | 0.04956 | -0.52121 |
| PECAPJ41CLONED2 | -0.16824 | -0.01751 | -0.14522 | -0.12457 | -0.19462 | -0.50642 | -0.12782 | 0.00083 | -0.27538 | -0.24262 | -0.05845 | 0.0702 | -0.23997 | 0.0896 | -0.05547 | -0.1798 | 0.04879 | 0.2013 | -0.09904 | 0.03559 | 0.02689 | -0.22616 | 0.141696 | -0.11973 | 0.18372 | 0.18219 | -0.27395 |
| JHH5 | -0.10492 | -0.03458 | -0.64867 | -0.14615 | -0.09714 | -0.20353 | -0.06514 | -0.0869 | -0.14318 | -0.16426 | -0.431 | 0.10047 | -0.19964 | -0.22898 | -0.21786 | -0.0985 | -0.0894 | 0.25167 | -0.11761 | 0.17108 | 0.07582 | -0.16804 | 0.150394 | -0.00708 | 0.06818 | -0.03284 | -0.25026 |
| PECAPJ49 | -0.12832 | 0.068903 | -0.75209 | -0.06675 | -0.13857 | -0.14566 | 0.001891 | -0.23116 | -0.2564 | -0.10449 | -0.16398 | 0.08852 | -0.24044 | -0.02532 | -0.10129 | 0.006773 | -0.03499 | 0.16507 | 0.047652 | 0.02992 | 0.08841 | -0.29929 | 0.097192 | -0.02434 | 0.03589 | 0.010577 | -0.43269 |
| SNU601 | -0.22784 | -0.02001 | -0.35698 | -0.11963 | -0.05114 | -0.06801 | -0.01652 | -0.00116 | -0.26478 | -0.19888 | -0.13782 | 0.14146 | -0.03103 | -0.02842 | -0.09713 | -0.16889 | 0.09274 | -0.02085 | 0.003073 | -0.003 | 0.14935 | -0.08941 | 0.208565 | -0.04541 | 0.19451 | -0.01093 | -0.2719 |
| GB1 | -0.11972 | 0.015575 | -0.4566 | 0.01505 | -0.10205 | -0.25187 | 0.117716 | -0.10613 | -0.1084 | 0.10812 | -0.1264 | 0.09466 | -0.2208 | -0.17525 | -0.12229 | -0.0473 | -0.15427 | 0.22262 | 0.005382 | 0.09166 | 0.13947 | -0.08462 | 0.236014 | -0.0935 | 0.17958 | 0.094512 | -0.31946 |
| HEPG2 | -0.1536 | 0.027866 | -0.68107 | -0.33234 | -0.17838 | -0.22322 | -0.03377 | -0.28216 | -0.04127 | 0.11095 | -0.03184 | 0.05781 | -0.16467 | 0.02926 | 0.094457 | 0.016747 | -0.15055 | 0.21583 | -0.02654 | 0.16302 | 0.11468 | -0.16076 | 0.085613 | -0.04976 | 0.03631 | -0.00509 | -0.41021 |
| A253 | -0.03382 | 0.069848 | 0.03398 | -0.01113 | -0.20365 | -0.13486 | 0.001533 | -0.18978 | -0.23629 | 0.01606 | -0.08176 | 0.09363 | -0.14253 | -0.07447 | -0.06515 | -0.01658 | 0.0205 | 0.17261 | -0.16061 | 0.14767 | 0.02848 | -0.20418 | -0.02969 | -0.03462 | 0.06783 | 0.000701 | -0.34571 |
| UBLC1 | -0.05356 | 0.058715 | -0.20935 | -0.09066 | -0.00745 | -0.04334 | -0.12936 | -0.25521 | -0.08789 | -0.08539 | -0.09527 | 0.09629 | -0.27224 | -0.12496 | -0.00935 | 0.104385 | 0.04636 | -0.07546 | -0.06223 | 0.11412 | 0.03663 | -0.62257 | 0.151167 | -0.02226 | 0.13355 | 0.000857 | -0.58572 |
| GSS | -0.16027 | -0.09945 | -0.63365 | -0.17069 | -0.21162 | -0.27199 | -0.16328 | 0.07978 | -0.00328 | -0.00979 | 0.017499 | 0.27601 | -0.40518 | 0.02453 | -0.1513 | -0.00365 | 0.07487 | 0.02147 | -0.13018 | -0.0402 | 0.18745 | -0.01694 | 0.060002 | -0.14872 | 0.04837 | -0.04907 | -0.25458 |
| NCIH1703 | -0.11695 | -0.06859 | -0.03079 | -0.07527 | -0.16256 | -0.00375 | 0.046339 | -0.03015 | -0.00391 | -0.07298 | -0.11828 | 0.03869 | -0.74202 | 0.0657 | -0.09767 | -0.07299 | -0.05452 | 0.19774 | -0.11674 | 0.05219 | 0.03131 | -0.15624 | 0.067528 | 0.02505 | 0.15221 | -0.02661 | -0.35115 |
| SJSA1 | -0.01324 | 0.022468 | -0.54715 | -0.06349 | -0.20196 | -0.00158 | -0.00681 | -0.27692 | -0.13477 | -0.04621 | -0.08123 | 0.09751 | -0.13348 | -0.09414 | -0.1488 | -0.02317 | -0.0584 | 0.1797 | -0.08035 | 0.0362 | 0.00747 | -0.18207 | 0.048637 | -0.0746 | 0.23284 | 0.069125 | -0.15253 |
| JMSU1 | -0.1393 | 0.01456 | -0.32193 | -0.063 | -0.10536 | -0.05931 | 0.035776 | -0.17924 | -0.0157 | -0.03193 | -0.04867 | 0.06776 | -0.19869 | -0.02378 | -0.13855 | -0.08442 | 0.00481 | -0.12673 | -0.07935 | 0.1404 | 0.07241 | -0.25411 | 0.089925 | -0.01032 | 0.1354 | -0.08183 | -0.40189 |
| L428 | -0.23783 | -0.00805 | -0.13452 | -0.07597 | -0.25198 | -0.03189 | -0.20271 | -0.36608 | 0.104069 | -0.14309 | -0.04888 | 0.12791 | 0.07514 | -0.35396 | -0.0477 | 0.033485 | -0.13002 | -0.09431 | -0.00098 | 0.12653 | 0.12536 | -0.18902 | 0.2666 | -0.22495 | 0.13199 | -0.0397 | -0.2012 |
| GI1 | -0.09518 | -0.06516 | -0.14362 | -0.06159 | -0.07056 | -0.19761 | -0.02299 | -0.1434 | -0.05269 | -0.0052 | -0.57293 | 0.23107 | -0.32162 | -0.24399 | -0.11271 | -0.12712 | -0.22314 | 0.08227 | 0.061883 | 0.20128 | 0.19232 | -0.13418 | 0.080947 | -0.18311 | 0.25185 | -0.09477 | -0.14123 |
| A427 | -0.06556 | 0.026655 | 0.01405 | -0.05975 | 0.01167 | -0.18845 | 0.067873 | -0.14834 | -0.1405 | 0.12285 | -0.08558 | 0.08809 | -0.25403 | 0.18483 | 0.014942 | -0.07373 | -0.06487 | -0.0636 | -0.31931 | 0.06936 | -0.07511 | -0.14289 | -0.0699 | -0.10881 | 0.08708 | 0.023065 | -0.71994 |
| MKN74 | -0.11607 | -0.06964 | -0.21166 | -0.02457 | -0.24337 | -0.0947 | -0.02065 | -0.20871 | -0.12292 | -0.27255 | -0.02068 | 0.00931 | -0.16755 | 0.04594 | -0.22255 | -0.16135 | -0.02845 | 0.36884 | -0.10689 | 0.07016 | 0.26531 | -0.19646 | 0.199234 | -0.06844 | 0.16068 | -0.05932 | -0.25863 |
| LNZ308 | -0.08326 | 0.017933 | -0.12741 | -0.13219 | -0.18922 | -0.20769 | -0.03743 | -0.10147 | 0.030409 | -0.06372 | -0.01633 | 0.09933 | -0.3092 | -0.15261 | -0.09717 | -0.07831 | -0.05858 | -0.13867 | -0.05867 | 0.07617 | 0.07349 | -0.0263 | -0.0415 | -0.07198 | 0.10374 | 0.070454 | -0.34746 |
| YD38 | -0.09248 | 0.174869 | -0.37662 | 0.01858 | -0.22326 | -0.3513 | -0.14129 | 0.17738 | -0.08605 | -0.10917 | 0.021311 | 0.15646 | -0.25208 | -0.27414 | 0.086411 | -0.11612 | 0.26322 | 0.123 | 0.042711 | -0.00999 | 0.06778 | -0.19179 | 0.183839 | -0.07485 | -0.1134 | -0.03 | -0.47217 |
| MM1S | 0.00388 | -0.1658 | -0.90984 | -0.46937 | -0.0464 | -0.19778 | 0.108219 | -0.11393 | -0.00835 | -0.10689 | -0.03082 | 0.08373 | -0.21326 | -0.13942 | 0.057269 | -0.1597 | -0.22551 | -0.0096 | -0.06739 | 0.2015 | 0.02492 | -0.27648 | 0.095209 | 0.12228 | 0.14705 | 0.454337 | -0.6594 |
| SH10TC | -0.00275 | -0.06336 | -0.26898 | 0.04996 | -0.36613 | -0.54919 | 0.050203 | -0.31816 | -0.13188 | -0.16933 | 0.149242 | 0.1862 | -0.19429 | -0.01458 | -0.17423 | -0.10502 | -0.05901 | -0.06975 | -0.12786 | 0.20984 | 0.11308 | -0.35013 | 0.041536 | 0.16495 | 0.17804 | 0.037583 | -0.27563 |
| WM983B | -0.2545 | -0.15674 | -0.68776 | -0.12337 | -0.38505 | -0.18704 | -0.06578 | -0.0894 | 0.131496 | -0.01227 | -0.07854 | 0.17504 | -0.38155 | -0.25408 | -0.06332 | -0.22961 | -0.23713 | 0.49943 | -6.2E-05 | 0.31722 | 0.18531 | -0.15003 | 0.019444 | -0.21674 | 0.16588 | 0.159083 | -0.33065 |
| NCIH1648 | 0.0672 | 0.047076 | -0.14265 | -0.10871 | -0.06798 | -0.25067 | -0.09141 | -0.03231 | 0.176284 | -0.04127 | -0.01665 | 0.12958 | -0.24819 | -0.08742 | -0.26334 | 0.030851 | 0.04768 | 0.17056 | 0.188699 | 0.22886 | 0.06169 | 0.04722 | 0.229075 | -0.16865 | 0.06546 | 0.041445 | -0.40031 |
| NCIH526 | -0.22843 | 0.065939 | -1.03471 | -0.17073 | -0.10833 | -0.3146 | -0.10655 | -0.06995 | -0.31934 | -0.02976 | 0.073146 | 0.24247 | -0.40848 | -0.15732 | -0.1999 | 0.02618 | -0.08715 | 0.17087 | -0.03667 | 0.11367 | 0.09553 | -0.36794 | 0.074718 | -0.10172 | 0.06116 | 0.113411 | -0.31747 |
| MDAMB231 | -0.11799 | -0.00284 | -0.0471 | -0.10659 | -0.29001 | -0.21452 | -0.13621 | -0.06916 | -0.00877 | -0.10537 | -0.0392 | 0.0843 | -0.23662 | -0.14029 | -0.19478 | -0.26755 | 0.01742 | 0.13795 | -0.02798 | 0.00541 | 0.16511 | -0.18297 | 0.125798 | -0.15977 | -0.0224 | -0.05054 | -0.3902 |
| LK2 | -0.19618 | -0.02038 | -0.72007 | -0.07621 | -0.11558 | -0.06111 | -0.1495 | -0.34851 | -0.19117 | -0.03004 | -0.11765 | 0.14501 | -0.23985 | 0.00209 | -0.11602 | -0.06191 | 0.04286 | 0.07695 | -0.22903 | 0.06997 | 0.00626 | -0.28768 | 0.093804 | -0.07391 | 0.15523 | 0.04295 | -0.4324 |
| P31FUJ | -0.08325 | 0.024278 | -0.15295 | -0.00127 | -0.14282 | -0.23144 | 0.028447 | -0.18978 | 0.029715 | 0.0237 | -0.15918 | 0.13369 | -0.16266 | -0.04895 | -0.04581 | -0.09756 | 0.00195 | 0.10899 | 0.082968 | 0.04822 | 0.09255 | -0.00087 | 0.020505 | -0.07694 | 0.15139 | -0.0287 | -0.42102 |
| BICR56 | -0.06319 | -0.00484 | -0.31456 | 0.021 | -0.06773 | -0.18954 | -0.05181 | -0.18628 | -0.27755 | 0.01888 | -0.16263 | 0.16712 | -0.0995 | -0.03174 | -0.09534 | -0.08405 | -0.0928 | 0.01791 | -0.08614 | 0.06432 | 0.12467 | -0.18005 | 0.066308 | -0.07509 | 0.22944 | -0.08178 | -0.22232 |
| KIJK | -0.22319 | 0.065445 | -0.28052 | 0.07276 | 0.01363 | -0.02215 | 0.185379 | -0.20556 | -0.02734 | 0.11343 | 0.111553 | 0.03645 | -0.08792 | 0.0632 | -0.03872 | -0.26566 | -0.01226 | 0.12835 | -0.02721 | -0.10709 | 0.15193 | -0.08966 | -0.11409 | -0.01231 | 0.11345 | 0.028614 | -0.32031 |
| RERFLCAD2 | -0.0171 | 0.014778 | 0.02326 | -0.13772 | -0.10034 | -0.38714 | -0.13676 | -0.05109 | -0.17073 | -0.19764 | -0.09125 | 0.01782 | -0.3909 | -0.04867 | -0.17075 | -0.12942 | -0.04737 | -0.04693 | -0.06297 | 0.00017 | 0.20628 | -0.11268 | 0.112175 | -0.05106 | 0.00667 | 0.031553 | -0.47485 |
| NCIH727 | -0.07895 | 0.012463 | -0.42957 | -0.07359 | -0.03642 | -0.25604 | -0.1095 | -0.19647 | -0.16207 | -0.00051 | 3.46E-05 | 0.047 | -0.10486 | 0.01427 | -0.1477 | -0.01406 | -0.015 | 0.13145 | 0.046567 | 0.12847 | 0.11821 | -0.14439 | 0.293996 | 0.07039 | 0.04712 | 0.023394 | -0.44884 |
| ONS76 | -0.12702 | 0.084114 | -0.20141 | 0.03021 | -0.14474 | -0.2559 | -0.04652 | -0.18164 | -0.11921 | -0.20226 | -0.09003 | -0.00768 | -0.4093 | -0.02853 | 0.018561 | -0.03103 | -0.0601 | 0.31093 | 0.057559 | 0.11547 | 0.03445 | -0.37828 | 0.088824 | -0.12487 | 0.05591 | -0.12856 | -0.39474 |
| KYSE30 | -0.0862 | -0.14135 | -0.11175 | -0.08484 | -0.30413 | -0.11672 | -0.02969 | -0.17079 | -0.15245 | -0.12774 | -0.09998 | -0.08284 | -0.12007 | -0.06484 | -0.09405 | 0.00787 | 0.17728 | 0.17161 | -0.1045 | 0.18893 | 0.09099 | -0.44302 | 0.134022 | -0.21797 | 0.16243 | -0.11561 | -0.15247 |
| HSC3 | -0.19095 | 0.046574 | -0.16392 | 0.02675 | -0.0975 | -0.21926 | -0.00874 | -0.04894 | -0.10639 | -0.17654 | -0.09083 | 0.1839 | -0.20052 | -0.23012 | -0.07469 | -0.11775 | -0.12754 | 0.09436 | -0.1697 | 0.02517 | 0.14112 | -0.22987 | 0.054662 | -0.00023 | 0.162 | 0.066937 | -0.6702 |
| NCIH2023 | -0.24004 | 0.009935 | -0.54242 | -0.10279 | -0.07747 | -0.22072 | -0.01174 | -0.21033 | -0.02365 | -0.08542 | -0.22566 | 0.1344 | -0.22555 | -0.0967 | -0.11713 | -0.12464 | -0.01137 | -0.00299 | -0.06059 | 0.10072 | 0.04644 | -0.09931 | -0.0134 | -0.06698 | 0.10363 | 0.010076 | -0.4945 |
| SEM | -0.88257 | 0.049479 | -0.13173 | -0.2187 | -0.18856 | -0.08435 | -0.06334 | -0.26643 | -0.04201 | -0.11093 | 0.02991 | 0.15358 | -0.05268 | -0.1452 | -0.24755 | 0.02433 | -0.00845 | 0.36999 | -0.05935 | 0.04996 | 0.19833 | -0.36618 | -0.0328 | -0.13236 | 0.10658 | 0.152275 | -0.40939 |
| CAMA1 | -0.1869 | -0.05713 | 0.03111 | 0.05592 | -0.17501 | -0.09152 | 0.011715 | -0.27864 | -0.04348 | -0.0813 | 0.001264 | 0.24285 | -0.10916 | -0.18389 | 0.023855 | -0.07506 | 0.0103 | 0.21536 | -0.06476 | 0.19171 | 0.2189 | -0.08163 | 0.057776 | -0.05266 | 0.18683 | 0.026063 | -0.59285 |
| KYSE70 | -0.16921 | 0.14198 | -0.14488 | -0.07674 | -0.22142 | -0.15591 | 0.06677 | -0.11569 | -0.17302 | -0.07809 | -0.08164 | 0.16618 | -0.08513 | -0.17099 | -0.12871 | -0.05611 | -0.04978 | 0.19804 | -0.03698 | 0.14195 | 0.10785 | -0.39046 | 0.069131 | -0.2682 | 0.19155 | 0.034554 | -0.4214 |
| NCIH2126 | -0.12442 | 0.030158 | -0.5087 | -0.0865 | -0.1383 | -0.28341 | -0.13258 | -0.03258 | 0.091151 | 0.05912 | 0.025456 | 0.13247 | -0.3113 | -0.01691 | -0.02184 | 0.044783 | 0.02958 | -0.047 | 0.067717 | 0.01607 | 0.16378 | -0.54065 | 0.239564 | 0.02128 | 0.06561 | -0.00894 | -0.3582 |
| LXF289 | -0.03261 | -0.05288 | -0.34437 | -0.03679 | -0.063 | -0.06785 | 0.128398 | -0.07548 | -0.15093 | -0.14243 | -0.03332 | 0.0667 | -0.2541 | -0.20938 | -0.00909 | -0.02234 | -0.03861 | -0.00696 | 0.014482 | 0.01871 | 0.05017 | -0.271 | 0.039199 | 0.00245 | 0.17009 | -0.04964 | -0.30065 |
| A2058 | -0.09243 | -0.02802 | -0.13806 | -0.05937 | -0.12635 | -0.0738 | -0.02145 | -0.04804 | -0.15381 | 0.02559 | 0.00672 | 0.04102 | -0.2076 | -0.1734 | -0.08274 | -0.02403 | -0.1795 | 0.05109 | 0.001855 | 0.11062 | 0.05992 | -0.09858 | 0.118747 | -0.05062 | 0.08387 | 0.045851 | -0.38685 |
| SHP77 | -0.28569 | 0.033327 | -0.13098 | 0.0877 | 0.00281 | -0.33965 | -0.34828 | -0.07216 | -0.10247 | -0.17334 | -0.14101 | 0.10085 | -0.6991 | 0.0485 | -0.22825 | -0.092 | -0.43343 | 0.38817 | 0.03492 | 0.13328 | 0.08848 | -0.16841 | -0.05367 | 0.03162 | 0.26609 | -0.02858 | -0.21872 |
| RERFLCAD1 | -0.08812 | 0.06988 | -0.3599 | -0.05859 | -0.12113 | -0.22131 | -0.11314 | -0.11094 | -0.07755 | -0.20109 | -0.29163 | 0.16335 | -0.29589 | 0.11603 | -0.15364 | -0.08356 | 0.07001 | 0.09512 | -0.0369 | 0.14648 | 0.15398 | -0.28829 | 0.256521 | -0.13282 | 0.17757 | 0.001556 | -0.37268 |
| BFTC909 | -0.14 | 0.005154 | -0.1076 | 0.04158 | -0.06375 | -0.1594 | 0.083414 | -0.18111 | 0.077885 | 0.0765 | -0.08388 | 0.1144 | -0.04745 | -0.17036 | -0.02734 | 0.058765 | -0.04456 | 0.0111 | -0.06693 | 0.12689 | 0.11685 | -0.215 | 0.227118 | -0.07091 | 0.25264 | -0.06852 | -0.17534 |
| KATOIII | -0.17764 | 0.035637 | -0.11416 | 0.06384 | -0.17279 | 0.00446 | -0.02803 | -0.08769 | 0.006433 | -0.06558 | -0.12475 | 0.02733 | -0.01502 | -0.18877 | -0.10414 | -0.06018 | -0.05482 | -0.04226 | 0.024688 | 0.09976 | 0.09844 | -0.15672 | -0.01534 | -0.04149 | 0.07 | -0.00786 | -0.31946 |
| BICR22 | -0.01375 | 0.040795 | -0.17312 | -0.03735 | -0.00393 | -0.26174 | -0.04637 | -0.11008 | -0.05736 | -0.02325 | 0.005485 | 0.14129 | -0.40053 | -0.2813 | -0.00253 | -0.13002 | -0.0462 | 0.09462 | -0.08051 | 0.18556 | 0.07409 | -0.24016 | 0.016511 | -0.13183 | 0.13851 | 0.003028 | -0.27939 |
| MCAS | -0.13541 | 0.011418 | -0.1997 | -0.16149 | -0.06702 | -0.15444 | -0.01687 | -0.24984 | 0.023329 | -0.11436 | -0.06715 | 0.13966 | -0.08014 | -0.06859 | -0.15937 | 0.001732 | 0.04376 | 0.05411 | 0.027103 | 0.11732 | 0.10096 | -0.17028 | 0.137199 | -0.04913 | 0.2456 | 0.001136 | -0.36689 |
| HS695T | -0.08654 | -0.00166 | -0.12755 | -0.14119 | 0.08909 | -0.01937 | 0.087474 | -0.18806 | -0.51483 | 0.03153 | 0.10655 | 0.05829 | -0.21593 | -0.0783 | 0.063984 | -0.10706 | -0.05497 | 0.17885 | 0.124633 | 0.12147 | -0.15846 | 0.043202 | 0.014512 | 0.2001 | 0.06022 | 0.047191 | -0.23931 |
| NCIH446 | -0.21732 | -0.03807 | -0.28391 | -0.14826 | -0.19527 | -0.36372 | 0.108894 | 0.06281 | -0.29487 | -0.05138 | 0.066197 | 0.12347 | -0.32923 | 0.31038 | -0.26689 | -0.30276 | -0.04786 | 0.36491 | -0.01234 | 0.19778 | 0.10367 | -0.14654 | -0.06032 | -0.09447 | 0.01115 | -0.01343 | -0.57654 |
| HS936T | -0.23056 | 0.083973 | -0.46368 | 0.1095 | -0.08644 | -0.13755 | -0.15948 | 0.04808 | -0.24527 | 0.12446 | -0.04915 | 0.07508 | -0.11715 | -0.14916 | 0.048109 | -0.08935 | -0.2322 | -0.00851 | -0.01542 | 0.05172 | -0.00774 | 0.04791 | 0.161158 | -0.00407 | 0.10567 | 0.09073 | -0.55464 |
| BFTC905 | -0.12133 | 0.039043 | -0.61073 | -0.0404 | -0.11286 | -0.02003 | -0.0758 | -0.12563 | -0.00184 | -0.00493 | -0.1039 | 0.11596 | -0.31705 | -0.19164 | -0.0098 | -0.0867 | 0.02984 | 0.09611 | -0.18776 | 0.05944 | -0.01789 | -0.09104 | 0.080651 | -0.02331 | 0.17002 | -0.00447 | -0.33351 |
| NB1 | -0.20837 | 0.031565 | -0.24716 | 0.0109 | -0.1377 | -0.40206 | -0.03456 | -0.1684 | -0.12083 | -0.15574 | -0.06417 | 0.05865 | -0.33792 | 0.15036 | -0.04291 | -0.1165 | 0.19075 | -0.03467 | -0.14873 | 0.11676 | 0.04741 | -0.33272 | 0.066136 | 0.1231 | -0.0266 | -0.04816 | -0.45419 |
| COLO679 | -0.25876 | 0.137387 | -0.17425 | -0.02013 | -0.02321 | -0.31317 | -0.06503 | -0.0548 | -0.13875 | -0.07177 | -0.09595 | 0.04917 | -0.17277 | -0.0813 | 0.031887 | 0.072276 | -0.09721 | 0.01912 | 0.013851 | 0.15425 | 0.28212 | -0.26488 | 0.069615 | 0.09127 | 0.34221 | 0.227878 | -0.16124 |

| Cell line | FLT3 | FLT4 | IGF1R | INSR | INSRR | KDR | KIT | LMR1 | LMR2 | LTK | MET | MUSK | PDGFRA | PDGFRB | PTK7 | RET | RON | ROR1 | ROR2 | ROS1 | STYK1 | TIE1 | TIE2 | TRKA | TRKB | TRKC | TYRO3 |
|---|---|---|---|---|---|---|---|---|---|---|---|---|---|---|---|---|---|---|---|---|---|---|---|---|---|---|---|
| SNU738 | -0.01628 | 0.092742 | -0.16536 | -0.05597 | -0.13336 | 0.10684 | -0.15069 | 0.00276 | -0.1034 | 0.02321 | 0.041044 | 0.05343 | -0.25995 | -0.09488 | 0.045839 | -0.13212 | -0.08312 | 0.24545 | -0.0347 | 0.16659 | 0.18487 | -0.27495 | 0.204382 | -0.08313 | 0.12491 | -0.00161 | -0.42569 |
| KYSE410 | -0.08977 | 0.031887 | -0.05056 | -0.04393 | -0.04479 | -0.16827 | 0.131668 | -0.08624 | -0.06365 | -0.10582 | -0.02089 | 0.14184 | -0.04188 | -0.05026 | -0.08888 | 0.03536 | -0.02564 | 0.10098 | -0.03593 | 0.14498 | 0.08508 | -0.09527 | 0.09451 | 0.05096 | 0.12537 | 0.101628 | -0.30839 |
| SKMEL30 | -0.05331 | 0.163011 | -0.46968 | 0.11883 | -0.00577 | 0.03158 | -0.15721 | -0.13499 | -0.11326 | 0.38331 | 0.038806 | 0.0379 | -0.2595 | 0.06619 | -0.09188 | 0.139771 | 0.00172 | -0.00011 | 0.270766 | 0.21043 | 0.22294 | 0.062809 | 0.269801 | 0.03695 | 0.19315 | -0.13676 | -0.46785 |
| SKOV3 | 0.02153 | -0.00709 | -0.24999 | -0.07275 | -0.16411 | -0.25642 | 0.08477 | -0.21466 | -0.15179 | -0.15408 | -0.13513 | 0.16143 | -0.02255 | -0.13331 | -0.19374 | -0.06742 | 0.01049 | 0.14063 | -0.2068 | 0.07551 | 0.29039 | -0.25985 | 0.061098 | -0.10412 | 0.20645 | -0.10976 | -0.51316 |
| RPMI8226 | -0.04029 | 0.046563 | -0.53556 | -0.11031 | -0.13359 | 0.02769 | 0.063696 | 0.01195 | 0.297926 | 0.15213 | -0.0714 | 0.09453 | 0.07949 | -0.02288 | 0.004529 | -0.14391 | -0.08711 | 0.10663 | -0.00587 | 0.11444 | 0.08412 | -0.18579 | 0.086687 | -0.13675 | -0.00666 | 0.051905 | -0.23453 |
| LN18 | -0.14172 | 0.064031 | -0.12484 | -0.12111 | -0.27089 | -0.24801 | -0.01632 | -0.33725 | -0.07999 | -0.03721 | -0.37687 | 0.1659 | -0.15611 | -0.31289 | 0.050818 | -0.19943 | -0.1918 | 0.12491 | 0.032711 | 0.14445 | 0.03123 | -0.15867 | -0.04246 | -0.00765 | 0.182 | 0.024821 | -0.3264 |
| SW403 | -0.1426 | 0.159943 | -0.44735 | 0.09625 | -0.11478 | -0.21888 | 0.151303 | -0.43086 | 0.015473 | -0.14564 | -0.03317 | 0.14039 | -0.03796 | -0.15423 | -0.11133 | -0.38222 | -0.15232 | 0.13514 | -0.42396 | 0.13801 | 0.33959 | -0.33379 | 0.002873 | -0.01412 | 0.28038 | -0.34746 | -0.64207 |
| EJM | -0.0845 | 0.021141 | -0.23169 | -0.13861 | -0.33476 | -0.20218 | 0.172577 | -0.01989 | -0.22565 | -0.07921 | -0.02852 | 0.12683 | -0.4352 | -0.08028 | -0.0846 | -0.03673 | -0.26169 | -0.04951 | 0.111523 | 0.03539 | 0.17467 | -0.04144 | -0.02392 | 0.02314 | -0.11018 | 0.039073 | -0.28951 |
| SKMEL24 | -0.11712 | 0.009642 | -0.12604 | -0.03238 | -0.15183 | -0.22817 | -0.06354 | -0.1874 | 0.027682 | 0.06558 | -0.00721 | 0.12792 | 0.04669 | -0.08113 | -0.07645 | -0.04536 | -0.05041 | 0.05266 | -0.00279 | 0.17257 | 0.17079 | -0.25366 | -0.0677 | -0.07839 | 0.08067 | -0.05505 | -0.16759 |
| KYSE140 | -0.1285 | -0.02577 | -0.13031 | -0.02372 | -0.05514 | -0.18864 | -0.01353 | -0.18898 | -0.18492 | -0.0341 | -0.08657 | 0.12322 | -0.30745 | -0.12153 | 0.030845 | -0.13505 | -0.13504 | 0.03912 | -0.09398 | 0.07797 | 0.10397 | -0.25273 | 0.033871 | -0.01352 | 0.0946 | -5.7E-05 | -0.26045 |
| KYSE510 | -0.23034 | 0.089944 | -0.37197 | -0.02315 | -0.22505 | -0.13792 | 0.055771 | -0.1005 | -0.13533 | 0.10254 | -0.05962 | 0.00475 | -0.20061 | -0.06882 | -0.05861 | -0.07619 | -0.02861 | 0.16111 | 0.00612 | 0.04403 | 0.03437 | -0.06401 | 0.023694 | -0.141 | 0.15965 | -0.19941 | -0.24401 |
| WM793 | -0.02997 | 0.070694 | -0.16939 | 0.02571 | -0.18388 | -0.19204 | 0.016319 | -0.24497 | -0.07801 | -0.10818 | -0.00335 | 0.19729 | -0.21496 | -0.27166 | -0.08862 | -0.09896 | -0.09837 | 0.17896 | -0.08134 | 0.15942 | 0.12038 | -0.12471 | 0.124335 | -0.13313 | 0.12502 | -0.14942 | -0.41539 |
| HUNS1 | -0.21847 | 0.071844 | -0.08309 | 0.07356 | -0.21676 | -0.161 | 0.109109 | -0.15098 | -0.12186 | 0.09466 | -0.04 | -0.04678 | -0.02038 | -0.15885 | -0.17667 | -0.15068 | 0.01863 | 0.27508 | 0.056613 | 0.21355 | -0.00719 | -0.22925 | 0.059084 | -0.02094 | 0.07761 | 0.06672 | -0.62886 |
| HEC50B | -0.18831 | 0.020987 | -0.08559 | -0.09238 | -0.17382 | -0.08017 | -0.07009 | -0.27053 | -0.1673 | -0.24978 | -0.05718 | 0.09004 | -0.08659 | -0.14187 | -0.16136 | -0.21052 | -0.19792 | 0.31398 | -0.05269 | 0.05152 | 0.0525 | -0.08318 | 0.056542 | -0.09343 | 0.00288 | 0.061645 | -0.376 |
| CAL27 | -0.17537 | 0.213342 | 0.0541 | -0.02983 | -0.07539 | -0.13716 | -0.04345 | -0.08104 | -0.09351 | -0.18359 | -0.07927 | 0.0881 | -0.09012 | -0.06823 | -0.06591 | -0.12951 | -0.05679 | 0.17612 | -0.04302 | 0.10688 | 0.09269 | -0.07356 | 0.016293 | -0.0951 | 0.04582 | 0.088359 | -0.36106 |
| RH30 | -0.17455 | 0.080354 | -0.86516 | -0.00149 | -0.00229 | -0.21041 | -0.33625 | -0.17157 | -0.08803 | 0.26126 | -0.09524 | 0.22798 | -0.47397 | -0.19119 | -0.1474 | -0.17103 | -0.18076 | -0.09022 | 0.045284 | 0.01715 | 0.09826 | -0.09936 | 0.104034 | 0.16314 | 0.29742 | -0.35852 | -0.38185 |
| UMUC1 | -0.18717 | 0.032789 | -1.20196 | -0.01777 | -0.21699 | -0.38142 | -0.1332 | -0.27533 | -0.35252 | -0.0031 | -0.14541 | 0.17008 | -0.07587 | -0.06341 | -0.18873 | 0.003257 | -0.05582 | 0.23164 | -0.01557 | 0.21362 | -0.07749 | -0.04041 | 0.17231 | -0.06211 | 0.21528 | 0.124347 | -0.42205 |
| GCT | -0.11467 | -0.07786 | -0.31969 | -0.19725 | -0.15356 | -0.11504 | 0.101778 | -0.1192 | -0.27139 | -0.02183 | -0.07113 | 0.1363 | -0.23167 | -0.04649 | -0.01702 | -0.13333 | -0.09617 | 0.02317 | 0.006069 | 0.17663 | 0.06307 | -0.16958 | 0.103306 | -0.00817 | 0.10404 | -0.03446 | -0.41399 |
| YD15 | -0.22204 | -0.03211 | -0.24331 | -0.07306 | -0.20926 | 0.06777 | -0.00191 | -0.25609 | -0.07048 | -0.04958 | -0.16053 | 0.05918 | -0.31 | -0.1619 | -0.1822 | -0.12237 | -0.16386 | 0.03346 | -0.04126 | 0.1445 | 0.0351 | -0.05129 | 0.180404 | 0.05383 | 0.06087 | 0.022541 | -0.38581 |
| NCIH322 | -0.08589 | -0.01857 | -0.49104 | -0.13904 | -0.00794 | -0.17893 | -0.09757 | -0.055 | -0.2893 | 0.01147 | -0.0813 | 0.09125 | -0.21772 | 0.01946 | -0.08577 | -0.05416 | -0.15601 | 0.12629 | -0.02615 | -0.00021 | 0.15071 | -0.15055 | 0.054911 | 0.04442 | 0.1083 | 0.021093 | -0.38552 |
| AMO1 | -0.08794 | 0.043533 | -1.08018 | -0.44518 | -0.16885 | 0.08734 | -0.05691 | -0.07762 | -0.14788 | 0.16105 | -0.05743 | 0.09614 | 0.10132 | 0.04343 | 0.011871 | 0.00138 | -0.08133 | 0.19169 | 0.150624 | 0.17345 | 0.00032 | -0.32632 | 0.164068 | -0.11799 | 0.08136 | 0.103308 | -0.40003 |
| SCABER | -0.1527 | -0.02995 | -0.20606 | -0.05245 | -0.22263 | -0.22467 | -0.02411 | -0.19779 | -0.13453 | -0.29042 | -0.0435 | 0.20585 | -0.23973 | 0.00028 | -0.09179 | -0.01227 | -0.02499 | -0.04545 | -0.10305 | 0.04997 | 0.08586 | -0.27468 | 0.080115 | 0.04689 | 0.20424 | 0.061206 | -0.38297 |
| HCC366 | -0.15276 | 0.080069 | -0.42054 | -0.13636 | -0.27157 | -0.23289 | -0.09659 | -0.07338 | -0.14831 | -0.04578 | -0.10635 | 0.2963 | -0.42014 | -0.02264 | -0.1263 | -0.02758 | 0.13792 | 0.28602 | 0.186911 | 0.27949 | 0.03858 | -0.21934 | 0.007942 | -0.28738 | 0.21228 | 0.063789 | -0.5089 |
| NCIH2087 | -0.2133 | -0.02424 | -0.43507 | -0.14827 | -0.14679 | -0.1578 | -0.08426 | -0.06504 | -0.07301 | -0.10983 | 0.077191 | 0.06384 | -0.17276 | -0.04793 | 0.037845 | -0.00358 | 0.00065 | 0.12531 | -0.0795 | 0.08458 | 0.12399 | -0.2686 | 0.052761 | -0.03963 | 0.17646 | 0.011042 | -0.77949 |
| HARA | -0.07587 | -0.0591 | -0.37331 | -0.05859 | -0.1599 | -0.20283 | 0.016522 | -0.15752 | -0.2948 | -0.01612 | -0.12184 | 0.18166 | -0.26886 | -0.1298 | -0.02775 | -0.267 | -0.15991 | 0.16185 | -0.16896 | 0.13807 | 0.14798 | -0.23357 | 0.079037 | -0.0905 | 0.17651 | 0.005182 | -0.33619 |
| NCIH1373 | -0.04763 | 0.135607 | -0.16237 | -0.04369 | -0.12333 | 0.12123 | -0.10136 | -0.08962 | -0.22739 | -0.22707 | -0.08157 | -0.0245 | -0.12815 | -0.23761 | -0.18807 | -0.2939 | -0.05226 | 0.32445 | -0.26219 | 0.13615 | 0.27037 | -0.33147 | 0.052081 | -0.03751 | -0.03092 | 0.114517 | -0.41197 |
| FADU | -0.15861 | 0.166095 | -0.07865 | 0.11985 | -0.2542 | -0.15415 | -0.19157 | -0.28184 | -0.15872 | 0.03359 | -0.07298 | 0.07778 | -0.18879 | -0.02472 | -0.04986 | -0.06783 | -0.05407 | 0.04755 | -0.10581 | 0.21897 | 0.14356 | -0.19092 | 0.132868 | -0.1666 | -0.03743 | -0.07973 | -0.16944 |
| HGC27 | -0.12842 | -0.0301 | -0.00172 | -0.12513 | -0.14916 | -0.26794 | -0.02653 | -0.21992 | -0.053 | 0.03769 | -0.0018 | 0.10361 | -0.41059 | -0.2098 | -0.0242 | -0.16977 | -0.11629 | 0.19863 | 0.070263 | 0.08181 | 0.09201 | -0.26992 | -0.04428 | 0.04611 | 0.16133 | -0.05758 | -0.20545 |
| JHH7 | -0.42793 | 0.053225 | -0.83222 | -0.09131 | -0.08001 | -0.21792 | -0.08606 | -0.11032 | -0.39585 | -0.06697 | -0.02726 | 0.20065 | -0.0499 | 0.12087 | -0.06109 | -0.1137 | -0.11727 | 0.10785 | -0.0365 | 0.09926 | -0.07436 | -0.06534 | 0.159788 | -0.0495 | 0.03602 | -0.05947 | -0.35834 |
| MDAMB468 | -0.23454 | -0.10226 | -0.08898 | 0.03412 | -0.06526 | 0.0222 | -0.08678 | -0.12668 | 0.060107 | -0.00149 | -0.08294 | 0.19578 | -0.13836 | 0.00481 | -0.02173 | -0.11893 | 0.02598 | 0.24618 | 0.002088 | 0.06879 | -0.03462 | -0.19816 | 0.287237 | -0.06828 | 0.07769 | 0.106227 | -0.28053 |
| MORCPR | -0.11526 | 0.206128 | -0.20412 | 0.1345 | -0.01696 | -0.16507 | 0.078052 | -0.13561 | -0.28725 | -0.26797 | -0.61448 | 0.13171 | 0.02615 | 0.03052 | -0.18786 | -0.05777 | 0.04065 | -0.0748 | 0.039415 | 0.13318 | 0.07054 | -0.11694 | 0.01752 | 0.05195 | 0.31304 | -0.10234 | -0.41664 |
| NCIH661 | -0.01583 | -0.06466 | -0.13499 | -0.14966 | -0.06396 | -0.10845 | 0.050438 | -0.11461 | -0.22709 | 0.05319 | -0.0581 | 0.01468 | -0.06069 | -0.03932 | 0.027379 | -0.30665 | -0.06606 | 0.13601 | 0.015425 | 0.08829 | 0.04635 | -0.07439 | 0.178543 | -0.026 | 0.1381 | 0.069683 | -0.09882 |
| OCIMY5 | 0.00166 | 0.082281 | -0.37678 | -0.14364 | -0.0989 | 0.02321 | 0.167211 | 0.06758 | -0.0231 | -0.22898 | -0.06171 | 0.04279 | 0.08376 | -0.42494 | -0.10366 | 0.036526 | -0.09261 | 0.24395 | -0.00887 | 0.19006 | 0.06269 | -0.06502 | 0.093407 | -0.11309 | 0.19797 | -0.04448 | -0.55562 |
| KYSE150 | -0.13482 | -0.01924 | -0.21651 | -0.04794 | -0.34321 | -0.43249 | -0.0956 | -0.16717 | -0.01613 | 0.18114 | 0.046379 | 0.11977 | -0.22576 | -0.10739 | -0.08699 | -0.2337 | -0.09893 | 0.1025 | -0.12958 | 0.02995 | 0.05162 | -0.3556 | 0.076588 | -0.08658 | 0.13059 | -0.02471 | -0.36881 |
| CAL51 | -0.08933 | -0.15201 | -0.07088 | 0.11703 | -0.09178 | -0.11729 | -0.01196 | -0.20824 | -0.20117 | -0.06872 | -0.14015 | 0.21149 | -0.09107 | -0.00544 | -0.00846 | -0.11769 | 0.06032 | 0.14761 | 0.195398 | 0.19991 | 0.03292 | 0.016775 | 0.043029 | 0.02462 | 0.18342 | 0.019195 | -0.17209 |
| KNS62 | -0.16446 | -0.06184 | -0.18642 | -0.06041 | -0.17421 | -0.17467 | -0.04192 | -0.14454 | -0.14931 | -0.03102 | -5.5E-05 | 0.11341 | -0.28934 | -0.1788 | -0.09762 | -0.15989 | 0.01728 | 0.03175 | -0.10109 | 0.09544 | 0.02767 | -0.35868 | 0.083802 | -0.11061 | 0.12543 | 0.019617 | -0.24865 |
| HCC1954 | -0.11886 | -0.04328 | -0.13107 | 0.04867 | -0.17314 | -0.36047 | -0.14637 | -0.03564 | -0.05843 | -0.11778 | -0.05009 | 0.15832 | -0.30656 | -0.26348 | -0.05402 | -0.36817 | -0.16628 | 0.1875 | -0.07327 | 0.27378 | 0.27117 | -0.19481 | 0.262203 | -0.00814 | 0.14734 | -0.037 | -0.35073 |
| NCIH358 | 0.03968 | -0.02836 | -0.53211 | -0.11764 | -0.27355 | -0.10341 | -0.07631 | -0.52638 | -0.16292 | -0.12023 | -0.03052 | 0.129 | -0.19497 | -0.25854 | -0.17954 | -0.21947 | 0.07058 | 0.10359 | -0.09924 | 0.2264 | 0.08903 | -0.19885 | 0.144813 | -0.04062 | 0.13049 | -0.01895 | -0.46867 |
| HOP62 | -0.09219 | 0.061539 | -0.24506 | -0.0216 | -0.00405 | -0.07011 | 0.054636 | -0.05508 | -0.36906 | -0.08936 | -0.05341 | 0.02499 | -0.1289 | -0.06453 | -0.19447 | -0.02448 | 0.06894 | -0.03736 | 0.000954 | 0.11662 | -0.02468 | -0.16792 | -0.09433 | -0.00376 | 0.18943 | -0.02321 | -0.29922 |
| KMBC2 | -0.1358 | -0.04644 | -0.22548 | -0.16593 | -0.04391 | -0.07136 | -0.14957 | -0.21409 | 0.000167 | -0.0382 | -0.03779 | -0.0016 | -0.07193 | -0.14817 | -0.09316 | -0.22214 | -0.04187 | -0.07591 | 0.048266 | 0.05697 | 0.01623 | -0.11627 | 0.094484 | -0.0379 | 0.04944 | -0.03142 | -0.37031 |
| DBTRG05MG | -0.13516 | -0.06683 | -0.45848 | 0.0222 | 0.01211 | -0.0273 | 0.099717 | -0.14544 | -0.14175 | 0.08907 | -0.19305 | 0.26069 | -0.08881 | -0.07943 | -0.06428 | 0.007288 | 0.09299 | 0.00918 | 0.028587 | 0.07375 | 0.09295 | -0.10068 | 0.108514 | 0.01944 | 0.17036 | -0.12536 | -0.22404 |
| COLO684 | 0.02131 | 0.130655 | -0.01362 | 0.10056 | 0.15566 | -0.03473 | 0.162664 | -0.22541 | 0.004998 | -0.19716 | -0.01854 | 0.15735 | -0.05764 | 0.08599 | 0.000164 | -0.01359 | -0.191 | 0.19852 | 0.008593 | 0.01718 | 0.29593 | -0.13303 | 0.095682 | -0.0102 | 0.19204 | -0.02323 | -0.01322 |
| KYSE450 | -0.16483 | 0.026996 | -0.17312 | -0.04307 | -0.15356 | -0.09078 | 0.01001 | -0.19087 | -0.25062 | -0.02558 | -0.03664 | 0.07463 | -0.12598 | -0.00187 | -0.06156 | -0.0929 | 0.02908 | 0.17757 | -0.11427 | 0.0802 | -0.00846 | -0.10757 | 0.02838 | -0.05302 | 0.20191 | -0.10281 | -0.35158 |
| NCIH1048 | -0.14219 | 0.024974 | -0.18445 | -0.10843 | -0.08787 | -0.13047 | -0.04624 | -0.14447 | -0.11247 | 0.00947 | -0.09761 | 0.0699 | -0.21892 | -0.058 | 0.007356 | -0.12356 | 0.04226 | -0.06347 | -0.0615 | 0.05006 | 0.06377 | -0.15427 | 0.257957 | -0.11285 | 0.2105 | -0.02822 | -0.34066 |
| CHAGOK1 | -0.20355 | 0.035411 | -0.37921 | 0.03636 | -0.08982 | -0.20689 | -0.02033 | -0.18837 | -0.12677 | -0.10506 | -0.03706 | 0.04545 | -0.11493 | -0.02084 | -0.09309 | -0.11701 | -0.05506 | 0.11541 | -0.04748 | 0.13491 | 0.07639 | -0.09281 | 0.095121 | -0.06237 | 0.10903 | 0.024622 | -0.34492 |
| NCIH1568 | -0.07645 | 0.075742 | -0.29141 | -0.05767 | -0.14657 | -0.29614 | 0.09696 | -0.18032 | -0.31233 | -0.09709 | -0.09163 | 0.10721 | -0.18515 | 0.01455 | -0.03294 | -0.11049 | -0.14896 | 0.02543 | -0.06289 | 0.01205 | 0.15308 | -0.45978 | 0.175862 | -0.03656 | 0.04935 | -0.13862 | -0.72003 |
| NCIH510 | -0.17206 | -0.06356 | -0.02229 | -0.08544 | -0.14263 | -0.09933 | -0.11395 | -0.15301 | -0.07069 | -0.05673 | -0.08279 | 0.13914 | -0.26852 | -0.34085 | -0.0272 | 0.007429 | -0.07706 | -0.00577 | -0.0117 | 0.2143 | 0.13934 | -0.47244 | 0.074938 | -0.0307 | 0.15493 | 0.036374 | -0.44793 |
| HCC515 | -0.06017 | -0.03146 | -0.51298 | -0.09216 | -0.08164 | -0.23519 | -0.09134 | -0.18097 | 0.019984 | -0.05613 | -0.06067 | 0.1339 | -0.28016 | -0.19312 | -0.09938 | -0.30861 | -0.01537 | 0.08414 | -0.12237 | 0.13168 | 0.27754 | -0.53915 | -0.09548 | -0.1056 | 0.16983 | 0.083145 | -0.51186 |
| KYSE270 | -0.26038 | -0.02011 | -0.16378 | 0.12226 | -0.07259 | 0.01781 | 0.030649 | -0.04013 | 0.058408 | 0.03785 | -0.09017 | 0.12486 | -0.30268 | -0.096 | -0.08135 | -0.13509 | 0.15744 | 0.25907 | -0.0158 | -0.0516 | 0.02693 | -0.12047 | -0.00904 | -0.20196 | 0.15793 | 0.178695 | -0.38457 |
| MDAMB415 | -0.08537 | -0.06605 | -0.12041 | -0.02051 | -0.0847 | -0.32259 | -0.24477 | -0.07822 | -0.03931 | -0.17753 | 0.014318 | 0.19484 | -0.26182 | -0.27983 | -0.06625 | -0.3369 | -0.03988 | 0.19856 | -0.13161 | 0.23527 | 0.27165 | -0.03327 | 0.092753 | -0.03291 | 0.09465 | -0.19598 | -0.52041 |
| HCC15 | -0.21107 | -0.13558 | -0.39694 | -0.03943 | -0.36968 | -0.33352 | -0.15356 | -0.01264 | -0.23558 | -0.16538 | -0.06901 | 0.1167 | -0.32698 | -0.3526 | -0.45943 | -0.47112 | 0.01331 | 0.12205 | 0.13411 | -0.04145 | 0.01056 | -0.27252 | -0.07191 | -0.29903 | 0.13955 | -0.05652 | -1.10575 |
| MFE296 | -0.10848 | 0.344291 | -0.08183 | -0.0109 | 0.17724 | -0.13248 | 0.175306 | 0.17579 | 0.016973 | -0.03301 | -0.05819 | 0.14678 | 0.15928 | 0.04098 | -0.01853 | 0.012672 | -0.03309 | 0.24361 | 0.147429 | 0.11239 | 0.39676 | 0.014654 | -0.07765 | 0.02171 | 0.16333 | 0.09908 | -0.31234 |
| AGS | -0.06024 | 0.075807 | -0.14496 | 0.00394 | -0.02317 | -0.07004 | 0.051743 | -0.17988 | 0.060311 | -0.14272 | -0.14447 | 0.08948 | -0.13682 | -0.11482 | -0.06577 | -0.04045 | 0.09445 | 0.27739 | -0.06489 | 0.2679 | 0.06557 | -0.0186 | -0.01119 | -0.10066 | 0.20162 | -0.00327 | -0.27725 |
| MELJUSO | -0.18325 | 0.014974 | -0.80542 | -0.01037 | -0.07651 | -0.1841 | 0.042544 | -0.02896 | -0.07519 | 0.01862 | -0.05973 | 0.12954 | -0.16568 | -0.1338 | -0.11245 | -0.23713 | -0.21762 | 0.17695 | -0.18555 | 0.17461 | 0.28012 | -0.1245 | 0.073306 | -0.22596 | 0.32978 | -0.04843 | -0.47973 |
| IGR1 | 0.07446 | 0.03365 | -0.30817 | -0.12338 | -0.21263 | -0.19212 | -0.08216 | -0.09437 | -0.15073 | 0.03919 | -0.076 | 0.1173 | -0.0824 | 0.05694 | 0.038295 | -0.18436 | -0.0505 | 0.14171 | 0.023798 | 0.12667 | 0.0237 | -0.2211 | 0.157947 | 0.02221 | 0.18195 | -0.03437 | -0.36765 |
| MDAMB435S | -0.27687 | 0.079391 | -0.78746 | -0.08302 | -0.24094 | -0.15134 | -0.0277 | -0.03678 | -0.1503 | 0.09278 | -0.10644 | 0.04753 | -0.15578 | -0.13708 | -0.0324 | -0.11478 | -0.17599 | 0.12929 | -0.04506 | 0.20886 | -0.01756 | -0.16385 | 0.054552 | -0.0433 | 0.12743 | -0.12388 | -0.29915 |
| TOV21G | -0.02359 | 0.004389 | 0.11172 | 0.04452 | -0.08726 | -0.14346 | 0.013722 | -0.34531 | -0.11705 | -0.07 | 0.001896 | 0.04749 | -0.2024 | -0.06513 | -0.0304 | -0.09942 | -0.0323 | 0.02821 | -0.02168 | 0.12499 | 0.11208 | -0.08144 | 0.151691 | -0.02711 | 0.26629 | -0.04187 | -0.32909 |
| NCIH2009 | -0.13444 | 0.073814 | -0.34144 | -0.00082 | -0.07354 | -0.12349 | 0.065179 | -0.21776 | -0.00103 | -0.04382 | -0.08212 | 0.1698 | -0.25138 | -0.16048 | -0.11091 | -0.13803 | 0.02364 | -0.16338 | 0.022131 | 0.17937 | 0.13618 | 0.053941 | 0.137418 | -0.18861 | 0.15476 | 0.061673 | -0.34515 |
| SF172 | -0.24177 | 0.085203 | -0.40642 | -0.12173 | -0.13543 | -0.23639 | -0.01402 | -0.18771 | -0.04935 | -0.05169 | -0.06769 | 0.13007 | -0.11458 | -0.10786 | -0.04989 | -0.04026 | 0.02703 | 0.05233 | -0.02101 | 0.03503 | 0.15438 | -0.2423 | 0.210622 | -0.06844 | 0.15604 | -0.05022 | -0.25415 |
| NCIH1793 | -0.05929 | -0.00979 | -0.22152 | -0.08326 | -0.14239 | -0.15656 | 0.036153 | -0.32554 | -0.24216 | 0.01241 | -0.02874 | 0.07382 | -0.24704 | -0.06258 | -0.04219 | -0.05224 | -0.11025 | 0.1954 | -0.03489 | 0.12178 | 0.15864 | -0.19403 | 0.146679 | -0.02779 | 0.17699 | -0.03367 | -0.32573 |
| KMM1 | -0.21394 | -0.07444 | -0.70463 | -0.15774 | -0.03208 | -0.10799 | -0.06064 | 0.01895 | -0.1947 | 0.08363 | -0.04186 | 0.08489 | -0.13598 | -0.0491 | 0.067166 | -0.08723 | 0.00463 | 0.09358 | -0.03601 | 0.09332 | 0.04705 | -0.18747 | 0.071975 | -0.12226 | 0.16884 | -0.06274 | -0.28971 |
| SW1271 | -0.14558 | 0.020637 | -0.19973 | -0.20432 | -0.06426 | -0.24199 | 0.036339 | -0.00797 | -0.06579 | 0.10652 | -0.02267 | 0.0646 | -0.19691 | 0.0232 | -0.00868 | -0.19451 | -0.11437 | 0.01309 | -0.00167 | 0.1159 | 0.00896 | -0.22595 | -0.03378 | 0.05768 | -0.05204 | 0.009962 | -0.47407 |
| NCIH1869 | -0.11425 | 0.019326 | 0.15253 | -0.11819 | -0.28603 | -0.42628 | -0.09329 | -0.12153 | -0.12865 | -0.20658 | -0.0105 | 0.21036 | -0.25187 | -0.1768 | -0.16237 | -0.17894 | -0.00762 | 0.02549 | -0.20028 | 0.07796 | 0.06592 | -0.22548 | 0.047608 | -0.03467 | 0.11221 | 0.113558 | -0.48316 |
| 647V | -0.17651 | 0.200997 | -0.09946 | -0.07883 | -0.15192 | -0.11961 | 0.036613 | -0.1115 | -0.2502 | -0.07228 | -0.09399 | -0.04691 | -0.31248 | -0.17544 | -0.11646 | -0.1275 | 0.0692 | 0.06967 | -0.09598 | 0.04968 | -0.02586 | -0.17272 | 0.123922 | -0.10895 | 0.11174 | 0.032921 | -0.31901 |
| SNU719 | -0.16938 | 0.094686 | -0.59731 | -0.18771 | -0.18772 | -0.17822 | -0.08818 | -0.04716 | 0.024197 | 0.06153 | -0.01431 | 0.05119 | -0.22255 | 0.0931 | -0.01025 | -0.07843 | 0.01898 | 0.00694 | -0.10397 | 0.07034 | 0.12233 | -0.15848 | 0.095792 | -0.02939 | 0.19429 | 0.049432 | -0.36615 |
| WM88 | 0.00712 | 0.213989 | -0.18862 | -0.17116 | 0.15662 | -0.06581 | 0.156505 | -0.1038 | -0.24568 | -0.09845 | 0.005917 | 0.05462 | -0.24953 | -0.33774 | 0.059257 | 0.116784 | -0.35554 | 0.21259 | 0.085583 | 0.27759 | -0.23212 | -0.04178 | 0.05394 | -0.05714 | -0.17359 | -0.18746 | -0.53786 |
| NCIH23 | -0.31479 | 0.054195 | -0.27495 | 0.07208 | -0.22422 | -0.05463 | -0.0639 | -0.17868 | -0.09649 | -0.20106 | -0.04339 | 0.10925 | -0.4239 | 0.0837 | -0.06501 | -0.0778 | -0.13912 | 0.074 | -0.08131 | 0.08688 | 0.02833 | -0.10898 | 0.171875 | -0.19866 | 0.13634 | 0.130067 | -0.37903 |
| HCC1359 | -0.01796 | -0.02409 | -0.18229 | -0.13108 | -0.15952 | -0.22513 | -0.04783 | -0.12007 | -0.29115 | -0.14283 | -0.07407 | 0.04434 | -0.30396 | 0.03203 | -0.12876 | -0.03938 | -0.14721 | 0.17178 | -0.02754 | 0.11837 | 0.04288 | -0.15446 | -0.03393 | -0.09929 | 0.18635 | -0.07878 | -0.4346 |
| FTC133 | -0.1523 | 0.139605 | -0.1138 | -0.06213 | -0.12316 | -0.1245 | -0.03412 | -0.1169 | 0.011897 | -0.07372 | 0.09987 | 0.02502 | -0.32619 | -0.06937 | -0.06275 | -0.16516 | -0.12689 | 0.20349 | -0.09418 | 0.02545 | 0.17463 | -0.18345 | 0.142745 | -0.04307 | -0.01152 | 0.001208 | -0.32306 |
| 5637 | -0.34044 | 0.00182 | -0.60362 | -0.10092 | -0.091 | -0.27133 | -0.07461 | -0.23063 | -0.1261 | -0.07011 | -0.03324 | 0.11061 | -0.34312 | -0.06064 | -0.17952 | -0.06609 | 0.02818 | 0.21206 | -0.05008 | 0.14737 | -0.02424 | -0.06234 | 0.064163 | -0.10049 | 0.06729 | 0.163562 | -0.85686 |
| ES2 | -0.00867 | 0.038301 | -0.05293 | -0.09756 | -0.12662 | -0.19376 | 0.095606 | -0.10666 | -0.22059 | -0.04853 | -0.07312 | 0.15502 | -0.14368 | -0.01025 | -0.06862 | -0.00383 | -0.05778 | 0.05406 | -0.07023 | 0.05719 | 0.06743 | -0.2246 | 0.130591 | -0.03811 | 0.23919 | -0.03064 | -0.28896 |
| SNU349 | -0.11834 | -0.1295 | -0.13498 | 0.09383 | -0.23381 | -0.13664 | 0.082953 | -0.25604 | -0.05632 | -0.18035 | -0.03939 | 0.20637 | -0.46686 | -0.27035 | -0.2705 | -0.21412 | -0.3226 | 0.12349 | -0.07141 | 0.06798 | 0.24136 | -0.1943 | 0.214729 | -0.08852 | 0.09611 | 0.104062 | -0.27815 |
| MDAMB453 | -0.16225 | -0.07417 | -0.05912 | -0.01899 | -0.06811 | 0.00287 | 0.147454 | -0.19065 | 0.055735 | -0.06195 | 0.031995 | 0.1956 | -0.08577 | -0.26176 | 0.006679 | -0.03097 | -0.04477 | 0.08474 | -0.08807 | 0.09995 | 0.12292 | -0.126 | -0.01694 | 0.03513 | 0.14381 | 0.057991 | -0.68642 |
| NUGC3 | -0.13698 | 0.075517 | -0.13655 | -0.05268 | -0.13157 | -0.13738 | -0.06647 | -0.23857 | -0.19397 | -0.1602 | -0.03943 | 0.12095 | -0.21846 | 0.00626 | -0.11972 | -0.10441 | -0.01086 | 0.0493 | 0.010911 | 0.14886 | 0.12705 | -0.0912 | 0.186441 | 0.0336 | 0.17436 | 0.065331 | -0.37913 |

| Cell line | FLT3 | FLT4 | IGF1R | INSR | INSRR | KDR | KIT | LMR1 | LMR2 | LTK | MET | MUSK | PDGFRA | PDGFRB | PTK7 | RET | RON | ROR1 | ROR2 | ROS1 | STYK1 | TIE1 | TIE2 | TRKA | TRKB | TRKC | TYRO3 |
|---|---|---|---|---|---|---|---|---|---|---|---|---|---|---|---|---|---|---|---|---|---|---|---|---|---|---|---|
| NCIH2286 | -0.16563 | -1.6E-05 | -0.05763 | -0.05324 | -0.14846 | -0.18727 | -0.04488 | -0.16501 | -0.15336 | -0.26475 | -0.59868 | 0.03991 | -0.1651 | -0.11461 | -0.13189 | -0.02418 | -0.09501 | 0.07098 | -0.06498 | 0.12772 | 0.29297 | -0.1948 | 0.097787 | -0.10482 | 0.14962 | -0.091 | -0.3149 |
| ESS1 | -0.12165 | -0.01178 | -0.1104 | -0.10021 | -0.17541 | -0.20103 | 0.061769 | -0.11 | -0.003 | -0.21783 | -0.10807 | 0.10949 | -0.14516 | -0.11551 | -0.0215 | -0.12338 | -0.23635 | 0.02246 | 0.026592 | 0.1123 | 0.03589 | -0.15783 | 0.028322 | -0.01656 | 0.13512 | -0.07268 | -0.4616 |
| HT | -0.08602 | -0.00651 | -0.02555 | -0.04947 | 0.02853 | -0.06593 | -0.03596 | -0.11764 | -0.0965 | -0.04064 | -0.09551 | 0.10484 | -0.19625 | -0.15761 | -0.1405 | -0.11915 | -0.09437 | 0.10858 | 0.156824 | 0.15696 | 0.05469 | -0.049 | 0.015914 | -0.06547 | 0.09052 | -0.02474 | -0.55327 |
| IPC298 | -0.08816 | -0.00349 | -0.62506 | -0.16532 | -0.12193 | -0.05662 | 0.009199 | -0.1137 | 0.075992 | 0.08769 | -0.14415 | 0.04231 | -0.11195 | -0.18773 | -0.0376 | -0.02963 | -0.0997 | 0.3575 | -0.10478 | -0.09339 | 0.22276 | -0.00878 | 0.047229 | 0.0369 | -0.00657 | 0.04194 | -0.23578 |
| NCIH1573 | -0.08965 | 0.134042 | -0.51212 | -0.14299 | -0.20505 | -0.04193 | -0.20863 | -0.14111 | -0.6563 | 0.11563 | 0.043647 | 0.01192 | 0.03569 | 0.52351 | -0.00668 | 0.068302 | 0.1191 | -0.48738 | -0.31652 | 0.1078 | -0.06061 | 0.073765 | -0.37009 | -0.48177 | 0.24292 | 0.202462 | -0.77536 |
| TE4 | -0.14025 | -0.01538 | -0.14581 | -0.12423 | -0.14095 | -0.24988 | -0.11242 | -0.14545 | -0.18041 | -0.02141 | -0.05642 | 0.15775 | -0.27186 | -0.00842 | -0.06766 | -0.08843 | -0.07465 | -0.00282 | -0.09796 | 0.09397 | 0.05487 | -0.21902 | 0.148289 | -0.00517 | 0.07868 | 0.070813 | -0.4248 |
| IM95 | -0.12413 | 0.170702 | 0.0364 | -0.11338 | -0.01709 | -0.07973 | 0.032975 | -0.00293 | 0.059449 | 0.10073 | -0.62117 | -0.01798 | -0.02591 | -0.00861 | -0.07516 | -0.06401 | 0.10731 | 0.18819 | 0.291959 | -0.00314 | 0.13646 | 0.078369 | -0.04629 | 0.13452 | 0.09272 | -0.0039 | -0.38288 |
| CMLT1 | -0.31503 | 0.198192 | -0.12004 | 0.0297 | 0.04326 | -0.57006 | -0.00261 | -0.20503 | -0.22742 | 0.0226 | -0.03994 | 0.36871 | -0.10831 | 0.26862 | -0.16765 | -0.20335 | -0.11717 | 0.34667 | -0.20941 | 0.29268 | -0.04277 | 0.108566 | -0.23259 | 0.04516 | 0.26497 | 0.126483 | -0.29095 |
| NCIH157DM | -0.10236 | 0.027478 | -0.12852 | -0.09564 | -0.25913 | -0.20681 | 0.169434 | -0.17686 | -0.23419 | 0.09279 | -0.09709 | 0.13518 | -0.13212 | -0.04987 | -0.0791 | 0.018919 | -0.01971 | 0.18633 | -0.03376 | 0.09319 | 0.03015 | -0.11444 | 0.043491 | -0.021 | 0.16984 | -0.13347 | -0.20539 |
| RCHACV | -0.01996 | 0.043411 | -0.18641 | 0.00474 | -0.18875 | -0.08744 | 0.047764 | -0.39205 | 0.147629 | 0.04865 | -0.12672 | 0.17251 | -0.12597 | -0.28803 | -0.03641 | -0.11916 | 0.00946 | 0.08222 | -0.21427 | 0.14215 | 0.24621 | 0.000819 | 0.10323 | 0.06331 | 0.31101 | -0.00199 | -0.31327 |
| NCIH2172 | -0.09319 | 0.009572 | -0.3861 | -0.16034 | -0.16692 | -0.27053 | 0.010466 | -0.19277 | 0.077987 | -0.05905 | -0.14787 | 0.18549 | -0.36948 | -0.17956 | -0.12359 | 0.002747 | -0.04866 | -0.03664 | -0.03787 | 0.22495 | 0.11938 | -0.13466 | 0.116549 | 0.03999 | 0.1275 | 0.03477 | -0.17373 |
| HT55 | 0.01813 | 0.045263 | -0.98232 | -0.28109 | -0.18398 | -0.29755 | 0.070362 | -0.1808 | -0.1231 | -0.28825 | -0.05741 | -0.07904 | -0.19006 | -0.27061 | -0.052 | -0.07032 | -0.05123 | 0.2892 | -0.10803 | 0.17607 | 0.29569 | -0.24807 | 0.056041 | -0.00793 | 0.31031 | -0.17448 | -0.53169 |
| JHUEM1 | -0.41063 | -0.22234 | -0.15764 | -0.05909 | 0.03133 | 0.06779 | 0.188025 | -0.34869 | -0.42081 | -0.07369 | 0.038807 | 0.18586 | -0.4124 | -0.35698 | 0.030371 | -0.19055 | 0.00995 | 0.43246 | 0.04218 | 0.12541 | -0.00295 | -0.12245 | 0.035843 | 0.1078 | 0.19696 | 0.136386 | -0.46013 |
| NCIH2110 | 0.0386 | 0.133422 | -0.98587 | -0.21842 | -0.19003 | -0.2122 | 0.001206 | -0.12389 | -0.33875 | 0.03766 | 0.07993 | 0.22426 | -0.24929 | -0.10194 | 0.087345 | -0.13835 | -0.21532 | 0.19406 | -0.0512 | 0.17117 | -0.01106 | -0.10763 | 0.075263 | -0.09322 | 0.22623 | 0.038882 | -0.37852 |
| SNU1 | -0.25346 | -0.03587 | -0.47783 | -0.19796 | -0.03013 | -0.11259 | -0.05668 | -0.21764 | -0.03723 | 0.05749 | -0.08431 | 0.16206 | -1.02105 | -0.13733 | -0.09887 | -0.03813 | -0.16471 | 0.10621 | -0.01956 | 0.10198 | 0.02755 | -0.20949 | 0.083114 | -0.07737 | 0.16314 | 0.015971 | -0.45111 |
| MDAMB361 | -0.19952 | -0.07733 | -0.17913 | -0.03219 | -0.14201 | -0.02386 | -0.16657 | -0.01924 | 0.029728 | -0.03609 | -0.02894 | 0.08956 | -0.0642 | 0.04416 | -0.08751 | -0.21599 | -0.03985 | -0.01181 | -0.10079 | 0.15856 | 0.07173 | -0.22313 | 0.143834 | 0.10585 | 0.07915 | 0.080295 | -0.07964 |
| MDST8 | 0.03982 | 0.150862 | -0.80449 | 0.02091 | -0.15217 | -0.02424 | -0.09315 | -0.01847 | -0.18281 | 0.08244 | -0.06742 | 0.17782 | -0.2787 | -0.1783 | -0.01872 | -0.06779 | -0.05317 | 0.13279 | -0.07618 | 0.13461 | 0.0986 | -0.28542 | 0.150726 | -0.01684 | 0.2325 | -0.11278 | -0.30723 |
| EFO27 | -0.14896 | 0.00859 | -0.11027 | 0.00073 | -0.17651 | -0.28226 | 0.024211 | -0.1811 | -0.1189 | 0.00277 | -0.09571 | 0.01736 | -0.19411 | -0.10715 | -0.08794 | -0.31019 | -0.05286 | 0.21319 | 0.006238 | 0.0617 | 0.16381 | -0.1346 | 0.172696 | -0.08937 | 0.12235 | 0.009416 | -0.40786 |
| PF382 | -0.11003 | -0.02384 | -0.07839 | -0.00013 | -0.17884 | -0.17166 | -0.05431 | -0.20461 | -0.18448 | -0.04936 | -0.057 | 0.06094 | -0.11619 | -0.06052 | -0.05823 | -0.16337 | -0.09327 | 0.18266 | -0.17652 | 0.16906 | 0.08882 | -0.12937 | 0.046357 | -0.08695 | 0.1816 | -0.035 | -0.29833 |
| NALM6 | -0.14149 | -0.06049 | -0.20343 | -0.00223 | -0.27497 | -0.16216 | -0.062 | -0.11865 | -0.22121 | 0.05239 | 0.058394 | -0.03751 | -0.06732 | -0.03221 | -0.03372 | -0.03538 | -0.18459 | 0.18692 | -0.09263 | 0.14289 | 0.26481 | -0.1481 | -0.06166 | -0.05547 | 0.0618 | -0.0295 | -0.28109 |
| SKUT1 | -0.13201 | -0.02466 | 0.12604 | -0.11632 | -0.13203 | -0.15232 | 0.043325 | -0.52602 | -0.17375 | 0.05035 | -0.10513 | 0.21473 | -0.09167 | -0.27824 | -0.07085 | -0.24582 | -0.0654 | 0.03 | -0.07223 | 0.11394 | 0.19475 | -0.28848 | -0.00858 | -0.08413 | 0.13136 | 0.197124 | -0.4554 |
| HEC1B | -0.10247 | 0.033905 | -0.13737 | -0.10124 | -0.27957 | -0.16043 | 0.052867 | -0.04632 | -0.0756 | -0.06972 | -0.06682 | 0.05754 | -0.17585 | -0.16648 | -0.23013 | -0.21199 | -0.15783 | 0.06214 | -0.13315 | 0.11388 | 0.10932 | -0.12218 | 0.010986 | -0.35371 | 0.07707 | -0.08876 | -0.31107 |
| RKO | -0.13119 | 0.087282 | -0.13377 | -0.00549 | -0.12344 | -0.08462 | 0.089073 | -0.15569 | -0.10322 | -0.0479 | -0.1742 | 0.03101 | -0.18804 | -0.0901 | -0.07372 | -0.13508 | -0.07377 | 0.06876 | -0.06219 | 0.067 | 0.16676 | -0.12021 | 0.146857 | -0.04943 | 0.14439 | -0.03911 | -0.29987 |
| NAMALWA | -0.0573 | 0.020728 | -0.19337 | -0.08046 | -0.04053 | -0.11652 | -0.0407 | -0.25116 | 0.106258 | -0.09936 | -0.06634 | 0.17991 | -0.22794 | -0.18662 | -0.12941 | -0.24958 | 0.03341 | 0.07237 | -0.0687 | 0.23242 | 0.14356 | -0.07064 | 0.073762 | -0.01035 | 0.16869 | 0.04531 | -0.2827 |
| NCIH650 | -0.048 | -0.25376 | -0.66751 | -0.14715 | -0.1432 | -0.0527 | -0.09131 | -0.10924 | -0.41172 | -0.01627 | -0.22154 | 0.33471 | -0.14319 | -0.35031 | -0.21232 | -0.11411 | -0.22959 | -0.01537 | 0.088659 | -0.05123 | 0.06408 | -0.49436 | -0.07125 | -0.16046 | 0.4125 | 0.094724 | -0.71449 |
| HEC265 | -0.13961 | 0.060528 | -0.05723 | -0.05984 | -0.12606 | -0.19812 | -0.12029 | -0.14544 | -0.35561 | 0.19665 | -0.09265 | -0.02665 | -0.11469 | -0.05491 | -0.03165 | -0.18566 | 0.00936 | 0.26663 | -0.04543 | 0.13609 | -0.01629 | -0.16949 | 0.103077 | -0.03775 | 0.0509 | 0.058565 | -0.40314 |
| OVK18 | -0.09658 | 0.001142 | -0.09937 | -0.06582 | -0.18771 | -0.20957 | -0.06337 | -0.22845 | -0.07902 | -0.12648 | -0.05938 | 0.06333 | -0.20081 | -0.18816 | -0.14154 | -0.06324 | 0.10363 | 0.11239 | 0.056739 | 0.04389 | 0.1402 | -0.12843 | 0.062779 | -0.14769 | 0.13388 | -0.02681 | -0.46604 |
| 2313287 | -0.14991 | 0.120217 | -0.02423 | 0.01199 | 0.08579 | -0.15059 | 0.031918 | 0.05139 | -0.21698 | -0.01751 | -0.10457 | 0.10815 | -0.13766 | -0.03293 | -0.07338 | -0.08226 | 0.07363 | 0.06592 | 0.024342 | 0.09975 | 0.14009 | -0.13323 | -0.04707 | -0.12636 | 0.05742 | 0.084416 | -0.28589 |
| LOVO | -0.2278 | -0.00067 | -0.17567 | 0.01703 | -0.04917 | -0.1959 | 0.162369 | -0.11492 | 0.25864 | -0.04637 | -0.04145 | 0.04143 | -0.25424 | -0.19705 | -0.08957 | -0.27488 | -0.11728 | 0.22201 | -0.10068 | 0.08282 | 0.05706 | 0.133277 | 0.054658 | -0.05573 | 0.21694 | -0.20932 | -0.64003 |
| SUPT1 | -0.09352 | 0.013262 | -0.0872 | -0.1062 | -0.21196 | -0.25181 | -0.11698 | -0.07052 | -0.10078 | 0.00039 | 0.010158 | -0.06303 | -0.37455 | -0.02968 | -0.06991 | -0.13181 | -0.16011 | 0.20581 | -0.12464 | 0.22498 | 0.23879 | -0.17517 | 0.000313 | 0.02756 | 0.0154 | 0.079194 | -0.41773 |
| 22RV1 | -0.09261 | -0.05785 | -0.32987 | -0.62618 | -0.22507 | -0.38251 | 0.186326 | -0.50604 | -0.09792 | -0.16227 | 0.026732 | 0.58407 | -0.28832 | 0.04833 | 0.034815 | -0.25426 | -0.10335 | 0.07307 | 0.181508 | -0.04846 | 0.3847 | -0.16166 | 0.125305 | 0.14679 | 0.05242 | -0.16013 | -0.35115 |
| LS180 | -0.05838 | 0.121901 | -0.12 | -0.04969 | -0.09461 | -0.16141 | 0.05678 | -0.15714 | -0.10207 | -0.07577 | -0.10655 | 0.08654 | -0.25968 | -0.06565 | -0.08063 | -0.07957 | -0.0104 | 0.10698 | -0.14632 | 0.32188 | 0.21238 | -0.05867 | 0.145411 | -0.17095 | 0.01739 | 0.020211 | -0.49616 |
| SW48 | -0.17102 | 0.017145 | -0.36261 | -0.1257 | -0.0292 | -0.01829 | -0.06116 | -0.01326 | -0.07002 | 0.09096 | -0.05137 | 0.09782 | -0.14432 | 0.08971 | 0.09811 | -0.13254 | -0.01075 | 0.22868 | -0.01215 | 0.0613 | 0.03336 | -0.2201 | -0.02519 | -0.08976 | 0.11824 | 0.045124 | -0.3586 |
| SNUC4 | -0.1943 | -0.08605 | -0.26173 | -0.11564 | -0.03738 | -0.19158 | -0.12167 | -0.23573 | -0.1593 | 0.19245 | 0.029071 | 0.04354 | -0.14455 | 0.04804 | -0.09725 | -0.0667 | 0.07123 | 0.09019 | 0.04576 | 0.02939 | 0.08938 | -0.29851 | 0.142567 | 0.10675 | 0.19075 | 0.029585 | -0.21734 |
| REH | -0.13499 | 0.035363 | -0.37558 | -0.02751 | -0.14556 | -0.21411 | 0.075363 | -0.07329 | -0.16976 | 0.04791 | -0.05041 | 0.03414 | -0.03533 | -0.04743 | -0.11237 | 0.009281 | -0.0885 | -0.02252 | -0.05167 | 0.10527 | 0.1258 | -0.14948 | 0.104873 | 0.0255 | 0.17024 | 0.008483 | -0.30704 |
| ISHIKAWAHERAKLIO0 | -0.22838 | 0.047499 | -0.00031 | 0.05407 | 0.13402 | -0.231 | 0.109169 | 0.05989 | -0.57729 | 0.2848 | 0.026331 | 0.11507 | -0.14685 | -0.20732 | -0.01582 | -0.06881 | -0.00023 | 0.08754 | -0.12388 | 0.01787 | 0.1473 | -0.08527 | -0.04028 | 0.02808 | 0.17504 | 0.25316 | -0.35392 |
| OC314 | -0.43135 | 0.048115 | -0.04122 | -0.12288 | -0.01774 | -0.38379 | -0.22967 | -0.47743 | 0.118362 | -0.25282 | 0.16605 | 0.30255 | -0.2906 | -0.18657 | -0.04215 | -0.29194 | -0.09627 | 0.06583 | 0.146586 | 0.26096 | 0.16066 | -0.18436 | 0.080102 | -0.0246 | 0.20916 | 0.094831 | -0.42996 |
| CCK81 | -0.24996 | 0.010982 | -0.30093 | -0.00265 | -0.10793 | -0.35977 | 0.072052 | -0.20671 | 0.053411 | -0.28403 | -0.10297 | 0.18758 | -0.14006 | -0.20169 | -0.20954 | -0.14235 | -0.08483 | 0.44358 | -0.42076 | 0.19277 | 0.32058 | -0.23349 | 0.068494 | 0.03545 | 0.31233 | -0.11879 | -0.67642 |
| RL952 | -0.21008 | -0.03213 | -0.10241 | -0.05201 | -0.14386 | -0.10588 | -0.00474 | -0.10364 | -0.12363 | -0.01969 | -0.06833 | 0.07924 | -0.14226 | -0.22496 | 0.017375 | -0.08083 | -0.04155 | 0.10886 | -0.09443 | 0.07378 | 0.08276 | -0.1878 | 0.070273 | 0.03307 | 0.12972 | 0.033327 | -0.30288 |
| IGROV1 | -0.18409 | -0.40451 | 0.2259 | -0.17312 | -0.21595 | -0.36083 | 0.182419 | -0.03126 | -0.20116 | 0.26322 | 0.21096 | 0.30747 | 0.0753 | -0.18118 | -0.19536 | -0.08536 | -0.00375 | -0.04887 | 0.049378 | 0.15519 | 0.20431 | -0.04461 | 0.147489 | -0.05755 | 0.34973 | 0.22869 | -0.87399 |
| COLO792 | -0.04768 | 0.086114 | -0.4203 | -0.13801 | -0.00774 | -0.18123 | -0.13124 | -0.26852 | -0.22862 | 0.15129 | -0.16909 | 0.11081 | -0.19037 | 0.04251 | 0.103464 | 0.014366 | -0.15023 | 0.02839 | 0.086527 | 0.21159 | 0.05178 | -0.14738 | -0.01073 | 0.01005 | 0.1537 | -0.03699 | -0.909 |
| KM12 | -0.08963 | 0.109352 | -0.00417 | -0.11007 | -0.21866 | -0.34562 | -0.04297 | -0.03228 | -0.13442 | -0.10678 | -0.03703 | 0.00882 | -0.26864 | -0.04037 | -0.12355 | -0.30487 | -0.07218 | 0.05235 | -0.04248 | 0.11569 | 0.07488 | -0.14601 | -0.12343 | -0.1591 | 0.17759 | -0.04848 | -0.21434 |
| SNUC5 | -0.23353 | 0.188056 | 0.01008 | 0.18827 | 0.0027 | -0.11206 | -0.1168 | 0.01177 | -0.07886 | 0.00931 | 0.063267 | 0.24217 | -0.21371 | 0.08794 | 0.061613 | -0.16149 | -0.05902 | 0.11971 | -0.07517 | 0.00938 | 0.14437 | -0.13995 | 0.162856 | -0.00275 | 0.15527 | -0.06224 | -0.22289 |
| HCT116 | -0.19801 | 0.191672 | -0.14313 | -0.03311 | -0.18451 | 0.03066 | -0.00631 | -0.10465 | -0.25862 | 0.00121 | -0.21041 | 0.02264 | -0.15598 | 0.06563 | 0.079458 | -0.04631 | -0.02061 | 0.08364 | 0.073514 | -0.08138 | -0.0466 | -0.00916 | 0.062936 | 0.10174 | 0.06886 | 0.034805 | -0.50462 |
| 639V | -0.13559 | 0.163601 | -0.26488 | -0.09875 | -0.1218 | -0.01736 | 0.075171 | -0.20644 | -0.0175 | -0.05932 | -0.26153 | 0.0461 | -0.15472 | -0.11747 | -0.10925 | -0.05721 | -0.19525 | 0.08094 | 0.11908 | 0.12291 | 0.02395 | -0.00402 | 0.076066 | -0.05614 | 0.14132 | -0.06489 | -0.36024 |
| SNGM | -0.17164 | 0.125258 | 0.02732 | 0.0089 | -0.03336 | -0.11978 | 0.051039 | -0.28375 | -0.10847 | -0.03835 | -0.00955 | 0.13792 | -0.20464 | -0.07642 | 0.013298 | -0.14105 | -0.02795 | 0.12118 | 0.027528 | 0.10798 | 0.01724 | -0.08618 | 0.140859 | -0.14046 | 0.17342 | 0.007942 | -0.43757 |
| HCC2450 | 0.02856 | 0.042318 | -0.1767 | -0.12152 | -0.10568 | -0.26105 | -0.03724 | -0.15997 | -0.15117 | -0.0553 | -0.004 | 0.0831 | -0.34332 | 0.02494 | -0.15704 | -0.08257 | -0.04175 | 0.06627 | -0.06952 | 0.11725 | -0.01427 | -0.31945 | 0.108229 | 0.01946 | 0.19987 | -0.01986 | -0.32155 |
| HUCCT1 | -0.24629 | 0.069514 | -0.11187 | 0.00781 | -0.17807 | -0.21163 | -0.10752 | -0.01873 | -0.08248 | -0.07304 | 0.012061 | 0.09267 | -0.30116 | -0.06693 | -0.20198 | -0.04026 | 0.03345 | 0.19928 | -0.05142 | 0.17368 | 0.22091 | -0.10313 | 0.129966 | -0.12453 | 0.02514 | -0.12544 | -0.34888 |
| LNCAPCLONEFGC | -0.10373 | 0.111151 | -0.07755 | -0.04986 | 0.03316 | -0.01071 | -0.02663 | -0.17133 | -0.12839 | -0.04096 | -0.10422 | 0.11505 | -0.08256 | -0.13337 | 0.03854 | 0.012636 | 0.05319 | -0.02274 | 0.014849 | 0.13959 | 0.16416 | -0.11015 | 0.079397 | 0.04749 | 0.03733 | -0.00428 | -0.26411 |
| EN | -0.13802 | -0.10578 | -0.1346 | 0.12146 | -0.34784 | -0.41825 | 0.149312 | -0.14638 | -0.30103 | -0.14548 | 0.033587 | 0.236 | -0.18603 | -0.0787 | -0.06725 | 0.020204 | 0.00621 | -0.04618 | 0.050078 | 0.1648 | 0.24523 | -0.24374 | 0.164619 | -0.0079 | 0.52233 | -0.14075 | -0.58197 |
| DU145 | -0.22767 | 0.02641 | -0.18048 | -0.23359 | -0.36575 | -0.24845 | -0.03402 | -0.21519 | -0.10602 | -0.00488 | -0.03728 | 0.16809 | -0.46818 | -0.33807 | -0.26605 | -0.26102 | -0.09238 | -0.07813 | -0.10444 | 0.13895 | 0.13784 | -0.27279 | 0.00451 | -0.12662 | 0.22776 | -0.03398 | -0.43355 |
| NCIH1155 | -0.08917 | 0.017607 | -0.1038 | -0.0399 | -0.13553 | -0.0234 | 0.076469 | -0.21597 | -0.02284 | -0.00901 | -0.14255 | 0.09965 | 0.01945 | -0.12699 | -0.04273 | -0.0681 | -0.0187 | 0.06505 | -0.00418 | 0.06506 | 0.10078 | -0.09971 | 0.073248 | -0.12488 | 0.17454 | 0.067588 | -0.44139 |
| KCL22 | -0.01723 | -0.02229 | -0.09059 | -0.09314 | -0.07056 | -0.14428 | -0.00033 | -0.1781 | -0.21843 | 0.00548 | -0.10716 | 0.0379 | 0.00353 | -0.185 | 0.007266 | -0.13356 | -0.17243 | 0.23761 | -0.00682 | 0.03436 | -0.01397 | -0.22879 | -0.03625 | 0.08705 | 0.15366 | -0.00795 | -0.31779 |
| HEC6 | 0.06791 | -0.05342 | -0.06099 | 0.04927 | -0.2392 | -0.10523 | -0.23713 | 0.00332 | -0.34859 | 0.0013 | 0.091235 | 0.07255 | -0.5297 | -0.18061 | -0.01408 | -0.05685 | -0.00074 | 0.16233 | -0.05618 | 0.16824 | -0.00528 | -0.10742 | 0.033838 | 0.11977 | -0.08035 | -0.05629 | -0.2445 |
| LS411N | -0.27467 | -0.19317 | -0.29324 | -0.12127 | -0.26858 | -0.21119 | -0.08242 | -0.08492 | -0.34119 | -0.12219 | 0.055203 | 0.04338 | -0.14066 | 0.17074 | -0.29909 | -0.05144 | -0.10571 | 0.13401 | -0.25877 | 0.31451 | 0.20098 | -0.05887 | 0.038887 | -0.32611 | 0.08459 | -0.07637 | -0.45941 |
| HT115 | -0.25414 | -0.064 | -0.52593 | -0.08317 | -0.16911 | -0.4244 | -0.17828 | 0.03765 | -0.13022 | -0.18888 | 0.021968 | 0.11741 | -0.34749 | 0.05731 | -0.06267 | -0.05737 | 0.00914 | 0.24447 | -0.1781 | 0.17745 | 0.11281 | -0.16555 | 0.176016 | -0.04683 | 0.16613 | -0.05605 | -0.38215 |
| MFE319 | -0.17411 | 0.025654 | -0.20538 | 0.01741 | 0.03501 | -0.20455 | 0.074551 | -0.09798 | -0.19905 | 0.05219 | -0.02799 | 0.05818 | -0.11556 | -0.19623 | 0.040335 | -0.04728 | -0.03487 | -0.02973 | -0.08532 | 0.06994 | 0.13151 | -0.11306 | 0.041173 | 0.02153 | 0.2176 | 0.094354 | -0.66134 |
| SNU81 | -0.37628 | 0.116125 | 0.18987 | 0.06373 | -0.15974 | -0.13181 | -0.23236 | -0.26772 | -0.0352 | -0.13247 | -0.04072 | 0.11578 | -0.29621 | 0.11119 | -0.28162 | -0.33853 | 0.00362 | 0.11773 | -0.1338 | 0.15144 | 0.10048 | -0.07713 | 0.164297 | -0.03555 | -0.00498 | 0.078265 | -0.75279 |
| JHUEM7 | -0.12084 | -0.09915 | -0.07073 | 0.08653 | -0.08721 | -0.05992 | 0.107158 | -0.2181 | -0.27269 | -0.00427 | 0.019261 | 0.21766 | -0.36898 | 0.01127 | 0.018812 | 0.003706 | -0.04111 | 0.20563 | -0.04012 | 0.1881 | 0.16174 | -0.16537 | -0.15215 | 0.14468 | 0.15689 | 0.151171 | -0.2905 |
| HEC59 | -0.18639 | 0.014698 | -0.18237 | -0.05152 | -0.25077 | 0.09682 | 0.142525 | 0.01941 | -0.24916 | -0.06692 | -0.15375 | 0.21653 | -0.20841 | -0.03608 | 0.108669 | -0.10087 | -0.09055 | 0.55048 | -0.05803 | 0.03541 | 0.0595 | -0.24292 | 0.13618 | -0.03207 | 0.18369 | 0.042795 | -0.34114 |
| JURKAT | 0.0359 | -0.09081 | -0.01697 | -0.23017 | 0.0036 | -0.12372 | 0.046086 | -0.21715 | -0.18468 | -0.18478 | -0.05255 | 0.0521 | 0.0373 | -0.39548 | -8.3E-05 | -0.13985 | -0.17996 | 0.30848 | -0.3719 | 0.09189 | 0.00793 | -0.19602 | -0.04291 | 0.15025 | 0.27737 | 0.167086 | -0.44782 |
| HEC251 | -0.14005 | 0.022456 | 0.086 | -0.01995 | -0.17064 | 0.06358 | 0.140971 | -0.13865 | -0.30954 | -0.11427 | -0.19872 | 0.24431 | -0.19185 | -0.08869 | 0.102503 | -0.11435 | -0.04979 | -0.09186 | -0.08019 | 0.09863 | 0.03857 | -0.23784 | 0.044942 | -0.06674 | 0.15097 | -0.17529 | -0.46566 |
| HCT15 | -0.18871 | 0.153925 | -0.09316 | 0.04099 | -0.13037 | -0.08723 | 0.063 | -0.08551 | -0.13725 | 0.01334 | -0.01601 | 0.05798 | -0.17185 | 0.04695 | 0.008454 | -0.24609 | -0.18913 | 0.06513 | -0.08174 | 0.254 | 0.21754 | -0.00536 | 0.032745 | -0.021 | 0.01967 | 0.056582 | -0.15782 |
| 143B | -0.17956 | -0.07688 | -0.2171 | -0.14598 | -0.05138 | -0.12531 | 0.013962 | -0.05291 | -0.15475 | -0.05538 | -0.05507 | 0.15449 | -0.23574 | -0.01508 | -0.15854 | -0.05955 | -0.03674 | 0.12769 | -0.01381 | 0.16863 | 0.16493 | -0.1879 | 0.004898 | -0.027 | 0.16143 | 0.09207 | -0.23519 |
| BECKER | -0.1578 | 0.136811 | -0.03975 | 0.00436 | -0.06455 | -0.16383 | 0.205552 | 0.09128 | -0.03596 | 0.06151 | -0.99528 | 0.25206 | -0.41436 | -0.14698 | -0.15709 | -0.22985 | -0.062 | 0.09409 | -0.00073 | 0.14864 | 0.16276 | 0.147947 | -0.07817 | 0.28344 | 0.00371 | 0.180912 | -0.4114 |
| BT16 | -0.18187 | -0.16448 | -0.20307 | 0.10421 | -0.14543 | -0.3733 | -0.01639 | -0.53477 | -0.40961 | 0.2937 | -0.05008 | 0.17008 | -0.48365 | -0.24078 | -0.18979 | -0.13762 | -0.17253 | 0.06152 | -0.05777 | 0.19984 | -0.01169 | -0.56398 | -0.13055 | -0.23049 | 0.33492 | 0.338014 | -0.33715 |
| CHLA06ATRT | -0.11938 | 0.021972 | -0.04499 | -0.09461 | -0.11618 | -0.16202 | -0.06252 | -0.11138 | -0.03732 | 0.0357 | -0.85478 | 0.04886 | -0.15876 | -0.12523 | -0.0919 | -0.06805 | -0.00246 | 0.09641 | 0.032775 | 0.08609 | 0.06174 | -0.06477 | 0.100098 | -0.09252 | 0.27346 | -0.07647 | -0.2692 |
| CHLA10 | -0.12297 | 0.008397 | -0.07175 | -0.64681 | -0.07072 | -0.16363 | 0.013047 | -0.19272 | -0.01613 | -0.11012 | -0.16159 | 0.11825 | -0.37779 | -0.29486 | -0.04014 | -0.06641 | -0.10552 | 0.01188 | -0.0314 | 0.11635 | 0.07543 | -0.26407 | 0.041546 | -0.02291 | 0.28633 | 0.051942 | -0.39716 |
| CHLA266 | -0.21624 | 0.009874 | -0.11492 | -0.09362 | -0.09813 | 0.05174 | -0.06397 | -0.04919 | -0.16563 | -0.12667 | -0.00794 | 0.20347 | -0.18787 | -0.13182 | 0.042619 | -0.01092 | -0.25445 | 0.3568 | 0.151115 | 0.1182 | 0.06153 | -0.13104 | 0.160893 | -0.20619 | 0.35508 | 0.192643 | -0.45368 |
| CHLA57 | 0.37073 | -0.03524 | 0.09711 | -0.21628 | -0.15743 | -0.25275 | 0.320115 | -0.21779 | -0.2411 | -0.34874 | -0.04349 | 0.03172 | -0.0427 | -0.28645 | -0.06951 | -0.14555 | -0.02355 | 0.47954 | 0.172066 | 0.2855 | 0.41052 | -0.14632 | 0.207881 | 0.03927 | 0.18941 | 0.197913 | -0.40332 |
| CMK115 | -0.23269 | -0.18972 | -0.03153 | -0.11226 | -0.0801 | -0.12981 | -0.24734 | -0.32514 | -0.14802 | 0.25495 | -0.05388 | 0.09357 | -0.21351 | -0.41669 | 0.002297 | 0.007556 | 0.00821 | 0.07221 | 0.214922 | 0.32924 | 0.23016 | -0.19844 | 0.203731 | 0.08892 | 0.45883 | -0.02304 | -0.55277 |
| COGE352 | -0.06765 | -0.03602 | -0.12736 | -0.60572 | -0.03537 | -0.23012 | -0.10833 | -0.13633 | -0.05629 | 0.15219 | -0.11568 | 0.08723 | -0.19341 | -0.0605 | -0.10253 | -0.04066 | -0.04143 | 0.16799 | -0.0549 | 0.1099 | 0.20257 | -0.2278 | 0.066534 | -0.08185 | 0.14457 | -0.01487 | -0.3626 |
| COLO205 | 0.2004 | -0.03026 | -1.83421 | -0.10111 | -0.01016 | 0.16541 | 0.243339 | -0.1721 | -0.09218 | 0.12947 | -0.04175 | 0.12764 | -0.1219 | 0.04124 | -0.08039 | 0.445582 | -0.06876 | 0.13442 | 0.043308 | 0.11923 | -0.13819 | -0.24273 | -0.02304 | 0.15472 | 0.18144 | 0.077529 | -0.28744 |
| CHL1DM | -0.09916 | -0.09746 | -0.75556 | -0.36301 | -0.15668 | -0.05201 | -0.02496 | -0.25281 | -0.05149 | -0.12462 | -0.18069 | 0.13992 | -0.20625 | -0.21001 | -0.02931 | -0.11774 | -0.16701 | -0.00161 | -0.20561 | 0.25172 | 0.07977 | -0.14487 | 0.049442 | -0.01794 | 0.2794 | 0.050806 | -0.32945 |

| Cell line | FLT3 | FLT4 | IGF1R | INSR | INSRR | KDR | KIT | LMR1 | LMR2 | LTK | MET | MUSK | PDGFRA | PDGFRB | PTK7 | RET | RON | ROR1 | ROR2 | ROS1 | STYK1 | TIE1 | TIE2 | TRKA | TRKB | TRKC | TYRO3 |
|---|---|---|---|---|---|---|---|---|---|---|---|---|---|---|---|---|---|---|---|---|---|---|---|---|---|---|---|
| COV504 | -0.01647 | 0.034782 | -0.66189 | -0.19357 | -0.11736 | -0.19718 | -0.01139 | -0.21385 | -0.02962 | -0.05986 | 0.028749 | 0.1388 | -0.12815 | -0.35563 | -0.16633 | -0.24815 | 0.0698 | -0.05416 | 0.000897 | 0.06301 | 0.23194 | -0.18984 | 0.030715 | -0.06504 | 0.08888 | 0.016123 | -0.41848 |
| CW9019 | -0.08556 | -0.05816 | -0.23541 | -0.11897 | 0.04071 | -0.22807 | -0.00505 | 0.01494 | -0.08552 | 0.00844 | -0.08503 | 0.13308 | -0.18661 | -0.21793 | -0.01745 | 0.016089 | -0.04 | 0.0569 | 0.080648 | 0.14405 | 0.15143 | -0.2556 | -0.07037 | -0.01717 | 0.06676 | 0.018189 | -0.43034 |
| D425 | 0.11988 | 0.154675 | -0.77539 | -0.30924 | -0.08846 | -0.11036 | -0.07603 | -0.21615 | -0.15578 | -0.20012 | -0.06922 | 0.19843 | -0.21136 | -0.04983 | -0.25373 | -0.06943 | 0.17268 | 0.18143 | -0.04149 | 0.03913 | 0.42273 | -0.05727 | -0.00362 | 0.00061 | 0.21303 | 0.022298 | -0.38842 |
| D458 | -0.01534 | 0.037083 | -0.95015 | -0.23019 | 0.02429 | -0.18167 | 0.101489 | -0.23256 | 0.046329 | -0.05559 | -0.02156 | 0.1079 | -0.15096 | -0.02797 | -0.10783 | -0.14833 | 0.08994 | 0.19538 | 0.077915 | 0.18667 | 0.20526 | -0.28448 | 0.142067 | -0.13023 | 0.02317 | -0.34136 | -0.4523 |
| DLD1 | -0.20647 | 0.141984 | -0.41352 | -0.00422 | -0.07582 | -0.20271 | 0.030824 | -0.17342 | -0.04261 | -0.13955 | -0.15143 | 0.09168 | -0.25515 | -0.18705 | -0.07295 | -0.27776 | 0.06473 | 0.09803 | -0.1127 | 0.06273 | 0.18415 | -0.18293 | 0.115867 | -0.07192 | 0.15071 | -0.07208 | -0.47932 |
| DOV13 | -0.26999 | -0.1457 | -0.30995 | -0.12915 | -0.27906 | -0.24075 | 0.029093 | -0.1886 | -0.19149 | 0.12094 | -0.13852 | 0.08405 | -0.16601 | -0.11556 | 0.043774 | -0.31713 | 0.03176 | -0.00236 | -0.0369 | 0.14387 | 0.13932 | -0.41307 | 0.194312 | 0.00568 | 0.12199 | -0.07211 | -0.57951 |
| EVSAT | -0.11763 | 0.033394 | 0.04377 | 0.04858 | -0.01068 | 0.0337 | -0.04161 | -0.16636 | -0.12575 | -0.19012 | -0.05821 | -0.04562 | -0.18652 | -0.06111 | -0.15449 | -0.28751 | -0.0312 | 0.08738 | -0.04775 | 0.31556 | 0.01484 | -0.09724 | 0.103103 | 0.03669 | 0.00698 | 0.086026 | -0.57781 |
| F5 | -0.2678 | 0.025272 | 0.13981 | 0.02439 | 0.02415 | -0.05324 | -0.10445 | -0.18699 | -0.00557 | -0.28107 | 0.009596 | 0.15757 | 0.05734 | 0.17356 | 0.003329 | -0.06391 | -0.11457 | 0.13523 | -0.06276 | 0.0832 | 0.08258 | -0.06834 | 0.09728 | 0.04966 | 0.14174 | 0.142798 | -0.20834 |
| NCIH292 | -0.05417 | 0.019693 | -0.58182 | -0.20967 | -0.09329 | -0.09199 | 0.045242 | -0.18108 | -0.13101 | 0.12814 | -0.01505 | -0.03072 | -0.071 | -0.07694 | -0.08337 | -0.24383 | -0.05354 | 0.14896 | 0.004963 | 0.02414 | -0.09171 | -0.30301 | 0.168912 | 0.10464 | 0.01797 | 0.014619 | -0.45711 |
| HCC2998 | -0.26104 | 0.17253 | -0.36991 | -0.23143 | -0.2269 | -0.1305 | -0.1188 | -0.0799 | -0.27726 | 0.03934 | 0.05317 | -0.17813 | -0.28066 | -0.207 | -0.08803 | -0.15236 | -0.08279 | 0.14806 | -0.17523 | 0.00867 | 0.25481 | -0.17078 | 0.223271 | -0.30136 | 0.15018 | -0.02956 | -0.13668 |
| JR | -0.01947 | -0.17625 | -0.96929 | -0.47278 | -0.13248 | -0.17831 | 0.063539 | -0.44137 | -0.05007 | 0.00787 | -0.12849 | 0.06998 | 0.03628 | -0.06862 | -0.11885 | -0.01568 | -0.13541 | 0.20539 | 0.072718 | 0.04074 | 0.23338 | -0.07716 | -0.01727 | 0.06938 | 0.2666 | 0.004771 | -0.52236 |
| KCIMOH1 | -0.05876 | 0.023503 | -0.68239 | -0.26435 | -0.17245 | -0.02898 | -0.1677 | -0.32879 | 0.004111 | 0.06889 | -0.13166 | 0.2376 | -0.16079 | -0.00596 | 0.051774 | -0.08365 | -0.00502 | -0.11407 | -0.07058 | 0.05507 | 0.29373 | -0.19581 | 0.080559 | -0.06727 | 0.37012 | 0.011044 | -0.28486 |
| KD | -0.0946 | -0.10035 | 0.10508 | -0.09925 | -0.33202 | -0.04551 | 0.18724 | -0.26488 | 0.006779 | 0.12668 | -0.1006 | 0.29251 | -0.35405 | -1.51065 | 0.179084 | -0.16839 | 0.01281 | 0.05622 | 0.031154 | 0.11031 | 0.16573 | 0.008406 | -0.10378 | -0.1099 | 0.15333 | -0.02749 | -0.65064 |
| KP1N | -0.15634 | 0.06724 | -0.31909 | -0.0566 | -0.17016 | -0.12308 | -0.01714 | -0.07026 | 0.005766 | 0.06758 | 0.029216 | 0.11779 | -0.31063 | -0.06802 | -0.18605 | -0.25546 | -0.03268 | 0.08238 | -0.04176 | 0.0201 | 0.00937 | -0.12341 | 0.259083 | 0.05517 | 0.09997 | -0.07241 | -0.21505 |
| MAC2A | 0.11415 | 0.063842 | -0.53489 | -0.00085 | -0.36108 | -0.31498 | 0.071804 | -0.15147 | -0.29643 | -0.16948 | -0.07949 | -0.03331 | -0.17461 | -0.45604 | -0.04562 | -0.15497 | -0.20375 | 0.21627 | -0.08622 | 0.0728 | 0.16975 | -0.1353 | 0.226948 | -0.01252 | 0.48339 | -0.11794 | -0.42182 |
| MOGGUVW | -0.00866 | 0.350416 | 0.02297 | -0.06733 | 0.18943 | 0.07641 | 0.176328 | -0.00508 | -0.11767 | -0.00331 | -1.25269 | 0.16875 | 0.05879 | 0.12158 | -0.00704 | 0.06629 | -0.01961 | 0.24144 | 0.002114 | 0.28661 | 0.13187 | 7.27E-05 | -0.02961 | 0.16286 | 0.01515 | 0.248715 | -0.39427 |
| MON | -0.28575 | -0.15055 | -0.20771 | -0.01904 | -0.17516 | -0.33855 | 0.198333 | -0.5166 | -0.32216 | 0.16862 | -1.04709 | 0.18148 | -0.55382 | -0.43107 | -0.01814 | -0.0927 | -0.10607 | -0.23102 | 0.256071 | 0.34757 | 0.13985 | 0.008766 | -0.09056 | -0.15634 | 0.25722 | 0.279986 | -0.3488 |
| MONOMAC1 | -0.61084 | 0.025039 | -0.35469 | 0.08727 | -0.23904 | -0.48124 | 0.193641 | -0.12577 | -0.05843 | 0.07318 | -0.14451 | 0.13489 | 0.01168 | 0.11402 | 0.150377 | 0.032855 | -0.18118 | 0.45965 | -0.06946 | -0.08929 | 0.16772 | -0.06112 | 0.005566 | 0.10028 | 0.22147 | 0.062498 | -0.15538 |
| MYLA | -0.13131 | 0.018595 | -0.6917 | -0.07193 | -0.10208 | -0.1583 | 0.055376 | -0.24824 | -0.13662 | -0.04024 | 0.011178 | 0.2204 | -0.1523 | -0.25673 | -0.04977 | -0.09205 | -0.18233 | 0.08106 | -0.01417 | 0.00403 | 0.22202 | -0.15433 | 0.148899 | -0.08967 | 0.08513 | 0.005181 | -0.46149 |
| NCIH1993 | -0.18443 | -0.05093 | -0.19735 | -0.17779 | -0.25246 | -0.12896 | 0.008103 | -0.20505 | -0.21103 | -0.02782 | -1.71886 | 0.19273 | -0.24601 | -0.0928 | -0.30964 | -0.222 | -0.05297 | 0.19521 | 0.051268 | -0.07518 | -0.06731 | -0.4036 | -0.02207 | -0.03945 | 0.08764 | -0.06679 | -0.53709 |
| OC316 | -0.20422 | -0.0084 | 0.02287 | -0.15838 | -0.00745 | 0.07806 | -0.03243 | -0.0605 | -0.10761 | 0.04932 | 0.012197 | 0.02804 | -0.29287 | -0.17722 | -0.07125 | -0.16404 | -0.08694 | 0.2378 | 0.024375 | 0.23458 | 0.11985 | -0.12625 | 0.07069 | -0.10678 | -0.00348 | 0.128838 | -0.24314 |
| OVCAR5 | -0.0288 | 0.011202 | -0.50081 | -0.0963 | -0.21115 | -0.04636 | 0.023283 | -0.00092 | -0.11921 | -0.07341 | -0.15523 | 0.1563 | -0.13067 | -0.09728 | -0.12074 | -0.10452 | -0.01299 | -0.02997 | -0.10733 | 0.12141 | 0.10649 | -0.15848 | 0.05007 | -0.03755 | 0.08884 | -0.01032 | -0.34182 |
| CCLFPEDS0001T | -0.0809 | 0.029932 | -0.14735 | -0.03957 | -0.27537 | -0.48687 | -0.03547 | -0.28174 | -0.20875 | -0.10275 | -0.22368 | 0.09599 | -0.22021 | -0.29501 | -0.16462 | -0.31102 | -0.14314 | 0.02521 | -0.06373 | 0.0641 | 0.22116 | -0.04904 | 0.14389 | -0.20362 | 0.1763 | 0.062744 | -0.59277 |
| CCLFPEDS0003T | 0.05013 | 0.076394 | -0.48916 | -0.08362 | 0.02069 | -0.00733 | 0.184672 | -0.23435 | -0.22697 | 0.02676 | -0.02495 | 0.19648 | -0.14218 | 0.24004 | -0.05252 | -0.37803 | -0.0657 | -0.19177 | 0.072206 | 0.23576 | -0.0244 | -0.04118 | 0.041571 | -0.10751 | 0.02216 | 0.195533 | -0.2569 |
| U251MGDM | -0.11829 | 0.042051 | -0.20786 | -0.08833 | -0.14971 | -0.10825 | -0.06588 | -0.18281 | -0.03972 | -0.09551 | -0.09201 | 0.1509 | -0.06772 | -0.19327 | -0.14511 | -0.13818 | -0.13047 | 0.11998 | -0.05029 | 0.13996 | 0.10099 | -0.20229 | 0.105268 | -0.11156 | 0.18945 | -0.0866 | -0.30529 |
| RT11284 | -0.36785 | 0.069039 | -0.34579 | 0.0069 | -0.09354 | -0.27229 | 0.022698 | -0.00402 | -0.10144 | 0.18449 | -0.05705 | 0.13343 | -0.08544 | -0.0989 | 0.060887 | -0.0681 | -0.04456 | 0.07948 | 0.036622 | 0.09918 | 0.1093 | -0.22957 | -0.04302 | -0.10624 | 0.26151 | -0.06379 | -0.58441 |
| SCMCRM2 | -0.1852 | -0.13318 | -0.68428 | -0.15029 | -0.09904 | -0.04402 | 0.044837 | -0.50647 | -0.31366 | -0.04809 | -0.03138 | -0.02943 | -0.52805 | -0.06876 | -0.02981 | 0.00831 | -0.11828 | 0.07352 | -0.01731 | 0.27522 | 0.00991 | -0.08001 | 0.102753 | 0.05312 | 0.05995 | 0.083005 | -0.4672 |
| SHSY5Y | 0.01444 | -0.10788 | -0.47687 | -0.36436 | -0.11955 | -0.0799 | -0.07496 | -0.24484 | 0.05118 | 0.01977 | 0.006113 | 0.20027 | -0.39433 | -0.29734 | -0.05088 | -0.07446 | -0.03693 | -0.00024 | 0.110801 | 0.15334 | 0.25535 | -0.16107 | 0.026894 | 0.14429 | 0.12018 | 0.037745 | -0.48947 |
| SKMEL2 | -0.07776 | 0.152219 | -0.65647 | -0.02057 | -0.13518 | -0.1714 | 0.150914 | -0.09997 | -0.07538 | 0.07602 | -0.06445 | 0.09241 | -0.21654 | 0.04357 | -0.11486 | -0.03023 | -0.14983 | 0.10585 | 0.092851 | 0.09947 | 0.13845 | -0.15872 | 0.015997 | 0.02205 | 0.19849 | 0.081485 | -0.18359 |
| SKNEP1 | -0.06192 | 0.028043 | -1.03274 | -0.05398 | -0.235 | -0.29557 | -0.18789 | 0.05731 | -0.17984 | -0.19908 | -0.06726 | -0.04335 | -0.35169 | -0.33496 | 0.011266 | -0.04811 | 0.03321 | 0.23876 | -0.05003 | 0.10221 | 0.18546 | -0.03901 | 0.111813 | 0.04645 | 0.14887 | 0.107489 | -0.72073 |
| SKPNDW | 0.01776 | 0.210717 | -0.581 | -0.06765 | -0.08082 | -0.01393 | 0.069106 | -0.06089 | 0.104328 | 0.09458 | 0.087691 | 0.12732 | 0.10506 | -0.00731 | 0.035371 | -0.04064 | -0.33271 | 0.12885 | -0.00807 | 0.21569 | 0.20424 | 0.095221 | 0.096193 | 0.05286 | 0.26739 | 0.037218 | -0.22112 |
| SKRC31 | -0.00942 | -0.12816 | -0.31312 | -0.01855 | -0.10069 | -0.11977 | -0.03147 | -0.26968 | -0.25124 | -0.07046 | -0.1108 | 0.09474 | -0.12384 | -0.204 | -0.22742 | -0.00042 | -0.2293 | 0.05938 | -0.08375 | 0.17781 | 0.23041 | -0.25325 | 0.172012 | -0.22157 | 0.01 | -0.1026 | -0.37633 |
| SMSCTR | -0.59314 | -0.02998 | -1.0708 | -0.01604 | -0.00283 | -0.06233 | 0.057552 | -0.29197 | -0.182 | 0.03146 | 0.07454 | 0.17723 | -0.06778 | -0.05326 | -0.0731 | 0.022086 | -0.05349 | 0.00392 | 0.060188 | 0.32982 | -0.00072 | 0.036919 | 0.221357 | 0.01246 | 0.19217 | 0.142225 | -0.30204 |
| SMZ1 | -0.08216 | 0.1384 | -0.10326 | 0.12033 | -0.08809 | -0.0846 | -0.11596 | -0.09116 | -0.18012 | -0.11965 | -0.17975 | 0.15472 | -0.08976 | 0.15988 | -0.41924 | 0.00912 | -0.04719 | -0.07775 | -0.11659 | 0.16293 | 0.26996 | -0.05752 | 0.164033 | -0.0574 | 0.07622 | 0.015919 | -0.18606 |
| TC32 | -0.08596 | 0.088529 | -0.67205 | -0.17386 | -0.12639 | -0.09613 | 0.033549 | 0.03006 | -0.09043 | 0.14574 | -0.0079 | 0.07688 | 0.0729 | -0.10957 | -0.06335 | -0.11982 | -0.18299 | 0.16484 | 0.085453 | 0.16712 | 0.02931 | -0.48096 | 0.005851 | 0.02204 | 0.21981 | 0.038053 | -0.45656 |
| TTC549 | -0.19618 | 0.184601 | -0.6665 | -0.16697 | -0.12398 | 0.09953 | 0.214447 | -0.53635 | -0.07623 | -0.02893 | -0.41526 | 0.31751 | -0.18566 | -0.28444 | -0.09486 | -0.03052 | 0.04718 | -0.00812 | -0.10441 | 0.18043 | 0.2848 | -0.24632 | 0.119795 | -0.016 | 0.26019 | 0.100674 | -0.521 |
| TTC642 | -0.03788 | -0.14987 | -0.07478 | -0.22562 | -0.08504 | -0.30998 | -0.00323 | -0.08384 | -0.21311 | 0.18383 | -0.19719 | 0.10179 | -0.23605 | 0.10657 | 0.145772 | -0.21831 | 0.06243 | 0.37456 | 0.072149 | 0.2664 | 0.2859 | 0.057981 | -0.12133 | 0.29002 | 0.14671 | 0.14921 | -0.46976 |
| UPCISCC152 | -0.32258 | -0.11763 | -0.26494 | -0.03838 | -0.21615 | -0.26558 | 0.020601 | -0.02861 | -0.31843 | 0.16639 | 0.021448 | 0.09276 | -0.26522 | 0.03425 | -0.00104 | -0.04731 | -0.02694 | 0.1945 | -0.24495 | 0.07589 | 0.01931 | -0.29109 | 0.0848 | -0.01312 | 0.07769 | -0.05239 | -0.3149 |
| UPCISCC154 | -0.17047 | 0.004466 | -0.01722 | -0.03666 | -0.25207 | -0.09411 | -0.14202 | -0.19132 | -0.16575 | -0.16195 | -0.12997 | 0.20884 | -0.23801 | -0.17468 | -0.1556 | -0.14034 | 0.06919 | 0.0936 | 0.017124 | 0.13517 | 0.12534 | -0.31835 | 0.072087 | -0.09623 | 0.14476 | 0.090305 | -0.39465 |
| UW228 | -0.0991 | -0.03401 | -0.08487 | -0.04792 | -0.13741 | -0.08681 | -0.09349 | -0.15532 | -0.20643 | 0.04473 | 0.082305 | 0.15646 | -0.12061 | -0.35955 | -0.02424 | -0.10802 | -0.15022 | 0.00156 | -0.06514 | 0.20071 | 0.20019 | 0.008295 | 0.081677 | -0.11015 | 0.25651 | -0.05453 | -0.38496 |
| VMRCLCD | -0.11223 | -0.03218 | 0.14132 | 0.03364 | -0.01402 | -0.02989 | 0.086955 | -0.34293 | -0.11845 | -0.09072 | -0.33659 | 0.24421 | -0.1324 | -0.14312 | -0.11121 | -0.11017 | -0.08011 | 0.10765 | 0.036921 | 0.02836 | 0.051 | -0.20873 | -0.00081 | -0.09404 | 0.16196 | 0.095953 | -0.2702 |
| WM2664 | -0.07659 | 0.000821 | -0.18037 | -0.0526 | -0.02563 | -0.20528 | 0.046295 | -0.15532 | -0.22169 | 0.01606 | -0.03208 | 0.18936 | -0.12832 | -0.12669 | -0.06054 | -0.1222 | -0.06063 | 0.096 | 0.070695 | 0.1476 | 0.20355 | -0.16987 | 0.12908 | 0.01379 | 0.14191 | 0.008967 | -0.46159 |
| 127399 | -0.11761 | 0.154244 | -0.36889 | 0.03003 | -0.10252 | -0.31371 | 0.000804 | -0.13932 | -0.08551 | 0.02032 | 0.042127 | 0.25656 | -0.27888 | -0.28526 | 0.052623 | -0.39435 | -0.25789 | 0.08257 | -0.016 | 0.05461 | 0.23312 | -0.23767 | 0.03943 | 0.04401 | 0.34566 | 0.135895 | -0.17643 |
| SW982 | 0.0576 | 0.080663 | -0.16179 | 0.0297 | -0.10499 | 0.05118 | -0.20662 | -0.21019 | -0.32452 | 0.26755 | -0.0656 | 0.1164 | -0.03331 | -0.23851 | 0.025481 | -0.10835 | -0.2668 | 0.13585 | -0.10332 | 0.08346 | 0.04776 | -0.16854 | -0.026 | -0.04461 | 0.35573 | 0.094571 | -0.29951 |
| SYO1 | -0.33071 | -0.00059 | -0.02136 | -0.12644 | -0.13057 | -0.14869 | -0.06378 | -0.27282 | -0.05774 | 0.15447 | 0.050387 | 0.12426 | -0.52945 | -0.01641 | -0.05898 | -0.27069 | 0.04931 | 0.05214 | -0.06791 | 0.0418 | 0.0114 | -0.11063 | 0.039486 | 0.01125 | 0.21184 | -0.0312 | -0.20468 |
| YAMATO | -0.43903 | 0.170392 | -0.22334 | -0.05836 | -0.1768 | 0.06104 | 0.226007 | -0.16701 | -0.01117 | -0.07848 | -0.39083 | 0.01978 | 0.06117 | 0.01595 | 0.088616 | -0.19543 | -0.03553 | -0.28561 | -0.17952 | 0.20983 | 0.24962 | -0.12731 | 0.219465 | -0.0606 | 0.02526 | 0.112598 | -0.40041 |
| BIN67 | -0.147 | -0.08435 | -0.07354 | -0.12071 | -0.13975 | -0.10281 | -0.06153 | -0.11284 | -0.07998 | -0.09176 | -0.08537 | 0.08753 | -0.03323 | -0.05473 | -0.12483 | -0.2239 | -0.04168 | -0.06786 | -0.06208 | 0.05919 | 0.11736 | -0.13372 | 0.046514 | 0.10648 | 0.10839 | -0.0516 | -0.28691 |
| SCCOHT1 | -0.10217 | 0.087573 | -0.12593 | -0.09583 | 0.05425 | 0.13826 | -0.04163 | -0.30286 | -0.21171 | 0.00604 | -1.09901 | 0.04544 | -0.16403 | -0.09306 | 0.00921 | -0.07957 | -0.08116 | 0.18958 | -0.0172 | 0.31136 | 0.10073 | -0.14815 | -0.06836 | -0.06037 | -0.03754 | 0.194603 | -0.48299 |
| SCS214 | -0.04134 | -0.16685 | -0.29203 | 0.02828 | -0.12448 | -0.04193 | -0.15993 | -0.07975 | 0.005702 | -0.25642 | -0.16332 | 0.14894 | -0.48093 | 0.16102 | -0.12366 | 0.015057 | -0.30504 | 0.16218 | 0.01768 | 0.09001 | 0.05637 | -0.15102 | 0.175568 | -0.1079 | 0.33606 | 0.144553 | -0.28781 |
| TC106 | -0.25591 | 0.114747 | -0.97197 | -0.14734 | -0.09151 | -0.16006 | 0.076124 | -0.14936 | -0.04708 | -0.07437 | -0.2053 | -0.00966 | -0.2758 | -0.50434 | -0.09788 | -0.08557 | -0.11783 | 0.17401 | -0.11807 | 0.07997 | 0.06964 | -0.06195 | 0.123949 | -0.15086 | 0.15126 | 0.012492 | -0.6768 |
| COGAR359 | -0.0994 | 0.131649 | -0.41739 | -0.09174 | -0.07859 | 0.07268 | 0.122919 | -0.26906 | -0.0872 | -0.04522 | -0.10987 | 0.1058 | -0.1034 | 0.02834 | -0.01648 | 0.12621 | -0.21217 | 0.04072 | 0.219634 | -0.01891 | 0.05722 | -0.02832 | 0.183961 | -0.07839 | 0.24013 | 0.016784 | -0.36181 |
| Y79 | -0.14977 | 0.02818 | -0.55311 | -0.08655 | -0.07604 | -0.01793 | 0.067742 | -0.28425 | -0.06837 | -0.00257 | -0.01235 | 0.07498 | -0.14787 | -0.1334 | -0.10432 | -0.0747 | -0.08548 | 0.16429 | -0.03732 | 0.16326 | 0.09687 | -0.05328 | 0.037619 | -0.11769 | 0.1136 | -0.05772 | -0.61029 |
| CHLA15 | -0.15143 | 0.003029 | -0.11435 | -1.12689 | -0.28628 | -0.06931 | -0.04209 | -0.36046 | -0.03436 | -0.12138 | 0.006547 | 0.07517 | -0.28292 | -0.28764 | -0.05396 | -0.20733 | -0.18769 | 0.02902 | 0.022486 | 0.32024 | 0.05276 | -0.17226 | 0.040747 | -0.19033 | 0.17269 | 0.057034 | -0.39456 |
| COGN278 | -0.1352 | 0.021675 | 0.16671 | -0.29479 | -0.1593 | -0.21173 | 0.036415 | -0.19611 | -0.21854 | 0.11907 | -0.08495 | -0.07051 | -0.25165 | 0.01626 | 0.029807 | -0.10813 | -0.08468 | 0.31383 | 0.058799 | 0.18413 | 0.06113 | -0.59139 | -0.00746 | 0.09501 | 0.26772 | 0.184387 | -0.76502 |
| COGN305 | -0.01964 | 0.040298 | -0.03491 | -0.9004 | -0.13287 | -0.27118 | 0.056388 | -0.15674 | 0.037209 | -0.16613 | 0.372455 | 0.10436 | -0.03327 | 0.09693 | -0.18486 | -0.06861 | -0.30211 | -0.0011 | -0.05601 | 0.37623 | -0.06917 | 0.056222 | 0.210045 | 0.0131 | 0.0736 | 0.109281 | -0.34051 |
| NB1643 | -0.19169 | -0.13736 | 0.03518 | -0.09413 | 0.06945 | -0.15662 | -0.01424 | -0.1199 | -0.26937 | 0.18187 | -0.0355 | 0.04057 | -0.14901 | 0.07047 | 0.01699 | 0.017256 | -0.05661 | 0.37145 | -0.17872 | 0.15123 | -0.23774 | -0.34613 | -0.03649 | 0.03562 | -0.02583 | 0.088436 | -0.70927 |
| 8305C | -0.0467 | 0.047234 | -0.4061 | -0.2346 | -0.02349 | -0.29702 | -0.09022 | -0.06271 | 0.105163 | -0.14746 | -0.09303 | 0.23144 | -0.3166 | 0.02268 | -0.0484 | -0.0544 | -0.08436 | -0.16771 | -0.03786 | 0.20815 | 0.12117 | 0.014762 | 0.125228 | 0.03998 | 0.10582 | 0.101973 | -0.37061 |
| 8505C | -0.13594 | 0.075587 | -0.01207 | -0.09899 | -0.3042 | -0.21068 | -0.31672 | -0.08761 | 0.09326 | 0.06719 | -0.11291 | -0.01842 | -0.22421 | -0.07597 | -0.10437 | -0.1982 | -0.046 | 0.25843 | 0.019615 | 0.3021 | 0.09998 | -0.25024 | 0.115441 | -0.21509 | 0.1224 | 0.10041 | -0.2804 |
| HA1E | 0.10066 | 0.07106 | -0.0496 | -0.08275 | -0.09789 | -0.0508 | -0.08882 | -0.33062 | -0.01687 | 0.08724 | -0.14822 | 0.11572 | -0.09347 | -0.51129 | -0.12049 | -0.20152 | 0.03357 | 0.18301 | -0.12166 | 0.24784 | 0.09766 | -0.36634 | -0.06566 | 0.00171 | 0.06078 | 0.096716 | -0.29656 |
| PLCPRF5 | -0.14605 | -0.0073 | -0.09218 | -0.03358 | -0.1326 | -0.12267 | 0.079417 | -0.25398 | -0.22517 | 0.0245 | -0.06002 | 0.04875 | -0.26661 | -0.22167 | -0.15322 | -0.19483 | -0.07364 | 0.17248 | -0.01217 | 0.07991 | 0.10489 | -0.25388 | 0.007969 | -0.04116 | -0.07391 | 0.147112 | -0.57119 |
| CME1 | -0.29649 | -0.0127 | -0.19396 | -0.00819 | 0.02531 | -0.25631 | -0.23628 | 0.03578 | -0.34561 | -0.21917 | 0.042278 | 0.09196 | -0.63306 | 0.14026 | 0.062504 | -0.04197 | -0.06757 | 0.0045 | -0.12722 | 0.01319 | 0.16908 | -0.6369 | 0.049889 | -0.0739 | 0.04201 | -0.04583 | -0.48882 |
| A431 | -0.15335 | 0.02099 | -0.46573 | -0.1392 | -0.29639 | -0.26963 | -0.1184 | -0.37882 | 0.009378 | -0.02331 | -0.09583 | 0.22114 | -0.36542 | -0.16873 | -0.08783 | -0.03238 | -0.19045 | 0.1182 | -0.06277 | 0.04583 | 0.10667 | -0.29068 | 0.053938 | -0.10482 | 0.07191 | 0.011699 | -0.32619 |
| ANGMCSS | -0.03972 | -0.01918 | -0.0982 | -0.06434 | -0.08829 | -0.20032 | 0.042536 | -0.1449 | -0.11907 | -0.05331 | -0.09553 | 0.08828 | -0.32629 | -0.38417 | -0.06979 | -0.03563 | -0.12491 | 0.06666 | 0.017201 | 0.10114 | 0.12991 | -0.0976 | 0.156332 | -0.08635 | 0.13529 | 0.058549 | -0.38536 |
| BICR10 | -0.12474 | 0.015006 | 0.01797 | -0.05384 | -0.07447 | -0.26165 | -0.07315 | -0.18295 | -0.20764 | -0.18836 | -0.09139 | 0.09345 | -0.22723 | -0.05231 | 0.041223 | -0.18549 | 0.01041 | 0.04559 | -0.04165 | -0.00926 | 0.15208 | -0.33477 | -0.10207 | 0.15199 | 0.12686 | -0.03727 | -0.25523 |
| BICR78 | -0.23341 | 0.343706 | 0.13741 | 0.06088 | -0.00575 | -0.26754 | -0.06234 | 0.11018 | -0.37811 | 0.07287 | 0.104139 | 0.11907 | -0.16156 | 0.21062 | -0.2752 | 0.03723 | -0.01813 | 0.06238 | -0.04904 | 0.04324 | -0.01154 | -0.08642 | 0.478495 | -0.11581 | 0.11792 | 0.114767 | -0.55327 |
| C33A | -0.21356 | -0.00931 | -0.0645 | -0.07768 | 0.04766 | -0.19671 | -0.08503 | -0.22493 | -0.13576 | -0.0742 | -0.02617 | 0.05956 | -0.57604 | 0.16461 | -0.04097 | -0.27934 | -0.37109 | -0.02107 | 0.040668 | -0.06801 | 0.02011 | -0.17654 | 0.144577 | 0.04675 | 0.12988 | -0.01838 | -0.49942 |
| C4I | -0.15207 | -0.04199 | -0.09173 | -0.09097 | -0.17825 | 0.01476 | -0.01933 | -0.14484 | 0.10347 | -0.23523 | -0.16278 | 0.12126 | -0.26388 | -0.22819 | -0.13444 | -0.0729 | -0.03533 | 0.01258 | -0.05217 | 0.12436 | 0.09355 | -0.15994 | 0.081066 | -0.14023 | 0.10591 | 0.064831 | -0.28906 |
| C4II | -0.04648 | -0.02803 | -0.12849 | -0.05292 | -0.08203 | -0.26762 | -0.03777 | -0.40978 | -0.05148 | 0.1708 | 0.002498 | 0.09379 | -0.10384 | -0.09412 | -0.16317 | -0.24207 | -0.02914 | 0.09707 | 0.057396 | 0.18385 | 0.12174 | -0.13503 | 0.133263 | 0.00687 | 0.18048 | -0.04242 | -0.37685 |
| CASKI | -0.32981 | -0.08172 | -0.00165 | -0.01986 | -0.18849 | -0.18521 | -0.10437 | -0.18874 | 0.063388 | -0.08871 | -0.08304 | 0.07689 | -0.2443 | 0.03154 | -0.05581 | -0.14181 | -0.02671 | 0.15548 | -0.05502 | 0.07781 | 0.014 | -0.17957 | 0.05618 | -0.19341 | 0.03544 | -0.09047 | -0.39129 |
| CHP134 | -0.05897 | -0.03584 | -0.30186 | -0.01244 | -0.32787 | -0.02978 | -0.25023 | -0.49373 | -0.04909 | -0.18471 | -0.13544 | 0.1523 | -0.05982 | -0.36189 | -0.12009 | -0.10444 | -0.02512 | -0.0055 | -0.20844 | 0.18854 | 0.26018 | -0.30142 | 0.106062 | 0.03775 | 0.10804 | 0.067628 | -0.35784 |
| COV413A | -0.18635 | 0.023794 | -0.11929 | -0.10408 | -0.11578 | -0.14841 | 0.02057 | -0.12922 | 0.051005 | -0.02042 | -0.12527 | 0.16174 | -0.06179 | -0.10093 | -0.11622 | 0.007697 | -0.0016 | 0.08908 | -0.05243 | 0.13174 | 0.14712 | -0.17595 | 0.103639 | -0.14914 | 0.08237 | 0.070249 | -0.36044 |
| DOTC24510 | -0.1561 | -0.01119 | -0.15042 | -0.08952 | -0.14105 | -0.12318 | -0.04362 | -0.13928 | -0.21449 | -0.05368 | -0.17609 | 0.11529 | -0.25317 | -0.14583 | -0.10948 | -0.03859 | -0.05958 | 0.00839 | -0.06831 | 0.1693 | 0.22034 | -0.18275 | 0.070375 | -0.02791 | 0.21214 | -0.02016 | -0.34457 |
| GIMEN | 0.41762 | 0.022636 | -0.28003 | -0.41869 | -0.22945 | 0.14715 | -0.12087 | 0.00131 | -0.71923 | -0.10869 | -0.16778 | 0.09637 | -0.0051 | -0.11332 | 0.050191 | -0.00028 | 0.10497 | 0.20115 | -0.28334 | 0.54182 | 0.24521 | 0.014554 | 0.258623 | 0.04294 | -0.08143 | 0.11129 | -0.33068 |
| GP5D | -0.04949 | 0.017498 | -0.18687 | -0.19301 | -0.11811 | -0.27883 | -0.06132 | -0.30482 | -0.18462 | 0.0331 | 0.051834 | 0.18133 | -0.24 | 0.09541 | -0.0552 | -2.3E-05 | -0.1598 | 0.15962 | -0.01139 | 0.09476 | 0.15655 | 0.016114 | -0.0183 | 0.00853 | 0.08879 | 0.044562 | -0.42814 |

| Cell line | FLT3 | FLT4 | IGF1R | INSR | INSRR | KDR | KIT | LMR1 | LMR2 | LTK | MET | MUSK | PDGFRA | PDGFRB | PTK7 | RET | RON | ROR1 | ROR2 | ROS1 | STYK1 | TIE1 | TIE2 | TRKA | TRKB | TRKC | TYRO3 |
|---|---|---|---|---|---|---|---|---|---|---|---|---|---|---|---|---|---|---|---|---|---|---|---|---|---|---|---|
| H103 | -0.26567 | 0.020092 | -0.29392 | -0.01376 | -0.18177 | -0.15864 | -0.07755 | -0.09385 | -0.06766 | -0.13354 | -0.06958 | 0.2063 | -0.41545 | -0.02522 | 0.031755 | -0.09412 | -0.04779 | 0.18572 | 0.020331 | 0.11305 | 0.03955 | -0.21704 | 0.106882 | -0.02055 | 0.11209 | 0.031423 | -0.30379 |
| H157 | -0.02185 | -0.01556 | -0.23331 | -0.12305 | -0.09885 | -0.22009 | -0.05772 | -0.20573 | -0.20608 | -0.051 | -0.02904 | 0.05431 | -0.19944 | -0.18146 | -0.15156 | -0.12951 | 0.02522 | 0.15284 | -0.08649 | 0.14381 | 0.125 | -0.3623 | 0.128327 | 0.00083 | 0.10673 | -0.11766 | -0.4221 |
| JOPACA1 | -0.07917 | -0.05291 | -0.52086 | -0.05789 | -0.1575 | -0.14691 | -0.08545 | -0.08774 | 0.00091 | -0.02252 | -0.11451 | 0.10709 | -0.07098 | -0.23278 | -0.09259 | -0.03191 | 0.00308 | -0.03253 | 0.013573 | -0.01358 | 0.08196 | -0.17362 | 0.033021 | -0.13654 | 0.07761 | 0.031306 | -0.48972 |
| LAN2 | -0.06187 | -0.06538 | -0.45911 | -0.35546 | -0.27765 | -0.24254 | -0.00771 | -0.26628 | -0.01097 | 0.04986 | 0.16713 | 0.1182 | -0.11031 | -0.31221 | -0.01935 | -0.34203 | -0.05703 | 0.28604 | -0.25493 | 0.30745 | 0.11135 | -0.23784 | -0.21274 | -0.01733 | 0.12969 | 0.099791 | -0.78462 |
| MB1 | 0.013 | -0.05861 | 0.00369 | -0.10054 | -0.06129 | -0.16432 | -0.04738 | -0.00341 | -0.26611 | -0.15562 | -0.06028 | 0.10795 | -0.18643 | -0.19128 | -0.07175 | -0.10009 | -0.06146 | 0.18508 | -0.04749 | -0.09245 | 0.07088 | -0.22404 | 0.152126 | 0.02765 | 0.14539 | -0.03491 | -0.29064 |
| MS751 | -0.20138 | 0.069536 | -0.16876 | -0.092 | -0.2099 | -0.19764 | -0.18948 | -0.197 | 0.060678 | -0.0041 | 0.004828 | 0.17852 | -0.2166 | -0.31449 | -0.1803 | -0.10541 | 0.00568 | 0.08736 | -0.04148 | 0.1822 | 0.04452 | -0.18686 | 0.098878 | -0.04264 | 0.11464 | -0.06034 | -0.26295 |
| NGP | -0.20787 | -0.05387 | -0.62574 | -0.26891 | 0.01671 | -0.09199 | -0.09234 | -0.20393 | -0.13556 | 0.23931 | -0.127 | 0.1097 | -0.28605 | -0.14249 | 0.082664 | -0.09831 | 0.08804 | 0.12317 | 0.132985 | 0.21659 | 0.14959 | -0.28628 | 0.008769 | 0.10136 | 0.16106 | -0.02901 | -0.66411 |
| NMB | -0.14734 | 0.066525 | -0.53152 | 0.03069 | -0.28854 | -0.26419 | 0.062544 | -0.0275 | -0.22577 | 0.12116 | -0.03892 | -0.05347 | -0.47871 | 0.00153 | -0.13077 | -0.22918 | -0.11506 | 0.14363 | -0.17805 | 0.07241 | 0.11261 | -0.02107 | 0.066363 | 0.0732 | 0.34702 | 0.018609 | -0.41405 |
| OACM51 | -0.08751 | 0.024282 | -0.75676 | 0.01803 | -0.10218 | -0.11749 | -0.08217 | -0.15845 | -0.23452 | -0.00266 | -0.0156 | 0.13151 | -1.0715 | -0.16295 | -0.16067 | -0.03901 | -0.16292 | 0.04526 | -0.03693 | 0.10797 | 0.09079 | -0.07879 | 0.129211 | -0.04615 | 0.12621 | -0.00653 | -0.41011 |
| OCIP5X | -0.07125 | -0.10218 | 0.00835 | 0.01443 | -0.28134 | -0.39157 | 0.005402 | -0.07076 | -0.07919 | 0.09238 | 0.023473 | 0.04205 | -0.20174 | 0.05324 | -0.02979 | 0.127688 | 0.13596 | 0.14231 | -0.04351 | 0.30345 | 0.12689 | -0.06576 | 0.227459 | -0.10194 | 0.05953 | -0.04445 | -0.25133 |
| PA1 | -0.21218 | -0.16873 | -0.26968 | -0.27625 | -0.47536 | -0.18185 | -0.02878 | -0.21817 | -0.43683 | -0.0091 | -0.36567 | 0.20983 | -0.31353 | 0.16553 | -0.1074 | -0.47504 | -0.07016 | 0.28953 | 0.712384 | -0.09167 | 0.28544 | -0.12552 | 0.11861 | 0.11542 | 0.19506 | 0.194957 | -0.37179 |
| PACADD119 | -0.17891 | 0.050897 | -0.51568 | -0.09758 | -0.20248 | -0.26734 | 0.082025 | -0.09698 | 0.001246 | 0.02161 | -0.1259 | 0.16673 | -0.0623 | -0.05081 | -0.01875 | 0.034336 | -0.0642 | 0.06725 | 0.170077 | 0.19959 | 0.31412 | 0.068065 | 0.090008 | -0.09716 | 0.25487 | 0.02992 | -0.15365 |
| PACADD137 | -0.14712 | -0.05524 | -0.11723 | -0.43769 | 0.05338 | -0.14677 | 0.064396 | -0.13565 | 0.079738 | 0.12541 | -0.07743 | 0.10093 | -0.06477 | -0.05412 | -0.06544 | 0.044535 | -0.07988 | -0.02263 | 0.106382 | 0.20461 | 0.16363 | -0.11901 | 0.068178 | -0.13131 | -0.05135 | -0.00743 | -0.5376 |
| PACADD161 | -0.14219 | 0.126389 | -0.59851 | -0.1107 | -0.1917 | -0.24343 | -0.1059 | -0.24225 | -0.15815 | 0.0879 | 0.017724 | 0.17387 | -0.16855 | -0.13008 | -0.02757 | -0.14246 | 0.04828 | 0.07834 | -0.04617 | 0.12078 | 0.02665 | -0.15445 | 0.194102 | -0.05216 | 0.19913 | 0.003385 | -0.45896 |
| PACADD165 | -0.24894 | -0.03493 | -0.56184 | -0.14058 | -0.10742 | -0.10696 | -0.00463 | -0.10542 | -0.14307 | -0.06373 | -0.05686 | 0.06197 | -0.1117 | 0.08348 | -0.02477 | -0.0678 | -0.01285 | 0.081 | 0.075731 | 0.09766 | 0.08895 | -0.27476 | 0.061383 | -0.11458 | 0.1353 | -0.04587 | -0.37641 |
| PACADD188 | -0.15002 | -0.0721 | -0.0448 | -0.54288 | -0.35865 | -0.09033 | 0.237391 | -0.11975 | -0.36158 | 0.13155 | -0.10787 | -0.02213 | -0.01031 | 0.23812 | 0.031357 | -0.16204 | -0.39451 | 0.10128 | 0.007891 | 0.11537 | 0.23161 | -0.01628 | 0.086546 | 0.23359 | 0.54637 | 0.018581 | -0.35313 |
| RPMI2650 | -0.08096 | 0.046597 | -0.25788 | -0.13895 | 0.02424 | -0.03386 | 0.011908 | -0.13504 | -0.26679 | 0.04591 | -0.06754 | 0.11794 | -0.13408 | -0.1185 | -0.02622 | -0.0162 | -0.01348 | -0.04239 | 0.058498 | 0.27277 | 0.13139 | -0.21895 | 0.121643 | -0.03464 | -0.0099 | -0.06715 | -0.39428 |
| SCLC22H | -0.17762 | -0.00963 | -0.71877 | -0.28657 | -0.0985 | -0.14154 | -0.09507 | -0.13929 | -0.16631 | 0.02401 | -0.0989 | 0.16349 | -0.18042 | -0.21151 | -0.01621 | -0.05273 | -0.02797 | -0.11337 | -0.01853 | 0.18215 | 0.23205 | -0.19308 | 0.260196 | -0.02113 | 0.22722 | 0.061302 | -0.3077 |
| SUM102PT | -0.11682 | 0.032159 | -0.21917 | -0.09443 | -0.12475 | -0.19258 | -0.02556 | -0.13375 | 0.147619 | -0.00087 | -0.05206 | 0.12249 | -0.20626 | -0.14284 | 0.038014 | -0.02165 | 0.00851 | 0.06242 | 0.028668 | 0.1925 | 0.09981 | -0.18707 | 0.101421 | -0.02566 | 0.17823 | 0.000775 | -0.32703 |
| SUM1315MO2 | -0.05532 | 0.073357 | -0.16828 | -0.24059 | -0.16169 | -0.0422 | -0.0545 | -0.26198 | 0.030008 | -0.12008 | -0.2154 | 0.12756 | -0.28007 | -0.17231 | -0.07969 | -0.12797 | -0.18508 | 0.00574 | 0.052953 | 0.05787 | 0.26168 | -0.28735 | 0.075015 | -0.15391 | 0.1445 | 0.052966 | -0.3479 |
| SUM149PT | 0.14068 | 0.30985 | -0.37577 | -0.03845 | 0.15558 | -0.10305 | 0.272543 | -0.16933 | 0.040511 | 0.06477 | -0.09657 | 0.18681 | -0.21513 | 0.05246 | -0.36126 | 0.107813 | 0.05109 | 0.3899 | 0.106665 | 0.18136 | 0.10163 | -0.11163 | 0.3936 | -0.0508 | 0.09227 | 0.101544 | -0.18501 |
| SUM159PT | 0.10158 | 0.024091 | -0.38921 | -0.09766 | -0.34444 | -0.16164 | -0.12026 | -0.10969 | -0.05174 | -0.12253 | -0.0389 | 0.06862 | -0.30012 | -0.14597 | -0.16291 | -0.12858 | -0.12619 | 0.21269 | -0.21643 | 0.08531 | 0.12353 | -0.30632 | 0.074657 | 0.02543 | 0.10688 | -0.05714 | -0.35705 |
| SUM185PE | -0.18334 | 0.031391 | -0.2233 | -0.07137 | -0.19245 | -0.1884 | 0.313338 | -0.08198 | -0.33379 | -0.03056 | 0.072649 | 0.19204 | -0.39648 | 0.16357 | -0.12037 | -0.37323 | 0.03721 | 0.03564 | 0.078489 | -0.17357 | 0.02405 | 0.022937 | 0.171451 | 0.16077 | 0.25257 | 0.199705 | -0.12361 |
| SUM190PT | -0.15454 | -0.00099 | -0.08164 | -0.19204 | -0.12453 | -0.09563 | -0.19275 | -0.26234 | -0.20586 | 0.06636 | -0.19883 | 0.1688 | -0.34329 | -0.123 | -0.13975 | -0.34329 | -0.10695 | 0.03263 | -0.04265 | 0.23326 | 0.12471 | -0.17297 | 0.042175 | -0.06954 | 0.06891 | 0.052683 | -0.31206 |
| SUM229PE | 0.01984 | 0.078205 | -0.61842 | -0.04076 | -0.0448 | -0.3569 | -0.05476 | -0.27354 | 0.004151 | -0.01573 | -0.14319 | 0.05961 | 0.02869 | -0.1824 | 0.01014 | -0.27768 | 0.05668 | 0.20715 | 0.127289 | -0.04054 | 0.41089 | 0.081107 | 0.037002 | -0.1513 | 0.26636 | 0.178036 | -0.37219 |
| SUM52PE | -0.16933 | 0.230176 | -0.29831 | -0.0661 | -0.14864 | -0.13048 | -0.12557 | -0.44838 | 0.068873 | -0.01473 | -0.08658 | 0.17646 | -0.26276 | 0.0237 | -0.09575 | -0.2732 | -0.06087 | 0.42604 | -0.12548 | 0.19522 | -0.11873 | -0.28745 | 0.309668 | -0.17537 | 0.17312 | -0.26047 | -0.30246 |
| SW156 | 0.0706 | 0.061656 | -0.23599 | -0.26816 | -0.26393 | -0.18841 | -0.24936 | -0.3008 | 0.071742 | -0.00617 | -0.12419 | 0.204 | -0.10606 | -0.20953 | 0.091592 | -0.20147 | -0.1626 | -0.12188 | -0.13184 | 0.197 | 0.17996 | -0.34819 | -0.18791 | -0.13486 | 0.31724 | 0.121415 | -0.14115 |
| SW626 | -0.23537 | -0.14985 | -0.58998 | -0.23632 | -0.29042 | -0.20565 | 0.042442 | -0.13427 | -0.22224 | 0.0255 | -0.072 | 0.31079 | -0.16437 | -0.24476 | 0.004047 | -0.17207 | -0.01962 | 0.10148 | 0.067267 | 0.21042 | 0.01996 | -0.20271 | 0.125845 | -0.01442 | 0.09219 | -0.0869 | -0.36644 |
| SW954 | -0.16836 | 0.058226 | -0.15269 | 0.11736 | -0.11012 | -0.34495 | -0.05964 | -0.29387 | 0.056111 | -0.0366 | -0.12169 | 0.21664 | -0.29961 | -0.30692 | -0.2591 | 0.011192 | -0.0861 | -0.11297 | 0.019946 | 0.19806 | 0.34503 | -0.0415 | -0.06028 | -0.10417 | 0.1113 | -0.01583 | -0.26052 |
| SW756 | -0.08861 | -0.02676 | -0.30366 | -0.07145 | -0.21572 | -0.18158 | -0.08194 | -0.15647 | -0.07739 | 0.0143 | 0.021132 | 0.05754 | -0.22151 | -0.00116 | -0.12856 | -0.14632 | -0.25713 | 0.20466 | -0.06653 | 0.14023 | 0.07276 | -0.13192 | 0.12672 | -0.06922 | 0.1319 | 0.051925 | -0.60378 |
| TO14 | -0.16107 | -0.07315 | -0.33229 | -0.17629 | -0.16501 | -0.20714 | -0.11071 | -0.22621 | -0.28727 | 0.0198 | -0.12197 | 0.18492 | -0.25254 | -0.01422 | -0.0924 | -0.00699 | -0.08032 | 0.06368 | -0.07192 | 0.20505 | 0.04662 | -0.16974 | 0.11196 | -0.01366 | 0.17336 | 0.042506 | -0.27205 |
| UMUC13 | 0.01459 | 0.053422 | -0.22565 | -0.33185 | -0.10793 | -0.24033 | -0.04876 | -0.12295 | -0.03322 | 0.14475 | -0.23816 | 0.1059 | -0.11789 | 0.02835 | -0.09094 | -0.06434 | -0.11257 | 0.18846 | -0.02312 | 0.14252 | 0.1238 | -0.19284 | 0.125643 | -0.02523 | 0.02473 | -0.05216 | -0.25706 |
| UMUC14 | -0.079 | 0.045579 | -0.59047 | -0.07122 | -0.18895 | -0.10861 | 0.193989 | -0.26303 | -0.24689 | -0.10483 | 0.061732 | 0.13485 | -0.19175 | -0.30624 | -0.17217 | -0.00138 | -0.15738 | 0.10808 | -0.11785 | 0.07126 | 0.03344 | -0.30914 | 0.070738 | 0.10059 | 0.09978 | 0.019643 | -0.29131 |
| UMUC16 | -0.22312 | 0.023408 | -0.11577 | -0.0859 | -0.02084 | -0.27231 | 0.083645 | -0.06323 | -0.10047 | -0.16105 | -0.228 | 0.1313 | -0.16665 | -0.12984 | 0.012155 | -0.19222 | -0.18815 | 0.31709 | 0.075956 | -0.01785 | 0.05082 | -0.38712 | -0.00233 | 0.1332 | -0.05398 | 0.147127 | -0.64369 |
| UMUC5 | -0.06568 | 0.151252 | -0.20012 | -0.02053 | -0.17223 | -0.40113 | -0.0861 | -0.24776 | -0.22569 | 0.0437 | 0.041673 | 0.11711 | -0.11823 | -0.13014 | -0.0671 | -0.36612 | -0.08154 | -0.05055 | -0.14961 | 0.0738 | 0.0062 | -0.02471 | 0.201698 | -0.0189 | 0.2297 | -0.14734 | -0.36923 |
| UMUC10 | -0.1854 | 0.00545 | -0.15486 | -0.11348 | -0.13593 | -0.10582 | -0.07838 | -0.11564 | -0.113 | 0.03563 | -0.03561 | 0.18332 | -0.16183 | -0.08955 | -0.05918 | -0.08004 | -0.14621 | 0.01631 | -0.18776 | 0.17813 | 0.07335 | -0.11524 | 0.108578 | -0.13343 | 0.1736 | -0.01503 | -0.24057 |
| UMUC11 | -0.02386 | 0.014583 | -0.08172 | -0.01561 | -0.10372 | -0.00798 | 0.180135 | -0.14804 | -0.06257 | 0.15814 | 0.04243 | 0.0823 | -0.30836 | -0.09998 | -0.10924 | 0.02319 | -0.12225 | 0.14482 | 0.075924 | 0.28624 | 0.03786 | -0.12078 | 0.03564 | -0.07981 | 0.27592 | 0.018819 | -0.25335 |
| UMUC6 | -0.16047 | 0.077265 | -0.46656 | -0.06035 | 0.02821 | -0.02681 | -0.13948 | -0.12483 | -0.12601 | -0.00226 | -0.00108 | 0.0531 | -0.18554 | -0.11156 | -0.02488 | -0.12764 | 0.03492 | 0.08331 | 0.095152 | 0.09914 | 0.02782 | -0.33146 | 0.098414 | -0.17568 | 0.06873 | 0.02729 | -0.3488 |
| UMUC7 | -0.02565 | 0.096012 | -0.02658 | 0.0139 | -0.3527 | -0.30737 | 0.08701 | 0.0165 | -0.17535 | -0.16198 | -0.17562 | 0.08016 | -0.1688 | 0.05554 | -0.05692 | -0.2141 | -0.12536 | 0.06308 | -0.22552 | 0.00731 | -0.12769 | -0.52314 | 0.054242 | 0.01212 | 0.11388 | -0.12185 | -0.41976 |
| UMUC9 | -0.24494 | -0.01919 | 0.02491 | -0.08254 | -0.14165 | -0.35801 | 0.136393 | -0.34339 | 0.0303 | -0.29087 | -0.01784 | -0.0108 | -0.2555 | 0.01468 | -0.21123 | 0.057444 | -0.15194 | 0.39115 | -0.26616 | 0.11098 | -0.0354 | -0.23581 | -0.14196 | -0.19001 | 0.13546 | 0.131925 | -0.48806 |
| UWB1289 | -0.11593 | 0.038168 | -0.11963 | 0.04084 | -0.16292 | -0.17313 | -0.00829 | -0.17609 | -0.18946 | 0.02974 | 0.019217 | 0.12765 | -0.14476 | -0.21837 | -0.12167 | -0.09479 | -0.035 | 0.15093 | -0.09685 | 0.1244 | 0.13676 | -0.06618 | 0.111852 | -0.04336 | 0.04175 | -0.00262 | -0.21274 |
| VP229 | -0.02902 | 0.170262 | -0.25434 | -0.12129 | -0.14078 | -0.22528 | -0.12472 | -0.12402 | -0.10648 | 0.06885 | -0.15977 | 0.26319 | -0.37177 | -0.22298 | -0.11387 | -0.11806 | 0.01782 | 0.00321 | -0.23294 | 0.23168 | 0.13273 | -0.48636 | 0.073315 | 0.00644 | 0.08555 | 0.023245 | -0.26925 |
| WERIRB1 | -0.34168 | 0.067661 | -0.32109 | -0.22051 | -0.17936 | 0.06416 | 0.147654 | -0.23201 | 0.03234 | 0.02289 | 0.011931 | 0.06457 | -0.03184 | -0.14545 | -0.04691 | -0.20073 | -0.12436 | 0.0305 | -0.06252 | 0.29764 | 0.22665 | -0.05291 | 0.063402 | 0.01823 | 0.30847 | 0.131814 | -0.42397 |
| WPE1NA22 | -0.24626 | -0.06432 | -0.69629 | 0.03594 | -0.20492 | -0.21643 | -0.10385 | -0.18667 | -0.10979 | -0.02915 | -0.07277 | 0.15102 | -0.19361 | -0.13629 | -0.04211 | -0.13187 | 0.01582 | -0.0289 | 0.002332 | 0.24543 | 0.08519 | -0.24665 | 0.086502 | -0.01797 | 0.16135 | 0.023516 | -0.48095 |
| TC138 | -0.27869 | 0.136282 | -0.50555 | -0.00956 | -0.09916 | -0.43165 | 0.036189 | -0.10421 | 0.010987 | -0.12487 | 0.065926 | 0.05702 | -0.06951 | 0.13521 | 0.139766 | -0.12184 | -0.35393 | 0.06367 | -0.01889 | 0.11318 | -0.07962 | -0.20755 | 0.176065 | 0.20106 | 0.30123 | 0.205818 | -0.36854 |
| TC205 | -0.18602 | 0.026297 | -0.5512 | 0.13076 | 0.05145 | -0.11933 | -0.18 | -0.10222 | -0.2068 | 0.04912 | 0.139766 | -0.03079 | -0.04572 | -0.05576 | -0.16945 | -0.0374 | -0.22518 | 0.09588 | 0.029992 | 0.16901 | 0.0234 | -0.35965 | 0.002412 | 0.04693 | 0.09051 | 0.027767 | -0.6853 |
| CCLFPEDS0008T | 0.0278 | -0.09677 | -0.2177 | -0.25562 | -0.23434 | -0.11626 | 0.109671 | -0.07995 | 0.067752 | -0.04129 | -0.33825 | 0.33927 | -0.23747 | -0.19742 | -0.20074 | -0.15249 | -0.24849 | 0.05248 | -0.12029 | 0.22611 | 0.12506 | -0.22649 | 0.113513 | -0.12487 | 0.03684 | -0.04436 | -0.22589 |
| A388 | -0.00494 | 0.005995 | -0.14069 | -0.08907 | -0.15462 | -0.10721 | 0.026088 | -0.14224 | -0.13081 | -0.14679 | -0.07015 | 0.1007 | -0.04997 | 0.04143 | -0.18457 | 0.035979 | -0.1197 | 0.03569 | -0.04418 | 0.03969 | 0.09219 | -0.20957 | 0.15264 | -0.08247 | 0.20003 | 0.015874 | -0.34786 |
| ASH3 | 0.05529 | 0.03524 | -0.21904 | -0.05708 | -0.11427 | -0.05274 | -0.04516 | -0.00122 | -0.09501 | -0.16672 | -0.05707 | 0.1279 | -0.23622 | -0.00691 | -0.12978 | -0.10011 | -0.05113 | -0.01266 | 0.004597 | 0.12036 | 0.10466 | -0.28571 | 0.206919 | 0.03144 | 0.11247 | -0.0705 | -0.21263 |
| BLUE1 | -0.02847 | -0.21628 | 0.02057 | 0.06728 | -0.25922 | -0.10299 | 0.001814 | -0.17368 | 0.052105 | -0.02911 | -0.11664 | 0.01442 | -0.4166 | -0.22884 | -0.32729 | -0.17807 | -0.20716 | 0.21728 | 0.055956 | 0.19625 | -0.06025 | 0.039762 | 0.020788 | -0.00193 | 0.27491 | -0.14219 | -0.36444 |
| BOKU | -0.08113 | 0.032422 | -0.41043 | -0.08758 | -0.25773 | 0.09478 | -0.03434 | -0.10293 | -0.22914 | -0.0039 | -0.11956 | 0.09509 | -0.24747 | -0.04823 | -0.13344 | -0.23757 | 0.10411 | 0.22031 | -0.01949 | 0.10727 | 0.21518 | -0.12165 | 0.227238 | -0.1626 | 0.10503 | 0.052658 | -0.29839 |
| BPH1 | -0.2348 | -0.02535 | -0.29491 | 0.07803 | -0.2185 | -0.05391 | -0.01064 | -0.06252 | -0.16399 | -0.13741 | -0.08161 | -0.06355 | -0.45659 | -0.10824 | -0.05036 | -0.08389 | -0.15671 | -0.2536 | -0.16345 | 0.24245 | 0.06738 | -0.17307 | 0.006282 | -0.04768 | 0.16392 | 0.158739 | -0.52942 |
| C10 | -0.0946 | 0.024597 | -0.1553 | -0.0411 | -0.10545 | -0.1113 | -0.11701 | -0.37915 | -0.21673 | -0.09168 | 0.011394 | 0.04541 | -0.05226 | -0.25666 | -0.15108 | -0.05574 | 0.1074 | 0.39178 | -0.00992 | 0.07846 | -0.41257 | -0.63148 | -0.00967 | -0.08337 | 0.33034 | -0.01676 | -0.36241 |
| C75 | 0.01805 | 0.020942 | -0.57689 | -0.12057 | 0.00032 | -0.40944 | -0.06466 | -0.0451 | -0.38569 | 0.02188 | -0.27983 | 0.18054 | -0.32762 | -0.17506 | -0.16007 | 0.111801 | -0.26594 | -0.00698 | 0.00609 | -0.19251 | 0.19469 | 0.188954 | 0.087781 | -0.15201 | 0.29207 | 0.266049 | -0.66148 |
| C80 | -0.30817 | -0.05092 | -0.27753 | -0.07104 | -0.05805 | -0.27616 | 0.075153 | -0.35298 | -0.3239 | 0.01606 | -0.10256 | 0.22142 | -0.27145 | -0.04634 | -0.0466 | -0.15664 | 0.16244 | 0.08006 | -0.16874 | 0.06217 | 0.17514 | -0.30702 | 0.069657 | 0.01125 | 0.14548 | 0.251968 | -0.37638 |
| C84 | -0.30988 | 0.011655 | -0.21898 | -0.07468 | -0.05128 | -0.10122 | 0.26007 | 0.14908 | 0.042737 | 0.04326 | -0.07864 | 0.12754 | 0.20106 | 0.0761 | -0.28425 | -0.28585 | 0.16909 | 0.2345 | -0.02716 | 0.24095 | 0.02472 | -0.21583 | -0.0829 | -0.17369 | 0.22291 | -0.12798 | -0.16735 |
| C99 | -0.33142 | -0.06216 | -0.57029 | 0.024 | 0.06414 | -0.10144 | -0.11856 | -0.47542 | -0.50229 | -0.00748 | -0.06264 | 0.01831 | -0.08295 | 0.0149 | -0.02935 | 0.068816 | -0.19092 | 0.23278 | -0.047 | 0.2306 | 0.0527 | -0.36203 | -0.06463 | -0.05438 | 0.224 | -0.16212 | -0.93759 |
| CHLA90 | -0.07112 | 0.006652 | -0.30862 | -0.16301 | -0.07273 | -0.06211 | -0.1327 | -0.4371 | -0.04718 | -0.09929 | -0.19556 | 0.22693 | -0.08257 | -0.52176 | -0.1005 | -0.06122 | -0.0057 | -0.29156 | -0.09678 | 0.17888 | 0.19902 | -0.18807 | 0.113486 | 0.04917 | 0.28838 | -0.00903 | -0.15366 |
| EGI1 | -0.03886 | 0.052452 | -0.44129 | -0.03778 | -0.1513 | -0.10115 | 0.080494 | -0.21286 | -5.3E-05 | -0.06629 | -0.11285 | 0.16621 | -0.10656 | -0.02388 | -0.13821 | 0.03417 | -0.08728 | 0.0863 | 0.007763 | 0.13555 | 0.05695 | -0.02515 | 0.11163 | -0.08426 | 0.14108 | 0.00322 | -0.46289 |
| EMTOKA | -0.03197 | -0.02671 | -0.08982 | -0.03977 | 0.10547 | -0.3295 | 0.037177 | -0.16455 | -0.1802 | -0.12864 | -0.01589 | -0.10992 | -0.33308 | 0.17498 | -0.11014 | -0.10009 | -0.20506 | -0.18783 | -0.12598 | 0.17745 | 0.09999 | -0.04984 | 0.134688 | 0.05274 | 0.13725 | 0.025279 | -0.24852 |
| ESO26 | -0.1568 | -0.0303 | -0.05238 | -0.1076 | -0.12366 | -0.05548 | -0.11815 | -0.22012 | -0.06626 | -0.15059 | -0.17629 | 0.08037 | -0.15195 | -0.24777 | -0.15806 | -0.09497 | -0.00765 | 0.01079 | -0.14231 | 0.15563 | 0.14648 | -0.15862 | 0.058883 | -0.14421 | 0.22717 | 0.141515 | -0.25772 |
| ESO51 | -0.02492 | 0.015185 | -0.31548 | -0.11488 | -0.13206 | -0.15739 | 0.000644 | -0.12748 | -0.76932 | 0.05086 | -0.57738 | 0.14122 | -0.09404 | -0.14494 | -0.0152 | -0.08784 | 0.05297 | -0.09198 | -0.06999 | 0.22191 | 0.2723 | -0.11771 | 0.034016 | -0.07511 | 0.26692 | -0.20048 | -0.31084 |
| FARAGE | -0.07003 | 0.02332 | -0.12284 | 0.04124 | -0.11095 | -0.05168 | 0.039127 | -0.13671 | -0.14712 | 0.02204 | -0.1145 | 0.19672 | -0.10676 | 0.07943 | -0.14698 | -0.12133 | -0.29113 | 0.21636 | -0.08394 | 0.18629 | 0.32679 | -0.4415 | 0.198301 | -0.06027 | 0.09403 | 0.08317 | -0.35847 |
| FLO1 | -0.19015 | 0.019761 | -0.30246 | -0.17935 | -0.12823 | -0.17138 | 0.084561 | -0.24593 | -0.21037 | -0.01282 | -0.11324 | 0.17745 | -0.19094 | -0.16321 | -0.10549 | -0.09993 | 0.08783 | 0.00258 | 0.113725 | 0.21259 | 0.13354 | -0.29891 | -0.11469 | -0.05168 | 0.20196 | 0.075801 | -0.58786 |
| H357 | 0.13324 | 0.106456 | -0.59168 | -0.0654 | -0.0137 | -0.0351 | -0.06045 | -0.31066 | -0.35498 | -0.07542 | -0.11036 | 0.17004 | -0.40529 | 0.07373 | -0.40373 | -0.07168 | -0.09735 | 0.55179 | 0.082767 | -0.02418 | 0.22084 | -0.57659 | 0.223544 | -0.22508 | 0.05738 | -0.04966 | -0.64457 |
| H376 | -0.15669 | 0.043088 | -0.49037 | -0.21511 | -0.09753 | -0.11135 | -0.1062 | -0.11494 | -0.25763 | 0.09008 | -0.06035 | 0.05964 | 0.12613 | -0.12304 | -0.00041 | -0.00573 | -0.06142 | -0.03022 | -0.06775 | 0.11441 | 0.05527 | -0.0825 | 0.206919 | -0.02522 | 0.04196 | -0.13636 | -0.33722 |
| H413 | -0.01529 | 0.048008 | -0.27342 | -0.09959 | -0.03056 | -0.12376 | -0.02854 | -0.05728 | 0.060974 | -0.13201 | -0.00091 | 0.10531 | -0.01675 | -0.11612 | 0.078592 | -0.16426 | -0.02526 | -0.01556 | 0.181768 | 0.28243 | -0.0506 | -0.35347 | 0.06128 | 0.02665 | 0.04346 | 0.008852 | -0.32327 |
| HCA1 | -0.07308 | -0.05154 | -0.21703 | -0.0515 | -0.25749 | -0.19837 | -0.06718 | -0.25887 | -0.13216 | 0.03573 | -0.05438 | 0.0696 | -0.19177 | -0.1402 | -0.16141 | -0.04524 | -0.09591 | 0.04374 | -0.00968 | 0.14494 | 0.07509 | -0.18116 | 0.136864 | -0.07737 | 0.0034 | -0.10669 | -0.31437 |
| HCS2 | -0.18317 | 0.05696 | -0.39699 | -0.04399 | -0.14379 | -0.22915 | -0.09601 | -0.19074 | -0.05607 | 0.04571 | -0.09826 | 0.0426 | -0.35499 | -0.08279 | -0.08751 | -0.06371 | 0.03731 | 0.06864 | -0.01931 | 0.07744 | 0.15416 | -0.12188 | 0.143458 | -0.15171 | 0.29305 | -0.0928 | -0.35815 |
| HEC1 | -0.16265 | -0.00397 | -0.03848 | -0.16769 | -0.25071 | -0.23935 | -0.10333 | -0.12805 | -0.12605 | -0.15525 | -0.10083 | 0.15077 | -0.24539 | -0.08632 | -0.16774 | -0.14324 | -0.11329 | 0.04358 | 0.013225 | 0.09576 | 0.26455 | -0.12582 | 0.070911 | -0.07426 | 0.15106 | 0.044087 | -0.46281 |
| HEC116 | -0.23023 | 0.122284 | -0.1482 | -0.0562 | -0.27355 | -0.17781 | 0.025023 | -0.14038 | 0.098347 | -0.21267 | -0.10345 | 0.12536 | -0.26528 | -0.17761 | -0.08963 | -0.05511 | -0.00586 | 0.03036 | -0.10248 | -0.00558 | 0.13622 | -0.21315 | 0.027837 | -0.06515 | 0.19449 | -0.1 | -0.36011 |
| HG3 | -0.16814 | 0.077811 | -0.12908 | 0.12554 | -0.04129 | -0.10772 | -0.09852 | -0.19995 | 0.00892 | -0.03694 | -0.09501 | 0.20886 | -0.13267 | 0.47027 | -0.19517 | 0.034142 | -0.0947 | 0.33271 | -0.28454 | 0.10542 | -0.03046 | -0.31723 | -0.00975 | -0.06072 | -0.07007 | -0.04606 | -0.33717 |
| HKA1 | -0.01085 | -0.00931 | -0.50187 | -0.08559 | -0.06475 | 0.03633 | -0.04669 | -0.46743 | -0.20581 | 0.09541 | 0.002664 | 0.0875 | -0.10761 | -0.13784 | -0.05024 | 0.02714 | 0.10357 | 0.14449 | -0.09056 | 0.16514 | 0.14899 | -0.45356 | 0.011389 | 0.04197 | 0.17162 | -0.02336 | -0.19157 |
| HMY1 | -0.01273 | 0.006421 | -0.35949 | -0.13027 | -0.23199 | 0.01375 | -0.1008 | -0.11439 | -0.16666 | -0.12007 | -0.03633 | 0.05868 | -0.32798 | -0.11884 | -0.00057 | -0.09968 | 0.04241 | -0.07831 | -0.04414 | 0.17993 | 0.10809 | -0.33503 | 0.138722 | 0.00602 | 0.24581 | 0.007124 | -0.37737 |

| Cell line | FLT3 | FLT4 | IGF1R | INSR | INSRR | KDR | KIT | LMR1 | LMR2 | LTK | MET | MUSK | PDGFRA | PDGFRB | PTK7 | RET | RON | ROR1 | ROR2 | ROS1 | STYK1 | TIE1 | TIE2 | TRKA | TRKB | TRKC | TYRO3 |
|---|---|---|---|---|---|---|---|---|---|---|---|---|---|---|---|---|---|---|---|---|---|---|---|---|---|---|---|
| HSC1 | -0.10915 | -0.01503 | -0.33317 | -0.15722 | -0.1296 | -0.30534 | -0.14355 | -0.07091 | -0.07626 | -0.0217 | 0.01689 | 0.12903 | -0.05972 | -0.1222 | 0.017073 | -0.20326 | -0.07542 | 0.22164 | -0.11223 | 0.05697 | 0.12844 | -0.09988 | 0.100613 | -0.01481 | 0.12499 | 0.017533 | -0.25239 |
| HSC5 | -0.16192 | -0.10778 | -0.01917 | -0.02315 | -0.11237 | -0.17944 | 0.080921 | -0.19643 | -0.04934 | -0.09148 | -0.06603 | 0.02126 | -0.20941 | -0.25923 | -0.14288 | -0.12545 | -0.10416 | 0.12383 | 0.068664 | 0.20095 | 0.11346 | -0.19453 | 0.074796 | -0.10863 | 0.02521 | 0.09992 | -0.61946 |
| HT3 | -0.23742 | 0.135503 | -0.39346 | -0.18137 | -0.02128 | -0.09008 | 0.067644 | -0.13141 | -0.07046 | 0.0009 | -0.17633 | -0.18822 | -0.05215 | -0.2123 | 0.056063 | -0.27829 | -0.00822 | 0.19304 | -0.12181 | 0.12608 | -0.00491 | -0.19526 | 0.219023 | 0.03151 | -0.03287 | 0.016133 | -0.3662 |
| HUO9 | 0.16903 | 0.037293 | -0.4763 | -0.0921 | -0.00757 | 0.06366 | 0.01484 | -0.14441 | 0.057592 | 0.12743 | -0.02876 | 0.11859 | -0.84658 | 0.14766 | 0.030196 | -0.17343 | -0.2183 | -0.09769 | -0.09141 | 0.08251 | 0.10895 | -0.16552 | 0.206891 | 0.09712 | 0.10025 | 0.116877 | -0.3112 |
| IHH4 | -0.17201 | 0.024841 | -0.17301 | -0.15982 | -0.15995 | -0.07033 | 0.00109 | -0.20282 | -0.02775 | -0.07433 | -0.08423 | 0.04156 | -0.03838 | -0.18569 | -0.08076 | -0.00337 | -0.1079 | 0.07571 | 0.022884 | 0.15538 | 0.13024 | -0.03707 | 0.091965 | -0.00559 | 0.22481 | 0.205776 | -0.4353 |
| JAR | -0.11275 | 0.146165 | -0.07692 | -0.11905 | -0.15273 | -0.12059 | -0.08325 | -0.08043 | 0.065817 | -0.09573 | 0.028186 | 0.08896 | -0.05471 | -0.08198 | -0.12137 | -0.14949 | -0.11657 | 0.18515 | -0.14654 | -0.00673 | 0.26394 | -0.2871 | 0.085763 | -0.03112 | 0.08332 | -0.00407 | -0.60094 |
| JEG3 | -0.16821 | -0.23267 | -0.38614 | -0.09073 | -0.03897 | -0.13281 | 0.141124 | -0.45739 | -0.32179 | 0.16507 | -0.03737 | 0.0526 | -0.29974 | -0.31629 | -0.13118 | -0.07261 | 0.19215 | 0.07173 | 0.0381 | 0.21344 | 0.20422 | 0.046359 | 0.122582 | -0.06846 | -0.02629 | 0.115514 | -0.5194 |
| JMURTK2 | -0.04113 | 0.071957 | 0.03089 | 0.07357 | -0.09348 | -0.16391 | -0.02365 | -0.19496 | -0.03761 | -0.06676 | -0.07282 | 0.04924 | -0.09145 | -0.10394 | -0.07637 | -0.06527 | -0.07321 | 0.16049 | 0.125165 | 0.17987 | 0.0391 | -0.05354 | 0.089308 | 0.00228 | 0.14547 | 0.029784 | -0.27066 |
| KARPAS1718 | -0.18263 | -0.04026 | -0.29354 | -0.15509 | -0.15331 | -0.15035 | -0.12365 | -0.24319 | 0.014367 | -0.09078 | -0.16221 | 0.0301 | -0.08678 | -0.15782 | -0.15033 | -0.00556 | -0.08047 | 0.12132 | -0.06886 | 0.26763 | 0.19999 | 0.06948 | -0.04368 | 0.00381 | 0.43437 | -0.16283 | -0.42234 |
| KKU100 | -0.29565 | -0.11872 | -0.31208 | -0.19282 | 0.14858 | -0.21141 | -0.06063 | -0.24469 | 0.136915 | -0.11061 | -0.04612 | 0.09201 | -0.15541 | -0.24755 | 0.016449 | -0.06898 | 0.11715 | 0.32339 | -0.09598 | 0.08371 | 0.06919 | -0.13561 | 0.098081 | 0.01781 | 0.00041 | 0.074365 | -0.57224 |
| KKU213 | 0.0904 | 0.199523 | -0.74479 | -0.07497 | -0.14561 | -0.18442 | 0.087246 | -0.11187 | -0.17306 | -0.1626 | -0.03843 | 0.09372 | -0.19216 | 0.10711 | -0.24398 | -0.18005 | -0.20505 | 0.07681 | -0.07318 | 0.01472 | 0.1583 | -0.16787 | 0.231365 | -0.02549 | -0.01547 | -0.06006 | -0.40462 |
| KML1 | -0.02652 | -0.12837 | -0.27426 | 0.18076 | -0.05752 | -0.1368 | 0.104962 | 0.08026 | -0.51123 | 0.16851 | -0.07967 | 0.27238 | 0.08306 | 0.04771 | -0.05645 | -0.1128 | -0.14224 | 0.1016 | -0.24318 | -0.08004 | 0.20183 | -0.1093 | 0.229313 | -0.12736 | 0.24132 | 0.16197 | -0.56219 |
| KON | -0.10026 | 0.182449 | -0.48222 | -0.00975 | -0.00964 | -0.00126 | -0.03103 | -0.25414 | -0.05996 | -0.19233 | -0.06402 | 0.11664 | -0.30061 | -0.00064 | -0.15201 | -0.11502 | 0.09825 | 0.19057 | -0.08978 | 0.13607 | 0.08316 | -0.18401 | 0.059234 | -0.01173 | 0.01724 | -0.07539 | -0.43418 |
| KOSC2 | -0.15993 | 0.030807 | -0.16637 | -0.14975 | -0.2907 | -0.04683 | -0.0293 | -0.23448 | -0.10816 | -0.25855 | -0.00752 | -0.01707 | -0.02184 | -0.42344 | -0.08429 | 0.077322 | -0.12419 | 0.17245 | -0.10995 | 0.27535 | -0.01368 | 0.039553 | 0.033246 | -0.09349 | 0.0757 | -0.06301 | -0.35245 |
| KYAE1 | -0.06819 | 0.229764 | -0.01101 | -0.01513 | 0.15022 | 0.11577 | 0.45522 | 0.04778 | 0.183607 | 0.00016 | -0.26087 | 0.02254 | 0.1546 | 0.21571 | -0.01003 | 0.332028 | 0.07701 | 0.19214 | -0.03864 | 0.20711 | 0.18228 | 0.010268 | -0.09602 | 0.08884 | -0.02688 | 0.164786 | 0.1318 |
| LO68 | -0.11708 | 0.032911 | -0.49678 | -0.26767 | -0.30388 | -0.019 | -0.13215 | -0.2669 | 0.043265 | -0.26753 | -0.08974 | 0.00161 | -0.26213 | -0.1348 | -0.11366 | -0.06322 | -0.11773 | 0.06171 | -0.02494 | 0.25666 | 0.13588 | -0.18686 | 0.091191 | -0.17139 | 0.19344 | 0.14554 | -0.3738 |
| LS | -0.07921 | 0.101186 | -0.50972 | 0.00041 | 0.02586 | -0.26988 | -0.22555 | -0.35992 | -0.2984 | 0.13821 | 0.049018 | -0.08383 | -0.07453 | -0.11206 | 0.068597 | -0.20114 | -0.01071 | 0.13768 | -0.05109 | 0.08559 | -0.01692 | -0.01293 | 0.051054 | 0.06808 | 0.05897 | 0.139184 | -0.01057 |
| LU135 | -0.05546 | 0.069809 | -0.32309 | -0.15536 | -0.12615 | -0.19328 | 0.041225 | -0.14309 | -0.06712 | -0.0492 | -0.12871 | 0.07528 | -0.28296 | -0.15822 | -0.15009 | 0.019413 | -0.05345 | 0.17332 | -0.07607 | 0.01371 | 0.15722 | -0.15337 | 0.124824 | -0.0385 | 0.14519 | 0.00063 | -0.33786 |
| MCC13 | -0.0299 | -0.10267 | -0.10307 | -0.10719 | -0.35462 | -0.08838 | -0.03505 | -0.32585 | -0.23171 | -0.03163 | -0.10654 | 0.41723 | -0.07877 | -0.3303 | -0.15301 | -0.15062 | -0.23994 | -0.14084 | 0.108142 | 0.25532 | 0.09057 | -0.1953 | 0.053007 | -0.04254 | 0.26363 | 0.109765 | -0.55639 |
| MCC142 | -0.06248 | 0.003928 | -0.21712 | -0.15957 | -0.1133 | -0.01932 | 0.036582 | -0.10342 | -0.09993 | 0.11918 | -0.09574 | 0.08032 | -0.15885 | -0.13027 | -0.10826 | -0.15103 | -0.08248 | 0.25902 | 0.006381 | 0.23011 | 0.00519 | -0.09338 | 0.105501 | -0.0431 | 0.06705 | 0.038503 | -0.2767 |
| MCC26 | 0.07861 | 0.008761 | -0.64692 | -0.02574 | -0.07088 | -0.36383 | -0.14021 | 0.02662 | -0.10446 | 0.11164 | -0.06798 | 0.07921 | -0.35332 | -0.18275 | 0.063379 | -0.04014 | -0.19422 | 0.04634 | -0.03053 | 0.12999 | 0.07464 | -0.00686 | 0.058855 | 0.04307 | 0.349 | 0.192742 | -0.35429 |
| MEL202 | -0.12731 | 0.036631 | -0.29385 | -0.14048 | -0.02573 | -0.08269 | -0.11405 | -0.2207 | -0.08985 | 0.00442 | -0.13254 | 0.15271 | -0.07838 | -0.05137 | -0.09976 | -0.08587 | -0.10782 | 0.12778 | 0.033777 | 0.06817 | 0.14504 | -0.14791 | 0.093541 | -0.04509 | 0.16593 | 0.095602 | -0.50869 |
| MERO14 | -0.00471 | -0.12696 | -0.27719 | 0.01821 | -0.12676 | 0.01911 | 0.033744 | -0.28635 | -0.09942 | 0.15842 | -0.02579 | -0.24242 | -0.0012 | -0.22261 | -0.19444 | -0.12279 | -0.04456 | 0.40422 | 0.138473 | 0.27823 | 0.09088 | -0.34867 | 0.268753 | -0.06602 | 0.13484 | -0.12655 | -0.4586 |
| MERO25 | -0.15442 | 0.032311 | -0.09177 | -0.04849 | -0.09215 | 0.02416 | -0.01178 | -0.51626 | -0.09376 | -0.07331 | -0.14802 | 0.00778 | -0.26641 | -0.29915 | -0.04978 | -0.06274 | -0.04964 | 0.02871 | 0.034811 | 0.10727 | 0.14115 | -0.19535 | 0.021673 | -0.07449 | 0.15084 | 0.005073 | -0.38714 |
| MERO41 | -0.11591 | -0.12158 | -0.24994 | -0.06672 | -0.10483 | 0.00688 | -0.04775 | -0.18208 | 0.011623 | -0.21501 | -0.20424 | 0.08267 | -0.32836 | -0.17904 | -0.0115 | -0.1466 | 0.06435 | 0.12768 | -0.03109 | -0.05138 | 0.0534 | -0.18458 | 0.069017 | -0.00593 | 0.0438 | 0.141911 | -0.47069 |
| MERO48A | -0.03947 | -0.02787 | -0.07718 | -0.08415 | -0.14345 | -0.20703 | -0.14219 | -0.47534 | -0.30095 | -0.04167 | -0.00708 | -0.04397 | -0.5796 | -0.46201 | -0.14482 | -0.04051 | 0.09758 | -0.05367 | 0.092645 | 0.15649 | -0.09696 | -0.20416 | 0.116279 | -0.10955 | 0.0288 | 0.143271 | -0.29871 |
| MERO82 | -0.19908 | 0.049477 | -0.08637 | 0.10788 | -0.02073 | -0.09218 | 0.045031 | -0.40199 | -0.24053 | 0.0057 | -0.16847 | 0.00193 | -0.43266 | -0.36848 | -0.21086 | -0.05073 | -0.11448 | 0.02636 | 0.085119 | 0.08771 | 0.02309 | -0.14085 | 0.184198 | -0.1338 | 0.19268 | 0.148 | -0.47518 |
| MERO83 | -0.01062 | 0.098253 | -0.29218 | -0.14999 | -0.23315 | -0.08357 | 0.072913 | -0.20891 | -0.08943 | -0.06866 | -0.14494 | 0.20362 | -0.06466 | -0.00264 | -0.24019 | -0.04985 | -0.1715 | -0.23297 | 0.075589 | 0.13007 | 0.07466 | -0.11847 | 0.198824 | -0.14217 | 0.41485 | -0.12313 | -0.30737 |
| MERO95 | -0.07481 | 0.046895 | -0.24936 | -0.0585 | -0.11392 | -0.14255 | -0.00859 | -0.17416 | -4.4E-05 | 0.12236 | -0.03223 | 0.15627 | -0.20669 | -0.14495 | -0.01849 | -0.24494 | 0.06182 | 0.15774 | -0.00031 | 0.14251 | 0.03615 | -0.19935 | 0.054697 | -0.09041 | 0.05357 | -0.01585 | -0.13983 |
| MM127 | -0.04327 | 0.033034 | -0.28506 | -0.03262 | -0.01947 | -0.20972 | -0.02746 | -0.19636 | -0.10116 | 0.13707 | -0.09643 | 0.1719 | -0.19761 | 0.06898 | 0.011956 | -0.11277 | -0.00803 | 0.03942 | 0.158313 | 0.30629 | 0.12648 | -0.29044 | 0.222741 | 0.03215 | 0.0276 | 0.005843 | -0.25442 |
| MM370 | -0.14422 | -0.00145 | -0.0989 | -0.16179 | -0.14847 | -0.06469 | -0.12939 | -0.08933 | 0.089343 | -0.15231 | -0.13061 | 0.27313 | -0.3146 | 0.01406 | 0.020653 | -0.10104 | -0.11809 | 0.05489 | -0.00845 | 0.24836 | -0.03317 | -0.19185 | 0.278244 | -0.10533 | 0.07243 | 0.003803 | -0.33606 |
| MM383 | -0.25172 | 0.045527 | -0.48234 | 0.06626 | -0.26414 | -0.02298 | 0.051944 | -0.10782 | 0.228682 | 0.19289 | 0.008781 | 0.25035 | -0.25164 | -0.22895 | -0.07012 | 0.063376 | 0.11448 | -0.05058 | -0.02688 | 0.12984 | 0.26684 | -0.11919 | 0.098036 | -0.08053 | -0.00319 | 0.1257 | -0.23401 |
| MM386 | 0.05572 | 0.09434 | -0.34201 | -0.21059 | -0.13251 | -0.31386 | -0.01835 | -0.1329 | 0.158325 | -0.08058 | -0.02874 | 0.24788 | -0.24317 | -0.07511 | -0.15706 | 0.082332 | -0.02052 | 0.17799 | -0.13446 | 0.20143 | 0.09788 | -0.19545 | 0.004162 | -0.04341 | 0.2266 | -0.1032 | -0.26209 |
| MM426 | -0.17326 | -0.0088 | -0.15884 | -0.05277 | -0.01733 | -0.18257 | -0.01202 | -0.26281 | 0.139495 | 0.16313 | 0.057596 | 0.19306 | -0.12819 | 0.00638 | 0.010237 | -0.0613 | -0.13806 | 0.19963 | 0.079134 | 0.33546 | -0.00436 | -0.07192 | -0.00375 | 0.03879 | 0.17694 | 0.087949 | -0.5346 |
| MOLM14 | -0.85339 | 0.195795 | -0.1037 | 0.01882 | -0.2183 | -0.03751 | -0.0073 | -0.19707 | -0.22725 | 0.01564 | -0.03534 | -0.05812 | -0.17033 | -0.32993 | -0.3272 | 0.1241 | 0.14341 | 0.14235 | 0.161245 | 0.17118 | 0.0656 | -0.01423 | -0.25527 | -0.07722 | -0.05952 | 0.077092 | -0.48825 |
| MUTZ8 | -0.0016 | -0.0617 | -0.47795 | -0.46116 | -0.06947 | -0.06148 | 0.265368 | -0.50595 | -0.22602 | 0.1125 | -0.11899 | -0.06336 | -0.58822 | -0.00568 | -0.2733 | 0.127868 | 0.06249 | -0.09935 | 0.019737 | -0.02456 | 0.16656 | -0.3885 | 0.21494 | 0.09062 | -0.01213 | 0.086704 | -0.483 |
| NH12 | -0.23924 | -0.37105 | -1.28597 | -0.3488 | -0.27953 | -0.16611 | -0.01241 | -0.19424 | -0.05453 | 0.04528 | 0.009609 | 0.268 | -0.16021 | -0.17477 | -0.23796 | 0.036343 | 0.06881 | -0.12459 | 0.129025 | -0.03179 | 0.18592 | -0.37742 | 0.193492 | 0.06767 | 0.29373 | -0.27022 | -0.51968 |
| NO10 | -0.05379 | -0.12709 | 0.0562 | 0.00999 | -0.08683 | -0.06421 | -0.01174 | -0.13949 | -0.12897 | -0.00141 | -0.09225 | 0.04952 | -0.2045 | -0.01081 | 0.046625 | 0.040005 | 0.07593 | 0.07514 | -0.0255 | 0.17354 | 0.12912 | -0.10802 | 0.191099 | -0.09648 | 0.17435 | 0.071962 | -0.45058 |
| NO11 | -0.10157 | 0.074688 | 0.01762 | 0.02026 | -0.00471 | -0.09318 | 0.183796 | 0.02649 | 0.051813 | 0.18679 | -0.32525 | 0.08474 | -0.38982 | -0.01114 | 0.019371 | -0.08981 | -0.01438 | 0.13471 | -0.14173 | 0.18284 | 0.19301 | -0.2036 | -0.02835 | -0.05958 | 0.17546 | 0.219206 | -0.20581 |
| NOZ | 0.02898 | 0.125779 | -0.25182 | 0.05673 | -0.03425 | -0.17233 | -0.10498 | -0.27461 | -0.16347 | -0.02859 | -0.08186 | 0.08281 | -0.21512 | -0.20703 | -0.06587 | -0.01307 | -0.00919 | -0.0763 | 0.014078 | 0.16578 | 0.11499 | -0.01063 | 0.093773 | -0.00329 | 0.13095 | -0.04052 | -0.32595 |
| NP2 | -0.09091 | -0.02831 | -0.2345 | -0.11415 | -0.15106 | -0.04409 | -0.01849 | -0.31232 | -0.14738 | -0.10854 | -0.08484 | 0.06567 | -0.14415 | -0.20169 | -0.203 | -0.02402 | -0.10908 | 0.06223 | 0.08293 | 0.13705 | 0.14751 | -0.25344 | 0.082489 | -0.03844 | 0.05412 | 0.02629 | -0.48668 |
| NP3 | -0.12327 | 0.171819 | -0.01501 | -0.08267 | -0.09139 | -0.19526 | -0.10257 | -0.12919 | -0.10261 | -0.03386 | -0.07355 | 0.10558 | -0.41276 | -0.07829 | -0.15247 | -0.15332 | -0.18986 | 0.26112 | -0.10032 | 0.02758 | 0.06361 | -0.12642 | 0.201247 | -0.0715 | 0.03873 | 0.013522 | -0.27884 |
| NP5 | 0.0246 | -0.13981 | 0.17145 | -0.04864 | -0.18803 | -0.19021 | -0.22574 | -0.10453 | -0.08345 | -0.11198 | 0.028907 | -0.13734 | -0.61183 | -0.46084 | 0.053988 | -0.27563 | -0.15407 | -0.22409 | -0.11053 | 0.05149 | 0.04317 | -0.53974 | 0.069364 | -0.08107 | -0.01931 | -0.07679 | -0.74159 |
| NP8 | -0.05345 | 0.144834 | -0.09076 | -0.06167 | 0.02529 | 0.24753 | 0.234627 | 0.07252 | -0.18447 | 0.00498 | -0.09418 | 0.22386 | -0.19356 | -0.11533 | -0.07813 | -0.09754 | -0.18014 | 0.04319 | 0.013707 | -0.00469 | -0.03059 | -0.04391 | 0.13148 | -0.20702 | 0.28424 | 0.095487 | -0.2655 |
| OCILY18 | -0.08078 | -0.00185 | -0.08973 | 0.02973 | -0.1174 | -0.0901 | 0.067807 | -0.11735 | -0.0082 | -0.08776 | -0.21743 | 0.0482 | -0.08023 | 0.14678 | 0.013984 | -0.11235 | 0.06411 | 0.07177 | -0.01125 | 0.11622 | 0.10057 | -0.05261 | 0.146299 | 0.04789 | 0.05973 | 0.005578 | -0.18514 |
| OCIM2 | -0.00582 | 0.228572 | -0.36096 | 0.14156 | -0.17247 | -0.22855 | 0.162863 | -0.30693 | -0.01925 | 0.01012 | -0.15611 | 0.29587 | -0.03964 | -0.13281 | -0.34764 | -0.0597 | 0.07187 | 0.00874 | -0.11075 | -0.0338 | 0.15149 | -0.01361 | -0.22691 | -0.11428 | 0.06071 | -0.08989 | -0.22993 |
| OCUG1 | -0.15854 | 0.044874 | -0.62966 | -0.2235 | 0.02233 | 0.11048 | 0.018875 | -0.17211 | -0.04642 | 0.0775 | 0.043115 | -0.09928 | -0.11729 | 0.02578 | 0.090329 | -0.08862 | 0.06935 | 0.27839 | -0.05299 | -0.06212 | 0.19986 | 0.007615 | 0.06893 | -0.04608 | 0.18443 | -0.23145 | -0.45406 |
| ONDA7 | -0.1495 | -0.06049 | -0.11532 | -0.06118 | -0.13125 | -0.14442 | 0.050836 | -0.12344 | -0.09268 | 0.06002 | -0.03974 | 0.09753 | -0.21766 | -0.40835 | -0.08649 | -0.12791 | -0.06228 | 0.01052 | 0.052136 | 0.2705 | 0.10759 | -0.02679 | 0.090315 | -0.02051 | 0.04194 | 0.031639 | -0.31275 |
| ONDA8 | -0.1433 | -0.05473 | -0.20136 | -0.11574 | -0.1535 | -0.11593 | -0.0547 | -0.05375 | -0.11338 | 0.15427 | -0.15848 | 0.08463 | -0.43024 | -0.24132 | -0.02429 | -0.21699 | -0.05378 | -0.08916 | -0.1172 | 0.16528 | 0.19546 | -0.34962 | 0.237786 | -0.0145 | 0.12222 | -0.04191 | -0.30417 |
| ONDA9 | 0.07348 | 0.066175 | 0.03425 | -0.03676 | -0.22918 | 0.03304 | -0.05323 | -0.1398 | -0.12135 | 0.1744 | -0.00246 | -0.03508 | -0.16986 | 0.06754 | 0.018056 | -0.12615 | -0.1527 | -0.0505 | 0.069358 | 0.10312 | 0.19202 | -0.29295 | 0.163692 | -0.02973 | 0.26909 | 0.252243 | -0.28701 |
| OSC19 | 0.02779 | 0.290529 | 0.07248 | 0.07501 | 0.01384 | -0.05682 | 0.329303 | 0.08439 | -0.09207 | -0.02417 | -0.12882 | 0.13461 | -0.17133 | 0.31663 | -0.06244 | -0.10577 | -0.04667 | 0.25783 | 0.105444 | 0.31825 | 0.23388 | -0.12214 | -0.32759 | 0.07549 | -0.03799 | 0.26746 | -0.25148 |
| OSC20 | -0.19734 | -0.01308 | -0.19291 | -0.1502 | -0.35847 | -0.16686 | -0.16293 | -0.22027 | -0.16028 | -0.12161 | 0.0684 | -0.01669 | -0.31796 | -0.16303 | -0.14117 | -0.06699 | -0.0369 | 0.10084 | -0.07121 | 0.14589 | 0.11655 | -0.3518 | 0.084914 | -0.03203 | 0.11673 | 0.076879 | -0.52134 |
| P4E6 | -0.07842 | -0.07919 | -0.64862 | -0.106 | -0.17335 | 0.0084 | -0.04119 | -0.23166 | -0.00546 | -0.14699 | -0.18965 | 0.13583 | -0.09025 | -0.07326 | -0.03599 | -0.10036 | -0.06097 | -0.14701 | -0.067 | 0.15624 | 0.12168 | -0.13479 | 0.150515 | -0.03337 | -0.01733 | 0.040614 | -0.33768 |
| PEA1 | 0.16359 | -0.10976 | -0.25331 | -0.08594 | 0.10232 | -0.26678 | -0.05554 | -0.15639 | -0.10336 | 0.02785 | -0.02684 | 0.08568 | -0.05154 | -0.03062 | -0.01777 | -0.10132 | -0.05106 | 0.08539 | -0.09349 | 0.07724 | 0.15395 | -0.21654 | -0.01432 | 0.1543 | 0.27487 | 0.04891 | -0.46715 |
| PEO1 | -0.12686 | -0.03167 | -0.49975 | -0.05416 | -0.06141 | -0.14828 | -0.10536 | -0.1922 | -0.15265 | -0.074 | -0.07586 | 0.11301 | -0.20451 | -0.18702 | -0.07139 | -0.08314 | -0.07553 | 0.05057 | -0.0002 | 0.12577 | 0.10803 | -0.181 | 0.146683 | 0.00054 | 0.13319 | -0.02344 | -0.28232 |
| PEO4 | -0.07479 | -0.13397 | -0.2251 | -0.12539 | 0.04176 | -0.05617 | 0.179759 | -0.08201 | -0.19204 | -0.13704 | -0.14191 | 0.05865 | -0.02148 | -0.16778 | -0.0221 | -0.03412 | 0.09739 | 0.0008 | -0.1159 | 0.28055 | 0.09349 | -0.31624 | 0.192114 | 0.18539 | 0.03408 | 0.105641 | -0.18812 |
| PGA1 | -0.19282 | 0.120997 | -0.10245 | -0.09082 | -0.00534 | 0.13186 | -0.10137 | -0.19766 | 0.175981 | -0.0215 | -0.19942 | 0.08617 | -0.08582 | -0.12393 | -0.0987 | 0.01883 | 0.00939 | 0.04625 | 0.106682 | 0.15212 | 0.01421 | -0.12613 | 0.141534 | 0.04228 | 0.15591 | -0.03668 | -0.49131 |
| ROS50 | -0.29945 | -0.25313 | -0.19227 | -0.19864 | -0.29355 | -0.08455 | 0.001863 | -0.07942 | 0.049625 | -0.22683 | -0.10194 | -0.1661 | -0.128 | -0.3632 | -0.16564 | -0.12834 | -0.30034 | 0.18664 | -0.09246 | -0.11938 | -0.04976 | -0.12992 | -0.01013 | -0.37129 | 0.25364 | -0.04706 | -0.66758 |
| SAT | -0.13831 | 0.126128 | -0.13855 | -0.09479 | -0.26177 | -0.25129 | -0.21699 | -0.28645 | 0.005945 | -0.04964 | -0.08423 | 0.14686 | -0.16646 | -0.23657 | -0.09599 | -0.02907 | -0.01013 | -0.06589 | -0.08259 | 0.13798 | 0.0266 | -0.27273 | 0.074098 | -0.08427 | 0.06163 | 0.167649 | -0.49506 |
| SEKI | -0.15432 | 0.039136 | -0.46516 | -0.08725 | -0.11925 | 0.0173 | 0.036002 | -0.19845 | -0.1628 | -0.02074 | -0.01515 | 0.10635 | -0.09656 | -0.15876 | 0.020316 | -0.06535 | -0.22049 | 0.19009 | -0.2396 | 0.10604 | -0.01751 | 0.00308 | -0.07309 | -0.02012 | 0.02191 | 0.033549 | -0.35353 |
| SHI1 | -0.20245 | -0.29257 | -0.68469 | -0.42006 | -0.09721 | -0.0523 | -0.04747 | -0.13129 | -0.02134 | 0.04382 | -0.10632 | 0.19884 | -0.08901 | -0.16439 | -0.08018 | -0.17311 | -0.07008 | 0.09352 | 0.319393 | 0.09565 | 0.10734 | -0.1729 | 0.08259 | -0.05922 | 0.09627 | 0.065655 | -0.28663 |
| SHMAC4 | -0.07442 | 0.033903 | -0.51734 | -0.04414 | -0.11967 | -0.07347 | -0.02137 | -0.23206 | -0.04486 | -0.01093 | -0.17295 | 0.15281 | -0.201 | -0.10808 | -0.09039 | -0.0365 | -0.04456 | -0.08114 | 0.021114 | 0.15144 | 0.08032 | -0.19731 | 0.160259 | -0.07818 | 0.18732 | -0.05336 | -0.28674 |
| SHMAC5 | -0.15046 | -0.03607 | -0.08921 | -0.17816 | 0.02074 | -0.0889 | -0.19613 | -0.40032 | -0.08046 | -0.15782 | -0.01038 | 0.27495 | -0.37628 | -0.10655 | -0.10859 | -0.10702 | 0.03755 | 0.08284 | -0.02326 | 0.0436 | 0.16062 | -0.16729 | 0.169275 | -0.1002 | -0.00812 | 0.032327 | -0.3421 |
| SISO | -0.24312 | 0.008019 | -0.95333 | -0.04463 | -0.21414 | -0.07933 | -0.02514 | -0.17547 | -0.09082 | 0.02014 | -0.12101 | 0.04879 | 0.01365 | -0.15348 | -0.10086 | -0.08186 | -0.09367 | 0.02341 | 0.036342 | 0.04439 | 0.04483 | -0.08049 | 0.058468 | -0.09869 | -0.05406 | -0.02157 | -0.2449 |
| SKGI | -0.17424 | 0.028285 | -0.21342 | 0.00199 | -0.00722 | -0.36996 | -0.17007 | -0.09377 | -0.05159 | -0.04971 | -0.06657 | 0.02218 | -0.34346 | -0.0649 | -0.08408 | 0.125454 | 0.12399 | 0.113 | -0.15851 | 0.11321 | 0.20716 | -0.37035 | 0.000426 | 0.02873 | -0.16409 | 0.255753 | -0.27023 |
| SKGII | -0.10262 | 0.069585 | -0.02269 | -0.07861 | -0.07446 | -0.16669 | -0.10749 | -0.05717 | -0.15689 | -0.14035 | -0.0908 | 0.04035 | -0.229 | -0.05837 | -0.06547 | -0.08107 | -0.05919 | 0.20921 | -0.03285 | 0.07762 | 0.11669 | -0.10803 | 0.094349 | -0.04887 | 0.12506 | -0.12635 | -0.34399 |
| SKGT2 | -0.03911 | 0.087966 | -0.31612 | 0.01127 | -0.07629 | -0.14131 | -0.01645 | -0.02856 | -0.2967 | -0.10163 | -0.08679 | 0.02338 | -0.26976 | -0.08206 | -0.08431 | -0.18936 | -0.02718 | 0.10059 | 0.09113 | 0.04493 | 0.09763 | -0.14111 | 0.037448 | -0.01985 | 0.1599 | 0.127054 | -0.33129 |
| SKGT4 | -0.24313 | 0.007594 | -0.17976 | -0.14775 | -0.12646 | -0.00974 | -0.01417 | -0.36893 | -0.11965 | -0.04255 | 0.006583 | 0.13846 | -0.07993 | -0.20743 | -0.08038 | -0.09454 | -0.01782 | 0.17238 | 0.021635 | 0.24764 | 0.07244 | -0.08018 | 0.095222 | -0.03054 | 0.15367 | 0.047518 | -0.37606 |
| SKN | 0.08614 | -0.04283 | -0.26068 | 0.0304 | -0.19551 | -0.16434 | -0.07514 | -0.11385 | -0.29486 | 0.04276 | -0.10416 | 0.12982 | -0.16098 | -0.18784 | -0.12969 | -0.13724 | 0.0032 | 0.25032 | 0.123359 | 0.03253 | 0.08871 | -0.10669 | -0.00682 | -0.05691 | 0.10417 | -0.04223 | -0.19145 |
| SKNO1 | -0.01832 | -0.04425 | 0.0062 | -0.11628 | -0.25071 | -0.31823 | -1.18271 | -0.52103 | 0.045619 | 0.02785 | -0.15075 | 0.13222 | -0.27556 | -0.2417 | -0.06846 | -0.06064 | -0.26011 | 0.04418 | -0.07009 | 0.1871 | 0.25952 | -0.15589 | 0.02489 | -0.06895 | 0.28461 | 0.103607 | -0.58058 |
| SNU638 | -0.05783 | 0.093615 | -0.18115 | -0.17031 | -0.13334 | -0.06415 | 0.001627 | -0.08062 | -0.02843 | -0.13719 | -1.6943 | 0.10193 | 0.0016 | -0.16448 | -0.08803 | -0.13803 | -0.05404 | -0.16796 | 0.024444 | 0.17031 | 0.12665 | -0.12623 | 0.077667 | -0.0684 | 0.09498 | 0.004536 | -0.24359 |
| SUSA | -0.04672 | -0.00564 | -0.51162 | -0.24841 | -0.01515 | -0.00107 | -0.13432 | -0.24073 | 0.156479 | -0.16149 | -0.07869 | -0.08369 | 0.23938 | -0.35832 | -0.23674 | -0.26897 | 0.05666 | -0.02319 | -0.01264 | 0.18498 | 0.15466 | -0.11381 | -0.02903 | -0.12991 | 0.30885 | 0.226495 | -0.37719 |
| TANOUE | -0.12602 | -0.05105 | -0.25558 | 0.08558 | -0.25073 | -0.06271 | -0.04771 | -0.29784 | 0.085352 | -0.07016 | -0.1024 | 0.14591 | -0.05639 | -0.23774 | -0.0176 | -0.01928 | 0.01975 | -0.04762 | 0.081573 | 0.18202 | 0.09602 | -0.16313 | -0.05527 | 0.00478 | 0.09171 | 0.083572 | -0.4589 |
| TASK1 | -0.04899 | 0.051557 | -0.24568 | -0.56812 | -0.01308 | 0.06436 | -0.09236 | -0.15432 | 0.019817 | -0.17005 | -0.0915 | 0.05026 | -0.3962 | -0.22703 | -0.10595 | -0.06444 | -0.10135 | 0.09934 | -0.00133 | 0.23762 | 0.01929 | -0.21693 | -0.04443 | 0.0484 | 0.09787 | 0.012935 | -0.49192 |

| Cell line | FLT3 | FLT4 | IGF1R | INSR | INSRR | KDR | KIT | LMR1 | LMR2 | LTK | MET | MUSK | PDGFRA | PDGFRB | PTK7 | RET | RON | ROR1 | ROR2 | ROS1 | STYK1 | TIE1 | TIE2 | TRKA | TRKB | TRKC | TYRO3 |
|---|---|---|---|---|---|---|---|---|---|---|---|---|---|---|---|---|---|---|---|---|---|---|---|---|---|---|---|
| TFK1 | -0.04676 | 0.030876 | -0.68418 | -0.04937 | -0.06133 | -0.03514 | -0.03648 | -0.1803 | 0.020667 | 0.05753 | -0.08348 | 0.15501 | -0.03501 | -0.10735 | -0.11872 | -0.05098 | 0.00709 | 0.05996 | -0.03989 | 0.07433 | 0.10618 | -0.17456 | 0.081506 | -0.06227 | 0.08487 | -0.03339 | -0.20141 |
| TGW | -0.18071 | -0.04949 | -0.86326 | -0.04101 | -0.15758 | -0.07897 | 0.01455 | -0.43006 | -0.12466 | 0.00745 | -0.03934 | 0.21138 | -0.20802 | -0.11991 | -0.10937 | -0.04646 | -0.03945 | -0.13475 | -0.05653 | 0.14478 | 0.20816 | -0.16164 | 0.038752 | -0.07602 | 0.15863 | 0.036048 | -0.47288 |
| TR146 | -0.16124 | -0.03903 | 0.00532 | -0.22812 | -0.16452 | -0.08367 | -0.03335 | -0.2145 | -0.13177 | -0.02305 | -0.15434 | 0.05553 | -0.12128 | -0.29884 | 0.072331 | -0.10847 | -0.01462 | -0.09561 | -0.13179 | 0.11421 | 0.14513 | -0.08882 | 0.19432 | 0.0474 | 0.11439 | 0.141454 | -0.65515 |
| U2904 | -0.01707 | 0.055495 | -0.04599 | -0.15121 | 0.1542 | -0.20237 | -0.01355 | -0.00888 | 0.007719 | 0.07904 | -0.07248 | 0.0985 | -0.11567 | 0.05194 | 0.059952 | -0.06117 | 0.04364 | 0.0851 | 0.1418 | 0.19948 | 0.07062 | -0.33061 | 0.187386 | 0.12282 | 0.05683 | 0.209137 | -0.27298 |
| UHO1 | -0.2212 | 0.086154 | -0.15172 | 0.06684 | -0.04941 | -0.07541 | -0.08024 | -0.30354 | -0.23434 | 0.07406 | -0.09202 | 0.09705 | -0.19777 | -0.08358 | 0.061948 | 0.045132 | 0.09203 | 0.05636 | 0.067855 | 0.09269 | 0.19197 | -0.08404 | 0.150311 | -0.09219 | 0.16008 | 0.093631 | -0.27332 |
| UMRC3 | 0.01327 | -0.04908 | -0.1736 | -0.0658 | -0.13819 | 0.07566 | -0.047 | -0.10656 | -0.09909 | -0.13364 | -0.183 | 0.12773 | -0.2946 | -0.48874 | -0.21756 | -0.09656 | -0.04619 | 0.03287 | 0.097117 | 0.09057 | 0.02441 | -0.17044 | 0.025464 | -0.00029 | 0.07763 | 0.075811 | -0.55614 |
| UMRC7 | -0.07655 | 0.006173 | -0.53954 | -0.15606 | -0.31327 | -0.01154 | -0.14194 | 0.00436 | -0.02994 | -0.1239 | -0.15804 | 0.17351 | -0.1987 | -0.39116 | -0.08127 | -0.12688 | -0.23448 | -0.16067 | -0.06871 | 0.11128 | 0.00773 | -0.18134 | 0.254568 | 0.00407 | 0.29944 | -0.04021 | -0.31482 |
| UPCISCC026 | -0.14069 | 0.053307 | -0.11559 | -0.0832 | -0.10559 | -0.08857 | 0.028225 | -0.26316 | -0.04562 | -0.19864 | -0.1156 | 0.02267 | -0.18708 | -0.16023 | -0.03993 | -0.01472 | -0.00768 | 0.13222 | -0.12765 | 0.01453 | 0.11198 | -0.08162 | 0.041967 | -0.10385 | 0.17615 | -0.00389 | -0.35084 |
| UPCISCC029A | 0.04017 | 0.006949 | -0.55333 | -0.13279 | -0.18294 | -0.05835 | -0.01635 | -0.3454 | -0.01851 | -0.04276 | -0.15718 | 0.10373 | -0.19101 | -0.50992 | 0.001576 | -0.02375 | -0.05423 | -0.07753 | 0.146211 | 0.05407 | 0.12118 | -0.07681 | 0.050239 | -0.08544 | 0.16583 | -0.113 | -0.37012 |
| UPCISCC040 | -0.16248 | 0.050847 | -0.16071 | -0.09009 | -0.15196 | 0.02122 | 0.018833 | -0.15366 | -0.11598 | -0.09319 | -0.1264 | 0.064 | -0.05255 | -0.1472 | -0.10321 | -0.11004 | -0.04535 | -0.03137 | -0.01279 | 0.16388 | 0.07678 | -0.36905 | 0.02714 | -0.07273 | 0.02892 | 0.065655 | -0.56018 |
| UPCISCC074 | -0.12141 | -0.01593 | -0.20535 | -0.05514 | -0.14863 | -0.09763 | -0.06908 | -0.22011 | -0.0142 | -0.10521 | -0.1449 | 0.12077 | -0.18654 | -0.16929 | -0.17302 | -0.08955 | -0.03296 | -0.02055 | -0.05213 | 0.16486 | 0.1171 | -0.14873 | 0.056526 | -0.04365 | 0.09396 | 0.029505 | -0.32438 |
| UPCISCC111 | -0.20185 | 0.049806 | -0.17666 | -0.03382 | -0.1199 | 0.01379 | -0.17508 | -0.16127 | -0.29572 | -0.03884 | -0.15693 | 0.17073 | -0.25126 | -0.12255 | -0.09211 | -0.11292 | -0.09903 | -0.09174 | -0.06905 | 0.19375 | 0.19553 | -0.12089 | 0.094361 | -0.02152 | 0.12968 | -0.02866 | -0.36812 |
| UPCISCC116 | -0.10461 | 0.131881 | -0.73674 | -0.14004 | -0.03714 | -0.15114 | 0.03554 | -0.1415 | -0.26355 | -0.07086 | 0.076448 | 0.13253 | -0.10304 | -0.19419 | -0.11045 | -0.01141 | 0.01829 | 0.10781 | -0.07973 | 0.01086 | 0.11153 | -0.34951 | 0.269779 | 0.06541 | 0.16031 | 0.115083 | -0.29641 |
| UPCISCC131 | -0.05369 | -0.06411 | -0.12086 | -0.00579 | -0.19758 | -0.06568 | -0.00379 | -0.35721 | 0.080167 | -0.14554 | -0.16859 | 0.13242 | -0.10027 | -0.29264 | -0.04645 | -0.1146 | -0.02501 | 0.13128 | 0.032113 | 0.13942 | 0.15738 | -0.42656 | 0.184871 | -0.02016 | 0.09653 | 0.067109 | -0.44702 |
| UPCISCC200 | -0.31971 | 0.072216 | -0.07163 | -0.10703 | -0.03597 | -0.12629 | -0.06793 | -0.28456 | -0.29984 | -0.20088 | 0.01764 | 0.07399 | -0.32552 | -0.20613 | 0.024687 | -0.2839 | 0.0552 | 0.02896 | -0.07908 | 0.07872 | 0.03553 | -0.39675 | 0.470694 | -0.01586 | 0.03644 | 0.142157 | -0.78417 |
| VAESBJ | -0.14009 | 0.040137 | -0.09479 | -0.21902 | -0.14192 | -0.01831 | -0.09746 | -0.39937 | -0.31056 | -0.13352 | -0.34318 | 0.01474 | -0.32387 | -0.30402 | -0.05903 | -0.10916 | -0.03718 | 0.03941 | -0.01787 | 0.18428 | 0.07474 | -0.01856 | 0.114241 | -0.06927 | 0.01375 | 0.029402 | -0.54752 |
| WAOSEL | -0.08648 | -0.11445 | -0.06555 | -0.09156 | -0.11903 | -0.01036 | 0.072857 | -0.21529 | -0.14144 | -0.11508 | -0.17771 | -0.1648 | 0.01771 | -0.29535 | -0.20926 | -0.04423 | 0.15351 | 0.13526 | 0.031923 | 0.08387 | 0.23229 | -0.14499 | 0.178504 | -0.16726 | 0.22976 | 0.172496 | -0.71773 |
| WSUNHL | -0.09372 | 0.094338 | -0.12656 | -0.08555 | -0.10281 | -0.02552 | 0.088793 | -0.14709 | -0.02599 | -0.08281 | -0.10888 | 0.19926 | 0.03994 | -0.12737 | -0.11949 | -0.01317 | -0.09611 | 0.004 | -0.10951 | 0.30014 | 0.06062 | -0.23844 | -0.04854 | -0.13471 | 0.10749 | 0.0267 | -0.42275 |
| PFSK1 | -0.26617 | 0.587639 | -0.6744 | -0.24442 | -0.38947 | -0.52575 | -0.1282 | -0.08581 | -0.61967 | 0.40282 | -0.15859 | -0.19013 | -0.03232 | -0.05802 | -0.06798 | -0.12437 | 0.08757 | -0.0536 | -0.51562 | -0.04541 | 0.18153 | -0.2666 | 0.136006 | -0.33794 | -0.0497 | -0.16372 | -0.23266 |
| CAL72 | -0.17541 | 0.227454 | -0.14656 | 0.07841 | -0.13863 | -0.37431 | 0.058844 | -0.06608 | -0.02139 | 0.08335 | -0.15032 | 0.15516 | 0.01356 | 0.03346 | -0.06597 | -0.06506 | -0.17777 | 0.04412 | -0.0224 | 0.09015 | 0.07584 | -0.20472 | 0.005038 | 0.07912 | 0.11698 | 0.153711 | -0.24061 |
| OCIC4P | -0.21888 | -0.03427 | -0.09956 | -0.17832 | -0.05628 | 0.01474 | 0.004272 | -0.1179 | -0.10115 | -0.00815 | -0.17865 | 0.09541 | -0.17358 | -0.14845 | -0.03463 | -0.11947 | -0.04932 | 0.10516 | -0.05348 | 0.17227 | 0.09362 | -0.1938 | 0.078244 | -0.10262 | 0.21722 | 0.070655 | -0.31922 |
| SEMK2 | -1.53891 | -0.09799 | -0.23959 | -0.09789 | -0.31938 | -0.15326 | 0.063342 | -0.1897 | -0.23235 | 0.01111 | 0.027939 | 0.07976 | -0.16029 | -0.08896 | -0.22912 | -0.13877 | -0.08269 | 0.32305 | -0.03143 | 0.09126 | 0.13527 | -0.26997 | 0.046331 | 0.01085 | 0.0515 | -0.02141 | -0.52589 |
| HB1119 | -1.38089 | -0.07494 | -0.27698 | -0.00332 | -0.07439 | -0.14295 | 0.133955 | -0.13996 | -0.09171 | 0.06671 | -0.06512 | 0.05775 | -0.10444 | -0.04688 | -0.06667 | -0.02623 | -0.03074 | -0.02745 | -0.01554 | 0.09474 | 0.10208 | -0.11991 | -0.05863 | -0.02382 | -0.00081 | 0.11335 | -0.34156 |
| CTV1DM | -0.35252 | -0.22239 | -0.19156 | 0.01074 | -0.23975 | -0.27173 | 0.043004 | -0.28478 | 0.000219 | 0.02747 | -0.00566 | 0.12496 | 0.01709 | -0.01128 | -0.14807 | -0.30894 | -0.10782 | 0.10704 | 0.103947 | 0.21949 | 0.10845 | -0.29946 | 0.06045 | 0.1954 | 0.17676 | 0.045746 | -0.66278 |
| RH28 | -0.05061 | 0.076961 | -0.45246 | 0.01019 | -0.24649 | -0.04813 | 0.166188 | -0.04111 | -0.00635 | -0.21313 | -0.10354 | 0.11279 | -0.02408 | 0.29591 | -0.03121 | -0.01647 | -0.21384 | 0.1276 | 0.021422 | 0.22846 | 0.05126 | -0.11891 | 0.188298 | -0.14348 | 0.23297 | -0.02648 | -0.48658 |
| RHJT | -0.1001 | 0.214437 | -0.94419 | -0.11516 | -0.08655 | -0.03234 | -0.03191 | -0.14022 | -0.11748 | -0.1389 | -0.19195 | 0.12541 | -0.09786 | -0.00577 | -0.04277 | -0.10375 | -0.11068 | -0.03064 | 0.07627 | 0.05861 | 0.13511 | -0.07058 | 0.059876 | 0.01923 | 0.26405 | -0.00506 | -0.49692 |
| TTC442 | 0.00401 | -0.01499 | -0.56072 | 0.00867 | -0.26132 | -0.14914 | -0.01669 | -0.15632 | -0.21895 | -0.00528 | -0.14178 | 0.29217 | -0.23828 | -0.13751 | -0.07335 | -0.08162 | -0.16664 | 0.19187 | 0.079549 | 0.15516 | 0.10167 | 0.009904 | 0.102603 | -0.08959 | 0.32223 | -0.0127 | -0.17154 |
| RH4 | -0.08006 | -0.01612 | -0.86999 | -0.05082 | -0.16277 | -0.06244 | -0.00439 | -0.15388 | -0.04627 | -0.11037 | -0.10087 | 0.09162 | -0.20716 | -0.0663 | -0.27242 | -0.09128 | -0.12882 | 0.01244 | -0.02802 | 0.1252 | 0.0303 | -0.13777 | 0.151982 | -0.12569 | 0.11705 | 0.014007 | -0.3251 |
| SNU1544 | -0.52284 | -0.28219 | -0.05738 | -0.12128 | 0.02078 | 0.00263 | -0.12448 | -0.18042 | -0.13019 | -0.10058 | 0.141873 | 0.03669 | -0.26811 | -0.23571 | -0.02086 | -0.03624 | 0.06299 | 0.15667 | -0.06711 | 0.14526 | 0.16311 | -0.37009 | -0.00402 | -0.02991 | 0.028 | 0.188308 | -0.42408 |
| LPS6 | -0.0772 | -0.19964 | -0.1798 | 0.01116 | -0.23249 | -0.14232 | -0.14062 | -0.02478 | -0.17846 | 0.13077 | -0.06318 | 0.16838 | -0.53953 | 0.09682 | 0.067639 | -0.09375 | -0.10172 | -0.01557 | 0.016892 | -0.01773 | 0.09224 | -0.19925 | 0.168861 | -0.06494 | 0.0942 | 0.1107 | -0.58745 |
| LPS27 | -0.0405 | 0.049666 | -0.20453 | 0.07775 | -0.12732 | -0.25988 | -0.10099 | -0.28125 | -0.10118 | -0.0348 | -0.10969 | 0.07099 | -0.34108 | -0.1695 | -0.08698 | -0.10359 | -0.07228 | 0.13618 | 0.069555 | 0.03671 | -0.19634 | -0.12236 | 0.076804 | -0.00474 | 0.10874 | 0.008646 | -0.34014 |
| 93T449 | -0.05352 | 0.001512 | -0.21113 | 0.03341 | -0.18217 | -0.06185 | 0.04315 | -0.27669 | 0.088909 | -0.0794 | -0.1131 | 0.16991 | -0.07756 | -0.1355 | -0.08951 | -0.03402 | -0.20592 | 0.12179 | -0.0982 | 0.22769 | 0.08218 | -0.10165 | 0.056491 | 0.03119 | 0.17585 | -0.00457 | -0.33956 |
| 94T778 | -0.14187 | 0.095883 | -0.12259 | -0.11735 | -0.18378 | -0.07441 | -0.02518 | -0.28654 | -0.17968 | -0.04063 | -0.12679 | 0.15902 | -0.17164 | -0.15071 | -0.20233 | -0.15864 | -0.01163 | -0.13703 | 0.003873 | 0.0686 | 0.18096 | -0.0867 | 0.115323 | -0.11187 | 0.09662 | 0.05547 | -0.30006 |
| 95T1000 | -0.10681 | -0.06191 | -0.14771 | -0.06347 | -0.00614 | -0.05465 | -0.10479 | -0.34822 | -0.255 | 0.16283 | -0.03743 | 0.08412 | -0.1509 | 0.03258 | -0.2462 | -0.02144 | -0.03754 | 0.33554 | -0.03295 | 0.10849 | 0.20141 | -0.28409 | 0.132939 | 0.0182 | 0.12057 | -0.11823 | -0.19882 |
| LPS141 | -0.27026 | 0.048213 | -0.30221 | -0.16732 | -0.11894 | -0.12471 | 0.066227 | -0.13138 | 0.187133 | 0.27558 | 0.038179 | -0.13391 | -0.19713 | -0.21782 | -0.0058 | -0.02326 | -0.12913 | -0.07255 | 0.164806 | 0.20403 | 0.13823 | -0.15552 | 0.193812 | -0.07707 | 0.18049 | -0.03576 | -0.08115 |
| LPS853 | -0.18843 | 0.016601 | -0.354 | 0.01127 | -0.11679 | -0.09274 | 0.001305 | -0.05659 | 0.057808 | 0.02246 | -0.10924 | -0.03195 | -0.18239 | -0.07375 | -0.00082 | -0.05945 | -0.13675 | 0.06337 | -0.01296 | 0.07522 | 0.09299 | -0.08682 | 0.08937 | -0.12455 | 0.10766 | -0.06067 | -0.30669 |
| LPS510 | -0.24106 | 0.062019 | -0.23718 | -0.13798 | -0.13166 | 0.0056 | 0.049849 | -0.17501 | -0.07231 | 0.06148 | -0.26895 | -0.0782 | -0.28769 | -0.28445 | 0.116904 | -0.1369 | 0.0176 | -0.14514 | -0.0676 | 0.37555 | 0.26878 | -0.00543 | 0.21076 | 0.00772 | 0.05849 | -0.2052 | -0.20521 |
| OS252 | -0.00088 | 0.020839 | -0.09095 | -0.22129 | -0.14646 | -0.29179 | 0.189673 | -0.05122 | -0.14056 | 0.20865 | 0.013285 | 0.1094 | -0.10874 | 0.14741 | 0.15066 | 0.11157 | -0.07275 | 0.14869 | 0.015824 | 0.14721 | 0.0489 | -0.1533 | 0.168464 | 0.20807 | 0.17716 | -0.04921 | -0.14837 |
| MFM223 | -0.08816 | 0.014368 | -0.12267 | -0.07375 | -0.10155 | -0.02196 | -0.17024 | -0.22666 | -0.07999 | 0.02319 | -0.15031 | 0.14578 | -0.16647 | -0.06727 | 0.000487 | 0.057976 | -0.09369 | 0.01595 | 0.01255 | 0.18612 | 0.10429 | -0.19319 | 0.099044 | -0.02367 | 0.13818 | 0.010515 | -0.23187 |
| COLO824 | -0.01372 | -0.02528 | -0.83729 | -0.07737 | -0.11648 | 0.03346 | -0.02055 | -0.19392 | 0.084048 | 0.02591 | -0.12347 | 0.11018 | -0.16694 | -0.21634 | -0.2123 | -0.28928 | -0.00737 | 0.09953 | -0.09762 | 0.27116 | 0.03207 | -0.16451 | 0.086381 | 0.03761 | 0.14192 | -0.09279 | -0.19719 |
| ICC10 | 0.00785 | 0.007649 | -0.0676 | 0.02555 | -0.01507 | -0.1173 | -0.08418 | -0.11895 | -0.25519 | -0.01449 | -0.10543 | 0.07237 | -0.31283 | -0.17615 | -0.00089 | -0.09321 | -0.0809 | 0.1748 | -0.05967 | 0.07017 | 0.2314 | -0.14377 | 0.145201 | -0.01739 | 0.10246 | 0.008481 | -0.27701 |
| ICC106 | -0.20335 | 0.059743 | -0.11125 | 0.01888 | -0.04781 | -0.05079 | -0.07831 | -0.23818 | -0.21724 | 0.039 | -0.18113 | 0.12717 | -0.17467 | -0.10482 | -0.04524 | -0.17886 | -0.03836 | 0.1067 | 0.132936 | 0.25204 | 0.15593 | -0.30265 | 0.140913 | -0.01374 | 0.11185 | 0.075906 | -0.27425 |
| ICC108 | -0.05357 | -0.03522 | -0.09899 | -0.13167 | -0.01071 | -0.22401 | -0.10718 | -0.0429 | -0.12598 | -0.20372 | -0.04622 | 0.12673 | -0.32192 | -0.13417 | -0.21242 | 0.007564 | 0.09109 | -0.046 | 0.051576 | 0.10587 | 0.07832 | -0.1674 | 0.216787 | -0.16753 | 0.09676 | -0.02427 | -0.24028 |
| ICC12 | 0.01836 | 0.079504 | -0.6567 | -0.10598 | -0.15382 | -0.21028 | 0.041101 | -0.24117 | -0.30436 | -0.19144 | 0.084829 | 0.06101 | -0.06493 | -0.0531 | -0.12683 | -0.12719 | -0.14014 | 0.06017 | -0.01427 | 0.14161 | -0.07663 | -0.0208 | 0.176445 | -0.11417 | 0.11933 | 0.020692 | -0.45913 |
| ICC137 | -0.19755 | 0.032299 | -0.38433 | -0.18981 | -0.25267 | -0.18527 | -0.27162 | -0.02525 | -0.05988 | -0.21453 | -0.10474 | 0.02831 | -0.19354 | -0.08013 | -0.15781 | -0.12213 | 0.02585 | 0.1879 | -0.15336 | 0.16815 | 0.12869 | -0.18497 | 0.057264 | -0.08267 | 0.03208 | 0.052135 | -0.38216 |
| ICC15 | -0.07566 | -0.01083 | -0.2924 | -0.19682 | -0.0438 | -0.11178 | -0.03855 | -0.1414 | -0.00753 | -0.06776 | -0.10012 | 0.08302 | -0.18122 | -0.04495 | -0.08625 | -0.08545 | -0.05136 | -0.02801 | -0.2083 | 0.13462 | 0.02572 | -0.18067 | 0.213874 | 0.05108 | 0.09171 | 0.045704 | -0.27837 |
| ICC2 | 0.14686 | -0.00704 | -0.21523 | -0.05496 | -0.11625 | -0.64449 | -0.08122 | -0.15177 | -0.49373 | 0.23208 | -0.20012 | 0.11419 | 0.06366 | 0.17507 | -0.18414 | -0.32987 | -0.12291 | 0.12835 | 0.101702 | 0.17493 | 0.24475 | -0.26831 | 0.090553 | 0.0028 | 0.31171 | 0.055847 | -0.32635 |
| ICC3 | -0.119 | 0.07527 | -0.28473 | 0.11635 | -0.04665 | -0.12205 | 0.160685 | -0.18234 | 0.019277 | -0.06066 | 0.037333 | 0.20115 | -0.17571 | -0.17834 | 0.23314 | 0.007279 | 0.02221 | 0.39782 | -0.06899 | 0.37183 | 0.04504 | -0.20005 | 0.036195 | 0.20943 | 0.06843 | 0.067517 | -0.32272 |
| ICC4 | -0.10142 | -0.06836 | -0.31168 | -0.2667 | -0.19969 | 0.06643 | 0.03017 | -0.46487 | -0.06738 | -0.12562 | -0.16622 | 0.09325 | -0.35148 | -0.11468 | -0.18378 | -0.30601 | -0.05891 | 0.15593 | -0.1515 | 0.30464 | 0.17473 | -0.13061 | 0.000109 | -0.1466 | 0.07194 | -0.03493 | -0.16721 |
| ICC8 | -0.10985 | -0.09799 | -0.17335 | -0.06648 | -0.01146 | -0.07074 | -0.1276 | -0.35241 | 0.025429 | -0.07998 | -0.08419 | 0.13788 | -0.09867 | -0.10168 | -0.1559 | -0.1411 | -0.00146 | -0.00801 | 0.006352 | 0.18053 | 0.06965 | -0.18646 | 0.058026 | -0.02172 | 0.13825 | 0.041551 | -0.25664 |
| ICC9 | -0.16198 | 0.02561 | -0.44738 | -0.33816 | -0.07533 | -0.13523 | -0.09351 | -0.11093 | -0.22903 | -0.06613 | -0.01227 | 0.07314 | -0.10062 | -0.19268 | -0.11886 | -0.12898 | -0.04711 | 0.0751 | -0.05294 | 0.11226 | 0.05579 | -0.2106 | 0.156222 | -0.05729 | 0.0076 | -0.03576 | -0.289 |
| G415 | -0.07312 | -0.03895 | -0.23185 | -0.13008 | -0.23607 | -0.12927 | -0.08453 | -0.18792 | -0.20687 | -0.06198 | -0.02267 | 0.17549 | -0.1797 | -0.1874 | -0.05816 | -0.03937 | 0.01714 | 0.02561 | 0.033843 | 0.08868 | 0.11348 | -0.13558 | 0.152156 | -0.15508 | 0.13627 | -0.10676 | -0.25892 |
| HKGZCC | -0.01796 | -0.02089 | -0.31268 | -0.06026 | -0.0807 | -0.04864 | 0.082865 | -0.29095 | 0.021784 | -0.08489 | -0.01704 | 0.10581 | -0.0434 | -0.05506 | -0.16572 | -0.06287 | -0.03871 | 0.08389 | 0.00146 | 0.10538 | 0.11445 | -0.15394 | 0.080463 | -0.07996 | 0.11394 | -0.01897 | -0.22754 |
| KMCH1 | -0.10965 | 0.082384 | -0.42768 | -0.052 | -0.71382 | -0.3709 | 0.026196 | -0.05004 | 0.035659 | 0.04959 | 0.06925 | 0.13551 | -0.29086 | -0.23594 | -0.32374 | -0.28513 | 0.03848 | 0.0771 | -0.11555 | 0.07793 | 0.04896 | -0.18502 | 0.142392 | -0.11349 | -0.23089 | -0.14056 | -0.30099 |
| RBE | -0.06417 | 0.120629 | -0.48551 | -0.04763 | -0.0964 | -0.15141 | -0.22573 | -0.17463 | -0.03892 | -0.09037 | 0.016998 | 0.19182 | -0.23845 | -0.07721 | -0.09397 | -0.07686 | -0.02144 | -0.0548 | 0.012178 | 0.18305 | 0.18771 | -0.14094 | 0.105081 | -0.14285 | 0.08102 | 0.0071 | -0.21303 |
| SG231 | -0.24341 | 0.079908 | -0.19019 | -0.16578 | -0.12503 | 0.05232 | 0.08462 | -0.2531 | -0.57615 | -0.269 | -0.01773 | 0.15815 | -0.26656 | -0.07318 | -0.04732 | -0.06379 | -0.15215 | 0.20005 | -0.19997 | 0.20578 | 0.02809 | -0.35275 | 0.196187 | -0.02842 | 0.11177 | -0.24547 | -0.42008 |
| SSP25 | -0.06546 | 0.040466 | -0.43305 | -0.08162 | -0.16929 | -0.14389 | 0.037064 | -0.13712 | -0.28852 | -0.0752 | -0.1652 | 0.07578 | -0.20594 | -0.14356 | -0.11978 | -0.08778 | -0.08206 | 0.06601 | -0.00101 | 0.11275 | 0.04381 | -0.12073 | 0.086909 | 0.03866 | 0.06089 | 0.022014 | -0.25305 |
| TGBC1TKB | -0.16699 | 0.017113 | -0.06052 | -0.14898 | -0.14723 | 0.04718 | -0.03319 | -0.14182 | -0.1364 | 0.0828 | 0.043382 | 0.03865 | -0.11141 | -0.04481 | -0.34872 | -0.12509 | 0.0751 | -0.20882 | 0.201496 | 0.23027 | 0.15338 | -0.32586 | 0.055544 | -0.11868 | 0.11309 | -0.23998 | -0.13846 |
| TGBC52TKB | 0.03109 | -0.11692 | -0.59677 | -0.13042 | -0.09219 | 0.00124 | -0.04471 | -0.10492 | 0.134898 | 0.0493 | -0.04539 | 0.14134 | -0.05081 | -0.19287 | -0.04545 | 0.003242 | -0.02813 | 0.06391 | 0.104212 | 0.2242 | -0.01315 | -0.10914 | 0.140969 | -0.00442 | 0.08011 | -0.0497 | -0.13037 |
| TKKK | -0.12466 | -0.08855 | -0.26778 | -0.07029 | -0.12535 | -0.13043 | 0.053778 | -0.26708 | 0.081722 | -0.17039 | -0.16008 | 0.3378 | 0.01319 | -0.22674 | -0.13997 | 0.025272 | 0.11942 | -0.11433 | 0.032263 | 0.13594 | 0.29598 | -0.39065 | 0.140425 | 0.20567 | 0.32193 | 0.045323 | -0.27397 |
| YSCCC | -0.04788 | -0.01896 | -0.08531 | -0.04123 | 0.01574 | -0.1919 | 0.079475 | -0.17345 | -0.04451 | 0.06285 | -0.07016 | 0.02051 | -0.13455 | -0.07055 | 0.098828 | -0.14344 | -0.03666 | -0.08538 | -0.1914 | 0.06435 | 0.08993 | -0.34904 | -0.01137 | -0.03709 | 0.01477 | 0.062759 | -0.46014 |
| CCLP1 | -0.09827 | 0.068795 | -0.69758 | -0.03439 | -0.04317 | -0.1638 | 0.039731 | -0.17991 | -0.02299 | -0.09649 | -0.14264 | 0.0701 | -0.16312 | -0.0003 | -0.04482 | -0.06348 | 0.0113 | 0.13141 | 0.020256 | 0.04563 | 0.13549 | -0.09947 | 0.187025 | -0.12502 | 0.08825 | 0.003271 | -0.27122 |
| CCSW1 | -0.13503 | 0.085633 | -0.44865 | -0.08974 | -0.08616 | -0.2433 | -0.04814 | -0.16509 | -0.32191 | -0.00865 | -0.4461 | 0.07578 | -0.13886 | 0.10083 | -0.04637 | -0.11303 | 0.06146 | 0.11571 | -0.03206 | 0.05567 | 0.09467 | -0.17238 | 0.131845 | -0.0984 | 0.17921 | 0.004183 | -0.54321 |
| GB2 | -0.06101 | 0.07939 | -0.29885 | -0.07461 | -0.22995 | -0.01234 | 0.073515 | -0.38933 | -0.08319 | -0.11881 | -0.13198 | 0.03965 | -0.01844 | -0.15789 | -0.03882 | 0.006655 | 0.0407 | 0.11502 | 0.077795 | 0.0979 | -0.00157 | -0.38123 | 0.245075 | -0.05749 | 0.04007 | -0.02929 | -0.39323 |
| MM253 | -0.10243 | -0.10755 | -0.25426 | -0.00761 | -0.15557 | 0.21876 | -0.09794 | -0.64459 | 0.023273 | 0.02502 | -0.12401 | 0.22817 | -0.20991 | -0.18321 | 0.016616 | 0.011344 | 0.02411 | 0.09194 | -0.08855 | 0.14158 | 0.18276 | -0.32873 | 0.225571 | 0.04 | 0.38066 | -0.07131 | -0.97618 |
| MM485 | 0.05698 | 0.213518 | -0.37891 | -0.21239 | -0.04045 | -0.00508 | -0.02556 | -0.17457 | 0.093834 | -0.034 | -0.05707 | 0.22808 | -0.28793 | 0.07207 | -0.28332 | 0.071578 | -0.19168 | -0.18534 | -0.32978 | 0.12728 | 0.14454 | -0.01755 | 0.079147 | 0.08256 | 0.29031 | 0.028794 | -0.21018 |
| NO36 | -0.08158 | -0.09435 | -0.36297 | -0.21281 | -0.14824 | -0.03651 | -0.02734 | -0.17637 | -0.22316 | 0.1079 | -0.07175 | 0.20683 | -0.07551 | -0.2328 | 0.05374 | 0.017753 | -0.06176 | -0.12598 | -0.00105 | 0.00337 | 0.15939 | -0.12092 | 0.001386 | 0.0409 | 0.18816 | -0.06213 | -0.42 |
| NZM3 | -0.12386 | 0.092291 | -0.25015 | -0.04267 | -0.30217 | -0.01992 | -0.11254 | -0.09805 | -0.09039 | 0.11658 | -0.16944 | -0.01867 | -0.09678 | -0.23991 | 0.036248 | -0.09638 | -0.12236 | -0.03665 | -0.11702 | 0.15002 | 0.1505 | -0.27298 | 0.037191 | 0.00215 | 0.09609 | -0.04116 | -0.34824 |
| NZM42 | -0.15394 | 0.056438 | -0.16638 | -0.26731 | -0.12826 | 0.07939 | -0.06012 | -0.27271 | 0.027998 | -0.00806 | -0.06796 | 0.20935 | -0.27439 | -0.1692 | 0.105286 | 0.030551 | -0.07989 | -0.13482 | 0.124513 | 0.24597 | 0.0441 | -0.21443 | 0.058797 | 0.07919 | 0.12931 | -0.03732 | -0.17445 |
| NZM7 | -0.16196 | -0.04239 | -0.27946 | -0.25298 | -0.06766 | -0.00365 | -0.06596 | -0.19759 | -0.10822 | -0.00974 | -0.19488 | 0.10967 | -0.11804 | -0.15139 | -0.02745 | -0.11405 | -0.10699 | -0.07654 | -0.04789 | 0.0627 | 0.11587 | -0.21983 | 0.096826 | -0.01718 | 0.06501 | -0.14067 | -0.33816 |
| NZOV9 | 0.10273 | -0.00741 | -0.05367 | -0.19498 | 0.02786 | 0.31248 | 0.095912 | -0.23569 | -0.19051 | -0.13521 | 0.058444 | 0.24321 | -0.34177 | -0.17138 | -0.1383 | 0.055655 | 0.05809 | 0.05446 | 0.211208 | 0.1291 | 0.35584 | -0.07549 | 0.122785 | -0.07217 | 0.26019 | 0.063158 | -0.07555 |
| ONE58 | -0.1116 | -0.00621 | -0.70145 | -0.13041 | -0.1922 | -0.04525 | -0.11132 | -0.2358 | -0.12406 | -0.04743 | -0.1319 | 0.15846 | -0.09998 | -0.17894 | -0.16952 | -0.09927 | -0.0907 | 0.08481 | -0.05483 | 0.21782 | 0.23773 | -0.15542 | 0.300264 | -0.04997 | 0.13111 | -0.00523 | -0.23006 |
| NALM16 | -0.22021 | -0.20383 | -0.41812 | -0.09974 | -0.16246 | -0.04955 | 0.024155 | -0.34508 | -0.28003 | 0.02862 | -0.16914 | 0.08331 | -0.17456 | -0.0914 | -0.07666 | -0.09686 | -0.0782 | 0.07071 | 0.10039 | 0.21987 | 0.1581 | -0.35069 | 0.016393 | -0.02693 | 0.28625 | -0.00805 | -0.55876 |
| ECC2 | -0.10119 | 0.074483 | -0.28278 | -0.17215 | -0.14275 | -0.17179 | -0.07356 | -0.14786 | -0.12751 | -0.21735 | 0.18177 | 0.01905 | -0.18845 | 0.00428 | -0.08281 | 0.157117 | 0.0392 | 0.01065 | 0.169994 | 0.02654 | 0.18786 | -0.13034 | 0.016142 | 0.06808 | 0.07655 | 0.031648 | -0.35013 |

| Cell line | FLT3 | FLT4 | IGF1R | INSR | INSRR | KDR | KIT | LMR1 | LMR2 | LTK | MET | MUSK | PDGFRA | PDGFRB | PTK7 | RET | RON | ROR1 | ROR2 | ROS1 | STYK1 | TIE1 | TIE2 | TRKA | TRKB | TRKC | TYRO3 |
|---|---|---|---|---|---|---|---|---|---|---|---|---|---|---|---|---|---|---|---|---|---|---|---|---|---|---|---|
| 9505BIK | -0.05312 | 0.030845 | -0.12172 | -0.05574 | -0.05194 | -0.15435 | 0.020794 | -0.26626 | 0.118652 | -0.16518 | 2.55E-05 | 0.12256 | -0.0927 | 0.11369 | -0.08997 | -0.04145 | -0.03092 | 0.15139 | -0.07807 | 0.03719 | 0.30272 | 0.003171 | -0.00744 | 0.13389 | 0.27042 | -0.07285 | -0.25886 |
| A375SKINCJ1 | -0.17461 | 0.045533 | -0.56894 | -0.0802 | -0.083 | -0.11484 | -0.11267 | -0.10302 | -0.00698 | 0.21877 | 0.006728 | 0.21048 | -0.10124 | -0.13529 | 0.047526 | -0.13818 | 0.05065 | 0.32682 | 0.072549 | 0.1143 | 0.08515 | 0.058598 | 0.072214 | -0.15047 | 0.24794 | -0.19995 | -0.41686 |
| A375SKINCJ2 | -0.16931 | 0.136935 | -0.26105 | -0.05577 | -0.13745 | 0.02672 | 0.075639 | -0.13207 | -0.14437 | 0.11219 | -0.05954 | 0.03651 | -0.19226 | -0.09076 | -0.1418 | -0.08253 | 0.00123 | 0.12113 | -0.03287 | 0.07538 | 0.03447 | -0.07147 | 0.08195 | 0.00068 | 0.05563 | -0.07144 | -0.29877 |
| A375SKINCJ3 | 0.0488 | 0.224228 | -0.47855 | -0.3987 | -0.28308 | 0.18729 | -0.05428 | -0.00571 | -0.22053 | -0.1721 | -0.25419 | 0.18795 | -0.09471 | -0.23488 | 0.055791 | 0.002819 | -0.26244 | 0.37096 | 0.021 | 0.05251 | 0.14207 | -0.15402 | -0.08812 | -0.11541 | -0.0053 | 0.0101 | -0.35277 |
| SKMEL19 | -0.13927 | -0.11244 | -0.4482 | -0.26655 | 0.0646 | -0.16241 | -0.21128 | -0.07894 | 0.098088 | 0.22303 | -0.09811 | 0.15315 | -0.51264 | 0.19415 | 0.047983 | -0.05787 | -0.16664 | 0.268 | -0.00431 | -0.16531 | -0.0387 | -0.16084 | 0.250682 | 0.02172 | 0.07131 | -0.22077 | -0.26212 |
| MEL270 | 0.02576 | 0.154618 | -0.92386 | -0.07312 | -0.40405 | -0.06174 | 0.394588 | -0.34314 | 0.131161 | -0.02642 | 0.22645 | 0.10018 | 0.27873 | 0.12533 | -0.01811 | -0.22991 | -0.00367 | -0.48503 | -0.16672 | -0.013 | 0.20063 | 0.094622 | 0.471418 | 0.05851 | 0.43036 | -0.25845 | -0.09083 |
| MEL285 | -0.14817 | -0.00814 | -0.13067 | -0.06448 | -0.01099 | -0.05983 | -0.03098 | -0.2755 | -0.02821 | -0.07983 | -0.12959 | 0.15076 | -0.1791 | -0.19959 | -0.03568 | -0.07015 | -0.04634 | -0.00877 | 0.080378 | 0.09733 | -0.05171 | -0.20166 | 0.164968 | 0.02721 | -0.02351 | 0.041016 | -0.42852 |
| MEL290 | -0.01032 | 0.10838 | -0.64612 | -0.14284 | -0.10263 | -0.12229 | -0.01255 | -0.2381 | 0.075839 | -0.13071 | -0.08699 | 0.13888 | -0.14232 | -0.34558 | -0.12766 | -0.12009 | -0.00034 | -0.0149 | 0.014143 | 0.22621 | 0.08464 | -0.17024 | 0.007381 | -0.00843 | 0.12076 | 0.039854 | -0.43733 |
| OMM25 | -0.24155 | 0.036722 | -0.30878 | -0.05755 | -0.0854 | -0.10023 | 0.073015 | -0.21372 | -0.02923 | 0.06991 | -0.16591 | 0.12039 | -0.26043 | -0.06787 | -0.04047 | -0.00266 | -0.10838 | -0.0887 | 0.06069 | 0.14472 | 0.09717 | -0.21385 | 0.090234 | -0.04508 | 0.11617 | -0.04937 | -0.41325 |
| HOKUG | -0.17488 | 0.076382 | -0.62679 | -0.07653 | -0.08455 | -0.0428 | -0.08951 | -0.09288 | -0.08315 | 0.13696 | -0.13909 | 0.17079 | -0.25019 | -0.125 | -0.07793 | -0.06913 | -0.00686 | -0.01602 | -0.09147 | 0.17519 | 0.06885 | -0.30987 | 0.060846 | -0.15716 | 0.13225 | 0.002808 | -0.27325 |
| SKGIIIA | -0.08931 | 0.055693 | -0.0755 | -0.0738 | -0.22242 | -0.11993 | -0.02709 | -0.17145 | -0.08053 | -0.05913 | -0.17726 | 0.12926 | -0.12742 | -0.21369 | -0.05557 | -0.04601 | -0.07098 | -0.03518 | -0.00455 | 0.08398 | 0.04464 | -0.20129 | 0.093846 | -0.13508 | 0.07036 | 0.090848 | -0.28382 |
| T3M3 | -0.12355 | -0.07977 | 0.00171 | -0.00829 | -0.00482 | 0.01762 | -0.07728 | -0.28692 | -0.11564 | -0.09724 | -0.05684 | 0.3285 | -0.37451 | -0.17838 | -0.10856 | -0.15027 | 0.00142 | 0.13041 | -0.03536 | 0.11156 | 0.19365 | -0.3502 | 0.165076 | -0.14269 | 0.31157 | -0.01118 | -0.76007 |
| TGBC18TKB | -0.15795 | 0.09573 | -0.38865 | -0.08074 | -0.08895 | -0.01972 | 0.063125 | -0.20498 | -0.09402 | -0.04492 | -0.14147 | 0.20305 | -0.03628 | -0.1603 | -0.18092 | 0.022636 | -0.00419 | -0.01754 | 0.04666 | 0.06446 | 0.14451 | -0.16796 | 0.101267 | 0.00601 | 0.06227 | -0.07612 | -0.27897 |
| ECC4 | 0.06743 | -0.0617 | -0.46061 | -0.16806 | -0.15531 | -0.06422 | 0.112767 | -0.29723 | -0.1535 | -0.01716 | -0.14833 | 0.29676 | -0.24346 | -0.28789 | -0.15332 | -0.16102 | -0.15534 | 0.23766 | -0.195 | -0.00565 | 0.04032 | -0.20521 | -0.02294 | -0.00561 | 0.40792 | 0.28788 | -0.35154 |
| TT1TKB | -0.0788 | 0.096315 | -0.05012 | -0.27737 | 0.0157 | 0.08037 | -0.04106 | -0.22048 | -0.04405 | -0.10199 | -0.07733 | 0.05078 | -0.32201 | -0.02807 | -0.1146 | 0.052737 | 0.00032 | -0.12633 | -0.1828 | 0.04211 | 0.24111 | 0.008705 | 0.003317 | -0.1888 | 0.10365 | 0.005415 | -0.38393 |
| HHUA | -0.2673 | 0.097425 | -0.1982 | -0.08106 | -0.13169 | 0.06633 | 0.051509 | -0.34527 | -0.06634 | -0.06775 | -0.05687 | 0.16977 | -0.21255 | -0.21259 | -0.15207 | -0.14193 | 0.01784 | -0.04319 | 0.094446 | 0.13419 | 0.15389 | -0.26535 | 0.10862 | -0.0395 | 0.07126 | -0.03002 | -0.23729 |
| HOUAI | -0.18494 | 0.051144 | -0.17126 | -0.12819 | 0.02135 | 0.03952 | -0.00899 | -0.37238 | -0.26115 | -0.07461 | -0.09514 | 0.33699 | -0.28024 | -0.14863 | -0.10984 | 0.01154 | 0.18504 | 0.09497 | 0.084965 | 0.21555 | 0.07597 | -0.14603 | 0.204765 | -0.03223 | 0.03158 | -0.09212 | -0.12318 |
| SAS | -0.0705 | 0.043851 | -0.15756 | -0.05252 | -0.17325 | -0.10021 | 0.027792 | -0.31841 | -0.11351 | -0.05832 | -0.10302 | 0.05677 | -0.21787 | -0.23063 | -0.16297 | -0.16409 | -0.03475 | 0.00816 | -0.11194 | 0.07702 | 0.19515 | -0.31864 | 0.04965 | -0.02173 | 0.13571 | 0.141238 | -0.42062 |
| LCAM1 | -0.13994 | -0.04374 | -0.65011 | -0.04946 | -0.29182 | -0.23917 | -0.21065 | -0.41511 | -0.0069 | -0.29573 | -0.23198 | 0.23488 | -0.41666 | -0.39601 | -0.24724 | -0.17373 | 0.09513 | 0.13771 | -0.02706 | 0.12057 | 0.10895 | -0.1174 | -0.02177 | 0.15982 | 0.18259 | 0.14907 | -0.48593 |
| PK8 | -0.08973 | -0.28688 | -0.64043 | -0.12082 | -0.18817 | 0.13332 | -0.20357 | -0.25914 | -0.37033 | 0.06045 | 0.022787 | 0.16736 | -0.44568 | -0.30562 | -0.08811 | -0.08908 | 0.13209 | 0.08753 | -0.18287 | 0.23382 | 0.12692 | -0.16571 | 0.103518 | 0.00285 | 0.1587 | 0.086548 | -0.58307 |
| HOTHC | -0.09456 | -0.07175 | -0.19427 | -0.11593 | -0.10482 | -0.10802 | -0.09529 | -0.23002 | 0.018524 | -0.06598 | -0.20108 | 0.20878 | -0.22035 | -0.25008 | -0.19024 | -0.03386 | -0.11856 | -0.03421 | -0.03407 | 0.06675 | 0.08973 | -0.16694 | 0.057394 | -0.06924 | 0.06626 | -0.00099 | -0.32677 |
| T3M5 | 0.00432 | 0.025317 | -0.41714 | -0.28531 | -0.24426 | 0.01671 | -0.04662 | -0.27217 | -0.15578 | -0.01801 | -0.05418 | 0.20113 | -0.08766 | -0.14907 | -0.0923 | -0.07908 | -0.07028 | -0.19928 | -0.05807 | 0.21214 | 0.237 | -0.44379 | -0.02778 | -0.08897 | 0.14315 | 0.152385 | -0.25098 |
| CA922 | -0.11205 | 0.024782 | -0.13937 | -0.02305 | -0.2367 | -0.03249 | -0.07444 | -0.20032 | -0.02338 | -0.00307 | -0.11546 | 0.10337 | -0.08009 | -0.08088 | -0.07455 | -0.05 | -0.02548 | -0.02019 | -0.02892 | 0.17821 | 0.18938 | -0.22092 | 0.034776 | -0.1367 | 0.20387 | 0.075397 | -0.27353 |
| HSQ89 | -0.1013 | 0.019314 | -0.21233 | -0.0764 | -0.30213 | -0.089 | -0.0882 | -0.1607 | 0.010607 | -0.22049 | -0.15829 | 0.21488 | -0.3014 | -0.18727 | -0.1044 | -0.03777 | -0.01093 | -0.08168 | -0.06017 | 0.09447 | 0.02377 | -0.20919 | 0.089137 | -0.14957 | 0.14143 | -0.09768 | -0.31443 |
| HO1U1 | -0.12085 | 0.001135 | 0.03515 | -0.09279 | -0.00507 | -0.13698 | 0.045461 | -0.9389 | 0.016173 | -0.18304 | -0.1812 | 0.25709 | -0.14578 | -0.06666 | 0.039938 | -0.15287 | -0.01219 | 0.17759 | -0.05857 | 0.17694 | 0.01476 | -0.30085 | 0.08694 | 0.08489 | 0.19063 | 0.024887 | -0.3438 |
| HTMMT | -0.06626 | -0.01476 | -0.155 | -0.12925 | -0.11437 | -0.15678 | -0.08008 | -0.15533 | -0.00429 | -0.04614 | -0.13341 | 0.23626 | -0.26125 | -0.26057 | -0.06241 | -0.07815 | -0.06282 | -0.00274 | -0.07126 | 0.08894 | 0.13155 | -0.15754 | 0.071447 | -0.13072 | 0.15 | 0.020614 | -0.43943 |
| RMSYM | -0.10411 | -0.03947 | -0.6939 | -0.16708 | -0.16795 | -0.12179 | -0.03894 | -0.20322 | 0.064841 | 0.08447 | -0.33524 | 0.1286 | -0.16676 | -0.2333 | -0.02455 | -0.08356 | -0.10927 | 0.0029 | -0.18151 | 0.2174 | 0.04152 | -0.1673 | 0.09715 | -0.08999 | 0.08826 | 0.063584 | -0.38445 |
| LU134A | -0.15156 | -0.00782 | 0.06351 | -0.13969 | -0.18043 | -0.07734 | -0.24934 | -0.11549 | 0.05935 | -0.07058 | -0.05876 | 0.08953 | -0.23689 | -0.22173 | 0.082299 | -0.02646 | -0.04456 | 0.18905 | -0.02426 | 0.02542 | 0.16303 | -0.246 | -0.05535 | -0.06188 | 0.14225 | 0.174111 | -0.666 |
| P30OHK | -0.07469 | -0.05988 | -0.09181 | -0.12044 | -0.08466 | -0.01724 | 0.021056 | -0.23728 | 0.150189 | 0.11287 | -0.11152 | 0.18325 | -0.27175 | -0.21388 | -0.14285 | -0.1741 | 0.03246 | 0.10046 | -0.07101 | 0.27845 | 0.38004 | -0.21319 | 0.355246 | -0.06046 | 0.23728 | 0.010245 | -0.32319 |
| P2URK562 | -0.15128 | 0.029141 | -0.26556 | -0.13374 | 0.01856 | 0.00809 | -0.0535 | -0.26566 | -0.2422 | -0.03371 | -0.11917 | 0.05284 | -0.55861 | 0.09483 | -0.16498 | -0.49917 | 0.00621 | 0.11565 | -0.03346 | 0.06068 | 0.27808 | -0.11282 | 0.14369 | 0.03746 | 0.24151 | 0.217317 | -0.52193 |
| SLVL | 0.03292 | 0.093715 | 0.10525 | -0.0187 | -0.02983 | 0.02245 | -0.01681 | -0.14757 | 0.019863 | -0.01039 | -0.03427 | 0.01443 | -0.04882 | -0.43287 | -0.03461 | -0.19985 | 0.11241 | 0.16677 | -0.17822 | 0.0849 | 0.05173 | -0.11558 | 0.406522 | 0.02764 | 0.34308 | 0.084443 | -0.54403 |
| HSSCH2 | 0.06335 | -0.03911 | -0.6524 | -0.0991 | -0.13241 | -0.12628 | -0.07343 | -0.12082 | 0.093082 | -0.0482 | -0.26021 | 0.16305 | -0.02592 | -0.3332 | -0.09738 | -0.10947 | -0.21186 | -0.06014 | 0.076953 | 0.10894 | 0.14623 | -0.09014 | 0.135197 | 0.02117 | 0.13636 | -0.17908 | -0.20762 |
| NOS1 | -0.2264 | 0.086567 | -0.43173 | -0.1777 | -0.24523 | 0.20843 | 0.070053 | -0.15223 | 0.064593 | -0.04638 | -0.30294 | 0.06888 | -0.45181 | -0.09278 | 0.050723 | -0.12357 | 0.00529 | -0.30044 | -0.03445 | -0.06351 | 0.22724 | -0.2993 | 0.117249 | -0.17199 | 0.29855 | 0.136737 | -0.35896 |
| HSOS1 | -0.11759 | 0.046391 | -0.379 | -0.39581 | -0.16212 | -0.10927 | 0.128446 | -0.24467 | 0.076254 | -0.08794 | -0.16269 | 0.12131 | -0.8751 | -0.5453 | -0.03201 | -0.21039 | -0.15181 | 0.08445 | -0.15301 | 0.03803 | 0.14699 | -0.07901 | 0.223276 | 0.00799 | 0.20676 | 0.141932 | -0.23619 |
| HKBMM | -0.15329 | 0.00753 | -0.10425 | -0.12047 | -0.11637 | -0.21581 | -0.03269 | -0.28624 | -0.05559 | -0.02047 | -0.12507 | 0.0625 | -0.59322 | -0.2568 | -0.12908 | -0.00859 | -0.06086 | -0.0978 | 0.032665 | 0.14543 | 0.07832 | -0.17366 | 0.082756 | -0.09745 | 0.19715 | -0.03288 | -0.42845 |
| LU165 | -0.13708 | 0.1164 | -0.31106 | -0.03173 | -0.20051 | -0.35595 | -0.16286 | -0.20171 | -0.0201 | -0.13902 | -0.28077 | 0.17966 | -0.3105 | -0.16282 | -0.01624 | -0.0765 | -0.03621 | -0.11135 | -0.29909 | 0.28683 | 0.3012 | -0.15743 | 0.034024 | 0.12034 | 0.34053 | -0.06085 | -0.29273 |
| TN2 | -0.14507 | -0.0775 | -0.89361 | -0.2099 | -0.14294 | -0.14626 | -0.05781 | -0.24652 | -0.02869 | -0.00078 | -0.1302 | 0.15372 | -0.10654 | -0.19959 | -0.06354 | -0.03838 | -0.09797 | -0.03068 | 0.00392 | 0.1463 | 0.13846 | -0.2147 | 0.042369 | -0.07497 | 0.16406 | 0.024079 | -0.24173 |
| NB69 | -0.06516 | -0.12272 | -0.67299 | -0.04075 | -0.04948 | -0.12654 | -0.14197 | -0.50943 | -0.26752 | -0.06024 | 0.082197 | 0.30844 | -0.3076 | -0.32085 | -0.12128 | -0.03252 | -0.11004 | -0.01232 | -0.07861 | 0.17064 | 0.05939 | -0.18729 | 0.073592 | -0.25665 | 0.32482 | -0.05471 | -0.3693 |
| MMAC | -0.05468 | 0.103847 | -0.19056 | -0.10527 | -0.03643 | -0.10468 | 0.064559 | -0.24333 | 0.079003 | -0.03494 | -0.16402 | 0.13973 | -0.18671 | -0.02844 | 0.000739 | -0.06834 | -0.00345 | 0.01782 | 0.152273 | 0.14593 | 0.11384 | -0.24946 | 0.078854 | -0.058 | 0.0478 | 0.159695 | -0.30706 |
| ES4 | -0.21599 | -0.05322 | -0.37977 | -0.12917 | 0.03643 | -0.27444 | -0.07073 | -0.31752 | -0.11913 | 0.25759 | -0.01731 | 0.2403 | -0.20206 | -0.02241 | -0.13003 | 0.003456 | -0.07837 | 0.05364 | 0.097499 | 0.0581 | 0.09656 | -0.15478 | 0.092543 | -0.0735 | 0.17648 | -0.0951 | -0.57904 |
| ES5 | 0.09257 | -0.20353 | -1.90787 | -0.19734 | -0.00112 | -0.40467 | 0.13327 | -0.16725 | -0.34137 | -0.06905 | -0.08242 | 0.03311 | -0.38034 | -0.3479 | -0.01809 | -0.00021 | 0.04282 | 0.06319 | 0.017476 | 0.23735 | 0.03826 | -0.44247 | 0.054809 | 0.06037 | 0.22476 | -0.12307 | -0.34042 |
| ES8 | 0.00476 | -0.00702 | -0.67178 | -0.18506 | -0.08011 | -0.22676 | -0.15412 | -0.41009 | 0.046513 | -0.21712 | -0.10137 | 0.11602 | -0.34784 | -0.1219 | 0.045215 | -0.11566 | -0.03314 | 0.13899 | 0.118234 | 0.12764 | 0.06348 | 0.032493 | 0.242329 | -0.27709 | 0.15264 | 0.075434 | -0.53949 |
| EW1 | 0.04088 | 0.139675 | -1.5034 | -0.12069 | -0.0404 | -0.02169 | 0.051062 | -0.32084 | -0.11521 | 0.13787 | 0.213145 | 0.2219 | -0.22523 | -0.08591 | -0.09864 | -0.0413 | 0.11307 | 0.16867 | 0.069936 | 0.2156 | 0.03546 | -0.18647 | -0.00706 | -0.03965 | 0.0375 | 0.10308 | -0.38394 |
| EW16 | -0.10286 | -0.11686 | -0.90533 | -0.18575 | -0.12644 | -0.11241 | -0.27022 | -0.14949 | -0.08202 | -0.11503 | -0.23834 | 0.23839 | -0.51311 | -0.23279 | -0.06508 | -0.08595 | 0.0231 | 0.11341 | -0.11104 | 0.21555 | 0.0596 | -0.3266 | -0.07646 | 0.08294 | 0.18588 | -0.00504 | -0.3271 |
| EW22 | 0.11932 | -0.01614 | -1.27155 | -0.1568 | -0.19077 | -0.05557 | 0.069716 | -0.05539 | -0.089 | -0.00394 | -0.08962 | 0.03431 | -0.02289 | -0.16368 | -0.03498 | -0.04759 | -0.04462 | -0.01816 | -0.10208 | 0.08973 | 0.08236 | -0.15794 | -0.07116 | 0.0642 | 0.16551 | 0.036317 | -0.31658 |
| EW7 | -0.14025 | -0.14186 | -0.25684 | -0.10075 | 0.02393 | -0.05647 | 0.047972 | -0.17734 | -0.15156 | 0.16414 | 0.157459 | 0.18563 | 0.00041 | -0.01583 | 0.060241 | -0.22531 | 0.1031 | -0.00592 | 0.127144 | 0.30199 | 0.11989 | -0.27586 | 0.110826 | 0.06771 | 0.25035 | -0.16296 | -0.37662 |
| HEY | -0.16875 | 0.016031 | 0.05955 | -0.07469 | -0.28042 | -0.30727 | 0.062153 | -0.14887 | -0.19682 | -0.05348 | 0.042338 | 0.14907 | -0.17631 | -0.10252 | -0.21156 | -0.1481 | -0.06079 | -0.04115 | 0.067155 | 0.02019 | 0.16925 | -0.32466 | 0.127544 | -0.05276 | 0.18626 | -0.04729 | -0.68764 |
| HSC39 | -0.08396 | 0.074344 | -0.82435 | -0.06441 | -0.04458 | -0.17546 | 0.178375 | -0.3015 | -0.07608 | -0.12463 | -0.05209 | 0.0677 | 0.02753 | -0.14767 | -0.04667 | -0.08708 | 0.00396 | 0.18753 | 0.124601 | -0.02117 | 0.15615 | -0.17233 | 0.056743 | 0.07046 | 0.12138 | -0.35324 | -0.34937 |
| ISTMEL1 | 0.0012 | -0.02639 | -0.18413 | -0.09361 | -0.08669 | -0.19215 | -0.08616 | 0.12277 | 0.174473 | 0.12614 | 0.037105 | 0.11069 | -0.30235 | -0.07678 | -0.09095 | -0.18531 | -0.06679 | 0.14292 | -0.24213 | 0.28549 | 0.01363 | -0.00764 | 0.136626 | -0.05127 | -0.01751 | -0.10405 | -0.05377 |
| LB1047RCC | 0.00537 | 0.110733 | 0.01747 | -0.07967 | 0.09862 | -0.22857 | -0.02301 | -0.2108 | 0.085905 | -0.09816 | 0.032747 | 0.03534 | -0.14583 | -0.14596 | -0.09741 | -0.16136 | -0.03402 | 0.05322 | 0.074508 | 0.01384 | 0.12842 | -0.16953 | -0.13143 | 0.00658 | 0.1812 | -0.07527 | -0.3508 |
| LC1SQ | -0.0739 | -0.25139 | -0.49492 | -0.05658 | -0.4464 | -0.04095 | 0.005677 | -0.17969 | -0.26768 | -0.03388 | 0.090237 | 0.0887 | -0.14591 | -0.12431 | -0.3822 | -0.23931 | 0.07735 | 0.08811 | -0.09671 | 0.0546 | -0.05162 | -0.13527 | 0.131471 | 0.17039 | 0.25992 | -0.10946 | -0.0097 |
| MCIXC | -0.00456 | -0.11088 | -0.59274 | -0.07697 | -0.21139 | 0.07604 | 0.057193 | -0.298 | -0.41804 | 0.06175 | -0.21353 | 0.15834 | -0.17977 | -0.34248 | -0.11755 | -0.1554 | -0.2053 | -0.21366 | -0.11868 | 0.14836 | 0.16325 | -0.18102 | 0.001199 | -0.29919 | 0.32673 | 0.094805 | -0.37607 |
| MKN28 | -0.14281 | 0.194098 | -0.66259 | 0.00655 | -0.25967 | -0.29802 | 0.014286 | -0.07313 | 0.001676 | -0.07648 | -0.04783 | -0.11765 | -0.20078 | -0.00173 | 0.059715 | 0.052818 | 0.08703 | 0.13953 | -0.04468 | 0.15284 | 0.03937 | -0.32838 | 0.068683 | -0.09017 | 0.24411 | 0.114175 | -0.2987 |
| OCUBM | -0.3518 | 0.010775 | 0.17264 | 0.03433 | -0.0135 | -0.34782 | -0.35854 | 0.12492 | 0.081052 | 0.08176 | 0.074467 | 0.09917 | -0.41238 | -0.10629 | -0.17858 | -0.40264 | -0.25185 | -0.02805 | -0.19767 | -0.16732 | 0.23783 | -0.11882 | 0.181852 | 0.09206 | 0.06604 | 0.09809 | -0.28278 |
| OVCA420 | -0.18064 | -0.13604 | -0.48867 | 0.11414 | -0.30836 | -0.19789 | -0.00434 | -0.29746 | -0.21973 | 0.01559 | 0.055224 | 0.17981 | -0.02572 | -0.25286 | -0.14391 | -0.25799 | 0.07432 | 0.06948 | -0.15729 | 0.02948 | 0.03675 | -0.43447 | 0.139594 | -0.13179 | 0.38307 | 0.161228 | -0.4063 |
| OVMIU | -0.23554 | -0.18465 | -0.02601 | -0.06888 | -0.3656 | -0.30107 | -0.02186 | -0.54909 | -0.12922 | -0.32233 | -0.00203 | -0.09992 | -0.3732 | -0.18964 | 0.094421 | -0.27749 | -0.10345 | 0.19683 | -0.16776 | 0.1255 | 0.16173 | -0.37387 | 0.016455 | -0.13458 | 0.32448 | 0.016886 | -0.23097 |
| PL4 | -0.11192 | 0.019542 | -0.97118 | 0.14242 | 0.06645 | -0.08672 | 0.293422 | -0.15987 | -0.01402 | -0.03734 | -0.24582 | 0.22203 | -0.32892 | 0.14416 | -0.12987 | -0.06543 | 0.03603 | 0.13817 | 0.136772 | 0.29487 | 0.09332 | -0.09928 | 0.303245 | -0.20703 | 0.07553 | -0.13235 | 0.0408 |
| RCCFG2 | -0.10071 | -0.04357 | -0.12031 | -0.00425 | -0.36823 | -0.03136 | -0.14484 | -0.25464 | -0.24283 | -0.12454 | -0.13123 | 0.04503 | -0.15534 | -0.27078 | -0.03339 | -0.08013 | 0.09559 | 0.05352 | -0.03962 | 0.13365 | 0.02572 | -0.23533 | 0.059253 | -0.01118 | 0.16745 | 0.000808 | -0.29683 |
| SCH | -0.01479 | 0.351251 | -0.0985 | 0.25992 | 0.11244 | -0.20785 | 0.110343 | 0.05699 | 0.025934 | 0.03445 | -0.04908 | -0.09519 | 0.07132 | 0.18111 | 0.055716 | 0.108705 | 0.20888 | 0.27525 | 0.111995 | 0.19705 | 0.16739 | 0.120284 | -0.15329 | 0.14098 | 0.0059 | 0.033961 | -0.27372 |
| TMK1 | 0.10633 | 0.318145 | -0.4068 | -0.01285 | 0.10765 | 0.17355 | 0.143171 | 0.02972 | 0.154407 | 0.13389 | -0.18697 | 0.09692 | 0.19823 | -0.14309 | -0.05985 | 0.049551 | 0.07083 | 0.29904 | -0.03613 | 0.20354 | 0.13945 | -0.00863 | -0.10794 | -0.04208 | 0.03087 | 0.024568 | 0.03956 |
| ARH77 | -0.04626 | 0.265001 | 0.05266 | 0.12379 | 0.09596 | -0.018 | 0.333357 | -0.01611 | -0.11196 | 0.04593 | -0.0821 | 0.05902 | -0.0025 | 0.00083 | 0.012905 | 0.164312 | 0.00771 | 0.25683 | -0.02348 | 0.21175 | 0.16174 | 0.133929 | -0.05321 | 0.03496 | 0.07093 | 0.042921 | -0.17392 |
| BB30HNC | 0.13674 | 0.050095 | -0.233 | -0.04504 | -0.38788 | 0.02516 | -0.19285 | -0.16324 | -0.17494 | -0.28765 | -0.02481 | 0.16029 | -0.20948 | -0.11623 | -0.15626 | -0.19658 | 0.08732 | 0.22894 | 0.016248 | 0.13555 | 0.0882 | -0.25848 | -0.01698 | -0.19739 | 0.05045 | -0.29045 | -0.78615 |
| BE2M17 | -0.10051 | -0.24178 | -1.94394 | -0.26962 | -0.15215 | -0.22272 | 0.00135 | -0.27629 | -0.15419 | -0.02492 | -0.00417 | 0.28627 | -0.29499 | -0.24442 | -0.05897 | -0.20799 | -0.04719 | 0.05861 | -0.0925 | 0.0912 | 0.0936 | -0.36632 | 0.011085 | 0.04173 | 0.18705 | 0.024485 | -0.61142 |
| D423MG | -0.19071 | -0.00024 | 0.05056 | -0.31392 | -0.10298 | 0.11599 | -0.41277 | -0.1734 | -0.0109 | -0.06741 | -0.24588 | 0.06635 | -0.05798 | -0.30439 | 0.091948 | 0.212601 | -0.21923 | 0.11306 | -0.00847 | 0.25972 | 0.05364 | -0.27222 | 0.152564 | -0.10856 | 0.21218 | 0.003274 | -0.26087 |
| D502MG | 0.00673 | -0.05867 | -0.94965 | 0.05332 | -0.30399 | 0.06325 | -0.08957 | -0.76207 | -0.09276 | 0.00517 | 0.044074 | 0.15241 | -0.40975 | -0.09404 | -0.04927 | 0.246671 | -0.09908 | 0.34687 | 0.123353 | -0.07863 | 0.08276 | -0.19959 | 0.438317 | -0.23598 | 0.35784 | -0.12934 | -0.60688 |
| D542MG | -0.11338 | 0.011524 | -0.11581 | -0.20069 | -0.03804 | -0.15707 | -0.10832 | -0.36906 | -0.12088 | -0.27972 | -0.05321 | 0.0632 | -0.55686 | -0.4489 | -0.09234 | -0.06459 | 0.02498 | 0.07392 | -0.05394 | 0.26022 | -0.03339 | -0.52753 | 0.075778 | -0.00135 | 0.23565 | 0.117042 | -0.44945 |
| DIFI | -0.09544 | 0.067668 | -0.64031 | 0.13818 | -0.29194 | -0.03288 | -0.0939 | -0.28086 | -0.01363 | -0.17199 | 0.053512 | -0.02622 | -0.0945 | -0.12062 | -0.15193 | 0.018419 | -0.07361 | 0.01109 | 0.040318 | 0.39942 | 0.17788 | -0.12747 | 0.203464 | 0.0183 | 0.13415 | -0.04209 | -0.32514 |
| DOK | -0.07095 | 0.290539 | -0.54701 | -0.12578 | -0.02549 | -0.28628 | 0.240287 | -0.04563 | -0.01108 | -0.14324 | -0.11049 | 0.07824 | 0.06875 | 0.10365 | 0.037587 | 0.057711 | 0.12569 | 0.24588 | 0.009534 | 0.33815 | 0.37506 | -0.05663 | 0.006669 | -0.04845 | 0.13258 | 0.099566 | -0.43201 |
| H3118 | 0.0353 | -0.01322 | -1.22443 | -0.08604 | 0.03475 | -0.11288 | 0.109408 | -0.18117 | -0.44968 | 0.17038 | -0.2359 | 0.10162 | -0.03575 | -0.27716 | -0.16285 | -0.02817 | 0.01405 | 0.0969 | -0.11862 | 0.1558 | 0.11748 | -0.28847 | 0.007553 | 0.021 | 0.11629 | 0.425556 | -0.16333 |
| HCE4 | -0.11723 | 0.194192 | -0.05831 | 0.02375 | 0.03961 | -0.33435 | 0.17181 | 0.09153 | -0.05778 | -0.19159 | -0.28503 | 0.30427 | -0.02676 | -0.19167 | -0.04483 | -0.1675 | -0.09652 | 0.32624 | -0.15263 | 0.24353 | 0.26591 | -0.08732 | -0.01756 | 0.07906 | 0.04394 | 0.294934 | -0.37887 |
| IM9 | -0.04675 | 0.257362 | -0.09484 | 0.2099 | -0.07719 | -0.00303 | 0.281145 | -0.34685 | 0.000452 | -0.17596 | -0.11703 | 0.24015 | -0.02279 | -0.22957 | 0.162115 | -0.35029 | 0.18559 | 0.0749 | -0.00594 | 0.22932 | -0.09782 | -0.13973 | -0.07125 | -0.09953 | 0.30624 | 0.05235 | -0.21784 |
| JHU011 | -0.26396 | 0.039798 | -0.63269 | -0.05401 | -0.20086 | -0.01577 | 0.093977 | -0.13909 | -0.02267 | -0.15961 | -0.05376 | 0.10591 | -0.30286 | -0.12821 | -0.05274 | -0.15114 | 0.04758 | 0.20909 | 0.058997 | 0.17806 | 0.05597 | -0.18314 | 0.07952 | -0.05326 | 0.14602 | 0.05263 | -0.37632 |
| JHU022 | -0.08509 | 0.139452 | -0.1886 | -0.19696 | -0.25057 | -0.13729 | 0.049797 | -0.26981 | -0.34661 | 0.19514 | -0.18303 | 0.14646 | -0.14353 | -0.08592 | 0.015704 | -0.02521 | -0.04161 | 0.20804 | 0.049381 | -0.00012 | 0.04262 | -0.22293 | 0.085375 | -0.19347 | 0.11971 | 0.189513 | -0.08205 |
| JHU029 | -0.17058 | 0.023396 | -0.21704 | 0.02009 | -0.18904 | -0.22758 | -0.1026 | -0.27137 | -0.07135 | -0.15373 | 0.015743 | 0.0648 | -0.10992 | -0.12915 | -0.11946 | -0.04482 | -0.0032 | 0.11264 | -0.04891 | -0.0653 | 0.12294 | -0.24861 | 0.049644 | 0.03708 | 0.2283 | 0.072249 | -0.32354 |

| Cell line | FLT3 | FLT4 | IGF1R | INSR | INSRR | KDR | KIT | LMR1 | LMR2 | LTK | MET | MUSK | PDGFRA | PDGFRB | PTK7 | RET | RON | ROR1 | ROR2 | ROS1 | STYK1 | TIE1 | TIE2 | TRKA | TRKB | TRKC | TYRO3 |
|---|---|---|---|---|---|---|---|---|---|---|---|---|---|---|---|---|---|---|---|---|---|---|---|---|---|---|---|
| KINGS1 | -0.11744 | -0.07208 | -0.19872 | -0.13628 | -0.19218 | 0.0663 | -0.22654 | -0.41364 | -0.06879 | -0.24714 | -1.57687 | 0.01342 | -0.48445 | -0.119 | -0.13437 | 0.009032 | -0.12792 | 0.25673 | 0.137159 | 0.20141 | 0.08109 | -0.14612 | 0.072918 | 0.18283 | -0.04115 | -0.11439 | -0.41185 |
| KPNYS | -0.18318 | -0.1177 | -0.24105 | -0.05837 | -0.05797 | -0.14343 | -0.25703 | -0.43124 | -0.09267 | -0.12305 | -0.04759 | 0.17526 | -0.16597 | -0.34041 | -0.2167 | -0.02479 | -0.06677 | 0.07105 | -0.08117 | 0.14747 | 0.05899 | -0.24605 | 0.060002 | -0.02455 | 0.2331 | 0.01331 | -0.24398 |
| KYSE220 | -0.13724 | 0.137353 | -0.38281 | -0.18008 | -0.04363 | -0.05117 | 0.074372 | -0.1234 | -0.07261 | -0.1309 | -0.11278 | 0.05891 | -0.04225 | -0.05789 | -0.06769 | -0.17467 | -0.04974 | 0.0962 | -0.06353 | 0.02885 | 0.09409 | -0.22325 | -0.08859 | -0.13388 | -0.04923 | 0.053925 | -0.21312 |
| LB771HNC | 0.02192 | 0.170999 | -1.81856 | -0.0139 | -0.1405 | -0.2692 | 0.053807 | -0.12635 | -0.16516 | -0.43206 | -0.08266 | 0.07476 | -0.37655 | 0.00083 | -0.34488 | -0.05646 | -0.04122 | 0.18945 | -0.12822 | 0.14758 | 0.15979 | -0.07456 | -0.11229 | -0.00435 | -0.08148 | 0.119392 | -0.39845 |
| LNZTA3WT4 | -0.06446 | 0.022149 | 0.08524 | -0.06812 | -0.03624 | -0.16997 | -0.01833 | -0.18657 | -0.0396 | -0.2781 | 0.012378 | 0.13216 | -0.4417 | -0.25478 | -0.05985 | -0.06235 | -0.17864 | 0.15004 | -0.13681 | 0.07095 | -0.00989 | -0.16968 | -0.02147 | 0.03907 | 0.31511 | 0.17732 | -0.52244 |
| NB10 | -0.17224 | -0.13695 | -1.2167 | 0.01179 | -0.23138 | -0.15263 | 0.016103 | -0.69638 | -0.16468 | 0.00838 | 0.012471 | 0.2692 | -0.23183 | -0.11577 | -0.19589 | -0.21897 | 0.01635 | 0.08047 | 0.070345 | 0.03792 | 0.19681 | -0.1159 | 0.034127 | -0.04541 | 0.49657 | -0.22018 | -0.97196 |
| NB13 | -0.04034 | -0.31359 | -0.94852 | -0.15991 | -0.4797 | -0.16419 | 0.008031 | -0.35886 | -0.27274 | -0.05103 | -0.16481 | 0.26753 | -0.04854 | -0.37461 | -0.18149 | -0.22474 | -0.00618 | 0.05383 | -0.13122 | 0.15496 | 0.09851 | -0.41111 | 0.040046 | 0.06786 | 0.27463 | -0.27501 | -0.43752 |
| NB17 | -0.08094 | -0.27364 | -0.80012 | -0.00764 | -0.05065 | -0.32569 | -0.12707 | -0.70654 | -0.21981 | 0.01526 | -0.04851 | 0.10299 | -0.21161 | -0.23835 | -0.17367 | -0.06265 | -0.13988 | 0.27134 | -0.10447 | 0.09971 | 0.08565 | -0.25548 | 0.071939 | 0.03886 | 0.34028 | -0.13556 | -0.38643 |
| NB5 | -0.20442 | -0.22479 | -0.31177 | -0.13437 | -0.16547 | -0.16348 | -0.17621 | -0.40693 | -0.05323 | 0.01271 | -0.03486 | 0.09331 | -0.23462 | -0.2126 | -0.08976 | 0.034113 | -0.07591 | 0.07504 | -0.08013 | 0.02976 | 0.15017 | -0.33524 | 0.15174 | 0.08108 | 0.30317 | -0.11055 | -0.45233 |
| NB6 | -0.27911 | -0.04204 | -0.14096 | -0.09032 | -0.19071 | -0.2909 | -0.01813 | -0.33753 | -0.14634 | 0.00721 | -0.03156 | -0.00465 | -0.27177 | -0.34289 | -0.24287 | -0.27691 | -0.06813 | 0.25236 | 0.066883 | -0.02049 | 0.12436 | -0.25193 | 0.202332 | -0.06893 | 0.22375 | 0.031386 | -0.72864 |
| NB7 | -0.26508 | 0.157014 | -0.27529 | -0.21178 | -0.16503 | 0.08355 | -0.04631 | -0.33306 | -0.12928 | -0.04163 | -0.03788 | 0.16184 | -0.11467 | -0.18795 | -0.10345 | -0.10882 | 0.00142 | -0.02946 | 0.066767 | 0.15383 | 0.10248 | -0.31818 | 0.209906 | -0.12321 | 0.16746 | -0.10122 | -0.25204 |
| NTERA2CLD1 | -0.1527 | -0.05212 | -0.57112 | -0.08639 | -0.0015 | 0.10044 | -0.12385 | -0.31379 | 0.057181 | 0.15267 | -0.12224 | 0.0128 | 0.02792 | -0.17185 | -0.10271 | -0.08468 | -0.04209 | 0.22019 | 0.09572 | 0.1839 | -0.18282 | 0.063734 | -0.01527 | 0.11502 | 0.24487 | -0.08485 | -0.35568 |
| PCI15A | -0.20191 | 0.030766 | -0.23794 | 0.027 | -0.33124 | -0.17795 | 0.270053 | -0.15747 | -0.04238 | -0.16169 | 0.018585 | 0.25528 | -0.24794 | -0.24501 | -0.23434 | -0.33985 | -0.08062 | 0.15512 | 0.025012 | -0.07307 | 0.11806 | -0.16597 | 0.173242 | -0.09964 | 0.14042 | 0.092256 | -0.48311 |
| PCI30 | -0.04757 | 0.036477 | -0.0983 | -0.04732 | -0.09805 | 0.06608 | -0.02864 | -0.28307 | -0.16186 | 0.10256 | -0.16985 | 0.09405 | -0.10971 | -0.04968 | -0.12064 | -0.02296 | -0.15202 | 0.08487 | -0.07938 | 0.13905 | 0.02554 | -0.20649 | 0.017115 | 0.01141 | 0.11638 | -0.06261 | -0.35404 |
| PCI38 | -0.19237 | -0.11957 | -0.65154 | -0.27625 | -0.12169 | -0.14502 | -0.07244 | -0.07434 | -0.14692 | -0.01313 | -0.09716 | 0.05595 | -0.26513 | -0.16063 | 0.017492 | -0.1263 | -0.07381 | 0.06105 | -0.07855 | 0.10549 | 0.18554 | -0.21191 | -0.0021 | -0.1303 | 0.09572 | 0.037031 | -0.17817 |
| PCI4B | -0.11828 | 0.157851 | -0.03337 | 0.04478 | -0.02931 | -0.27909 | 0.043914 | -0.10907 | -0.25711 | -0.18855 | -0.0308 | -0.12121 | -0.15317 | -0.12541 | -0.08536 | -0.27654 | -0.06019 | 0.17074 | -0.08762 | 0.04138 | 0.04364 | -0.14446 | 0.270099 | 0.00785 | 0.23461 | 0.131323 | -0.95737 |
| PCI6A | -0.06628 | 0.143731 | -0.28621 | -0.11429 | -0.02031 | -0.03011 | 0.166534 | -0.23465 | -0.06621 | -0.00655 | -0.04719 | 0.20423 | -0.06983 | -0.04825 | -0.03401 | 0.099564 | 0.09767 | 0.2895 | 0.11414 | -0.01235 | 0.04337 | -0.13231 | -0.01585 | -0.16019 | 0.13273 | 0.239827 | -0.34314 |
| SKMG1 | -0.13793 | -0.0923 | 0.05201 | -0.16422 | -0.0941 | -0.16031 | -0.07623 | -0.22483 | -0.10382 | -0.00696 | 0.071253 | -0.0538 | -0.2878 | -0.21669 | -0.2022 | -0.16353 | -0.05941 | -0.02221 | -0.07123 | -0.14196 | 0.01707 | -0.22359 | 0.076596 | 0.0686 | 0.31989 | -0.06513 | -0.40985 |
| SKN3 | -0.12087 | 0.334574 | -0.41477 | 0.04546 | 0.2066 | -0.19495 | 0.114501 | 0.05984 | -0.12228 | -0.01258 | -0.25983 | -0.01486 | 0.00351 | 0.09231 | -0.06627 | 0.021783 | 0.05899 | -0.00254 | 0.192406 | 0.07736 | 0.19183 | 0.067504 | 0.068196 | 0.10044 | -0.00505 | 0.231748 | -0.12114 |
| 21NT | -0.16938 | 0.088896 | 0.08679 | -0.1356 | -0.32399 | -0.21435 | -0.09673 | -0.11399 | 0.089231 | -0.08236 | -0.0932 | 0.18201 | -0.39991 | -0.1351 | -0.1313 | -0.08505 | 0.10295 | 0.22728 | -0.11403 | 0.14818 | 0.11755 | -0.12557 | 0.120625 | -0.00088 | 0.06275 | 0.003577 | -0.45693 |
| CCLFUPGI0005T | -0.12057 | -0.03592 | -0.14055 | 0.00151 | -0.10911 | -0.17667 | -0.17977 | -0.16499 | -0.04975 | 0.04115 | -0.06824 | 0.19737 | -1.63677 | -0.16296 | 0.02191 | -0.01821 | -0.06318 | -0.06946 | -0.07781 | 0.19021 | 0.2002 | -0.09917 | 0.03874 | -0.06544 | 0.07548 | -0.08631 | -0.44702 |
| HT144SKINFV1 | -0.03198 | -0.0305 | -0.09326 | 0.01994 | -0.06415 | -0.00519 | 0.073947 | -0.1895 | -0.07937 | -0.00418 | -0.0019 | 0.13443 | -0.3528 | -0.19551 | 0.125185 | -0.0657 | -0.06628 | -0.05736 | 0.07947 | 0.31816 | -0.00477 | -0.1627 | 0.106885 | -0.05266 | 0.09136 | 0.011909 | -0.42324 |
| HT144SKINFV3 | -0.21313 | 0.052633 | -0.19083 | -0.11298 | -0.22611 | -0.1196 | -0.05059 | -0.36529 | -0.14643 | -0.05183 | -0.11993 | 0.21536 | -0.28873 | -0.35904 | -0.18311 | -0.14275 | 0.05926 | -0.11433 | 0.023518 | 0.28147 | 0.09472 | -0.31785 | 0.039991 | 0.01185 | 0.09359 | 0.032651 | -0.41639 |
| HT144SKINFV2 | -0.20453 | 0.090174 | 0.00361 | -0.01247 | -0.08735 | 0.15252 | 0.025751 | -0.21076 | -0.35667 | 0.03963 | -0.00229 | 0.23157 | -0.32209 | -0.04995 | 0.066016 | 0.139655 | -0.11889 | 0.08493 | 0.141995 | 0.19415 | 0.10946 | -0.06312 | 0.10874 | -0.042 | 0.07544 | 0.160189 | -0.28436 |
| RVH421SKINFV1 | -0.09424 | -0.03869 | -0.09531 | -0.12046 | -0.04442 | -0.09067 | -0.11831 | -0.08752 | -0.11927 | -0.12782 | -0.13699 | 0.10049 | -0.14286 | -0.25335 | -0.10819 | -0.04901 | -0.05123 | -0.01913 | 0.009953 | 0.18824 | 0.19819 | -0.13939 | 0.160745 | 0.03111 | 0.07425 | 0.202753 | -0.34695 |
| RPE1SS48 | 0.00464 | -0.02892 | -0.07326 | -0.10225 | 0.04192 | 0.01366 | 0.029579 | -0.26042 | -0.20673 | 0.00113 | -0.12423 | 0.07497 | -0.14597 | -0.2208 | -0.11147 | 0.118885 | -0.00206 | -0.00186 | 0.111398 | 0.13604 | 0.12407 | -0.21562 | 0.07107 | 0.02024 | 0.0053 | -0.00093 | -0.25619 |
| RPE1SS77 | -0.06042 | 0.079025 | -0.10687 | -0.14837 | -0.06844 | -0.03541 | -0.02986 | -0.43923 | -0.04115 | -0.05545 | -0.09023 | 0.10494 | -0.25693 | -0.29763 | -0.29728 | -0.03777 | -0.1672 | -0.12553 | 0.12112 | 0.18026 | 0.14524 | -0.23261 | 0.122245 | 0.05208 | 0.11487 | 0.010263 | -0.37008 |
| RPE1SS6 | -0.02827 | 0.074886 | -0.06881 | -0.06996 | -0.09723 | 0.04715 | -0.10092 | -0.29914 | -0.15279 | -0.08647 | -0.1268 | 0.22693 | -0.17572 | -0.28412 | -0.08931 | -0.03196 | -0.04733 | 0.06164 | 0.005851 | 0.17795 | 0.09962 | -0.31619 | 0.170104 | -0.00828 | 0.06214 | -0.0717 | -0.19613 |
| RPE1SS119 | 0.0312 | 0.105985 | -0.09637 | 0.02228 | -0.17015 | 0.09941 | -0.10864 | -0.223 | -0.04569 | -0.0561 | -0.04601 | 0.11619 | -0.04077 | -0.21181 | -0.13783 | 0.0287 | -0.02795 | 0.03673 | 0.028331 | 0.29302 | 0.15448 | -0.21131 | 0.133629 | -0.0137 | 0.0843 | 0.120255 | -0.21998 |
| RPE1SS51 | -0.03688 | 0.217748 | -0.21254 | -0.01787 | -0.11748 | -0.03806 | -0.11733 | -0.20034 | -0.0214 | -0.07003 | -0.41221 | -0.01866 | -0.19535 | -0.22336 | -0.09779 | -0.02233 | 0.08608 | -0.10631 | -0.11378 | 0.25544 | 0.16067 | -0.28091 | 0.081509 | -0.06057 | 0.12609 | 0.036866 | -0.17817 |
| PSS008 | 0.00773 | -0.26737 | -0.34653 | -0.11925 | -0.23714 | -0.0566 | -0.05795 | -0.5008 | 0.013445 | -0.10916 | -0.15276 | 0.09777 | -0.24203 | -0.06353 | 0.040528 | -0.09631 | -0.04865 | -0.14027 | -0.18074 | 0.2524 | 0.32209 | -0.1842 | -0.0191 | 0.21417 | 0.37061 | 0.268558 | -0.34243 |
| MAVER1 | -0.07884 | -0.0452 | -0.13723 | -0.1883 | -0.22255 | 0.06657 | 0.029417 | -0.47749 | -0.09595 | -0.14359 | 0.037169 | 0.06003 | -0.21335 | -0.51639 | -0.16846 | -0.0457 | -0.02716 | 0.0792 | 0.203058 | 0.14071 | -0.11282 | -0.14013 | 0.126627 | -0.07488 | 0.01753 | 0.137533 | -0.32435 |
| MESOV | -0.01295 | 0.020091 | -0.24329 | -0.01632 | -0.11459 | -0.0211 | -0.10485 | -0.07696 | -0.27884 | -0.0373 | -0.12173 | 0.07606 | -0.25439 | -0.22337 | -0.11669 | 0.007349 | -0.06556 | -0.05942 | 0.049814 | -0.03765 | 0.13499 | -0.19142 | 0.094902 | -0.00775 | 0.12789 | -0.13049 | -0.41728 |
| WM3211 | -0.07865 | 0.023748 | -0.57481 | -0.0616 | -0.16783 | -0.08587 | -0.09939 | -0.14939 | -0.23811 | -0.0704 | -0.16732 | 0.10437 | -0.36051 | -0.06567 | -0.06281 | -0.11576 | 0.03118 | 0.00508 | -0.07486 | 0.20343 | 0.09216 | -0.2598 | 0.065416 | 0.05431 | 0.21749 | -0.05424 | -0.72698 |
| M040416 | 0.01952 | 0.035861 | -0.1936 | -0.1055 | -0.08444 | -0.28669 | 0.208541 | -0.05364 | -0.02419 | -0.09892 | -0.07948 | 0.18829 | -0.05873 | 0.08096 | 0.001279 | -0.20536 | -0.07871 | 0.21673 | 0.021976 | 0.11295 | -0.06279 | -0.01482 | -0.0116 | 0.00875 | 0.62281 | 0.049866 | -0.57626 |
| M140325 | -0.04077 | 0.109179 | -0.43787 | -0.03135 | -0.15971 | -0.04982 | -0.08374 | -0.13295 | 0.068759 | -0.1225 | -0.17584 | 0.06698 | -0.58989 | -0.29254 | -0.12888 | -0.13543 | -0.11931 | -0.14632 | 0.106628 | 0.12428 | 0.04822 | -0.10121 | 0.058433 | 0.04168 | 0.07383 | 0.012081 | -0.41592 |
| MM160113 | -0.30096 | -0.02017 | -0.57069 | -0.01698 | -0.33904 | -0.19105 | 0.136481 | -0.46999 | 0.123484 | -0.23246 | -0.08667 | 0.09345 | -0.01513 | -0.19943 | 0.177886 | -0.11575 | 0.24243 | 0.01444 | -0.02538 | 0.20878 | 0.11949 | -0.06436 | 0.097706 | 0.16164 | 0.28322 | 0.280047 | -0.18227 |
| SNU739 | -0.10055 | 0.120032 | -0.56605 | -0.1336 | -0.148 | 0.05036 | 0.00015 | -0.29985 | 0.018757 | -0.11463 | -0.26732 | 0.17506 | -0.58576 | -0.71656 | -0.0701 | -0.0854 | -0.09856 | -0.06255 | -0.01447 | 0.06978 | 0.14276 | -0.14551 | 0.172392 | -0.0646 | 0.13371 | -0.0531 | -0.31492 |
| SNU1327 | -0.14528 | -0.02316 | -0.29709 | 0.00061 | 0.04924 | -0.13832 | -0.0739 | -0.47424 | -0.14681 | -0.18263 | 0.021067 | 0.17374 | 0.11779 | -0.19356 | -0.01199 | 0.065973 | 0.08048 | 0.24345 | -0.06427 | 0.07307 | 0.21348 | -0.04372 | 0.248656 | -0.0765 | -0.00638 | -0.25785 | -0.4595 |
| SNU2535 | -0.05459 | -0.03294 | -0.03214 | -0.19373 | -0.11054 | -0.01314 | -0.12032 | -0.18035 | -0.16122 | -0.15041 | -0.05347 | 0.12729 | -0.35604 | -0.34945 | 0.003778 | -0.01591 | 0.04468 | -0.05959 | 0.175184 | 0.19096 | 0.17954 | -0.14932 | 0.036252 | -0.00949 | 0.14764 | 0.005004 | -0.40236 |
| CCC5 | -0.09886 | 0.001872 | -0.18179 | -0.02731 | -0.09525 | -0.17097 | -0.03027 | -0.53922 | -0.05204 | -0.13761 | 0.031884 | 0.1931 | -0.26438 | -0.39168 | 0.097247 | -0.11581 | -0.13973 | 0.04754 | -0.09845 | 0.1819 | -0.04268 | -0.10495 | 0.102176 | 0.06303 | 0.17354 | 0.172856 | -0.43684 |
| ETCC016 | -0.19789 | -0.05758 | -0.33408 | -0.07783 | -0.04778 | -0.02428 | -0.05345 | -0.31899 | -0.17026 | -0.11781 | -0.05796 | 0.25001 | 0.00566 | -0.17924 | -0.07543 | -8.4E-05 | -0.00099 | 0.05938 | 0.084689 | 0.13661 | 0.1118 | -0.0673 | 0.146075 | -0.03133 | 0.15886 | 0.115256 | -0.30355 |
| JVE015 | -0.12911 | -0.04797 | -0.14287 | 0.075 | -0.08715 | -0.08029 | -0.245 | -0.22287 | 0.0387 | 0.22756 | -0.12655 | 0.07479 | -0.39152 | -0.26992 | -0.13493 | -0.10429 | -0.23623 | 0.12782 | 0.210873 | 0.11915 | 0.10215 | -0.53082 | 0.162406 | -0.00561 | 0.17625 | -0.23307 | -0.38606 |
| JVE127 | -0.08243 | 0.099299 | -0.76743 | -0.16535 | -0.12919 | -0.02354 | -0.12204 | -0.19232 | -0.05134 | 0.0068 | -0.07644 | 0.19592 | -0.20494 | -0.18174 | -0.00782 | -0.09411 | -0.00594 | 0.01096 | -0.01347 | -0.00543 | 0.24851 | -0.09503 | 0.032311 | -0.02331 | 0.24335 | -0.02685 | -0.41558 |
| JVE253 | -0.12104 | 0.050095 | -1.08044 | -0.17192 | -0.1183 | -0.16919 | -0.03351 | -0.12632 | -0.01865 | 0.01567 | -0.13135 | 0.18751 | -0.1206 | -0.21118 | -0.06985 | 0.004402 | -0.01169 | 0.06003 | 0.036534 | 0.21104 | 0.21704 | -0.17172 | 0.033326 | -0.07604 | 0.26253 | 0.169986 | -0.31746 |
| KP363T | -0.07382 | -0.10406 | -0.12405 | -0.03797 | -0.08471 | -0.15738 | 0.104469 | -0.14454 | -0.22627 | 0.0425 | -0.08842 | 0.09353 | -0.09206 | -0.14243 | -0.24383 | -0.05234 | 0.00244 | -0.03398 | 0.002685 | 0.13286 | 0.2191 | -0.07228 | 0.010461 | -0.01619 | 0.13459 | -0.00853 | -0.32178 |
| MAPACHS77 | -0.14383 | 0.028769 | -0.33927 | -0.04521 | -0.15608 | -0.00602 | -0.08468 | -0.24023 | -0.17395 | -0.1445 | -0.12724 | 0.15488 | -0.16834 | -0.19046 | -0.17977 | -0.08852 | -0.07541 | 0.13195 | -0.09673 | 0.15654 | 0.19227 | -0.23676 | 0.141974 | -0.08508 | 0.04965 | -0.06618 | -0.35409 |
| 170MGBA | -0.13073 | -0.02245 | -0.15824 | -0.0502 | -0.10566 | -0.17129 | -0.22724 | -0.28899 | -0.07285 | -0.13531 | -0.30396 | 0.07405 | -1.05872 | -0.40381 | -0.06827 | 0.001681 | -0.02632 | 0.06829 | 0.114026 | 0.11572 | 0.00592 | -0.18078 | 0.027587 | -0.08074 | 0.10103 | 0.050527 | -0.2884 |
| WM3772F | -0.28544 | -0.09246 | -0.09723 | -0.04488 | 0.01197 | 0.10118 | -0.03256 | -0.49224 | -0.32052 | 0.0171 | -0.12905 | 0.29875 | -0.09642 | 0.0212 | 0.015899 | -0.09741 | 0.0654 | -0.17807 | 0.026593 | 0.10813 | 0.22841 | -0.25052 | -0.00384 | -0.03094 | -0.02239 | 0.027493 | -0.4693 |
| S462 | -0.05101 | -0.10824 | -0.19027 | -0.23723 | -0.08127 | 0.01796 | 0.021994 | -0.54925 | 0.009103 | -0.13633 | -0.33004 | 0.10586 | -0.28761 | -0.0961 | -0.09039 | 0.055545 | -0.01602 | 0.24644 | -0.08512 | 0.14548 | 0.01428 | -0.1954 | 0.069949 | -0.04968 | 0.18835 | 0.008684 | -0.50515 |
| MPNST724 | -0.07397 | 0.060401 | -0.04934 | -0.10499 | -0.12148 | -0.04088 | -0.06242 | -0.18903 | -0.01872 | -0.03818 | -0.30327 | 0.07804 | -0.28403 | -0.31752 | -0.14181 | -0.08545 | -0.0356 | -0.0772 | -0.00231 | 0.13332 | 0.14472 | -0.0656 | 0.109212 | -0.01383 | 0.12133 | -0.14628 | -0.40766 |
| NCCLMS1C1 | -0.17888 | -0.04863 | -0.10759 | -0.18436 | -0.0572 | -0.07748 | -0.00236 | -0.22784 | -0.09838 | -0.12566 | -0.14866 | 0.09371 | -0.3612 | -0.34493 | -0.07956 | -0.07298 | 0.01678 | 0.00902 | -0.02969 | 0.24106 | 0.13302 | -0.19715 | 0.07986 | 0.0318 | -0.00248 | 0.073996 | -0.32394 |
| NCCMPNST1C1 | -0.11478 | -0.00605 | -0.23224 | -0.08921 | -0.05435 | -0.09772 | -0.117 | -0.11847 | 0.120043 | -0.03948 | -0.24465 | 0.16669 | -0.27469 | -0.40117 | -0.18949 | -0.05488 | -0.10584 | 0.06855 | -0.06682 | 0.10457 | 0.28552 | -0.05306 | 0.070632 | -0.03061 | 0.18183 | -0.08003 | -0.30957 |
| NCCMPNST2C1 | -0.11129 | 0.062953 | -0.34211 | -0.32911 | -0.10476 | 0.03247 | -0.09683 | -0.19644 | -0.04909 | -0.23701 | -0.45979 | 0.26078 | -0.57795 | -0.34749 | 0.131471 | 0.014274 | 0.13564 | -0.02108 | 0.004717 | 0.04369 | 0.0979 | -0.11445 | 0.111267 | -0.08263 | 0.09451 | 0.072648 | -0.1496 |
| PSS131R | -0.11997 | 0.004727 | -0.36156 | 0.03935 | -0.23083 | 0.19034 | 0.011076 | -0.08746 | 0.0502 | -0.00935 | -0.15885 | 0.0765 | -0.06774 | -0.86298 | -0.03451 | -0.19104 | -0.14832 | -0.17104 | -0.30464 | 0.10468 | 0.10885 | -0.10839 | -0.09231 | -0.01559 | 0.26248 | 0.031138 | -0.22402 |
| YUHOIN0650 | -0.06293 | 0.046783 | -0.15915 | -0.13327 | -0.11405 | 0.0621 | 0.019384 | -0.2084 | -0.12938 | -0.14708 | -0.08314 | 0.17118 | -0.20296 | -0.0819 | -0.09622 | -0.10326 | -0.05079 | 0.02232 | -0.0021 | 0.1791 | 0.15128 | -0.14376 | 0.010019 | -0.1173 | 0.05243 | 0.033989 | -0.40652 |
| SKNMM | -0.19759 | -0.03231 | -0.69502 | 0.06137 | -0.10088 | -0.01031 | -0.21894 | -0.20839 | -0.02987 | -0.06184 | -0.18391 | 0.11569 | -0.20588 | -0.23988 | -0.06776 | -0.11118 | -0.09629 | 0.08102 | 0.020496 | 0.13561 | 0.13764 | -0.14764 | -0.03333 | -0.09206 | 0.13907 | 0.027044 | -0.34415 |
| UPMD1 | 0.01061 | 0.13707 | -0.10895 | -0.12977 | -0.13234 | -0.06081 | -0.13829 | -0.45114 | 0.029962 | -0.08766 | -0.09061 | 0.15078 | -0.30052 | -0.29137 | -0.13145 | -0.04194 | -0.26954 | -0.06466 | -0.02363 | 0.22596 | 0.14851 | -0.21882 | 0.038054 | -0.01387 | 0.02353 | -0.00015 | -0.20746 |
| average | -0.11589 | 0.010923 | -0.30765 | -0.07952 | -0.12499 | -0.14881 | -0.01766 | -0.18011 | -0.11166 | -0.03301 | -0.09592 | 0.10676 | -0.20884 | -0.12742 | -0.07862 | -0.09275 | -0.05628 | 0.09439 | -0.03156 | 0.1253 | 0.11147 | -0.17236 | 0.075093 | -0.03994 | 0.14395 | 0.015012 | -0.37519 |
