## Supplement Table S5B for "Single-molecule behavior and cell-growth regulation in human RTKs"

| R | EC | IDRE | TM | IDRC | Linker | Tail | IDRE pos | IDRC pos | RK cluste | GxxxG | CRAC |
|---|---|---|---|---|---|---|---|---|---|---|---|
| NIHOVCAR3 | -0.1032 | 0.0449 | -0.1114 | 0.0174 | 0.0707 | -0.1032 | 0.108 | 0.0295 | -0.1831 | -0.043 | 0.1228 |
| HEL | -0.1121 | -0.0001 | 0.0663 | 0.1371 | -0.0284 | 0.0664 | -0.0031 | -0.0595 | -0.1379 | -0.278 | 0.3684 |
| HEL9217 | -0.1336 | 0.018 | -0.0349 | 0.0933 | 0.1533 | -0.1674 | 0.1714 | 0.0208 | -0.1673 | -0.2925 | 0.1593 |
| LS513 | -0.1184 | 0.1511 | -0.0163 | 0.0403 | 0.1794 | -0.1441 | 0.0885 | -0.0173 | -0.1481 | -0.0664 | 0.0179 |
| C2BBE1 | -0.0082 | 0.112 | 0.0921 | -0.1035 | 0.1567 | -0.1745 | -0.0294 | -0.2071 | -0.3928 | -0.1191 | 0.0704 |
| 253J | 0.0937 | 0.1127 | -0.0333 | -0.0583 | 0.0935 | -0.1825 | 0.0096 | -0.099 | -0.226 | -0.0518 | 0.1229 |
| HCC827 | 0.1303 | -0.0143 | -0.1147 | 0.0526 | -0.0449 | -0.3953 | 0.1077 | -0.0151 | -0.1309 | -0.1691 | 0.2223 |
| ONCODG1 | -0.097 | 0.1251 | -0.2424 | -0.008 | 0.0532 | -0.3655 | 0.1745 | -0.0454 | -0.3119 | -0.2005 | 0.2635 |
| HS294T | 0.081 | -0.0139 | 0.1427 | -0.034 | 0.1256 | -0.1002 | 0.012 | -0.0224 | -0.1105 | -0.089 | 0.0665 |
| NCIH1581 | -0.014 | -0.0379 | 0.1453 | -0.2794 | 0.0859 | -0.0886 | -0.1328 | -0.1692 | -0.3021 | -0.0494 | 0.1532 |
| SKBR3 | -0.0528 | 0.1201 | 0.055 | -0.1099 | 0.1882 | -0.2555 | 0.1235 | -0.2408 | -0.3637 | -0.043 | 0.0589 |
| T24 | 0.1153 | 0.0138 | -0.0429 | -0.0108 | 0.179 | -0.2061 | -0.0856 | 0 | -0.1434 | -0.1283 | 0.1976 |
| MCF7 | -0.1334 | 0.1324 | 0.0508 | 0.0587 | 0.1253 | -0.1447 | 0.0347 | 0.0221 | -0.1529 | 0.0881 | 0.1886 |
| NCIH1693 | -0.01 | 0.0284 | 0.152 | 0.1111 | 0.0128 | -0.1786 | -0.1474 | 0.0186 | -0.0328 | 0.0317 | 0.1723 |
| PATU8988S | -0.1777 | 0.1401 | -0.0109 | 0.0959 | 0.0753 | -0.1041 | 0.1052 | -0.0006 | -0.1862 | -0.187 | 0.0112 |
| PATU8988T | 0.1596 | 0.01 | 0.2163 | 0.0355 | -0.0086 | -0.0737 | -0.1314 | -0.0516 | -0.0671 | -0.0089 | 0.2332 |
| OPM2 | 0.1167 | 0.0061 | -0.0297 | 0.0681 | -0.0991 | -0.0399 | 0.0672 | -0.0149 | -0.0838 | 0.0023 | 0.2291 |
| CH157MN | -0.0513 | -0.0044 | 0.1166 | -0.0774 | 0.1248 | -0.0472 | -0.0396 | -0.0767 | -0.2055 | -0.1779 | 0.236 |
| KPL1 | 0.1316 | -0.1371 | 0.257 | 0.1524 | -0.1887 | -0.1851 | -0.0263 | 0.0979 | 0.0058 | -0.1818 | 0.1662 |
| HCC827GR5 | -0.0188 | 0.0609 | 0.028 | 0.0703 | -0.1068 | -0.2022 | 0.0888 | -0.0356 | -0.1214 | -0.0723 | 0.171 |
| PC14 | -0.0657 | 0.1262 | 0.1372 | -0.024 | 0.1389 | -0.1393 | 0.1063 | -0.1286 | -0.3005 | -0.1604 | 0.0569 |
| NCIH1650 | 0.0595 | -0.0033 | 0.0969 | 0.1343 | -0.0961 | -0.1038 | 0.0003 | 0.0891 | -0.0226 | 0.0231 | 0.3034 |
| U343 | -0.0049 | 0.0872 | 0.0139 | 0.1599 | -0.1764 | -0.1264 | 0.0024 | 0.057 | -0.0039 | 0.1367 | 0.2176 |
| S117 | -0.1532 | 0.246 | -0.0353 | 0.1876 | 0.103 | -0.3394 | 0.1933 | 0.1036 | 0.0229 | -0.0218 | 0.1796 |
| SKNMC | -0.1214 | 0.2516 | -0.1645 | 0.0794 | 0.1587 | -0.3899 | 0.2715 | -0.0169 | -0.2364 | 0.0755 | 0.0501 |
| U118MG | -0.0132 | 0.1173 | -0.0003 | 0.1796 | -0.041 | -0.0783 | 0.0622 | 0.0142 | -0.0454 | 0.0891 | 0.1313 |
| RDES | 0.0659 | 0.1156 | -0.0607 | 0.0075 | 0.0136 | -0.0847 | 0.1377 | -0.0161 | -0.1363 | 0.0127 | 0.2256 |
| PANC0203 | 0.1265 | 0.0926 | 0.102 | -0.0071 | 0.0621 | -0.241 | -0.0354 | -0.095 | -0.1851 | -0.1327 | 0.136 |
| MV411 | 0.0607 | 0.1044 | 0.1993 | 0.2282 | -0.0491 | -0.0487 | 0.1213 | 0.2102 | 0.1566 | -0.1835 | 0.0711 |
| GCIY | 0.0724 | 0.1258 | 0.0085 | 0.1007 | 0.0356 | -0.3516 | 0.0953 | 0.0391 | -0.1472 | -0.0113 | 0.1659 |
| TOV112D | 0.0813 | -0.0564 | 0.0215 | 0.0787 | -0.071 | -0.296 | 0.0631 | -0.0004 | -0.1166 | -0.0243 | 0.2372 |
| A673 | -0.0269 | 0.0893 | 0.0083 | -0.0259 | 0.0081 | -0.0595 | 0.2653 | -0.2087 | -0.2472 | -0.1866 | -0.1323 |
| KARPAS299 | -0.008 | 0.12 | -0.0117 | -0.0199 | 0.0839 | -0.3321 | 0.1287 | -0.1454 | -0.3223 | -0.1655 | 0.208 |
| HT1080 | 0.0283 | -0.1162 | 0.1956 | 0.151 | -0.1949 | 0.0189 | -0.1056 | -0.0653 | -0.1087 | -0.2096 | 0.1387 |
| D283MED | 0.0022 | 0.0685 | 0.0204 | -0.0475 | -0.0912 | -0.1921 | 0.1082 | -0.0074 | -0.0943 | -0.1304 | 0.0916 |
| PANC1005 | -0.0406 | 0.1211 | 0.0811 | -0.1663 | 0.176 | 0.008 | -0.0476 | -0.1753 | -0.2252 | 0.0017 | 0.0714 |
| HS683 | -0.019 | 0.0499 | -0.0045 | 0.1623 | -0.2256 | -0.1632 | 0.2006 | 0.1576 | -0.0008 | 0.0394 | -0.0014 |
| 697 | -0.1667 | 0.3091 | -0.166 | 0.015 | 0.145 | -0.3036 | 0.1479 | -0.1178 | -0.2971 | -0.0523 | 0.1794 |
| KU812 | 0.0712 | 0.0782 | 0.0168 | -0.0815 | 0.0161 | -0.1587 | -0.1459 | -0.1285 | -0.1764 | -0.173 | 0.0954 |
| U87MG | -0.0853 | 0.1237 | -0.073 | 0.1077 | -0.144 | -0.271 | 0.305 | 0.0373 | -0.1161 | -0.1245 | -0.0266 |
| NCO2 | -0.2648 | 0.1749 | -0.1231 | 0.2542 | -0.1974 | -0.221 | 0.1383 | 0.0657 | -0.1885 | -0.0582 | 0.1717 |
| MJ | -0.0146 | 0.009 | 0.0383 | 0.1423 | 0.051 | -0.2541 | 0.2425 | 0.0642 | 0.0112 | -0.3234 | 0.0122 |
| MHHNB11 | -0.0248 | -0.0183 | -0.0066 | 0.1585 | 0.0065 | -0.2241 | 0.1279 | 0.1522 | -0.0015 | -0.162 | 0.3443 |
| G292CLONEA141B1 | 0.0726 | 0.0176 | -0.1162 | 0.0348 | -0.0629 | 0.0563 | -0.0574 | -0.0171 | -0.1068 | 0.1548 | 0.0914 |
| T3M4 | -0.022 | 0.1518 | -0.2475 | -0.1274 | 0.2657 | -0.3351 | 0.157 | -0.23 | -0.3812 | -0.1179 | 0.0316 |
| ACCMESO1 | 0.0251 | 0.0152 | -0.0133 | 0.0545 | -0.0911 | -0.1107 | 0.0096 | 0.0429 | -0.0211 | 0.0918 | 0.1839 |
| PC3 | 0.1044 | 0.2585 | 0.1881 | 0.082 | 0.0924 | -0.2773 | -0.0666 | 0.006 | -0.0297 | 0.177 | 0.1993 |
| NCIH2452 | 0.1121 | 0.1341 | -0.0125 | 0.1927 | -0.127 | -0.1578 | -0.0206 | 0.0752 | -0.0598 | 0.046 | 0.3582 |
| PANC0504 | 0.0315 | -0.0173 | 0.0764 | 0.1154 | 0.0923 | -0.1007 | -0.0373 | 0.1127 | -0.0745 | -0.1639 | 0.1387 |
| HPAFII | 0.0834 | 0.0628 | -0.1136 | 0.0503 | -0.0179 | -0.2465 | -0.1401 | 0.0041 | -0.1492 | -0.1184 | 0.0961 |
| D341Med | -0.0451 | 0.0966 | -0.175 | 0.097 | 0.0375 | -0.2165 | 0.0847 | 0.0567 | -0.1525 | -0.1582 | 0.0669 |
| ZR751 | 0.0254 | 0.0862 | 0.0647 | -0.0995 | 0.0363 | -0.2072 | -0.0768 | -0.1819 | -0.3635 | -0.2197 | 0.0551 |
| GAMG | 0.0016 | 0.0727 | -0.0758 | 0.0764 | -0.171 | -0.0657 | 0.0476 | -0.002 | -0.1137 | 0.1977 | 0.0509 |
| SIMA | 0.1137 | 0.0497 | -0.0248 | 0.1568 | -0.0572 | -0.2272 | 0.0493 | 0.1636 | 0.0084 | 0.0214 | 0.2316 |
| KE37 | 0.1695 | -0.0825 | 0.0937 | 0.2001 | -0.1677 | -0.2204 | 0.0213 | 0.1025 | 0.0223 | -0.2647 | 0.3562 |
| CAOV4 | 0.2436 | 0.064 | 0.0869 | 0.0172 | 0.1317 | -0.3236 | -0.1684 | 0.0586 | -0.0959 | 0.029 | 0.4755 |
| KP3 | 0.066 | -0.0419 | 0.1008 | -0.0935 | 0.0654 | -0.1997 | -0.0362 | -0.18 | -0.2503 | -0.1924 | 0.1455 |
| HCC1187 | 0.0948 | 0.0075 | 0.1703 | 0.1984 | -0.1269 | -0.1716 | -0.1165 | 0.032 | 0.0536 | 0.0228 | 0.5152 |
| OCIAML2 | -0.0229 | 0.2251 | 0.0558 | 0.162 | 0.0691 | -0.2166 | 0.0813 | 0.0488 | -0.0886 | -0.0178 | 0.2481 |
| SU8686 | 0.0923 | 0.1115 | 0.0668 | -0.0361 | -0.1612 | -0.0256 | -0.1265 | -0.1999 | -0.2237 | -0.0405 | 0.1756 |
| VCAP | -0.2311 | 0.2496 | -0.0071 | 0.1165 | 0.1834 | -0.1081 | 0.1769 | -0.0015 | -0.1521 | 0.0179 | 0.1736 |
| HUPT3 | -0.0213 | 0.0835 | 0.0706 | -0.1956 | 0.2687 | -0.1569 | 0.0059 | -0.1216 | -0.3274 | -0.1697 | 0.0318 |
| CHP212 | -0.0582 | -0.052 | 0.0058 | 0.102 | -0.1017 | -0.0618 | 0.0209 | -0.0127 | 0.0337 | -0.3068 | 0.0342 |
| COV434 | -0.0399 | -0.0127 | -0.0355 | 0.0859 | 0.1057 | -0.3267 | 0.1758 | 0.0186 | -0.2298 | -0.1992 | 0.1155 |
| OCILY19 | 0.1144 | 0.0506 | -0.0908 | 0.0563 | -0.0304 | -0.1724 | 0.0262 | 0.045 | -0.0607 | -0.0813 | 0.2341 |
| SLR20 | -0.0964 | -0.0157 | 0.152 | 0.1477 | -0.0505 | -0.0926 | 0.0424 | 0.0924 | -0.0379 | -0.1924 | 0.0563 |
| LN319 | -0.0312 | 0.1418 | 0.0563 | -0.0188 | 0.2124 | -0.2456 | 0.1972 | -0.1638 | -0.3505 | -0.1378 | 0.0822 |
| JHOS2 | 0.1202 | 0.0505 | 0.2001 | -0.1686 | 0.0897 | -0.2092 | -0.0682 | -0.3192 | -0.3311 | -0.062 | 0.1013 |
| HS729 | 0.0399 | 0.1487 | 0.0314 | 0.0046 | 0.0228 | -0.0309 | -0.0203 | -0.026 | -0.0973 | 0.1637 | -0.0608 |
| 8MGBA | -0.0323 | -0.0308 | -0.2859 | -0.0191 | 0.1774 | -0.3059 | 0.129 | 0.0152 | -0.2587 | -0.1382 | 0.2546 |
| CFPAC1 | -0.1016 | 0.1659 | -0.0321 | -0.1224 | 0.3291 | -0.1421 | 0.0824 | -0.2149 | -0.4373 | -0.1207 | -0.013 |
| PANC0327 | 0.1065 | 0.0466 | 0.0979 | 0.0033 | -0.0085 | -0.0947 | -0.0414 | -0.1027 | -0.1901 | 0.0832 | 0.2191 |
| SNU308 | -0.0561 | 0.0083 | -0.0267 | -0.178 | 0.1891 | -0.2378 | 0.1133 | -0.2228 | -0.3684 | -0.1196 | -0.0104 |
| CAL29 | 0.0469 | 0.075 | 0.0381 | -0.1272 | 0.137 | -0.3677 | 0.0338 | -0.1222 | -0.3087 | 0.0043 | 0.2376 |
| HCC2429 | 0.0173 | -0.0713 | 0.119 | 0.0874 | -0.14 | -0.0574 | -0.0159 | -0.0542 | -0.0786 | -0.0422 | 0.2354 |
| RERFGC1B | -0.1716 | 0.1929 | -0.2176 | 0.0551 | 0.0581 | -0.1022 | 0.2219 | 0.1353 | -0.018 | 0.1058 | 0.1274 |
| SKLMS1 | 0.0845 | 0.037 | 0.0144 | 0.0384 | -0.0365 | -0.2378 | -0.0163 | -0.0194 | -0.1295 | 0.0601 | 0.0257 |
| THP1 | -0.0933 | 0.023 | 0.0475 | -0.0677 | -0.0444 | -0.0636 | -0.1297 | -0.1659 | -0.282 | -0.162 | 0.0815 |
| T47D | 0.019 | 0.0597 | 0.0529 | 0.0644 | 0.0864 | -0.2074 | -0.0782 | -0.0855 | -0.2629 | -0.2375 | 0.1302 |
| HS578T | 0.1266 | -0.0483 | 0.122 | 0.0785 | -0.134 | -0.2027 | -0.1323 | 0.0582 | -0.0258 | -0.024 | 0.2831 |
| SKNSH | -0.0695 | 0.1587 | -0.015 | 0.0236 | 0.0997 | 0.0164 | 0.1287 | 0.0952 | -0.0646 | -0.1228 | 0.036 |
| HCC2935 | 0.11 | -0.0155 | 0.0599 | -0.011 | -0.1289 | 0.0618 | -0.0975 | -0.0816 | -0.0308 | 0.0414 | 0.2307 |
| JM1 | -0.0919 | 0.0241 | -0.0312 | 0.08 | 0.0388 | -0.0949 | 0.0964 | -0.0397 | -0.1747 | -0.0883 | 0.1285 |
| M059K | -0.0605 | 0.0956 | -0.0945 | -0.0792 | 0.146 | -0.0911 | 0.0237 | -0.0382 | -0.2717 | 0.0616 | 0.0477 |
| NCIH2052 | -0.1689 | 0.1872 | 0.133 | -0.1162 | -0.0002 | 0.0758 | 0.1827 | -0.1605 | -0.2222 | -0.0439 | 0.0963 |
| SW1990 | 0.1078 | 0.1181 | 0.0167 | 0.0385 | 0.1063 | -0.2928 | 0.0105 | 0.0149 | -0.0943 | -0.0131 | 0.111 |
| OSRC2 | -0.0206 | 0.007 | 0.1492 | 0.0599 | -0.1221 | -0.1555 | -0.1132 | 0.1276 | -0.072 | -0.1102 | 0.051 |
| BT12 | 0.1026 | 0.0083 | -0.0658 | 0.0402 | -0.036 | -0.1245 | -0.0142 | 0.0495 | -0.1068 | 0.0345 | 0.0705 |
| CORL105 | -0.0243 | 0.0946 | 0.1872 | -0.0777 | 0.194 | -0.208 | -0.0015 | -0.1009 | -0.2382 | -0.2852 | -0.037 |
| SW579 | 0.0629 | -0.0421 | 0.1213 | 0.0377 | -0.009 | 0.0018 | -0.2246 | -0.003 | -0.0415 | 0.0829 | 0.1356 |
| PANC1 | 0.0537 | 0.1403 | 0.0971 | 0.0346 | 0.0542 | -0.115 | 0.018 | -0.0561 | -0.1092 | -0.1239 | 0.1231 |
| NOMO1 | -0.0794 | 0.0328 | 0.1957 | 0.0966 | -0.1009 | 0.0263 | 0.0154 | 0.074 | -0.0095 | -0.0381 | 0.2521 |
| RD | -0.0547 | 0.0212 | -0.1266 | 0.0986 | -0.0532 | -0.2188 | 0.1208 | 0.0097 | -0.1322 | -0.2063 | 0.1706 |
| CAL62 | 0.0793 | -0.059 | -0.0747 | 0.1541 | -0.2343 | -0.2051 | 0.0434 | 0.0648 | -0.0528 | -0.1207 | 0.2576 |
| LOUNH91 | -0.0936 | 0.1176 | 0.1525 | 0.0066 | 0.2 | -0.2549 | 0.0693 | -0.0067 | -0.2589 | -0.1506 | 0.1111 |
| HS766T | -0.005 | 0.012 | 0.1499 | -0.1415 | 0.1183 | -0.1358 | -0.1014 | -0.1132 | -0.1939 | -0.1203 | 0.0921 |
| SCC9 | 0.0111 | 0.1012 | 0.1231 | -0.0024 | 0.1222 | -0.1544 | 0.0271 | -0.1595 | -0.3044 | -0.1655 | -0.0304 |
| SNU869 | -0.0554 | 0.0521 | -0.1375 | 0.031 | 0.0247 | -0.0808 | 0.0261 | 0.0623 | -0.1145 | -0.1464 | 0.0326 |
| L363 | -0.0586 | 0.0979 | -0.0274 | 0.0695 | 0.023 | -0.1537 | 0.0352 | 0.0696 | -0.1294 | -0.0832 | 0.2126 |
| CORL311 | 0.0267 | 0.0137 | 0.0336 | 0.0631 | 0.0554 | -0.2537 | 0.0219 | 0.0776 | -0.0081 | 0.0443 | 0.1502 |
| SCC25 | 0.0928 | -0.015 | 0.1721 | -0.0038 | 0.0917 | -0.1448 | -0.0601 | -0.0767 | -0.18 | -0.0846 | 0.1415 |
| RCC10RGB | 0.006 | 0.0014 | -0.0365 | -0.0338 | -0.0314 | -0.1624 | 0.0876 | -0.1336 | -0.2537 | -0.2463 | 0.0029 |
| HDMYZ | -0.1007 | -0.0433 | -0.0252 | 0.0885 | -0.0466 | -0.157 | -0.0884 | 0.119 | 0.1649 | 0.0015 | 0.1318 |
| BHT101 | 0.2263 | 0.1043 | 0.1749 | 0.0795 | 0.0821 | -0.3214 | 0.0442 | 0.1366 | 0.0306 | -0.0304 | 0.2145 |
| MFE280 | -0.0246 | -0.0131 | -0.0233 | -0.2069 | 0.2492 | -0.2145 | 0.07 | -0.2063 | -0.4125 | -0.2233 | 0.034 |
| SET2 | -0.1622 | 0.1575 | -0.0057 | 0.1009 | 0.1097 | 0.0074 | 0.0547 | -0.0942 | -0.192 | -0.3251 | 0.0536 |
| EOL1 | 0.17 | -0.0074 | -0.0466 | 0.0224 | -0.1778 | -0.245 | 0.1868 | -0.145 | -0.186 | -0.125 | 0.2405 |
| NMCG1 | -0.1495 | 0.1367 | 0.1443 | -0.0724 | 0.1125 | -0.1369 | -0.0421 | -0.0826 | -0.0984 | 0.0123 | 0.0401 |
| COLO320 | -0.1394 | 0.0827 | -0.2916 | -0.0287 | 0.2135 | -0.1935 | 0.1141 | -0.0339 | -0.2305 | -0.2118 | 0.0012 |
| LP1 | -0.0839 | -0.0884 | 0.0083 | 0.0988 | -0.1944 | -0.0004 | 0.002 | 0.0946 | 0.033 | -0.1545 | 0.0921 |
| C8166 | 0.0093 | -0.0347 | 0.0448 | 0.1709 | 0.0183 | -0.2307 | 0.2472 | 0.1203 | -0.0722 | -0.1507 | 0.2718 |
| DETROIT562 | -0.0346 | 0.1271 | 0.2099 | -0.128 | 0.1491 | -0.2152 | -0.0114 | -0.2648 | -0.3889 | 0.0676 | 0.1084 |
| SNU1079 | 0.0454 | -0.0584 | 0.1486 | 0.0418 | -0.0839 | -0.2075 | 0.027 | -0.1333 | -0.2356 | -0.1043 | 0.0523 |
| DAOY | 0.0655 | 0.0182 | 0.0302 | 0.0767 | -0.1767 | -0.275 | 0.0176 | 0.0038 | -0.094 | 0.13 | 0.1114 |
| CAL120 | 0.2138 | -0.0107 | -0.111 | 0.0099 | -0.0482 | -0.1019 | -0.0462 | 0.0123 | -0.0583 | 0.086 | 0.0363 |
| HUPT4 | -0.0384 | 0.0831 | 0.1284 | -0.1106 | 0.0905 | -0.0403 | 0.0205 | -0.2674 | -0.3692 | -0.1277 | -0.0514 |
| LN382 | 0.0207 | 0.1659 | -0.0288 | 0.0582 | -0.0705 | -0.1516 | 0.0304 | -0.0293 | -0.1082 | 0.0652 | 0.2618 |
| JHESOAD1 | 0.0499 | 0.0726 | 0.0405 | -0.0322 | 0.1151 | -0.2523 | -0.0484 | -0.0934 | -0.2985 | -0.1151 | 0.0958 |
| JHH6 | -0.0622 | -0.04 | -0.0731 | 0.1008 | -0.0505 | -0.2252 | 0.095 | -0.0365 | -0.2217 | -0.0423 | 0.1461 |
| A375 | -0.0525 | 0.0267 | 0.0315 | 0.1607 | -0.0309 | -0.0631 | -0.0397 | 0.091 | -0.028 | -0.177 | 0.2017 |
| SNU398 | 0.1876 | 0.1126 | 0.019 | -0.0567 | 0.0818 | -0.3132 | 0.0471 | -0.0237 | 0.0049 | 0.023 | 0.2093 |
| ASPC1 | -0.0753 | 0.1833 | -0.1022 | 0.0078 | 0.0725 | -0.269 | 0.1346 | 0.004 | -0.081 | 0.0275 | 0.0148 |
| HCC1937 | 0.1223 | 0.0377 | 0.2945 | -0.0749 | 0.0912 | -0.3102 | -0.0904 | -0.1094 | -0.2127 | -0.0214 | 0.0691 |
| SUPM2 | 0.1025 | 0.1763 | 0.0964 | 0.1625 | 0.0991 | -0.3096 | 0.0537 | 0.1521 | 0.2169 | -0.0746 | 0.2676 |
| KPNYN | -0.1611 | 0.0092 | -0.0115 | 0.1126 | -0.0183 | -0.0227 | 0.0388 | 0.1514 | -0.0075 | -0.1315 | 0.1471 |
| BICR31 | -0.0407 | 0.0967 | 0.2522 | -0.0086 | 0.0922 | -0.188 | 0.073 | -0.1076 | -0.2578 | -0.0619 | 0.1932 |
| KALS1 | -0.0157 | -0.0814 | 0.0595 | -0.0064 | -0.0828 | 0.0497 | -0.1002 | -0.0101 | -0.1353 | -0.0754 | 0.2845 |
| U251MG | 0.1449 | -0.0355 | 0.2243 | -0.1409 | 0.0232 | -0.1294 | -0.1907 | -0.0545 | -0.1455 | 0.0325 | 0.1063 |

| R | EC | IDRE | TM | IDRC | Linker | Tail | IDRE pos | IDRC pos | RK cluste | GxxxG | CRAC |
|---|---|---|---|---|---|---|---|---|---|---|---|
| CAKI2 | -0.123 | 0.0929 | -0.0864 | -0.0474 | 0.0122 | -0.3507 | 0.2375 | -0.2168 | -0.5172 | -0.1001 | 0.133 |
| PANC0403 | -0.0561 | 0.149 | 0.1211 | -0.1128 | 0.2083 | -0.0938 | 0.0117 | -0.2131 | -0.3507 | -0.1786 | 0.0271 |
| JHOM1 | -0.0073 | 0.003 | 0.1096 | 0.0638 | 0.0232 | -0.0379 | -0.0715 | 0.0446 | -0.1014 | -0.2258 | 0.0548 |
| SCC4 | -0.0142 | 0.1866 | 0.0354 | 0.006 | 0.0653 | -0.2639 | 0.1551 | -0.1334 | -0.3337 | -0.01 | 0.1978 |
| DANG | 0.0337 | 0.0686 | -0.0544 | -0.0625 | 0.0914 | -0.2374 | 0.0387 | -0.1713 | -0.2423 | -0.2197 | 0.1094 |
| DKMG | -0.0665 | 0.1108 | -0.0343 | 0.2421 | -0.2298 | -0.1029 | -0.0119 | 0.1196 | 0.0566 | 0.1482 | 0.2644 |
| SLR23 | 0.0081 | 0.1527 | 0.1044 | -0.1274 | 0.0917 | -0.2109 | -0.0898 | -0.1085 | -0.1909 | -0.0555 | -0.0047 |
| OCUM1 | -0.0172 | 0.1501 | -0.0756 | 0.0917 | 0.0333 | -0.3411 | 0.1684 | 0.0025 | -0.0447 | 0.138 | -0.0255 |
| AU565 | 0.0098 | 0.1338 | 0.0733 | -0.2162 | 0.264 | -0.2767 | 0.0786 | -0.2773 | -0.3984 | -0.0121 | -0.0568 |
| CL11 | 0.1212 | 0.0671 | 0.2295 | -0.0247 | 0.0147 | -0.1418 | -0.213 | -0.0862 | -0.2118 | -0.0217 | 0.1263 |
| KMRC20 | 0.1202 | 0.0695 | 0.0741 | 0.0615 | -0.1538 | -0.2728 | 0.079 | -0.0859 | -0.2486 | -0.1019 | 0.149 |
| NCIH2887 | 0.1034 | 0.0245 | 0.1862 | -0.0358 | -0.0424 | -0.1224 | -0.1402 | 0.0443 | -0.1059 | -0.0701 | 0.1198 |
| LS1034 | 0.0088 | 0.0804 | -0.1999 | 0.1277 | 0.0232 | -0.2789 | 0.2217 | 0.1877 | -0.0439 | -0.1534 | 0.253 |
| COLO201 | 0.0008 | 0.0642 | -0.1241 | 0.1751 | 0.0021 | -0.2956 | 0.1642 | 0.0096 | -0.133 | -0.1507 | 0.1894 |
| LMSU | 0.1741 | 0.0279 | -0.006 | 0.1731 | -0.1827 | -0.188 | -0.011 | 0.0848 | 0.0432 | 0.1132 | 0.2177 |
| COV318 | 0.2037 | 0.0166 | 0.2398 | 0.1658 | -0.0835 | -0.1569 | -0.0108 | 0.1683 | 0.163 | -0.157 | 0.1218 |
| CORL279 | 0.041 | -0.0539 | 0.1208 | 0.1136 | -0.1727 | 0.2016 | -0.2445 | 0.0071 | -0.0009 | -0.1263 | 0.1441 |
| DU4475 | 0.0159 | 0.106 | 0.2126 | 0.1372 | -0.0162 | -0.244 | 0.1468 | 0.0893 | -0.04 | 0.0134 | 0.0655 |
| KELLY | -0.1649 | 0.1349 | -0.1238 | 0.1736 | 0.0139 | -0.215 | 0.2187 | 0.145 | -0.049 | -0.1047 | 0.2347 |
| SKNAS | 0.0202 | 0.1575 | -0.1627 | 0.1103 | 0.0158 | -0.2505 | 0.248 | 0.078 | -0.0325 | -0.0775 | 0.0972 |
| RERFLCAI | -0.014 | 0.0678 | -0.0318 | 0.0228 | -0.0156 | -0.0679 | 0.0332 | -0.068 | -0.188 | -0.1073 | 0.1787 |
| UOK101 | 0.1047 | 0.1016 | -0.0708 | 0.009 | 0.0384 | -0.1813 | -0.132 | -0.0344 | -0.0629 | -0.0514 | -0.0087 |
| KASUMI1 | -0.1031 | 0.2772 | -0.0704 | 0.1207 | 0.1933 | -0.1615 | 0.0085 | 0.1245 | 0.0249 | 0.1405 | 0.2576 |
| CALU6 | -0.0145 | 0.0729 | -0.0824 | 0.0503 | -0.0274 | -0.2659 | 0.0694 | 0.0513 | -0.1266 | -0.1191 | 0.1561 |
| KP4 | -0.1854 | 0.1593 | 0.0504 | 0.1059 | -0.2364 | 0.0664 | 0.0273 | -0.0078 | -0.0279 | 0.0065 | 0.1369 |
| SNU213 | -0.0345 | 0.0966 | 0.0219 | -0.1141 | 0.144 | -0.3066 | 0.0464 | -0.2292 | -0.4285 | -0.1416 | 0.0612 |
| AM38 | -0.0953 | 0.2463 | 0.0133 | 0.0453 | 0.0835 | -0.284 | 0.251 | 0.026 | -0.0648 | 0.1015 | 0.2048 |
| SUDHL10 | 0.2147 | -0.0592 | 0.1014 | 0.1644 | -0.1841 | -0.2175 | -0.0655 | 0.0345 | -0.0383 | -0.1309 | 0.4325 |
| SLR24 | -0.0331 | 0.042 | 0.1639 | -0.0623 | 0.1102 | -0.2219 | -0.0198 | -0.1173 | -0.3092 | -0.0995 | 0.1044 |
| SF539 | 0.1117 | -0.0528 | 0.1218 | 0.1871 | -0.2365 | -0.0388 | -0.2285 | 0.0409 | 0.0093 | 0.0582 | 0.2699 |
| HS852T | -0.0661 | 0.1653 | 0.0514 | 0.035 | 0.0112 | -0.0832 | 0.0844 | 0.0019 | -0.163 | 0.1103 | 0.3004 |
| HCC38 | 0.1326 | 0.0961 | 0.1018 | 0.0169 | -0.0119 | -0.0531 | -0.0286 | -0.0539 | -0.0962 | 0.05 | 0.149 |
| HCC1419 | -0.0954 | 0.1308 | -0.0228 | -0.2025 | 0.2795 | -0.3487 | 0.1506 | -0.271 | -0.4382 | -0.0655 | -0.0115 |
| COV362 | -0.0466 | -0.047 | 0.1354 | 0.118 | -0.1168 | -0.2122 | 0.059 | 0.0872 | -0.108 | -0.0487 | 0.3058 |
| EWS502 | -0.0052 | 0.223 | -0.0887 | 0.1178 | 0.0049 | -0.1417 | 0.1972 | 0.0789 | -0.032 | -0.0466 | 0.1924 |
| SNU840 | 0.0606 | 0.0641 | 0.0947 | -0.0559 | 0.1276 | -0.2445 | -0.0574 | -0.0645 | -0.2303 | -0.0851 | 0.2143 |
| KP2 | 0.1439 | -0.0111 | -0.0159 | 0.0754 | -0.1042 | -0.2056 | -0.0461 | -0.0785 | -0.0815 | -0.0633 | 0.1296 |
| NCIH1755 | -0.0438 | 0.0375 | -0.0634 | -0.0881 | -0.0748 | -0.0623 | -0.0149 | -0.1124 | -0.2868 | -0.0262 | 0.1097 |
| SNU1033 | -0.0244 | -0.0445 | -0.1232 | 0.1185 | -0.0423 | -0.2267 | 0.0379 | 0.2376 | 0.0648 | -0.0528 | 0.2907 |
| BT549 | 0.1924 | 0.0534 | 0.0715 | 0.0442 | -0.1097 | -0.1867 | -0.02 | 0.0236 | -0.0462 | 0.0801 | 0.1858 |
| NCIH209 | 0.0337 | 0.0985 | 0.0141 | 0.072 | 0.0961 | -0.2483 | 0.0295 | 0.0543 | -0.1551 | -0.2058 | 0.1553 |
| OV90 | -0.0746 | 0.0847 | -0.0725 | 0.1471 | 0.0091 | -0.2047 | 0.0545 | 0.0787 | -0.0617 | -0.1709 | 0.1773 |
| NCIH841 | 0.1469 | -0.0177 | 0.0594 | 0.0722 | -0.2135 | -0.0848 | 0.0892 | 0.0252 | -0.0573 | -0.146 | 0.3068 |
| KLE | -0.1313 | 0.0221 | -0.1992 | -0.1328 | 0.0729 | -0.2284 | 0.1269 | -0.0964 | -0.3201 | -0.0785 | 0.0602 |
| NB4 | -0.1643 | 0.116 | -0.1205 | 0.0896 | 0.0443 | -0.1648 | 0.0666 | -0.073 | -0.2269 | -0.1804 | 0.0998 |
| EM2 | -0.2775 | 0.1234 | -0.2494 | 0.0325 | 0.1075 | -0.1182 | 0.1667 | 0.0101 | -0.2716 | -0.1807 | 0.1503 |
| OUMS23 | -0.0757 | -0.0121 | -0.0049 | 0.0068 | -0.0377 | -0.349 | 0.2901 | -0.0432 | -0.1949 | -0.0134 | 0.293 |
| SNU1077 | -0.0832 | 0.0726 | -0.0714 | 0.085 | 0.0446 | -0.1988 | 0.1149 | -0.0136 | -0.2234 | -0.1011 | 0.1308 |
| SNU5 | 0.2181 | 0.0584 | -0.0653 | 0.1528 | 0.0911 | -0.2334 | -0.0434 | 0.0979 | -0.0884 | -0.1552 | 0.0868 |
| WM115 | 0.2363 | -0.0237 | 0.0313 | -0.0057 | 0.0921 | -0.4054 | 0.0782 | -0.1258 | -0.1792 | 0.0486 | 0.2716 |
| ECGI10 | -0.027 | 0.1035 | 0.0336 | -0.1401 | 0.2106 | -0.3181 | 0.0633 | -0.2526 | -0.4482 | -0.1017 | 0.0411 |
| PK1 | -0.0534 | 0.1902 | 0.0519 | -0.154 | 0.1987 | -0.1911 | 0.105 | -0.2575 | -0.4018 | -0.067 | 0.0469 |
| EFO21 | -0.0021 | -0.0344 | 0.0897 | -0.0147 | 0.0058 | -0.154 | -0.1226 | -0.0067 | -0.1076 | -0.1225 | 0.1265 |
| IMR32 | 0.1616 | 0.1501 | -0.0844 | 0.1887 | 0.066 | -0.4051 | 0.0778 | 0.1389 | 0.0391 | -0.0832 | 0.1301 |
| NCIH2122 | -0.0886 | 0.1576 | -0.193 | 0.0752 | 0.1382 | -0.2193 | 0.0897 | 0.0379 | 0.0054 | -0.0108 | 0.0823 |
| SKNBE2 | -0.0633 | 0.1563 | -0.2391 | 0.0973 | 0.1093 | -0.1253 | 0.1047 | 0.061 | -0.0632 | -0.183 | 0.1708 |
| KMRC3 | -0.1024 | 0.1088 | 0.0478 | -0.1741 | 0.1527 | 0.0738 | -0.0627 | -0.1829 | -0.268 | -0.0334 | -0.1198 |
| KARPAS422 | -0.032 | 0.0301 | 0.0062 | 0.2529 | -0.0582 | -0.1019 | 0.0169 | 0.0606 | -0.0994 | -0.2335 | 0.3781 |
| SNU886 | -0.0417 | -0.0212 | -0.0642 | 0.0748 | -0.0608 | -0.1534 | 0.0806 | -0.0318 | -0.2142 | 0.0584 | 0.1229 |
| TUHR14TKB | 0.0105 | -0.1128 | 0.0169 | 0.1297 | 0.0044 | -0.2407 | -0.0043 | 0.0581 | -0.1408 | -0.0686 | 0.1014 |
| TE10 | -0.203 | 0.0956 | -0.0347 | 0.0302 | 0.0376 | -0.1382 | 0.1419 | -0.0825 | -0.3101 | -0.0028 | 0.1249 |
| MPP89 | 0.0054 | 0.0651 | -0.0367 | -0.0361 | 0.1485 | -0.1092 | -0.093 | -0.1305 | -0.2162 | -0.0436 | 0.1126 |
| PSN1 | 0.0252 | 0.022 | -0.0966 | 0.0949 | -0.0193 | -0.1824 | 0.1307 | 0.1183 | -0.0094 | -0.0232 | 0.0745 |
| HT144 | 0.1305 | 0.0292 | 0.2691 | 0.0916 | -0.0706 | -0.2999 | 0.0243 | 0.0562 | -0.0003 | -0.0761 | 0.1544 |
| 42MGBA | 0.0353 | 0.0207 | 0.2057 | 0.1833 | -0.1445 | -0.099 | -0.0804 | 0.093 | 0.1183 | 0.1196 | 0.1497 |
| JHOC5 | -0.0111 | 0.0894 | 0.0696 | -0.1371 | 0.1834 | -0.3001 | 0.0494 | -0.1955 | -0.4105 | -0.0516 | -0.0062 |
| SNU620 | -0.0067 | 0.1715 | -0.0703 | -0.0312 | 0.2478 | -0.149 | 0.047 | -0.0696 | -0.27 | -0.0352 | 0.0821 |
| JURLMK1 | -0.1328 | 0.0648 | -0.0992 | -0.0461 | 0.1833 | -0.0586 | 0.0559 | -0.0209 | -0.2571 | -0.0986 | 0.2282 |
| CCFSTTG1 | 0.1527 | 0.1322 | 0.0693 | 0.0746 | -0.0281 | -0.2689 | 0.031 | 0.0959 | -0.0593 | 0.0762 | 0.1704 |
| EFM19 | -0.0272 | 0.1997 | 0.1861 | -0.0538 | 0.0834 | -0.0742 | -0.0646 | 0.0577 | -0.0243 | -0.0252 | 0.126 |
| ISTMES2 | 0.1028 | 0.1269 | 0.0479 | -0.0152 | 0.0092 | -0.0504 | 0.0058 | -0.0599 | -0.153 | 0.0571 | 0.1245 |
| YAPC | -0.0439 | 0.2073 | 0.0069 | -0.0037 | 0.1694 | -0.1353 | 0.0821 | -0.0108 | -0.172 | -0.0878 | 0.0157 |
| DB | -0.0803 | 0.141 | -0.1146 | 0.0282 | 0.1286 | -0.2426 | 0.2084 | 0.0186 | -0.1499 | -0.1704 | 0.3229 |
| MSTO211H | 0.0032 | 0.095 | -0.0885 | 0.1138 | -0.1548 | 0.0005 | 0.004 | -0.0184 | -0.0721 | -0.1271 | 0.1079 |
| OCIAML3 | -0.0805 | 0.0857 | -0.103 | 0.22 | 0.0729 | -0.1956 | 0.1971 | 0.2754 | 0.1126 | -0.2553 | 0.2064 |
| NCIH3122 | -0.0892 | -0.0401 | 0.032 | 0.0718 | 0.0872 | -0.2684 | 0.1898 | 0.0732 | -0.0553 | -0.1333 | 0.2051 |
| HCC461 | 0.0856 | 0.164 | 0.187 | 0.0567 | 0.0737 | -0.1531 | -0.0728 | -0.0317 | -0.1026 | 0.027 | 0.0849 |
| SKNFI | -0.1369 | 0.2093 | -0.1836 | 0.1355 | -0.013 | -0.0909 | 0.1776 | 0.0579 | -0.1049 | -0.0445 | 0.0922 |
| NCIH522 | 0.0414 | 0.1168 | 0.0327 | 0.1478 | 0.0104 | -0.1667 | 0.0694 | 0.0296 | -0.0892 | -0.1412 | 0.192 |
| SNU668 | -0.0779 | 0.1515 | -0.0656 | -0.033 | 0.115 | -0.3453 | 0.1273 | -0.0337 | -0.1529 | 0.0329 | 0.1562 |
| JVM3 | 0.1124 | 0.0978 | -0.116 | 0.271 | -0.0923 | -0.168 | 0.1949 | 0.0882 | 0.0138 | -0.1031 | 0.1295 |
| RPMI7951 | -0.1507 | 0.0623 | -0.0283 | -0.12 | 0.1045 | -0.1241 | -0.0902 | -0.2668 | -0.3239 | -0.0286 | 0.1899 |
| COLO678 | 0.0379 | 0.0278 | 0.1226 | 0.0126 | 0.1246 | -0.0877 | -0.0612 | -0.0674 | -0.1733 | -0.2645 | 0.1507 |
| HCC1428 | 0.0172 | 0.0618 | 0.0627 | -0.0612 | -0.0367 | -0.2068 | 0.1489 | -0.0739 | -0.22 | -0.0411 | 0.2413 |
| CAPAN1 | 0.0389 | 0.0541 | 0.1095 | -0.1212 | 0.1095 | -0.1194 | -0.0634 | -0.2241 | -0.3228 | -0.1388 | 0.1096 |
| NCIH82 | -0.0403 | 0.1149 | -0.1347 | 0.0463 | 0.0591 | -0.3273 | -0.0007 | -0.0689 | -0.2127 | -0.1182 | 0.0552 |
| MKN45 | -0.0532 | 0.0754 | -0.0969 | 0.0443 | 0.1016 | -0.397 | 0.2097 | 0.0483 | -0.0916 | -0.0576 | 0.2436 |
| MG63 | 0.0716 | -0.0139 | -0.0552 | 0.0836 | -0.0098 | -0.1015 | 0.0042 | 0.0741 | -0.1076 | -0.0741 | 0.0591 |
| SKHEP1 | 0.0938 | 0.1787 | 0.0696 | 0.0098 | 0.1788 | -0.0926 | -0.0442 | -0.1201 | -0.1554 | 0.0838 | 0.0348 |
| MOLM13 | 0.0458 | 0.1792 | 0.1954 | 0.1527 | 0.034 | -0.0241 | 0.1938 | 0.1226 | 0.0897 | -0.0525 | 0.0984 |
| SKMM2 | -0.1101 | 0.0703 | -0.1295 | 0.1497 | 0.0889 | -0.1312 | -0.0123 | 0.099 | -0.0815 | -0.0373 | 0.2014 |
| U2OS | -0.0918 | 0.2143 | -0.2113 | -0.04 | 0.0486 | 0.0148 | 0.0321 | -0.1281 | -0.2872 | 0.1067 | 0.1184 |
| SUDHL4 | 0.2045 | 0.0478 | 0.0612 | 0.1475 | -0.0982 | -0.1777 | -0.0779 | 0.0053 | -0.0577 | -0.0659 | 0.2831 |
| SKNDZ | 0.0482 | 0.1348 | 0.0746 | 0.1573 | -0.0695 | -0.1785 | 0.0473 | 0.0997 | -0.0579 | -0.1273 | 0.3009 |
| NCIH226 | -0.0141 | -0.121 | -0.1297 | 0.0832 | -0.1373 | -0.0559 | 0.0794 | 0.0809 | 0.0323 | 0.096 | 0.066 |
| SNU1105 | 0.0053 | -0.0551 | -0.0517 | 0.0878 | -0.1779 | -0.0648 | 0.004 | 0.0288 | -0.0172 | 0.184 | 0.0876 |
| SNU626 | -0.1062 | 0.069 | -0.0618 | 0.0324 | -0.0345 | -0.1651 | 0.0198 | -0.0141 | -0.1902 | -0.117 | -0.033 |
| RL | 0.0602 | 0.1488 | 0.0578 | 0.1146 | -0.0209 | -0.3103 | 0.1311 | 0.0032 | -0.0944 | -0.0476 | 0.1049 |
| HCC1143 | 0.0789 | 0.1265 | 0.1457 | 0.1012 | -0.0522 | -0.251 | -0.0101 | -0.0033 | -0.0901 | -0.0289 | 0.0773 |
| G402 | 0.2926 | -0.1063 | -0.018 | 0.0859 | -0.0813 | -0.0698 | -0.051 | 0.0577 | 0.0575 | 0.0684 | 0.0414 |
| SF295 | -0.1158 | 0.0866 | -0.0847 | 0.1465 | -0.227 | -0.2013 | 0.1094 | 0.0417 | -0.131 | -0.0368 | 0.1608 |
| SNU478 | 0.146 | -0.0922 | 0.1001 | 0.0856 | -0.0843 | -0.1823 | 0.0399 | 0.0756 | -0.056 | -0.055 | 0.1107 |
| NCIH647 | -0.1607 | 0.1071 | -0.2371 | -0.0732 | 0.1282 | -0.4086 | 0.1814 | -0.1033 | -0.1878 | 0.0405 | -0.015 |
| T84 | -0.1051 | 0.0423 | -0.1462 | 0.2101 | 0.0153 | -0.1955 | 0.0523 | 0.104 | -0.1116 | -0.1645 | 0.1849 |
| OE33 | -0.1141 | 0.064 | -0.0624 | 0.0009 | -0.2062 | -0.1767 | 0.107 | -0.0759 | -0.193 | -0.1216 | 0.0094 |
| SKRC20 | -0.155 | 0.0295 | 0.0743 | -0.0016 | 0.0343 | -0.2279 | 0.1111 | -0.0674 | -0.2894 | -0.1437 | 0.0833 |
| TF1 | 0.0378 | -0.0478 | -0.0767 | 0.2152 | -0.1322 | -0.3829 | 0.0812 | 0.1019 | -0.0673 | -0.1314 | 0.2822 |
| H4 | 0.0525 | 0.2021 | -0.0676 | 0.1476 | -0.0652 | -0.3732 | 0.1282 | 0.1687 | 0.0627 | 0.1266 | 0.2423 |
| LUDLU1 | -0.0482 | 0.1376 | -0.012 | 0.0293 | 0.1431 | -0.1863 | 0.0134 | -0.0169 | -0.2265 | -0.0972 | 0.0645 |
| MHHES1 | -0.0043 | 0.1464 | -0.1008 | 0.2004 | 0.0104 | -0.1259 | 0.0837 | 0.1951 | 0.0486 | -0.0721 | 0.1274 |
| HLF | 0.111 | 0.0084 | -0.0939 | 0.1127 | -0.1763 | -0.1073 | 0.048 | -0.0022 | -0.0707 | 0.098 | 0.1398 |
| NCIH520 | 0.003 | 0.1098 | 0.0249 | 0.1976 | 0.0355 | -0.2535 | 0.1111 | 0.2004 | 0.0666 | -0.0596 | 0.1945 |
| J82 | 0.0233 | -0.0067 | -0.0497 | -0.0904 | -0.002 | -0.0095 | -0.0385 | -0.1711 | -0.2236 | 0.0142 | 0.1304 |
| TEN | -0.045 | 0.1354 | 0.0437 | -0.0721 | 0.1835 | -0.2938 | -0.0011 | -0.1761 | -0.3651 | -0.1353 | 0.0484 |
| RI1 | 0.195 | -0.0545 | 0.0638 | 0.248 | -0.0847 | -0.2702 | 0.0376 | 0.1571 | 0.0905 | -0.1242 | 0.2358 |
| COLO800 | -0.0987 | 0.0874 | 0.0384 | -0.0527 | 0.2208 | -0.0935 | 0.0183 | 0.0272 | -0.0776 | -0.2301 | -0.0131 |
| BL70 | -0.0708 | -0.0619 | 0.028 | 0.0758 | -0.0739 | -0.0463 | 0.1719 | 0.0395 | -0.1286 | -0.1622 | 0.4035 |
| NCIH747 | 0.1214 | 0.0231 | 0.0549 | 0.0329 | 0.0874 | -0.3262 | -0.0348 | 0.0283 | -0.132 | -0.1046 | 0.2 |
| K029AX | -0.2366 | 0.106 | -0.1648 | -0.0707 | 0.1803 | -0.0316 | 0.2403 | -0.2256 | -0.2991 | -0.0514 | -0.0413 |
| MEC1 | 0.0435 | 0.0562 | 0.2759 | 0.0774 | 0.0012 | -0.1924 | -0.0125 | -0.0186 | -0.1883 | -0.2786 | 0.2122 |
| U937 | -0.0435 | 0.048 | -0.1204 | -0.0047 | 0.0727 | -0.1781 | 0.0602 | -0.0795 | -0.2662 | -0.1851 | 0.1736 |
| SNU685 | -0.0659 | 0.0618 | -0.1489 | 0.1724 | -0.0544 | -0.1063 | -0.0005 | -0.0395 | -0.1612 | -0.0442 | 0.1507 |
| TE5 | -0.0493 | 0.0477 | 0.0538 | 0.0005 | 0.0825 | -0.2424 | 0.0903 | -0.0068 | -0.1853 | -0.1214 | 0.0573 |
| SAOS2 | 0.0391 | 0.0318 | 0.0042 | -0.0279 | 0.0059 | -0.1806 | -0.0606 | -0.1272 | -0.1988 | -0.0686 | 0.1067 |
| 769P | -0.0417 | 0.0955 | 0.0773 | -0.0992 | 0.0538 | -0.2767 | 0.0044 | -0.1348 | -0.3694 | -0.0938 | 0.1024 |
| NCIH1944 | 0.0028 | 0.1218 | -0.0446 | 0.0537 | 0.0844 | -0.3145 | 0.101 | -0.0112 | -0.16 | -0.1063 | 0.0934 |
| BICR6 | 0.0748 | 0.1678 | 0.0933 | -0.0683 | 0.1444 | -0.2622 | -0.0032 | -0.1193 | -0.2393 | 0.0467 | 0.1968 |
| NCIH838 | -0.032 | 0.0867 | -0.1316 | 0.0398 | -0.0193 | -0.1182 | 0.1127 | 0.0381 | -0.1343 | 0.0465 | 0.1643 |

| R | EC | IDRE | TM | IDRC | Linker | Tail | IDRE pos | IDRC pos | RK cluste | GxxxG | CRAC |
|---|---|---|---|---|---|---|---|---|---|---|---|
| SNU449 | -0.0011 | 0.0414 | 0.0253 | -0.0478 | 0.0115 | -0.1451 | 0.0384 | -0.139 | -0.2589 | -0.139 | 0.0863 |
| SW837 | -0.1178 | 0.0803 | 0.0512 | -0.0631 | 0.1508 | -0.1223 | -0.0072 | -0.0956 | -0.2685 | -0.1831 | 0.0608 |
| SNU475 | 0.1349 | 0.0657 | -0.018 | 0.2257 | -0.141 | -0.3539 | 0.0567 | 0.1245 | -0.0215 | 0.0807 | 0.2586 |
| TC71 | -0.1213 | 0.1464 | -0.1365 | 0.1068 | 0.0058 | -0.0554 | 0.1672 | 0.0397 | -0.0643 | -0.0317 | 0.0536 |
| UACC62 | 0.0055 | 0.0874 | 0.1927 | 0.1508 | -0.0906 | -0.341 | 0.0378 | 0.0197 | -0.1168 | -0.0183 | 0.252 |
| KMS20 | -0.1313 | 0.136 | -0.0787 | 0.1115 | 0.1765 | -0.208 | 0.2429 | 0.0721 | -0.1513 | -0.2133 | 0.2527 |
| NCIN87 | -0.152 | 0.1489 | -0.0568 | -0.1197 | 0.1757 | -0.1839 | 0.1481 | -0.2334 | -0.4352 | -0.0711 | 0.0506 |
| TYKNU | 0.0198 | 0.013 | 0.0215 | 0.2018 | -0.1208 | -0.1935 | 0.0101 | 0.2715 | 0.1769 | -0.1631 | 0.243 |
| NCIH1694 | 0.0083 | 0.1765 | -0.0885 | -0.0817 | 0.2273 | -0.3562 | 0.0572 | -0.0439 | -0.2158 | -0.0046 | 0.1152 |
| CAKI1 | 0.0394 | -0.029 | 0.048 | -0.0306 | -0.0714 | -0.1886 | 0.0598 | -0.1124 | -0.248 | -0.2029 | 0.0865 |
| NCIH1915 | 0.0989 | 0.0159 | 0.1061 | 0.1021 | -0.0192 | -0.1978 | -0.0743 | 0.0856 | 0.0504 | -0.0533 | 0.228 |
| EFE184 | 0.0104 | 0.1106 | -0.022 | -0.0617 | 0.1638 | -0.4281 | 0.2479 | -0.0914 | -0.2575 | -0.1468 | 0.1299 |
| OCIMY7 | 0.0141 | 0.0183 | 0.1314 | 0.0458 | -0.048 | -0.1526 | 0.2136 | -0.0016 | -0.0829 | -0.2351 | -0.0086 |
| SW1088 | -0.05 | 0.2 | 0.0092 | 0.0494 | -0.1107 | -0.1622 | 0.0796 | 0.0595 | -0.0721 | 0.1143 | 0.0896 |
| LU65 | 0.016 | 0.0396 | 0.1102 | 0.089 | 0.0819 | -0.0849 | -0.0983 | 0.0201 | -0.0293 | -0.1535 | 0.1322 |
| SH4 | 0.0577 | 0.0161 | 0.1949 | 0.1596 | -0.0261 | -0.1806 | -0.006 | 0.1141 | 0.0128 | -0.0333 | 0.0231 |
| RERFLCSQ1 | -0.0175 | 0.2116 | 0.0339 | -0.0917 | 0.2275 | -0.2768 | 0.1284 | -0.1262 | -0.3635 | -0.074 | 0.0798 |
| LU99 | 0.0937 | 0.1039 | 0.0424 | 0.0687 | -0.0281 | -0.2293 | 0.1729 | -0.0784 | -0.227 | 0.0133 | 0.1644 |
| KNS60 | 0.0999 | 0.0023 | 0.0343 | -0.1045 | -0.1008 | -0.1858 | 0.013 | -0.307 | -0.2864 | 0.0873 | 0.1648 |
| NCIH1666 | -0.0485 | 0.0695 | 0.1615 | -0.0092 | 0.1105 | -0.1167 | -0.0097 | -0.0478 | -0.2063 | -0.115 | 0.0224 |
| MELHO | 0.1839 | 0.0026 | 0.0704 | 0.0346 | 0.1429 | -0.3259 | -0.0424 | -0.0821 | -0.0997 | -0.1055 | 0.2467 |
| TE8 | 0.0186 | 0.1531 | -0.0515 | -0.0343 | 0.1249 | -0.1761 | 0.1133 | -0.0597 | -0.2859 | -0.0456 | 0.0028 |
| HCC95 | -0.0563 | 0.1037 | 0.0257 | -0.0308 | 0.1392 | -0.1688 | 0.0407 | -0.0345 | -0.2032 | 0.0308 | 0.1366 |
| LN428 | 0.1181 | -0.0043 | 0.0446 | 0.0951 | 0.0385 | -0.2325 | -0.0466 | 0.1214 | -0.0237 | -0.0568 | 0.1137 |
| BCPAP | 0.0633 | 0.1708 | 0.2163 | 0.0082 | 0.0027 | -0.1949 | 0.0151 | -0.1113 | -0.1093 | -0.1378 | 0.1668 |
| CJM | 0.0338 | 0.0601 | -0.0158 | 0.0554 | -0.0192 | -0.2755 | 0.2954 | 0.0457 | -0.1186 | 0.0119 | 0.0587 |
| TUHR10TKB | -0.035 | -0.0161 | 0.0212 | -0.0181 | -0.0348 | -0.0831 | -0.1071 | -0.0241 | -0.1365 | 0.0912 | 0.2774 |
| SNU8 | -0.0656 | 0.1922 | 0.0849 | -0.1774 | 0.3125 | -0.2661 | 0.0262 | -0.2415 | -0.4409 | 0.0254 | -0.0157 |
| SNU1196 | -0.1561 | 0.327 | -0.1331 | 0.1168 | 0.1033 | -0.1896 | 0.2366 | 0.0197 | -0.1518 | 0.0238 | 0.0618 |
| NCIH460 | 0.0546 | -0.0352 | 0.1263 | -0.0257 | -0.007 | -0.2513 | -0.0133 | 0.1001 | -0.0468 | -0.1374 | 0.1957 |
| CAS1 | 0.0197 | 0.0932 | -0.035 | 0.0319 | 0.005 | -0.2007 | 0.0132 | -0.1 | -0.1914 | 0.0855 | 0.2516 |
| SNU216 | -0.0487 | 0.0276 | -0.02 | -0.1434 | 0.1882 | -0.3127 | 0.0486 | -0.2485 | -0.4372 | -0.1597 | 0.0483 |
| HCC56 | -0.2359 | 0.135 | -0.0176 | 0.1283 | -0.0493 | -0.0189 | -0.0256 | 0.1294 | -0.0988 | -0.0902 | 0.1497 |
| PK45H | -0.0424 | 0.1465 | 0.0542 | -0.0295 | 0.0792 | -0.2334 | -0.0506 | -0.169 | -0.1737 | -0.0099 | 0.3361 |
| YH13 | 0.0011 | 0.051 | -0.0698 | 0.2128 | -0.3345 | -0.1004 | 0.0342 | 0.0948 | 0.0554 | 0.0693 | 0.1397 |
| SW1463 | 0.18 | 0.0524 | 0.1151 | 0.0501 | -0.0721 | -0.4322 | 0.048 | 0.0265 | -0.0552 | -0.1794 | 0.2642 |
| LI7 | 0.013 | 0.0007 | -0.0487 | 0.0916 | -0.164 | 0.0301 | 0.0423 | 0.0404 | -0.0393 | 0.096 | -0.0042 |
| HSC2 | -0.0194 | 0.1065 | 0.1347 | -0.0474 | 0.157 | -0.2213 | 0.0368 | -0.1317 | -0.3093 | -0.116 | 0.0538 |
| RT112 | 0.0157 | -0.0188 | -0.0802 | 0.0998 | -0.028 | -0.0561 | 0.1205 | 0.1373 | -0.009 | -0.067 | 0.297 |
| HUH1 | -0.0222 | 0.0788 | 0.1508 | 0.0378 | 0.0216 | -0.2617 | 0.099 | -0.0201 | -0.152 | -0.1768 | 0.0598 |
| JHH4 | 0.0654 | 0.1222 | 0.0112 | -0.0799 | 0.1836 | -0.2442 | 0.0574 | -0.2471 | -0.3616 | -0.0148 | 0.1435 |
| MALME3M | -0.1026 | 0.0794 | 0.0618 | 0.0869 | 0.0275 | -0.3038 | 0.1592 | -0.0196 | -0.0651 | -0.0656 | 0.0822 |
| SNU387 | -0.0956 | 0.114 | -0.2187 | 0.0292 | 0.0133 | 0.0131 | 0.0771 | -0.0157 | -0.196 | 0.0113 | 0.0597 |
| KNS81 | 0.0224 | 0.0839 | 0.1419 | -0.0423 | 0.0701 | -0.094 | 0.0507 | -0.0543 | -0.0877 | 0.0306 | 0.0862 |
| HUH7 | 0.0428 | -0.0354 | 0.3139 | -0.1441 | -0.0242 | -0.0873 | -0.0018 | 0.0183 | -0.0638 | -0.0597 | 0.2248 |
| NCIH2170 | -0.01 | 0.0718 | 0.0183 | -0.1165 | 0.1897 | -0.2317 | -0.0105 | -0.1352 | -0.2762 | -0.1514 | 0.0854 |
| SNU182 | -0.0497 | 0.0657 | -0.0421 | 0.1066 | 0.1164 | -0.4363 | 0.1056 | 0.0515 | -0.1625 | 0.041 | 0.2036 |
| VMRCRCW | -0.1195 | 0.028 | 0.0222 | 0.1665 | 0.0009 | -0.1808 | 0.1759 | 0.0823 | -0.1577 | -0.3937 | 0.0917 |
| GSU | -0.0625 | 0.1662 | 0.1003 | 0.013 | 0.1875 | -0.1568 | -0.0246 | -0.054 | -0.1891 | -0.1226 | 0.1427 |
| KU1919 | 0.1107 | -0.0235 | 0.1356 | 0.1655 | -0.0946 | -0.2895 | 0.0125 | -0.0565 | -0.0601 | -0.2424 | 0.2127 |
| F36P | 0.113 | -0.003 | 0.1334 | -0.2721 | 0.0702 | 0.0187 | 0.096 | -0.3581 | -0.2694 | -0.1371 | 0.1086 |
| TE11 | -0.005 | 0.1417 | 0.0733 | -0.0167 | 0.1411 | -0.3352 | 0.1402 | -0.1714 | -0.3695 | -0.0581 | 0.1664 |
| SW1116 | 0.0283 | 0.0355 | -0.0341 | -0.0931 | 0.1442 | -0.2698 | -0.0449 | -0.1617 | -0.3679 | -0.1082 | 0.0401 |
| SF767 | 0.0443 | 0.2049 | -0.0148 | -0.2113 | 0.2852 | -0.3645 | 0.1488 | -0.1899 | -0.4172 | 0.0862 | 0.1102 |
| NCIH716 | 0.1173 | 0.0412 | 0.1945 | -0.056 | -0.0493 | 0.0587 | -0.1777 | -0.0233 | -0.0276 | -0.0159 | 0.1311 |
| SNU423 | 0.049 | 0.0966 | 0.0097 | -0.081 | 0.0524 | -0.0449 | 0.0284 | -0.1704 | -0.1801 | -0.0013 | 0.0574 |
| TUHR4TKB | 0.1564 | 0.0543 | -0.1987 | 0.0234 | 0.1469 | -0.3356 | 0.0501 | 0.0533 | -0.2236 | -0.0125 | 0.098 |
| NCIH1792 | -0.0769 | 0.1316 | -0.0538 | 0.0577 | 0.1188 | -0.0658 | 0.0548 | 0.0707 | -0.1063 | -0.0238 | -0.0045 |
| EW8 | -0.2798 | 0.0858 | -0.0165 | 0.0503 | 0.0494 | -0.0448 | 0.1869 | -0.0353 | -0.3241 | -0.1704 | -0.0123 |
| SNU46 | 0.0708 | 0.0922 | -0.189 | 0.1313 | 0.0214 | -0.3304 | 0.2343 | 0.1277 | -0.0865 | 0.1008 | 0.4071 |
| LS123 | 0.0295 | -0.0497 | 0.0252 | -0.2558 | 0.0835 | -0.0334 | -0.1355 | -0.3411 | -0.3721 | 0.0048 | -0.1138 |
| TCCPAN2 | -0.0082 | 0.1673 | -0.0949 | 0.1882 | 0.0272 | -0.2397 | 0.2287 | 0.1536 | 0.0188 | -0.0092 | 0.1154 |
| BICR16 | -0.093 | 0.0669 | 0.0221 | -0.0069 | 0.0746 | -0.2633 | 0.1812 | -0.158 | -0.3092 | -0.108 | 0.0489 |
| SNB75 | -0.0757 | 0.0644 | -0.0889 | -0.1269 | 0.1844 | -0.0991 | -0.1262 | -0.3002 | -0.4155 | -0.0729 | 0.2621 |
| RKN | -0.009 | 0.0321 | -0.1365 | -0.0209 | 0.0543 | -0.2584 | 0.0849 | -0.0076 | -0.1512 | -0.1091 | 0.2135 |
| KE39 | -0.1512 | 0.0774 | -0.0863 | 0.0523 | 0.1103 | -0.136 | 0.1025 | 0.0039 | -0.2024 | -0.0881 | -0.0393 |
| NCIH1299 | -0.0311 | -0.0381 | -0.0899 | 0.0299 | -0.0302 | -0.2043 | 0.0055 | -0.0705 | -0.2013 | -0.2418 | 0.2402 |
| CALU1 | 0.0128 | 0.2514 | 0.0237 | -0.0846 | -0.0407 | -0.2116 | 0.2232 | -0.225 | -0.3159 | -0.0363 | 0.2308 |
| INA6 | -0.1168 | 0.0933 | -0.213 | 0.3028 | -0.1603 | 0.0339 | 0.1411 | 0.2986 | 0.0555 | 0.0379 | 0.2637 |
| NCIH1092 | -0.027 | 0.1768 | -0.0544 | 0.0518 | 0.0179 | -0.2207 | 0.1657 | -0.0459 | -0.1998 | -0.1576 | 0.2566 |
| CAL78 | -0.0701 | -0.0269 | 0.0108 | -0.0829 | 0.0157 | -0.1277 | 0.0849 | -0.206 | -0.3421 | -0.1296 | 0.0824 |
| SNU410 | -0.0149 | 0.0694 | -0.0882 | -0.0579 | 0.0326 | -0.2199 | 0.137 | -0.0832 | -0.231 | -0.0472 | 0.1301 |
| CAL33 | 0.0003 | 0.165 | -0.002 | -0.0953 | 0.2223 | -0.3087 | 0.1711 | -0.175 | -0.4298 | -0.1078 | 0.1191 |
| 59M | -0.1097 | 0.0846 | -0.1335 | -0.0258 | 0.0365 | -0.0306 | 0.1227 | -0.0308 | -0.2558 | -0.0714 | 0.0319 |
| NCIH2030 | 0.1184 | 0.1228 | 0.0998 | 0.0127 | 0.1189 | -0.3062 | 0.0148 | -0.0242 | -0.2484 | -0.0137 | 0.099 |
| UMUC3 | 0.0922 | 0.1074 | 0.0991 | 0.06 | -0.1113 | -0.165 | -0.014 | -0.0893 | -0.1872 | 0.0579 | 0.3047 |
| KURAMOCHI | -0.0188 | 0.0513 | -0.0745 | -0.0587 | 0.0926 | -0.5073 | 0.0848 | -0.1216 | -0.3415 | -0.187 | 0.2054 |
| NCIH2171 | 0.0589 | 0.0548 | 0.1518 | 0.0082 | 0.0221 | -0.244 | -0.0448 | -0.0012 | -0.1425 | -0.2236 | 0.2819 |
| OVISE | 0.0598 | 0.0889 | 0.0599 | 0.0371 | 0.0601 | -0.3371 | 0.0824 | -0.0529 | -0.2116 | -0.1992 | 0.0957 |
| ABC1 | 0.0085 | 0.0621 | -0.1894 | 0.0324 | 0.0177 | -0.1864 | 0.0588 | 0.0838 | -0.0751 | 0.0401 | 0.1643 |
| SNU61 | -0.0323 | 0.1759 | 0.0693 | 0.015 | 0.1798 | -0.1687 | 0.0422 | 0.0113 | -0.2057 | -0.0794 | -0.0024 |
| NCIH2004RT | 0.0154 | 0.0775 | 0.2545 | 0.1907 | -0.0693 | -0.0767 | -0.0885 | 0.1228 | 0.1128 | 0.181 | 0.2764 |
| BXPC3 | -0.0952 | 0.1286 | -0.0582 | -0.0982 | 0.2145 | -0.1644 | 0.056 | -0.1466 | -0.2883 | -0.056 | 0.0886 |
| SNU761 | -0.0432 | 0.0784 | 0.1745 | -0.0747 | 0.1043 | -0.1347 | -0.0523 | -0.1327 | -0.3067 | -0.0843 | 0.124 |
| KMS34 | -0.0572 | 0.0414 | -0.0782 | 0.0282 | -0.0335 | -0.0728 | 0.1197 | -0.0738 | -0.2269 | -0.0842 | 0.3179 |
| HEYA8 | 0.105 | -0.147 | 0.1518 | 0.0649 | -0.0956 | -0.0796 | -0.0849 | 0.0365 | -0.0443 | -0.1944 | 0.2817 |
| OE21 | 0.0382 | 0.1963 | 0.2189 | -0.1113 | 0.1265 | -0.2127 | 0.119 | -0.2626 | -0.3714 | 0.0352 | 0.0353 |
| VMCUB1 | 0.003 | 0.0509 | -0.1058 | -0.1276 | 0.1867 | -0.2497 | 0.0731 | -0.0711 | -0.3267 | -0.1566 | -0.0084 |
| HSC4 | 0.032 | -0.1035 | 0.1021 | -0.0312 | 0.0169 | -0.1817 | -0.022 | -0.04 | -0.2221 | -0.1739 | 0.0458 |
| HT1197 | 0.1729 | 0.0373 | -0.1725 | 0.0763 | -0.0544 | -0.3606 | 0.1558 | 0.008 | -0.1864 | -0.0641 | 0.3035 |
| BHY | -0.0406 | 0.0756 | 0.1121 | -0.0533 | 0.1626 | -0.1507 | 0.0084 | -0.1122 | -0.3035 | -0.1071 | -0.0156 |
| SNU1076 | 0.0425 | 0.0472 | 0.0874 | -0.0764 | 0.0908 | -0.2358 | 0.0262 | -0.0803 | -0.2451 | 0.0118 | 0.008 |
| IGR39 | -0.0153 | 0.0524 | 0.0112 | -0.0656 | 0.1495 | -0.1853 | -0.0472 | -0.0375 | -0.2393 | -0.0854 | 0.1567 |
| K562 | -0.0276 | 0.1292 | 0.1497 | 0.2253 | -0.0157 | -0.1106 | 0.1048 | 0.1121 | -0.0622 | -0.1611 | 0.2003 |
| HT29 | -0.1633 | 0.1987 | -0.1475 | -0.0121 | 0.1719 | -0.1434 | 0.1184 | -0.0467 | -0.2574 | -0.0536 | 0.0271 |
| UACC893 | 0.0109 | 0.1097 | 0.0106 | -0.1697 | 0.1929 | -0.1128 | -0.0094 | -0.2049 | -0.3163 | -0.0172 | -0.0761 |
| SIHA | 0.1442 | -0.0319 | -0.0307 | 0.0649 | -0.0797 | -0.2054 | 0.0723 | -0.0074 | -0.1037 | 0.0251 | 0.13 |
| AML193 | -0.0145 | 0.066 | -0.0468 | 0.0827 | -0.1253 | 0.0304 | 0.0176 | 0.0002 | -0.1452 | -0.0833 | 0.1487 |
| A172 | 0.0946 | 0.0007 | 0.0113 | 0.0526 | -0.1109 | -0.0944 | 0.0053 | -0.0591 | -0.0724 | 0.1225 | 0.0682 |
| NCIH1836 | -0.0789 | 0.0224 | 0.1557 | 0.0398 | 0.065 | -0.1284 | -0.1366 | -0.0409 | -0.1565 | -0.1433 | -0.0802 |
| TDOTT | -0.0409 | 0.0968 | -0.0226 | -0.0719 | 0.1134 | -0.2072 | 0.1113 | -0.1042 | -0.2343 | -0.0275 | -0.0351 |
| HCC78 | 0.0484 | 0.121 | 0.009 | -0.0131 | 0.1359 | -0.2533 | -0.0202 | -0.0434 | -0.2778 | -0.2434 | 0.1617 |
| EBC1 | -0.0588 | 0.006 | -0.1167 | 0.1861 | -0.3455 | -0.2224 | 0.214 | 0.1222 | 0.0597 | -0.0916 | 0.3157 |
| RCM1 | -0.0526 | 0.1465 | 0.0355 | 0.0382 | 0.1847 | -0.2058 | 0.1362 | 0.0774 | -0.1544 | -0.1566 | 0.0082 |
| ST486 | 0.2434 | 0.0067 | 0.0668 | 0.0842 | 0.0318 | -0.2437 | -0.0342 | 0.0335 | -0.0064 | -0.185 | 0.181 |
| YKG1 | -0.0296 | 0.1488 | 0.019 | -0.0275 | -0.1861 | -0.0463 | 0.1079 | -0.0694 | -0.1579 | 0.0777 | 0.1045 |
| T98G | 0.041 | 0.1113 | -0.123 | 0.1388 | -0.0106 | -0.018 | -0.0233 | 0.0631 | -0.0624 | 0.0188 | 0.1733 |
| MDAMB436 | 0.0486 | 0.0107 | -0.0533 | 0.2152 | -0.1187 | -0.1577 | -0.0055 | 0.0951 | -0.0338 | -0.1985 | 0.0988 |
| KMS27 | -0.1388 | 0.1761 | -0.0235 | 0.1181 | 0.0356 | -0.1698 | 0.2971 | 0.0251 | -0.1605 | 0.0217 | 0.1822 |
| JHH2 | 0.0468 | -0.0577 | -0.1341 | 0.0015 | -0.0825 | 0.0544 | 0.0396 | -0.007 | -0.1781 | 0.0254 | 0.1285 |
| UACC257 | 0.1668 | 0.095 | 0.0138 | 0.0099 | 0.0224 | -0.2178 | -0.0739 | -0.0318 | -0.0399 | 0.038 | -0.0822 |
| C32 | 0.1385 | 0.1203 | 0.0992 | -0.0692 | 0.1792 | -0.4153 | 0.1658 | 0.0932 | 0.0397 | 0.0705 | -0.1687 |
| SNU16 | 0.0363 | 0.0433 | -0.0622 | -0.18 | 0.0838 | 0.0215 | -0.0925 | -0.1283 | -0.2538 | 0.1484 | 0.2489 |
| JHOS4 | 0.0662 | 0.1371 | 0.078 | 0.0278 | 0.0099 | -0.3027 | 0.0221 | 0.0068 | -0.0647 | 0.0027 | 0.1556 |
| EPLC272H | 0.1008 | 0.0193 | 0.0658 | -0.1113 | 0.0993 | -0.3472 | 0.0188 | -0.0734 | -0.2392 | 0.0357 | 0.2744 |
| NCIH1975 | -0.062 | 0.0908 | 0.1408 | -0.0957 | 0.1925 | -0.1281 | -0.0142 | -0.1669 | -0.3717 | -0.139 | -0.0264 |
| KMS26 | -0.1793 | 0.197 | -0.0808 | 0.0772 | 0.0679 | -0.0949 | 0.2562 | -0.0279 | -0.1944 | -0.1285 | 0.112 |
| NCIH1437 | -0.1725 | 0.0836 | -0.0805 | 0.1763 | 0.0081 | -0.1314 | 0.084 | 0.0954 | -0.0426 | -0.1086 | 0.1736 |
| LN235 | 0.1719 | 0.078 | -0.0404 | 0.0192 | -0.0076 | -0.3492 | 0.0022 | 0.0434 | -0.0466 | 0.0972 | 0.2841 |
| TM31 | -0.0171 | 0.0372 | -0.142 | 0.0422 | -0.104 | -0.0811 | -0.0174 | -0.0973 | -0.1865 | 0.0016 | 0.1463 |
| BC3C | 0.0142 | 0.1053 | 0.1202 | 0.0436 | 0.0586 | -0.218 | -0.0686 | -0.0112 | -0.1615 | -0.0014 | 0.2825 |
| LN229 | -0.0027 | 0.1293 | -0.0113 | -0.146 | 0.2173 | -0.2276 | 0.0491 | -0.2255 | -0.3829 | -0.106 | 0.0052 |
| LCLC97TM1 | -0.0975 | 0.0447 | -0.0355 | 0.0128 | 0.093 | -0.2025 | 0.0793 | -0.0822 | -0.209 | -0.1119 | 0.1197 |
| TTC709 | 0.0674 | -0.0514 | 0.1397 | -0.0596 | 0.1456 | -0.2485 | -0.053 | -0.0298 | -0.2679 | -0.0735 | 0.2824 |
| PATU8902 | 0.0255 | 0.0085 | -0.044 | 0.0796 | 0.0347 | -0.1532 | 0.0464 | 0.1051 | -0.0486 | -0.2161 | 0.125 |
| SLR26 | -0.0529 | 0.0298 | -0.0158 | 0.0508 | -0.2189 | -0.1012 | -0.0749 | -0.0065 | -0.079 | 0.0409 | 0.0677 |
| MIAPACA2 | -0.0371 | 0.0994 | 0.0334 | 0.0374 | 0.0254 | -0.1616 | 0.0964 | 0.0291 | -0.1047 | -0.0938 | 0.1441 |
| M07E | -0.0687 | -0.0245 | 0.2021 | 0.1075 | 0.0115 | -0.0965 | -0.0214 | -0.0602 | -0.1504 | -0.3091 | 0.2134 |

| R | EC | IDRE | TM | IDRC | Linker | Tail | IDRE pos | IDRC pos | RK cluste | GxxxG | CRAC |
|---|---|---|---|---|---|---|---|---|---|---|---|
| TE6 | -0.0588 | 0.2101 | 0.067 | -0.1256 | 0.2502 | -0.3402 | 0.1763 | -0.2328 | -0.4473 | -0.0906 | 0.1077 |
| PECAPJ34CLONEC12 | 0.0353 | 0.0572 | 0.1592 | -0.0789 | 0.1118 | -0.2204 | 0.0757 | -0.1308 | -0.2966 | -0.167 | 0.0178 |
| KYM1 | -0.074 | 0.1221 | -0.0418 | 0.1941 | -0.1911 | -0.1536 | 0.101 | 0.0289 | -0.042 | 0.0739 | 0.2031 |
| COV644 | -0.0609 | 0.0411 | 0.1949 | -0.1067 | 0.1668 | -0.1334 | -0.0034 | -0.1376 | -0.3285 | -0.1854 | 0.0246 |
| SF126 | -0.0211 | 0.08 | -0.1036 | 0.0724 | -0.0734 | -0.1109 | 0.0667 | 0.0871 | -0.0706 | 0.1274 | 0.0759 |
| RVH421 | 0.0465 | -0.09 | -0.0571 | 0.0606 | 0.0253 | -0.3748 | 0.1788 | 0.06 | -0.1314 | -0.1723 | 0.1695 |
| HS746T | 0.0257 | 0.0649 | 0.1255 | -0.0124 | 0.0559 | -0.258 | 0.0583 | 0.0826 | -0.0377 | -0.0854 | 0.162 |
| SNU1041 | -0.0301 | 0.0034 | 0.0867 | -0.0562 | 0.1177 | -0.2621 | 0.1458 | -0.1547 | -0.2637 | -0.1417 | 0.2864 |
| PECAPJ15 | -0.0093 | 0.1524 | 0.0039 | -0.0323 | 0.1644 | -0.3602 | 0.1296 | -0.0825 | -0.3413 | 0.0068 | 0.1412 |
| JHH1 | -0.01 | 0.1161 | 0.1477 | -0.086 | 0.1153 | -0.119 | 0.037 | -0.1402 | -0.3322 | -0.1212 | 0.0664 |
| MDAMB157 | 0.0904 | 0.0305 | -0.0312 | 0.1398 | -0.0599 | -0.2359 | 0.0524 | 0.0948 | -0.0567 | -0.0163 | 0.2489 |
| KNS42 | -0.1026 | 0.1208 | -0.0376 | 0.1372 | 0.022 | -0.1991 | 0.0865 | 0.0329 | -0.136 | -0.0735 | 0.3396 |
| SNU201 | 0.0276 | 0.1402 | 0.064 | 0.1378 | -0.0375 | -0.1624 | -0.0117 | 0.1356 | 0.0006 | -0.0448 | 0.2008 |
| HCC1806 | 0.111 | 0.0674 | 0.0968 | -0.0819 | 0.1206 | -0.4119 | 0.0216 | -0.1051 | -0.214 | 0.0003 | 0.1835 |
| LCLC103H | 0.0401 | 0.0394 | 0.0248 | -0.0858 | 0.1151 | -0.1682 | 0.0456 | -0.1518 | -0.2416 | -0.0207 | 0.0833 |
| YD8 | 0.0181 | 0.0846 | 0.0238 | -0.1426 | 0.1252 | -0.1257 | 0.0099 | -0.21 | -0.3506 | 0.0318 | -0.0737 |
| HS944T | -0.0549 | 0.1395 | -0.0571 | 0.0337 | 0.0563 | -0.066 | 0.0794 | -0.0902 | -0.1618 | -0.0762 | 0.2196 |
| FU97 | 0.1654 | 0.0491 | 0.2447 | 0.2022 | -0.1006 | -0.1916 | -0.0973 | 0.1633 | 0.1721 | 0.0585 | 0.4101 |
| LN340 | 0.1463 | 0.1908 | -0.1451 | 0.0293 | -0.0707 | -0.2646 | 0.1621 | 0.0367 | -0.0116 | -0.0851 | 0.0245 |
| KYSE520 | 0.0796 | -0.0501 | 0.1245 | 0.0313 | -0.0817 | -0.1647 | -0.0404 | 0.0598 | -0.0938 | -0.0271 | 0.2379 |
| NCIH441 | -0.0442 | 0.1068 | 0.0923 | 0.0365 | 0.1459 | -0.1682 | 0.0273 | -0.0106 | -0.0638 | -0.0588 | 0.0257 |
| NCIH211 | 0.0761 | 0.0965 | -0.0657 | 0.0853 | -0.0366 | -0.3514 | 0.1496 | 0.0762 | -0.127 | -0.0429 | 0.191 |
| JL1 | 0.0083 | 0.1454 | 0.2068 | 0.0811 | 0.0074 | -0.1106 | 0.059 | -0.042 | -0.183 | 0.0678 | 0.0141 |
| OVMANA | -0.0064 | 0.189 | 0.0331 | -0.0498 | 0.2135 | -0.4202 | 0.0993 | -0.0818 | -0.3155 | -0.1075 | 0.0811 |
| TE1 | -0.0089 | 0.0788 | 0.0232 | 0.0827 | 0.0661 | -0.2152 | 0.0742 | 0.0301 | -0.0898 | -0.1056 | 0.1303 |
| NCIH28 | -0.2211 | 0.0672 | 0.0175 | -0.0528 | 0.1533 | -0.208 | 0.1384 | -0.0369 | -0.2492 | -0.0675 | 0.0476 |
| 786O | -0.0104 | 0.0167 | 0.1579 | -0.046 | 0.0117 | -0.1829 | -0.1485 | -0.0825 | -0.173 | -0.0802 | 0.2444 |
| SW620 | -0.0006 | 0.0119 | 0.0265 | 0.0473 | -0.0323 | -0.1253 | -0.009 | -0.0029 | -0.0316 | -0.2017 | 0.3372 |
| SUIT2 | -0.1992 | 0.156 | -0.0079 | 0.0086 | 0.0909 | -0.0288 | 0.0376 | -0.0611 | -0.2966 | -0.0148 | 0.1814 |
| JJN3 | 0.113 | -0.0787 | 0.1671 | 0.0255 | 0.0164 | -0.2627 | 0.1115 | 0.007 | -0.0983 | -0.1171 | 0.092 |
| RAJI | 0.0293 | 0.1155 | 0.0141 | 0.2218 | -0.0273 | -0.2586 | 0.1135 | 0.2962 | 0.1331 | -0.0362 | -0.0036 |
| SF268 | 0.0616 | -0.0987 | 0.0067 | -0.0754 | -0.069 | -0.1887 | 0.1121 | -0.1098 | -0.2742 | -0.2225 | 0.0693 |
| SUDHL8 | 0.2357 | -0.0004 | -0.0531 | -0.0318 | -0.0311 | -0.3304 | 0.0241 | 0.1562 | 0.0404 | -0.0443 | 0.2356 |
| A2780 | 0.3325 | -0.1118 | 0.0483 | 0.1127 | -0.2302 | -0.312 | 0.0054 | 0.0605 | 0.0331 | -0.0542 | 0.2312 |
| KMS18 | 0.0238 | -0.0117 | -0.0801 | 0.0224 | -0.0094 | 0.0495 | 0.1628 | -0.0151 | -0.0941 | 0.0374 | 0.1851 |
| SUDHL5 | -0.0176 | 0.0153 | -0.0329 | 0.0436 | -0.0322 | -0.0748 | -0.0125 | 0.1288 | 0.0144 | -0.0565 | 0.4058 |
| WM1799 | -0.0322 | 0.1726 | 0.0252 | -0.1105 | 0.1658 | -0.3542 | 0.0823 | -0.1373 | -0.3373 | -0.0903 | 0.0394 |
| CORL23 | 0.1282 | 0.0685 | 0.024 | 0.4064 | -0.0989 | -0.1607 | 0.109 | 0.2662 | 0.2524 | -0.142 | 0.1198 |
| OVTOKO | -0.053 | 0.0265 | 0.0856 | -0.1431 | 0.117 | -0.201 | -0.0898 | -0.1536 | -0.2898 | -0.2324 | 0.1647 |
| SUDHL1 | 0.0365 | 0.1647 | 0.0701 | 0.1085 | 0.0834 | -0.1871 | 0.0727 | -0.0056 | -0.1307 | -0.2271 | 0.1816 |
| SKMES1 | -0.0419 | 0.1449 | 0.0793 | -0.0088 | 0.1963 | -0.2374 | 0.0579 | -0.0696 | -0.2738 | -0.1201 | 0.1324 |
| NCIH1355 | 0.0509 | 0.0059 | -0.0501 | -0.0347 | 0.0252 | -0.0101 | 0.0014 | -0.1383 | -0.2006 | -0.0273 | 0.1371 |
| HCC44 | 0.074 | 0.1571 | -0.0835 | -0.0726 | 0.0767 | -0.0317 | -0.1824 | -0.1622 | -0.1919 | -0.0214 | 0.1553 |
| HCC70 | 0.1849 | 0.0835 | 0.1337 | -0.0094 | 0.0472 | -0.3579 | 0.05 | 0.087 | -0.0168 | 0.0843 | 0.3473 |
| HUH6 | -0.1104 | 0.1667 | 0.0294 | 0.0985 | 0.0391 | 0.0589 | 0.0767 | 0.03 | -0.0538 | 0.0213 | 0.0161 |
| LN443 | 0.1228 | 0.0273 | 0.0326 | 0.1862 | -0.1466 | -0.1751 | -0.0356 | 0.1118 | 0.037 | 0.1101 | 0.1999 |
| LN464 | -0.0473 | -0.0268 | 0.0829 | 0.1268 | -0.0291 | -0.0635 | -0.1455 | 0.0662 | -0.0847 | -0.2933 | 0.0451 |
| SW1573 | -0.0733 | 0.1323 | 0.0291 | -0.0472 | 0.2467 | -0.2051 | 0.1423 | -0.0541 | -0.1472 | -0.0262 | 0.0225 |
| SW948 | 0.1029 | -0.1714 | -0.2275 | 0.0354 | -0.1408 | -0.2483 | -0.0074 | -0.0479 | -0.1526 | -0.2084 | 0.3326 |
| A549 | -0.0021 | 0.0592 | 0.0232 | -0.0361 | 0.1221 | -0.2345 | 0.0623 | -0.144 | -0.3103 | -0.1181 | 0.0272 |
| SNU1066 | 0.095 | 0.1508 | 0.096 | -0.06 | 0.1594 | -0.3445 | 0.069 | -0.1553 | -0.3241 | -0.0094 | 0.1583 |
| SNU503 | 0.0474 | -0.0746 | 0.1026 | -0.0653 | -0.0038 | -0.1774 | -0.0457 | -0.071 | -0.2436 | -0.1933 | 0.0822 |
| KMRC1 | -0.0628 | 0.0094 | 0.2036 | -0.0282 | -0.0412 | 0.1143 | -0.243 | 0.0038 | -0.0565 | 0.1995 | 0.1284 |
| L33 | -0.0417 | 0.0838 | -0.1261 | -0.0274 | 0.1607 | -0.286 | 0.1344 | -0.0483 | -0.2858 | -0.1117 | 0.1629 |
| OV7 | -0.0385 | 0.005 | -0.0064 | 0.1019 | -0.0716 | -0.2397 | 0.034 | 0.0027 | -0.1705 | -0.051 | 0.2445 |
| KYSE180 | 0.0551 | 0.0609 | 0.1188 | -0.1248 | 0.206 | -0.3551 | 0.0932 | -0.202 | -0.3455 | -0.087 | 0.1219 |
| TE9 | 0.009 | 0.0845 | 0.0505 | -0.0367 | 0.1241 | -0.2899 | -0.0231 | -0.1218 | -0.2396 | -0.0938 | 0.1439 |
| CORL47 | -0.1557 | 0.0439 | -0.1352 | 0.0687 | 0.0935 | -0.2722 | 0.2792 | 0.0783 | -0.1974 | -0.1502 | -0.0746 |
| OVCAR8 | -0.0259 | 0.0401 | 0.1682 | -0.1128 | 0.0688 | -0.2522 | 0.0033 | -0.1134 | -0.2691 | -0.1015 | 0.1067 |
| A3KAW | -0.1694 | -0.0224 | -0.0807 | 0.0474 | 0.0161 | -0.0356 | 0.0748 | 0.1027 | -0.0569 | -0.0472 | 0.1821 |
| DMS53 | 0.2039 | -0.194 | 0.1407 | 0.0229 | -0.1125 | -0.1199 | -0.0593 | 0.0154 | -0.082 | -0.081 | 0.1496 |
| HCC1395 | 0.0172 | -0.0366 | 0.1471 | 0.0452 | -0.0704 | -0.2541 | -0.0053 | 0.1368 | -0.0202 | -0.0928 | 0.2077 |
| NCIH2882 | -0.1846 | 0.0754 | -0.2169 | 0.0759 | 0.0156 | -0.3615 | 0.1622 | -0.097 | -0.272 | 0.0006 | 0.212 |
| RMUGS | -0.0435 | 0.031 | 0.0653 | -0.0771 | 0.0162 | -0.2209 | 0.0752 | -0.0928 | -0.18 | -0.1944 | 0.0934 |
| L1236 | 0.1089 | 0.0654 | -0.1578 | 0.1358 | -0.0317 | -0.2565 | -0.0693 | 0.2026 | 0.0632 | -0.0375 | 0.1192 |
| OAW42 | 0.0044 | 0.134 | -0.0282 | 0.0404 | 0.1011 | -0.414 | 0.0669 | 0.1128 | -0.0441 | -0.0972 | 0.2736 |
| EKVX | -0.0833 | 0.1147 | 0.0248 | -0.0044 | 0.04 | -0.1618 | 0.1434 | -0.0955 | -0.3099 | -0.2254 | 0.075 |
| KMRC2 | -0.0511 | 0.0901 | 0.1115 | -0.0223 | 0.1553 | -0.2429 | 0.1371 | -0.0722 | -0.1646 | -0.1506 | 0.0572 |
| JIMT1 | 0.0751 | 0.0665 | 0.096 | -0.0357 | 0.0863 | -0.2712 | -0.0236 | -0.1246 | -0.2681 | -0.1817 | 0.1521 |
| CAOV3 | 0.1523 | -0.0893 | 0.2047 | -0.0649 | 0.0472 | -0.0829 | -0.0696 | -0.02 | -0.0278 | -0.011 | 0.2404 |
| KMS11 | 0.008 | 0.0403 | -0.1443 | 0.0892 | -0.0309 | -0.0078 | 0.2249 | 0.061 | -0.0112 | 0.0811 | 0.1569 |
| TT2609C02 | 0.077 | 0.1488 | 0.0112 | -0.0223 | 0.1682 | -0.2034 | -0.0912 | -0.0671 | -0.2126 | -0.1194 | -0.0097 |
| COLO680N | -0.1236 | -0.0175 | 0.1246 | -0.0533 | 0.0377 | -0.1094 | -0.057 | 0.022 | -0.128 | -0.0641 | 0.2041 |
| NCIH2291 | 0.0655 | -0.0044 | 0.3023 | -0.1214 | 0.0361 | -0.297 | 0.1135 | -0.2271 | -0.2156 | 0.075 | 0.0995 |
| RMGI | 0.0197 | 0.05 | 0.0699 | -0.0224 | 0.126 | -0.2691 | 0.0417 | -0.0979 | -0.3183 | -0.1923 | 0.1098 |
| TCCSUP | 0.1458 | -0.0154 | -0.1071 | 0.0798 | -0.1157 | -0.1113 | 0.0935 | 0.0556 | -0.0634 | 0.0852 | 0.0969 |
| HMC18 | -0.0613 | 0.0475 | 0.2157 | 0.1271 | 0.0105 | -0.021 | -0.138 | 0.0982 | 0.0552 | -0.018 | 0.1341 |
| SNUC1 | -0.073 | 0.0622 | -0.1719 | -0.0473 | 0.1849 | -0.05 | 0.0761 | 0.0753 | -0.059 | -0.0313 | 0.0411 |
| HT1376 | -0.1576 | 0.1752 | -0.0682 | -0.077 | 0.288 | -0.2029 | 0.1216 | -0.0841 | -0.3293 | -0.0332 | 0.1283 |
| HCC202 | 0.0212 | 0.0511 | 0.0107 | -0.062 | 0.1021 | -0.2515 | -0.0732 | -0.0638 | -0.1583 | 0.0385 | 0.1499 |
| PECAPJ41CLONED2 | -0.0016 | 0.1093 | 0.1573 | -0.0217 | 0.1398 | -0.1648 | 0.0482 | -0.0711 | -0.1441 | -0.0077 | 0.1861 |
| JHH5 | -0.1125 | 0.1355 | 0.0367 | -0.0638 | 0.0375 | -0.1287 | 0.0812 | -0.0916 | -0.2517 | -0.0876 | 0.0725 |
| PECAPJ49 | -0.0537 | 0.0726 | 0.0923 | -0.1019 | 0.1424 | -0.2015 | 0.0701 | -0.1985 | -0.3383 | -0.1255 | 0.0333 |
| SNU601 | -0.0754 | 0.2235 | -0.0003 | 0.0911 | 0.0838 | -0.2491 | 0.2808 | 0.0876 | 0.016 | 0.0059 | 0.0281 |
| GB1 | -0.0415 | 0.1087 | -0.0596 | 0.1636 | -0.151 | -0.1342 | 0.0185 | 0.0451 | -0.0561 | -0.0686 | 0.2948 |
| HEPG2 | -0.0862 | 0.1542 | -0.1027 | 0.0889 | -0.0493 | -0.1544 | 0.0519 | 0.0261 | -0.0968 | 0.0382 | 0.1187 |
| A253 | 0.0443 | 0.0332 | 0.1236 | -0.0791 | 0.1135 | -0.2938 | 0.0209 | -0.1769 | -0.3759 | -0.0919 | 0.0864 |
| UBLC1 | 0.0407 | -0.0559 | 0.0777 | 0.0114 | 0.0148 | -0.1835 | -0.0665 | -0.077 | -0.2183 | -0.1248 | 0.1093 |
| GSS | -0.1711 | 0.0396 | 0.2408 | -0.2145 | 0.2848 | -0.0007 | -0.155 | -0.1047 | -0.1771 | -0.055 | 0.0686 |
| NCIH1703 | -0.038 | 0.0508 | 0.1882 | -0.0164 | 0.0679 | -0.0661 | -0.0685 | -0.0199 | -0.0836 | 0.0935 | 0.1089 |
| SJSA1 | -0.0356 | 0.0653 | -0.1428 | 0.04 | 0.0787 | -0.2629 | 0.2016 | 0.0219 | -0.161 | -0.0576 | 0.2012 |
| JMSU1 | 0.1276 | 0.0049 | -0.0608 | 0.0401 | -0.1041 | -0.0263 | -0.0707 | -0.0397 | -0.1177 | 0.0974 | -0.0061 |
| L428 | 0.0759 | 0.0913 | 0.1251 | 0.0713 | 0.1 | -0.1302 | 0.0226 | 0.1151 | 0.0465 | -0.307 | 0.0242 |
| GI1 | -0.0134 | 0.0558 | 0.0196 | 0.169 | -0.2522 | -0.0798 | 0.0742 | 0.0342 | 0.0074 | -0.1014 | 0.1903 |
| A427 | 0.1513 | -0.0305 | 0.0876 | 0.0999 | -0.0937 | -0.1105 | -0.1165 | 0.0474 | 0.0468 | -0.0801 | 0.0159 |
| MKN74 | -0.0665 | 0.021 | 0.1486 | -0.0376 | 0.0786 | -0.1971 | 0.0683 | -0.1147 | -0.2222 | -0.1868 | 0.0382 |
| LNZ308 | -0.0189 | 0.1481 | 0.0651 | 0.0572 | 0.0814 | -0.0991 | -0.1326 | 0.0381 | -0.0964 | 0.0318 | 0.1931 |
| YD38 | -0.0126 | 0.0984 | 0.0354 | 0.0119 | 0.1135 | -0.0291 | -0.0829 | -0.0962 | -0.1511 | -0.0256 | 0.1602 |
| MM1S | -0.1619 | 0.1913 | -0.0385 | 0.102 | 0.081 | -0.118 | 0.0477 | 0.0504 | -0.1165 | 0.0011 | 0.2168 |
| SH10TC | -0.0232 | 0.0495 | -0.1076 | -0.0166 | 0.1522 | -0.3526 | 0.0254 | -0.0008 | -0.2416 | -0.1975 | 0.221 |
| WM983B | -0.1438 | 0.107 | -0.0348 | 0.274 | -0.072 | 0.0155 | 0.1258 | 0.1611 | -0.041 | -0.167 | 0.2939 |
| NCIH1648 | 0.0654 | 0.0224 | 0.19 | -0.0182 | 0.0307 | 0.0822 | -0.0467 | -0.0321 | -0.1124 | -0.1524 | 0.0379 |
| NCIH526 | -0.0455 | 0.051 | -0.1137 | 0.1119 | -0.0055 | -0.127 | 0.0048 | 0.123 | 0.0308 | 0.0352 | 0.2358 |
| MDAMB231 | -0.0304 | -0.0366 | 0.2632 | -0.0689 | -0.0417 | 0.0156 | -0.1143 | -0.0313 | -0.0806 | 0.0009 | 0.1653 |
| LK2 | 0.1017 | 0.0093 | -0.1338 | 0.1536 | -0.1235 | -0.1542 | 0.101 | 0.1342 | 0.0235 | 0.0721 | 0.0579 |
| P31FUJ | -0.012 | 0.1222 | 0.099 | 0.0798 | -0.0003 | -0.1256 | 0.1165 | -0.035 | -0.0891 | -0.2924 | 0.2056 |
| BICR56 | -0.0372 | 0.1373 | 0.0183 | -0.1504 | 0.2057 | -0.3166 | 0.0921 | -0.208 | -0.3971 | 0.0048 | -0.0302 |
| KIJK | -0.1444 | 0.0754 | 0.1085 | 0.2578 | -0.0577 | -0.2167 | 0.1548 | 0.1458 | -0.0117 | -0.2542 | 0.1308 |
| RERFLCAD2 | -0.0042 | -0.0355 | 0.023 | -0.0196 | -0.0152 | -0.116 | -0.113 | 0.0248 | 0.0579 | 0.0111 | 0.1507 |
| NCIH727 | 0.0283 | 0.1512 | 0.0913 | 0.0215 | 0.0707 | -0.2866 | 0.0407 | -0.0301 | -0.1592 | -0.179 | 0.194 |
| ONS76 | 0.1056 | 0.0283 | -0.0676 | -0.009 | 0.0355 | -0.125 | 0.037 | -0.0127 | -0.0687 | -0.1064 | 0.1998 |
| KYSE30 | 0.0333 | 0.0474 | 0.0145 | -0.0402 | 0.1188 | -0.2091 | -0.0263 | -0.2176 | -0.4039 | -0.1024 | 0.0975 |
| HSC3 | -0.0616 | 0.0324 | 0.1569 | -0.0445 | 0.0964 | -0.1326 | -0.0703 | -0.0839 | -0.2343 | -0.19 | 0.1921 |
| NCIH2023 | -0.0313 | 0.2125 | -0.0049 | 0.1632 | 0.0598 | -0.1425 | 0.0777 | 0.0737 | -0.0708 | -0.1121 | 0.1353 |
| SEM | -0.0547 | 0.0233 | 0.1026 | 0.1564 | -0.066 | -0.0766 | 0.1193 | 0.1014 | -0.0046 | -0.2331 | 0.2985 |
| CAMA1 | 0.1632 | -0.0142 | 0.1971 | 0.1321 | -0.1495 | -0.2231 | -0.0539 | -0.0113 | -0.0775 | -0.222 | 0.2958 |
| KYSE70 | 0.0561 | -0.0301 | -0.081 | 0.1052 | -0.013 | -0.146 | -0.0659 | 0.0787 | -0.0945 | -0.1696 | 0.317 |
| NCIH2126 | -0.1824 | 0.0252 | 0.051 | 0.0053 | 0.1503 | -0.0383 | 0.0017 | -0.0448 | -0.1812 | -0.1712 | -0.0156 |
| LXF289 | -0.0026 | 0.0769 | -0.117 | -0.0212 | 0.1008 | -0.1965 | 0.0237 | -0.1432 | -0.2856 | 0.0217 | 0.1326 |
| A2058 | 0.0136 | 0.1219 | 0.1015 | -0.0412 | 0.1009 | -0.2523 | -0.0423 | -0.1245 | -0.2205 | -0.0516 | 0.1711 |
| SHP77 | -0.0204 | -0.001 | 0.1298 | -0.0333 | -0.1065 | -0.0269 | 0.0157 | 0.0362 | -0.0422 | 0.0342 | 0.2159 |
| RERFLCAD1 | 0.0016 | 0.1245 | -0.0667 | -0.0051 | 0.0251 | -0.0961 | 0.0155 | -0.0503 | -0.1212 | -0.0639 | 0.19 |
| BFTC909 | 0.039 | 0.0265 | -0.0234 | -0.0754 | 0.0738 | -0.1306 | 0.0643 | -0.058 | -0.2322 | 0.0122 | -0.015 |
| KATOIII | 0.0987 | 0.057 | 0.0573 | -0.3209 | 0.1385 | 0.0191 | -0.1622 | -0.2921 | -0.3811 | 0.0865 | 0.1588 |
| BICR22 | 0.0412 | 0.0595 | 0.1104 | -0.1225 | 0.1733 | -0.1878 | -0.0008 | -0.2025 | -0.3444 | -0.0214 | 0.0712 |
| MCAS | 0.014 | 0.0758 | 0.2398 | -0.1936 | 0.1443 | -0.1806 | 0.0117 | -0.3207 | -0.4195 | -0.1951 | 0.1011 |
| HS695T | 0.1003 | 0.0803 | 0.2112 | -0.1416 | 0.1263 | -0.5234 | 0.1701 | -0.0733 | -0.1576 | 0.0495 | 0.1657 |
| NCIH446 | -0.0032 | -0.183 | 0.0511 | 0.1518 | -0.0693 | -0.0701 | 0.1023 | 0.2602 | 0.0896 | -0.1727 | 0.1953 |

| R | EC | IDRE | TM | IDRC | Linker | Tail | IDRE pos | IDRC pos | RK cluste | GxxxG | CRAC |
|---|---|---|---|---|---|---|---|---|---|---|---|
| BFTC905 | -0.0188 | 0.2322 | -0.0456 | 0.1827 | 0.0012 | -0.1055 | 0.1113 | 0.0484 | -0.0165 | -0.0454 | 0.1913 |
| NB1 | -0.1286 | -0.0036 | -0.1281 | 0.1079 | 0.0201 | -0.1815 | 0.1908 | 0.132 | 0.0479 | -0.1619 | 0.1826 |
| COLO679 | -0.1002 | 0.1857 | -0.0577 | -0.0386 | 0.0582 | -0.2674 | 0.1962 | -0.1569 | -0.3525 | 0.0184 | 0.2171 |
| SNU738 | 0.0945 | -0.0411 | -0.029 | 0.0084 | 0.0066 | -0.0643 | -0.0709 | -0.1373 | -0.1871 | -0.011 | 0.1314 |
| KYSE410 | -0.0425 | 0.1362 | -0.0147 | -0.2045 | 0.2476 | -0.2115 | 0.0366 | -0.2764 | -0.4261 | -0.1045 | 0.0088 |
| SKMEL30 | 0.0886 | 0.0197 | 0.0272 | 0.2961 | -0.1403 | -0.165 | 0.1416 | 0.0442 | 0.0466 | -0.338 | -0.0428 |
| SKOV3 | -0.0411 | -0.0261 | -0.0512 | -0.187 | 0.1441 | -0.1963 | -0.0694 | -0.1546 | -0.3367 | -0.1209 | 0.0526 |
| RPMI8226 | -0.1719 | 0.146 | 0.0152 | -0.0235 | 0.0259 | 0.1761 | -0.0117 | -0.0324 | -0.2167 | -0.0996 | 0.0175 |
| LN18 | -0.0234 | 0.1709 | -0.0296 | 0.1199 | -0.157 | -0.2597 | 0.212 | 0.0232 | -0.1446 | -0.0886 | 0.3124 |
| SW403 | -0.0119 | -0.0465 | 0.0191 | 0.0058 | -0.0017 | -0.1504 | -0.0285 | 0.0236 | -0.0737 | -0.3221 | 0.0397 |
| EJM | -0.1578 | -0.0407 | 0.0526 | -0.0111 | -0.0264 | -0.1762 | 0.0006 | 0.0012 | -0.091 | -0.0726 | 0.1711 |
| SKMEL24 | -0.0355 | 0.0233 | -0.0265 | 0.0329 | 0.1107 | -0.1974 | 0.1757 | -0.001 | -0.2921 | -0.2935 | 0.0237 |
| KYSE140 | 0.0001 | 0.1668 | 0.1419 | -0.0884 | 0.1718 | -0.326 | 0.0585 | -0.1535 | -0.3376 | 0.0237 | 0.1314 |
| KYSE510 | -0.0638 | 0.1601 | 0.0234 | -0.0111 | 0.1403 | -0.2163 | 0.164 | -0.0873 | -0.2594 | -0.0824 | 0.0117 |
| WM793 | 0.1082 | -0.049 | 0.0949 | -0.0947 | 0.0121 | -0.2244 | -0.0204 | -0.0573 | -0.1403 | -0.1983 | 0.1787 |
| HUNS1 | 0.1442 | 0.0873 | 0.0673 | 0.0533 | 0.0129 | -0.1478 | 0.1259 | -0.0152 | -0.1734 | -0.1734 | 0.3399 |
| HEC50B | 0.0431 | 0.0805 | 0.2123 | 0.008 | -0.029 | -0.2487 | 0.053 | 0.0491 | -0.0663 | -0.0687 | 0.2217 |
| CAL27 | 0.0352 | 0.1445 | 0.1217 | -0.183 | 0.2122 | -0.2094 | 0.0502 | -0.2625 | -0.3974 | -0.0282 | 0.0484 |
| RH30 | -0.1241 | 0.1286 | 0.1652 | 0.1244 | 0.0247 | -0.1404 | 0.082 | 0.0937 | 0.0294 | -0.0343 | -0.0714 |
| UMUC1 | 0.0562 | 0.1781 | -0.1817 | -0.0162 | 0.0375 | -0.2077 | 0.0974 | -0.023 | -0.1573 | 0.0678 | 0.2432 |
| GCT | 0.036 | 0.0319 | 0.1472 | -0.0572 | 0.1121 | -0.3233 | 0.0407 | -0.0802 | -0.128 | -0.0769 | 0.0782 |
| YD15 | 0.0283 | 0.1961 | 0.1853 | -0.014 | 0.0906 | -0.2206 | -0.0265 | -0.069 | -0.2134 | -0.0414 | 0.0156 |
| NCIH322 | -0.1815 | 0.112 | 0.0727 | 0.0257 | 0.1076 | -0.2894 | 0.166 | -0.0278 | -0.2185 | -0.1224 | 0.0488 |
| AMO1 | -0.1448 | 0.0974 | -0.1559 | 0.0472 | 0.1717 | -0.1199 | 0.1496 | -0.019 | -0.2165 | -0.1419 | 0.1124 |
| SCABER | -0.0293 | 0.1583 | 0.0237 | -0.0846 | 0.2429 | -0.2666 | 0.0232 | -0.1451 | -0.3405 | -0.07 | 0.1133 |
| HCC366 | 0.0688 | -0.1318 | 0.0394 | 0.0441 | -0.0225 | -0.0628 | -0.1006 | -0.0549 | -0.1735 | -0.1401 | 0.2734 |
| NCIH2087 | -0.0021 | 0.0925 | -0.0074 | 0.0938 | 0.0925 | -0.0762 | 0.0153 | 0.0298 | -0.0912 | -0.1174 | 0.1688 |
| HARA | -0.0372 | 0.1458 | 0.0308 | -0.0649 | 0.1472 | -0.3218 | 0.1803 | -0.1899 | -0.3455 | -0.0385 | 0.0785 |
| NCIH1373 | 0.0252 | -0.0582 | -0.0422 | 0.1328 | -0.0484 | -0.1363 | -0.0574 | 0.0873 | 0.0053 | -0.0537 | 0.2196 |
| FADU | 0.2134 | -0.0405 | -0.0022 | 0.0228 | -0.0298 | -0.3298 | -0.024 | 0.0432 | -0.0842 | -0.0806 | 0.0414 |
| HGC27 | -0.056 | 0.0967 | 0.0202 | 0.0699 | -0.0106 | -0.2112 | 0.0842 | -0.0445 | -0.1905 | -0.0274 | 0.3754 |
| JHH7 | 0.1144 | 0.1155 | 0.1289 | 0.015 | -0.0337 | -0.1986 | 0.1958 | -0.0017 | 0.0346 | 0.0361 | 0.0882 |
| MDAMB468 | 0.0397 | 0.0772 | 0.2612 | 0.1588 | -0.1348 | -0.0745 | 0.1276 | 0.1365 | 0.0744 | 0.0505 | 0.1429 |
| MORCPR | 0.0616 | 0.1089 | -0.2698 | -0.0121 | -0.0926 | -0.2313 | 0.1675 | -0.0406 | -0.1279 | -0.1621 | 0.0944 |
| NCIH661 | -0.0039 | 0.1797 | 0.0754 | -0.087 | 0.0827 | -0.3339 | 0.169 | -0.2659 | -0.3385 | 0.0112 | 0.0601 |
| OCIMY5 | -0.0083 | 0.0582 | -0.1473 | -0.0606 | -0.0537 | -0.0024 | 0.0449 | -0.0538 | -0.2187 | -0.1444 | 0.1751 |
| KYSE150 | -0.1548 | 0.0557 | 0.1341 | 0.1663 | 0.0579 | -0.1687 | 0.0792 | 0.1569 | -0.041 | -0.1486 | 0.1522 |
| CAL51 | 0.1502 | 0.1806 | 0.0087 | 0.0154 | -0.016 | -0.3896 | 0.1424 | 0.0234 | -0.181 | -0.0489 | 0.3271 |
| KNS62 | 0.0641 | -0.0378 | 0.0286 | 0.1709 | 0.0189 | -0.2433 | 0.1182 | -0.0018 | -0.0658 | -0.1487 | 0.1623 |
| HCC1954 | 0.0663 | 0.0958 | 0.3835 | -0.0294 | 0.0421 | -0.0479 | -0.0309 | 0.0247 | -0.0496 | -0.006 | 0.2426 |
| NCIH358 | 0.1227 | 0.1157 | 0.0717 | 0.0608 | 0.1194 | -0.3644 | 0.0992 | 0.0122 | -0.1432 | -0.1515 | 0.2006 |
| HOP62 | 0.1521 | -0.0355 | -0.0381 | -0.0568 | 0.1429 | -0.3506 | 0.1502 | -0.0519 | -0.1587 | -0.0425 | 0.0377 |
| KMBC2 | -0.0455 | 0.1557 | 0.2177 | -0.0431 | 0.1525 | -0.1778 | 0.0375 | -0.0648 | -0.2477 | -0.0205 | 0.0383 |
| DBTRG05MG | 0.0455 | -0.1068 | 0.1034 | -0.0239 | -0.1615 | -0.1605 | 0.11 | -0.038 | -0.0233 | 0.0033 | 0.1151 |
| COLO684 | -0.048 | -0.0669 | -0.0685 | -0.0911 | 0.0222 | -0.2199 | 0.0869 | 0.07 | -0.0339 | -0.0856 | -0.0784 |
| KYSE450 | 0.1006 | 0.1015 | 0.0934 | -0.0541 | 0.1084 | -0.3272 | 0.1261 | -0.192 | -0.3776 | -0.0716 | 0.0924 |
| NCIH1048 | 0.1369 | 0.089 | 0.0559 | -0.164 | 0.0862 | -0.1332 | -0.022 | -0.1373 | -0.2143 | 0.1364 | 0.2233 |
| CHAGOK1 | 0.109 | 0.1709 | -0.0314 | 0.1081 | 0.0047 | -0.2645 | 0.1258 | 0.047 | -0.0818 | -0.1113 | 0.1965 |
| NCIH1568 | 0.0061 | -0.0697 | -0.0229 | -0.0221 | -0.0403 | -0.2599 | -0.1147 | -0.0047 | -0.15 | -0.1458 | 0.2517 |
| NCIH510 | 0.0686 | -0.0869 | 0.0156 | 0.1531 | 0.0399 | -0.1628 | -0.0031 | 0.1174 | -0.0296 | -0.1784 | 0.1926 |
| HCC515 | -0.0731 | -0.095 | 0.0728 | 0.1574 | -0.0508 | -0.0571 | -0.0845 | 0.0883 | -0.1325 | -0.255 | 0.2184 |
| KYSE270 | -0.0633 | 0.0678 | 0.2336 | 0.0423 | 0.0339 | -0.0868 | 0.011 | -0.1707 | -0.2853 | -0.1113 | 0.1944 |
| MDAMB415 | 0.1736 | 0.0081 | 0.336 | -0.2007 | 0.017 | 0.0231 | -0.2003 | -0.0616 | -0.109 | 0.1222 | 0.2586 |
| HCC15 | -0.046 | 0.0284 | 0.2022 | 0.1382 | -0.005 | -0.0123 | -0.0121 | 0.0865 | 0.066 | -0.1852 | 0.3321 |
| MFE296 | 0.022 | -0.0363 | 0.0608 | -0.3399 | 0.0524 | 0.071 | -0.1667 | -0.3387 | -0.3689 | -0.0213 | 0.167 |
| AGS | 0.1748 | 0.0247 | 0.0058 | 0.0946 | -0.0963 | -0.0876 | 0.1066 | 0.0293 | -0.0998 | -0.2116 | 0.2036 |
| MELJUSO | -0.1038 | 0.025 | -0.0474 | 0.1275 | -0.0996 | -0.0528 | 0.05 | 0.0785 | -0.0432 | -0.08 | 0.1127 |
| IGR1 | 0.0355 | 0.0451 | -0.0192 | 0.1685 | -0.185 | -0.1438 | 0.0864 | 0.0595 | 0.0182 | -0.0279 | 0.1879 |
| MDAMB435S | -0.0485 | 0.1898 | -0.1501 | 0.1466 | -0.0781 | -0.0755 | 0.2295 | 0.0481 | -0.0892 | -0.1049 | 0.0687 |
| TOV21G | 0.1985 | -0.0301 | 0.0846 | 0.0954 | -0.0973 | -0.4018 | 0.013 | -0.0605 | -0.1161 | -0.2307 | 0.2599 |
| NCIH2009 | 0.1372 | 0.0622 | 0.1828 | 0.0879 | 0.0051 | -0.1717 | -0.0813 | -0.0119 | 0.0067 | -0.2011 | 0.1171 |
| SF172 | -0.054 | 0.1706 | 0.0608 | -0.0774 | 0.158 | -0.1843 | 0.0764 | -0.163 | -0.2952 | -0.0596 | 0.1043 |
| NCIH1793 | 0.0709 | 0.0395 | 0.0578 | 0.0167 | 0.022 | -0.3779 | 0.0999 | -0.0832 | -0.2546 | -0.0055 | 0.1516 |
| KMM1 | -0.033 | 0.0971 | -0.0981 | 0.1091 | -0.042 | -0.1205 | 0.0715 | 0.0166 | -0.1117 | -0.0146 | 0.0299 |
| SW1271 | -0.073 | 0.0147 | 0.1169 | 0.0906 | -0.0146 | -0.079 | -0.0395 | 0.1339 | 0.0163 | 0.0672 | 0.1391 |
| NCIH1869 | -0.0027 | 0.032 | 0.1384 | -0.0959 | 0.1658 | -0.1915 | -0.0931 | -0.0268 | -0.2005 | -0.2103 | 0.2627 |
| 647V | 0.1158 | 0.1015 | 0.1227 | -0.2114 | 0.1612 | -0.2342 | 0.063 | -0.2923 | -0.3078 | 0.1084 | 0.0815 |
| SNU719 | -0.1545 | 0.2192 | 0.0413 | 0.0556 | 0.1888 | -0.095 | 0.144 | 0.0073 | -0.2049 | -0.0186 | -0.022 |
| WM88 | 0.2014 | -0.1257 | -0.0288 | -0.0916 | -0.0209 | -0.1585 | 0.0755 | 0.0275 | 0.0144 | -0.1147 | -0.0365 |
| NCIH23 | 0.0483 | 0.1043 | 0.0427 | 0.2138 | 0.011 | -0.187 | 0.133 | 0.2026 | 0.1803 | -0.0805 | 0.1613 |
| HCC1359 | 0.1072 | -0.0433 | -0.0698 | 0.0841 | -0.1056 | -0.0983 | 0.0409 | 0.0755 | -0.01 | 0.0883 | 0.0705 |
| FTC133 | 0.0023 | 0.1089 | -0.0053 | -0.0013 | 0.0457 | -0.0718 | -0.0919 | -0.0644 | -0.1478 | 0.0137 | 0.1617 |
| 5637 | 0.0683 | 0.0928 | 0.1543 | 0.001 | 0.0717 | -0.1722 | -0.0317 | -0.0581 | -0.2093 | -0.0715 | 0.1623 |
| ES2 | 0.0678 | -0.0439 | -0.0294 | -0.0875 | 0.0068 | -0.2975 | 0.0656 | -0.2131 | -0.2402 | -0.1903 | 0.2281 |
| SNU349 | 0.0658 | -0.0831 | 0.2707 | -0.1793 | -0.0167 | -0.0818 | -0.0958 | -0.1138 | -0.1236 | 0.1031 | 0.1364 |
| MDAMB453 | 0.0286 | 0.0771 | 0.2637 | -0.1048 | 0.1204 | -0.1202 | 0.0149 | -0.1608 | -0.2087 | -0.0802 | 0.0999 |
| NUGC3 | 0.0241 | 0.1187 | 0.1671 | -0.0574 | 0.1521 | -0.2103 | 0.0497 | -0.1397 | -0.2992 | -0.1333 | 0.0462 |
| NCIH2286 | -0.0402 | 0.1138 | -0.0514 | 0.0246 | -0.0853 | -0.1675 | 0.0831 | 0.0091 | -0.1168 | -0.0926 | 0.1316 |
| ESS1 | 0.0455 | 0.0753 | 0.0764 | -0.1349 | -0.015 | -0.0502 | -0.0297 | -0.1571 | -0.2171 | -0.0281 | 0.0998 |
| HT | 0.1489 | -0.0888 | 0.2319 | 0.0235 | -0.0533 | -0.1423 | -0.0175 | 0.0205 | -0.0636 | -0.2936 | 0.2774 |
| IPC298 | -0.3089 | 0.1937 | 0.037 | -0.0678 | 0.0134 | -0.0547 | 0.081 | -0.1983 | -0.3728 | 0.0355 | 0.0904 |
| NCIH1573 | 0.2476 | -0.1805 | -0.0182 | 0.1601 | -0.1212 | -0.2105 | -0.0948 | -0.0309 | 0.0509 | -0.1127 | -0.0318 |
| TE4 | -0.0365 | 0.0956 | -0.0912 | -0.0971 | 0.2015 | -0.2324 | -0.0023 | -0.1309 | -0.2795 | -0.1351 | 0.0481 |
| IM95 | -0.1094 | 0.079 | 0.1745 | -0.1483 | -0.021 | -0.0329 | 0.0996 | -0.2667 | -0.4007 | -0.1498 | -0.1432 |
| CMLT1 | 0.0424 | 0.1169 | -0.1084 | -0.0724 | 0.1433 | -0.284 | 0.1897 | 0.0077 | -0.1522 | 0.002 | 0.1621 |
| NCIH157DM | 0.0104 | -0.0618 | 0.0023 | -0.0311 | -0.0267 | -0.3488 | 0.2126 | 0.0187 | -0.2108 | -0.0989 | 0.1478 |
| RCHACV | 0.0845 | 0.1772 | 0.0532 | 0.1125 | -0.0012 | -0.1812 | -0.0177 | -0.0887 | -0.1997 | -0.1688 | 0.131 |
| NCIH2172 | -0.0297 | 0.1017 | 0.1 | 0.1008 | -0.0007 | -0.0756 | 0.0256 | 0.112 | -0.0192 | -0.1134 | 0.0916 |
| HT55 | -0.0928 | 0.0807 | 0.0091 | -0.0347 | 0.0556 | -0.1285 | 0.0683 | -0.028 | -0.2211 | -0.1171 | 0.106 |
| JHUEM1 | 0.0573 | 0.0821 | 0.1239 | 0.1812 | -0.0483 | -0.4346 | 0.2283 | 0.153 | -0.0407 | -0.0392 | 0.2991 |
| NCIH2110 | -0.0464 | 0.1375 | -0.1316 | 0.1011 | 0.1012 | -0.2625 | 0.1523 | 0.1026 | -0.0745 | -0.0823 | 0.0929 |
| SNU1 | -0.0075 | 0.0787 | 0.1501 | 0.1387 | -0.0379 | -0.107 | -0.0395 | 0.0905 | 0.0565 | 0.049 | 0.2263 |
| MDAMB361 | -0.0472 | 0.1907 | 0.0234 | -0.1806 | 0.256 | -0.1618 | 0.1184 | -0.3021 | -0.3898 | 0.0386 | -0.0811 |
| MDST8 | -0.0571 | 0.0645 | -0.2511 | 0.1448 | -0.0405 | -0.1058 | 0.1652 | 0.1469 | 0.0048 | -0.0089 | 0.1466 |
| EFO27 | -0.052 | 0.0778 | 0.1866 | -0.0179 | 0.0635 | -0.167 | 0.0244 | -0.0663 | -0.2705 | -0.1833 | 0.0497 |
| PF382 | 0.1803 | 0.0517 | 0.0104 | 0.0835 | 0.0265 | -0.2774 | 0.0507 | 0.0715 | -0.0668 | -0.186 | 0.2492 |
| NALM6 | 0.0458 | -0.0034 | -0.0735 | 0.0466 | 0.0764 | -0.2476 | 0.0056 | -0.0328 | -0.2638 | -0.158 | 0.1112 |
| SKUT1 | 0.0963 | 0.0132 | 0.0861 | 0.1337 | -0.0469 | -0.4388 | 0.063 | 0.0034 | -0.1696 | -0.1855 | 0.3847 |
| HEC1B | -0.0479 | 0.043 | -0.0008 | -0.1275 | 0.1315 | -0.1477 | -0.0365 | -0.1501 | -0.3446 | -0.1576 | 0.0701 |
| RKO | -0.0235 | 0.064 | 0.1307 | -0.0024 | 0.0294 | -0.2703 | 0.0863 | -0.0149 | -0.1167 | -0.2047 | 0.0341 |
| NAMALWA | 0.1177 | 0.1437 | 0.2494 | 0.1654 | -0.0336 | -0.1327 | -0.0077 | 0.0598 | -0.0473 | -0.1213 | 0.1857 |
| NCIH650 | -0.1672 | -0.0763 | -0.1281 | 0.1969 | -0.1325 | -0.2005 | 0.0781 | 0.1071 | -0.0715 | -0.2498 | 0.4107 |
| HEC265 | 0.1042 | 0.0175 | 0.2137 | 0.0694 | -0.0453 | -0.3359 | 0.1494 | -0.0187 | -0.1265 | -0.0831 | 0.1202 |
| OVK18 | 0.0775 | -0.0769 | 0.2285 | 0.111 | -0.0945 | -0.1667 | -0.0565 | 0.0564 | -0.03 | -0.3051 | 0.2988 |
| 2313287 | 0.1045 | -0.1591 | 0.286 | 0.009 | -0.0547 | -0.1686 | -0.0532 | 0.0015 | -0.0738 | -0.1157 | 0.1625 |
| LOVO | 0.0008 | 0.0284 | 0.2934 | -0.1576 | 0.0112 | 0.0616 | -0.1016 | -0.1775 | -0.2534 | -0.2204 | 0.0434 |
| SUPT1 | 0.1017 | 0.0099 | 0.121 | 0.0322 | 0.0443 | -0.0727 | -0.1585 | -0.0505 | -0.0819 | -0.0312 | 0.2419 |
| 22RV1 | -0.2239 | -0.0103 | 0.1932 | -0.0033 | 0.0538 | -0.3013 | 0.1867 | 0.0771 | 0.0063 | -0.1594 | 0.0266 |
| LS180 | 0.1823 | -0.0519 | 0.1614 | -0.0172 | -0.023 | -0.1868 | -0.1282 | 0.0112 | -0.1355 | -0.181 | 0.1474 |
| SW48 | -0.0945 | 0.0418 | 0.0924 | 0.1921 | -0.03 | -0.0743 | 0.1458 | 0.1287 | -0.0767 | -0.0898 | 0.0728 |
| SNUC4 | -0.1101 | 0.0897 | 0.1188 | -0.0192 | 0.1903 | -0.3019 | 0.1319 | -0.0889 | -0.3132 | -0.1358 | -0.0693 |
| REH | -0.0081 | 0.2085 | -0.1507 | -0.0011 | 0.1828 | -0.2288 | 0.0764 | -0.0679 | -0.1907 | -0.111 | 0.143 |
| ISHIKAWAHERAKLIO02ER | 0.023 | -0.1065 | -0.0125 | 0.0956 | 0.0605 | -0.4177 | 0.0984 | 0.0832 | -0.079 | 0.0082 | 0.1906 |
| OC314 | 0.0524 | 0.1059 | 0.2408 | 0.0057 | 0.103 | -0.2229 | 0.051 | -0.0593 | -0.1277 | -0.0645 | 0.2751 |
| CCK81 | -0.067 | -0.0167 | 0.0912 | -0.1642 | 0.1495 | -0.1072 | 0.0134 | -0.0055 | -0.1932 | -0.2158 | 0.0718 |
| RL952 | -0.0291 | 0.2179 | 0.0739 | -0.0273 | 0.1257 | -0.2613 | 0.0811 | -0.2747 | -0.389 | -0.0987 | 0.1531 |
| IGROV1 | -0.0013 | -0.0173 | 0.2568 | 0.0698 | 0.0632 | -0.1971 | -0.1247 | -0.0083 | -0.1666 | -0.264 | 0.1388 |
| COLO792 | 0.0773 | 0.0278 | 0.0438 | 0.1675 | -0.0924 | -0.2776 | 0.0789 | 0.0222 | -0.057 | -0.1772 | 0.144 |
| KM12 | -0.0257 | 0.0086 | -0.0375 | 0.1453 | 0.0097 | -0.0945 | 0.0767 | 0.1841 | 0.0891 | -0.1484 | 0.2403 |
| SNUC5 | 0.009 | -0.0364 | 0.1519 | -0.0242 | -0.0705 | -0.1407 | 0.022 | -0.0683 | -0.0573 | -0.1366 | 0.0787 |
| HCT116 | 0.0302 | 0.0936 | -0.0484 | 0.0132 | -0.1279 | -0.1886 | 0.0698 | -0.0214 | -0.0379 | 0.035 | 0.0794 |
| 639V | 0.0518 | 0.1259 | 0.0182 | -0.0038 | -0.0871 | -0.1122 | 0.0807 | -0.1155 | -0.252 | -0.0443 | 0.0577 |
| SNGM | 0.216 | 0.0101 | 0.2229 | 0.0127 | -0.0396 | -0.315 | 0.024 | -0.0602 | -0.1175 | -0.1162 | 0.2931 |
| HCC2450 | 0.0169 | 0.0565 | 0.055 | -0.0362 | 0.0928 | -0.2651 | -0.0201 | -0.1051 | -0.2039 | -0.0723 | 0.1342 |
| HUCCT1 | 0.0367 | -0.032 | 0.2323 | -0.1614 | 0.1345 | -0.0781 | -0.1324 | -0.2243 | -0.3202 | -0.1581 | 0.0376 |
| LNCAPCLONEFGC | 0.1962 | -0.0203 | 0.1133 | 0.0442 | -0.1088 | -0.3193 | 0.03 | 0.0046 | -0.0346 | -0.0324 | 0.0618 |
| EN | -0.0109 | -0.0186 | -0.0164 | 0.0001 | 0.0902 | -0.2999 | 0.1897 | -0.0168 | -0.1868 | -0.2593 | 0.0554 |
| DU145 | -0.0449 | 0.0451 | 0.1856 | 0.0664 | 0.068 | -0.1497 | -0.0723 | 0.0323 | -0.1494 | -0.1635 | 0.1327 |

| R | EC | IDRE | TM | IDRC | Linker | Tail | IDRE pos | IDRC pos | RK cluste | GxxxG | CRAC |
|---|---|---|---|---|---|---|---|---|---|---|---|
| KCL22 | -0.0321 | 0.112 | -0.0353 | 0.0188 | 0.0153 | -0.3417 | 0.2907 | -0.0412 | -0.2468 | -0.0123 | 0.2974 |
| HEC6 | 0.1368 | -0.0567 | 0.0447 | 0.0688 | -0.0147 | -0.2347 | -0.044 | 0.0645 | 0.0599 | 0.0569 | 0.0936 |
| LS411N | 0.182 | -0.1918 | -0.0092 | 0.0867 | -0.0103 | -0.1382 | -0.0215 | 0.2367 | 0.147 | -0.2195 | -0.0057 |
| HT115 | -0.0603 | 0.1267 | -0.0245 | -0.0725 | 0.2707 | -0.1211 | 0.1078 | -0.0771 | -0.2521 | -0.0867 | 0.0366 |
| MFE319 | -0.0662 | 0.0732 | -0.0179 | -0.0272 | 0.1433 | -0.2953 | -0.0013 | -0.1254 | -0.3276 | -0.1866 | 0.1757 |
| SNU81 | 0.1011 | -0.0068 | 0.2114 | -0.013 | 0.0499 | -0.1282 | 0.0091 | -0.0547 | -0.0576 | -0.1182 | 0.0803 |
| JHUEM7 | 0.0549 | -0.0653 | 0.0861 | 0.0491 | -0.1194 | -0.4084 | -0.0061 | -0.1094 | -0.242 | -0.1051 | 0.3973 |
| HEC59 | -0.1753 | 0.0636 | -0.0606 | 0.0481 | -0.0277 | -0.2297 | 0.2523 | 0.0417 | -0.1297 | -0.0388 | 0.2616 |
| JURKAT | 0.028 | 0.168 | -0.1971 | 0.175 | -0.0965 | -0.2655 | 0.113 | 0.03 | -0.1334 | -0.051 | 0.3 |
| HEC251 | 0.1162 | -0.1493 | 0.0519 | -0.0133 | 0.0821 | -0.2984 | 0.1642 | 0.0994 | 0.0223 | -0.2664 | 0.0416 |
| HCT15 | 0.1376 | 0.0431 | 0.0097 | 0.1121 | -0.1833 | -0.1957 | 0.0214 | 0.1097 | 0.0641 | 0.0056 | 0.1186 |
| 143B | -0.0168 | 0.0195 | 0.1094 | 0.0845 | 0.0659 | -0.215 | 0.0304 | 0.0561 | -0.0964 | -0.0372 | 0.2456 |
| BECKER | -0.0875 | 0.0268 | 0.186 | -0.0056 | -0.2236 | 0.0248 | 0.0793 | -0.0516 | 0.0045 | 0.0045 | 0.0284 |
| BT16 | 0.0558 | -0.1017 | -0.0701 | 0.266 | -0.0066 | -0.4046 | 0.1075 | 0.1012 | -0.0661 | -0.3256 | 0.239 |
| CHLA06ATRT | -0.0955 | 0.1644 | 0.0602 | 0.1317 | -0.2379 | -0.0598 | 0.2268 | 0.0071 | -0.062 | -0.0867 | 0.0907 |
| CHLA10 | -0.0586 | 0.2147 | 0.1383 | 0.1294 | 0.123 | -0.1351 | 0.0444 | 0.0949 | -0.0498 | -0.0249 | 0.1627 |
| CHLA266 | 0.0694 | -0.0156 | 0.0371 | -0.0456 | 0.0046 | -0.1113 | -0.0052 | -0.0878 | -0.2164 | -0.0504 | 0.3996 |
| CHLA57 | -0.0152 | -0.0372 | 0.0215 | -0.0735 | 0.0251 | -0.29 | 0.1046 | 0.044 | -0.2386 | -0.056 | 0.3069 |
| CMK115 | 0.1036 | 0.0672 | 0.1715 | 0.2861 | -0.079 | -0.2594 | 0.0572 | 0.1509 | 0.0186 | -0.167 | 0.1542 |
| COGE352 | -0.1195 | 0.068 | 0.1095 | 0.093 | 0.0251 | -0.131 | -0.016 | 0.0141 | -0.2062 | 0.0122 | 0.0968 |
| COLO205 | -0.083 | 0.02 | -0.2056 | 0.0274 | 0.0241 | -0.0525 | 0.097 | 0.0248 | -0.0803 | -0.0678 | -0.0173 |
| CHL1DM | 0.0128 | 0.1972 | -0.0661 | -0.0108 | 0.0964 | -0.1601 | 0.1287 | -0.1067 | -0.2916 | -0.0248 | 0.102 |
| COV504 | -0.0406 | 0.0613 | 0.0427 | 0.0362 | 0.0811 | -0.1043 | -0.07 | 0.0815 | -0.0987 | -0.0103 | 0.0907 |
| CW9019 | 0.01 | -0.0269 | 0.1313 | -0.1658 | 0.0618 | -0.091 | -0.0599 | -0.3024 | -0.338 | -0.0709 | 0.2728 |
| D425 | -0.1507 | -0.0614 | 0.0127 | 0.1618 | 0.0317 | -0.191 | 0.0303 | 0.1362 | -0.0659 | -0.18 | 0.1785 |
| D458 | -0.0068 | -0.0984 | 0.0831 | -0.0263 | -0.0338 | -0.0074 | -0.0343 | 0.0242 | -0.0177 | -0.2654 | 0.0187 |
| DLD1 | -0.0406 | 0.0416 | 0.1072 | 0.0527 | 0.0355 | -0.1526 | 0.0512 | 0.0194 | -0.1065 | -0.2242 | 0.1129 |
| DOV13 | -0.0582 | 0.1307 | 0.0592 | 0.1008 | 0.0757 | -0.2703 | 0.0675 | 0.0271 | -0.1412 | -0.0458 | 0.1104 |
| EVSAT | 0.322 | 0.0364 | 0.2524 | -0.1236 | 0.0016 | -0.1513 | -0.0508 | -0.0867 | -0.1494 | 0.1064 | 0.2141 |
| F5 | 0.0795 | 0.0862 | 0.1716 | -0.036 | -0.0215 | -0.1962 | 0.1352 | -0.0488 | -0.0376 | -0.1749 | 0.1461 |
| NCIH292 | -0.0784 | 0.1439 | 0.0747 | 0.0837 | 0.0073 | -0.1995 | 0.1292 | 0.0223 | -0.1334 | -0.0008 | 0.0451 |
| HCC2998 | -0.0319 | 0.0268 | -0.1662 | 0.178 | -0.0072 | -0.1453 | 0.0951 | 0.1271 | 0.0905 | 0.0811 | 0.1065 |
| JR | -0.1291 | 0.1112 | 0.1758 | 0.0525 | -0.0681 | -0.1717 | 0.1269 | 0.0135 | -0.1754 | -0.061 | 0.1775 |
| KCIMOH1 | -0.1257 | 0.099 | 0.1351 | 0.0695 | 0.0821 | -0.2028 | 0.1422 | -0.0947 | -0.2218 | -0.0525 | 0.0568 |
| KD | 0.0729 | 0.0545 | 0.0925 | 0.2085 | -0.0798 | -0.1065 | -0.0115 | 0.1493 | 0.0362 | -0.0401 | 0.1503 |
| KP1N | -0.0293 | 0.1935 | 0.112 | 0.1377 | 0.0013 | -0.0526 | 0.183 | 0.039 | 0.0463 | 0.015 | -0.0926 |
| MAC2A | -0.0572 | 0.1978 | 0.0274 | -0.0243 | 0.1532 | -0.2365 | 0.1345 | -0.0836 | -0.1736 | -0.1122 | 0.1492 |
| MOGGUVW | -0.0185 | 0.0073 | -0.0716 | -0.0268 | -0.2139 | -0.053 | 0.06 | -0.0556 | -0.0807 | -0.0542 | 0.0335 |
| MON | -0.0383 | 0.1482 | 0.0367 | 0.2422 | -0.1199 | -0.3398 | 0.2111 | 0.2141 | 0.0684 | -0.007 | 0.1858 |
| MONOMAC1 | -0.2429 | 0.2048 | -0.0615 | 0.0046 | 0.04 | -0.1372 | 0.3404 | -0.0533 | -0.2055 | -0.0562 | 0.0906 |
| MYLA | -0.0718 | 0.0505 | -0.0061 | 0.0387 | 0.0442 | -0.2089 | 0.0624 | 0.1038 | -0.0411 | -0.1065 | 0.1621 |
| NCIH1993 | -0.191 | 0.0476 | -0.0209 | 0.0114 | -0.2024 | -0.0683 | 0.1314 | 0.0553 | -0.0921 | -0.0693 | 0.075 |
| OC316 | 0.0421 | 0.0619 | 0.1417 | -0.1205 | 0.1488 | -0.2097 | 0.0497 | -0.1794 | -0.3554 | -0.0024 | 0.0788 |
| OVCAR5 | -0.0965 | 0.0484 | -0.052 | -0.022 | 0.0976 | -0.127 | 0.005 | -0.0694 | -0.2568 | -0.144 | 0.034 |
| CCLFPEDS0001T | 0.0128 | 0.0907 | 0.1904 | 0.0409 | -0.0838 | -0.2717 | -0.0117 | -0.0244 | -0.0863 | -0.1486 | 0.2715 |
| CCLFPEDS0003T | 0.0378 | 0.0728 | 0.0508 | 0.1676 | -0.0982 | -0.3551 | 0.0654 | 0.1111 | 0.0202 | -0.1042 | 0.1478 |
| U251MGDM | -0.0056 | -0.0093 | 0.0658 | -0.0122 | -0.0927 | -0.1421 | 0.0257 | 0.0298 | -0.1261 | -0.1585 | 0.1897 |
| RT11284 | -0.0483 | 0.0716 | -0.1026 | 0.1443 | -0.0053 | -0.0988 | 0.192 | 0.0517 | -0.1313 | -0.0461 | 0.216 |
| SCMCRM2 | 0.0718 | 0.0286 | 0.0311 | 0.1569 | -0.0312 | -0.4038 | 0.0981 | 0.1886 | 0.0629 | -0.0009 | 0.1147 |
| SHSY5Y | -0.2536 | 0.0294 | 0.0315 | 0.088 | 0.0785 | -0.145 | 0.0987 | 0.1454 | 0.0024 | -0.1771 | 0.2213 |
| SKMEL2 | -0.181 | 0.0972 | -0.1433 | 0.1173 | -0.0082 | -0.1398 | 0.1028 | 0.0222 | -0.1337 | -0.1253 | 0.1356 |
| SKNEP1 | -0.1121 | 0.2105 | -0.1261 | 0.0124 | 0.0771 | -0.0934 | 0.0649 | -0.066 | -0.1808 | -0.0026 | 0.2015 |
| SKPNDW | -0.1539 | 0.0914 | -0.1905 | 0.0496 | -0.0449 | -0.0597 | 0.0964 | -0.0654 | -0.2113 | -0.296 | 0.0057 |
| SKRC31 | 0.0915 | -0.0989 | -0.085 | 0.0901 | -0.0748 | -0.297 | -0.0601 | 0.1805 | -0.0041 | -0.1551 | 0.0985 |
| SMSCTR | -0.0063 | 0.2352 | -0.0878 | 0.1694 | 0.084 | -0.2848 | 0.2689 | 0.1726 | -0.0159 | -0.069 | 0.0275 |
| SMZ1 | 0.12 | -0.0797 | 0.0469 | -0.0818 | -0.0805 | -0.1705 | 0.0024 | -0.0682 | -0.0208 | -0.2275 | 0.0441 |
| TC32 | -0.1257 | -0.0557 | -0.1379 | 0.028 | -0.0102 | -0.0166 | 0.2319 | 0.0684 | -0.095 | -0.0604 | 0.1128 |
| TTC549 | 0.0267 | -0.0136 | -0.0005 | 0.1496 | -0.1404 | -0.2555 | 0.0687 | 0.0726 | -0.057 | -0.1962 | 0.1671 |
| TTC642 | -0.1375 | 0.0191 | 0.1502 | 0.2013 | -0.0738 | -0.2557 | 0.1666 | 0.1032 | -0.0729 | 0.0144 | 0.1962 |
| UPCISCC152 | -0.062 | 0.1651 | -0.0757 | 0.0048 | 0.2506 | -0.2867 | 0.1457 | -0.0503 | -0.2405 | -0.0693 | 0.0241 |
| UPCISCC154 | 0.0387 | 0.0787 | 0.1656 | -0.0748 | 0.1827 | -0.2535 | 0.0691 | -0.1566 | -0.2978 | -0.0782 | 0.1328 |
| UW228 | 0.2241 | -0.002 | 0.0899 | -0.1142 | 0.0308 | -0.219 | -0.1056 | -0.1257 | -0.1987 | 0.1406 | 0.2278 |
| VMRCLCD | 0.0913 | -0.0905 | 0.0678 | 0.084 | -0.1552 | -0.3159 | 0.155 | 0.0835 | -0.0203 | -0.2532 | 0.321 |
| WM2664 | 0.0529 | 0.0203 | 0.0966 | -0.0426 | -0.0008 | -0.3381 | 0.028 | -0.12 | -0.189 | -0.0949 | 0.3033 |
| 127399 | -0.2131 | 0.2936 | -0.1723 | 0.0022 | 0.1534 | -0.2513 | 0.118 | -0.1429 | -0.2949 | -0.0731 | 0.2732 |
| SW982 | 0.0197 | 0.0315 | -0.0537 | 0.1265 | -0.0697 | -0.3827 | 0.1394 | 0.0396 | -0.1242 | -0.0093 | 0.2061 |
| SYO1 | 0.0535 | 0.0458 | 0.0816 | 0.2434 | -0.0903 | -0.2032 | 0.0161 | 0.1504 | 0.0125 | 0.1278 | 0.2478 |
| YAMATO | 0.0448 | 0.1467 | -0.0145 | 0.0505 | -0.0936 | -0.1404 | 0.097 | -0.0529 | -0.005 | -0.1088 | -0.0113 |
| BIN67 | -0.0703 | 0.2055 | 0.0105 | -0.1211 | 0.2104 | -0.2287 | 0.1537 | -0.1905 | -0.3227 | -0.0496 | -0.0846 |
| SCCOHT1 | 0.1319 | -0.0335 | 0.0642 | 0.0467 | -0.3204 | -0.1092 | 0.0046 | -0.0166 | -0.0734 | -0.0123 | 0.1368 |
| SCS214 | -0.1703 | 0.0862 | -0.0351 | -0.0615 | 0.0919 | -0.1313 | 0.0419 | 0.0145 | -0.1707 | -0.0539 | 0.1478 |
| TC106 | -0.0641 | 0.2149 | -0.163 | -0.0031 | 0.1164 | -0.0716 | 0.0694 | -0.0715 | -0.1511 | -0.0617 | 0.138 |
| COGAR359 | -0.1028 | 0.075 | -0.0268 | -0.0212 | -0.046 | -0.2344 | 0.1819 | -0.1408 | -0.1734 | -0.1793 | 0.146 |
| Y79 | 0.0507 | 0.0185 | 0.0807 | 0.1236 | -0.057 | -0.1831 | 0.0218 | 0.0702 | -0.0496 | -0.2474 | 0.16 |
| CHLA15 | -0.1592 | 0.0805 | 0.0524 | 0.0942 | 0.0684 | -0.1506 | 0.1036 | 0.1851 | 0.0011 | -0.0587 | 0.2709 |
| COGN278 | 0.0296 | -0.0387 | -0.0025 | 0.2279 | -0.1353 | -0.1722 | -0.0453 | 0.0712 | -0.0542 | -0.1553 | 0.3803 |
| COGN305 | 0.1005 | 0.0742 | 0.1415 | -0.1246 | 0.0933 | -0.0555 | -0.0792 | -0.0349 | -0.1089 | -0.0407 | 0.1756 |
| NB1643 | 0.1038 | -0.2018 | 0.014 | 0.0545 | -0.1412 | -0.1471 | -0.0451 | 0.069 | 0.0313 | -0.1132 | 0.3445 |
| 8305C | -0.0858 | 0.2194 | 0.064 | -0.0771 | 0.1928 | -0.0636 | -0.0672 | 0.0051 | -0.0306 | -0.0373 | 0.1129 |
| 8505C | 0.0595 | 0.0441 | 0.0542 | 0.2368 | -0.1121 | 0.0383 | 0.0102 | 0.1317 | -0.0003 | -0.082 | 0.2231 |
| HA1E | 0.1543 | -0.0176 | -0.085 | 0.0411 | 0.0193 | -0.1226 | -0.0539 | -0.0733 | -0.2166 | 0.0261 | 0.1673 |
| PLCPRF5 | 0.0325 | 0.0245 | 0.1145 | -0.0531 | 0.1157 | -0.2807 | -0.1007 | -0.1754 | -0.3383 | -0.1649 | 0.2062 |
| CME1 | 0.0638 | -0.1415 | 0.036 | -0.0543 | 0.0955 | -0.1448 | -0.1236 | -0.0446 | -0.0246 | 0.0096 | 0.1856 |
| A431 | -0.1026 | 0.1752 | 0.0765 | -0.1236 | 0.1898 | -0.2379 | 0.0501 | -0.2341 | -0.4235 | -0.027 | 0.0823 |
| ANGMCSS | 0.0445 | 0.095 | 0.1127 | 0.0129 | 0.0198 | -0.2396 | -0.0458 | -0.0427 | -0.1509 | 0.1206 | 0.2953 |
| BICR10 | -0.0139 | -0.0396 | -0.1307 | 0.1387 | -0.0743 | -0.2425 | 0.1076 | 0.209 | 0.0032 | 0.0582 | 0.1483 |
| BICR78 | 0.0756 | 0.013 | 0.0911 | -0.1608 | 0.1514 | -0.2155 | -0.019 | -0.1723 | -0.2116 | -0.0467 | -0.0633 |
| C33A | -0.1217 | 0.1128 | 0.0729 | -0.1618 | 0.1602 | -0.2674 | 0.0913 | -0.1409 | -0.215 | -0.0344 | 0.0566 |
| C4I | -0.0108 | 0.1166 | 0.1446 | 0.004 | 0.1722 | -0.0625 | -0.0294 | -0.0388 | -0.2164 | -0.221 | 0.0796 |
| C4II | 0.0359 | 0.1386 | 0.1338 | 0.1168 | 0.0563 | -0.3097 | 0.0294 | -0.0687 | -0.2585 | -0.3048 | 0.135 |
| CASKI | 0.0492 | -0.003 | 0.2201 | -0.09 | 0.0735 | -0.0644 | 0.0133 | -0.0693 | -0.1685 | -0.2385 | 0.0722 |
| CHP134 | 0.0454 | 0.0166 | -0.1104 | 0.1282 | 0.0516 | -0.2664 | 0.1209 | -0.0192 | -0.1207 | -0.3132 | 0.0342 |
| COV413A | -0.0489 | 0.112 | 0.134 | -0.1137 | 0.2018 | -0.1433 | -0.0076 | -0.1929 | -0.3987 | -0.1974 | 0.0345 |
| DOTC24510 | -0.0311 | 0.1379 | 0.0385 | -0.0927 | 0.2413 | -0.317 | 0.0612 | -0.2015 | -0.4195 | -0.1429 | 0.017 |
| GIMEN | 0.1701 | -0.073 | -0.1374 | -0.0235 | 0.1583 | -0.3709 | -0.0202 | 0.0376 | -0.1063 | -0.0186 | -0.0085 |
| GP5D | -0.0073 | 0.053 | 0.0783 | 0.0104 | 0.0652 | -0.3524 | 0.0593 | 0.0031 | -0.1575 | -0.154 | 0.201 |
| H103 | -0.0564 | 0.1227 | 0.0992 | -0.0256 | 0.1374 | -0.1784 | 0.0707 | -0.1381 | -0.2875 | -0.0646 | 0.126 |
| H157 | 0.087 | 0.1131 | 0.076 | -0.0374 | 0.0964 | -0.2525 | -0.0227 | -0.1247 | -0.2785 | 0.0157 | 0.1401 |
| JOPACA1 | -0.1052 | 0.1521 | 0.0375 | -0.1156 | 0.1932 | -0.0805 | -0.0365 | -0.2053 | -0.3599 | -0.021 | 0.0933 |
| LAN2 | 0.0158 | -0.0694 | 0.1768 | 0.1982 | -0.02 | -0.0805 | -0.0105 | 0.1464 | -0.0117 | -0.1988 | 0.2393 |
| MB1 | -0.1213 | 0.0506 | -0.0829 | 0.0296 | -0.0335 | -0.2214 | 0.1118 | 0.1189 | 0.0015 | 0.0454 | 0.2716 |
| MS751 | 0.1238 | 0.1124 | 0.2284 | 0.0394 | 0.1086 | -0.0636 | 0.0348 | 0.019 | -0.0525 | -0.1486 | 0.0873 |
| NGP | -0.0491 | 0.0187 | 0.0443 | 0.2492 | -0.0841 | -0.1985 | 0.1669 | 0.0883 | -0.0931 | -0.1466 | 0.1679 |
| NMB | -0.1529 | 0.2192 | -0.1012 | 0.0707 | 0.1841 | -0.2218 | 0.2304 | 0.0983 | -0.0816 | 0.0234 | 0.1495 |
| OACM51 | 0.0152 | 0.0953 | 0.0181 | 0.1272 | 0.0527 | -0.1821 | -0.0286 | 0.0855 | 0.061 | 0.054 | 0.1993 |
| OCIP5X | 0.1819 | -0.1211 | 0.0316 | 0.024 | 0.1326 | -0.1692 | -0.1866 | 0.0686 | -0.1058 | -0.2492 | 0.072 |
| PA1 | -0.2911 | 0.1123 | 0.106 | 0.0876 | -0.0099 | -0.3106 | 0.3679 | 0.0456 | -0.1264 | -0.0173 | 0.299 |
| PACADD119 | -0.1701 | 0.1738 | -0.0001 | -0.0866 | 0.2417 | -0.1978 | 0.1579 | -0.2097 | -0.4222 | -0.1176 | 0.0812 |
| PACADD137 | -0.0305 | -0.0103 | 0.1906 | -0.0471 | 0.0594 | -0.0952 | -0.1616 | -0.0807 | -0.2847 | -0.2775 | 0.0116 |
| PACADD161 | -0.0144 | 0.152 | -0.0446 | 0.044 | 0.1439 | -0.2878 | 0.1982 | 0.0169 | -0.1623 | -0.0051 | 0.2008 |
| PACADD165 | 0.0552 | 0.0004 | -0.0626 | 0.1 | -0.1002 | -0.0459 | 0.0523 | 0.0113 | -0.0862 | 0.0228 | 0.011 |
| PACADD188 | -0.1576 | 0.1247 | -0.0212 | -0.0083 | 0.0606 | -0.2954 | 0.2939 | -0.0423 | -0.2715 | -0.0915 | -0.075 |
| RPMI2650 | 0.2036 | -0.0655 | 0.0178 | 0.0763 | -0.1018 | -0.2652 | -0.1269 | 0.0309 | -0.0624 | -0.0835 | 0.0816 |
| SCLC22H | -0.0188 | 0.1926 | -0.1136 | 0.2181 | 0.0568 | -0.186 | 0.0972 | 0.1262 | 0.03 | 0.0092 | 0.0611 |
| SUM102PT | 0.0181 | 0.1468 | 0.1625 | 0.0254 | 0.0811 | -0.0358 | -0.003 | -0.1506 | -0.2486 | -0.1777 | 0.2071 |
| SUM1315MO2 | -0.0095 | 0.0077 | -0.0253 | 0.1448 | -0.1765 | -0.043 | -0.0141 | -0.0018 | -0.0782 | 0.0197 | 0.0961 |
| SUM149PT | 0.0527 | -0.1103 | 0.2492 | -0.21 | -0.0192 | -0.106 | -0.0484 | -0.3694 | -0.3455 | -0.2287 | 0.1123 |
| SUM159PT | -0.0649 | -0.054 | 0.078 | -0.033 | -0.0304 | -0.0412 | -0.1744 | -0.1691 | -0.2195 | -0.1872 | 0.0876 |
| SUM185PE | -0.1386 | 0.179 | 0.0471 | -0.0422 | 0.1283 | -0.3181 | 0.36 | -0.1025 | -0.1058 | 0.0864 | 0.2755 |
| SUM190PT | 0.0167 | 0.1083 | 0.0472 | -0.0315 | 0.1708 | -0.2678 | 0.0219 | -0.0895 | -0.2199 | -0.0892 | 0.0573 |
| SUM229PE | -0.1443 | 0.1461 | 0.1028 | 0.0648 | -0.0262 | -0.1568 | 0.1485 | -0.1255 | -0.2805 | -0.1398 | 0.1729 |
| SUM52PE | 0.182 | 0.0344 | 0.0515 | -0.1702 | 0.1003 | -0.0578 | 0.2025 | -0.0934 | -0.1244 | 0.1111 | 0.2037 |
| SW156 | -0.0458 | 0.045 | 0.0694 | 0.0048 | 0.0461 | -0.1144 | 0.0377 | -0.1279 | -0.1586 | -0.1636 | 0.1685 |
| SW626 | -0.02 | 0.0803 | 0.0974 | 0.0507 | 0.0706 | -0.1905 | 0.1288 | 0.0543 | -0.076 | -0.115 | 0.1824 |
| SW954 | -0.0688 | 0.0803 | 0.0911 | -0.1519 | 0.2751 | -0.2265 | -0.0785 | -0.1592 | -0.3498 | -0.1873 | 0.0421 |
| SW756 | -0.0008 | 0.0389 | 0.1449 | -0.007 | 0.0517 | -0.1398 | -0.0942 | 0.0026 | -0.1332 | -0.1526 | 0.262 |
| TO14 | 0.012 | 0.0389 | 0.0604 | 0.1399 | -0.0538 | -0.3756 | 0.1377 | 0.0993 | -0.0157 | -0.1892 | 0.1225 |
| UMUC13 | 0.0179 | 0.0242 | -0.0608 | 0.073 | -0.1296 | -0.0189 | 0.0041 | 0.0284 | -0.0764 | 0.1303 | -0.0251 |

| R | EC | IDRE | TM | IDRC | Linker | Tail | IDRE pos | IDRC pos | RK cluste | GxxxG | CRAC |
|---|---|---|---|---|---|---|---|---|---|---|---|
| UMUC16 | -0.2013 | 0.0315 | -0.0067 | -0.0236 | -0.08 | -0.1057 | 0.0401 | 0.0094 | -0.1428 | -0.2208 | 0.4018 |
| UMUC5 | 0.0338 | 0.207 | 0.0195 | 0.0245 | 0.1441 | -0.3487 | 0.1446 | -0.0157 | -0.0819 | -0.0002 | 0.0204 |
| UMUC10 | -0.0188 | 0.0696 | 0.0728 | -0.1088 | 0.151 | -0.2585 | 0.0434 | -0.1074 | -0.3404 | -0.0516 | -0.0171 |
| UMUC11 | 0.1537 | -0.0231 | 0.0485 | 0.1235 | -0.0723 | -0.1236 | 0.0964 | 0.0727 | -0.0459 | 0.0798 | 0.0499 |
| UMUC6 | 0.0504 | 0.01 | 0.0458 | 0.1778 | 0.0031 | -0.1655 | 0.0679 | 0.0903 | -0.026 | -0.0513 | 0.2293 |
| UMUC7 | -0.1013 | 0.0417 | -0.0891 | -0.0776 | 0.097 | -0.1547 | 0.0198 | -0.0601 | -0.2404 | -0.1011 | 0.0482 |
| UMUC9 | 0.0468 | -0.0074 | -0.0065 | -0.1301 | 0.1254 | -0.1658 | -0.0577 | -0.0894 | -0.3165 | -0.0442 | 0.2274 |
| UWB1289 | 0.073 | 0.103 | 0.0864 | -0.1519 | 0.1261 | -0.3147 | 0.0314 | -0.2586 | -0.4356 | 0.0064 | 0.0669 |
| VP229 | 0.0311 | -0.0092 | -0.0429 | 0.0032 | 0.1574 | -0.2005 | 0.0144 | -0.0528 | -0.2336 | -0.1101 | 0.0788 |
| WERIRB1 | -0.0209 | 0.1529 | -0.0306 | 0.1051 | 0.0842 | -0.148 | 0.0331 | 0.0119 | -0.1707 | -0.1354 | 0.1063 |
| WPE1NA22 | 0.0123 | 0.1154 | 0.121 | 0.0728 | 0.1319 | -0.1721 | 0.0999 | -0.0335 | -0.1834 | -0.2382 | 0.0359 |
| TC138 | -0.2043 | 0.3062 | -0.1947 | 0.0402 | 0.1061 | -0.1623 | 0.3178 | 0.0923 | -0.0682 | 0.0452 | 0.0901 |
| TC205 | 0.0762 | -0.0673 | -0.0326 | 0.0169 | -0.0396 | -0.1175 | -0.0003 | -0.0157 | -0.0721 | -0.1585 | 0.185 |
| CCLFPEDS0008T | -0.11 | 0.1279 | 0.0483 | -0.0529 | 0.056 | -0.0319 | 0.0322 | 0.0924 | -0.0962 | 0.094 | -0.0032 |
| A388 | -0.0604 | 0.0613 | 0.1295 | -0.1224 | 0.1546 | -0.2146 | 0.0353 | -0.1933 | -0.3772 | -0.1821 | 0.0412 |
| ASH3 | -0.0803 | 0.021 | 0.1241 | -0.079 | 0.1042 | -0.1573 | 0.0159 | -0.1252 | -0.2016 | -0.2093 | 0.0258 |
| BLUE1 | 0.0795 | 0.1462 | 0.2191 | -0.0229 | -0.0627 | -0.0071 | -0.0407 | -0.1959 | -0.2644 | -0.2025 | 0.242 |
| BOKU | 0.0008 | 0.1205 | -0.0959 | 0.0142 | 0.1009 | -0.1982 | 0.0526 | -0.0472 | -0.1667 | 0.0448 | 0.2449 |
| BPH1 | -0.0083 | 0.1499 | 0.0164 | -0.0725 | 0.2268 | -0.2038 | -0.0445 | -0.1716 | -0.3372 | -0.0192 | 0.0268 |
| C10 | 0.1542 | 0.0528 | 0.0639 | 0.1099 | -0.084 | -0.2181 | 0.2104 | 0.0742 | -0.0101 | 0.0729 | 0.33 |
| C75 | -0.2262 | 0.181 | -0.023 | -0.156 | 0.1948 | -0.2874 | 0.0244 | -0.2693 | -0.4082 | -0.1017 | 0.0953 |
| C80 | -0.023 | 0.0735 | 0.0579 | -0.0106 | 0.1464 | -0.445 | 0.1584 | 0.0096 | -0.1932 | 0.1185 | 0.1357 |
| C84 | -0.0555 | 0.0263 | 0.1333 | -0.1693 | 0.2481 | 0.0235 | 0.101 | -0.1686 | -0.4268 | -0.1669 | -0.0888 |
| C99 | 0.1139 | 0.0252 | -0.0746 | 0.0281 | 0.0271 | -0.3844 | 0.0182 | -0.0009 | -0.2062 | -0.1941 | 0.1539 |
| CHLA90 | 0.0838 | 0.1593 | -0.1256 | 0.1262 | -0.0548 | -0.2404 | -0.0017 | 0.0137 | -0.1477 | -0.077 | 0.0066 |
| EGI1 | 0.0802 | 0.0117 | 0.1532 | -0.1495 | -0.0041 | -0.1124 | 0.0362 | -0.2697 | -0.3181 | -0.1547 | 0.1455 |
| EMTOKA | 0.0813 | 0.1589 | 0.022 | -0.0225 | 0.05 | -0.2779 | -0.0219 | -0.1433 | -0.228 | -0.0707 | -0.0064 |
| ESO26 | -0.0206 | 0.1361 | 0.0081 | -0.1691 | 0.2571 | -0.1956 | 0.016 | -0.2598 | -0.4434 | -0.1292 | -0.0082 |
| ESO51 | 0.1288 | -0.0243 | -0.2041 | 0.1085 | -0.1662 | -0.4262 | 0.1934 | 0.0822 | 0.0109 | 0.0632 | 0.1138 |
| FARAGE | 0.0088 | -0.1956 | 0.1354 | 0.0304 | -0.1063 | -0.1862 | -0.0102 | -0.0167 | -0.15 | -0.1999 | 0.124 |
| FLO1 | 0.0954 | -0.0567 | 0.113 | 0.0219 | 0.0014 | -0.2342 | 0.001 | -0.095 | -0.1434 | -0.1382 | 0.3104 |
| H357 | -0.1024 | -0.0495 | -0.0569 | -0.0601 | 0.1214 | -0.2763 | 0.0125 | -0.1306 | -0.3298 | -0.1506 | 0.1588 |
| H376 | 0.0703 | 0.1604 | -0.0359 | -0.0495 | 0.0813 | -0.2298 | 0.1041 | -0.0421 | -0.1879 | 0.0862 | -0.0789 |
| H413 | 0.0173 | 0.177 | 0.0521 | -0.1769 | 0.1921 | -0.1381 | 0.0774 | -0.2963 | -0.4221 | -0.0449 | 0.0476 |
| HCA1 | 0.0781 | 0.029 | 0.1807 | -0.1339 | 0.1607 | -0.2736 | -0.0076 | -0.1019 | -0.2964 | -0.1301 | 0.0536 |
| HCS2 | 0.0072 | 0.0803 | 0.1188 | 0.0624 | 0.0238 | -0.1238 | 0.0131 | -0.0465 | -0.1916 | -0.0893 | 0.1038 |
| HEC1 | -0.0605 | 0.0424 | 0.1714 | -0.0031 | 0.0696 | -0.2074 | -0.0676 | -0.0403 | -0.1641 | -0.1832 | 0.2127 |
| HEC116 | -0.1184 | 0.0564 | -0.0307 | 0.1288 | 0.0394 | -0.0277 | 0.0755 | 0.1074 | -0.0421 | -0.3114 | 0.0392 |
| HG3 | 0.0784 | -0.0566 | -0.0217 | -0.2466 | 0.1668 | -0.1072 | -0.0495 | -0.1612 | -0.2445 | -0.0992 | -0.0124 |
| HKA1 | 0.0434 | 0.0131 | -0.1031 | -0.0126 | 0.1471 | -0.3503 | 0.1103 | -0.1405 | -0.3047 | -0.0768 | 0.0824 |
| HMY1 | 0.0391 | 0.1064 | -0.0685 | 0.1949 | 0.0398 | -0.2256 | 0.0549 | 0.0736 | -0.0259 | -0.0572 | 0.1922 |
| HSC1 | -0.1169 | 0.2538 | 0.1354 | -0.0996 | 0.2267 | -0.1999 | 0.13 | -0.0826 | -0.2308 | 0.1633 | 0.0213 |
| HSC5 | 0.0252 | 0.0859 | 0.1686 | -0.1226 | 0.1376 | -0.2254 | -0.0447 | -0.2177 | -0.3954 | -0.0988 | 0.2029 |
| HT3 | 0.0645 | 0.2246 | -0.0042 | 0.2092 | -0.195 | -0.0473 | 0.0431 | 0.19 | 0.1145 | 0.063 | 0.1247 |
| HUO9 | -0.1612 | 0.1432 | -0.0803 | 0.1557 | 0.0837 | -0.1367 | -0.0495 | 0.0573 | 0.0143 | 0.052 | 0.1376 |
| IHH4 | 0.0218 | 0.1622 | 0.1284 | -0.0165 | 0.083 | -0.1802 | -0.0151 | -0.1395 | -0.229 | -0.209 | 0.2135 |
| JAR | -0.074 | -0.0055 | 0.0849 | 0.0594 | 0.0375 | 0.0319 | -0.0322 | -0.0551 | -0.0675 | -0.1343 | 0.1008 |
| JEG3 | 0.1883 | 0.0763 | 0.0893 | 0.0826 | 0.0556 | -0.4443 | -0.018 | 0.0382 | -0.1301 | -0.0514 | 0.2089 |
| JMURTK2 | 0.1988 | 0.0127 | 0.0294 | -0.0235 | -0.1325 | -0.0274 | -0.0326 | -0.0202 | -0.1009 | 0.1276 | 0.1063 |
| KARPAS1718 | 0.0596 | 0.1579 | 0.1277 | 0.1066 | 0.0411 | -0.1323 | 0.1289 | 0.0015 | -0.1231 | -0.213 | -0.0124 |
| KKU100 | -0.0213 | 0.1688 | 0.2893 | -0.0911 | 0.1164 | -0.1189 | -0.0731 | -0.0483 | -0.1677 | -0.0161 | 0.1482 |
| KKU213 | -0.1288 | 0.0689 | -0.0239 | -0.2136 | 0.2117 | -0.1763 | -0.0252 | -0.0861 | -0.2031 | -0.0862 | -0.0249 |
| KML1 | -0.0843 | -0.096 | -0.0774 | 0.0624 | -0.0747 | -0.3363 | 0.0183 | 0.0059 | -0.1491 | -0.1565 | 0.1614 |
| KON | 0.0763 | 0.0108 | 0.0976 | -0.0704 | 0.0287 | -0.1686 | -0.0474 | -0.0395 | -0.2077 | 0.0152 | 0.0742 |
| KOSC2 | 0.1466 | 0.1745 | -0.0145 | -0.1086 | 0.1882 | -0.2022 | 0.0107 | -0.1111 | -0.2746 | -0.0092 | 0.0399 |
| KYAE1 | -0.0736 | 0.1061 | -0.033 | -0.3739 | 0.3017 | -0.121 | 0.003 | -0.409 | -0.5754 | -0.1268 | -0.1536 |
| LO68 | 0.08 | 0.1324 | -0.1663 | 0.1677 | -0.0422 | -0.0101 | 0.0512 | 0.1081 | 0.0189 | 0.0975 | 0.1045 |
| LS | 0.0026 | 0.2569 | -0.1588 | 0.0691 | 0.1108 | -0.4228 | 0.3291 | 0.0476 | -0.1389 | 0.0545 | 0.1354 |
| LU135 | -0.0662 | 0.0984 | 0.0209 | -0.0011 | 0.0453 | -0.1248 | 0.0315 | 0.0329 | -0.1146 | 0.0197 | 0.2282 |
| MCC13 | 0.083 | 0.0116 | 0.0866 | 0.0488 | 0.0915 | -0.3289 | 0.0975 | 0.0425 | -0.054 | -0.1941 | 0.2011 |
| MCC142 | 0.1188 | 0.0155 | -0.0044 | 0.1498 | -0.2074 | -0.0811 | 0.0648 | 0.0601 | -0.0699 | 0.0839 | 0.2697 |
| MCC26 | -0.2141 | 0.198 | -0.1202 | 0.1792 | 0.0246 | -0.1656 | 0.1345 | 0.1067 | -0.0296 | 0.1212 | 0.1749 |
| MEL202 | -0.0521 | 0.1191 | 0.0946 | 0.0741 | -0.0342 | -0.2715 | 0.0708 | -0.0727 | -0.2324 | -0.0931 | 0.1637 |
| MERO14 | 0.1067 | -0.0173 | -0.1928 | 0.1887 | -0.1661 | -0.0901 | 0.0645 | 0.1294 | -0.1089 | -0.0386 | 0.0252 |
| MERO25 | 0.206 | 0.0417 | 0.1569 | 0.0993 | -0.0668 | -0.3271 | 0.063 | 0.0343 | -0.0769 | -0.0787 | 0.2564 |
| MERO41 | 0.0696 | 0.0633 | 0.1039 | 0.1193 | -0.159 | -0.0147 | 0.0277 | -0.0248 | 0.0105 | 0.1411 | 0.1592 |
| MERO48A | 0.2948 | 0.0186 | 0.0663 | 0.0965 | -0.075 | -0.2499 | -0.0674 | 0.0366 | 0.0345 | 0.1488 | 0.2141 |
| MERO82 | 0.1929 | 0.0161 | 0.0802 | 0.1261 | -0.1267 | -0.2177 | 0.072 | 0.0771 | -0.0031 | 0.1119 | 0.1736 |
| MERO83 | 0.0548 | 0.0936 | 0.017 | 0.0145 | -0.1008 | -0.1088 | 0.0447 | -0.0809 | -0.1688 | 0.0158 | -0.1122 |
| MERO95 | 0.1184 | 0.0094 | 0.0362 | 0.123 | -0.1144 | -0.0313 | 0.0265 | 0.0142 | -0.0419 | 0.0607 | 0.0367 |
| MM127 | 0.0918 | -0.0018 | 0.0451 | 0.1843 | -0.097 | -0.2319 | 0.0434 | 0.1069 | 0.028 | -0.0882 | 0.2181 |
| MM370 | 0.0997 | 0.0878 | 0.2466 | -0.0081 | -0.1369 | 0.0026 | -0.063 | -0.1267 | -0.0385 | -0.071 | 0.1511 |
| MM383 | -0.1093 | 0.1656 | -0.0003 | 0.0363 | 0.3141 | -0.0875 | 0.023 | -0.0975 | -0.2253 | -0.1452 | 0.1222 |
| MM386 | -0.1058 | 0.0076 | 0.0422 | -0.0255 | 0.0278 | -0.0226 | -0.0791 | -0.1203 | -0.2573 | -0.226 | 0.1748 |
| MM426 | 0.0795 | 0.0788 | 0.159 | 0.0253 | -0.0359 | -0.1597 | 0.0842 | -0.0353 | -0.2265 | -0.0965 | 0.1435 |
| MOLM14 | 0.1663 | 0.084 | 0.0838 | 0.1657 | 0.0127 | -0.104 | 0.009 | 0.0424 | -0.0057 | -0.1327 | 0.2184 |
| MUTZ8 | -0.0518 | 0.0782 | -0.0664 | 0.1065 | 0.0472 | -0.2779 | -0.0576 | 0.0191 | -0.0229 | -0.081 | 0.1769 |
| NH12 | -0.2614 | 0.0778 | 0.0273 | 0.0892 | 0.066 | -0.1143 | 0.1214 | 0.0622 | -0.1215 | -0.1473 | 0.0245 |
| NO10 | 0.1488 | -0.0466 | 0.1056 | 0.0916 | -0.0447 | -0.2898 | -0.1622 | 0.0013 | -0.1097 | -0.0534 | 0.2642 |
| NO11 | 0.0623 | -0.0262 | 0.0445 | 0.0443 | -0.1444 | 0.0267 | 0.0132 | -0.1893 | -0.1304 | 0.1179 | 0.1772 |
| NOZ | 0.2146 | 0.0257 | -0.0143 | -0.0896 | 0.0505 | -0.3271 | -0.0591 | -0.171 | -0.1085 | -0.1395 | 0.0744 |
| NP2 | 0.1328 | -0.0744 | 0.0123 | 0.061 | -0.1892 | -0.1758 | -0.0273 | 0.0143 | -0.1179 | -0.0018 | 0.1957 |
| NP3 | 0.1166 | -0.0219 | 0.0326 | 0.1072 | -0.183 | -0.0228 | -0.0408 | 0.1129 | 0.0625 | 0.138 | 0.144 |
| NP5 | 0.0616 | -0.0393 | 0.0738 | 0.117 | 0.0435 | -0.0612 | -0.223 | 0.0996 | 0.1142 | -0.1089 | 0.1948 |
| NP8 | 0.0068 | 0.0333 | -0.0186 | 0.1454 | 0.015 | -0.1249 | 0.1052 | 0.1043 | 0.0255 | -0.0711 | -0.015 |
| OCILY18 | 0.0309 | 0.2205 | 0.0013 | -0.0537 | -0.0431 | -0.179 | 0.0285 | -0.1571 | -0.2576 | -0.13 | 0.083 |
| OCIM2 | -0.0197 | -0.0246 | -0.0972 | -0.1352 | 0.1125 | -0.1651 | -0.0549 | -0.1626 | -0.2089 | -0.2781 | 0.0326 |
| OCUG1 | -0.1486 | 0.189 | 0.046 | 0.0107 | 0.1736 | -0.183 | 0.0791 | -0.0291 | -0.2111 | -0.0249 | -0.0068 |
| ONDA7 | 0.1656 | 0.1206 | 0.0928 | 0.0147 | 0.0388 | -0.2092 | 0.006 | -0.0374 | -0.1906 | 0.0135 | 0.1934 |
| ONDA8 | 0.0321 | 0.019 | -0.0544 | 0.1594 | -0.0841 | -0.0672 | 0.0414 | 0.1137 | 0.0464 | 0.1177 | -0.0389 |
| ONDA9 | -0.0192 | 0.0639 | -0.0135 | 0.2887 | -0.2106 | -0.1954 | 0.1265 | 0.1215 | 0.0106 | -0.0494 | 0.1844 |
| OSC19 | 0.0069 | -0.0985 | 0.1006 | -0.2008 | 0.1407 | -0.1397 | -0.0304 | -0.3133 | -0.4521 | -0.1321 | 0.0463 |
| OSC20 | 0.0431 | 0.052 | 0.1234 | -0.0464 | 0.2162 | -0.1722 | 0.0622 | -0.0026 | -0.2101 | -0.0611 | 0.0888 |
| P4E6 | -0.0164 | 0.1496 | 0.0713 | 0.0058 | 0.1012 | -0.1446 | 0.0577 | -0.0959 | -0.1975 | -0.1081 | -0.0412 |
| PEA1 | -0.0509 | 0.1659 | 0.0103 | 0.0114 | 0.1472 | -0.2617 | 0.0114 | -0.097 | -0.231 | 0.0371 | 0.2472 |
| PEO1 | -0.0017 | 0.226 | 0.0044 | -0.0652 | 0.1886 | -0.2831 | 0.0872 | -0.1728 | -0.3368 | 0.0296 | 0.1132 |
| PEO4 | 0.0537 | 0.1132 | -0.12 | -0.0061 | 0.119 | -0.3156 | 0.0924 | -0.0571 | -0.1933 | -0.0645 | -0.0085 |
| PGA1 | 0.0537 | 0.176 | 0.1293 | 0.1524 | -0.0645 | 0.0144 | 0.061 | 0.0039 | -0.0516 | -0.2243 | 0.109 |
| ROS50 | -0.0311 | -0.0204 | 0.1619 | 0.1424 | -0.1718 | 0.166 | 0.0868 | 0.0641 | 0.1056 | -0.1448 | 0.2858 |
| SAT | 0.031 | 0.1155 | 0.1099 | -0.0243 | 0.2177 | -0.195 | 0.0154 | -0.1238 | -0.2974 | -0.1308 | 0.0379 |
| SEKI | 0.0739 | 0.1889 | -0.0409 | 0.2034 | -0.0772 | -0.2196 | 0.0066 | 0.076 | 0.0145 | -0.1081 | 0.1741 |
| SHI1 | -0.2154 | 0.1924 | 0.027 | 0.2176 | -0.034 | -0.0655 | 0.2387 | 0.1067 | -0.0184 | -0.1697 | 0.226 |
| SHMAC4 | -0.0068 | 0.1179 | 0.0742 | -0.004 | 0.0891 | -0.1875 | 0.0593 | -0.1119 | -0.2747 | -0.1605 | 0.0168 |
| SHMAC5 | 0.0567 | 0.0642 | 0.2689 | -0.0356 | 0.1688 | -0.2502 | -0.0547 | -0.0756 | -0.1534 | -0.0908 | 0.0692 |
| SISO | -0.1575 | 0.204 | -0.0151 | -0.0178 | 0.172 | -0.1606 | 0.1396 | -0.0449 | -0.2448 | -0.1241 | -0.0773 |
| SKGI | 0.0051 | 0.0592 | 0.0277 | 0.0397 | 0.1324 | -0.1077 | -0.1858 | -0.0065 | -0.1308 | 0.0224 | 0.2671 |
| SKGII | 0.04 | 0.0497 | 0.1541 | -0.2259 | 0.1469 | -0.1983 | -0.006 | -0.2797 | -0.3915 | -0.0402 | 0.0444 |
| SKGT2 | -0.0687 | 0.1325 | -0.0016 | 0.0209 | 0.0677 | -0.3074 | 0.1384 | -0.0904 | -0.2117 | 0.0136 | 0.2158 |
| SKGT4 | 0.0777 | 0.1623 | 0.1964 | -0.0806 | 0.1746 | -0.2996 | 0.0927 | -0.2342 | -0.422 | -0.1496 | 0.1473 |
| SKN | -0.0515 | 0.0113 | -0.1558 | -0.1451 | 0.1197 | -0.2724 | 0.1077 | -0.0979 | -0.2781 | -0.0553 | 0.2362 |
| SKNO1 | 0.0601 | 0.0529 | 0.0759 | 0.2325 | -0.0533 | -0.1386 | -0.0037 | 0.16 | 0.0568 | -0.1214 | 0.1874 |
| SNU638 | -0.1249 | 0.1405 | -0.0963 | 0.047 | -0.2862 | 0.0216 | 0.1553 | -0.035 | -0.0691 | -0.033 | -0.0009 |
| SUSA | -0.0384 | 0.0418 | 0.0513 | 0.0857 | -0.0122 | -0.0318 | 0.171 | -0.0189 | -0.201 | -0.0199 | 0.0464 |
| TANOUE | 0.0968 | 0.0737 | 0.013 | 0.1728 | 0.0624 | -0.1482 | 0.0421 | 0.0955 | -0.0704 | -0.3234 | 0.2238 |
| TASK1 | -0.0395 | 0.1031 | -0.0254 | 0.0434 | 0.2084 | -0.0797 | 0.0041 | 0.0335 | -0.1194 | -0.0859 | 0.1483 |
| TFK1 | -0.1001 | 0.1249 | 0.0079 | -0.0316 | 0.1499 | -0.1304 | 0.0987 | -0.0372 | -0.2315 | -0.1215 | -0.0184 |
| TGW | -0.0744 | 0.041 | -0.102 | 0.1817 | 0.0551 | -0.3178 | 0.1325 | 0.1599 | -0.0104 | -0.2475 | 0.1379 |
| TR146 | 0.0431 | 0.1903 | 0.1885 | 0.0972 | 0.0967 | -0.2735 | -0.0468 | -0.0367 | -0.1891 | -0.1097 | 0.117 |
| U2904 | -0.0628 | 0.2081 | -0.0779 | 0.0475 | 0.2105 | -0.1859 | 0.1422 | -0.0406 | -0.1392 | 0.0133 | 0.163 |
| UHO1 | 0.1099 | 0.0671 | -0.0405 | 0.2335 | -0.0773 | -0.4679 | 0.1824 | 0.2004 | 0.0504 | -0.1015 | 0.3056 |
| UMRC3 | 0.1014 | -0.0697 | 0.0713 | 0.1658 | -0.219 | -0.0123 | -0.0247 | 0.0553 | 0.0359 | -0.0401 | 0.1987 |
| UMRC7 | -0.0572 | 0.1828 | 0.0495 | -0.0869 | 0.1382 | -0.0144 | -0.0159 | -0.0327 | -0.0559 | 0.1139 | -0.0651 |
| UPCISCC026 | -0.0004 | 0.1075 | 0.1126 | -0.001 | 0.0956 | -0.2464 | -0.0015 | -0.1294 | -0.3076 | -0.1141 | 0.0899 |
| UPCISCC029A | 0.0677 | 0.0688 | -0.1027 | 0.0608 | -0.0403 | -0.0808 | 0.0203 | 0.0064 | -0.0916 | 0.076 | 0.0434 |
| UPCISCC040 | 0.057 | 0.0625 | 0.0752 | -0.0452 | 0.1311 | -0.2079 | 0.0134 | -0.156 | -0.3096 | -0.1364 | 0.1093 |
| UPCISCC074 | 0.0112 | 0.1598 | 0.1712 | -0.0789 | 0.2211 | -0.1831 | 0.0252 | -0.1663 | -0.3419 | -0.1297 | 0.0325 |
| UPCISCC111 | 0.0694 | 0.1282 | 0.0847 | -0.0741 | 0.1719 | -0.3238 | 0.0188 | -0.1337 | -0.2977 | 0.0458 | -0.0163 |
| UPCISCC116 | -0.076 | 0.1528 | 0.0234 | -0.1338 | 0.2549 | -0.2763 | 0.1723 | -0.2155 | -0.3633 | -0.0311 | -0.0295 |

| R | EC | IDRE | TM | IDRC | Linker | Tail | IDRE pos | IDRC pos | RK cluste | GxxxG | CRAC |
|---|---|---|---|---|---|---|---|---|---|---|---|
| UPCISCC200 | 0.1099 | 0.1113 | 0.1105 | -0.0021 | 0.0894 | -0.2898 | 0.0005 | -0.0453 | -0.0958 | 0.0078 | 0.122 |
| VAESBJ | 0.2082 | 0.1137 | 0.0684 | 0.1391 | -0.1544 | -0.3212 | -0.0305 | 0.0539 | 0.0402 | -0.0422 | 0.2305 |
| WAOSEL | 0.1101 | 0.0553 | 0.1641 | 0.1549 | -0.0066 | -0.1394 | 0.0289 | 0.0715 | 0.0208 | -0.1813 | 0.2325 |
| WSUNHL | 0.1453 | -0.0046 | -0.0424 | -0.1028 | 0.1667 | -0.165 | -0.0525 | -0.0301 | -0.2437 | -0.2888 | 0.0697 |
| PFSK1 | -0.0676 | 0.0844 | -0.2835 | 0.0787 | 0.1514 | -0.2561 | 0.149 | 0.1149 | 0.0273 | -0.0442 | -0.1654 |
| CAL72 | -0.0353 | -0.0253 | -0.1101 | 0.1211 | -0.184 | -0.0289 | 0.1342 | 0.1177 | 0.0215 | -0.006 | -0.0133 |
| OCIC4P | 0.0745 | 0.0539 | 0.1023 | 0.1217 | -0.0062 | -0.1897 | 0.0584 | -0.0308 | -0.1977 | -0.1513 | 0.1382 |
| SEMK2 | 0.0064 | 0.1605 | 0.2253 | 0.1212 | 0.0347 | -0.0862 | 0.1464 | 0.12 | 0.053 | -0.0715 | 0.1113 |
| HB1119 | -0.0016 | 0.1914 | 0.1732 | 0.1291 | 0.0536 | -0.08 | 0.1147 | 0.1007 | 0.0282 | -0.0593 | 0.0457 |
| CTV1DM | -0.0836 | 0.0728 | 0.1132 | 0.1819 | 0.0007 | -0.1705 | 0.2362 | 0.1948 | 0.012 | -0.2247 | 0.1457 |
| RH28 | -0.0395 | 0.0629 | -0.0243 | -0.0774 | 0.0109 | -0.0906 | 0.0811 | -0.1563 | -0.2634 | -0.3212 | 0.1588 |
| RHJT | -0.1021 | 0.1728 | 0.0167 | 0.0131 | -0.0418 | -0.1312 | 0.1889 | -0.1063 | -0.1351 | -0.1101 | 0.1211 |
| TTC442 | -0.0548 | 0.1601 | 0.0078 | -0.0983 | 0.1109 | -0.2943 | 0.2045 | -0.2839 | -0.3722 | -0.1181 | 0.1165 |
| RH4 | -0.038 | 0.042 | 0.0625 | 0.0353 | -0.0567 | -0.0587 | 0.0611 | 0.0044 | -0.06 | -0.0667 | 0.1386 |
| SNU1544 | -0.0261 | 0.0069 | 0.1452 | 0.0513 | 0.148 | -0.2343 | -0.0558 | 0.0723 | -0.205 | -0.0897 | 0.1643 |
| LPS6 | -0.0548 | -0.057 | 0.1392 | 0.0596 | -0.0402 | -0.148 | 0.0782 | 0.0033 | -0.0746 | 0.151 | 0.0491 |
| LPS27 | 0.1459 | 0.1117 | 0.0823 | 0.1086 | -0.1514 | -0.1487 | 0.1059 | 0.0236 | 0.0015 | 0.1388 | 0.1279 |
| 93T449 | 0.1421 | 0.009 | -0.0791 | -0.0339 | -0.0849 | -0.0271 | 0.017 | -0.136 | -0.2015 | 0.0316 | 0.0062 |
| 94T778 | 0.1473 | -0.0201 | 0.102 | -0.1064 | -0.0631 | -0.1787 | -0.1146 | -0.1698 | -0.2147 | 0.1544 | 0.1398 |
| 95T1000 | 0.1334 | -0.0984 | 0.01 | -0.0822 | -0.0469 | -0.1953 | -0.0638 | -0.1428 | -0.2415 | 0.013 | 0.026 |
| LPS141 | -0.1084 | 0.1934 | 0.0935 | 0.1753 | 0.0632 | -0.056 | 0.1322 | 0.1027 | -0.1266 | -0.0552 | -0.0564 |
| LPS853 | -0.0547 | 0.1838 | 0.0096 | 0.1464 | -0.0705 | 0.0093 | 0.0171 | 0.0154 | -0.1265 | -0.1786 | 0.0611 |
| LPS510 | 0.2197 | -0.0219 | 0.0886 | 0.0531 | -0.1644 | -0.058 | -0.0369 | -0.0935 | -0.0684 | -0.0065 | -0.1173 |
| OS252 | -0.0191 | 0.0965 | -0.0282 | -0.1679 | 0.1356 | -0.2417 | 0.1488 | -0.1614 | -0.3282 | 0.0114 | -0.0025 |
| MFM223 | 0.146 | 0.0071 | 0.1097 | -0.2338 | 0.0627 | -0.0971 | -0.1566 | -0.1846 | -0.2945 | 0.0334 | 0.2165 |
| COLO824 | -0.0577 | 0.1751 | 0.0923 | -0.054 | 0.1503 | -0.1048 | 0.1278 | -0.0503 | -0.1913 | -0.0504 | -0.0023 |
| ICC10 | 0.0772 | 0.0932 | 0.1359 | -0.0299 | 0.0707 | -0.2529 | -0.0616 | -0.0833 | -0.2211 | 0.0876 | 0.2307 |
| ICC106 | 0.132 | 0.0286 | 0.0733 | -0.0469 | -0.0058 | -0.2606 | 0.0008 | -0.1275 | -0.2774 | -0.0457 | 0.262 |
| ICC108 | -0.0075 | 0.0873 | 0.141 | -0.1075 | 0.1918 | -0.1439 | -0.1035 | -0.1007 | -0.2889 | -0.0877 | 0.058 |
| ICC12 | 0.0908 | 0.1197 | -0.166 | -0.0286 | 0.0013 | -0.3024 | 0.034 | 0.0125 | -0.1177 | 0.0377 | 0.1918 |
| ICC137 | 0.0372 | 0.1649 | -0.0411 | -0.2378 | 0.2868 | -0.032 | -0.0125 | -0.2245 | -0.3772 | 0.1027 | 0.1135 |
| ICC15 | -0.0079 | 0.1871 | 0.1435 | -0.0582 | 0.1445 | -0.1656 | -0.0222 | -0.1701 | -0.3142 | -0.0921 | -0.0112 |
| ICC2 | -0.1453 | 0.0528 | -0.0066 | -0.047 | 0.1642 | -0.4241 | 0.2498 | -0.1005 | -0.3682 | -0.0616 | 0.1124 |
| ICC3 | 0.1318 | 0.131 | -0.0267 | 0.0113 | 0.0988 | -0.1935 | 0.0573 | -0.0683 | -0.2115 | 0.0361 | 0.227 |
| ICC4 | 0.0296 | 0.0643 | 0.0956 | 0.0003 | 0.1624 | -0.2897 | 0.0029 | 0.0341 | -0.2178 | -0.1162 | 0.0722 |
| ICC8 | 0.1519 | 0.1009 | 0.1553 | 0.0437 | -0.0176 | -0.2448 | 0.0696 | 0.0554 | -0.0647 | 0.0558 | 0.2414 |
| ICC9 | 0.0173 | 0.1091 | 0.0878 | -0.099 | 0.1525 | -0.1755 | 0.0437 | -0.0468 | -0.1771 | 0.0679 | 0.0153 |
| G415 | 0.1173 | -0.0256 | 0.1245 | 0.021 | 0.0067 | -0.2827 | 0.0262 | -0.009 | -0.0792 | -0.125 | 0.1523 |
| HKGZCC | -0.0424 | 0.1107 | 0.1376 | -0.2003 | 0.2549 | -0.2673 | 0.1045 | -0.2839 | -0.5098 | -0.1535 | -0.002 |
| KMCH1 | -0.0801 | 0.0945 | 0.1571 | -0.1287 | 0.1744 | -0.0027 | -0.032 | -0.0909 | -0.241 | -0.0955 | 0.0063 |
| RBE | -0.0117 | 0.0982 | 0.0407 | -0.0268 | 0.0777 | -0.1896 | -0.0134 | -0.1111 | -0.2288 | -0.0522 | 0.1422 |
| SG231 | 0.1875 | -0.0893 | 0.0018 | -0.0519 | 0.022 | -0.4221 | 0.1497 | 0.0249 | 0.0267 | 0.0042 | 0.0421 |
| SSP25 | -0.0315 | 0.2003 | 0.004 | -0.1647 | 0.1842 | -0.3155 | 0.0947 | -0.249 | -0.4021 | 0.0593 | 0.0537 |
| TGBC1TKB | 0.0852 | -0.0191 | 0.2741 | -0.1554 | 0.2458 | -0.1812 | -0.0778 | -0.1382 | -0.343 | -0.1468 | -0.1347 |
| TGBC52TKB | -0.0194 | 0.1698 | 0.1083 | -0.0993 | 0.155 | -0.0558 | 0.0433 | -0.2369 | -0.3489 | -0.2091 | -0.0953 |
| TKKK | -0.0835 | 0.0614 | 0.0728 | -0.0498 | 0.1854 | -0.2062 | 0.1231 | -0.0887 | -0.3141 | -0.242 | 0.0143 |
| YSCCC | -0.0373 | -0.03 | 0.0725 | 0.1256 | -0.1441 | -0.2267 | 0.0191 | -0.1215 | -0.2416 | -0.1874 | 0.1141 |
| CCLP1 | 0.0107 | 0.0526 | -0.1575 | 0.0383 | -0.0313 | -0.0478 | 0.0896 | 0.0543 | -0.0654 | 0.0782 | 0.0047 |
| CCSW1 | -0.0606 | 0.17 | -0.0166 | -0.0634 | 0.0921 | -0.3069 | 0.1472 | -0.1949 | -0.3148 | -0.0261 | 0.051 |
| GB2 | 0.0825 | 0.0981 | 0.0679 | -0.0155 | 0.0656 | -0.2018 | 0.1063 | -0.0915 | -0.2412 | -0.1942 | 0.0812 |
| MM253 | 0.0928 | -0.0362 | 0.1111 | -0.0195 | 0.1173 | -0.2903 | 0.0216 | -0.129 | -0.2263 | -0.3726 | 0.0815 |
| MM485 | 0.0019 | 0.1354 | 0.0456 | -0.185 | 0.2685 | -0.0691 | -0.1533 | -0.1305 | -0.2041 | -0.0238 | -0.1586 |
| NO36 | -0.0469 | 0.1344 | -0.0023 | 0.1291 | 0.0601 | -0.2852 | 0.1024 | 0.0401 | -0.1366 | -0.0852 | -0.0054 |
| NZM3 | -0.0388 | 0.104 | -0.0905 | 0.1911 | 0.0403 | -0.1364 | 0.0423 | 0.0735 | -0.1015 | -0.185 | 0.0021 |
| NZM42 | 0.0881 | 0.1214 | 0.1747 | 0.0398 | 0.0054 | -0.2218 | 0.1168 | -0.0309 | -0.1573 | -0.0523 | 0.0746 |
| NZM7 | -0.0534 | 0.1465 | 0.0415 | 0.041 | 0.0009 | -0.2563 | 0.0995 | -0.0306 | -0.1408 | -0.1213 | 0.037 |
| NZOV9 | 0.113 | -0.1751 | 0.1022 | -0.0125 | 0.0322 | -0.218 | -0.1069 | -0.0487 | -0.1203 | -0.0673 | 0.1273 |
| ONE58 | 0.0838 | 0.0871 | -0.0732 | 0.103 | -0.0866 | -0.0955 | 0.0481 | 0.062 | -0.0706 | 0.0957 | 0.0123 |
| NALM16 | 0.053 | -0.0748 | 0.0173 | 0.3204 | -0.1372 | -0.3052 | 0.1714 | 0.2009 | 0.0496 | -0.3173 | 0.198 |
| ECC2 | -0.0242 | 0.0737 | 0.1098 | -0.2138 | 0.2676 | -0.2144 | -0.082 | -0.2368 | -0.3933 | -0.0518 | 0.0862 |
| 9505BIK | -0.173 | 0.1857 | 0.1044 | -0.1375 | 0.2798 | -0.2375 | 0.1249 | -0.0807 | -0.3895 | -0.1554 | 0.005 |
| A375SKINCJ1 | -0.0482 | 0.0699 | 0.1077 | 0.0642 | 0.0152 | -0.1125 | 0.0932 | -0.0414 | -0.1424 | -0.2366 | 0.0644 |
| A375SKINCJ2 | 0.0852 | 0.088 | 0.1306 | -0.0418 | 0.1557 | -0.2517 | 0.0991 | -0.1215 | -0.2064 | -0.0202 | 0.0307 |
| A375SKINCJ3 | -0.1723 | -0.0423 | -0.104 | 0.0341 | -0.064 | -0.0516 | 0.0806 | 0.0543 | -0.1064 | -0.1352 | 0.1563 |
| SKMEL19 | -0.3166 | 0.0369 | 0.2488 | -0.0868 | 0.1574 | -0.0711 | 0.1312 | 0.0096 | -0.0668 | 0.0419 | -0.139 |
| MEL270 | -0.1219 | 0.2186 | 0.0259 | -0.0194 | 0.0897 | -0.1722 | 0.2327 | -0.0881 | -0.1144 | -0.0164 | -0.1998 |
| MEL285 | 0.1719 | 0.0266 | 0.0426 | 0.0539 | -0.1097 | -0.0641 | -0.0356 | 0.0016 | -0.014 | 0.0371 | 0.0615 |
| MEL290 | 0.0542 | 0.0836 | -0.034 | 0.0182 | -0.0098 | -0.0437 | -0.0296 | 0.0106 | -0.1013 | 0.1055 | 0.0413 |
| OMM25 | -0.0341 | 0.0784 | 0.063 | 0.128 | -0.0146 | -0.2269 | 0.0268 | 0.0506 | -0.0859 | -0.2252 | 0.0976 |
| HOKUG | 0.003 | 0.0441 | 0.0609 | 0.0231 | 0.1102 | -0.1107 | -0.0086 | -0.0687 | -0.2036 | -0.1366 | 0.0166 |
| SKGIIIA | -0.0003 | 0.0458 | 0.118 | 0.0442 | 0.1253 | -0.228 | 0.0817 | -0.0176 | -0.1299 | -0.2228 | 0.0772 |
| T3M3 | 0.0432 | -0.1261 | 0.1569 | 0.1237 | -0.0376 | -0.255 | -0.0607 | 0.0332 | -0.0275 | -0.3369 | 0.1845 |
| TGBC18TKB | -0.0404 | 0.0778 | 0.0126 | -0.1558 | 0.226 | -0.1707 | 0.0264 | -0.1978 | -0.3879 | -0.2031 | -0.0682 |
| ECC4 | -0.1132 | 0.0128 | -0.1174 | 0.0661 | 0.1914 | -0.2515 | 0.1555 | 0.0528 | -0.1059 | -0.1235 | 0.2837 |
| TT1TKB | 0.0024 | 0.023 | 0.1635 | -0.0032 | 0.0665 | -0.1894 | -0.1563 | -0.0911 | -0.2444 | -0.1729 | 0.0119 |
| HHUA | 0.1087 | -0.014 | 0.0574 | 0.0378 | 0.0485 | -0.2624 | 0.0316 | -0.0165 | -0.1299 | -0.1575 | 0.1381 |
| HOUAI | 0.2182 | 0.0044 | 0.1593 | 0.0498 | 0.0846 | -0.419 | 0.0878 | 0.0852 | -0.032 | -0.1524 | 0.0196 |
| SAS | 0.0032 | 0.0293 | 0.1468 | 0.0053 | 0.111 | -0.2183 | -0.0239 | -0.0865 | -0.2781 | -0.1606 | 0.0969 |
| LCAM1 | 0.0154 | 0.1651 | 0.0332 | 0.1054 | 0.0218 | -0.1159 | 0.0895 | 0.0231 | -0.0611 | 0.0113 | 0.1878 |
| PK8 | 0.1237 | 0.0416 | 0.0543 | 0.2257 | -0.0405 | -0.3074 | -0.0266 | 0.1394 | 0.0165 | 0.0709 | 0.1505 |
| HOTHC | -0.0093 | 0.054 | 0.0663 | 0.0608 | -0.0118 | -0.1328 | -0.0488 | 0.0201 | -0.0659 | -0.1571 | 0.1545 |
| T3M5 | -0.034 | -0.0758 | -0.0747 | 0.0895 | 0.2091 | -0.2756 | 0.0353 | 0.0925 | -0.1341 | -0.2306 | 0.0957 |
| CA922 | 0.0466 | -0.0407 | 0.0686 | 0.1656 | -0.044 | -0.2003 | 0.0811 | 0.0187 | -0.1268 | -0.2691 | 0.1345 |
| HSQ89 | 0.0839 | 0.0522 | 0.0501 | 0.0748 | -0.0158 | -0.0178 | -0.0536 | 0.0729 | 0.0277 | -0.1075 | 0.0232 |
| HO1U1 | 0.2116 | 0.101 | 0.0449 | 0.0421 | -0.0617 | -0.4677 | 0.1966 | -0.0245 | -0.1734 | -0.0838 | 0.2155 |
| HTMMT | 0.0183 | 0.0051 | 0.0935 | 0.1687 | -0.0939 | -0.1275 | -0.0993 | 0.0418 | -0.0727 | -0.1638 | 0.1832 |
| RMSYM | -0.0194 | 0.1947 | -0.1151 | 0.2035 | -0.0237 | -0.0604 | 0.0709 | 0.1629 | 0.0031 | 0.0223 | -0.0247 |
| LU134A | -0.0201 | 0.0135 | 0.2052 | 0.1371 | 0.0011 | -0.0292 | -0.025 | 0.0003 | -0.0138 | -0.1174 | 0.2281 |
| P30OHK | 0.1005 | 0.0628 | 0.1146 | 0.059 | 0.0038 | -0.0661 | -0.1315 | -0.0185 | -0.1636 | -0.048 | 0.175 |
| P2URK562 | 0.0647 | 0.0934 | 0.1448 | 0.1947 | -0.033 | -0.2232 | 0.0002 | 0.041 | 0.0075 | -0.0504 | 0.3 |
| SLVL | 0.1511 | 0.1817 | -0.0065 | 0.1667 | -0.1804 | -0.0828 | -0.0709 | -0.0203 | -0.078 | 0.0282 | 0.3284 |
| HSSCH2 | -0.0492 | 0.1115 | -0.0865 | 0.0472 | -0.1003 | 0.008 | 0.0545 | 0.0179 | -0.1256 | 0.0661 | -0.0737 |
| NOS1 | -0.172 | 0.1383 | -0.0195 | 0.1625 | -0.0119 | -0.0656 | 0.1173 | 0.0467 | 0.0159 | -0.0983 | 0.0311 |
| HSOS1 | -0.1231 | 0.2857 | 0.0445 | 0.1491 | 0.1036 | -0.1223 | 0.006 | 0.0884 | 0.005 | 0.1 | 0.2071 |
| HKBMM | 0.1332 | 0.0585 | 0.1522 | 0.1179 | 0.0075 | -0.1739 | -0.0117 | 0.0715 | 0.0585 | 0.0052 | 0.19 |
| LU165 | 0.0063 | 0.139 | 0.0335 | 0.0279 | 0.1204 | -0.1873 | 0.0868 | -0.0765 | -0.158 | -0.135 | -0.0819 |
| TN2 | -0.099 | 0.131 | -0.131 | 0.1741 | -0.0352 | -0.1188 | 0.1412 | 0.1377 | -0.0285 | -0.0731 | 0.0988 |
| NB69 | 0.0882 | -0.0396 | 0.0369 | -0.0142 | 0.1185 | -0.3739 | 0.0399 | -0.0203 | -0.1764 | -0.1191 | 0.2368 |
| MMAC | -0.0334 | -0.0153 | -0.0045 | 0.0888 | -0.0395 | -0.1485 | -0.0462 | 0.0311 | -0.1603 | -0.2597 | 0.4143 |
| ES4 | -0.0007 | 0.0635 | 0.1859 | -0.0125 | 0.1086 | -0.2941 | 0.0913 | -0.0708 | -0.2433 | -0.1698 | 0.079 |
| ES5 | -0.0911 | 0.1315 | -0.2195 | 0.151 | 0.0783 | -0.1613 | 0.1785 | 0.159 | 0.0067 | -0.0309 | 0.0945 |
| ES8 | 0.0531 | 0.1314 | 0.0232 | 0.1335 | -0.1521 | -0.129 | 0.0204 | 0.0108 | 0.0286 | -0.0827 | 0.3519 |
| EW1 | -0.0254 | 0.0471 | -0.1255 | 0.1933 | 0.0657 | -0.1705 | 0.1362 | 0.1537 | 0.0215 | -0.0837 | 0.0488 |
| EW16 | -0.1221 | 0.1476 | -0.1271 | 0.1561 | 0.0909 | -0.1496 | 0.1209 | 0.086 | -0.0991 | -0.1006 | 0.1791 |
| EW22 | -0.153 | 0.1169 | -0.2365 | 0.0596 | 0.1243 | -0.0616 | 0.213 | 0.0111 | -0.1014 | -0.0158 | -0.0449 |
| EW7 | 0.2075 | -0.0043 | 0.0503 | 0.0787 | 0.0305 | -0.2612 | 0.0612 | -0.028 | -0.1539 | -0.1558 | 0.21 |
| HEY | -0.0342 | -0.1212 | 0.2341 | -0.0739 | 0.045 | -0.2325 | -0.087 | -0.0898 | -0.1497 | -0.2489 | 0.2529 |
| HSC39 | -0.0027 | 0.1813 | 0.0438 | -0.282 | 0.1589 | -0.1391 | 0.0733 | -0.2667 | -0.4241 | 0.1028 | 0.0797 |
| ISTMEL1 | 0.0422 | 0.0956 | 0.0456 | -0.0221 | 0.1228 | 0.1296 | -0.1256 | -0.0244 | -0.0218 | -0.0777 | -0.129 |
| LB1047RCC | -0.0608 | 0.0654 | 0.1837 | -0.1481 | 0.125 | -0.1894 | 0.0471 | -0.2196 | -0.349 | -0.0637 | 0.0948 |
| LC1SQ | -0.0707 | 0.1592 | 0.1918 | -0.0599 | 0.2547 | -0.2345 | 0.2408 | -0.0776 | -0.2822 | -0.0823 | -0.1073 |
| MCIXC | 0.0609 | -0.1131 | -0.0921 | 0.3495 | -0.1887 | -0.3259 | 0.1254 | 0.2863 | 0.1341 | -0.2726 | 0.0657 |
| MKN28 | -0.0576 | 0.1129 | -0.1111 | 0.1491 | 0.0561 | -0.0608 | 0.149 | 0.0444 | -0.1267 | -0.1425 | 0.1129 |
| OCUBM | -0.087 | 0.1777 | 0.1205 | -0.1989 | 0.24 | -0.1059 | 0.0341 | -0.3083 | -0.3472 | 0.0609 | -0.0419 |
| OVCA420 | -0.0651 | 0.1594 | -0.0329 | 0.0271 | 0.1983 | -0.3379 | 0.203 | -0.1277 | -0.3088 | -0.0898 | 0.1366 |
| OVMIU | 0.0117 | 0.155 | 0.0253 | 0.1129 | 0.0578 | -0.3503 | 0.1744 | -0.0058 | -0.2041 | -0.1837 | 0.2293 |
| PL4 | 0.0566 | 0.037 | -0.0858 | -0.0269 | -0.0036 | -0.0853 | 0.0658 | -0.0171 | -0.0719 | -0.0114 | 0.0146 |
| RCCFG2 | 0.2134 | 0.0016 | -0.0256 | 0.1314 | -0.1374 | -0.2301 | 0.1297 | 0.105 | 0.0481 | 0.1102 | 0.0809 |
| SCH | 0.0471 | 0.1453 | -0.0615 | 0.0219 | -0.0305 | -0.0806 | 0.0227 | -0.0731 | -0.1103 | -0.0387 | 0.0238 |
| TMK1 | -0.0426 | -0.0719 | -0.0688 | -0.1029 | 0.0066 | 0.0442 | 0.04 | -0.1846 | -0.3305 | -0.2162 | -0.1708 |
| ARH77 | 0.2118 | -0.1571 | -0.0093 | -0.1065 | -0.0916 | -0.1832 | -0.0542 | -0.0857 | -0.1164 | -0.2617 | 0.0571 |
| BB30HNC | 0.0622 | -0.1086 | 0.1732 | -0.1124 | 0.0512 | -0.1115 | -0.0344 | -0.1203 | -0.2388 | -0.2011 | 0.091 |
| BE2M17 | -0.1446 | 0.1426 | -0.1272 | 0.1343 | 0.0646 | -0.125 | 0.1203 | 0.1127 | -0.0552 | -0.0475 | 0.1072 |
| D423MG | 0.1011 | 0.0694 | 0.02 | 0.0414 | -0.1173 | -0.0936 | -0.0718 | -0.0965 | -0.2205 | -0.0105 | 0.0967 |
| D502MG | -0.0101 | 0.0081 | 0.0675 | -0.0034 | 0.009 | -0.31 | 0.1126 | -0.0604 | -0.1961 | -0.0576 | 0.1169 |
| D542MG | 0.2304 | 0.0159 | 0.1284 | 0.0852 | -0.0974 | -0.2017 | -0.0397 | 0.0645 | -0.0245 | 0.1583 | 0.3807 |
| DIFI | 0.0998 | 0.1186 | -0.0602 | 0.05 | 0.0993 | -0.152 | 0.0143 | -0.1226 | -0.262 | -0.2686 | 0.1004 |
| DOK | -0.0215 | 0.1061 | 0.1227 | -0.1878 | 0.1631 | -0.1611 | 0.0198 | -0.3331 | -0.4814 | -0.0945 | 0.0018 |

| R | EC | IDRE | TM | IDRC | Linker | Tail | IDRE pos | IDRC pos | RK cluste | GxxxG | CRAC |
|---|---|---|---|---|---|---|---|---|---|---|---|
| HCE4 | -0.0193 | 0.039 | -0.0787 | 0.0276 | -0.001 | -0.067 | -0.0174 | -0.0626 | -0.1048 | -0.0199 | 0.0935 |
| IM9 | 0.2379 | 0.113 | -0.1211 | 0.1578 | -0.0499 | -0.2586 | 0.1208 | 0.1343 | 0.0115 | -0.1603 | 0.3317 |
| JHU011 | -0.0427 | 0.1206 | 0.1088 | 0.0149 | 0.1242 | -0.1074 | 0.0971 | -0.0626 | -0.2514 | -0.122 | 0.0127 |
| JHU022 | -0.0743 | 0.1102 | 0.0232 | -0.012 | 0.1187 | -0.3835 | 0.2123 | -0.0719 | -0.291 | -0.0802 | 0.0959 |
| JHU029 | -0.0639 | 0.1124 | 0.1246 | -0.0248 | 0.0879 | -0.17 | 0.0767 | -0.0954 | -0.3032 | -0.112 | 0.0717 |
| KINGS1 | -0.0748 | 0.0944 | -0.0067 | -0.0105 | -0.2394 | -0.0516 | 0.0912 | 0.0252 | -0.0146 | 0.0351 | 0.0774 |
| KPNYS | 0.1224 | 0.0498 | 0.0004 | 0.131 | 0.0097 | -0.2744 | 0.1423 | 0.2791 | 0.0628 | -0.056 | 0.1991 |
| KYSE220 | -0.1495 | 0.054 | -0.0735 | 0.0013 | 0.0575 | -0.1265 | -0.0481 | 0.0849 | -0.1592 | -0.1075 | 0.133 |
| LB771HNC | -0.0861 | 0.1146 | -0.0915 | -0.0587 | 0.1494 | -0.0723 | 0.0426 | -0.0004 | -0.1231 | -0.0017 | 0.0622 |
| LNZTA3WT4 | 0.114 | 0.0556 | -0.0374 | 0.0752 | -0.054 | -0.0521 | -0.0064 | 0.044 | -0.0214 | 0.1455 | 0.1746 |
| NB10 | 0.0069 | 0.0451 | 0.054 | 0.169 | -0.0742 | -0.2497 | 0.1612 | 0.0548 | -0.0173 | -0.2277 | 0.1698 |
| NB13 | -0.0747 | 0.1841 | -0.173 | 0.1344 | 0.0946 | -0.2612 | 0.2774 | 0.1131 | -0.0846 | -0.1367 | 0.0499 |
| NB17 | 0.0009 | 0.114 | -0.1346 | 0.1423 | -0.0092 | -0.3683 | 0.2272 | 0.2217 | -0.0182 | 0.0426 | 0.2732 |
| NB5 | -0.0692 | 0.1707 | 0.0481 | 0.1706 | 0.0409 | -0.2537 | 0.0441 | 0.1266 | -0.0147 | -0.144 | 0.1333 |
| NB6 | -0.0197 | 0.1176 | 0.1069 | 0.2029 | -0.0397 | -0.2072 | 0.0773 | 0.1457 | 0.0031 | -0.0952 | 0.3997 |
| NB7 | 0.045 | 0.0758 | -0.1229 | 0.1636 | 0.0209 | -0.3031 | 0.167 | 0.0669 | -0.0396 | -0.157 | 0.2006 |
| NTERA2CLD1 | -0.024 | 0.295 | 0.0142 | 0.075 | 0.0972 | -0.1916 | 0.2082 | -0.0187 | -0.2063 | -0.2123 | 0.0049 |
| PCI15A | -0.1259 | 0.0925 | 0.2433 | -0.0501 | 0.1379 | -0.1602 | 0.0752 | -0.0806 | -0.2423 | -0.1271 | 0.1594 |
| PCI30 | 0.1969 | -0.1209 | -0.0776 | 0.1368 | -0.2828 | -0.2008 | 0.0705 | 0.0425 | -0.0174 | -0.0477 | 0.0354 |
| PCI38 | -0.147 | 0.1913 | 0.0692 | 0.0344 | 0.1135 | -0.1959 | 0.1552 | -0.0845 | -0.2948 | 0.083 | 0.0474 |
| PCI4B | 0.0463 | 0.1426 | 0.1483 | -0.0221 | 0.0761 | -0.2528 | 0.0121 | -0.1281 | -0.2479 | -0.0463 | 0.1365 |
| PCI6A | -0.0525 | 0.1306 | 0.105 | -0.1 | 0.2001 | -0.2831 | 0.1858 | -0.2392 | -0.4066 | -0.0549 | 0.1639 |
| SKMG1 | -0.0631 | 0.0876 | 0.2273 | 0.0536 | 0.0011 | -0.2185 | 0.0877 | -0.0557 | -0.0117 | -0.044 | 0.181 |
| SKN3 | -0.1082 | 0.0953 | -0.0503 | 0.0247 | -0.0829 | -0.1262 | 0.0854 | 0.019 | -0.0637 | -0.0283 | -0.048 |
| 21NT | 0.0116 | 0.1573 | 0.3389 | -0.0815 | 0.191 | -0.0669 | -0.0299 | -0.0958 | -0.1874 | -0.0529 | 0.0664 |
| CCLFUPGI0005T | 0.07 | 0.0168 | 0.1789 | 0.032 | 0.1322 | -0.1133 | -0.2009 | -0.0308 | -0.0705 | -0.0142 | 0.1733 |
| HT144SKINFV1 | 0.154 | -0.0581 | 0.0615 | 0.1721 | 0.0403 | -0.2535 | -0.0418 | 0.1493 | 0.0353 | -0.1654 | 0.2005 |
| HT144SKINFV3 | 0.1577 | 0.0806 | 0.079 | 0.1989 | 0.0663 | -0.3125 | 0.0731 | 0.1712 | 0.0336 | -0.1282 | 0.1559 |
| HT144SKINFV2 | 0.1173 | -0.0374 | 0.1338 | 0.1045 | 0.008 | -0.4654 | 0.0726 | 0.0572 | 0.0069 | -0.1011 | 0.1799 |
| RVH421SKINFV1 | 0.0929 | 0.1682 | 0.0232 | 0.162 | -0.0603 | -0.2208 | -0.0258 | -0.0289 | -0.1212 | -0.0017 | 0.2151 |
| RPE1SS48 | 0.1675 | -0.0923 | 0.0522 | -0.0402 | -0.0668 | -0.3611 | -0.0638 | -0.1245 | -0.1801 | -0.0914 | 0.1745 |
| RPE1SS77 | 0.1372 | 0.062 | 0.2582 | -0.1715 | 0.1144 | -0.2582 | -0.0641 | -0.201 | -0.2073 | -0.0352 | 0.0721 |
| RPE1SS6 | 0.1815 | -0.1081 | 0.1372 | -0.2029 | 0.07 | -0.2264 | -0.0095 | -0.2253 | -0.1338 | -0.1613 | 0.0641 |
| RPE1SS119 | 0.1893 | -0.129 | 0.1743 | -0.1294 | 0.1289 | -0.186 | 0.0357 | -0.1053 | -0.1597 | -0.2764 | 0.0563 |
| RPE1SS51 | 0.0767 | 0.0079 | -0.0259 | 0.0255 | -0.0511 | -0.1179 | 0.1213 | -0.0987 | -0.1584 | -0.1243 | 0.1115 |
| PSS008 | -0.0639 | 0.2322 | -0.0336 | 0.1006 | 0.1621 | -0.3922 | 0.1193 | -0.0546 | -0.3046 | -0.0825 | 0.1122 |
| MAVER1 | 0.1288 | 0.1175 | 0.1116 | 0.1304 | 0.0054 | -0.3238 | 0.0754 | 0.1906 | 0.0281 | -0.0076 | 0.2351 |
| MESOV | -0.0546 | 0.0004 | -0.0192 | 0.041 | 0.077 | -0.2632 | -0.0377 | -0.0257 | -0.0389 | -0.1599 | 0.0446 |
| WM3211 | 0.1277 | -0.0048 | -0.0829 | 0.1247 | -0.1162 | -0.0727 | -0.0186 | 0.0318 | -0.029 | 0.0394 | 0.0286 |
| M040416 | -0.0561 | 0.0651 | 0.1342 | 0.1012 | -0.0807 | -0.1163 | 0.3183 | -0.0916 | -0.202 | -0.198 | 0.2761 |
| M140325 | -0.0128 | 0.132 | 0.0637 | 0.1026 | 0.0243 | -0.0455 | 0.0188 | -0.0343 | -0.0525 | -0.1192 | 0.0891 |
| MM160113 | -0.0564 | 0.2422 | -0.1261 | 0.1215 | 0.1866 | -0.2615 | 0.3596 | -0.0454 | -0.2192 | -0.1333 | 0.1638 |
| SNU739 | 0.025 | 0.1348 | -0.0221 | -0.0076 | -0.0003 | -0.0903 | -0.0743 | -0.082 | -0.1776 | -0.0035 | 0.1862 |
| SNU1327 | 0.16 | 0.0577 | 0.0832 | -0.2159 | 0.1184 | -0.3513 | -0.03 | -0.0861 | -0.2508 | -0.1807 | 0.1344 |
| SNU2535 | 0.1363 | 0.0526 | 0.1658 | 0.2037 | -0.024 | -0.2226 | -0.0316 | 0.1227 | 0.1032 | -0.1564 | 0.1198 |
| CCC5 | 0.1646 | 0.1721 | 0.0251 | 0.1325 | 0.0493 | -0.3819 | 0.0247 | 0.074 | -0.0879 | -0.0768 | 0.2848 |
| ETCC016 | 0.0758 | 0.1646 | 0.0561 | 0.074 | 0.0216 | -0.3822 | 0.137 | -0.0272 | -0.1672 | -0.0736 | 0.1425 |
| JVE015 | 0.0059 | 0.0477 | 0.1209 | 0.1433 | -0.088 | -0.004 | 0.0568 | 0.1167 | 0.0636 | -0.1235 | 0.1154 |
| JVE127 | -0.172 | 0.18 | -0.1028 | 0.1432 | 0.0941 | -0.1812 | 0.1841 | 0.0647 | -0.0444 | -0.1036 | 0.0575 |
| JVE253 | -0.1214 | 0.1805 | -0.1697 | 0.2156 | 0.0378 | -0.1 | 0.1694 | 0.0883 | -0.1042 | -0.108 | 0.1301 |
| KP363T | 0.1046 | -0.0908 | 0.101 | -0.0123 | 0.0447 | -0.2771 | -0.0419 | 0.0055 | -0.1574 | -0.1339 | 0.1684 |
| MAPACHS77 | 0.0654 | -0.0107 | 0.0175 | -0.0705 | 0.1065 | -0.2123 | -0.0585 | -0.1342 | -0.2797 | -0.1584 | 0.0734 |
| 170MGBA | 0.0856 | 0.0955 | 0.1536 | 0.1394 | -0.0396 | -0.1526 | -0.0631 | 0.0477 | 0.0343 | 0.1012 | 0.2792 |
| WM3772F | 0.203 | -0.1866 | 0.1725 | 0.1642 | -0.2415 | -0.4306 | -0.0185 | 0.0942 | 0.0597 | -0.2129 | 0.1416 |
| S462 | 0.0948 | 0.0312 | 0.0874 | 0.0494 | -0.0361 | -0.2617 | -0.0309 | -0.0395 | -0.2613 | -0.145 | 0.1502 |
| MPNST724 | 0.1297 | 0.0257 | -0.0472 | 0.1014 | -0.1463 | -0.0199 | -0.0568 | 0.048 | -0.0194 | 0.08 | -0.0093 |
| NCCLMS1C1 | 0.2141 | 0.0646 | 0.0691 | 0.1952 | -0.1039 | -0.1507 | -0.0672 | 0.1769 | 0.0417 | 0.0563 | 0.1431 |
| NCCMPNST1C1 | -0.0965 | 0.124 | -0.0066 | 0.085 | 0.0725 | -0.0035 | -0.0291 | 0.0772 | -0.0776 | -0.0284 | 0.0494 |
| NCCMPNST2C1 | -0.0573 | 0.2673 | -0.1506 | 0.1342 | 0.0125 | -0.1542 | 0.0112 | 0.0018 | -0.0585 | 0.1079 | 0.1936 |
| PSS131R | -0.0091 | 0.2262 | -0.048 | 0.1054 | 0.0624 | -0.0341 | 0.0115 | -0.0389 | -0.1259 | -0.0305 | -0.0013 |
| YUHOIN0650 | 0.0827 | -0.0102 | 0.0737 | -0.1242 | 0.1695 | -0.2626 | -0.103 | -0.1425 | -0.2517 | -0.14 | 0.1661 |
| SKNMM | -0.0601 | 0.1393 | -0.1013 | 0.2425 | 0.0172 | -0.1196 | 0.1756 | 0.1618 | -0.0049 | -0.1697 | 0.1015 |
| UPMD1 | 0.1586 | 0.0266 | -0.0478 | 0.1252 | -0.0372 | -0.2347 | -0.0409 | 0.1761 | -0.0045 | -0.0934 | 0.1225 |
