## Supplement Table S5C for "Single-molecule behavior and cell-growth regulation in human RTKs"

| R | EG1 | EG2 | EG3 | EG4 | EG5 |
|---|---|---|---|---|---|
| NIHOVCAR3 | -0.2861 | 0.1654 | -0.0799 | -0.1269 | 0.2419 |
| HEL | -0.1999 | 0.1509 | -0.3206 | -0.0451 | 0.3501 |
| HEL9217 | -0.269 | 0.0062 | -0.2273 | -0.0161 | 0.4145 |
| LS513 | -0.3444 | 0.1531 | -0.3089 | 0.1581 | 0.2051 |
| C2BBE1 | -0.6384 | 0.2921 | -0.0582 | 0.0482 | 0.1293 |
| 253J | -0.4688 | 0.2248 | -0.0011 | -0.016 | 0.0999 |
| HCC827 | -0.3026 | 0.3082 | -0.0894 | -0.0855 | 0.0735 |
| ONCODG1 | -0.4792 | 0.2435 | -0.2238 | 0.1397 | 0.1379 |
| HS294T | -0.193 | 0.1074 | -0.24 | 0.0185 | 0.238 |
| NCIH1581 | -0.1748 | 0.321 | -0.2173 | -0.0837 | 0.1032 |
| SKBR3 | -0.6318 | 0.2116 | -0.0057 | 0.0397 | 0.1623 |
| T24 | -0.2907 | 0.3126 | -0.0533 | -0.1085 | 0.0508 |
| MCF7 | -0.3258 | 0.3596 | -0.0098 | -0.256 | 0.1466 |
| NCIH1693 | -0.3265 | 0.2849 | -0.0734 | -0.105 | 0.118 |
| PATU8988S | -0.4072 | 0.2587 | -0.2509 | 0.0204 | 0.2352 |
| PATU8988T | -0.1393 | 0.2491 | -0.1785 | -0.0672 | 0.0948 |
| OPM2 | -0.1228 | 0.2662 | -0.0725 | -0.1743 | 0.0797 |
| CH157MN | -0.1711 | 0.2193 | -0.3442 | -0.1009 | 0.3485 |
| KPL1 | -0.1565 | 0.2211 | -0.0177 | -0.0726 | -0.0206 |
| HCC827GR5 | -0.2707 | 0.231 | -0.228 | -0.0017 | 0.1755 |
| PC14 | -0.5633 | 0.174 | -0.074 | 0.0447 | 0.2181 |
| NCIH1650 | -0.217 | 0.3185 | -0.0335 | -0.299 | 0.1884 |
| U343 | -0.1555 | 0.3448 | -0.0379 | -0.298 | 0.1252 |
| S117 | -0.2213 | 0.2391 | -0.1729 | -0.1203 | 0.2117 |
| SKNMC | -0.2762 | 0.161 | -0.2087 | 0.0246 | 0.2006 |
| U118MG | -0.1189 | 0.1768 | -0.1333 | -0.2607 | 0.3233 |
| RDES | -0.1507 | 0.321 | -0.1816 | -0.1948 | 0.175 |
| PANC0203 | -0.3181 | 0.2509 | -0.1982 | 0.0458 | 0.1041 |
| MV411 | -0.0514 | 0.0518 | 0.0348 | -0.2225 | 0.1939 |
| GCIY | -0.4266 | 0.265 | -0.0644 | 0.0257 | 0.0493 |
| TOV112D | -0.3736 | 0.3666 | 0.0463 | -0.2101 | 0.064 |
| A673 | -0.2793 | 0.0311 | -0.2302 | 0.1088 | 0.2607 |
| KARPAS299 | -0.4539 | 0.2529 | -0.1226 | -0.0105 | 0.1775 |
| HT1080 | -0.2378 | 0.2282 | -0.1062 | -0.0555 | 0.0947 |
| D283MED | -0.0684 | 0.2295 | -0.2751 | 0.0433 | 0.0422 |
| PANC1005 | -0.3326 | 0.2894 | -0.1701 | -0.1275 | 0.2391 |
| HS683 | -0.1748 | 0.2721 | -0.1551 | -0.1928 | 0.2109 |
| 697 | -0.315 | 0.1461 | -0.0954 | -0.0856 | 0.2497 |
| KU812 | -0.2097 | 0.3736 | -0.2193 | 0.0102 | -0.0287 |
| U87MG | -0.2196 | 0.1965 | -0.2618 | 0.0639 | 0.1377 |
| NCO2 | -0.2364 | 0.0354 | -0.2904 | 0.1876 | 0.2011 |
| MJ | -0.3476 | -0.1531 | -0.0014 | 0.1358 | 0.2307 |
| MHHNB11 | -0.0597 | 0.2592 | -0.2625 | -0.1367 | 0.1939 |
| G292CLONEA141B1 | -0.0761 | 0.1715 | -0.0306 | -0.2832 | 0.223 |
| T3M4 | -0.5531 | 0.229 | -0.06 | 0.1185 | 0.0605 |
| ACCMESO1 | -0.23 | 0.3989 | -0.111 | -0.1954 | 0.079 |
| PC3 | -0.1473 | 0.1137 | 0.0643 | -0.1788 | 0.1165 |
| NCIH2452 | -0.0663 | 0.2786 | -0.0629 | -0.3159 | 0.1779 |
| PANC0504 | -0.3398 | 0.1515 | -0.1221 | 0.0188 | 0.1714 |
| HPAFII | -0.2968 | 0.1146 | 0.0869 | -0.0295 | 0.0249 |
| D341Med | -0.1163 | 0.1607 | -0.3309 | 0.0852 | 0.1516 |
| ZR751 | -0.5845 | 0.2455 | -0.1213 | 0.1524 | 0.0881 |
| GAMG | -0.1211 | 0.318 | -0.1364 | -0.2299 | 0.1524 |
| SIMA | -0.0763 | 0.2943 | -0.136 | -0.1874 | 0.0995 |
| KE37 | -0.0059 | 0.0315 | -0.1024 | -0.032 | 0.1103 |
| CAOV4 | -0.122 | 0.2549 | -0.0422 | -0.2477 | 0.1417 |
| KP3 | -0.4733 | 0.3189 | -0.226 | 0.0792 | 0.1281 |
| HCC1187 | -0.0888 | 0.4365 | -0.0645 | -0.3515 | 0.0758 |
| OCIAML2 | -0.2235 | 0.1424 | -0.1631 | -0.0276 | 0.1971 |
| SU8686 | -0.2958 | 0.3168 | -0.213 | 0.0099 | 0.0782 |
| VCAP | -0.2535 | 0.1334 | -0.3114 | -0.0331 | 0.3801 |
| HUPT3 | -0.5231 | 0.3329 | -0.203 | 0.0167 | 0.1929 |
| CHP212 | 0.0525 | -0.1673 | -0.3022 | 0.2949 | 0.1082 |
| COV434 | -0.4245 | 0.245 | -0.1878 | -0.0095 | 0.2304 |
| OCILY19 | -0.0299 | 0.2513 | -0.2682 | -0.08 | 0.1252 |
| SLR20 | -0.1928 | 0.2504 | -0.2875 | -0.0111 | 0.1752 |
| LN319 | -0.5843 | 0.162 | 0.0192 | -0.0322 | 0.2358 |
| JHOS2 | -0.5073 | 0.3053 | -0.0409 | -0.0496 | 0.1216 |
| HS729 | -0.1706 | 0.2358 | -0.1301 | -0.1597 | 0.1827 |
| 8MGBA | -0.2837 | 0.2436 | -0.2329 | -0.0246 | 0.2017 |
| CFPAC1 | -0.6915 | 0.2656 | -0.0065 | -0.0932 | 0.2955 |
| PANC0327 | -0.3582 | 0.3649 | -0.0663 | -0.112 | 0.0593 |
| SNU308 | -0.611 | 0.2222 | -0.0632 | 0.14 | 0.0844 |
| CAL29 | -0.5587 | 0.3881 | -0.0566 | 0.0069 | 0.0254 |
| HCC2429 | -0.0092 | 0.1636 | -0.2075 | -0.1762 | 0.2453 |
| RERFGC1B | -0.152 | 0.2588 | -0.2729 | -0.0488 | 0.1675 |
| SKLMS1 | -0.2303 | 0.2079 | -0.1061 | -0.0775 | 0.1344 |
| THP1 | -0.3754 | 0.2091 | -0.2062 | 0.1032 | 0.1274 |
| T47D | -0.5092 | 0.0795 | -0.0861 | 0.0842 | 0.2454 |
| HS578T | -0.0731 | 0.3316 | -0.0624 | -0.2474 | 0.053 |
| SKNSH | -0.1599 | 0.2239 | -0.248 | -0.1051 | 0.2451 |
| HCC2935 | -0.0082 | 0.2532 | -0.2296 | -0.1861 | 0.1881 |
| JM1 | -0.2961 | 0.236 | -0.1809 | -0.0714 | 0.2174 |
| M059K | -0.3563 | 0.3332 | -0.1339 | -0.2614 | 0.3231 |
| NCIH2052 | -0.3355 | 0.2494 | -0.3312 | -0.0265 | 0.3302 |
| SW1990 | -0.4862 | 0.2438 | 0.0078 | 0.0715 | -0.0138 |
| OSRC2 | -0.2854 | 0.3507 | -0.1584 | 0.0225 | -0.031 |
| BT12 | -0.1811 | 0.1903 | -0.0538 | -0.1304 | 0.1263 |
| CORL105 | -0.5071 | 0.1005 | -0.1287 | 0.1961 | 0.1416 |
| SW579 | -0.1198 | 0.2268 | 0.0019 | -0.238 | 0.1135 |
| PANC1 | -0.3388 | 0.1377 | -0.1015 | -0.0126 | 0.1989 |
| NOMO1 | -0.0046 | 0.1848 | -0.1424 | -0.2864 | 0.2784 |
| RD | -0.1953 | 0.191 | -0.329 | 0.1134 | 0.1396 |
| CAL62 | -0.1357 | 0.3963 | -0.3018 | 0.0119 | -0.0192 |
| LOUNH91 | -0.5134 | 0.2071 | -0.0495 | -0.056 | 0.2395 |
| HS766T | -0.24 | 0.3684 | -0.3016 | -0.0127 | 0.104 |
| SCC9 | -0.6345 | 0.0451 | 0.0787 | 0.1004 | 0.1787 |
| SNU869 | -0.2504 | 0.2558 | -0.1511 | -0.0199 | 0.0808 |
| L363 | -0.2751 | 0.2748 | -0.2192 | -0.0224 | 0.1487 |
| CORL311 | -0.1133 | 0.3093 | -0.2683 | -0.0582 | 0.0975 |
| SCC25 | -0.5022 | 0.3829 | -0.0064 | -0.1615 | 0.1302 |
| RCC10RGB | -0.3032 | 0.3576 | -0.2479 | 0.0631 | 0.0181 |
| HDMYZ | -0.0735 | 0.0744 | -0.2559 | 0.1542 | 0.0583 |
| BHT101 | -0.2627 | 0.2232 | -0.119 | -0.0523 | 0.1252 |
| MFE280 | -0.5439 | 0.0752 | -0.006 | 0.0764 | 0.201 |
| SET2 | -0.2795 | -0.0761 | -0.2524 | 0.1304 | 0.3663 |
| EOL1 | -0.2668 | 0.1406 | -0.1645 | 0.0561 | 0.1358 |
| NMCG1 | -0.1362 | 0.2147 | -0.2292 | -0.0935 | 0.2071 |
| COLO320 | -0.2783 | 0.0525 | -0.1044 | 0.0552 | 0.1721 |
| LP1 | 0.0471 | 0.2447 | -0.4033 | 0.0894 | 0.0287 |
| C8166 | -0.2555 | 0.1405 | -0.0551 | -0.1997 | 0.3028 |
| DETROIT562 | -0.6052 | 0.369 | -0.12 | -0.0321 | 0.1814 |
| SNU1079 | -0.5338 | 0.1772 | 0.0365 | 0.0664 | 0.061 |
| DAOY | -0.2498 | 0.4311 | -0.0966 | -0.1924 | 0.0419 |
| CAL120 | -0.177 | 0.2415 | 0.0181 | -0.1637 | 0.0375 |
| HUPT4 | -0.4963 | 0.1841 | -0.1928 | 0.0985 | 0.2234 |
| LN382 | -0.2245 | 0.4354 | -0.135 | -0.2482 | 0.1215 |
| JHESOAD1 | -0.4816 | 0.2732 | -0.0999 | 0.0172 | 0.1219 |
| A375 | -0.1395 | 0.2596 | -0.3143 | -0.007 | 0.1535 |
| SNU398 | -0.2538 | 0.1921 | -0.1428 | 0.0437 | 0.0677 |
| ASPC1 | -0.2697 | 0.2165 | -0.3246 | 0.1627 | 0.1035 |
| HCC1937 | -0.4432 | 0.4837 | -0.0771 | -0.0613 | -0.0494 |
| SUPM2 | -0.1597 | 0.092 | -0.1126 | 0.0602 | 0.0581 |
| KPNYN | -0.0051 | 0.2063 | -0.3509 | -0.0919 | 0.2499 |
| BICR31 | -0.6119 | 0.2587 | -0.0248 | -0.0903 | 0.2656 |
| KALS1 | -0.1543 | 0.3289 | -0.293 | -0.1958 | 0.282 |
| U251MG | -0.2398 | 0.358 | -0.1629 | -0.0742 | 0.0437 |
| DEL | -0.1676 | 0.2205 | -0.1499 | -0.1122 | 0.1632 |

| R | EG1 | EG2 | EG3 | EG4 | EG5 |
|---|---|---|---|---|---|
| CAKI2 | -0.5937 | 0.2899 | -0.1306 | 0.0421 | 0.1813 |
| PANC0403 | -0.5416 | 0.2159 | -0.195 | 0.0768 | 0.2472 |
| JHOM1 | -0.1904 | 0.1952 | -0.1569 | -0.1277 | 0.2277 |
| SCC4 | -0.5479 | 0.3288 | -0.0796 | -0.0869 | 0.2046 |
| DANG | -0.4599 | 0.1073 | -0.1091 | 0.1385 | 0.1482 |
| DKMG | -0.0102 | 0.2077 | -0.0814 | -0.2346 | 0.1407 |
| SLR23 | -0.4515 | 0.4372 | -0.2059 | 0.0746 | -0.0195 |
| OCUM1 | -0.2647 | 0.2722 | -0.2741 | 0.1722 | -0.0164 |
| AU565 | -0.655 | 0.2535 | 0.0414 | 0.0266 | 0.1029 |
| CL11 | -0.4011 | 0.1315 | 0.1154 | -0.0424 | 0.0619 |
| KMRC20 | -0.458 | 0.3717 | -0.0741 | -0.0536 | 0.0607 |
| NCIH2887 | -0.0944 | 0.235 | -0.05 | -0.1571 | 0.0508 |
| LS1034 | -0.0892 | 0.1714 | -0.1652 | -0.0541 | 0.112 |
| COLO201 | -0.2484 | 0.1913 | -0.1541 | -0.0381 | 0.1672 |
| LMSU | -0.0288 | 0.2913 | -0.0386 | -0.2975 | 0.0962 |
| COV318 | 0.0531 | 0.107 | -0.0374 | -0.122 | 0.0311 |
| CORL279 | -0.0225 | 0.0811 | -0.0341 | -0.2218 | 0.2138 |
| DU4475 | -0.201 | 0.3147 | -0.0292 | -0.1518 | 0.0141 |
| KELLY | -0.1486 | 0.2192 | -0.3277 | -0.0161 | 0.2234 |
| SKNAS | -0.1047 | 0.2488 | -0.308 | 0.017 | 0.1086 |
| RERFLCAI | -0.2447 | 0.2725 | -0.3512 | -0.0181 | 0.2585 |
| UOK101 | -0.2804 | 0.256 | 0.0108 | -0.0921 | 0.0184 |
| KASUMI1 | -0.1862 | 0.143 | 0.0144 | -0.2849 | 0.2801 |
| CALU6 | -0.2873 | 0.4221 | -0.2726 | 0.0088 | 0.0282 |
| KP4 | -0.0831 | 0.266 | -0.3147 | -0.1264 | 0.2431 |
| SNU213 | -0.6137 | 0.2411 | -0.1402 | 0.1758 | 0.1046 |
| AM38 | -0.1896 | 0.2785 | -0.2539 | -0.1072 | 0.218 |
| SUDHL10 | -0.0173 | 0.1538 | -0.0568 | -0.1877 | 0.1226 |
| SLR24 | -0.5538 | 0.3686 | -0.0918 | -0.0049 | 0.09 |
| SF539 | -0.1321 | 0.3423 | 0.0618 | -0.216 | -0.0782 |
| HS852T | -0.3177 | 0.4725 | -0.238 | -0.2556 | 0.2564 |
| HCC38 | -0.3727 | 0.268 | 0.0062 | -0.1696 | 0.1571 |
| HCC1419 | -0.7155 | 0.2053 | 0.0111 | 0.093 | 0.1474 |
| COV362 | -0.1644 | 0.3717 | -0.2017 | -0.1805 | 0.1375 |
| EWS502 | -0.2919 | 0.1728 | -0.1637 | -0.0642 | 0.2526 |
| SNU840 | -0.4552 | 0.261 | -0.0622 | 0.0025 | 0.0954 |
| KP2 | -0.1916 | 0.2298 | -0.0467 | 0.0544 | -0.1184 |
| NCIH1755 | -0.2484 | 0.2323 | -0.2593 | -0.0631 | 0.2591 |
| SNU1033 | -0.0733 | 0.2405 | -0.2125 | -0.0827 | 0.1115 |
| BT549 | -0.0961 | 0.3887 | -0.1269 | -0.2319 | 0.0581 |
| NCIH209 | -0.3986 | 0.2314 | -0.1136 | -0.0175 | 0.1618 |
| OV90 | -0.16 | 0.1074 | -0.3185 | 0.1203 | 0.1821 |
| NCIH841 | -0.1772 | 0.2778 | -0.217 | -0.0526 | 0.1132 |
| KLE | -0.3668 | 0.3133 | -0.2903 | 0.067 | 0.1419 |
| NB4 | -0.3382 | 0.1909 | -0.2183 | 0.0191 | 0.2269 |
| EM2 | -0.2766 | 0.1042 | -0.1979 | -0.0068 | 0.2817 |
| OUMS23 | -0.2667 | 0.2179 | -0.2437 | -0.0309 | 0.2341 |
| SNU1077 | -0.4894 | 0.2677 | -0.1203 | -0.0032 | 0.1754 |
| SNU5 | -0.3824 | -0.0678 | 0.1798 | 0.0557 | 0.0757 |
| WM115 | -0.3825 | 0.2149 | -0.0363 | -0.0326 | 0.1071 |
| ECGI10 | -0.7634 | 0.1976 | 0.0096 | 0.1234 | 0.154 |
| PK1 | -0.6907 | 0.2666 | -0.0539 | 0.0441 | 0.189 |
| EFO21 | -0.1552 | 0.3346 | -0.3351 | 0.0387 | 0.0588 |
| IMR32 | -0.0168 | 0.0955 | -0.1124 | -0.0243 | 0.0549 |
| NCIH2122 | -0.1006 | 0.1372 | -0.2944 | 0.0683 | 0.1471 |
| SKNBE2 | -0.0737 | 0.107 | -0.3642 | 0.0143 | 0.2894 |
| KMRC3 | -0.2716 | 0.2967 | -0.3724 | 0.0016 | 0.2512 |
| KARPAS422 | -0.2208 | 0.0369 | -0.0488 | -0.1064 | 0.2739 |
| SNU886 | -0.3415 | 0.3149 | -0.1239 | -0.2258 | 0.2824 |
| TUHR14TKB | -0.4277 | 0.2284 | 0.0144 | -0.0347 | 0.0748 |
| TE10 | -0.4653 | 0.3361 | -0.2134 | -0.0287 | 0.2127 |
| MPP89 | -0.2973 | 0.2328 | -0.0843 | -0.1484 | 0.2101 |
| PSN1 | -0.1578 | 0.3104 | -0.1378 | -0.0908 | 0.0311 |
| HT144 | -0.1255 | 0.3126 | -0.1824 | -0.0375 | -0.0068 |
| 42MGBA | -0.057 | 0.2787 | -0.0345 | -0.2857 | 0.1099 |
| JHOC5 | -0.7171 | 0.3056 | -0.0015 | 0.068 | 0.0884 |
| SNU620 | -0.3376 | 0.1005 | -0.1263 | -0.0327 | 0.2824 |
| JURLMK1 | -0.2392 | 0.0732 | -0.1866 | -0.1325 | 0.4165 |
| CCFSTTG1 | -0.1715 | 0.3015 | 0.0145 | -0.2644 | 0.0892 |
| EFM19 | -0.1962 | 0.2115 | -0.1396 | -0.0712 | 0.135 |
| ISTMES2 | -0.3071 | 0.253 | -0.0597 | -0.1657 | 0.1909 |
| YAPC | -0.3785 | 0.1874 | -0.2579 | 0.0824 | 0.226 |
| DB | -0.1364 | 0.0641 | -0.1731 | -0.0728 | 0.2788 |
| MSTO211H | -0.0526 | 0.2289 | -0.2726 | 0.0898 | -0.0216 |
| OCIAML3 | -0.0202 | -0.0865 | -0.2545 | -0.0054 | 0.3602 |
| NCIH3122 | -0.1997 | 0.2068 | -0.1997 | -0.057 | 0.1864 |
| HCC461 | -0.4281 | 0.3564 | -0.0298 | -0.1452 | 0.1137 |
| SKNFI | -0.1122 | 0.1722 | -0.2939 | 0.0868 | 0.0986 |
| NCIH522 | -0.4325 | 0.1989 | -0.0213 | -0.1304 | 0.2492 |
| SNU668 | -0.3249 | 0.2835 | -0.1792 | -0.0824 | 0.1991 |
| JVM3 | -0.1429 | 0.0025 | -0.1213 | 0.0181 | 0.192 |
| RPMI7951 | -0.2977 | 0.2845 | -0.2949 | -0.0506 | 0.2607 |
| COLO678 | -0.4006 | 0.1747 | -0.0544 | -0.0002 | 0.1414 |
| HCC1428 | -0.1576 | 0.2032 | -0.1212 | -0.2458 | 0.2935 |
| CAPAN1 | -0.4889 | 0.1941 | -0.1204 | 0.1032 | 0.1308 |
| NCIH82 | -0.3215 | 0.1881 | -0.1254 | 0.0711 | 0.0682 |
| MKN45 | -0.2672 | 0.2921 | -0.1531 | -0.1492 | 0.2008 |
| MG63 | -0.1803 | 0.2531 | -0.1615 | -0.1299 | 0.1701 |
| SKHEP1 | -0.2529 | 0.2068 | -0.0418 | -0.1626 | 0.1804 |
| MOLM13 | -0.0836 | 0.1625 | -0.025 | -0.2764 | 0.2237 |
| SKMM2 | -0.1086 | 0.0891 | -0.2295 | 0.0565 | 0.1487 |
| U2OS | -0.2603 | 0.3932 | -0.257 | -0.1588 | 0.2098 |
| SUDHL4 | -0.1515 | 0.2291 | 0.0012 | -0.0915 | -0.0299 |
| SKNDZ | -0.0778 | 0.2758 | -0.1908 | -0.1762 | 0.1613 |
| NCIH226 | -0.0535 | 0.317 | -0.09 | -0.2639 | 0.1007 |
| SNU1105 | -0.0219 | 0.4151 | -0.1531 | -0.3527 | 0.1436 |
| SNU626 | -0.2017 | 0.1611 | -0.1223 | 0.057 | 0.0296 |
| RL | -0.2172 | 0.1662 | -0.0499 | -0.0965 | 0.1325 |
| HCC1143 | -0.4102 | 0.2202 | 0.0921 | -0.0873 | 0.0522 |
| G402 | -0.2907 | 0.2994 | 0.0315 | -0.1291 | 0.002 |
| SF295 | -0.1747 | 0.2636 | -0.2933 | -0.0123 | 0.1573 |
| SNU478 | -0.3844 | 0.1846 | 0.0191 | 0.0509 | -0.0094 |
| NCIH647 | -0.3406 | 0.2787 | -0.2034 | 0.1058 | 0.0296 |
| T84 | -0.2187 | 0.0898 | -0.037 | -0.1026 | 0.2037 |
| OE33 | -0.3276 | 0.2614 | -0.2696 | 0.1247 | 0.0836 |
| SKRC20 | -0.5471 | 0.2326 | 0.0418 | -0.0812 | 0.1726 |
| TF1 | -0.237 | 0.2294 | -0.0981 | -0.0447 | 0.073 |
| H4 | -0.0756 | 0.273 | -0.1783 | -0.1522 | 0.1234 |
| LUDLU1 | -0.4241 | 0.3605 | -0.1461 | -0.0258 | 0.0909 |
| MHHES1 | -0.0761 | 0.1725 | -0.221 | -0.0891 | 0.1969 |
| HLF | -0.1302 | 0.27 | -0.1013 | -0.217 | 0.157 |
| NCIH520 | -0.178 | 0.2314 | -0.1702 | -0.1481 | 0.2193 |
| J82 | -0.3295 | 0.3469 | -0.1352 | -0.149 | 0.1686 |
| TEN | -0.6149 | 0.1449 | 0.0134 | 0.0829 | 0.1509 |
| RI1 | -0.065 | 0.1123 | -0.0634 | -0.0943 | 0.0981 |
| COLO800 | -0.271 | 0.1422 | -0.1596 | -0.0592 | 0.2598 |
| BL70 | -0.0976 | 0.0395 | -0.1372 | -0.2139 | 0.3987 |
| NCIH747 | -0.3971 | 0.2721 | -0.0763 | -0.0192 | 0.0846 |
| K029AX | -0.1851 | 0.2292 | -0.3211 | -0.056 | 0.2747 |
| MEC1 | -0.2991 | 0.0788 | -0.1505 | 0.0206 | 0.244 |
| U937 | -0.3834 | 0.2699 | -0.2528 | 0.0061 | 0.2263 |
| SNU685 | -0.1926 | 0.2155 | -0.1648 | -0.1041 | 0.1904 |
| TE5 | -0.5117 | 0.2983 | -0.0875 | -0.034 | 0.1608 |
| SAOS2 | -0.2986 | 0.2773 | -0.2179 | 0.1063 | 0.0176 |
| 769P | -0.5695 | 0.3231 | -0.0802 | 0.0169 | 0.1099 |
| NCIH1944 | -0.4568 | 0.3463 | -0.1457 | -0.027 | 0.1274 |
| BICR6 | -0.5819 | 0.3487 | -0.0508 | -0.0628 | 0.1514 |
| NCIH838 | -0.1948 | 0.3099 | -0.2419 | -0.131 | 0.2044 |

| R | EG1 | EG2 | EG3 | EG4 | EG5 |
|---|---|---|---|---|---|
| SNU449 | -0.2271 | 0.2285 | -0.2411 | -0.0675 | 0.2357 |
| SW837 | -0.4389 | 0.2762 | -0.2848 | 0.0768 | 0.2097 |
| SNU475 | -0.2984 | 0.3039 | 0.0754 | -0.1935 | 0.03 |
| TC71 | -0.087 | 0.214 | -0.3031 | 0.0294 | 0.1132 |
| UACC62 | -0.2687 | 0.3054 | -0.1668 | -0.1285 | 0.1793 |
| KMS20 | -0.2444 | 0.1325 | -0.1559 | -0.1612 | 0.3617 |
| NCIN87 | -0.6896 | 0.1925 | -0.0392 | 0.0376 | 0.255 |
| TYKNU | -0.0476 | 0.0792 | -0.1293 | 0.0379 | 0.0392 |
| NCIH1694 | -0.3274 | 0.2443 | -0.1654 | -0.0507 | 0.1909 |
| CAKI1 | -0.4659 | 0.3001 | -0.1153 | 0.1233 | -0.0176 |
| NCIH1915 | -0.1407 | 0.2675 | -0.1622 | -0.097 | 0.0941 |
| EFE184 | -0.4304 | 0.2358 | -0.0846 | 0.0601 | 0.063 |
| OCIMY7 | -0.2471 | 0.0096 | -0.1078 | 0.0885 | 0.1613 |
| SW1088 | -0.1658 | 0.3277 | -0.1923 | -0.1999 | 0.1945 |
| LU65 | -0.2476 | 0.252 | -0.0846 | -0.0364 | 0.0345 |
| SH4 | -0.2403 | 0.2445 | -0.061 | -0.1227 | 0.1093 |
| RERFLCSQ1 | -0.7097 | 0.1665 | 0.0561 | 0.0445 | 0.1911 |
| LU99 | -0.3409 | 0.2168 | -0.0729 | -0.1097 | 0.2002 |
| KNS60 | -0.1063 | 0.2619 | -0.1322 | -0.1895 | 0.1499 |
| NCIH1666 | -0.4595 | 0.2927 | -0.0477 | -0.0382 | 0.0972 |
| MELHO | -0.2942 | 0.1753 | -0.0707 | -0.0561 | 0.1496 |
| TE8 | -0.5039 | 0.2815 | -0.0443 | -0.0423 | 0.1385 |
| HCC95 | -0.4511 | 0.2668 | -0.1054 | -0.0774 | 0.2188 |
| LN428 | -0.2933 | 0.2616 | -0.147 | 0.0064 | 0.0698 |
| BCPAP | -0.2656 | 0.2584 | -0.2825 | 0.0585 | 0.1325 |
| CJM | -0.3141 | 0.333 | -0.129 | -0.039 | 0.0443 |
| TUHR10TKB | -0.1262 | 0.3009 | -0.0379 | -0.2708 | 0.1198 |
| SNU8 | -0.7706 | 0.3216 | -0.0133 | 0.0469 | 0.1424 |
| SNU1196 | -0.4343 | 0.2719 | -0.1623 | -0.0367 | 0.2145 |
| NCIH460 | -0.1099 | 0.3025 | -0.2721 | -0.0329 | 0.0778 |
| CAS1 | -0.2637 | 0.3168 | -0.1217 | -0.2501 | 0.2544 |
| SNU216 | -0.7125 | 0.2079 | 0.0159 | 0.129 | 0.0979 |
| HCC56 | -0.2117 | 0.2242 | -0.263 | 0.0195 | 0.1553 |
| PK45H | -0.1919 | 0.3644 | -0.3713 | -0.0452 | 0.1822 |
| YH13 | -0.0363 | 0.2747 | -0.1326 | -0.2276 | 0.134 |
| SW1463 | -0.2181 | 0.1886 | -0.139 | 0.0954 | -0.0131 |
| LI7 | -0.0611 | 0.177 | -0.0638 | -0.195 | 0.143 |
| HSC2 | -0.5715 | 0.212 | -0.058 | 0.0814 | 0.1285 |
| RT112 | -0.0637 | 0.3252 | -0.1619 | -0.256 | 0.1622 |
| HUH1 | -0.3877 | 0.1979 | -0.1726 | 0.0641 | 0.1565 |
| JHH4 | -0.5602 | 0.1573 | 0.0169 | -0.0419 | 0.2377 |
| MALME3M | -0.2548 | 0.2582 | -0.2495 | 0.0555 | 0.096 |
| SNU387 | -0.1614 | 0.2047 | -0.2121 | -0.1041 | 0.2282 |
| KNS81 | -0.2239 | 0.4897 | -0.1753 | -0.1767 | 0.0278 |
| HUH7 | -0.1167 | 0.3201 | -0.226 | -0.0901 | 0.082 |
| NCIH2170 | -0.5177 | 0.1938 | -0.0387 | 0.0367 | 0.142 |
| SNU182 | -0.4119 | 0.3444 | -0.1405 | -0.1251 | 0.2037 |
| VMRCRCW | -0.2127 | -0.0473 | -0.0938 | 0.1037 | 0.1649 |
| GSU | -0.4135 | 0.3682 | -0.0924 | -0.155 | 0.166 |
| KU1919 | -0.3424 | 0.1407 | -0.0285 | 0.0337 | 0.0738 |
| F36P | -0.069 | 0.1768 | -0.339 | 0.0408 | 0.162 |
| TE11 | -0.6577 | 0.2992 | -0.0331 | -0.0154 | 0.1801 |
| SW1116 | -0.5886 | 0.259 | -0.0432 | 0.0779 | 0.0818 |
| SF767 | -0.6787 | 0.349 | -0.0484 | -0.0056 | 0.1484 |
| NCIH716 | -0.0812 | 0.113 | -0.0995 | -0.07 | 0.1172 |
| SNU423 | -0.1817 | 0.1735 | -0.265 | -0.067 | 0.2844 |
| TUHR4TKB | -0.4207 | 0.2833 | -0.0257 | -0.0515 | 0.074 |
| NCIH1792 | -0.3081 | 0.2525 | -0.169 | -0.0995 | 0.228 |
| EW8 | -0.3897 | 0.2345 | -0.3273 | 0.0811 | 0.2571 |
| SNU46 | -0.1824 | 0.1646 | -0.0043 | -0.2277 | 0.2114 |
| LS123 | -0.4093 | 0.0964 | -0.0162 | 0.1633 | 0.0056 |
| TCCPAN2 | -0.159 | 0.2554 | -0.2011 | -0.0558 | 0.1113 |
| BICR16 | -0.5101 | 0.2887 | -0.0875 | -0.0306 | 0.1656 |
| SNB75 | -0.3324 | 0.0906 | -0.2823 | 0.0559 | 0.3465 |
| RKN | -0.2809 | 0.4042 | -0.2987 | -0.0545 | 0.1382 |
| KE39 | -0.3884 | 0.3299 | -0.2194 | 0.0005 | 0.1422 |
| NCIH1299 | -0.2503 | 0.2962 | -0.2772 | 0.0451 | 0.0943 |
| CALU1 | -0.3451 | 0.3587 | -0.2863 | -0.0545 | 0.2133 |
| INA6 | -0.0594 | 0.1512 | -0.1324 | -0.1692 | 0.2077 |
| NCIH1092 | -0.2909 | 0.1386 | -0.2446 | 0.0399 | 0.2516 |
| CAL78 | -0.3512 | 0.369 | -0.302 | -0.0415 | 0.2083 |
| SNU410 | -0.2407 | 0.3343 | -0.3565 | -0.0458 | 0.23 |
| CAL33 | -0.6605 | 0.287 | -0.0321 | -0.0015 | 0.1777 |
| 59M | -0.2827 | 0.182 | -0.1926 | -0.1183 | 0.3262 |
| NCIH2030 | -0.4921 | 0.3003 | -0.0803 | 0.0586 | 0.0361 |
| UMUC3 | -0.3534 | 0.2584 | -0.0473 | -0.1514 | 0.1875 |
| KURAMOCHI | -0.5733 | 0.304 | -0.1116 | 0.0355 | 0.1423 |
| NCIH2171 | -0.225 | 0.2316 | -0.2231 | -0.066 | 0.2115 |
| OVISE | -0.5513 | 0.2496 | -0.1067 | 0.1169 | 0.0873 |
| ABC1 | -0.1905 | 0.4095 | -0.2983 | -0.0069 | 0.0207 |
| SNU61 | -0.4818 | 0.2437 | -0.1738 | 0.0614 | 0.1763 |
| NCIH2004RT | -0.0052 | 0.2178 | -0.0649 | -0.3252 | 0.2113 |
| BXPC3 | -0.548 | 0.2573 | -0.1248 | 0.0482 | 0.1716 |
| SNU761 | -0.535 | 0.2269 | -0.0554 | 0.0002 | 0.1773 |
| KMS34 | -0.1572 | 0.0993 | -0.2045 | -0.0406 | 0.2529 |
| HEYA8 | -0.1472 | 0.2681 | -0.1572 | -0.1168 | 0.1147 |
| OE21 | -0.6253 | 0.2877 | -0.0193 | 0.0289 | 0.1076 |
| VMCUB1 | -0.5663 | 0.3823 | -0.0792 | -0.0398 | 0.1105 |
| HSC4 | -0.4282 | 0.2186 | 0.0665 | -0.0435 | 0.0426 |
| HT1197 | -0.299 | 0.3262 | -0.1115 | 0.0137 | -0.0348 |
| BHY | -0.5741 | 0.2942 | -0.0218 | 0.0171 | 0.0833 |
| SNU1076 | -0.5411 | 0.3777 | 0.0159 | -0.0293 | -0.008 |
| IGR39 | -0.386 | 0.3762 | -0.1997 | -0.078 | 0.1618 |
| K562 | -0.1635 | 0.1252 | -0.1295 | -0.1573 | 0.2855 |
| HT29 | -0.435 | 0.2172 | -0.2034 | 0.0824 | 0.1786 |
| UACC893 | -0.5585 | 0.194 | 0.0651 | -0.0062 | 0.1123 |
| SIHA | -0.263 | 0.2479 | -0.0113 | -0.1887 | 0.1443 |
| AML193 | -0.1144 | 0.1419 | -0.1104 | -0.1422 | 0.2009 |
| A172 | -0.0945 | 0.3079 | -0.0673 | -0.2544 | 0.1032 |
| NCIH1836 | -0.2566 | 0.1507 | -0.1633 | 0.0267 | 0.1504 |
| TDOTT | -0.5052 | 0.3646 | -0.1065 | 0.0566 | 0.0088 |
| HCC78 | -0.4048 | 0.1731 | -0.0534 | -0.0519 | 0.202 |
| EBC1 | -0.0121 | 0.1959 | -0.2712 | -0.003 | 0.0866 |
| RCM1 | -0.4069 | 0.1677 | -0.167 | 0.1196 | 0.1322 |
| ST486 | -0.1431 | 0.0022 | -0.0265 | -0.0742 | 0.2 |
| YKG1 | -0.1248 | 0.3814 | -0.303 | -0.1753 | 0.1976 |
| T98G | -0.0409 | 0.2531 | -0.0172 | -0.3354 | 0.1628 |
| MDAMB436 | -0.1815 | 0.2452 | -0.0921 | -0.089 | 0.064 |
| KMS27 | -0.2144 | 0.1947 | -0.3093 | -0.0366 | 0.2951 |
| JHH2 | -0.2602 | 0.2667 | -0.0086 | -0.2786 | 0.2207 |
| UACC257 | -0.1849 | 0.2806 | -0.1613 | 0.1201 | -0.1319 |
| C32 | -0.2848 | 0.2416 | -0.0597 | 0.0549 | -0.057 |
| SNU16 | 0.0347 | 0.093 | -0.0145 | -0.2983 | 0.2298 |
| JHOS4 | -0.3861 | 0.2756 | 0.0162 | -0.0736 | 0.0416 |
| EPLC272H | -0.3733 | 0.4199 | -0.1142 | -0.1541 | 0.1087 |
| NCIH1975 | -0.6198 | 0.1355 | 0.0288 | 0.1018 | 0.1271 |
| KMS26 | -0.3333 | 0.1645 | -0.2214 | 0.0019 | 0.2723 |
| NCIH1437 | -0.1124 | 0.1621 | -0.2815 | -0.0484 | 0.2464 |
| LN235 | -0.1753 | 0.3942 | -0.0485 | -0.2926 | 0.0932 |
| TM31 | -0.2674 | 0.2681 | -0.1823 | -0.0278 | 0.1195 |
| BC3C | -0.3714 | 0.4062 | -0.1009 | -0.1549 | 0.1089 |
| LN229 | -0.6418 | 0.228 | -0.0293 | 0.0715 | 0.1408 |
| LCLC97TM1 | -0.2771 | 0.2121 | -0.2273 | -0.0226 | 0.2211 |
| TTC709 | -0.1943 | 0.0203 | -0.006 | -0.1568 | 0.2864 |
| PATU8902 | -0.1354 | 0.183 | -0.3122 | 0.0727 | 0.1368 |
| SLR26 | -0.2393 | 0.375 | -0.1432 | -0.0368 | -0.0348 |
| MIAPACA2 | -0.1569 | 0.3652 | -0.3182 | -0.0821 | 0.1465 |
| M07E | -0.2853 | 0.0579 | -0.2121 | 0 | 0.3403 |

| R | EG1 | EG2 | EG3 | EG4 | EG5 |
|---|---|---|---|---|---|
| TE6 | -0.7215 | 0.1798 | -0.0416 | 0.0731 | 0.2514 |
| PECAPJ34CLONEC12 | -0.587 | 0.1881 | 0.0129 | 0.032 | 0.1465 |
| KYM1 | -0.1322 | 0.2399 | -0.109 | -0.237 | 0.2182 |
| COV644 | -0.4807 | 0.2638 | -0.1388 | 0.0237 | 0.1623 |
| SF126 | -0.1541 | 0.271 | -0.0734 | -0.2843 | 0.2184 |
| RVH421 | -0.3106 | 0.3048 | -0.138 | -0.0218 | 0.0601 |
| HS746T | -0.2659 | 0.3387 | -0.2686 | -0.0514 | 0.1604 |
| SNU1041 | -0.5175 | 0.1846 | -0.149 | -0.0076 | 0.3105 |
| PECAPJ15 | -0.6139 | 0.3312 | -0.0094 | -0.0856 | 0.1738 |
| JHH1 | -0.5389 | 0.2487 | -0.0401 | 0.0101 | 0.1319 |
| MDAMB157 | -0.1193 | 0.2585 | -0.0843 | -0.2233 | 0.1514 |
| KNS42 | -0.2262 | 0.3361 | -0.2069 | -0.2026 | 0.2432 |
| SNU201 | -0.2319 | 0.2261 | -0.1337 | -0.0918 | 0.1607 |
| HCC1806 | -0.5331 | 0.4439 | -0.1382 | 0.0109 | 0.0301 |
| LCLC103H | -0.3581 | 0.242 | -0.1803 | -0.1155 | 0.3 |
| YD8 | -0.5774 | 0.2064 | 0.0175 | -0.0066 | 0.1602 |
| HS944T | -0.2188 | 0.1781 | -0.2442 | -0.0351 | 0.2479 |
| FU97 | -0.0431 | 0.2263 | 0.0028 | -0.3056 | 0.138 |
| LN340 | -0.1488 | 0.1847 | -0.1951 | 0.0153 | 0.0905 |
| KYSE520 | -0.3688 | 0.3586 | -0.0636 | -0.0839 | 0.0387 |
| NCIH441 | -0.3026 | 0.2105 | -0.2279 | 0.0411 | 0.1693 |
| NCIH211 | -0.2965 | 0.2628 | -0.0804 | -0.1428 | 0.1695 |
| JL1 | -0.3266 | 0.1898 | -0.0491 | -0.2118 | 0.3074 |
| OVMANA | -0.6235 | 0.168 | 0.0941 | 0.1052 | 0.028 |
| TE1 | -0.3999 | 0.2642 | -0.2096 | 0.0211 | 0.1829 |
| NCIH28 | -0.4302 | 0.4018 | -0.3399 | 0.0735 | 0.1373 |
| 786O | -0.2716 | 0.361 | -0.2265 | -0.1043 | 0.1583 |
| SW620 | -0.1579 | 0.2076 | -0.3116 | 0.0417 | 0.1608 |
| SUIT2 | -0.3068 | 0.3835 | -0.2733 | -0.0806 | 0.1793 |
| JJN3 | -0.1931 | 0.252 | -0.1871 | 0.0307 | 0.0271 |
| RAJI | -0.1738 | 0.1519 | 0.0174 | -0.0739 | 0.0261 |
| SF268 | -0.4511 | 0.3112 | -0.1257 | 0.0313 | 0.0741 |
| SUDHL8 | -0.0196 | 0.2867 | -0.1672 | -0.0224 | -0.0818 |
| A2780 | -0.0741 | 0.2768 | -0.0357 | -0.1499 | -0.0264 |
| KMS18 | -0.1217 | 0.1372 | -0.0995 | -0.136 | 0.1926 |
| SUDHL5 | -0.0937 | 0.2607 | -0.0946 | -0.302 | 0.23 |
| WM1799 | -0.4888 | 0.2647 | -0.2422 | 0.1044 | 0.1805 |
| CORL23 | -0.1503 | -0.0367 | 0.0456 | -0.0217 | 0.1132 |
| OVTOKO | -0.4696 | 0.1945 | -0.1038 | 0.054 | 0.1558 |
| SUDHL1 | -0.3383 | 0.1263 | -0.1579 | -0.0531 | 0.3112 |
| SKMES1 | -0.614 | 0.2212 | -0.0561 | -0.0014 | 0.237 |
| NCIH1355 | -0.2092 | 0.0669 | -0.0099 | -0.1563 | 0.2528 |
| HCC44 | -0.2208 | 0.2789 | -0.2368 | -0.0101 | 0.1132 |
| HCC70 | -0.1361 | 0.2857 | 0.0225 | -0.253 | 0.0612 |
| HUH6 | -0.1601 | 0.2489 | -0.2896 | -0.0976 | 0.2535 |
| LN443 | -0.1182 | 0.2566 | 0.0496 | -0.3277 | 0.1345 |
| LN464 | -0.2849 | 0.097 | -0.1154 | 0.0465 | 0.1527 |
| SW1573 | -0.3474 | 0.3822 | -0.2025 | -0.115 | 0.1745 |
| SW948 | -0.1932 | 0.1553 | -0.1312 | 0.1594 | -0.0748 |
| A549 | -0.5151 | 0.2804 | -0.057 | 0.0083 | 0.1035 |
| SNU1066 | -0.6722 | 0.317 | 0.0343 | 0.0163 | 0.0693 |
| SNU503 | -0.5203 | 0.2917 | -0.0415 | 0.037 | 0.0482 |
| KMRC1 | -0.0961 | 0.391 | -0.1267 | -0.2234 | 0.0462 |
| L33 | -0.5563 | 0.2304 | -0.033 | 0.0111 | 0.1533 |
| OV7 | -0.1627 | 0.2642 | -0.199 | -0.14 | 0.1963 |
| KYSE180 | -0.5987 | 0.3195 | -0.0169 | 0.0204 | 0.0654 |
| TE9 | -0.5508 | 0.3853 | -0.0709 | -0.0122 | 0.0586 |
| CORL47 | -0.2515 | 0.1811 | -0.0928 | -0.0985 | 0.1851 |
| OVCAR8 | -0.4445 | 0.3523 | -0.2065 | 0.0013 | 0.1428 |
| A3KAW | -0.2018 | 0.1385 | -0.1378 | -0.1009 | 0.2429 |
| DMS53 | -0.1429 | 0.2014 | -0.0366 | -0.1557 | 0.1011 |
| HCC1395 | -0.141 | 0.3895 | -0.2394 | -0.0738 | 0.0238 |
| NCIH2882 | -0.2679 | 0.1809 | -0.2001 | -0.0301 | 0.2274 |
| RMUGS | -0.3178 | 0.3226 | -0.254 | 0.0441 | 0.0899 |
| L1236 | -0.1897 | 0.1761 | -0.0667 | 0.068 | -0.0611 |
| OAW42 | -0.1914 | 0.2124 | -0.0943 | -0.1293 | 0.1502 |
| EKVX | -0.5369 | 0.2617 | -0.1864 | 0.0617 | 0.2065 |
| KMRC2 | -0.3984 | 0.2454 | -0.1664 | -0.0012 | 0.1822 |
| JIMT1 | -0.5433 | 0.3397 | -0.0765 | 0.0024 | 0.0886 |
| CAOV3 | -0.1074 | 0.3326 | -0.0861 | -0.2576 | 0.1094 |
| KMS11 | -0.036 | 0.184 | -0.1591 | -0.118 | 0.1295 |
| TT2609C02 | -0.3371 | 0.1348 | -0.1017 | 0.0246 | 0.1595 |
| COLO680N | -0.3029 | 0.3463 | -0.1976 | -0.1309 | 0.1941 |
| NCIH2291 | -0.2672 | 0.4508 | -0.2853 | -0.0161 | 0.0267 |
| RMGI | -0.6391 | 0.2354 | -0.0436 | 0.0468 | 0.1732 |
| TCCSUP | -0.1444 | 0.2702 | 0.0056 | -0.2592 | 0.106 |
| HMC18 | -0.1201 | 0.2114 | -0.1491 | -0.1215 | 0.1509 |
| SNUC1 | -0.1281 | 0.1942 | -0.2607 | -0.0084 | 0.1593 |
| HT1376 | -0.5244 | 0.3511 | -0.1821 | -0.0379 | 0.2152 |
| HCC202 | -0.422 | 0.3109 | 0.0148 | -0.057 | 0.0129 |
| PECAPJ41CLONED2 | -0.374 | 0.1628 | -0.0372 | -0.0834 | 0.211 |
| JHH5 | -0.4166 | 0.2577 | -0.2858 | 0.0813 | 0.2097 |
| PECAPJ49 | -0.5759 | 0.3014 | -0.1482 | 0.1048 | 0.1063 |
| SNU601 | -0.3243 | 0.2873 | -0.2302 | -0.0337 | 0.1919 |
| GB1 | -0.1336 | 0.228 | -0.2312 | -0.084 | 0.1835 |
| HEPG2 | -0.1196 | 0.3388 | -0.3051 | -0.028 | 0.0754 |
| A253 | -0.6762 | 0.3157 | 0.0194 | 0.0302 | 0.0725 |
| UBLC1 | -0.4848 | 0.3021 | -0.0738 | 0.0596 | 0.0218 |
| GSS | -0.2714 | 0.2616 | -0.2128 | -0.1148 | 0.2556 |
| NCIH1703 | -0.2927 | 0.2891 | -0.1536 | -0.1279 | 0.1974 |
| SJSA1 | -0.1349 | 0.1607 | -0.3375 | 0.0362 | 0.2246 |
| JMSU1 | -0.1438 | 0.2372 | -0.01 | -0.1941 | 0.0819 |
| L428 | -0.2843 | 0.0923 | -0.0589 | 0.0701 | 0.0744 |
| GI1 | -0.1589 | 0.1982 | -0.1757 | -0.1334 | 0.2292 |
| A427 | -0.1266 | 0.2028 | -0.0186 | -0.0819 | -0.0107 |
| MKN74 | -0.297 | 0.2496 | -0.2553 | 0.0348 | 0.1609 |
| LNZ308 | -0.2682 | 0.3035 | -0.1673 | -0.1508 | 0.2061 |
| YD38 | -0.485 | 0.2368 | 0.0034 | -0.0423 | 0.1233 |
| MM1S | -0.1938 | 0.1471 | -0.3425 | 0.0527 | 0.2634 |
| SH10TC | -0.3378 | 0.2385 | -0.099 | -0.028 | 0.1119 |
| WM983B | -0.2429 | 0.1564 | -0.2129 | -0.1136 | 0.3409 |
| NCIH1648 | -0.2728 | 0.0921 | -0.144 | -0.0043 | 0.2346 |
| NCIH526 | -0.056 | 0.1643 | -0.1988 | -0.0718 | 0.1507 |
| MDAMB231 | -0.1693 | 0.4316 | -0.2228 | -0.2123 | 0.1371 |
| LK2 | -0.0814 | 0.2678 | -0.1389 | -0.1449 | 0.0849 |
| P31FUJ | -0.2503 | 0.0869 | -0.1801 | -0.0162 | 0.2744 |
| BICR56 | -0.6899 | 0.4035 | -0.0444 | -0.0087 | 0.1007 |
| KIJK | -0.2644 | 0.0866 | -0.0659 | 0.0235 | 0.1257 |
| RERFLCAD2 | -0.1249 | 0.2824 | -0.2061 | -0.049 | 0.0595 |
| NCIH727 | -0.3806 | 0.1319 | -0.1759 | 0.1056 | 0.1751 |
| ONS76 | -0.1586 | 0.26 | -0.1122 | -0.1716 | 0.1461 |
| KYSE30 | -0.6159 | 0.2323 | 0.0593 | 0.056 | 0.0481 |
| HSC3 | -0.4419 | 0.2221 | -0.1703 | -0.0015 | 0.2382 |
| NCIH2023 | -0.2963 | 0.2106 | -0.2159 | -0.0563 | 0.2611 |
| SEM | -0.0178 | 0.1477 | -0.2244 | -0.1231 | 0.2248 |
| CAMA1 | -0.236 | 0.2248 | -0.0435 | -0.1536 | 0.1431 |
| KYSE70 | -0.2588 | 0.1784 | -0.1406 | -0.0417 | 0.1773 |
| NCIH2126 | -0.4325 | 0.2353 | -0.1559 | -0.0159 | 0.2204 |
| LXF289 | -0.4024 | 0.3097 | -0.1178 | -0.1182 | 0.2018 |
| A2058 | -0.4272 | 0.3131 | -0.2007 | 0.0371 | 0.1252 |
| SHP77 | -0.0797 | 0.258 | -0.1257 | -0.2966 | 0.2486 |
| RERFLCAD1 | -0.269 | 0.2764 | -0.2577 | -0.0682 | 0.2325 |
| BFTC909 | -0.3499 | 0.3947 | -0.1023 | -0.1749 | 0.1299 |
| KATOIII | -0.1299 | 0.1364 | 0.0396 | -0.1909 | 0.1206 |
| BICR22 | -0.6749 | 0.3188 | 0.0873 | -0.0841 | 0.1276 |
| MCAS | -0.5606 | 0.3501 | -0.2118 | 0.0342 | 0.1895 |
| HS695T | -0.3839 | 0.3214 | -0.1289 | 0.0764 | -0.0268 |
| NCIH446 | -0.1207 | 0.1112 | -0.1386 | -0.0678 | 0.1813 |

| R | EG1 | EG2 | EG3 | EG4 | EG5 |
|---|---|---|---|---|---|
| BFTC905 | -0.2149 | 0.1702 | -0.2016 | 0.0175 | 0.1521 |
| NB1 | -0.0934 | 0.1654 | -0.2465 | 0.0626 | 0.0725 |
| COLO679 | -0.4848 | 0.3273 | -0.0938 | -0.1498 | 0.249 |
| SNU738 | -0.3075 | 0.1902 | -0.0545 | -0.1091 | 0.186 |
| KYSE410 | -0.6878 | 0.1463 | 0.0337 | 0.1153 | 0.1409 |
| SKMEL30 | -0.2643 | -0.1667 | 0.0134 | 0.1818 | 0.1241 |
| SKOV3 | -0.4719 | 0.3003 | -0.1311 | 0.0395 | 0.0949 |
| RPMI8226 | -0.2276 | 0.1549 | -0.2005 | -0.0013 | 0.1956 |
| LN18 | -0.2561 | 0.3279 | -0.2053 | -0.1298 | 0.1884 |
| SW403 | -0.0454 | 0.0931 | -0.1937 | 0.0521 | 0.0725 |
| EJM | -0.2052 | 0.3237 | -0.3187 | 0.0189 | 0.1081 |
| SKMEL24 | -0.4873 | 0.1617 | -0.0187 | -0.0519 | 0.2326 |
| KYSE140 | -0.6521 | 0.3942 | 0.0463 | -0.1467 | 0.1477 |
| KYSE510 | -0.5537 | 0.3365 | -0.0737 | -0.0095 | 0.109 |
| WM793 | -0.2805 | 0.2823 | -0.1479 | 0.0111 | 0.0362 |
| HUNS1 | -0.2115 | 0.2593 | -0.1333 | -0.1673 | 0.1974 |
| HEC50B | -0.2988 | 0.2931 | -0.2042 | -0.0062 | 0.113 |
| CAL27 | -0.7243 | 0.2086 | -0.004 | 0.0848 | 0.1739 |
| RH30 | -0.2212 | 0.242 | -0.123 | -0.0717 | 0.1048 |
| UMUC1 | -0.1113 | 0.2617 | -0.261 | -0.0621 | 0.1409 |
| GCT | -0.2555 | 0.2969 | -0.3234 | 0.0401 | 0.1487 |
| YD15 | -0.5729 | 0.3049 | -0.1231 | 0.0958 | 0.0857 |
| NCIH322 | -0.5176 | 0.3024 | -0.2296 | 0.0228 | 0.2397 |
| AMO1 | -0.2863 | 0.12 | -0.3421 | 0.1649 | 0.226 |
| SCABER | -0.6681 | 0.2645 | -0.0725 | 0.0283 | 0.2126 |
| HCC366 | -0.2606 | 0.2419 | -0.1475 | -0.1192 | 0.2078 |
| NCIH2087 | -0.2088 | 0.2385 | -0.2263 | -0.1124 | 0.2487 |
| HARA | -0.6557 | 0.2107 | -0.0206 | 0.0419 | 0.1912 |
| NCIH1373 | -0.1439 | 0.0561 | -0.1626 | 0.0803 | 0.1113 |
| FADU | -0.4142 | 0.3849 | 0.0815 | -0.0601 | -0.1295 |
| HGC27 | -0.2945 | 0.38 | -0.0549 | -0.3319 | 0.2353 |
| JHH7 | -0.1814 | 0.3073 | -0.1648 | -0.0869 | 0.0722 |
| MDAMB468 | -0.2746 | 0.3168 | -0.0682 | -0.1944 | 0.1462 |
| MORCPR | -0.1282 | 0.1393 | -0.2963 | 0.1252 | 0.1018 |
| NCIH661 | -0.6391 | 0.2563 | -0.0247 | 0.0833 | 0.0928 |
| OCIMY5 | -0.2223 | 0.2639 | -0.3162 | 0.0608 | 0.13 |
| KYSE150 | -0.3488 | 0.3039 | -0.1142 | -0.0931 | 0.1411 |
| CAL51 | -0.3142 | 0.2096 | 0.0142 | -0.1762 | 0.1766 |
| KNS62 | -0.3795 | 0.2505 | 0.0235 | -0.1227 | 0.1097 |
| HCC1954 | -0.2892 | 0.267 | 0.0227 | -0.3134 | 0.2466 |
| NCIH358 | -0.3339 | 0.2244 | -0.173 | 0.0421 | 0.1199 |
| HOP62 | -0.3198 | 0.3944 | -0.1396 | -0.1052 | 0.0705 |
| KMBC2 | -0.6065 | 0.3703 | -0.085 | 0.0158 | 0.0929 |
| DBTRG05MG | 0.0176 | 0.3888 | -0.3143 | -0.0762 | -0.0015 |
| COLO684 | -0.0843 | 0.1118 | -0.0712 | -0.0731 | 0.0954 |
| KYSE450 | -0.6338 | 0.2441 | 0.0167 | 0.0691 | 0.0761 |
| NCIH1048 | -0.1666 | 0.2918 | -0.0225 | -0.2335 | 0.0984 |
| CHAGOK1 | -0.3367 | 0.2423 | -0.1222 | -0.0331 | 0.1363 |
| NCIH1568 | -0.2209 | 0.2555 | -0.1965 | 0.0412 | 0.0394 |
| NCIH510 | -0.2403 | 0.2119 | -0.0471 | -0.2069 | 0.2214 |
| HCC515 | -0.2623 | 0.1907 | -0.1109 | -0.1149 | 0.2188 |
| KYSE270 | -0.6106 | 0.2878 | -0.004 | -0.0462 | 0.1658 |
| MDAMB415 | -0.0311 | 0.3133 | 0.0178 | -0.4273 | 0.1631 |
| HCC15 | -0.0236 | 0.1876 | -0.2727 | -0.1444 | 0.2607 |
| MFE296 | -0.073 | 0.1299 | -0.0826 | -0.1658 | 0.1843 |
| AGS | -0.265 | 0.2276 | -0.0886 | -0.0773 | 0.1198 |
| MELJUSO | -0.0536 | 0.2781 | -0.3198 | -0.0627 | 0.1461 |
| IGR1 | -0.0209 | 0.1554 | -0.2237 | -0.0573 | 0.1456 |
| MDAMB435S | -0.159 | 0.18 | -0.2472 | 0.0242 | 0.1442 |
| TOV21G | -0.3077 | 0.2586 | -0.0289 | -0.0095 | -0.0183 |
| NCIH2009 | -0.1781 | 0.154 | -0.171 | 0.0449 | 0.0833 |
| SF172 | -0.4797 | 0.2995 | -0.1569 | -0.024 | 0.1969 |
| NCIH1793 | -0.4293 | 0.2624 | -0.0739 | -0.0461 | 0.1427 |
| KMM1 | -0.28 | 0.2792 | -0.1114 | -0.0481 | 0.0683 |
| SW1271 | -0.2381 | 0.4261 | -0.1874 | -0.1614 | 0.0956 |
| NCIH1869 | -0.3881 | 0.211 | -0.1122 | -0.0404 | 0.1992 |
| 647V | -0.4716 | 0.4796 | -0.1658 | -0.0388 | 0.037 |
| SNU719 | -0.4301 | 0.2038 | -0.2238 | 0.04 | 0.2563 |
| WM88 | -0.2102 | 0.1859 | -0.1238 | 0.1599 | -0.1023 |
| NCIH23 | -0.1363 | 0.1527 | -0.2064 | 0.0042 | 0.138 |
| HCC1359 | -0.0462 | 0.225 | -0.0955 | -0.2191 | 0.1437 |
| FTC133 | -0.3314 | 0.2793 | -0.1178 | -0.1249 | 0.1932 |
| 5637 | -0.422 | 0.3375 | -0.1519 | -0.0411 | 0.1353 |
| ES2 | -0.3254 | 0.2481 | -0.2027 | 0.0144 | 0.151 |
| SNU349 | -0.0834 | 0.3737 | -0.2364 | -0.2515 | 0.1961 |
| MDAMB453 | -0.4845 | 0.1767 | 0.0383 | -0.0487 | 0.1552 |
| NUGC3 | -0.5369 | 0.1426 | -0.0085 | 0.0586 | 0.1512 |
| NCIH2286 | -0.1989 | 0.2772 | -0.2789 | -0.0854 | 0.2262 |
| ESS1 | -0.2676 | 0.3885 | -0.2687 | -0.1011 | 0.1669 |
| HT | -0.2657 | 0.1639 | -0.1584 | -0.0553 | 0.2292 |
| IPC298 | -0.2751 | 0.2991 | -0.3656 | 0.0215 | 0.2221 |
| NCIH1573 | 0.0186 | 0.0847 | -0.0201 | 0.1845 | -0.2815 |
| TE4 | -0.5431 | 0.1431 | -0.0213 | 0.0807 | 0.143 |
| IM95 | -0.5794 | 0.1992 | -0.0871 | 0.0823 | 0.1746 |
| CMLT1 | -0.1997 | 0.2045 | -0.09 | -0.1343 | 0.1648 |
| NCIH157DM | -0.3192 | 0.4327 | -0.1902 | -0.0419 | 0.0122 |
| RCHACV | -0.2155 | 0.0156 | -0.0888 | -0.0029 | 0.2171 |
| NCIH2172 | -0.1634 | 0.194 | -0.2094 | -0.1208 | 0.2559 |
| HT55 | -0.2818 | 0.3459 | -0.3223 | 0.0279 | 0.1294 |
| JHUEM1 | -0.3334 | 0.2284 | -0.0274 | -0.1028 | 0.1306 |
| NCIH2110 | -0.2793 | 0.195 | -0.21 | 0.0578 | 0.1331 |
| SNU1 | -0.1284 | 0.2842 | -0.1607 | -0.1818 | 0.1619 |
| MDAMB361 | -0.6836 | 0.1233 | 0.0906 | 0.0805 | 0.1428 |
| MDST8 | -0.0611 | 0.1898 | -0.2802 | -0.0365 | 0.1707 |
| EFO27 | -0.543 | 0.1691 | -0.0024 | 0.0585 | 0.1227 |
| PF382 | -0.2494 | 0.1087 | -0.0089 | -0.0675 | 0.1378 |
| NALM6 | -0.3271 | 0.0843 | -0.0125 | -0.0403 | 0.1864 |
| SKUT1 | -0.3824 | 0.242 | -0.0637 | -0.0688 | 0.1476 |
| HEC1B | -0.623 | 0.3309 | -0.0949 | 0.112 | 0.0462 |
| RKO | -0.2955 | 0.2238 | -0.2505 | 0.0769 | 0.1342 |
| NAMALWA | -0.2692 | 0.1558 | -0.0353 | -0.1236 | 0.1923 |
| NCIH650 | -0.0984 | 0.1867 | -0.3264 | -0.0253 | 0.2319 |
| HEC265 | -0.405 | 0.231 | -0.0104 | 0.0224 | 0.0188 |
| OVK18 | -0.1271 | 0.1819 | -0.1487 | -0.0483 | 0.1032 |
| 2313287 | -0.4455 | 0.3053 | 0.0495 | -0.1426 | 0.0941 |
| LOVO | -0.2479 | 0.2324 | -0.2153 | 0.0249 | 0.1169 |
| SUPT1 | -0.2299 | 0.1229 | -0.0289 | -0.1544 | 0.2272 |
| 22RV1 | -0.1215 | 0.1522 | -0.3231 | 0.0461 | 0.199 |
| LS180 | -0.3858 | 0.3098 | -0.088 | -0.0023 | 0.0325 |
| SW48 | -0.3639 | 0.2122 | -0.1013 | -0.0506 | 0.1827 |
| SNUC4 | -0.6444 | 0.2636 | -0.0021 | 0.0326 | 0.1228 |
| REH | -0.2234 | 0.1194 | -0.2138 | 0.012 | 0.2269 |
| ISHIKAWAHERAKLIO02ER | -0.3212 | 0.1982 | 0.0139 | -0.0966 | 0.1047 |
| OC314 | -0.2866 | 0.2227 | -0.0424 | -0.1484 | 0.1713 |
| CCK81 | -0.2546 | 0.2701 | -0.3025 | -0.0037 | 0.2026 |
| RL952 | -0.6674 | 0.3075 | -0.0276 | -0.0267 | 0.1852 |
| IGROV1 | -0.2168 | -0.0207 | 0.0033 | -0.0281 | 0.19 |
| COLO792 | -0.2445 | 0.1674 | -0.2 | 0.0663 | 0.1185 |
| KM12 | -0.1176 | 0.2537 | -0.1442 | -0.1398 | 0.1223 |
| SNUC5 | -0.2863 | 0.243 | 0.0037 | -0.1307 | 0.0851 |
| HCT116 | -0.0741 | 0.2027 | -0.2733 | 0.1034 | 0.0043 |
| 639V | -0.3816 | 0.2449 | -0.2031 | 0.0132 | 0.1926 |
| SNGM | -0.3599 | 0.2728 | -0.013 | -0.0854 | 0.0697 |
| HCC2450 | -0.4287 | 0.2946 | -0.1239 | 0.0314 | 0.0741 |
| HUCCT1 | -0.5444 | 0.2593 | -0.0548 | 0.0243 | 0.1237 |
| LNCAPCLONEFGC | -0.3037 | 0.311 | -0.0379 | -0.0527 | -0.0165 |
| EN | -0.3402 | 0.1609 | -0.0789 | -0.0659 | 0.2131 |
| DU145 | -0.3176 | 0.292 | -0.16 | -0.0637 | 0.146 |

| R | EG1 | EG2 | EG3 | EG4 | EG5 |
|---|---|---|---|---|---|
| KCL22 | -0.2827 | 0.3047 | -0.145 | -0.1649 | 0.2076 |
| HEC6 | -0.2553 | 0.3149 | 0.0341 | -0.1361 | -0.0315 |
| LS411N | -0.0122 | 0.1675 | -0.1535 | 0.0553 | -0.0674 |
| HT115 | -0.4996 | 0.2586 | -0.111 | -0.0831 | 0.2706 |
| MFE319 | -0.5412 | 0.3222 | -0.1311 | 0.0223 | 0.1372 |
| SNU81 | -0.3007 | 0.2727 | -0.1527 | -0.0027 | 0.0791 |
| JHUEM7 | -0.3479 | 0.3409 | -0.095 | -0.1217 | 0.1161 |
| HEC59 | -0.2901 | 0.3087 | -0.3239 | 0.0029 | 0.2012 |
| JURKAT | -0.1118 | 0.1177 | -0.1613 | 0.011 | 0.1043 |
| HEC251 | -0.2523 | 0.1386 | -0.1061 | 0.0891 | 0.0333 |
| HCT15 | -0.0937 | 0.2699 | -0.0604 | -0.093 | -0.0452 |
| 143B | -0.3161 | 0.3242 | -0.1997 | -0.1722 | 0.2729 |
| BECKER | -0.0843 | 0.213 | -0.3097 | -0.0492 | 0.2063 |
| BT16 | -0.351 | 0.0398 | 0.1426 | -0.0767 | 0.1318 |
| CHLA06ATRT | -0.2055 | 0.2286 | -0.254 | -0.0376 | 0.2012 |
| CHLA10 | -0.2419 | 0.1238 | -0.1826 | -0.0753 | 0.3002 |
| CHLA266 | -0.247 | 0.1314 | -0.1355 | -0.1163 | 0.2943 |
| CHLA57 | -0.2499 | 0.241 | -0.1643 | -0.0969 | 0.1938 |
| CMK115 | -0.1013 | 0.0285 | 0.0314 | -0.1432 | 0.1651 |
| COGE352 | -0.2567 | 0.2625 | -0.1706 | -0.1409 | 0.2319 |
| COLO205 | 0.0206 | 0.0971 | -0.3217 | 0.1123 | 0.0866 |
| CHL1DM | -0.3673 | 0.1983 | -0.1942 | -0.0199 | 0.2577 |
| COV504 | -0.2217 | 0.2711 | -0.1956 | -0.1027 | 0.183 |
| CW9019 | -0.3778 | 0.433 | -0.1901 | -0.2313 | 0.2602 |
| D425 | -0.169 | 0.0573 | -0.2505 | 0.062 | 0.2347 |
| D458 | -0.0171 | 0.1505 | -0.26 | 0.0944 | 0.016 |
| DLD1 | -0.3614 | 0.2041 | -0.1747 | 0.0508 | 0.15 |
| DOV13 | -0.4002 | 0.3571 | -0.1258 | -0.0984 | 0.1389 |
| EVSAT | -0.2306 | 0.3142 | -0.0451 | -0.2244 | 0.1303 |
| F5 | -0.3282 | 0.2686 | -0.1129 | -0.0068 | 0.0659 |
| NCIH292 | -0.3903 | 0.3405 | -0.201 | 0.0452 | 0.065 |
| HCC2998 | -0.0247 | 0.2398 | -0.1949 | -0.0181 | -0.0087 |
| JR | -0.1797 | 0.2889 | -0.349 | -0.0201 | 0.1996 |
| KCIMOH1 | -0.3726 | 0.2352 | -0.2218 | 0.0124 | 0.2159 |
| KD | -0.0768 | 0.2196 | 0.0046 | -0.2711 | 0.1266 |
| KP1N | -0.2541 | 0.1557 | -0.1186 | -0.0485 | 0.1824 |
| MAC2A | -0.2281 | 0.1874 | -0.2991 | 0.0279 | 0.2296 |
| MOGGUVW | -0.0758 | 0.0887 | -0.3296 | 0.132 | 0.1439 |
| MON | -0.1093 | 0.2971 | -0.1818 | -0.1449 | 0.1168 |
| MONOMAC1 | -0.1114 | 0.1154 | -0.1675 | -0.1475 | 0.2884 |
| MYLA | -0.1808 | 0.1845 | -0.2719 | 0.023 | 0.1799 |
| NCIH1993 | -0.195 | 0.2261 | -0.3387 | 0.064 | 0.1688 |
| OC316 | -0.7011 | 0.2597 | 0.0701 | -0.0093 | 0.138 |
| OVCAR5 | -0.4553 | 0.252 | -0.2979 | 0.152 | 0.1744 |
| CCLFPEDS0001T | -0.2473 | 0.2397 | -0.1991 | 0.006 | 0.1141 |
| CCLFPEDS0003T | -0.25 | 0.2187 | -0.1062 | 0.1156 | -0.0777 |
| U251MGDM | -0.1989 | 0.4215 | -0.3298 | -0.0853 | 0.1327 |
| RT11284 | -0.3522 | 0.2774 | -0.0822 | -0.1792 | 0.2333 |
| SCMCRM2 | -0.0858 | 0.2691 | -0.2547 | 0.1096 | -0.08 |
| SHSY5Y | -0.0979 | 0.1568 | -0.3721 | 0.066 | 0.2059 |
| SKMEL2 | -0.1896 | 0.2249 | -0.318 | 0.0165 | 0.1985 |
| SKNEP1 | -0.2226 | 0.2987 | -0.3283 | -0.0119 | 0.1882 |
| SKPNDW | -0.2207 | -0.001 | -0.33 | 0.2662 | 0.1798 |
| SKRC31 | -0.2508 | 0.2765 | -0.1532 | 0.0506 | -0.0158 |
| SMSCTR | -0.2038 | 0.1515 | -0.1711 | 0.043 | 0.1048 |
| SMZ1 | -0.2042 | 0.1231 | -0.1933 | 0.1394 | 0.0488 |
| TC32 | -0.0804 | 0.3003 | -0.3273 | -0.0897 | 0.179 |
| TTC549 | -0.068 | 0.1285 | -0.2847 | 0.0618 | 0.132 |
| TTC642 | -0.088 | 0.2074 | -0.2599 | -0.1488 | 0.275 |
| UPCISCC152 | -0.5364 | 0.2131 | 0.03 | -0.0827 | 0.1985 |
| UPCISCC154 | -0.6535 | 0.2841 | -0.0206 | -0.064 | 0.2339 |
| UW228 | -0.1349 | 0.3898 | -0.1079 | -0.2678 | 0.1032 |
| VMRCLCD | -0.1672 | 0.0493 | -0.0676 | -0.0646 | 0.1991 |
| WM2664 | -0.2132 | 0.3447 | -0.2735 | -0.1288 | 0.2108 |
| 127399 | -0.3794 | 0.2201 | -0.1297 | -0.1557 | 0.3299 |
| SW982 | -0.2848 | 0.3086 | -0.1117 | -0.0737 | 0.0707 |
| SYO1 | -0.1188 | 0.2508 | 0.1033 | -0.3379 | 0.0982 |
| YAMATO | -0.1485 | 0.2629 | -0.2756 | 0.1408 | -0.0465 |
| BIN67 | -0.6189 | 0.3255 | -0.0606 | -0.0693 | 0.2158 |
| SCCOHT1 | -0.0478 | 0.1726 | -0.1735 | -0.0286 | 0.0638 |
| SCS214 | -0.3012 | 0.2618 | -0.3407 | 0.0362 | 0.2353 |
| TC106 | -0.1566 | 0.2645 | -0.3809 | 0.0133 | 0.2039 |
| COGAR359 | -0.2649 | 0.3224 | -0.4082 | 0.1539 | 0.088 |
| Y79 | -0.1187 | 0.2369 | -0.2871 | 0.0492 | 0.0731 |
| CHLA15 | -0.0709 | 0.2484 | -0.3727 | 0.0257 | 0.1422 |
| COGN278 | -0.1239 | 0.1332 | -0.0813 | -0.0843 | 0.1224 |
| COGN305 | -0.1856 | 0.1932 | -0.2267 | 0.085 | 0.0603 |
| NB1643 | 0.0291 | 0.2402 | -0.1655 | -0.0839 | -0.0005 |
| 8305C | -0.1675 | 0.2188 | -0.3817 | -0.0073 | 0.2803 |
| 8505C | -0.2462 | 0.139 | -0.0208 | -0.1362 | 0.1936 |
| HA1E | -0.237 | 0.1956 | 0.037 | -0.2447 | 0.1934 |
| PLCPRF5 | -0.5826 | 0.2491 | -0.0377 | 0.0302 | 0.1353 |
| CME1 | -0.258 | 0.3594 | -0.027 | -0.2346 | 0.0962 |
| A431 | -0.7061 | 0.3662 | -0.0824 | 0.0117 | 0.164 |
| ANGMCSS | -0.3463 | 0.4461 | -0.1591 | -0.2056 | 0.167 |
| BICR10 | -0.1442 | 0.2482 | -0.0269 | -0.2553 | 0.156 |
| BICR78 | -0.4765 | 0.2076 | -0.0483 | 0.1544 | -0.0197 |
| C33A | -0.5043 | 0.3735 | -0.1939 | 0.004 | 0.145 |
| C4I | -0.5025 | 0.1577 | -0.0824 | 0.0098 | 0.2417 |
| C4II | -0.5425 | 0.2014 | 0.0565 | -0.0148 | 0.1125 |
| CASKI | -0.4332 | 0.3525 | -0.0687 | -0.0736 | 0.0806 |
| CHP134 | -0.2068 | 0.0165 | -0.103 | -0.0147 | 0.2377 |
| COV413A | -0.6759 | 0.2575 | -0.0617 | 0.0485 | 0.1914 |
| DOTC24510 | -0.7343 | 0.1937 | -0.0156 | 0.0791 | 0.2132 |
| GIMEN | -0.1917 | 0.0738 | -0.1599 | 0.1875 | 0.0032 |
| GP5D | -0.3833 | 0.2222 | -0.1817 | 0.0428 | 0.1621 |
| H103 | -0.6326 | 0.287 | -0.0658 | -0.0442 | 0.2405 |
| H157 | -0.4892 | 0.3737 | -0.0161 | -0.1627 | 0.1422 |
| JOPACA1 | -0.4623 | 0.3772 | -0.2557 | -0.0414 | 0.2261 |
| LAN2 | -0.0556 | 0.2863 | -0.1992 | -0.1645 | 0.1316 |
| MB1 | -0.1525 | 0.3796 | -0.2464 | -0.2216 | 0.212 |
| MS751 | -0.274 | 0.2099 | -0.0534 | -0.1922 | 0.2354 |
| NGP | -0.2401 | 0.2343 | -0.1524 | -0.1491 | 0.2402 |
| NMB | -0.1935 | 0.2597 | -0.281 | -0.0975 | 0.2558 |
| OACM51 | -0.1051 | 0.1848 | -0.1646 | -0.1017 | 0.1613 |
| OCIP5X | -0.1963 | -0.0083 | 0.0867 | 0.006 | 0.043 |
| PA1 | -0.2369 | 0.2424 | -0.3589 | -0.1231 | 0.4077 |
| PACADD119 | -0.5821 | 0.1998 | -0.2228 | 0.086 | 0.3074 |
| PACADD137 | -0.5332 | 0.1769 | -0.0255 | 0.1022 | 0.0831 |
| PACADD161 | -0.3325 | 0.246 | -0.1968 | -0.0049 | 0.1733 |
| PACADD165 | -0.1725 | 0.2677 | -0.0823 | -0.182 | 0.129 |
| PACADD188 | -0.332 | 0.2048 | -0.2973 | 0.1014 | 0.1966 |
| RPMI2650 | -0.2952 | 0.2342 | -0.032 | 0.0155 | -0.0267 |
| SCLC22H | -0.1788 | 0.1162 | -0.1779 | -0.079 | 0.266 |
| SUM102PT | -0.4042 | 0.2129 | -0.1009 | -0.1313 | 0.2973 |
| SUM1315MO2 | -0.1349 | 0.166 | -0.1393 | -0.128 | 0.2033 |
| SUM149PT | -0.4131 | 0.1392 | -0.1928 | 0.2177 | 0.0816 |
| SUM159PT | -0.1997 | 0.2624 | -0.2511 | 0.0478 | 0.066 |
| SUM185PE | -0.2242 | 0.2705 | -0.2764 | -0.0675 | 0.2271 |
| SUM190PT | -0.5499 | 0.1766 | 0.0076 | 0.0137 | 0.1593 |
| SUM229PE | -0.3576 | 0.1181 | -0.1707 | 0.0276 | 0.2553 |
| SUM52PE | -0.0406 | 0.2055 | -0.1027 | -0.2381 | 0.1878 |
| SW156 | -0.2132 | 0.2136 | -0.1895 | -0.0749 | 0.1981 |
| SW626 | -0.1654 | 0.2346 | -0.2719 | -0.0607 | 0.2125 |
| SW954 | -0.5961 | 0.184 | -0.0677 | 0.0824 | 0.1812 |
| SW756 | -0.3266 | 0.2682 | -0.2453 | 0.018 | 0.1703 |
| TO14 | -0.2246 | 0.2152 | -0.2196 | 0.0443 | 0.1017 |
| UMUC13 | -0.1166 | 0.1781 | -0.0507 | -0.1778 | 0.1459 |

| R | EG1 | EG2 | EG3 | EG4 | EG5 |
|---|---|---|---|---|---|
| UMUC16 | -0.1929 | 0.3172 | -0.3594 | -0.0346 | 0.2064 |
| UMUC5 | -0.3075 | 0.3034 | -0.1145 | -0.0357 | 0.0513 |
| UMUC10 | -0.6308 | 0.3821 | -0.0774 | 0.035 | 0.068 |
| UMUC11 | -0.215 | 0.2737 | 0.0111 | -0.2554 | 0.1388 |
| UMUC6 | -0.3174 | 0.2719 | -0.0579 | -0.1039 | 0.1083 |
| UMUC7 | -0.5783 | 0.3358 | -0.0383 | 0.0495 | 0.0248 |
| UMUC9 | -0.4135 | 0.4348 | -0.1157 | -0.0991 | 0.0606 |
| UWB1289 | -0.6921 | 0.3045 | 0.0597 | 0.0132 | 0.0727 |
| VP229 | -0.5779 | 0.2126 | 0.0519 | -0.0657 | 0.1854 |
| WERIRB1 | -0.2648 | 0.1311 | -0.1707 | 0.0175 | 0.193 |
| WPE1NA22 | -0.4395 | 0.1404 | -0.0794 | 0.019 | 0.2047 |
| TC138 | -0.2271 | 0.2616 | -0.2839 | -0.0302 | 0.2039 |
| TC205 | -0.1406 | 0.296 | -0.1948 | -0.0421 | 0.0372 |
| CCLFPEDS0008T | -0.1981 | 0.395 | -0.3077 | -0.1565 | 0.2156 |
| A388 | -0.6069 | 0.1937 | -0.144 | 0.1617 | 0.167 |
| ASH3 | -0.4444 | 0.1362 | -0.1456 | 0.2058 | 0.0709 |
| BLUE1 | -0.1401 | 0.0606 | -0.0575 | -0.1401 | 0.2437 |
| BOKU | -0.2352 | 0.2053 | -0.2525 | -0.0596 | 0.2668 |
| BPH1 | -0.6921 | 0.3009 | -0.0295 | 0.0811 | 0.0902 |
| C10 | -0.0589 | 0.3935 | -0.0934 | -0.2331 | -0.0031 |
| C75 | -0.6262 | 0.298 | -0.2469 | 0.1326 | 0.2104 |
| C80 | -0.3373 | 0.4429 | -0.1175 | -0.2412 | 0.1623 |
| C84 | -0.5038 | 0.1435 | -0.0234 | -0.0263 | 0.2377 |
| C99 | -0.3492 | 0.2859 | -0.1076 | 0.046 | -0.0014 |
| CHLA90 | -0.2618 | 0.1017 | 0.0017 | -0.0235 | 0.0935 |
| EGI1 | -0.3429 | 0.2921 | -0.3387 | 0.1006 | 0.1587 |
| EMTOKA | -0.4792 | 0.2431 | 0.0132 | 0.0367 | 0.0157 |
| ESO26 | -0.7304 | 0.1527 | 0.0723 | 0.0823 | 0.1603 |
| ESO51 | -0.1084 | 0.2873 | -0.1373 | -0.0392 | -0.0359 |
| FARAGE | -0.2971 | 0.1278 | -0.1153 | -0.0412 | 0.2271 |
| FLO1 | -0.1737 | 0.3136 | -0.2746 | -0.0674 | 0.1492 |
| H357 | -0.5581 | 0.2378 | -0.1928 | 0.1256 | 0.1798 |
| H376 | -0.4073 | 0.3211 | -0.1072 | 0.0271 | 0.0218 |
| H413 | -0.7236 | 0.2765 | 0.0249 | 0.0286 | 0.1389 |
| HCA1 | -0.5232 | 0.3562 | -0.08 | -0.0148 | 0.0816 |
| HCS2 | -0.3432 | 0.2921 | -0.0921 | -0.1119 | 0.1482 |
| HEC1 | -0.3514 | 0.2249 | -0.2609 | -0.0032 | 0.2689 |
| HEC116 | -0.3167 | 0.1546 | -0.1883 | 0.0388 | 0.1973 |
| HG3 | -0.3485 | 0.2229 | -0.108 | -0.0029 | 0.1156 |
| HKA1 | -0.4727 | 0.2216 | -0.0096 | 0.0221 | 0.072 |
| HMY1 | -0.2247 | 0.1912 | -0.1679 | -0.0297 | 0.1563 |
| HSC1 | -0.4087 | 0.3296 | -0.1667 | -0.1272 | 0.2449 |
| HSC5 | -0.6938 | 0.3584 | -0.0592 | -0.0097 | 0.1644 |
| HT3 | -0.0301 | 0.2557 | -0.1312 | -0.0595 | -0.0388 |
| HUO9 | -0.1613 | 0.1003 | -0.2272 | 0.0251 | 0.2043 |
| IHH4 | -0.2482 | 0.0883 | -0.3158 | 0.129 | 0.2463 |
| JAR | -0.1668 | 0.2073 | -0.2074 | -0.1334 | 0.2568 |
| JEG3 | -0.3073 | 0.2348 | 0.0171 | -0.0581 | 0.0131 |
| JMURTK2 | -0.0595 | 0.1972 | 0.038 | -0.2973 | 0.1335 |
| KARPAS1718 | -0.2176 | 0.0634 | -0.2063 | 0.0527 | 0.2264 |
| KKU100 | -0.3301 | 0.4121 | -0.163 | -0.1515 | 0.1344 |
| KKU213 | -0.3351 | 0.2483 | -0.426 | 0.2508 | 0.1181 |
| KML1 | -0.2832 | 0.1886 | -0.1429 | 0.0934 | 0.0355 |
| KON | -0.4089 | 0.4101 | -0.1358 | -0.0387 | 0.0354 |
| KOSC2 | -0.4644 | 0.3262 | -0.0981 | 0.0256 | 0.0465 |
| KYAE1 | -0.738 | 0.1904 | -0.0242 | 0.1983 | 0.0954 |
| LO68 | -0.0699 | 0.2017 | -0.2053 | -0.1592 | 0.2258 |
| LS | -0.1745 | 0.1415 | -0.0336 | -0.084 | 0.0992 |
| LU135 | -0.1378 | 0.269 | -0.2963 | -0.1403 | 0.2728 |
| MCC13 | -0.2092 | 0.1687 | -0.2365 | -0.0341 | 0.2422 |
| MCC142 | -0.1741 | 0.2254 | -0.0954 | -0.1725 | 0.1749 |
| MCC26 | -0.149 | 0.3277 | -0.3094 | -0.1123 | 0.2034 |
| MEL202 | -0.4289 | 0.2544 | -0.2594 | 0.0349 | 0.2462 |
| MERO14 | -0.1953 | 0.1777 | 0.0245 | -0.1127 | 0.0503 |
| MERO25 | -0.2438 | 0.2871 | -0.053 | -0.1138 | 0.0513 |
| MERO41 | -0.081 | 0.2318 | -0.1262 | -0.2429 | 0.2167 |
| MERO48A | -0.0644 | 0.2292 | 0.0393 | -0.2114 | 0.008 |
| MERO82 | -0.172 | 0.3503 | -0.0031 | -0.298 | 0.0956 |
| MERO83 | -0.2933 | 0.2945 | -0.0962 | -0.0732 | 0.0743 |
| MERO95 | -0.1693 | 0.1644 | 0.0664 | -0.2502 | 0.1572 |
| MM127 | -0.1477 | 0.0652 | -0.0618 | -0.0754 | 0.1767 |
| MM370 | -0.2061 | 0.4131 | -0.2012 | -0.1653 | 0.1059 |
| MM383 | -0.4914 | 0.1799 | -0.0471 | -0.0828 | 0.2797 |
| MM386 | -0.2133 | 0.1541 | -0.222 | 0.0107 | 0.1951 |
| MM426 | -0.3119 | 0.3111 | -0.0975 | -0.1677 | 0.1759 |
| MOLM14 | -0.0413 | 0.1499 | -0.0139 | -0.187 | 0.0985 |
| MUTZ8 | -0.0681 | 0.15 | -0.2476 | 0.0985 | 0.0329 |
| NH12 | -0.1869 | 0.2533 | -0.3608 | 0.0399 | 0.1852 |
| NO10 | -0.3294 | 0.3214 | -0.0512 | -0.0693 | 0.0216 |
| NO11 | -0.1367 | 0.1635 | -0.0379 | -0.1904 | 0.1749 |
| NOZ | -0.2145 | 0.1874 | -0.1786 | 0.1299 | -0.0129 |
| NP2 | -0.1358 | 0.351 | -0.2121 | -0.1416 | 0.1068 |
| NP3 | -0.0433 | 0.1659 | -0.0132 | -0.2649 | 0.1694 |
| NP5 | -0.0726 | 0.1428 | -0.0206 | -0.0964 | 0.0322 |
| NP8 | -0.2169 | 0.0198 | -0.0916 | -0.0044 | 0.2182 |
| OCILY18 | -0.3933 | 0.1046 | -0.1239 | 0.136 | 0.125 |
| OCIM2 | -0.1217 | 0.0938 | -0.2338 | 0.1445 | 0.0591 |
| OCUG1 | -0.3902 | 0.1814 | -0.2244 | 0.1751 | 0.1034 |
| ONDA7 | -0.3544 | 0.29 | -0.0264 | -0.226 | 0.2183 |
| ONDA8 | -0.1064 | 0.2923 | -0.0456 | -0.3048 | 0.1608 |
| ONDA9 | -0.2175 | 0.1494 | -0.0772 | -0.0346 | 0.1082 |
| OSC19 | -0.6563 | 0.1793 | -0.0231 | 0.0881 | 0.1743 |
| OSC20 | -0.4452 | 0.2351 | -0.0956 | -0.068 | 0.2263 |
| P4E6 | -0.4819 | 0.2284 | -0.1586 | 0.1037 | 0.1295 |
| PEA1 | -0.3301 | 0.1971 | -0.1424 | -0.1238 | 0.298 |
| PEO1 | -0.6098 | 0.2842 | -0.0451 | -0.0355 | 0.1982 |
| PEO4 | -0.5113 | 0.1474 | 0.0038 | 0.1043 | 0.0666 |
| PGA1 | -0.1301 | 0.0275 | -0.2395 | 0.0616 | 0.2286 |
| ROS50 | 0.1558 | 0.1266 | -0.2867 | -0.0784 | 0.1454 |
| SAT | -0.6705 | 0.1403 | 0.0343 | 0.0677 | 0.1878 |
| SEKI | -0.1364 | 0.0436 | -0.13 | 0.1097 | 0.0536 |
| SHI1 | -0.1328 | 0.1284 | -0.295 | -0.0983 | 0.3622 |
| SHMAC4 | -0.5035 | 0.2417 | -0.12 | 0.0621 | 0.1379 |
| SHMAC5 | -0.4417 | 0.0837 | -0.0084 | 0.0575 | 0.1491 |
| SISO | -0.4477 | 0.2494 | -0.2371 | 0.1368 | 0.1281 |
| SKGI | -0.305 | 0.1923 | 0.0516 | -0.232 | 0.2124 |
| SKGII | -0.6143 | 0.2473 | -0.0569 | 0.0608 | 0.1429 |
| SKGT2 | -0.5242 | 0.3385 | -0.1701 | -0.0087 | 0.1833 |
| SKGT4 | -0.6832 | 0.1837 | -0.0021 | 0.0854 | 0.1694 |
| SKN | -0.3206 | 0.2504 | -0.1693 | -0.0962 | 0.2348 |
| SKNO1 | -0.1265 | 0.0958 | -0.0255 | -0.1797 | 0.2116 |
| SNU638 | -0.0999 | 0.1458 | -0.3274 | 0.1135 | 0.1209 |
| SUSA | -0.2337 | 0.3311 | -0.2644 | -0.0663 | 0.1593 |
| TANOUE | -0.2239 | 0.021 | -0.1005 | -0.0292 | 0.2579 |
| TASK1 | -0.21 | 0.1454 | -0.3215 | 0.0052 | 0.3074 |
| TFK1 | -0.4029 | 0.2827 | -0.1836 | 0.0098 | 0.1529 |
| TGW | -0.2155 | 0.1912 | -0.2705 | 0.0989 | 0.1102 |
| TR146 | -0.5171 | 0.1523 | -0.0396 | 0.0638 | 0.1539 |
| U2904 | -0.3651 | 0.0626 | -0.0784 | -0.0737 | 0.3357 |
| UHO1 | -0.1867 | 0.1643 | -0.023 | -0.0904 | 0.0809 |
| UMRC3 | -0.0947 | 0.2145 | -0.1635 | -0.1377 | 0.1636 |
| UMRC7 | -0.181 | 0.3071 | -0.3508 | -0.0259 | 0.1905 |
| UPCISCC026 | -0.6006 | 0.2793 | -0.0651 | 0.0834 | 0.0851 |
| UPCISCC029A | -0.1204 | 0.1929 | -0.0987 | -0.1561 | 0.1575 |
| UPCISCC040 | -0.6062 | 0.2631 | -0.0741 | 0.0464 | 0.1551 |
| UPCISCC074 | -0.6536 | 0.22 | -0.0149 | 0.0231 | 0.1957 |
| UPCISCC111 | -0.6703 | 0.2952 | 0.0031 | 0.009 | 0.1291 |
| UPCISCC116 | -0.5824 | 0.3152 | -0.1507 | 0.0251 | 0.1877 |

| R | EG1 | EG2 | EG3 | EG4 | EG5 |
|---|---|---|---|---|---|
| UPCISCC200 | -0.4905 | 0.2523 | -0.0111 | 0.0265 | 0.0494 |
| VAESBJ | -0.1986 | 0.2109 | -0.1294 | 0.0706 | -0.0302 |
| WAOSEL | -0.138 | 0.129 | -0.1427 | -0.0267 | 0.1333 |
| WSUNHL | -0.3787 | 0.1512 | -0.1619 | 0.1211 | 0.1234 |
| PFSK1 | -0.209 | 0.241 | -0.1355 | 0.1156 | -0.0975 |
| CAL72 | -0.0621 | 0.1701 | -0.1014 | -0.15 | 0.1382 |
| OCIC4P | -0.4667 | 0.1059 | -0.0403 | -0.0111 | 0.2511 |
| SEMK2 | -0.0773 | 0.189 | -0.103 | -0.2047 | 0.1915 |
| HB1119 | -0.1014 | 0.1833 | -0.0402 | -0.2061 | 0.1516 |
| CTV1DM | -0.0573 | 0.2045 | -0.2356 | -0.0976 | 0.1768 |
| RH28 | -0.423 | 0.2141 | -0.3384 | 0.2429 | 0.1309 |
| RHJT | -0.2483 | 0.292 | -0.4058 | 0.0982 | 0.1669 |
| TTC442 | -0.5411 | 0.2534 | -0.2163 | 0.0409 | 0.2706 |
| RH4 | -0.1327 | 0.3252 | -0.3612 | 0.0129 | 0.1082 |
| SNU1544 | -0.4803 | 0.3286 | 0.0462 | -0.1585 | 0.1144 |
| LPS6 | -0.2782 | 0.3144 | -0.1194 | -0.1902 | 0.1975 |
| LPS27 | -0.1811 | 0.2861 | -0.019 | -0.238 | 0.1151 |
| 93T449 | -0.1585 | 0.1823 | -0.0815 | -0.1598 | 0.1799 |
| 94T778 | -0.1697 | 0.3789 | -0.155 | -0.1831 | 0.09 |
| 95T1000 | -0.1401 | 0.1554 | -0.0388 | -0.1355 | 0.1251 |
| LPS141 | -0.3897 | 0.2429 | -0.0211 | -0.1159 | 0.161 |
| LPS853 | -0.3225 | 0.2227 | -0.1502 | -0.0271 | 0.168 |
| LPS510 | -0.1945 | 0.2179 | 0.0266 | -0.0955 | -0.0116 |
| OS252 | -0.3989 | 0.2484 | -0.0837 | -0.0646 | 0.1672 |
| MFM223 | -0.1259 | 0.1839 | -0.0128 | -0.197 | 0.1296 |
| COLO824 | -0.3447 | 0.4044 | -0.2342 | -0.0582 | 0.1193 |
| ICC10 | -0.4097 | 0.246 | 0.0716 | -0.1999 | 0.1714 |
| ICC106 | -0.4194 | 0.2557 | 0.0058 | -0.113 | 0.1374 |
| ICC108 | -0.5673 | 0.2105 | 0.0133 | 0.0439 | 0.0977 |
| ICC12 | -0.2819 | 0.3694 | -0.2866 | 0.1444 | -0.059 |
| ICC137 | -0.381 | 0.2193 | -0.1211 | -0.1274 | 0.2917 |
| ICC15 | -0.6153 | 0.1989 | -0.0481 | 0.1042 | 0.1351 |
| ICC2 | -0.5512 | 0.1954 | -0.1072 | 0.0197 | 0.2496 |
| ICC3 | -0.3454 | 0.2298 | -0.0244 | -0.2046 | 0.2469 |
| ICC4 | -0.4973 | 0.2308 | -0.0966 | 0.111 | 0.0672 |
| ICC8 | -0.2459 | 0.4364 | -0.1386 | -0.2224 | 0.1094 |
| ICC9 | -0.4348 | 0.3513 | -0.1477 | -0.002 | 0.0823 |
| G415 | -0.2894 | 0.3233 | -0.1728 | -0.0384 | 0.081 |
| HKGZCC | -0.7865 | 0.2698 | -0.0712 | 0.1432 | 0.1558 |
| KMCH1 | -0.4441 | 0.3639 | -0.1143 | -0.0544 | 0.1004 |
| RBE | -0.4729 | 0.3679 | -0.1458 | -0.0005 | 0.0871 |
| SG231 | -0.1207 | 0.3035 | -0.2298 | 0.0731 | -0.0762 |
| SSP25 | -0.6826 | 0.3448 | -0.1311 | 0.0914 | 0.1302 |
| TGBC1TKB | -0.6166 | 0.3211 | 0.0857 | -0.0567 | 0.0585 |
| TGBC52TKB | -0.5189 | 0.0851 | -0.1454 | 0.1895 | 0.1886 |
| TKKK | -0.623 | 0.1185 | -0.0616 | 0.0235 | 0.3236 |
| YSCCC | -0.4674 | 0.2606 | -0.0158 | 0.0498 | 0.0049 |
| CCLP1 | -0.0986 | 0.1918 | -0.1601 | -0.1119 | 0.1567 |
| CCSW1 | -0.589 | 0.3014 | -0.1607 | 0.1118 | 0.1196 |
| GB2 | -0.4576 | 0.1768 | -0.0677 | 0.0809 | 0.0997 |
| MM253 | -0.3149 | 0.0358 | -0.202 | 0.1856 | 0.1658 |
| MM485 | -0.1986 | 0.0125 | -0.2221 | 0.097 | 0.2315 |
| NO36 | -0.4012 | 0.2098 | -0.1031 | -0.0409 | 0.2004 |
| NZM3 | -0.286 | 0.1134 | -0.1306 | 0.0368 | 0.1629 |
| NZM42 | -0.3772 | 0.3136 | -0.1041 | -0.0652 | 0.1089 |
| NZM7 | -0.359 | 0.3264 | -0.2529 | 0.0605 | 0.0936 |
| NZOV9 | -0.1523 | 0.0249 | -0.0717 | -0.037 | 0.1872 |
| ONE58 | -0.168 | 0.1897 | -0.0858 | -0.148 | 0.1699 |
| NALM16 | -0.0586 | 0.1164 | -0.2098 | -0.0399 | 0.1759 |
| ECC2 | -0.5309 | 0.2879 | -0.1887 | 0.0529 | 0.1885 |
| 9505BIK | -0.467 | 0.2109 | -0.163 | -0.0431 | 0.3046 |
| A375SKINCJ1 | -0.3297 | 0.1542 | -0.1321 | 0.028 | 0.162 |
| A375SKINCJ2 | -0.3299 | 0.2089 | -0.1721 | 0.0538 | 0.1187 |
| A375SKINCJ3 | -0.1908 | 0.1561 | -0.4419 | 0.1276 | 0.2687 |
| SKMEL19 | -0.3036 | 0.3353 | -0.2699 | -0.0411 | 0.1783 |
| MEL270 | -0.2957 | 0.2381 | -0.1978 | 0.1885 | -0.0564 |
| MEL285 | -0.1063 | 0.1801 | -0.0184 | -0.205 | 0.135 |
| MEL290 | -0.1986 | 0.3332 | -0.1733 | -0.1492 | 0.1352 |
| OMM25 | -0.3475 | 0.237 | -0.2077 | 0.0808 | 0.1078 |
| HOKUG | -0.4699 | 0.2227 | -0.0846 | 0.027 | 0.1385 |
| SKGIIIA | -0.5291 | 0.1601 | -0.1108 | 0.1125 | 0.1712 |
| T3M3 | -0.2401 | 0.1653 | -0.1578 | 0.0184 | 0.1287 |
| TGBC18TKB | -0.6704 | 0.1342 | -0.0015 | 0.1396 | 0.1498 |
| ECC4 | -0.0866 | 0.0209 | -0.3456 | 0.0496 | 0.3262 |
| TT1TKB | -0.5706 | 0.1889 | -0.0281 | 0.1994 | -0.0097 |
| HHUA | -0.4146 | 0.2354 | -0.0628 | 0.0132 | 0.0832 |
| HOUAI | -0.4267 | 0.1658 | 0.0547 | 0.0029 | 0.0547 |
| SAS | -0.5297 | 0.1888 | -0.0256 | 0.0509 | 0.1259 |
| LCAM1 | -0.0062 | 0.1136 | -0.2958 | -0.16 | 0.3638 |
| PK8 | -0.1523 | 0.3059 | -0.1178 | -0.1285 | 0.0537 |
| HOTHC | -0.2533 | 0.2735 | -0.2824 | -0.057 | 0.2375 |
| T3M5 | -0.3482 | 0.1979 | -0.1889 | 0.0201 | 0.196 |
| CA922 | -0.3005 | 0.2067 | -0.1725 | -0.0035 | 0.1657 |
| HSQ89 | -0.0792 | 0.2335 | -0.1746 | -0.1503 | 0.1594 |
| HO1U1 | -0.3588 | 0.2527 | 0.0047 | -0.038 | 0.0188 |
| HTMMT | -0.3357 | 0.2457 | -0.104 | -0.1251 | 0.2161 |
| RMSYM | -0.1845 | 0.1974 | -0.1617 | -0.0914 | 0.186 |
| LU134A | -0.1747 | 0.1491 | -0.1591 | -0.1169 | 0.2538 |
| P30OHK | -0.1648 | 0.086 | -0.0266 | -0.2129 | 0.2843 |
| P2URK562 | -0.1837 | 0.0017 | -0.0758 | -0.0484 | 0.2476 |
| SLVL | -0.152 | 0.0239 | 0.0281 | -0.0454 | 0.0975 |
| HSSCH2 | -0.1753 | 0.2122 | -0.1387 | -0.1325 | 0.1879 |
| NOS1 | -0.1603 | 0.2194 | -0.3289 | 0.0402 | 0.1696 |
| HSOS1 | -0.1525 | 0.1978 | -0.23 | -0.1136 | 0.2578 |
| HKBMM | -0.0822 | 0.2345 | -0.1517 | -0.167 | 0.1561 |
| LU165 | -0.3209 | 0.1044 | -0.1446 | -0.0017 | 0.2515 |
| TN2 | -0.1104 | 0.2307 | -0.2941 | -0.0466 | 0.1871 |
| NB69 | -0.2902 | 0.2873 | -0.2004 | 0.0182 | 0.0822 |
| MMAC | -0.3072 | 0.0164 | -0.127 | 0.098 | 0.2023 |
| ES4 | -0.5309 | 0.2089 | -0.026 | -0.0342 | 0.2014 |
| ES5 | -0.0338 | 0.1223 | -0.211 | -0.0202 | 0.1331 |
| ES8 | 0.0426 | 0.2264 | -0.3419 | 0.0196 | 0.0659 |
| EW1 | -0.1061 | 0.138 | -0.1931 | 0.0182 | 0.1042 |
| EW16 | -0.3006 | 0.2314 | -0.2421 | -0.0657 | 0.2797 |
| EW22 | -0.0839 | 0.0889 | -0.3505 | 0.1165 | 0.187 |
| EW7 | -0.1783 | 0.1279 | 0.0484 | -0.1856 | 0.1459 |
| HEY | -0.2357 | 0.3554 | -0.3066 | -0.0164 | 0.1231 |
| HSC39 | -0.3198 | 0.1962 | -0.0224 | -0.093 | 0.138 |
| ISTMEL1 | -0.2932 | 0.0328 | 0.0576 | 0.0302 | 0.0674 |
| LB1047RCC | -0.5483 | 0.3188 | -0.0608 | -0.0445 | 0.1491 |
| LC1SQ | -0.5757 | 0.1585 | -0.0262 | 0.1002 | 0.1322 |
| MCIXC | -0.1288 | 0.0203 | -0.0812 | 0.1131 | 0.0195 |
| MKN28 | -0.313 | 0.1992 | -0.1388 | -0.006 | 0.1505 |
| OCUBM | -0.6431 | 0.1897 | 0.0877 | -0.0478 | 0.1953 |
| OVCA420 | -0.5532 | 0.1366 | -0.0192 | 0.0547 | 0.183 |
| OVMIU | -0.4544 | 0.2549 | -0.0092 | -0.047 | 0.1029 |
| PL4 | -0.1681 | 0.1341 | -0.118 | 0.0116 | 0.0807 |
| RCCFG2 | -0.0077 | 0.3586 | -0.1413 | -0.2107 | 0.0216 |
| SCH | -0.223 | 0.2163 | 0.0169 | -0.0695 | -0.0106 |
| TMK1 | -0.4411 | 0.1199 | -0.0965 | 0.173 | 0.0725 |
| ARH77 | -0.1775 | 0.1206 | -0.0477 | 0.0976 | -0.0654 |
| BB30HNC | -0.4253 | 0.3782 | -0.1647 | 0.0286 | 0.0322 |
| BE2M17 | -0.1325 | 0.2165 | -0.2928 | -0.0448 | 0.2125 |
| D423MG | -0.3752 | 0.2739 | -0.0651 | -0.0175 | 0.0554 |
| D502MG | -0.2944 | 0.3185 | -0.2733 | 0.0871 | 0.0504 |
| D542MG | -0.0772 | 0.3884 | -0.0596 | -0.3414 | 0.1003 |
| DIFI | -0.4421 | 0.0824 | -0.0487 | 0.1307 | 0.1097 |
| DOK | -0.6517 | 0.2014 | -0.0772 | 0.1579 | 0.1258 |

| R | EG1 | EG2 | EG3 | EG4 | EG5 |
|---|---|---|---|---|---|
| HCE4 | -0.2403 | 0.1603 | -0.1302 | -0.1261 | 0.2666 |
| IM9 | -0.2064 | 0.0827 | 0.0594 | -0.0251 | 0.0205 |
| JHU011 | -0.5003 | 0.2114 | -0.1021 | 0.0298 | 0.1841 |
| JHU022 | -0.6444 | 0.1349 | 0.0059 | 0.1499 | 0.1135 |
| JHU029 | -0.4321 | 0.2252 | -0.0716 | -0.0122 | 0.1418 |
| KINGS1 | 0.0484 | 0.2241 | -0.4051 | -0.008 | 0.1583 |
| KPNYS | -0.006 | 0.1607 | -0.0374 | -0.2423 | 0.1494 |
| KYSE220 | -0.3638 | 0.3268 | -0.217 | 0.0727 | 0.047 |
| LB771HNC | -0.2216 | 0.238 | -0.2945 | 0.0555 | 0.1395 |
| LNZTA3WT4 | -0.0157 | 0.1727 | -0.0533 | -0.3104 | 0.2352 |
| NB10 | 0.0065 | 0.1288 | -0.3091 | 0.0452 | 0.1258 |
| NB13 | -0.1753 | 0.1161 | -0.1887 | -0.0002 | 0.1871 |
| NB17 | 0.0937 | 0.24 | -0.2067 | -0.2004 | 0.1279 |
| NB5 | -0.1325 | 0.2499 | -0.2274 | -0.0504 | 0.1198 |
| NB6 | -0.0633 | 0.1292 | -0.1717 | -0.1741 | 0.2769 |
| NB7 | -0.2002 | 0.067 | -0.1814 | 0.1211 | 0.1107 |
| NTERA2CLD1 | -0.3875 | 0.1012 | -0.1662 | 0.1576 | 0.143 |
| PCI15A | -0.508 | 0.2998 | -0.1807 | -0.0206 | 0.2352 |
| PCI30 | -0.0886 | 0.2547 | -0.0959 | -0.0917 | 0.0007 |
| PCI38 | -0.5657 | 0.397 | -0.1121 | -0.0899 | 0.1839 |
| PCI4B | -0.5451 | 0.2382 | -0.0977 | 0.1024 | 0.1016 |
| PCI6A | -0.6608 | 0.3411 | -0.1333 | 0.0295 | 0.1906 |
| SKMG1 | -0.0478 | 0.2496 | -0.2497 | -0.0173 | 0.0504 |
| SKN3 | -0.2862 | 0.2309 | -0.1595 | 0.1064 | -0.0028 |
| 21NT | -0.4241 | 0.2266 | -0.0894 | -0.1143 | 0.2662 |
| CCLFUPGI0005T | -0.3295 | 0.1754 | 0.0765 | -0.1946 | 0.179 |
| HT144SKINFV1 | -0.3214 | 0.1395 | -0.0014 | -0.0068 | 0.0791 |
| HT144SKINFV3 | -0.2438 | 0.2431 | -0.0856 | -0.1332 | 0.1494 |
| HT144SKINFV2 | -0.4024 | 0.2261 | -0.0867 | 0.0676 | 0.0482 |
| RVH421SKINFV1 | -0.4067 | 0.0639 | -0.0331 | -0.0191 | 0.2558 |
| RPE1SS48 | -0.2472 | 0.235 | -0.1392 | 0.0437 | 0.0169 |
| RPE1SS77 | -0.2399 | 0.2834 | -0.2498 | -0.0405 | 0.1679 |
| RPE1SS6 | -0.0935 | 0.1712 | -0.2732 | 0.0535 | 0.1037 |
| RPE1SS119 | -0.1925 | 0.1042 | -0.209 | 0.0643 | 0.1591 |
| RPE1SS51 | -0.201 | 0.2264 | -0.1611 | -0.0184 | 0.0863 |
| PSS008 | -0.4709 | 0.187 | -0.1559 | 0.0254 | 0.248 |
| MAVER1 | -0.1948 | 0.316 | -0.1625 | -0.0301 | 0.0069 |
| MESOV | -0.3044 | 0.241 | -0.237 | 0.0817 | 0.104 |
| WM3211 | -0.0944 | 0.1482 | -0.1024 | -0.1156 | 0.144 |
| M040416 | -0.2462 | 0.015 | -0.2359 | 0.0492 | 0.327 |
| M140325 | -0.3108 | 0.1309 | -0.1656 | 0.0077 | 0.229 |
| MM160113 | -0.3289 | 0.0046 | -0.1515 | 0.0578 | 0.2974 |
| SNU739 | -0.2636 | 0.2931 | -0.1547 | -0.1117 | 0.1576 |
| SNU1327 | -0.2792 | 0.2656 | -0.1064 | 0.0218 | -0.0013 |
| SNU2535 | -0.2637 | 0.0795 | 0.0054 | -0.0582 | 0.1518 |
| CCC5 | -0.3536 | 0.1409 | -0.0344 | 0.0195 | 0.1025 |
| ETCC016 | -0.4831 | 0.1793 | -0.0864 | 0.0471 | 0.1701 |
| JVE015 | -0.0845 | 0.074 | 0.0107 | -0.257 | 0.2555 |
| JVE127 | -0.236 | 0.2311 | -0.3523 | 0.0337 | 0.2377 |
| JVE253 | -0.2319 | 0.1098 | -0.2408 | -0.0225 | 0.3072 |
| KP363T | -0.2654 | 0.2079 | -0.1589 | -0.0188 | 0.1449 |
| MAPACHS77 | -0.5071 | 0.2247 | -0.1402 | 0.1046 | 0.1304 |
| 170MGBA | -0.1711 | 0.2783 | -0.0381 | -0.2608 | 0.1607 |
| WM3772F | -0.1853 | 0.2097 | -0.0243 | 0.0506 | -0.1207 |
| S462 | -0.31 | 0.1167 | -0.104 | 0.0434 | 0.1414 |
| MPNST724 | -0.1079 | 0.2029 | -0.017 | -0.2005 | 0.107 |
| NCCLMS1C1 | -0.1641 | 0.1467 | 0.0676 | -0.2428 | 0.1621 |
| NCCMPNST1C1 | -0.1512 | 0.1075 | -0.2179 | -0.1663 | 0.3935 |
| NCCMPNST2C1 | -0.2704 | 0.2028 | -0.1825 | -0.0423 | 0.203 |
| PSS131R | -0.1895 | 0.1536 | -0.1295 | -0.0438 | 0.1482 |
| YUHOIN0650 | -0.4447 | 0.2837 | -0.2569 | 0.0878 | 0.1659 |
| SKNMM | -0.2095 | 0.1042 | -0.2025 | -0.0519 | 0.2924 |
| UPMD1 | -0.1971 | 0.1321 | -0.0511 | -0.0451 | 0.0976 |
