## Supplement Table S5D for "Single-molecule behavior and cell-growth regulation in human RTKs"

| resting | EC | IDRE | TM | IDRC | Linker | Tail | IDRE posi | IDRC pos | RK cluste | GxxxG | CRAC |
|---|---|---|---|---|---|---|---|---|---|---|---|
| D1 | 0.2966 | 0.0106 | 0.0132 | 0.1085 | 0.3068 | -0.0369 | -0.051 | 0.1126 | 0.0764 | -0.0356 | 0.088 |
| D2 | 0.2671 | 0.0263 | -0.0145 | 0.1277 | 0.2958 | -0.2455 | 0.0224 | 0.0863 | 0.0426 | -0.0346 | 0.1755 |
| D3 | -0.1861 | -0.0704 | -0.1085 | 0.1105 | 0.078 | -0.0313 | 0.2073 | 0.1027 | 0.1016 | -0.0828 | 0.1983 |
| F1 | -0.078 | 0.193 | 0.0355 | -0.1126 | -0.0433 | 0.0887 | -0.1394 | -0.0179 | -0.0217 | 0.1062 | -0.2104 |
| F2 | 0.0933 | 0.1827 | 0.135 | -0.1121 | 0.0526 | -0.0886 | -0.1035 | 0.0179 | -0.0018 | 0.1526 | -0.1746 |
| F3 | 0.2304 | 0.0989 | 0.1703 | -0.0987 | 0.1725 | -0.3468 | -0.0375 | -0.0295 | 0.0017 | 0.0703 | 0.032 |
| I1 | 0.0493 | 0.3119 | 0.0151 | 0.0286 | -0.0122 | -0.1395 | -0.0328 | 0.1145 | 0.0895 | 0.1976 | -0.1747 |
| I2 | 0.0966 | 0.2585 | -0.0408 | 0.0429 | 0.0135 | -0.2713 | 0.0381 | 0.1443 | 0.0489 | 0.2117 | -0.0372 |
| I3 | 0.0278 | 0.3202 | -0.0362 | 0.0215 | -0.0075 | -0.3124 | 0.1161 | 0.1074 | 0.0535 | 0.2907 | -0.0031 |
| L1 | -0.0356 | 0.0843 | -0.173 | -0.0956 | 0.2236 | -0.1667 | -0.0437 | -0.0318 | -0.2768 | 0.0292 | -0.0914 |
| L2 | -0.0436 | -0.1971 | -0.2495 | -0.037 | -0.0631 | -0.3268 | -0.0005 | -0.0315 | -0.151 | -0.0624 | 0.1258 |
| P1 | -0.3009 | 0.0314 | -0.1074 | -0.0383 | -0.2777 | 0.2819 | 0.0146 | -0.0326 | -0.0084 | 0.086 | -0.087 |
| P2 | -0.1104 | 0.3252 | -0.0313 | -0.0014 | 0.1881 | 0.3238 | -0.0831 | 0.0206 | -0.0164 | 0.2192 | -0.2433 |
| tau13 | -0.2804 | 0.0311 | 0.0328 | -0.0739 | -0.2412 | 0.4086 | -0.0328 | -0.1096 | 0.0146 | 0.0568 | -0.0944 |
| tau21 | 0.2058 | 0.0932 | 0.1732 | 0.0227 | 0.2474 | 0.0435 | -0.1013 | -0.0577 | 0.0399 | -0.0159 | -0.0565 |
| tau23 | -0.2369 | 0.2502 | 0.1548 | -0.1311 | 0.0626 | 0.5994 | -0.1007 | -0.1441 | -0.0342 | 0.1674 | -0.2393 |
| tau31 | 0.2211 | -0.1775 | 0.246 | -0.0244 | 0.0072 | -0.2336 | 0.0086 | -0.0999 | 0.0313 | -0.1936 | 0.1441 |
| tau32 | 0.2138 | -0.1582 | 0.214 | -0.0307 | 0.0614 | -0.2325 | 0.0453 | -0.0849 | -0.0087 | -0.1909 | 0.1602 |

| response | EC | IDRE | TM | IDRC | Linker | Tail | IDRE posi | IDRC pos | RK cluste | GxxxG | CRAC |
|---|---|---|---|---|---|---|---|---|---|---|---|
| rD1(1) | 0.0211 | 0.1139 | 0.0549 | -0.2026 | 0.1268 | -0.0548 | 0.0699 | -0.1748 | -0.2272 | 0.1456 | 0.0351 |
| rD1(2) | 0.0359 | 0.0978 | 0.038 | 0.0334 | 0.0317 | 0.1173 | 0.0725 | 0.0241 | -0.0087 | 0.0502 | 0.1262 |
| rD1(3) | 0.009 | 0.0545 | 0.0719 | 0.0574 | -0.0083 | 0.05 | 0.0583 | 0.03 | -0.0213 | 0.0237 | 0.13 |
| rD2(1) | 0.0116 | 0.0546 | 0.0063 | -0.0433 | -0.0992 | 0.0146 | 0.0204 | -0.1099 | -0.2159 | 0.0695 | -0.0121 |
| rD2(2) | 0.0451 | -0.0118 | -0.0051 | 0.0436 | -0.1041 | 0.2213 | 0.0354 | -0.0087 | -0.036 | -0.0023 | 0.0716 |
| rD2(3) | -0.1045 | 0.0431 | 0.0481 | -0.067 | -0.0653 | 0.1438 | 0.0944 | -0.0545 | -0.1196 | -0.0391 | 0.0584 |
| rD3(1) | 0.1205 | 0.0571 | 0.0643 | -0.187 | 0.0195 | 0.1669 | 0.0648 | -0.2517 | -0.2456 | 0.1706 | -0.0541 |
| rD3(2) | -0.1059 | -0.0096 | -0.0314 | 0.0296 | -0.0864 | 0.2709 | 0.0207 | -0.0214 | -0.1207 | 0.0049 | 0.0188 |
| rD3(3) | -0.0436 | -0.0062 | -0.0075 | 0.061 | -0.081 | 0.2157 | 0.0821 | -0.0001 | -0.0609 | -0.0332 | -0.0092 |
| rF1(1) | -0.09 | -0.0792 | -0.044 | 0.1271 | 0.0199 | -0.1263 | -0.0192 | 0.1235 | 0.1619 | -0.039 | 0.0282 |
| rF1(2) | -0.0107 | 0.0456 | 0.008 | 0.0325 | 0.1085 | -0.1757 | -0.0153 | 0.0271 | 0.0717 | 0.1096 | 0.0152 |
| rF1(3) | -0.0114 | 0.0891 | -0.0131 | 0.0634 | 0.1433 | -0.1957 | 0.0156 | 0.0322 | 0.0717 | 0.1159 | -0.011 |
| rF2(1) | -0.2879 | -0.0564 | -0.0491 | 0.172 | -0.0563 | -0.1413 | 0.0706 | 0.0989 | 0.0595 | -0.0186 | 0.0341 |
| rF2(2) | -0.199 | 0.0857 | -0.0136 | 0.2171 | -0.0731 | -0.121 | 0.0952 | 0.0962 | 0.0423 | 0.1985 | 0.1849 |
| rF2(3) | -0.2086 | 0.188 | -0.0243 | 0.3093 | -0.0242 | -0.1451 | 0.1247 | 0.1452 | 0.0912 | 0.1894 | 0.1261 |
| rF3(1) | -0.3342 | 0.2633 | -0.0357 | 0.0295 | 0.0569 | -0.0179 | 0.2529 | -0.1091 | -0.2382 | 0.1971 | 0.0716 |
| rF3(2) | -0.2138 | 0.2394 | -0.1333 | 0.2286 | -0.1491 | 0.1219 | 0.247 | 0.0354 | -0.0926 | 0.1923 | 0.241 |
| rF3(3) | -0.1889 | 0.2607 | -0.0817 | 0.2372 | -0.0979 | 0.0396 | 0.2123 | 0.0307 | -0.0744 | 0.138 | 0.1766 |
| rI1(1) | -0.1956 | 0.0222 | 0.0251 | 0.1242 | 0.099 | -0.272 | 0.1635 | 0.0428 | 0.0166 | 0.0785 | 0.0697 |
| rI1(2) | -0.0128 | 0.2202 | 0.058 | 0.0954 | -0.0439 | -0.2482 | 0.1948 | -0.0121 | -0.0311 | 0.2259 | 0.0148 |
| rI1(3) | -0.0394 | 0.219 | -0.0019 | 0.0885 | 0.0819 | -0.3187 | 0.2343 | 0.0272 | -0.0013 | 0.274 | -0.016 |
| rI2(1) | -0.3003 | 0.0967 | 0.1378 | -0.0316 | 0.0924 | -0.0038 | 0.2233 | -0.1106 | -0.2308 | 0.0509 | 0.019 |
| rI2(2) | -0.1092 | 0.2693 | 0.1219 | 0.127 | -0.0262 | -0.0193 | 0.2639 | -0.0341 | -0.1303 | 0.2206 | 0.0804 |
| rI2(3) | -0.1465 | 0.289 | 0.1334 | 0.1703 | -0.0051 | -0.0866 | 0.2445 | 0.0273 | -0.0843 | 0.1993 | 0.0755 |
| rI3(1) | -0.3472 | -0.0773 | 0.0429 | -0.0066 | 0.0085 | 0.0326 | 0.1541 | -0.1046 | -0.286 | -0.0932 | 0.1228 |
| rI3(2) | -0.1473 | 0.2816 | -0.0733 | 0.0046 | -0.0154 | 0.0396 | 0.2754 | -0.1126 | -0.2999 | 0.1208 | 0.0215 |
| rI3(3) | -0.1529 | 0.3005 | -0.0449 | 0.0176 | 0.0107 | -0.0076 | 0.292 | -0.1107 | -0.2531 | 0.086 | -0.0155 |
| rL1(1) | 0.2102 | -0.061 | -0.1066 | 0.1519 | -0.0534 | -0.1129 | 0.1452 | 0.1449 | 0.1298 | -0.0525 | 0.2045 |
| rL1(2) | 0.1564 | -0.0505 | 0.0171 | 0.118 | -0.1186 | -0.0754 | 0.0637 | 0.1684 | 0.1244 | -0.0302 | 0.0463 |
| rL1(3) | 0.0442 | 0.0354 | -0.0188 | 0.1005 | -0.0691 | -0.0237 | 0.1933 | 0.1143 | 0.0507 | 0.0938 | 0.1725 |
| rL2(1) | 0.0167 | 0.096 | 0.0756 | -0.1187 | 0.1441 | 0.0025 | -0.0097 | 0.0046 | -0.0727 | 0.1663 | -0.0906 |
| rL2(2) | -0.0145 | 0.0547 | 0.1118 | 0.0097 | -0.0985 | 0.0951 | 0.0567 | 0.0312 | -0.0324 | 0.1781 | -0.0502 |
| rL2(3) | 0.0251 | -0.0997 | 0.023 | 0.0159 | -0.1493 | 0.0642 | 0.049 | -0.018 | -0.092 | -0.0015 | -0.0144 |
| rP1(1) | 0.0755 | -0.1582 | 0.0242 | 0.1117 | 0.0358 | -0.0136 | -0.2061 | 0.1439 | 0.2364 | -0.1545 | -0.0455 |
| rP1(2) | 0.0675 | -0.0402 | 0.0616 | -0.0781 | 0.1695 | -0.1487 | -0.1142 | -0.0273 | 0.0568 | -0.0285 | -0.1174 |
| rP1(3) | 0.0989 | -0.0311 | 0.0179 | -0.0637 | 0.1282 | -0.1739 | -0.056 | -0.0189 | 0.0622 | -0.0087 | -0.1129 |
| rP2(1) | 0.0926 | -0.3146 | -0.0183 | 0.1682 | -0.1901 | -0.1876 | -0.1254 | 0.1568 | 0.2329 | -0.1379 | 0.0112 |
| rP2(2) | 0.1482 | -0.1988 | 0.0507 | 0.0086 | -0.0041 | -0.2244 | -0.0817 | 0.0705 | 0.1206 | -0.0787 | -0.0692 |
| rP2(3) | 0.1023 | -0.1745 | 0.0152 | -0.0549 | 0.0054 | 0.0105 | -0.1532 | 0.0056 | 0.0971 | -0.0728 | -0.1053 |
| rtau13(1) | 0.0075 | -0.1188 | -0.0472 | 0.1096 | -0.052 | -0.0455 | -0.1272 | 0.1114 | 0.2229 | -0.1049 | -0.036 |
| rtau13(2) | 0.0394 | -0.0214 | 0.002 | -0.0989 | 0.0727 | -0.1424 | -0.0924 | -0.0608 | 0.0519 | 0.0021 | -0.1066 |
| rtau13(3) | 0.0464 | -0.0039 | -0.0261 | -0.1165 | 0.0644 | -0.1552 | -0.0213 | -0.0616 | 0.0453 | 0.0209 | -0.1309 |
| rtau21(1) | 0.1232 | 0.0212 | -0.0927 | -0.0197 | -0.2331 | -0.365 | 0.199 | -0.0679 | -0.1303 | 0.1729 | 0.0477 |
| rtau21(2) | 0.0974 | -0.1464 | -0.0166 | -0.133 | -0.2834 | -0.0808 | 0.0352 | -0.183 | -0.1509 | 0.1071 | 0.053 |
| rtau21(3) | -0.0148 | -0.145 | -0.0596 | -0.1199 | -0.2631 | -0.0514 | 0.0663 | -0.0842 | -0.0839 | 0.0457 | -0.0126 |
| rtau23(1) | 0.0613 | -0.1709 | -0.0667 | 0.1055 | -0.0987 | -0.0777 | -0.1528 | 0.1742 | 0.2975 | -0.0799 | 0.0121 |
| rtau23(2) | 0.1106 | -0.0515 | -0.0119 | -0.0595 | 0.0341 | -0.1993 | -0.0935 | 0.0219 | 0.1323 | -0.0274 | -0.0835 |
| rtau23(3) | 0.0803 | -0.0258 | -0.0248 | -0.1403 | 0.0752 | -0.0882 | -0.1016 | -0.0488 | 0.1044 | -0.0026 | -0.117 |
| rtau31(1) | -0.0393 | 0.2368 | -0.0472 | -0.1779 | 0.0348 | -0.0348 | 0.2471 | -0.1858 | -0.2729 | 0.2136 | 0.0262 |
| rtau31(2) | -0.1155 | 0.0509 | -0.0812 | -0.0488 | -0.204 | 0.2263 | 0.0561 | -0.1186 | -0.1476 | 0.1113 | 0.1248 |
| rtau31(3) | -0.1634 | 0.0594 | -0.0359 | -0.0438 | -0.1362 | 0.1383 | 0.098 | -0.0945 | -0.1569 | 0.0588 | 0.0996 |
| rtau32(1) | 0.001 | 0.2758 | -0.0169 | -0.2326 | 0.1422 | 0.0984 | 0.1028 | -0.1323 | -0.2295 | 0.2574 | 0.044 |
| rtau32(2) | -0.0602 | 0.0641 | -0.0686 | -0.0186 | -0.151 | 0.2562 | 0.0233 | -0.027 | -0.0739 | 0.0299 | 0.1073 |
| rtau32(3) | -0.0931 | 0.0581 | -0.0298 | -0.0772 | -0.0723 | 0.1768 | 0.0407 | -0.0817 | -0.1249 | -0.0005 | 0.1584 |
