## Supplement Table S5E for "Single-molecule behavior and cell-growth regulation in human RTKs"

| Resting | EG1 | EG2 | EG3 | EG4 | EG5 |
|---|---|---|---|---|---|
| D1 | 0.0483 | -0.2885 | 0.3422 | -0.2341 | 0.1744 |
| D2 | -0.0205 | -0.2134 | 0.327 | -0.2091 | 0.1318 |
| D3 | 0.0053 | -0.2203 | -0.0705 | 0.0309 | 0.253 |
| F1 | 0.0429 | 0.1698 | -0.0784 | 0.0351 | -0.1583 |
| F2 | -0.0137 | 0.0802 | 0.1629 | -0.0431 | -0.1863 |
| F3 | -0.0825 | -0.0874 | 0.3484 | -0.0539 | -0.1474 |
| I1 | 0.0277 | 0.179 | 0.0573 | -0.105 | -0.1379 |
| I2 | 0.0557 | 0.2108 | 0.132 | -0.1769 | -0.1829 |
| I3 | 0.0082 | 0.1984 | 0.1587 | -0.176 | -0.1672 |
| L1 | -0.1151 | 0.3276 | -0.0367 | -0.2123 | 0.0197 |
| L2 | -0.0764 | 0.1305 | -0.1034 | 0.1176 | -0.1077 |
| P1 | 0.0283 | 0.2354 | -0.3011 | 0.0983 | -0.0618 |
| P2 | -0.0221 | 0.0543 | -0.0089 | -0.2218 | 0.2151 |
| tau13 | -0.0176 | 0.0812 | -0.2408 | 0.1384 | 0.0176 |
| tau21 | -0.0728 | -0.314 | 0.3163 | -0.0842 | 0.1386 |
| tau23 | -0.0954 | -0.0117 | -0.1024 | -0.072 | 0.2562 |
| tau31 | -0.0535 | -0.2793 | 0.1757 | 0.186 | -0.0679 |
| tau32 | -0.0753 | -0.2322 | 0.1594 | 0.132 | -0.0245 |

| Respons | EG1 | EG2 | EG3 | EG4 | EG5 |
|---|---|---|---|---|---|
| rD1(1) | -0.3986 | 0.1469 | 0.1532 | -0.1274 | 0.1013 |
| rD1(2) | -0.0619 | 0.0885 | 0.1074 | -0.376 | 0.2616 |
| rD1(3) | -0.0963 | 0.1173 | 0.0909 | -0.3181 | 0.2074 |
| rD2(1) | -0.2999 | 0.1328 | 0.2148 | -0.2206 | 0.0926 |
| rD2(2) | -0.0312 | 0.0846 | 0.1635 | -0.3766 | 0.19 |
| rD2(3) | -0.049 | 0.2148 | -0.0982 | -0.3121 | 0.2615 |
| rD3(1) | -0.2176 | 0.2137 | 0.2132 | -0.315 | 0.0645 |
| rD3(2) | -0.0053 | 0.0789 | 0.1471 | -0.4262 | 0.2503 |
| rD3(3) | 0.0434 | 0.0804 | 0.1356 | -0.3731 | 0.1696 |
| rF1(1) | 0.2909 | -0.1972 | -0.121 | 0.1289 | -0.0146 |
| rF1(2) | 0.0651 | -0.1373 | -0.0387 | 0.1777 | -0.0636 |
| rF1(3) | 0.0228 | -0.1643 | 0.0145 | 0.1578 | -0.0401 |
| rF2(1) | 0.2285 | -0.1246 | -0.2398 | 0.0537 | 0.1558 |
| rF2(2) | 0.1273 | -0.0338 | -0.1805 | -0.1166 | 0.2606 |
| rF2(3) | 0.0604 | -0.0867 | -0.0408 | -0.1848 | 0.2931 |
| rF3(1) | -0.1026 | 0.1141 | -0.1504 | -0.3349 | 0.4748 |
| rF3(2) | 0.1023 | 0.0513 | -0.0442 | -0.515 | 0.4974 |
| rF3(3) | 0.0554 | -0.0328 | 0.0597 | -0.4584 | 0.4455 |
| rI1(1) | -0.0197 | -0.1909 | -0.1671 | 0.1648 | 0.188 |
| rI1(2) | -0.0319 | -0.0864 | 0.0758 | 0.0562 | -0.0309 |
| rI1(3) | -0.098 | -0.0661 | 0.1295 | -0.029 | 0.0327 |
| rI2(1) | -0.2614 | -0.0398 | -0.0646 | -0.1647 | 0.4576 |
| rI2(2) | -0.1685 | 0.1112 | 0.0928 | -0.4294 | 0.3823 |
| rI2(3) | -0.1918 | 0.0055 | 0.1416 | -0.4046 | 0.4269 |
| rI3(1) | -0.0525 | -0.0996 | -0.1888 | -0.1258 | 0.4622 |
| rI3(2) | -0.1957 | 0.1424 | -0.0603 | -0.2778 | 0.3538 |
| rI3(3) | -0.2174 | 0.049 | 0.0213 | -0.256 | 0.3556 |
| rL1(1) | 0.1788 | -0.1753 | 0.0595 | -0.0616 | 0.0675 |
| rL1(2) | 0.1484 | 0.0088 | 0.0907 | -0.1728 | -0.0046 |
| rL2(1) | 0.113 | 0.1183 | 0.0834 | -0.2859 | 0.0417 |
| rL2(2) | 0.1546 | 0.3607 | -0.0533 | -0.3805 | 0.0137 |
| rL1(3) | 0.1592 | 0.2027 | -0.0865 | -0.3337 | 0.1502 |
| rL2(3) | 0.0702 | 0.2455 | 0.0101 | -0.2973 | 0.0284 |
| rP1(1) | 0.2174 | -0.2697 | 0.0618 | 0.2287 | -0.1876 |
| rP1(2) | -0.0649 | -0.1392 | 0.0325 | 0.3442 | -0.2327 |
| rP1(3) | -0.0424 | -0.1197 | 0.0191 | 0.3424 | -0.2516 |
| rP2(1) | 0.1938 | -0.1108 | -0.0717 | 0.3692 | -0.3536 |
| rP2(2) | 0.026 | 0.0421 | -0.1658 | 0.3968 | -0.3335 |
| rP2(3) | 0.0004 | 0.0432 | -0.176 | 0.3665 | -0.2739 |
| rtau13(1) | 0.2813 | -0.2814 | -0.048 | 0.2771 | -0.1615 |
| rtau13(2) | 0.0087 | -0.1831 | -0.0501 | 0.4121 | -0.2295 |
| rtau13(3) | 0.0227 | -0.1626 | -0.0763 | 0.4122 | -0.2333 |
| rtau21(1) | -0.0433 | 0.2283 | -0.1284 | 0.0648 | -0.1435 |
| rtau21(2) | 0.1734 | 0.1366 | -0.281 | 0.0942 | -0.0732 |
| rtau21(3) | 0.1815 | 0.1134 | -0.3146 | 0.1133 | -0.0429 |
| rtau23(1) | 0.2877 | -0.1761 | -0.0656 | 0.2878 | -0.2652 |
| rtau23(2) | 0.0465 | -0.1416 | -0.0691 | 0.4317 | -0.2987 |
| rtau23(3) | 0.022 | -0.1424 | -0.1123 | 0.4319 | -0.2388 |
| rtau31(1) | -0.1928 | 0.2354 | -0.1161 | -0.2943 | 0.3331 |
| rtau31(2) | 0.157 | 0.0103 | -0.0232 | -0.3442 | 0.2923 |
| rtau31(3) | 0.1008 | -0.012 | -0.051 | -0.28 | 0.3079 |
| rtau32(1) | -0.2315 | 0.2593 | 0.0507 | -0.396 | 0.2803 |
| rtau32(2) | 0.1129 | -0.0289 | 0.0939 | -0.3731 | 0.2753 |
| rtau32(3) | 0.0963 | -0.0356 | 0.0402 | -0.3408 | 0.3106 |
