## Supplement Table S5F for "Single-molecule behavior and cell-growth regulation in human RTKs"

| R | EG1 | EG2 | EG3 | EG4 | EG5 |
|---|---|---|---|---|---|
| EC | 0.0748 | -0.0627 | 0.537 | -0.0245 | -0.496 |
| IDRE | -0.2087 | 0.0627 | -0.1262 | -0.076 | 0.284 |
| TM | -0.071 | 0.1738 | 0.0017 | -0.1567 | 0.0447 |
| IDRC | 0.2978 | -0.2802 | 0.258 | -0.2107 | 0.0616 |
| Linker | -0.3783 | -0.029 | -0.074 | 0.0341 | 0.312 |
| Tail | 0.2356 | -0.0866 | -0.1381 | -0.1586 | 0.247 |
| IDRE posi | -0.0779 | 0.0254 | -0.1549 | -0.0771 | 0.2658 |
| IDRC posi | 0.4572 | -0.1079 | 0.1396 | -0.2622 | -0.0391 |
| RK cluster | 0.6226 | -0.1098 | 0.1036 | -0.1655 | -0.2165 |
| GxxxG | 0.0827 | 0.359 | 0.0315 | -0.3869 | -0.0153 |
| CRAC | 0.1261 | 0.0123 | 0.0123 | -0.3004 | 0.2263 |
