## Supplement Table S6A for "Single-molecule behavior and cell-growth regulation in human RTKs"

resting state

| R | D1 | D2 | D3 | F1 | F2 | F3 | I1 | I2 | I3 | L1 | L2 | P1 | P2 | tau13 | tau21 | tau23 | tau31 | tau32 |
|---|---|---|---|---|---|---|---|---|---|---|---|---|---|---|---|---|---|---|
| D1 | 1 | 0.9106 | 0.3052 | -0.5406 | -0.1619 | 0.2717 | -0.1182 | -0.0436 | -0.1196 | -0.0017 | -0.2599 | -0.8717 | 0.1549 | -0.7074 | 0.8401 | -0.1517 | 0.3758 | 0.4066 |
| D2 | 0.9106 | 1 | 0.2985 | -0.5041 | -0.0513 | 0.4584 | -0.0148 | 0.1658 | 0.1132 | 0.0851 | -0.031 | -0.9288 | 0.0694 | -0.8471 | 0.7958 | -0.4299 | 0.3836 | 0.3613 |
| D3 | 0.3052 | 0.2985 | 1 | -0.7441 | -0.6489 | -0.2156 | -0.6017 | -0.463 | -0.3393 | -0.5408 | -0.0869 | -0.2013 | -0.476 | -0.0001 | 0.101 | -0.0427 | 0.4948 | 0.5782 |
| F1 | -0.5406 | -0.5041 | -0.7441 | 1 | 0.8139 | 0.2592 | 0.7241 | 0.5873 | 0.5422 | 0.4492 | 0.1684 | 0.4579 | 0.5255 | 0.2194 | -0.3662 | 0.1825 | -0.6579 | -0.7315 |
| F2 | -0.1619 | -0.0513 | -0.6489 | 0.8139 | 1 | 0.6362 | 0.804 | 0.7884 | 0.7401 | 0.4631 | 0.1649 | -0.0182 | 0.5704 | -0.223 | 0.0409 | -0.0765 | -0.4505 | -0.5727 |
| F3 | 0.2717 | 0.4584 | -0.2156 | 0.2592 | 0.6362 | 1 | 0.5034 | 0.6362 | 0.6558 | 0.2005 | 0.2165 | -0.5738 | 0.0571 | -0.6029 | 0.4156 | -0.4703 | 0.2439 | 0.1085 |
| I1 | -0.1182 | -0.0148 | -0.6017 | 0.7241 | 0.804 | 0.5034 | 1 | 0.8668 | 0.8034 | 0.4086 | -0.0324 | -0.022 | 0.5913 | -0.1909 | 0.0692 | -0.0285 | -0.4408 | -0.5483 |
| I2 | -0.0436 | 0.1658 | -0.463 | 0.5873 | 0.7884 | 0.6362 | 0.8668 | 1 | 0.9456 | 0.4574 | 0.2205 | -0.1638 | 0.4042 | -0.4238 | 0.0277 | -0.3793 | -0.3887 | -0.5041 |
| I3 | -0.1196 | 0.1132 | -0.3393 | 0.5422 | 0.7401 | 0.6558 | 0.8034 | 0.9456 | 1 | 0.3424 | 0.2704 | -0.0891 | 0.2949 | -0.3306 | -0.0485 | -0.3475 | -0.3398 | -0.4362 |
| L1 | -0.0017 | 0.0851 | -0.5408 | 0.4492 | 0.4631 | 0.2005 | 0.4086 | 0.4574 | 0.3424 | 1 | 0.4175 | -0.0648 | 0.4941 | -0.4124 | -0.1106 | -0.2024 | -0.6077 | -0.5374 |
| L2 | -0.2599 | -0.031 | -0.0869 | 0.1684 | 0.1649 | 0.2165 | -0.0324 | 0.2205 | 0.2704 | 0.4175 | 1 | 0.0656 | -0.1977 | -0.2359 | -0.3752 | -0.5942 | -0.226 | -0.2513 |
| P1 | -0.8717 | -0.9288 | -0.2013 | 0.4579 | -0.0182 | -0.5738 | -0.022 | -0.1638 | -0.0891 | -0.0648 | 0.0656 | 1 | 0.0157 | 0.8867 | -0.8335 | 0.4866 | -0.5317 | -0.4853 |
| P2 | 0.1549 | 0.0694 | -0.476 | 0.5255 | 0.5704 | 0.0571 | 0.5913 | 0.4042 | 0.2949 | 0.4941 | -0.1977 | 0.0157 | 1 | -0.0898 | 0.2701 | 0.4544 | -0.6986 | -0.7221 |
| tau13 | -0.7074 | -0.8471 | -0.0001 | 0.2194 | -0.223 | -0.6029 | -0.1909 | -0.4238 | -0.3306 | -0.4124 | -0.2359 | 0.8867 | -0.0898 | 1 | -0.577 | 0.6807 | -0.1955 | -0.1767 |
| tau21 | 0.8401 | 0.7958 | 0.101 | -0.3662 | 0.0409 | 0.4156 | 0.0692 | 0.0277 | -0.0485 | -0.1106 | -0.3752 | -0.8335 | 0.2701 | -0.577 | 1 | -0.033 | 0.4446 | 0.3372 |
| tau23 | -0.1517 | -0.4299 | -0.0427 | 0.1825 | -0.0765 | -0.4703 | -0.0285 | -0.3793 | -0.3475 | -0.2024 | -0.5942 | 0.4866 | 0.4544 | 0.6807 | -0.033 | 1 | -0.2833 | -0.1991 |
| tau31 | 0.3758 | 0.3836 | 0.4948 | -0.6579 | -0.4505 | 0.2439 | -0.4408 | -0.3887 | -0.3398 | -0.6077 | -0.226 | -0.5317 | -0.6986 | -0.1955 | 0.4446 | -0.2833 | 1 | 0.9432 |
| tau32 | 0.4066 | 0.3613 | 0.5782 | -0.7315 | -0.5727 | 0.1085 | -0.5483 | -0.5041 | -0.4362 | -0.5374 | -0.2513 | -0.4853 | -0.7221 | -0.1767 | 0.3372 | -0.1991 | 0.9432 | 1 |

response state

| R | rD1(1) | rD1(2) | rD1(3) | rD2(1) | rD2(2) | rD2(3) | rD3(1) | rD3(2) | rD3(3) | rF1(1) | rF1(2) | rF1(3) | rF2(1) | rF2(2) | rF2(3) | rF3(1) | rF3(2) | rF3(3) | rI1(1) | rI1(2) | rI1(3) | rI2(1) | rI2(2) | rI2(3) | rI3(1) | rI3(2) | rI3(3) |
|---|---|---|---|---|---|---|---|---|---|---|---|---|---|---|---|---|---|---|---|---|---|---|---|---|---|---|---|
| rD1(1) | 1 | 0.7059 | 0.7062 | 0.7349 | 0.5895 | 0.582 | 0.6528 | 0.4754 | 0.4463 | -0.7452 | -0.5616 | -0.4898 | -0.5376 | -0.3312 | -0.1533 | 0.2233 | 0.2862 | 0.363 | -0.4776 | -0.4609 | -0.312 | 0.1833 | 0.3015 | 0.3151 | 0.0701 | 0.4964 | 0.483 |
| rD1(2) | 0.7059 | 1 | 0.8851 | 0.6692 | 0.8633 | 0.8165 | 0.6268 | 0.774 | 0.7559 | -0.6749 | -0.8083 | -0.7429 | -0.4474 | -0.3874 | -0.1837 | 0.3681 | 0.5845 | 0.6478 | -0.5801 | -0.6544 | -0.5486 | 0.1505 | 0.4306 | 0.4361 | 0.1498 | 0.5074 | 0.4909 |
| rD1(3) | 0.7062 | 0.8851 | 1 | 0.599 | 0.8011 | 0.7586 | 0.4902 | 0.6678 | 0.6548 | -0.595 | -0.7198 | -0.6692 | -0.3872 | -0.3453 | -0.1441 | 0.3294 | 0.5376 | 0.5678 | -0.4843 | -0.5517 | -0.534 | 0.1385 | 0.3347 | 0.3306 | 0.1696 | 0.5178 | 0.4243 |
| rD2(1) | 0.7349 | 0.6692 | 0.599 | 1 | 0.7217 | 0.7802 | 0.7341 | 0.6573 | 0.5671 | -0.7514 | -0.6138 | -0.5605 | -0.4322 | -0.2322 | -0.1097 | 0.4218 | 0.4575 | 0.486 | -0.5839 | -0.3435 | -0.2864 | 0.3269 | 0.4883 | 0.4733 | 0.1986 | 0.5991 | 0.5824 |
| rD2(2) | 0.5895 | 0.8633 | 0.8011 | 0.7217 | 1 | 0.8524 | 0.6076 | 0.8202 | 0.7782 | -0.6242 | -0.7981 | -0.7636 | -0.3867 | -0.3704 | -0.2504 | 0.4021 | 0.6344 | 0.6371 | -0.6604 | -0.5857 | -0.5884 | 0.1048 | 0.3625 | 0.3216 | 0.1599 | 0.5276 | 0.4724 |
| rD2(3) | 0.582 | 0.8165 | 0.7586 | 0.7802 | 0.8524 | 1 | 0.6305 | 0.8047 | 0.7747 | -0.6026 | -0.7607 | -0.739 | -0.2908 | -0.2982 | -0.1918 | 0.5025 | 0.6102 | 0.6142 | -0.5578 | -0.5441 | -0.512 | 0.2888 | 0.42 | 0.3942 | 0.2876 | 0.5891 | 0.5385 |
| rD3(1) | 0.6528 | 0.6268 | 0.4902 | 0.7341 | 0.6076 | 0.6305 | 1 | 0.7205 | 0.6434 | -0.6379 | -0.5639 | -0.5428 | -0.412 | -0.315 | -0.2212 | 0.3989 | 0.4054 | 0.4061 | -0.5079 | -0.3444 | -0.3103 | 0.2511 | 0.4519 | 0.3454 | 0.1059 | 0.4994 | 0.4454 |
| rD3(2) | 0.4754 | 0.774 | 0.6678 | 0.6573 | 0.8202 | 0.8047 | 0.7205 | 1 | 0.9179 | -0.484 | -0.7593 | -0.7622 | -0.2169 | -0.3428 | -0.2638 | 0.4876 | 0.6429 | 0.6284 | -0.5098 | -0.5181 | -0.5336 | 0.2771 | 0.4537 | 0.3847 | 0.3182 | 0.5172 | 0.4591 |
| rD3(3) | 0.4463 | 0.7559 | 0.6548 | 0.5671 | 0.7782 | 0.7747 | 0.6434 | 0.9179 | 1 | -0.5023 | -0.8036 | -0.8013 | -0.2883 | -0.4461 | -0.3582 | 0.3584 | 0.5804 | 0.5706 | -0.5393 | -0.5353 | -0.5547 | 0.1931 | 0.3927 | 0.2945 | 0.2193 | 0.4327 | 0.3515 |
| rF1(1) | -0.7452 | -0.6749 | -0.595 | -0.7514 | -0.6242 | -0.6026 | -0.6379 | -0.484 | -0.5023 | 1 | 0.7649 | 0.683 | 0.8425 | 0.5656 | 0.3944 | -0.0944 | -0.2132 | -0.2692 | 0.7229 | 0.453 | 0.3598 | -0.0639 | -0.3614 | -0.3396 | 0.0891 | -0.5094 | -0.4928 |
| rF1(2) | -0.5616 | -0.8083 | -0.7198 | -0.6138 | -0.7981 | -0.7607 | -0.5639 | -0.7593 | -0.8036 | 0.7649 | 1 | 0.9703 | 0.5372 | 0.6781 | 0.5277 | -0.2522 | -0.4223 | -0.4646 | 0.6753 | 0.6814 | 0.6429 | -0.0907 | -0.2676 | -0.2725 | -0.1529 | -0.4584 | -0.4479 |
| rF1(3) | -0.4898 | -0.7429 | -0.6692 | -0.5605 | -0.7636 | -0.739 | -0.5428 | -0.7622 | -0.8013 | 0.683 | 0.9703 | 1 | 0.4469 | 0.6512 | 0.6063 | -0.2679 | -0.4272 | -0.4105 | 0.6543 | 0.6719 | 0.7147 | -0.0982 | -0.2379 | -0.1851 | -0.1726 | -0.4068 | -0.3449 |
| rF2(1) | -0.5376 | -0.4474 | -0.3872 | -0.4322 | -0.3867 | -0.2908 | -0.412 | -0.2169 | -0.2883 | 0.8425 | 0.5372 | 0.4469 | 1 | 0.7006 | 0.5491 | 0.3573 | 0.1608 | 0.1149 | 0.5915 | 0.2536 | 0.2068 | 0.2733 | -0.0611 | -0.0159 | 0.4056 | -0.2438 | -0.2153 |
| rF2(2) | -0.3312 | -0.3874 | -0.3453 | -0.2322 | -0.3704 | -0.2982 | -0.315 | -0.3428 | -0.4461 | 0.5656 | 0.6781 | 0.6512 | 0.7006 | 1 | 0.8513 | 0.2807 | 0.2448 | 0.1606 | 0.4362 | 0.3813 | 0.4207 | 0.2158 | 0.182 | 0.1571 | 0.1559 | -0.0943 | -0.1248 |
| rF2(3) | -0.1533 | -0.1837 | -0.1441 | -0.1097 | -0.2504 | -0.1918 | -0.2212 | -0.2638 | -0.3582 | 0.3944 | 0.5277 | 0.6063 | 0.5491 | 0.8513 | 1 | 0.324 | 0.2793 | 0.3541 | 0.3722 | 0.2789 | 0.4534 | 0.1995 | 0.2234 | 0.3654 | 0.1398 | 0.0118 | 0.1256 |
| rF3(1) | 0.2233 | 0.3681 | 0.3294 | 0.4218 | 0.4021 | 0.5025 | 0.3989 | 0.4876 | 0.3584 | -0.0944 | -0.2522 | -0.2679 | 0.3573 | 0.2807 | 0.324 | 1 | 0.8008 | 0.7997 | -0.1045 | -0.1374 | -0.1109 | 0.6434 | 0.6172 | 0.6313 | 0.5991 | 0.5855 | 0.588 |
| rF3(2) | 0.2862 | 0.5845 | 0.5376 | 0.4575 | 0.6344 | 0.6102 | 0.4054 | 0.6429 | 0.5804 | -0.2132 | -0.4223 | -0.4272 | 0.1608 | 0.2448 | 0.2793 | 0.8008 | 1 | 0.9234 | -0.3162 | -0.2865 | -0.2841 | 0.3645 | 0.5632 | 0.5156 | 0.4129 | 0.6193 | 0.5391 |
| rF3(3) | 0.363 | 0.6478 | 0.5678 | 0.486 | 0.6371 | 0.6142 | 0.4061 | 0.6284 | 0.5706 | -0.2692 | -0.4646 | -0.4105 | 0.1149 | 0.1606 | 0.3541 | 0.7997 | 0.9234 | 1 | -0.3294 | -0.3491 | -0.2478 | 0.3758 | 0.5607 | 0.6314 | 0.3972 | 0.6076 | 0.6592 |
| rI1(1) | -0.4776 | -0.5801 | -0.4843 | -0.5839 | -0.6604 | -0.5578 | -0.5079 | -0.5098 | -0.5393 | 0.7229 | 0.6753 | 0.6543 | 0.5915 | 0.4362 | 0.3722 | -0.1045 | -0.3162 | -0.3294 | 1 | 0.6543 | 0.6167 | 0.2228 | -0.1328 | -0.1035 | 0.2511 | -0.2646 | -0.2253 |
| rI1(2) | -0.4609 | -0.6544 | -0.5517 | -0.3435 | -0.5857 | -0.5441 | -0.3444 | -0.5181 | -0.5353 | 0.453 | 0.6814 | 0.6719 | 0.2536 | 0.3813 | 0.2789 | -0.1374 | -0.2865 | -0.3491 | 0.6543 | 1 | 0.8484 | 0.1076 | 0.0875 | -0.0167 | 0.0103 | -0.0147 | -0.0833 |
| rI1(3) | -0.312 | -0.5486 | -0.534 | -0.2864 | -0.5884 | -0.512 | -0.3103 | -0.5336 | -0.5547 | 0.3598 | 0.6429 | 0.7147 | 0.2068 | 0.4207 | 0.4534 | -0.1109 | -0.2841 | -0.2478 | 0.6167 | 0.8484 | 1 | 0.1316 | 0.143 | 0.1981 | 0.0013 | -0.0318 | 0.0502 |
| rI2(1) | 0.1833 | 0.1505 | 0.1385 | 0.3269 | 0.1048 | 0.2888 | 0.2511 | 0.2771 | 0.1931 | -0.0639 | -0.0907 | -0.0982 | 0.2733 | 0.2158 | 0.1995 | 0.6434 | 0.3645 | 0.3758 | 0.2228 | 0.1076 | 0.1316 | 1 | 0.6762 | 0.64 | 0.7838 | 0.4251 | 0.4467 |
| rI2(2) | 0.3015 | 0.4306 | 0.3347 | 0.4883 | 0.3625 | 0.42 | 0.4519 | 0.4537 | 0.3927 | -0.3614 | -0.2676 | -0.2379 | -0.0611 | 0.182 | 0.2234 | 0.6172 | 0.5632 | 0.5607 | -0.1328 | 0.0875 | 0.143 | 0.6762 | 1 | 0.865 | 0.3848 | 0.5778 | 0.5166 |
| rI2(3) | 0.3151 | 0.4361 | 0.3306 | 0.4733 | 0.3216 | 0.3942 | 0.3454 | 0.3847 | 0.2945 | -0.3396 | -0.2725 | -0.1851 | -0.0159 | 0.1571 | 0.3654 | 0.6313 | 0.5156 | 0.6314 | -0.1035 | -0.0167 | 0.1981 | 0.64 | 0.865 | 1 | 0.4006 | 0.4584 | 0.6029 |
| rI3(1) | 0.0701 | 0.1498 | 0.1696 | 0.1986 | 0.1599 | 0.2876 | 0.1059 | 0.3182 | 0.2193 | 0.0891 | -0.1529 | -0.1726 | 0.4056 | 0.1559 | 0.1398 | 0.5991 | 0.4129 | 0.3972 | 0.2511 | 0.0103 | 0.0013 | 0.7838 | 0.3848 | 0.4006 | 1 | 0.4272 | 0.4453 |
| rI3(2) | 0.4964 | 0.5074 | 0.5178 | 0.5991 | 0.5276 | 0.5891 | 0.4994 | 0.5172 | 0.4327 | -0.5094 | -0.4584 | -0.4068 | -0.2438 | -0.0943 | 0.0118 | 0.5855 | 0.6193 | 0.6076 | -0.2646 | -0.0147 | -0.0318 | 0.4251 | 0.5778 | 0.4584 | 0.4272 | 1 | 0.8706 |
| rI3(3) | 0.483 | 0.4909 | 0.4243 | 0.5824 | 0.4724 | 0.5385 | 0.4454 | 0.4591 | 0.3515 | -0.4928 | -0.4479 | -0.3449 | -0.2153 | -0.1248 | 0.1256 | 0.588 | 0.5391 | 0.6592 | -0.2253 | -0.0833 | 0.0502 | 0.4467 | 0.5166 | 0.6029 | 0.4453 | 0.8706 | 1 |
| rL1(1) | -0.1253 | -0.0143 | -0.0348 | -0.1035 | -0.0309 | -0.0565 | -0.1477 | -0.147 | -0.0941 | -0.0407 | -0.0672 | -0.0325 | -0.0136 | -0.0166 | 0.0334 | 0.0199 | 0.0587 | 0.1057 | -0.0414 | -0.0285 | 0.0818 | -0.1039 | -0.0955 | 0.0279 | 0.052 | 0.0401 | 0.0707 |
| rL1(2) | 0.3318 | 0.3755 | 0.3422 | 0.3177 | 0.3218 | 0.3573 | 0.152 | 0.242 | 0.2757 | -0.2522 | -0.4027 | -0.3662 | 0.0231 | -0.0454 | 0.1008 | 0.3186 | 0.4013 | 0.4934 | -0.3392 | -0.41 | -0.2499 | 0.0908 | 0.0449 | 0.1881 | 0.2109 | 0.3594 | 0.4478 |
| rL1(3) | 0.362 | 0.6127 | 0.611 | 0.4384 | 0.6165 | 0.5739 | 0.4408 | 0.5544 | 0.573 | -0.3603 | -0.6014 | -0.581 | -0.0879 | -0.1954 | -0.0907 | 0.3357 | 0.5226 | 0.5228 | -0.4897 | -0.628 | -0.5796 | 0.0331 | 0.1065 | 0.1349 | 0.2088 | 0.2945 | 0.2236 |
| rL2(1) | 0.0939 | 0.1674 | 0.1444 | 0.062 | 0.2155 | 0.1332 | 0.1416 | 0.1643 | 0.2102 | -0.1497 | -0.235 | -0.2072 | -0.0618 | -0.0943 | -0.0789 | 0.15 | 0.2166 | 0.2358 | -0.2761 | -0.2913 | -0.2345 | -0.0027 | -0.0376 | -0.0096 | 0.0252 | 0.0828 | 0.0356 |
| rL2(2) | 0.3331 | 0.535 | 0.5535 | 0.3739 | 0.5136 | 0.4983 | 0.4435 | 0.5092 | 0.5222 | -0.3614 | -0.5324 | -0.5173 | -0.0947 | -0.1027 | -0.0226 | 0.3629 | 0.5275 | 0.4982 | -0.5133 | -0.5347 | -0.4792 | -0.0425 | 0.116 | 0.1198 | 0.1147 | 0.3168 | 0.213 |
| rL2(3) | 0.3411 | 0.5341 | 0.5544 | 0.3162 | 0.4682 | 0.5028 | 0.2889 | 0.3756 | 0.4326 | -0.3098 | -0.4831 | -0.4666 | -0.0558 | -0.0876 | 0.0458 | 0.3949 | 0.5662 | 0.573 | -0.39 | -0.4536 | -0.4115 | 0.0134 | 0.1217 | 0.1837 | 0.1986 | 0.4297 | 0.385 |
| rP1(1) | -0.7551 | -0.7409 | -0.6815 | -0.7779 | -0.6549 | -0.7527 | -0.697 | -0.6016 | -0.5652 | 0.8453 | 0.7668 | 0.7179 | 0.5092 | 0.3576 | 0.1908 | -0.4703 | -0.504 | -0.5399 | 0.5787 | 0.4461 | 0.331 | -0.2714 | -0.501 | -0.5153 | -0.1935 | -0.6781 | -0.6729 |
| rP1(2) | -0.5119 | -0.837 | -0.7696 | -0.6229 | -0.8276 | -0.8345 | -0.5531 | -0.8116 | -0.8102 | 0.625 | 0.9059 | 0.882 | 0.3124 | 0.3705 | 0.2442 | -0.4546 | -0.6956 | -0.6882 | 0.5855 | 0.6124 | 0.5513 | -0.1665 | -0.3785 | -0.392 | -0.2613 | -0.5569 | -0.5235 |
| rP1(3) | -0.5223 | -0.8548 | -0.7916 | -0.5965 | -0.8239 | -0.8359 | -0.5654 | -0.848 | -0.827 | 0.5824 | 0.8979 | 0.8855 | 0.2643 | 0.3938 | 0.2374 | -0.4871 | -0.6803 | -0.7036 | 0.56 | 0.6549 | 0.6057 | -0.1707 | -0.3585 | -0.389 | -0.2759 | -0.5348 | -0.514 |
| rP2(1) | -0.2498 | -0.4774 | -0.389 | -0.3669 | -0.499 | -0.4728 | -0.4456 | -0.5574 | -0.4863 | 0.3937 | 0.4453 | 0.419 | 0.2588 | 0.2037 | 0.1218 | -0.5792 | -0.5459 | -0.5591 | 0.3914 | 0.2043 | 0.2246 | -0.3342 | -0.5396 | -0.4621 | -0.1478 | -0.5278 | -0.4916 |
| rP2(2) | -0.1431 | -0.4796 | -0.3906 | -0.2265 | -0.4984 | -0.4023 | -0.3859 | -0.6828 | -0.6661 | 0.2166 | 0.5071 | 0.5328 | 0.0465 | 0.2331 | 0.1697 | -0.5358 | -0.6314 | -0.6304 | 0.3153 | 0.3083 | 0.3734 | -0.29 | -0.4278 | -0.3884 | -0.2256 | -0.321 | -0.3066 |
| rP2(3) | -0.085 | -0.3951 | -0.262 | -0.1939 | -0.4107 | -0.3251 | -0.2866 | -0.5774 | -0.5624 | 0.1268 | 0.4035 | 0.4292 | -0.0786 | 0.0904 | 0.1171 | -0.5865 | -0.6121 | -0.634 | 0.2701 | 0.254 | 0.2686 | -0.3752 | -0.481 | -0.4479 | -0.2792 | -0.2763 | -0.2853 |
| rtau13(1) | -0.7708 | -0.712 | -0.6636 | -0.7681 | -0.6357 | -0.6718 | -0.6649 | -0.5164 | -0.4769 | 0.8797 | 0.6892 | 0.6224 | 0.5752 | 0.3129 | 0.1416 | -0.4073 | -0.4324 | -0.4832 | 0.6519 | 0.4617 | 0.3093 | -0.2535 | -0.5261 | -0.5282 | -0.1306 | -0.6252 | -0.6261 |
| rtau13(2) | -0.5886 | -0.8673 | -0.8064 | -0.6414 | -0.8262 | -0.805 | -0.5707 | -0.7699 | -0.7697 | 0.6634 | 0.8592 | 0.8056 | 0.351 | 0.3155 | 0.143 | -0.4398 | -0.6731 | -0.6993 | 0.6465 | 0.6719 | 0.5379 | -0.1658 | -0.413 | -0.4369 | -0.2368 | -0.5606 | -0.5485 |
| rtau13(3) | -0.5942 | -0.8711 | -0.8452 | -0.6214 | -0.8289 | -0.7996 | -0.57 | -0.7819 | -0.7636 | 0.61 | 0.8319 | 0.7868 | 0.2945 | 0.3024 | 0.1035 | -0.4565 | -0.6672 | -0.7073 | 0.6118 | 0.6818 | 0.5715 | -0.1574 | -0.3834 | -0.422 | -0.2492 | -0.5363 | -0.5187 |
| rtau21(1) | 0.4291 | 0.3408 | 0.3485 | 0.4679 | 0.2307 | 0.4045 | 0.3329 | 0.1737 | 0.2075 | -0.5138 | -0.4175 | -0.3928 | -0.3466 | -0.2672 | -0.1774 | 0.0592 | 0.1055 | 0.117 | -0.2015 | -0.1174 | -0.0772 | -0.0286 | 0.1289 | 0.1952 | -0.0018 | 0.3291 | 0.3032 |
| rtau21(2) | 0.2196 | 0.295 | 0.3229 | 0.3345 | 0.3682 | 0.4497 | 0.187 | 0.2438 | 0.2545 | -0.3277 | -0.4752 | -0.5046 | -0.1697 | -0.2608 | -0.3262 | 0.0983 | 0.1951 | 0.1126 | -0.2141 | -0.2241 | -0.2912 | -0.0438 | -0.022 | 0 | 0.1565 | 0.2402 | 0.1557 |
| rtau21(3) | 0.1187 | 0.2104 | 0.2201 | 0.1987 | 0.2691 | 0.4158 | 0.1062 | 0.2733 | 0.2752 | -0.1891 | -0.4455 | -0.492 | -0.0563 | -0.3321 | -0.3299 | 0.1008 | 0.1141 | 0.0965 | -0.0679 | -0.2154 | -0.2762 | -0.0558 | -0.1118 | -0.0283 | 0.1246 | 0.1721 | 0.1702 |
| rtau23(1) | -0.6676 | -0.7209 | -0.6658 | -0.756 | -0.7055 | -0.7059 | -0.6762 | -0.6201 | -0.5478 | 0.7797 | 0.6829 | 0.6209 | 0.4549 | 0.27 | 0.1125 | -0.5998 | -0.5703 | -0.6208 | 0.6218 | 0.427 | 0.3385 | -0.365 | -0.6215 | -0.6023 | -0.2425 | -0.7022 | -0.6933 |
| rtau23(2) | -0.5437 | -0.8496 | -0.7871 | -0.5998 | -0.8409 | -0.7697 | -0.5823 | -0.8247 | -0.8153 | 0.5769 | 0.8217 | 0.7915 | 0.2448 | 0.2694 | 0.1275 | -0.5517 | -0.7604 | -0.7673 | 0.6182 | 0.6685 | 0.5869 | -0.2356 | -0.457 | -0.459 | -0.3148 | -0.5446 | -0.5169 |
| rtau23(3) | -0.4926 | -0.7944 | -0.7304 | -0.5621 | -0.7844 | -0.7271 | -0.5184 | -0.7888 | -0.7782 | 0.4979 | 0.782 | 0.7548 | 0.1351 | 0.2133 | 0.07 | -0.5954 | -0.7483 | -0.7916 | 0.5802 | 0.6529 | 0.549 | -0.2714 | -0.4658 | -0.5055 | -0.3476 | -0.5032 | -0.5086 |
| rtau31(1) | 0.4712 | 0.5548 | 0.4657 | 0.6049 | 0.513 | 0.6627 | 0.6113 | 0.5711 | 0.5219 | -0.5931 | -0.5511 | -0.5263 | -0.3495 | -0.2056 | -0.1165 | 0.5615 | 0.5364 | 0.5279 | -0.3832 | -0.2187 | -0.1797 | 0.3301 | 0.583 | 0.5433 | 0.144 | 0.6356 | 0.5854 |
| rtau31(2) | 0.1909 | 0.4873 | 0.4233 | 0.3567 | 0.558 | 0.5777 | 0.4005 | 0.7204 | 0.7132 | -0.267 | -0.6138 | -0.6713 | -0.0325 | -0.2384 | -0.257 | 0.4927 | 0.6546 | 0.5766 | -0.329 | -0.3419 | -0.4322 | 0.2558 | 0.3255 | 0.2855 | 0.3206 | 0.3665 | 0.2894 |
| rtau31(3) | 0.1007 | 0.4015 | 0.2503 | 0.2348 | 0.4482 | 0.4692 | 0.2953 | 0.653 | 0.6444 | -0.1653 | -0.5409 | -0.6001 | 0.0769 | -0.2133 | -0.2472 | 0.5237 | 0.5671 | 0.5528 | -0.2318 | -0.3263 | -0.3641 | 0.3087 | 0.349 | 0.3405 | 0.3424 | 0.2961 | 0.2904 |
| rtau32(1) | 0.4916 | 0.5466 | 0.4551 | 0.5602 | 0.4807 | 0.5631 | 0.6085 | 0.574 | 0.5313 | -0.6146 | -0.5191 | -0.4914 | -0.4534 | -0.2707 | -0.1641 | 0.482 | 0.4807 | 0.4784 | -0.459 | -0.2196 | -0.1949 | 0.3106 | 0.5713 | 0.4937 | 0.0675 | 0.5784 | 0.5147 |
| rtau32(2) | 0.2528 | 0.5665 | 0.4652 | 0.3826 | 0.5868 | 0.5859 | 0.4348 | 0.7604 | 0.7661 | -0.3627 | -0.6985 | -0.7298 | -0.129 | -0.3293 | -0.2814 | 0.4782 | 0.6412 | 0.6262 | -0.431 | -0.4387 | -0.47 | 0.2896 | 0.3759 | 0.3619 | 0.2756 | 0.3788 | 0.3633 |
| rtau32(3) | 0.235 | 0.5528 | 0.44 | 0.2983 | 0.5551 | 0.5526 | 0.4169 | 0.7522 | 0.7623 | -0.2735 | -0.6403 | -0.689 | -0.0636 | -0.314 | -0.3248 | 0.4962 | 0.634 | 0.6108 | -0.373 | -0.4278 | -0.4812 | 0.2709 | 0.3414 | 0.2901 | 0.3015 | 0.367 | 0.3141 |

| R | rL1(1) | rL1(2) | rL1(3) | rL2(1) | rL2(2) | rL2(3) | rP1(1) | rP1(2) | rP1(3) | rP2(1) | rP2(2) | rP2(3) | rtau13(1) | rtau13(2) | rtau13(3) | rtau21(1) | rtau21(2) | rtau21(3) | rtau23(1) | rtau23(2) | rtau23(3) | rtau31(1) | rtau31(2) | rtau31(3) | rtau32(1) | rtau32(2) | rtau32(3) |
|---|---|---|---|---|---|---|---|---|---|---|---|---|---|---|---|---|---|---|---|---|---|---|---|---|---|---|---|
| rD1(1) | -0.1253 | 0.3318 | 0.362 | 0.0939 | 0.3331 | 0.3411 | -0.7551 | -0.5119 | -0.5223 | -0.2498 | -0.1431 | -0.085 | -0.7708 | -0.5886 | -0.5942 | 0.4291 | 0.2196 | 0.1187 | -0.6676 | -0.5437 | -0.4926 | 0.4712 | 0.1909 | 0.1007 | 0.4916 | 0.2528 | 0.235 |
| rD1(2) | -0.0143 | 0.3755 | 0.6127 | 0.1674 | 0.535 | 0.5341 | -0.7409 | -0.837 | -0.8548 | -0.4774 | -0.4796 | -0.3951 | -0.712 | -0.8673 | -0.8711 | 0.3408 | 0.295 | 0.2104 | -0.7209 | -0.8496 | -0.7944 | 0.5548 | 0.4873 | 0.4015 | 0.5466 | 0.5665 | 0.5528 |
| rD1(3) | -0.0348 | 0.3422 | 0.611 | 0.1444 | 0.5535 | 0.5544 | -0.6815 | -0.7696 | -0.7916 | -0.389 | -0.3906 | -0.262 | -0.6636 | -0.8064 | -0.8452 | 0.3485 | 0.3229 | 0.2201 | -0.6658 | -0.7871 | -0.7304 | 0.4657 | 0.4233 | 0.2503 | 0.4551 | 0.4652 | 0.44 |
| rD2(1) | -0.1035 | 0.3177 | 0.4384 | 0.062 | 0.3739 | 0.3162 | -0.7779 | -0.6229 | -0.5965 | -0.3669 | -0.2265 | -0.1939 | -0.7681 | -0.6414 | -0.6214 | 0.4679 | 0.3345 | 0.1987 | -0.756 | -0.5998 | -0.5621 | 0.6049 | 0.3567 | 0.2348 | 0.5602 | 0.3826 | 0.2983 |
| rD2(2) | -0.0309 | 0.3218 | 0.6165 | 0.2155 | 0.5136 | 0.4682 | -0.6549 | -0.8276 | -0.8239 | -0.499 | -0.4984 | -0.4107 | -0.6357 | -0.8262 | -0.8289 | 0.2307 | 0.3682 | 0.2691 | -0.7055 | -0.8409 | -0.7844 | 0.513 | 0.558 | 0.4482 | 0.4807 | 0.5868 | 0.5551 |
| rD2(3) | -0.0565 | 0.3573 | 0.5739 | 0.1332 | 0.4983 | 0.5028 | -0.7527 | -0.8345 | -0.8359 | -0.4728 | -0.4023 | -0.3251 | -0.6718 | -0.805 | -0.7996 | 0.4045 | 0.4497 | 0.4158 | -0.7059 | -0.7697 | -0.7271 | 0.6627 | 0.5777 | 0.4692 | 0.5631 | 0.5859 | 0.5526 |
| rD3(1) | -0.1477 | 0.152 | 0.4408 | 0.1416 | 0.4435 | 0.2889 | -0.697 | -0.5531 | -0.5654 | -0.4456 | -0.3859 | -0.2866 | -0.6649 | -0.5707 | -0.57 | 0.3329 | 0.187 | 0.1062 | -0.6762 | -0.5823 | -0.5184 | 0.6113 | 0.4005 | 0.2953 | 0.6085 | 0.4348 | 0.4169 |
| rD3(2) | -0.147 | 0.242 | 0.5544 | 0.1643 | 0.5092 | 0.3756 | -0.6016 | -0.8116 | -0.848 | -0.5574 | -0.6828 | -0.5774 | -0.5164 | -0.7699 | -0.7819 | 0.1737 | 0.2438 | 0.2733 | -0.6201 | -0.8247 | -0.7888 | 0.5711 | 0.7204 | 0.653 | 0.574 | 0.7604 | 0.7522 |
| rD3(3) | -0.0941 | 0.2757 | 0.573 | 0.2102 | 0.5222 | 0.4326 | -0.5652 | -0.8102 | -0.827 | -0.4863 | -0.6661 | -0.5624 | -0.4769 | -0.7697 | -0.7636 | 0.2075 | 0.2545 | 0.2752 | -0.5478 | -0.8153 | -0.7782 | 0.5219 | 0.7132 | 0.6444 | 0.5313 | 0.7661 | 0.7623 |
| rF1(1) | -0.0407 | -0.2522 | -0.3603 | -0.1497 | -0.3614 | -0.3098 | 0.8453 | 0.625 | 0.5824 | 0.3937 | 0.2166 | 0.1268 | 0.8797 | 0.6634 | 0.61 | -0.5138 | -0.3277 | -0.1891 | 0.7797 | 0.5769 | 0.4979 | -0.5931 | -0.267 | -0.1653 | -0.6146 | -0.3627 | -0.2735 |
| rF1(2) | -0.0672 | -0.4027 | -0.6014 | -0.235 | -0.5324 | -0.4831 | 0.7668 | 0.9059 | 0.8979 | 0.4453 | 0.5071 | 0.4035 | 0.6892 | 0.8592 | 0.8319 | -0.4175 | -0.4752 | -0.4455 | 0.6829 | 0.8217 | 0.782 | -0.5511 | -0.6138 | -0.5409 | -0.5191 | -0.6985 | -0.6403 |
| rF1(3) | -0.0325 | -0.3662 | -0.581 | -0.2072 | -0.5173 | -0.4666 | 0.7179 | 0.882 | 0.8855 | 0.419 | 0.5328 | 0.4292 | 0.6224 | 0.8056 | 0.7868 | -0.3928 | -0.5046 | -0.492 | 0.6209 | 0.7915 | 0.7548 | -0.5263 | -0.6713 | -0.6001 | -0.4914 | -0.7298 | -0.689 |
| rF2(1) | -0.0136 | 0.0231 | -0.0879 | -0.0618 | -0.0947 | -0.0558 | 0.5092 | 0.3124 | 0.2643 | 0.2588 | 0.0465 | -0.0786 | 0.5752 | 0.351 | 0.2945 | -0.3466 | -0.1697 | -0.0563 | 0.4549 | 0.2448 | 0.1351 | -0.3495 | -0.0325 | 0.0769 | -0.4534 | -0.129 | -0.0636 |
| rF2(2) | -0.0166 | -0.0454 | -0.1954 | -0.0943 | -0.1027 | -0.0876 | 0.3576 | 0.3705 | 0.3938 | 0.2037 | 0.2331 | 0.0904 | 0.3129 | 0.3155 | 0.3024 | -0.2672 | -0.2608 | -0.3321 | 0.27 | 0.2694 | 0.2133 | -0.2056 | -0.2384 | -0.2133 | -0.2707 | -0.3293 | -0.314 |
| rF2(3) | 0.0334 | 0.1008 | -0.0907 | -0.0789 | -0.0226 | 0.0458 | 0.1908 | 0.2442 | 0.2374 | 0.1218 | 0.1697 | 0.1171 | 0.1416 | 0.143 | 0.1035 | -0.1774 | -0.3262 | -0.3299 | 0.1125 | 0.1275 | 0.07 | -0.1165 | -0.257 | -0.2472 | -0.1641 | -0.2814 | -0.3248 |
| rF3(1) | 0.0199 | 0.3186 | 0.3357 | 0.15 | 0.3629 | 0.3949 | -0.4703 | -0.4546 | -0.4871 | -0.5792 | -0.5358 | -0.5865 | -0.4073 | -0.4398 | -0.4565 | 0.0592 | 0.0983 | 0.1008 | -0.5998 | -0.5517 | -0.5954 | 0.5615 | 0.4927 | 0.5237 | 0.482 | 0.4782 | 0.4962 |
| rF3(2) | 0.0587 | 0.4013 | 0.5226 | 0.2166 | 0.5275 | 0.5662 | -0.504 | -0.6956 | -0.6803 | -0.5459 | -0.6314 | -0.6121 | -0.4324 | -0.6731 | -0.6672 | 0.1055 | 0.1951 | 0.1141 | -0.5703 | -0.7604 | -0.7483 | 0.5364 | 0.6546 | 0.5671 | 0.4807 | 0.6412 | 0.634 |
| rF3(3) | 0.1057 | 0.4934 | 0.5228 | 0.2358 | 0.4982 | 0.573 | -0.5399 | -0.6882 | -0.7036 | -0.5591 | -0.6304 | -0.634 | -0.4832 | -0.6993 | -0.7073 | 0.117 | 0.1126 | 0.0965 | -0.6208 | -0.7673 | -0.7916 | 0.5279 | 0.5766 | 0.5528 | 0.4784 | 0.6262 | 0.6108 |
| rI1(1) | -0.0414 | -0.3392 | -0.4897 | -0.2761 | -0.5133 | -0.39 | 0.5787 | 0.5855 | 0.56 | 0.3914 | 0.3153 | 0.2701 | 0.6519 | 0.6465 | 0.6118 | -0.2015 | -0.2141 | -0.0679 | 0.6218 | 0.6182 | 0.5802 | -0.3832 | -0.329 | -0.2318 | -0.459 | -0.431 | -0.373 |
| rI1(2) | -0.0285 | -0.41 | -0.628 | -0.2913 | -0.5347 | -0.4536 | 0.4461 | 0.6124 | 0.6549 | 0.2043 | 0.3083 | 0.254 | 0.4617 | 0.6719 | 0.6818 | -0.1174 | -0.2241 | -0.2154 | 0.427 | 0.6685 | 0.6529 | -0.2187 | -0.3419 | -0.3263 | -0.2196 | -0.4387 | -0.4278 |
| rI1(3) | 0.0818 | -0.2499 | -0.5796 | -0.2345 | -0.4792 | -0.4115 | 0.331 | 0.5513 | 0.6057 | 0.2246 | 0.3734 | 0.2686 | 0.3093 | 0.5379 | 0.5715 | -0.0772 | -0.2912 | -0.2762 | 0.3385 | 0.5869 | 0.549 | -0.1797 | -0.4322 | -0.3641 | -0.1949 | -0.47 | -0.4812 |
| rI2(1) | -0.1039 | 0.0908 | 0.0331 | -0.0027 | -0.0425 | 0.0134 | -0.2714 | -0.1665 | -0.1707 | -0.3342 | -0.29 | -0.3752 | -0.2535 | -0.1658 | -0.1574 | -0.0286 | -0.0438 | -0.0558 | -0.365 | -0.2356 | -0.2714 | 0.3301 | 0.2558 | 0.3087 | 0.3106 | 0.2896 | 0.2709 |
| rI2(2) | -0.0955 | 0.0449 | 0.1065 | -0.0376 | 0.116 | 0.1217 | -0.501 | -0.3785 | -0.3585 | -0.5396 | -0.4278 | -0.481 | -0.5261 | -0.413 | -0.3834 | 0.1289 | -0.022 | -0.1118 | -0.6215 | -0.457 | -0.4658 | 0.583 | 0.3255 | 0.349 | 0.5713 | 0.3759 | 0.3414 |
| rI2(3) | 0.0279 | 0.1881 | 0.1349 | -0.0096 | 0.1198 | 0.1837 | -0.5153 | -0.392 | -0.389 | -0.4621 | -0.3884 | -0.4479 | -0.5282 | -0.4369 | -0.422 | 0.1952 | 0 | -0.0283 | -0.6023 | -0.459 | -0.5055 | 0.5433 | 0.2855 | 0.3405 | 0.4937 | 0.3619 | 0.2901 |
| rI3(1) | 0.052 | 0.2109 | 0.2088 | 0.0252 | 0.1147 | 0.1986 | -0.1935 | -0.2613 | -0.2759 | -0.1478 | -0.2256 | -0.2792 | -0.1306 | -0.2368 | -0.2492 | -0.0018 | 0.1565 | 0.1246 | -0.2425 | -0.3148 | -0.3476 | 0.144 | 0.3206 | 0.3424 | 0.0675 | 0.2756 | 0.3015 |
| rI3(2) | 0.0401 | 0.3594 | 0.2945 | 0.0828 | 0.3168 | 0.4297 | -0.6781 | -0.5569 | -0.5348 | -0.5278 | -0.321 | -0.2763 | -0.6252 | -0.5606 | -0.5363 | 0.3291 | 0.2402 | 0.1721 | -0.7022 | -0.5446 | -0.5032 | 0.6356 | 0.3665 | 0.2961 | 0.5784 | 0.3788 | 0.367 |
| rI3(3) | 0.0707 | 0.4478 | 0.2236 | 0.0356 | 0.213 | 0.385 | -0.6729 | -0.5235 | -0.514 | -0.4916 | -0.3066 | -0.2853 | -0.6261 | -0.5485 | -0.5187 | 0.3032 | 0.1557 | 0.1702 | -0.6933 | -0.5169 | -0.5086 | 0.5854 | 0.2894 | 0.2904 | 0.5147 | 0.3633 | 0.3141 |
| rL1(1) | 1 | 0.5054 | 0.1709 | 0.1834 | 0.2255 | 0.5206 | -0.1158 | -0.1337 | -0.092 | 0.0989 | 0.0551 | -0.0132 | -0.1223 | -0.1522 | -0.1188 | 0.1718 | 0.1766 | 0.1215 | -0.0321 | -0.091 | -0.1315 | -0.0108 | 0.0502 | 0.064 | -0.0496 | 0.0694 | 0.0588 |
| rL1(2) | 0.5054 | 1 | 0.4301 | 0.213 | 0.4994 | 0.7506 | -0.4387 | -0.5081 | -0.4979 | -0.0031 | -0.0662 | -0.1187 | -0.4144 | -0.5604 | -0.5464 | 0.2846 | 0.2878 | 0.2559 | -0.3238 | -0.4737 | -0.5436 | 0.1413 | 0.2352 | 0.2285 | 0.1224 | 0.3399 | 0.2901 |
| rL1(3) | 0.1709 | 0.4301 | 1 | 0.5519 | 0.8681 | 0.6168 | -0.4805 | -0.6743 | -0.6705 | -0.2098 | -0.3776 | -0.3311 | -0.4406 | -0.6887 | -0.6865 | 0.3274 | 0.4064 | 0.2883 | -0.4335 | -0.7204 | -0.7092 | 0.3148 | 0.5135 | 0.3965 | 0.2288 | 0.4813 | 0.4528 |
| rL2(1) | 0.1834 | 0.213 | 0.5519 | 1 | 0.6077 | 0.228 | -0.1215 | -0.2408 | -0.211 | -0.2403 | -0.2502 | -0.3683 | -0.1478 | -0.2618 | -0.222 | 0.043 | 0.1326 | 0.0558 | -0.2204 | -0.2915 | -0.3342 | 0.1323 | 0.2789 | 0.2939 | 0.1237 | 0.2797 | 0.3087 |
| rL2(2) | 0.2255 | 0.4994 | 0.8681 | 0.6077 | 1 | 0.6418 | -0.4976 | -0.6426 | -0.6266 | -0.2377 | -0.3143 | -0.3002 | -0.4759 | -0.6797 | -0.6713 | 0.299 | 0.3464 | 0.1959 | -0.465 | -0.686 | -0.6879 | 0.3094 | 0.4314 | 0.3169 | 0.276 | 0.4149 | 0.3956 |
| rL2(3) | 0.5206 | 0.7506 | 0.6168 | 0.228 | 0.6418 | 1 | -0.5441 | -0.6258 | -0.6247 | -0.1197 | -0.2204 | -0.1654 | -0.4932 | -0.6703 | -0.681 | 0.4235 | 0.3841 | 0.2967 | -0.4183 | -0.633 | -0.6311 | 0.3283 | 0.3621 | 0.2426 | 0.2612 | 0.3799 | 0.3672 |
| rP1(1) | -0.1158 | -0.4387 | -0.4805 | -0.1215 | -0.4976 | -0.5441 | 1 | 0.7888 | 0.7693 | 0.4275 | 0.3066 | 0.2222 | 0.9327 | 0.7847 | 0.7533 | -0.6615 | -0.4866 | -0.4084 | 0.8562 | 0.7211 | 0.6769 | -0.7601 | -0.4694 | -0.3685 | -0.6689 | -0.4943 | -0.4154 |
| rP1(2) | -0.1337 | -0.5081 | -0.6743 | -0.2408 | -0.6426 | -0.6258 | 0.7888 | 1 | 0.974 | 0.4537 | 0.5399 | 0.4525 | 0.673 | 0.9414 | 0.9152 | -0.4437 | -0.5382 | -0.4746 | 0.6809 | 0.9076 | 0.8755 | -0.6135 | -0.7211 | -0.5999 | -0.5293 | -0.7557 | -0.6907 |
| rP1(3) | -0.092 | -0.4979 | -0.6705 | -0.211 | -0.6266 | -0.6247 | 0.7693 | 0.974 | 1 | 0.4696 | 0.5937 | 0.4674 | 0.6536 | 0.9233 | 0.9322 | -0.3945 | -0.4649 | -0.4818 | 0.6745 | 0.9137 | 0.89 | -0.5976 | -0.7314 | -0.6413 | -0.5336 | -0.774 | -0.7249 |
| rP2(1) | 0.0989 | -0.0031 | -0.2098 | -0.2403 | -0.2377 | -0.1197 | 0.4275 | 0.4537 | 0.4696 | 1 | 0.7806 | 0.727 | 0.4151 | 0.4265 | 0.3966 | 0.1458 | 0.0663 | 0.0185 | 0.6743 | 0.5348 | 0.505 | -0.7286 | -0.5522 | -0.5677 | -0.8368 | -0.6389 | -0.6373 |
| rP2(2) | 0.0551 | -0.0662 | -0.3776 | -0.2502 | -0.3143 | -0.2204 | 0.3066 | 0.5399 | 0.5937 | 0.7806 | 1 | 0.8792 | 0.2328 | 0.4556 | 0.4607 | 0.149 | 0.0614 | -0.0657 | 0.4377 | 0.6298 | 0.6029 | -0.5196 | -0.7859 | -0.7933 | -0.6283 | -0.8643 | -0.8727 |
| rP2(3) | -0.0132 | -0.1187 | -0.3311 | -0.3683 | -0.3002 | -0.1654 | 0.2222 | 0.4525 | 0.4674 | 0.727 | 0.8792 | 1 | 0.1826 | 0.3997 | 0.3651 | 0.1784 | 0.0792 | 0.0361 | 0.4175 | 0.5542 | 0.6059 | -0.4404 | -0.6484 | -0.7633 | -0.5115 | -0.7472 | -0.8217 |
| rtau13(1) | -0.1223 | -0.4144 | -0.4406 | -0.1478 | -0.4759 | -0.4932 | 0.9327 | 0.673 | 0.6536 | 0.4151 | 0.2328 | 0.1826 | 1 | 0.7737 | 0.7431 | -0.5044 | -0.2795 | -0.1496 | 0.9097 | 0.7076 | 0.6674 | -0.6307 | -0.2717 | -0.1898 | -0.6152 | -0.3524 | -0.2731 |
| rtau13(2) | -0.1522 | -0.5604 | -0.6887 | -0.2618 | -0.6797 | -0.6703 | 0.7847 | 0.9414 | 0.9233 | 0.4265 | 0.4556 | 0.3997 | 0.7737 | 1 | 0.9809 | -0.3424 | -0.3468 | -0.2417 | 0.7584 | 0.9539 | 0.931 | -0.5172 | -0.5262 | -0.4218 | -0.4878 | -0.6186 | -0.5602 |
| rtau13(3) | -0.1188 | -0.5464 | -0.6865 | -0.222 | -0.6713 | -0.681 | 0.7533 | 0.9152 | 0.9322 | 0.3966 | 0.4607 | 0.3651 | 0.7431 | 0.9809 | 1 | -0.3132 | -0.3069 | -0.2341 | 0.7326 | 0.9505 | 0.9315 | -0.4736 | -0.5135 | -0.4022 | -0.4518 | -0.5971 | -0.5441 |
| rtau21(1) | 0.1718 | 0.2846 | 0.3274 | 0.043 | 0.299 | 0.4235 | -0.6615 | -0.4437 | -0.3945 | 0.1458 | 0.149 | 0.1784 | -0.5044 | -0.3424 | -0.3132 | 1 | 0.7501 | 0.662 | -0.3277 | -0.2178 | -0.2031 | 0.4733 | 0.2364 | 0.1138 | 0.2128 | 0.1153 | 0.0526 |
| rtau21(2) | 0.1766 | 0.2878 | 0.4064 | 0.1326 | 0.3464 | 0.3841 | -0.4866 | -0.5382 | -0.4649 | 0.0663 | 0.0614 | 0.0792 | -0.2795 | -0.3468 | -0.3069 | 0.7501 | 1 | 0.84 | -0.2197 | -0.2748 | -0.2494 | 0.3995 | 0.468 | 0.3237 | 0.119 | 0.2748 | 0.2297 |
| rtau21(3) | 0.1215 | 0.2559 | 0.2883 | 0.0558 | 0.1959 | 0.2967 | -0.4084 | -0.4746 | -0.4818 | 0.0185 | -0.0657 | 0.0361 | -0.1496 | -0.2417 | -0.2341 | 0.662 | 0.84 | 1 | -0.107 | -0.1868 | -0.1955 | 0.4231 | 0.5457 | 0.5054 | 0.1605 | 0.3832 | 0.3266 |
| rtau23(1) | -0.0321 | -0.3238 | -0.4335 | -0.2204 | -0.465 | -0.4183 | 0.8562 | 0.6809 | 0.6745 | 0.6743 | 0.4377 | 0.4175 | 0.9097 | 0.7584 | 0.7326 | -0.3277 | -0.2197 | -0.107 | 1 | 0.7699 | 0.7427 | -0.7138 | -0.3769 | -0.3337 | -0.6903 | -0.4482 | -0.408 |
| rtau23(2) | -0.091 | -0.4737 | -0.7204 | -0.2915 | -0.686 | -0.633 | 0.7211 | 0.9076 | 0.9137 | 0.5348 | 0.6298 | 0.5542 | 0.7076 | 0.9539 | 0.9505 | -0.2178 | -0.2748 | -0.1868 | 0.7699 | 1 | 0.9684 | -0.5211 | -0.6282 | -0.5438 | -0.5133 | -0.6921 | -0.6637 |
| rtau23(3) | -0.1315 | -0.5436 | -0.7092 | -0.3342 | -0.6879 | -0.6311 | 0.6769 | 0.8755 | 0.89 | 0.505 | 0.6029 | 0.6059 | 0.6674 | 0.931 | 0.9315 | -0.2031 | -0.2494 | -0.1955 | 0.7427 | 0.9684 | 1 | -0.4804 | -0.5982 | -0.574 | -0.4616 | -0.6789 | -0.6549 |
| rtau31(1) | -0.0108 | 0.1413 | 0.3148 | 0.1323 | 0.3094 | 0.3283 | -0.7601 | -0.6135 | -0.5976 | -0.7286 | -0.5196 | -0.4404 | -0.6307 | -0.5172 | -0.4736 | 0.4733 | 0.3995 | 0.4231 | -0.7138 | -0.5211 | -0.4804 | 1 | 0.6359 | 0.5711 | 0.8968 | 0.6102 | 0.554 |
| rtau31(2) | 0.0502 | 0.2352 | 0.5135 | 0.2789 | 0.4314 | 0.3621 | -0.4694 | -0.7211 | -0.7314 | -0.5522 | -0.7859 | -0.6484 | -0.2717 | -0.5262 | -0.5135 | 0.2364 | 0.468 | 0.5457 | -0.3769 | -0.6282 | -0.5982 | 0.6359 | 1 | 0.9116 | 0.5866 | 0.9314 | 0.8887 |
| rtau31(3) | 0.064 | 0.2285 | 0.3965 | 0.2939 | 0.3169 | 0.2426 | -0.3685 | -0.5999 | -0.6413 | -0.5677 | -0.7933 | -0.7633 | -0.1898 | -0.4218 | -0.4022 | 0.1138 | 0.3237 | 0.5054 | -0.3337 | -0.5438 | -0.574 | 0.5711 | 0.9116 | 1 | 0.5263 | 0.8867 | 0.9031 |
| rtau32(1) | -0.0496 | 0.1224 | 0.2288 | 0.1237 | 0.276 | 0.2612 | -0.6689 | -0.5293 | -0.5336 | -0.8368 | -0.6283 | -0.5115 | -0.6152 | -0.4878 | -0.4518 | 0.2128 | 0.119 | 0.1605 | -0.6903 | -0.5133 | -0.4616 | 0.8968 | 0.5866 | 0.5263 | 1 | 0.6572 | 0.5967 |
| rtau32(2) | 0.0694 | 0.3399 | 0.4813 | 0.2797 | 0.4149 | 0.3799 | -0.4943 | -0.7557 | -0.774 | -0.6389 | -0.8643 | -0.7472 | -0.3524 | -0.6186 | -0.5971 | 0.1153 | 0.2748 | 0.3832 | -0.4482 | -0.6921 | -0.6789 | 0.6102 | 0.9314 | 0.8867 | 0.6572 | 1 | 0.9464 |
| rtau32(3) | 0.0588 | 0.2901 | 0.4528 | 0.3087 | 0.3956 | 0.3672 | -0.4154 | -0.6907 | -0.7249 | -0.6373 | -0.8727 | -0.8217 | -0.2731 | -0.5602 | -0.5441 | 0.0526 | 0.2297 | 0.3266 | -0.408 | -0.6637 | -0.6549 | 0.554 | 0.8887 | 0.9031 | 0.5967 | 0.9464 | 1 |
