## Supplement Table S6B for "Single-molecule behavior and cell-growth regulation in human RTKs"

| R | EC | IDRE | TM | IDRC | Linker | TK | Tail | IDRE pos | IDRC pos | RK cluste | GxxxG | CRAC |
|---|---|---|---|---|---|---|---|---|---|---|---|---|
| EC | 1 | -0.2479 | 0.0522 | -0.0028 | -0.1393 | -0.0034 | -0.2627 | -0.2313 | 0.0183 | 0.1628 | -0.0414 | 0.0876 |
| IDRE | -0.2479 | 1 | -0.0208 | 0.019 | 0.3065 | -0.1523 | -0.0618 | 0.3788 | -0.0826 | -0.0888 | 0.4635 | -0.0647 |
| TM | 0.0522 | -0.0208 | 1 | -0.2267 | -0.0289 | -0.1397 | 0.1358 | -0.145 | -0.1955 | -0.0069 | 0.0531 | -0.1055 |
| IDRC | -0.0028 | 0.019 | -0.2267 | 1 | -0.4927 | -0.2529 | -0.1048 | 0.2867 | 0.6521 | 0.5487 | -0.0734 | 0.2675 |
| Linker | -0.1393 | 0.3065 | -0.0289 | -0.4927 | 1 | 0.0907 | -0.0499 | -0.0371 | -0.1979 | -0.3183 | 0.0705 | -0.2144 |
| TK | -0.0034 | -0.1523 | -0.1397 | -0.2529 | 0.0907 | 1 | -0.1134 | -0.0294 | -0.2624 | -0.2707 | -0.2988 | -0.1061 |
| Tail | -0.2627 | -0.0618 | 0.1358 | -0.1048 | -0.0499 | -0.1134 | 1 | -0.2814 | -0.0901 | 0.036 | -0.077 | -0.1717 |
| IDRE posi | -0.2313 | 0.3788 | -0.145 | 0.2867 | -0.0371 | -0.0294 | -0.2814 | 1 | 0.1803 | 0.0888 | 0.1566 | -0.0586 |
| IDRC posi | 0.0183 | -0.0826 | -0.1955 | 0.6521 | -0.1979 | -0.2624 | -0.0901 | 0.1803 | 1 | 0.7865 | 0.0232 | 0.1119 |
| RK cluster | 0.1628 | -0.0888 | -0.0069 | 0.5487 | -0.3183 | -0.2707 | 0.036 | 0.0888 | 0.7865 | 1 | 0.107 | 0.0146 |
| GxxxG | -0.0414 | 0.4635 | 0.0531 | -0.0734 | 0.0705 | -0.2988 | -0.077 | 0.1566 | 0.0232 | 0.107 | 1 | 0.005 |
| CRAC | 0.0876 | -0.0647 | -0.1055 | 0.2675 | -0.2144 | -0.1061 | -0.1717 | -0.0586 | 0.1119 | 0.0146 | 0.005 | 1 |
