## Supplement Table S7 for "Single-molecule behavior and cell-growth regulation in human RTKs"

**Supplement Table S6. RTK sequences around the TM-JM region**

EGFR: TNGPKIPSIATGMVGALLLLLVVALGIGLFMRRRHIVRKRTLRRLLQERELVEPLTPSGEAPNQALLRILKETE

ERBB2: GCPAEQRASPLTSIISAVVGILLVVVLGVVFGILIKRRQQKIRKYTMRRLLQETELVEPLTPSGAMPNQAQMRILKETEL

ERBB3: MALTVIAGLVVIFMMLGGTFLYWRGRRIQNKRAMRRYLERGESIEPLDPSEKANKVLARIFKETE

ERBB4: LPQHARTPLIAAGVIGGLFILVIVGLTFAVYVRRKSIKKKRALRRFLETELVEPLTPSGTAPNQAQLRILKETE

INSR: NIAKIIIGPLIFVFLFSVVIGSIYLFLRKRQPDGPLGPLYASSNPEYLSASDVFPCSVYVPDEWEVSREK

IGF1R: YENFIHLIIALPVAVLLIVGGLVIMLYVFHRKRNNSRLGNGVLYASVNPEYFSAADVYVPDEWEVAREK

INSRR: LGPEEEDAGGLHVLLTATPVGLTLLIVLAALGFFYGKKRNRTLYASVNPEYFSASDMYVPDEWEVPREQ

PDGFRA: ELKLVAPTLRSELTVAAAVLVLLVIVIISLIVLVVIWKQKPRYEIRWRVIESISPDGHEYIYVDPMQLPYDSRWEFPRDG

PDGFRB: PFKVVVISAILALVVLTIISLIILIMLWQKKPRYEIRWKVIESVSSDGHEYIYVDPMQLPYDSTWELPRDQ

KIT: KEQIHPHTLFTPLLIGFVIVAGMMCIIVMILTYKYLQKPMYEVQWKVVEEINGNNYVYIDPTQLPYDHKWEFPRNR

CSF1R: AHTHPPDEFLFTPVVVACMSIMALLLLLLLLLYKYKQKPKYQVRWKIIESYEGNSYTFIDPTQLPYNEKWEFPRNN

FLT3: ISFYATIGVCLLFIVVLTLLICHKYKKQFRYESQLQMVQVTGSSDNEYFYVDFREYEYDLKWEFPREN

FLT1: TSDKSNLELITLTCTCVAATLFWLLLTLFIRKMKRSSSEIKTDYLSIIMDPDEVPLDEQCERLPYDASKWEFARER

KDR: QEKTNLEIIILVGTAVIAMFFWLLLVIILRTVKRANGGELKTGYLSIVMDPDELPLDEHCERLPYDASKWEFPRDR

FLT4: SVAVEGSEDKGSMEIVILVGTGVIAVFFWVLLLLIFCNMRRPAHADIKTGYLSIIMDPGEVPLEEQCEYLSYDASQWEFPRER

FGFR1: LEALEERPAVMTSPLYLEIIIYCTGAFLISCMVGSVIVYKMKSGTKKSDFHSQMAVHKLAKSIPLRRQVTVSADSSASMNSGVLLVRPSRLSSSGTPMLAGVSEYELPEDPRWELPRDR

FGFR2: PGREKEITASPDYLEIAIYCIGVFLIACMVVTVILCRMKNTTKKPDFSSQPAVHKLTKRIPLRRQVTVSAESSSSMNSNTPLVRITTRLSSTADTPMLAGVSEYELPEDPKWEFPRDK

FGFR3: LPAEEELVEADEAGSVYAGILSYGVGFFLFILVVAAVTLCRLRSPPKKGLGSPTVHKISRFPLKRQVSLESNASMSSNTPLVRIARLSSGEGPTLANVSELELPADPKWELSRAR

FGFR4: PTWTAAAPEARYTDIILYASGSLALAVLLLLAGLYRGQALHGRHPRPPATVQKLSRFPLARQFSLESGSSGKSSSSLVRGVRLSSSGPALLAGLVSLDLPLDPLWEFPRDR

PTK7: DKPVPEESEGPGSPPPYKMIQTIGLSVGAAVAYIIAVLGLMFYCKKRCKAKRLQKQPEGEEPEMECLNGGPLQNGQPSAEIQEEVALTSLGSGPAATNKRHSTSDKMHFPRSS

TRKA: DNPFEFNPEDPIPVSFSPVDTNSTSGDPVEKKDETPFGVSVAVGLAVFACLFLSTLLLVLNKCGRRNKFGINRPAVLAPEDGLAMSLHFMTLGGSSLSPTEGKGSGLQGHIIENPQYFSD
ACVHHIKRRD

TRKB: IGDTTNRSNEIPSTDVTDKTGREHLSVYAVVVIASVVGFCLLVMLFLLKLARHSKFGMKGPASVISNDDDSASPLHHISNGSNTPSSSEGGPDAVIIGMTKIPVIENPQYFGITNSQLKP
DTFVQHIKRHN

TRKC: FILFDEVSPTPPITVTHKPEEDTFGVSIAVGLAAFACVLLVVLFVMINKYGRRSKFGMKGPVAVISGEEDSASPLHHINHGITTPSSLDAGPDTVVIGMTRIPVIENPQYFRQGHNCHKP
DTYVQHIKRRD

ROR1: DSKEKNKMEILYILVPSVAIPLAIALLFFFICVCRNNQKSSSAPVQRQPKHVRGQNVEMSMLNAYKPKSKAKELPLSA

ROR2: PSCSPRDSSKMGILYILVPSIAIPLVIACLFFLVCMCRNKQKASASTPQRRQLMASPSQDMEMPLINQHKQAKLKEISLSA

MUSK: KPSVDIPNLPSSSSSSFSVSPTYSMTVIISIMSSFAIFVLLTITTLYCCRRRKQWKNKKRESAAVTLTTLPSELLLDRLHPNPMYQRMPLLLNPKLLSLEYPRNN

MET: QPDQNFTGLIAGVVSISTALLLLLGFFLWLKKRKQIKDLGSELVRYDARVHTPHLDRLVSARSVSPTTEMVSNESVDYRATFPEDQFPNSSQNGSCRQVQYPLTDMSPILTSGDSDISSP
LLQNTVHIDLSALNPELVQAVQHVVIGPSSLI

RON: VRPGPDGVPQSTLLGILLPLLLLVAALATALVFSYWWRRKQLVLPPNLNDLASLDQTAGATPLPILYSGSDYRSGLALPAIDGLDSTTCVHGASFSDSEDESCVPLLRKESIQLRDLDSA
LLAEVKDVLIPHERVV

AXL: HQLVKEPSTPAFSWPWWYVLLGAVVAAACVLILALFLVHRRKKETRYGEVFEPTVERGELVVRYRVRKSYSRRTTEATLNSLGISEELKEKLRDVMVDRHKVALGKTLGEGEFGAVMEG
QL

TYRO3: VSSHDRAGQQGPPHSRTSWVPVVLGVLTALVTAAALALILLRKRRKETRFGQAFDSVMARGEPAVHFRAARSFNRERPERIEATLDSLGISDELKEKLEDVLIPEQQ

TIE1: TVEESTLGNGLQAEGPVQESRAAEEGLDQQLILAVVGSVSATCLTILAALLTLVCIRRSCLHRRRTFTYQSGSGEETILQFSSGTLTLTRRPKLQPEPLSYPVLEWED

TIE2: SNPAFSHELVTLPESQAPADLGGGKMLLIAILGSAGMTCLTVLLAFLIILQLKRANVQRRMAQAFQNVREEPAVQFNSGTLALNRKVKNNPDPTIYPVLDWND

EPHA1: RTSPPVSRGLTGGEIVAVIFGLLLGAALLLGILVFRSRRAQRQRQQRQRDRATDVDREDKLWLKPYVDLQAYEDPAQGALDFTRELDPAW

EPHA2: HEFQTLSPEGSGNLAVIGGVAVGVVLLLVLAGVGFFIHRRRKNQRARQSPEDVYFSKSEQLKPLKTYVDPHTYEDPNQAVLKFTTEIHPSC

EPHA3: TSPDSFSISGESSQVVMIAISAAVAIILLTVVIYVLIGRFCGYKSKHGADEKRLHFGNGHLKLPGLRTYVDPHTYEDPTQAVHEFAKELDATN

EPHA4: FSEPLEVTTNTVPSRIIGDGANSTVLLVSVSGSVVLVVILIAAFVISRRRSKYSKAKQEADEEKHLNQGVRTYVDPFTYEDPNQAVREFAKEIDASC

EPHA6: GDETSDMAAEQGQILVIATAAVGGFTLLVILTLFFLITGRCQWYIKAKMKSEEKRRNHLQNGHLRFPGIKTYIDPDTYEDPSLAVHEFAKEIDPSR

EPHA7: DVATLEEATGKMFEATAVSSEQNPVIIIAVVAVAGTIILVFMVFGFIIGRRHCGYSKADQEGDEELYFHFKFPGTKTYIDPETYEDPNRAVHQFAKELDASC

EPHB1: DDYKSELREQLPLIAGSAAAGVVFVVSLVAISIVCSRKRAYSKEAVYSDKLQHYSTGRGSPGMKIYIDPFTYEDPNEAVREFAKEIDVSF

EPHB2: KLPLIIGSSAAGLVFLIAVVVIAIVCNRRGFERADSEYTDKLQHYTSGHMTPGMKIYIDPFTYEDPNEAVREFAKEIDISC

EPHB3: SRPAEFETTSERGSGAQQLQEQLPLIVGSATAGLVFVVAVVVIAIVCLRKQRHGSDSEYTEKLQQYIAPGMKVYIDPFTYEDPNEAVREFAKEIDVSC

EPHB4: YGPFGQEHHSQTQLDESEGWREQLALIAGTAVVGVVLVLVVIVVAVLCLRKQSNGREAEYSDKHGQYLIGHGTKVYIDPFTYEDPNEAVREFAKEIDVSY

EPHB6: GELSSQLPERLSLVIGSILGALAFLLLAAITVLAVVFQRKRRGTGYTEQLQQYSSPGLGVKYYIDPSTYEDPCQAIRELAREVDPAY

RET: TVIAAAVLFSFIVSVLLSAFCIHCYHKFAHKPPISSAEMTFRRPAQAFPVSYSSSGARRPSLDSMENQVSVDAFKILEDPKWEFPRKN

DDR1: FSSLELEPRGQQPVAKAEGSPTAILIGCLVAIILLLLLIIALMLWRLHWRRLLSKAERRVLEEELTVHLSVPGDTILINNRPGPREPPPYQEPRPRGNPPHSAPCVPNGSALLLSNPAYR
LLLATYARPPRGPGPPTPAWAKPTNTQAYSGDYMEPEKPGAPLLPPPPQNSVPHYAEADIVTLQGVTGGNTYAVPALPPGAVGDGPPRVDFPRSR

DDR2: AMYNNSEALPTSPMAPTTYDPMLKVDDSNTRILIGCLVAIIFILLAIIVIILWRQFWQKMLEKASRRMLDDEMTVSLSLPSDSSMFNNNRSSSPSEQGSNSTYDRIFPLRPDYQEPSRLI
RKLPEFAPGEEESGCSGVVKPVQPSGPEGVPHYAEADIVNLQGVTGGNTYSVPAVTMDLLSGKDVAVEEFPRKL

ROS1: IPETSFILTIIVGIFLVVTIPLTFVWHRRLKNQKSAKEGVTVLINEDKELAELRGLAAGVGLANACYAIHTLPTQEEIENLPAFPREK

LMR1: DGAPLSELSWPSSLAVVAVSFSGLFAVIVLMLACLCCKKGGIGFKEFENAEGDEYAADLAQGSPATAAQNGPDVYVLPLTEVSLPMAKQPGRSVQLLKSTDVGRHS

LMR2: GSAGAAPLPQTGAGEAPPAAEVSSSFVILCVCSLIIILIVLIANCVSCCKDPEIDFKEFEDNFDDEIDFTPPAEDTPSVQSPAEVFTLSVPNISLPAPSQFQPSVEGLKSQVARHS

LTK: HKPPGPLVLMVAVVATSTLSLLMVCGVLILVKQKKWQGLQEMRLPSPELELSKLRTSAIRTAPNPYYCQVGLGPAQSWPLPPGVTEVSPANVT

ALK: SCIVSPTPEPHLPLSLILSVVTSALVAALVLAFSGIMIVYRRKHQELQAMQMELQSPEYKLSKLRTSTIMTDYNPNYCFAGKTSSISDLKEVPRKN

STYK1: VIIVPTLLVTIFLILLGVILWLFIREQRTQQQRSGPQGIAPVPPPRDLSWEAGHGGNVALPLKETSVENFLGATTPALAKLQVPREQL

The amino acid sequences from the IDRE to the linker before the Kinase domain are listed.

TM , IDRE and IDRC, sequence in the kinase domain, R and K, G/Ax(x)xG/A in TM, **Y/F in CRAC/CARC (L/V (X)(1-5)-Y-(X)(1-5)-R/K)**
