## Supplement Table S8A for "Single-molecule behavior and cell-growth regulation in human RTKs"

| | pR | | | | | | | | | p-values in t-test | | | | |
|---|---|---|---|---|---|---|---|---|---|---|---|---|---|---|
| | TM | Tail | RK cluste | GxxxG | CRAC | determ | F-value | p-value | | TM | Tail | RK cluste | GxxxG | CRAC |
| KURAMOCHI | 0.01392 | -0.4911 | -0.3048 | -0.1935 | 0.12792 | 0.41463 | 6.51654 | 0.00012 | | 0.46219 | 0.00017 | 0.01659 | 0.09136 | 0.19054 |
| SKUT1 | 0.18886 | -0.4185 | -0.1351 | -0.2149 | 0.33583 | 0.39192 | 5.92947 | 0.00026 | | 0.09686 | 0.00138 | 0.17727 | 0.06902 | 0.00916 |
| CAKI2 | -0.0341 | -0.3194 | -0.4997 | -0.0698 | 0.08222 | 0.39131 | 5.91455 | 0.00027 | | 0.40814 | 0.01265 | 0.00013 | 0.31689 | 0.28719 |
| HS695T | 0.2944 | -0.5397 | -0.1384 | 0.00666 | 0.10602 | 0.38431 | 5.74257 | 0.00034 | | 0.02002 | 3.1E-05 | 0.17154 | 0.4819 | 0.23421 |
| KYAE1 | -0.0331 | -0.1314 | -0.5604 | -0.0744 | -0.1711 | 0.37519 | 5.52453 | 0.00046 | | 0.4107 | 0.184 | 1.4E-05 | 0.30582 | 0.11987 |
| JHUEM7 | 0.17923 | -0.3731 | -0.2195 | -0.1217 | 0.35598 | 0.37515 | 5.52349 | 0.00046 | | 0.10893 | 0.00414 | 0.06486 | 0.20248 | 0.00603 |
| HKGZCC | 0.17787 | -0.2886 | -0.4836 | -0.1333 | -0.025 | 0.36867 | 5.37252 | 0.00057 | | 0.11072 | 0.02215 | 0.00022 | 0.18053 | 0.43221 |
| SKGT4 | 0.25688 | -0.3096 | -0.3954 | -0.1454 | 0.12771 | 0.35062 | 4.96727 | 0.00102 | | 0.0374 | 0.01521 | 0.00246 | 0.15944 | 0.19092 |
| ONCODG1 | -0.1713 | -0.3125 | -0.2848 | -0.1859 | 0.19684 | 0.33372 | 4.60802 | 0.00172 | | 0.11969 | 0.01439 | 0.02366 | 0.10042 | 0.08761 |
| HCC1187 | 0.2418 | -0.1176 | 0.0527 | -0.0073 | 0.51981 | 0.33184 | 4.56921 | 0.00182 | | 0.04708 | 0.21042 | 0.35956 | 0.48015 | 6.5E-05 |
| TE6 | 0.12104 | -0.3353 | -0.4271 | -0.0775 | 0.06951 | 0.32775 | 4.4853 | 0.00206 | | 0.20371 | 0.00925 | 0.0011 | 0.29833 | 0.31754 |
| WM3772F | 0.26174 | -0.4762 | 0.10682 | -0.2753 | 0.08724 | 0.32755 | 4.48135 | 0.00207 | | 0.03463 | 0.00027 | 0.23253 | 0.02776 | 0.27557 |
| CAOV4 | 0.16928 | -0.2656 | -0.0925 | 0.00718 | 0.44901 | 0.32324 | 4.39415 | 0.00236 | | 0.12246 | 0.03253 | 0.26371 | 0.48049 | 0.00061 |
| BT16 | 0.02204 | -0.4046 | -0.0157 | -0.357 | 0.17389 | 0.32099 | 4.34908 | 0.00253 | | 0.44027 | 0.00197 | 0.45724 | 0.00589 | 0.11605 |
| MCAS | 0.28513 | -0.2016 | -0.3921 | -0.1843 | 0.1032 | 0.31565 | 4.24344 | 0.00296 | | 0.02353 | 0.08236 | 0.00267 | 0.10239 | 0.2402 |
| ICC2 | 0.05605 | -0.4148 | -0.3473 | -0.0596 | 0.05247 | 0.31296 | 4.19079 | 0.0032 | | 0.35103 | 0.00152 | 0.00725 | 0.34208 | 0.36016 |
| SW1463 | 0.20906 | -0.4408 | -0.0171 | -0.2237 | 0.21188 | 0.31158 | 4.16393 | 0.00333 | | 0.0747 | 0.00076 | 0.45354 | 0.06119 | 0.07194 |
| SF767 | 0.02878 | -0.335 | -0.4169 | 0.10322 | 0.06124 | 0.31122 | 4.15694 | 0.00337 | | 0.42219 | 0.00932 | 0.00144 | 0.24016 | 0.33797 |
| HCC1419 | 0.01783 | -0.3507 | -0.4194 | -0.0483 | -0.0635 | 0.30957 | 4.12504 | 0.00353 | | 0.45161 | 0.00675 | 0.00135 | 0.37088 | 0.33228 |
| FU97 | 0.31183 | -0.1689 | 0.17344 | 0.00829 | 0.41136 | 0.30771 | 4.08923 | 0.00373 | | 0.01458 | 0.12295 | 0.11667 | 0.47747 | 0.00166 |
| ECGI10 | 0.07873 | -0.3197 | -0.4271 | -0.0847 | 0.00111 | 0.30448 | 4.02761 | 0.00409 | | 0.29537 | 0.01256 | 0.0011 | 0.28131 | 0.49697 |
| MM253 | 0.18059 | -0.3304 | -0.1722 | -0.3895 | 0.04824 | 0.304 | 4.01846 | 0.00415 | | 0.10717 | 0.01022 | 0.11841 | 0.00284 | 0.37102 |
| TOV21G | 0.17565 | -0.4071 | -0.075 | -0.2644 | 0.21092 | 0.30294 | 3.99825 | 0.00428 | | 0.11368 | 0.00185 | 0.30429 | 0.0332 | 0.07287 |
| T3M4 | -0.2073 | -0.3074 | -0.3612 | -0.0917 | -0.0373 | 0.30159 | 3.97276 | 0.00445 | | 0.07652 | 0.01583 | 0.00539 | 0.26542 | 0.39951 |
| SNU216 | 0.02746 | -0.3118 | -0.4108 | -0.1412 | 0.00437 | 0.29933 | 3.93022 | 0.00475 | | 0.42571 | 0.0146 | 0.00169 | 0.1665 | 0.48811 |
| UHO1 | 0.05363 | -0.4494 | 0.0794 | -0.1486 | 0.2337 | 0.29859 | 3.91649 | 0.00485 | | 0.35719 | 0.0006 | 0.29381 | 0.15406 | 0.05303 |
| C4II | 0.20594 | -0.3369 | -0.2123 | -0.3194 | 0.10354 | 0.29811 | 3.90753 | 0.00491 | | 0.07785 | 0.00897 | 0.07149 | 0.01263 | 0.23947 |
| UWB1289 | 0.12792 | -0.3094 | -0.4264 | 0.02128 | 0.03336 | 0.29651 | 3.87761 | 0.00514 | | 0.19054 | 0.01527 | 0.00113 | 0.44229 | 0.41001 |
| HT144SKINFV2 | 0.22129 | -0.4883 | 0.04091 | -0.1554 | 0.11953 | 0.29437 | 3.83807 | 0.00546 | | 0.06325 | 0.00019 | 0.39009 | 0.14318 | 0.20666 |
| MEC1 | 0.3432 | -0.2198 | -0.1492 | -0.2989 | 0.21431 | 0.29382 | 3.82784 | 0.00555 | | 0.00789 | 0.06454 | 0.15311 | 0.01849 | 0.06961 |
| OVK18 | 0.30306 | -0.1825 | 0.01041 | -0.3378 | 0.30099 | 0.29237 | 3.80115 | 0.00578 | | 0.01714 | 0.10469 | 0.4717 | 0.0088 | 0.0178 |
| DOTC24510 | 0.08483 | -0.3286 | -0.3928 | -0.1305 | -0.0241 | 0.29044 | 3.76587 | 0.0061 | | 0.28112 | 0.01058 | 0.00262 | 0.18572 | 0.43472 |
| JHUEM1 | 0.21234 | -0.4264 | -0.0187 | -0.0825 | 0.2489 | 0.29008 | 3.75915 | 0.00616 | | 0.07149 | 0.00112 | 0.44926 | 0.28656 | 0.0423 |
| HO1U1 | 0.1287 | -0.4624 | -0.1462 | -0.1113 | 0.15234 | 0.28956 | 3.74964 | 0.00625 | | 0.18907 | 0.00041 | 0.1581 | 0.22323 | 0.14802 |
| OVMANA | 0.09782 | -0.4282 | -0.2874 | -0.1151 | 0.02263 | 0.28805 | 3.72234 | 0.00652 | | 0.25186 | 0.00107 | 0.02262 | 0.21558 | 0.43867 |
| SNU213 | 0.06987 | -0.3074 | -0.4038 | -0.1259 | 0.02233 | 0.28797 | 3.72086 | 0.00653 | | 0.31667 | 0.01583 | 0.00201 | 0.19442 | 0.43948 |
| CAL33 | 0.04823 | -0.2936 | -0.4105 | -0.0895 | 0.08018 | 0.28621 | 3.68898 | 0.00686 | | 0.37103 | 0.0203 | 0.00169 | 0.27045 | 0.29197 |
| HSC5 | 0.22039 | -0.2149 | -0.3796 | -0.0874 | 0.19525 | 0.28395 | 3.64835 | 0.0073 | | 0.06404 | 0.06907 | 0.00357 | 0.27518 | 0.08941 |
| KYSE180 | 0.17921 | -0.3603 | -0.3228 | -0.0901 | 0.08411 | 0.27887 | 3.55777 | 0.00839 | | 0.10896 | 0.00549 | 0.01183 | 0.26897 | 0.28278 |
| PCI6A | 0.15591 | -0.2694 | -0.3933 | -0.0425 | 0.14004 | 0.27781 | 3.53909 | 0.00864 | | 0.14236 | 0.03063 | 0.00259 | 0.38588 | 0.1686 |
| HEY | 0.30754 | -0.2491 | -0.1129 | -0.2736 | 0.24563 | 0.27703 | 3.52533 | 0.00882 | | 0.01579 | 0.0422 | 0.21985 | 0.02858 | 0.04446 |
| DOK | 0.14553 | -0.1699 | -0.4672 | -0.0653 | -0.0049 | 0.27628 | 3.51209 | 0.009 | | 0.1592 | 0.1216 | 0.00036 | 0.32785 | 0.48673 |
| SNGM | 0.30193 | -0.3172 | -0.0921 | -0.1482 | 0.27255 | 0.27512 | 3.49177 | 0.00929 | | 0.0175 | 0.01317 | 0.26457 | 0.15483 | 0.02907 |
| SNU8 | 0.11146 | -0.2697 | -0.4347 | 0.04544 | -0.0441 | 0.27476 | 3.48549 | 0.00938 | | 0.2229 | 0.03045 | 0.0009 | 0.37828 | 0.38163 |
| TGBC1TKB | 0.29831 | -0.2458 | -0.3143 | -0.1472 | -0.1401 | 0.27455 | 3.4818 | 0.00944 | | 0.01868 | 0.04434 | 0.01394 | 0.1564 | 0.16845 |
| VMRCLCD | 0.15554 | -0.3109 | 0.01866 | -0.2888 | 0.2852 | 0.27306 | 3.45586 | 0.00982 | | 0.14294 | 0.01483 | 0.44937 | 0.02208 | 0.0235 |
| TE11 | 0.13119 | -0.322 | -0.3533 | -0.0527 | 0.13032 | 0.27286 | 3.45232 | 0.00988 | | 0.18445 | 0.01203 | 0.00638 | 0.35948 | 0.18605 |
| MMAC | 0.06515 | -0.1025 | -0.1344 | -0.2587 | 0.4068 | 0.27218 | 3.44051 | 0.01006 | | 0.32825 | 0.24167 | 0.17854 | 0.03635 | 0.00186 |
| EFE184 | 0.0499 | -0.4276 | -0.2258 | -0.1585 | 0.0658 | 0.27192 | 3.43604 | 0.01013 | | 0.36674 | 0.00109 | 0.05939 | 0.13831 | 0.32665 |
| PLCPRF5 | 0.17765 | -0.2746 | -0.3125 | -0.163 | 0.18319 | 0.26781 | 3.36504 | 0.0113 | | 0.111 | 0.02809 | 0.01441 | 0.13159 | 0.10384 |
| 8MGBA | -0.228 | -0.2423 | -0.2415 | -0.1198 | 0.19311 | 0.26752 | 3.36011 | 0.01139 | | 0.05759 | 0.04672 | 0.04726 | 0.20605 | 0.09186 |
| KYSE450 | 0.14515 | -0.3293 | -0.3584 | -0.0665 | 0.05676 | 0.26664 | 3.34501 | 0.01166 | | 0.15985 | 0.01042 | 0.00572 | 0.32482 | 0.34924 |
| HT | 0.3001 | -0.158 | -0.0257 | -0.3204 | 0.28385 | 0.2665 | 3.34269 | 0.0117 | | 0.01808 | 0.13906 | 0.43043 | 0.01241 | 0.02405 |
| CAL29 | 0.10219 | -0.3371 | -0.2992 | 0.00399 | 0.1948 | 0.26649 | 3.34241 | 0.01171 | | 0.24237 | 0.00894 | 0.01839 | 0.48914 | 0.08991 |
| T3M3 | 0.23309 | -0.2886 | 0.02219 | -0.3746 | 0.16107 | 0.26548 | 3.32525 | 0.01203 | | 0.0535 | 0.02214 | 0.43985 | 0.004 | 0.13445 |
| SNU46 | -0.1246 | -0.242 | -0.0942 | 0.09711 | 0.35329 | 0.26527 | 3.32164 | 0.01209 | | 0.19674 | 0.04697 | 0.25979 | 0.25342 | 0.00639 |
| JHOC5 | 0.10668 | -0.3107 | -0.3938 | -0.0389 | -0.0423 | 0.26455 | 3.30934 | 0.01233 | | 0.23282 | 0.0149 | 0.00256 | 0.39547 | 0.38633 |
| HOUAI | 0.2304 | -0.4724 | 0.00872 | -0.2017 | -0.0364 | 0.26436 | 3.30613 | 0.01239 | | 0.05561 | 0.00031 | 0.47628 | 0.08231 | 0.40207 |
| DETROIT562 | 0.24243 | -0.2103 | -0.3897 | 0.07973 | 0.10312 | 0.26429 | 3.30494 | 0.01241 | | 0.04664 | 0.07347 | 0.00282 | 0.29303 | 0.24038 |
| KE37 | 0.1743 | -0.2109 | 0.05808 | -0.2981 | 0.33902 | 0.26378 | 3.29619 | 0.01258 | | 0.1155 | 0.07288 | 0.34591 | 0.01875 | 0.00859 |
| CAL51 | 0.0883 | -0.3528 | -0.1648 | -0.0645 | 0.27853 | 0.26232 | 3.27152 | 0.01307 | | 0.27314 | 0.00645 | 0.12885 | 0.32983 | 0.02632 |
| JEG3 | 0.16942 | -0.4438 | -0.1062 | -0.0839 | 0.15253 | 0.2623 | 3.27116 | 0.01308 | | 0.12226 | 0.0007 | 0.23379 | 0.28328 | 0.14772 |
| SUDHL10 | 0.17844 | -0.1805 | -0.0202 | -0.1543 | 0.4214 | 0.26058 | 3.24226 | 0.01368 | | 0.10997 | 0.1073 | 0.44528 | 0.14497 | 0.00128 |
| SSP25 | 0.03894 | -0.2985 | -0.3995 | 0.07697 | 0.01196 | 0.26017 | 3.23521 | 0.01383 | | 0.39525 | 0.01862 | 0.00223 | 0.29957 | 0.4675 |
| SUM149PT | 0.28959 | -0.1292 | -0.3171 | -0.2207 | 0.12642 | 0.26012 | 3.23439 | 0.01385 | | 0.02178 | 0.18809 | 0.0132 | 0.06377 | 0.19336 |
| NCIH650 | -0.0548 | -0.1455 | -0.0449 | -0.2552 | 0.38182 | 0.25995 | 3.23161 | 0.01391 | | 0.35411 | 0.15926 | 0.37968 | 0.03839 | 0.00339 |
| KARPAS299 | 0.05475 | -0.313 | -0.2958 | -0.1616 | 0.16511 | 0.25974 | 3.22814 | 0.01399 | | 0.35433 | 0.01426 | 0.01954 | 0.13358 | 0.12846 |
| HCC70 | 0.20866 | -0.3284 | -0.0132 | 0.04779 | 0.31283 | 0.25831 | 3.20408 | 0.01452 | | 0.0751 | 0.01062 | 0.46409 | 0.37218 | 0.01432 |
| C80 | 0.11828 | -0.4336 | -0.1884 | 0.09862 | 0.076 | 0.25822 | 3.20256 | 0.01456 | | 0.20914 | 0.00093 | 0.09741 | 0.2501 | 0.30187 |
| BICR56 | 0.05031 | -0.3209 | -0.3862 | 0.0191 | -0.0745 | 0.25818 | 3.20189 | 0.01457 | | 0.36569 | 0.01229 | 0.00307 | 0.44817 | 0.30549 |
| NCIH2291 | 0.35013 | -0.3184 | -0.2087 | 0.05378 | 0.08449 | 0.25787 | 3.19681 | 0.01469 | | 0.00683 | 0.01287 | 0.07502 | 0.3568 | 0.28191 |
| KYSE140 | 0.1927 | -0.3212 | -0.3287 | 0.02341 | 0.10126 | 0.25688 | 3.18025 | 0.01507 | | 0.09233 | 0.01221 | 0.01056 | 0.43656 | 0.24438 |
| A253 | 0.17287 | -0.3011 | -0.3555 | -0.0866 | 0.05858 | 0.2565 | 3.17385 | 0.01522 | | 0.11745 | 0.01775 | 0.0061 | 0.27712 | 0.34464 |
| NCIH1155 | 0.12336 | -0.2115 | -0.054 | -0.3619 | 0.26455 | 0.2563 | 3.17064 | 0.0153 | | 0.1992 | 0.07229 | 0.35623 | 0.0053 | 0.03311 |
| HCC1937 | 0.34777 | -0.3455 | -0.1937 | -0.0459 | 0.04951 | 0.25521 | 3.15246 | 0.01574 | | 0.00717 | 0.00752 | 0.09114 | 0.37695 | 0.36773 |
| C32 | 0.13805 | -0.4751 | 0.05892 | 0.02149 | -0.2367 | 0.25481 | 3.14579 | 0.0159 | | 0.17208 | 0.00028 | 0.3438 | 0.44173 | 0.05075 |
| HEL | 0.10765 | 0.10442 | -0.1184 | -0.265 | 0.40072 | 0.25166 | 3.09386 | 0.01725 | | 0.23079 | 0.23761 | 0.20894 | 0.03289 | 0.00216 |
| C75 | 0.02176 | -0.2729 | -0.3903 | -0.0824 | 0.05682 | 0.25105 | 3.08393 | 0.01752 | | 0.44101 | 0.02889 | 0.00278 | 0.2868 | 0.34909 |
| SNB75 | -0.0585 | -0.0345 | -0.4153 | -0.0293 | 0.25623 | 0.25049 | 3.07472 | 0.01777 | | 0.34474 | 0.4071 | 0.0015 | 0.42081 | 0.03777 |
| HCC1806 | 0.16571 | -0.4057 | -0.1981 | -0.0192 | 0.13425 | 0.25017 | 3.06938 | 0.01792 | | 0.12758 | 0.00191 | 0.08617 | 0.44787 | 0.17888 |
| SNU1066 | 0.15256 | -0.3339 | -0.3117 | -0.0105 | 0.12164 | 0.25008 | 3.06796 | 0.01796 | | 0.14767 | 0.00951 | 0.01462 | 0.47155 | 0.20254 |
| NALM16 | 0.09703 | -0.3232 | 0.09804 | -0.3586 | 0.1531 | 0.2493 | 3.05522 | 0.01832 | | 0.25358 | 0.01175 | 0.25136 | 0.0057 | 0.14681 |
| MFE280 | 0.01468 | -0.2172 | -0.3832 | -0.1999 | 0.00482 | 0.24913 | 3.05251 | 0.0184 | | 0.46013 | 0.06689 | 0.00328 | 0.08428 | 0.48688 |
| PECAPJ15 | 0.05682 | -0.3389 | -0.3315 | 0.01273 | 0.09377 | 0.24876 | 3.04643 | 0.01858 | | 0.34909 | 0.0086 | 0.01 | 0.46542 | 0.26081 |
| CAMA1 | 0.27141 | -0.2284 | -0.0448 | -0.2506 | 0.28708 | 0.24846 | 3.04156 | 0.01872 | | 0.02962 | 0.05725 | 0.37998 | 0.04122 | 0.02275 |
| WM2664 | 0.17296 | -0.3177 | -0.1683 | -0.1119 | 0.26998 | 0.24842 | 3.04085 | 0.01874 | | 0.11733 | 0.01305 | 0.12383 | 0.22205 | 0.03032 |
| NCIH2882 | -0.1627 | -0.3036 | -0.2658 | 0.01355 | 0.14648 | 0.24836 | 3.03998 | 0.01877 | | 0.13197 | 0.01697 | 0.03247 | 0.4632 | 0.15761 |
| PSS008 | 0.02652 | -0.3828 | -0.2825 | -0.0834 | 0.05376 | 0.24823 | 3.03778 | 0.01883 | | 0.42822 | 0.00331 | 0.0246 | 0.28432 | 0.35687 |
| AU565 | 0.10066 | -0.292 | -0.3861 | 0.0018 | -0.0907 | 0.24712 | 3.01982 | 0.01937 | | 0.24567 | 0.0209 | 0.00307 | 0.49511 | 0.26761 |
| RL952 | 0.12336 | -0.249 | -0.3719 | -0.0852 | 0.12922 | 0.24703 | 3.01828 | 0.01941 | | 0.1992 | 0.04224 | 0.00425 | 0.28018 | 0.1881 |
| WM115 | 0.10233 | -0.3728 | -0.1717 | 0.03177 | 0.22073 | 0.24658 | 3.01105 | 0.01964 | | 0.24207 | 0.00417 | 0.11904 | 0.41423 | 0.06375 |
| SH10TC | -0.0367 | -0.327 | -0.2112 | -0.199 | 0.16504 | 0.24607 | 3.00272 | 0.01989 | | 0.40127 | 0.01091 | 0.07255 | 0.08524 | 0.12856 |
| OSC19 | 0.1268 | -0.1419 | -0.4356 | -0.1034 | 0.04222 | 0.24513 | 2.98753 | 0.02037 | | 0.19265 | 0.16531 | 0.00088 | 0.23982 | 0.38667 |
| A431 | 0.11145 | -0.2282 | -0.4147 | -0.0064 | 0.06091 | 0.24367 | 2.96395 | 0.02114 | | 0.22292 | 0.0574 | 0.00152 | 0.48254 | 0.3388 |
| WM1799 | 0.0758 | -0.3617 | -0.3141 | -0.0885 | -0.0097 | 0.2436 | 2.96294 | 0.02117 | | 0.30235 | 0.00533 | 0.01397 | 0.27268 | 0.47354 |
| IGROV1 | 0.31718 | -0.2344 | -0.1273 | -0.2859 | 0.13532 | 0.24313 | 2.95526 | 0.02143 | | 0.01319 | 0.05251 | 0.19163 | 0.02321 | 0.17695 |
| NCIH647 | -0.1982 | -0.3894 | -0.178 | 0.04055 | -0.1004 | 0.2427 | 2.94835 | 0.02166 | | 0.08613 | 0.00284 | 0.11051 | 0.39103 | 0.24634 |
| SNU1327 | 0.151 | -0.3631 | -0.2173 | -0.1939 | 0.09213 | 0.24203 | 2.93767 | 0.02203 | | 0.15018 | 0.00517 | 0.06687 | 0.09095 | 0.26448 |
| OE21 | 0.2476 | -0.225 | -0.3667 | 0.0438 | 0.02787 | 0.24082 | 2.9183 | 0.02271 | | 0.04315 | 0.06003 | 0.00477 | 0.38252 | 0.42462 |
| ESO26 | 0.03406 | -0.1982 | -0.4248 | -0.1006 | -0.0319 | 0.24062 | 2.91515 | 0.02282 | | 0.40813 | 0.08612 | 0.00118 | 0.2457 | 0.41386 |
| SW948 | -0.162 | -0.188 | -0.1296 | -0.2019 | 0.28616 | 0.24058 | 2.91451 | 0.02285 | | 0.13307 | 0.09785 | 0.18747 | 0.08212 | 0.02312 |
| 9505BIK | 0.14267 | -0.258 | -0.3635 | -0.1438 | -0.0183 | 0.24002 | 2.90555 | 0.02317 | | 0.16405 | 0.03674 | 0.00512 | 0.16211 | 0.45038 |
| EPLC272H | 0.1303 | -0.3135 | -0.2335 | 0.0284 | 0.23754 | 0.23947 | 2.89682 | 0.02349 | | 0.1861 | 0.01412 | 0.0532 | 0.42319 | 0.05015 |
| NCIH2171 | 0.22516 | -0.2436 | -0.11 | -0.2439 | 0.26663 | 0.23896 | 2.88877 | 0.02379 | | 0.05992 | 0.04584 | 0.22588 | 0.04566 | 0.03202 |
| JHU022 | 0.08268 | -0.3844 | -0.2681 | -0.0857 | 0.04292 | 0.23837 | 2.87929 | 0.02415 | | 0.2861 | 0.0032 | 0.03127 | 0.279 | 0.38482 |
| MFE319 | 0.04214 | -0.2804 | -0.3 | -0.179 | 0.13731 | 0.23786 | 2.87133 | 0.02445 | | 0.38686 | 0.0255 | 0.01811 | 0.10919 | 0.1734 |
| HT1197 | -0.1033 | -0.3035 | -0.1728 | -0.0647 | 0.24332 | 0.23748 | 2.86522 | 0.02469 | | 0.2399 | 0.01699 | 0.11758 | 0.32938 | 0.04603 |
| SNUC4 | 0.15953 | -0.3424 | -0.2832 | -0.1398 | -0.1065 | 0.23739 | 2.86383 | 0.02474 | | 0.13678 | 0.00801 | 0.0243 | 0.16901 | 0.23328 |
| SUPM2 | 0.1718 | -0.3126 | 0.24038 | -0.1346 | 0.22915 | 0.23685 | 2.85534 | 0.02507 | | 0.11893 | 0.01437 | 0.04808 | 0.17818 | 0.05662 |
| TTC442 | 0.05793 | -0.2841 | -0.3514 | -0.1059 | 0.07945 | 0.23661 | 2.85145 | 0.02523 | | 0.34628 | 0.02395 | 0.00665 | 0.23456 | 0.29368 |
| A388 | 0.17062 | -0.2342 | -0.3496 | -0.1719 | 0.02497 | 0.23658 | 2.85097 | 0.02524 | | 0.12057 | 0.05267 | 0.0069 | 0.11878 | 0.43238 |
| SNU1041 | 0.15325 | -0.2396 | -0.2425 | -0.1436 | 0.26565 | 0.23648 | 2.84948 | 0.0253 | | 0.14656 | 0.04861 | 0.04658 | 0.1624 | 0.03253 |
| OVMIU | 0.10044 | -0.3413 | -0.1727 | -0.1977 | 0.18476 | 0.23601 | 2.84211 | 0.0256 | | 0.24615 | 0.00821 | 0.11774 | 0.08662 | 0.10188 |
| UACC62 | 0.26411 | -0.3386 | -0.1009 | -0.0487 | 0.22337 | 0.23532 | 2.83119 | 0.02604 | | 0.03335 | 0.00865 | 0.24522 | 0.36978 | 0.06144 |
| C99 | -0.0039 | -0.3776 | -0.1721 | -0.205 | 0.09218 | 0.23492 | 2.82486 | 0.0263 | | 0.48948 | 0.00374 | 0.11851 | 0.07886 | 0.26436 |
| TEN | 0.09032 | -0.3016 | -0.3402 | -0.127 | 0.01174 | 0.23454 | 2.81895 | 0.02655 | | 0.26856 | 0.0176 | 0.00838 | 0.19227 | 0.4681 |
| HCC1954 | 0.42198 | -0.0581 | -0.0455 | -0.0294 | 0.27798 | 0.2345 | 2.8183 | 0.02658 | | 0.00126 | 0.34595 | 0.37808 | 0.42044 | 0.02656 |

| | | | | | | | | | | | | | |
|---|---|---|---|---|---|---|---|---|---|---|---|---|---|
| KCL22 | 0.03042 | -0.2946 | -0.2384 | -0.0123 | 0.25355 | 0.23397 | 2.81004 | 0.02692 | 0.4178 | 0.01994 | 0.04954 | 0.46645 | 0.03938 |
| TTC709 | 0.20103 | -0.2263 | -0.2542 | -0.0758 | 0.26879 | 0.23391 | 2.80899 | 0.02697 | 0.08302 | 0.05893 | 0.03898 | 0.3024 | 0.03092 |
| OAW42 | 0.05446 | -0.3942 | -0.0188 | -0.1295 | 0.21252 | 0.23325 | 2.79862 | 0.02741 | 0.35508 | 0.00253 | 0.44892 | 0.18759 | 0.07132 |
| NCIH661 | 0.11989 | -0.3338 | -0.3275 | 0.01407 | 0.02012 | 0.23275 | 2.79081 | 0.02775 | 0.20595 | 0.00953 | 0.01081 | 0.46179 | 0.44543 |
| UMUC16 | 0.05394 | -0.0565 | -0.123 | -0.2169 | 0.4007 | 0.23209 | 2.7805 | 0.0282 | 0.3564 | 0.34983 | 0.19991 | 0.06721 | 0.00216 |
| HCC827 | -0.0382 | -0.3739 | -0.1001 | -0.1859 | 0.15646 | 0.23149 | 2.77122 | 0.02862 | 0.39724 | 0.00406 | 0.24691 | 0.10048 | 0.1415 |
| CORL105 | 0.23184 | -0.2645 | -0.1945 | -0.2968 | -0.0537 | 0.23138 | 2.76944 | 0.0287 | 0.05448 | 0.03313 | 0.09023 | 0.01918 | 0.35708 |
| SNU182 | 0.02566 | -0.4088 | -0.1522 | 0.02378 | 0.13817 | 0.23114 | 2.76569 | 0.02887 | 0.43053 | 0.00177 | 0.14823 | 0.43557 | 0.17188 |
| 769P | 0.12442 | -0.2747 | -0.3507 | -0.0844 | 0.07386 | 0.23063 | 2.75786 | 0.02923 | 0.19717 | 0.02803 | 0.00675 | 0.28221 | 0.30701 |
| COV413A | 0.16606 | -0.1609 | -0.3731 | -0.1788 | 0.03076 | 0.23039 | 2.75407 | 0.0294 | 0.12708 | 0.13472 | 0.00414 | 0.10949 | 0.41691 |
| MCIXC | -0.0259 | -0.3531 | 0.18062 | -0.3177 | 0.0013 | 0.22835 | 2.72255 | 0.0309 | 0.42995 | 0.00641 | 0.10713 | 0.01306 | 0.49647 |
| ES4 | 0.24401 | -0.3248 | -0.2108 | -0.1855 | 0.05292 | 0.22785 | 2.71479 | 0.03128 | 0.04555 | 0.01139 | 0.07295 | 0.10101 | 0.35901 |
| NB69 | 0.11227 | -0.3618 | -0.1506 | -0.1377 | 0.18939 | 0.22719 | 2.70457 | 0.03179 | 0.22124 | 0.00532 | 0.15082 | 0.1727 | 0.09623 |
| MDAMB415 | 0.35548 | 0.0407 | -0.1251 | 0.11828 | 0.30432 | 0.22718 | 2.70447 | 0.03179 | 0.0061 | 0.39064 | 0.19589 | 0.20912 | 0.01675 |
| M07E | 0.25736 | -0.1135 | -0.1136 | -0.3204 | 0.22429 | 0.22692 | 2.70052 | 0.03199 | 0.03711 | 0.21872 | 0.21858 | 0.0124 | 0.06066 |
| KYSE410 | 0.01274 | -0.2077 | -0.41 | -0.0772 | -0.0191 | 0.22633 | 2.69131 | 0.03246 | 0.46538 | 0.07603 | 0.00172 | 0.29895 | 0.44811 |
| BLUE1 | 0.25646 | -0.0008 | -0.2461 | -0.1912 | 0.27346 | 0.22617 | 2.68899 | 0.03258 | 0.03764 | 0.49773 | 0.04411 | 0.09405 | 0.02864 |
| CCLFPEDS0001T | 0.26369 | -0.2756 | -0.0591 | -0.1788 | 0.25369 | 0.22565 | 2.68092 | 0.03299 | 0.03357 | 0.02763 | 0.34329 | 0.10952 | 0.0393 |
| KU1919 | 0.21118 | -0.3071 | -0.0207 | -0.276 | 0.18388 | 0.22482 | 2.66825 | 0.03366 | 0.07262 | 0.01591 | 0.44377 | 0.02746 | 0.10298 |
| COLO679 | -0.0132 | -0.2186 | -0.3516 | 0.039 | 0.18309 | 0.22364 | 2.65015 | 0.03463 | 0.46407 | 0.06569 | 0.00662 | 0.39511 | 0.10398 |
| PACADD119 | 0.03319 | -0.1844 | -0.4065 | -0.0904 | 0.05939 | 0.22357 | 2.64915 | 0.03469 | 0.41046 | 0.10227 | 0.00187 | 0.26846 | 0.34261 |
| NCIH1793 | 0.11801 | -0.3686 | -0.2404 | -0.015 | 0.10432 | 0.22322 | 2.64378 | 0.03498 | 0.20966 | 0.00458 | 0.04806 | 0.45932 | 0.23781 |
| IM95 | 0.17472 | -0.0749 | -0.3817 | -0.1234 | -0.1314 | 0.22322 | 2.64373 | 0.03498 | 0.11493 | 0.30443 | 0.0034 | 0.19918 | 0.18399 |
| HCA1 | 0.23012 | -0.3003 | -0.2698 | -0.1367 | 0.0309 | 0.22316 | 2.64288 | 0.03503 | 0.05584 | 0.018 | 0.0304 | 0.17442 | 0.41654 |
| HARA | 0.07713 | -0.3164 | -0.3307 | -0.0317 | 0.03723 | 0.2226 | 2.63425 | 0.03551 | 0.29918 | 0.01337 | 0.01015 | 0.41429 | 0.39977 |
| NB6 | 0.17826 | -0.1747 | 0.01796 | -0.122 | 0.3888 | 0.22232 | 2.63 | 0.03575 | 0.1102 | 0.11494 | 0.45125 | 0.20193 | 0.00288 |
| HSC39 | 0.05584 | -0.1079 | -0.4356 | 0.13772 | 0.07274 | 0.22212 | 2.62701 | 0.03592 | 0.35158 | 0.2302 | 0.00088 | 0.17267 | 0.3097 |
| TF1 | 0.00306 | -0.3555 | -0.0411 | -0.1556 | 0.22288 | 0.22199 | 2.62497 | 0.03604 | 0.49167 | 0.00609 | 0.38961 | 0.14282 | 0.06186 |
| CW9019 | 0.16949 | -0.0568 | -0.3336 | -0.05 | 0.28608 | 0.22175 | 2.62146 | 0.03624 | 0.12216 | 0.34922 | 0.00958 | 0.36654 | 0.02314 |
| CCC5 | 0.10411 | -0.3613 | -0.0664 | -0.1042 | 0.23524 | 0.22146 | 2.61706 | 0.03649 | 0.23826 | 0.00537 | 0.32506 | 0.23808 | 0.05186 |
| SKNMC | -0.1234 | -0.3624 | -0.2324 | 0.07911 | -0.0222 | 0.22139 | 2.61595 | 0.03655 | 0.19916 | 0.00524 | 0.05407 | 0.29449 | 0.43995 |
| NCIN87 | -0.033 | -0.1625 | -0.426 | -0.0364 | 0.02559 | 0.22105 | 2.61084 | 0.03685 | 0.41086 | 0.13227 | 0.00114 | 0.40201 | 0.43072 |
| ZR751 | 0.1069 | -0.2198 | -0.3333 | -0.2068 | 0.03449 | 0.22094 | 2.60911 | 0.03695 | 0.23236 | 0.0646 | 0.00964 | 0.07701 | 0.40701 |
| VMRCRCW | 0.07805 | -0.2068 | -0.1076 | -0.4025 | 0.06802 | 0.22078 | 2.60666 | 0.03709 | 0.29699 | 0.07699 | 0.23085 | 0.00207 | 0.32119 |
| SKGII | 0.18437 | -0.2058 | -0.3806 | -0.0253 | 0.03418 | 0.22075 | 2.60629 | 0.03711 | 0.10237 | 0.07805 | 0.00349 | 0.43145 | 0.40783 |
| C84 | 0.13022 | -0.0012 | -0.411 | -0.1296 | -0.0686 | 0.22047 | 2.60204 | 0.03736 | 0.18624 | 0.49686 | 0.00167 | 0.18747 | 0.3198 |
| RMGI | 0.12401 | -0.2763 | -0.2884 | -0.1897 | 0.08062 | 0.22018 | 2.59753 | 0.03763 | 0.19796 | 0.0273 | 0.02225 | 0.0958 | 0.29093 |
| 127399 | -0.1214 | -0.1885 | -0.2868 | -0.0516 | 0.23242 | 0.22011 | 2.5965 | 0.03769 | 0.20298 | 0.0973 | 0.02286 | 0.3623 | 0.05402 |
| HT144 | 0.33503 | -0.333 | 0.02521 | -0.1229 | 0.13285 | 0.21988 | 2.59312 | 0.03789 | 0.00931 | 0.0097 | 0.43174 | 0.20008 | 0.18142 |
| DEL | 0.17833 | -0.3351 | -0.0196 | -0.0858 | 0.26333 | 0.2198 | 2.59184 | 0.03797 | 0.11011 | 0.0093 | 0.44687 | 0.27884 | 0.03377 |
| ECC2 | 0.14522 | -0.2109 | -0.382 | -0.0352 | 0.07106 | 0.21937 | 2.58537 | 0.03836 | 0.15973 | 0.07293 | 0.00338 | 0.40506 | 0.31378 |
| CAL27 | 0.15191 | -0.2111 | -0.3881 | -0.0112 | 0.03386 | 0.21888 | 2.57797 | 0.03881 | 0.14871 | 0.07265 | 0.00293 | 0.46955 | 0.40868 |
| D542MG | 0.18054 | -0.1499 | -0.0383 | 0.13943 | 0.37383 | 0.21875 | 2.57604 | 0.03893 | 0.10723 | 0.15192 | 0.39699 | 0.16966 | 0.00407 |
| CHLA266 | 0.08686 | -0.0489 | -0.2159 | -0.0377 | 0.4037 | 0.21862 | 2.57398 | 0.03905 | 0.27644 | 0.36937 | 0.06809 | 0.39855 | 0.00201 |
| NCIH358 | 0.1466 | -0.3671 | -0.1123 | -0.1763 | 0.15557 | 0.21828 | 2.56899 | 0.03936 | 0.15741 | 0.00473 | 0.22107 | 0.11285 | 0.1429 |
| OUMS23 | 0.06336 | -0.3098 | -0.1846 | -0.0221 | 0.24932 | 0.21713 | 2.5516 | 0.04046 | 0.33269 | 0.01515 | 0.1021 | 0.43998 | 0.04203 |
| OVCA420 | 0.02265 | -0.3219 | -0.2893 | -0.0853 | 0.08839 | 0.21711 | 2.55133 | 0.04047 | 0.43861 | 0.01204 | 0.02191 | 0.28006 | 0.27295 |
| HKA1 | -0.0535 | -0.334 | -0.286 | -0.0692 | 0.02394 | 0.21698 | 2.54932 | 0.0406 | 0.35742 | 0.00951 | 0.02316 | 0.31833 | 0.43514 |
| BICR31 | 0.30077 | -0.1916 | -0.2445 | -0.0674 | 0.19596 | 0.21695 | 2.549 | 0.04062 | 0.01787 | 0.09359 | 0.04521 | 0.32269 | 0.0886 |
| CHLA57 | 0.0861 | -0.2504 | -0.2269 | -0.057 | 0.27657 | 0.21667 | 2.5447 | 0.0409 | 0.2782 | 0.04134 | 0.05845 | 0.34869 | 0.0272 |
| OC314 | 0.30313 | -0.2201 | -0.1124 | -0.0869 | 0.27133 | 0.21666 | 2.54453 | 0.04091 | 0.01712 | 0.06427 | 0.22102 | 0.27642 | 0.02966 |
| GP5D | 0.15175 | -0.3548 | -0.1271 | -0.1766 | 0.15884 | 0.21607 | 2.53581 | 0.04148 | 0.14896 | 0.00618 | 0.19203 | 0.11246 | 0.13783 |
| TGBC18TKB | 0.03618 | -0.1922 | -0.3601 | -0.1808 | -0.0912 | 0.21589 | 2.53308 | 0.04166 | 0.40252 | 0.09288 | 0.00552 | 0.10686 | 0.26658 |
| CFPAC1 | -0.0152 | -0.1367 | -0.423 | -0.085 | -0.0314 | 0.21557 | 2.5282 | 0.04198 | 0.45865 | 0.17444 | 0.00123 | 0.28078 | 0.4151 |
| HEC265 | 0.27789 | -0.3636 | -0.1005 | -0.1156 | 0.08913 | 0.21455 | 2.51304 | 0.04299 | 0.0266 | 0.00511 | 0.24611 | 0.21452 | 0.27126 |
| HCC15 | 0.25223 | -0.0044 | 0.08514 | -0.2098 | 0.35777 | 0.21434 | 2.50991 | 0.04321 | 0.0402 | 0.48811 | 0.28041 | 0.07397 | 0.0058 |
| KYSE30 | 0.05047 | -0.1951 | -0.3892 | -0.0788 | 0.07539 | 0.21419 | 2.5076 | 0.04336 | 0.36529 | 0.08959 | 0.00285 | 0.29513 | 0.30333 |
| SW954 | 0.13352 | -0.2434 | -0.3213 | -0.1788 | 0.01998 | 0.21399 | 2.50473 | 0.04356 | 0.1802 | 0.04598 | 0.01219 | 0.1095 | 0.4458 |
| MERO25 | 0.23057 | -0.3262 | -0.055 | -0.1113 | 0.22609 | 0.21387 | 2.50289 | 0.04369 | 0.05548 | 0.01108 | 0.35366 | 0.22324 | 0.05913 |
| RERFLCSQ1 | 0.07659 | -0.2715 | -0.3473 | -0.062 | 0.04663 | 0.21231 | 2.47973 | 0.04531 | 0.30046 | 0.02958 | 0.00724 | 0.33604 | 0.37518 |
| OVISE | 0.12372 | -0.3553 | -0.1757 | -0.2146 | 0.05141 | 0.21203 | 2.47563 | 0.04561 | 0.19851 | 0.00612 | 0.11356 | 0.06933 | 0.36288 |
| JHOS2 | 0.24069 | -0.2188 | -0.3167 | -0.0582 | 0.09404 | 0.21193 | 2.47406 | 0.04572 | 0.04786 | 0.06546 | 0.01331 | 0.34564 | 0.26021 |
| H357 | -0.0045 | -0.2547 | -0.3077 | -0.1377 | 0.11976 | 0.21186 | 2.47302 | 0.0458 | 0.48788 | 0.0387 | 0.01575 | 0.17277 | 0.20622 |
| ES2 | 0.03681 | -0.2773 | -0.2122 | -0.1918 | 0.18838 | 0.21184 | 2.4727 | 0.04582 | 0.40088 | 0.02687 | 0.07165 | 0.09332 | 0.09745 |
| KARPAS422 | 0.06628 | -0.0616 | -0.0771 | -0.2354 | 0.3768 | 0.21178 | 2.47191 | 0.04588 | 0.32547 | 0.33717 | 0.2993 | 0.05173 | 0.00381 |
| HCC78 | 0.06779 | -0.2497 | -0.2445 | -0.2407 | 0.13075 | 0.21154 | 2.46839 | 0.04613 | 0.32176 | 0.04177 | 0.04518 | 0.04784 | 0.18526 |
| NCIH1869 | 0.20094 | -0.1855 | -0.1728 | -0.2181 | 0.25569 | 0.21103 | 2.46077 | 0.04669 | 0.08311 | 0.101 | 0.11751 | 0.06613 | 0.03809 |
| TO14 | 0.13442 | -0.4006 | 0.02328 | -0.23 | 0.06864 | 0.21013 | 2.44753 | 0.04768 | 0.17857 | 0.00217 | 0.43693 | 0.05596 | 0.31966 |
| LS | -0.1028 | -0.3907 | -0.1311 | 0.04357 | 0.05912 | 0.2101 | 2.44704 | 0.04771 | 0.24115 | 0.00276 | 0.18458 | 0.38312 | 0.34328 |
| BL70 | 0.07836 | 0.0067 | -0.1178 | -0.1554 | 0.41545 | 0.20989 | 2.44389 | 0.04795 | 0.29626 | 0.48179 | 0.21016 | 0.14323 | 0.0015 |
| SW1116 | 0.00435 | -0.2648 | -0.3485 | -0.0915 | 0.00068 | 0.20946 | 2.43725 | 0.04846 | 0.48817 | 0.03298 | 0.00706 | 0.26595 | 0.49815 |
| UPCISCC116 | 0.05339 | -0.284 | -0.3498 | -0.018 | -0.0674 | 0.20935 | 2.43603 | 0.04855 | 0.3578 | 0.024 | 0.00687 | 0.45119 | 0.3227 |
| ICC106 | 0.1304 | -0.2308 | -0.2671 | -0.043 | 0.24019 | 0.20869 | 2.42626 | 0.0493 | 0.18592 | 0.0553 | 0.03177 | 0.38467 | 0.04822 |
| UPCISCC111 | 0.11996 | -0.3361 | -0.2887 | 0.04472 | -0.0574 | 0.20792 | 2.41505 | 0.05018 | 0.20581 | 0.00911 | 0.02211 | 0.38013 | 0.34756 |
| 697 | -0.1176 | -0.2583 | -0.2866 | -0.0359 | 0.12696 | 0.20777 | 2.41285 | 0.05036 | 0.21049 | 0.03656 | 0.02292 | 0.40317 | 0.19233 |
| SCC4 | 0.08285 | -0.2338 | -0.3274 | 0.00181 | 0.17117 | 0.20775 | 2.41243 | 0.05039 | 0.2857 | 0.05293 | 0.01082 | 0.49506 | 0.11981 |
| JIMT1 | 0.15502 | -0.2764 | -0.239 | -0.1863 | 0.12544 | 0.20685 | 2.39936 | 0.05144 | 0.14376 | 0.02726 | 0.04908 | 0.10002 | 0.19522 |
| LB1047RCC | 0.22043 | -0.1962 | -0.3358 | -0.055 | 0.08955 | 0.20685 | 2.39925 | 0.05145 | 0.06401 | 0.08837 | 0.00916 | 0.35359 | 0.2703 |
| UPCISCC154 | 0.21531 | -0.259 | -0.2801 | -0.0802 | 0.11552 | 0.20637 | 2.39233 | 0.05201 | 0.06867 | 0.03615 | 0.02562 | 0.29189 | 0.21464 |
| ESO51 | -0.1488 | -0.3988 | 0.01972 | 0.03818 | 0.02916 | 0.2063 | 2.39129 | 0.0521 | 0.1538 | 0.00227 | 0.44652 | 0.39725 | 0.42118 |
| HGC27 | 0.07934 | -0.1554 | -0.1869 | -0.0254 | 0.35998 | 0.20586 | 2.3848 | 0.05264 | 0.29394 | 0.14321 | 0.09928 | 0.43122 | 0.00553 |
| C2BBE1 | 0.12485 | -0.1754 | -0.3759 | -0.0993 | 0.05944 | 0.2058 | 2.38394 | 0.05271 | 0.19635 | 0.11402 | 0.00389 | 0.24856 | 0.34248 |
| OVTOKO | 0.13917 | -0.2025 | -0.2593 | -0.2284 | 0.14951 | 0.20544 | 2.37879 | 0.05314 | 0.17013 | 0.08148 | 0.03599 | 0.05726 | 0.1526 |
| LN319 | 0.10036 | -0.2477 | -0.3281 | -0.1273 | 0.05568 | 0.20359 | 2.35184 | 0.05544 | 0.24632 | 0.0431 | 0.01069 | 0.19165 | 0.35197 |
| MCC13 | 0.16178 | -0.3401 | -0.0186 | -0.2276 | 0.16119 | 0.20347 | 2.35014 | 0.05559 | 0.13338 | 0.0084 | 0.44951 | 0.05786 | 0.13427 |
| PA1 | 0.17459 | -0.2868 | -0.1148 | -0.0377 | 0.27001 | 0.20346 | 2.34996 | 0.05561 | 0.1151 | 0.02288 | 0.21614 | 0.39844 | 0.03031 |
| LC1SQ | 0.21963 | -0.2838 | -0.2593 | -0.0874 | -0.1286 | 0.20283 | 2.34078 | 0.05642 | 0.06472 | 0.02406 | 0.03602 | 0.27522 | 0.18923 |
| MKN45 | -0.0246 | -0.3662 | -0.0729 | -0.0775 | 0.17956 | 0.20267 | 2.33849 | 0.05662 | 0.43338 | 0.00483 | 0.30933 | 0.29819 | 0.1085 |
| PACADD137 | 0.22348 | -0.1347 | -0.2493 | -0.2731 | 0.01705 | 0.20236 | 2.33405 | 0.05702 | 0.06135 | 0.17804 | 0.04204 | 0.02881 | 0.45373 |
| RVH421 | 0.01413 | -0.369 | -0.0991 | -0.1914 | 0.11006 | 0.20215 | 2.33101 | 0.0573 | 0.46163 | 0.00454 | 0.24899 | 0.09383 | 0.22577 |
| KYSE270 | 0.27128 | -0.0853 | -0.2723 | -0.1042 | 0.21281 | 0.20141 | 2.32028 | 0.05827 | 0.02968 | 0.27998 | 0.0292 | 0.2381 | 0.07104 |
| FLO1 | 0.18067 | -0.2154 | -0.1224 | -0.1527 | 0.29502 | 0.2011 | 2.31587 | 0.05868 | 0.10706 | 0.06858 | 0.20105 | 0.1474 | 0.0198 |
| PK1 | 0.07991 | -0.1863 | -0.3902 | -0.044 | 0.02926 | 0.20087 | 2.31249 | 0.05899 | 0.2926 | 0.09998 | 0.00279 | 0.38199 | 0.42091 |
| TKKK | 0.11398 | -0.2311 | -0.2796 | -0.2359 | -0.0081 | 0.20074 | 2.31064 | 0.05917 | 0.21776 | 0.05505 | 0.02585 | 0.05135 | 0.47808 |
| LOVO | 0.30838 | 0.02641 | -0.231 | -0.2104 | 0.08485 | 0.20071 | 2.31027 | 0.0592 | 0.01555 | 0.42853 | 0.05516 | 0.07334 | 0.28106 |
| LN229 | 0.01891 | -0.2281 | -0.365 | -0.0854 | -0.0262 | 0.20038 | 2.30543 | 0.05965 | 0.4487 | 0.05745 | 0.00496 | 0.27989 | 0.42905 |
| SKBR3 | 0.09167 | -0.2524 | -0.3512 | -0.0298 | 0.03045 | 0.20035 | 2.30504 | 0.05969 | 0.2655 | 0.04007 | 0.00667 | 0.41937 | 0.41774 |
| COV434 | 0.02549 | -0.3276 | -0.1969 | -0.205 | 0.06583 | 0.1998 | 2.2971 | 0.06044 | 0.43099 | 0.01079 | 0.08757 | 0.0788 | 0.32657 |
| H413 | 0.07144 | -0.1271 | -0.4161 | -0.0141 | 0.03947 | 0.19944 | 2.29192 | 0.06094 | 0.31285 | 0.19207 | 0.00147 | 0.46162 | 0.39386 |
| JHH4 | 0.05048 | -0.218 | -0.3555 | 0.00318 | 0.11655 | 0.19901 | 2.28577 | 0.06153 | 0.36526 | 0.06619 | 0.0061 | 0.49135 | 0.21258 |
| UPCISCC074 | 0.20587 | -0.2048 | -0.3204 | -0.1222 | 0.02429 | 0.19895 | 2.28495 | 0.06161 | 0.07793 | 0.07902 | 0.0124 | 0.20137 | 0.4342 |
| SKMEL24 | 0.01514 | -0.2132 | -0.2539 | -0.2835 | -0.0062 | 0.19891 | 2.28436 | 0.06167 | 0.45887 | 0.07064 | 0.03916 | 0.0242 | 0.48306 |
| COV644 | 0.22707 | -0.1626 | -0.3024 | -0.1778 | 0.02591 | 0.19887 | 2.28378 | 0.06172 | 0.05832 | 0.13219 | 0.01734 | 0.11087 | 0.42987 |
| SKN | -0.1057 | -0.2184 | -0.2696 | -0.0386 | 0.19167 | 0.19838 | 2.27678 | 0.06241 | 0.23491 | 0.06579 | 0.03049 | 0.39603 | 0.09352 |
| RPE1SS77 | 0.30485 | -0.287 | -0.19 | -0.0534 | 0.05801 | 0.19827 | 2.27519 | 0.06257 | 0.01659 | 0.02279 | 0.09551 | 0.35773 | 0.34609 |
| ETCC016 | 0.12211 | -0.3852 | -0.1437 | -0.0948 | 0.09177 | 0.19815 | 2.27353 | 0.06273 | 0.20161 | 0.00314 | 0.16223 | 0.25854 | 0.26528 |
| SNU308 | 0.0042 | -0.2414 | -0.3482 | -0.1009 | -0.0458 | 0.19809 | 2.27266 | 0.06282 | 0.48858 | 0.04734 | 0.00711 | 0.24517 | 0.37737 |
| COLO320 | -0.2649 | -0.1737 | -0.2051 | -0.1889 | -0.0527 | 0.19805 | 2.27202 | 0.06288 | 0.03292 | 0.11627 | 0.07876 | 0.09684 | 0.35962 |
| OC316 | 0.17412 | -0.2087 | -0.3486 | 0.00925 | 0.06635 | 0.19751 | 2.26427 | 0.06365 | 0.11574 | 0.0751 | 0.00705 | 0.47486 | 0.32528 |
| KATOIII | 0.05767 | 0.06685 | -0.3997 | 0.13044 | 0.1815 | 0.19702 | 2.25737 | 0.06435 | 0.34693 | 0.32405 | 0.00222 | 0.18583 | 0.10599 |
| HUNS1 | 0.12658 | -0.1151 | -0.1547 | -0.1742 | 0.3366 | 0.19695 | 2.25635 | 0.06445 | 0.19307 | 0.21557 | 0.14431 | 0.1157 | 0.00902 |
| ISHIKAWAHERAKLIO02E | 0.05632 | -0.4025 | -0.0639 | -0.0196 | 0.12841 | 0.19678 | 2.25389 | 0.0647 | 0.35036 | 0.00207 | 0.33136 | 0.44681 | 0.18962 |
| NCIH2004RT | 0.28601 | -0.0569 | 0.09647 | 0.14965 | 0.29461 | 0.19654 | 2.25052 | 0.06505 | 0.02317 | 0.34878 | 0.25482 | 0.15237 | 0.01995 |
| MJ | 0.09572 | -0.3018 | 0.06146 | -0.3582 | -0.0286 | 0.19654 | 2.25047 | 0.06505 | 0.25649 | 0.01754 | 0.33743 | 0.00575 | 0.42269 |
| RPE1SS48 | 0.11906 | -0.358 | -0.1566 | -0.1091 | 0.12838 | 0.19608 | 2.24386 | 0.06573 | 0.20759 | 0.00578 | 0.14122 | 0.22771 | 0.18967 |
| PEO1 | 0.04339 | -0.2604 | -0.3328 | 0.04251 | 0.07771 | 0.19607 | 2.24382 | 0.06574 | 0.3836 | 0.03536 | 0.00973 | 0.3859 | 0.29781 |
| PCI15A | 0.28971 | -0.1736 | -0.2222 | -0.1329 | 0.16404 | 0.19521 | 2.23158 | 0.06701 | 0.02173 | 0.1164 | 0.06242 | 0.18127 | 0.13002 |
| PECAPJ34CLONEC12 | 0.20045 | -0.2508 | -0.2682 | -0.1682 | 0.00058 | 0.19486 | 2.22658 | 0.06755 | 0.08364 | 0.04107 | 0.03121 | 0.12396 | 0.49843 |
| SLR24 | 0.20822 | -0.2316 | -0.2903 | -0.0978 | 0.09127 | 0.19454 | 2.222 | 0.06803 | 0.07554 | 0.05468 | 0.02152 | 0.25193 | 0.26641 |

| | | | | | | | | | | | | | |
|---|---|---|---|---|---|---|---|---|---|---|---|---|---|
| LOUNH91 | 0.20565 | -0.2711 | -0.2321 | -0.158 | 0.0904 | 0.19438 | 2.21979 | 0.06827 | 0.07815 | 0.02977 | 0.05426 | 0.13915 | 0.26838 |
| DB | -0.0497 | -0.195 | -0.1291 | -0.1704 | 0.28692 | 0.19406 | 2.21526 | 0.06876 | 0.36715 | 0.08966 | 0.18825 | 0.12084 | 0.02281 |
| SCABER | 0.0678 | -0.2546 | -0.3258 | -0.0588 | 0.08181 | 0.19379 | 2.21139 | 0.06918 | 0.32174 | 0.03878 | 0.01118 | 0.34416 | 0.28815 |
| EN | 0.04128 | -0.3196 | -0.1462 | -0.2704 | 0.00836 | 0.19305 | 2.20101 | 0.07032 | 0.38911 | 0.01259 | 0.15809 | 0.03009 | 0.47728 |
| SHMAC5 | 0.31926 | -0.2882 | -0.1292 | -0.1164 | 0.05582 | 0.19223 | 2.18936 | 0.07162 | 0.01268 | 0.02229 | 0.1881 | 0.21285 | 0.35163 |
| CHP134 | -0.0571 | -0.2839 | -0.076 | -0.3238 | -0.0178 | 0.19192 | 2.18501 | 0.07211 | 0.34845 | 0.02404 | 0.30195 | 0.0116 | 0.4517 |
| BHT101 | 0.24345 | -0.3306 | 0.04956 | -0.075 | 0.18304 | 0.19187 | 2.18431 | 0.07219 | 0.04593 | 0.01018 | 0.36762 | 0.30424 | 0.10404 |
| UMUC10 | 0.10481 | -0.2726 | -0.3245 | -0.0432 | -0.0479 | 0.19159 | 2.18042 | 0.07264 | 0.23678 | 0.02906 | 0.01145 | 0.38418 | 0.37193 |
| VMCUB1 | -0.0723 | -0.2489 | -0.3026 | -0.1393 | -0.0536 | 0.19091 | 2.17076 | 0.07375 | 0.31067 | 0.04229 | 0.0173 | 0.16997 | 0.35718 |
| NCIH1975 | 0.16238 | -0.1519 | -0.3516 | -0.1215 | -0.0296 | 0.1907 | 2.16784 | 0.07409 | 0.13248 | 0.14867 | 0.00661 | 0.20275 | 0.41993 |
| OVCAR8 | 0.21762 | -0.266 | -0.2478 | -0.1075 | 0.08813 | 0.19068 | 2.16752 | 0.07412 | 0.06653 | 0.03237 | 0.043 | 0.23118 | 0.27353 |
| HEC251 | 0.11444 | -0.3407 | 0.06814 | -0.306 | -0.0044 | 0.1905 | 2.16508 | 0.07441 | 0.21683 | 0.00829 | 0.32088 | 0.01624 | 0.48815 |
| IHH4 | 0.18431 | -0.1789 | -0.2016 | -0.212 | 0.20622 | 0.19041 | 2.16381 | 0.07456 | 0.10244 | 0.10936 | 0.08242 | 0.07178 | 0.07757 |
| M040416 | 0.18708 | -0.102 | -0.18 | -0.1979 | 0.28194 | 0.19037 | 2.16324 | 0.07463 | 0.09902 | 0.24277 | 0.10792 | 0.0864 | 0.02484 |
| ROS50 | 0.1814 | 0.18305 | 0.1118 | -0.154 | 0.33548 | 0.18972 | 2.15413 | 0.0757 | 0.10613 | 0.10402 | 0.22219 | 0.1454 | 0.00922 |
| MDAMB361 | 0.02882 | -0.1639 | -0.3893 | 0.06657 | -0.1009 | 0.1897 | 2.15377 | 0.07575 | 0.42207 | 0.13019 | 0.00285 | 0.32475 | 0.24516 |
| RH28 | 0.01742 | -0.0821 | -0.23 | -0.3046 | 0.15141 | 0.18948 | 2.15071 | 0.07611 | 0.45272 | 0.28744 | 0.05595 | 0.01667 | 0.14952 |
| HUCCT1 | 0.25779 | -0.1049 | -0.2996 | -0.148 | 0.05191 | 0.18936 | 2.14904 | 0.07631 | 0.03687 | 0.23667 | 0.01825 | 0.15503 | 0.36159 |
| COGN278 | 0.06226 | -0.1293 | -0.0366 | -0.1665 | 0.36602 | 0.18915 | 2.14607 | 0.07667 | 0.33541 | 0.18794 | 0.40134 | 0.12646 | 0.00485 |
| KPL1 | 0.3157 | -0.2183 | 0.03699 | -0.2201 | 0.16261 | 0.18878 | 2.14091 | 0.07729 | 0.01356 | 0.06593 | 0.40039 | 0.06429 | 0.13214 |
| L33 | -0.076 | -0.2538 | -0.2683 | -0.0991 | 0.11567 | 0.18875 | 2.14059 | 0.07733 | 0.30192 | 0.03922 | 0.03118 | 0.24905 | 0.21434 |
| ICC10 | 0.18331 | -0.2266 | -0.2237 | 0.08328 | 0.21394 | 0.18832 | 2.13447 | 0.07808 | 0.1037 | 0.05871 | 0.06118 | 0.28471 | 0.06996 |
| 22RV1 | 0.25172 | -0.354 | 0.0428 | -0.2045 | -0.0072 | 0.18799 | 2.12988 | 0.07864 | 0.04051 | 0.00629 | 0.38514 | 0.07933 | 0.4804 |
| PFSK1 | -0.2695 | -0.2664 | 0.04429 | -0.0539 | -0.2399 | 0.18788 | 2.12841 | 0.07883 | 0.03059 | 0.03212 | 0.38127 | 0.3565 | 0.04842 |
| UPCISCC026 | 0.15825 | -0.2546 | -0.2864 | -0.1118 | 0.06761 | 0.18747 | 2.12264 | 0.07954 | 0.13874 | 0.03876 | 0.02303 | 0.22214 | 0.32219 |
| IM9 | -0.0514 | -0.2169 | 0.03393 | -0.1794 | 0.28942 | 0.18746 | 2.12257 | 0.07955 | 0.36289 | 0.06719 | 0.40848 | 0.10873 | 0.02185 |
| CCSW1 | 0.0237 | -0.2995 | -0.302 | -0.0181 | 0.0066 | 0.18739 | 2.1216 | 0.07967 | 0.43581 | 0.01829 | 0.01746 | 0.45085 | 0.48205 |
| PK45H | 0.11306 | -0.1891 | -0.1693 | -0.0139 | 0.31809 | 0.18674 | 2.11255 | 0.08081 | 0.21963 | 0.09653 | 0.12243 | 0.46212 | 0.01296 |
| CALU1 | 0.06885 | -0.1749 | -0.3099 | -0.0213 | 0.21266 | 0.18639 | 2.10769 | 0.08143 | 0.31916 | 0.1147 | 0.01511 | 0.44228 | 0.07119 |
| NB17 | -0.0701 | -0.3205 | -0.0126 | 0.02195 | 0.21081 | 0.18623 | 2.10543 | 0.08172 | 0.31616 | 0.01238 | 0.46572 | 0.44051 | 0.07298 |
| ANGMCSS | 0.16224 | -0.1992 | -0.1587 | 0.11225 | 0.27995 | 0.18616 | 2.10445 | 0.08185 | 0.13269 | 0.08503 | 0.13802 | 0.22127 | 0.02569 |
| HEC1 | 0.23156 | -0.2143 | -0.1365 | -0.1984 | 0.20329 | 0.18613 | 2.10402 | 0.08191 | 0.05469 | 0.06963 | 0.17478 | 0.08587 | 0.08061 |
| MELHO | 0.14179 | -0.3165 | -0.0765 | -0.1302 | 0.20907 | 0.18607 | 2.1032 | 0.08201 | 0.16557 | 0.01336 | 0.30067 | 0.18624 | 0.07469 |
| H4 | -0.0073 | -0.3359 | 0.06207 | 0.09362 | 0.18248 | 0.18583 | 2.09984 | 0.08244 | 0.48018 | 0.00914 | 0.3359 | 0.26115 | 0.10475 |
| SG231 | 0.06091 | -0.4397 | 0.04739 | -0.0378 | -0.0275 | 0.18567 | 2.09769 | 0.08272 | 0.33879 | 0.00079 | 0.37322 | 0.39827 | 0.42557 |
| IMR32 | -0.0164 | -0.4045 | 0.06557 | -0.1208 | 0.05853 | 0.18546 | 2.09473 | 0.08311 | 0.4555 | 0.00197 | 0.32722 | 0.20424 | 0.34477 |
| NUGC3 | 0.20735 | -0.2332 | -0.2756 | -0.1329 | 0.03266 | 0.18538 | 2.09356 | 0.08326 | 0.07642 | 0.05341 | 0.02764 | 0.18124 | 0.41185 |
| NCIH727 | 0.15753 | -0.2904 | -0.129 | -0.1967 | 0.16357 | 0.1851 | 2.08977 | 0.08376 | 0.13985 | 0.02146 | 0.18853 | 0.08772 | 0.13072 |
| 647V | 0.15141 | -0.2238 | -0.312 | 0.11619 | 0.06302 | 0.18476 | 2.08502 | 0.08438 | 0.14952 | 0.06109 | 0.01453 | 0.2133 | 0.33353 |
| NCIH157DM | 0.06374 | -0.3421 | -0.1879 | -0.109 | 0.09903 | 0.18449 | 2.08122 | 0.08489 | 0.33173 | 0.00807 | 0.09808 | 0.22803 | 0.24922 |
| HUH7 | 0.35724 | -0.0979 | -0.0527 | -0.0818 | 0.24688 | 0.18444 | 2.08066 | 0.08496 | 0.00587 | 0.25177 | 0.35966 | 0.28822 | 0.04362 |
| TGW | -0.0352 | -0.3214 | 0.02901 | -0.2739 | 0.07993 | 0.18421 | 2.07739 | 0.0854 | 0.40508 | 0.01217 | 0.42157 | 0.02845 | 0.29256 |
| RPE1SS119 | 0.2246 | -0.2272 | -0.1192 | -0.2933 | 0.04414 | 0.18398 | 2.07424 | 0.08583 | 0.06039 | 0.05823 | 0.20729 | 0.02042 | 0.38166 |
| MFE296 | 0.0627 | 0.11192 | -0.3779 | 0.02339 | 0.19827 | 0.18381 | 2.07186 | 0.08615 | 0.33433 | 0.22195 | 0.00371 | 0.43662 | 0.08602 |
| SAS | 0.19403 | -0.234 | -0.2522 | -0.1623 | 0.08164 | 0.18364 | 2.0695 | 0.08647 | 0.09079 | 0.05279 | 0.04024 | 0.1326 | 0.28854 |
| S462 | 0.14242 | -0.2626 | -0.2369 | -0.148 | 0.12433 | 0.18322 | 2.06375 | 0.08725 | 0.16448 | 0.03416 | 0.05062 | 0.15507 | 0.19735 |
| TUHR4TKB | -0.1555 | -0.3015 | -0.2138 | -0.0047 | 0.03297 | 0.18318 | 2.06313 | 0.08733 | 0.14299 | 0.01763 | 0.07013 | 0.48714 | 0.41103 |
| TR146 | 0.24417 | -0.2943 | -0.1645 | -0.1282 | 0.09527 | 0.18283 | 2.05838 | 0.08799 | 0.04544 | 0.02007 | 0.12932 | 0.19003 | 0.25747 |
| KLE | -0.1706 | -0.1958 | -0.309 | -0.0515 | 0.01334 | 0.18228 | 2.05296 | 0.08874 | 0.12062 | 0.08883 | 0.01539 | 0.36263 | 0.46375 |
| MHHNB11 | 0.06314 | -0.1929 | 0.02095 | -0.184 | 0.31847 | 0.18224 | 2.05022 | 0.08912 | 0.33323 | 0.0921 | 0.44318 | 0.10285 | 0.01287 |
| LN235 | 0.02124 | -0.3043 | -0.0472 | 0.07654 | 0.2344 | 0.18163 | 2.04179 | 0.09031 | 0.44242 | 0.01677 | 0.37383 | 0.30058 | 0.05249 |
| SCMCRM2 | 0.09599 | -0.4144 | 0.0827 | -0.047 | 0.05267 | 0.18157 | 2.04103 | 0.09041 | 0.25587 | 0.00154 | 0.28607 | 0.3742 | 0.35965 |
| NCIH1299 | -0.0325 | -0.1757 | -0.173 | -0.2361 | 0.21026 | 0.18125 | 2.03666 | 0.09104 | 0.41219 | 0.11357 | 0.11724 | 0.05119 | 0.07351 |
| EGI1 | 0.1899 | -0.1124 | -0.2997 | -0.1421 | 0.15132 | 0.18109 | 2.03442 | 0.09135 | 0.09562 | 0.22098 | 0.0182 | 0.16495 | 0.14966 |
| PC3 | 0.23495 | -0.2659 | -0.0369 | 0.14715 | 0.17821 | 0.18058 | 2.02747 | 0.09236 | 0.05208 | 0.03239 | 0.40066 | 0.1565 | 0.11027 |
| T47D | 0.10422 | -0.2126 | -0.231 | -0.2352 | 0.10921 | 0.18042 | 2.0252 | 0.09268 | 0.23803 | 0.0712 | 0.05517 | 0.05189 | 0.22754 |
| SW982 | 0.01103 | -0.3568 | -0.1106 | -0.0263 | 0.14778 | 0.1804 | 2.02501 | 0.09271 | 0.47001 | 0.00592 | 0.22458 | 0.42883 | 0.15545 |
| TANOUE | 0.0726 | -0.147 | -0.0317 | -0.3362 | 0.20834 | 0.18031 | 2.02381 | 0.09289 | 0.31004 | 0.15671 | 0.41451 | 0.00909 | 0.07542 |
| HSC2 | 0.17479 | -0.2371 | -0.2881 | -0.1129 | 0.03628 | 0.18019 | 2.02208 | 0.09314 | 0.11483 | 0.05045 | 0.02237 | 0.21991 | 0.40228 |
| COV318 | 0.2916 | -0.1995 | 0.19288 | -0.209 | 0.11656 | 0.17969 | 2.01531 | 0.09413 | 0.02103 | 0.08463 | 0.09212 | 0.07473 | 0.21257 |
| ICC137 | -0.0419 | 0.01947 | -0.3957 | 0.14813 | 0.11746 | 0.1789 | 2.00446 | 0.09574 | 0.38748 | 0.44718 | 0.00244 | 0.15487 | 0.21076 |
| TGBC52TKB | 0.11852 | -0.0906 | -0.3234 | -0.1873 | -0.0927 | 0.17872 | 2.00205 | 0.09611 | 0.20866 | 0.26792 | 0.0117 | 0.09874 | 0.26323 |
| YUHOIN0650 | 0.12962 | -0.2591 | -0.2282 | -0.1431 | 0.13936 | 0.17821 | 1.99503 | 0.09717 | 0.18736 | 0.03612 | 0.0574 | 0.16335 | 0.16978 |
| PACADD188 | 0.01167 | -0.3162 | -0.2487 | -0.0893 | -0.124 | 0.17813 | 1.99402 | 0.09732 | 0.4683 | 0.01344 | 0.04244 | 0.27093 | 0.19794 |
| SUM185PE | 0.10631 | -0.2823 | -0.1058 | 0.06917 | 0.23945 | 0.17798 | 1.99189 | 0.09765 | 0.2336 | 0.02468 | 0.23461 | 0.31838 | 0.04875 |
| HEC50B | 0.27396 | -0.2562 | -0.0477 | -0.0989 | 0.20781 | 0.17792 | 1.99113 | 0.09776 | 0.0284 | 0.03782 | 0.37249 | 0.24947 | 0.07595 |
| EM2 | -0.2213 | -0.0697 | -0.2566 | -0.1474 | 0.1195 | 0.17772 | 1.98847 | 0.09817 | 0.06326 | 0.317 | 0.03756 | 0.15602 | 0.20673 |
| PECAPJ49 | 0.12682 | -0.2135 | -0.3177 | -0.1147 | 0.01523 | 0.17708 | 1.97976 | 0.09952 | 0.1926 | 0.0704 | 0.01306 | 0.21626 | 0.45865 |
| 2313287 | 0.33726 | -0.1951 | -0.0515 | -0.144 | 0.16605 | 0.1768 | 1.97584 | 0.10013 | 0.0089 | 0.08955 | 0.36258 | 0.16182 | 0.12708 |
| EFO27 | 0.22592 | -0.1955 | -0.2428 | -0.1846 | 0.04446 | 0.1765 | 1.97189 | 0.10075 | 0.05927 | 0.08912 | 0.0464 | 0.10207 | 0.38081 |
| TMK1 | -0.0828 | 0.02387 | -0.3107 | -0.1759 | -0.17 | 0.17648 | 1.97159 | 0.1008 | 0.2859 | 0.43533 | 0.01491 | 0.11334 | 0.12149 |
| JURKAT | -0.1414 | -0.202 | -0.1257 | -0.0469 | 0.25248 | 0.1764 | 1.97048 | 0.10097 | 0.1662 | 0.08199 | 0.19467 | 0.37458 | 0.04004 |
| EVSAT | 0.28858 | -0.1393 | -0.1559 | 0.09597 | 0.22246 | 0.17504 | 1.95199 | 0.10393 | 0.02217 | 0.16997 | 0.14238 | 0.25593 | 0.06223 |
| NO10 | 0.17125 | -0.2748 | -0.0941 | -0.0748 | 0.2368 | 0.17461 | 1.94622 | 0.10487 | 0.1197 | 0.028 | 0.26014 | 0.30483 | 0.05069 |
| NCIH2030 | 0.14797 | -0.3082 | -0.2351 | -0.0205 | 0.06525 | 0.17425 | 1.94137 | 0.10567 | 0.15513 | 0.01561 | 0.05199 | 0.44441 | 0.32799 |
| NCIH1694 | -0.0381 | -0.3345 | -0.2042 | -0.0067 | 0.05671 | 0.17315 | 1.92655 | 0.10814 | 0.39739 | 0.00941 | 0.07967 | 0.48172 | 0.34938 |
| PANC0403 | 0.14516 | -0.109 | -0.3292 | -0.1596 | 0.02931 | 0.17254 | 1.91842 | 0.10952 | 0.15982 | 0.22803 | 0.01046 | 0.13664 | 0.42077 |
| HSC3 | 0.2042 | -0.1339 | -0.2106 | -0.1895 | 0.19463 | 0.17253 | 1.9182 | 0.10956 | 0.07966 | 0.17942 | 0.07317 | 0.09605 | 0.0901 |
| BICR16 | 0.06328 | -0.2668 | -0.2886 | -0.1011 | 0.0145 | 0.1725 | 1.91783 | 0.10962 | 0.33288 | 0.03193 | 0.02218 | 0.2447 | 0.46063 |
| 21NT | 0.36401 | -0.0992 | -0.1761 | -0.0614 | 0.09065 | 0.17228 | 1.91487 | 0.11013 | 0.00507 | 0.24888 | 0.11308 | 0.3375 | 0.26781 |
| KML1 | -0.014 | -0.3246 | -0.121 | -0.1683 | 0.10678 | 0.17185 | 1.90905 | 0.11113 | 0.46203 | 0.01143 | 0.2037 | 0.12386 | 0.23261 |
| MAVER1 | 0.17867 | -0.3191 | 0.04299 | -0.0472 | 0.19876 | 0.17155 | 1.90512 | 0.11182 | 0.10967 | 0.01271 | 0.38465 | 0.37366 | 0.08549 |
| SNU475 | 0.04435 | -0.3192 | -0.0186 | 0.0547 | 0.20846 | 0.17091 | 1.89653 | 0.11333 | 0.38111 | 0.01268 | 0.44946 | 0.35446 | 0.0753 |
| UPCISCC131 | 0.13869 | -0.1717 | -0.2808 | -0.1324 | 0.15044 | 0.17081 | 1.89517 | 0.11356 | 0.17097 | 0.11906 | 0.02532 | 0.18233 | 0.15109 |
| LN18 | 0.03386 | -0.2186 | -0.1304 | -0.0947 | 0.28084 | 0.17076 | 1.8945 | 0.11368 | 0.40868 | 0.06562 | 0.18585 | 0.25872 | 0.02531 |
| NCIH747 | 0.12123 | -0.321 | -0.1086 | -0.1249 | 0.15984 | 0.17072 | 1.89396 | 0.11378 | 0.20333 | 0.01227 | 0.2289 | 0.19618 | 0.13631 |
| SNU16 | -0.0599 | 0.09936 | -0.2817 | 0.188 | 0.26287 | 0.17068 | 1.89344 | 0.11387 | 0.3414 | 0.2485 | 0.02494 | 0.09791 | 0.03402 |
| COV362 | 0.19502 | -0.1894 | -0.0972 | -0.0647 | 0.29562 | 0.17066 | 1.89311 | 0.11393 | 0.08966 | 0.09619 | 0.25317 | 0.3293 | 0.01959 |
| BICR22 | 0.1398 | -0.1851 | -0.3368 | -0.0074 | 0.05912 | 0.17056 | 1.89179 | 0.11416 | 0.16901 | 0.10142 | 0.00898 | 0.47999 | 0.34328 |
| CMK115 | 0.23553 | -0.2868 | 0.05087 | -0.2077 | 0.13003 | 0.17047 | 1.89067 | 0.11436 | 0.05164 | 0.02284 | 0.36426 | 0.07606 | 0.18659 |
| SEM | 0.1571 | -0.0656 | 0.02113 | -0.2503 | 0.30475 | 0.17036 | 1.88914 | 0.11464 | 0.14051 | 0.32719 | 0.44271 | 0.04141 | 0.01662 |
| LS1034 | -0.1381 | -0.2378 | -0.0217 | -0.1631 | 0.19869 | 0.17017 | 1.8866 | 0.11509 | 0.172 | 0.04994 | 0.44115 | 0.13144 | 0.08556 |
| SKMEL30 | 0.06946 | -0.2186 | 0.0954 | -0.3683 | -0.0725 | 0.17 | 1.88434 | 0.1155 | 0.31768 | 0.06561 | 0.25719 | 0.00461 | 0.31021 |
| NCIH82 | -0.0871 | -0.3186 | -0.1892 | -0.1179 | -0.0053 | 0.16991 | 1.88311 | 0.11572 | 0.27597 | 0.01283 | 0.09646 | 0.20998 | 0.48557 |
| PCI4B | 0.19639 | -0.255 | -0.2335 | -0.052 | 0.11708 | 0.16988 | 1.88279 | 0.11578 | 0.08812 | 0.0385 | 0.05316 | 0.36135 | 0.21152 |
| SW620 | 0.08576 | -0.0963 | -0.0094 | -0.2143 | 0.33089 | 0.16942 | 1.87656 | 0.1169 | 0.27897 | 0.25524 | 0.47433 | 0.06962 | 0.01011 |
| SNU840 | 0.14978 | -0.2307 | -0.2143 | -0.0889 | 0.1941 | 0.16912 | 1.87261 | 0.11763 | 0.15217 | 0.05539 | 0.06964 | 0.27179 | 0.09072 |
| PF382 | 0.07985 | -0.2662 | -0.0376 | -0.2077 | 0.21352 | 0.16903 | 1.87139 | 0.11785 | 0.29274 | 0.03224 | 0.39889 | 0.07604 | 0.07036 |
| UMUC9 | 0.03201 | -0.1239 | -0.3125 | -0.0231 | 0.21418 | 0.16896 | 1.87048 | 0.11802 | 0.41357 | 0.19824 | 0.01441 | 0.43736 | 0.06974 |
| BICR6 | 0.14015 | -0.2394 | -0.2372 | 0.04535 | 0.17373 | 0.16894 | 1.87018 | 0.11807 | 0.16841 | 0.04878 | 0.05043 | 0.37851 | 0.11627 |
| UPCISCC040 | 0.11786 | -0.2078 | -0.2889 | -0.1281 | 0.09088 | 0.16893 | 1.87004 | 0.1181 | 0.20996 | 0.07599 | 0.02203 | 0.19012 | 0.26729 |
| JHESOAD1 | 0.08551 | -0.2509 | -0.2781 | -0.1095 | 0.06637 | 0.16876 | 1.86774 | 0.11852 | 0.27955 | 0.041 | 0.0265 | 0.22687 | 0.32523 |
| SUDHL5 | 0.01495 | -0.0125 | 0.0157 | -0.062 | 0.40531 | 0.16864 | 1.86627 | 0.11879 | 0.45939 | 0.46594 | 0.45738 | 0.33618 | 0.00193 |
| KMRC20 | 0.12742 | -0.2694 | -0.2285 | -0.1055 | 0.12007 | 0.16836 | 1.86242 | 0.1195 | 0.19147 | 0.03063 | 0.05717 | 0.23527 | 0.20561 |
| MEL202 | 0.1493 | -0.2682 | -0.2131 | -0.0995 | 0.13699 | 0.16815 | 1.85973 | 0.12 | 0.15295 | 0.03119 | 0.07081 | 0.24809 | 0.17396 |
| KP3 | 0.1511 | -0.2046 | -0.2231 | -0.1929 | 0.13056 | 0.16806 | 1.85851 | 0.12023 | 0.15002 | 0.07922 | 0.06164 | 0.09206 | 0.18561 |
| EKVX | 0.06284 | -0.1665 | -0.2817 | -0.2117 | 0.05815 | 0.16788 | 1.85603 | 0.1207 | 0.33399 | 0.1264 | 0.02495 | 0.07212 | 0.34572 |
| P31FUJ | 0.1542 | -0.1336 | -0.0534 | -0.3061 | 0.20128 | 0.16769 | 1.85362 | 0.12115 | 0.14505 | 0.18009 | 0.35765 | 0.01621 | 0.08274 |
| U2OS | -0.2218 | 0.08777 | -0.3104 | 0.15791 | 0.11387 | 0.16766 | 1.85316 | 0.12124 | 0.06284 | 0.27437 | 0.01497 | 0.13927 | 0.21797 |
| BIN67 | 0.03104 | -0.2451 | -0.308 | -0.0366 | -0.1187 | 0.16762 | 1.85262 | 0.12134 | 0.41617 | 0.04484 | 0.01566 | 0.4014 | 0.20831 |
| SUM102PT | 0.1975 | -0.0281 | -0.2317 | -0.1667 | 0.22728 | 0.1674 | 1.84971 | 0.12189 | 0.08687 | 0.42392 | 0.05456 | 0.12613 | 0.05814 |
| PEA1 | 0.05965 | -0.2209 | -0.2302 | 0.04046 | 0.21866 | 0.16714 | 1.84632 | 0.12253 | 0.34195 | 0.06359 | 0.05581 | 0.39128 | 0.06559 |
| SNU761 | 0.20834 | -0.1363 | -0.2942 | -0.075 | 0.12724 | 0.16705 | 1.84508 | 0.12277 | 0.07542 | 0.17516 | 0.02009 | 0.30422 | 0.19181 |
| GCT | 0.20368 | -0.348 | -0.1036 | -0.1037 | 0.0419 | 0.167 | 1.84447 | 0.12289 | 0.0802 | 0.00713 | 0.23939 | 0.23921 | 0.3875 |
| P2URK562 | 0.2079 | -0.2093 | 0.02096 | -0.0812 | 0.2861 | 0.16691 | 1.84323 | 0.12312 | 0.07587 | 0.07447 | 0.44316 | 0.28951 | 0.02314 |
| CCLFPEDS0003T | 0.11901 | -0.3674 | 0.04831 | -0.1444 | 0.09732 | 0.1669 | 1.84307 | 0.12315 | 0.20769 | 0.00471 | 0.37083 | 0.16105 | 0.25296 |
| SKGT2 | 0.05259 | -0.2759 | -0.2051 | 0.0106 | 0.17686 | 0.16644 | 1.83699 | 0.12432 | 0.35983 | 0.02748 | 0.07874 | 0.47119 | 0.11206 |
| HUPT4 | 0.13515 | -0.0609 | -0.3546 | -0.1014 | -0.0419 | 0.16585 | 1.82918 | 0.12584 | 0.17725 | 0.33879 | 0.00621 | 0.24397 | 0.38753 |
| UMUC3 | 0.14279 | -0.1211 | -0.1927 | 0.06011 | 0.30146 | 0.16553 | 1.82493 | 0.12667 | 0.16385 | 0.20366 | 0.09233 | 0.34081 | 0.01765 |

| | | | | | | | | | | | | | |
|---|---|---|---|---|---|---|---|---|---|---|---|---|---|
| TE9 | 0.10447 | -0.2849 | -0.2197 | -0.0983 | 0.10966 | 0.16551 | 1.82466 | 0.12672 | 0.2375 | 0.02362 | 0.06462 | 0.25086 | 0.2266 |
| OCIAML3 | -0.0436 | -0.1878 | 0.14695 | -0.284 | 0.16886 | 0.16513 | 1.81962 | 0.12772 | 0.38313 | 0.09812 | 0.15683 | 0.02399 | 0.12305 |
| SKRC20 | 0.11814 | -0.2348 | -0.2661 | -0.1399 | 0.06 | 0.1644 | 1.81009 | 0.12962 | 0.20941 | 0.05217 | 0.03231 | 0.1689 | 0.34108 |
| RKO | 0.18466 | -0.3097 | -0.0797 | -0.2298 | 0.0027 | 0.16429 | 1.8086 | 0.12992 | 0.102 | 0.01517 | 0.293 | 0.05608 | 0.49265 |
| SKMES1 | 0.12744 | -0.2358 | -0.2534 | -0.1185 | 0.10961 | 0.16421 | 1.80758 | 0.13013 | 0.19143 | 0.0514 | 0.03948 | 0.20876 | 0.22672 |
| BC3C | 0.17401 | -0.1903 | -0.1564 | -0.0099 | 0.27053 | 0.16412 | 1.80631 | 0.13038 | 0.1159 | 0.0951 | 0.14165 | 0.47309 | 0.03005 |
| JHH1 | 0.17581 | -0.1281 | -0.3159 | -0.1069 | 0.06814 | 0.16365 | 1.8002 | 0.13162 | 0.11346 | 0.19024 | 0.01351 | 0.23229 | 0.32089 |
| WSUNHL | -0.001 | -0.1712 | -0.2083 | -0.2799 | 0.04462 | 0.163 | 1.79158 | 0.13339 | 0.49724 | 0.11972 | 0.07547 | 0.02573 | 0.3804 |
| HUH1 | 0.20399 | -0.2946 | -0.1193 | -0.1977 | 0.03345 | 0.16294 | 1.79086 | 0.13354 | 0.07987 | 0.01995 | 0.20712 | 0.08666 | 0.40976 |
| OS252 | -0.0042 | -0.234 | -0.3223 | 0.02825 | -0.0386 | 0.16287 | 1.78999 | 0.13372 | 0.48852 | 0.05282 | 0.01196 | 0.42361 | 0.39616 |
| DANG | -0.005 | -0.2331 | -0.212 | -0.2151 | 0.07296 | 0.16224 | 1.7817 | 0.13545 | 0.4863 | 0.05347 | 0.0718 | 0.06888 | 0.30916 |
| ICC4 | 0.1466 | -0.3058 | -0.1927 | -0.1271 | 0.03856 | 0.16213 | 1.78028 | 0.13574 | 0.15742 | 0.01629 | 0.0923 | 0.1921 | 0.39626 |
| GIMEN | -0.0944 | -0.3712 | -0.089 | -0.0322 | -0.0807 | 0.16141 | 1.77075 | 0.13776 | 0.25948 | 0.00432 | 0.27159 | 0.41295 | 0.29074 |
| H157 | 0.11747 | -0.2372 | -0.273 | 0.01984 | 0.11563 | 0.16135 | 1.77002 | 0.13792 | 0.21074 | 0.05037 | 0.02888 | 0.44617 | 0.21442 |
| UW228 | 0.12799 | -0.1815 | -0.2094 | 0.14117 | 0.21244 | 0.16108 | 1.7665 | 0.13867 | 0.1904 | 0.10602 | 0.07433 | 0.16664 | 0.07139 |
| KKU100 | 0.3246 | -0.1315 | -0.1602 | -0.0271 | 0.16236 | 0.1609 | 1.76409 | 0.13919 | 0.01144 | 0.18393 | 0.13579 | 0.42675 | 0.13252 |
| TYKNU | 0.08256 | -0.1905 | 0.20305 | -0.205 | 0.21709 | 0.16074 | 1.76209 | 0.13962 | 0.28639 | 0.09491 | 0.08086 | 0.07886 | 0.06702 |
| HEYA8 | 0.2034 | -0.0724 | -0.0221 | -0.2099 | 0.29212 | 0.16071 | 1.76159 | 0.13973 | 0.0805 | 0.31058 | 0.44013 | 0.0739 | 0.02084 |
| A549 | 0.05939 | -0.2411 | -0.2895 | -0.1088 | -0.0031 | 0.16052 | 1.75918 | 0.14025 | 0.34261 | 0.04759 | 0.0218 | 0.22836 | 0.49145 |
| KMS34 | -0.042 | -0.0094 | -0.2251 | -0.0602 | 0.31547 | 0.16042 | 1.75783 | 0.14054 | 0.38711 | 0.47434 | 0.05999 | 0.34065 | 0.01362 |
| HUPT3 | 0.10164 | -0.1685 | -0.3043 | -0.1556 | 0.0188 | 0.16026 | 1.75581 | 0.14098 | 0.24356 | 0.12359 | 0.01676 | 0.1428 | 0.44898 |
| DIFI | -0.0201 | -0.1472 | -0.2308 | -0.2546 | 0.07765 | 0.16024 | 1.75548 | 0.14105 | 0.44554 | 0.15641 | 0.05532 | 0.03875 | 0.29794 |
| VAESBJ | 0.136 | -0.3154 | 0.05835 | -0.0809 | 0.19026 | 0.16023 | 1.75544 | 0.14106 | 0.17573 | 0.01365 | 0.34522 | 0.2902 | 0.09519 |
| RI1 | 0.13138 | -0.2701 | 0.11584 | -0.1654 | 0.2024 | 0.16011 | 1.75376 | 0.14143 | 0.18411 | 0.03024 | 0.214 | 0.12805 | 0.08155 |
| NCIH1092 | 0.00199 | -0.1872 | -0.1799 | -0.154 | 0.22807 | 0.15993 | 1.75141 | 0.14194 | 0.49459 | 0.09888 | 0.10811 | 0.14539 | 0.0575 |
| MFM223 | 0.13694 | -0.0623 | -0.3002 | 0.05232 | 0.22436 | 0.1598 | 1.7498 | 0.14229 | 0.17405 | 0.33532 | 0.01805 | 0.36054 | 0.06059 |
| KMS20 | -0.0204 | -0.1789 | -0.1254 | -0.2137 | 0.22275 | 0.15969 | 1.7483 | 0.14262 | 0.44457 | 0.10932 | 0.19523 | 0.07022 | 0.06197 |
| KNS42 | 0.01917 | -0.1481 | -0.1273 | -0.0739 | 0.31846 | 0.15966 | 1.74799 | 0.14269 | 0.44798 | 0.15494 | 0.19174 | 0.30688 | 0.01287 |
| YSCCC | 0.12252 | -0.2347 | -0.2134 | -0.1896 | 0.09076 | 0.15953 | 1.74627 | 0.14307 | 0.20082 | 0.05226 | 0.0705 | 0.096 | 0.26757 |
| SUM229PE | 0.14582 | -0.1485 | -0.2625 | -0.1317 | 0.16728 | 0.15925 | 1.74255 | 0.1439 | 0.15871 | 0.15422 | 0.03422 | 0.18346 | 0.1253 |
| SCC9 | 0.15022 | -0.184 | -0.2792 | -0.1576 | -0.0413 | 0.15924 | 1.74253 | 0.1439 | 0.15145 | 0.10282 | 0.02601 | 0.13976 | 0.38906 |
| KP363T | 0.16113 | -0.2818 | -0.1321 | -0.1507 | 0.13964 | 0.15884 | 1.7373 | 0.14507 | 0.13435 | 0.0249 | 0.18275 | 0.15071 | 0.16931 |
| GCIY | 0.06731 | -0.338 | -0.1333 | -0.0272 | 0.117 | 0.15875 | 1.73612 | 0.14533 | 0.32294 | 0.00877 | 0.18054 | 0.42639 | 0.21169 |
| TT1TKB | 0.20159 | -0.2229 | -0.216 | -0.1776 | -0.001 | 0.15865 | 1.73477 | 0.14563 | 0.08242 | 0.06187 | 0.06806 | 0.11105 | 0.49721 |
| SKOV3 | -0.0204 | -0.1857 | -0.3198 | -0.1 | 0.02373 | 0.15853 | 1.73323 | 0.14598 | 0.44462 | 0.10073 | 0.01254 | 0.24715 | 0.43571 |
| BCPAP | 0.27011 | -0.2132 | -0.0849 | -0.1603 | 0.16071 | 0.15814 | 1.72824 | 0.14711 | 0.03026 | 0.07065 | 0.28087 | 0.13562 | 0.13499 |
| HEC1B | 0.029 | -0.1411 | -0.3256 | -0.1354 | 0.05432 | 0.15814 | 1.72815 | 0.14713 | 0.4216 | 0.16682 | 0.01122 | 0.17679 | 0.35544 |
| NCIH211 | -0.0046 | -0.3277 | -0.1111 | -0.0566 | 0.13616 | 0.15801 | 1.72649 | 0.1475 | 0.48747 | 0.01076 | 0.22354 | 0.34953 | 0.17544 |
| 786O | 0.21003 | -0.1713 | -0.1594 | -0.0886 | 0.23988 | 0.15777 | 1.72339 | 0.14821 | 0.07375 | 0.11964 | 0.13701 | 0.27237 | 0.04844 |
| CAPAN1 | 0.14156 | -0.1183 | -0.306 | -0.1232 | 0.10933 | 0.15748 | 1.71967 | 0.14906 | 0.16597 | 0.209 | 0.01626 | 0.19951 | 0.2273 |
| KIJK | 0.1683 | -0.244 | 0.02721 | -0.2854 | 0.10763 | 0.15744 | 1.71904 | 0.14921 | 0.12384 | 0.04556 | 0.42637 | 0.02342 | 0.23083 |
| CORL47 | -0.1036 | -0.2859 | -0.17 | -0.1479 | -0.1314 | 0.15739 | 1.71841 | 0.14935 | 0.23938 | 0.02321 | 0.12138 | 0.1553 | 0.18415 |
| U937 | -0.0781 | -0.1466 | -0.2458 | -0.1666 | 0.14456 | 0.15686 | 1.71162 | 0.15092 | 0.29694 | 0.15735 | 0.04437 | 0.12624 | 0.16084 |
| HT1376 | -0.0362 | -0.17 | -0.3238 | -0.0102 | 0.10006 | 0.15674 | 1.71007 | 0.15128 | 0.40238 | 0.12147 | 0.01163 | 0.47217 | 0.24696 |
| NCIH1568 | 0.03926 | -0.235 | -0.1281 | -0.1534 | 0.21808 | 0.15664 | 1.70881 | 0.15158 | 0.39443 | 0.05204 | 0.19027 | 0.14634 | 0.06611 |
| ICC15 | 0.16918 | -0.1879 | -0.297 | -0.0836 | -0.0209 | 0.15664 | 1.70879 | 0.15158 | 0.1226 | 0.09796 | 0.01912 | 0.28392 | 0.44327 |
| NCIH322 | 0.12134 | -0.307 | -0.1927 | -0.1319 | 0.01234 | 0.15653 | 1.70732 | 0.15193 | 0.20311 | 0.01594 | 0.09235 | 0.18314 | 0.46648 |
| SAT | 0.14557 | -0.2108 | -0.2757 | -0.1254 | 0.02173 | 0.15631 | 1.70452 | 0.15258 | 0.15914 | 0.07302 | 0.02759 | 0.19535 | 0.4411 |
| HEL9217 | 0.01631 | -0.164 | -0.132 | -0.2926 | 0.1363 | 0.15627 | 1.70398 | 0.15271 | 0.4557 | 0.13013 | 0.18295 | 0.02067 | 0.17519 |
| IPC298 | 0.04414 | -0.025 | -0.3807 | 0.07151 | 0.09598 | 0.15615 | 1.70245 | 0.15307 | 0.38165 | 0.43222 | 0.00348 | 0.31268 | 0.25589 |
| EW7 | 0.11186 | -0.2537 | -0.1286 | -0.1684 | 0.181 | 0.15593 | 1.69957 | 0.15375 | 0.22206 | 0.03931 | 0.18929 | 0.1237 | 0.10664 |
| PC14 | 0.16998 | -0.155 | -0.2783 | -0.1519 | 0.05308 | 0.15592 | 1.69943 | 0.15378 | 0.12148 | 0.14384 | 0.02643 | 0.14875 | 0.35861 |
| SW756 | 0.19811 | -0.1299 | -0.1136 | -0.1623 | 0.2631 | 0.15571 | 1.6967 | 0.15443 | 0.0862 | 0.1868 | 0.21847 | 0.13263 | 0.03389 |
| HT144SKINFV3 | 0.14495 | -0.3277 | 0.06275 | -0.1684 | 0.11483 | 0.15544 | 1.69325 | 0.15525 | 0.16018 | 0.01077 | 0.33421 | 0.12366 | 0.21604 |
| SNU1544 | 0.19687 | -0.2366 | -0.1867 | -0.0992 | 0.14764 | 0.15542 | 1.693 | 0.15531 | 0.08758 | 0.05086 | 0.09954 | 0.2489 | 0.15568 |
| OCUBM | 0.12424 | -0.1112 | -0.3506 | 0.0835 | -0.0432 | 0.1554 | 1.69276 | 0.15536 | 0.19752 | 0.22343 | 0.00675 | 0.28421 | 0.38401 |
| NALM6 | -0.0278 | -0.2343 | -0.2407 | -0.1492 | 0.07233 | 0.15516 | 1.68966 | 0.15611 | 0.42489 | 0.0526 | 0.04788 | 0.15315 | 0.3107 |
| SF268 | 0.04762 | -0.195 | -0.2447 | -0.2141 | 0.04546 | 0.15502 | 1.68779 | 0.15656 | 0.37261 | 0.08965 | 0.04511 | 0.06977 | 0.37822 |
| CA922 | 0.12434 | -0.2164 | -0.0894 | -0.2834 | 0.11314 | 0.15468 | 1.6835 | 0.15759 | 0.19733 | 0.06767 | 0.27053 | 0.02425 | 0.21945 |
| SKPNDW | -0.1719 | -0.0537 | -0.1813 | -0.2715 | -0.0177 | 0.15454 | 1.68161 | 0.15805 | 0.11877 | 0.35694 | 0.10629 | 0.02956 | 0.45209 |
| MDAMB453 | 0.29784 | -0.1414 | -0.1939 | -0.0867 | 0.11027 | 0.15395 | 1.67407 | 0.15989 | 0.01883 | 0.16629 | 0.091 | 0.27685 | 0.22535 |
| LS123 | 0.0136 | -0.0381 | -0.3734 | 0.04167 | -0.1137 | 0.15372 | 1.67116 | 0.16061 | 0.46304 | 0.39738 | 0.00411 | 0.38811 | 0.21829 |
| ECC4 | -0.0571 | -0.2086 | -0.0886 | -0.1283 | 0.24385 | 0.15358 | 1.66932 | 0.16106 | 0.34836 | 0.0752 | 0.27239 | 0.18986 | 0.04566 |
| NB10 | 0.118 | -0.2624 | 0.01843 | -0.2568 | 0.13823 | 0.1535 | 1.66831 | 0.16131 | 0.20969 | 0.03428 | 0.45 | 0.03745 | 0.17176 |
| L428 | 0.16952 | -0.1807 | 0.09037 | -0.3396 | 0.01141 | 0.15348 | 1.66799 | 0.16139 | 0.12212 | 0.10708 | 0.26844 | 0.00848 | 0.46898 |
| MKN74 | 0.19081 | -0.2271 | -0.1923 | -0.194 | 0.02306 | 0.15295 | 1.66123 | 0.16307 | 0.09454 | 0.05832 | 0.09279 | 0.09088 | 0.4375 |
| D502MG | 0.12066 | -0.3117 | -0.1778 | -0.0694 | 0.07905 | 0.15289 | 1.66047 | 0.16326 | 0.20445 | 0.01461 | 0.1108 | 0.31779 | 0.29463 |
| C10 | 0.1178 | -0.176 | -0.0132 | 0.05295 | 0.31208 | 0.15288 | 1.66031 | 0.1633 | 0.21008 | 0.11323 | 0.46419 | 0.35893 | 0.01452 |
| JHU029 | 0.15904 | -0.1786 | -0.2854 | -0.104 | 0.06246 | 0.15284 | 1.65977 | 0.16344 | 0.13753 | 0.10972 | 0.02341 | 0.23859 | 0.33493 |
| COLO201 | -0.0651 | -0.2712 | -0.1089 | -0.1571 | 0.1383 | 0.1526 | 1.6568 | 0.16418 | 0.32835 | 0.02972 | 0.2282 | 0.14045 | 0.17164 |
| WM793 | 0.15346 | -0.2305 | -0.1104 | -0.2131 | 0.15795 | 0.15226 | 1.65235 | 0.1653 | 0.14624 | 0.05552 | 0.22504 | 0.07074 | 0.13919 |
| BPH1 | 0.04117 | -0.1973 | -0.3297 | -0.0013 | 0.00205 | 0.1521 | 1.65033 | 0.16582 | 0.38941 | 0.08714 | 0.01036 | 0.49642 | 0.49442 |
| T3M5 | -0.0196 | -0.2796 | -0.0991 | -0.2407 | 0.04822 | 0.15194 | 1.64825 | 0.16635 | 0.44693 | 0.02583 | 0.24903 | 0.04785 | 0.37106 |
| NCIH2170 | 0.06187 | -0.2326 | -0.2526 | -0.1458 | 0.05636 | 0.15167 | 1.64487 | 0.16721 | 0.3364 | 0.05391 | 0.03996 | 0.15872 | 0.35024 |
| SNU2535 | 0.22263 | -0.2563 | 0.1342 | -0.2028 | 0.09829 | 0.15128 | 1.63986 | 0.1685 | 0.06208 | 0.03771 | 0.17896 | 0.08116 | 0.25082 |
| MORCPR | -0.231 | -0.2015 | -0.1063 | -0.1541 | 0.03772 | 0.15106 | 1.637 | 0.16924 | 0.05517 | 0.08247 | 0.23373 | 0.14514 | 0.39848 |
| A2058 | 0.15199 | -0.2444 | -0.2067 | -0.0571 | 0.14851 | 0.15102 | 1.63653 | 0.16936 | 0.14859 | 0.0453 | 0.07709 | 0.34826 | 0.15424 |
| A2780 | 0.11482 | -0.3036 | 0.0517 | -0.0902 | 0.1909 | 0.15099 | 1.63614 | 0.16946 | 0.21605 | 0.01698 | 0.36213 | 0.26892 | 0.09444 |
| C8166 | 0.10871 | -0.2141 | -0.0493 | -0.1689 | 0.24805 | 0.15073 | 1.63288 | 0.17031 | 0.22857 | 0.06977 | 0.36818 | 0.12293 | 0.04286 |
| MM160113 | -0.0765 | -0.2331 | -0.1996 | -0.1264 | 0.11922 | 0.15072 | 1.63276 | 0.17034 | 0.30061 | 0.05347 | 0.08461 | 0.19337 | 0.20727 |
| BB30HNC | 0.21019 | -0.1315 | -0.2126 | -0.2001 | 0.09472 | 0.15068 | 1.63213 | 0.1705 | 0.07359 | 0.18385 | 0.07127 | 0.08403 | 0.25869 |
| NB7 | -0.0598 | -0.2828 | -0.0133 | -0.1749 | 0.14676 | 0.15049 | 1.6298 | 0.17111 | 0.34152 | 0.02449 | 0.46395 | 0.11468 | 0.15715 |
| CASKI | 0.25449 | -0.0982 | -0.1382 | -0.2452 | 0.08539 | 0.15026 | 1.62683 | 0.17189 | 0.03881 | 0.25096 | 0.17175 | 0.04476 | 0.27983 |
| LAN2 | 0.22653 | -0.0858 | 0.0129 | -0.2201 | 0.2494 | 0.15026 | 1.62676 | 0.17191 | 0.05876 | 0.2788 | 0.46496 | 0.06433 | 0.04198 |
| SUDHL8 | 0.01254 | -0.3085 | 0.05703 | -0.0757 | 0.18345 | 0.15013 | 1.62518 | 0.17232 | 0.46592 | 0.01552 | 0.34857 | 0.30262 | 0.10352 |
| SNU668 | -0.0138 | -0.3189 | -0.1456 | 0.02416 | 0.10194 | 0.15001 | 1.62362 | 0.17273 | 0.46258 | 0.01277 | 0.15905 | 0.43456 | 0.2429 |
| KMBC2 | 0.24792 | -0.199 | -0.2368 | -0.0238 | 0.0338 | 0.14981 | 1.62116 | 0.17338 | 0.04294 | 0.08519 | 0.05068 | 0.43549 | 0.40884 |
| MON | 0.10005 | -0.3365 | 0.08433 | -0.0479 | 0.1376 | 0.14966 | 1.61924 | 0.17389 | 0.24698 | 0.00905 | 0.28228 | 0.37183 | 0.17289 |
| WAOSEL | 0.22002 | -0.1475 | 0.04681 | -0.2105 | 0.23075 | 0.14948 | 1.61691 | 0.17451 | 0.06437 | 0.15584 | 0.37471 | 0.07328 | 0.05534 |
| SKMG1 | 0.28123 | -0.2335 | 0.00451 | -0.0782 | 0.17088 | 0.14926 | 1.61414 | 0.17525 | 0.02514 | 0.05317 | 0.48773 | 0.29664 | 0.1202 |
| RPMI7951 | -0.0004 | -0.0815 | -0.3235 | -0.0012 | 0.18057 | 0.1492 | 1.61338 | 0.17546 | 0.49894 | 0.28895 | 0.01169 | 0.49684 | 0.10719 |
| SKGIIIA | 0.17077 | -0.2572 | -0.0944 | -0.2418 | 0.05365 | 0.14907 | 1.61169 | 0.17591 | 0.12036 | 0.03722 | 0.25936 | 0.04706 | 0.35715 |
| NAMALWA | 0.2977 | -0.1504 | -0.027 | -0.1468 | 0.19238 | 0.14901 | 1.61088 | 0.17613 | 0.01888 | 0.15112 | 0.42704 | 0.15711 | 0.0927 |
| G415 | 0.1854 | -0.2967 | -0.0527 | -0.1526 | 0.12239 | 0.14886 | 1.60898 | 0.17664 | 0.10108 | 0.0192 | 0.35949 | 0.14757 | 0.20109 |
| EOL1 | 0.00986 | -0.2142 | -0.1679 | -0.1251 | 0.20779 | 0.14883 | 1.6087 | 0.17671 | 0.4732 | 0.06972 | 0.12442 | 0.19595 | 0.07598 |
| EBC1 | -0.0567 | -0.178 | 0.07359 | -0.1116 | 0.27862 | 0.14877 | 1.60794 | 0.17692 | 0.34937 | 0.11049 | 0.30766 | 0.22271 | 0.02628 |
| ICC8 | 0.20802 | -0.2297 | -0.0617 | 0.03253 | 0.22462 | 0.14856 | 1.60517 | 0.17767 | 0.07575 | 0.05616 | 0.33673 | 0.4122 | 0.06037 |
| NB1643 | 0.07152 | -0.1114 | 0.04506 | -0.1321 | 0.33292 | 0.14844 | 1.60364 | 0.17808 | 0.31265 | 0.22299 | 0.37926 | 0.18287 | 0.00971 |
| NCIH1581 | 0.17256 | -0.076 | -0.2969 | -0.0335 | 0.16284 | 0.1481 | 1.59945 | 0.17922 | 0.11788 | 0.30195 | 0.01914 | 0.40968 | 0.1318 |
| YD8 | 0.02624 | -0.1273 | -0.3508 | 0.05861 | -0.088 | 0.14798 | 1.59792 | 0.17964 | 0.42898 | 0.19176 | 0.00673 | 0.34457 | 0.27392 |
| PEO4 | -0.0822 | -0.3149 | -0.1746 | -0.0653 | -0.0684 | 0.1478 | 1.59564 | 0.18027 | 0.2872 | 0.01376 | 0.11512 | 0.3278 | 0.32034 |
| KNS60 | 0.06294 | -0.1502 | -0.2937 | 0.10304 | 0.14941 | 0.14777 | 1.59526 | 0.18037 | 0.33374 | 0.15155 | 0.02026 | 0.24056 | 0.15277 |
| HOP62 | 0.00994 | -0.3546 | -0.1397 | -0.0553 | -0.0198 | 0.14772 | 1.59457 | 0.18056 | 0.47298 | 0.00621 | 0.16924 | 0.35289 | 0.44623 |
| KYSE70 | -0.0273 | -0.1012 | -0.0773 | -0.1691 | 0.29875 | 0.1477 | 1.59427 | 0.18064 | 0.42622 | 0.2444 | 0.29868 | 0.12267 | 0.01853 |
| CH157MN | 0.15503 | -0.0314 | -0.1888 | -0.1696 | 0.25058 | 0.14767 | 1.59394 | 0.18073 | 0.14375 | 0.41509 | 0.09696 | 0.12199 | 0.04123 |
| COLO678 | 0.16553 | -0.099 | -0.1423 | -0.2665 | 0.15454 | 0.1474 | 1.59048 | 0.18169 | 0.12784 | 0.24923 | 0.16468 | 0.03211 | 0.14452 |
| CAL78 | 0.03758 | -0.1169 | -0.3273 | -0.1059 | 0.07155 | 0.14694 | 1.58466 | 0.18331 | 0.39883 | 0.21183 | 0.01084 | 0.23438 | 0.31259 |
| HCC1395 | 0.20861 | -0.2602 | 0.00128 | -0.1249 | 0.18564 | 0.14692 | 1.58439 | 0.18338 | 0.07515 | 0.03549 | 0.49651 | 0.19618 | 0.10079 |
| HCC1428 | 0.11138 | -0.1786 | -0.2119 | -0.0392 | 0.22567 | 0.1466 | 1.58038 | 0.1845 | 0.22305 | 0.10978 | 0.07192 | 0.39457 | 0.05949 |
| NCIH460 | 0.18839 | -0.2594 | -0.0209 | -0.166 | 0.17221 | 0.14648 | 1.57888 | 0.18492 | 0.09743 | 0.03596 | 0.44335 | 0.12718 | 0.11837 |
| TE4 | -0.0553 | -0.2235 | -0.2589 | -0.1217 | 0.00829 | 0.1462 | 1.57532 | 0.18593 | 0.35299 | 0.06129 | 0.03623 | 0.20234 | 0.47745 |
| NCIH209 | 0.07149 | -0.249 | -0.1243 | -0.216 | 0.12298 | 0.14567 | 1.56872 | 0.1878 | 0.31273 | 0.04223 | 0.19731 | 0.068 | 0.19993 |
| SKNDZ | 0.1337 | -0.1565 | -0.0402 | -0.1435 | 0.28945 | 0.14561 | 1.56789 | 0.18804 | 0.17987 | 0.14138 | 0.39197 | 0.16256 | 0.02183 |
| OCIM2 | -0.0608 | -0.1712 | -0.1744 | -0.2694 | 0.00063 | 0.14557 | 1.56743 | 0.18817 | 0.33908 | 0.11972 | 0.11542 | 0.03059 | 0.49828 |
| MAPACHS77 | 0.05787 | -0.2145 | -0.2562 | -0.1508 | 0.04719 | 0.14556 | 1.56726 | 0.18822 | 0.34644 | 0.06947 | 0.0378 | 0.15054 | 0.37373 |
| GB2 | 0.11219 | -0.2139 | -0.2128 | -0.1942 | 0.06041 | 0.14475 | 1.55707 | 0.19116 | 0.22139 | 0.06997 | 0.07101 | 0.09062 | 0.34004 |
| JOPACA1 | 0.05224 | -0.0578 | -0.3599 | 0.00983 | 0.09405 | 0.14471 | 1.55657 | 0.1913 | 0.36073 | 0.3467 | 0.00554 | 0.47327 | 0.26019 |
| NCIH2452 | 0.03452 | -0.0976 | -0.0656 | 0.04194 | 0.3458 | 0.14467 | 1.55606 | 0.19145 | 0.40691 | 0.25243 | 0.3272 | 0.38738 | 0.00747 |
| SNU503 | 0.14475 | -0.1923 | -0.216 | -0.193 | 0.06851 | 0.14452 | 1.5542 | 0.19199 | 0.16051 | 0.09276 | 0.06799 | 0.09201 | 0.31998 |

| | | | | | | | | | | | | | |
|---|---|---|---|---|---|---|---|---|---|---|---|---|---|
| ES8 | 0.0772 | -0.0893 | 0.03799 | -0.0994 | 0.34469 | 0.14392 | 1.54668 | 0.19419 | 0.29902 | 0.27093 | 0.39775 | 0.2483 | 0.00765 |
| MM386 | 0.0721 | -0.0079 | -0.2372 | -0.2059 | 0.1855 | 0.14319 | 1.53756 | 0.19689 | 0.31125 | 0.47863 | 0.0504 | 0.07785 | 0.10096 |
| JJN3 | 0.22063 | -0.2896 | -0.0719 | -0.1438 | 0.06732 | 0.14304 | 1.53558 | 0.19748 | 0.06383 | 0.02178 | 0.31169 | 0.16221 | 0.32291 |
| RCC10RGB | -0.0041 | -0.176 | -0.2218 | -0.2358 | -0.0233 | 0.143 | 1.53508 | 0.19763 | 0.48882 | 0.11323 | 0.06281 | 0.05144 | 0.43675 |
| LS180 | 0.21328 | -0.2034 | -0.1076 | -0.1972 | 0.13753 | 0.14298 | 1.53493 | 0.19768 | 0.07059 | 0.08047 | 0.23079 | 0.08722 | 0.173 |
| TOV112D | 0.08091 | -0.2717 | -0.1049 | -0.0393 | 0.20078 | 0.14298 | 1.53484 | 0.1977 | 0.29025 | 0.02946 | 0.2365 | 0.39426 | 0.08328 |
| HSC1 | 0.15246 | -0.1981 | -0.2404 | 0.16566 | 0.00607 | 0.14291 | 1.53403 | 0.19795 | 0.14783 | 0.08616 | 0.04805 | 0.12765 | 0.4835 |
| HT144SKINFV1 | 0.12564 | -0.2592 | 0.06442 | -0.1998 | 0.16928 | 0.14267 | 1.53105 | 0.19884 | 0.19485 | 0.03608 | 0.33006 | 0.08437 | 0.12246 |
| NOZ | 0.042 | -0.3383 | -0.0793 | -0.1594 | 0.02273 | 0.14259 | 1.53003 | 0.19915 | 0.38722 | 0.00871 | 0.29403 | 0.13704 | 0.43841 |
| CAKI1 | 0.09087 | -0.197 | -0.2199 | -0.1997 | 0.06642 | 0.14231 | 1.52645 | 0.20023 | 0.26731 | 0.08744 | 0.06449 | 0.08446 | 0.32513 |
| CL11 | 0.2637 | -0.1497 | -0.2038 | -0.026 | 0.13149 | 0.14209 | 1.52376 | 0.20105 | 0.03357 | 0.15225 | 0.08011 | 0.42954 | 0.1839 |
| WPE1NA22 | 0.16301 | -0.2042 | -0.1488 | -0.2467 | 0.02143 | 0.14169 | 1.51869 | 0.20259 | 0.13154 | 0.07968 | 0.1537 | 0.04372 | 0.4419 |
| SUDHL1 | 0.12535 | -0.1906 | -0.0999 | -0.2386 | 0.16474 | 0.14162 | 1.51786 | 0.20285 | 0.1954 | 0.0948 | 0.24731 | 0.04937 | 0.129 |
| H103 | 0.13566 | -0.1715 | -0.2761 | -0.0561 | 0.11518 | 0.14157 | 1.51719 | 0.20305 | 0.17633 | 0.1194 | 0.02741 | 0.35096 | 0.21533 |
| UPCISCC152 | -0.038 | -0.2828 | -0.2232 | -0.065 | -0.0249 | 0.14152 | 1.51665 | 0.20322 | 0.39773 | 0.0245 | 0.06155 | 0.32864 | 0.43267 |
| HS852T | 0.07948 | -0.0256 | -0.179 | 0.12174 | 0.3064 | 0.14087 | 1.50851 | 0.20573 | 0.2936 | 0.4308 | 0.10919 | 0.20235 | 0.01613 |
| SHMAC4 | 0.10839 | -0.2056 | -0.2499 | -0.1554 | -0.0027 | 0.14014 | 1.49938 | 0.20858 | 0.22924 | 0.07818 | 0.04166 | 0.14324 | 0.49269 |
| OCUM1 | -0.0455 | -0.3393 | -0.0443 | 0.11949 | -0.0886 | 0.13991 | 1.49651 | 0.20949 | 0.37817 | 0.00854 | 0.38124 | 0.20674 | 0.27257 |
| PCI38 | 0.09027 | -0.185 | -0.2981 | 0.09575 | 0.02905 | 0.1397 | 1.49391 | 0.21031 | 0.26866 | 0.10153 | 0.01873 | 0.25642 | 0.42146 |
| YD15 | 0.22021 | -0.2474 | -0.1975 | -0.051 | -0.0005 | 0.13965 | 1.49328 | 0.21051 | 0.0642 | 0.04326 | 0.08684 | 0.36396 | 0.49851 |
| PANC0203 | 0.154 | -0.2477 | -0.1615 | -0.1432 | 0.11278 | 0.13963 | 1.49302 | 0.21059 | 0.14538 | 0.04306 | 0.13383 | 0.16317 | 0.2202 |
| SW403 | 0.06309 | -0.1807 | -0.031 | -0.3361 | 0.01744 | 0.13961 | 1.49278 | 0.21067 | 0.33336 | 0.10702 | 0.41618 | 0.00911 | 0.45266 |
| HCC515 | 0.11608 | -0.05 | -0.1059 | -0.2548 | 0.22493 | 0.13942 | 1.49046 | 0.2114 | 0.21352 | 0.36651 | 0.23447 | 0.03865 | 0.06011 |
| SNU1079 | 0.18766 | -0.2273 | -0.215 | -0.109 | 0.03677 | 0.13901 | 1.48533 | 0.21304 | 0.09832 | 0.05812 | 0.06895 | 0.22802 | 0.40098 |
| CORL23 | 0.07016 | -0.1789 | 0.27823 | -0.1897 | 0.09336 | 0.13878 | 1.48251 | 0.21395 | 0.31596 | 0.10942 | 0.02645 | 0.09582 | 0.26173 |
| BHY | 0.13569 | -0.1707 | -0.2857 | -0.0967 | -0.026 | 0.13841 | 1.47796 | 0.21542 | 0.17628 | 0.12042 | 0.0233 | 0.25422 | 0.42969 |
| ICC12 | -0.1195 | -0.2558 | -0.1152 | 0.03599 | 0.13676 | 0.13833 | 1.47694 | 0.21575 | 0.20669 | 0.03805 | 0.21526 | 0.40304 | 0.17438 |
| NCIH2009 | 0.23458 | -0.2045 | 0.03914 | -0.234 | 0.10732 | 0.13789 | 1.47151 | 0.21751 | 0.05236 | 0.07931 | 0.39472 | 0.05278 | 0.23149 |
| FARAGE | 0.1854 | -0.2044 | -0.1202 | -0.2132 | 0.11129 | 0.13762 | 1.46814 | 0.21862 | 0.10108 | 0.07945 | 0.20537 | 0.07069 | 0.22324 |
| COLO792 | 0.10478 | -0.2881 | -0.0257 | -0.2027 | 0.10696 | 0.13736 | 1.46494 | 0.21967 | 0.23685 | 0.02234 | 0.4303 | 0.08121 | 0.23223 |
| F5 | 0.22624 | -0.2194 | -0.0083 | -0.2035 | 0.13345 | 0.13727 | 1.46385 | 0.22003 | 0.05901 | 0.06495 | 0.4774 | 0.08036 | 0.18032 |
| Y79 | 0.13669 | -0.1975 | -0.0148 | -0.269 | 0.1421 | 0.13724 | 1.46348 | 0.22015 | 0.1745 | 0.08689 | 0.45976 | 0.0308 | 0.16503 |
| ST486 | 0.12873 | -0.2525 | 0.02428 | -0.2147 | 0.15192 | 0.13721 | 1.46304 | 0.2203 | 0.18902 | 0.04002 | 0.43425 | 0.06928 | 0.1487 |
| UMUC1 | -0.1464 | -0.1403 | -0.1649 | 0.0814 | 0.20561 | 0.1372 | 1.46297 | 0.22032 | 0.15769 | 0.16816 | 0.12871 | 0.28911 | 0.07819 |
| HS578T | 0.17783 | -0.1833 | -0.0169 | -0.0471 | 0.27087 | 0.1371 | 1.46173 | 0.22073 | 0.11077 | 0.10367 | 0.45414 | 0.37407 | 0.02988 |
| RKN | -0.0837 | -0.2215 | -0.1347 | -0.1081 | 0.16916 | 0.13695 | 1.4599 | 0.22134 | 0.28365 | 0.06307 | 0.17806 | 0.22981 | 0.12263 |
| LXF289 | -0.09 | -0.1533 | -0.2869 | 0.04485 | 0.10071 | 0.13691 | 1.45941 | 0.2215 | 0.26938 | 0.14656 | 0.0228 | 0.3798 | 0.24556 |
| NCIH1944 | 0.00756 | -0.3122 | -0.1369 | -0.1163 | 0.04318 | 0.13616 | 1.4501 | 0.22461 | 0.47945 | 0.01449 | 0.17414 | 0.21304 | 0.38416 |
| 170MGBA | 0.19619 | -0.1268 | 0.02799 | 0.07663 | 0.27732 | 0.13562 | 1.44349 | 0.22684 | 0.08834 | 0.1927 | 0.4243 | 0.30036 | 0.02686 |
| EMTOKA | 0.05918 | -0.2921 | -0.2085 | -0.0737 | -0.0469 | 0.13553 | 1.44241 | 0.22721 | 0.34314 | 0.02084 | 0.07523 | 0.30729 | 0.37443 |
| ICC108 | 0.16989 | -0.154 | -0.2745 | -0.0794 | 0.05389 | 0.13548 | 1.44177 | 0.22743 | 0.1216 | 0.14543 | 0.02816 | 0.29373 | 0.35653 |
| MM426 | 0.19966 | -0.1627 | -0.2108 | -0.0978 | 0.14019 | 0.13505 | 1.43645 | 0.22925 | 0.0845 | 0.13196 | 0.07297 | 0.25192 | 0.16834 |
| NCIH1048 | 0.08139 | -0.0869 | -0.2297 | 0.14884 | 0.21954 | 0.13466 | 1.43169 | 0.23088 | 0.28912 | 0.27641 | 0.05617 | 0.1537 | 0.0648 |
| S117 | 0.02542 | -0.3267 | 0.03869 | -0.053 | 0.12585 | 0.13464 | 1.43145 | 0.23096 | 0.43117 | 0.01097 | 0.39592 | 0.35869 | 0.19444 |
| SET2 | 0.01617 | -0.0027 | -0.1595 | -0.3094 | 0.05875 | 0.13425 | 1.42659 | 0.23265 | 0.4561 | 0.49255 | 0.13682 | 0.01526 | 0.34422 |
| SF172 | 0.09451 | -0.1754 | -0.2844 | -0.0481 | 0.0885 | 0.13411 | 1.42492 | 0.23323 | 0.25915 | 0.11402 | 0.02383 | 0.37134 | 0.27269 |
| TTC642 | 0.2029 | -0.2516 | -0.064 | -0.0098 | 0.17535 | 0.13373 | 1.42019 | 0.23488 | 0.08102 | 0.04058 | 0.3312 | 0.47336 | 0.11407 |
| CAS1 | 0.00281 | -0.1474 | -0.1995 | 0.09416 | 0.229 | 0.13333 | 1.4153 | 0.2366 | 0.49236 | 0.15616 | 0.08465 | 0.25994 | 0.05675 |
| SNU349 | 0.29232 | -0.083 | -0.1309 | 0.09439 | 0.15444 | 0.13288 | 1.40985 | 0.23853 | 0.02077 | 0.28535 | 0.18499 | 0.25943 | 0.14467 |
| DU145 | 0.23063 | -0.1676 | -0.1249 | -0.1759 | 0.13096 | 0.13269 | 1.40753 | 0.23935 | 0.05543 | 0.12486 | 0.19633 | 0.11329 | 0.18487 |
| HCC2450 | 0.1042 | -0.2608 | -0.1869 | -0.0785 | 0.10355 | 0.13258 | 1.40612 | 0.23985 | 0.23807 | 0.03514 | 0.09922 | 0.29601 | 0.23946 |
| HHUA | 0.11366 | -0.2693 | -0.1023 | -0.1739 | 0.1062 | 0.13252 | 1.40548 | 0.24008 | 0.2184 | 0.03066 | 0.24207 | 0.1161 | 0.23385 |
| UMUC5 | 0.06611 | -0.363 | -0.0653 | -0.0245 | -0.0339 | 0.13251 | 1.40531 | 0.24014 | 0.32589 | 0.00518 | 0.32798 | 0.43356 | 0.40863 |
| OV7 | 0.04624 | -0.2071 | -0.1602 | -0.0534 | 0.2164 | 0.13229 | 1.40265 | 0.2411 | 0.37619 | 0.07663 | 0.13579 | 0.35789 | 0.06766 |
| RMUGS | 0.11413 | -0.2349 | -0.15 | -0.2028 | 0.06835 | 0.13214 | 1.40084 | 0.24175 | 0.21744 | 0.05215 | 0.15177 | 0.08109 | 0.32039 |
| LNCAPCLONEFGC | 0.16545 | -0.3427 | -0.0144 | -0.0662 | 0.02094 | 0.1321 | 1.40026 | 0.24195 | 0.12796 | 0.00796 | 0.46093 | 0.32577 | 0.44322 |
| RD | -0.0756 | -0.1982 | -0.1053 | -0.2069 | 0.13119 | 0.13194 | 1.39835 | 0.24264 | 0.30276 | 0.08613 | 0.23567 | 0.07685 | 0.18445 |
| SUM190PT | 0.09075 | -0.2763 | -0.1996 | -0.094 | 0.02276 | 0.13186 | 1.39739 | 0.24299 | 0.26758 | 0.02731 | 0.08458 | 0.26028 | 0.43832 |
| 94T778 | 0.1264 | -0.1532 | -0.2272 | 0.15956 | 0.12933 | 0.13177 | 1.39631 | 0.24338 | 0.19339 | 0.1466 | 0.05818 | 0.13674 | 0.1879 |
| JR | 0.21992 | -0.1712 | -0.1629 | -0.0692 | 0.17401 | 0.13172 | 1.39566 | 0.24361 | 0.06446 | 0.11979 | 0.13176 | 0.31819 | 0.1159 |
| KMRC2 | 0.15885 | -0.2669 | -0.1366 | -0.1651 | 0.03096 | 0.13168 | 1.39517 | 0.24379 | 0.13781 | 0.03186 | 0.17457 | 0.12844 | 0.41638 |
| 5637 | 0.19669 | -0.1707 | -0.1961 | -0.0749 | 0.15696 | 0.13165 | 1.39478 | 0.24393 | 0.08777 | 0.12046 | 0.08841 | 0.30449 | 0.14072 |
| SNU1033 | -0.0675 | -0.1826 | 0.07495 | -0.0725 | 0.25151 | 0.13151 | 1.39312 | 0.24453 | 0.3224 | 0.1046 | 0.30438 | 0.31025 | 0.04064 |
| TTC549 | 0.05949 | -0.2569 | -0.0261 | -0.217 | 0.13068 | 0.13151 | 1.39307 | 0.24455 | 0.34235 | 0.03738 | 0.42942 | 0.0671 | 0.18539 |
| NCIH3122 | 0.09376 | -0.2627 | -0.031 | -0.1561 | 0.17107 | 0.13112 | 1.38834 | 0.24627 | 0.26084 | 0.03411 | 0.4163 | 0.1421 | 0.11994 |
| NCIH841 | 0.10753 | -0.0565 | -0.0427 | -0.153 | 0.3098 | 0.13099 | 1.38679 | 0.24684 | 0.23105 | 0.35001 | 0.38551 | 0.14692 | 0.01515 |
| PACADD161 | 0.00653 | -0.2564 | -0.1544 | -0.0095 | 0.15975 | 0.13071 | 1.38334 | 0.24811 | 0.48225 | 0.03765 | 0.14475 | 0.47429 | 0.13644 |
| JURLMK1 | -0.0728 | -0.0065 | -0.2532 | -0.0693 | 0.22348 | 0.13056 | 1.3815 | 0.24879 | 0.30952 | 0.48224 | 0.03958 | 0.31809 | 0.06134 |
| SJSA1 | -0.0943 | -0.2222 | -0.1501 | -0.0545 | 0.15554 | 0.13048 | 1.3805 | 0.24916 | 0.25959 | 0.06247 | 0.15163 | 0.35507 | 0.14295 |
| A375SKINCJ2 | 0.1679 | -0.2689 | -0.1925 | -0.0293 | 0.00516 | 0.1301 | 1.3759 | 0.25086 | 0.12442 | 0.03084 | 0.09257 | 0.4209 | 0.48595 |
| OMM25 | 0.11663 | -0.2479 | -0.0509 | -0.2454 | 0.06933 | 0.13001 | 1.37483 | 0.25126 | 0.21242 | 0.04295 | 0.36408 | 0.04462 | 0.31797 |
| ASH3 | 0.16121 | -0.1871 | -0.171 | -0.214 | 0.01424 | 0.12908 | 1.3636 | 0.25547 | 0.13423 | 0.099 | 0.12 | 0.06987 | 0.46131 |
| SNU398 | 0.07722 | -0.2961 | 0.01428 | -0.0062 | 0.16641 | 0.12896 | 1.36206 | 0.25605 | 0.29896 | 0.01943 | 0.46121 | 0.48306 | 0.12656 |
| RPE1SS6 | 0.185 | -0.2543 | -0.1048 | -0.1797 | 0.04236 | 0.12866 | 1.35842 | 0.25743 | 0.10159 | 0.03893 | 0.23686 | 0.10836 | 0.38629 |
| CTV1DM | 0.16686 | -0.1918 | 0.04532 | -0.2539 | 0.13095 | 0.12826 | 1.35359 | 0.25927 | 0.12591 | 0.09336 | 0.37859 | 0.0392 | 0.1849 |
| HCC366 | 0.07875 | -0.0298 | -0.162 | -0.1306 | 0.27961 | 0.12782 | 1.34826 | 0.26132 | 0.29533 | 0.41959 | 0.13305 | 0.18552 | 0.02584 |
| C33A | 0.1135 | -0.2744 | -0.2004 | -0.0402 | 0.02458 | 0.12749 | 1.34427 | 0.26286 | 0.21873 | 0.02818 | 0.08374 | 0.39196 | 0.43343 |
| OCIC4P | 0.14887 | -0.1944 | -0.1748 | -0.1561 | 0.12389 | 0.1274 | 1.34324 | 0.26326 | 0.15365 | 0.09041 | 0.11485 | 0.14206 | 0.19819 |
| COGAR359 | 0.02466 | -0.2272 | -0.1471 | -0.1829 | 0.11269 | 0.12736 | 1.34276 | 0.26344 | 0.43322 | 0.05821 | 0.15654 | 0.1042 | 0.22038 |
| 143B | 0.16293 | -0.1986 | -0.0858 | -0.0531 | 0.23019 | 0.12731 | 1.34216 | 0.26368 | 0.13166 | 0.08564 | 0.27898 | 0.35843 | 0.05578 |
| HSC4 | 0.14067 | -0.2023 | -0.1954 | -0.1762 | 0.02961 | 0.12651 | 1.33244 | 0.26747 | 0.16751 | 0.08167 | 0.08922 | 0.11298 | 0.41996 |
| C4I | 0.17504 | -0.081 | -0.1904 | -0.2166 | 0.088 | 0.12644 | 1.33164 | 0.26779 | 0.1145 | 0.28999 | 0.09505 | 0.06748 | 0.27382 |
| FADU | 0.04842 | -0.3441 | -0.0603 | -0.1032 | -0.0111 | 0.12629 | 1.32986 | 0.26849 | 0.37056 | 0.00775 | 0.34038 | 0.24029 | 0.46973 |
| MV411 | 0.23303 | -0.0907 | 0.18416 | -0.223 | 0.07848 | 0.1262 | 1.32877 | 0.26892 | 0.05355 | 0.26769 | 0.10263 | 0.06179 | 0.29598 |
| SNU1076 | 0.1165 | -0.2454 | -0.2365 | 0.01217 | -0.0184 | 0.12602 | 1.3266 | 0.26978 | 0.21268 | 0.04464 | 0.05088 | 0.46693 | 0.44998 |
| EW8 | -0.0065 | -0.0464 | -0.3072 | -0.1406 | -0.0158 | 0.12593 | 1.32542 | 0.27025 | 0.48226 | 0.37584 | 0.01588 | 0.16756 | 0.4572 |
| LU99 | 0.08299 | -0.2071 | -0.2228 | 0.01613 | 0.14079 | 0.12494 | 1.31353 | 0.27501 | 0.28538 | 0.07668 | 0.06195 | 0.45621 | 0.1673 |
| TE10 | -0.0122 | -0.1047 | -0.3105 | 0.02244 | 0.11009 | 0.12487 | 1.31271 | 0.27534 | 0.46676 | 0.23696 | 0.01497 | 0.43919 | 0.22571 |
| HS746T | 0.18216 | -0.2676 | -0.0165 | -0.1145 | 0.13604 | 0.12431 | 1.30604 | 0.27804 | 0.10516 | 0.03154 | 0.45511 | 0.21661 | 0.17566 |
| DU4475 | 0.25513 | -0.2712 | -0.0272 | -0.0184 | 0.04634 | 0.1243 | 1.30591 | 0.27809 | 0.03843 | 0.02973 | 0.42654 | 0.45014 | 0.37595 |
| SKRC31 | -0.0298 | -0.3001 | 0.02508 | -0.1795 | 0.04438 | 0.12378 | 1.29965 | 0.28065 | 0.41942 | 0.01809 | 0.4321 | 0.10857 | 0.38103 |
| LU134A | 0.24261 | -0.0296 | -0.0004 | -0.1337 | 0.24926 | 0.12321 | 1.29277 | 0.28348 | 0.04651 | 0.4199 | 0.49888 | 0.17981 | 0.04207 |
| KMRC1 | 0.19159 | 0.13692 | -0.0849 | 0.20811 | 0.17233 | 0.12311 | 1.29165 | 0.28395 | 0.09362 | 0.17409 | 0.28101 | 0.07565 | 0.1182 |
| SUIT2 | 0.00703 | 0.01494 | -0.3017 | 0.01731 | 0.18902 | 0.12305 | 1.29089 | 0.28426 | 0.48088 | 0.45944 | 0.01756 | 0.453 | 0.09667 |
| JHOS4 | 0.13207 | -0.3 | -0.0524 | -0.0224 | 0.11885 | 0.12293 | 1.28946 | 0.28485 | 0.18283 | 0.0181 | 0.36039 | 0.43919 | 0.20801 |
| NTERA2CLD1 | 0.05077 | -0.2124 | -0.1753 | -0.2124 | -0.0226 | 0.12256 | 1.28506 | 0.28669 | 0.36451 | 0.07145 | 0.11415 | 0.0714 | 0.43883 |
| MERO48A | 0.1102 | -0.2245 | 0.0276 | 0.12175 | 0.18611 | 0.12233 | 1.28227 | 0.28786 | 0.2255 | 0.06044 | 0.42535 | 0.20233 | 0.10021 |
| ONDA7 | 0.13543 | -0.1895 | -0.1866 | 0.0108 | 0.17777 | 0.12229 | 1.28177 | 0.28807 | 0.17675 | 0.09607 | 0.09966 | 0.47066 | 0.11085 |
| ICC3 | 0.01126 | -0.1485 | -0.214 | 0.04597 | 0.2056 | 0.12204 | 1.27883 | 0.2893 | 0.46941 | 0.15431 | 0.06988 | 0.3769 | 0.0782 |
| PATU8988T | 0.25206 | -0.0644 | -0.0644 | -0.0216 | 0.24979 | 0.12203 | 1.27869 | 0.28936 | 0.0403 | 0.33014 | 0.33007 | 0.44142 | 0.04173 |
| RL | 0.11082 | -0.3172 | -0.0757 | -0.0701 | 0.06354 | 0.12198 | 1.27809 | 0.28962 | 0.22421 | 0.01319 | 0.30268 | 0.31615 | 0.33224 |
| UPCISCC200 | 0.16009 | -0.2943 | -0.0838 | -0.0148 | 0.08966 | 0.12182 | 1.27626 | 0.29039 | 0.13592 | 0.02006 | 0.28352 | 0.45985 | 0.27006 |
| NZM42 | 0.21627 | -0.2413 | -0.1408 | -0.0676 | 0.05839 | 0.12138 | 1.27091 | 0.29265 | 0.06778 | 0.04741 | 0.16729 | 0.32217 | 0.34512 |
| HEC59 | -0.0095 | -0.1876 | -0.122 | -0.0409 | 0.23039 | 0.12137 | 1.27079 | 0.29271 | 0.47427 | 0.09835 | 0.2018 | 0.3902 | 0.05563 |
| KALS1 | 0.08202 | 0.09234 | -0.1362 | -0.0597 | 0.31128 | 0.12095 | 1.26583 | 0.29482 | 0.28765 | 0.26401 | 0.17543 | 0.34186 | 0.01473 |
| REH | -0.1089 | -0.1983 | -0.1748 | -0.1023 | 0.10057 | 0.12083 | 1.26447 | 0.29541 | 0.2282 | 0.08599 | 0.11482 | 0.24221 | 0.24586 |
| HT1080 | 0.22318 | 0.00377 | -0.087 | -0.2126 | 0.16524 | 0.12067 | 1.26247 | 0.29626 | 0.0616 | 0.48974 | 0.27614 | 0.07119 | 0.12827 |
| K029AX | -0.1711 | -0.0082 | -0.298 | -0.0108 | -0.0564 | 0.12047 | 1.2601 | 0.29728 | 0.11994 | 0.4777 | 0.01878 | 0.47059 | 0.35024 |
| AM38 | 0.06061 | -0.2544 | -0.0668 | 0.085 | 0.16809 | 0.12045 | 1.25988 | 0.29738 | 0.33954 | 0.03886 | 0.3242 | 0.28071 | 0.12415 |
| SW837 | 0.08152 | -0.1288 | -0.2458 | -0.1713 | 0.05172 | 0.12045 | 1.25985 | 0.29739 | 0.28881 | 0.18887 | 0.04436 | 0.11957 | 0.36207 |
| F36P | 0.14883 | 0.02151 | -0.2586 | -0.1164 | 0.13231 | 0.12024 | 1.25742 | 0.29844 | 0.15372 | 0.44168 | 0.0364 | 0.21292 | 0.1824 |
| INA6 | -0.2027 | 0.10856 | 0.04095 | 0.05132 | 0.26007 | 0.11964 | 1.25032 | 0.30153 | 0.0812 | 0.22888 | 0.38997 | 0.36311 | 0.03556 |
| UACC893 | 0.01467 | -0.1186 | -0.3113 | 0.00661 | -0.0904 | 0.11875 | 1.23977 | 0.30616 | 0.46015 | 0.20857 | 0.01473 | 0.48203 | 0.26832 |
| CMLT1 | -0.0634 | -0.2504 | -0.1454 | 0.00108 | 0.11449 | 0.11868 | 1.23892 | 0.30654 | 0.33265 | 0.04133 | 0.15939 | 0.49706 | 0.21673 |
| KYSE510 | 0.05438 | -0.2237 | -0.2425 | -0.0765 | -0.0171 | 0.11867 | 1.23881 | 0.30659 | 0.35528 | 0.06112 | 0.04657 | 0.30068 | 0.45358 |
| WM983B | -0.0021 | 0.05684 | -0.0303 | -0.1608 | 0.30468 | 0.11859 | 1.23786 | 0.30701 | 0.4944 | 0.34904 | 0.41801 | 0.13485 | 0.01664 |
| SUDHL4 | 0.1142 | -0.1513 | -0.0468 | -0.0799 | 0.27022 | 0.11834 | 1.23485 | 0.30835 | 0.2173 | 0.1497 | 0.3747 | 0.29254 | 0.0302 |
| ARH77 | 0.03486 | -0.2009 | -0.0804 | -0.2705 | 0.02881 | 0.11826 | 1.23397 | 0.30874 | 0.40601 | 0.08316 | 0.29141 | 0.03006 | 0.42211 |
| KCIMOH1 | 0.17027 | -0.2157 | -0.2074 | -0.0561 | 0.04105 | 0.11804 | 1.23133 | 0.30991 | 0.12106 | 0.06827 | 0.07633 | 0.3508 | 0.38972 |

| | | | | | | | | | | | | | |
|---|---|---|---|---|---|---|---|---|---|---|---|---|---|
| NB4 | -0.0859 | -0.1465 | -0.2055 | -0.1655 | 0.06937 | 0.11791 | 1.22972 | 0.31063 | 0.27867 | 0.15752 | 0.07826 | 0.12792 | 0.3179 |
| NOMO1 | 0.22241 | 0.04137 | -0.0085 | -0.0472 | 0.28299 | 0.11783 | 1.22884 | 0.31103 | 0.06227 | 0.38889 | 0.47676 | 0.37369 | 0.0244 |
| CAOV3 | 0.24283 | -0.0736 | -0.0241 | -0.0282 | 0.25384 | 0.11781 | 1.22854 | 0.31116 | 0.04636 | 0.30769 | 0.43463 | 0.42372 | 0.03921 |
| BOKU | -0.0577 | -0.1428 | -0.1709 | 0.05407 | 0.21653 | 0.11779 | 1.22835 | 0.31125 | 0.34693 | 0.16376 | 0.12016 | 0.35606 | 0.06754 |
| PK8 | 0.10473 | -0.3006 | 0.02223 | 0.03924 | 0.10936 | 0.11769 | 1.22716 | 0.31178 | 0.23695 | 0.01792 | 0.43974 | 0.39448 | 0.22723 |
| SKNBE2 | -0.2042 | -0.086 | -0.0448 | -0.1747 | 0.13604 | 0.11762 | 1.22637 | 0.31214 | 0.0797 | 0.27844 | 0.37992 | 0.11502 | 0.17565 |
| SLVL | 0.02923 | -0.0249 | -0.0852 | 0.03223 | 0.32831 | 0.11725 | 1.22197 | 0.31412 | 0.42098 | 0.43262 | 0.28033 | 0.413 | 0.01064 |
| RCHACV | 0.09765 | -0.1817 | -0.176 | -0.1697 | 0.11349 | 0.11677 | 1.21637 | 0.31666 | 0.25222 | 0.10575 | 0.11323 | 0.12184 | 0.21875 |
| UMUC14 | -0.2058 | -0.1022 | -0.078 | 0.07562 | 0.18224 | 0.1162 | 1.20956 | 0.31977 | 0.07796 | 0.24236 | 0.2972 | 0.30279 | 0.10505 |
| OCIMY7 | 0.16894 | -0.1973 | -0.0471 | -0.2541 | -0.0227 | 0.11616 | 1.20907 | 0.31999 | 0.12294 | 0.08707 | 0.37389 | 0.03904 | 0.43852 |
| SYO1 | 0.12378 | -0.1724 | 0.00477 | 0.10627 | 0.23068 | 0.11593 | 1.20645 | 0.3212 | 0.19839 | 0.11815 | 0.48702 | 0.23368 | 0.05539 |
| CHL1DM | -0.0418 | -0.1306 | -0.2881 | -0.0022 | 0.07935 | 0.11583 | 1.20527 | 0.32175 | 0.3877 | 0.18555 | 0.02235 | 0.49404 | 0.29392 |
| MALME3M | 0.11384 | -0.3178 | -0.0437 | -0.0917 | 0.04069 | 0.11579 | 1.20472 | 0.322 | 0.21804 | 0.01304 | 0.38277 | 0.26553 | 0.39066 |
| DLD1 | 0.15338 | -0.1715 | -0.0753 | -0.238 | 0.10192 | 0.11549 | 1.20123 | 0.32361 | 0.14635 | 0.11935 | 0.3036 | 0.04981 | 0.24295 |
| SCC25 | 0.21099 | -0.1503 | -0.1656 | -0.0904 | 0.14082 | 0.11545 | 1.20083 | 0.3238 | 0.0728 | 0.15129 | 0.1278 | 0.26849 | 0.16724 |
| CHAGOK1 | 0.02525 | -0.2483 | -0.0615 | -0.126 | 0.15801 | 0.11498 | 1.19526 | 0.32639 | 0.43164 | 0.04271 | 0.3372 | 0.19416 | 0.13911 |
| CALU6 | -0.0309 | -0.2483 | -0.106 | -0.1258 | 0.11236 | 0.11451 | 1.18968 | 0.329 | 0.41665 | 0.0427 | 0.23421 | 0.19454 | 0.22104 |
| MAC2A | 0.07655 | -0.2297 | -0.1539 | -0.1181 | 0.12069 | 0.11438 | 1.18819 | 0.3297 | 0.30057 | 0.05619 | 0.14555 | 0.20952 | 0.20439 |
| KPNYS | 0.05835 | -0.2647 | 0.07992 | -0.0888 | 0.15902 | 0.11432 | 1.18747 | 0.33004 | 0.34522 | 0.03301 | 0.29258 | 0.27193 | 0.13755 |
| TE5 | 0.09703 | -0.2551 | -0.1621 | -0.129 | 0.02678 | 0.11429 | 1.1871 | 0.33022 | 0.25358 | 0.03846 | 0.13295 | 0.18856 | 0.42753 |
| NCIH1650 | 0.13765 | -0.068 | -0.0249 | 0.01173 | 0.30657 | 0.11426 | 1.18674 | 0.33039 | 0.1728 | 0.32112 | 0.43246 | 0.46813 | 0.01608 |
| MDAMB231 | 0.28195 | 0.01399 | -0.0815 | -0.0053 | 0.19866 | 0.11383 | 1.18171 | 0.33277 | 0.02484 | 0.46199 | 0.28884 | 0.48557 | 0.0856 |
| BXPC3 | -0.0321 | -0.1415 | -0.2806 | -0.0355 | 0.06517 | 0.11377 | 1.18105 | 0.33308 | 0.4133 | 0.16606 | 0.02543 | 0.4044 | 0.32819 |
| IGR39 | 0.0508 | -0.1664 | -0.2267 | -0.0774 | 0.13716 | 0.11373 | 1.18057 | 0.33331 | 0.36444 | 0.12657 | 0.05866 | 0.29861 | 0.17367 |
| K562 | 0.19573 | -0.1146 | -0.0407 | -0.177 | 0.20276 | 0.11364 | 1.17951 | 0.33381 | 0.08886 | 0.21656 | 0.39055 | 0.11187 | 0.08117 |
| OCILY18 | 0.03606 | -0.1741 | -0.2392 | -0.12 | 0.06104 | 0.11351 | 1.17799 | 0.33453 | 0.40284 | 0.11579 | 0.04895 | 0.20565 | 0.33846 |
| NB13 | -0.1321 | -0.253 | -0.061 | -0.1426 | -0.0059 | 0.1133 | 1.17555 | 0.3357 | 0.18271 | 0.03972 | 0.33848 | 0.16415 | 0.48388 |
| D341Med | -0.1383 | -0.2013 | -0.1302 | -0.1525 | 0.02045 | 0.11314 | 1.17362 | 0.33662 | 0.1716 | 0.08278 | 0.18626 | 0.14776 | 0.44455 |
| CAL62 | -0.0207 | -0.172 | -0.0362 | -0.1301 | 0.22705 | 0.11291 | 1.17097 | 0.33789 | 0.44384 | 0.11868 | 0.40258 | 0.18649 | 0.05834 |
| KOSC2 | 0.0107 | -0.1916 | -0.2682 | 0.00413 | 0.01198 | 0.11267 | 1.16814 | 0.33925 | 0.47093 | 0.09359 | 0.03121 | 0.48878 | 0.46744 |
| PANC0327 | 0.12329 | -0.0586 | -0.2003 | 0.09243 | 0.22448 | 0.11255 | 1.16683 | 0.33989 | 0.19935 | 0.3446 | 0.08385 | 0.2638 | 0.06049 |
| UBLC1 | 0.11809 | -0.1859 | -0.1988 | -0.1246 | 0.09337 | 0.11246 | 1.16573 | 0.34042 | 0.20952 | 0.10043 | 0.0854 | 0.19687 | 0.26169 |
| BICR78 | 0.116 | -0.2435 | -0.1954 | -0.0502 | -0.0897 | 0.11243 | 1.16537 | 0.34059 | 0.21368 | 0.04587 | 0.08925 | 0.36588 | 0.26985 |
| SNU410 | -0.0518 | -0.1913 | -0.2221 | -0.0359 | 0.0952 | 0.11202 | 1.16056 | 0.34292 | 0.36183 | 0.09399 | 0.06257 | 0.40333 | 0.25762 |
| H3118 | 0.01666 | -0.208 | -0.2438 | 0.00169 | 0.05434 | 0.11197 | 1.15995 | 0.34322 | 0.45476 | 0.07578 | 0.0457 | 0.49541 | 0.35537 |
| THP1 | 0.07015 | -0.0603 | -0.2654 | -0.1424 | 0.08317 | 0.11186 | 1.15877 | 0.3438 | 0.31598 | 0.34042 | 0.03268 | 0.16446 | 0.28498 |
| OCIMY5 | -0.1305 | 0.04398 | -0.2116 | -0.1123 | 0.17256 | 0.11183 | 1.15836 | 0.344 | 0.18564 | 0.38206 | 0.07216 | 0.22123 | 0.11788 |
| JHH6 | -0.0344 | -0.1962 | -0.213 | -0.0333 | 0.11202 | 0.11171 | 1.157 | 0.34466 | 0.40731 | 0.08829 | 0.07087 | 0.41004 | 0.22175 |
| OCIAML2 | 0.10692 | -0.1914 | -0.081 | -0.0307 | 0.22784 | 0.11169 | 1.15672 | 0.3448 | 0.23233 | 0.0938 | 0.28996 | 0.41707 | 0.05769 |
| TE8 | -0.0318 | -0.1682 | -0.2768 | -0.0271 | -0.0253 | 0.11158 | 1.15544 | 0.34542 | 0.41406 | 0.12396 | 0.02708 | 0.42674 | 0.43146 |
| T84 | -0.1001 | -0.1658 | -0.0911 | -0.163 | 0.148 | 0.1114 | 1.15332 | 0.34646 | 0.24689 | 0.1275 | 0.26684 | 0.13158 | 0.15509 |
| A673 | 0.0131 | -0.0908 | -0.2236 | -0.1697 | -0.1424 | 0.11127 | 1.15184 | 0.34719 | 0.46439 | 0.26757 | 0.06125 | 0.12193 | 0.16455 |
| MDAMB468 | 0.28925 | -0.088 | 0.07512 | 0.01959 | 0.15712 | 0.11116 | 1.15052 | 0.34784 | 0.02191 | 0.27379 | 0.30397 | 0.44686 | 0.14048 |
| PECAPJ41CLONED2 | 0.1977 | -0.1566 | -0.138 | -0.0164 | 0.18211 | 0.1108 | 1.14642 | 0.34986 | 0.08666 | 0.14123 | 0.17214 | 0.45555 | 0.10522 |
| NCIH1915 | 0.15935 | -0.192 | 0.06437 | -0.0845 | 0.21132 | 0.1108 | 1.14635 | 0.3499 | 0.13705 | 0.0932 | 0.33019 | 0.28192 | 0.07248 |
| KNS62 | 0.08414 | -0.2437 | -0.0403 | -0.1683 | 0.13079 | 0.1106 | 1.1441 | 0.35101 | 0.28272 | 0.04574 | 0.39156 | 0.12391 | 0.18519 |
| TUHR10TKB | 0.04816 | -0.0285 | -0.15 | 0.1011 | 0.2793 | 0.11057 | 1.1437 | 0.35121 | 0.37122 | 0.42295 | 0.15179 | 0.24471 | 0.02598 |
| NCO2 | -0.0825 | -0.1841 | -0.1791 | -0.0495 | 0.13425 | 0.11055 | 1.14345 | 0.35133 | 0.28652 | 0.10267 | 0.10907 | 0.36784 | 0.17888 |
| NO36 | 0.0378 | -0.3026 | -0.1142 | -0.098 | -0.0512 | 0.11048 | 1.14236 | 0.35187 | 0.39827 | 0.01728 | 0.21732 | 0.25138 | 0.36335 |
| OCIP5X | 0.07578 | -0.1884 | -0.0713 | -0.2604 | 0.04994 | 0.11032 | 1.14076 | 0.35267 | 0.3024 | 0.09739 | 0.31311 | 0.03539 | 0.36665 |
| LN464 | 0.11586 | -0.0932 | -0.0489 | -0.3016 | 0.04356 | 0.1101 | 1.13827 | 0.35391 | 0.21397 | 0.26216 | 0.36932 | 0.01759 | 0.38316 |
| KYSE520 | 0.1705 | -0.1479 | -0.0865 | -0.0395 | 0.23193 | 0.10993 | 1.13625 | 0.35492 | 0.12074 | 0.15522 | 0.27737 | 0.39385 | 0.05441 |
| NB5 | 0.10373 | -0.2643 | 0.01248 | -0.1717 | 0.09949 | 0.10987 | 1.13552 | 0.35528 | 0.23907 | 0.03322 | 0.46607 | 0.11902 | 0.24821 |
| U87MG | -0.0358 | -0.2866 | -0.0905 | -0.1346 | -0.0776 | 0.10962 | 1.13262 | 0.35673 | 0.40363 | 0.02295 | 0.26818 | 0.17819 | 0.29803 |
| MM127 | 0.10209 | -0.2237 | 0.04644 | -0.1167 | 0.19036 | 0.10957 | 1.13212 | 0.35699 | 0.24259 | 0.06117 | 0.37566 | 0.21226 | 0.09507 |
| DOV13 | 0.10651 | -0.2716 | -0.1255 | -0.0593 | 0.07715 | 0.1087 | 1.12197 | 0.36212 | 0.23319 | 0.02952 | 0.19507 | 0.34286 | 0.29912 |
| SNU449 | 0.05724 | -0.1416 | -0.2409 | -0.1276 | 0.0722 | 0.10831 | 1.11754 | 0.36438 | 0.34802 | 0.16592 | 0.04775 | 0.19123 | 0.31101 |
| CHP212 | 0.03816 | -0.0904 | 0.07145 | -0.3236 | 0.02323 | 0.10829 | 1.11725 | 0.36453 | 0.3973 | 0.2683 | 0.31282 | 0.01167 | 0.43705 |
| A375SKINCJ1 | 0.14466 | -0.1367 | -0.1113 | -0.2432 | 0.05904 | 0.10816 | 1.11571 | 0.36531 | 0.16068 | 0.1745 | 0.22316 | 0.04611 | 0.34349 |
| CCK81 | 0.12525 | -0.1232 | -0.1659 | -0.2145 | 0.06738 | 0.10783 | 1.1119 | 0.36727 | 0.19558 | 0.1995 | 0.12724 | 0.06941 | 0.32276 |
| U251MG | 0.25248 | -0.1377 | -0.1429 | 0.02325 | 0.11124 | 0.1078 | 1.11163 | 0.36741 | 0.04004 | 0.17272 | 0.16372 | 0.43701 | 0.22334 |
| CCFSTTG1 | 0.11512 | -0.2542 | -0.0574 | 0.05601 | 0.13942 | 0.10774 | 1.11088 | 0.3678 | 0.21545 | 0.039 | 0.34758 | 0.35114 | 0.16969 |
| NCIH28 | 0.0495 | -0.2074 | -0.2352 | -0.0611 | 0.02094 | 0.10772 | 1.11071 | 0.36788 | 0.36777 | 0.0764 | 0.05192 | 0.3384 | 0.44322 |
| HCC1143 | 0.19006 | -0.2692 | -0.0743 | -0.052 | 0.05245 | 0.10751 | 1.10824 | 0.36915 | 0.09544 | 0.03074 | 0.30587 | 0.36132 | 0.3602 |
| MB1 | -0.036 | -0.1732 | 0.00046 | 0.03278 | 0.23792 | 0.10744 | 1.10746 | 0.36956 | 0.40299 | 0.11704 | 0.49875 | 0.41153 | 0.04986 |
| NCIH23 | 0.09101 | -0.1926 | 0.19892 | -0.1221 | 0.13549 | 0.10744 | 1.10742 | 0.36958 | 0.267 | 0.09242 | 0.08531 | 0.20161 | 0.17665 |
| KELLY | -0.0731 | -0.1786 | -0.034 | -0.1119 | 0.19736 | 0.10717 | 1.10426 | 0.37121 | 0.30891 | 0.10973 | 0.40825 | 0.22192 | 0.08704 |
| NCIH520 | 0.08082 | -0.2472 | 0.08364 | -0.0927 | 0.15983 | 0.10686 | 1.10076 | 0.37304 | 0.29045 | 0.04342 | 0.28388 | 0.26328 | 0.13633 |
| H376 | -0.0229 | -0.2334 | -0.1876 | 0.0901 | -0.1191 | 0.10686 | 1.10069 | 0.37307 | 0.43804 | 0.05324 | 0.09844 | 0.26906 | 0.20751 |
| SNU1077 | -0.0334 | -0.1765 | -0.209 | -0.091 | 0.10049 | 0.10647 | 1.09625 | 0.37539 | 0.40987 | 0.1126 | 0.0748 | 0.267 | 0.24604 |
| NZM7 | 0.08516 | -0.2739 | -0.116 | -0.1345 | 0.00131 | 0.10644 | 1.09589 | 0.37558 | 0.28035 | 0.02845 | 0.21371 | 0.17836 | 0.49643 |
| RBE | 0.07847 | -0.1747 | -0.2187 | -0.047 | 0.12386 | 0.1064 | 1.09545 | 0.37581 | 0.29599 | 0.11497 | 0.06555 | 0.37417 | 0.19825 |
| LS411N | 0.02548 | -0.1735 | 0.18103 | -0.2534 | -0.0341 | 0.10618 | 1.09286 | 0.37716 | 0.43101 | 0.11657 | 0.1066 | 0.03946 | 0.4079 |
| SISO | 0.00276 | -0.1794 | -0.2246 | -0.1135 | -0.1039 | 0.10586 | 1.08921 | 0.37909 | 0.4925 | 0.10865 | 0.06036 | 0.21882 | 0.23862 |
| DAOY | 0.0664 | -0.2587 | -0.0978 | 0.11663 | 0.07484 | 0.10586 | 1.08921 | 0.37909 | 0.32517 | 0.03633 | 0.25188 | 0.21242 | 0.30466 |
| T24 | 0.00584 | -0.1834 | -0.1253 | -0.1301 | 0.16916 | 0.10561 | 1.08636 | 0.38059 | 0.48411 | 0.10363 | 0.19557 | 0.18639 | 0.12262 |
| GSU | 0.14105 | -0.1564 | -0.1711 | -0.1245 | 0.13384 | 0.10538 | 1.08365 | 0.38203 | 0.16685 | 0.14159 | 0.11987 | 0.19711 | 0.17962 |
| HS766T | 0.18487 | -0.1487 | -0.1754 | -0.1232 | 0.08925 | 0.10496 | 1.07888 | 0.38457 | 0.10175 | 0.15392 | 0.11401 | 0.19945 | 0.27098 |
| MUTZ8 | -0.0118 | -0.2613 | -0.0047 | -0.1006 | 0.13135 | 0.10489 | 1.0781 | 0.38498 | 0.46807 | 0.03487 | 0.48712 | 0.24569 | 0.18416 |
| 42MGBA | 0.23189 | -0.1012 | 0.11203 | 0.08672 | 0.15468 | 0.10448 | 1.0734 | 0.3875 | 0.05444 | 0.2444 | 0.22172 | 0.27677 | 0.1443 |
| MDAMB157 | 0.02029 | -0.2017 | -0.0494 | -0.0287 | 0.21728 | 0.1043 | 1.07131 | 0.38862 | 0.44496 | 0.08227 | 0.36799 | 0.42235 | 0.06685 |
| VP229 | -0.0079 | -0.191 | -0.2166 | -0.1014 | 0.04879 | 0.10424 | 1.07059 | 0.38901 | 0.47851 | 0.0943 | 0.06745 | 0.244 | 0.36961 |
| HPAFII | -0.0712 | -0.2325 | -0.1293 | -0.1189 | 0.05117 | 0.10371 | 1.06448 | 0.39231 | 0.31337 | 0.05393 | 0.18789 | 0.20789 | 0.36347 |
| SHSY5Y | 0.08253 | -0.1367 | 0.02579 | -0.1958 | 0.20714 | 0.10302 | 1.0566 | 0.3966 | 0.28646 | 0.17451 | 0.4302 | 0.08875 | 0.07664 |
| KYSE150 | 0.18268 | -0.1818 | -0.0169 | -0.1712 | 0.14134 | 0.10281 | 1.05421 | 0.3979 | 0.10449 | 0.10566 | 0.45406 | 0.11974 | 0.16634 |
| MESOV | 0.02853 | -0.2806 | -0.0092 | -0.182 | 0.00048 | 0.10279 | 1.05401 | 0.39801 | 0.42285 | 0.0254 | 0.47505 | 0.10533 | 0.49868 |
| GB1 | -0.0148 | -0.0879 | -0.0496 | -0.0706 | 0.27926 | 0.10264 | 1.05229 | 0.39895 | 0.45986 | 0.27408 | 0.36752 | 0.31477 | 0.026 |
| KARPAS1718 | 0.16001 | -0.1719 | -0.0914 | -0.2248 | -0.0226 | 0.10261 | 1.05192 | 0.39916 | 0.13605 | 0.11873 | 0.26601 | 0.06022 | 0.43881 |
| HT29 | -0.1319 | -0.1193 | -0.2508 | -0.0289 | -0.0034 | 0.10256 | 1.05139 | 0.39945 | 0.18322 | 0.20719 | 0.04107 | 0.42187 | 0.49065 |
| OSC20 | 0.15774 | -0.1779 | -0.1971 | -0.0625 | 0.07809 | 0.10227 | 1.04811 | 0.40125 | 0.13952 | 0.11065 | 0.08732 | 0.33481 | 0.29691 |
| HCC202 | 0.05155 | -0.2301 | -0.155 | 0.03407 | 0.11791 | 0.10197 | 1.04459 | 0.4032 | 0.36251 | 0.05586 | 0.14372 | 0.40812 | 0.20987 |
| D425 | 0.06437 | -0.1866 | -0.0402 | -0.1943 | 0.15477 | 0.1017 | 1.04155 | 0.40489 | 0.33017 | 0.09956 | 0.39196 | 0.09052 | 0.14415 |
| AMO1 | -0.1296 | -0.0893 | -0.2025 | -0.1207 | 0.08696 | 0.10165 | 1.04101 | 0.40518 | 0.18744 | 0.27081 | 0.08144 | 0.20435 | 0.2762 |
| SW1990 | 0.06433 | -0.289 | -0.0812 | -0.0304 | 0.06946 | 0.10147 | 1.03897 | 0.40632 | 0.33028 | 0.02199 | 0.28959 | 0.41777 | 0.31766 |
| LCLC97TM1 | 0.00402 | -0.1887 | -0.1921 | -0.1066 | 0.09108 | 0.10106 | 1.03428 | 0.40894 | 0.48906 | 0.09708 | 0.09297 | 0.23306 | 0.26684 |
| GSS | 0.25468 | -0.0168 | -0.1706 | -0.052 | 0.09535 | 0.10095 | 1.03301 | 0.40964 | 0.0387 | 0.45435 | 0.12059 | 0.36136 | 0.2573 |
| MM370 | 0.27092 | -0.0088 | -0.03 | -0.0838 | 0.17897 | 0.10092 | 1.03274 | 0.4098 | 0.02986 | 0.47597 | 0.41899 | 0.28358 | 0.10928 |
| HEC116 | -0.0045 | -0.0454 | -0.0074 | -0.314 | 0.03257 | 0.10077 | 1.03095 | 0.4108 | 0.48785 | 0.37842 | 0.4799 | 0.014 | 0.41208 |
| MS751 | 0.26026 | -0.0935 | -0.031 | -0.1668 | 0.09994 | 0.10052 | 1.02808 | 0.41241 | 0.03546 | 0.26147 | 0.41636 | 0.12598 | 0.24723 |
| SW626 | 0.14747 | -0.19 | -0.0564 | -0.1322 | 0.1668 | 0.10047 | 1.02759 | 0.41269 | 0.15596 | 0.09549 | 0.35006 | 0.1826 | 0.126 |
| JHU011 | 0.13104 | -0.124 | -0.234 | -0.1135 | 0.00924 | 0.10037 | 1.02637 | 0.41338 | 0.18473 | 0.19802 | 0.05281 | 0.21866 | 0.47489 |
| PANC0813 | 0.07686 | -0.1607 | -0.1874 | -0.1513 | 0.0868 | 0.10017 | 1.02421 | 0.4146 | 0.29982 | 0.13502 | 0.0986 | 0.14962 | 0.27658 |
| U251MGDM | 0.11115 | -0.1351 | -0.1055 | -0.1644 | 0.18057 | 0.10011 | 1.02352 | 0.41499 | 0.22354 | 0.1774 | 0.23535 | 0.12946 | 0.1072 |
| SLR23 | 0.13518 | -0.2325 | -0.1746 | -0.0618 | -0.0275 | 0.10005 | 1.02282 | 0.41538 | 0.17719 | 0.05397 | 0.11507 | 0.33659 | 0.42555 |
| TT2609C02 | 0.04158 | -0.218 | -0.1913 | -0.1177 | -0.0393 | 0.09993 | 1.02142 | 0.41618 | 0.38834 | 0.0662 | 0.09392 | 0.21023 | 0.39421 |
| AGS | 0.04744 | -0.0736 | -0.077 | -0.2125 | 0.19821 | 0.09974 | 1.01923 | 0.41742 | 0.37309 | 0.30773 | 0.29947 | 0.07131 | 0.08609 |
| RVH421SKINFV1 | 0.0693 | -0.194 | -0.1156 | -0.0088 | 0.19081 | 0.09951 | 1.01664 | 0.41889 | 0.31805 | 0.09086 | 0.21444 | 0.47594 | 0.09454 |
| NCIH1836 | 0.17718 | -0.1746 | -0.1315 | -0.1516 | -0.0888 | 0.09944 | 1.01585 | 0.41934 | 0.11163 | 0.11505 | 0.18386 | 0.14916 | 0.27196 |
| TDOTT | -0.0015 | -0.2122 | -0.2236 | -0.0195 | -0.0684 | 0.09932 | 1.01452 | 0.4201 | 0.49582 | 0.07164 | 0.06126 | 0.44717 | 0.32035 |
| CORL279 | 0.11896 | 0.20949 | 0.00214 | -0.1177 | 0.19313 | 0.0993 | 1.01429 | 0.42023 | 0.20778 | 0.07427 | 0.49417 | 0.21023 | 0.09183 |
| NCIH1666 | 0.1877 | -0.1407 | -0.1879 | -0.1158 | 0.02141 | 0.09929 | 1.0142 | 0.42029 | 0.09827 | 0.16751 | 0.09801 | 0.21413 | 0.44195 |
| CHLA15 | 0.1011 | -0.1259 | 0.01065 | -0.0761 | 0.26018 | 0.09923 | 1.01344 | 0.42072 | 0.24471 | 0.19434 | 0.47105 | 0.30152 | 0.0355 |
| OV90 | -0.0223 | -0.1895 | -0.0378 | -0.181 | 0.1439 | 0.09919 | 1.01303 | 0.42095 | 0.43967 | 0.09607 | 0.39835 | 0.10664 | 0.16196 |
| KASUMI1 | -0.0382 | -0.1063 | 0.01093 | 0.13203 | 0.23449 | 0.09908 | 1.01175 | 0.42169 | 0.39721 | 0.23373 | 0.47028 | 0.18291 | 0.05243 |
| SNU620 | -0.0483 | -0.1237 | -0.2653 | -0.0141 | 0.05968 | 0.09886 | 1.00927 | 0.42311 | 0.37083 | 0.19857 | 0.03272 | 0.46164 | 0.34188 |
| NCIH1755 | -0.051 | -0.0267 | -0.2882 | 0.00478 | 0.10395 | 0.09885 | 1.00915 | 0.42318 | 0.3638 | 0.42765 | 0.0223 | 0.487 | 0.23862 |
| NB1 | -0.0808 | -0.1617 | 0.07007 | -0.1783 | 0.14616 | 0.09861 | 1.0064 | 0.42475 | 0.29052 | 0.1335 | 0.31617 | 0.11019 | 0.15814 |
| HKBMM | 0.19667 | -0.1742 | 0.0663 | -0.0267 | 0.17996 | 0.09814 | 1.00113 | 0.4278 | 0.08781 | 0.11568 | 0.32542 | 0.42787 | 0.10798 |
| SF539 | 0.15013 | -0.0068 | 0.00132 | 0.04814 | 0.28429 | 0.09808 | 1.00047 | 0.42818 | 0.15159 | 0.48162 | 0.49641 | 0.37128 | 0.02387 |

| | | | | | | | | | | | | | |
|---|---|---|---|---|---|---|---|---|---|---|---|---|---|
| JL1 | 0.22098 | -0.1258 | -0.1843 | 0.06599 | 0.01811 | 0.09805 | 1.00016 | 0.42836 | 0.06352 | 0.1945 | 0.10249 | 0.32619 | 0.45085 |
| RAJI | 0.05284 | -0.2856 | 0.15266 | -0.0771 | -0.0489 | 0.0979 | 0.99846 | 0.42935 | 0.35919 | 0.02332 | 0.14751 | 0.29922 | 0.36938 |
| SNU81 | 0.24852 | -0.1575 | -0.0364 | -0.14 | 0.08065 | 0.09786 | 0.99794 | 0.42964 | 0.04255 | 0.13982 | 0.40204 | 0.16861 | 0.29086 |
| NGP | 0.09435 | -0.196 | -0.0704 | -0.1599 | 0.14605 | 0.0976 | 0.99499 | 0.43136 | 0.25952 | 0.0886 | 0.3153 | 0.13622 | 0.15833 |
| SKGI | 0.06189 | -0.0637 | -0.1349 | 0.02735 | 0.26455 | 0.09752 | 0.99409 | 0.43188 | 0.33634 | 0.33176 | 0.17768 | 0.42601 | 0.03311 |
| OVCAR5 | -0.0289 | -0.1217 | -0.2393 | -0.1263 | 0.01415 | 0.0971 | 0.98939 | 0.43463 | 0.42178 | 0.20235 | 0.04885 | 0.19351 | 0.46156 |
| M059K | -0.0902 | -0.0562 | -0.2806 | 0.0919 | 0.0322 | 0.09709 | 0.98926 | 0.43471 | 0.26874 | 0.35075 | 0.0254 | 0.265 | 0.41308 |
| LN382 | 0.00647 | -0.1008 | -0.1154 | 0.06827 | 0.2465 | 0.09657 | 0.98337 | 0.43817 | 0.48241 | 0.24546 | 0.2148 | 0.32058 | 0.04387 |
| SW156 | 0.10923 | -0.1086 | -0.1388 | -0.1637 | 0.16422 | 0.09649 | 0.98251 | 0.43867 | 0.2275 | 0.22873 | 0.17074 | 0.13045 | 0.12976 |
| MEL270 | 0.03112 | -0.2144 | -0.1007 | -0.0226 | -0.2318 | 0.09595 | 0.97642 | 0.44227 | 0.41595 | 0.06953 | 0.24556 | 0.43873 | 0.05452 |
| KM12 | 0.00601 | -0.0728 | 0.10633 | -0.1668 | 0.22768 | 0.09559 | 0.97234 | 0.4447 | 0.48366 | 0.30945 | 0.23356 | 0.12599 | 0.05782 |
| SMZ1 | 0.08939 | -0.1988 | 0.0133 | -0.2491 | 0.0204 | 0.09539 | 0.97009 | 0.44604 | 0.27067 | 0.08544 | 0.46387 | 0.04218 | 0.44467 |
| DKMG | -0.008 | -0.0489 | 0.03969 | 0.13935 | 0.25392 | 0.09535 | 0.96969 | 0.44628 | 0.47833 | 0.36927 | 0.3933 | 0.16981 | 0.03916 |
| NZOV9 | 0.1472 | -0.2222 | -0.1042 | -0.0816 | 0.10664 | 0.09508 | 0.96665 | 0.44809 | 0.15641 | 0.06246 | 0.23816 | 0.28861 | 0.23291 |
| RT112 | -0.0454 | -0.0047 | -0.0064 | -0.0657 | 0.29181 | 0.09503 | 0.96603 | 0.44846 | 0.3784 | 0.4873 | 0.48267 | 0.32683 | 0.02095 |
| 95T1000 | 0.03293 | -0.1895 | -0.2368 | 0.02202 | 0.00023 | 0.09484 | 0.96398 | 0.44969 | 0.41113 | 0.09605 | 0.05066 | 0.44031 | 0.49938 |
| NCIH510 | 0.06549 | -0.1566 | -0.0052 | -0.1942 | 0.17366 | 0.09476 | 0.96307 | 0.45024 | 0.32741 | 0.14125 | 0.48578 | 0.09059 | 0.11637 |
| SUM159PT | 0.102 | -0.0452 | -0.1999 | -0.1752 | 0.09436 | 0.09474 | 0.96285 | 0.45037 | 0.24278 | 0.37896 | 0.08429 | 0.11434 | 0.25949 |
| SKMEL19 | 0.24975 | -0.1236 | -0.0615 | 0.02635 | -0.1331 | 0.09464 | 0.96171 | 0.45106 | 0.04176 | 0.19868 | 0.33733 | 0.42869 | 0.18095 |
| RCM1 | 0.07204 | -0.2272 | -0.1279 | -0.1641 | -0.0206 | 0.0946 | 0.96123 | 0.45135 | 0.3114 | 0.05824 | 0.19061 | 0.12988 | 0.44425 |
| HTMMT | 0.13853 | -0.1276 | -0.0509 | -0.1765 | 0.17753 | 0.09435 | 0.95847 | 0.45301 | 0.17124 | 0.19115 | 0.36423 | 0.11259 | 0.11117 |
| SNU387 | -0.2262 | 0.06359 | -0.206 | 0.05002 | 0.04952 | 0.09419 | 0.9567 | 0.45408 | 0.05906 | 0.33212 | 0.07785 | 0.36643 | 0.36772 |
| SNU1 | 0.18597 | -0.093 | 0.05517 | 0.02491 | 0.22899 | 0.09403 | 0.95491 | 0.45516 | 0.10039 | 0.26243 | 0.35328 | 0.43256 | 0.05675 |
| L1236 | -0.1151 | -0.2368 | 0.07616 | -0.0581 | 0.06555 | 0.09369 | 0.95101 | 0.45753 | 0.21544 | 0.05071 | 0.3015 | 0.34597 | 0.32725 |
| CHLA90 | -0.0948 | -0.2358 | -0.1311 | -0.0758 | -0.0416 | 0.09353 | 0.94931 | 0.45856 | 0.25841 | 0.05143 | 0.18456 | 0.30226 | 0.38821 |
| LN340 | -0.1073 | -0.2635 | 0.00833 | -0.1005 | -0.0317 | 0.09296 | 0.94291 | 0.46247 | 0.2315 | 0.03369 | 0.47736 | 0.24612 | 0.41442 |
| 253J | -0.0003 | -0.1609 | -0.2172 | -0.0414 | 0.09856 | 0.09274 | 0.94037 | 0.46402 | 0.49913 | 0.13466 | 0.06689 | 0.38879 | 0.25023 |
| COLO680N | 0.16239 | -0.0969 | -0.119 | -0.0685 | 0.20667 | 0.09265 | 0.93939 | 0.46462 | 0.13247 | 0.25382 | 0.20762 | 0.31998 | 0.07711 |
| LUDLU1 | 0.02014 | -0.1817 | -0.2107 | -0.0899 | 0.03892 | 0.0926 | 0.93886 | 0.46495 | 0.44537 | 0.1057 | 0.07305 | 0.26946 | 0.39531 |
| NCIH2110 | -0.089 | -0.2481 | -0.0571 | -0.0908 | 0.0422 | 0.0925 | 0.93776 | 0.46562 | 0.27147 | 0.04281 | 0.34841 | 0.26747 | 0.3867 |
| LU165 | 0.05829 | -0.2202 | -0.1331 | -0.1402 | -0.1109 | 0.09223 | 0.93469 | 0.46752 | 0.34538 | 0.06419 | 0.18104 | 0.16825 | 0.22404 |
| SMSCTR | -0.0464 | -0.2901 | 0.00415 | -0.0892 | -0.0268 | 0.09203 | 0.93249 | 0.46887 | 0.37572 | 0.02159 | 0.48871 | 0.2712 | 0.42746 |
| SEMK2 | 0.2588 | -0.1111 | 0.06795 | -0.1017 | 0.11905 | 0.09201 | 0.93227 | 0.46901 | 0.03628 | 0.22353 | 0.32135 | 0.24346 | 0.20761 |
| UMUC7 | -0.0649 | -0.1406 | -0.2271 | -0.0843 | 0.02097 | 0.09165 | 0.92829 | 0.47147 | 0.32878 | 0.16768 | 0.05831 | 0.28241 | 0.44313 |
| SNU5 | -0.0204 | -0.2331 | -0.0632 | -0.1655 | 0.04639 | 0.09105 | 0.92154 | 0.47567 | 0.4448 | 0.05346 | 0.3332 | 0.12784 | 0.37579 |
| NCIH446 | 0.09184 | -0.0685 | 0.11084 | -0.1957 | 0.19262 | 0.09085 | 0.9193 | 0.47707 | 0.26512 | 0.31996 | 0.22417 | 0.08892 | 0.09242 |
| SUPT1 | 0.15568 | -0.0506 | -0.0788 | -0.0362 | 0.25099 | 0.09082 | 0.91902 | 0.47724 | 0.14272 | 0.36483 | 0.29519 | 0.40245 | 0.04097 |
| KU812 | 0.05479 | -0.1608 | -0.1529 | -0.1723 | 0.0767 | 0.09054 | 0.91588 | 0.47921 | 0.35423 | 0.13491 | 0.14705 | 0.11828 | 0.30021 |
| SIMA | 0.02292 | -0.1962 | 0.01244 | 0.00274 | 0.20017 | 0.09052 | 0.91572 | 0.47931 | 0.4379 | 0.08838 | 0.46621 | 0.49255 | 0.08394 |
| LCLC103H | 0.05192 | -0.1563 | -0.2355 | -0.0106 | 0.0654 | 0.09015 | 0.91151 | 0.48196 | 0.36155 | 0.1417 | 0.05168 | 0.47108 | 0.32764 |
| KMRC3 | 0.02679 | 0.06218 | -0.2684 | -0.0008 | -0.1024 | 0.0901 | 0.91102 | 0.48227 | 0.42751 | 0.33563 | 0.03109 | 0.49782 | 0.24194 |
| NCIH522 | 0.08004 | -0.1566 | -0.0694 | -0.151 | 0.17526 | 0.08988 | 0.90859 | 0.4838 | 0.2923 | 0.14123 | 0.31785 | 0.15024 | 0.1142 |
| NCIH1693 | 0.19219 | -0.1749 | -0.0287 | 0.01032 | 0.16285 | 0.08978 | 0.90749 | 0.48449 | 0.09292 | 0.11463 | 0.42238 | 0.47195 | 0.13179 |
| SKNAS | -0.1219 | -0.2323 | -0.0163 | -0.0874 | 0.04516 | 0.0897 | 0.90657 | 0.48507 | 0.20208 | 0.05413 | 0.45583 | 0.27518 | 0.379 |
| SU8686 | 0.08609 | 0.00144 | -0.2236 | -0.022 | 0.18833 | 0.08969 | 0.90647 | 0.48514 | 0.2782 | 0.49608 | 0.06128 | 0.44032 | 0.09751 |
| JHH5 | 0.06231 | -0.1237 | -0.2397 | -0.0751 | 0.06169 | 0.08957 | 0.90516 | 0.48597 | 0.3353 | 0.19849 | 0.0486 | 0.30412 | 0.33684 |
| MERO82 | 0.11855 | -0.2006 | -0.007 | 0.09012 | 0.15133 | 0.08955 | 0.90485 | 0.48616 | 0.20859 | 0.08348 | 0.48107 | 0.269 | 0.14965 |
| KMCH1 | 0.16489 | -0.0189 | -0.2309 | -0.0811 | 0.02418 | 0.08948 | 0.90406 | 0.48666 | 0.12878 | 0.44874 | 0.05525 | 0.28985 | 0.43449 |
| HA1E | -0.0627 | -0.0767 | -0.2214 | 0.04651 | 0.15055 | 0.0891 | 0.89995 | 0.48927 | 0.33426 | 0.30019 | 0.06313 | 0.3755 | 0.15091 |
| CJM | 0.02195 | -0.2717 | -0.1091 | 0.0014 | 0.01594 | 0.08839 | 0.892 | 0.49434 | 0.4405 | 0.02949 | 0.22778 | 0.49618 | 0.45672 |
| RPMI2650 | 0.06463 | -0.2731 | -0.0417 | -0.1037 | 0.04262 | 0.08833 | 0.89132 | 0.49478 | 0.32955 | 0.02879 | 0.38814 | 0.23904 | 0.38562 |
| TUHR14TKB | 0.05979 | -0.2385 | -0.1245 | -0.0771 | 0.06894 | 0.08823 | 0.89028 | 0.49545 | 0.34162 | 0.04946 | 0.19695 | 0.29917 | 0.31892 |
| NCIH2887 | 0.22108 | -0.1346 | -0.0924 | -0.0829 | 0.12174 | 0.08782 | 0.88571 | 0.49838 | 0.06343 | 0.17827 | 0.26378 | 0.28561 | 0.20234 |
| SKNEP1 | -0.1034 | -0.0392 | -0.1848 | 0.01875 | 0.18642 | 0.08763 | 0.88366 | 0.4997 | 0.23971 | 0.39455 | 0.10179 | 0.44914 | 0.09983 |
| MOLM14 | 0.12767 | -0.0962 | 0.01148 | -0.1492 | 0.21592 | 0.0876 | 0.88324 | 0.49997 | 0.191 | 0.25542 | 0.46881 | 0.15315 | 0.06811 |
| MCC142 | 0.0225 | -0.0284 | -0.0821 | 0.08791 | 0.2679 | 0.08756 | 0.88284 | 0.50023 | 0.43902 | 0.42308 | 0.28757 | 0.27403 | 0.03137 |
| NCIH2052 | 0.13446 | 0.08608 | -0.224 | -0.0211 | 0.12865 | 0.0875 | 0.88214 | 0.50069 | 0.17849 | 0.27823 | 0.06092 | 0.4427 | 0.18918 |
| RPMI8226 | -0.0031 | 0.18868 | -0.2176 | -0.0619 | 0.05304 | 0.08744 | 0.88148 | 0.50111 | 0.49158 | 0.09708 | 0.06652 | 0.33621 | 0.35869 |
| KON | 0.12437 | -0.1663 | -0.2036 | 0.01728 | 0.06168 | 0.08729 | 0.87986 | 0.50216 | 0.19727 | 0.12678 | 0.08031 | 0.45309 | 0.33687 |
| MM383 | 0.02742 | -0.0738 | -0.2102 | -0.1305 | 0.11618 | 0.08694 | 0.87601 | 0.50465 | 0.42583 | 0.30724 | 0.07361 | 0.18577 | 0.21332 |
| PGA1 | 0.15486 | -0.0016 | -0.0277 | -0.2303 | 0.12664 | 0.08688 | 0.87535 | 0.50508 | 0.14401 | 0.49566 | 0.42519 | 0.05569 | 0.19295 |
| MCF7 | 0.07891 | -0.1117 | -0.1607 | 0.09162 | 0.1796 | 0.08669 | 0.87321 | 0.50647 | 0.29496 | 0.22232 | 0.13496 | 0.26562 | 0.10845 |
| CCLFUPGI0005T | 0.21386 | -0.1116 | -0.0646 | -0.0281 | 0.17774 | 0.08666 | 0.8729 | 0.50667 | 0.07004 | 0.22253 | 0.32961 | 0.42393 | 0.11088 |
| SH4 | 0.22892 | -0.216 | 0.02901 | -0.0652 | 0.0101 | 0.08638 | 0.86982 | 0.50868 | 0.05681 | 0.06805 | 0.42158 | 0.32808 | 0.47253 |
| SHI1 | 0.06701 | -0.0496 | -0.0004 | -0.1781 | 0.22544 | 0.08624 | 0.86834 | 0.50965 | 0.32366 | 0.36754 | 0.49892 | 0.11036 | 0.05968 |
| HCS2 | 0.15017 | -0.1273 | -0.1779 | -0.0885 | 0.10082 | 0.08606 | 0.86632 | 0.51096 | 0.15153 | 0.19164 | 0.11062 | 0.27261 | 0.24533 |
| HCC2429 | 0.15194 | -0.0366 | -0.0749 | -0.0463 | 0.24643 | 0.08603 | 0.86601 | 0.51117 | 0.14866 | 0.40142 | 0.30453 | 0.37603 | 0.04392 |
| OCILY19 | -0.0465 | -0.1352 | -0.05 | -0.0849 | 0.20715 | 0.08596 | 0.86524 | 0.51167 | 0.37562 | 0.17713 | 0.36636 | 0.28094 | 0.07662 |
| PATU8902 | -0.0009 | -0.1526 | -0.0204 | -0.2261 | 0.10008 | 0.0858 | 0.86341 | 0.51287 | 0.49757 | 0.14763 | 0.44461 | 0.05909 | 0.24692 |
| 59M | -0.1306 | -0.003 | -0.2528 | -0.0377 | 0.02144 | 0.08557 | 0.86096 | 0.51448 | 0.18551 | 0.49179 | 0.03983 | 0.39846 | 0.44189 |
| HCC95 | 0.05475 | -0.1453 | -0.2034 | 0.03786 | 0.12016 | 0.08484 | 0.85291 | 0.51978 | 0.35434 | 0.15954 | 0.08049 | 0.39811 | 0.20542 |
| SNU886 | -0.0431 | -0.1165 | -0.2198 | 0.0747 | 0.10119 | 0.08453 | 0.84947 | 0.52205 | 0.38428 | 0.21263 | 0.06457 | 0.30498 | 0.24452 |
| HCC827GR5 | 0.07276 | -0.1889 | -0.1077 | -0.0799 | 0.14825 | 0.08444 | 0.84847 | 0.52272 | 0.30966 | 0.09682 | 0.23073 | 0.29253 | 0.15467 |
| ABC1 | -0.1607 | -0.1368 | -0.078 | 0.04583 | 0.12474 | 0.08411 | 0.84485 | 0.52512 | 0.13507 | 0.17434 | 0.29707 | 0.37725 | 0.19656 |
| SEKI | 0.00915 | -0.208 | 0.03385 | -0.1289 | 0.13953 | 0.08404 | 0.8441 | 0.52562 | 0.47513 | 0.07574 | 0.4087 | 0.18864 | 0.1695 |
| SLR20 | 0.18589 | -0.1242 | -0.0103 | -0.211 | 0.05578 | 0.0839 | 0.84254 | 0.52666 | 0.10048 | 0.19768 | 0.47188 | 0.07276 | 0.35172 |
| SKNO1 | 0.12171 | -0.139 | 0.07583 | -0.1475 | 0.17596 | 0.08368 | 0.84021 | 0.52822 | 0.20239 | 0.17041 | 0.30228 | 0.15589 | 0.11326 |
| ASPC1 | -0.0721 | -0.2614 | -0.0736 | 0.0193 | -0.0367 | 0.08365 | 0.83982 | 0.52848 | 0.3112 | 0.03482 | 0.30758 | 0.44765 | 0.40107 |
| HT55 | 0.03931 | -0.1186 | -0.2065 | -0.1067 | 0.09329 | 0.08363 | 0.83958 | 0.52864 | 0.39428 | 0.2085 | 0.0773 | 0.2328 | 0.26187 |
| LMSU | 0.03044 | -0.1529 | 0.03593 | 0.09499 | 0.19368 | 0.08304 | 0.83319 | 0.53291 | 0.41776 | 0.14712 | 0.4032 | 0.25809 | 0.0912 |
| SNU61 | 0.09498 | -0.1843 | -0.1898 | -0.0782 | -0.0209 | 0.08297 | 0.83234 | 0.53348 | 0.25811 | 0.10241 | 0.0958 | 0.29661 | 0.44345 |
| HCC461 | 0.21649 | -0.1641 | -0.0978 | 0.0129 | 0.08094 | 0.08287 | 0.83131 | 0.53418 | 0.06758 | 0.12993 | 0.25191 | 0.46495 | 0.29017 |
| CORL311 | 0.07764 | -0.2424 | -0.0029 | 0.02124 | 0.11665 | 0.08258 | 0.82809 | 0.53634 | 0.29797 | 0.04664 | 0.49222 | 0.4424 | 0.21239 |
| D458 | 0.10569 | -0.0393 | 0.0136 | -0.2756 | 0.02426 | 0.08243 | 0.82653 | 0.53739 | 0.23492 | 0.39431 | 0.46304 | 0.02763 | 0.43428 |
| L363 | 0.01351 | -0.1241 | -0.119 | -0.0817 | 0.19491 | 0.08234 | 0.82553 | 0.53806 | 0.46329 | 0.1978 | 0.2078 | 0.28838 | 0.08979 |
| TFK1 | 0.02764 | -0.1412 | -0.2139 | -0.1108 | -0.036 | 0.08226 | 0.82467 | 0.53865 | 0.42523 | 0.16662 | 0.07003 | 0.2243 | 0.40296 |
| NCCMPNST2C1 | -0.1266 | -0.0975 | -0.0704 | 0.11387 | 0.16391 | 0.08224 | 0.82444 | 0.5388 | 0.193 | 0.25246 | 0.31539 | 0.21798 | 0.13021 |
| COGE352 | 0.13384 | -0.1247 | -0.2039 | 0.01684 | 0.09242 | 0.08217 | 0.8236 | 0.53937 | 0.17962 | 0.1967 | 0.07998 | 0.45427 | 0.26382 |
| OCUG1 | 0.06875 | -0.1918 | -0.201 | -0.0217 | -0.0294 | 0.08143 | 0.8156 | 0.54479 | 0.31941 | 0.09333 | 0.08309 | 0.44127 | 0.42047 |
| HT115 | -0.0049 | -0.1136 | -0.2409 | -0.0695 | 0.02045 | 0.08137 | 0.81494 | 0.54524 | 0.48657 | 0.21858 | 0.04771 | 0.3176 | 0.44453 |
| SAOS2 | 0.03832 | -0.1695 | -0.1868 | -0.0642 | 0.08464 | 0.08135 | 0.81474 | 0.54537 | 0.3969 | 0.1221 | 0.09937 | 0.33068 | 0.28157 |
| UMUC6 | 0.09163 | -0.1457 | -0.0161 | -0.0667 | 0.21447 | 0.08132 | 0.81437 | 0.54563 | 0.2656 | 0.15894 | 0.45618 | 0.32444 | 0.06946 |
| HMY1 | -0.0211 | -0.2009 | -0.0135 | -0.0709 | 0.15605 | 0.08116 | 0.81258 | 0.54684 | 0.4427 | 0.08321 | 0.46335 | 0.31414 | 0.14214 |
| OE33 | -0.0356 | -0.1781 | -0.1743 | -0.1147 | -0.0218 | 0.08107 | 0.81166 | 0.54747 | 0.40415 | 0.11036 | 0.11555 | 0.21627 | 0.44088 |
| SNUC5 | 0.18993 | -0.1649 | -0.0344 | -0.1561 | 0.07168 | 0.08099 | 0.81075 | 0.54809 | 0.09559 | 0.12873 | 0.40734 | 0.14206 | 0.31228 |
| BICR10 | -0.094 | -0.2088 | 0.00364 | 0.04618 | 0.10221 | 0.08076 | 0.80826 | 0.54979 | 0.26039 | 0.07499 | 0.49011 | 0.37635 | 0.24233 |
| HS944T | -0.029 | -0.0241 | -0.1577 | -0.0607 | 0.21501 | 0.08061 | 0.80663 | 0.5509 | 0.42156 | 0.43461 | 0.13953 | 0.33933 | 0.06895 |
| MDST8 | -0.2316 | -0.0553 | 0.0037 | -0.0018 | 0.1126 | 0.08053 | 0.80581 | 0.55146 | 0.05467 | 0.35304 | 0.48994 | 0.49516 | 0.22055 |
| P4E6 | 0.09263 | -0.1688 | -0.1785 | -0.1066 | -0.0573 | 0.08016 | 0.80176 | 0.55424 | 0.26336 | 0.12308 | 0.10987 | 0.23297 | 0.34792 |
| JHOM1 | 0.13597 | -0.0609 | -0.0746 | -0.23 | 0.06094 | 0.08005 | 0.80055 | 0.55507 | 0.17579 | 0.33888 | 0.3053 | 0.05591 | 0.33872 |
| P30OHK | 0.14215 | -0.0515 | -0.1588 | -0.0434 | 0.18364 | 0.07989 | 0.79886 | 0.55623 | 0.16495 | 0.36272 | 0.13783 | 0.38347 | 0.10328 |
| TE1 | 0.06881 | -0.2135 | -0.0704 | -0.1186 | 0.10248 | 0.07975 | 0.79727 | 0.55732 | 0.31926 | 0.07034 | 0.31531 | 0.20841 | 0.24175 |
| NP5 | 0.11022 | -0.0576 | 0.12852 | -0.1339 | 0.19535 | 0.07899 | 0.78904 | 0.563 | 0.22544 | 0.34711 | 0.1894 | 0.17943 | 0.08929 |
| A375 | 0.06952 | -0.0519 | -0.0088 | -0.1847 | 0.20114 | 0.07897 | 0.7888 | 0.56317 | 0.31751 | 0.36159 | 0.47599 | 0.10193 | 0.0829 |
| KKU213 | -0.0015 | -0.1848 | -0.1871 | -0.08 | -0.0536 | 0.07884 | 0.78743 | 0.56412 | 0.49601 | 0.10188 | 0.09897 | 0.29238 | 0.35721 |
| KMS27 | 0.00947 | -0.1353 | -0.1608 | 0.02721 | 0.16217 | 0.07869 | 0.78577 | 0.56527 | 0.47425 | 0.17696 | 0.13486 | 0.42638 | 0.13279 |
| SNU626 | -0.0405 | -0.1728 | -0.1716 | -0.1095 | -0.0639 | 0.07859 | 0.78468 | 0.56603 | 0.39126 | 0.11751 | 0.11923 | 0.22692 | 0.33141 |
| NCIH1648 | 0.19651 | 0.05986 | -0.0984 | -0.1481 | 0.07105 | 0.07856 | 0.78435 | 0.56626 | 0.08798 | 0.34143 | 0.25067 | 0.155 | 0.31379 |
| 639V | 0.03697 | -0.1023 | -0.2458 | -0.028 | 0.0478 | 0.0781 | 0.77936 | 0.56972 | 0.40043 | 0.24215 | 0.04434 | 0.42418 | 0.37216 |
| SUM52PE | 0.06854 | -0.0169 | -0.1392 | 0.11998 | 0.20943 | 0.0778 | 0.7761 | 0.57199 | 0.3199 | 0.4541 | 0.17012 | 0.20577 | 0.07434 |
| BT549 | 0.108 | -0.1655 | -0.049 | 0.06599 | 0.16915 | 0.07759 | 0.77383 | 0.57358 | 0.23007 | 0.12795 | 0.36898 | 0.32618 | 0.12264 |
| JHH2 | -0.1391 | 0.10804 | -0.1915 | 0.06087 | 0.13492 | 0.07751 | 0.77302 | 0.57414 | 0.17031 | 0.22998 | 0.09378 | 0.33889 | 0.17767 |
| RERFLCAI | -0.0046 | -0.0378 | -0.1794 | -0.0917 | 0.17476 | 0.07751 | 0.77296 | 0.57418 | 0.48743 | 0.39819 | 0.10868 | 0.26554 | 0.11488 |
| MYLA | 0.04151 | -0.2003 | -0.0225 | -0.1224 | 0.13299 | 0.07709 | 0.76848 | 0.57732 | 0.3885 | 0.08382 | 0.43908 | 0.20114 | 0.18116 |
| HG3 | -0.0071 | -0.1091 | -0.2314 | -0.0823 | -0.0281 | 0.07693 | 0.76678 | 0.57851 | 0.48073 | 0.22786 | 0.05482 | 0.28697 | 0.42398 |
| HOTHC | 0.10842 | -0.1341 | -0.0443 | -0.1692 | 0.14437 | 0.07681 | 0.76543 | 0.57946 | 0.22919 | 0.17909 | 0.38113 | 0.12258 | 0.16115 |
| HMC18 | 0.23884 | -0.0324 | 0.06013 | -0.0404 | 0.15301 | 0.07677 | 0.76503 | 0.57974 | 0.04919 | 0.41247 | 0.34075 | 0.39154 | 0.14694 |
| ICC9 | 0.10714 | -0.1788 | -0.1771 | 0.0674 | -0.0019 | 0.07671 | 0.76433 | 0.58023 | 0.23185 | 0.10943 | 0.11177 | 0.32271 | 0.49486 |
| NCIH2122 | -0.1623 | -0.1937 | 0.01283 | -0.0186 | 0.03179 | 0.07669 | 0.7642 | 0.58033 | 0.13267 | 0.09113 | 0.46515 | 0.44948 | 0.41418 |

| | | | | | | | | | | | | | |
|---|---|---|---|---|---|---|---|---|---|---|---|---|---|
| SF295 | -0.0469 | -0.1711 | -0.1233 | -0.035 | 0.12847 | 0.07651 | 0.76223 | 0.58171 | 0.37446 | 0.1199 | 0.19935 | 0.40578 | 0.1895 |
| SKMEL2 | -0.1116 | -0.1109 | -0.1197 | -0.1156 | 0.10706 | 0.07651 | 0.76221 | 0.58173 | 0.22253 | 0.22412 | 0.2063 | 0.21443 | 0.23202 |
| EFO21 | 0.1298 | -0.1589 | -0.0884 | -0.1327 | 0.11483 | 0.07638 | 0.76083 | 0.5827 | 0.18703 | 0.13775 | 0.27286 | 0.18171 | 0.21604 |
| LU135 | 0.05419 | -0.0885 | -0.1165 | 0.02137 | 0.2203 | 0.07622 | 0.75911 | 0.5839 | 0.35576 | 0.27263 | 0.21267 | 0.44207 | 0.06412 |
| SKLMS1 | 0.04187 | -0.2364 | -0.1263 | 0.05329 | -0.0089 | 0.07613 | 0.75816 | 0.58458 | 0.38758 | 0.05101 | 0.19366 | 0.35806 | 0.47571 |
| OSRC2 | 0.18462 | -0.1817 | -0.0511 | -0.1287 | 0.04062 | 0.07571 | 0.75361 | 0.58779 | 0.10206 | 0.10579 | 0.36375 | 0.18912 | 0.39086 |
| MM485 | 0.04031 | -0.0973 | -0.1967 | -0.0116 | -0.1681 | 0.07565 | 0.75295 | 0.58825 | 0.39167 | 0.25302 | 0.08781 | 0.46861 | 0.12407 |
| PATU8988S | 0.01257 | -0.1144 | -0.1628 | -0.1791 | -0.0038 | 0.07553 | 0.75165 | 0.58918 | 0.46584 | 0.21697 | 0.13184 | 0.10916 | 0.48968 |
| TM31 | -0.1265 | -0.0328 | -0.1909 | 0.02556 | 0.12998 | 0.07528 | 0.74891 | 0.59111 | 0.19324 | 0.41154 | 0.09448 | 0.4308 | 0.1867 |
| LN443 | 0.06701 | -0.1472 | 0.03039 | 0.0911 | 0.18076 | 0.07526 | 0.74869 | 0.59127 | 0.32365 | 0.15634 | 0.41789 | 0.26678 | 0.10695 |
| DMS53 | 0.17761 | -0.1232 | -0.0685 | -0.0934 | 0.14866 | 0.07518 | 0.74792 | 0.59182 | 0.11106 | 0.19952 | 0.32004 | 0.2617 | 0.154 |
| RERFGC1B | -0.2073 | -0.0473 | -0.0316 | 0.11606 | 0.09725 | 0.07517 | 0.74776 | 0.59193 | 0.0765 | 0.37344 | 0.41473 | 0.21355 | 0.2531 |
| COLO684 | -0.0434 | -0.2426 | -0.0129 | -0.0999 | -0.124 | 0.07503 | 0.74629 | 0.59298 | 0.38362 | 0.04655 | 0.46484 | 0.24725 | 0.19806 |
| NP2 | 0.0506 | -0.1484 | -0.1143 | -0.0046 | 0.17721 | 0.07488 | 0.74462 | 0.59416 | 0.36494 | 0.15437 | 0.21702 | 0.48759 | 0.1116 |
| MDAMB436 | -0.013 | -0.1596 | -0.0067 | -0.2098 | 0.07114 | 0.07477 | 0.74344 | 0.595 | 0.46473 | 0.13662 | 0.48175 | 0.074 | 0.31358 |
| RT11284 | -0.0734 | -0.0522 | -0.1293 | -0.0334 | 0.20134 | 0.07469 | 0.74261 | 0.59559 | 0.30819 | 0.36094 | 0.18789 | 0.40988 | 0.08269 |
| UPMD1 | -0.0018 | -0.2293 | 0.01464 | -0.113 | 0.08331 | 0.0746 | 0.7416 | 0.59631 | 0.49514 | 0.05648 | 0.46023 | 0.21981 | 0.28464 |
| HOKUG | 0.08425 | -0.1242 | -0.1847 | -0.1309 | 0.00754 | 0.07448 | 0.74035 | 0.5972 | 0.28246 | 0.19752 | 0.10201 | 0.18499 | 0.47949 |
| EW16 | -0.091 | -0.1155 | -0.0875 | -0.0961 | 0.15144 | 0.07433 | 0.7387 | 0.59837 | 0.26706 | 0.21465 | 0.27494 | 0.25562 | 0.14946 |
| NO11 | 0.04741 | 0.06974 | -0.1499 | 0.13581 | 0.19564 | 0.07419 | 0.73721 | 0.59943 | 0.37316 | 0.31698 | 0.15194 | 0.17607 | 0.08896 |
| U343 | 0.04044 | -0.0858 | -0.0174 | 0.12875 | 0.20675 | 0.07407 | 0.73594 | 0.60035 | 0.39131 | 0.27881 | 0.45286 | 0.18897 | 0.07703 |
| RDES | -0.0352 | -0.0354 | -0.1412 | 0.02583 | 0.21777 | 0.07385 | 0.73361 | 0.60201 | 0.40506 | 0.40453 | 0.16655 | 0.43009 | 0.0664 |
| EJM | 0.09526 | -0.1662 | -0.0778 | -0.0829 | 0.15416 | 0.07375 | 0.73254 | 0.60277 | 0.25749 | 0.12686 | 0.2977 | 0.28569 | 0.14512 |
| WERIRB1 | 0.00304 | -0.1383 | -0.153 | -0.1302 | 0.08572 | 0.07324 | 0.72704 | 0.60671 | 0.49174 | 0.17173 | 0.14691 | 0.18622 | 0.27906 |
| G292CLONEA141B1 | -0.1316 | 0.10988 | -0.1328 | 0.184 | 0.09742 | 0.07307 | 0.72526 | 0.60799 | 0.18375 | 0.22614 | 0.18148 | 0.10283 | 0.25274 |
| LN428 | 0.08961 | -0.2362 | -0.0073 | -0.0794 | 0.08305 | 0.07304 | 0.72491 | 0.60824 | 0.27017 | 0.05111 | 0.48016 | 0.29385 | 0.28524 |
| SHP77 | 0.15291 | -0.0039 | -0.0476 | 0.02973 | 0.23186 | 0.07303 | 0.7248 | 0.60832 | 0.14711 | 0.48944 | 0.37276 | 0.41964 | 0.05446 |
| LPS6 | 0.15559 | -0.1483 | -0.0838 | 0.14008 | 0.04057 | 0.073 | 0.72452 | 0.60852 | 0.14286 | 0.15451 | 0.28351 | 0.16853 | 0.39099 |
| HCC2935 | 0.07022 | 0.10155 | -0.043 | 0.04882 | 0.25591 | 0.07287 | 0.72306 | 0.60957 | 0.31582 | 0.24376 | 0.3847 | 0.36951 | 0.03797 |
| U2904 | -0.0452 | -0.1503 | -0.1381 | 0.01821 | 0.13438 | 0.07283 | 0.72262 | 0.60989 | 0.37888 | 0.15136 | 0.17207 | 0.45057 | 0.17863 |
| COLO800 | 0.06475 | -0.1233 | -0.0469 | -0.2379 | -0.0256 | 0.07274 | 0.72167 | 0.61057 | 0.32924 | 0.19932 | 0.37451 | 0.04987 | 0.43071 |
| EW22 | -0.2402 | -0.0379 | -0.1012 | 0.0052 | -0.0753 | 0.07271 | 0.72141 | 0.61076 | 0.04818 | 0.39811 | 0.24453 | 0.48586 | 0.30347 |
| TC205 | 0.00709 | -0.0998 | -0.0537 | -0.1617 | 0.17016 | 0.07246 | 0.71872 | 0.61269 | 0.48073 | 0.24763 | 0.35713 | 0.13351 | 0.12122 |
| COGN305 | 0.16902 | -0.0458 | -0.1042 | -0.043 | 0.18732 | 0.07245 | 0.71863 | 0.61275 | 0.12283 | 0.37722 | 0.23801 | 0.38474 | 0.09873 |
| KE39 | -0.0722 | -0.1366 | -0.1891 | -0.0743 | -0.0672 | 0.07225 | 0.71645 | 0.61433 | 0.31109 | 0.17465 | 0.09663 | 0.30605 | 0.32311 |
| NMB | -0.0654 | -0.1894 | -0.079 | 0.02019 | 0.11112 | 0.07216 | 0.7155 | 0.61501 | 0.32759 | 0.09622 | 0.29468 | 0.44524 | 0.22359 |
| MCC26 | -0.0957 | -0.1169 | -0.0411 | 0.12091 | 0.14469 | 0.07207 | 0.71407 | 0.61604 | 0.25649 | 0.21182 | 0.38972 | 0.20396 | 0.16061 |
| KMS26 | -0.056 | -0.0728 | -0.1816 | -0.1121 | 0.09678 | 0.07198 | 0.71353 | 0.61643 | 0.3512 | 0.30951 | 0.1059 | 0.22149 | 0.25414 |
| J82 | -0.0436 | 0.0311 | -0.2316 | 0.04297 | 0.13435 | 0.07181 | 0.71172 | 0.61774 | 0.38303 | 0.41601 | 0.05465 | 0.3847 | 0.17869 |
| SNU685 | -0.1264 | -0.0633 | -0.1589 | -0.0261 | 0.12893 | 0.07172 | 0.71083 | 0.61838 | 0.19343 | 0.33295 | 0.13778 | 0.42949 | 0.18865 |
| NCIH1573 | 0.01449 | -0.2382 | 0.07551 | -0.1396 | -0.0715 | 0.07171 | 0.71073 | 0.61845 | 0.46064 | 0.04967 | 0.30305 | 0.16944 | 0.31262 |
| SNU201 | 0.10756 | -0.1503 | 0.01087 | -0.0642 | 0.18652 | 0.07162 | 0.70973 | 0.61918 | 0.23097 | 0.15137 | 0.47047 | 0.33058 | 0.09971 |
| NCIH1703 | 0.20427 | -0.0632 | -0.091 | 0.0869 | 0.12051 | 0.07148 | 0.70826 | 0.62024 | 0.07959 | 0.33299 | 0.26711 | 0.27634 | 0.20475 |
| NCIH526 | -0.082 | -0.0779 | 0.02678 | 0.02958 | 0.21325 | 0.07138 | 0.70722 | 0.621 | 0.28766 | 0.29723 | 0.42752 | 0.42004 | 0.07062 |
| CHLA10 | 0.17506 | -0.1331 | -0.0418 | -0.0408 | 0.15909 | 0.07118 | 0.705 | 0.6226 | 0.11447 | 0.18088 | 0.38774 | 0.39045 | 0.13745 |
| BFTC909 | -0.0132 | -0.1243 | -0.2303 | 0.02814 | -0.0345 | 0.07087 | 0.70171 | 0.62499 | 0.46415 | 0.19749 | 0.05569 | 0.42391 | 0.40695 |
| SNU1196 | -0.1113 | -0.1623 | -0.1506 | 0.03324 | 0.0242 | 0.07074 | 0.7003 | 0.62601 | 0.22319 | 0.13259 | 0.15078 | 0.41031 | 0.43444 |
| OACM51 | 0.05711 | -0.1593 | 0.06125 | 0.03123 | 0.17688 | 0.07071 | 0.70003 | 0.62621 | 0.34835 | 0.13714 | 0.33796 | 0.41566 | 0.11202 |
| JHH7 | 0.1646 | -0.2101 | 0.04159 | 0.00643 | 0.06884 | 0.07069 | 0.69982 | 0.62637 | 0.12919 | 0.07363 | 0.38831 | 0.48252 | 0.31918 |
| D283MED | 0.06113 | -0.1974 | -0.0726 | -0.1414 | 0.06591 | 0.0705 | 0.69784 | 0.6278 | 0.33825 | 0.08701 | 0.30998 | 0.16616 | 0.32637 |
| PANC0504 | 0.11361 | -0.1045 | -0.0534 | -0.1729 | 0.13435 | 0.07016 | 0.69416 | 0.63048 | 0.2185 | 0.23747 | 0.3578 | 0.11737 | 0.17869 |
| HEC6 | 0.08211 | -0.2357 | 0.06519 | 0.02714 | 0.06073 | 0.07013 | 0.69389 | 0.63068 | 0.28744 | 0.05151 | 0.32815 | 0.42657 | 0.33926 |
| SNU739 | 0.00223 | -0.0524 | -0.1795 | 0.01062 | 0.18 | 0.07006 | 0.69308 | 0.63127 | 0.49394 | 0.36023 | 0.10857 | 0.47113 | 0.10792 |
| ES5 | -0.1948 | -0.1289 | 0.01271 | -0.0321 | 0.05183 | 0.06954 | 0.68754 | 0.63531 | 0.08995 | 0.18868 | 0.46546 | 0.41328 | 0.36178 |
| NCIH838 | -0.1115 | -0.0688 | -0.1413 | 0.06155 | 0.14243 | 0.06804 | 0.67172 | 0.64689 | 0.22291 | 0.31921 | 0.16641 | 0.3372 | 0.16447 |
| MPP89 | -0.0147 | -0.0845 | -0.2118 | -0.0271 | 0.09972 | 0.06796 | 0.67083 | 0.64754 | 0.46007 | 0.28182 | 0.07202 | 0.42659 | 0.2477 |
| MOLM13 | 0.21818 | -0.044 | 0.09958 | -0.0787 | 0.11275 | 0.06786 | 0.66979 | 0.64831 | 0.06603 | 0.38192 | 0.248 | 0.2954 | 0.22026 |
| GAMG | -0.0804 | -0.0267 | -0.1368 | 0.21431 | 0.03877 | 0.06774 | 0.66853 | 0.64924 | 0.29153 | 0.42765 | 0.17426 | 0.06961 | 0.3957 |
| LU65 | 0.14592 | -0.0946 | -0.0088 | -0.1683 | 0.13235 | 0.06771 | 0.6682 | 0.64948 | 0.15855 | 0.25888 | 0.47607 | 0.1239 | 0.18233 |
| TCCPAN2 | -0.0555 | -0.223 | 0.02823 | -0.0268 | 0.07093 | 0.06769 | 0.66796 | 0.64965 | 0.35234 | 0.06173 | 0.42365 | 0.42739 | 0.31409 |
| MONOMAC1 | -0.0374 | -0.1163 | -0.1981 | -0.0424 | 0.06974 | 0.06768 | 0.66781 | 0.64976 | 0.39922 | 0.2131 | 0.0862 | 0.3863 | 0.31698 |
| PANC1 | 0.13216 | -0.1194 | -0.0918 | -0.1309 | 0.11858 | 0.06741 | 0.66498 | 0.65185 | 0.18267 | 0.20696 | 0.26533 | 0.18502 | 0.20855 |
| MM1S | -0.0076 | -0.077 | -0.1175 | 0.00708 | 0.20441 | 0.06739 | 0.66476 | 0.65201 | 0.4792 | 0.29937 | 0.21067 | 0.48073 | 0.07944 |
| SKMM2 | -0.0978 | -0.0871 | -0.0783 | -0.0314 | 0.17745 | 0.06738 | 0.6647 | 0.65206 | 0.25187 | 0.27596 | 0.2965 | 0.41531 | 0.11127 |
| 8505C | 0.07563 | 0.06315 | 0.00331 | -0.0827 | 0.24227 | 0.06735 | 0.66439 | 0.65228 | 0.30276 | 0.33322 | 0.49099 | 0.28595 | 0.04675 |
| TC106 | -0.1453 | -0.029 | -0.1484 | -0.041 | 0.12007 | 0.06728 | 0.66362 | 0.65285 | 0.15964 | 0.4215 | 0.15441 | 0.38998 | 0.20561 |
| NZM3 | -0.0646 | -0.1437 | -0.0767 | -0.1843 | -0.0273 | 0.06728 | 0.66357 | 0.65288 | 0.32953 | 0.16226 | 0.30021 | 0.10247 | 0.42611 |
| ONDA9 | 0.03104 | -0.1787 | 0.02224 | -0.068 | 0.15696 | 0.06702 | 0.66092 | 0.65484 | 0.41615 | 0.10966 | 0.43972 | 0.3212 | 0.14073 |
| HCC44 | -0.0704 | 0.01134 | -0.1954 | 0.00337 | 0.15269 | 0.06666 | 0.65712 | 0.65764 | 0.31527 | 0.46918 | 0.08924 | 0.49084 | 0.14746 |
| ESS1 | 0.09198 | -0.0377 | -0.2153 | -0.0133 | 0.10628 | 0.06665 | 0.65694 | 0.65777 | 0.26482 | 0.39854 | 0.06871 | 0.46376 | 0.23366 |
| NCIH292 | 0.10466 | -0.2061 | -0.1247 | -0.009 | 0.02265 | 0.0666 | 0.65646 | 0.65813 | 0.2371 | 0.07771 | 0.1967 | 0.47543 | 0.43862 |
| RH30 | 0.18525 | -0.1865 | 0.04534 | -0.0629 | -0.0842 | 0.06627 | 0.65298 | 0.6607 | 0.10127 | 0.09979 | 0.37854 | 0.33374 | 0.28251 |
| TC138 | -0.1758 | -0.1231 | -0.0713 | 0.0524 | 0.05118 | 0.06604 | 0.65058 | 0.66247 | 0.11344 | 0.19978 | 0.31312 | 0.36032 | 0.36346 |
| NCIH716 | 0.20469 | 0.05841 | -0.0285 | -0.0201 | 0.16326 | 0.06577 | 0.64764 | 0.66465 | 0.07914 | 0.34507 | 0.42281 | 0.44559 | 0.13117 |
| SW1573 | 0.05798 | -0.2114 | -0.1357 | -0.031 | -0.0056 | 0.06572 | 0.64712 | 0.66503 | 0.34615 | 0.07237 | 0.17618 | 0.41622 | 0.48487 |
| ONS76 | -0.0301 | -0.096 | -0.0567 | -0.1071 | 0.18155 | 0.06561 | 0.64596 | 0.66589 | 0.41855 | 0.25595 | 0.34941 | 0.23204 | 0.10594 |
| RCCFG2 | 0.00421 | -0.2182 | 0.04595 | 0.08808 | 0.04279 | 0.06549 | 0.64468 | 0.66684 | 0.48854 | 0.06601 | 0.37694 | 0.27364 | 0.38516 |
| KYSE220 | -0.0433 | -0.1035 | -0.147 | -0.098 | 0.11327 | 0.06527 | 0.64241 | 0.66852 | 0.38393 | 0.23963 | 0.15675 | 0.25153 | 0.21919 |
| HDMYZ | 0.00874 | -0.1487 | 0.17192 | -0.0294 | 0.1048 | 0.06524 | 0.64205 | 0.66879 | 0.47623 | 0.15401 | 0.11876 | 0.42062 | 0.2368 |
| SNU478 | 0.13926 | -0.1892 | -0.0418 | -0.073 | 0.09391 | 0.06519 | 0.64162 | 0.66911 | 0.16997 | 0.0964 | 0.38786 | 0.30912 | 0.2605 |
| JVE253 | -0.1451 | -0.0662 | -0.0941 | -0.0959 | 0.10531 | 0.06511 | 0.64076 | 0.66974 | 0.15987 | 0.32572 | 0.26003 | 0.25611 | 0.23572 |
| D423MG | 0.03805 | -0.0748 | -0.2194 | 0.00469 | 0.09101 | 0.0649 | 0.63851 | 0.67141 | 0.3976 | 0.30464 | 0.06492 | 0.48723 | 0.26701 |
| PANC1005 | 0.08455 | 0.02145 | -0.2291 | 0.02291 | 0.08719 | 0.06487 | 0.63822 | 0.67163 | 0.28176 | 0.44184 | 0.05667 | 0.43792 | 0.27568 |
| UACC257 | 0.0324 | -0.2399 | -0.0316 | 0.0218 | -0.1196 | 0.06462 | 0.63555 | 0.67361 | 0.41255 | 0.04843 | 0.41462 | 0.44091 | 0.20644 |
| MIAPACA2 | 0.07246 | -0.1542 | -0.0898 | -0.1006 | 0.12711 | 0.0645 | 0.63435 | 0.6745 | 0.31037 | 0.1451 | 0.2698 | 0.24582 | 0.19206 |
| HSOS1 | 0.07314 | -0.0912 | -0.0036 | 0.08854 | 0.19872 | 0.06439 | 0.63319 | 0.67537 | 0.30873 | 0.26664 | 0.49021 | 0.27261 | 0.08553 |
| EFM19 | 0.21359 | -0.0826 | -0.0174 | -0.0417 | 0.13481 | 0.06433 | 0.63254 | 0.67584 | 0.0703 | 0.28623 | 0.45282 | 0.3879 | 0.17787 |
| SNU601 | 0.0346 | -0.2584 | 0.02776 | -0.0187 | -0.0129 | 0.06433 | 0.63251 | 0.67586 | 0.40672 | 0.03649 | 0.42492 | 0.44921 | 0.46484 |
| NIHOVCAR3 | -0.091 | -0.0684 | -0.1802 | -0.0247 | 0.10418 | 0.06405 | 0.62953 | 0.67808 | 0.26695 | 0.32033 | 0.1077 | 0.43304 | 0.23812 |
| SCS214 | -0.0056 | -0.1055 | -0.1641 | -0.0449 | 0.13176 | 0.06396 | 0.62864 | 0.67875 | 0.4848 | 0.23534 | 0.12995 | 0.37979 | 0.18342 |
| KP2 | 0.02457 | -0.1949 | -0.068 | -0.0728 | 0.10011 | 0.06281 | 0.61659 | 0.68771 | 0.43346 | 0.08978 | 0.32127 | 0.30957 | 0.24686 |
| NCIH2126 | 0.06449 | -0.0563 | -0.1612 | -0.1616 | -0.0153 | 0.06257 | 0.61402 | 0.68963 | 0.32987 | 0.35032 | 0.13421 | 0.13363 | 0.4585 |
| NCIH1373 | -0.003 | -0.1068 | 0.01304 | -0.0641 | 0.20107 | 0.06236 | 0.61182 | 0.69127 | 0.49178 | 0.23256 | 0.46456 | 0.33074 | 0.08298 |
| KYM1 | -0.0111 | -0.1136 | -0.0482 | 0.06995 | 0.18281 | 0.06223 | 0.61052 | 0.69224 | 0.46978 | 0.2186 | 0.37118 | 0.31647 | 0.10434 |
| NCIH1355 | -0.0403 | 0.02665 | -0.2037 | -0.002 | 0.14038 | 0.0619 | 0.60704 | 0.69483 | 0.39181 | 0.42788 | 0.08022 | 0.49451 | 0.16801 |
| SIHA | 0.00296 | -0.1834 | -0.1008 | 0.02115 | 0.10015 | 0.06158 | 0.60371 | 0.69732 | 0.49195 | 0.10359 | 0.24545 | 0.44266 | 0.24678 |
| HCC2998 | -0.1469 | -0.1109 | 0.08485 | 0.07096 | 0.07039 | 0.06146 | 0.60243 | 0.69827 | 0.15691 | 0.22412 | 0.28107 | 0.31401 | 0.31541 |
| LNZ308 | 0.09289 | -0.0729 | -0.0992 | 0.03095 | 0.19165 | 0.06082 | 0.59577 | 0.70325 | 0.26277 | 0.30942 | 0.2488 | 0.41641 | 0.09355 |
| NCIH2286 | -0.0155 | -0.1506 | -0.1031 | -0.0929 | 0.1061 | 0.06064 | 0.59388 | 0.70466 | 0.45783 | 0.15086 | 0.24036 | 0.26273 | 0.23404 |
| KYO1 | 0.10357 | -0.2034 | 0.02705 | -0.1253 | 0.04536 | 0.06055 | 0.59301 | 0.70531 | 0.23942 | 0.08049 | 0.42682 | 0.19556 | 0.37848 |
| VCAP | 0.01794 | -0.0746 | -0.1546 | 0.02687 | 0.16479 | 0.06054 | 0.59286 | 0.70543 | 0.45132 | 0.30518 | 0.14446 | 0.42728 | 0.12892 |
| RPE1SS51 | 0.00388 | -0.1058 | -0.1433 | -0.1178 | 0.09644 | 0.06047 | 0.59215 | 0.70596 | 0.48946 | 0.23466 | 0.16291 | 0.21013 | 0.25488 |
| LPS27 | 0.106 | -0.1334 | -0.0078 | 0.12318 | 0.11567 | 0.06044 | 0.59187 | 0.70616 | 0.23426 | 0.18033 | 0.47872 | 0.19955 | 0.21433 |
| OPM2 | -0.0072 | 0.00472 | -0.0886 | 0.01134 | 0.23038 | 0.06026 | 0.58998 | 0.70758 | 0.48034 | 0.48716 | 0.2724 | 0.46917 | 0.05563 |
| KMS18 | -0.0772 | 0.10203 | -0.1076 | 0.0599 | 0.19571 | 0.05981 | 0.58527 | 0.7111 | 0.29896 | 0.24271 | 0.231 | 0.34132 | 0.08888 |
| RERFLCAD1 | -0.0377 | -0.0605 | -0.116 | -0.0551 | 0.17765 | 0.05967 | 0.58379 | 0.7122 | 0.39864 | 0.33981 | 0.21366 | 0.35353 | 0.11101 |
| HS294T | 0.16991 | -0.1159 | -0.0958 | -0.097 | 0.06642 | 0.0595 | 0.58199 | 0.71355 | 0.12157 | 0.21386 | 0.25638 | 0.25356 | 0.32511 |
| COLO824 | 0.10859 | -0.1179 | -0.1813 | -0.0458 | -0.0082 | 0.05939 | 0.58084 | 0.71441 | 0.22882 | 0.20983 | 0.10624 | 0.37743 | 0.47768 |
| A375SKINCJ3 | -0.0798 | -0.0217 | -0.0951 | -0.1232 | 0.14621 | 0.05903 | 0.57716 | 0.71716 | 0.29295 | 0.44114 | 0.25789 | 0.1996 | 0.15807 |
| JVM3 | -0.0792 | -0.1507 | 0.02949 | -0.1141 | 0.09539 | 0.05902 | 0.57706 | 0.71723 | 0.29435 | 0.15066 | 0.4203 | 0.21744 | 0.2572 |
| RHJT | 0.04985 | -0.1236 | -0.1201 | -0.11 | 0.10746 | 0.05841 | 0.57069 | 0.722 | 0.36686 | 0.19867 | 0.20547 | 0.226 | 0.2312 |
| MERO41 | 0.11349 | 0.01017 | -0.0061 | 0.13569 | 0.17229 | 0.05816 | 0.56808 | 0.72395 | 0.21875 | 0.47235 | 0.48339 | 0.17629 | 0.11825 |
| JM1 | -0.0059 | -0.0738 | -0.1656 | -0.0765 | 0.11797 | 0.05803 | 0.56678 | 0.72492 | 0.48404 | 0.30722 | 0.12767 | 0.3007 | 0.20975 |
| SKNMM | -0.0686 | -0.1115 | 0.0164 | -0.1768 | 0.07573 | 0.0579 | 0.56547 | 0.7259 | 0.31971 | 0.2228 | 0.45546 | 0.11212 | 0.30251 |
| NCIH1437 | -0.0442 | -0.107 | -0.0293 | -0.1121 | 0.15158 | 0.05735 | 0.55972 | 0.7302 | 0.38144 | 0.23222 | 0.4208 | 0.22166 | 0.14924 |
| MERO83 | 0.01803 | -0.1258 | -0.1648 | 0.0234 | -0.1296 | 0.05673 | 0.55329 | 0.735 | 0.45106 | 0.19451 | 0.12895 | 0.4366 | 0.18735 |
| MKN28 | -0.0894 | -0.0374 | -0.1136 | -0.1289 | 0.09933 | 0.05619 | 0.54769 | 0.73919 | 0.27055 | 0.39923 | 0.21845 | 0.18865 | 0.24857 |
| EWS502 | -0.0537 | -0.1084 | -0.0256 | -0.0502 | 0.16871 | 0.05572 | 0.54289 | 0.74276 | 0.35712 | 0.22932 | 0.43079 | 0.36584 | 0.12327 |
| NMCG1 | 0.16756 | -0.1505 | -0.0925 | 0.00157 | 0.03328 | 0.05524 | 0.53789 | 0.74649 | 0.1249 | 0.15102 | 0.26368 | 0.49574 | 0.41021 |

| | | | | | | | | | | | | | |
|---|---|---|---|---|---|---|---|---|---|---|---|---|---|
| JVE015 | 0.14449 | -0.0155 | 0.07842 | -0.1414 | 0.12757 | 0.0547 | 0.53231 | 0.75065 | 0.16096 | 0.458 | 0.29613 | 0.16631 | 0.1912 |
| SNU738 | -0.0127 | -0.0334 | -0.1886 | 0.00659 | 0.12702 | 0.05443 | 0.52958 | 0.75268 | 0.46542 | 0.40977 | 0.09719 | 0.48207 | 0.19223 |
| SKNFI | -0.1682 | -0.0552 | -0.1019 | -0.0292 | 0.06666 | 0.05403 | 0.52544 | 0.75577 | 0.12402 | 0.35321 | 0.24295 | 0.42105 | 0.32452 |
| COLO205 | -0.2033 | -0.0336 | -0.0743 | -0.0515 | -0.0431 | 0.05375 | 0.52264 | 0.75784 | 0.08058 | 0.40945 | 0.30591 | 0.36269 | 0.38432 |
| LNZTA3WT4 | -0.0268 | -0.0062 | -0.0399 | 0.14983 | 0.17051 | 0.05374 | 0.52245 | 0.75799 | 0.42756 | 0.48318 | 0.39283 | 0.15208 | 0.12073 |
| YAPC | 0.029 | -0.1404 | -0.1578 | -0.0833 | -0.0026 | 0.05362 | 0.5212 | 0.75891 | 0.42159 | 0.16789 | 0.13945 | 0.28471 | 0.49285 |
| SNU869 | -0.1205 | -0.0695 | -0.0986 | -0.1348 | 0.01009 | 0.05354 | 0.52046 | 0.75947 | 0.20467 | 0.3176 | 0.25024 | 0.17782 | 0.47257 |
| MERO14 | -0.1841 | -0.0637 | -0.1054 | -0.0225 | -0.0035 | 0.05348 | 0.51979 | 0.75996 | 0.10275 | 0.33196 | 0.23558 | 0.43908 | 0.49052 |
| SNU719 | 0.05069 | -0.1006 | -0.1997 | -0.0075 | -0.0309 | 0.05337 | 0.51871 | 0.76077 | 0.36471 | 0.24582 | 0.0845 | 0.47968 | 0.41641 |
| AML193 | -0.0364 | 0.06289 | -0.1433 | -0.062 | 0.15804 | 0.05308 | 0.51572 | 0.76298 | 0.40193 | 0.33385 | 0.1629 | 0.33619 | 0.13906 |
| CME1 | 0.07097 | -0.1242 | -0.0219 | -0.0022 | 0.17212 | 0.053 | 0.51493 | 0.76357 | 0.31399 | 0.19756 | 0.44064 | 0.49395 | 0.11848 |
| GI1 | 0.05404 | -0.0647 | 0.01939 | -0.1122 | 0.18521 | 0.05299 | 0.5148 | 0.76366 | 0.35614 | 0.32944 | 0.44739 | 0.22135 | 0.10132 |
| JVE127 | -0.0711 | -0.1756 | -0.027 | -0.1106 | 0.02076 | 0.05298 | 0.51464 | 0.76378 | 0.31379 | 0.11378 | 0.42696 | 0.22467 | 0.4437 |
| HCT15 | 0.04667 | -0.1907 | 0.0721 | -0.0198 | 0.08981 | 0.05295 | 0.51437 | 0.76398 | 0.37509 | 0.09462 | 0.31125 | 0.44641 | 0.26972 |
| NCIH2087 | 0.02321 | -0.0574 | -0.079 | -0.1155 | 0.16314 | 0.05251 | 0.50987 | 0.76732 | 0.43712 | 0.34767 | 0.29471 | 0.21476 | 0.13135 |
| DBTRG05MG | 0.13688 | -0.1622 | -0.0163 | -0.0153 | 0.10199 | 0.05226 | 0.50732 | 0.7692 | 0.17415 | 0.13275 | 0.45561 | 0.45857 | 0.2428 |
| YD38 | 0.05312 | -0.0031 | -0.1516 | -0.0133 | 0.16753 | 0.05205 | 0.50515 | 0.77081 | 0.35848 | 0.49152 | 0.14918 | 0.46398 | 0.12495 |
| NCIH2023 | 0.03153 | -0.1338 | -0.0547 | -0.1188 | 0.11705 | 0.05195 | 0.50409 | 0.77159 | 0.41486 | 0.17966 | 0.3544 | 0.20801 | 0.21158 |
| UMRC3 | 0.09556 | 0.00542 | 0.03867 | -0.0499 | 0.20944 | 0.05176 | 0.50217 | 0.77301 | 0.25682 | 0.48527 | 0.39597 | 0.36676 | 0.07433 |
| KMS11 | -0.1398 | 0.04513 | -0.026 | 0.09407 | 0.14979 | 0.05126 | 0.49705 | 0.77679 | 0.16894 | 0.37908 | 0.42951 | 0.26013 | 0.15215 |
| FTC133 | 0.01441 | -0.0392 | -0.1513 | 0.02527 | 0.15862 | 0.05111 | 0.49556 | 0.77789 | 0.46085 | 0.39464 | 0.14963 | 0.43158 | 0.13817 |
| HB1119 | 0.19736 | -0.1065 | 0.04149 | -0.0826 | 0.04799 | 0.05095 | 0.49393 | 0.77909 | 0.08704 | 0.2333 | 0.38856 | 0.28623 | 0.37166 |
| NH12 | 0.05294 | -0.1276 | -0.1007 | -0.1492 | 0.01036 | 0.05048 | 0.48914 | 0.78261 | 0.35896 | 0.19116 | 0.24559 | 0.15315 | 0.47184 |
| NCIH441 | 0.12216 | -0.1875 | -0.0483 | -0.0746 | 0.00749 | 0.05047 | 0.48901 | 0.7827 | 0.20153 | 0.09851 | 0.37087 | 0.30517 | 0.47962 |
| 93T449 | -0.0831 | -0.0039 | -0.2081 | 0.05799 | -0.0005 | 0.05043 | 0.4886 | 0.78301 | 0.28509 | 0.48926 | 0.07568 | 0.34612 | 0.49875 |
| IGR1 | 0.01748 | -0.1213 | 0.02472 | -0.0417 | 0.16872 | 0.05043 | 0.48857 | 0.78303 | 0.45255 | 0.20313 | 0.43306 | 0.38815 | 0.12325 |
| NCCLMS1C1 | 0.1003 | -0.141 | 0.04181 | 0.035 | 0.12873 | 0.05032 | 0.48747 | 0.78384 | 0.24644 | 0.16693 | 0.38773 | 0.40566 | 0.18902 |
| LCAM1 | 0.06374 | -0.0911 | -0.0607 | 0.00649 | 0.17971 | 0.05019 | 0.48619 | 0.78478 | 0.33173 | 0.26689 | 0.33944 | 0.48235 | 0.10831 |
| HCC38 | 0.11965 | -0.0348 | -0.1019 | 0.05112 | 0.15686 | 0.04976 | 0.48174 | 0.78804 | 0.20642 | 0.40615 | 0.24304 | 0.3636 | 0.14089 |
| EW1 | -0.0968 | -0.1643 | 0.03681 | -0.0951 | 0.01036 | 0.04943 | 0.47837 | 0.7905 | 0.25402 | 0.12971 | 0.40086 | 0.25775 | 0.47184 |
| HEPG2 | -0.0796 | -0.1211 | -0.0989 | 0.04319 | 0.09078 | 0.04888 | 0.47281 | 0.79456 | 0.29328 | 0.20354 | 0.24945 | 0.38413 | 0.26752 |
| ISTMES2 | 0.06012 | -0.0254 | -0.1608 | 0.06848 | 0.12848 | 0.04866 | 0.47062 | 0.79616 | 0.34076 | 0.43122 | 0.13479 | 0.32007 | 0.18948 |
| KD | 0.12555 | -0.1049 | 0.04518 | -0.0604 | 0.14517 | 0.04865 | 0.47048 | 0.79626 | 0.19501 | 0.23668 | 0.37893 | 0.34002 | 0.15981 |
| ACCMESO1 | 0.01009 | -0.0746 | -0.0303 | 0.08788 | 0.17215 | 0.04849 | 0.46885 | 0.79745 | 0.47257 | 0.30514 | 0.41816 | 0.2741 | 0.11845 |
| HS936T | 0.01789 | -0.1826 | 0.0198 | 0.09064 | 0.04077 | 0.04836 | 0.46751 | 0.79842 | 0.45144 | 0.10456 | 0.4463 | 0.26783 | 0.39047 |
| LPS853 | 0.02368 | 0.00875 | -0.1097 | -0.1678 | 0.06752 | 0.04828 | 0.46675 | 0.79897 | 0.43585 | 0.47621 | 0.22658 | 0.12453 | 0.32241 |
| KNS81 | 0.16235 | -0.0961 | -0.0869 | 0.02342 | 0.08794 | 0.04799 | 0.46374 | 0.80116 | 0.13253 | 0.25572 | 0.27625 | 0.43655 | 0.27397 |
| SW1088 | 0.02947 | -0.1431 | -0.0795 | 0.10987 | 0.06876 | 0.04794 | 0.46324 | 0.80152 | 0.42033 | 0.16332 | 0.29351 | 0.22617 | 0.31938 |
| LO68 | -0.1673 | 0.03679 | 0.00349 | 0.10835 | 0.09259 | 0.04776 | 0.46147 | 0.8028 | 0.12528 | 0.40092 | 0.49051 | 0.22934 | 0.26344 |
| HLF | -0.0785 | -0.0648 | -0.082 | 0.10534 | 0.1211 | 0.04739 | 0.45763 | 0.80558 | 0.29585 | 0.3292 | 0.28771 | 0.23565 | 0.20359 |
| T98G | -0.1119 | 0.03113 | -0.0703 | 0.03384 | 0.16769 | 0.04729 | 0.45664 | 0.8063 | 0.22205 | 0.41591 | 0.31551 | 0.40871 | 0.12473 |
| SKHEP1 | 0.07846 | -0.0853 | -0.1619 | 0.09027 | 0.03036 | 0.04716 | 0.45532 | 0.80725 | 0.29602 | 0.28004 | 0.13315 | 0.26866 | 0.41797 |
| WM88 | -0.0042 | -0.181 | 0.03597 | -0.1319 | -0.0679 | 0.04693 | 0.45306 | 0.80887 | 0.48865 | 0.10662 | 0.40309 | 0.18313 | 0.3214 |
| PCI30 | -0.0477 | -0.1993 | -0.004 | -0.06 | -0.0035 | 0.04653 | 0.44895 | 0.81183 | 0.37235 | 0.08485 | 0.48901 | 0.34101 | 0.49056 |
| SNU1105 | -0.0499 | -0.0289 | -0.0377 | 0.18809 | 0.07694 | 0.04646 | 0.44821 | 0.81236 | 0.36671 | 0.42174 | 0.39848 | 0.09779 | 0.29963 |
| SUSA | 0.05877 | -0.024 | -0.2 | -0.0037 | 0.05142 | 0.04643 | 0.44799 | 0.81252 | 0.34418 | 0.43497 | 0.08409 | 0.4898 | 0.36284 |
| YKG1 | 0.02668 | -0.0186 | -0.1685 | 0.09234 | 0.10614 | 0.04625 | 0.44612 | 0.81386 | 0.42781 | 0.44963 | 0.12355 | 0.264 | 0.23397 |
| SW579 | 0.13021 | 0.01856 | -0.0523 | 0.08221 | 0.15285 | 0.04554 | 0.43897 | 0.81897 | 0.18627 | 0.44964 | 0.36049 | 0.2872 | 0.1472 |
| SF126 | -0.0955 | -0.0753 | -0.0839 | 0.13542 | 0.05343 | 0.04549 | 0.43843 | 0.81936 | 0.25687 | 0.30355 | 0.28337 | 0.17678 | 0.3577 |
| TASK1 | 0.00072 | -0.0577 | -0.1109 | -0.0792 | 0.1405 | 0.04547 | 0.43823 | 0.8195 | 0.49805 | 0.34674 | 0.22405 | 0.29422 | 0.16781 |
| LS513 | 0.00544 | -0.1455 | -0.136 | -0.0633 | -0.0043 | 0.04515 | 0.435 | 0.8218 | 0.4852 | 0.15926 | 0.17565 | 0.33292 | 0.48843 |
| SW1271 | 0.13861 | -0.07 | 0.0121 | 0.05247 | 0.14131 | 0.04512 | 0.43471 | 0.82201 | 0.1711 | 0.31645 | 0.46713 | 0.36014 | 0.16639 |
| BFTC905 | -0.0139 | -0.0767 | -0.011 | -0.0503 | 0.17707 | 0.04505 | 0.43403 | 0.82249 | 0.46231 | 0.3003 | 0.47006 | 0.36583 | 0.11177 |
| HS729 | 0.01841 | -0.0267 | -0.1138 | 0.17315 | -0.0627 | 0.04464 | 0.42983 | 0.82546 | 0.45003 | 0.42765 | 0.21808 | 0.11706 | 0.33436 |
| SCLC22H | -0.0875 | -0.1719 | 0.03561 | -0.0033 | 0.02184 | 0.0443 | 0.42641 | 0.82788 | 0.2749 | 0.11877 | 0.40403 | 0.49093 | 0.44079 |
| MDAMB435S | -0.1334 | -0.0536 | -0.0789 | -0.0938 | 0.04703 | 0.04417 | 0.42514 | 0.82878 | 0.18047 | 0.35715 | 0.29508 | 0.26075 | 0.37415 |
| UOK101 | -0.0476 | -0.1851 | -0.0498 | -0.0576 | -0.0445 | 0.04342 | 0.41762 | 0.83406 | 0.37259 | 0.10141 | 0.36705 | 0.3472 | 0.38066 |
| NP3 | 0.04081 | 0.00544 | 0.04644 | 0.13052 | 0.1479 | 0.04341 | 0.41751 | 0.83414 | 0.39034 | 0.4852 | 0.37567 | 0.1857 | 0.15525 |
| JAR | 0.09988 | 0.03042 | -0.0556 | -0.1319 | 0.11806 | 0.04282 | 0.41153 | 0.83832 | 0.24736 | 0.41782 | 0.35229 | 0.18319 | 0.20957 |
| MHHES1 | -0.0707 | -0.1075 | 0.05946 | -0.0835 | 0.10104 | 0.04244 | 0.40779 | 0.84091 | 0.31455 | 0.23117 | 0.34242 | 0.28425 | 0.24484 |
| LK2 | -0.1171 | -0.1301 | 0.01999 | 0.06601 | 0.02262 | 0.04227 | 0.40601 | 0.84215 | 0.21152 | 0.18647 | 0.44579 | 0.32613 | 0.43869 |
| NCIH2172 | 0.12813 | -0.087 | -0.003 | -0.1271 | 0.0908 | 0.04217 | 0.40508 | 0.84279 | 0.19014 | 0.27601 | 0.49194 | 0.19214 | 0.26748 |
| JMURTK2 | 0.03274 | 0.00209 | -0.1172 | 0.13804 | 0.11112 | 0.04216 | 0.40499 | 0.84285 | 0.41165 | 0.49433 | 0.21134 | 0.1721 | 0.22358 |
| A3KAW | -0.0606 | 0.0021 | -0.0558 | -0.0388 | 0.17712 | 0.04209 | 0.4042 | 0.84339 | 0.33947 | 0.49429 | 0.35168 | 0.39568 | 0.11171 |
| PSN1 | -0.0679 | -0.1691 | -0.0009 | -0.0327 | 0.03853 | 0.04104 | 0.39372 | 0.85059 | 0.32157 | 0.12274 | 0.49766 | 0.4118 | 0.39633 |
| TC32 | -0.1278 | 0.01875 | -0.0935 | -0.0426 | 0.1041 | 0.04051 | 0.3884 | 0.85421 | 0.19073 | 0.44912 | 0.26139 | 0.38555 | 0.23829 |
| HCT116 | -0.0214 | -0.1743 | -0.0353 | 0.02628 | 0.04761 | 0.03995 | 0.38282 | 0.85799 | 0.44199 | 0.11555 | 0.40483 | 0.42887 | 0.37266 |
| SCCOHT1 | 0.09182 | -0.0981 | -0.0692 | -0.018 | 0.13075 | 0.0398 | 0.38134 | 0.85899 | 0.26517 | 0.2512 | 0.31828 | 0.45117 | 0.18527 |
| KPNYN | 0.01254 | -0.0096 | 0.00509 | -0.1342 | 0.14738 | 0.03936 | 0.37698 | 0.86191 | 0.46592 | 0.47386 | 0.48615 | 0.17898 | 0.15612 |
| HCC56 | 0.00159 | 0.00421 | -0.0925 | -0.0808 | 0.15239 | 0.03915 | 0.37481 | 0.86336 | 0.49569 | 0.48856 | 0.26358 | 0.29045 | 0.14794 |
| BE2M17 | -0.1032 | -0.0989 | -0.0488 | -0.0448 | 0.08024 | 0.03892 | 0.37254 | 0.86487 | 0.24011 | 0.24944 | 0.3697 | 0.37998 | 0.29182 |
| CCLP1 | -0.162 | -0.0184 | -0.0756 | 0.09359 | -0.0149 | 0.03859 | 0.36924 | 0.86706 | 0.13302 | 0.45006 | 0.3028 | 0.26121 | 0.45954 |
| TN2 | -0.1065 | -0.097 | -0.0189 | -0.0733 | 0.07154 | 0.03844 | 0.36776 | 0.86804 | 0.23326 | 0.25362 | 0.44871 | 0.30837 | 0.31259 |
| TCCSUP | -0.0942 | -0.0761 | -0.0722 | 0.09172 | 0.07449 | 0.03817 | 0.36505 | 0.86983 | 0.25978 | 0.30175 | 0.31099 | 0.2654 | 0.30548 |
| HSSCH2 | -0.1032 | 0.02002 | -0.1352 | 0.08794 | -0.0796 | 0.03774 | 0.36085 | 0.87258 | 0.24019 | 0.4457 | 0.1771 | 0.27396 | 0.29333 |
| SNU423 | 0.01769 | -0.0299 | -0.1812 | 0.01456 | 0.05674 | 0.03739 | 0.35737 | 0.87485 | 0.45198 | 0.41907 | 0.10632 | 0.46046 | 0.34929 |
| RERFLCAD2 | 0.05221 | -0.1021 | 0.06061 | -0.0067 | 0.13783 | 0.03726 | 0.35602 | 0.87573 | 0.36081 | 0.24249 | 0.33954 | 0.48178 | 0.17249 |
| MSTO211H | -0.0747 | 0.02214 | -0.0626 | -0.1153 | 0.10527 | 0.03715 | 0.35496 | 0.87641 | 0.30494 | 0.43998 | 0.33448 | 0.21513 | 0.23581 |
| HUO9 | -0.0563 | -0.1066 | 0.01125 | 0.04498 | 0.113 | 0.03713 | 0.35478 | 0.87653 | 0.35051 | 0.23309 | 0.46941 | 0.37946 | 0.21974 |
| BECKER | 0.19065 | 0.00674 | 0.00544 | -0.006 | 0.04965 | 0.03703 | 0.35381 | 0.87716 | 0.09473 | 0.48166 | 0.4852 | 0.48377 | 0.36738 |
| A427 | 0.11233 | -0.1353 | 0.06348 | -0.1033 | 0.00409 | 0.0361 | 0.3446 | 0.88308 | 0.22111 | 0.17699 | 0.33239 | 0.24001 | 0.48888 |
| YAMATO | 0.00974 | -0.1581 | 0.01442 | -0.1229 | -0.037 | 0.03577 | 0.34132 | 0.88516 | 0.47353 | 0.13901 | 0.46084 | 0.20015 | 0.40039 |
| RH4 | 0.08746 | -0.05 | -0.0521 | -0.0704 | 0.14031 | 0.03566 | 0.3402 | 0.88587 | 0.27506 | 0.36654 | 0.36113 | 0.31547 | 0.16812 |
| SW48 | 0.11593 | -0.0828 | -0.0638 | -0.0959 | 0.07217 | 0.03563 | 0.33987 | 0.88608 | 0.21382 | 0.28587 | 0.33169 | 0.2561 | 0.31107 |
| LP1 | 0.02747 | -0.0021 | 0.04917 | -0.1618 | 0.09476 | 0.03558 | 0.33943 | 0.88635 | 0.4257 | 0.49428 | 0.36863 | 0.13329 | 0.2586 |
| LPS141 | 0.10023 | -0.0794 | -0.1165 | -0.0539 | -0.0575 | 0.03478 | 0.33154 | 0.89131 | 0.24659 | 0.29377 | 0.21277 | 0.35641 | 0.34726 |
| SNUC1 | -0.166 | -0.0233 | -0.0577 | -0.0182 | 0.02051 | 0.03451 | 0.32888 | 0.89296 | 0.12716 | 0.43687 | 0.34698 | 0.45063 | 0.44436 |
| YH13 | -0.0504 | -0.0699 | 0.04934 | 0.06071 | 0.12137 | 0.03443 | 0.32805 | 0.89347 | 0.36554 | 0.3167 | 0.36819 | 0.33929 | 0.20305 |
| ISTMEL1 | 0.0237 | 0.10312 | -0.0165 | -0.0687 | -0.1082 | 0.0341 | 0.32476 | 0.8955 | 0.4358 | 0.24038 | 0.45533 | 0.31952 | 0.22966 |
| KMM1 | -0.0837 | -0.105 | -0.1078 | -0.0067 | 0.00463 | 0.03314 | 0.31534 | 0.90124 | 0.28371 | 0.23637 | 0.23044 | 0.4818 | 0.48742 |
| M140325 | 0.08641 | -0.0498 | -0.0382 | -0.124 | 0.09081 | 0.03265 | 0.31055 | 0.90412 | 0.27747 | 0.36691 | 0.39733 | 0.19794 | 0.26746 |
| A172 | 0.02045 | -0.0747 | -0.0837 | 0.1243 | 0.05817 | 0.03253 | 0.30938 | 0.90481 | 0.44455 | 0.30509 | 0.28374 | 0.1974 | 0.34567 |
| BT12 | -0.0496 | -0.1023 | -0.1085 | 0.04067 | 0.04909 | 0.03245 | 0.30858 | 0.90529 | 0.3674 | 0.24214 | 0.22905 | 0.39071 | 0.36882 |
| CAL120 | -0.1048 | -0.076 | -0.0664 | 0.09276 | 0.01272 | 0.03168 | 0.301 | 0.90976 | 0.23688 | 0.30195 | 0.32527 | 0.26307 | 0.46545 |
| LPS510 | 0.08805 | -0.0896 | -0.0616 | -0.0109 | -0.1224 | 0.03164 | 0.30063 | 0.90998 | 0.27373 | 0.27008 | 0.33713 | 0.47044 | 0.20101 |
| HT3 | 0.01039 | -0.0285 | 0.10868 | 0.04807 | 0.11907 | 0.03163 | 0.30046 | 0.91008 | 0.47176 | 0.42289 | 0.22864 | 0.37145 | 0.20756 |
| KP4 | 0.05414 | 0.08853 | -0.0344 | 0.01332 | 0.15826 | 0.03132 | 0.29749 | 0.9118 | 0.3559 | 0.27263 | 0.40721 | 0.46382 | 0.13872 |
| UPCISCC029A | -0.0983 | -0.0528 | -0.1001 | 0.08774 | 0.02498 | 0.03128 | 0.29711 | 0.91203 | 0.25085 | 0.35927 | 0.24687 | 0.27444 | 0.43236 |
| NCIH226 | -0.1266 | -0.0237 | 0.02111 | 0.09835 | 0.04774 | 0.03101 | 0.29444 | 0.91357 | 0.19308 | 0.43588 | 0.44277 | 0.25069 | 0.37232 |
| UMUC13 | -0.0724 | -0.0002 | -0.0918 | 0.14408 | -0.0322 | 0.031 | 0.29428 | 0.91366 | 0.31045 | 0.49953 | 0.26522 | 0.16165 | 0.41308 |
| JMSU1 | -0.0684 | -0.0057 | -0.13 | 0.11453 | -0.013 | 0.03085 | 0.29281 | 0.9145 | 0.32034 | 0.48447 | 0.18657 | 0.21663 | 0.46475 |
| U118MG | 0.01449 | -0.05 | -0.055 | 0.08969 | 0.12458 | 0.03075 | 0.29189 | 0.91503 | 0.46065 | 0.36647 | 0.35381 | 0.26998 | 0.19687 |
| COV504 | 0.06442 | -0.0964 | -0.0947 | -0.0114 | 0.08239 | 0.02974 | 0.28203 | 0.92061 | 0.33007 | 0.25501 | 0.25865 | 0.4689 | 0.28678 |
| MG63 | -0.0355 | -0.0913 | -0.0978 | -0.0689 | 0.0415 | 0.02931 | 0.27779 | 0.92295 | 0.40438 | 0.26634 | 0.25187 | 0.31894 | 0.38855 |
| LB771HNC | -0.0815 | -0.0477 | -0.1239 | 0.01196 | 0.04713 | 0.02906 | 0.27533 | 0.92431 | 0.28885 | 0.37239 | 0.19821 | 0.46749 | 0.37388 |
| KP1N | 0.11409 | -0.0867 | 0.05192 | -0.0028 | -0.0962 | 0.02861 | 0.27098 | 0.92668 | 0.21752 | 0.27676 | 0.36155 | 0.49234 | 0.25535 |
| HS683 | 0.01394 | -0.1682 | 0.00308 | 0.02552 | -0.029 | 0.02842 | 0.26912 | 0.92768 | 0.46212 | 0.12402 | 0.49162 | 0.43091 | 0.42157 |
| CHLA06ATRT | 0.0824 | -0.0606 | -0.0508 | -0.0907 | 0.09019 | 0.02777 | 0.26283 | 0.93104 | 0.28677 | 0.33955 | 0.36431 | 0.26767 | 0.26885 |
| ONE58 | -0.0696 | -0.0767 | -0.0791 | 0.10203 | -0.0075 | 0.02768 | 0.26189 | 0.93154 | 0.31744 | 0.30022 | 0.29453 | 0.24271 | 0.47955 |
| UMUC11 | 0.06433 | -0.1187 | -0.0494 | 0.07235 | 0.03664 | 0.02766 | 0.26167 | 0.93166 | 0.33029 | 0.20835 | 0.36797 | 0.31063 | 0.4013 |
| MEL290 | -0.0347 | -0.0197 | -0.114 | 0.11782 | 0.03537 | 0.02749 | 0.26001 | 0.93252 | 0.40656 | 0.44644 | 0.21771 | 0.21005 | 0.40467 |
| SKN3 | -0.0387 | -0.1341 | -0.0548 | -0.0303 | -0.0742 | 0.02679 | 0.25325 | 0.93603 | 0.39589 | 0.17917 | 0.35412 | 0.41811 | 0.30619 |
| HCE4 | -0.0648 | -0.0412 | -0.104 | -0.0089 | 0.08119 | 0.02653 | 0.2507 | 0.93733 | 0.32907 | 0.3894 | 0.23853 | 0.47582 | 0.2896 |
| TC71 | -0.1278 | -0.0313 | -0.0624 | -0.0208 | 0.0358 | 0.02576 | 0.24327 | 0.94107 | 0.19081 | 0.41556 | 0.33511 | 0.4436 | 0.40354 |
| NP8 | 0.0011 | -0.1398 | 0.04027 | -0.086 | -0.039 | 0.02516 | 0.23745 | 0.94393 | 0.497 | 0.16902 | 0.39175 | 0.27842 | 0.39501 |
| ONDA8 | -0.0577 | -0.0618 | 0.03702 | 0.11229 | -0.0567 | 0.02442 | 0.23033 | 0.94734 | 0.34696 | 0.3366 | 0.40032 | 0.22119 | 0.34929 |
| 8305C | 0.086 | -0.0585 | -0.0248 | -0.0443 | 0.11253 | 0.02434 | 0.22955 | 0.94772 | 0.27842 | 0.34486 | 0.43288 | 0.38127 | 0.2207 |
| UMRC7 | 0.03868 | -0.0192 | -0.0666 | 0.11786 | -0.0639 | 0.0235 | 0.22145 | 0.95148 | 0.39593 | 0.44803 | 0.32456 | 0.20996 | 0.33139 |
| HCC1359 | -0.0591 | -0.074 | -0.0179 | 0.08735 | 0.05142 | 0.02292 | 0.21581 | 0.95403 | 0.34328 | 0.30659 | 0.4515 | 0.27532 | 0.36285 |

| | | | | | | | | | | | | | |
|---|---|---|---|---|---|---|---|---|---|---|---|---|---|
| MELJUSO | -0.0277 | -0.0358 | -0.0353 | -0.078 | 0.10453 | 0.02275 | 0.21417 | 0.95476 | 0.42519 | 0.40355 | 0.40484 | 0.297 | 0.23738 |
| CCLFPEDS0008T | 0.04565 | -0.0268 | -0.1057 | 0.10082 | -0.002 | 0.02272 | 0.21385 | 0.9549 | 0.37773 | 0.42735 | 0.23485 | 0.24533 | 0.49462 |
| SCH | -0.0501 | -0.071 | -0.105 | -0.0303 | 0.00805 | 0.02174 | 0.20449 | 0.95896 | 0.36628 | 0.31403 | 0.2364 | 0.41804 | 0.47812 |
| NCIH1993 | -0.0031 | -0.0584 | -0.0841 | -0.065 | 0.06624 | 0.02127 | 0.1999 | 0.96089 | 0.49162 | 0.34501 | 0.28292 | 0.32874 | 0.32557 |
| SLR26 | -0.0013 | -0.0855 | -0.0813 | 0.04283 | 0.05383 | 0.02049 | 0.19243 | 0.96394 | 0.49648 | 0.27953 | 0.28936 | 0.38506 | 0.35667 |
| SKNSH | -0.0074 | 0.01721 | -0.0535 | -0.1156 | 0.03952 | 0.01946 | 0.1826 | 0.96778 | 0.47988 | 0.45329 | 0.35763 | 0.21453 | 0.39372 |
| PSS131R | -0.0452 | -0.0263 | -0.1233 | -0.0169 | -0.0087 | 0.01913 | 0.1794 | 0.96898 | 0.37878 | 0.42873 | 0.19923 | 0.45408 | 0.47644 |
| HSQ89 | 0.06363 | -0.0329 | 0.0416 | -0.1179 | 0.02426 | 0.01816 | 0.17015 | 0.97235 | 0.332 | 0.41129 | 0.38829 | 0.20983 | 0.4343 |
| RMSYM | -0.1143 | -0.0509 | 0.00225 | 0.02441 | -0.0456 | 0.01791 | 0.16773 | 0.9732 | 0.21715 | 0.36412 | 0.49388 | 0.43389 | 0.37778 |
| NCIH1792 | -0.0475 | -0.06 | -0.1026 | -0.0148 | -0.0182 | 0.01785 | 0.16718 | 0.97339 | 0.37297 | 0.34119 | 0.24142 | 0.45973 | 0.45054 |
| PL4 | -0.0765 | -0.0737 | -0.0691 | -0.0056 | -0.0051 | 0.01781 | 0.16685 | 0.97351 | 0.30064 | 0.30733 | 0.31857 | 0.48466 | 0.48624 |
| SUM1315MO2 | -0.0149 | -0.0201 | -0.0818 | 0.02723 | 0.0921 | 0.01702 | 0.15928 | 0.97607 | 0.45957 | 0.44553 | 0.28815 | 0.42634 | 0.26454 |
| LI7 | -0.0617 | 0.04844 | -0.0531 | 0.10872 | -0.0021 | 0.017 | 0.15911 | 0.97613 | 0.3369 | 0.37051 | 0.35861 | 0.22856 | 0.49422 |
| NOS1 | -0.0019 | -0.0715 | 0.02959 | -0.1069 | 0.01874 | 0.01629 | 0.15236 | 0.9783 | 0.49481 | 0.31282 | 0.42001 | 0.23226 | 0.44917 |
| MOGGUVW | -0.0618 | -0.0418 | -0.075 | -0.0462 | 0.02109 | 0.0159 | 0.14867 | 0.97945 | 0.33663 | 0.3877 | 0.30433 | 0.37617 | 0.44281 |
| SNU638 | -0.1011 | 0.03578 | -0.0692 | -0.0175 | -0.0043 | 0.01587 | 0.14839 | 0.97953 | 0.24468 | 0.4036 | 0.3183 | 0.45257 | 0.48835 |
| PACADD165 | -0.0606 | -0.0318 | -0.089 | 0.03313 | 0.00024 | 0.01369 | 0.12774 | 0.98536 | 0.33945 | 0.41409 | 0.27144 | 0.4106 | 0.49935 |
| CAL72 | -0.1101 | -0.02 | 0.02232 | -0.0039 | -0.0286 | 0.01359 | 0.12672 | 0.98562 | 0.22572 | 0.44587 | 0.43951 | 0.48937 | 0.42255 |
| WM3211 | -0.0766 | -0.0559 | -0.0322 | 0.04257 | 0.01113 | 0.01334 | 0.12442 | 0.98621 | 0.30046 | 0.35137 | 0.41296 | 0.38574 | 0.46975 |
| G402 | -0.0095 | -0.061 | 0.05296 | 0.05842 | 0.02882 | 0.01266 | 0.11799 | 0.98777 | 0.4743 | 0.33859 | 0.35891 | 0.34506 | 0.42208 |
| MEL285 | 0.0549 | -0.0588 | -0.0157 | 0.03101 | 0.05727 | 0.011 | 0.10235 | 0.99117 | 0.35395 | 0.34408 | 0.45739 | 0.41625 | 0.34795 |
| MPNST724 | -0.0526 | -0.0079 | -0.0284 | 0.08532 | -0.0162 | 0.01017 | 0.09453 | 0.99265 | 0.35988 | 0.47841 | 0.4232 | 0.27999 | 0.45597 |
| MERO95 | 0.03981 | -0.0239 | -0.048 | 0.06165 | 0.03721 | 0.00931 | 0.08642 | 0.99403 | 0.39298 | 0.43537 | 0.37176 | 0.33696 | 0.39981 |
| NCCMPNST1C1 | -0.0015 | 0.00682 | -0.0765 | -0.0198 | 0.0516 | 0.00904 | 0.08391 | 0.99443 | 0.49584 | 0.48145 | 0.30061 | 0.44619 | 0.36239 |
| HUH6 | 0.02164 | 0.06573 | -0.0599 | 0.03144 | 0.03038 | 0.00889 | 0.0825 | 0.99465 | 0.44133 | 0.32682 | 0.34143 | 0.41509 | 0.41791 |
| KINGS1 | 0.00388 | -0.0366 | -0.0179 | 0.03364 | 0.07156 | 0.00884 | 0.0821 | 0.99471 | 0.48945 | 0.40133 | 0.45136 | 0.40924 | 0.31255 |

| | | | | | | |
|---|---|---|---|---|---|---|
| average | | | | | | |
| p < 0.01 | 0.14375 | -0.3262 | -0.2432 | -0.1332 | 0.13003 | 0.31209 |
| p < 0.05 | 0.12088 | -0.2762 | -0.2235 | -0.1144 | 0.13584 | 0.25426 |
| p < 0.1 | 0.11413 | -0.2582 | -0.2095 | -0.1107 | 0.1252 | 0.2308 |
| total | 0.06959 | -0.1735 | -0.134 | -0.0781 | 0.10783 | 0.13644 |
