## Supplement Table S8B for "Single-molecule behavior and cell-growth regulation in human RTKs"

| | pR | | | | | | | p-values in t-test | | | |
|---|---|---|---|---|---|---|---|---|---|---|---|
| | EG1 | EG2 | EG3 | EG4 | determ | F-value | p-value | EG1 | EG2 | EG3 | EG4 |
| HKGZCC | -0.8062 | 0.089864 | -0.17892 | -0.03439 | 0.66612 | 23.4418 | 1.06E-10 | 8.08538E-13 | 0.26743 | 0.10689 | 0.4063 |
| SNU8 | -0.7846 | 0.141262 | -0.12281 | -0.09121 | 0.64735 | 21.5695 | 3.74E-10 | 7.83643E-12 | 0.16391 | 0.19776 | 0.26435 |
| ECGI10 | -0.7839 | 0.034428 | -0.11386 | -0.04685 | 0.59842 | 17.5098 | 7.35E-09 | 8.34781E-12 | 0.40619 | 0.21555 | 0.37332 |
| A431 | -0.7362 | 0.159402 | -0.19431 | -0.13517 | 0.59264 | 17.094 | 1.02E-08 | 5.56818E-10 | 0.13442 | 0.08816 | 0.17466 |
| BICR56 | -0.6886 | 0.238717 | -0.11441 | -0.09423 | 0.57729 | 16.0468 | 2.37E-08 | 1.62339E-08 | 0.04751 | 0.21442 | 0.25754 |
| CFPAC1 | -0.7903 | -0.02353 | -0.23812 | -0.32478 | 0.57208 | 15.7082 | 3.14E-08 | 4.39768E-12 | 0.43558 | 0.04793 | 0.01069 |
| HSC5 | -0.725 | 0.152958 | -0.17632 | -0.15071 | 0.5698 | 15.5626 | 3.54E-08 | 1.30558E-09 | 0.14446 | 0.11032 | 0.14808 |
| DOTC24510 | -0.7875 | -0.01534 | -0.1804 | -0.13019 | 0.56784 | 15.4391 | 3.93E-08 | 5.83477E-12 | 0.45788 | 0.10498 | 0.18376 |
| KYAE1 | -0.7303 | 0.074917 | -0.09424 | 0.057681 | 0.5669 | 15.3801 | 4.13E-08 | 8.79659E-10 | 0.30255 | 0.25754 | 0.34536 |
| TE6 | -0.7953 | -0.05642 | -0.23102 | -0.16786 | 0.56103 | 15.0173 | 5.61E-08 | 2.61148E-12 | 0.34857 | 0.05324 | 0.12196 |
| H413 | -0.7394 | 0.108496 | -0.08989 | -0.10115 | 0.55995 | 14.9513 | 5.93E-08 | 4.30385E-10 | 0.22663 | 0.26737 | 0.2423 |
| JHOC5 | -0.7074 | 0.171245 | -0.07085 | -0.02896 | 0.55777 | 14.8196 | 6.64E-08 | 4.63449E-09 | 0.11721 | 0.31243 | 0.42087 |
| SSP25 | -0.6971 | 0.169227 | -0.20608 | -0.04884 | 0.5567 | 14.7559 | 7.01E-08 | 9.36433E-09 | 0.12003 | 0.07553 | 0.36813 |
| KYSE140 | -0.6779 | 0.194361 | -0.07992 | -0.23361 | 0.54928 | 14.3197 | 1.02E-07 | 3.17959E-08 | 0.08811 | 0.29056 | 0.05126 |
| CAL27 | -0.758 | 0.027412 | -0.1402 | -0.09174 | 0.54754 | 14.219 | 1.12E-07 | 9.18418E-11 | 0.42506 | 0.16576 | 0.26316 |
| SF767 | -0.7029 | 0.158162 | -0.15523 | -0.13381 | 0.54051 | 13.8218 | 1.59E-07 | 6.33726E-09 | 0.13632 | 0.14087 | 0.17711 |
| ESO26 | -0.7567 | -0.00605 | -0.06939 | -0.08169 | 0.53997 | 13.792 | 1.63E-07 | 1.03408E-10 | 0.48338 | 0.31603 | 0.28638 |
| PCI6A | -0.708 | 0.118669 | -0.25534 | -0.14572 | 0.53808 | 13.6874 | 1.79E-07 | 4.44313E-09 | 0.20588 | 0.03675 | 0.1563 |
| HCC1419 | -0.7363 | 0.045623 | -0.10751 | -0.06276 | 0.52922 | 13.2088 | 2.75E-07 | 5.49917E-10 | 0.37653 | 0.22871 | 0.33251 |
| PK1 | -0.7348 | 0.061155 | -0.1915 | -0.13392 | 0.52824 | 13.1567 | 2.88E-07 | 6.19872E-10 | 0.33655 | 0.09139 | 0.17691 |
| C75 | -0.6863 | 0.069031 | -0.36062 | -0.08972 | 0.52749 | 13.1174 | 2.99E-07 | 1.88311E-08 | 0.31691 | 0.00505 | 0.26775 |
| OC316 | -0.7182 | 0.095973 | -0.0535 | -0.12729 | 0.52589 | 13.0331 | 3.23E-07 | 2.16344E-09 | 0.25366 | 0.35605 | 0.18919 |
| BPH1 | -0.6853 | 0.166095 | -0.09445 | -0.02121 | 0.52539 | 13.0073 | 3.31E-07 | 2.00353E-08 | 0.1245 | 0.25707 | 0.44188 |
| BICR22 | -0.6887 | 0.15073 | -0.03183 | -0.17147 | 0.5245 | 12.9609 | 3.45E-07 | 1.61892E-08 | 0.14805 | 0.41315 | 0.1169 |
| UWB1289 | -0.6763 | 0.182755 | -0.01025 | -0.0542 | 0.5224 | 12.8523 | 3.81E-07 | 3.51647E-08 | 0.10199 | 0.47183 | 0.35427 |
| SNU216 | -0.7081 | 0.086715 | -0.06464 | 0.006154 | 0.5223 | 12.8469 | 3.83E-07 | 4.43012E-09 | 0.27466 | 0.32781 | 0.48308 |
| NCIN87 | -0.7677 | -0.04929 | -0.23195 | -0.19625 | 0.52167 | 12.8147 | 3.95E-07 | 3.87561E-11 | 0.36695 | 0.05253 | 0.08598 |
| RERFLCSQ1 | -0.7534 | -0.01938 | -0.10646 | -0.13544 | 0.51948 | 12.7027 | 4.38E-07 | 1.364E-10 | 0.44686 | 0.23091 | 0.17418 |
| RL952 | -0.7113 | 0.09646 | -0.16777 | -0.18093 | 0.51379 | 12.4166 | 5.71E-07 | 3.54231E-09 | 0.25258 | 0.1221 | 0.1043 |
| SCABER | -0.7261 | 0.040945 | -0.22475 | -0.16581 | 0.50753 | 12.1095 | 7.63E-07 | 1.20848E-09 | 0.38885 | 0.0583 | 0.12492 |
| COV413A | -0.7223 | 0.052121 | -0.19954 | -0.13288 | 0.50747 | 12.1066 | 7.65E-07 | 1.5971E-09 | 0.35962 | 0.08237 | 0.17881 |
| A253 | -0.6614 | 0.191826 | -0.04182 | -0.04185 | 0.50571 | 12.0214 | 8.29E-07 | 8.5212E-08 | 0.09101 | 0.38654 | 0.38645 |
| UPCISCC154 | -0.7236 | 0.039535 | -0.20069 | -0.25019 | 0.50431 | 11.9542 | 8.84E-07 | 1.45022E-09 | 0.39258 | 0.08114 | 0.03985 |
| SNU1066 | -0.6562 | 0.195253 | -0.02762 | -0.04906 | 0.50126 | 11.8092 | 1.02E-06 | 1.14939E-07 | 0.0871 | 0.4245 | 0.36757 |
| UPCISCC111 | -0.6852 | 0.130926 | -0.09937 | -0.10657 | 0.49664 | 11.5931 | 1.25E-06 | 2.02532E-08 | 0.18239 | 0.24617 | 0.23068 |
| TE11 | -0.6998 | 0.093912 | -0.16812 | -0.16848 | 0.49655 | 11.5889 | 1.26E-06 | 7.79237E-09 | 0.25827 | 0.12159 | 0.12108 |
| CAL33 | -0.7011 | 0.086182 | -0.16546 | -0.15651 | 0.49332 | 11.4402 | 1.45E-06 | 7.15226E-09 | 0.27589 | 0.12542 | 0.13887 |
| UMUC10 | -0.6173 | 0.24765 | -0.11466 | -0.03464 | 0.49169 | 11.3657 | 1.56E-06 | 8.99912E-07 | 0.04146 | 0.21393 | 0.40563 |
| SKGT4 | -0.7178 | 0.011261 | -0.13525 | -0.08748 | 0.4853 | 11.0789 | 2.07E-06 | 2.21959E-09 | 0.46907 | 0.17451 | 0.2729 |
| H103 | -0.7076 | 0.036705 | -0.24148 | -0.24178 | 0.48471 | 11.0528 | 2.12E-06 | 4.5722E-09 | 0.40011 | 0.04557 | 0.04536 |
| DETROIT562 | -0.652 | 0.147921 | -0.23768 | -0.1815 | 0.48351 | 10.9997 | 2.23E-06 | 1.45669E-07 | 0.15264 | 0.04825 | 0.10358 |
| BIN67 | -0.6823 | 0.086511 | -0.21798 | -0.23811 | 0.48016 | 10.8533 | 2.58E-06 | 2.42313E-08 | 0.27513 | 0.06417 | 0.04794 |
| KYSE410 | -0.7074 | 0.004268 | -0.0845 | -0.04122 | 0.47953 | 10.8259 | 2.65E-06 | 4.65549E-09 | 0.48827 | 0.27979 | 0.38812 |
| MDAMB361 | -0.7045 | -0.01537 | -0.04116 | -0.06768 | 0.47058 | 10.4441 | 3.90E-06 | 5.65383E-09 | 0.45781 | 0.38829 | 0.32024 |
| PECAPJ15 | -0.656 | 0.124135 | -0.14445 | -0.21292 | 0.46342 | 10.1479 | 5.27E-06 | 1.16001E-07 | 0.1952 | 0.15845 | 0.06882 |
| UPCISCC074 | -0.704 | 0.019146 | -0.16608 | -0.15478 | 0.46324 | 10.1407 | 5.31E-06 | 5.86328E-09 | 0.4475 | 0.12452 | 0.14158 |
| PACADD119 | -0.6954 | -0.08478 | -0.41799 | -0.20765 | 0.46312 | 10.1358 | 5.33E-06 | 1.04249E-08 | 0.27915 | 0.00126 | 0.07394 |
| SAT | -0.7155 | -0.03741 | -0.12099 | -0.11608 | 0.46244 | 10.1079 | 5.49E-06 | 2.62789E-09 | 0.39822 | 0.20131 | 0.21106 |
| HEC1B | -0.5989 | 0.22443 | -0.11124 | 0.039261 | 0.46234 | 10.1038 | 5.51E-06 | 2.16894E-06 | 0.05857 | 0.22092 | 0.39331 |
| HARA | -0.7036 | 0.015424 | -0.16693 | -0.1374 | 0.46223 | 10.0995 | 5.54E-06 | 6.03965E-09 | 0.45767 | 0.12329 | 0.17067 |
| PCI38 | -0.6168 | 0.168011 | -0.23341 | -0.22484 | 0.46203 | 10.0913 | 5.58E-06 | 9.20875E-07 | 0.12175 | 0.05141 | 0.05823 |
| C2BBE1 | -0.6558 | 0.128346 | -0.14784 | -0.07881 | 0.46092 | 10.0463 | 5.85E-06 | 1.17723E-07 | 0.1872 | 0.15278 | 0.29322 |
| TGBC18TKB | -0.6959 | -0.01227 | -0.11926 | -0.03165 | 0.46068 | 10.0369 | 5.90E-06 | 1.00919E-08 | 0.4663 | 0.2047 | 0.41365 |
| BICR31 | -0.7015 | -0.00552 | -0.22901 | -0.29663 | 0.46035 | 10.0232 | 5.99E-06 | 6.93776E-09 | 0.48483 | 0.05483 | 0.01823 |
| KMBC2 | -0.6076 | 0.218748 | -0.14027 | -0.06999 | 0.4603 | 10.0213 | 6.00E-06 | 1.44108E-06 | 0.06348 | 0.16563 | 0.31457 |
| DOK | -0.6663 | 0.059582 | -0.16004 | 0.002349 | 0.45895 | 9.96691 | 6.35E-06 | 6.40999E-08 | 0.34053 | 0.13345 | 0.49354 |
| AU565 | -0.6576 | 0.118703 | -0.04851 | -0.07109 | 0.45691 | 9.88541 | 6.90E-06 | 1.06198E-07 | 0.20581 | 0.36898 | 0.31185 |
| TKKK | -0.7415 | -0.16169 | -0.3037 | -0.26632 | 0.45524 | 9.81905 | 7.40E-06 | 3.64892E-10 | 0.13098 | 0.01601 | 0.03078 |
| OSC19 | -0.6955 | 0.003901 | -0.15569 | -0.08979 | 0.45282 | 9.72363 | 8.17E-06 | 1.03938E-08 | 0.48928 | 0.14015 | 0.2676 |
| MCAS | -0.6149 | 0.126615 | -0.31638 | -0.14138 | 0.45121 | 9.66083 | 8.72E-06 | 1.00975E-06 | 0.19046 | 0.0126 | 0.1637 |
| SNUC4 | -0.658 | 0.111015 | -0.09844 | -0.08421 | 0.45071 | 9.64142 | 8.90E-06 | 1.03792E-07 | 0.22139 | 0.24821 | 0.28047 |
| OCUBM | -0.6941 | -0.00445 | -0.08487 | -0.20485 | 0.44789 | 9.53202 | 9.98E-06 | 1.13753E-08 | 0.48778 | 0.27893 | 0.07678 |
| RMGI | -0.679 | 0.049005 | -0.17096 | -0.11819 | 0.44738 | 9.51219 | 1.02E-05 | 2.97568E-08 | 0.3677 | 0.1176 | 0.20684 |
| TGBC1TKB | -0.5993 | 0.207089 | 0.021461 | -0.09147 | 0.44304 | 9.34682 | 1.21E-05 | 2.13168E-06 | 0.07451 | 0.4412 | 0.26377 |
| LN229 | -0.6649 | 0.068755 | -0.13405 | -0.07226 | 0.44115 | 9.27539 | 1.31E-05 | 6.97173E-08 | 0.31759 | 0.17667 | 0.30901 |
| NCIH661 | -0.6377 | 0.128931 | -0.09263 | -0.02189 | 0.44101 | 9.27007 | 1.32E-05 | 3.17566E-07 | 0.1861 | 0.26114 | 0.44004 |
| PEO1 | -0.6649 | 0.067744 | -0.19188 | -0.19864 | 0.44039 | 9.24675 | 1.35E-05 | 6.9712E-08 | 0.32009 | 0.09095 | 0.08334 |
| SNU213 | -0.6203 | 0.107644 | -0.19298 | 0.033624 | 0.43959 | 9.21675 | 1.39E-05 | 7.75619E-07 | 0.22842 | 0.08968 | 0.40834 |
| UPCISCC116 | -0.6342 | 0.100503 | -0.26681 | -0.14635 | 0.43754 | 9.14026 | 1.51E-05 | 3.8185E-07 | 0.2437 | 0.03053 | 0.15525 |
| OE21 | -0.6325 | 0.142015 | -0.10005 | -0.0735 | 0.43615 | 9.08873 | 1.60E-05 | 4.16256E-07 | 0.16261 | 0.24469 | 0.30597 |
| HCC1806 | -0.5076 | 0.326303 | -0.13271 | -0.01859 | 0.43359 | 8.99467 | 1.77E-05 | 8.41629E-05 | 0.01037 | 0.17912 | 0.44903 |
| SKMES1 | -0.6887 | -0.01246 | -0.23112 | -0.20836 | 0.43332 | 8.98494 | 1.79E-05 | 1.61861E-08 | 0.46579 | 0.05317 | 0.07324 |
| KYSE270 | -0.6489 | 0.096248 | -0.13385 | -0.17783 | 0.43263 | 8.95943 | 1.84E-05 | 1.73096E-07 | 0.25305 | 0.17704 | 0.10832 |
| CCSW1 | -0.6052 | 0.143364 | -0.22102 | -0.02502 | 0.4324 | 8.95126 | 1.85E-05 | 1.61374E-06 | 0.1603 | 0.06148 | 0.43153 |
| CAKI2 | -0.6412 | 0.085639 | -0.24587 | -0.12858 | 0.43219 | 8.94344 | 1.87E-05 | 2.62192E-07 | 0.27715 | 0.04261 | 0.18675 |
| BICR6 | -0.615 | 0.155557 | -0.15939 | -0.17709 | 0.43133 | 8.9124 | 1.93E-05 | 1.00581E-06 | 0.14036 | 0.13444 | 0.10929 |
| VMCUB1 | -0.5796 | 0.214348 | -0.14953 | -0.1249 | 0.42698 | 8.75522 | 2.29E-05 | 5.15943E-06 | 0.06749 | 0.15 | 0.19373 |
| A388 | -0.6462 | 0.021043 | -0.24523 | -0.03105 | 0.42656 | 8.74049 | 2.33E-05 | 2.00767E-07 | 0.44234 | 0.04303 | 0.41525 |
| KYSE450 | -0.6242 | 0.1325 | -0.04685 | -0.01734 | 0.42602 | 8.72101 | 2.38E-05 | 6.36711E-07 | 0.1795 | 0.37333 | 0.45242 |
| SKBR3 | -0.6667 | 0.038869 | -0.1324 | -0.11366 | 0.42569 | 8.70917 | 2.41E-05 | 6.26049E-08 | 0.39435 | 0.17968 | 0.21595 |
| JHU022 | -0.6532 | 0.016881 | -0.08485 | 0.007392 | 0.42386 | 8.6444 | 2.58E-05 | 1.35726E-07 | 0.45368 | 0.27899 | 0.47968 |
| UPCISCC040 | -0.6394 | 0.085097 | -0.18079 | -0.10268 | 0.41864 | 8.46124 | 3.16E-05 | 2.89454E-07 | 0.27841 | 0.10448 | 0.23899 |
| SKGII | -0.6405 | 0.082315 | -0.15753 | -0.08175 | 0.4178 | 8.43222 | 3.26E-05 | 2.72387E-07 | 0.28492 | 0.1373 | 0.28626 |
| TTC442 | -0.6386 | -0.01356 | -0.38387 | -0.20752 | 0.41663 | 8.39165 | 3.41E-05 | 3.01685E-07 | 0.46278 | 0.00296 | 0.07408 |
| KYSE180 | -0.5863 | 0.20038 | -0.06486 | -0.0427 | 0.41527 | 8.34489 | 3.59E-05 | 3.8406E-06 | 0.08147 | 0.32725 | 0.38423 |
| HT1376 | -0.5948 | 0.107126 | -0.31329 | -0.21517 | 0.41472 | 8.32577 | 3.67E-05 | 2.62103E-06 | 0.22951 | 0.01337 | 0.06672 |
| HS852T | -0.4251 | 0.170429 | -0.38981 | -0.40613 | 0.41218 | 8.23916 | 4.04E-05 | 0.001044251 | 0.11834 | 0.00257 | 0.00171 |
| PECAPJ49 | -0.5863 | 0.153872 | -0.20066 | -0.0184 | 0.41181 | 8.22652 | 4.09E-05 | 3.84681E-06 | 0.14301 | 0.08117 | 0.44953 |
| 647V | -0.4543 | 0.348975 | -0.15991 | -0.05993 | 0.41044 | 8.18028 | 4.31E-05 | 0.000459535 | 0.0065 | 0.13366 | 0.33965 |
| SCC9 | -0.6776 | -0.10537 | -0.07886 | -0.08489 | 0.41042 | 8.17958 | 4.31E-05 | 3.24506E-08 | 0.23323 | 0.29309 | 0.27889 |
| CAL29 | -0.5288 | 0.285969 | -0.06463 | -0.0173 | 0.40997 | 8.16421 | 4.39E-05 | 3.94392E-05 | 0.02205 | 0.32782 | 0.45255 |
| UPCISCC026 | -0.5982 | 0.153117 | -0.11838 | -0.01505 | 0.40846 | 8.11339 | 4.64E-05 | 2.24328E-06 | 0.14421 | 0.20645 | 0.4587 |
| NCIH28 | -0.4677 | 0.208569 | -0.37626 | -0.06779 | 0.40792 | 8.09514 | 4.74E-05 | 0.000307738 | 0.07304 | 0.00354 | 0.31998 |
| NCIH322 | -0.6011 | 0.049434 | -0.36997 | -0.19339 | 0.40662 | 8.05181 | 4.98E-05 | 1.95705E-06 | 0.36658 | 0.00409 | 0.08921 |
| SNU308 | -0.6074 | 0.108614 | -0.11636 | 0.025782 | 0.40625 | 8.03932 | 5.05E-05 | 1.45128E-06 | 0.22638 | 0.21049 | 0.42947 |
| SCC4 | -0.611 | 0.097989 | -0.22403 | -0.24073 | 0.40573 | 8.02219 | 5.14E-05 | 1.22309E-06 | 0.2492 | 0.05891 | 0.04609 |
| ICC15 | -0.6375 | 0.050367 | -0.14443 | -0.04398 | 0.40465 | 7.98623 | 5.35E-05 | 3.20513E-07 | 0.36416 | 0.15849 | 0.38084 |
| TE9 | -0.5385 | 0.257616 | -0.10208 | -0.05993 | 0.40385 | 7.95997 | 5.52E-05 | 2.73718E-05 | 0.03545 | 0.24029 | 0.33964 |
| SLR24 | -0.5575 | 0.219712 | -0.14331 | -0.08216 | 0.40332 | 7.94227 | 5.63E-05 | 1.30073E-05 | 0.06263 | 0.16038 | 0.28529 |
| KURAMOCHI | -0.6024 | 0.127514 | -0.2002 | -0.09917 | 0.40299 | 7.93146 | 5.70E-05 | 1.83853E-06 | 0.18876 | 0.08166 | 0.2466 |
| JOPACA1 | -0.5431 | 0.119169 | -0.3799 | -0.22723 | 0.40261 | 7.919 | 5.78E-05 | 2.29932E-05 | 0.20489 | 0.00325 | 0.05626 |
| HUPT3 | -0.5821 | 0.11045 | -0.31209 | -0.15682 | 0.40203 | 7.89969 | 5.90E-05 | 4.61893E-06 | 0.22256 | 0.01368 | 0.13839 |
| UMUC7 | -0.5465 | 0.245239 | -0.04977 | 0.013554 | 0.40099 | 7.86565 | 6.13E-05 | 2.01104E-05 | 0.04303 | 0.36571 | 0.46278 |
| YD15 | -0.5729 | 0.172812 | -0.16466 | -0.00678 | 0.40055 | 7.85146 | 6.23E-05 | 6.86196E-06 | 0.11505 | 0.12659 | 0.48135 |
| CW9019 | -0.4827 | 0.136218 | -0.35506 | -0.39218 | 0.39955 | 7.8188 | 6.47E-05 | 0.000192756 | 0.17278 | 0.0057 | 0.00243 |
| KYSE30 | -0.5933 | 0.145178 | 0.008826 | -0.00223 | 0.39953 | 7.81809 | 6.47E-05 | 2.80437E-06 | 0.15722 | 0.47575 | 0.49386 |
| H357 | -0.6077 | 0.045709 | -0.29379 | -0.06792 | 0.3981 | 7.77141 | 6.82E-05 | 1.4363E-06 | 0.37631 | 0.01919 | 0.31964 |
| ZR751 | -0.5848 | 0.124118 | -0.1651 | 0.031389 | 0.3971 | 7.7392 | 7.08E-05 | 4.09804E-06 | 0.19523 | 0.12594 | 0.41434 |
| OVMANA | -0.5899 | 0.110388 | 0.052112 | 0.050329 | 0.39654 | 7.72104 | 7.23E-05 | 3.26644E-06 | 0.22269 | 0.35964 | 0.36426 |
| 769P | -0.5822 | 0.16811 | -0.1499 | -0.08416 | 0.39651 | 7.72015 | 7.23E-05 | 4.59244E-06 | 0.12161 | 0.1494 | 0.2806 |
| SKGT2 | -0.5781 | 0.122413 | -0.27862 | -0.1665 | 0.39333 | 7.618 | 8.13E-05 | 5.49196E-06 | 0.19853 | 0.02504 | 0.12391 |
| C33A | -0.5401 | 0.180165 | -0.26719 | -0.12396 | 0.39094 | 7.54211 | 8.86E-05 | 2.58275E-05 | 0.10528 | 0.03034 | 0.19554 |
| PANC0403 | -0.6271 | -0.02464 | -0.34864 | -0.16154 | 0.38988 | 7.50844 | 9.21E-05 | 5.50037E-07 | 0.43256 | 0.00654 | 0.1312 |
| NCIH1975 | -0.6375 | 0.006553 | -0.07752 | -0.03875 | 0.38986 | 7.50801 | 9.21E-05 | 3.19414E-07 | 0.48199 | 0.29629 | 0.39467 |
| SW954 | -0.6435 | 0.002212 | -0.19626 | -0.09991 | 0.38909 | 7.48374 | 9.48E-05 | 2.31891E-07 | 0.49392 | 0.08597 | 0.245 |
| TEN | -0.6452 | -0.0047 | -0.10842 | -0.073 | 0.38853 | 7.46606 | 9.67E-05 | 2.11675E-07 | 0.48708 | 0.2268 | 0.30721 |
| KYSE510 | -0.5672 | 0.17942 | -0.14407 | -0.10205 | 0.38604 | 7.38795 | 0.000106 | 8.74087E-06 | 0.10625 | 0.1591 | 0.24033 |

| | | | | | | | | | | | |
|---|---|---|---|---|---|---|---|---|---|---|---|
| EKVX | -0.6018 | 0.043541 | -0.30977 | -0.1367 | 0.38383 | 7.31949 | 0.000114 | 1.88999E-06 | 0.382 | 0.01429 | 0.17192 |
| SW1116 | -0.5853 | 0.139763 | -0.09856 | -0.01607 | 0.38372 | 7.31594 | 0.000115 | 4.00602E-06 | 0.16651 | 0.24795 | 0.45591 |
| ECC2 | -0.5871 | 0.07832 | -0.29748 | -0.12724 | 0.38367 | 7.31432 | 0.000115 | 3.70689E-06 | 0.29438 | 0.01795 | 0.18928 |
| SNU1076 | -0.4954 | 0.304094 | 0.01885 | -0.01387 | 0.38365 | 7.31386 | 0.000115 | 0.000127128 | 0.0159 | 0.44831 | 0.46192 |
| MFE319 | -0.5701 | 0.145862 | -0.21155 | -0.10416 | 0.38095 | 7.23073 | 0.000127 | 7.72535E-06 | 0.15607 | 0.07013 | 0.23581 |
| LN319 | -0.6607 | -0.05822 | -0.1708 | -0.22919 | 0.38071 | 7.22324 | 0.000128 | 8.89563E-08 | 0.34398 | 0.11783 | 0.05469 |
| SCC25 | -0.5305 | 0.199214 | -0.10769 | -0.22878 | 0.3803 | 7.21086 | 0.00013 | 3.7008E-05 | 0.08272 | 0.22832 | 0.05501 |
| LB1047RCC | -0.5828 | 0.133813 | -0.16548 | -0.16204 | 0.3795 | 7.18633 | 0.000134 | 4.47543E-06 | 0.17711 | 0.12539 | 0.13045 |
| PCI15A | -0.5899 | 0.050919 | -0.32792 | -0.2204 | 0.37875 | 7.16344 | 0.000137 | 3.264E-06 | 0.36273 | 0.01004 | 0.06202 |
| BHY | -0.5728 | 0.166324 | -0.08287 | -0.06065 | 0.37855 | 7.15725 | 0.000138 | 6.90869E-06 | 0.12417 | 0.28362 | 0.33783 |
| PLCPRF5 | -0.6074 | 0.089756 | -0.13643 | -0.09688 | 0.37844 | 7.15411 | 0.000139 | 1.45545E-06 | 0.26767 | 0.1724 | 0.25166 |
| SLR23 | -0.4068 | 0.360068 | -0.14695 | 0.07015 | 0.37657 | 7.09725 | 0.000148 | 0.001687085 | 0.00511 | 0.15425 | 0.31416 |
| VP229 | -0.6288 | 0.021496 | -0.10522 | -0.20885 | 0.37616 | 7.08506 | 0.00015 | 5.04931E-07 | 0.4411 | 0.23354 | 0.07276 |
| IM95 | -0.6246 | 0.019413 | -0.20634 | -0.09416 | 0.37599 | 7.07969 | 0.000151 | 6.23455E-07 | 0.44678 | 0.07527 | 0.25771 |
| HCC1937 | -0.3837 | 0.420357 | -0.02182 | -0.0004 | 0.3751 | 7.05301 | 0.000156 | 0.002973599 | 0.00119 | 0.44022 | 0.49891 |
| JIMT1 | -0.5471 | 0.197993 | -0.13021 | -0.0758 | 0.37433 | 7.02999 | 0.00016 | 1.96463E-05 | 0.08406 | 0.18373 | 0.30043 |
| COLO679 | -0.5756 | 0.06171 | -0.27032 | -0.3244 | 0.37324 | 6.99719 | 0.000167 | 6.11298E-06 | 0.33515 | 0.02881 | 0.01077 |
| ICC2 | -0.6372 | -0.0427 | -0.28128 | -0.20432 | 0.36899 | 6.87094 | 0.000193 | 3.25602E-07 | 0.3842 | 0.02392 | 0.07733 |
| SNU1041 | -0.6374 | -0.09927 | -0.36228 | -0.27694 | 0.36763 | 6.83087 | 0.000203 | 3.21361E-07 | 0.24638 | 0.00486 | 0.02577 |
| BXPC3 | -0.5942 | 0.067526 | -0.23367 | -0.11582 | 0.36655 | 6.79925 | 0.00021 | 2.6966E-06 | 0.32063 | 0.05121 | 0.21158 |
| H157 | -0.5247 | 0.182566 | -0.1248 | -0.24013 | 0.36599 | 6.78293 | 0.000215 | 4.58529E-05 | 0.10222 | 0.19392 | 0.04651 |
| PECAPJ34CLONEC12 | -0.6172 | 0.03279 | -0.10537 | -0.10536 | 0.36378 | 6.71843 | 0.000232 | 9.01729E-07 | 0.41058 | 0.23322 | 0.23324 |
| HCA1 | -0.525 | 0.216565 | -0.12735 | -0.08187 | 0.36322 | 6.70226 | 0.000236 | 4.54183E-05 | 0.06544 | 0.18907 | 0.28596 |
| YD8 | -0.6155 | 0.036368 | -0.11254 | -0.14489 | 0.3619 | 6.66393 | 0.000247 | 9.84262E-07 | 0.40101 | 0.21824 | 0.1577 |
| TE10 | -0.5389 | 0.097304 | -0.33592 | -0.20647 | 0.36103 | 6.63908 | 0.000255 | 2.7014E-05 | 0.25072 | 0.00854 | 0.07513 |
| PC14 | -0.6322 | -0.03478 | -0.2303 | -0.15894 | 0.36001 | 6.60974 | 0.000264 | 4.23491E-07 | 0.40526 | 0.05381 | 0.13513 |
| TT1TKB | -0.5217 | 0.156621 | -0.01448 | 0.150417 | 0.35769 | 6.54344 | 0.000285 | 5.10777E-05 | 0.1387 | 0.46025 | 0.14855 |
| HSC2 | -0.5937 | 0.065833 | -0.14705 | -0.05452 | 0.35733 | 6.53314 | 0.000289 | 2.7566E-06 | 0.32483 | 0.15408 | 0.35343 |
| HT115 | -0.6004 | -0.00949 | -0.30084 | -0.29584 | 0.35547 | 6.48023 | 0.000308 | 2.02482E-06 | 0.47393 | 0.01688 | 0.01849 |
| WM1799 | -0.544 | 0.06642 | -0.33325 | -0.08357 | 0.35546 | 6.47998 | 0.000308 | 2.21816E-05 | 0.32337 | 0.00902 | 0.28197 |
| JHH4 | -0.6394 | -0.06344 | -0.17409 | -0.23775 | 0.3552 | 6.47268 | 0.000311 | 2.8962E-07 | 0.33082 | 0.11331 | 0.0482 |
| OVISE | -0.5538 | 0.127975 | -0.15296 | 0.00685 | 0.3544 | 6.45011 | 0.000319 | 1.5095E-05 | 0.18789 | 0.14447 | 0.48117 |
| OVCAR5 | -0.51 | 0.061176 | -0.37233 | -0.04434 | 0.3518 | 6.37721 | 0.000348 | 7.73467E-05 | 0.3365 | 0.00388 | 0.3799 |
| TDOTT | -0.4708 | 0.280527 | -0.09087 | 0.032576 | 0.35162 | 6.37202 | 0.000351 | 0.000279704 | 0.02423 | 0.26514 | 0.41115 |
| KKU213 | -0.3701 | 0.102646 | -0.42902 | 0.075134 | 0.35112 | 6.35825 | 0.000356 | 0.004077264 | 0.23906 | 0.00094 | 0.30203 |
| M059K | -0.4951 | 0.00803 | -0.36028 | -0.46852 | 0.34975 | 6.31987 | 0.000373 | 0.00012861 | 0.47793 | 0.00508 | 0.0003 |
| SKRC20 | -0.5938 | 0.047345 | -0.10316 | -0.20873 | 0.34848 | 6.28473 | 0.00039 | 2.74415E-06 | 0.37202 | 0.23795 | 0.07288 |
| LC1SQ | -0.5994 | 0.020783 | -0.12491 | -0.04433 | 0.34717 | 6.24844 | 0.000407 | 2.1209E-06 | 0.44305 | 0.1937 | 0.37992 |
| L33 | -0.5924 | 0.060844 | -0.14687 | -0.12619 | 0.34704 | 6.24507 | 0.000409 | 2.92102E-06 | 0.33734 | 0.15439 | 0.19126 |
| RH28 | -0.4578 | 0.065573 | -0.36999 | 0.05832 | 0.34704 | 6.24489 | 0.000409 | 0.000414665 | 0.32548 | 0.00409 | 0.34373 |
| KP3 | -0.5028 | 0.150427 | -0.27924 | -0.05572 | 0.34464 | 6.17911 | 0.000443 | 9.91902E-05 | 0.14854 | 0.02477 | 0.35038 |
| PCI4B | -0.5554 | 0.107708 | -0.1571 | -0.01593 | 0.34211 | 6.10999 | 0.000482 | 1.41657E-05 | 0.22828 | 0.13795 | 0.45628 |
| PA1 | -0.4284 | -0.13032 | -0.60442 | -0.44419 | 0.34154 | 6.09467 | 0.000491 | 0.000954451 | 0.18351 | 1.7E-06 | 0.00062 |
| HUCCT1 | -0.5661 | 0.106872 | -0.14075 | -0.0909 | 0.3414 | 6.09092 | 0.000493 | 9.13064E-06 | 0.23004 | 0.16479 | 0.26506 |
| ICC108 | -0.5739 | 0.088996 | -0.0665 | -0.0542 | 0.34106 | 6.08167 | 0.000499 | 6.58421E-06 | 0.26941 | 0.32319 | 0.35427 |
| SW837 | -0.513 | 0.052465 | -0.38985 | -0.12879 | 0.34075 | 6.07317 | 0.000504 | 6.97152E-05 | 0.35873 | 0.00257 | 0.18637 |
| ANGMCSS | -0.4056 | 0.220005 | -0.25711 | -0.29238 | 0.33962 | 6.04282 | 0.000523 | 0.001736745 | 0.06237 | 0.03573 | 0.01968 |
| T3M4 | -0.5417 | 0.132827 | -0.09506 | 0.031336 | 0.33947 | 6.03864 | 0.000526 | 2.42523E-05 | 0.1789 | 0.25569 | 0.41448 |
| TE5 | -0.555 | 0.10848 | -0.19573 | -0.16483 | 0.33909 | 6.0285 | 0.000532 | 1.43811E-05 | 0.22666 | 0.08656 | 0.12634 |
| UPCISCC152 | -0.5973 | 0.011502 | -0.13285 | -0.23248 | 0.33807 | 6.00113 | 0.00055 | 2.33602E-06 | 0.46841 | 0.17886 | 0.05212 |
| RBE | -0.4814 | 0.221344 | -0.18369 | -0.07658 | 0.33589 | 5.94283 | 0.000591 | 0.00020094 | 0.0612 | 0.10081 | 0.29855 |
| UPCISCC131 | -0.5994 | 0.007558 | -0.18352 | -0.14987 | 0.33431 | 5.90094 | 0.000622 | 2.11388E-06 | 0.47923 | 0.10103 | 0.14944 |
| BICR16 | -0.5561 | 0.097066 | -0.19952 | -0.16662 | 0.33424 | 5.89892 | 0.000624 | 1.37895E-05 | 0.25124 | 0.08239 | 0.12374 |
| SF172 | -0.5441 | 0.080876 | -0.27897 | -0.18932 | 0.33357 | 5.88123 | 0.000638 | 2.20748E-05 | 0.28831 | 0.02489 | 0.09396 |
| C80 | -0.3949 | 0.221214 | -0.22067 | -0.3136 | 0.33277 | 5.86003 | 0.000655 | 0.002269198 | 0.06131 | 0.06179 | 0.01329 |
| SNU761 | -0.5851 | 0.039087 | -0.18347 | -0.15493 | 0.33204 | 5.84078 | 0.00067 | 4.05054E-06 | 0.39377 | 0.10108 | 0.14134 |
| JHH1 | -0.5653 | 0.092045 | -0.13566 | -0.10822 | 0.33188 | 5.83653 | 0.000674 | 9.45711E-06 | 0.26247 | 0.17377 | 0.22721 |
| 9505BIK | -0.5878 | -0.07392 | -0.3687 | -0.29713 | 0.33186 | 5.83621 | 0.000674 | 3.58453E-06 | 0.30497 | 0.00421 | 0.01807 |
| UACC893 | -0.5733 | 0.064444 | -0.0372 | -0.10262 | 0.3309 | 5.81077 | 0.000696 | 6.76263E-06 | 0.3283 | 0.39878 | 0.23911 |
| OVCAR8 | -0.4838 | 0.165157 | -0.27546 | -0.12399 | 0.32993 | 5.78543 | 0.000718 | 0.000186098 | 0.12586 | 0.02642 | 0.19549 |
| HUPT4 | -0.573 | -0.031 | -0.32817 | -0.12532 | 0.32961 | 5.77698 | 0.000725 | 6.843E-06 | 0.41539 | 0.00999 | 0.19293 |
| CAL78 | -0.4313 | 0.126733 | -0.40236 | -0.21173 | 0.32859 | 5.75039 | 0.00075 | 0.000881616 | 0.19024 | 0.00189 | 0.06996 |
| SNU1544 | -0.5022 | 0.168916 | -0.05373 | -0.21283 | 0.32799 | 5.73489 | 0.000764 | 0.000101107 | 0.12047 | 0.35548 | 0.06891 |
| ES4 | -0.5937 | 0.005859 | -0.17933 | -0.20048 | 0.32794 | 5.73346 | 0.000766 | 2.75299E-06 | 0.48389 | 0.10637 | 0.08136 |
| UMUC9 | -0.4129 | 0.29505 | -0.139 | -0.12354 | 0.32737 | 5.71878 | 0.00078 | 0.00144077 | 0.01876 | 0.16786 | 0.19635 |
| ONCODG1 | -0.5133 | 0.083264 | -0.28521 | -0.02121 | 0.32715 | 5.71291 | 0.000785 | 6.88752E-05 | 0.28269 | 0.02234 | 0.44189 |
| LOUNH91 | -0.5971 | -0.02551 | -0.22785 | -0.24931 | 0.32654 | 5.69724 | 0.000801 | 2.35206E-06 | 0.4302 | 0.05576 | 0.04041 |
| OVCA420 | -0.6048 | -0.03661 | -0.15946 | -0.12118 | 0.32606 | 5.68483 | 0.000813 | 1.64244E-06 | 0.40035 | 0.13433 | 0.20094 |
| NCIH2052 | -0.4795 | -0.06366 | -0.52149 | -0.30764 | 0.32589 | 5.68049 | 0.000818 | 0.000212998 | 0.33026 | 5.2E-05 | 0.01488 |
| JHOS2 | -0.5309 | 0.144809 | -0.12819 | -0.14166 | 0.32583 | 5.67885 | 0.000819 | 3.64906E-05 | 0.15784 | 0.1875 | 0.16322 |
| YUHOIN0650 | -0.4958 | 0.092861 | -0.33336 | -0.08263 | 0.32522 | 5.66311 | 0.000835 | 0.000125297 | 0.26063 | 0.009 | 0.28418 |
| EW8 | -0.492 | -0.01777 | -0.46075 | -0.16711 | 0.32483 | 5.65301 | 0.000846 | 0.000142329 | 0.45127 | 0.00038 | 0.12304 |
| TGBC52TKB | -0.5759 | -0.08155 | -0.2633 | -0.0301 | 0.32446 | 5.64353 | 0.000856 | 6.03526E-06 | 0.28673 | 0.03234 | 0.41782 |
| SUM190PT | -0.5896 | 0.013624 | -0.11961 | -0.12962 | 0.32397 | 5.63076 | 0.00087 | 3.31207E-06 | 0.46259 | 0.20402 | 0.18481 |
| MEL202 | -0.5226 | 0.006451 | -0.39868 | -0.19052 | 0.32258 | 5.59517 | 0.000909 | 4.94906E-05 | 0.48227 | 0.00207 | 0.09254 |
| SNU503 | -0.5051 | 0.191972 | -0.07072 | -0.01582 | 0.32111 | 5.55761 | 0.000953 | 9.17973E-05 | 0.09084 | 0.31278 | 0.45658 |
| HSC1 | -0.5033 | 0.066775 | -0.32459 | -0.30476 | 0.32059 | 5.54438 | 0.000969 | 9.76063E-05 | 0.32249 | 0.01073 | 0.0157 |
| SKGIIIA | -0.5765 | -0.00876 | -0.22236 | -0.06969 | 0.32042 | 5.54016 | 0.000974 | 5.89832E-06 | 0.47592 | 0.06032 | 0.31528 |
| HGC27 | -0.3929 | 0.11409 | -0.22882 | -0.442 | 0.31837 | 5.48822 | 0.001039 | 0.002380407 | 0.21508 | 0.05498 | 0.00066 |
| A549 | -0.5287 | 0.139483 | -0.12658 | -0.08459 | 0.31798 | 5.4783 | 0.001052 | 3.95427E-05 | 0.167 | 0.19053 | 0.27959 |
| NCIH1944 | -0.4872 | 0.17264 | -0.21527 | -0.13059 | 0.31724 | 5.45965 | 0.001077 | 0.000166445 | 0.11528 | 0.06663 | 0.18302 |
| MM383 | -0.5975 | -0.07869 | -0.25765 | -0.30358 | 0.31674 | 5.44694 | 0.001095 | 2.31264E-06 | 0.29349 | 0.03543 | 0.01605 |
| C4II | -0.5587 | 0.070082 | -0.04411 | -0.10889 | 0.31633 | 5.43676 | 0.001109 | 1.24087E-05 | 0.31433 | 0.38051 | 0.22581 |
| JHH5 | -0.4924 | 0.037906 | -0.39062 | -0.12552 | 0.31631 | 5.43614 | 0.00111 | 0.000140497 | 0.39691 | 0.00252 | 0.19255 |
| TE8 | -0.5364 | 0.112745 | -0.14408 | -0.15119 | 0.31478 | 5.39783 | 0.001164 | 2.9722E-05 | 0.21783 | 0.15908 | 0.14731 |
| CALU1 | -0.4282 | 0.114658 | -0.39392 | -0.22532 | 0.31396 | 5.37727 | 0.001195 | 0.000960024 | 0.21393 | 0.00232 | 0.05783 |
| MAPACHS77 | -0.5351 | 0.074331 | -0.21335 | -0.03965 | 0.31382 | 5.37367 | 0.0012 | 3.11444E-05 | 0.30397 | 0.06842 | 0.39229 |
| MFE280 | -0.6055 | -0.09913 | -0.16322 | -0.12144 | 0.31359 | 5.36816 | 0.001209 | 1.59262E-06 | 0.24669 | 0.12869 | 0.20043 |
| SNU1077 | -0.542 | 0.07274 | -0.23314 | -0.15568 | 0.31326 | 5.35987 | 0.001221 | 2.39806E-05 | 0.30783 | 0.05162 | 0.14016 |
| SNU61 | -0.5355 | 0.053161 | -0.27605 | -0.1105 | 0.31216 | 5.33254 | 0.001264 | 3.07439E-05 | 0.35694 | 0.02616 | 0.22245 |
| KMRC20 | -0.454 | 0.245238 | -0.10623 | -0.09124 | 0.31181 | 5.32386 | 0.001278 | 0.000464477 | 0.04303 | 0.2314 | 0.26429 |
| GSU | -0.4671 | 0.159447 | -0.20374 | -0.25545 | 0.3103 | 5.28643 | 0.00134 | 0.000313274 | 0.13436 | 0.07793 | 0.03669 |
| SNU182 | -0.485 | 0.110979 | -0.2714 | -0.26717 | 0.30957 | 5.26838 | 0.001371 | 0.000178769 | 0.22146 | 0.02829 | 0.03035 |
| TE4 | -0.5749 | 7.02E-05 | -0.12959 | -0.06769 | 0.30951 | 5.26694 | 0.001374 | 6.31739E-06 | 0.49981 | 0.18487 | 0.32022 |
| SHMAC4 | -0.5357 | 0.081827 | -0.20333 | -0.07639 | 0.30913 | 5.25755 | 0.00139 | 3.05063E-05 | 0.28607 | 0.07836 | 0.29899 |
| EFO27 | -0.5643 | 0.036587 | -0.09867 | -0.06575 | 0.30901 | 5.25464 | 0.001395 | 9.83707E-06 | 0.40042 | 0.24771 | 0.32505 |
| KMCH1 | -0.4616 | 0.207753 | -0.1693 | -0.12655 | 0.30686 | 5.20192 | 0.001492 | 0.000369791 | 0.07385 | 0.11992 | 0.19059 |
| IGR39 | -0.4395 | 0.169053 | -0.28503 | -0.19707 | 0.30555 | 5.16987 | 0.001554 | 0.000702347 | 0.12027 | 0.02241 | 0.08507 |
| SISO | -0.4791 | 0.09567 | -0.28794 | -0.01469 | 0.30462 | 5.14734 | 0.001599 | 0.000215641 | 0.25434 | 0.0213 | 0.45967 |
| COV644 | -0.5272 | 0.080082 | -0.23736 | -0.12506 | 0.30455 | 5.1456 | 0.001602 | 4.17927E-05 | 0.29019 | 0.04848 | 0.19342 |
| C4I | -0.5883 | -0.06626 | -0.25558 | -0.20447 | 0.30423 | 5.13787 | 0.001618 | 3.52001E-06 | 0.32377 | 0.03662 | 0.07717 |
| EPLC272H | -0.4005 | 0.245421 | -0.1757 | -0.20468 | 0.30415 | 5.13577 | 0.001622 | 0.001976406 | 0.04291 | 0.11115 | 0.07696 |
| KON | -0.3956 | 0.295503 | -0.13496 | -0.05843 | 0.30354 | 5.12108 | 0.001653 | 0.002229962 | 0.01861 | 0.17503 | 0.34346 |
| NCIH2030 | -0.4728 | 0.208359 | -0.09173 | 0.010124 | 0.30321 | 5.11296 | 0.00167 | 0.000262903 | 0.07324 | 0.26318 | 0.47218 |
| T47D | -0.5962 | -0.13079 | -0.26132 | -0.15471 | 0.30264 | 5.09934 | 0.0017 | 2.45211E-06 | 0.18264 | 0.0334 | 0.14168 |
| NUGC3 | -0.5734 | -0.00683 | -0.12593 | -0.09058 | 0.30263 | 5.09913 | 0.0017 | 6.73204E-06 | 0.48123 | 0.19177 | 0.26579 |
| BB30HNC | -0.4092 | 0.272763 | -0.15524 | -0.00782 | 0.30253 | 5.0967 | 0.001705 | 0.001585561 | 0.02765 | 0.14085 | 0.47852 |
| JHU011 | -0.5566 | 0.021507 | -0.22571 | -0.13985 | 0.30191 | 5.08176 | 0.001738 | 1.35057E-05 | 0.44107 | 0.05751 | 0.16637 |
| SAS | -0.5537 | 0.04961 | -0.11939 | -0.07383 | 0.30191 | 5.08168 | 0.001738 | 1.5182E-05 | 0.36612 | 0.20445 | 0.30517 |
| CORL105 | -0.541 | -0.03236 | -0.21313 | 0.015669 | 0.3016 | 5.07417 | 0.001755 | 2.49551E-05 | 0.41173 | 0.06863 | 0.457 |
| COGAR359 | -0.2898 | 0.184822 | -0.39127 | 0.032518 | 0.3011 | 5.06221 | 0.001782 | 0.020601381 | 0.09941 | 0.00248 | 0.41131 |
| PACADD137 | -0.5349 | 0.073982 | -0.08557 | 8.41E-05 | 0.3005 | 5.04763 | 0.001815 | 3.13984E-05 | 0.30481 | 0.2773 | 0.49977 |
| PSS008 | -0.5623 | -0.04813 | -0.31844 | -0.19881 | 0.30039 | 5.04515 | 0.001821 | 1.07052E-05 | 0.36999 | 0.0121 | 0.08316 |
| PATU8988S | -0.4969 | 0.018499 | -0.38333 | -0.19122 | 0.29943 | 5.02204 | 0.001876 | 0.000120863 | 0.44927 | 0.003 | 0.09172 |
| SKOV3 | -0.4844 | 0.162001 | -0.17816 | -0.0549 | 0.29841 | 4.9977 | 0.001935 | 0.00018236 | 0.13051 | 0.10789 | 0.35248 |
| SNU886 | -0.4605 | 0.025635 | -0.32029 | -0.40765 | 0.29708 | 4.96589 | 0.002016 | 0.000382529 | 0.42987 | 0.01168 | 0.00165 |
| KMRC3 | -0.3799 | 0.035903 | -0.49171 | -0.21856 | 0.29662 | 4.95516 | 0.002044 | 0.003250587 | 0.40225 | 0.00014 | 0.06365 |
| P4E6 | -0.5115 | 0.077979 | -0.22718 | -0.03949 | 0.29627 | 4.94682 | 0.002066 | 7.33883E-05 | 0.29519 | 0.0563 | 0.39271 |
| FADU | -0.3158 | 0.405581 | 0.166392 | 0.070541 | 0.29624 | 4.94593 | 0.002068 | 0.012749132 | 0.00174 | 0.12407 | 0.3132 |

| | | | | | | | | | | | |
|---|---|---|---|---|---|---|---|---|---|---|---|
| UBLC1 | -0.4588 | 0.22099 | -0.07537 | 0.023338 | 0.29613 | 4.94335 | 0.002075 | 0.000403043 | 0.06151 | 0.30146 | 0.4361 |
| C84 | -0.5874 | -0.07433 | -0.20587 | -0.22665 | 0.29602 | 4.94078 | 0.002082 | 3.66239E-06 | 0.30398 | 0.07574 | 0.05673 |
| SNU719 | -0.5289 | -0.04137 | -0.37852 | -0.19571 | 0.29593 | 4.93867 | 0.002087 | 3.93446E-05 | 0.38772 | 0.00336 | 0.08659 |
| COLO824 | -0.3796 | 0.224794 | -0.27868 | -0.14572 | 0.2955 | 4.92851 | 0.002115 | 0.003276376 | 0.05827 | 0.02501 | 0.15631 |
| HCC461 | -0.4536 | 0.191309 | -0.11317 | -0.20278 | 0.29517 | 4.92072 | 0.002136 | 0.000468696 | 0.09161 | 0.21697 | 0.07893 |
| NCIH2170 | -0.551 | 0.040846 | -0.14254 | -0.09816 | 0.29508 | 4.91851 | 0.002142 | 1.69094E-05 | 0.38911 | 0.1617 | 0.24883 |
| SNU1079 | -0.5241 | 0.091569 | -0.01934 | -0.00615 | 0.29488 | 4.91394 | 0.002155 | 4.68088E-05 | 0.26355 | 0.44696 | 0.48311 |
| HCC95 | -0.529 | 0.037859 | -0.25567 | -0.24648 | 0.2948 | 4.91198 | 0.00216 | 3.91277E-05 | 0.39704 | 0.03657 | 0.04222 |
| ICC9 | -0.4437 | 0.212124 | -0.18134 | -0.07337 | 0.29441 | 4.90277 | 0.002186 | 0.000623923 | 0.06958 | 0.10377 | 0.30629 |
| EGI1 | -0.3982 | 0.105159 | -0.39223 | -0.06722 | 0.29337 | 4.87824 | 0.002256 | 0.002091494 | 0.23367 | 0.00242 | 0.32138 |
| CAKI1 | -0.421 | 0.250534 | -0.07702 | 0.103166 | 0.29294 | 4.86808 | 0.002286 | 0.001165444 | 0.03964 | 0.2975 | 0.23794 |
| JHESOAD1 | -0.5073 | 0.119278 | -0.17493 | -0.09441 | 0.2928 | 4.86473 | 0.002295 | 8.49051E-05 | 0.20467 | 0.11217 | 0.25716 |
| KOSC2 | -0.4526 | 0.220529 | -0.11403 | -0.02249 | 0.29271 | 4.86262 | 0.002302 | 0.000483658 | 0.06191 | 0.2152 | 0.43839 |
| SW1573 | -0.4105 | 0.163731 | -0.29731 | -0.23446 | 0.29236 | 4.85446 | 0.002326 | 0.001533714 | 0.12795 | 0.01801 | 0.05062 |
| LUDLU1 | -0.4383 | 0.212532 | -0.18686 | -0.09792 | 0.29233 | 4.85386 | 0.002328 | 0.000727353 | 0.0692 | 0.09691 | 0.24936 |
| BC3C | -0.3988 | 0.234419 | -0.16536 | -0.20546 | 0.29186 | 4.84268 | 0.002362 | 0.002058891 | 0.05065 | 0.12556 | 0.07616 |
| SUIT2 | -0.3754 | 0.161017 | -0.3569 | -0.21415 | 0.29172 | 4.83944 | 0.002371 | 0.003610905 | 0.13198 | 0.00548 | 0.06767 |
| NCIH2291 | -0.2604 | 0.334423 | -0.24595 | -0.03476 | 0.29105 | 4.82382 | 0.00242 | 0.033921038 | 0.00881 | 0.04256 | 0.40531 |
| A2058 | -0.4587 | 0.14812 | -0.25699 | -0.08314 | 0.29083 | 4.81859 | 0.002436 | 0.000403374 | 0.15232 | 0.0358 | 0.28298 |
| SNU1196 | -0.5112 | 0.04531 | -0.29707 | -0.21369 | 0.29036 | 4.80773 | 0.00247 | 7.4273E-05 | 0.37735 | 0.01809 | 0.0681 |
| ICC4 | -0.4937 | 0.128972 | -0.12921 | 0.020149 | 0.29 | 4.79933 | 0.002497 | 0.000134735 | 0.18603 | 0.18557 | 0.44477 |
| U2OS | -0.3482 | 0.144609 | -0.3681 | -0.2965 | 0.28919 | 4.78038 | 0.002559 | 0.006608701 | 0.15818 | 0.00427 | 0.01827 |
| RKN | -0.3304 | 0.209685 | -0.34446 | -0.15965 | 0.28913 | 4.77893 | 0.002564 | 0.00955514 | 0.07194 | 0.00715 | 0.13405 |
| 5637 | -0.4591 | 0.159482 | -0.22639 | -0.14756 | 0.28802 | 4.75338 | 0.00265 | 0.000398646 | 0.1343 | 0.05695 | 0.15324 |
| CCLFPEDS0008T | -0.2938 | 0.141498 | -0.41254 | -0.29991 | 0.28797 | 4.75209 | 0.002655 | 0.019191651 | 0.1635 | 0.00145 | 0.01717 |
| TR146 | -0.5564 | -0.00124 | -0.15252 | -0.08921 | 0.28788 | 4.74994 | 0.002662 | 1.35781E-05 | 0.4966 | 0.14517 | 0.26893 |
| 127399 | -0.5199 | -0.08653 | -0.36235 | -0.39931 | 0.28736 | 4.73791 | 0.002704 | 5.45684E-05 | 0.27508 | 0.00486 | 0.00203 |
| KLE | -0.4116 | 0.13513 | -0.34079 | -0.0765 | 0.28709 | 4.7317 | 0.002726 | 0.00149147 | 0.17473 | 0.00772 | 0.29873 |
| HSC3 | -0.5304 | -0.01268 | -0.32209 | -0.20943 | 0.28672 | 4.72329 | 0.002755 | 3.71674E-05 | 0.46519 | 0.01127 | 0.07219 |
| 143B | -0.4322 | 0.040444 | -0.3726 | -0.36122 | 0.28637 | 4.71506 | 0.002785 | 0.000861361 | 0.39017 | 0.00385 | 0.00498 |
| KKU100 | -0.3738 | 0.218958 | -0.2345 | -0.22532 | 0.28599 | 4.70641 | 0.002816 | 0.003744918 | 0.06329 | 0.05059 | 0.05783 |
| SNU410 | -0.3405 | 0.082222 | -0.46246 | -0.23378 | 0.28501 | 4.68379 | 0.0029 | 0.007770062 | 0.28513 | 0.00036 | 0.05113 |
| IPC298 | -0.3682 | 0.060702 | -0.46334 | -0.17895 | 0.28498 | 4.68304 | 0.002903 | 0.004258607 | 0.3377 | 0.00035 | 0.10686 |
| SKMEL24 | -0.5695 | -0.05594 | -0.19818 | -0.24044 | 0.28471 | 4.67699 | 0.002926 | 7.92433E-06 | 0.3498 | 0.08385 | 0.04629 |
| U937 | -0.4703 | 0.034319 | -0.37778 | -0.19366 | 0.28391 | 4.65863 | 0.002996 | 0.000283742 | 0.40648 | 0.00342 | 0.0889 |
| CASKI | -0.4413 | 0.214382 | -0.11767 | -0.12286 | 0.28389 | 4.65802 | 0.002999 | 0.000667685 | 0.06746 | 0.20787 | 0.19766 |
| COV434 | -0.5104 | 0.011477 | -0.32978 | -0.20832 | 0.28338 | 4.64636 | 0.003045 | 7.64234E-05 | 0.46848 | 0.00967 | 0.07328 |
| DOV13 | -0.4409 | 0.172072 | -0.20871 | -0.19148 | 0.28301 | 4.63804 | 0.003078 | 0.000675777 | 0.11606 | 0.0729 | 0.09142 |
| RHJT | -0.3151 | 0.098599 | -0.45159 | -0.0761 | 0.28282 | 4.63365 | 0.003095 | 0.012916056 | 0.24786 | 0.0005 | 0.29969 |
| CAPAN1 | -0.5185 | 0.049934 | -0.19803 | -0.04099 | 0.2828 | 4.63318 | 0.003097 | 5.72445E-05 | 0.36528 | 0.08401 | 0.38873 |
| KNS81 | -0.2209 | 0.364204 | -0.16012 | -0.15002 | 0.28238 | 4.62355 | 0.003136 | 0.061566702 | 0.00466 | 0.13334 | 0.1492 |
| SF268 | -0.4545 | 0.186933 | -0.15758 | -0.04256 | 0.2817 | 4.60807 | 0.0032 | 0.000456542 | 0.09682 | 0.13722 | 0.38459 |
| U251MGDM | -0.2518 | 0.227763 | -0.36466 | -0.17673 | 0.28145 | 4.60233 | 0.003224 | 0.038840438 | 0.05583 | 0.00461 | 0.10977 |
| 2313287 | -0.4596 | 0.166563 | -0.03517 | -0.18377 | 0.28008 | 4.57123 | 0.003357 | 0.000392838 | 0.12382 | 0.40421 | 0.10071 |
| BFTC909 | -0.3899 | 0.208784 | -0.18308 | -0.2381 | 0.27922 | 4.5518 | 0.003443 | 0.002564955 | 0.07282 | 0.10158 | 0.04795 |
| KE39 | -0.4317 | 0.147994 | -0.28512 | -0.12403 | 0.27896 | 4.54583 | 0.00347 | 0.00087384 | 0.15252 | 0.02238 | 0.19541 |
| LXF289 | -0.4753 | 0.085029 | -0.25204 | -0.26068 | 0.27809 | 4.52624 | 0.00356 | 0.000243524 | 0.27857 | 0.03872 | 0.03374 |
| 21NT | -0.5284 | -0.03123 | -0.28036 | -0.31416 | 0.27795 | 4.52318 | 0.003574 | 4.0029E-05 | 0.41476 | 0.0243 | 0.01315 |
| OSC20 | -0.5274 | 0.006932 | -0.25384 | -0.24634 | 0.27745 | 4.5118 | 0.003628 | 4.15174E-05 | 0.48095 | 0.03764 | 0.0423 |
| KARPAS299 | -0.5103 | 0.05942 | -0.23663 | -0.16276 | 0.27736 | 4.50977 | 0.003637 | 7.65957E-05 | 0.34094 | 0.04902 | 0.12939 |
| SUM102PT | -0.526 | -0.06652 | -0.31395 | -0.35341 | 0.2765 | 4.49055 | 0.00373 | 4.36873E-05 | 0.32314 | 0.0132 | 0.00591 |
| CALU6 | -0.2797 | 0.310608 | -0.23714 | -0.0184 | 0.27593 | 4.47772 | 0.003793 | 0.024596811 | 0.01407 | 0.04864 | 0.44953 |
| A375SKINCJ3 | -0.3144 | -0.0888 | -0.56029 | -0.14427 | 0.27529 | 4.46333 | 0.003865 | 0.013096548 | 0.26986 | 1.2E-05 | 0.15876 |
| PK45H | -0.2709 | 0.143627 | -0.43646 | -0.19154 | 0.27503 | 4.45759 | 0.003894 | 0.028546648 | 0.15985 | 0.00077 | 0.09135 |
| UPCISCC200 | -0.4782 | 0.159907 | -0.0477 | -0.02439 | 0.27475 | 4.45133 | 0.003926 | 0.000222228 | 0.13366 | 0.37109 | 0.43323 |
| YD38 | -0.5111 | 0.089494 | -0.09456 | -0.13795 | 0.2746 | 4.44795 | 0.003944 | 7.44435E-05 | 0.26827 | 0.25682 | 0.16969 |
| HT29 | -0.4934 | 0.03043 | -0.30117 | -0.09757 | 0.27448 | 4.44519 | 0.003958 | 0.000135713 | 0.41692 | 0.01678 | 0.25012 |
| NCIH2126 | -0.5126 | 0.011726 | -0.29675 | -0.2041 | 0.27396 | 4.43377 | 0.004018 | 7.06403E-05 | 0.46779 | 0.01819 | 0.07755 |
| SCS214 | -0.399 | 0.020895 | -0.45408 | -0.18003 | 0.27381 | 4.43036 | 0.004036 | 0.002049514 | 0.44274 | 0.00046 | 0.10545 |
| NCIH1666 | -0.4741 | 0.154133 | -0.11425 | -0.11214 | 0.27267 | 4.40497 | 0.004172 | 0.000252524 | 0.1426 | 0.21475 | 0.21907 |
| NCIH157DM | -0.3009 | 0.331549 | -0.15956 | -0.04047 | 0.27157 | 4.38051 | 0.004308 | 0.016850113 | 0.00933 | 0.13418 | 0.39009 |
| ESS1 | -0.3329 | 0.174692 | -0.34344 | -0.21791 | 0.27122 | 4.37294 | 0.004352 | 0.009091124 | 0.11249 | 0.00731 | 0.06423 |
| TOV112D | -0.3778 | 0.238611 | -0.01394 | -0.20548 | 0.27117 | 4.37174 | 0.004358 | 0.003415526 | 0.04758 | 0.46173 | 0.07614 |
| ETCC016 | -0.5334 | 0.007245 | -0.20226 | -0.11522 | 0.27106 | 4.36933 | 0.004372 | 3.31864E-05 | 0.48009 | 0.07947 | 0.21278 |
| LCLC103H | -0.4849 | -0.04574 | -0.3787 | -0.34458 | 0.27063 | 4.35977 | 0.004427 | 0.000179413 | 0.37622 | 0.00334 | 0.00713 |
| PEO4 | -0.5062 | 0.063686 | -0.04951 | 0.015943 | 0.26968 | 4.33886 | 0.004551 | 8.81722E-05 | 0.33019 | 0.36639 | 0.45625 |
| RPMI7951 | -0.4089 | 0.018731 | -0.43809 | -0.26407 | 0.2696 | 4.33703 | 0.004562 | 0.001597309 | 0.44864 | 0.00073 | 0.03193 |
| SW1990 | -0.4417 | 0.203097 | 0.017005 | 0.062912 | 0.26893 | 4.32225 | 0.004651 | 0.000660361 | 0.0786 | 0.45335 | 0.33213 |
| NCIH292 | -0.3937 | 0.217212 | -0.20979 | -0.02475 | 0.2687 | 4.31737 | 0.004681 | 0.002334439 | 0.06486 | 0.07184 | 0.43227 |
| SNB75 | -0.4851 | -0.2018 | -0.49582 | -0.26333 | 0.26821 | 4.30645 | 0.004749 | 0.000178177 | 0.07996 | 0.00013 | 0.03233 |
| HEC59 | -0.3713 | 0.084771 | -0.41399 | -0.1739 | 0.26746 | 4.28997 | 0.004853 | 0.003969014 | 0.27917 | 0.0014 | 0.11356 |
| LN382 | -0.2697 | 0.247515 | -0.20225 | -0.28286 | 0.26583 | 4.25445 | 0.005086 | 0.029108869 | 0.04155 | 0.07948 | 0.02327 |
| ICC8 | -0.2833 | 0.257779 | -0.1956 | -0.254 | 0.26575 | 4.25277 | 0.005097 | 0.023114764 | 0.03535 | 0.08671 | 0.03754 |
| NCIH522 | -0.5274 | -0.03966 | -0.21327 | -0.3108 | 0.2653 | 4.2429 | 0.005164 | 4.14703E-05 | 0.39226 | 0.0685 | 0.01402 |
| KALS1 | -0.2875 | 0.037022 | -0.45335 | -0.38592 | 0.26494 | 4.23519 | 0.005217 | 0.021449692 | 0.39927 | 0.00047 | 0.00282 |
| HEC1 | -0.4627 | -0.03469 | -0.41776 | -0.2375 | 0.26455 | 4.22657 | 0.005277 | 0.000358476 | 0.40549 | 0.00127 | 0.04838 |
| VCAP | -0.4295 | -0.19451 | -0.54522 | -0.35599 | 0.26447 | 4.22489 | 0.005289 | 0.000927399 | 0.08793 | 2.1E-05 | 0.00559 |
| HOKUG | -0.505 | 0.066418 | -0.17595 | -0.10192 | 0.26421 | 4.21913 | 0.005329 | 9.20339E-05 | 0.32338 | 0.11081 | 0.24063 |
| RERFLCAI | -0.3588 | 0.011002 | -0.48073 | -0.23899 | 0.26396 | 4.21387 | 0.005366 | 0.005248028 | 0.46978 | 0.0002 | 0.04731 |
| TE1 | -0.4632 | 0.064136 | -0.30945 | -0.14494 | 0.26396 | 4.21378 | 0.005367 | 0.00035297 | 0.32906 | 0.01438 | 0.15762 |
| MCF7 | -0.3761 | 0.167946 | -0.12333 | -0.31037 | 0.26358 | 4.20558 | 0.005425 | 0.003550268 | 0.12184 | 0.19675 | 0.01413 |
| PANC0813 | -0.4633 | -0.0258 | -0.39755 | -0.15503 | 0.26356 | 4.2052 | 0.005428 | 0.000351437 | 0.42943 | 0.00213 | 0.14118 |
| ICC12 | -0.2299 | 0.337811 | -0.17949 | 0.154321 | 0.26335 | 4.20049 | 0.005462 | 0.054125594 | 0.00821 | 0.10616 | 0.1423 |
| HT55 | -0.3267 | 0.170754 | -0.35612 | -0.0933 | 0.26332 | 4.19996 | 0.005466 | 0.010288955 | 0.11789 | 0.00557 | 0.25964 |
| MDAMB453 | -0.5271 | 0.01694 | -0.09222 | -0.17045 | 0.26311 | 4.19542 | 0.005499 | 4.20661E-05 | 0.45352 | 0.26208 | 0.11832 |
| OVTOKO | -0.5136 | 0.030574 | -0.20465 | -0.09781 | 0.26306 | 4.1944 | 0.005506 | 6.82871E-05 | 0.41653 | 0.07699 | 0.24961 |
| NZM7 | -0.3795 | 0.18353 | -0.27324 | -0.03884 | 0.26291 | 4.19097 | 0.005531 | 0.003280692 | 0.10101 | 0.02743 | 0.39442 |
| BICR78 | -0.4297 | 0.179214 | -0.02257 | 0.127089 | 0.2627 | 4.1865 | 0.005564 | 0.000921233 | 0.10651 | 0.43819 | 0.18957 |
| ICC137 | -0.5018 | -0.0571 | -0.3255 | -0.34585 | 0.26216 | 4.17491 | 0.00565 | 0.000102554 | 0.34683 | 0.01054 | 0.00695 |
| CH157MN | -0.3372 | -0.10181 | -0.54619 | -0.37661 | 0.26138 | 4.15802 | 0.005778 | 0.008321485 | 0.24085 | 2E-05 | 0.00351 |
| OCIC4P | -0.56 | -0.11451 | -0.22975 | -0.22756 | 0.26097 | 4.14932 | 0.005845 | 1.17677E-05 | 0.21422 | 0.05424 | 0.056 |
| EMTOKA | -0.4504 | 0.179326 | -0.00191 | 0.012396 | 0.25981 | 4.1244 | 0.006041 | 0.000515853 | 0.10637 | 0.49474 | 0.46596 |
| SNU1105 | -0.0941 | 0.214066 | -0.2339 | -0.37651 | 0.25951 | 4.11787 | 0.006094 | 0.257815899 | 0.06775 | 0.05104 | 0.00352 |
| MDAMB231 | -0.2268 | 0.232228 | -0.28377 | -0.27094 | 0.25936 | 4.11476 | 0.006119 | 0.05661693 | 0.05231 | 0.02291 | 0.02851 |
| ICC10 | -0.4664 | 0.058843 | -0.07872 | -0.29213 | 0.25833 | 4.09268 | 0.006301 | 0.000320112 | 0.3424 | 0.29342 | 0.01977 |
| TFK1 | -0.4506 | 0.102369 | -0.26534 | -0.12675 | 0.25797 | 4.08487 | 0.006366 | 0.000512515 | 0.23966 | 0.03128 | 0.19021 |
| YAPC | -0.4657 | -0.03042 | -0.3816 | -0.13906 | 0.25752 | 4.07538 | 0.006447 | 0.00032741 | 0.41694 | 0.00312 | 0.16774 |
| SKMEL19 | -0.372 | 0.123817 | -0.35342 | -0.18518 | 0.25748 | 4.07458 | 0.006454 | 0.003905144 | 0.19581 | 0.00591 | 0.09896 |
| YSCCC | -0.4339 | 0.201639 | -0.01625 | 0.031199 | 0.25714 | 4.06727 | 0.006517 | 0.000821791 | 0.08013 | 0.4554 | 0.41485 |
| ONDA7 | -0.4394 | 0.056547 | -0.1929 | -0.35173 | 0.25667 | 4.05729 | 0.006604 | 0.000705401 | 0.34825 | 0.08977 | 0.00613 |
| HCC2450 | -0.4339 | 0.173862 | -0.15614 | -0.04248 | 0.25526 | 4.02723 | 0.006873 | 0.000822167 | 0.11362 | 0.13944 | 0.38481 |
| SNU840 | -0.4692 | 0.130549 | -0.12432 | -0.08168 | 0.25519 | 4.02575 | 0.006887 | 0.000293536 | 0.18309 | 0.19484 | 0.28641 |
| LS513 | -0.4235 | -0.04102 | -0.4053 | -0.06696 | 0.25418 | 4.00457 | 0.007084 | 0.001090913 | 0.38865 | 0.00175 | 0.32202 |
| YKG1 | -0.2169 | 0.144867 | -0.39476 | -0.29761 | 0.25412 | 4.0033 | 0.007096 | 0.065168265 | 0.15775 | 0.00228 | 0.01791 |
| MB1 | -0.2498 | 0.132176 | -0.36141 | -0.34309 | 0.25332 | 3.9864 | 0.007258 | 0.040101538 | 0.18009 | 0.00496 | 0.00736 |
| HCC1187 | -0.1209 | 0.284448 | -0.11057 | -0.31639 | 0.25325 | 3.98485 | 0.007273 | 0.20139563 | 0.02264 | 0.22231 | 0.0126 |
| COLO680N | -0.3795 | 0.119985 | -0.30888 | -0.26295 | 0.25199 | 3.95827 | 0.007535 | 0.003280969 | 0.20328 | 0.01453 | 0.03252 |
| OVMIU | -0.4724 | 0.119811 | -0.08844 | -0.12345 | 0.25178 | 3.95391 | 0.007579 | 0.000266569 | 0.20362 | 0.27067 | 0.19652 |
| HCC1954 | -0.3938 | 0.016158 | -0.17648 | -0.43863 | 0.25165 | 3.95124 | 0.007606 | 0.002330986 | 0.45566 | 0.11011 | 0.00072 |
| 253J | -0.4841 | 0.098499 | -0.07958 | -0.09872 | 0.25073 | 3.93195 | 0.007804 | 0.000184099 | 0.24808 | 0.29139 | 0.2476 |
| CAS1 | -0.3743 | 0.04915 | -0.29655 | -0.40051 | 0.25055 | 3.92822 | 0.007843 | 0.003700768 | 0.36732 | 0.01826 | 0.00197 |
| UMUC16 | -0.2843 | 0.087364 | -0.44613 | -0.20517 | 0.24999 | 3.91645 | 0.007968 | 0.022709586 | 0.27316 | 0.00058 | 0.07645 |
| HKA1 | -0.4734 | 0.117971 | -0.06432 | -0.04725 | 0.24952 | 3.90662 | 0.008073 | 0.000258051 | 0.20727 | 0.32859 | 0.37228 |
| KYSE220 | -0.36 | 0.22063 | -0.20809 | 0.01065 | 0.24899 | 3.89563 | 0.008193 | 0.005116649 | 0.06182 | 0.07351 | 0.47074 |
| SW1271 | -0.269 | 0.260596 | -0.22315 | -0.19848 | 0.24896 | 3.89504 | 0.008199 | 0.029435715 | 0.03379 | 0.05965 | 0.08352 |
| J82 | -0.3909 | 0.140631 | -0.23949 | -0.25349 | 0.24877 | 3.86977 | 0.008481 | 0.002502195 | 0.165 | 0.04696 | 0.03785 |
| ASH3 | -0.4467 | 0.051494 | -0.17068 | 0.084436 | 0.24772 | 3.86924 | 0.008487 | 0.000573739 | 0.36124 | 0.118 | 0.27995 |
| PANC1005 | -0.43 | 0.039659 | -0.32266 | -0.29991 | 0.24764 | 3.86754 | 0.008507 | 0.000913524 | 0.39225 | 0.01115 | 0.01717 |
| KCIMOH1 | -0.455 | 0.015207 | -0.34509 | -0.17999 | 0.24741 | 3.86269 | 0.008562 | 0.000450264 | 0.45826 | 0.00706 | 0.10551 |

| | | | | | | | | | | | |
|---|---|---|---|---|---|---|---|---|---|---|---|
| RT11284 | -0.4451 | 0.034759 | -0.24879 | -0.3316 | 0.24626 | 3.83898 | 0.008838 | 0.000600009 | 0.40531 | 0.04073 | 0.00932 |
| NCIH1793 | -0.4697 | 0.094335 | -0.17081 | -0.15763 | 0.24625 | 3.83863 | 0.008842 | 0.000289743 | 0.25732 | 0.11781 | 0.13713 |
| KNS42 | -0.3339 | 0.073285 | -0.35485 | -0.35684 | 0.24589 | 3.8313 | 0.00893 | 0.008895085 | 0.30651 | 0.00573 | 0.00548 |
| SET2 | -0.4464 | -0.34879 | -0.48778 | -0.22759 | 0.24476 | 3.8079 | 0.009214 | 0.000578029 | 0.00652 | 0.00016 | 0.05597 |
| DANG | -0.5008 | -0.03223 | -0.20289 | -0.03111 | 0.24471 | 3.80692 | 0.009226 | 0.000106053 | 0.41207 | 0.07882 | 0.4151 |
| H376 | -0.3871 | 0.236032 | -0.10167 | 0.000234 | 0.24437 | 3.79989 | 0.009314 | 0.002739505 | 0.04945 | 0.24116 | 0.49936 |
| HOP62 | -0.3314 | 0.255409 | -0.16561 | -0.13646 | 0.24419 | 3.79618 | 0.00936 | 0.009355588 | 0.03671 | 0.1252 | 0.17235 |
| 639V | -0.4514 | 0.041163 | -0.31203 | -0.15913 | 0.24362 | 3.78449 | 0.009508 | 0.00050071 | 0.38827 | 0.01369 | 0.13484 |
| HCC202 | -0.3962 | 0.234962 | 0.001451 | -0.05188 | 0.24324 | 3.77669 | 0.009609 | 0.002197751 | 0.05024 | 0.49601 | 0.36025 |
| HEL9217 | -0.4616 | -0.32191 | -0.50601 | -0.37399 | 0.24321 | 3.776 | 0.009617 | 0.000370631 | 0.01131 | 8.9E-05 | 0.00373 |
| PACADD188 | -0.4076 | 0.006424 | -0.38946 | -0.09981 | 0.24229 | 3.75735 | 0.009861 | 0.00165209 | 0.48234 | 0.00259 | 0.24522 |
| KMRC2 | -0.4615 | 0.049794 | -0.27485 | -0.16024 | 0.242 | 3.75142 | 0.00994 | 0.000371124 | 0.36565 | 0.0267 | 0.13315 |
| WM2664 | -0.3053 | 0.105585 | -0.38186 | -0.27599 | 0.24147 | 3.74054 | 0.010087 | 0.015553214 | 0.23277 | 0.00311 | 0.02619 |
| SUM149PT | -0.4233 | 0.045393 | -0.21639 | 0.083454 | 0.24107 | 3.7323 | 0.010199 | 0.001095562 | 0.37713 | 0.06561 | 0.28224 |
| HS766T | -0.2751 | 0.208478 | -0.31979 | -0.09996 | 0.24055 | 3.72169 | 0.010346 | 0.026605404 | 0.07313 | 0.01179 | 0.24488 |
| 8305C | -0.2988 | -0.04847 | -0.52194 | -0.25037 | 0.24049 | 3.72051 | 0.010362 | 0.017526669 | 0.36908 | 5.1E-05 | 0.03974 |
| DAOY | -0.2522 | 0.306857 | -0.10919 | -0.1736 | 0.23925 | 3.69536 | 0.010719 | 0.038621851 | 0.0151 | 0.22517 | 0.11397 |
| AMO1 | -0.3806 | -0.08353 | -0.44792 | -0.08033 | 0.23894 | 3.68894 | 0.010812 | 0.003201222 | 0.28206 | 0.00055 | 0.28959 |
| 786O | -0.3321 | 0.159799 | -0.30341 | -0.21267 | 0.23877 | 3.68557 | 0.010861 | 0.009226239 | 0.13382 | 0.0161 | 0.06907 |
| H3118 | -0.4386 | 0.126806 | -0.1796 | -0.04936 | 0.23876 | 3.68537 | 0.010864 | 0.000721152 | 0.1901 | 0.10601 | 0.36677 |
| PANC0327 | -0.3612 | 0.240934 | -0.09901 | -0.1316 | 0.23858 | 3.68167 | 0.010919 | 0.004986545 | 0.04595 | 0.24695 | 0.18116 |
| ICC106 | -0.4578 | 0.093271 | -0.10379 | -0.20061 | 0.23792 | 3.66837 | 0.011116 | 0.000414292 | 0.25971 | 0.23659 | 0.08122 |
| CHL1DM | -0.4716 | -0.04685 | -0.35629 | -0.23956 | 0.23792 | 3.66831 | 0.011117 | 0.000272594 | 0.37332 | 0.00555 | 0.04691 |
| RCC10RGB | -0.2892 | 0.267714 | -0.20973 | 0.029084 | 0.23728 | 3.65537 | 0.011313 | 0.020815886 | 0.03008 | 0.0719 | 0.42055 |
| HEY | -0.2809 | 0.183148 | -0.33883 | -0.11938 | 0.23716 | 3.65295 | 0.01135 | 0.024058303 | 0.10149 | 0.00804 | 0.20447 |
| JVE127 | -0.3401 | -0.00517 | -0.46516 | -0.18389 | 0.23672 | 3.64403 | 0.011487 | 0.007835076 | 0.48579 | 0.00033 | 0.10057 |
| EW16 | -0.4214 | -0.03812 | -0.41148 | -0.29141 | 0.23665 | 3.64275 | 0.011507 | 0.001151884 | 0.39634 | 0.00149 | 0.02002 |
| OCUG1 | -0.4134 | 0.061477 | -0.25846 | 0.034109 | 0.23641 | 3.6379 | 0.011582 | 0.001423529 | 0.33574 | 0.03497 | 0.40704 |
| KYSE520 | -0.3603 | 0.252274 | -0.08068 | -0.09353 | 0.23634 | 3.63642 | 0.011605 | 0.005079228 | 0.03858 | 0.28877 | 0.25912 |
| ABC1 | -0.1865 | 0.306548 | -0.25155 | -0.02303 | 0.23627 | 3.63492 | 0.011629 | 0.097342931 | 0.01519 | 0.03902 | 0.43692 |
| SNU349 | -0.1778 | 0.140034 | -0.34104 | -0.35047 | 0.23593 | 3.6282 | 0.011735 | 0.108322853 | 0.16604 | 0.00768 | 0.00629 |
| HS695T | -0.3406 | 0.274546 | -0.08046 | 0.077814 | 0.23528 | 3.61517 | 0.011943 | 0.007759549 | 0.02684 | 0.2893 | 0.29559 |
| JL1 | -0.4596 | -0.09277 | -0.2811 | -0.41961 | 0.23518 | 3.61306 | 0.011977 | 0.000393016 | 0.26084 | 0.02399 | 0.00121 |
| HS746T | -0.3279 | 0.140588 | -0.33819 | -0.1768 | 0.2347 | 3.60336 | 0.012135 | 0.010044663 | 0.16507 | 0.00815 | 0.10968 |
| GB2 | -0.4737 | 0.06076 | -0.132 | -0.02965 | 0.23407 | 3.5908 | 0.012343 | 0.000255487 | 0.33755 | 0.18042 | 0.41901 |
| HEL | -0.3646 | -0.15703 | -0.52882 | -0.33831 | 0.23398 | 3.58902 | 0.012373 | 0.004615293 | 0.13807 | 3.9E-05 | 0.00813 |
| MM370 | -0.2447 | 0.242177 | -0.24215 | -0.21031 | 0.23396 | 3.58855 | 0.01238 | 0.043373713 | 0.04509 | 0.04511 | 0.07133 |
| RMUGS | -0.3395 | 0.183465 | -0.2711 | -0.04726 | 0.23384 | 3.58618 | 0.01242 | 0.00792437 | 0.1011 | 0.02844 | 0.37225 |
| SNU601 | -0.3981 | 0.075281 | -0.33278 | -0.1918 | 0.23382 | 3.58591 | 0.012425 | 0.002097022 | 0.30167 | 0.0091 | 0.09105 |
| JURLMK1 | -0.4351 | -0.27064 | -0.47547 | -0.45854 | 0.23375 | 3.58434 | 0.012451 | 0.000795347 | 0.02866 | 0.00024 | 0.00041 |
| TUHR4TKB | -0.4265 | 0.16501 | -0.07863 | -0.10141 | 0.23343 | 3.57805 | 0.012557 | 0.001006523 | 0.12607 | 0.29365 | 0.24173 |
| JHU029 | -0.4718 | 0.065689 | -0.16826 | -0.13273 | 0.23231 | 3.55556 | 0.012945 | 0.000271196 | 0.32519 | 0.12139 | 0.17908 |
| WPE1NA22 | -0.511 | -0.05073 | -0.22406 | -0.16558 | 0.2321 | 3.55145 | 0.013017 | 7.47499E-05 | 0.36321 | 0.05888 | 0.12524 |
| JHUEM7 | -0.3809 | 0.177258 | -0.16644 | -0.18813 | 0.23167 | 3.54293 | 0.013168 | 0.003176121 | 0.10907 | 0.12399 | 0.09537 |
| GCT | -0.3123 | 0.116861 | -0.37223 | -0.10152 | 0.23081 | 3.52576 | 0.013478 | 0.013612579 | 0.20949 | 0.00388 | 0.24149 |
| UMRC7 | -0.2651 | 0.091986 | -0.42678 | -0.18503 | 0.23075 | 3.52452 | 0.013501 | 0.031405902 | 0.26261 | 0.001 | 0.09914 |
| ICC3 | -0.4459 | -0.01351 | -0.2139 | -0.36155 | 0.23008 | 3.51131 | 0.013744 | 0.000587322 | 0.46292 | 0.0679 | 0.00494 |
| NZM42 | -0.4043 | 0.161398 | -0.16796 | -0.14167 | 0.22985 | 3.50672 | 0.01383 | 0.001797317 | 0.13141 | 0.12182 | 0.1632 |
| MIAPACA2 | -0.2202 | 0.172408 | -0.36643 | -0.18653 | 0.22943 | 3.49848 | 0.013985 | 0.062194753 | 0.1156 | 0.00443 | 0.09731 |
| MDAMB415 | -0.1126 | 0.118426 | -0.11458 | -0.4467 | 0.22943 | 3.49844 | 0.013986 | 0.218142941 | 0.20636 | 0.21409 | 0.00057 |
| SKNEP1 | -0.3023 | 0.087122 | -0.40723 | -0.17308 | 0.2291 | 3.49183 | 0.014112 | 0.016430319 | 0.27372 | 0.00167 | 0.11469 |
| GCIY | -0.4192 | 0.170079 | -0.08962 | -0.02486 | 0.22904 | 3.49078 | 0.014132 | 0.001222886 | 0.11883 | 0.26798 | 0.43196 |
| PEA1 | -0.458 | -0.07957 | -0.34729 | -0.34879 | 0.22816 | 3.47346 | 0.014468 | 0.000411709 | 0.2914 | 0.00674 | 0.00652 |
| HOTHC | -0.356 | 0.028382 | -0.40994 | -0.24829 | 0.22788 | 3.46784 | 0.014579 | 0.005584838 | 0.42244 | 0.00156 | 0.04105 |
| OE33 | -0.3454 | 0.140127 | -0.2785 | 0.01558 | 0.22678 | 3.44625 | 0.015012 | 0.007012616 | 0.16588 | 0.02509 | 0.45724 |
| D502MG | -0.2977 | 0.211373 | -0.25522 | 0.017887 | 0.22629 | 3.43655 | 0.015211 | 0.01787279 | 0.0703 | 0.03683 | 0.45094 |
| EFE184 | -0.4297 | 0.136198 | -0.11643 | -0.01233 | 0.2262 | 3.43485 | 0.015247 | 0.000920987 | 0.17281 | 0.21036 | 0.46615 |
| OS252 | -0.4542 | 0.06405 | -0.19778 | -0.19214 | 0.22603 | 3.43145 | 0.015317 | 0.000461278 | 0.32928 | 0.08429 | 0.09065 |
| SW756 | -0.3891 | 0.0772 | -0.32772 | -0.13614 | 0.22577 | 3.42629 | 0.015425 | 0.002611371 | 0.29706 | 0.01008 | 0.17291 |
| RERFLCAD1 | -0.3679 | 0.034592 | -0.38654 | -0.25194 | 0.22535 | 3.4181 | 0.015597 | 0.004284021 | 0.40575 | 0.00278 | 0.03878 |
| TC106 | -0.2495 | 0.047817 | -0.46107 | -0.16887 | 0.22511 | 3.41339 | 0.015697 | 0.040301686 | 0.37079 | 0.00038 | 0.12053 |
| HCC38 | -0.4248 | 0.08748 | -0.11899 | -0.25804 | 0.22479 | 3.40709 | 0.015832 | 0.00105247 | 0.27289 | 0.20525 | 0.03521 |
| NO36 | -0.4734 | 0.007399 | -0.23931 | -0.20439 | 0.22451 | 3.40167 | 0.015949 | 0.000257955 | 0.47966 | 0.04709 | 0.07725 |
| TMK1 | -0.4444 | 0.037346 | -0.13323 | 0.059719 | 0.2237 | 3.38587 | 0.016296 | 0.000611491 | 0.3984 | 0.17816 | 0.34018 |
| U2904 | -0.5097 | -0.21535 | -0.32652 | -0.34612 | 0.2237 | 3.38583 | 0.016297 | 7.81925E-05 | 0.06656 | 0.01033 | 0.0069 |
| JR | -0.2686 | 0.070441 | -0.43253 | -0.18883 | 0.22336 | 3.37929 | 0.016442 | 0.029646285 | 0.31345 | 0.00085 | 0.09453 |
| 59M | -0.4288 | -0.11369 | -0.40903 | -0.36947 | 0.22299 | 3.37205 | 0.016605 | 0.00094545 | 0.21589 | 0.00159 | 0.00414 |
| GP5D | -0.4372 | 0.047312 | -0.2711 | -0.11136 | 0.22259 | 3.36423 | 0.016783 | 0.000749372 | 0.37211 | 0.02843 | 0.22067 |
| MCC26 | -0.2422 | 0.098022 | -0.40435 | -0.25782 | 0.22227 | 3.35809 | 0.016924 | 0.045067768 | 0.24913 | 0.00179 | 0.03533 |
| SNU668 | -0.4023 | 0.066548 | -0.2983 | -0.23277 | 0.2222 | 3.35675 | 0.016954 | 0.001888569 | 0.32305 | 0.01768 | 0.0519 |
| CCK81 | -0.3392 | 0.053184 | -0.39822 | -0.17984 | 0.22184 | 3.34974 | 0.017117 | 0.007974656 | 0.35688 | 0.00209 | 0.1057 |
| KMS26 | -0.4477 | -0.08493 | -0.38924 | -0.23681 | 0.221 | 3.3335 | 0.0175 | 0.000556311 | 0.27879 | 0.0026 | 0.04888 |
| RCM1 | -0.4436 | 0.027981 | -0.23587 | -0.03052 | 0.22091 | 3.33169 | 0.017543 | 0.000626732 | 0.42352 | 0.04957 | 0.41669 |
| LN235 | -0.2098 | 0.237328 | -0.11178 | -0.28977 | 0.22055 | 3.32474 | 0.01771 | 0.07186063 | 0.04851 | 0.21981 | 0.02062 |
| LS180 | -0.3728 | 0.218695 | -0.09499 | -0.03006 | 0.22 | 3.31413 | 0.017968 | 0.003835119 | 0.06353 | 0.25585 | 0.41793 |
| NCIH209 | -0.4511 | 0.054866 | -0.21715 | -0.15399 | 0.21987 | 3.31155 | 0.018031 | 0.000504502 | 0.35255 | 0.06491 | 0.14283 |
| GSS | -0.3821 | 0.004705 | -0.36933 | -0.30528 | 0.21857 | 3.28655 | 0.018656 | 0.003090538 | 0.48706 | 0.00415 | 0.01555 |
| LN18 | -0.3333 | 0.109992 | -0.31047 | -0.25715 | 0.21856 | 3.28632 | 0.018662 | 0.00900101 | 0.22351 | 0.0141 | 0.03571 |
| THP1 | -0.412 | 0.064441 | -0.263 | -0.03798 | 0.21843 | 3.28393 | 0.018723 | 0.001473972 | 0.3283 | 0.0325 | 0.39671 |
| D542MG | -0.1228 | 0.227149 | -0.12609 | -0.33062 | 0.21808 | 3.27712 | 0.018898 | 0.197714327 | 0.05633 | 0.19146 | 0.00951 |
| MM426 | -0.3784 | 0.106602 | -0.21561 | -0.27324 | 0.21802 | 3.27623 | 0.018921 | 0.003369229 | 0.23061 | 0.06632 | 0.02743 |
| KMS27 | -0.3497 | -0.07917 | -0.47655 | -0.28414 | 0.21735 | 3.26316 | 0.019261 | 0.006399159 | 0.29235 | 0.00023 | 0.02276 |
| KYSE150 | -0.3946 | 0.128332 | -0.2013 | -0.18964 | 0.21726 | 3.26128 | 0.019311 | 0.002287795 | 0.18722 | 0.08049 | 0.09358 |
| LPS6 | -0.3584 | 0.092119 | -0.2499 | -0.30811 | 0.21711 | 3.25845 | 0.019385 | 0.005297329 | 0.2623 | 0.04003 | 0.01475 |
| SKN | -0.4168 | 0.012303 | -0.3186 | -0.27383 | 0.21699 | 3.25614 | 0.019447 | 0.001303052 | 0.46621 | 0.01207 | 0.02716 |
| LPS141 | -0.4425 | 0.064565 | -0.14353 | -0.22324 | 0.21699 | 3.25611 | 0.019447 | 0.000646327 | 0.32799 | 0.16002 | 0.05958 |
| KCL22 | -0.3678 | 0.076579 | -0.27801 | -0.29892 | 0.2169 | 3.25451 | 0.01949 | 0.004300119 | 0.29855 | 0.0253 | 0.01749 |
| NCIH1869 | -0.4607 | 0.009261 | -0.24556 | -0.20299 | 0.21651 | 3.24697 | 0.019691 | 0.000379936 | 0.47455 | 0.04282 | 0.0787 |
| LU135 | -0.2676 | -0.00302 | -0.44872 | -0.33847 | 0.21649 | 3.2467 | 0.019699 | 0.030154461 | 0.4917 | 0.00054 | 0.0081 |
| SUDHL1 | -0.4724 | -0.14582 | -0.36992 | -0.30998 | 0.21627 | 3.24249 | 0.019812 | 0.000266225 | 0.15614 | 0.0041 | 0.01424 |
| K029AX | -0.3122 | -0.03591 | -0.4698 | -0.28013 | 0.21608 | 3.23881 | 0.019912 | 0.01365034 | 0.40224 | 0.00029 | 0.0244 |
| ACCMESO1 | -0.253 | 0.252223 | -0.14976 | -0.20811 | 0.21608 | 3.23879 | 0.019913 | 0.038157964 | 0.03861 | 0.14962 | 0.07349 |
| NCIH1650 | -0.2972 | 0.102587 | -0.17495 | -0.3775 | 0.21591 | 3.23551 | 0.020002 | 0.018037124 | 0.23918 | 0.11215 | 0.00344 |
| TUHR14TKB | -0.4333 | 0.121101 | -0.04768 | -0.09018 | 0.21544 | 3.22655 | 0.020248 | 0.000835398 | 0.20109 | 0.37115 | 0.26671 |
| HUH1 | -0.4384 | 0.032611 | -0.25947 | -0.09128 | 0.21535 | 3.22476 | 0.020298 | 0.000725105 | 0.41106 | 0.03441 | 0.26421 |
| SUSA | -0.2977 | 0.135429 | -0.33405 | -0.18652 | 0.21509 | 3.21979 | 0.020436 | 0.017880672 | 0.17419 | 0.00887 | 0.09732 |
| ASPC1 | -0.3022 | 0.089039 | -0.33754 | 0.025234 | 0.21442 | 3.20707 | 0.020795 | 0.016471842 | 0.26931 | 0.00826 | 0.43095 |
| NCIH747 | -0.41 | 0.14778 | -0.12692 | -0.08766 | 0.21439 | 3.20646 | 0.020812 | 0.00155122 | 0.15288 | 0.18989 | 0.27247 |
| FTC133 | -0.4053 | 0.067854 | -0.24521 | -0.25784 | 0.21432 | 3.20517 | 0.020849 | 0.001751997 | 0.31982 | 0.04305 | 0.03532 |
| UMUC3 | -0.4227 | 0.055941 | -0.18516 | -0.27175 | 0.21375 | 3.1943 | 0.021161 | 0.001114862 | 0.3498 | 0.09899 | 0.02813 |
| NH12 | -0.2678 | 0.053734 | -0.43053 | -0.13362 | 0.21368 | 3.19297 | 0.021199 | 0.030051847 | 0.35546 | 0.0009 | 0.17746 |
| TASK1 | -0.352 | -0.12775 | -0.49586 | -0.2652 | 0.2134 | 3.18775 | 0.021351 | 0.006087946 | 0.18832 | 0.00013 | 0.03135 |
| HSC4 | -0.4172 | 0.138724 | 0.018789 | -0.06825 | 0.21315 | 3.18305 | 0.021489 | 0.00128963 | 0.16833 | 0.44848 | 0.31883 |
| CAL62 | -0.1154 | 0.32762 | -0.22287 | 0.025267 | 0.21309 | 3.18178 | 0.021526 | 0.212392836 | 0.0101 | 0.05989 | 0.43086 |
| NB4 | -0.4289 | -0.02835 | -0.351 | -0.18482 | 0.21272 | 3.17489 | 0.02173 | 0.000942948 | 0.42254 | 0.00622 | 0.0994 |
| JHH2 | -0.3538 | 0.036236 | -0.18085 | -0.39129 | 0.21229 | 3.16667 | 0.021976 | 0.005860762 | 0.40136 | 0.1044 | 0.00248 |
| RH4 | -0.1782 | 0.171062 | -0.37009 | -0.08545 | 0.21182 | 3.15779 | 0.022244 | 0.107887267 | 0.11746 | 0.00408 | 0.27759 |
| HCC78 | -0.4776 | -0.02274 | -0.20137 | -0.21359 | 0.21122 | 3.14635 | 0.022595 | 0.000226621 | 0.43772 | 0.08042 | 0.06819 |
| HHUA | -0.4256 | 0.120013 | -0.1151 | -0.06339 | 0.2111 | 3.14416 | 0.022663 | 0.00103149 | 0.20322 | 0.21304 | 0.33094 |
| EJM | -0.2451 | 0.16997 | -0.33652 | -0.08111 | 0.21104 | 3.14303 | 0.022698 | 0.043146741 | 0.11898 | 0.00843 | 0.28775 |
| LNZ308 | -0.3536 | 0.076765 | -0.29444 | -0.28754 | 0.21076 | 3.13781 | 0.022861 | 0.00587989 | 0.2981 | 0.01897 | 0.02145 |
| CME1 | -0.2877 | 0.207514 | -0.09716 | -0.2511 | 0.21036 | 3.13016 | 0.023102 | 0.021385311 | 0.07408 | 0.25103 | 0.03929 |
| WM983B | -0.3996 | -0.14548 | -0.4367 | -0.37901 | 0.21032 | 3.12952 | 0.023122 | 0.002020349 | 0.15671 | 0.00076 | 0.00332 |
| KMS20 | -0.4116 | -0.18071 | -0.40804 | -0.43105 | 0.20973 | 3.11831 | 0.02348 | 0.001489978 | 0.10458 | 0.00163 | 0.00089 |
| NCIH647 | -0.3296 | 0.196404 | -0.1837 | 0.049413 | 0.20957 | 3.11541 | 0.023573 | 0.009711315 | 0.08581 | 0.1008 | 0.36664 |
| NCCMPNST1C1 | -0.342 | -0.22554 | -0.48209 | -0.46251 | 0.20943 | 3.11271 | 0.023661 | 0.007523867 | 0.05765 | 0.0002 | 0.00036 |
| DIFI | -0.4646 | -0.02156 | -0.12493 | -0.00295 | 0.20931 | 3.1104 | 0.023736 | 0.000338633 | 0.44094 | 0.19367 | 0.49189 |
| HCS2 | -0.3931 | 0.113472 | -0.18942 | -0.20923 | 0.20892 | 3.10311 | 0.023974 | 0.002373912 | 0.21634 | 0.09383 | 0.07239 |
| NCIH2023 | -0.4078 | -0.03983 | -0.3761 | -0.26847 | 0.20877 | 3.10032 | 0.024066 | 0.001641861 | 0.39181 | 0.00355 | 0.02971 |

| | | | | | | | | | | | |
|---|---|---|---|---|---|---|---|---|---|---|---|
| RVH421SKINFV1 | -0.5071 | -0.15135 | -0.2278 | -0.23738 | 0.20866 | 3.09824 | 0.024135 | 8.56471E-05 | 0.14705 | 0.0558 | 0.04847 |
| NCIH1792 | -0.4017 | 0.01928 | -0.31307 | -0.27026 | 0.20846 | 3.09449 | 0.024259 | 0.001916542 | 0.44714 | 0.01342 | 0.02884 |
| NMB | -0.3102 | 0.003062 | -0.42326 | -0.29314 | 0.20836 | 3.09267 | 0.02432 | 0.014170163 | 0.49158 | 0.0011 | 0.01941 |
| NCIH1755 | -0.3626 | -0.02116 | -0.4087 | -0.27152 | 0.20827 | 3.09089 | 0.024379 | 0.004825984 | 0.44202 | 0.00161 | 0.02824 |
| SUM185PE | -0.3237 | 0.034274 | -0.39698 | -0.24666 | 0.20823 | 3.09018 | 0.024402 | 0.010913807 | 0.4066 | 0.00216 | 0.0421 |
| SHI1 | -0.3089 | -0.18431 | -0.51821 | -0.38672 | 0.20822 | 3.08994 | 0.024411 | 0.014535113 | 0.10004 | 5.8E-05 | 0.00277 |
| HTMMT | -0.421 | 0.023378 | -0.25235 | -0.27804 | 0.20808 | 3.08743 | 0.024495 | 0.001165593 | 0.43599 | 0.03853 | 0.02529 |
| HCC1143 | -0.4055 | 0.132516 | 0.031465 | -0.10774 | 0.20782 | 3.08246 | 0.024662 | 0.001742676 | 0.17947 | 0.41414 | 0.22822 |
| SKPNDW | -0.2963 | -0.14254 | -0.40195 | 0.032135 | 0.20674 | 3.06224 | 0.025356 | 0.018355298 | 0.16171 | 0.00191 | 0.41234 |
| NCIH1703 | -0.3717 | 0.072303 | -0.2767 | -0.26363 | 0.20593 | 3.0471 | 0.025888 | 0.003928934 | 0.3089 | 0.02587 | 0.03217 |
| SHMAC5 | -0.4844 | -0.05164 | -0.12423 | -0.08953 | 0.20552 | 3.03952 | 0.026159 | 0.000182217 | 0.36087 | 0.19501 | 0.26818 |
| UW228 | -0.1776 | 0.225998 | -0.16639 | -0.28078 | 0.2052 | 3.03356 | 0.026374 | 0.108581213 | 0.05727 | 0.12407 | 0.02413 |
| SKUT1 | -0.4289 | 0.074363 | -0.16665 | -0.17812 | 0.20489 | 3.02788 | 0.02658 | 0.000941762 | 0.30389 | 0.12369 | 0.10793 |
| M07E | -0.4384 | -0.22267 | -0.43557 | -0.29768 | 0.20456 | 3.02173 | 0.026805 | 0.000724939 | 0.06006 | 0.00078 | 0.01788 |
| COV362 | -0.2225 | 0.184689 | -0.26744 | -0.24864 | 0.20404 | 3.01211 | 0.027162 | 0.060218083 | 0.09957 | 0.03022 | 0.04083 |
| PACADD161 | -0.3961 | 0.05732 | -0.2918 | -0.15508 | 0.20399 | 3.01119 | 0.027196 | 0.002204756 | 0.34628 | 0.01988 | 0.14111 |
| NCIH2286 | -0.2999 | 0.040225 | -0.39825 | -0.25859 | 0.20395 | 3.01045 | 0.027224 | 0.017164177 | 0.39075 | 0.00209 | 0.0349 |
| SUM229PE | -0.4614 | -0.10812 | -0.33584 | -0.20361 | 0.20395 | 3.01033 | 0.027228 | 0.000372055 | 0.22742 | 0.00855 | 0.07806 |
| NCIH727 | -0.4414 | -0.03404 | -0.27673 | -0.07801 | 0.20392 | 3.00975 | 0.02725 | 0.000666931 | 0.40722 | 0.02586 | 0.29512 |
| OCUM1 | -0.2359 | 0.227531 | -0.20314 | 0.136896 | 0.20364 | 3.0046 | 0.027443 | 0.049520292 | 0.05602 | 0.07855 | 0.17157 |
| OMM25 | -0.3762 | 0.101874 | -0.24872 | -0.0368 | 0.20357 | 3.00327 | 0.027493 | 0.003542733 | 0.24072 | 0.04078 | 0.39986 |
| MKN74 | -0.3569 | 0.069905 | -0.32814 | -0.11594 | 0.2032 | 2.99643 | 0.027753 | 0.005470414 | 0.31476 | 0.01 | 0.21134 |
| KNS62 | -0.4068 | 0.110955 | -0.06797 | -0.18327 | 0.20305 | 2.99377 | 0.027855 | 0.001686521 | 0.22151 | 0.31953 | 0.10134 |
| JHH6 | -0.3604 | 0.058558 | -0.32655 | -0.22873 | 0.20294 | 2.99163 | 0.027937 | 0.005065423 | 0.34313 | 0.01032 | 0.05505 |
| JHOS4 | -0.3778 | 0.184507 | -0.02001 | -0.08877 | 0.20293 | 2.99144 | 0.027944 | 0.00341566 | 0.0998 | 0.44514 | 0.26994 |
| EFO21 | -0.1735 | 0.21747 | -0.3106 | -0.02389 | 0.20286 | 2.99027 | 0.027989 | 0.114101558 | 0.06462 | 0.01407 | 0.43459 |
| NTERA2CLD1 | -0.4313 | -0.03294 | -0.24385 | -0.01297 | 0.20228 | 2.97951 | 0.028406 | 0.000883297 | 0.41017 | 0.04396 | 0.46438 |
| NCIH838 | -0.285 | 0.083125 | -0.35196 | -0.272 | 0.20204 | 2.97503 | 0.028581 | 0.022435582 | 0.28302 | 0.0061 | 0.02801 |
| LCAM1 | -0.1928 | -0.19726 | -0.52006 | -0.43204 | 0.20177 | 2.97001 | 0.028779 | 0.089855429 | 0.08486 | 5.4E-05 | 0.00086 |
| DBTRG05MG | 0.01706 | 0.307791 | -0.2466 | -0.05288 | 0.20151 | 2.96536 | 0.028964 | 0.453193981 | 0.01484 | 0.04214 | 0.35767 |
| T3M5 | -0.4222 | 0.00151 | -0.30347 | -0.15712 | 0.20151 | 2.96526 | 0.028967 | 0.001127355 | 0.49585 | 0.01608 | 0.13792 |
| IHH4 | -0.3559 | -0.12458 | -0.44324 | -0.1237 | 0.20135 | 2.96234 | 0.029084 | 0.005601767 | 0.19435 | 0.00063 | 0.19604 |
| MM1S | -0.3144 | -0.09171 | -0.47773 | -0.1929 | 0.2009 | 2.95405 | 0.029417 | 0.013085273 | 0.26323 | 0.00023 | 0.08977 |
| HT144SKINFV2 | -0.3963 | 0.14027 | -0.1064 | 0.005892 | 0.20083 | 2.95276 | 0.02947 | 0.002192707 | 0.16563 | 0.23104 | 0.4838 |
| SU8686 | -0.3132 | 0.188127 | -0.22959 | -0.0613 | 0.20054 | 2.9474 | 0.029688 | 0.013380016 | 0.09538 | 0.05437 | 0.33618 |
| DU145 | -0.3683 | 0.115129 | -0.24127 | -0.17302 | 0.20023 | 2.9417 | 0.029921 | 0.004250537 | 0.21297 | 0.04571 | 0.11476 |
| MKN45 | -0.35 | 0.071978 | -0.27902 | -0.28179 | 0.19928 | 2.92426 | 0.030648 | 0.006364268 | 0.30969 | 0.02487 | 0.02371 |
| UACC62 | -0.3403 | 0.099475 | -0.27286 | -0.24821 | 0.19922 | 2.92314 | 0.030695 | 0.007808734 | 0.24594 | 0.02761 | 0.0411 |
| TC138 | -0.3146 | 0.045489 | -0.38461 | -0.19984 | 0.19856 | 2.91104 | 0.031211 | 0.013047717 | 0.37688 | 0.00291 | 0.08205 |
| KP4 | -0.2018 | 0.018069 | -0.43985 | -0.30259 | 0.19825 | 2.90551 | 0.031449 | 0.079999745 | 0.45044 | 0.0007 | 0.01634 |
| NCIH1694 | -0.4005 | 0.042051 | -0.28092 | -0.20306 | 0.19813 | 2.90332 | 0.031544 | 0.001977298 | 0.38593 | 0.02407 | 0.07864 |
| SW1088 | -0.2531 | 0.105023 | -0.30502 | -0.31239 | 0.1975 | 2.89172 | 0.032051 | 0.038065932 | 0.23396 | 0.01563 | 0.0136 |
| DLD1 | -0.4107 | 0.042651 | -0.256 | -0.09509 | 0.19702 | 2.88298 | 0.032439 | 0.001523857 | 0.38434 | 0.03637 | 0.25564 |
| BCPAP | -0.3133 | 0.099259 | -0.32724 | -0.07431 | 0.19698 | 2.8822 | 0.032474 | 0.013353583 | 0.24642 | 0.01018 | 0.30401 |
| ES2 | -0.378 | 0.076572 | -0.2789 | -0.12183 | 0.19678 | 2.87864 | 0.032634 | 0.003399474 | 0.29857 | 0.02492 | 0.19966 |
| 8MGBA | -0.3656 | 0.032971 | -0.34271 | -0.19395 | 0.19634 | 2.87054 | 0.033 | 0.00451403 | 0.41009 | 0.00742 | 0.08857 |
| MDAMB468 | -0.3287 | 0.134558 | -0.16904 | -0.26617 | 0.19617 | 2.86757 | 0.033135 | 0.009884391 | 0.17576 | 0.12029 | 0.03086 |
| HUH6 | -0.2782 | -0.00363 | -0.42814 | -0.29114 | 0.19603 | 2.86489 | 0.033258 | 0.025225024 | 0.49002 | 0.00096 | 0.02012 |
| NCIH1299 | -0.2795 | 0.159212 | -0.29289 | -0.05038 | 0.19602 | 2.86486 | 0.033259 | 0.024663757 | 0.13471 | 0.0195 | 0.36412 |
| OCIMY5 | -0.272 | 0.105596 | -0.3518 | -0.07041 | 0.19531 | 2.8519 | 0.033858 | 0.027988563 | 0.23275 | 0.00612 | 0.31352 |
| D423MG | -0.3748 | 0.172234 | -0.09504 | -0.06094 | 0.19505 | 2.84717 | 0.03408 | 0.003659734 | 0.11584 | 0.25574 | 0.3371 |
| COGE352 | -0.3562 | 0.024089 | -0.31738 | -0.30311 | 0.19461 | 2.83924 | 0.034454 | 0.005556726 | 0.43406 | 0.01236 | 0.01619 |
| MERO82 | -0.208 | 0.200786 | -0.07787 | -0.29566 | 0.19445 | 2.83632 | 0.034593 | 0.073613547 | 0.08103 | 0.29547 | 0.01855 |
| HOUAI | -0.422 | 0.087635 | 1.02E-05 | -0.04575 | 0.19443 | 2.83587 | 0.034615 | 0.001136151 | 0.27253 | 0.49997 | 0.3762 |
| CJM | -0.3127 | 0.227602 | -0.13666 | -0.06651 | 0.19424 | 2.83243 | 0.034779 | 0.01351312 | 0.05596 | 0.17199 | 0.32316 |
| HEC265 | -0.3835 | 0.167318 | -0.02298 | -0.00049 | 0.19418 | 2.83144 | 0.034826 | 0.002988363 | 0.12274 | 0.43708 | 0.49866 |
| SW48 | -0.4299 | 0.02327 | -0.22395 | -0.1958 | 0.19398 | 2.82785 | 0.034999 | 0.000917137 | 0.43628 | 0.05898 | 0.08648 |
| OUMS23 | -0.3666 | -0.01279 | -0.37672 | -0.22677 | 0.19381 | 2.82477 | 0.035148 | 0.004411502 | 0.46488 | 0.0035 | 0.05664 |
| WSUNHL | -0.413 | 0.021969 | -0.22488 | -0.02171 | 0.19349 | 2.81893 | 0.035432 | 0.001437105 | 0.43982 | 0.0582 | 0.44052 |
| BOKU | -0.3544 | -0.04848 | -0.40941 | -0.27575 | 0.1932 | 2.81366 | 0.03569 | 0.005787965 | 0.36906 | 0.00158 | 0.02629 |
| HEC50B | -0.334 | 0.141978 | -0.25007 | -0.10323 | 0.1931 | 2.81195 | 0.035775 | 0.008888835 | 0.16267 | 0.03993 | 0.23779 |
| OSRC2 | -0.2475 | 0.300962 | -0.10046 | 0.043182 | 0.19307 | 2.81129 | 0.035807 | 0.041562692 | 0.01684 | 0.2438 | 0.38294 |
| U343 | -0.208 | 0.173117 | -0.12863 | -0.32154 | 0.19272 | 2.80504 | 0.036117 | 0.073646396 | 0.11463 | 0.18666 | 0.01139 |
| TC32 | -0.1663 | 0.095586 | -0.39926 | -0.22044 | 0.19253 | 2.80171 | 0.036284 | 0.124227601 | 0.25453 | 0.00204 | 0.06198 |
| JM1 | -0.3851 | 0.014668 | -0.31399 | -0.24094 | 0.19151 | 2.78319 | 0.037222 | 0.002873274 | 0.45974 | 0.01319 | 0.04594 |
| NCIH1092 | -0.398 | -0.08911 | -0.39123 | -0.19164 | 0.19148 | 2.78269 | 0.037248 | 0.002103927 | 0.26915 | 0.00248 | 0.09123 |
| NCIH1993 | -0.2668 | 0.045178 | -0.40017 | -0.10212 | 0.19126 | 2.77871 | 0.037453 | 0.030514502 | 0.3777 | 0.00199 | 0.24019 |
| PECAPJ41CLONED2 | -0.4538 | -0.03795 | -0.1957 | -0.24388 | 0.19117 | 2.77719 | 0.037531 | 0.000466815 | 0.39678 | 0.08659 | 0.04394 |
| KINGS1 | -0.0368 | 0.051851 | -0.44429 | -0.14421 | 0.191 | 2.77414 | 0.03769 | 0.399935006 | 0.36032 | 0.00061 | 0.15886 |
| SKGI | -0.3908 | -0.01588 | -0.12682 | -0.35086 | 0.191 | 2.77411 | 0.037691 | 0.002505529 | 0.45642 | 0.19007 | 0.00624 |
| AM38 | -0.2871 | 0.047764 | -0.37205 | -0.26691 | 0.19082 | 2.77088 | 0.037859 | 0.021593619 | 0.37093 | 0.0039 | 0.03048 |
| COLO678 | -0.4425 | 0.026231 | -0.15439 | -0.12383 | 0.19028 | 2.76117 | 0.03837 | 0.000645872 | 0.42825 | 0.14219 | 0.19579 |
| MM160113 | -0.4566 | -0.23085 | -0.35395 | -0.21897 | 0.19007 | 2.7575 | 0.038565 | 0.000429955 | 0.05338 | 0.00584 | 0.06328 |
| G415 | -0.3088 | 0.191067 | -0.2001 | -0.09819 | 0.18961 | 2.74728 | 0.039113 | 0.014551631 | 0.0919 | 0.08177 | 0.24877 |
| SNU620 | -0.4569 | -0.14345 | -0.32223 | -0.27028 | 0.18937 | 2.74486 | 0.039243 | 0.000425925 | 0.16015 | 0.01124 | 0.02883 |
| KU812 | -0.1788 | 0.317213 | -0.15027 | 0.032386 | 0.18928 | 2.7432 | 0.039333 | 0.107102371 | 0.0124 | 0.14879 | 0.41166 |
| RPE1SS77 | -0.3078 | 0.091069 | -0.32938 | -0.17568 | 0.18907 | 2.73957 | 0.039531 | 0.014823284 | 0.26468 | 0.00975 | 0.11117 |
| OCILY18 | -0.4274 | -0.01608 | -0.19631 | -0.01257 | 0.18904 | 2.73906 | 0.039559 | 0.000981781 | 0.45588 | 0.08592 | 0.46547 |
| 94T778 | -0.2029 | 0.22778 | -0.19312 | -0.209 | 0.18893 | 2.73699 | 0.039672 | 0.078790946 | 0.05582 | 0.08952 | 0.07262 |
| MESOV | -0.3345 | 0.107976 | -0.26889 | -0.03285 | 0.18887 | 2.73602 | 0.039725 | 0.008795193 | 0.22772 | 0.0295 | 0.41041 |
| LU99 | -0.4177 | 0.013069 | -0.21532 | -0.2532 | 0.1887 | 2.73293 | 0.039895 | 0.001271798 | 0.46411 | 0.06659 | 0.03802 |
| SAOS2 | -0.2847 | 0.20472 | -0.18574 | 0.060188 | 0.18865 | 2.73204 | 0.039944 | 0.022540761 | 0.07692 | 0.09828 | 0.33899 |
| ISTMES2 | -0.3817 | 0.048975 | -0.19756 | -0.28488 | 0.18862 | 2.73156 | 0.03997 | 0.003116887 | 0.36778 | 0.08453 | 0.02247 |
| MAC2A | -0.3287 | -0.03329 | -0.41682 | -0.18095 | 0.18834 | 2.72653 | 0.040249 | 0.009892916 | 0.40925 | 0.0013 | 0.10428 |
| RKO | -0.3418 | 0.070649 | -0.30335 | -0.06267 | 0.18798 | 2.72007 | 0.04061 | 0.007563362 | 0.31294 | 0.01612 | 0.33275 |
| C99 | -0.3217 | 0.226495 | 0.033926 | -0.18174 | 0.18765 | 2.71417 | 0.040942 | 0.011369683 | 0.05686 | 0.28158 | 0.40753 |
| NO10 | -0.3152 | 0.236388 | -0.0574 | -0.06823 | 0.18746 | 2.7109 | 0.041127 | 0.012893602 | 0.04919 | 0.34607 | 0.3189 |
| HCC1395 | -0.1424 | 0.28832 | -0.20751 | -0.07328 | 0.18745 | 2.71073 | 0.041137 | 0.161911749 | 0.02115 | 0.07409 | 0.30651 |
| SNGM | -0.3681 | 0.160093 | -0.06522 | -0.12176 | 0.18736 | 2.70913 | 0.041228 | 0.004272365 | 0.13337 | 0.32637 | 0.19981 |
| ECC4 | -0.2477 | -0.24069 | -0.5297 | -0.25003 | 0.18709 | 2.70428 | 0.041505 | 0.041410668 | 0.04611 | 3.8E-05 | 0.03996 |
| SNU475 | -0.2909 | 0.216002 | 0.035855 | -0.16386 | 0.18685 | 2.69994 | 0.041754 | 0.02021838 | 0.06596 | 0.40238 | 0.12776 |
| M040416 | -0.3955 | -0.24596 | -0.44384 | -0.25098 | 0.18604 | 2.68552 | 0.042594 | 0.002238187 | 0.04256 | 0.00062 | 0.03937 |
| LCLC97TM1 | -0.3695 | -0.00712 | -0.35353 | -0.20946 | 0.18599 | 2.68468 | 0.042643 | 0.004130228 | 0.48044 | 0.00589 | 0.07216 |
| KPNYN | -0.1333 | -0.03444 | -0.47377 | -0.28397 | 0.18593 | 2.68373 | 0.0427 | 0.177980282 | 0.40617 | 0.00026 | 0.02283 |
| FLO1 | -0.2371 | 0.129635 | -0.33413 | -0.17841 | 0.18562 | 2.67813 | 0.043031 | 0.048682146 | 0.18479 | 0.00886 | 0.10756 |
| NCIH1693 | -0.3621 | 0.131571 | -0.15092 | -0.17795 | 0.18549 | 2.67578 | 0.043171 | 0.004880648 | 0.1812 | 0.14773 | 0.10816 |
| MJ | -0.4396 | -0.30262 | -0.18295 | -0.10512 | 0.18512 | 2.66932 | 0.043558 | 0.000702037 | 0.01634 | 0.10174 | 0.23375 |
| L363 | -0.3304 | 0.099415 | -0.29006 | -0.146 | 0.18504 | 2.66786 | 0.043646 | 0.009554928 | 0.24608 | 0.02051 | 0.15585 |
| SKNBE2 | -0.2169 | -0.14381 | -0.51531 | -0.24297 | 0.18478 | 2.66329 | 0.043923 | 0.065150229 | 0.15953 | 6.4E-05 | 0.04455 |
| NCIH441 | -0.3664 | 0.032521 | -0.31318 | -0.11883 | 0.1842 | 2.65299 | 0.044552 | 0.004438536 | 0.4113 | 0.0134 | 0.20556 |
| PANC0203 | -0.3472 | 0.115784 | -0.23836 | -0.05846 | 0.18406 | 2.65055 | 0.044703 | 0.00675346 | 0.21165 | 0.04776 | 0.34338 |
| SKMEL2 | -0.2771 | 0.020763 | -0.40723 | -0.16191 | 0.18402 | 2.6499 | 0.044743 | 0.025684812 | 0.4431 | 0.00167 | 0.13065 |
| MEL270 | -0.2439 | 0.232216 | -0.11142 | 0.183465 | 0.18402 | 2.64982 | 0.044747 | 0.043911572 | 0.05232 | 0.22056 | 0.10109 |
| LP1 | 0.02867 | 0.170285 | -0.34058 | 0.038512 | 0.1838 | 2.64595 | 0.044988 | 0.421674876 | 0.11854 | 0.00776 | 0.3953 |
| NCIH2087 | -0.3207 | -0.00804 | -0.37456 | -0.2975 | 0.18325 | 2.63633 | 0.045589 | 0.011584562 | 0.4779 | 0.00368 | 0.01794 |
| BL70 | -0.2952 | -0.28315 | -0.42252 | -0.5009 | 0.18274 | 2.62723 | 0.046167 | 0.018723182 | 0.02316 | 0.00112 | 0.00011 |
| KELLY | -0.2521 | -0.00334 | -0.43455 | -0.20689 | 0.1825 | 2.62302 | 0.046436 | 0.038658444 | 0.49082 | 0.00081 | 0.07471 |
| U118MG | -0.2761 | -0.11551 | -0.35995 | -0.46821 | 0.18243 | 2.62186 | 0.04651 | 0.026144293 | 0.21219 | 0.00512 | 0.0003 |
| WM115 | -0.4081 | 0.085044 | -0.11302 | -0.11686 | 0.18229 | 2.61933 | 0.046674 | 0.001629369 | 0.27853 | 0.21727 | 0.20949 |
| HEPG2 | -0.1492 | 0.207659 | -0.30003 | -0.0859 | 0.18214 | 2.61678 | 0.046839 | 0.150551101 | 0.07394 | 0.01713 | 0.27655 |
| SHP77 | -0.2014 | 0.007412 | -0.29511 | -0.42851 | 0.18214 | 2.61674 | 0.046841 | 0.08034087 | 0.47963 | 0.01874 | 0.00095 |
| SLR26 | -0.203 | 0.323145 | -0.08544 | 0.004289 | 0.18183 | 2.61138 | 0.047189 | 0.078718982 | 0.01104 | 0.27761 | 0.48821 |
| SNU449 | -0.3309 | -0.00566 | -0.37588 | -0.25419 | 0.18165 | 2.60811 | 0.047403 | 0.009455222 | 0.48445 | 0.00357 | 0.03743 |
| CCLFUPGI0005T | -0.3962 | -0.00281 | -0.08075 | -0.29502 | 0.18131 | 2.60218 | 0.047794 | 0.002196236 | 0.49229 | 0.28862 | 0.01877 |
| BT549 | -0.1186 | 0.260679 | -0.14591 | -0.21584 | 0.1813 | 2.60209 | 0.0478 | 0.206021886 | 0.03374 | 0.15599 | 0.06611 |
| LS123 | -0.3807 | 0.071562 | -0.01718 | 0.111312 | 0.18115 | 2.59945 | 0.047974 | 0.003193723 | 0.3107 | 0.45287 | 0.22077 |
| RVH421 | -0.3176 | 0.192979 | -0.15616 | -0.06807 | 0.18049 | 2.5879 | 0.048746 | 0.012307407 | 0.08968 | 0.13941 | 0.31929 |
| U251MG | -0.2439 | 0.247759 | -0.16296 | -0.09102 | 0.18049 | 2.5878 | 0.048753 | 0.043927532 | 0.04139 | 0.12908 | 0.26479 |
| MPP89 | -0.3825 | 0.017896 | -0.23208 | -0.28934 | 0.18038 | 2.58588 | 0.048883 | 0.003058437 | 0.45091 | 0.05243 | 0.02077 |

| | | | | | | | | | | | |
|---|---|---|---|---|---|---|---|---|---|---|---|
| NB69 | -0.3102 | 0.161657 | -0.22282 | -0.05896 | 0.18001 | 2.57941 | 0.049322 | 0.01418906 | 0.13102 | 0.05993 | 0.3421 |
| JVE253 | -0.3721 | -0.15566 | -0.43213 | -0.2847 | 0.17965 | 2.57309 | 0.049755 | 0.003898277 | 0.1402 | 0.00086 | 0.02254 |
| RDES | -0.2291 | 0.115105 | -0.2812 | -0.29168 | 0.17936 | 2.56815 | 0.050096 | 0.054743273 | 0.21302 | 0.02395 | 0.01993 |
| SF126 | -0.2546 | 0.041523 | -0.23009 | -0.39333 | 0.17924 | 2.56595 | 0.050249 | 0.037171343 | 0.38732 | 0.05397 | 0.00236 |
| CHLA15 | -0.1386 | 0.083708 | -0.40599 | -0.10611 | 0.17923 | 2.56575 | 0.050263 | 0.168513679 | 0.28165 | 0.00172 | 0.23165 |
| SNU739 | -0.3244 | 0.106858 | -0.24616 | -0.21731 | 0.1792 | 2.56527 | 0.050296 | 0.010767325 | 0.23007 | 0.04242 | 0.06477 |
| KMRC1 | -0.1124 | 0.27188 | -0.13629 | -0.19932 | 0.17889 | 2.55992 | 0.050669 | 0.218495951 | 0.02807 | 0.17265 | 0.08261 |
| C8166 | -0.3916 | -0.12797 | -0.28221 | -0.40691 | 0.17886 | 2.55942 | 0.050705 | 0.002459874 | 0.18791 | 0.02354 | 0.00168 |
| NCIH358 | -0.3699 | 0.082338 | -0.231 | -0.07496 | 0.17884 | 2.55894 | 0.050738 | 0.004092808 | 0.28486 | 0.05326 | 0.30244 |
| CHAGOK1 | -0.3809 | 0.083629 | -0.20378 | -0.14272 | 0.17813 | 2.5466 | 0.051612 | 0.00317812 | 0.28183 | 0.07788 | 0.1614 |
| HCC1428 | -0.2965 | -0.07123 | -0.32703 | -0.43163 | 0.178 | 2.54433 | 0.051774 | 0.01827539 | 0.31152 | 0.01022 | 0.00087 |
| MEL290 | -0.2528 | 0.156127 | -0.24324 | -0.22443 | 0.17788 | 2.54223 | 0.051924 | 0.038234349 | 0.13947 | 0.04437 | 0.05857 |
| NCIH211 | -0.3608 | 0.07356 | -0.19697 | -0.24985 | 0.17781 | 2.54116 | 0.052002 | 0.005022054 | 0.30584 | 0.08518 | 0.04007 |
| EVSAT | -0.2799 | 0.144983 | -0.13835 | -0.27368 | 0.17775 | 2.54014 | 0.052075 | 0.024500962 | 0.15755 | 0.16899 | 0.02723 |
| HT1197 | -0.2581 | 0.284646 | -0.06051 | 0.040182 | 0.17734 | 2.53298 | 0.052594 | 0.035160945 | 0.02256 | 0.33817 | 0.39087 |
| CL11 | -0.4021 | 0.054837 | 0.042137 | -0.08433 | 0.17693 | 2.52582 | 0.053117 | 0.001897932 | 0.35263 | 0.3857 | 0.28019 |
| EWS502 | -0.3994 | -0.06292 | -0.32818 | -0.26657 | 0.17655 | 2.51919 | 0.053606 | 0.002030144 | 0.33211 | 0.00999 | 0.03065 |
| SNU423 | -0.314 | -0.08748 | -0.43321 | -0.29641 | 0.17654 | 2.51903 | 0.053618 | 0.013184871 | 0.27288 | 0.00084 | 0.0183 |
| A673 | -0.3919 | -0.18108 | -0.38706 | -0.15065 | 0.17654 | 2.519 | 0.05362 | 0.002441484 | 0.10411 | 0.00274 | 0.14817 |
| MS751 | -0.374 | -0.0201 | -0.22771 | -0.3427 | 0.1763 | 2.51482 | 0.053931 | 0.003726384 | 0.44489 | 0.05587 | 0.00742 |
| NGP | -0.3452 | -0.0047 | -0.30961 | -0.31626 | 0.17614 | 2.51207 | 0.054137 | 0.007039807 | 0.48707 | 0.01434 | 0.01263 |
| NCO2 | -0.3217 | -0.13066 | -0.38749 | -0.04241 | 0.17597 | 2.50919 | 0.054354 | 0.011366012 | 0.18288 | 0.00272 | 0.38498 |
| LB771HNC | -0.2764 | 0.077667 | -0.3422 | -0.08258 | 0.17593 | 2.50855 | 0.054402 | 0.026019716 | 0.29594 | 0.0075 | 0.28429 |
| LPS853 | -0.3842 | 0.04309 | -0.2509 | -0.1662 | 0.17569 | 2.50426 | 0.054725 | 0.002941151 | 0.38318 | 0.03941 | 0.12435 |
| SUDHL5 | -0.2049 | 0.024199 | -0.25594 | -0.41611 | 0.17539 | 2.49917 | 0.055112 | 0.076779654 | 0.43376 | 0.03641 | 0.00133 |
| HCC827 | -0.3171 | 0.185105 | -0.12845 | -0.12508 | 0.1752 | 2.49582 | 0.055368 | 0.012419222 | 0.09906 | 0.187 | 0.19338 |
| RT112 | -0.1422 | 0.128538 | -0.25553 | -0.32404 | 0.17518 | 2.49561 | 0.055384 | 0.162254164 | 0.18684 | 0.03664 | 0.01085 |
| HCC827GR5 | -0.3401 | 0.043762 | -0.3182 | -0.15477 | 0.17501 | 2.49252 | 0.055622 | 0.007827289 | 0.38142 | 0.01216 | 0.14159 |
| SHSY5Y | -0.1963 | -0.03878 | -0.45576 | -0.13314 | 0.17392 | 2.47381 | 0.057081 | 0.085952049 | 0.39459 | 0.00044 | 0.17832 |
| RD | -0.2521 | 0.040504 | -0.36946 | -0.04138 | 0.17382 | 2.47208 | 0.057217 | 0.038685352 | 0.39002 | 0.00414 | 0.38769 |
| UMUC5 | -0.3102 | 0.19878 | -0.13078 | -0.07028 | 0.1732 | 2.46142 | 0.058068 | 0.014166388 | 0.0832 | 0.18266 | 0.31384 |
| SLR20 | -0.2681 | 0.059333 | -0.36477 | -0.16115 | 0.17319 | 2.46129 | 0.058079 | 0.02990102 | 0.34116 | 0.0046 | 0.13178 |
| NP2 | -0.1803 | 0.192548 | -0.25146 | -0.19416 | 0.1729 | 2.45619 | 0.05849 | 0.105151016 | 0.09018 | 0.03907 | 0.08833 |
| CAL51 | -0.3809 | 0.026021 | -0.1281 | -0.27987 | 0.17265 | 2.45201 | 0.058829 | 0.003173803 | 0.42882 | 0.18766 | 0.02451 |
| EM2 | -0.4002 | -0.1399 | -0.37813 | -0.25117 | 0.17263 | 2.45171 | 0.058853 | 0.001991127 | 0.16627 | 0.00339 | 0.03925 |
| A375SKINCJ2 | -0.3655 | 0.071131 | -0.22926 | -0.06553 | 0.17136 | 2.42991 | 0.060656 | 0.004521038 | 0.31176 | 0.05463 | 0.32558 |
| HEC116 | -0.3939 | -0.03366 | -0.30406 | -0.14498 | 0.17119 | 2.4269 | 0.060909 | 0.002326691 | 0.40825 | 0.01591 | 0.15756 |
| MALME3M | -0.2846 | 0.12788 | -0.27243 | -0.04448 | 0.17105 | 2.42455 | 0.061108 | 0.02257562 | 0.18807 | 0.02781 | 0.37952 |
| EN | -0.4236 | -0.04113 | -0.23025 | -0.23331 | 0.171 | 2.42434 | 0.061126 | 0.001086779 | 0.38836 | 0.05385 | 0.05149 |
| HG3 | -0.3811 | 0.084586 | -0.1763 | -0.10318 | 0.17099 | 2.42359 | 0.061189 | 0.003159623 | 0.2796 | 0.11034 | 0.23792 |
| MM253 | -0.376 | -0.10253 | -0.29007 | -0.01303 | 0.17091 | 2.4221 | 0.061315 | 0.00356381 | 0.23931 | 0.02051 | 0.46422 |
| SNU5 | -0.3919 | -0.11308 | 0.08214 | -0.02651 | 0.17086 | 2.42124 | 0.061388 | 0.002438991 | 0.21714 | 0.28533 | 0.42749 |
| UMUC6 | -0.3487 | 0.129027 | -0.13101 | -0.16863 | 0.17084 | 2.42105 | 0.061405 | 0.006541485 | 0.18592 | 0.18224 | 0.12088 |
| C10 | -0.0527 | 0.312688 | -0.07123 | -0.16312 | 0.17081 | 2.42052 | 0.061449 | 0.358033727 | 0.01352 | 0.31151 | 0.12885 |
| NOS1 | -0.2352 | 0.039245 | -0.39305 | -0.11977 | 0.17079 | 2.42019 | 0.061478 | 0.050032782 | 0.39335 | 0.00237 | 0.2037 |
| HO1U1 | -0.3409 | 0.184426 | -0.01114 | -0.04348 | 0.17055 | 2.41598 | 0.061837 | 0.007700472 | 0.0999 | 0.46941 | 0.38215 |
| SF295 | -0.2422 | 0.083779 | -0.35527 | -0.14634 | 0.17037 | 2.41289 | 0.062102 | 0.045104386 | 0.28148 | 0.00567 | 0.15527 |
| HCC366 | -0.3475 | 0.026909 | -0.28019 | -0.26655 | 0.17017 | 2.40955 | 0.06239 | 0.006708009 | 0.42642 | 0.02437 | 0.03066 |
| LU165 | -0.4257 | -0.11603 | -0.31233 | -0.22125 | 0.17004 | 2.40735 | 0.06258 | 0.001028843 | 0.21115 | 0.01362 | 0.06128 |
| 697 | -0.4192 | -0.08165 | -0.27207 | -0.27928 | 0.16996 | 2.40598 | 0.062699 | 0.001221834 | 0.28649 | 0.02798 | 0.02476 |
| F5 | -0.3368 | 0.159839 | -0.14097 | -0.06247 | 0.16981 | 2.40339 | 0.062924 | 0.008381818 | 0.13376 | 0.16442 | 0.33323 |
| TTC642 | -0.2227 | -0.05333 | -0.42177 | -0.34644 | 0.16972 | 2.40184 | 0.063059 | 0.060015748 | 0.35651 | 0.00114 | 0.00686 |
| SH10TC | -0.3694 | 0.099789 | -0.16632 | -0.11784 | 0.16835 | 2.37848 | 0.065132 | 0.004144141 | 0.24526 | 0.12418 | 0.20752 |
| KASUMI1 | -0.316 | -0.10808 | -0.20949 | -0.44775 | 0.16795 | 2.37168 | 0.065748 | 0.012688616 | 0.22751 | 0.07213 | 0.00056 |
| CHP212 | -0.0072 | -0.21722 | -0.3236 | 0.115197 | 0.16774 | 2.36818 | 0.066067 | 0.480240138 | 0.06485 | 0.01094 | 0.21284 |
| T24 | -0.2945 | 0.206395 | -0.0821 | -0.12168 | 0.16773 | 2.36802 | 0.066081 | 0.018936687 | 0.07521 | 0.28543 | 0.19996 |
| SJSA1 | -0.2401 | -0.05037 | -0.44316 | -0.17068 | 0.16765 | 2.3667 | 0.066203 | 0.046558914 | 0.36415 | 0.00063 | 0.11799 |
| A375 | -0.2077 | 0.083691 | -0.36878 | -0.13919 | 0.16758 | 2.36549 | 0.066313 | 0.073858377 | 0.28169 | 0.0042 | 0.16751 |
| SW982 | -0.2992 | 0.187543 | -0.14382 | -0.11433 | 0.16757 | 2.36534 | 0.066327 | 0.017387183 | 0.09608 | 0.15951 | 0.2146 |
| CHLA10 | -0.3777 | -0.13907 | -0.38064 | -0.31611 | 0.16746 | 2.36349 | 0.066497 | 0.003421793 | 0.16772 | 0.0032 | 0.01266 |
| COV504 | -0.2988 | 0.069437 | -0.29854 | -0.23316 | 0.1674 | 2.36248 | 0.06659 | 0.017522388 | 0.31591 | 0.01761 | 0.0516 |
| HS683 | -0.2698 | 0.048265 | -0.28856 | -0.32166 | 0.16706 | 2.35673 | 0.067122 | 0.029045117 | 0.36962 | 0.02106 | 0.01137 |
| SKNSH | -0.2737 | -0.01672 | -0.38882 | -0.2892 | 0.16687 | 2.35344 | 0.067428 | 0.027223969 | 0.45413 | 0.00263 | 0.02083 |
| NCIH2171 | -0.3165 | 0.015779 | -0.34269 | -0.23198 | 0.16663 | 2.34933 | 0.067813 | 0.012562834 | 0.4567 | 0.00742 | 0.05251 |
| ONDA8 | -0.1809 | 0.103751 | -0.16268 | -0.35748 | 0.16594 | 2.3378 | 0.068905 | 0.104368172 | 0.23668 | 0.1295 | 0.00541 |
| M140325 | -0.4046 | -0.07735 | -0.3111 | -0.19475 | 0.16565 | 2.33274 | 0.069389 | 0.00178034 | 0.2967 | 0.01394 | 0.08766 |
| HS944T | -0.3295 | -0.05506 | -0.38801 | -0.24182 | 0.16552 | 2.3307 | 0.069585 | 0.009737801 | 0.35206 | 0.00268 | 0.04534 |
| SNU478 | -0.35 | 0.152998 | 0.022532 | 0.044511 | 0.16547 | 2.32974 | 0.069677 | 0.006362237 | 0.1444 | 0.43829 | 0.37945 |
| SNU81 | -0.3182 | 0.152607 | -0.18277 | -0.07115 | 0.16539 | 2.32844 | 0.069803 | 0.012152125 | 0.14502 | 0.10196 | 0.3117 |
| JHUEM1 | -0.3749 | 0.077138 | -0.12452 | -0.18733 | 0.16529 | 2.32677 | 0.069964 | 0.003650627 | 0.29721 | 0.19447 | 0.09634 |
| SW626 | -0.262 | 0.01744 | -0.38192 | -0.2291 | 0.16517 | 2.32473 | 0.070162 | 0.033034856 | 0.45216 | 0.0031 | 0.05476 |
| NCIH2882 | -0.3642 | -0.03667 | -0.3371 | -0.22035 | 0.16503 | 2.32241 | 0.070388 | 0.00465522 | 0.4002 | 0.00833 | 0.06206 |
| LNCAPCLONEFGC | -0.2719 | 0.258229 | -0.01683 | -0.02301 | 0.16472 | 2.31716 | 0.070901 | 0.02807796 | 0.0351 | 0.45383 | 0.43698 |
| NCIH2452 | -0.1527 | 0.079472 | -0.18979 | -0.38032 | 0.16432 | 2.31048 | 0.071559 | 0.14489727 | 0.29164 | 0.0934 | 0.00322 |
| CHLA06ATRT | -0.2932 | 0.021563 | -0.3589 | -0.20277 | 0.16398 | 2.30467 | 0.072138 | 0.019375347 | 0.44092 | 0.00524 | 0.07894 |
| GAMG | -0.1902 | 0.130566 | -0.22776 | -0.29689 | 0.16389 | 2.30322 | 0.072282 | 0.092858091 | 0.18306 | 0.05583 | 0.01814 |
| MELJUSO | -0.1247 | 0.104087 | -0.36732 | -0.17238 | 0.16389 | 2.30313 | 0.072291 | 0.194176504 | 0.23596 | 0.00434 | 0.11563 |
| UMUC11 | -0.2699 | 0.106375 | -0.1007 | -0.30317 | 0.16346 | 2.29597 | 0.073011 | 0.029025245 | 0.23109 | 0.24327 | 0.01617 |
| NCIH650 | -0.2101 | -0.03565 | -0.4402 | -0.22085 | 0.16328 | 2.29295 | 0.073317 | 0.071489545 | 0.40292 | 0.00069 | 0.06163 |
| MERO83 | -0.309 | 0.173618 | -0.13443 | -0.11708 | 0.16327 | 2.29283 | 0.073329 | 0.014501918 | 0.11395 | 0.176 | 0.20905 |
| NOMO1 | -0.1475 | -0.07381 | -0.33177 | -0.44729 | 0.16309 | 2.27822 | 0.074827 | 0.153342663 | 0.30523 | 0.00929 | 0.00056 |
| CHLA266 | -0.3794 | -0.12838 | -0.33886 | -0.34014 | 0.16233 | 2.27703 | 0.074951 | 0.003289083 | 0.18713 | 0.00804 | 0.00783 |
| CA922 | -0.3626 | 0.032301 | -0.26671 | -0.14747 | 0.16216 | 2.27408 | 0.075257 | 0.004827993 | 0.41189 | 0.03058 | 0.1534 |
| KYO1 | -0.284 | 0.172226 | 0.155777 | -0.07266 | 0.16147 | 2.26262 | 0.07646 | 0.022810155 | 0.11585 | 0.14001 | 0.30804 |
| G402 | -0.2693 | 0.234486 | 0.023278 | -0.09364 | 0.16132 | 2.26017 | 0.076719 | 0.029279072 | 0.0506 | 0.43626 | 0.25887 |
| CHLA57 | -0.3304 | 0.037142 | -0.28236 | -0.2385 | 0.161 | 2.2547 | 0.077302 | 0.009558424 | 0.39895 | 0.02348 | 0.04766 |
| HCC44 | -0.2621 | 0.130673 | -0.27596 | -0.10622 | 0.16075 | 2.25066 | 0.077736 | 0.032982486 | 0.18286 | 0.0262 | 0.23142 |
| TM31 | -0.3083 | 0.117142 | -0.23795 | -0.12433 | 0.16069 | 2.24956 | 0.077854 | 0.014685439 | 0.20892 | 0.04805 | 0.19483 |
| CAOV3 | -0.1554 | 0.175945 | -0.1542 | -0.27903 | 0.16037 | 2.24433 | 0.07842 | 0.140573053 | 0.11081 | 0.14249 | 0.02487 |
| PANC1 | -0.4151 | -0.04829 | -0.23687 | -0.18297 | 0.16035 | 2.24396 | 0.07846 | 0.001361072 | 0.36955 | 0.04884 | 0.10172 |
| NCIH1581 | -0.2144 | 0.171696 | -0.2527 | -0.14982 | 0.16003 | 2.2386 | 0.079044 | 0.067423228 | 0.11658 | 0.03832 | 0.14953 |
| NCCMPNST2C1 | -0.354 | -0.00014 | -0.30393 | -0.20761 | 0.15991 | 2.23661 | 0.079262 | 0.005835395 | 0.49962 | 0.01595 | 0.07399 |
| SYO1 | -0.1602 | 0.120292 | 0.003976 | -0.32633 | 0.15986 | 2.23577 | 0.079353 | 0.133254317 | 0.20267 | 0.48907 | 0.01037 |
| PANC0504 | -0.4018 | -0.0157 | -0.23144 | -0.1366 | 0.15985 | 2.23557 | 0.079375 | 0.001910613 | 0.45691 | 0.05292 | 0.1721 |
| NIHOVCAR3 | -0.3886 | -0.06035 | -0.25374 | -0.30189 | 0.15978 | 2.23435 | 0.07951 | 0.002646247 | 0.33858 | 0.03769 | 0.01656 |
| OCIAML3 | -0.2039 | -0.35219 | -0.48469 | -0.31892 | 0.15967 | 2.23264 | 0.079698 | 0.077747251 | 0.00607 | 0.00018 | 0.01199 |
| NCIH510 | -0.3357 | -0.00747 | -0.2117 | -0.34089 | 0.15957 | 2.23093 | 0.079886 | 0.008580099 | 0.47948 | 0.06999 | 0.00771 |
| COLO800 | -0.3838 | -0.09271 | -0.3307 | -0.26937 | 0.15953 | 2.2303 | 0.079956 | 0.002966816 | 0.26096 | 0.0095 | 0.02927 |
| S117 | -0.3132 | 0.021584 | -0.30324 | -0.27076 | 0.15949 | 2.22959 | 0.080035 | 0.013387188 | 0.44087 | 0.01615 | 0.0286 |
| HUNS1 | -0.2968 | 0.048814 | -0.26074 | -0.29171 | 0.15896 | 2.22074 | 0.081021 | 0.018181898 | 0.3682 | 0.03371 | 0.01992 |
| JAR | -0.2861 | -0.03909 | -0.36601 | -0.31954 | 0.15811 | 2.20664 | 0.082616 | 0.02201226 | 0.39376 | 0.00447 | 0.01185 |
| 170MGBA | -0.2406 | 0.092731 | -0.15677 | -0.32615 | 0.15801 | 2.20502 | 0.082802 | 0.04617127 | 0.26092 | 0.13847 | 0.0104 |
| BE2M17 | -0.2317 | 0.003125 | -0.39842 | -0.21779 | 0.1578 | 2.20156 | 0.083199 | 0.052741544 | 0.49141 | 0.00208 | 0.06434 |
| LN443 | -0.1782 | 0.096275 | -0.06694 | -0.35074 | 0.15737 | 2.19439 | 0.084028 | 0.107773034 | 0.25299 | 0.32208 | 0.00626 |
| SKNMC | -0.3581 | -0.03118 | -0.32266 | -0.15792 | 0.15733 | 2.19383 | 0.084093 | 0.005330819 | 0.4149 | 0.01115 | 0.13669 |
| RERFGC1B | -0.2265 | 0.071982 | -0.3472 | -0.1812 | 0.15665 | 2.18253 | 0.085417 | 0.056862659 | 0.30968 | 0.00675 | 0.10396 |
| HSOS1 | -0.2733 | -0.04732 | -0.38457 | -0.30633 | 0.15557 | 2.16476 | 0.087543 | 0.02738577 | 0.37208 | 0.00291 | 0.01525 |
| HCC15 | -0.1559 | -0.05765 | -0.42059 | -0.33081 | 0.15534 | 2.16091 | 0.08801 | 0.139789305 | 0.34543 | 0.00118 | 0.00948 |
| HA1E | -0.3183 | 0.001711 | -0.12338 | -0.34328 | 0.15518 | 2.15836 | 0.08832 | 0.012144453 | 0.4953 | 0.19665 | 0.00733 |
| SW620 | -0.2285 | 0.0369 | -0.37244 | -0.11094 | 0.15515 | 2.15782 | 0.088387 | 0.055276588 | 0.39959 | 0.00387 | 0.22153 |
| OC314 | -0.3527 | 0.040503 | -0.16849 | -0.25545 | 0.15515 | 2.15779 | 0.08839 | 0.006001309 | 0.39002 | 0.12107 | 0.03669 |
| CCFSTTG1 | -0.2042 | 0.167373 | -0.05887 | -0.26619 | 0.15503 | 2.15579 | 0.088635 | 0.077412584 | 0.12266 | 0.34233 | 0.03085 |
| OV7 | -0.2512 | 0.053508 | -0.31167 | -0.27134 | 0.15502 | 2.15561 | 0.088658 | 0.039247377 | 0.35604 | 0.01379 | 0.02832 |
| HCC56 | -0.2753 | 0.054342 | -0.32983 | -0.12197 | 0.15482 | 2.15241 | 0.08905 | 0.02650863 | 0.3539 | 0.00966 | 0.1994 |
| MKN28 | -0.3663 | 0.038458 | -0.22811 | -0.13588 | 0.15479 | 2.15179 | 0.089126 | 0.004445718 | 0.39544 | 0.05555 | 0.17338 |
| HCE4 | -0.359 | -0.08385 | -0.31292 | -0.32293 | 0.15433 | 2.14423 | 0.090062 | 0.005225572 | 0.28132 | 0.01346 | 0.01109 |
| LN428 | -0.3066 | 0.151237 | -0.17093 | -0.05646 | 0.15426 | 2.14309 | 0.090204 | 0.015167641 | 0.14723 | 0.11764 | 0.34847 |
| SIHA | -0.317 | 0.08174 | -0.12261 | -0.26046 | 0.15416 | 2.14146 | 0.090408 | 0.012455062 | 0.28627 | 0.19814 | 0.03386 |
| A375SKINCJ1 | -0.3877 | -0.00618 | -0.23192 | -0.12183 | 0.1541 | 2.14047 | 0.090532 | 0.002703776 | 0.48302 | 0.05255 | 0.19967 |

| | | | | | | | | | | | |
|---|---|---|---|---|---|---|---|---|---|---|---|
| MCC13 | -0.3177 | -0.05799 | -0.37744 | -0.23611 | 0.15402 | 2.13918 | 0.090694 | 0.012272214 | 0.34458 | 0.00344 | 0.04939 |
| CORL311 | -0.1548 | 0.166969 | -0.28846 | -0.12673 | 0.15395 | 2.13811 | 0.090827 | 0.141619989 | 0.12324 | 0.0211 | 0.19024 |
| SNU738 | -0.3796 | 0.003244 | -0.18968 | -0.24034 | 0.15389 | 2.13716 | 0.090947 | 0.00327859 | 0.49108 | 0.09353 | 0.04636 |
| WM793 | -0.2776 | 0.194009 | -0.14517 | -0.02377 | 0.15383 | 2.13613 | 0.091076 | 0.025490468 | 0.08851 | 0.15724 | 0.43493 |
| MHHNB11 | -0.1548 | 0.051524 | -0.35982 | -0.26687 | 0.15353 | 2.1312 | 0.0917 | 0.141511371 | 0.36116 | 0.00514 | 0.0305 |
| HEC6 | -0.2195 | 0.273111 | 0.051696 | -0.0693 | 0.15325 | 2.12664 | 0.092279 | 0.062795214 | 0.02749 | 0.36072 | 0.31626 |
| T98G | -0.1215 | 0.071212 | -0.1419 | -0.38102 | 0.15322 | 2.1261 | 0.092348 | 0.200243011 | 0.31156 | 0.1628 | 0.00317 |
| BECKER | -0.1839 | 0.005257 | -0.40683 | -0.21548 | 0.15322 | 2.12607 | 0.092351 | 0.100502375 | 0.48555 | 0.00168 | 0.06644 |
| NCIH2110 | -0.3263 | 0.048793 | -0.27051 | -0.07533 | 0.15311 | 2.12426 | 0.092584 | 0.010366976 | 0.36825 | 0.02872 | 0.30155 |
| NCIH1155 | -0.2828 | -0.00606 | -0.34623 | -0.05872 | 0.15269 | 2.11733 | 0.093473 | 0.023278606 | 0.48334 | 0.00689 | 0.34271 |
| MEC1 | -0.4016 | -0.13025 | -0.31103 | -0.19872 | 0.15254 | 2.11497 | 0.09378 | 0.001923877 | 0.18365 | 0.01396 | 0.08327 |
| NCIH2004RT | -0.1135 | 0.005098 | -0.21777 | -0.4162 | 0.15118 | 2.09277 | 0.0967 | 0.216363507 | 0.48599 | 0.06435 | 0.00132 |
| NCIH2172 | -0.2824 | -0.04877 | -0.36682 | -0.30974 | 0.15088 | 2.08781 | 0.097365 | 0.023446605 | 0.3683 | 0.00439 | 0.0143 |
| TUHR10TKB | -0.1781 | 0.142815 | -0.12434 | -0.29749 | 0.15075 | 2.08575 | 0.097642 | 0.107901593 | 0.16123 | 0.19481 | 0.01795 |
| SKNMM | -0.3438 | -0.14839 | -0.39023 | -0.29269 | 0.15075 | 2.08567 | 0.097653 | 0.007249451 | 0.15187 | 0.00254 | 0.01957 |
| TN2 | -0.1981 | 0.034391 | -0.37944 | -0.19678 | 0.15058 | 2.08293 | 0.098022 | 0.083898784 | 0.40629 | 0.00329 | 0.08539 |
| YAMATO | -0.1132 | 0.243907 | -0.1807 | 0.140793 | 0.15056 | 2.08259 | 0.098069 | 0.216895122 | 0.04392 | 0.1046 | 0.16472 |
| HT | -0.3632 | -0.05142 | -0.30554 | -0.23977 | 0.14973 | 2.06915 | 0.099906 | 0.004765459 | 0.36142 | 0.01547 | 0.04676 |
| SF539 | -0.0817 | 0.331543 | 0.110373 | -0.08528 | 0.14951 | 2.06557 | 0.1004 | 0.286357502 | 0.00934 | 0.22272 | 0.27798 |
| OV90 | -0.2414 | -0.05893 | -0.39477 | -0.07375 | 0.14919 | 2.06031 | 0.101133 | 0.045601829 | 0.34218 | 0.00228 | 0.30539 |
| RCCFG2 | -0.0183 | 0.265681 | -0.12847 | -0.16884 | 0.14914 | 2.05963 | 0.101227 | 0.449943689 | 0.0311 | 0.18696 | 0.12057 |
| HCC515 | -0.3547 | -0.02221 | -0.26003 | -0.27314 | 0.14903 | 2.05782 | 0.10148 | 0.005742675 | 0.43916 | 0.0341 | 0.02748 |
| KMM1 | -0.2936 | 0.166312 | -0.14166 | -0.09389 | 0.14895 | 2.05646 | 0.101671 | 0.01925022 | 0.12418 | 0.16322 | 0.25832 |
| P31FUJ | -0.3722 | -0.14778 | -0.3583 | -0.2515 | 0.1489 | 2.05573 | 0.101773 | 0.003886389 | 0.15288 | 0.00531 | 0.03905 |
| BT16 | -0.3917 | -0.07253 | 0.008516 | -0.16986 | 0.14885 | 2.05485 | 0.101897 | 0.002450898 | 0.30834 | 0.4766 | 0.11914 |
| KYM1 | -0.2343 | 0.017107 | -0.25801 | -0.35948 | 0.14838 | 2.04728 | 0.102967 | 0.050753781 | 0.45307 | 0.03523 | 0.00517 |
| TT2609C02 | -0.3933 | -0.01949 | -0.20596 | -0.12202 | 0.1479 | 2.0394 | 0.104092 | 0.002361597 | 0.44657 | 0.07565 | 0.19929 |
| NCIH520 | -0.2772 | 0.009488 | -0.30716 | -0.29724 | 0.14786 | 2.03879 | 0.10418 | 0.025663354 | 0.47393 | 0.01501 | 0.01803 |
| NCIH460 | -0.1415 | 0.177183 | -0.2759 | -0.09141 | 0.14723 | 2.02859 | 0.105657 | 0.163570525 | 0.10917 | 0.02623 | 0.26392 |
| LOVO | -0.289 | 0.091074 | -0.26192 | -0.08456 | 0.1471 | 2.02646 | 0.105968 | 0.020894408 | 0.26467 | 0.03307 | 0.27967 |
| 22RV1 | -0.2146 | -0.03691 | -0.41172 | -0.14129 | 0.14698 | 2.02458 | 0.106243 | 0.067296574 | 0.39955 | 0.00149 | 0.16386 |
| U87MG | -0.2736 | 0.046312 | -0.31499 | -0.07499 | 0.14685 | 2.02241 | 0.106561 | 0.027278318 | 0.37473 | 0.01294 | 0.30237 |
| EW22 | -0.1737 | -0.07733 | -0.42377 | -0.08068 | 0.1468 | 2.02175 | 0.106658 | 0.113893703 | 0.29674 | 0.00108 | 0.28877 |
| FARAGE | -0.3911 | -0.07831 | -0.26998 | -0.228 | 0.14674 | 2.02065 | 0.106821 | 0.002488635 | 0.29441 | 0.02897 | 0.05564 |
| HSC39 | -0.3662 | 0.045869 | -0.12646 | -0.18686 | 0.14629 | 2.01352 | 0.107876 | 0.00445827 | 0.37588 | 0.19077 | 0.09691 |
| TGW | -0.2556 | 0.06382 | -0.30016 | -0.02605 | 0.14589 | 2.00696 | 0.108857 | 0.036598841 | 0.32986 | 0.01709 | 0.42875 |
| SUM159PT | -0.2183 | 0.154801 | -0.25003 | -0.02373 | 0.14587 | 2.00671 | 0.108894 | 0.063840831 | 0.14154 | 0.03995 | 0.43502 |
| LPS27 | -0.2263 | 0.134813 | -0.10572 | -0.27003 | 0.14583 | 2.00602 | 0.108997 | 0.056985822 | 0.1753 | 0.23247 | 0.02895 |
| NCIH226 | -0.1012 | 0.170477 | -0.15042 | -0.27588 | 0.14579 | 2.00547 | 0.109079 | 0.242110633 | 0.11828 | 0.14855 | 0.02624 |
| SKN3 | -0.2627 | 0.184249 | -0.12357 | 0.07816 | 0.14576 | 2.00485 | 0.109174 | 0.032644219 | 0.10012 | 0.19628 | 0.29476 |
| HCC2935 | -0.1043 | 0.051357 | -0.32931 | -0.29695 | 0.14472 | 1.98816 | 0.111714 | 0.235476457 | 0.3616 | 0.00977 | 0.01812 |
| HUH7 | -0.1499 | 0.187752 | -0.24281 | -0.13578 | 0.14469 | 1.98775 | 0.111777 | 0.149400105 | 0.09583 | 0.04466 | 0.17356 |
| NCIH1437 | -0.2305 | -0.06645 | -0.41623 | -0.24994 | 0.1444 | 1.98305 | 0.112504 | 0.05364741 | 0.3233 | 0.00132 | 0.04001 |
| SNU1 | -0.2018 | 0.096393 | -0.25434 | -0.27098 | 0.14342 | 1.96739 | 0.114957 | 0.079964014 | 0.25273 | 0.03734 | 0.02849 |
| CCLFPEDS0001T | -0.287 | 0.099051 | -0.24696 | -0.09554 | 0.1433 | 1.9655 | 0.115257 | 0.021667925 | 0.24687 | 0.0419 | 0.25463 |
| SKNAS | -0.1526 | 0.110539 | -0.32844 | -0.08287 | 0.14322 | 1.96408 | 0.115482 | 0.145104411 | 0.22237 | 0.00994 | 0.2836 |
| NCIH82 | -0.3319 | 0.094558 | -0.15259 | -0.009 | 0.14299 | 1.96045 | 0.116062 | 0.009272859 | 0.25682 | 0.14505 | 0.47528 |
| D341Med | -0.1853 | 0.00712 | -0.38045 | -0.07202 | 0.14244 | 1.9517 | 0.117467 | 0.098793994 | 0.48043 | 0.00321 | 0.30959 |
| ISHIKAWAHERAKLIO02ER | -0.3504 | 0.073683 | -0.07162 | -0.1603 | 0.14164 | 1.93893 | 0.119551 | 0.00630561 | 0.30554 | 0.31056 | 0.13306 |
| F36P | -0.1471 | 0.011642 | -0.39506 | -0.1127 | 0.14154 | 1.93736 | 0.11981 | 0.154076368 | 0.46802 | 0.00226 | 0.21792 |
| SNU387 | -0.2664 | -0.01859 | -0.34719 | -0.27372 | 0.14132 | 1.93382 | 0.120394 | 0.030750765 | 0.44901 | 0.00675 | 0.02721 |
| SW156 | -0.2987 | 0.012209 | -0.30561 | -0.22657 | 0.14117 | 1.93135 | 0.120805 | 0.01755934 | 0.46647 | 0.01545 | 0.0568 |
| MYLA | -0.2594 | 0.00364 | -0.35618 | -0.14095 | 0.14115 | 1.93115 | 0.120837 | 0.034457816 | 0.48999 | 0.00556 | 0.16445 |
| TOV21G | -0.2746 | 0.218312 | -0.00834 | 0.009264 | 0.1411 | 1.93024 | 0.120989 | 0.026807311 | 0.06387 | 0.47709 | 0.47454 |
| A172 | -0.1403 | 0.161385 | -0.13442 | -0.27128 | 0.141 | 1.92863 | 0.121257 | 0.165560088 | 0.13143 | 0.17601 | 0.02835 |
| CCC5 | -0.3791 | 0.030269 | -0.10797 | -0.07581 | 0.14055 | 1.92146 | 0.122459 | 0.003312281 | 0.41735 | 0.22773 | 0.30039 |
| HS578T | -0.0947 | 0.219667 | -0.09102 | -0.2224 | 0.14047 | 1.92026 | 0.122662 | 0.256456367 | 0.06266 | 0.2648 | 0.06029 |
| SNUC5 | -0.308 | 0.124462 | -0.06423 | -0.16744 | 0.14007 | 1.91388 | 0.123743 | 0.014766779 | 0.19457 | 0.32883 | 0.12257 |
| MERO41 | -0.1863 | 0.011953 | -0.27035 | -0.36232 | 0.13998 | 1.9125 | 0.123978 | 0.097636397 | 0.46717 | 0.02879 | 0.00486 |
| NCIH1048 | -0.2044 | 0.152529 | -0.09534 | -0.25217 | 0.1396 | 1.90647 | 0.12501 | 0.077203441 | 0.14515 | 0.25508 | 0.03864 |
| MMAC | -0.3876 | -0.14658 | -0.25967 | -0.10718 | 0.13913 | 1.89895 | 0.12631 | 0.002709479 | 0.15488 | 0.0343 | 0.2294 |
| LMSU | -0.0761 | 0.153821 | -0.1063 | -0.29584 | 0.13912 | 1.89886 | 0.126325 | 0.299754412 | 0.14309 | 0.23125 | 0.01849 |
| NCIH841 | -0.2218 | 0.12981 | -0.26037 | -0.13647 | 0.13907 | 1.8981 | 0.126458 | 0.060767579 | 0.18446 | 0.03391 | 0.17233 |
| SKRC31 | -0.2234 | 0.230476 | -0.10837 | 0.049848 | 0.13888 | 1.89507 | 0.126984 | 0.059459589 | 0.05367 | 0.22688 | 0.36551 |
| PATU8902 | -0.1953 | 0.036431 | -0.354 | -0.06794 | 0.13867 | 1.89175 | 0.127565 | 0.086994518 | 0.40084 | 0.00583 | 0.3196 |
| SKHEP1 | -0.3263 | 0.020803 | -0.17516 | -0.27347 | 0.1386 | 1.89061 | 0.127765 | 0.01037866 | 0.44299 | 0.11186 | 0.02733 |
| LNZTA3WT4 | -0.1355 | -0.04931 | -0.2275 | -0.42663 | 0.13818 | 1.8839 | 0.128949 | 0.174018503 | 0.3669 | 0.05604 | 0.001 |
| UACC257 | -0.1028 | 0.325295 | -0.02319 | 0.20085 | 0.13816 | 1.88363 | 0.128996 | 0.238755699 | 0.01058 | 0.4365 | 0.08097 |
| ES8 | 0.00539 | 0.126522 | -0.32158 | -0.04372 | 0.13797 | 1.88064 | 0.129528 | 0.485170903 | 0.19064 | 0.01138 | 0.38152 |
| UMUC1 | -0.1752 | 0.095244 | -0.31688 | -0.16736 | 0.13753 | 1.87365 | 0.130777 | 0.111804559 | 0.25529 | 0.01248 | 0.12267 |
| HT144SKINFV3 | -0.302 | 0.073879 | -0.18533 | -0.22546 | 0.1375 | 1.87316 | 0.130865 | 0.016537625 | 0.30507 | 0.09878 | 0.05771 |
| SKNDZ | -0.1548 | 0.090243 | -0.27762 | -0.26648 | 0.13686 | 1.86301 | 0.132701 | 0.141601625 | 0.26656 | 0.02547 | 0.0307 |
| SNU1327 | -0.257 | 0.210385 | -0.0829 | 0.016633 | 0.13681 | 1.86231 | 0.132829 | 0.035773797 | 0.07126 | 0.28355 | 0.45436 |
| SG231 | -0.0722 | 0.299362 | -0.1211 | 0.118641 | 0.13608 | 1.85078 | 0.134948 | 0.30909659 | 0.01734 | 0.20108 | 0.20594 |
| HS936T | -0.2893 | 0.070496 | -0.25399 | -0.15821 | 0.13599 | 1.84936 | 0.135211 | 0.02077076 | 0.31331 | 0.03755 | 0.13624 |
| MG63 | -0.2539 | 0.065455 | -0.26142 | -0.2412 | 0.13589 | 1.84782 | 0.135497 | 0.037581228 | 0.32577 | 0.03334 | 0.04576 |
| KP363T | -0.3195 | 0.049673 | -0.23955 | -0.14014 | 0.13583 | 1.84681 | 0.135686 | 0.011847928 | 0.36596 | 0.04691 | 0.16585 |
| JHH7 | -0.2046 | 0.185346 | -0.1869 | -0.12499 | 0.13575 | 1.84558 | 0.135914 | 0.076997332 | 0.09876 | 0.09686 | 0.19356 |
| JHOM1 | -0.2929 | -0.02566 | -0.30327 | -0.29005 | 0.13539 | 1.84 | 0.13696 | 0.019506223 | 0.42979 | 0.01614 | 0.02051 |
| MAVER1 | -0.1833 | 0.243732 | -0.13356 | -0.02741 | 0.13526 | 1.83784 | 0.137365 | 0.101264094 | 0.04403 | 0.17757 | 0.42506 |
| NCIH1648 | -0.3725 | -0.11234 | -0.29849 | -0.20823 | 0.13515 | 1.83623 | 0.137669 | 0.003858189 | 0.21866 | 0.01762 | 0.07337 |
| NMCG1 | -0.2323 | 0.005964 | -0.344 | -0.24769 | 0.13493 | 1.83272 | 0.138333 | 0.052242577 | 0.48361 | 0.00722 | 0.04143 |
| HLF | -0.2009 | 0.089042 | -0.2037 | -0.29172 | 0.13414 | 1.82028 | 0.140714 | 0.080859941 | 0.2693 | 0.07797 | 0.01991 |
| REH | -0.3229 | -0.08474 | -0.34747 | -0.1899 | 0.13403 | 1.81857 | 0.141043 | 0.011094388 | 0.27924 | 0.00671 | 0.09327 |
| BICR10 | -0.2133 | 0.072727 | -0.1442 | -0.31803 | 0.13386 | 1.81596 | 0.141548 | 0.068473486 | 0.30786 | 0.15888 | 0.0122 |
| HS294T | -0.3006 | -0.10293 | -0.37689 | -0.19501 | 0.13381 | 1.81508 | 0.141719 | 0.016960063 | 0.23845 | 0.00349 | 0.08737 |
| TCCSUP | -0.1878 | 0.129425 | -0.07919 | -0.27715 | 0.13347 | 1.80987 | 0.142735 | 0.095772665 | 0.18518 | 0.29232 | 0.02568 |
| MERO25 | -0.2515 | 0.185929 | -0.08227 | -0.12586 | 0.13339 | 1.80858 | 0.142987 | 0.039065531 | 0.09804 | 0.28503 | 0.1919 |
| JEG3 | -0.2904 | 0.174812 | 0.003143 | -0.05276 | 0.13339 | 1.80855 | 0.142992 | 0.020397244 | 0.11233 | 0.49136 | 0.35796 |
| NALM6 | -0.3979 | -0.08047 | -0.15685 | -0.19174 | 0.13307 | 1.80351 | 0.143984 | 0.002109794 | 0.28928 | 0.13834 | 0.09111 |
| SNU869 | -0.2727 | 0.138004 | -0.18283 | -0.08482 | 0.13292 | 1.80123 | 0.144433 | 0.027668066 | 0.1696 | 0.10189 | 0.27906 |
| 42MGBA | -0.1092 | 0.133157 | -0.11378 | -0.29936 | 0.13278 | 1.79912 | 0.144853 | 0.225252961 | 0.1783 | 0.21572 | 0.01734 |
| GI1 | -0.2646 | -0.02446 | -0.3193 | -0.29544 | 0.13272 | 1.79817 | 0.145041 | 0.031671035 | 0.43305 | 0.0119 | 0.01863 |
| DU4475 | -0.1928 | 0.237022 | -0.03408 | -0.12031 | 0.1326 | 1.79625 | 0.145422 | 0.089885841 | 0.04873 | 0.40713 | 0.20264 |
| MON | -0.161 | 0.142181 | -0.23546 | -0.2053 | 0.13242 | 1.79337 | 0.145996 | 0.132018755 | 0.16232 | 0.04987 | 0.07632 |
| K562 | -0.2978 | -0.12639 | -0.32726 | -0.36165 | 0.13215 | 1.78922 | 0.146829 | 0.017843464 | 0.19088 | 0.01017 | 0.00493 |
| LAN2 | -0.119 | 0.121992 | -0.2608 | -0.23219 | 0.13169 | 1.78201 | 0.148284 | 0.205210469 | 0.19935 | 0.03368 | 0.05234 |
| NCIH3122 | -0.2802 | 0.016109 | -0.30442 | -0.2036 | 0.13168 | 1.78185 | 0.148317 | 0.024369253 | 0.4558 | 0.0158 | 0.07807 |
| GB1 | -0.2177 | 0.035101 | -0.32697 | -0.22033 | 0.1312 | 1.77435 | 0.149846 | 0.0643903 | 0.40439 | 0.01023 | 0.06208 |
| SNU638 | -0.1544 | 0.019637 | -0.35341 | -0.02493 | 0.13116 | 1.7737 | 0.14998 | 0.142174644 | 0.44617 | 0.00591 | 0.43178 |
| WERIRB1 | -0.3437 | -0.04883 | -0.28679 | -0.15638 | 0.13084 | 1.76879 | 0.15099 | 0.007263841 | 0.36815 | 0.02173 | 0.13907 |
| UOK101 | -0.2683 | 0.187394 | -0.006 | -0.08161 | 0.13068 | 1.76633 | 0.151499 | 0.029779755 | 0.09626 | 0.4835 | 0.28659 |
| TO14 | -0.2597 | 0.089482 | -0.25331 | -0.05743 | 0.13066 | 1.76603 | 0.151561 | 0.034287997 | 0.2683 | 0.03795 | 0.346 |
| SNU201 | -0.2967 | 0.051615 | -0.23209 | -0.20583 | 0.1304 | 1.76196 | 0.152407 | 0.018198175 | 0.36093 | 0.05242 | 0.07578 |
| HCC70 | -0.1571 | 0.177016 | -0.03055 | -0.23356 | 0.13035 | 1.76125 | 0.152555 | 0.137978094 | 0.10939 | 0.4166 | 0.05129 |
| MM386 | -0.2972 | -0.03228 | -0.32881 | -0.163 | 0.12964 | 1.75012 | 0.154893 | 0.018039695 | 0.41194 | 0.00986 | 0.12903 |
| KU1919 | -0.354 | 0.052791 | -0.08063 | -0.0405 | 0.12955 | 1.74879 | 0.155175 | 0.00583569 | 0.35789 | 0.28889 | 0.39003 |
| NCIH1568 | -0.2242 | 0.170388 | -0.18606 | -0.00518 | 0.12943 | 1.74688 | 0.155581 | 0.058755191 | 0.1184 | 0.09788 | 0.48577 |
| NB17 | 0.02071 | 0.088353 | -0.26384 | -0.25444 | 0.12867 | 1.73509 | 0.158107 | 0.443251148 | 0.27088 | 0.03206 | 0.03728 |
| SCMCRM2 | -0.0381 | 0.275278 | -0.13771 | 0.14802 | 0.12864 | 1.73473 | 0.158183 | 0.396523746 | 0.02651 | 0.17013 | 0.15248 |
| RPMI8226 | -0.3107 | -0.03209 | -0.31227 | -0.17196 | 0.12859 | 1.73396 | 0.15835 | 0.014031004 | 0.41245 | 0.01363 | 0.11621 |
| SNU685 | -0.2757 | 0.019808 | -0.28008 | -0.24059 | 0.12856 | 1.73349 | 0.158451 | 0.026315104 | 0.4457 | 0.02442 | 0.04618 |
| HS729 | -0.2514 | 0.04191 | -0.24659 | -0.27339 | 0.1285 | 1.73253 | 0.158659 | 0.039084729 | 0.3863 | 0.04214 | 0.02737 |
| KYSE70 | -0.3301 | 0.00085 | -0.25059 | -0.18471 | 0.12848 | 1.73226 | 0.158717 | 0.009617948 | 0.49766 | 0.03961 | 0.09954 |
| BHT101 | -0.3069 | 0.077247 | -0.19256 | -0.1467 | 0.12845 | 1.73172 | 0.158836 | 0.015083229 | 0.29695 | 0.09016 | 0.15467 |
| C32 | -0.2336 | 0.235429 | -0.00212 | 0.088883 | 0.12841 | 1.73107 | 0.158975 | 0.051284236 | 0.0499 | 0.49416 | 0.26967 |
| UMUC14 | -0.201 | 0.117182 | -0.19916 | -0.24026 | 0.12838 | 1.73063 | 0.159072 | 0.080789186 | 0.20884 | 0.08279 | 0.04641 |
| Y79 | -0.1472 | 0.129115 | -0.28405 | -0.02898 | 0.12819 | 1.72774 | 0.1597 | 0.153828154 | 0.18576 | 0.0228 | 0.42084 |
| LO68 | -0.1807 | -0.01897 | -0.33988 | -0.31082 | 0.12807 | 1.72593 | 0.160096 | 0.104560947 | 0.44798 | 0.00787 | 0.01401 |

| | | | | | | | | | | | |
|---|---|---|---|---|---|---|---|---|---|---|---|
| MELHO | -0.3485 | 0.020254 | -0.17364 | -0.17072 | 0.12791 | 1.72342 | 0.160644 | 0.006565602 | 0.44448 | 0.11392 | 0.11793 |
| COLO201 | -0.3153 | 0.019003 | -0.25327 | -0.17334 | 0.12757 | 1.71809 | 0.161816 | 0.012870541 | 0.44789 | 0.03798 | 0.11432 |
| AGS | -0.3063 | 0.084994 | -0.16431 | -0.15979 | 0.12743 | 1.71599 | 0.16228 | 0.015265676 | 0.27865 | 0.1271 | 0.13384 |
| P30OHK | -0.2984 | -0.15632 | -0.24514 | -0.4001 | 0.12742 | 1.71578 | 0.162326 | 0.017662057 | 0.13917 | 0.04309 | 0.002 |
| NAMALWA | -0.3475 | -0.02881 | -0.17945 | -0.25614 | 0.12706 | 1.71021 | 0.163565 | 0.006710767 | 0.42129 | 0.1062 | 0.03629 |
| TC71 | -0.1385 | 0.079527 | -0.3282 | -0.0781 | 0.12624 | 1.69762 | 0.166397 | 0.168736033 | 0.29151 | 0.00999 | 0.29491 |
| LU134A | -0.2918 | -0.08256 | -0.32556 | -0.30515 | 0.12617 | 1.6965 | 0.16665 | 0.019873632 | 0.28434 | 0.01052 | 0.01559 |
| PACADD165 | -0.2257 | 0.109326 | -0.16661 | -0.24238 | 0.12602 | 1.6943 | 0.16715 | 0.057550323 | 0.22489 | 0.12376 | 0.04495 |
| MOGGUVW | -0.144 | -0.04356 | -0.37339 | -0.03195 | 0.12592 | 1.69269 | 0.167517 | 0.159189563 | 0.38195 | 0.00378 | 0.41283 |
| CAOV4 | -0.1855 | 0.089265 | -0.14499 | -0.30018 | 0.12586 | 1.69186 | 0.167708 | 0.098545187 | 0.26879 | 0.15754 | 0.01709 |
| CAMA1 | -0.2914 | 0.064473 | -0.14713 | -0.23445 | 0.12569 | 1.68917 | 0.168324 | 0.020016185 | 0.32822 | 0.15395 | 0.05063 |
| YH13 | -0.1025 | 0.110938 | -0.21023 | -0.27922 | 0.12564 | 1.68833 | 0.168516 | 0.239473109 | 0.22155 | 0.07141 | 0.02478 |
| G292CLONEA141B1 | -0.185 | -0.04063 | -0.19999 | -0.39654 | 0.12554 | 1.68689 | 0.168846 | 0.099187443 | 0.38968 | 0.08189 | 0.00218 |
| KML1 | -0.2797 | 0.120738 | -0.14069 | 0.035414 | 0.12541 | 1.68483 | 0.16932 | 0.024581202 | 0.2018 | 0.1649 | 0.40355 |
| SCLC22H | -0.3019 | -0.11807 | -0.34993 | -0.28884 | 0.12532 | 1.68345 | 0.16964 | 0.016559592 | 0.20708 | 0.00637 | 0.02096 |
| RPMI2650 | -0.2588 | 0.205734 | -0.00412 | 0.034404 | 0.12525 | 1.68238 | 0.169886 | 0.034792672 | 0.07588 | 0.48866 | 0.40626 |
| MONOMAC1 | -0.2512 | -0.13636 | -0.35941 | -0.3572 | 0.12512 | 1.6804 | 0.170344 | 0.039255625 | 0.17252 | 0.00518 | 0.00544 |
| S462 | -0.3589 | -0.01949 | -0.19355 | -0.09286 | 0.12507 | 1.67972 | 0.170503 | 0.005242811 | 0.44658 | 0.08903 | 0.26064 |
| MDAMB157 | -0.188 | 0.084415 | -0.18584 | -0.29136 | 0.12499 | 1.67846 | 0.170796 | 0.095493072 | 0.28 | 0.09816 | 0.02004 |
| CORL47 | -0.3274 | -0.00317 | -0.21915 | -0.23201 | 0.12496 | 1.67803 | 0.170894 | 0.010148934 | 0.49129 | 0.06312 | 0.05248 |
| HCC2429 | -0.1347 | -0.06448 | -0.35701 | -0.33996 | 0.12465 | 1.67325 | 0.172011 | 0.175541097 | 0.3282 | 0.00546 | 0.00786 |
| D425 | -0.2768 | -0.13984 | -0.38255 | -0.16118 | 0.1246 | 1.6724 | 0.172209 | 0.02584617 | 0.16638 | 0.00306 | 0.13174 |
| MM485 | -0.3024 | -0.17264 | -0.3576 | -0.13345 | 0.12437 | 1.66896 | 0.173017 | 0.01638749 | 0.11529 | 0.00539 | 0.17777 |
| OCILY19 | -0.092 | 0.099421 | -0.31018 | -0.16639 | 0.12424 | 1.66692 | 0.173498 | 0.26247833 | 0.24606 | 0.01418 | 0.12407 |
| SNU46 | -0.2771 | -0.03683 | -0.17005 | -0.3469 | 0.12423 | 1.66683 | 0.17352 | 0.025680231 | 0.39978 | 0.11887 | 0.00679 |
| ONS76 | -0.2215 | 0.089795 | -0.20367 | -0.24985 | 0.1242 | 1.66625 | 0.173657 | 0.061056394 | 0.26758 | 0.078 | 0.04007 |
| FU97 | -0.1107 | 0.069598 | -0.10657 | -0.33817 | 0.12357 | 1.65663 | 0.175943 | 0.221946733 | 0.31552 | 0.23068 | 0.00815 |
| SH4 | -0.278 | 0.106635 | -0.13428 | -0.1829 | 0.1235 | 1.6556 | 0.176189 | 0.025301451 | 0.23054 | 0.17626 | 0.10181 |
| COLO792 | -0.2867 | 0.038515 | -0.25111 | -0.05646 | 0.12301 | 1.64811 | 0.177993 | 0.021772084 | 0.39529 | 0.03929 | 0.34848 |
| KARPAS1718 | -0.3173 | -0.12851 | -0.34114 | -0.16055 | 0.12281 | 1.64503 | 0.178737 | 0.012369352 | 0.18688 | 0.00767 | 0.13269 |
| SKMEL30 | -0.3078 | -0.2293 | -0.08724 | 0.020829 | 0.12221 | 1.63586 | 0.180979 | 0.014829174 | 0.0546 | 0.27345 | 0.44292 |
| NB6 | -0.2009 | -0.11646 | -0.35374 | -0.3661 | 0.12214 | 1.6349 | 0.181214 | 0.080961713 | 0.21029 | 0.00587 | 0.00446 |
| MOLM13 | -0.1922 | -0.04826 | -0.19606 | -0.39232 | 0.12157 | 1.62608 | 0.183397 | 0.09057172 | 0.36965 | 0.08619 | 0.00242 |
| KNS60 | -0.1752 | 0.08833 | -0.22241 | -0.2659 | 0.12154 | 1.62569 | 0.183492 | 0.111812833 | 0.27094 | 0.06028 | 0.03099 |
| A3KAW | -0.3112 | -0.08233 | -0.30012 | -0.28425 | 0.12142 | 1.62391 | 0.183936 | 0.013906672 | 0.28487 | 0.01711 | 0.02272 |
| SIMA | -0.1216 | 0.153594 | -0.18569 | -0.22038 | 0.12104 | 1.61812 | 0.185388 | 0.200130488 | 0.14345 | 0.09834 | 0.06204 |
| NZM3 | -0.3478 | -0.03901 | -0.23138 | -0.11627 | 0.12099 | 1.61725 | 0.185606 | 0.006660367 | 0.39398 | 0.05297 | 0.21067 |
| ROS50 | 0.06902 | -0.0148 | -0.34066 | -0.18292 | 0.12088 | 1.61565 | 0.186009 | 0.316931793 | 0.45938 | 0.00774 | 0.10177 |
| PSN1 | -0.1617 | 0.220168 | -0.13316 | -0.09182 | 0.12053 | 1.61027 | 0.187369 | 0.130988462 | 0.06222 | 0.17829 | 0.26299 |
| MCC142 | -0.2507 | 0.039817 | -0.21311 | -0.27569 | 0.12049 | 1.60965 | 0.187528 | 0.039561167 | 0.39184 | 0.06865 | 0.02632 |
| PK8 | -0.1682 | 0.198859 | -0.13521 | -0.13843 | 0.12016 | 1.60476 | 0.188774 | 0.121467415 | 0.08311 | 0.17459 | 0.16886 |
| TC205 | -0.1489 | 0.204081 | -0.18293 | -0.06244 | 0.11961 | 1.59632 | 0.190945 | 0.151053969 | 0.07757 | 0.10176 | 0.33331 |
| EOL1 | -0.3162 | 0.003767 | -0.23677 | -0.07887 | 0.11941 | 1.59339 | 0.191705 | 0.012650152 | 0.48964 | 0.04891 | 0.29307 |
| HSSCH2 | -0.2585 | 0.01922 | -0.25752 | -0.25861 | 0.11939 | 1.59303 | 0.191797 | 0.034960382 | 0.4473 | 0.0355 | 0.03488 |
| NB5 | -0.184 | 0.102552 | -0.27375 | -0.14066 | 0.11934 | 1.59225 | 0.192001 | 0.100447216 | 0.23926 | 0.0272 | 0.16496 |
| MORCPR | -0.1707 | 0.029617 | -0.31385 | 8.36E-05 | 0.11902 | 1.58736 | 0.193275 | 0.117956906 | 0.41911 | 0.01323 | 0.49977 |
| SUPT1 | -0.3291 | -0.08221 | -0.20191 | -0.30856 | 0.11901 | 1.5872 | 0.193315 | 0.00981276 | 0.28517 | 0.07984 | 0.01462 |
| HT144 | -0.1123 | 0.251858 | -0.13841 | -0.0207 | 0.11884 | 1.58475 | 0.193958 | 0.218711535 | 0.03883 | 0.16888 | 0.44327 |
| TCCPAN2 | -0.204 | 0.11363 | -0.24629 | -0.13701 | 0.11862 | 1.58135 | 0.194852 | 0.077662979 | 0.21602 | 0.04234 | 0.17136 |
| CCLFPEDS0003T | -0.1908 | 0.233735 | -0.02246 | 0.150255 | 0.11858 | 1.58072 | 0.195017 | 0.092238977 | 0.05116 | 0.43848 | 0.14882 |
| SNUC1 | -0.2001 | 0.027588 | -0.33111 | -0.14524 | 0.11852 | 1.57979 | 0.195265 | 0.081725337 | 0.42458 | 0.00942 | 0.15712 |
| NCIH1836 | -0.3142 | 0.000296 | -0.24729 | -0.11251 | 0.11805 | 1.57282 | 0.197112 | 0.013129385 | 0.49918 | 0.04169 | 0.21832 |
| KP1N | -0.3284 | -0.02108 | -0.23732 | -0.1941 | 0.1177 | 1.56749 | 0.198536 | 0.009949099 | 0.44223 | 0.04851 | 0.0884 |
| MDST8 | -0.1442 | 0.01505 | -0.35551 | -0.17525 | 0.11767 | 1.56696 | 0.19868 | 0.15881227 | 0.45869 | 0.00565 | 0.11174 |
| PFSK1 | -0.1428 | 0.266881 | -0.03 | 0.167523 | 0.11761 | 1.5661 | 0.19891 | 0.161274913 | 0.0305 | 0.41807 | 0.12245 |
| HMY1 | -0.2878 | 0.027531 | -0.25563 | -0.15785 | 0.11756 | 1.56536 | 0.199111 | 0.021335734 | 0.42474 | 0.03658 | 0.1368 |
| H4 | -0.1332 | 0.117906 | -0.23785 | -0.21624 | 0.11756 | 1.56528 | 0.19913 | 0.178161647 | 0.2074 | 0.04813 | 0.06574 |
| TTC709 | -0.3267 | -0.20981 | -0.2306 | -0.36214 | 0.1174 | 1.56291 | 0.19977 | 0.010287099 | 0.07182 | 0.05358 | 0.00488 |
| RPE1SS48 | -0.2368 | 0.171993 | -0.12306 | 0.016345 | 0.1168 | 1.5539 | 0.202216 | 0.048864648 | 0.11617 | 0.19728 | 0.45515 |
| HEYA8 | -0.1949 | 0.120897 | -0.21439 | -0.18343 | 0.11623 | 1.54533 | 0.204567 | 0.087446553 | 0.20149 | 0.06744 | 0.10114 |
| SKNFI | -0.1542 | 0.058103 | -0.30944 | -0.02444 | 0.11613 | 1.54379 | 0.204992 | 0.142445805 | 0.34428 | 0.01438 | 0.43311 |
| BFTC905 | -0.2767 | 0.014297 | -0.27886 | -0.12056 | 0.11599 | 1.5417 | 0.205572 | 0.025894748 | 0.46075 | 0.02494 | 0.20215 |
| RMSYM | -0.266 | 0.008966 | -0.27417 | -0.2277 | 0.11597 | 1.5414 | 0.205653 | 0.030957759 | 0.47536 | 0.027 | 0.05588 |
| HT144SKINFV1 | -0.3374 | 0.047633 | -0.06348 | -0.07399 | 0.11567 | 1.53697 | 0.206887 | 0.008286562 | 0.37127 | 0.33071 | 0.30479 |
| RH30 | -0.2581 | 0.108193 | -0.17967 | -0.14268 | 0.11562 | 1.53611 | 0.207128 | 0.035162889 | 0.22726 | 0.10592 | 0.16147 |
| RERFLCAD2 | -0.1459 | 0.175794 | -0.20937 | -0.08688 | 0.11527 | 1.53085 | 0.2086 | 0.156051575 | 0.11102 | 0.07225 | 0.27428 |
| MERO95 | -0.2371 | 0.005642 | -0.07159 | -0.31549 | 0.11523 | 1.53025 | 0.208768 | 0.048662384 | 0.48449 | 0.31063 | 0.01282 |
| KMS34 | -0.2752 | -0.1211 | -0.36066 | -0.25008 | 0.11515 | 1.5291 | 0.209092 | 0.026556015 | 0.20108 | 0.00504 | 0.03993 |
| KARPAS422 | -0.3447 | -0.18693 | -0.25443 | -0.31529 | 0.11515 | 1.52907 | 0.209101 | 0.007114647 | 0.09683 | 0.03729 | 0.01287 |
| 8505C | -0.3269 | -0.04307 | -0.16909 | -0.26626 | 0.11429 | 1.51625 | 0.212742 | 0.010250101 | 0.38324 | 0.12022 | 0.03081 |
| OCIAML2 | -0.3077 | -0.0431 | -0.28401 | -0.19201 | 0.11427 | 1.51594 | 0.212831 | 0.014848101 | 0.38316 | 0.02281 | 0.0908 |
| LU65 | -0.2463 | 0.171515 | -0.0939 | -0.05607 | 0.11417 | 1.51444 | 0.21326 | 0.042360317 | 0.11683 | 0.2583 | 0.34948 |
| MDAMB435S | -0.221 | 0.028251 | -0.30861 | -0.1089 | 0.11389 | 1.51013 | 0.214499 | 0.061538951 | 0.42279 | 0.01461 | 0.22578 |
| JJN3 | -0.1921 | 0.177324 | -0.16889 | -0.00185 | 0.11333 | 1.50176 | 0.216928 | 0.090646952 | 0.10898 | 0.1205 | 0.49492 |
| SNU16 | -0.0862 | -0.10789 | -0.19267 | -0.41325 | 0.11305 | 1.49762 | 0.218138 | 0.275895385 | 0.2279 | 0.09004 | 0.00143 |
| NCIH2122 | -0.1686 | -0.00783 | -0.34809 | -0.08009 | 0.11288 | 1.49518 | 0.218856 | 0.120976044 | 0.47847 | 0.00662 | 0.29017 |
| LN464 | -0.3415 | -0.04394 | -0.21137 | -0.10051 | 0.11276 | 1.49329 | 0.21941 | 0.007601409 | 0.38096 | 0.07031 | 0.24369 |
| JMURTK2 | -0.1236 | 0.050215 | -0.07529 | -0.32833 | 0.11202 | 1.48224 | 0.222689 | 0.196287708 | 0.36455 | 0.30164 | 0.00996 |
| DEL | -0.2387 | 0.045159 | -0.24689 | -0.22256 | 0.11194 | 1.48111 | 0.223028 | 0.047523849 | 0.37775 | 0.04195 | 0.06015 |
| NCIH1355 | -0.3232 | -0.1466 | -0.20711 | -0.33232 | 0.11177 | 1.47854 | 0.223797 | 0.011036644 | 0.15483 | 0.07449 | 0.00919 |
| HT1080 | -0.2683 | 0.10529 | -0.15835 | -0.12229 | 0.11142 | 1.47341 | 0.225342 | 0.029813932 | 0.2334 | 0.13603 | 0.19878 |
| SUM52PE | -0.1341 | 0.013922 | -0.2291 | -0.33369 | 0.11129 | 1.47134 | 0.22597 | 0.176614956 | 0.46178 | 0.05476 | 0.00894 |
| CTV1DM | -0.1439 | 0.021873 | -0.32513 | -0.22405 | 0.11118 | 1.46983 | 0.226428 | 0.159458468 | 0.44008 | 0.01062 | 0.05889 |
| SKLMS1 | -0.2817 | 0.058008 | -0.18958 | -0.17267 | 0.11045 | 1.45885 | 0.229781 | 0.023730989 | 0.34453 | 0.09364 | 0.11524 |
| CMLT1 | -0.2691 | 0.031279 | -0.20088 | -0.23971 | 0.1104 | 1.45813 | 0.230004 | 0.029402604 | 0.41464 | 0.08094 | 0.0468 |
| COGN305 | -0.2024 | 0.104819 | -0.2263 | 0.007767 | 0.10978 | 1.44905 | 0.232816 | 0.079371553 | 0.2344 | 0.05702 | 0.47865 |
| SNU398 | -0.2691 | 0.098064 | -0.16598 | -0.02814 | 0.1096 | 1.44627 | 0.233682 | 0.029397827 | 0.24904 | 0.12466 | 0.42311 |
| JVE015 | -0.2094 | -0.1431 | -0.19306 | -0.40633 | 0.10946 | 1.44422 | 0.234326 | 0.072193715 | 0.16075 | 0.08959 | 0.00171 |
| NCIH1915 | -0.1783 | 0.13675 | -0.20208 | -0.15135 | 0.10914 | 1.43949 | 0.23581 | 0.107696514 | 0.17183 | 0.07966 | 0.14704 |
| DB | -0.2693 | -0.16934 | -0.35636 | -0.29569 | 0.10912 | 1.43913 | 0.235923 | 0.029277162 | 0.11987 | 0.00554 | 0.01854 |
| HSQ89 | -0.1551 | 0.058435 | -0.26337 | -0.24635 | 0.10893 | 1.43641 | 0.236783 | 0.141080071 | 0.34344 | 0.0323 | 0.0423 |
| MSTO211H | -0.0374 | 0.197454 | -0.19795 | 0.082729 | 0.10855 | 1.43079 | 0.238568 | 0.398181495 | 0.08465 | 0.0841 | 0.28394 |
| KD | -0.1361 | 0.073314 | -0.09615 | -0.30359 | 0.10839 | 1.42847 | 0.239308 | 0.173067206 | 0.30644 | 0.25327 | 0.01605 |
| D283MED | -0.0849 | 0.147734 | -0.25016 | -0.0061 | 0.10827 | 1.42667 | 0.239884 | 0.278962701 | 0.15295 | 0.03988 | 0.48324 |
| NCCLMS1C1 | -0.2348 | -0.01217 | -0.07454 | -0.31455 | 0.10807 | 1.42369 | 0.240839 | 0.050336114 | 0.46657 | 0.30345 | 0.01305 |
| SEM | -0.1321 | -0.06078 | -0.35418 | -0.28427 | 0.10784 | 1.42028 | 0.241937 | 0.180257196 | 0.33749 | 0.00581 | 0.02271 |
| COLO320 | -0.3454 | -0.09428 | -0.218 | -0.11123 | 0.10776 | 1.41914 | 0.242306 | 0.007012053 | 0.25745 | 0.06415 | 0.22094 |
| T3M3 | -0.2878 | 0.028884 | -0.22587 | -0.09947 | 0.10767 | 1.41774 | 0.242758 | 0.021352127 | 0.42109 | 0.05737 | 0.24595 |
| TANOUE | -0.3393 | -0.18678 | -0.28261 | -0.24637 | 0.10764 | 1.41731 | 0.242897 | 0.007957271 | 0.097 | 0.02338 | 0.04229 |
| NOZ | -0.1913 | 0.157967 | -0.13059 | 0.103729 | 0.10742 | 1.41413 | 0.243928 | 0.091617002 | 0.13662 | 0.18301 | 0.23673 |
| HKBMM | -0.1562 | 0.061834 | -0.24274 | -0.2554 | 0.10684 | 1.40552 | 0.246743 | 0.139327407 | 0.33484 | 0.0447 | 0.03672 |
| TF1 | -0.2564 | 0.1233 | -0.1349 | -0.09565 | 0.10656 | 1.40143 | 0.248093 | 0.036163497 | 0.19681 | 0.17514 | 0.25438 |
| WM88 | -0.1414 | 0.227272 | -0.01695 | 0.203267 | 0.10656 | 1.40139 | 0.248104 | 0.163752229 | 0.05623 | 0.4535 | 0.07842 |
| OAW42 | -0.2539 | 0.049058 | -0.19281 | -0.22343 | 0.10611 | 1.39479 | 0.250295 | 0.037573103 | 0.36756 | 0.08988 | 0.05941 |
| HUO9 | -0.254 | -0.08194 | -0.34022 | -0.16079 | 0.10602 | 1.39343 | 0.250746 | 0.037551965 | 0.28579 | 0.00781 | 0.13232 |
| KM12 | -0.1714 | 0.103656 | -0.2101 | -0.2064 | 0.1056 | 1.38735 | 0.252783 | 0.116960506 | 0.23689 | 0.07154 | 0.0752 |
| PGA1 | -0.2377 | -0.1585 | -0.36909 | -0.15611 | 0.10535 | 1.38369 | 0.254014 | 0.048256811 | 0.1358 | 0.00417 | 0.1395 |
| COLO205 | -0.0256 | 0.00828 | -0.32196 | 0.004109 | 0.10432 | 1.36849 | 0.259194 | 0.430031262 | 0.47725 | 0.0113 | 0.4887 |
| RPE1SS51 | -0.2299 | 0.110502 | -0.19505 | -0.08859 | 0.10429 | 1.36801 | 0.259356 | 0.054117435 | 0.22245 | 0.08733 | 0.27035 |
| SW579 | -0.169 | 0.089278 | -0.08802 | -0.26866 | 0.10427 | 1.36773 | 0.259453 | 0.120358318 | 0.26876 | 0.27164 | 0.02962 |
| EFM19 | -0.2506 | 0.060291 | -0.2165 | -0.16873 | 0.10413 | 1.36581 | 0.260114 | 0.039617051 | 0.33874 | 0.06551 | 0.12074 |
| HCT116 | -0.0706 | 0.156397 | -0.21893 | 0.069743 | 0.10398 | 1.36362 | 0.260873 | 0.312954936 | 0.13905 | 0.06332 | 0.31516 |
| MHHES1 | -0.1715 | -0.01924 | -0.32946 | -0.2356 | 0.10351 | 1.35662 | 0.263305 | 0.116862497 | 0.44726 | 0.00974 | 0.04977 |
| SKMG1 | -0.0701 | 0.157051 | -0.23663 | -0.05641 | 0.10261 | 1.34348 | 0.267925 | 0.31439624 | 0.13803 | 0.04902 | 0.3486 |
| NB10 | -0.0587 | 0.00235 | -0.34291 | -0.07788 | 0.10238 | 1.34017 | 0.2691 | 0.342650042 | 0.49354 | 0.00739 | 0.29542 |
| PATU8988T | -0.1773 | 0.121665 | -0.21549 | -0.13069 | 0.10225 | 1.33827 | 0.269774 | 0.108967399 | 0.19999 | 0.06643 | 0.18283 |
| SNU1033 | -0.125 | 0.101709 | -0.25546 | -0.15635 | 0.10083 | 1.31763 | 0.277226 | 0.193498754 | 0.24108 | 0.03669 | 0.13912 |
| SEMK2 | -0.1698 | -0.002 | -0.23222 | -0.3132 | 0.10078 | 1.31692 | 0.277485 | 0.119161707 | 0.4945 | 0.05232 | 0.01339 |
| OPM2 | -0.1544 | 0.147082 | -0.11997 | -0.19369 | 0.10057 | 1.31377 | 0.27864 | 0.142198269 | 0.15404 | 0.20332 | 0.08887 |

| | | | | | | | | | | | |
|---|---|---|---|---|---|---|---|---|---|---|---|
| RPE1SS6 | -0.1396 | 0.053214 | -0.29716 | -0.05263 | 0.10053 | 1.31329 | 0.278818 | 0.166714817 | 0.3568 | 0.01806 | 0.35832 |
| UMRC3 | -0.1716 | 0.04012 | -0.25792 | -0.2411 | 0.10024 | 1.30906 | 0.280375 | 0.116693473 | 0.39103 | 0.03527 | 0.04583 |
| RPE1SS119 | -0.2595 | -0.04325 | -0.29026 | -0.09342 | 0.10013 | 1.30739 | 0.280994 | 0.034366567 | 0.38277 | 0.02044 | 0.25937 |
| HCC1359 | -0.1166 | 0.06416 | -0.18859 | -0.28157 | 0.09953 | 1.29874 | 0.284209 | 0.21008108 | 0.32901 | 0.09483 | 0.0238 |
| HPAFII | -0.2868 | 0.070665 | 0.04884 | -0.04276 | 0.09874 | 1.28725 | 0.288533 | 0.021717002 | 0.3129 | 0.36813 | 0.38406 |
| ONE58 | -0.2425 | 0.015626 | -0.20156 | -0.25391 | 0.09857 | 1.28482 | 0.289454 | 0.044885481 | 0.45711 | 0.08021 | 0.0376 |
| LK2 | -0.1188 | 0.144226 | -0.17649 | -0.17737 | 0.09852 | 1.28416 | 0.289707 | 0.205522666 | 0.15883 | 0.11009 | 0.10893 |
| 93T449 | -0.2389 | 0.00183 | -0.20616 | -0.27107 | 0.09833 | 1.28139 | 0.290764 | 0.047359452 | 0.49497 | 0.07545 | 0.02845 |
| SUM1315MO2 | -0.2291 | -0.02946 | -0.27017 | -0.26892 | 0.09808 | 1.27775 | 0.292155 | 0.054724884 | 0.41952 | 0.02888 | 0.02949 |
| TTC549 | -0.1306 | -0.00273 | -0.32855 | -0.07149 | 0.09765 | 1.27156 | 0.294534 | 0.182929167 | 0.49249 | 0.00992 | 0.31088 |
| NP3 | -0.1271 | -0.00276 | -0.14402 | -0.3367 | 0.09615 | 1.2499 | 0.303 | 0.189529412 | 0.49241 | 0.15918 | 0.0084 |
| JMSU1 | -0.1749 | 0.122471 | -0.07247 | -0.20977 | 0.09564 | 1.24266 | 0.305881 | 0.112264346 | 0.19842 | 0.3085 | 0.07186 |
| SMZ1 | -0.2136 | 0.058632 | -0.19085 | 0.056521 | 0.0956 | 1.24201 | 0.30614 | 0.068206262 | 0.34294 | 0.09215 | 0.34832 |
| CAL120 | -0.1826 | 0.160832 | -0.0153 | -0.14931 | 0.09533 | 1.23817 | 0.307674 | 0.102126662 | 0.13226 | 0.45801 | 0.15035 |
| NB13 | -0.2581 | -0.05601 | -0.29628 | -0.16378 | 0.09465 | 1.22842 | 0.311609 | 0.035196047 | 0.34963 | 0.01835 | 0.12787 |
| HMC18 | -0.1885 | 0.047705 | -0.23652 | -0.21837 | 0.09451 | 1.2264 | 0.312432 | 0.094977588 | 0.37109 | 0.04909 | 0.06382 |
| EBC1 | -0.0557 | 0.086184 | -0.28207 | -0.07788 | 0.09428 | 1.22311 | 0.31377 | 0.350372202 | 0.27589 | 0.0236 | 0.29544 |
| DKMG | -0.0818 | 0.05281 | -0.17514 | -0.28995 | 0.09408 | 1.22023 | 0.314951 | 0.286056615 | 0.35784 | 0.11189 | 0.02055 |
| SW1463 | -0.1946 | 0.159038 | -0.09927 | 0.079395 | 0.0938 | 1.21621 | 0.316601 | 0.087887166 | 0.13498 | 0.2464 | 0.29182 |
| MDAMB436 | -0.2005 | 0.142849 | -0.12311 | -0.11933 | 0.09339 | 1.21038 | 0.319009 | 0.081309544 | 0.16118 | 0.19719 | 0.20456 |
| NCIH1573 | 0.16204 | 0.288737 | 0.206144 | 0.37752 | 0.093 | 1.20486 | 0.321302 | 0.130456462 | 0.021 | 0.07546 | 0.00344 |
| ESO51 | -0.0816 | 0.254809 | -0.07997 | 0.003488 | 0.0929 | 1.20334 | 0.321936 | 0.286594303 | 0.03707 | 0.29045 | 0.49041 |
| SUDHL8 | 0.02397 | 0.290583 | -0.06734 | 0.055629 | 0.09286 | 1.2028 | 0.32216 | 0.434374985 | 0.02032 | 0.32109 | 0.3506 |
| CHP134 | -0.3132 | -0.17438 | -0.26863 | -0.21839 | 0.09278 | 1.20163 | 0.322652 | 0.013385417 | 0.11291 | 0.02963 | 0.06379 |
| L428 | -0.3007 | 0.014102 | -0.10516 | -0.01524 | 0.09192 | 1.18945 | 0.327781 | 0.01692027 | 0.46128 | 0.23368 | 0.45817 |
| T84 | -0.3067 | -0.08985 | -0.18978 | -0.25119 | 0.09179 | 1.18751 | 0.328606 | 0.015152276 | 0.26745 | 0.09341 | 0.03924 |
| OCIMY7 | -0.3111 | -0.11959 | -0.21217 | -0.0781 | 0.09168 | 1.1859 | 0.329291 | 0.013949133 | 0.20405 | 0.06954 | 0.2949 |
| KP2 | -0.1159 | 0.274539 | 0.056513 | 0.142251 | 0.09039 | 1.16768 | 0.337132 | 0.211350589 | 0.02684 | 0.34834 | 0.1622 |
| SNU2535 | -0.3215 | -0.05699 | -0.11541 | -0.17413 | 0.08975 | 1.15848 | 0.341151 | 0.011411483 | 0.34711 | 0.2124 | 0.11325 |
| SW948 | -0.1399 | 0.181448 | -0.04449 | 0.178808 | 0.08955 | 1.15569 | 0.342381 | 0.166285028 | 0.10364 | 0.3795 | 0.10704 |
| SMSCTR | -0.242 | 0.036815 | -0.21759 | -0.06112 | 0.0895 | 1.15498 | 0.342691 | 0.045198174 | 0.39982 | 0.06451 | 0.33663 |
| INA6 | -0.1617 | -0.04457 | -0.26814 | -0.30205 | 0.08947 | 1.1546 | 0.342863 | 0.130988551 | 0.37929 | 0.02987 | 0.01651 |
| NO11 | -0.2162 | -0.009 | -0.16778 | -0.28843 | 0.08937 | 1.15312 | 0.343512 | 0.06579474 | 0.47528 | 0.12207 | 0.02111 |
| HEC251 | -0.25 | 0.083097 | -0.10988 | 0.034313 | 0.08933 | 1.15264 | 0.343726 | 0.039948895 | 0.28308 | 0.22374 | 0.4065 |
| VAESBJ | -0.1678 | 0.190125 | -0.07819 | 0.076643 | 0.08894 | 1.1471 | 0.346183 | 0.122100595 | 0.093 | 0.29468 | 0.2984 |
| NP8 | -0.3124 | -0.15641 | -0.24426 | -0.19397 | 0.08788 | 1.1321 | 0.352913 | 0.013592493 | 0.13903 | 0.04368 | 0.08855 |
| RL | -0.2686 | 0.026554 | -0.14382 | -0.18459 | 0.08774 | 1.13016 | 0.35379 | 0.029632274 | 0.42738 | 0.15952 | 0.09969 |
| UPCISCC029A | -0.1922 | 0.027877 | -0.20206 | -0.24887 | 0.08732 | 1.12417 | 0.356514 | 0.090567802 | 0.4238 | 0.07969 | 0.04068 |
| ISTMEL1 | -0.3053 | -0.02731 | -0.00769 | -0.03747 | 0.08703 | 1.12004 | 0.358402 | 0.015549492 | 0.42533 | 0.47888 | 0.39808 |
| KIJK | -0.3088 | -0.03083 | -0.15111 | -0.09322 | 0.08674 | 1.11605 | 0.360231 | 0.014565841 | 0.41584 | 0.14743 | 0.25982 |
| NB7 | -0.2418 | -0.03452 | -0.23036 | -0.01067 | 0.08659 | 1.11389 | 0.361229 | 0.045370138 | 0.40596 | 0.05377 | 0.47068 |
| SCH | -0.2004 | 0.178962 | 0.021702 | -0.04012 | 0.08655 | 1.11333 | 0.361484 | 0.081478645 | 0.10684 | 0.44054 | 0.39104 |
| CCLP1 | -0.1716 | 0.027674 | -0.24978 | -0.21669 | 0.08646 | 1.11201 | 0.362096 | 0.116699347 | 0.42435 | 0.04011 | 0.06533 |
| RCHACV | -0.3106 | -0.15891 | -0.24124 | -0.19201 | 0.08645 | 1.11195 | 0.362123 | 0.014067565 | 0.13517 | 0.04573 | 0.0908 |
| OACM51 | -0.18 | 0.018522 | -0.25696 | -0.21342 | 0.08635 | 1.11053 | 0.362782 | 0.105490521 | 0.44921 | 0.03582 | 0.06836 |
| BT12 | -0.2321 | 0.050462 | -0.14199 | -0.20329 | 0.08578 | 1.1025 | 0.366516 | 0.052404834 | 0.36391 | 0.16265 | 0.0784 |
| P2URK562 | -0.297 | -0.19394 | -0.255 | -0.25105 | 0.0857 | 1.10142 | 0.367021 | 0.018114314 | 0.08858 | 0.03695 | 0.03933 |
| HB1119 | -0.1716 | 0.024941 | -0.15123 | -0.27927 | 0.08559 | 1.09976 | 0.367795 | 0.116762794 | 0.43175 | 0.14723 | 0.02476 |
| EW7 | -0.2396 | -0.01427 | -0.07695 | -0.25968 | 0.08536 | 1.09659 | 0.369282 | 0.046853744 | 0.46084 | 0.29765 | 0.03429 |
| PF382 | -0.3011 | -0.02288 | -0.11562 | -0.16854 | 0.08501 | 1.09166 | 0.371606 | 0.016804462 | 0.43734 | 0.21198 | 0.121 |
| NB1 | -0.1235 | 0.073218 | -0.25153 | -0.01891 | 0.08445 | 1.08386 | 0.375308 | 0.196432499 | 0.30667 | 0.03903 | 0.44814 |
| BLUE1 | -0.2547 | -0.14436 | -0.23744 | -0.31283 | 0.0844 | 1.08309 | 0.375676 | 0.037150328 | 0.1586 | 0.04843 | 0.01348 |
| NCIH23 | -0.1968 | 0.011586 | -0.27155 | -0.11767 | 0.08416 | 1.07969 | 0.377299 | 0.085337546 | 0.46818 | 0.02822 | 0.20787 |
| LN340 | -0.1839 | 0.074309 | -0.22518 | -0.06824 | 0.08397 | 1.07709 | 0.378544 | 0.100593405 | 0.30402 | 0.05795 | 0.31888 |
| AML193 | -0.209 | -0.04651 | -0.2454 | -0.27688 | 0.08377 | 1.07431 | 0.37988 | 0.072624593 | 0.37421 | 0.04292 | 0.02579 |
| PSS131R | -0.2511 | 0.00423 | -0.219 | -0.16082 | 0.08365 | 1.07265 | 0.38068 | 0.039267295 | 0.48837 | 0.06326 | 0.13227 |
| GIMEN | -0.1786 | 0.055715 | -0.12856 | 0.130601 | 0.08339 | 1.06896 | 0.382463 | 0.107310191 | 0.35038 | 0.1868 | 0.183 |
| A2780 | -0.0548 | 0.239027 | -0.00739 | -0.08361 | 0.08302 | 1.06376 | 0.384984 | 0.352693276 | 0.04729 | 0.4797 | 0.28188 |
| MFM223 | -0.1829 | 0.042777 | -0.11228 | -0.25351 | 0.08227 | 1.05328 | 0.390116 | 0.101807849 | 0.38401 | 0.21878 | 0.03783 |
| OCIM2 | -0.1427 | 0.027327 | -0.23098 | 0.051078 | 0.08131 | 1.04001 | 0.396686 | 0.161390165 | 0.42529 | 0.05328 | 0.36232 |
| MPNST724 | -0.1547 | 0.075602 | -0.09779 | -0.23629 | 0.08109 | 1.03683 | 0.398277 | 0.141737617 | 0.3009 | 0.24964 | 0.04926 |
| KPNYS | -0.0824 | 0.008902 | -0.14734 | -0.30314 | 0.0809 | 1.03426 | 0.399566 | 0.284738741 | 0.47554 | 0.1536 | 0.01618 |
| SKNO1 | -0.2257 | -0.09131 | -0.18702 | -0.313 | 0.08084 | 1.0334 | 0.399996 | 0.057529653 | 0.26414 | 0.09672 | 0.01344 |
| MEL285 | -0.1676 | 0.035562 | -0.12095 | -0.264 | 0.08058 | 1.02984 | 0.40179 | 0.1222845 | 0.40316 | 0.20138 | 0.03197 |
| HDMYZ | -0.0979 | 0.01271 | -0.24772 | 0.058702 | 0.08058 | 1.02974 | 0.401839 | 0.249470786 | 0.4651 | 0.04141 | 0.34276 |
| WM3772F | -0.109 | 0.260505 | 0.075983 | 0.141537 | 0.08014 | 1.02364 | 0.404925 | 0.225676637 | 0.03384 | 0.29998 | 0.16343 |
| VMRCRCW | -0.2812 | -0.16727 | -0.20396 | -0.07043 | 0.07954 | 1.0154 | 0.40912 | 0.023972645 | 0.12281 | 0.07769 | 0.31348 |
| LPS510 | -0.1736 | 0.180915 | 0.030116 | -0.05785 | 0.0795 | 1.01485 | 0.4094 | 0.114008476 | 0.10432 | 0.41777 | 0.34492 |
| UMUC13 | -0.1827 | 0.025403 | -0.15499 | -0.25408 | 0.07885 | 1.00585 | 0.414028 | 0.102042919 | 0.4305 | 0.14124 | 0.0375 |
| KMS18 | -0.2114 | -0.04364 | -0.23029 | -0.26522 | 0.07873 | 1.00411 | 0.414927 | 0.070276631 | 0.38173 | 0.05382 | 0.03134 |
| CHLA90 | -0.2898 | 0.00644 | -0.07237 | -0.09854 | 0.07871 | 1.00387 | 0.415054 | 0.020616308 | 0.4823 | 0.30874 | 0.248 |
| SKMM2 | -0.1767 | -0.04698 | -0.29818 | -0.08985 | 0.07837 | 0.99911 | 0.41752 | 0.109808463 | 0.37297 | 0.01772 | 0.26745 |
| MERO48A | -0.0636 | 0.174359 | 0.024649 | -0.15743 | 0.0783 | 0.9982 | 0.417995 | 0.330407988 | 0.11294 | 0.43254 | 0.13744 |
| MUTZ8 | -0.0798 | 0.092345 | -0.22117 | 0.04131 | 0.07813 | 0.99578 | 0.419258 | 0.290823272 | 0.26179 | 0.06135 | 0.38788 |
| D458 | -0.024 | 0.105981 | -0.21769 | 0.053112 | 0.07798 | 0.99372 | 0.420331 | 0.434238572 | 0.23193 | 0.06443 | 0.35706 |
| IGR1 | -0.0942 | 0.007737 | -0.29121 | -0.16816 | 0.07797 | 0.99368 | 0.420351 | 0.257522625 | 0.47873 | 0.02009 | 0.12154 |
| NCIH2009 | -0.2073 | 0.055765 | -0.2005 | -0.04089 | 0.07796 | 0.99346 | 0.420469 | 0.074314323 | 0.35025 | 0.08134 | 0.38901 |
| CORL279 | -0.1308 | -0.10465 | -0.19551 | -0.34488 | 0.07764 | 0.9891 | 0.422753 | 0.18262677 | 0.23476 | 0.08681 | 0.00709 |
| DMS53 | -0.1839 | 0.07912 | -0.10854 | -0.19917 | 0.0772 | 0.98295 | 0.425992 | 0.100516261 | 0.29247 | 0.22654 | 0.08277 |
| NCIH526 | -0.1292 | 0.010783 | -0.27555 | -0.18288 | 0.07693 | 0.97921 | 0.427974 | 0.185510085 | 0.47038 | 0.02638 | 0.10183 |
| HCT15 | -0.0632 | 0.248431 | -0.01199 | -0.0266 | 0.07617 | 0.96879 | 0.433526 | 0.331298276 | 0.04096 | 0.46706 | 0.42725 |
| HCC2998 | -0.0183 | 0.195939 | -0.14682 | -0.00527 | 0.07615 | 0.96855 | 0.433658 | 0.449799549 | 0.08633 | 0.15447 | 0.48552 |
| ONDA9 | -0.2565 | 0.03248 | -0.14619 | -0.11923 | 0.07606 | 0.96721 | 0.434376 | 0.036109517 | 0.41141 | 0.15552 | 0.20477 |
| LI7 | -0.1299 | 0.026863 | -0.16307 | -0.26382 | 0.07455 | 0.94654 | 0.445578 | 0.184245665 | 0.42654 | 0.12892 | 0.03207 |
| NALM16 | -0.1446 | -0.04685 | -0.30408 | -0.18223 | 0.07409 | 0.94019 | 0.449066 | 0.158201116 | 0.37331 | 0.0159 | 0.10265 |
| NCIH2887 | -0.1132 | 0.145288 | -0.07947 | -0.15617 | 0.07334 | 0.92991 | 0.454752 | 0.216822344 | 0.15704 | 0.29165 | 0.13941 |
| NB1643 | 0.02712 | 0.189805 | -0.13011 | -0.05927 | 0.0729 | 0.92387 | 0.458119 | 0.425845803 | 0.09338 | 0.1839 | 0.34131 |
| SNU626 | -0.2014 | 0.103739 | -0.11975 | 0.014696 | 0.07284 | 0.9231 | 0.458549 | 0.080342212 | 0.23671 | 0.20374 | 0.45966 |
| IGROV1 | -0.2979 | -0.16613 | -0.14723 | -0.1862 | 0.07275 | 0.92193 | 0.459207 | 0.017828109 | 0.12444 | 0.15379 | 0.09772 |
| PCI30 | -0.0822 | 0.200289 | -0.07614 | -0.06585 | 0.07164 | 0.9067 | 0.467799 | 0.285251504 | 0.08157 | 0.29961 | 0.32479 |
| MERO14 | -0.2062 | 0.100466 | -0.02027 | -0.12414 | 0.07161 | 0.90636 | 0.467992 | 0.075446854 | 0.24378 | 0.44443 | 0.19519 |
| KMS11 | -0.0998 | 0.043001 | -0.22751 | -0.19721 | 0.07097 | 0.89763 | 0.472971 | 0.245133555 | 0.38342 | 0.05603 | 0.08492 |
| MV411 | -0.1472 | -0.11201 | -0.12544 | -0.3279 | 0.07038 | 0.88951 | 0.477642 | 0.153854224 | 0.21934 | 0.1927 | 0.01005 |
| MFE296 | -0.1623 | -0.04292 | -0.21047 | -0.27922 | 0.06997 | 0.88397 | 0.480844 | 0.130122741 | 0.38363 | 0.07117 | 0.02478 |
| HT3 | -0.0078 | 0.232217 | -0.07279 | -0.00838 | 0.06965 | 0.87971 | 0.483321 | 0.47856433 | 0.05232 | 0.30771 | 0.47697 |
| SUDHL4 | -0.1245 | 0.204226 | 0.024557 | -0.03897 | 0.06925 | 0.87418 | 0.486543 | 0.194544189 | 0.07742 | 0.43279 | 0.39409 |
| KATOIII | -0.182 | 0.012398 | -0.06392 | -0.24136 | 0.06888 | 0.86917 | 0.489484 | 0.103002302 | 0.46595 | 0.32959 | 0.04565 |
| OVK18 | -0.1705 | 0.062058 | -0.19862 | -0.12464 | 0.06851 | 0.86418 | 0.492422 | 0.118303113 | 0.33428 | 0.08337 | 0.19423 |
| NCIH446 | -0.2047 | -0.05528 | -0.25221 | -0.2068 | 0.06753 | 0.85099 | 0.500243 | 0.076907924 | 0.3515 | 0.03861 | 0.0748 |
| L1236 | -0.1437 | 0.187037 | -0.00438 | 0.101802 | 0.06742 | 0.84941 | 0.501191 | 0.159729516 | 0.0967 | 0.48795 | 0.24088 |
| PC3 | -0.1959 | -0.00218 | -0.04116 | -0.22914 | 0.06695 | 0.84312 | 0.504956 | 0.08633466 | 0.49399 | 0.38827 | 0.05472 |
| CAL72 | -0.1284 | 0.025188 | -0.18887 | -0.22759 | 0.06554 | 0.82414 | 0.516447 | 0.187123402 | 0.43108 | 0.09449 | 0.05597 |
| VMRCLCD | -0.2568 | -0.11808 | -0.21023 | -0.2201 | 0.06554 | 0.82411 | 0.516463 | 0.035938033 | 0.20706 | 0.07141 | 0.06228 |
| KPL1 | -0.1339 | 0.19058 | 0.002293 | -0.03362 | 0.06548 | 0.82334 | 0.516935 | 0.177029363 | 0.09247 | 0.4937 | 0.40835 |
| UHO1 | -0.214 | 0.065816 | -0.08193 | -0.13509 | 0.06488 | 0.81523 | 0.521901 | 0.067834304 | 0.32487 | 0.28582 | 0.1748 |
| LS1034 | -0.1399 | 0.046828 | -0.21857 | -0.13642 | 0.06399 | 0.80332 | 0.529257 | 0.166209784 | 0.37338 | 0.06364 | 0.17241 |
| 95T1000 | -0.1936 | 0.023895 | -0.12917 | -0.20577 | 0.06371 | 0.79957 | 0.531588 | 0.088942932 | 0.43458 | 0.18565 | 0.07585 |
| EW1 | -0.1516 | 0.026588 | -0.23448 | -0.07826 | 0.06361 | 0.79823 | 0.532421 | 0.146720768 | 0.42729 | 0.0506 | 0.29453 |
| ES5 | -0.0997 | -0.00849 | -0.27131 | -0.13081 | 0.06221 | 0.77952 | 0.544158 | 0.245474711 | 0.47666 | 0.02834 | 0.18261 |
| WAOSEL | -0.196 | -0.00337 | -0.21764 | -0.13559 | 0.0613 | 0.76728 | 0.551927 | 0.086251778 | 0.49074 | 0.06447 | 0.17391 |
| UPMD1 | -0.2321 | 0.027154 | -0.11731 | -0.11747 | 0.06062 | 0.75828 | 0.557687 | 0.05238875 | 0.42576 | 0.20859 | 0.20828 |
| JVM3 | -0.2307 | -0.14937 | -0.24702 | -0.15499 | 0.05976 | 0.74681 | 0.565081 | 0.053501501 | 0.15026 | 0.04186 | 0.14125 |
| WM3211 | -0.1612 | 0.003358 | -0.19421 | -0.20817 | 0.05966 | 0.74545 | 0.565958 | 0.13170728 | 0.49077 | 0.08828 | 0.07343 |
| SUDHL10 | -0.079 | 0.024631 | -0.14146 | -0.24077 | 0.05839 | 0.72868 | 0.576885 | 0.292740878 | 0.43259 | 0.16357 | 0.04606 |
| LS | -0.2121 | 0.033338 | -0.10468 | -0.14653 | 0.05755 | 0.71752 | 0.584223 | 0.069615142 | 0.40911 | 0.2347 | 0.15495 |
| NCIH1373 | -0.19 | -0.04349 | -0.21592 | -0.04014 | 0.05678 | 0.70732 | 0.590975 | 0.093129411 | 0.38213 | 0.06603 | 0.39098 |
| PL4 | -0.1967 | 0.042058 | -0.15673 | -0.06235 | 0.05647 | 0.70329 | 0.593657 | 0.085496495 | 0.38591 | 0.13853 | 0.33355 |
| MM127 | -0.2273 | -0.08795 | -0.18806 | -0.20823 | 0.05518 | 0.68621 | 0.605098 | 0.056210157 | 0.2718 | 0.09547 | 0.07338 |
| ST486 | -0.235 | -0.1559 | -0.17858 | -0.22772 | 0.05491 | 0.68262 | 0.607515 | 0.050252374 | 0.13983 | 0.10734 | 0.05586 |

| | | | | | | | | | | | | |
|---|---|---|---|---|---|---|---|---|---|---|---|---|
| NZOV9 | -0.2369 | -0.12793 | -0.20417 | -0.19007 | 0.05456 | 0.67805 | 0.610602 | | 0.048843441 | 0.18798 | 0.07748 | 0.09307 |
| SCCOHT1 | -0.0769 | 0.085815 | -0.18709 | -0.0761 | 0.05313 | 0.65928 | 0.623358 | | 0.297667922 | 0.27674 | 0.09663 | 0.29969 |
| ARH77 | -0.1302 | 0.146666 | 0.013955 | 0.126589 | 0.05249 | 0.65093 | 0.629081 | | 0.183713318 | 0.15473 | 0.46169 | 0.19051 |
| A427 | -0.1114 | 0.168315 | -0.00621 | -0.0489 | 0.05236 | 0.64916 | 0.630293 | | 0.220602611 | 0.12132 | 0.48293 | 0.36797 |
| MOLM14 | -0.0888 | 0.040555 | -0.0886 | -0.21921 | 0.05198 | 0.64428 | 0.633653 | | 0.269843602 | 0.38988 | 0.27032 | 0.06307 |
| JURKAT | -0.1568 | 0.010587 | -0.20941 | -0.08344 | 0.05164 | 0.63975 | 0.636775 | | 0.138374939 | 0.47091 | 0.0722 | 0.28229 |
| RAJI | -0.1739 | 0.099183 | -0.00684 | -0.07539 | 0.05074 | 0.62801 | 0.644903 | | 0.113609885 | 0.24658 | 0.4812 | 0.30142 |
| COGN278 | -0.1774 | 0.00847 | -0.16067 | -0.16706 | 0.05025 | 0.62165 | 0.649327 | | 0.108926812 | 0.47672 | 0.1325 | 0.12311 |
| IM9 | -0.2011 | 0.049061 | 0.03064 | -0.0358 | 0.04829 | 0.59624 | 0.667117 | | 0.080694393 | 0.36755 | 0.41635 | 0.40252 |
| CMK115 | -0.1784 | -0.10773 | -0.10546 | -0.2463 | 0.04697 | 0.57906 | 0.67924 | | 0.107526705 | 0.22823 | 0.23303 | 0.04233 |
| LS411N | 0.02345 | 0.185225 | -0.0679 | 0.098308 | 0.0466 | 0.57435 | 0.682577 | | 0.435796849 | 0.09891 | 0.31971 | 0.2485 |
| SUPM2 | -0.1773 | 0.026677 | -0.13464 | -0.00801 | 0.04546 | 0.55956 | 0.693096 | | 0.108980078 | 0.42705 | 0.17562 | 0.47798 |
| SW403 | -0.0792 | 0.016253 | -0.20985 | -0.02632 | 0.04439 | 0.54577 | 0.702936 | | 0.292235738 | 0.4554 | 0.07178 | 0.42802 |
| OCIP5X | -0.2033 | -0.04043 | 0.034434 | -0.0333 | 0.04301 | 0.5281 | 0.715602 | | 0.078412615 | 0.39022 | 0.40618 | 0.40922 |
| SEKI | -0.1535 | -0.0079 | -0.14478 | 0.031145 | 0.04283 | 0.52578 | 0.717265 | | 0.143586572 | 0.47828 | 0.15789 | 0.415 |
| NCIH716 | -0.1353 | -0.00336 | -0.17087 | -0.15237 | 0.03828 | 0.46764 | 0.759142 | | 0.174504073 | 0.49075 | 0.11773 | 0.1454 |
| CORL23 | -0.1969 | -0.11817 | -0.05326 | -0.11441 | 0.03398 | 0.41332 | 0.798152 | | 0.085217221 | 0.20688 | 0.35667 | 0.21444 |
| COLO684 | -0.1269 | 0.012952 | -0.13141 | -0.13544 | 0.03139 | 0.38084 | 0.821193 | | 0.190002388 | 0.46443 | 0.1815 | 0.17417 |
| SLVL | -0.1905 | -0.058 | -0.0547 | -0.11755 | 0.03137 | 0.38057 | 0.821381 | | 0.092558477 | 0.34456 | 0.35299 | 0.2081 |
| RI1 | -0.1105 | 0.011178 | -0.12736 | -0.15297 | 0.03094 | 0.37516 | 0.825187 | | 0.222500328 | 0.46929 | 0.18906 | 0.14445 |
| MCIXC | -0.1289 | 0.000612 | -0.07934 | 0.063455 | 0.03024 | 0.3664 | 0.831321 | | 0.18608951 | 0.49832 | 0.29195 | 0.33077 |
| NP5 | -0.0835 | 0.087204 | -0.04164 | -0.09672 | 0.0287 | 0.34714 | 0.844685 | | 0.282041486 | 0.27353 | 0.38701 | 0.252 |
| COV318 | 0.03304 | 0.059843 | -0.05403 | -0.11399 | 0.02409 | 0.28999 | 0.883018 | | 0.409911756 | 0.33987 | 0.3547 | 0.21528 |
| TYKNU | -0.0641 | 0.031586 | -0.13286 | -0.00731 | 0.02246 | 0.26996 | 0.895844 | | 0.329169647 | 0.41381 | 0.17885 | 0.4799 |
| IMR32 | -0.0437 | 0.031966 | -0.13193 | -0.06535 | 0.02021 | 0.24231 | 0.912881 | | 0.381530962 | 0.41279 | 0.18055 | 0.32603 |
| KE37 | -0.0622 | -0.06212 | -0.16768 | -0.11922 | 0.0194 | 0.23243 | 0.918753 | | 0.333943086 | 0.33413 | 0.12222 | 0.20478 |

| average | EG1 | EG2 | EG3 | EG4 | determ |
|---|---|---|---|---|---|
| total | -0.3513 | 0.064998 | -0.22253 | -0.15759 | 0.21713 |
| p < 0,.01 | -0.5294 | 0.094098 | -0.21316 | -0.13848 | 0.35339 |
| p < 0.05 | -0.457 | 0.082575 | -0.23544 | -0.1529 | 0.29765 |
| p < 0.1 | -0.4257 | 0.074962 | -0.23822 | -0.16228 | 0.27312 |
