## Supplement Table S8C for "Single-molecule behavior and cell-growth regulation in human RTKs"

| | pR | | | | | | | | p-values in t-test | | | |
|---|---|---|---|---|---|---|---|---|---|---|---|---|
| | EG1 | EG2 | EG3 | EG4 | determ | F-value | p-value | | EG1 | EG2 | EG3 | EG4 |
| RK cluste | 0.68607 | 0.08412 | 0.25236 | 0.07154 | 0.43222 | 8.94459 | 1.87E-05 | | 1.9E-08 | 0.28069 | 0.03853 | 0.31075 |
| EC | 0.32425 | 0.34164 | 0.81447 | 0.41636 | 0.43003 | 8.865 | 2.03E-05 | | 0.0108 | 0.00759 | 3.1E-13 | 0.00132 |
| IDRC pos | 0.44216 | -0.0542 | 0.14096 | -0.1524 | 0.26764 | 4.29399 | 0.00483 | | 0.00065 | 0.35423 | 0.16444 | 0.14543 |
| IDRC | 0.2432 | -0.2695 | 0.15483 | -0.2038 | 0.23085 | 3.52654 | 0.01346 | | 0.04439 | 0.02919 | 0.1415 | 0.07782 |
| GxxxG | 0.08424 | 0.29511 | 0.03689 | -0.2619 | 0.2154 | 3.22579 | 0.02027 | | 0.2804 | 0.01874 | 0.39962 | 0.03308 |
| Linker | -0.5097 | -0.2688 | -0.3043 | -0.2486 | 0.21464 | 3.21126 | 0.02068 | | 7.8E-05 | 0.02953 | 0.01582 | 0.04084 |
| Tail | 0.09036 | -0.2631 | -0.3037 | -0.3289 | 0.1382 | 1.88432 | 0.12887 | | 0.26628 | 0.03246 | 0.01601 | 0.00984 |
| IDRE | -0.3388 | -0.1745 | -0.3235 | -0.3025 | 0.12359 | 1.65696 | 0.17586 | | 0.00805 | 0.11276 | 0.01097 | 0.01638 |
| CRAC | 1.60E-16 | -0.1687 | -0.1687 | -0.4117 | 0.1195 | 1.5947 | 0.19136 | | 0.5 | 0.1208 | 0.1208 | 0.00149 |
| IDRE pos | -0.2087 | -0.1895 | -0.3317 | -0.2874 | 0.08497 | 1.09109 | 0.37187 | | 0.07291 | 0.09369 | 0.00931 | 0.02151 |
| TM | -0.0886 | 0.10182 | -0.0339 | -0.1506 | 0.04753 | 0.58639 | 0.67406 | | 0.27037 | 0.24085 | 0.4075 | 0.14827 |
