## Supplement Table S9 for "Single-molecule behavior and cell-growth regulation in human RTKs"

| Parameter | Rave | SD | \|Rave\|/SD |
|---|---|---|---|
| *D* 1 | 0.0882 | 0.1931 | 0.45676 |
| *D* 2 | 0.0666 | 0.1936 | 0.34401 |
| *D* 3 | -0.0541 | 0.2017 | 0.26822 |
| *L* 1 | 0.2012 | 0.2039 | 0.98676 |
| *L* 2 | 0.0436 | 0.1885 | 0.2313 |
| *P* 1 | -0.0722 | 0.2637 | 0.2738 |
| *P* 2 | 0.2575 | 0.1957 | 1.31579 |
| *P* 3 | -0.1467 | 0.2469 | 0.59417 |
| *tau* 12 | -0.3238 | 0.1785 | 1.81401 |
| *tau* 13 | -0.235 | 0.2098 | 1.12011 |
| *tau* 21 | -0.1538 | 0.2453 | 0.62699 |
| *tau* 23 | -0.0559 | 0.2485 | 0.22495 |
| *tau* 31 | -0.3071 | 0.1973 | 1.55651 |
| *tau* 32 | -0.3292 | 0.1876 | 1.7548 |
